# Treatment of human oocytes with extracellular vesicles from follicular fluid during rescue in vitro maturation enhances maturation rates and modulates oocyte proteome and ultrastructure

**DOI:** 10.1101/2025.02.05.636623

**Authors:** Sofia Makieva, Mara D. Saenz-de-Juano, Carmen Almiñana, Stefan Bauersachs, Sandra Bernal-Ulloa, Min Xie, Ana G. Velasco, Natalia Cervantes, Maike Sachs, Susanne E. Ulbrich, Brigitte Leeners

**Author notes:** Authors contributed equally (Sofia Makieva, Mara D. Saenz-de-Juano). Correspondence: Sofia Makieva Department of Reproductive Endocrinology, University Hospital Zurich Frauenklinikstrasse 10, 8091 Zürich, Switzerland.

## Abstract

**Study question:** Could follicular fluid-derived extracellular vesicles (ffEVs) benefit human oocyte rescue *in vitro* maturation (rIVM)?

**Summary answer:** Supplementation of rIVM culture with ffEVs isolated from mature follicles enhanced oocyte maturation rates by >20%, inducing changes in oocyte protein profile and organelle distribution.

**What is already known:** IVM involves the culture of immature germinal vesicle (GV) oocytes under set laboratory conditions to allow for their transition to mature metaphase II (MII) stage, which is confirmed by the extrusion of the first polar body. Efficient IVM could circumvent aggressive controlled ovarian stimulation (COS), reduce the cost and broaden the repertoire of infertility treatments. Animal studies suggest that extracellular vesicles (EVs), membranous nanosized vesicles containing different molecular content (e.g. nucleic acids, proteins) and present in the ovarian follicular fluid could enhance oocyte maturation. The uptake of ffEVs by bovine, equine and feline oocytes, but not human, has been demonstrated.

**Study design, size, duration:** Women undergoing transvaginal oocyte retrieval after COS (n=83) were recruited to donate follicular fluid (n=54 single follicles) and/or immature GV oocytes (n=95). We aimed to: a) define differences in the protein cargo of ffEVs derived from human follicles containing mature (MII-ffEVs, n=10) versus immature (GV-ffEVs, n=5; metaphase I MI-ffEVs, n=5) oocytes, b) demonstrate the capacity of human GV oocytes to uptake MII-ffEVs and c) determine the effect of MII-ffEVs supplementation on oocyte maturation.

**Participants/materials, setting, methods:** ffEVs were isolated by ultracentrifugation. The protein content of ffEVs was analysed by mass spectrometry. The uptake of fluorescently-labelled MII-ffEVs by GV oocytes (n=15) was assessed by confocal microscopy. GVs were cultured for rIVM in a timelapse incubator with MII-ffEVs (n=45 GVs) or without (n=40 GVs) and extrusion of polar body denoted maturation. The impact of MII-ffEVs supplementation on IVM-matured oocytes was assessed through single-cell proteomics and intracellular organelles appearance on transmission electron microscopy (TEM).

**Main results and the role of chance:** We identified 1340 proteins in ffEVs, with proteins such as F12, IGKV1-39, FREM2, and C1QC being significantly enriched in MII-ffEVs. GV oocytes internalised MII-ffEVs, and their supplementation for 48 hours increased the oocyte maturation rate compared to control by 22.8±9.4% (77.8% vs 55% maturation rate respectively; p-value=0.0372). Proteomic analysis of ffEV-supplemented mature oocytes (n=6) revealed 56 differentially abundant proteins (DAPs) compared to not supplemented mature oocytes (n=5). Among them, 37 DAPs were in higher abundance in ffEVs- supplemented mature oocytes including Hyaluronan Synthase 1 (HAS1) that is associated with oocyte maturation (6.55 fold increase). Electron microscopy showed differences in oocyte organelle distribution and appearance, particularly that of endoplasmic reticulum (RE) and RE-mitochondria complexes. Functional enrichment analysis of differentially abundant proteins during ffEV-oocyte interaction revealed regulation of endoplasmic reticulum, steroid biosynthesis, and keratin organisation pathways.

**Large scale data:** **N/A**

**Limitations, reasons for caution:** This study utilised immature oocytes from COS cycles, therefore the results should be interpreted within the context of rIVM potential.

**Winder implications of the findings:** These results provide new insights into the role of ffEVs in enhancing oocyte maturation, offering potential improvements for clinical rIVM protocols and inspire the development of global IVM supplements based on ffEVs or associated specific cargo.

**Study funding/competing interest(s):** This work was funded by an EMDO research fellowship and a FAN research grant (Fonds zur Förderung des akademischen Nachwuchses) from the University of Zurich.

**What does it mean for the patients:** Infertility rates are rising, with 17% of couples worldwide needing help to get pregnant, often through treatments like in vitro fertilisation (IVF). IVF usually involves using hormones to stimulate the ovaries to produce multiple eggs, which can be tough on a woman’s health, both physically and emotionally, and can be very expensive. In vitro oocyte maturation (IVM) is a gentler alternative, where eggs are matured outside the body, reducing risks and costs. However, IVM isn’t as effective as IVF yet, mainly because the current methods are not perfect. Our research is exploring a new approach to improve IVM by adding extracellular vesicles from follicular fluid to the egg culture. This could help the eggs mature better, leading to higher success rates and giving more options to couples struggling with infertility.

## Introduction

Global fertility rates are falling below replacement thresholds in over half of all countries as of the year 2021; a decline that is anticipated to exert a significant influence on both economic and societal dynamics worldwide (Bhattacharjee *et al*., 2024). Presently, infertility poses a formidable challenge for one in six couples globally, necessitating recourse to medically assisted reproduction treatments (Harris, 2023). Of these, the most prominent intervention is controlled ovarian stimulation (COS) utilizing exogenous hormones aimed at growing multiple oocytes within small ovarian antral follicles (Macklon *et al*., 2006). After COS, mature oocytes are retrieved from large follicles to undergo *in vitro* fertilisation (IVF). Despite the rigor of protocols, approximately 30% of oocytes remain immature and ineligible for IVF (Mandelbaum *et al*., 2021). This percentage increases with milder COS regimes, a prerequisite for sensitive groups, such as cancer patients and subfertile women at risk of the fatal ovarian hyperstimulation syndrome (Nargund *et al*., 2022). Even milder stimulation approaches profoundly influence a couple’s experience undergoing infertility treatment (Njagi *et al*., 2023). For instance, COS necessitates the purchase of expensive hormones and frequent appointments to evaluate follicular growth. Furthermore, the process is not devoid of symptoms; it can induce physical discomfort and even be mentally taxing.

Several advancements aim to reduce the impact of COS. Among these is *in vitro* maturation (IVM), wherein multiple immature oocytes are retrieved from small antral follicles without prior COS and cultured extracorporeal in the laboratory (Gilchrist *et al*., 2024). Indeed, IVM enhances patient’s satisfaction by streamlining the treatment and reducing risks. This approach benefits not only patients who necessitate mild COS, comprising 15% of the IVF population, but also those with complete resistance to hormones (Vuong *et al*., 2020; Le *et al*., 2021). However, despite 80 years of the IVM concept, extensive livestock applications, and clinical success, healthcare providers remain wary of adopting a method they perceive as a significant departure from the current convention, namely COS (Rock and Menkin, 1944; Edwards, 1965; Merton *et al*., 2003; Roesner *et al*., 2017; Viana, 2020; Guo *et al*., 2023). Consequently, the clinical integration of IVM has been slow. It has particularly been hindered by the absence of standardisation and efficacy concerns, leading to its classification as non- experimental only in 2021 (De Vos *et al*., 2021; “In vitro maturation: a committee opinion,” 2021; Krisher, 2022; Makieva *et al*., 2024). Undermining IVM’s credibility further, oocytes matured *in vitro* are known to possess lower competence compared to their intracorporeally matured counterparts (De Vos *et al*., 2021). This disparity has also been highlighted by the only two clinical trials comparing the outcomes of IVM to COS, revealing lower live birth rates for the former (Vuong *et al*., 2020; Zheng *et al*., 2022). Nonetheless, IVM has achieved live birth rates exceeding 40% per IVF cycle in selected patients (Ho *et al*., 2019; MacKens *et al*., 2020; Mostinckx *et al*., 2024).

Currently, there are about ten different IVM variations in clinical usage. Refining these protocols is paramount for maximizing efficacy and addressing concerns (Gilchrist *et al*., 2024). In conventional or standard IVM (sIVM), immature oocytes with a germinal vesicle (GV) present in the cytoplasm are retrieved from small follicles and maintained surrounded by supportive cumulus cells, forming the cumulus-oocyte-complex (COC). Following exposure to hormones for up to 48 hours, the COCs are denuded, and oocyte maturity is confirmed by the presence of a polar body, denoting completion of nuclear maturation and oocyte arrest at the metaphase II stage of meiosis I. Conversely, rescue IVM (rIVM) is used to mature denuded GV oocytes retrieved after COS. The addition of hormonal supplements during rIVM is unnecessary due to the demonstrated unresponsiveness to stimulation; therefore, these oocytes are cultured in a standard IVF medium with the hope of maturing spontaneously within 24 hours. Indeed, most species’ oocytes, including humans —albeit at a lower frequency— could spontaneously resume meiosis once freed from follicles (Edwards, 1965). Given its seamless integration into conventional protocols and recent supportive findings regarding oocyte competence, rIVM may gain greater priority and acceptance in fertility clinics (Esbert *et al*., 2024). Successful implementation of rIVM has reportedly increased the pool of available oocytes for treatment while maintaining the integrity of conventional procedures and has led to healthy live births (Shani *et al*., 2023; Yuan *et al*., 2024).

Preclinical animal studies suggest a regulatory role for extracellular vesicles from follicular fluid (ffEVs) in governing oocyte maturation, highlighting their potential relevance to IVM (Pioltine *et al*., 2020a). The ffEVs, nano/micro-scale structures encapsulating nucleic acids and proteins, are released by various cells within the follicle and are considered indicators of oocyte competence and maturity (Santonocito *et al*., 2014). Studies on diverse animal species have documented the internalisation of ffEVs by oocytes, directly facilitating maturation (de Almeida Monteiro Melo Ferraz *et al*., 2020a; Uzbekova *et al*., 2020a). Our study aimed to investigate for the first time whether supplementation of human rIVM with ffEVs originating from mature MII oocyte-containing follicles could enhance GV oocyte maturation. Using follicular fluid and GV oocytes from women undergoing COS, our objectives included disclosing the “mature ffEVs” protein signature and determining whether the uptake of those ffEVs would increase the maturation rate of GV oocytes in an rIVM system.

## Methods

### Human Samples

Follicular fluid samples (n=54) and immature oocytes (n=95) were obtained from 83 women undergoing routine oocyte retrieval procedures for infertility treatment at the IVF center of University Hospital Zurich following written informed consent. The women recruited in our study had no previous diagnosed endocrine pathology and were undergoing fertility treatment for oocyte cryopreservation or unexplained infertility. The average age of the women was 36.4 ±2.7. All women received COS with gonadotrophins for up to 14 days to allow the simultaneous growth of multiple ovarian follicles before being triggered for superovulation. Oocyte retrieval took place 36 hours after triggering ovulation. Oocytes were denuded from cumulus cells for the purpose of intracytoplasmic sperm injection (ICSI) treatment or oocyte cryopreservation for social purposes. Oocytes that failed to mature despite COS were not used for treatment and, thus, were donated for research. The use of the aforementioned human material has been approved by the local ethics committee (BASEC-Nr. 2018-00797).

### Controlled ovarian stimulation (COS)

Prior stimulation women received a gestagen [10 mg/d] for 10 days up to 28 days, beginning at the second cycle day, in the short or antagonist protocols, and a GnRH-agonist [triptorelin, 0.1 mg/d] on cycle day 21 for the long protocol of COS. For COS, either short or long GnRH-agonist or GnRH-antagonist protocols were used with either hMG or recombinant FSH application. When at least three follicles with a diameter of ≥ 17 mm were observed during vaginal ultrasound, the final oocyte maturation was induced with either 6500 IE hCG or with the addition of a GnRH-agonist accompanied by about 1600 IE hCG within the GnRH-antagonist protocol. Ultrasound-guided oocyte retrieval was performed around 36 hours after administration of the hCG/GnRH-antagonist trigger.

### Oocyte handling

All cumulus-oocyte complexes (COCs) were cultured in a humidified incubator with conditions of 37°C, 20% O_2_, and 6% CO2 in suitable fertilisation media (Global for Fertilisation, CooperSurgical, Trumbull, USA or G-IVF Vitrolife, Gothenburg, Sweden) under oil overlay (OVOIL, Vitrolife, Gothenburg, Sweden). Two hours after retrieval, the oocytes were denuded using hyaluronidase enzyme (80 IU/mL). The denuded immature oocytes were subjected to cryopreservation using the Cryotop® method according to the Kitazato vitrification protocol (VT601, Kitazato, Fuji, Shizuoka, Japan) and stored in liquid nitrogen. In this method, an open vitrification carrier, which contains a polypropylene strip accompanied by a protective cover, is used. By aspirating the excess solution that is placed on the filmstrip, only a thin layer covering the cryopreserved cells ultimately remains. Using this minimal volume increases the cooling rate to 2300°C/min and the warming rate to 4210°C/min (Kuwayama, 2007). The oocytes were warmed for the experiments in this study using a thawing kit (VT602, Kitazato, Fuji, Shizuoka, Japan). Following warming, the immature oocytes were incubated in fertilisation media under an oil overlay for up to one hour before being used for experiments.

### Extracellular vesicle isolation from follicular fluid samples

The follicular fluid from single follicles was collected in preheated tubes kept at 37°C and placed in petri dishes for oocyte search. Once the COC was removed, the follicular fluid devoid of blood contamination was collected in tubes, centrifuged at 800 x g for 10 min, and the supernatant was stored at -80°C until further processing. Extracellular vesicles were isolated using differential centrifugation and ultracentrifugation. The follicular fluid samples were thawed on ice and centrifuged at 2000 g for 30 min at 4°C to remove cell debris. The supernatant was then transferred into 6.5 ml Ultraclear ultracentrifugation tubes (Beckman Coulter, Brea, USA) and centrifuged at 110,000 g at 4°C for 70 min in an Optimax XE90 ultracentrifuge (Beckman Coulter, Brea, USA) with a 50.2 Ti rotor (Beckman Coulter, Brea, USA). The pelleted ffEVs were washed with sterile filtered PBS (Gibco, Thermo Fisher Scientific, Waltham, USA) and ultracentrifuged again at 110,000 g at 4°C for 70 min. All procedures before and after ultracentrifugation were performed under the sterile hood to avoid external contamination of ffEVs. Depending on the downstream experiment, final ffEVs were collected in PBS or culture media.

### TEM analyses of ffEVs and oocytes

Extracellular vesicles and oocyte observations were performed by the Scientific Center for Optical and Electron Microscopy (ScopEM) of ETH Zurich (Zurich, Switzerland). For the TEM analysis of ffEVs, three microliters of the vortexed dispersion of EVs were placed on glow-discharged carbon-coated grids (Quantifoil, Germany) for 1 min. Negative contrast staining was performed in 2% sodium phosphotungstate at pH 7.2 for 1 second, followed by a second step for 15 sec. Excess moisture was drained with filter paper, and the air-dried grids were imaged using a TEM Morgagni 268 (Thermo Fisher) operated at 100 kV.

Oocyte ultrastructural analysis was done by TEM on oocytes fixed in 2.5% glutaraldehyde (EM grade; Polysciences Europe GmbH, Hirschberg an der Bergstrasse, Germany) and 2% formaldehyde (EM grade; Polysciences) in 0.1 M phosphate buffer (pH 7.4). To enable easy handling of the samples during the following washing and staining steps, the oocytes were embedded in freshly prepared low-gelling temperature agarose (4%; Carl Roth GmbH, Karlsruhe, Germany). After gelling on ice, small cubes containing one oocyte each were cut from the gelled block and washed three times in 0.15 M sodium cacodylate buffer. Then, the samples were postfixed with 2% osmium tetroxide (Polysciences) supplemented with 2.5% potassium hexacyanoferrate trihydrate and 2 mM calcium chloride. After washing again in cacodylate buffer, the samples were stained with 2% aqueous osmium tetroxide, 1% aqueous uranyl acetate (Polysciences) and Walton’s lead aspartate (Deerinck *et al*., 2010). Between the staining steps, the samples were washed in double distilled water. Then, the samples were dehydrated in increasing concentrations of ethanol and stepwise infiltrated with Epon (2x 30% Epon in ethanol, 2x 70% Epon, and 3x 100% Epon; Epoxy Embedding Kit, Sigma- Aldrich, Germany). Until 70% Epon, all steps were performed in a BioWave Pro+ Tissue Processor (Ted Pella Inc., Redding CA, USA). The samples were left in 100% Epon for 1 hour, 2 hours and 3 hours at RT on a shaker, then transferred into fresh Epon resin and polymerized at 60°C for 3 days. Thin sections of 50 nm were obtained with a diamond knife (Diatome Ltd., Switzerland) on a Leica UC7 ultramicrotome (Leica Microsystems, Heerbrugg, Switzerland), placed on formvar and carbon-coated TEM grids (Quantifoil, Großlöbichau, Germany), and stained with 2% uranyl acetate and Reynold’s lead citrate. The stained sections were then visualised using a Talos L120C TEM (Thermo Fisher Scientific, Eindhoven, The Netherlands) equipped with a Ceta-S camera at 80 kV. Image mosaics were recorded to cover the entire cross-section of each oocyte using the MAPS software (Thermo Fisher Scientific).

### Tunable Resistive Pulse Sensing (TRPS) and Western blotting

Particle concentration and size distribution of isolated ffEVs were performed using the qNano Gold system (Izon Science Ltd., Christchurch, New Zealand) and an NP400 Nanopore. The measurement was performed with undiluted samples and the CPC400 beads as the calibration standard. The number of particles analysed per sample was >1000. Data were processed with Izon Control Suite software version 3.3 (Izon Science). The protein concentration of ffEVs and western blotting was performed as previously described (Saenz-de-Juano *et al*., 2022). The protein concentration of ffEVs was measured using the Pierce™ Bicinchoninic Acid (BCA) Protein Assay Kit (Thermo Fisher Scientific, Waltham, USA) and the NanoDrop 2000 (Thermo Fisher Scientific, Waltham, USA). Samples of ffEVs diluted in RIPA buffer were mixed with 4x Laemmli Buffer (Bio-Rad, Hercules, USA) and incubated for 5 min at 95°C. If reducing conditions were necessary, 10% β-mercaptoethanol (Sigma-Aldrich) was added to the mix. Samples were loaded into a 4-20% Mini-PROTEAN TGX Stain-Free Precast Gel (Bio-Rad, Hercules, USA), and electrophoresis was performed at 200 V for 30 min. Proteins were transferred onto a 0.2 µm PVDF Trans-Blot Turbo Transfer Pack (Bio-Rad, Hercules, USA) using the Trans-Blot Turbo Transfer System (Bio-Rad, Hercules, USA) with transfer conditions of 1.3 A, 25 V, for 7 min. Immediately, the membrane was blocked with TBST containing 5% skim milk powder (Sigma-Aldrich) at room temperature for 1 hour. The membrane was then incubated overnight with rabbit anti-TSG-101 (1:500, PA531260, Thermo Fisher Scientific, Waltham, USA), mouse anti-CD63 (1:250, ab59479, Abcam, Cambridge, UK), and rabbit anti-MFGE8 (1:250, HPA002807, Sigma-Aldrich). After overnight incubation, the membrane was washed and incubated with the secondary antibodies: goat anti-rabbit IgG-HRP (1:10000, sc-2004, Santa Cruz Biotechnology, Dallas, USA) and goat anti-mouse IgM-HRP (1:10000, sc-2005, Santa Cruz Biotechnology, Dallas, USA). Precision Protein StrepTactin-HRP (Bio-Rad, Hercules, USA) was also added to visualise the ladder. Finally, Clarity™ ECL Substrate (Bio-Rad, Hercules, USA) was applied to the membrane, and bands were visualised with the ChemiDoc™ MP Imaging System (Bio- Rad, Hercules, USA).

### RNA extraction and miRNA qPCR

RNA cargo from ffEVs was isolated with the miRNeasy MicroKit (Qiagen, Hombrechtikon, Switzerland). The concentration of the RNA was determined using the Quantus™ Fluorometer and the QuantiFluor® RNA System kit (Promega, Dübendorf, Switzerland). The Agilent Pico Kit and the Agilent 2100 BioAnalyzer (Agilent Technologies, Basel, Switzerland) were utilised to assess the length of RNA fragments. We used the TaqMan™ Advanced miRNA Assays (Life Technologies, Zug, Switzerland) to evaluate the specific miRNA. A qualitative detection for the four miRNAs in four ffEVs samples from individual follicles was performed. Data were expressed as mean values of the quantification cycle (Cq). We used the TaqMan Advanced miRNA cDNA Synthesis Kit (A28007, Applied Biosystems) to generate cDNA from total RNA (2 µL of total RNA input), following the manufacturer’s instructions. The abundance of four miRNAs [hsa-let-7a-5p (478575_mir assay, Applied Biosystems), hsa-let-7b-5p (478576_mir, Applied Biosystems), hsa-miR-148a-3p (477814_mir, Applied Biosystems), and hsa-miR-223-3p (477983_mir assay, Applied Biosystems)] was measured using TaqMan Fast Advanced Master Mix (#4444556, Thermo Fisher Scientific, Life Technologies Europe BV, Zug, Switzerland). The real-time PCR reactions were performed in 384-well plates in a final volume of 10 μL. The PCR cycle parameters included 1 cycle of enzyme activation and cDNA denaturation at 95°C for 30 sec, followed by 40 cycles of denaturation at 95°C for 5 sec and annealing/extension at 60°C for 30 sec.

### Analysis of human ffEVs protein cargo by mass spectrometry

The proteome of ffEVs was analysed using liquid chromatography coupled to high-resolution tandem mass spectrometry (LC-MS/MS) at the Functional Genomic Centre Zurich (FGCZ, Zurich, Switzerland). The ffEVs samples were boiled for 10 min at 95°C followed by mechanical lysis using a tissue homogenizer (2×2 min cycles at 30Hz and high-intensity focused ultrasound (HIFU). Proteins were reduced and alkylated by adding Tris(2- carboxyethyl)phosphine and 2-chloroacetamide to a final concentration of 2 mM and 15 mM, respectively. Then, the samples were incubated for 30 min at 30°C (700 rpm, and light- protected). After dilution with pure ethanol to reach a final concentration of 60% EtOH (v/v), the KingFisher Flex System (Thermo Fisher Scientific, Waltham, USA) was used to bind the proteins to the carboxylated magnetic beads (hydrophobic and hydrophilic) and wash them. The enzymatic digestion was performed by adding trypsin in 50 mM triethylammonium bicarbonate buffer (TEAB) and incubating the samples overnight at 37°C. The remaining peptides were extracted from beads with water, and the two elutions were combined and dried down. The digested samples were dissolved in aqueous 3% acetonitrile with 0.1% formic acid, and the peptide concentration was estimated with the Lunatic UV/Vis absorbance spectrometer (Unchained Lab). Peptides were separated on an M-class UPLC and analysed on an Iontrap mass spectrometer (Thermo Fisher Scientific, Waltham, USA).

The acquired data were processed using the Fragpipe v17 (Kong *et al*., 2017). The spectra were searched against the *Homo sapiens* (SwissProt) protein background database, using Acetyl (Protein N-term) and Oxidation (M) as variable modifications and Carbamidomethyl (C) as fixed modifications. The protein identification results were imported for further analysis in the software Scaffold v5.2.0 (Proteome Software Inc., Portland, USA).

### Single oocyte proteomics analysis

For the oocyte sample preparation, 20 µl of 50 mM TEAB, 0.1% DDM were added for cell lysis using freeze-thaw cycles (Snap freeze samples in liquid nitrogen), followed by 10 min boiling at 95°C. This process was repeated 3 times. Proteins were reduced and alkylated by adding Tris(2-carboxyethyl) phosphine and chloroacetamid to a final concentration of 5 mM and 15 mM, respectively. The samples were incubated for 30 min at 30 °C (700 rpm and light-protected). Then, digestion was done in buffered trypsin solution at pH 8 (10 mM Tris/2 mM CaCl_2_). Samples were enzymatically digested using 2 µl of 4 µg/µl trypsin overnight plus an additional incubation for 3 h the next day with 2 µl of 4 µg/µl Trypsin. Digestion was stopped by acidifying the samples. Peptide concentration was estimated using the Lunatic UV/Vis absorbance spectrometer (Unchained Lab). Consequently, samples were loaded on Evotip (Evosep) following the manufacturer’s instructions. Samples were analysed on an Evosep One LC coupled to a TIMS TOF Pro mass spectrometer (Bruker), acquiring the data in diaPASEF scans.

Independent acquisition spectra were processed with DIA-NN (Demichev *et al*., 2020) using a library-free approach with the protein database *Homo sapiens* (SwissProt) with one sequence per gene. The modifications set for variable and fixed modifications were Oxidation (M) and Carbamidomethyl (C), respectively. The protein identification and quantification were performed using FragPipe (Demichev *et al*., 2022). To filter and normalise the data, a set of functions implemented in the R package [prolfqua] was used (Wolski *et al*., 2023). The resulting protein matrix was filtered to consider proteins with a minimum of 2 peptides/protein.

### Data mining and bioinformatics analysis of ffEVs protein cargo and oocyte proteomic profiles

Total identified proteins were filtered to select proteins present in at least 4 out of 5 samples in at least one group for oocyte data. For ffEVs data, the identified proteins were filtered to select proteins present in at least 4 out of 5 samples for GV-ffEVs and GVBD-ffEVs, and 9 out of 11 for MII-ffEVs data. Missing value imputation and statistical analysis were also performed with the DEP tool, a BioConductor R-tool for proteomics analysis (Zhang *et al*., 2018). The qri imputation method was used for oocytes, and the *man* imputation method was used for ffEVs. Subsequently, differential enrichment analysis was performed by applying empirical Bayes statistics on protein-wise linear models using limma (Smyth, 2004; Ritchie *et al*., 2015).

The data have been deposited to the ETH Research Collection repository and will be made publicly available with a dataset identifier upon publishing of the manuscript.

BioDBnet tool (Mudunuri *et al*., 2009) was used to convert Uniprot protein identifiers to gene symbols and NCBI Entrez Gene IDs. Then, human gene identifiers or symbols were used for subsequent functional annotation. The self-organizing tree algorithm software (SOTA, Multi Experiment Viewer software v.4.8.1, (Howe *et al*., 2011) was used to identify clusters of proteins/genes with similar expression profiles across experimental groups.

We used g:Profiler software (https://biit.cs.ut.ee/gprofiler/gost) with default parameters to obtain information about overrepresented biological functions and pathways in DAPs identified in ffEVs or single oocyte datasets. The protein-protein interactions between MII- ffEVs and treated-oocyte DAPs were obtained using STRING (Morris *et al*., 2018).

### Uptake of labelled ffEVs by oocytes

The ffEVs used for the uptake experiment were isolated from 50 ml of pooled follicular fluid from 10 patients. The ffEV sample was labelled using green Vybrant™ DiO solution dye (Thermo Fisher Scientific, Reinach, Switzerland). A total of nineteen GV oocytes from 4 different patients were thawed and distributed randomly in one of the three conditions as follows: A) 10 oocytes we co-incubated with G1 medium with labelled ffEVs, B) 5 oocytes were co-incubated with G1 medium with labelled PBS (control dye), and C) 4 oocytes were co-incubated with just G1 medium (control). GV oocytes were incubated for 24 hours, fixed, and individually evaluated under a confocal microscope Zeiss LSM 780 (Zeiss, Feldbach, Switzerland). To perform the ffEV uptake experiments, the ffEV pellet obtained after ultracentrifugation was resuspended in 250 µl of PBS. The resulting ffEV sample had a protein concentration of approximately 950 µg/ml. From this, 200 µl of ffEVs were labeled using green Vybrant™ DiO solution dye (Molecular Probes, Thermo Fisher Scientific, Waltham, USA). Briefly, one µl of DiO dye was mixed with 200 µl of the ffEV pellet diluted in PBS or 200 µl of PBS (used as a dye control). Both the dye-ffEV and PBS-ffEV solutions were vortexed and incubated for 20 min at 37°C in the dark. Subsequently, both solutions were centrifuged at 2000 x g for 5 min at 4°C to remove any potential dye aggregates. The resulting labeled ffEVs and labeled PBS solutions were resuspended in 50 µl of sterile G1- IVF medium and incubated with oocytes. GV oocytes were incubated for 24 hours at 37°C, 20% O_2_, and 6% CO_2_. The next day, oocytes were washed four times in PBS supplemented with 1% polyvinyl alcohol (PVA) and fixed in 4% paraformaldehyde (PFA) for 20 min at 37°C. Afterward, the oocytes were washed three times for 10 min each in PBS with 1% PVA and mounted on a microscope slide using VECTASHIELD PLUS antifade mounting medium with DAPI (H2000-10, Vector Labs, Newark, USA). Confocal images were acquired using a Zeiss LSM 780 confocal laser microscope (Carl Zeiss Microscopy GmbH, Oberkochen, Germany) with the objectives 40x 0.75 NA EC Plan-Neofluar Ph2 M27 and 63x 1.4 NA Oil Plan-Apochromat DIC M27. The acquired images were processed using Zen 2012 software (Carl Zeiss Microscopy GmbH, Oberkochen, Germany).

### In vitro maturation experiments

In total, three independent IVM experiments were performed. In each experiment, ffEVs for the in vitro treatment were obtained from a pool of follicular fluid (60-80 ml) from 12-17 different patients. In each experiment, oocytes at the GV stage were thawed and divided into the ffEV treatment group and control group. The first experiment used 31 oocytes from 17 patients (15 control, 16 ffEVs treatment),the second experiment used 31 oocytes from 22 patients (15 control, 16 ffEVs treatment) and the third experiment used 23 oocytes from 17 patients (10 control, 13 ffEVs treatment). The IVM culture was performed using a timelapse GERI Incubator (Genea Biomedx, Sydney, Australia) to record the time of the polar body extrusion. GV oocytes were incubated in individual microwells of a single GERI dish (Genea Biomedx) sharing 100 µl of G1-IVF medium supplemented with resuspended ffEVs pellet (ffEV treatment group) or in another GERI dish sharing 100 µl of G1-IVF medium (control group) during 48 hours, at 37°C and 6% CO_2_. After 48 hours, the maturation stage of the oocytes was evaluated individually under the microscope using morphological criteria. Additionally, the exact time of the polar body extrusion was noted using the GERI software (Genea Biomedx). The rIVM-matured oocytes were either individually snap-frozen in liquid nitrogen for proteomic analysis or fixed for TEM evaluation.

### Statistics

Data on maturation rates from three independent experiments were analysed using Fisher’s exact test in GraphPad Prism 9 v.9.2.0 (San Diego, California).

## Results

### Successful isolation of ffEVs from pools and individual follicles

Transmission electron microscopy (TEM) analysis confirmed the presence of heterogenic ffEVs populations in individual follicular fluid samples (Figure 1A, B). The size distribution evaluation with tunable resistive pulse sensing (TRPS) unveiled particle dimensions ranging from 50 to 700 nm (Supplementary Figure 1A). We successfully extracted measurable quantities of RNA from ffEVs derived from 1.5 – 2.0 mL of 4 patient-individual follicular fluid samples and obtained an average of 3.64 ± 2.12 ng RNA. The Bioanalyzer profiles showed that the majority of the RNA belonged to the small RNA population, with a minimal contribution from larger RNAs (Supplementary Figure 1B). Furthermore, within the isolated ffEVs, we identified the presence of previously reported follicular fluid EV-miRNAs (Martinez *et al*., 2018) (Supplementary Figure 1C). Immunoblotting results confirmed three known EV markers including transmembrane proteins CD63 (Supplementary Figure 1D), cytosolic protein tumor susceptibility gene 101 (TSG101, Supplementary Figure 1E), and milk fat globule-EGF factor 8 protein (MFGE8, Supplementary Figure 1F) in isolated ffEVs. Proteomic analysis by mass spectrometry of ffEVs derived from a pool of follicular fluid samples from different patients revealed that ffEVs also contained further exosomal protein markers such as CD81, CD9, FLOT1, FLOT2, and other proteins characteristic of EV biogenesis such as annexins, heat shock proteins, and Rab GTPases proteins (Supplementary Table 1).

**Fig 1.**
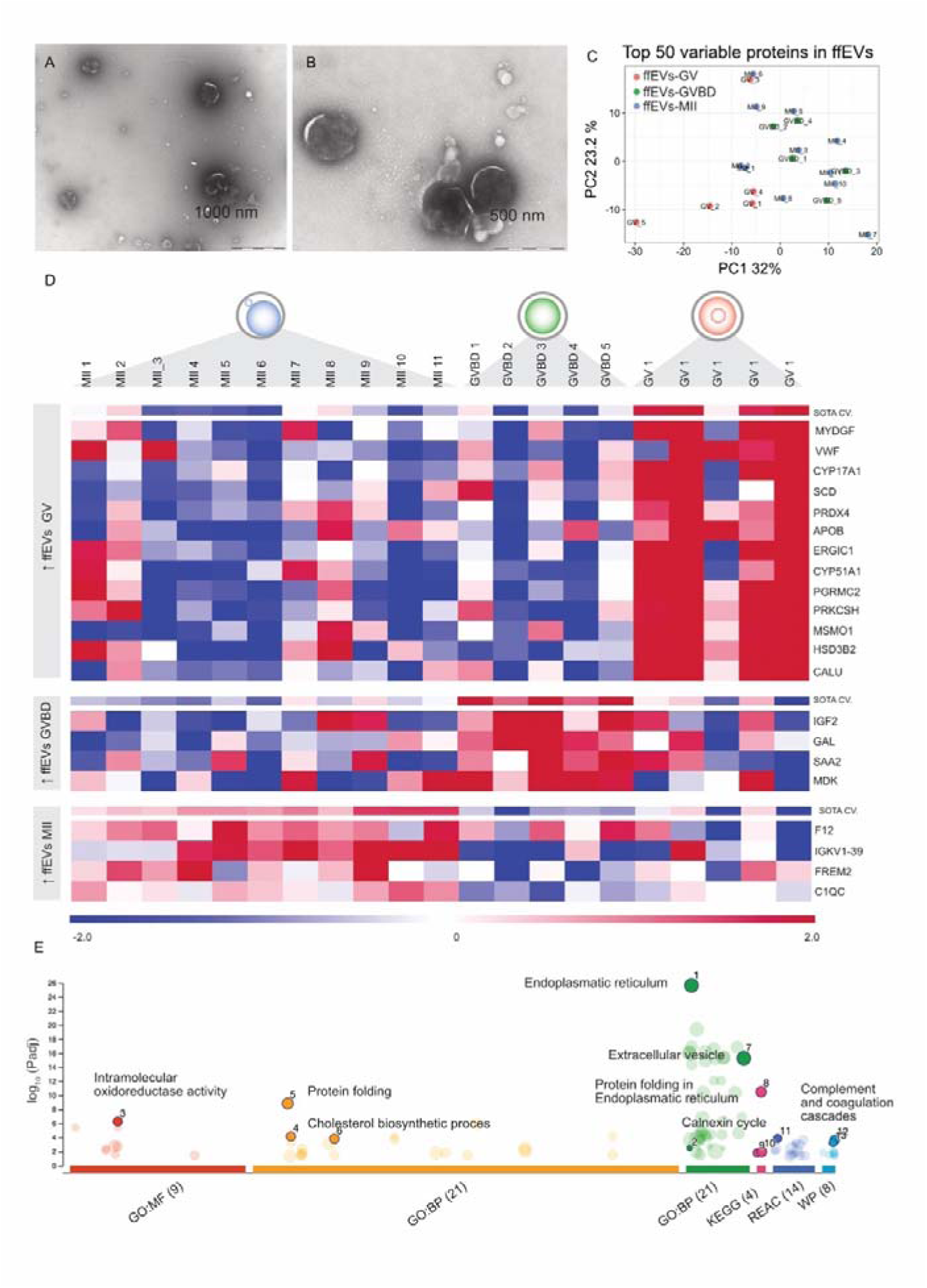
Follicular fluid EVs (ffEVs) protein cargo changed during oocyte maturation. (A, B) TEM analysis confirmed that ffEVs were successfully isolated and exhibited size heterogeneity. (C) Principal component analysis (PCA) based on the top 50 proteins with the greatest changes between all experimental groups (ffEVs-MII, ffEVs-GVBD, and ffEVs- GV). (D) Heatmap showing differential abundant proteins (DAPs) in ffEVs (FDR<0.1). Self- organizing tree algorithm (SOTA) analysis identified 3 clusters of proteins with similar expression profiles across different maturation stages. (E) GOSt multiquery plot of Manhattan showing gene ontology (GO) overrepresentation analysis results of 76 unique proteins in MII-ffEVs. BP: biological process; CC: cellular component; MF: molecular function; REAC: reactome; KEGG: Kyoto Encyclopedia of Genes and Genomes; WP: Wiki pathways.

### Differential protein cargo of human ffEVs derived from follicles containing mature or immature oocytes

We proceeded to analyse the proteomic cargo of ffEVs to identify proteins associated with maturation. Hence, we isolated ffEVs from single follicles containing a mature MII oocyte (n=11 patients; 1 ffEVs sample per patient), intermediate maturity GVBD (germinal vesicle breakdown) oocyte (n=5 patients; 1 ffEVs sample per patient), and immature GV oocyte (n=5 patients; 1 ffEVs sample per patient) to compare their proteomic cargo.

A total of 1340 proteins were identified across all ffEVs samples, regardless of their origin (Supplementary Table 2). Only proteins present in at least 9 out of 11 for the MII group and proteins present in 4 out of 5 GV or GVBD samples were selected for further statistical analysis, resulting in 789 filtered proteins (Supplementary Table 3).

A principal component analysis (PCA) based on the top 50 proteins with the greatest changes across the dataset showed a dynamic protein ffEV pattern, with GV-ffEVs samples separated from the rest of the samples based on principal component 1 (Figure 1C). To reveal differential abundant proteins (DAPs), three comparisons were statistically performed: MII versus (vs.) GV, GVBD vs. MII, and GVBD vs. GV, resulting in 14, 7, and 1 DAPs, respectively (False discovery range; FDR<0.1; Supplementary Table 4). The DAPs resulting from the three comparisons were further analysed for their expression profiles across samples using a Self-Organizing Tree Algorithm (SOTA) clustering analysis. This analysis provided three clear clusters (Figure 1D): cluster 1, with 13 proteins more abundant in GV-ffEVs; cluster 2, with 4 proteins more abundant in GVBD-ffEVs; and cluster 3, with 4 proteins more abundant in MII-ffEVs. The proteins exclusively associated with the MII-ffEVs cargo were the Coagulation Factor XII (F12), Complement C1q subcomponent subunit C (C1QC), FRAS1-related extracellular matrix protein 2 (FREM2), and Immunoglobulin Kappa Variable 1-39 (IGKV1-39).

When a cut-off of FDR<0.2 was used, 61, 15, and 14 DAPs were identified in the comparisons MII vs. GV, GVBD vs. MII, and GVBD vs. GV, respectively (Supplementary Table 5). A short-listing of the DAPs specific to the MII ffEVs identified 76 unique proteins. These proteins were used for the overexpression analysis using g:Profiler. A functional enrichment analysis highlighted that these DAPs were mainly related to pathways involving endoplasmic reticulum, oxidoreductase activity, complement cascade, extracellular vesicles, and steroid biosynthesis (Figure 1E; Supplementary Table 6).

### Immature oocytes uptake MII-ffEVs

Since we observed that certain proteins were enriched in ffEVs coming from mature follicles, we sought to examine if immature oocytes could take up the “maturity” signal by internalizing ffEVs. For studying the outcome of this internalisation, we co-incubated GV immature oocytes with fluorescently labelled ffEVs (coming from mature follicles) for 24 hours and performed confocal microscopy to identify any fluorescent signal inside the oocytes. Indeed, we found that oocytes had uptaken the ffEVs. Many localized in the zona pellucida (Figure 2A-J), the glycoprotein layer that functions as a mediator between the oocyte and the cumulus cells, which nourish the oocyte during *in vivo* oocyte maturation. It is possible that oocytes recruit the same mechanism to uptake ffEVs as the one used to internalize nutrients from cumulus cells.

**Fig 2.**
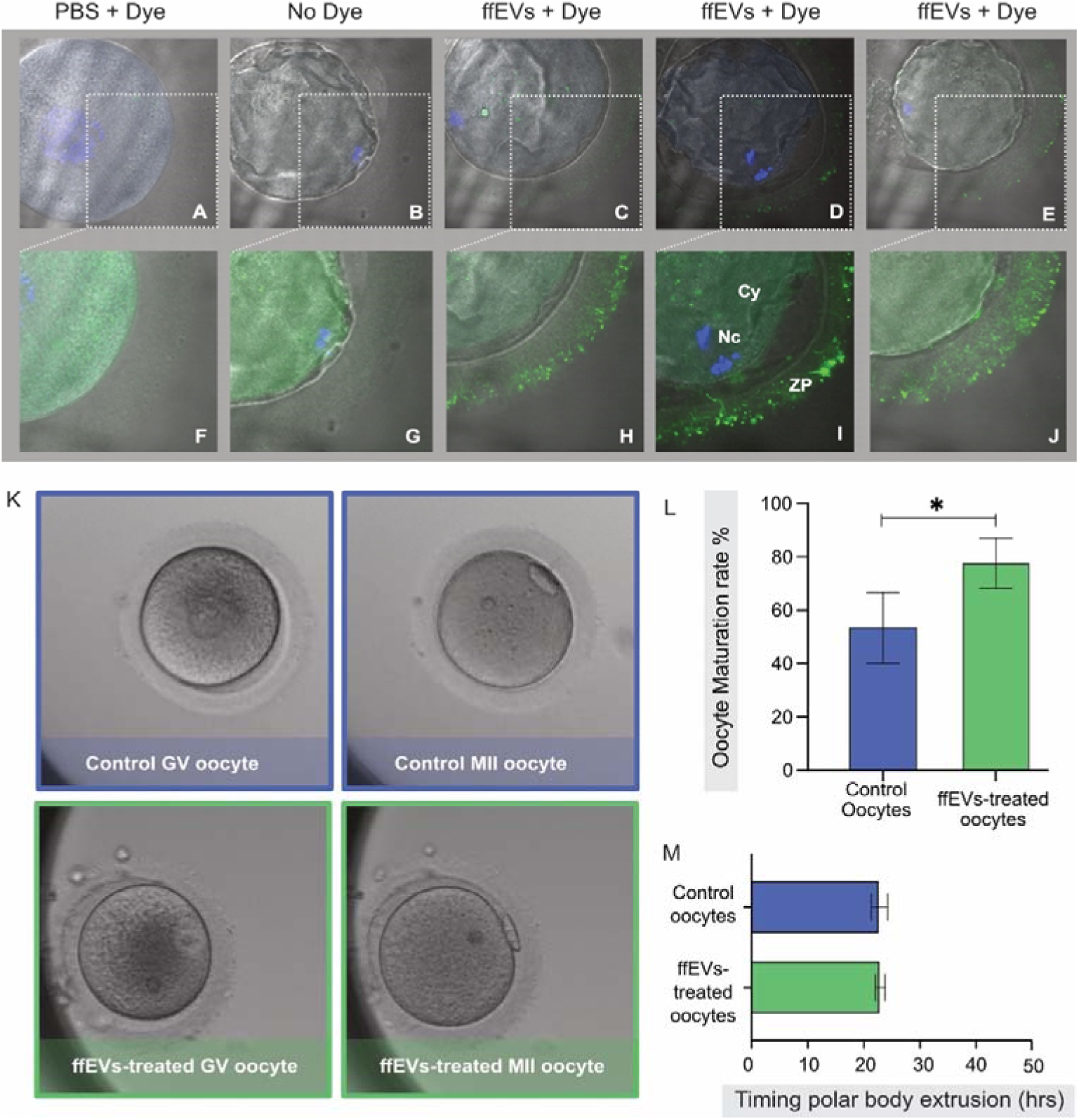
The uptake of MII-ffEVs by immature GV oocytes enhanced maturation rates. GV oocytes were cultured for 24 hours under three different treatment conditions: PBS + Dye (PBS labelled with Dye to act as control (A), no treatment (No Dye control; B) and with labelled MII-ffEVs (ffEVs+Dye, C, D, E). The insets in the top panel of oocyte images (A, B, C, D, E) are magnified in the bottom panel (F, G, H, I, J). The bottom panel was acquired during confocal imaging using a higher exposure on the green channel to allow for clear visualisation of the transzonal projections, which are structures in the ZP that function to maintain bidirectional communication between oocytes and cumulus cells. ZP: zona pellucida, Nc: nuclei, Cy: cytoplasm. (K) GV oocytes were cultured in a time-lapse Geri incubator for 48 hours with or without ffEVs originating from mature follicles. At the end of the culture, GV oocytes either arrested at the immature state progressed to an intermediate state of maturity (GVBD) or completely matured (MII). Mean ± SD of oocyte maturation rates (M) and polar body extrusion time (L) in ffEVs treated and control groups after three independent experiments. An asterisk (*) denotes significant differences (P<0.05) in the data.

### Incubation of GV oocytes with MII-ffEVs during rIVM enhances maturation rates

Following the confirmation that immature GV oocytes internalized ffEVs, we quantified the impact of this internalisation by examining the oocyte maturation rate. We cultured GV oocytes in a time-lapse incubator for 48 h with or without ffEVs. Using time-lapse imaging, we registered the timing of the polar body extrusion and the number of mature oocytes (MII) at the end of the culture in each group (Figure 2K). The maturation of each oocyte was confirmed under bright light microscopy before snap-freezing for single-oocyte proteomics or fixation for TEM. In three independent experiments, we found that the supplementation of culture media with ffEVs significantly increased the oocyte maturation rate by an average of 22.8±9.4% (p=0.037; Figure 2L). Specifically, treated ffEVs oocytes displayed 77.7±5.4% (average± SEM) maturation compared to 54.0±7.7% of controls. The timing of polar body extrusion was comparable between the two groups (22.9±0.8 and 22.8±1.4 h for ffEVs- treated oocytes and controls, respectively; Figure 2M).

### Treatment with MII-ffEVs changes the proteome of oocytes

Mature oocytes cultured with or without ffEVs were subjected to single oocyte proteomics. A total of 4587 proteins were identified across all samples (Supplementary Table 7). For the downstream analysis, these proteins were filtered, and only proteins present in at least 4 out of 5 control or treated oocyte samples were selected, resulting in 3884 filtered proteins (Supplementary Table 8). A PCA based on the top 50 proteins with the greatest changes across the dataset showed that samples were distributed in 2 groups (Figure 3A). Statistical analysis revealed 56 DAPs (FDR<0.1, Figure 3B; Supplementary Table 9). The highest difference was found in Hyaluronan Synthase 1 (HAS1) with an increased fold-change of 6.6 in ffEVs-treated oocytes. These DAPs were further analysed for their expression profiles across samples using SOTA clustering analysis, which provided two clear clusters (Figure 3C). Cluster 1, with 15 proteins, showed increased abundance in control mature oocytes, while cluster 2, with 44 proteins, represented increased abundance in ffEVs-treated mature oocytes.

**Fig 3.**
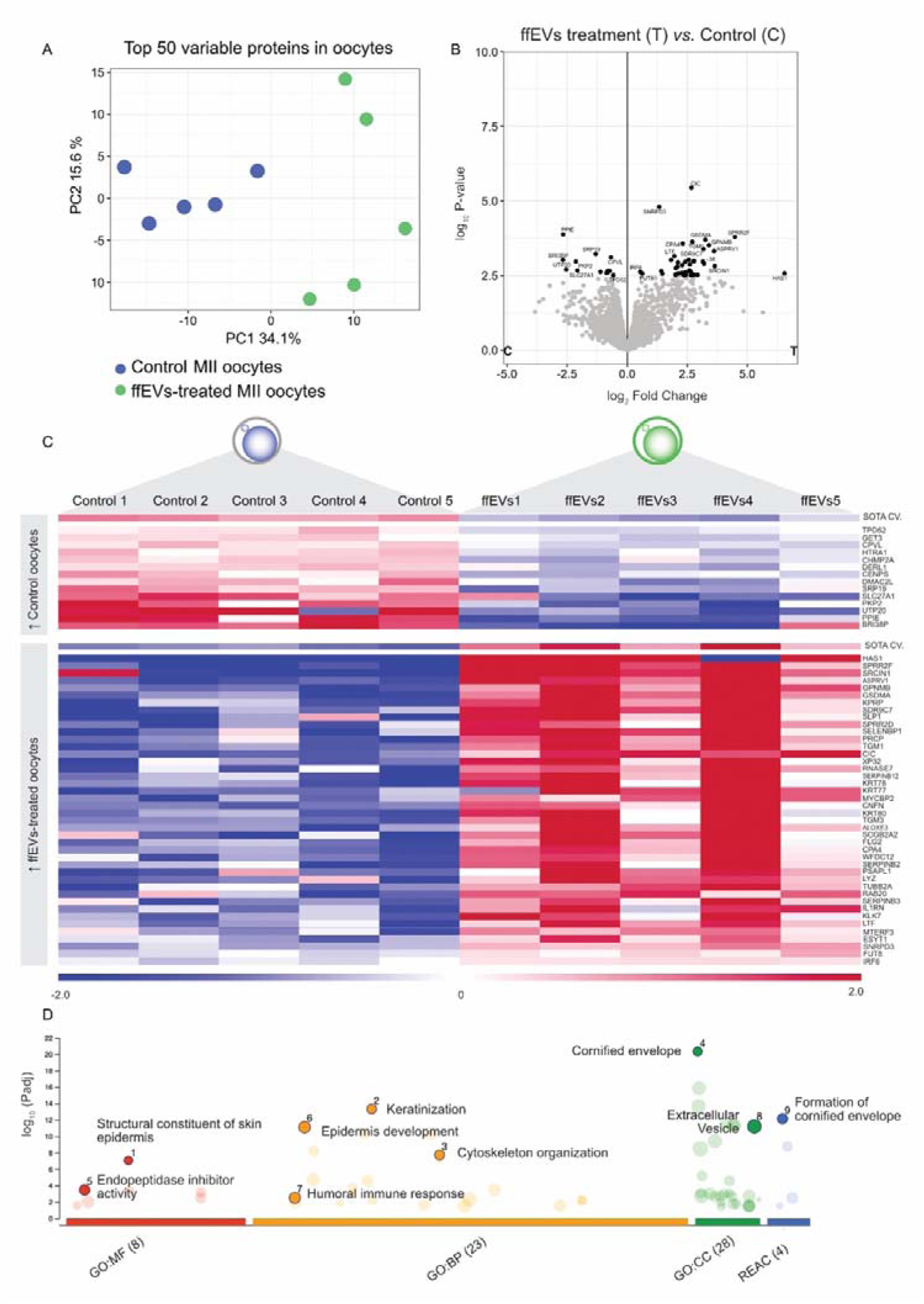
The ffEVs treatment modulated the oocyte proteome after maturation. (A) Principal component analysis (PCA) based on the top 50 proteins with the greatest change between all experimental groups (ffEVs-treated oocytes n=5, control oocytes n=5). (B) The volcano plot illustrates significantly differentially abundant proteins (DAPs) (FDR<0.1). The −log_10_ P-value is plotted against the log_2_ fold change: ffEVs-treated/control. (C) Heatmap showing DAPs in ffEVs (FDR<0.1). Self-organizing tree algorithm (SOTA) analysis identified two clusters of proteins with similar expression profiles. (D) GOSt multi-query plot of Manhattan showing gene ontology (GO) overrepresentation analysis results of 76 unique proteins in MII-ffEVs. BP: biological process; CC: cellular component; MF: molecular function; REAC: reactome.

With a cut-off of FDR<0.2, 93 DAPs emerged from the latter comparison (Supplementary Table 10). These DAPs were used for the GO overexpression analysis using g:Profiler functional enrichment and were mainly related to pathways involving keratinization, cytoskeleton organization, endopeptidase inhibitor activity, humoral immune response, and extracellular vesicle (Figure 3D; Supplementary Table 11).

To explore how the DAPs identified in ffEVs and among mature oocytes might be connected, we created a protein network using STRING and visualised possible interactions between proteins originating from ffEVs or oocytes (Figure 4). We observed two noteworthy clusters, one exclusively composed of proteins originating from the oocyte DAPs (cluster 1, green) and another composed of proteins originating from both ffEVs and oocytes (cluster 2, pink). Functional enrichment analysis revealed that cluster 1 was related to keratin and intermediate filament organization, while cluster 2 was related to steroid and lipid metabolism and extracellular vesicle transport.

**Fig 4.**
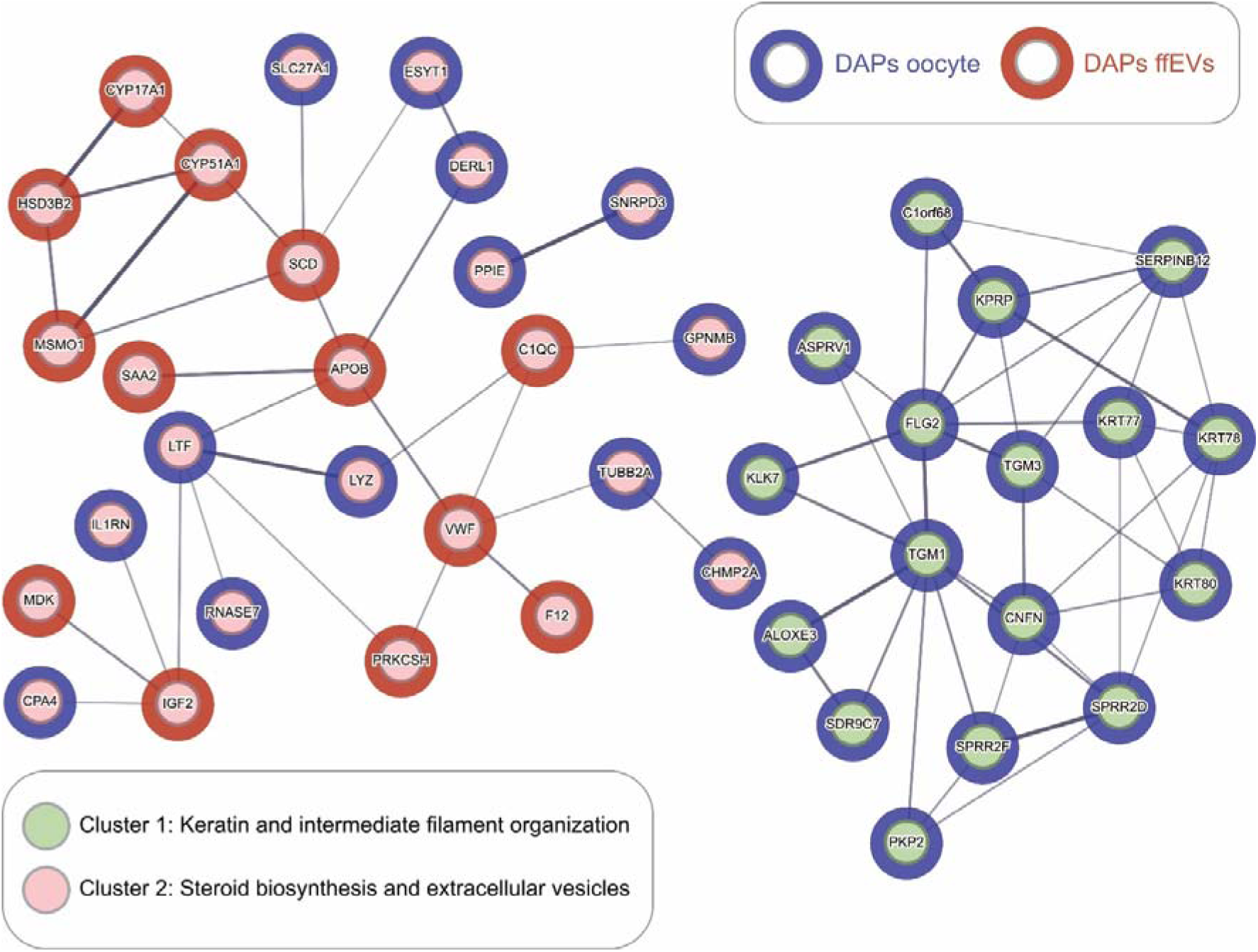
Oocyte and ffEVs DAPs communication network in rIVM. DAPs identified as specific to mature or immature ffEVs and DAPs identified in oocytes following rIVM with ffEVs were subjected to a pooled analysis to reveal possible interactions between proteins in the oocytes and proteins in ffEVs. Proteins are shown as nodes connected by lines that signify interacted, and line thickness indicates the strength of the data support. Blue nodes represent oocyte DAPs, and red nodes represent MII-ffEV DAPs. Two clusters can be observed (cluster 1- green, and cluster 2- pink). The green cluster is only composed of oocyte- originating proteins relevant to keratin and intermediate filament organization, while the pink cluster contains proteins from both the oocyte and the ffEVs relevant to steroid synthesis, lipid metabolism, and extracellular vesicles.

### Ultrastructure of mature oocytes is modified by MII-ffEVs treatment

TEM ultrastructural analysis of single oocytes revealed differences in oocyte organelle distribution and appearance between ffEVs-treated (Figure 5A,C) and control mature oocytes (Figure 5B,D). In particular, we observed differences in the appearance of smooth endoplasmic reticulum (SER) elements associated with mitochondria, called M-SER aggregates, and vesicles of SER and mitochondria, named MV complexes. While M-SER aggregates were more abundant in ffEVs-treated mature oocytes (Figure 5F), the MV complexes at various sizes were more abundant in control mature oocytes (Figure 5G). Treated oocytes had a more loose, microfilamentous microvilli architecture (Figure 5E), while in control oocytes, microvilli appeared slightly shorter in length, lesser in number, and more irregularly distributed (Figure 5H).

**Fig 5.**
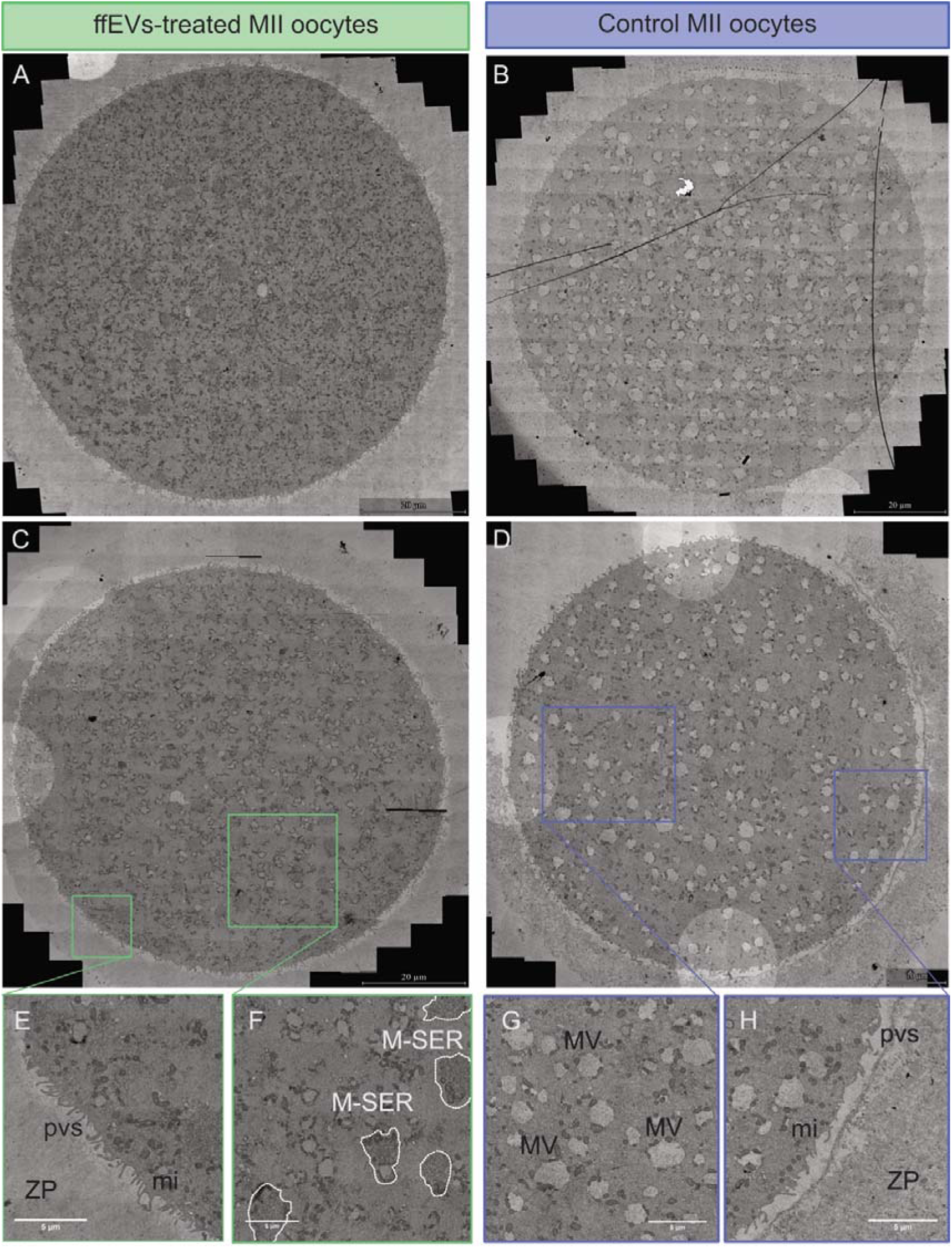
Cytoplasmic reorganization of oocytes treated or not with MII-ffEVs during *in vitro* maturation. TEM images of oocytes. Scale bars A-C: 20 µm, D: 10 µm and E-H: 5µm. M-SER: mitochondria smooth endoplasmic reticulum (highlighted by white dotted line); MV: mitochondria vesicle; pvs: perivitelline space; mi: microvilli; ZP: zona pellucida.

## Discussion

For the first time, we demonstrate here that EVs originating from human ovarian follicles can support oocyte maturation. Our findings can be delineated into three congruent trajectories. First, we have identified the proteins present in ffEVs originating from follicles containing immature and mature oocytes, which provide promising targets for the development of supplements aimed at enhancing global IVM solutions. Second, our data suggest that these ffEVs proteins may be functionally significant, as evidenced by the internalisation of ffEVs by immature oocytes and a concomitant increase in oocyte maturation rates enhanced by 20%. Third, the interaction between the follicular milieu and oocytes during maturation, mediated by ffEVs, appears to induce substantial changes in the maturing oocytes. These changes are reflected in both, the proteomic profile and ultrastructural alterations, with possible implications for oocyte competence. Below, we elaborate on these aspects and demonstrate how they collectively support the conclusion that a straightforward methodology—incorporating ffEVs, which can be readily isolated from the patient’s own sample on the day of oocyte retrieval for IVM —may be integrated into any IVM protocol. This approach has the potential to enhance the quality and quantity of oocytes available for treatment, thereby significantly increasing the likelihood of successful IVF, foremostly in the most hormone-vulnerable patients.

### The proteomic landscape of ffEVs derived from mature and immature follicles

All cells have the capability to secrete EVs. Studies in cattle have demonstrated that the proteomic cargo of ffEVs most likely originates from diverse cells in the ovarian microenvironment, including granulosa cells, theca cells, cumulus cells, next to the oocyte itself (Uzbekova *et al*., 2020a). In addition, ffEVs may originate from the blood plasma, as follicular fluid is formed in part via the transudation of plasma components. The role of ffEVs in the bidirectional oocyte-cumulus cell communication during follicular development is established as complementary to the traditional pathways mediated by transzonal projections (TZPs) and gap junctions (Machtinger *et al*., 2016). Our comprehensive proteomic profiling of ffEVs originating from follicles containing mature and immature oocytes revealed an intriguing protein signature relevant to follicular homeostasis, encompassing essential protein machinery involved in steroid biosynthesis, endoplasmic reticulum organization, oxidoreductase activity, and extracellular vesicle transport.

Despite the similar proteomic landscape between GV- and MII-ffEVs, we could highlight four proteins exclusively abundant in the MII-ffEVs cargo (F12, C1QC, IGKV1-39, and FREM-2). The roles of these proteins in the context of follicular biology suggest a link to oocyte competence. Particularly, the F12 protein has been found in lower abundance in follicular fluid of obese women, a condition linked to a low oocyte maturation rate (Liu *et al*., 2020). Reduced abundance of C1QC protein was previously reported in the follicular fluid of women with polycystic ovary syndrome, a disorder known to experience compromised oocyte quality (Pla *et al*., 2022). Similarly, reduced transcription of C1QC was noted in granulosa cells isolated from the follicular fluid of follicles containing oocytes that led to poor embryo development (Yu *et al*., 2023). A role for the complement pathway is yet to be established in the context of follicular competence; however, several seminal and recent studies point to its criticality in the facilitation of fertilisation, oocyte safeguarding, and embryo potential (Anderson *et al*., 1993; Reichhardt *et al*., 2019; Wang *et al*., 2022). This theory might also explain the higher availability of IGKV1-39 in ffEVs from MII-oocytes, as immunoglobulins are known activators of the complement components, while their inhibition in immature oocytes has been shown to block oocyte maturation (Wang *et al*., 2022). As IGKV1-39 is predominantly transferred via EVs, it is possible that the transcriptomically inactive immature oocytes receive components for immunoglobulin synthesis via ffEVs. With regards to FREM2, its transcript was found upregulated in human granulosa cells isolated from follicular fluid collected in the peri-ovulation window (Poulsen *et al*., 2020). FREM2 protein was additionally more abundant in mature feline oocytes (Turathum *et al*., 2018), while the critical role of FREM2 availability during early embryo development is also well established (Zhou *et al*., 2021). Thus, it is possible that the abundant availability of FREM2 in ffEVs coming from mature follicles in our study is a surrogate marker of oocyte health and potential. On the other side, proteins enriched in GV-ffEVs are associated with steroid biosynthesis and lipid metabolism, including CYP17A1 and CYP51A1. These proteins play a role in the activation of follicular fluid meiosis-activating sterol (FF-MAS), which promotes meiotic resumption when added to an IVM culture (Guo *et al*., 2020a). It is not unlikely that these proteins are introduced to the oocyte via ffEVs to facilitate GV oocyte maturation. Altogether, we pinpoint the proteins identified by our proteomic analysis in the GV- and MII-ffEVs for further investigation as mediators of follicle development, oocyte maturation, and competence.

### Supplementation of rescue IVM culture with ffEVs increases oocyte maturation rate

The biological activity of EVs relies on the capacity of target cells to internalize them. It is proposed that EVs enter cells and release their cargo via endocytosis, phagocytosis, micropinocytosis, or direct fusion with the plasma membrane (Mulcahy *et al*., 2014). EVs uptake studies in oocytes in different animal species have reported the presence of stained ffEVs in the cumulus cells (Hung *et al*., 2015; de Almeida Monteiro Melo Ferraz *et al*., 2020b) and in the interface of oocytes with the cumulus cells, in the zona pellucida, and the ooplasm (da Silveira *et al*., 2017a; Lange-Consiglio *et al*., 2017a; Uzbekova *et al*., 2020b; Gabryś *et al*., 2022a). We demonstrated for the first time that cumulus cells denuded human oocytes can uptake labelled ffEVs, which can be detected in the zona pellucida after 24 hours of co-incubation. Similar studies have been conducted in multiple animal models using COCs in the context of standard IVM. Two confocal studies in cat and cattle detected labelled ffEVs in the cumulus cells but not in the oocytes after 16-18 hours of co-incubation with COCs (Hung *et al*., 2015; de Almeida Monteiro Melo Ferraz *et al*., 2020c). However, in five other studies conducted using mice, canine, mare, and cattle COCs, positive ffEVs staining was detected in the interface of oocytes with the cumulus cells, in the zona pellucida, and the ooplasm after periods of incubation raging from 15 to 72 hours (da Silveira *et al*., 2017b; Lange-Consiglio *et al*., 2017b; Uzbekova *et al*., 2020a; Gabryś *et al*., 2022b; Fiorentino *et al*., 2024). Our confocal microscopy findings locating ffEVs in the ZP are intriguing in this regard, as they raise the possibility of ffEVs being transferred into the oocyte via the cytoplasmic TZPs. TZPs are thin cytoplasmic extensions originating from cumulus cells that traverse the 3–5 micron-thick zona pellucida to form gap-junctions and adherens-junctions with the oocyte, essential for communication between the oocyte and cumulus cells. Electron microscopy studies have detected EVs between the tip of the TZP and the oocyte plasma membrane (Macaulay *et al*., 2014; Marchais *et al*., 2022), suggesting a potential mechanism for transferring large cargo from granulosa cells to the oocyte. However, we have utilized denuded oocytes, which lack cumulus cells, and thus far, it is unknown whether TZPs’ tracks remain in the zona pellucida and could attract supplemented ffEVs. Our findings confirm that ffEVs likely transfer EVs cargo from the ovarian follicular fluid to the interior of the oocyte, but the precise mechanism mediating this internalisation and the relevance of this transfer of molecular information remains unknown.

Performing rIVM in the presence of ffEVs resulted in over 20% higher maturation rate, which is in line with a similar study in mares, whereby the supplementation of standard IVM culture with ffEVs resulted in a significant 25% increase in oocyte maturation rate (Gabryś *et al*., 2022c). A plethora of animal studies, but never human so far, have demonstrated the beneficial role of ffEV supplementation to standard IVM culture in terms of oocyte maturation and competence to produce good quality blastocysts (da Silveira *et al*., 2017a; Rodrigues *et al*., 2019; de Almeida Monteiro Melo Ferraz *et al*., 2020b; Gabryś *et al*., 2022c; Han *et al*., 2024). Currently, the mechanisms how ffEVs might contribute to improved oocyte maturation and competence in a standard IVM system is still not understood. However, it has been hypothesized that ffEVs might initially inhibit meiosis resumption in a similar way to the CNP peptide in the CAPA-IVM system (Pioltine *et al*., 2020b). However, the mechanism of ffEVs might be different from the one of the CNP peptide; the CNP-NPR2 modulates the cGMP levels, while the cargo of ffEVs might modulate the cAMP levels in the oocyte. More recently, a study suggested that ffEVs enhance IVM outcomes via the ERK1/2 pathway, which in turn results in an increase in the levels of ER and a decrease in ROS in the porcine oocytes (Han *et al*., 2024).

### The impact of the interaction of oocytes with ffEVs during oocyte maturation

For a deeper insight into how ffEVs contribute to improve oocyte maturation rates, we analysed oocytes subjected to rIVM with ffEVs using single-cell proteomics. Our methodology identified 4589 proteins present in the individual oocytes. The patient’s intrinsic variability and the prolonged *in vitro* culture might have hidden some significant differences between ffEVs treated and control oocytes. Nevertheless, we identified differential abundant proteins between ffEVs treated and control oocytes associated with keratinization, cytoskeleton organization, endopeptidase inhibitor activity, humoral immune responses, and, expectedly, extracellular vesicle transport. The organization of keratin and the cytoskeleton is critically important during oocyte maturation, with distinct differences in keratin distribution patterns observed between immature and mature oocytes (Kabashima *et al*., 2010). Moreover, the intermediate filaments formed by keratins in oocytes play a significant role in early embryonic divisions and development through a mechanism in which differential keratin regulation impacts cell lineage fate (Lim *et al*., 2020). Indeed, we found a plethora of keratins increased in oocytes during ffEVs-facilitated rIVM, including keratin 80 (KRT80). Interestingly, the reduction of KRT80 has been identified as a marker of poor oocyte competence, specifically aneuploidy, with sufficient levels of KRT80 potentially playing a crucial role in the segregation of oocyte chromosomes (Fragouli *et al*., 2010). We also identified keratins in ffEVs cargo. It is possible that ffEVs deposited some of their protein cargo to the oocytes upon internalisation, which could have contributed to the maturation processes and competence establishment. Amongst the proteins exhibiting higher expression in MII oocytes after ffEV-rIVM and also detected in ffEVs were skin-specific protein 32 (XP32), filaggrin-2 (FLG2), protein-glutamine gamma-glutamyltransferase E (TGM3). The literature identifies these three proteins as key maternal proteins participating in healthy embryo development (Iqbal *et al*., 2014; Foresta *et al*., 2016; Virant-Klun *et al*., 2016; Zha *et al*., 2023).

It is well established that EVs have the capacity to indirectly decrease the synthesis of proteins by decreasing their corresponding transcripts through their EV-miRNA cargo. It has been reported that the expression of most of the proteins with lower abundance in our dataset is regulated by miRNAs, of which miR-223-3p, miR-148a-3p, hsa-let-7a-5p, and hsa-let-7b- 5p have been found in the ffEVs in our study. Other studies have shown that injecting specific miRNAs in oocytes can exert changes in mRNA translation and, hence, impact protein availability (Kataruka *et al*., 2020). Notably, the protein with the largest drop in abundance after supplementation with ffEVs, BRI3-binding protein (BRI3BP), which is localized to the mitochondria and the ER, is downregulated to avoid developmental arrest at the morula stage (Nõmm *et al*., 2023).

Of particular interest is the presence of hyaluronan synthase 1 (HAS1), the most differentially abundant protein in MII oocytes undergoing ffEV-facilitated rIVM, intriguingly absent in the ffEVs themselves indicating no direct transfer from ffEVs. There is a substantial amount of literature on the role of hyaluronan synthesis during oocyte maturation and competence establishment (Fülöp *et al*., 1997; Kimura *et al*., 2002; Stock *et al*., 2002; Schoenfelder and Einspanier, 2003; McKenzie *et al*., 2004; Siiskonen *et al*., 2015). Yokoo M and Sato reported that hyaluronan synthesis and accumulation may be involved in the expansion of COC’s volume and play an important role in the progression of meiotic resumption and the induction of oocyte maturation (Yokoo and Sato, 2011). In addition, HAS1 has been associated with the endoplasmic reticulum (Siiskonen *et al*., 2015), explaining a hypothetical connection between the increased abundance of HAS1 and increased endoplasmic reticulum structures in ffEV-treated oocytes, which calls for further investigations.

To translate our findings into hypotheses elucidating the mechanisms of ffEV-facilitated rIVM, we explored the proteomic landscape of the ffEV-oocyte communication network, identifying potential candidates that merit further investigation. Hence, we observed two intriguing clusters, of which one is solely relevant to oocyte activity and another integrates the actions of both oocytes and ffEVs. It is tempting to hypothesize that ffEVs deliver cargo involved in steroid and lipid metabolism to GV oocytes, facilitating meiotic resumption via modulation of FF-MAS, which promotes meiotic resumption when added to IVM medium (Guo *et al*., 2020b). Concurrently, oocytes may fine-tune their keratin distribution and intermediate filament organization to prepare for maturation in an environment conducive to fertilisation. We highlight specific proteins that warrant exploration in mechanistic studies, particularly CYP17A1, HSD3B2, CYP51A1, MSMO1, SCD, SLC27A1, ESYT1, and APOB as involved proteins in interactions between the oocyte and ffEVs in steroid synthesis and lipid metabolism.

### Ultrastructure of oocytes after ffEV-rIVM

It was intriguing to observe such a striking difference in the ratio of MV/M-SER between the two groups of mature oocytes. M-SER is an ultrastructure exclusively observed in mature oocytes, and a moderate number of M-SER with MVs in the ooplasm is a common feature of healthy mature oocytes (Palmerini *et al*., 2014). The higher abundance of MVs - larger or dilated SER vesicles in the ooplasm – in oocytes not subjected to ffEV supplementation could be a sign of ER stress (Miglietta *et al*., 2023). Indeed, dilatation and vacuolization of ER cisterns were a result of the treatment of oocytes with an inductor of ER stress (Vašíčková *et al*., 2018). In contrast, the presence of numerous and large ooplasmic MV complexes in human mature oocytes, caused by swelling and coalescence of isolated SER vesicles, were stipulated to indicate cytoskeletal defects (Gualtieri *et al*., 2009). Dilatation and vesiculation of ER, associated with the uncoupling/loss of associated mitochondria, have also been observed in human-aged oocytes and correlated to oxidative stress (Oslowski and Urano, 2011; Bianchi *et al*., 2015). Similar to our study, Bianchi and colleagues additionally observed fewer M-SER aggregates and increased swelling of SER tubules in aged oocytes, but also associated with long *in vitro* culture (48 h) or the components of the *in vitro* milieu (Bianchi *et al*., 2015). The TEM observations regarding altered SER structures are in line with oocyte proteomic data pointing to several DAPs related to ER stress (BRI3BP, CANX, HSP90B1, PDIA4, PDIA6, PDIA3, HSPA5, ERLIN2, ERP44, UGGT1, ERMP1, TMX1). Overrepresentation of MVs in MII oocytes from rIVM without ffEVs could compromise the probability of fertilisation since SER plays a key role in calcium storage and release at oocyte activation during fertilisation (Whitaker, 2006). Moreover, changes in complexes of SER and associated mitochondria could induce a major impact on the oocyte in terms of energy accumulation, protein and lipid production, and production of nuclear membranes throughout early embryo development (Carroll, 1996). Besides the MV-M-SER ratio, we observed differences in the zona pellucida and microvilli morphology between the two groups of mature oocytes. While ffEV-supplemented oocytes had a loose, microfilamentous architecture, controls showed a more dense and granular texture. Regarding the microvilli pattern, literature findings imply that mature oocytes with normal morphology have numerous long and thin microvilli extending into the perivitelline space, an observation in line with the microvilli we noted in ffEVs-supplemented oocytes (Palmerini *et al*., 2014). These differences in microvilli morphology might affect the fertilisation a posteriori since microvilli are suggested to play a role during fusion between the oocyte and the sperm (Runge *et al*., 2007). A deep understanding of these ultrastructural changes will inform and ensure the biosafety and clinical utility of oocytes following r-IVM with ffEVs.

Our study is subject to certain limitations that may have introduced interpretational biases. Firstly, the oocytes were initially vitrified at the GV stage and subsequently warmed for experimentation, implying that potential cryodamage cannot be ruled out for both treated and untreated oocytes. Some ultrastructural observations may be attributable to the vitrification and warming procedures, as noted by Palmerini et al. (Palmerini *et al*., 2014). Additionally, we performed uptake assays using a system that employs a green fluorescent dye to indicate ffEV uptake by oocytes. However, the ooplasm and certain oocyte features, such as refractile bodies, tend to display autofluorescence, introducing an unspecific signal that is nevertheless distinguishable from the specific signal of ffEV uptake (Otsuki *et al*., 2007). Although proteomics and TEM studies provided insight into the potential competence of mature oocytes from rIVM, the ultimate test of competence—assessing fertilisation rate through insemination with ICSI—was not feasible due to legal restrictions. Finally, we invite other researchers to test our proposed ffEV-facilitated IVM system in alternative models, such as CAPA-IVM and standard IVM using unstimulated cumulus-oocyte complexes (COCs), rather than denuded GVs from stimulated cycles, which may have inherent developmental limitations.

In conclusion, we propose a novel approach adaptable to global IVM systems, where ffEVs should be further investigated as an adjunct to enhance oocyte maturation rates and potentially improve oocyte competence. This possible competence enhancement requires further validation through parthenogenic activation or insemination of oocytes from IVM. Our findings hold the promise of directly benefiting patients seeking reproductive medicine. Once optimized for clinical application, IVM could offer significant advantages to subfertile women at risk of OHSS, cancer patients undergoing fertility preservation, and the broader population undergoing IVF. This advancement could facilitate a more streamlined, cost- efficient, and safer approach to infertility treatment, ultimately contributing to heightening of women’s health.

## Authors’ roles

All authors significantly contributed to the study. SM and MDS conceptualized the study, implemented the experiments and authored the manuscript. CA, MX, AV, SB, MS and NC were involved in the implementation of the experiments. SB was involved in the analyses of the data. SU and BL were involved in the authoring of the manuscript and the inception of the study.

## Acknowledgements

This work was supported by an EMDO-Stiftung and a FAN grant from the University of Zurich. The authors are grateful to Dr. Miriam Susanna Lucas-Droste of the Scientific Center for Optical and Electron Microscopy at ETH Zurich (8093 Zurich, Switzerland) for her support with the electron microscopy imaging.

## Supplemental Data

**Supplementary Figure 1.**
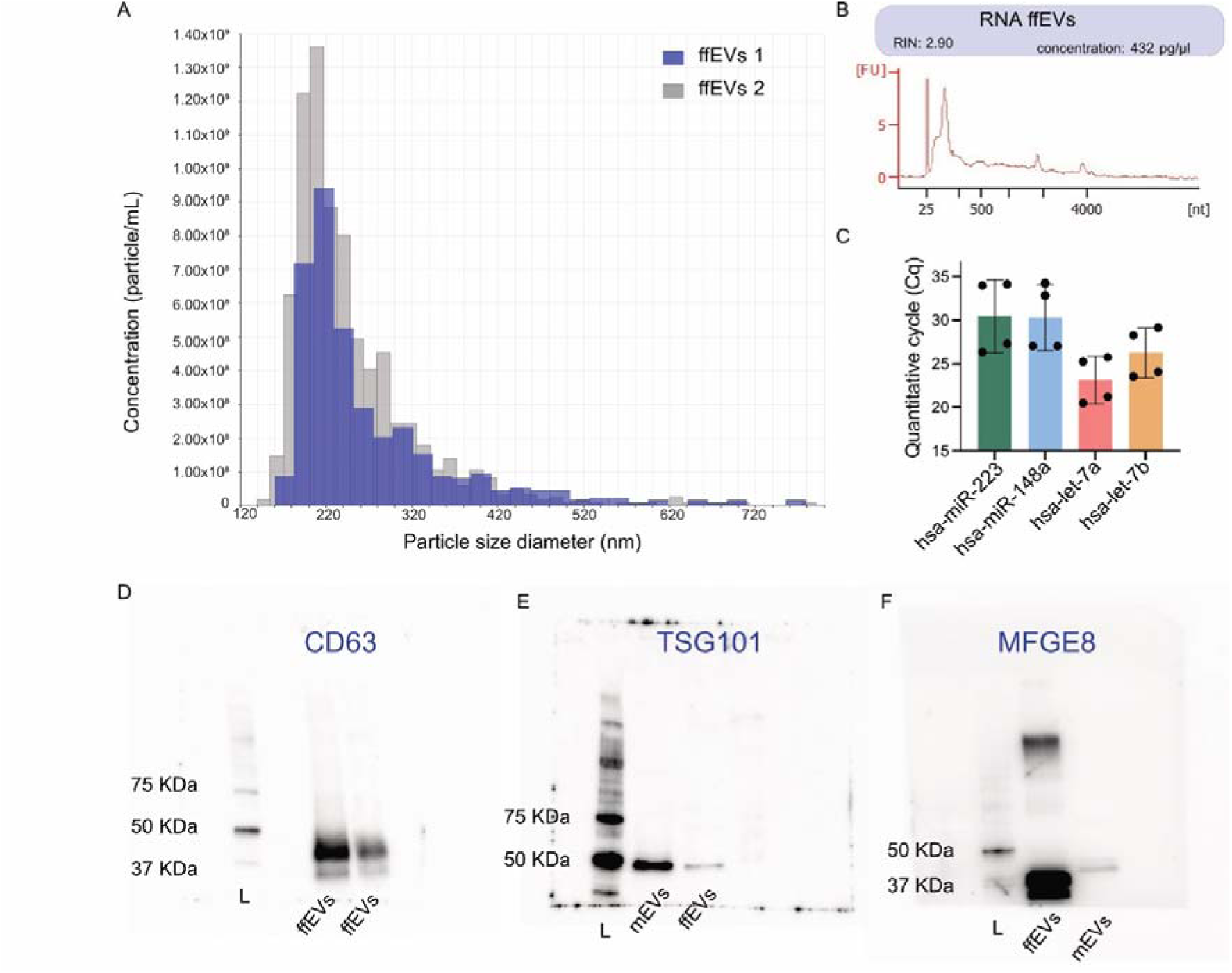
Size distribution evaluation unveiled particle dimensions ranging from 50 to 700 nm. (**A**). We successfully extracted measurable quantities of RNA from ffEVs derived from 1.5 – 2.0 mL of patient-individual follicular fluid. The Bioanalyzer profiles demonstrated that the majority of the RNA belonged to the small RNA population, with a minimal contribution from larger RNAs (**B**). Furthermore, within the isolated ffEVs, we identified the presence of typical follicular fluid miRNAs (**C**). Western blot analysis of the ffEVs samples confirmed their identity by detecting specific protein markers associated with EVs (**D-F**). Specifically, the transmembrane proteins CD63 and the cytosolic protein Tumor susceptibility gene 101 (TSG101) commonly enriched in EVs were present in our samples. L: Ladder. mEV: milk EVs.

## References

1. Almeida Monteiro Melo Ferraz M de, Fujihara M, Nagashima JB, Noonan MJ, Inoue-Murayama M, Songsasen N. Follicular extracellular vesicles enhance meiotic resumption of domestic cat vitrified oocytes. Scientific Reports 2020a;10:.

2. Almeida Monteiro Melo Ferraz M de, Fujihara M, Nagashima JB, Noonan MJ, Inoue- Murayama M, Songsasen N. Follicular extracellular vesicles enhance meiotic resumption of domestic cat vitrified oocytes. Scientific Reports 2020b;10:.

3. Almeida Monteiro Melo Ferraz M de, Fujihara M, Nagashima JB, Noonan MJ, Inoue- Murayama M, Songsasen N. Follicular extracellular vesicles enhance meiotic resumption of domestic cat vitrified oocytes. Scientific Reports 2020c;10:. Nature Research.

4. Anderson DJ, Abbott AF, Jack RM. The role of complement component C3b and its receptors in sperm-oocyte interaction. Proceedings of the National Academy of Sciences of the United States of America 1993;90:.

5. Bhattacharjee N V., Schumacher AE, Aali A, Abate YH, Abbasgholizadeh R, Abbasian M, Abbasi-Kangevari M, Abbastabar H, Abd ElHafeez S, Abd-Elsalam S, et al. Global fertility in 204 countries and territories, 1950–2021, with forecasts to 2100: a comprehensive demographic analysis for the Global Burden of Disease Study 2021. The Lancet 2024;403:.

6. Bianchi S, Macchiarelli G, Micara G, Linari A, Boninsegna C, Aragona C, Rossi G, Cecconi S, Nottola SA. Ultrastructural markers of quality are impaired in human metaphase II aged oocytes: a comparison between reproductive and in vitro aging. Journal of Assisted Reproduction and Genetics 2015;32:1343–1358. Springer New York LLC.

7. Carroll J. Development of oocyte banks and systems for the in-vitro development of oocytes: Future directions for the treatment of infertility. Human Reproduction 1996;11:159–168. Oxford University Press.

8. Deerinck TJ, Bushong E a., Thor a., Ellisman MH. NCMIR methods for 3D EM: A new protocol for preparation of biological specimens for serial block face scanning electron microscopy. Microscopy 2010;

9. Demichev V, Messner CB, Vernardis SI, Lilley KS, Ralser M. DIA-NN: neural networks and interference correction enable deep proteome coverage in high throughput. Nature Methods 2020;17:.

10. Demichev V, Szyrwiel L, Yu F, Teo GC, Rosenberger G, Niewienda A, Ludwig D, Decker J, Kaspar-Schoenefeld S, Lilley KS, et al. dia-PASEF data analysis using FragPipe and DIA-NN for deep proteomics of low sample amounts. Nature Communications 2022;13:.

11. Edwards RG. Maturation in vitro of mouse, sheep, cow, pig, rhesus monkey and human ovarian oocytes. Nature 1965;208:.

12. Esbert M, García C, Cutts G, Lara-Molina E, Garrido N, Ballestros A, Scott RT, Seli E, Wells D. Oocyte rescue in-vitro maturation does not adversely affect chromosome segregation during the first meiotic division. Reproductive BioMedicine Online 2024;48:.

13. Fiorentino G, Merico V, Zanoni M, Comincini S, Sproviero D, Garofalo M, Gagliardi S, Cereda C, Lin CJ, Innocenti F, et al. Extracellular vesicles secreted by cumulus cells contain microRNAs that are potential regulatory factors of mouse oocyte developmental competence. Molecular Human Reproduction 2024;30:1–16.

14. Foresta C, Ubaldi FM, Rienzi L, Franchin C, Pivato M, Romano S, Guidolin D, Caro R De, Ferlin A, Toni L De. Early protein profile of human embryonic secretome. Frontiers in Bioscience - Landmark 2016;21:620–634. Frontiers in Bioscience.

15. Fragouli E, Bianchi V, Patrizio P, Obradors A, Huang Z, Borini A, Delhanty JDA, Wells D. Transcriptomic profiling of human oocytes: Association of meiotic aneuploidy and altered oocyte gene expression. Mol Hum Reprod [Internet] 2010;16:570–582.

16. Fülöp C, Salustri A, Hascall VC. Coding sequence of a hyaluronan synthase homologue expressed during expansion of the mouse cumulus-oocyte complex. Archives of Biochemistry and Biophysics 1997;337:261–266. Academic Press Inc.

17. Gabryś J, Kij-Mitka B, Sawicki S, Kochan J, Nowak A, Łojko J, Karnas E, Bugno- Poniewierska M. Extracellular vesicles from follicular fluid may improve the nuclear maturation rate of in vitro matured mare oocytes. Theriogenology 2022a;188:116–124. Elsevier Inc.

18. Gabryś J, Kij-Mitka B, Sawicki S, Kochan J, Nowak A, Łojko J, Karnas E, Bugno- Poniewierska M. Extracellular vesicles from follicular fluid may improve the nuclear maturation rate of in vitro matured mare oocytes. Theriogenology 2022b;188:.

19. Gabryś J, Kij-Mitka B, Sawicki S, Kochan J, Nowak A, Łojko J, Karnas E, Bugno- Poniewierska M. Extracellular vesicles from follicular fluid may improve the nuclear maturation rate of in vitro matured mare oocytes. Theriogenology 2022c;188:116–124. Elsevier Inc.

20. Gilchrist RB, Ho TM, Vos M De, Sanchez F, Romero S, Ledger WL, Anckaert E, Vuong LN, Smitz J. A fresh start for IVM: capacitating the oocyte for development using pre-IVM. Human Reproduction Update 2024;30:.

21. Gualtieri R, Iaccarino M, Mollo V, Prisco M, Iaccarino S, Talevi R. Slow cooling of human oocytes: ultrastructural injuries and apoptotic status. Fertility and Sterility 2009;91:1023–1034.

22. Guo R, Wang X, Li Q, Sun X, Zhang J, Hao R. Follicular fluid meiosis-activating sterol (FF- MAS) promotes meiotic resumption via the MAPK pathway in porcine oocytes. Theriogenology 2020a;148:.

23. Guo R, Wang X, Li Q, Sun X, Zhang J, Hao R. Follicular fluid meiosis-activating sterol (FF- MAS) promotes meiotic resumption via the MAPK pathway in porcine oocytes. Theriogenology 2020b;148:.

24. Guo W, Zheng X, Zheng D, Yang Z, Yang S, Yang R, Li R, Qiao J. Effectiveness, Flexibility and Safety of Switching IVF to IVM as a Rescue Strategy in Unexpected Poor Ovarian Response for PCOS Infertility Patients. Journal of Clinical Medicine 2023;12:.

25. Han Y, Zhang J, Liang W, Lv Y, Luo X, Li C, Qu X, Zhang Y, Gu W, Chen X, et al. Follicular fluid exosome-derived miR-339-5p enhances in vitro maturation of porcine oocytes via targeting SFPQ, a regulator of the ERK1/2 pathway. Theriogenology 2024; Elsevier BV.

26. Harris E. Infertility Affects 1 in 6 People Globally. JAMA 2023;329:.

27. Ho VN, Mol BW, Vuong LN. Reply: IVM for expected high responders: Balancing the effectiveness and safety. Human Reproduction 2019;34:.

28. Howe EA, Sinha R, Schlauch D, Quackenbush J. RNA-Seq analysis in MeV. Bioinformatics 2011;27:.

29. Hung WT, Hong X, Christenson LK, McGinnis LK. Extracellular vesicles from bovine follicular fluid support cumulus expansion. Biology of Reproduction 2015;93:. Society for the Study of Reproduction.

30. In vitro maturation: a committee opinion Fertility and Sterility 2021;115:.

31. Iqbal K, Chitwood JL, Meyers-Brown GA, Roser JF, Ross PJ. RNA-seq transcriptome profiling of equine inner cell mass and trophectoderm. Biology of Reproduction 2014;90:. Society for the Study of Reproduction.

32. Kabashima K, Matsuzaki M, Suzuki H. Intermediate filament keratin dynamics during oocyte maturation requires maturation/M-phase promoting factor and mitogen-activated protein kinase kinase activities in the hamster. Reproduction in Domestic Animals 2010;45:.

33. Kataruka S, Modrak M, Kinterova V, Malik R, Zeitler DM, Horvat F, Kanka J, Meister G, Svoboda P. MicroRNA dilution during oocyte growth disables the microRNA pathway in mammalian oocytes. Nucleic Acids Research 2020;48:8050–8062. Oxford University Press.

34. Kimura N, Konno Y, Miyoshi K, Matsumoto H, Sato E. Expression of hyaluronan synthases and CD44 messenger RNAs in porcine cumulus-oocyte complexes during in vitro maturation. Biology of Reproduction 2002;66:707–717. Society for the Study of Reproduction.

35. Kong AT, Leprevost F V., Avtonomov DM, Mellacheruvu D, Nesvizhskii AI. MSFragger: Ultrafast and comprehensive peptide identification in mass spectrometry-based proteomics. Nature Methods 2017;14:513–520.

36. Krisher RL. Present state and future outlook for the application of in vitro oocyte maturation in human infertility treatment. Biology of Reproduction 2022;106:.

37. Kuwayama M. Highly efficient vitrification for cryopreservation of human oocytes and embryos: The Cryotop method. Theriogenology 2007;67:.

38. Lange-Consiglio A, Perrini C, Albini G, Modina S, Lodde V, Orsini E, Esposti P, Cremonesi F. Oviductal microvesicles and their effect on in vitro maturation of canine oocytes. Reproduction 2017a;154:167–180. BioScientifica Ltd.

39. Lange-Consiglio A, Perrini C, Albini G, Modina S, Lodde V, Orsini E, Esposti P, Cremonesi F. Oviductal microvesicles and their effect on in vitro maturation of canine oocytes. Reproduction 2017b;154:.

40. Le HL, Ho VNA, Le TTN, Tran VTT, Ma MPQ, Le AH, Nguyen LK, Ho TM, Vuong LN. Live birth after in vitro maturation in women with gonadotropin resistance ovary syndrome: report of two cases. Journal of Assisted Reproduction and Genetics 2021;38:.

41. Lim HYG, Alvarez YD, Gasnier M, Wang Y, Tetlak P, Bissiere S, Wang H, Biro M, Plachta N. Keratins are asymmetrically inherited fate determinants in the mammalian embryo. Nature 2020;585:404–409. Nature Research.

42. Liu X, Wang Y, Zhu P, Wang J, Liu J, Li N, Wang W, Zhang W, Zhang C, Wang Y, et al. Human follicular fluid proteome reveals association between overweight status and oocyte maturation abnormality. Clinical Proteomics 2020;17:. BioMed Central Ltd.

43. Macaulay AD, Gilbert I, Caballero J, Barreto R, Fournier E, Tossou P, Sirard MA, Clarke HJ, Khandjian ÉW, Richard FJ, et al. The gametic synapse: RNA transfer to the bovine oocyte. Biology of Reproduction 2014;91:. Society for the Study of Reproduction.

44. Machtinger R, Laurent LC, Baccarelli AA. Extracellular vesicles: Roles in gamete maturation, fertilisation and embryo implantation. Human Reproduction Update 2016;22:.

45. MacKens S, Pareyn S, Drakopoulos P, Deckers T, Mostinckx L, Blockeel C, Segers I, Verheyen G, Santos-Ribeiro S, Tournaye H, et al. Outcome of in-vitro oocyte maturation in patients with PCOS: Does phenotype have an impact? Human Reproduction 2020;35:.

46. Macklon NS, Stouffer RL, Giudice LC, Fauser BCJM. The science behind 25 years of ovarian stimulation for in vitro fertilisation. Endocrine Reviews 2006;27:.

47. Makieva S, Fraire-zamora JJ, Ammar OF, Liperis G, Sanchez F, Vuong LN, Gilchrist RB, Bortoletto P, Massarotti C. Road to in vitro maturation (IVM), from basic science to an informed clinical practice. 2024;00:1–6.

48. Mandelbaum RS, Awadalla MS, Smith MB, Violette CJ, Klooster BL, Danis RB, McGinnis LK, Ho JR, Bendikson KA, Paulson RJ, et al. Developmental potential of immature human oocytes aspirated after controlled ovarian stimulation. Journal of Assisted Reproduction and Genetics 2021;38:.

49. Marchais M, Gilbert I, Bastien A, Macaulay A, Robert C. Mammalian cumulus-oocyte complex communication: a dialog through long and short distance messaging. Journal of Assisted Reproduction and Genetics 2022;39:1011–1025. Springer.

50. Martinez RM, Liang L, Racowsky C, Dioni L, Mansur A, Adir M, Bollati V, Baccarelli AA, Hauser R, Machtinger R. Extracellular microRNAs profile in human follicular fluid and IVF outcomes. Scientific Reports 2018;8:. Nature Publishing Group.

51. McKenzie LJ, Pangas SA, Carson SA, Kovanci E, Cisneros P, Buster JE, Amato P, Matzuk MM. Human cumulus granulosa cell gene expression: A predictor of fertilisation and embryo selection in women undergoing IVF. Human Reproduction 2004;19:2869–2874. Oxford University Press.

52. Merton JS, Roos APW De, Mullaart E, Ruigh L De, Kaal L, Vos PLAM, Dieleman SJ. Factors affecting oocyte quality and quantity in commercial application of embryo technologies in the cattle breeding industry. Theriogenology 2003;59:.

53. Miglietta S, Cristiano L, Battaglione E, Macchiarelli G, Nottola SA, Marco MP De, Costanzi F, Schimberni M, Colacurci N, Caserta D, et al. Heavy Metals in Follicular Fluid Affect the Ultrastructure of the Human Mature Cumulus-Oocyte Complex. Cells 2023;12:.

54. Multidisciplinary Digital Publishing Institute (MDPI).

55. Morris JH, Huerta-Cepas J, Junge A, Szklarczyk D, Jensen LJ, Mering C von, Lyon D, Gable AL, Wyder S, Simonovic M, et al. STRING v11: protein–protein association networks with increased coverage, supporting functional discovery in genome-wide experimental datasets. Nucleic Acids Research 2018;47:D607–D613. Oxford University Press.

56. Mostinckx L, Goyens E, Mackens S, Roelens C, Boudry L, Uvin V, Segers I, Schoemans C, Drakopoulos P, Blockeel C, et al. Clinical outcomes from ART in predicted hyperresponders: in vitro maturation of oocytes versus conventional ovarian stimulation for IVF/ICSI. Human Reproduction 2024;39:.

57. Mudunuri U, Che A, Yi M, Stephens RM. bioDBnet: The biological database network. Bioinformatics 2009;25:.

58. Mulcahy LA, Pink RC, Carter DRF. Routes and mechanisms of extracellular vesicle uptake. Journal of Extracellular Vesicles 2014;3:. Co-Action Publishing.

59. Nargund G, Datta AK, Campbell S, Patrizio P, Chian RC, Ombelet W, Wolff M Von, Lindenberg S, Frydman R, Fauser BC. The case for mild stimulation for IVF: recommendations from The International Society for Mild Approaches in Assisted Reproduction. Reproductive BioMedicine Online 2022;45:.

60. Njagi P, Groot W, Arsenijevic J, Dyer S, Mburu G, Kiarie J. Financial costs of assisted reproductive technology for patients in low-and middle-income countries: A systematic review. Human Reproduction Open 2023;2023:.

61. Nõmm M, Ivask M, Pärn P, Reimann E, Kõks S, Jaakma Ü. Detecting Embryo Developmental Potential by Single Blastomere RNA-Seq. Genes 2023;14:. MDPI.

62. Oslowski CM, Urano F. Measuring ER stress and the unfolded protein response using mammalian tissue culture system. Methods in Enzymology 2011;490:71–92. Academic Press Inc.

63. Otsuki J, Nagai Y, Chiba K. Lipofuscin bodies in human oocytes as an indicator of oocyte quality. Journal of Assisted Reproduction and Genetics 2007;24:263–270.

64. Palmerini MG, Antinori M, Maione M, Cerusico F, Versaci C, Nottola SA, Macchiarelli G, Khalili MA, Antinori S. Ultrastructure of immature and mature human oocytes after cryotop vitrification. Journal of Reproduction and Development 2014;60:411–420. The Japanese Society of Animal Reproduction (JSAR).

65. Pioltine EM, Machado MF, Silveira JC da, Fontes PK, Botigelli RC, Quaglio AE V., Costa CB, Nogueira MFG. Can extracellular vesicles from bovine ovarian follicular fluid modulate the in-vitro oocyte meiosis progression similarly to the CNP-NPR2 system? Theriogenology 2020a;157:.

66. Pioltine EM, Machado MF, Silveira JC da, Fontes PK, Botigelli RC, Quaglio AE V., Costa CB, Nogueira MFG. Can extracellular vesicles from bovine ovarian follicular fluid modulate the in-vitro oocyte meiosis progression similarly to the CNP-NPR2 system? Theriogenology 2020b;157:.

67. Pla I, Sanchez A, Pors SE, Kristensen SG, Appelqvist R, Sahlin KB, Marko-Varga G, Andersen CY, Malm J. Proteomic Alterations in Follicular Fluid of Human Small Antral Follicles Collected from Polycystic Ovaries—A Pilot Study. Life 2022;12:.

68. Poulsen LC, Bøtkjær JA, Østrup O, Petersen KB, Yding Andersen C, Grøndahl ML, Englund ALM. Two waves of transcriptomic changes in periovulatory human granulosa cells. Human Reproduction 2020;35:1230–1245. Oxford University Press.

69. Reichhardt MP, Lundin K, Lokki AI, Recher G, Vuoristo S, Katayama S, Tapanainen JS, Kere J, Meri S, Tuuri T. Complement in Human Pre-implantation Embryos: Attack and Defense. Frontiers in Immunology 2019;10:.

70. Ritchie ME, Phipson B, Wu D, Hu Y, Law CW, Shi W, Smyth GK. Limma powers differential expression analyses for RNA-sequencing and microarray studies. Nucleic Acids Research 2015;43:.

71. Rock J, Menkin MF. In vitro fertilisation and cleavage of human ovarian eggs. Science 1944;100:.

72. Rodrigues TA, Tuna KM, Alli AA, Tribulo P, Hansen PJ, Koh J, Paula-Lopes FF. Follicular fluid exosomes act on the bovine oocyte to improve oocyte competence to support development and survival to heat shock. *Reproduction*, Fertility and Development 2019;31:888–897. CSIRO.

73. Roesner S, Wolff M Von, Elsaesser M, Roesner K, Reuner G, Pietz J, Bruckner T, Strowitzki T. Two-year development of children conceived by IVM: A prospective controlled single-blinded study. Human Reproduction 2017;32:.

74. Runge KE, Evans JE, He ZY, Gupta S, McDonald KL, Stahlberg H, Primakoff P, Myles DG. Oocyte CD9 is enriched on the microvillar membrane and required for normal microvillar shape and distribution. Developmental Biology 2007;304:317–325. Academic Press Inc.

75. Saenz-de-Juano M, Silvestrelli G, Bauersachs S, Ulbrich SE. Determining extracellular vesicles properties and miRNA cargo variability in bovine milk from healthy cows and cows undergoing subclinical mastitis. BMC Genomics 2022;23:1–15. BioMed Central.

76. Santonocito M, Vento M, Guglielmino MR, Battaglia R, Wahlgren J, Ragusa M, Barbagallo D, Borz?? P, Rizzari S, Maugeri M, et al. Molecular characterization of exosomes and their microRNA cargo in human follicular fluid: Bioinformatic analysis reveals that exosomal microRNAs control pathways involved in follicular maturation. Fertility and Sterility 2014;102:1751–1761. Elsevier Inc.

77. Schoenfelder M, Einspanier R. Expression of hyaluronan synthases and corresponding hyaluronan receptors is differentially regulated during oocyte maturation in cattle. Biology of Reproduction 2003;69:269–277.

78. Shani AK, Haham LM, Balakier H, Kuznyetsova I, Bashar S, Day EN, Librach CL. The developmental potential of mature oocytes derived from rescue in vitro maturation. Fertility and Sterility 2023;120:.

79. Siiskonen H, Oikari S, Pasonen-Seppänen S, Rilla K. Hyaluronan synthase 1: A mysterious enzyme with unexpected functions. Frontiers in Immunology 2015;6:. Frontiers Media S.A.

80. Silveira JC da, Andrade GM, Collado M del, Sampaio R V., Sangalli JR, Silva LA, Pinaffi FVL, Jardim IB, Cesar MC, Nogueira MFG, et al. Supplementation with small- extracellular vesicles from ovarian follicular fluid during in vitro production modulates bovine embryo development. PLoS ONE 2017a;12:. Public Library of Science.

81. Silveira JC da, Andrade GM, Collado M del, Sampaio R V., Sangalli JR, Silva LA, Pinaffi FVL, Jardim IB, Cesar MC, Nogueira MFG, et al. Supplementation with small- extracellular vesicles from ovarian follicular fluid during in vitro production modulates bovine embryo development. PLoS ONE 2017b;12:.

82. Smyth GK. Linear models and empirical bayes methods for assessing differential expression in microarray experiments. Statistical Applications in Genetics and Molecular Biology 2004;3:.

83. Stock AE, Bouchard N, Brown K, Spicer AP, Underhill CB, Doré M, Sirois J. Induction of hyaluronan synthase 2 by human chorionic gonadotropin in mural granulosa cells of equine preovulatory follicles. Endocrinology 2002;143:4375–4384. Endocrine Society.

84. Turathum B, Roytrakul S, Changsangfa C, Sroyraya M, Tanasawet S, Kitiyanant Y, Saikhun K. Missing and overexpressing proteins in domestic cat oocytes following vitrification and in vitro maturation as revealed by proteomic analysis. Biological Research 2018;51:. BioMed Central Ltd.

85. Uzbekova S, Almiñana C, Labas V, Teixeira-Gomes AP, Combes-Soia L, Tsikis G, Carvalho AV, Uzbekov R, Singina G. Protein Cargo of Extracellular Vesicles From Bovine Follicular Fluid and Analysis of Their Origin From Different Ovarian Cells. Frontiers in Veterinary Science 2020a;7:.

86. Uzbekova S, Almiñana C, Labas V, Teixeira-Gomes AP, Combes-Soia L, Tsikis G, Carvalho AV, Uzbekov R, Singina G. Protein Cargo of Extracellular Vesicles From Bovine Follicular Fluid and Analysis of Their Origin From Different Ovarian Cells. Frontiers in Veterinary Science 2020b;7:.

87. Vašíčková K, Moráň L, Gurín D, Vaňhara P. Alleviation of endoplasmic reticulum stress by tauroursodeoxycholic acid delays senescence of mouse ovarian surface epithelium. Cell and Tissue Research 2018;374:643–652. Springer Verlag.

88. Viana J. 2019 Statistics of embryo production and transfer in domestic farm animals Divergent trends for IVD and IVP embryos. Embryo Technology Newsletter 2020;38:.

89. Virant-Klun I, Leicht S, Hughes C, Krijgsveld J. Identification of maturation-specific proteins by single-cell proteomics of human oocytes. Molecular and Cellular Proteomics 2016;15:2616–2627. American Society for Biochemistry and Molecular Biology Inc.

90. Vos M De, Grynberg M, Ho TM, Yuan Y, Albertini DF, Gilchrist RB. Perspectives on the development and future of oocyte IVM in clinical practice. Journal of Assisted Reproduction and Genetics 2021;38:.

91. Vuong LN, Ho VNA, Ho TM, Dang VQ, Phung TH, Giang NH, Le AH, Pham TD, Wang R, Smitz J, et al. In-vitro maturation of oocytes versus conventional IVF in women with infertility and a high antral follicle count: A randomized non-inferiority controlled trial. Human Reproduction 2020;35:.

92. Wang Y, Luo FQ, He YH, Yang ZX, Wang X, Li CR, Cai BQ, Chen LJ, Wang Z Bin, Zhang CL, et al. Oocytes could rearrange immunoglobulin production to survive over adverse environmental stimuli. Frontiers in Immunology 2022;13:.

93. Whitaker M. Calcium at fertilisation and in early development. Physiological Reviews 2006;86:25–88.

94. Wolski WE, Nanni P, Grossmann J, d’Errico M, Schlapbach R, Panse C. prolfqua: A Comprehensive R-Package for Proteomics Differential Expression Analysis. Journal of Proteome Research 2023;22:.

95. Yokoo M, Sato E. Physiological function of hyaluronan in mammalian oocyte maturation. Reproductive Medicine and Biology 2011;10:221–229. John Wiley and Sons Ltd.

96. Yu EJ, Choi WY, Park MS, Eum JH, Lee DR, Lee WS, Lyu SW, Yoon SY. RNA sequencing-based transcriptome analysis of granulosa cells from follicular fluid: Genes involved in embryo quality during in vitro fertilisation and embryo transfer. PLoS ONE 2023;18:.

97. Yuan Y, Reed L, Swain JE, Schoolcraft WB, Katz-Jaffe MG. Rescue in vitro maturation and the transfer of a euploid blastocyst provided improved chances for patients with poor prognosis to conceive. Fertility and Sterility 2024;121:.

98. Zha D, Rayamajhi S, Sipes J, Russo A, Pathak HB, Li K, Sardiu ME, Bantis LE, Mitra A, Puri R V., et al. Proteomic Profiling of Fallopian Tube-Derived Extracellular Vesicles Using a Microfluidic Tissue-on-Chip System. Bioengineering 2023;10:. MDPI.

99. Zhang X, Smits AH, Tilburg GBA Van, Ovaa H, Huber W, Vermeulen M. Proteome-wide identification of ubiquitin interactions using UbIA-MS. Nature Protocols 2018;13:.

100. Zheng X, Guo W, Zeng L, Zheng D, Yang S, Xu Y, Wang L, Wang R, Mol BW, Li R, et al. In vitro maturation without gonadotropins versus in vitro fertilisation with hyperstimulation in women with polycystic ovary syndrome: A non-inferiority randomized controlled trial. Human Reproduction 2022;37:.

101. Zhou Y, Yang X, Liu Z, Zhang Y, Chen H, Zhang Y, Hu Y, Ma Y, Li Q. Excluding embryos with two novel mutations in FREM2 gene by the next-generation sequencing-based single nucleotide polymorphism haplotyping. Aging 2021;13:24786–24794. Impact Journals LLC.

