## Supplementary tables for "Treatment of human oocytes with extracellular vesicles from follicular fluid during rescue in vitro maturation enhances maturation rates and modulates oocyte proteome and ultrastructure"

- Supplementary Table 1 – Page 2
- Supplementary Table 2 – Page 44
- Supplementary Table 3 – Page 103
- Supplementary Table 4 – Page 128
- Supplementary Table 5 – Page 130
- Supplementary Table 6 – Page 134
- Supplementary Table 7 – Page 149
- Supplementary Table 8 – Page 371
- Supplementary Table 9 – Page 493
- Supplementary Table 10 – Page 495
- Supplementary Table 11 – Page 499

| **Supplementary Table 1: Identified Proteins (1263) in ffEVs coming from MII oocytes** | | | |
| --- | --- | --- | --- |
| Scaffold parameters - Protein Threshold: 1% FDR, Minimum peptides: 1, Peptide Threshold 0.1% FDR. | | | |
| **Description** | **Accession Number** | **Name** | **Molecular Weight** |
| Actin, alpha cardiac muscle 1 OS=Homo sapiens OX=9606 GN=ACTC1 PE=1 SV=1 | P68032 | ACTC_HUMAN (+1) | 42 kDa |
| Actin, cytoplasmic 1 OS=Homo sapiens OX=9606 GN=ACTB PE=1 SV=1 | P60709 | ACTB_HUMAN | 42 kDa |
| Actin, cytoplasmic 2 OS=Homo sapiens OX=9606 GN=ACTG1 PE=1 SV=1 | P63261 | ACTG_HUMAN | 42 kDa |
| ADP-ribosylation factor 3 OS=Homo sapiens OX=9606 GN=ARF3 PE=1 SV=2 | P61204 | ARF3_HUMAN (+1) | 21 kDa |
| ADP-ribosylation factor 4 OS=Homo sapiens OX=9606 GN=ARF4 PE=1 SV=3 | P18085 | ARF4_HUMAN | 21 kDa |
| ADP-ribosylation factor 5 OS=Homo sapiens OX=9606 GN=ARF5 PE=1 SV=2 | P84085 | ARF5_HUMAN | 21 kDa |
| Albumin OS=Bos taurus OX=9913 GN=ALB PE=1 SV=4 | P02769 | ALBU_BOVIN | 69 kDa |
| Albumin OS=Homo sapiens OX=9606 GN=ALB PE=1 SV=2 | P02768 | ALBU_HUMAN | 69 kDa |
| Alpha-actinin-1 OS=Homo sapiens OX=9606 GN=ACTN1 PE=1 SV=2 | P12814 | ACTN1_HUMAN | 103 kDa |
| Alpha-actinin-4 OS=Homo sapiens OX=9606 GN=ACTN4 PE=1 SV=2 | O43707 | ACTN4_HUMAN | 105 kDa |
| Cofilin-1 OS=Homo sapiens OX=9606 GN=CFL1 PE=1 SV=3 | P23528 | COF1_HUMAN | 19 kDa |
| Cofilin-2 OS=Homo sapiens OX=9606 GN=CFL2 PE=1 SV=1 | Q9Y281 | COF2_HUMAN | 19 kDa |
| Complement C4-A OS=Homo sapiens OX=9606 GN=C4A PE=1 SV=2 | P0C0L4 | CO4A_HUMAN | 193 kDa |
| Complement C4-B OS=Homo sapiens OX=9606 GN=C4B_2 PE=1 SV=2 | P0C0L5 | CO4B_HUMAN | 193 kDa |
| Eukaryotic initiation factor 4A-I OS=Homo sapiens OX=9606 GN=EIF4A1 PE=1 SV=1 | P60842 | IF4A1_HUMAN | 46 kDa |
| Eukaryotic initiation factor 4A-II OS=Homo sapiens OX=9606 GN=EIF4A2 PE=1 SV=2 | Q14240 | IF4A2_HUMAN | 46 kDa |
| Guanine nucleotide-binding protein G(I)/G(S)/G(T) subunit beta-1 OS=Homo sapiens OX=9606 GN=GNB1 PE=1 SV=3 | P62873 | GBB1_HUMAN | 37 kDa |
| Guanine nucleotide-binding protein G(I)/G(S)/G(T) subunit beta-2 OS=Homo sapiens OX=9606 GN=GNB2 PE=1 SV=3 | P62879 | GBB2_HUMAN | 37 kDa |
| Haptoglobin OS=Homo sapiens OX=9606 GN=HP PE=1 SV=1 | P00738 | HPT_HUMAN | 45 kDa |
| Haptoglobin-related protein OS=Homo sapiens OX=9606 GN=HPR PE=2 SV=2 | P00739 | HPTR_HUMAN | 39 kDa |
| Immunoglobulin heavy constant alpha 1 OS=Homo sapiens OX=9606 GN=IGHA1 PE=1 SV=2 | P01876 | IGHA1_HUMAN | 38 kDa |
| Immunoglobulin heavy constant alpha 2 OS=Homo sapiens OX=9606 GN=IGHA2 PE=1 SV=4 | P01877 | IGHA2_HUMAN | 37 kDa |
| Immunoglobulin heavy constant gamma 1 OS=Homo sapiens OX=9606 GN=IGHG1 PE=1 SV=1 | P01857 | IGHG1_HUMAN | 36 kDa |
| Immunoglobulin heavy constant gamma 2 OS=Homo sapiens OX=9606 GN=IGHG2 PE=1 SV=2 | P01859 | IGHG2_HUMAN | 36 kDa |
| Immunoglobulin heavy constant gamma 3 OS=Homo sapiens OX=9606 GN=IGHG3 PE=1 SV=2 | P01860 | IGHG3_HUMAN | 41 kDa |
| Immunoglobulin heavy constant gamma 4 OS=Homo sapiens OX=9606 GN=IGHG4 PE=1 SV=1 | P01861 | IGHG4_HUMAN | 36 kDa |
| Immunoglobulin lambda constant 2 OS=Homo sapiens OX=9606 GN=IGLC2 PE=1 SV=1 | P0DOY2 | IGLC2_HUMAN (+1) | 11 kDa |
| Immunoglobulin lambda variable 3-16 OS=Homo sapiens OX=9606 GN=IGLV3-16 PE=3 SV=2 | A0A075B6K0 | LV316_HUMAN (+2) | 12 kDa |
| Immunoglobulin lambda variable 3-9 OS=Homo sapiens OX=9606 GN=IGLV3-9 PE=3 SV=1 | A0A075B6K5 | LV39_HUMAN (+1) | 12 kDa |
| Immunoglobulin lambda-like polypeptide 5 OS=Homo sapiens OX=9606 GN=IGLL5 PE=2 SV=2 | B9A064 | IGLL5_HUMAN | 23 kDa |
| Keratin, type I cuticular Ha6 OS=Homo sapiens OX=9606 GN=KRT36 PE=1 SV=1 | O76013 | KRT36_HUMAN | 52 kDa |
| Keratin, type I cytoskeletal 10 OS=Homo sapiens OX=9606 GN=KRT10 PE=1 SV=6 | P13645 | K1C10_HUMAN | 59 kDa |
| Keratin, type I cytoskeletal 14 OS=Homo sapiens OX=9606 GN=KRT14 PE=1 SV=4 | P02533 | K1C14_HUMAN | 52 kDa |
| Keratin, type I cytoskeletal 16 OS=Homo sapiens OX=9606 GN=KRT16 PE=1 SV=4 | P08779 | K1C16_HUMAN | 51 kDa |
| Keratin, type I cytoskeletal 19 OS=Homo sapiens OX=9606 GN=KRT19 PE=1 SV=4 | P08727 | K1C19_HUMAN | 44 kDa |
| Keratin, type I microfibrillar 48 kDa, component 8C-1 OS=Ovis aries OX=9940 PE=1 SV=2 | P02534 | K1M1_SHEEP | 47 kDa |
| Keratin, type I microfibrillar, 47.6 kDa OS=Ovis aries OX=9940 PE=3 SV=2 | P25690 | K1M2_SHEEP | 46 kDa |
| Keratin, type II cuticular Hb5 OS=Homo sapiens OX=9606 GN=KRT85 PE=1 SV=1 | P78386 | KRT85_HUMAN | 56 kDa |
| Keratin, type II cytoskeletal 1 OS=Homo sapiens OX=9606 GN=KRT1 PE=1 SV=6 | P04264 | K2C1_HUMAN | 66 kDa |
| Keratin, type II cytoskeletal 1b OS=Homo sapiens OX=9606 GN=KRT77 PE=1 SV=3 | Q7Z794 | K2C1B_HUMAN | 62 kDa |
| Keratin, type II cytoskeletal 2 epidermal OS=Homo sapiens OX=9606 GN=KRT2 PE=1 SV=2 | P35908 | K22E_HUMAN | 65 kDa |
| Keratin, type II cytoskeletal 5 OS=Homo sapiens OX=9606 GN=KRT5 PE=1 SV=3 | P13647 | K2C5_HUMAN | 62 kDa |
| Keratin, type II cytoskeletal 6A OS=Homo sapiens OX=9606 GN=KRT6A PE=1 SV=3 | P02538 | K2C6A_HUMAN | 60 kDa |
| Keratin, type II cytoskeletal 6C OS=Homo sapiens OX=9606 GN=KRT6C PE=1 SV=3 | P48668 | K2C6C_HUMAN | 60 kDa |
| Keratin, type II cytoskeletal 79 OS=Homo sapiens OX=9606 GN=KRT79 PE=1 SV=2 | Q5XKE5 | K2C79_HUMAN | 58 kDa |
| Keratin, type II cytoskeletal 8 OS=Homo sapiens OX=9606 GN=KRT8 PE=1 SV=7 | P05787 | K2C8_HUMAN | 54 kDa |
| Keratin, type II microfibrillar, component 5 OS=Ovis aries OX=9940 PE=1 SV=1 | P25691 | K2M3_SHEEP | 55 kDa |
| Keratin, type II microfibrillar, component 7C OS=Ovis aries OX=9940 PE=1 SV=1 | P15241 | K2M2_SHEEP | 54 kDa |
| Lactotransferrin OS=Homo sapiens OX=9606 GN=LTF PE=1 SV=6 | P02788 | TRFL_HUMAN | 78 kDa |
| Peroxiredoxin-1 OS=Homo sapiens OX=9606 GN=PRDX1 PE=1 SV=1 | Q06830 | PRDX1_HUMAN | 22 kDa |
| Peroxiredoxin-4 OS=Homo sapiens OX=9606 GN=PRDX4 PE=1 SV=1 | Q13162 | PRDX4_HUMAN | 31 kDa |
| Proto-oncogene tyrosine-protein kinase Src OS=Homo sapiens OX=9606 GN=SRC PE=1 SV=3 | P12931 | SRC_HUMAN | 60 kDa |
| Ras-related protein Rab-10 OS=Homo sapiens OX=9606 GN=RAB10 PE=1 SV=1 | P61026 | RAB10_HUMAN | 23 kDa |
| Ras-related protein Rab-1A OS=Homo sapiens OX=9606 GN=RAB1A PE=1 SV=3 | P62820 | RAB1A_HUMAN | 23 kDa |
| Ras-related protein Rab-1B OS=Homo sapiens OX=9606 GN=RAB1B PE=1 SV=1 | Q9H0U4 | RAB1B_HUMAN | 22 kDa |
| Ras-related protein Rab-8A OS=Homo sapiens OX=9606 GN=RAB8A PE=1 SV=1 | P61006 | RAB8A_HUMAN | 24 kDa |
| Ras-related protein Rab-8B OS=Homo sapiens OX=9606 GN=RAB8B PE=1 SV=2 | Q92930 | RAB8B_HUMAN | 24 kDa |
| Ras-related protein Ral-A OS=Homo sapiens OX=9606 GN=RALA PE=1 SV=1 | P11233 | RALA_HUMAN | 24 kDa |
| Ras-related protein Ral-B OS=Homo sapiens OX=9606 GN=RALB PE=1 SV=1 | P11234 | RALB_HUMAN | 23 kDa |
| Serotransferrin OS=Homo sapiens OX=9606 GN=TF PE=1 SV=3 | P02787 | TRFE_HUMAN | 77 kDa |
| Tropomyosin alpha-3 chain OS=Homo sapiens OX=9606 GN=TPM3 PE=1 SV=2 | P06753 | TPM3_HUMAN | 33 kDa |
| Tropomyosin alpha-4 chain OS=Homo sapiens OX=9606 GN=TPM4 PE=1 SV=3 | P67936 | TPM4_HUMAN | 29 kDa |
| Tubulin alpha-1A chain OS=Homo sapiens OX=9606 GN=TUBA1A PE=1 SV=1 | Q71U36 | TBA1A_HUMAN | 50 kDa |
| Tubulin alpha-1B chain OS=Homo sapiens OX=9606 GN=TUBA1B PE=1 SV=1 | P68363 | TBA1B_HUMAN | 50 kDa |
| Tubulin beta chain OS=Homo sapiens OX=9606 GN=TUBB PE=1 SV=2 | P07437 | TBB5_HUMAN | 50 kDa |
| Tubulin beta-4A chain OS=Homo sapiens OX=9606 GN=TUBB4A PE=1 SV=2 | P04350 | TBB4A_HUMAN | 50 kDa |
| Tubulin beta-4B chain OS=Homo sapiens OX=9606 GN=TUBB4B PE=1 SV=1 | P68371 | TBB4B_HUMAN | 50 kDa |
| Tyrosine-protein kinase Yes OS=Homo sapiens OX=9606 GN=YES1 PE=1 SV=3 | P07947 | YES_HUMAN | 61 kDa |
| Vesicle-associated membrane protein 2 OS=Homo sapiens OX=9606 GN=VAMP2 PE=1 SV=3 | P63027 | VAMP2_HUMAN | 13 kDa |
| Vesicle-associated membrane protein 3 OS=Homo sapiens OX=9606 GN=VAMP3 PE=1 SV=3 | Q15836 | VAMP3_HUMAN | 11 kDa |
| (E3-independent) E2 ubiquitin-conjugating enzyme OS=Homo sapiens OX=9606 GN=UBE2O PE=1 SV=3 | Q9C0C9 | UBE2O_HUMAN | 141 kDa |
| 1,4-alpha-glucan-branching enzyme OS=Homo sapiens OX=9606 GN=GBE1 PE=1 SV=3 | Q04446 | GLGB_HUMAN | 80 kDa |
| 14-3-3 protein beta/alpha OS=Homo sapiens OX=9606 GN=YWHAB PE=1 SV=3 | P31946 | 1433B_HUMAN | 28 kDa |
| 14-3-3 protein epsilon OS=Homo sapiens OX=9606 GN=YWHAE PE=1 SV=1 | P62258 | 1433E_HUMAN | 29 kDa |
| 14-3-3 protein eta OS=Homo sapiens OX=9606 GN=YWHAH PE=1 SV=4 | Q04917 | 1433F_HUMAN | 28 kDa |
| 14-3-3 protein gamma OS=Homo sapiens OX=9606 GN=YWHAG PE=1 SV=2 | P61981 | 1433G_HUMAN | 28 kDa |
| 14-3-3 protein sigma OS=Homo sapiens OX=9606 GN=SFN PE=1 SV=1 | P31947 | 1433S_HUMAN | 28 kDa |
| 14-3-3 protein theta OS=Homo sapiens OX=9606 GN=YWHAQ PE=1 SV=1 | P27348 | 1433T_HUMAN | 28 kDa |
| 14-3-3 protein zeta/delta OS=Homo sapiens OX=9606 GN=YWHAZ PE=1 SV=1 | P63104 | 1433Z_HUMAN | 28 kDa |
| 17-beta-hydroxysteroid dehydrogenase type 1 OS=Homo sapiens OX=9606 GN=HSD17B1 PE=1 SV=3 | P14061 | DHB1_HUMAN | 35 kDa |
| 1-phosphatidylinositol 4,5-bisphosphate phosphodiesterase gamma-1 OS=Homo sapiens OX=9606 GN=PLCG1 PE=1 SV=1 | P19174 | PLCG1_HUMAN | 149 kDa |
| 2',3'-cyclic-nucleotide 3'-phosphodiesterase OS=Homo sapiens OX=9606 GN=CNP PE=1 SV=2 | P09543 | CN37_HUMAN | 48 kDa |
| 2,4-dienoyl-CoA reductase [(3E)-enoyl-CoA-producing], mitochondrial OS=Homo sapiens OX=9606 GN=DECR1 PE=1 SV=1 | Q16698 | DECR_HUMAN | 36 kDa |
| 26S proteasome non-ATPase regulatory subunit 1 OS=Homo sapiens OX=9606 GN=PSMD1 PE=1 SV=2 | Q99460 | PSMD1_HUMAN | 106 kDa |
| 26S proteasome non-ATPase regulatory subunit 10 OS=Homo sapiens OX=9606 GN=PSMD10 PE=1 SV=1 | O75832 | PSD10_HUMAN | 24 kDa |
| 26S proteasome non-ATPase regulatory subunit 11 OS=Homo sapiens OX=9606 GN=PSMD11 PE=1 SV=3 | O00231 | PSD11_HUMAN | 47 kDa |
| 26S proteasome non-ATPase regulatory subunit 12 OS=Homo sapiens OX=9606 GN=PSMD12 PE=1 SV=3 | O00232 | PSD12_HUMAN | 53 kDa |
| 26S proteasome non-ATPase regulatory subunit 13 OS=Homo sapiens OX=9606 GN=PSMD13 PE=1 SV=2 | Q9UNM6 | PSD13_HUMAN | 43 kDa |
| 26S proteasome non-ATPase regulatory subunit 14 OS=Homo sapiens OX=9606 GN=PSMD14 PE=1 SV=1 | O00487 | PSDE_HUMAN | 35 kDa |
| 26S proteasome non-ATPase regulatory subunit 2 OS=Homo sapiens OX=9606 GN=PSMD2 PE=1 SV=3 | Q13200 | PSMD2_HUMAN | 100 kDa |
| 26S proteasome non-ATPase regulatory subunit 3 OS=Homo sapiens OX=9606 GN=PSMD3 PE=1 SV=2 | O43242 | PSMD3_HUMAN | 61 kDa |
| 26S proteasome non-ATPase regulatory subunit 4 OS=Homo sapiens OX=9606 GN=PSMD4 PE=1 SV=1 | P55036 | PSMD4_HUMAN | 41 kDa |
| 26S proteasome non-ATPase regulatory subunit 6 OS=Homo sapiens OX=9606 GN=PSMD6 PE=1 SV=1 | Q15008 | PSMD6_HUMAN | 46 kDa |
| 26S proteasome non-ATPase regulatory subunit 7 OS=Homo sapiens OX=9606 GN=PSMD7 PE=1 SV=2 | P51665 | PSMD7_HUMAN | 37 kDa |
| 26S proteasome regulatory subunit 10B OS=Homo sapiens OX=9606 GN=PSMC6 PE=1 SV=1 | P62333 | PRS10_HUMAN | 44 kDa |
| 26S proteasome regulatory subunit 4 OS=Homo sapiens OX=9606 GN=PSMC1 PE=1 SV=1 | P62191 | PRS4_HUMAN | 49 kDa |
| 26S proteasome regulatory subunit 6A OS=Homo sapiens OX=9606 GN=PSMC3 PE=1 SV=3 | P17980 | PRS6A_HUMAN | 49 kDa |
| 26S proteasome regulatory subunit 6B OS=Homo sapiens OX=9606 GN=PSMC4 PE=1 SV=2 | P43686 | PRS6B_HUMAN | 47 kDa |
| 26S proteasome regulatory subunit 7 OS=Homo sapiens OX=9606 GN=PSMC2 PE=1 SV=3 | P35998 | PRS7_HUMAN | 49 kDa |
| 26S proteasome regulatory subunit 8 OS=Homo sapiens OX=9606 GN=PSMC5 PE=1 SV=1 | P62195 | PRS8_HUMAN | 46 kDa |
| 3-hydroxyacyl-CoA dehydrogenase type-2 OS=Homo sapiens OX=9606 GN=HSD17B10 PE=1 SV=3 | Q99714 | HCD2_HUMAN | 27 kDa |
| 40S ribosomal protein S11 OS=Homo sapiens OX=9606 GN=RPS11 PE=1 SV=3 | P62280 | RS11_HUMAN | 18 kDa |
| 40S ribosomal protein S12 OS=Homo sapiens OX=9606 GN=RPS12 PE=1 SV=3 | P25398 | RS12_HUMAN | 15 kDa |
| 40S ribosomal protein S13 OS=Homo sapiens OX=9606 GN=RPS13 PE=1 SV=2 | P62277 | RS13_HUMAN | 17 kDa |
| 40S ribosomal protein S14 OS=Homo sapiens OX=9606 GN=RPS14 PE=1 SV=3 | P62263 | RS14_HUMAN | 16 kDa |
| 40S ribosomal protein S15a OS=Homo sapiens OX=9606 GN=RPS15A PE=1 SV=2 | P62244 | RS15A_HUMAN | 15 kDa |
| 40S ribosomal protein S16 OS=Homo sapiens OX=9606 GN=RPS16 PE=1 SV=2 | P62249 | RS16_HUMAN | 16 kDa |
| 40S ribosomal protein S18 OS=Homo sapiens OX=9606 GN=RPS18 PE=1 SV=3 | P62269 | RS18_HUMAN | 18 kDa |
| 40S ribosomal protein S2 OS=Homo sapiens OX=9606 GN=RPS2 PE=1 SV=2 | P15880 | RS2_HUMAN | 31 kDa |
| 40S ribosomal protein S20 OS=Homo sapiens OX=9606 GN=RPS20 PE=1 SV=1 | P60866 | RS20_HUMAN | 13 kDa |
| 40S ribosomal protein S24 OS=Homo sapiens OX=9606 GN=RPS24 PE=1 SV=1 | P62847 | RS24_HUMAN | 15 kDa |
| 40S ribosomal protein S25 OS=Homo sapiens OX=9606 GN=RPS25 PE=1 SV=1 | P62851 | RS25_HUMAN | 14 kDa |
| 40S ribosomal protein S26 OS=Homo sapiens OX=9606 GN=RPS26 PE=1 SV=3 | P62854 | RS26_HUMAN | 13 kDa |
| 40S ribosomal protein S27-like OS=Homo sapiens OX=9606 GN=RPS27L PE=1 SV=3 | Q71UM5 | RS27L_HUMAN | 9 kDa |
| 40S ribosomal protein S28 OS=Homo sapiens OX=9606 GN=RPS28 PE=1 SV=1 | P62857 | RS28_HUMAN | 8 kDa |
| 40S ribosomal protein S3 OS=Homo sapiens OX=9606 GN=RPS3 PE=1 SV=2 | P23396 | RS3_HUMAN | 27 kDa |
| 40S ribosomal protein S3a OS=Homo sapiens OX=9606 GN=RPS3A PE=1 SV=2 | P61247 | RS3A_HUMAN | 30 kDa |
| 40S ribosomal protein S4, X isoform OS=Homo sapiens OX=9606 GN=RPS4X PE=1 SV=2 | P62701 | RS4X_HUMAN | 30 kDa |
| 40S ribosomal protein S5 OS=Homo sapiens OX=9606 GN=RPS5 PE=1 SV=4 | P46782 | RS5_HUMAN | 23 kDa |
| 40S ribosomal protein S6 OS=Homo sapiens OX=9606 GN=RPS6 PE=1 SV=1 | P62753 | RS6_HUMAN | 29 kDa |
| 40S ribosomal protein S8 OS=Homo sapiens OX=9606 GN=RPS8 PE=1 SV=2 | P62241 | RS8_HUMAN | 24 kDa |
| 40S ribosomal protein S9 OS=Homo sapiens OX=9606 GN=RPS9 PE=1 SV=3 | P46781 | RS9_HUMAN | 23 kDa |
| 40S ribosomal protein SA OS=Homo sapiens OX=9606 GN=RPSA PE=1 SV=4 | P08865 | RSSA_HUMAN | 33 kDa |
| 4F2 cell-surface antigen heavy chain OS=Homo sapiens OX=9606 GN=SLC3A2 PE=1 SV=3 | P08195 | 4F2_HUMAN | 68 kDa |
| 4-trimethylaminobutyraldehyde dehydrogenase OS=Homo sapiens OX=9606 GN=ALDH9A1 PE=1 SV=3 | P49189 | AL9A1_HUMAN | 54 kDa |
| 5'-3' exonuclease PLD3 OS=Homo sapiens OX=9606 GN=PLD3 PE=1 SV=1 | Q8IV08 | PLD3_HUMAN | 55 kDa |
| 5'-AMP-activated protein kinase subunit gamma-1 OS=Homo sapiens OX=9606 GN=PRKAG1 PE=1 SV=1 | P54619 | AAKG1_HUMAN | 38 kDa |
| 60 kDa heat shock protein, mitochondrial OS=Homo sapiens OX=9606 GN=HSPD1 PE=1 SV=2 | P10809 | CH60_HUMAN | 61 kDa |
| 60S acidic ribosomal protein P0-like OS=Homo sapiens OX=9606 GN=RPLP0P6 PE=5 SV=1 | Q8NHW5 | RLA0L_HUMAN | 34 kDa |
| 60S acidic ribosomal protein P2 OS=Homo sapiens OX=9606 GN=RPLP2 PE=1 SV=1 | P05387 | RLA2_HUMAN | 12 kDa |
| 60S ribosomal protein L10 OS=Homo sapiens OX=9606 GN=RPL10 PE=1 SV=5 | P27635 | RL10_HUMAN | 25 kDa |
| 60S ribosomal protein L10a OS=Homo sapiens OX=9606 GN=RPL10A PE=1 SV=2 | P62906 | RL10A_HUMAN | 25 kDa |
| 60S ribosomal protein L11 OS=Homo sapiens OX=9606 GN=RPL11 PE=1 SV=2 | P62913 | RL11_HUMAN | 20 kDa |
| 60S ribosomal protein L12 OS=Homo sapiens OX=9606 GN=RPL12 PE=1 SV=1 | P30050 | RL12_HUMAN | 18 kDa |
| 60S ribosomal protein L13a OS=Homo sapiens OX=9606 GN=RPL13A PE=1 SV=2 | P40429 | RL13A_HUMAN | 24 kDa |
| 60S ribosomal protein L14 OS=Homo sapiens OX=9606 GN=RPL14 PE=1 SV=4 | P50914 | RL14_HUMAN | 23 kDa |
| 60S ribosomal protein L15 OS=Homo sapiens OX=9606 GN=RPL15 PE=1 SV=2 | P61313 | RL15_HUMAN | 24 kDa |
| 60S ribosomal protein L17 OS=Homo sapiens OX=9606 GN=RPL17 PE=1 SV=3 | P18621 | RL17_HUMAN | 21 kDa |
| 60S ribosomal protein L18 OS=Homo sapiens OX=9606 GN=RPL18 PE=1 SV=2 | Q07020 | RL18_HUMAN | 22 kDa |
| 60S ribosomal protein L19 OS=Homo sapiens OX=9606 GN=RPL19 PE=1 SV=1 | P84098 | RL19_HUMAN | 23 kDa |
| 60S ribosomal protein L22 OS=Homo sapiens OX=9606 GN=RPL22 PE=1 SV=2 | P35268 | RL22_HUMAN | 15 kDa |
| 60S ribosomal protein L24 OS=Homo sapiens OX=9606 GN=RPL24 PE=1 SV=1 | P83731 | RL24_HUMAN | 18 kDa |
| 60S ribosomal protein L27a OS=Homo sapiens OX=9606 GN=RPL27A PE=1 SV=2 | P46776 | RL27A_HUMAN | 17 kDa |
| 60S ribosomal protein L28 OS=Homo sapiens OX=9606 GN=RPL28 PE=1 SV=3 | P46779 | RL28_HUMAN | 16 kDa |
| 60S ribosomal protein L3 OS=Homo sapiens OX=9606 GN=RPL3 PE=1 SV=2 | P39023 | RL3_HUMAN | 46 kDa |
| 60S ribosomal protein L4 OS=Homo sapiens OX=9606 GN=RPL4 PE=1 SV=5 | P36578 | RL4_HUMAN | 48 kDa |
| 60S ribosomal protein L5 OS=Homo sapiens OX=9606 GN=RPL5 PE=1 SV=3 | P46777 | RL5_HUMAN | 34 kDa |
| 60S ribosomal protein L7 OS=Homo sapiens OX=9606 GN=RPL7 PE=1 SV=1 | P18124 | RL7_HUMAN | 29 kDa |
| 60S ribosomal protein L7a OS=Homo sapiens OX=9606 GN=RPL7A PE=1 SV=2 | P62424 | RL7A_HUMAN | 30 kDa |
| 60S ribosomal protein L8 OS=Homo sapiens OX=9606 GN=RPL8 PE=1 SV=2 | P62917 | RL8_HUMAN | 28 kDa |
| 60S ribosomal protein L9 OS=Homo sapiens OX=9606 GN=RPL9P9 PE=1 SV=1 | P32969 | RL9_HUMAN | 22 kDa |
| 6-phosphogluconate dehydrogenase, decarboxylating OS=Homo sapiens OX=9606 GN=PGD PE=1 SV=3 | P52209 | 6PGD_HUMAN | 53 kDa |
| 6-phosphogluconolactonase OS=Homo sapiens OX=9606 GN=PGLS PE=1 SV=2 | O95336 | 6PGL_HUMAN | 28 kDa |
| Abl interactor 1 OS=Homo sapiens OX=9606 GN=ABI1 PE=1 SV=4 | Q8IZP0 | ABI1_HUMAN | 55 kDa |
| Acetoacetyl-CoA synthetase OS=Homo sapiens OX=9606 GN=AACS PE=1 SV=1 | Q86V21 | AACS_HUMAN | 75 kDa |
| Acetyl-CoA acetyltransferase, cytosolic OS=Homo sapiens OX=9606 GN=ACAT2 PE=1 SV=2 | Q9BWD1 | THIC_HUMAN | 41 kDa |
| Acetyl-CoA carboxylase 1 OS=Homo sapiens OX=9606 GN=ACACA PE=1 SV=2 | Q13085 | ACACA_HUMAN | 266 kDa |
| Acetyl-coenzyme A synthetase, cytoplasmic OS=Homo sapiens OX=9606 GN=ACSS2 PE=1 SV=1 | Q9NR19 | ACSA_HUMAN | 79 kDa |
| Actin-related protein 2 OS=Homo sapiens OX=9606 GN=ACTR2 PE=1 SV=1 | P61160 | ARP2_HUMAN | 45 kDa |
| Actin-related protein 2/3 complex subunit 1A OS=Homo sapiens OX=9606 GN=ARPC1A PE=1 SV=2 | Q92747 | ARC1A_HUMAN | 42 kDa |
| Actin-related protein 2/3 complex subunit 1B OS=Homo sapiens OX=9606 GN=ARPC1B PE=1 SV=3 | O15143 | ARC1B_HUMAN | 41 kDa |
| Actin-related protein 2/3 complex subunit 2 OS=Homo sapiens OX=9606 GN=ARPC2 PE=1 SV=1 | O15144 | ARPC2_HUMAN | 34 kDa |
| Actin-related protein 2/3 complex subunit 3 OS=Homo sapiens OX=9606 GN=ARPC3 PE=1 SV=3 | O15145 | ARPC3_HUMAN | 21 kDa |
| Actin-related protein 2/3 complex subunit 4 OS=Homo sapiens OX=9606 GN=ARPC4 PE=1 SV=3 | P59998 | ARPC4_HUMAN | 20 kDa |
| Actin-related protein 2/3 complex subunit 5 OS=Homo sapiens OX=9606 GN=ARPC5 PE=1 SV=3 | O15511 | ARPC5_HUMAN | 16 kDa |
| Actin-related protein 2/3 complex subunit 5-like protein OS=Homo sapiens OX=9606 GN=ARPC5L PE=1 SV=1 | Q9BPX5 | ARP5L_HUMAN | 17 kDa |
| Actin-related protein 3 OS=Homo sapiens OX=9606 GN=ACTR3 PE=1 SV=3 | P61158 | ARP3_HUMAN | 47 kDa |
| Activator of 90 kDa heat shock protein ATPase homolog 1 OS=Homo sapiens OX=9606 GN=AHSA1 PE=1 SV=1 | O95433 | AHSA1_HUMAN | 38 kDa |
| Acylamino-acid-releasing enzyme OS=Homo sapiens OX=9606 GN=APEH PE=1 SV=4 | P13798 | ACPH_HUMAN | 81 kDa |
| Acyl-coenzyme A thioesterase 8 OS=Homo sapiens OX=9606 GN=ACOT8 PE=1 SV=1 | O14734 | ACOT8_HUMAN | 36 kDa |
| Acyl-protein thioesterase 2 OS=Homo sapiens OX=9606 GN=LYPLA2 PE=1 SV=1 | O95372 | LYPA2_HUMAN | 25 kDa |
| Adenine phosphoribosyltransferase OS=Homo sapiens OX=9606 GN=APRT PE=1 SV=2 | P07741 | APT_HUMAN | 20 kDa |
| Adenomatous polyposis coli protein OS=Homo sapiens OX=9606 GN=APC PE=1 SV=2 | P25054 | APC_HUMAN | 312 kDa |
| Adenosylhomocysteinase OS=Homo sapiens OX=9606 GN=AHCY PE=1 SV=4 | P23526 | SAHH_HUMAN | 48 kDa |
| Adenylate cyclase type 9 OS=Homo sapiens OX=9606 GN=ADCY9 PE=1 SV=4 | O60503 | ADCY9_HUMAN | 151 kDa |
| Adenylosuccinate lyase OS=Homo sapiens OX=9606 GN=ADSL PE=1 SV=2 | P30566 | PUR8_HUMAN | 55 kDa |
| Adenylyl cyclase-associated protein 1 OS=Homo sapiens OX=9606 GN=CAP1 PE=1 SV=5 | Q01518 | CAP1_HUMAN | 52 kDa |
| Adhesion G protein-coupled receptor L2 OS=Homo sapiens OX=9606 GN=ADGRL2 PE=1 SV=2 | O95490 | AGRL2_HUMAN | 163 kDa |
| Adhesion G-protein coupled receptor G1 OS=Homo sapiens OX=9606 GN=ADGRG1 PE=1 SV=2 | Q9Y653 | AGRG1_HUMAN | 78 kDa |
| ADP/ATP translocase 2 OS=Homo sapiens OX=9606 GN=SLC25A5 PE=1 SV=7 | P05141 | ADT2_HUMAN | 33 kDa |
| ADP-ribosylation factor 6 OS=Homo sapiens OX=9606 GN=ARF6 PE=1 SV=2 | P62330 | ARF6_HUMAN | 20 kDa |
| ADP-ribosylation factor-like protein 15 OS=Homo sapiens OX=9606 GN=ARL15 PE=1 SV=1 | Q9NXU5 | ARL15_HUMAN | 23 kDa |
| ADP-ribosylation factor-like protein 3 OS=Homo sapiens OX=9606 GN=ARL3 PE=1 SV=2 | P36405 | ARL3_HUMAN | 20 kDa |
| ADP-sugar pyrophosphatase OS=Homo sapiens OX=9606 GN=NUDT5 PE=1 SV=1 | Q9UKK9 | NUDT5_HUMAN | 24 kDa |
| Afamin OS=Homo sapiens OX=9606 GN=AFM PE=1 SV=1 | P43652 | AFAM_HUMAN | 69 kDa |
| Aflatoxin B1 aldehyde reductase member 2 OS=Homo sapiens OX=9606 GN=AKR7A2 PE=1 SV=3 | O43488 | ARK72_HUMAN | 40 kDa |
| Aggrecan core protein OS=Homo sapiens OX=9606 GN=ACAN PE=1 SV=3 | P16112 | PGCA_HUMAN | 261 kDa |
| A-kinase anchor protein 7 isoforms alpha and beta OS=Homo sapiens OX=9606 GN=AKAP7 PE=1 SV=4 | O43687 | AKA7A_HUMAN (+1) | 11 kDa |
| AKT-interacting protein OS=Homo sapiens OX=9606 GN=AKTIP PE=1 SV=1 | Q9H8T0 | AKTIP_HUMAN | 33 kDa |
| Alanine--tRNA ligase, cytoplasmic OS=Homo sapiens OX=9606 GN=AARS1 PE=1 SV=2 | P49588 | SYAC_HUMAN | 107 kDa |
| Alcohol dehydrogenase class-3 OS=Homo sapiens OX=9606 GN=ADH5 PE=1 SV=4 | P11766 | ADHX_HUMAN | 40 kDa |
| Aldehyde dehydrogenase family 16 member A1 OS=Homo sapiens OX=9606 GN=ALDH16A1 PE=1 SV=2 | Q8IZ83 | A16A1_HUMAN | 85 kDa |
| Aldehyde dehydrogenase family 3 member A2 OS=Homo sapiens OX=9606 GN=ALDH3A2 PE=1 SV=1 | P51648 | AL3A2_HUMAN | 55 kDa |
| Aldo-keto reductase family 1 member A1 OS=Homo sapiens OX=9606 GN=AKR1A1 PE=1 SV=3 | P14550 | AK1A1_HUMAN | 37 kDa |
| Aldo-keto reductase family 1 member B1 OS=Homo sapiens OX=9606 GN=AKR1B1 PE=1 SV=3 | P15121 | ALDR_HUMAN | 36 kDa |
| Alpha-1-acid glycoprotein 1 OS=Homo sapiens OX=9606 GN=ORM1 PE=1 SV=2 | P02763 | A1AG1_HUMAN | 24 kDa |
| Alpha-1-antichymotrypsin OS=Homo sapiens OX=9606 GN=SERPINA3 PE=1 SV=2 | P01011 | AACT_HUMAN | 48 kDa |
| Alpha-1-antitrypsin OS=Homo sapiens OX=9606 GN=SERPINA1 PE=1 SV=3 | P01009 | A1AT_HUMAN | 47 kDa |
| Alpha-1B-glycoprotein OS=Homo sapiens OX=9606 GN=A1BG PE=1 SV=4 | P04217 | A1BG_HUMAN | 54 kDa |
| Alpha-2-antiplasmin OS=Homo sapiens OX=9606 GN=SERPINF2 PE=1 SV=3 | P08697 | A2AP_HUMAN | 55 kDa |
| Alpha-2-HS-glycoprotein OS=Homo sapiens OX=9606 GN=AHSG PE=1 SV=2 | P02765 | FETUA_HUMAN | 39 kDa |
| Alpha-2-macroglobulin OS=Homo sapiens OX=9606 GN=A2M PE=1 SV=3 | P01023 | A2MG_HUMAN | 163 kDa |
| Alpha-2-macroglobulin-like protein 1 OS=Homo sapiens OX=9606 GN=A2ML1 PE=1 SV=3 | A8K2U0 | A2ML1_HUMAN | 161 kDa |
| Alpha-aminoadipic semialdehyde dehydrogenase OS=Homo sapiens OX=9606 GN=ALDH7A1 PE=1 SV=5 | P49419 | AL7A1_HUMAN | 58 kDa |
| Alpha-centractin OS=Homo sapiens OX=9606 GN=ACTR1A PE=1 SV=1 | P61163 | ACTZ_HUMAN | 43 kDa |
| Alpha-crystallin B chain OS=Homo sapiens OX=9606 GN=CRYAB PE=1 SV=2 | P02511 | CRYAB_HUMAN | 20 kDa |
| Alpha-enolase OS=Homo sapiens OX=9606 GN=ENO1 PE=1 SV=2 | P06733 | ENOA_HUMAN | 47 kDa |
| Alpha-parvin OS=Homo sapiens OX=9606 GN=PARVA PE=1 SV=1 | Q9NVD7 | PARVA_HUMAN | 42 kDa |
| Alpha-S1-casein OS=Bos taurus OX=9913 GN=CSN1S1 PE=1 SV=2 | P02662 | CASA1_BOVIN | 25 kDa |
| Alpha-soluble NSF attachment protein OS=Homo sapiens OX=9606 GN=NAPA PE=1 SV=3 | P54920 | SNAA_HUMAN | 33 kDa |
| Aminoacyl tRNA synthase complex-interacting multifunctional protein 2 OS=Homo sapiens OX=9606 GN=AIMP2 PE=1 SV=2 | Q13155 | AIMP2_HUMAN | 35 kDa |
| Aminopeptidase B OS=Homo sapiens OX=9606 GN=RNPEP PE=1 SV=2 | Q9H4A4 | AMPB_HUMAN | 73 kDa |
| Angiotensin-converting enzyme OS=Homo sapiens OX=9606 GN=ACE PE=1 SV=1 | P12821 | ACE_HUMAN | 150 kDa |
| Angiotensinogen OS=Homo sapiens OX=9606 GN=AGT PE=1 SV=3 | P01019 | ANGT_HUMAN | 53 kDa |
| Anion exchange protein 2 OS=Homo sapiens OX=9606 GN=SLC4A2 PE=1 SV=4 | P04920 | B3A2_HUMAN | 137 kDa |
| Ankycorbin OS=Homo sapiens OX=9606 GN=RAI14 PE=1 SV=2 | Q9P0K7 | RAI14_HUMAN | 110 kDa |
| Ankyrin-2 OS=Homo sapiens OX=9606 GN=ANK2 PE=1 SV=4 | Q01484 | ANK2_HUMAN | 434 kDa |
| Annexin A1 OS=Homo sapiens OX=9606 GN=ANXA1 PE=1 SV=2 | P04083 | ANXA1_HUMAN | 39 kDa |
| Annexin A11 OS=Homo sapiens OX=9606 GN=ANXA11 PE=1 SV=1 | P50995 | ANX11_HUMAN | 54 kDa |
| Annexin A2 OS=Homo sapiens OX=9606 GN=ANXA2 PE=1 SV=2 | P07355 | ANXA2_HUMAN | 39 kDa |
| Annexin A4 OS=Homo sapiens OX=9606 GN=ANXA4 PE=1 SV=4 | P09525 | ANXA4_HUMAN | 36 kDa |
| Annexin A5 OS=Homo sapiens OX=9606 GN=ANXA5 PE=1 SV=2 | P08758 | ANXA5_HUMAN | 36 kDa |
| Annexin A6 OS=Homo sapiens OX=9606 GN=ANXA6 PE=1 SV=3 | P08133 | ANXA6_HUMAN | 76 kDa |
| Annexin A7 OS=Homo sapiens OX=9606 GN=ANXA7 PE=1 SV=3 | P20073 | ANXA7_HUMAN | 53 kDa |
| Anoctamin-6 OS=Homo sapiens OX=9606 GN=ANO6 PE=1 SV=2 | Q4KMQ2 | ANO6_HUMAN | 106 kDa |
| Antithrombin-III OS=Homo sapiens OX=9606 GN=SERPINC1 PE=1 SV=1 | P01008 | ANT3_HUMAN | 53 kDa |
| AP-1 complex subunit gamma-1 OS=Homo sapiens OX=9606 GN=AP1G1 PE=1 SV=5 | O43747 | AP1G1_HUMAN | 91 kDa |
| AP-2 complex subunit alpha-1 OS=Homo sapiens OX=9606 GN=AP2A1 PE=1 SV=3 | O95782 | AP2A1_HUMAN | 108 kDa |
| AP-2 complex subunit alpha-2 OS=Homo sapiens OX=9606 GN=AP2A2 PE=1 SV=2 | O94973 | AP2A2_HUMAN | 104 kDa |
| AP-2 complex subunit beta OS=Homo sapiens OX=9606 GN=AP2B1 PE=1 SV=1 | P63010 | AP2B1_HUMAN | 105 kDa |
| AP-2 complex subunit mu OS=Homo sapiens OX=9606 GN=AP2M1 PE=1 SV=2 | Q96CW1 | AP2M1_HUMAN | 50 kDa |
| Apolipoprotein A-I OS=Homo sapiens OX=9606 GN=APOA1 PE=1 SV=1 | P02647 | APOA1_HUMAN | 31 kDa |
| Apolipoprotein A-II OS=Homo sapiens OX=9606 GN=APOA2 PE=1 SV=1 | P02652 | APOA2_HUMAN | 11 kDa |
| Apolipoprotein A-IV OS=Homo sapiens OX=9606 GN=APOA4 PE=1 SV=4 | P06727 | APOA4_HUMAN | 45 kDa |
| Apolipoprotein A-V OS=Homo sapiens OX=9606 GN=APOA5 PE=1 SV=1 | Q6Q788 | APOA5_HUMAN | 41 kDa |
| Apolipoprotein C-I OS=Homo sapiens OX=9606 GN=APOC1 PE=1 SV=1 | P02654 | APOC1_HUMAN | 9 kDa |
| Apolipoprotein C-III OS=Homo sapiens OX=9606 GN=APOC3 PE=1 SV=1 | P02656 | APOC3_HUMAN | 11 kDa |
| Apolipoprotein C-IV OS=Homo sapiens OX=9606 GN=APOC4 PE=1 SV=1 | P55056 | APOC4_HUMAN | 15 kDa |
| Apolipoprotein E OS=Homo sapiens OX=9606 GN=APOE PE=1 SV=1 | P02649 | APOE_HUMAN | 36 kDa |
| Apolipoprotein F OS=Homo sapiens OX=9606 GN=APOF PE=1 SV=2 | Q13790 | APOF_HUMAN | 35 kDa |
| Apolipoprotein L1 OS=Homo sapiens OX=9606 GN=APOL1 PE=1 SV=5 | O14791 | APOL1_HUMAN | 44 kDa |
| Apoptosis regulator BAX OS=Homo sapiens OX=9606 GN=BAX PE=1 SV=1 | Q07812 | BAX_HUMAN | 21 kDa |
| Arf-GAP with GTPase, ANK repeat and PH domain-containing protein 3 OS=Homo sapiens OX=9606 GN=AGAP3 PE=1 SV=2 | Q96P47 | AGAP3_HUMAN | 95 kDa |
| Arginine--tRNA ligase, cytoplasmic OS=Homo sapiens OX=9606 GN=RARS1 PE=1 SV=2 | P54136 | SYRC_HUMAN | 75 kDa |
| Argininosuccinate synthase OS=Homo sapiens OX=9606 GN=ASS1 PE=1 SV=2 | P00966 | ASSY_HUMAN | 47 kDa |
| Aromatase OS=Homo sapiens OX=9606 GN=CYP19A1 PE=1 SV=3 | P11511 | CP19A_HUMAN | 58 kDa |
| Arrestin domain-containing protein 1 OS=Homo sapiens OX=9606 GN=ARRDC1 PE=1 SV=1 | Q8N5I2 | ARRD1_HUMAN | 46 kDa |
| Arsenite methyltransferase OS=Homo sapiens OX=9606 GN=AS3MT PE=1 SV=3 | Q9HBK9 | AS3MT_HUMAN | 42 kDa |
| Asparagine--tRNA ligase, cytoplasmic OS=Homo sapiens OX=9606 GN=NARS1 PE=1 SV=1 | O43776 | SYNC_HUMAN | 63 kDa |
| Aspartate aminotransferase, cytoplasmic OS=Homo sapiens OX=9606 GN=GOT1 PE=1 SV=3 | P17174 | AATC_HUMAN | 46 kDa |
| Aspartate--tRNA ligase, cytoplasmic OS=Homo sapiens OX=9606 GN=DARS1 PE=1 SV=2 | P14868 | SYDC_HUMAN | 57 kDa |
| Ataxin-10 OS=Homo sapiens OX=9606 GN=ATXN10 PE=1 SV=1 | Q9UBB4 | ATX10_HUMAN | 53 kDa |
| ATP synthase subunit alpha, mitochondrial OS=Homo sapiens OX=9606 GN=ATP5F1A PE=1 SV=1 | P25705 | ATPA_HUMAN | 60 kDa |
| ATP synthase subunit beta, mitochondrial OS=Homo sapiens OX=9606 GN=ATP5F1B PE=1 SV=3 | P06576 | ATPB_HUMAN | 57 kDa |
| ATPase GET3 OS=Homo sapiens OX=9606 GN=GET3 PE=1 SV=2 | O43681 | GET3_HUMAN | 39 kDa |
| ATP-binding cassette sub-family E member 1 OS=Homo sapiens OX=9606 GN=ABCE1 PE=1 SV=1 | P61221 | ABCE1_HUMAN | 67 kDa |
| ATP-binding cassette sub-family F member 3 OS=Homo sapiens OX=9606 GN=ABCF3 PE=1 SV=2 | Q9NUQ8 | ABCF3_HUMAN | 80 kDa |
| ATP-citrate synthase OS=Homo sapiens OX=9606 GN=ACLY PE=1 SV=3 | P53396 | ACLY_HUMAN | 121 kDa |
| ATP-dependent 6-phosphofructokinase, liver type OS=Homo sapiens OX=9606 GN=PFKL PE=1 SV=6 | P17858 | PFKAL_HUMAN | 85 kDa |
| ATP-dependent 6-phosphofructokinase, muscle type OS=Homo sapiens OX=9606 GN=PFKM PE=1 SV=2 | P08237 | PFKAM_HUMAN | 85 kDa |
| ATP-dependent 6-phosphofructokinase, platelet type OS=Homo sapiens OX=9606 GN=PFKP PE=1 SV=2 | Q01813 | PFKAP_HUMAN | 86 kDa |
| ATP-dependent RNA helicase DDX19A OS=Homo sapiens OX=9606 GN=DDX19A PE=1 SV=1 | Q9NUU7 | DD19A_HUMAN (+1) | 54 kDa |
| ATP-dependent RNA helicase DDX3X OS=Homo sapiens OX=9606 GN=DDX3X PE=1 SV=3 | O00571 | DDX3X_HUMAN | 73 kDa |
| Atrial natriuretic peptide receptor 2 OS=Homo sapiens OX=9606 GN=NPR2 PE=1 SV=1 | P20594 | ANPRB_HUMAN | 117 kDa |
| Atrial natriuretic peptide receptor 3 OS=Homo sapiens OX=9606 GN=NPR3 PE=1 SV=2 | P17342 | ANPRC_HUMAN | 60 kDa |
| Attractin OS=Homo sapiens OX=9606 GN=ATRN PE=1 SV=2 | O75882 | ATRN_HUMAN | 159 kDa |
| Axin interactor, dorsalization-associated protein OS=Homo sapiens OX=9606 GN=AIDA PE=1 SV=1 | Q96BJ3 | AIDA_HUMAN | 35 kDa |
| BAG family molecular chaperone regulator 2 OS=Homo sapiens OX=9606 GN=BAG2 PE=1 SV=1 | O95816 | BAG2_HUMAN | 24 kDa |
| Band 3 anion transport protein OS=Homo sapiens OX=9606 GN=SLC4A1 PE=1 SV=3 | P02730 | B3AT_HUMAN | 102 kDa |
| Band 4.1-like protein 2 OS=Homo sapiens OX=9606 GN=EPB41L2 PE=1 SV=1 | O43491 | E41L2_HUMAN | 113 kDa |
| Basement membrane-specific heparan sulfate proteoglycan core protein OS=Homo sapiens OX=9606 GN=HSPG2 PE=1 SV=4 | P98160 | PGBM_HUMAN | 469 kDa |
| Basigin OS=Homo sapiens OX=9606 GN=BSG PE=1 SV=2 | P35613 | BASI_HUMAN | 42 kDa |
| Beta-2-glycoprotein 1 OS=Homo sapiens OX=9606 GN=APOH PE=1 SV=3 | P02749 | APOH_HUMAN | 38 kDa |
| Beta-2-microglobulin OS=Homo sapiens OX=9606 GN=B2M PE=1 SV=1 | P61769 | B2MG_HUMAN | 14 kDa |
| Beta-2-syntrophin OS=Homo sapiens OX=9606 GN=SNTB2 PE=1 SV=1 | Q13425 | SNTB2_HUMAN | 58 kDa |
| Beta-centractin OS=Homo sapiens OX=9606 GN=ACTR1B PE=1 SV=1 | P42025 | ACTY_HUMAN | 42 kDa |
| Beta-enolase OS=Homo sapiens OX=9606 GN=ENO3 PE=1 SV=5 | P13929 | ENOB_HUMAN | 47 kDa |
| Bifunctional 3'-phosphoadenosine 5'-phosphosulfate synthase 2 OS=Homo sapiens OX=9606 GN=PAPSS2 PE=1 SV=2 | O95340 | PAPS2_HUMAN | 70 kDa |
| Bifunctional glutamate/proline--tRNA ligase OS=Homo sapiens OX=9606 GN=EPRS1 PE=1 SV=5 | P07814 | SYEP_HUMAN | 171 kDa |
| Bifunctional phosphoribosylaminoimidazole carboxylase/phosphoribosylaminoimidazole succinocarboxamide synthetase OS=Homo sapiens OX=9606 GN=PAICS PE=1 SV=3 | P22234 | PUR6_HUMAN | 47 kDa |
| Bifunctional purine biosynthesis protein ATIC OS=Homo sapiens OX=9606 GN=ATIC PE=1 SV=3 | P31939 | PUR9_HUMAN | 65 kDa |
| Bifunctional UDP-N-acetylglucosamine 2-epimerase/N-acetylmannosamine kinase OS=Homo sapiens OX=9606 GN=GNE PE=1 SV=1 | Q9Y223 | GLCNE_HUMAN | 79 kDa |
| Biliverdin reductase A OS=Homo sapiens OX=9606 GN=BLVRA PE=1 SV=2 | P53004 | BIEA_HUMAN | 33 kDa |
| Bleomycin hydrolase OS=Homo sapiens OX=9606 GN=BLMH PE=1 SV=1 | Q13867 | BLMH_HUMAN | 53 kDa |
| BolA-like protein 2 OS=Homo sapiens OX=9606 GN=BOLA2B PE=1 SV=1 | Q9H3K6 | BOLA2_HUMAN | 10 kDa |
| Bone marrow stromal antigen 2 OS=Homo sapiens OX=9606 GN=BST2 PE=1 SV=1 | Q10589 | BST2_HUMAN | 20 kDa |
| Bone morphogenetic protein receptor type-2 OS=Homo sapiens OX=9606 GN=BMPR2 PE=1 SV=2 | Q13873 | BMPR2_HUMAN | 115 kDa |
| Brain acid soluble protein 1 OS=Homo sapiens OX=9606 GN=BASP1 PE=1 SV=2 | P80723 | BASP1_HUMAN | 23 kDa |
| BRO1 domain-containing protein BROX OS=Homo sapiens OX=9606 GN=BROX PE=1 SV=1 | Q5VW32 | BROX_HUMAN | 46 kDa |
| Butyrophilin subfamily 2 member A1 OS=Homo sapiens OX=9606 GN=BTN2A1 PE=1 SV=3 | Q7KYR7 | BT2A1_HUMAN | 60 kDa |
| C-1-tetrahydrofolate synthase, cytoplasmic OS=Homo sapiens OX=9606 GN=MTHFD1 PE=1 SV=4 | P11586 | C1TC_HUMAN | 102 kDa |
| C4b-binding protein alpha chain OS=Homo sapiens OX=9606 GN=C4BPA PE=1 SV=2 | P04003 | C4BPA_HUMAN | 67 kDa |
| C4b-binding protein beta chain OS=Homo sapiens OX=9606 GN=C4BPB PE=1 SV=1 | P20851 | C4BPB_HUMAN | 28 kDa |
| CAD protein OS=Homo sapiens OX=9606 GN=CAD PE=1 SV=3 | P27708 | PYR1_HUMAN | 243 kDa |
| Cadherin-11 OS=Homo sapiens OX=9606 GN=CDH11 PE=1 SV=2 | P55287 | CAD11_HUMAN | 88 kDa |
| Cadherin-2 OS=Homo sapiens OX=9606 GN=CDH2 PE=1 SV=4 | P19022 | CADH2_HUMAN | 100 kDa |
| Calcineurin B homologous protein 1 OS=Homo sapiens OX=9606 GN=CHP1 PE=1 SV=3 | Q99653 | CHP1_HUMAN | 22 kDa |
| Calcyclin-binding protein OS=Homo sapiens OX=9606 GN=CACYBP PE=1 SV=2 | Q9HB71 | CYBP_HUMAN | 26 kDa |
| Caldesmon OS=Homo sapiens OX=9606 GN=CALD1 PE=1 SV=3 | Q05682 | CALD1_HUMAN | 93 kDa |
| Calmodulin-1 OS=Homo sapiens OX=9606 GN=CALM1 PE=1 SV=1 | P0DP23 | CALM1_HUMAN (+2) | 17 kDa |
| Calmodulin-like protein 3 OS=Homo sapiens OX=9606 GN=CALML3 PE=1 SV=2 | P27482 | CALL3_HUMAN | 17 kDa |
| Calmodulin-like protein 5 OS=Homo sapiens OX=9606 GN=CALML5 PE=1 SV=2 | Q9NZT1 | CALL5_HUMAN | 16 kDa |
| Calnexin OS=Homo sapiens OX=9606 GN=CANX PE=1 SV=2 | P27824 | CALX_HUMAN | 68 kDa |
| Calpain small subunit 1 OS=Homo sapiens OX=9606 GN=CAPNS1 PE=1 SV=1 | P04632 | CPNS1_HUMAN | 28 kDa |
| Calponin-3 OS=Homo sapiens OX=9606 GN=CNN3 PE=1 SV=1 | Q15417 | CNN3_HUMAN | 36 kDa |
| Calreticulin OS=Homo sapiens OX=9606 GN=CALR PE=1 SV=1 | P27797 | CALR_HUMAN | 48 kDa |
| Calretinin OS=Homo sapiens OX=9606 GN=CALB2 PE=2 SV=2 | P22676 | CALB2_HUMAN | 32 kDa |
| Calumenin OS=Homo sapiens OX=9606 GN=CALU PE=1 SV=2 | O43852 | CALU_HUMAN | 37 kDa |
| cAMP-dependent protein kinase catalytic subunit alpha OS=Homo sapiens OX=9606 GN=PRKACA PE=1 SV=2 | P17612 | KAPCA_HUMAN | 41 kDa |
| cAMP-dependent protein kinase type I-alpha regulatory subunit OS=Homo sapiens OX=9606 GN=PRKAR1A PE=1 SV=1 | P10644 | KAP0_HUMAN | 43 kDa |
| cAMP-dependent protein kinase type II-alpha regulatory subunit OS=Homo sapiens OX=9606 GN=PRKAR2A PE=1 SV=2 | P13861 | KAP2_HUMAN | 46 kDa |
| cAMP-dependent protein kinase type II-beta regulatory subunit OS=Homo sapiens OX=9606 GN=PRKAR2B PE=1 SV=3 | P31323 | KAP3_HUMAN | 46 kDa |
| Carbonyl reductase [NADPH] 3 OS=Homo sapiens OX=9606 GN=CBR3 PE=1 SV=3 | O75828 | CBR3_HUMAN (+1) | 31 kDa |
| Carboxypeptidase B2 OS=Homo sapiens OX=9606 GN=CPB2 PE=1 SV=2 | Q96IY4 | CBPB2_HUMAN | 48 kDa |
| Carcinoembryonic antigen-related cell adhesion molecule 5 OS=Homo sapiens OX=9606 GN=CEACAM5 PE=1 SV=4 | P06731 | CEAM5_HUMAN (+2) | 77 kDa |
| Cartilage intermediate layer protein 1 OS=Homo sapiens OX=9606 GN=CILP PE=1 SV=4 | O75339 | CILP1_HUMAN | 133 kDa |
| Casein kinase II subunit beta OS=Homo sapiens OX=9606 GN=CSNK2B PE=1 SV=1 | P67870 | CSK2B_HUMAN | 25 kDa |
| Caspase-9 OS=Homo sapiens OX=9606 GN=CASP9 PE=1 SV=3 | P55211 | CASP9_HUMAN | 46 kDa |
| Catalase OS=Homo sapiens OX=9606 GN=CAT PE=1 SV=3 | P04040 | CATA_HUMAN | 60 kDa |
| Catenin alpha-1 OS=Homo sapiens OX=9606 GN=CTNNA1 PE=1 SV=1 | P35221 | CTNA1_HUMAN | 100 kDa |
| Catenin beta-1 OS=Homo sapiens OX=9606 GN=CTNNB1 PE=1 SV=1 | P35222 | CTNB1_HUMAN | 85 kDa |
| Catenin delta-1 OS=Homo sapiens OX=9606 GN=CTNND1 PE=1 SV=1 | O60716 | CTND1_HUMAN | 108 kDa |
| Cathepsin D OS=Homo sapiens OX=9606 GN=CTSD PE=1 SV=1 | P07339 | CATD_HUMAN | 45 kDa |
| Cation-dependent mannose-6-phosphate receptor OS=Homo sapiens OX=9606 GN=M6PR PE=1 SV=1 | P20645 | MPRD_HUMAN | 31 kDa |
| CD166 antigen OS=Homo sapiens OX=9606 GN=ALCAM PE=1 SV=2 | Q13740 | CD166_HUMAN | 65 kDa |
| CD276 antigen OS=Homo sapiens OX=9606 GN=CD276 PE=1 SV=1 | Q5ZPR3 | CD276_HUMAN | 57 kDa |
| CD5 antigen-like OS=Homo sapiens OX=9606 GN=CD5L PE=1 SV=1 | O43866 | CD5L_HUMAN | 38 kDa |
| CD63 antigen OS=Homo sapiens OX=9606 GN=CD63 PE=1 SV=2 | P08962 | CD63_HUMAN | 26 kDa |
| CD81 antigen OS=Homo sapiens OX=9606 GN=CD81 PE=1 SV=1 | P60033 | CD81_HUMAN | 26 kDa |
| CD9 antigen OS=Homo sapiens OX=9606 GN=CD9 PE=1 SV=4 | P21926 | CD9_HUMAN | 25 kDa |
| CD99 antigen OS=Homo sapiens OX=9606 GN=CD99 PE=1 SV=1 | P14209 | CD99_HUMAN | 19 kDa |
| Cell cycle control protein 50A OS=Homo sapiens OX=9606 GN=TMEM30A PE=1 SV=1 | Q9NV96 | CC50A_HUMAN | 41 kDa |
| Ceruloplasmin OS=Homo sapiens OX=9606 GN=CP PE=1 SV=1 | P00450 | CERU_HUMAN | 122 kDa |
| Charged multivesicular body protein 4b OS=Homo sapiens OX=9606 GN=CHMP4B PE=1 SV=1 | Q9H444 | CHM4B_HUMAN | 25 kDa |
| Charged multivesicular body protein 6 OS=Homo sapiens OX=9606 GN=CHMP6 PE=1 SV=3 | Q96FZ7 | CHMP6_HUMAN | 23 kDa |
| Chloride intracellular channel protein 1 OS=Homo sapiens OX=9606 GN=CLIC1 PE=1 SV=4 | O00299 | CLIC1_HUMAN | 27 kDa |
| Chloride intracellular channel protein 3 OS=Homo sapiens OX=9606 GN=CLIC3 PE=1 SV=2 | O95833 | CLIC3_HUMAN | 27 kDa |
| Chloride intracellular channel protein 4 OS=Homo sapiens OX=9606 GN=CLIC4 PE=1 SV=4 | Q9Y696 | CLIC4_HUMAN | 29 kDa |
| Cholesterol side-chain cleavage enzyme, mitochondrial OS=Homo sapiens OX=9606 GN=CYP11A1 PE=1 SV=2 | P05108 | CP11A_HUMAN | 60 kDa |
| Chondroadherin-like protein OS=Homo sapiens OX=9606 GN=CHADL PE=1 SV=2 | Q6NUI6 | CHADL_HUMAN | 82 kDa |
| Chromobox protein homolog 1 OS=Homo sapiens OX=9606 GN=CBX1 PE=1 SV=1 | P83916 | CBX1_HUMAN (+1) | 21 kDa |
| Citrate synthase, mitochondrial OS=Homo sapiens OX=9606 GN=CS PE=1 SV=2 | O75390 | CISY_HUMAN | 52 kDa |
| CKLF-like MARVEL transmembrane domain-containing protein 4 OS=Homo sapiens OX=9606 GN=CMTM4 PE=1 SV=1 | Q8IZR5 | CKLF4_HUMAN | 26 kDa |
| CKLF-like MARVEL transmembrane domain-containing protein 6 OS=Homo sapiens OX=9606 GN=CMTM6 PE=1 SV=1 | Q9NX76 | CKLF6_HUMAN | 20 kDa |
| Clathrin heavy chain 1 OS=Homo sapiens OX=9606 GN=CLTC PE=1 SV=5 | Q00610 | CLH1_HUMAN | 192 kDa |
| Clathrin light chain A OS=Homo sapiens OX=9606 GN=CLTA PE=1 SV=1 | P09496 | CLCA_HUMAN | 27 kDa |
| Claudin domain-containing protein 1 OS=Homo sapiens OX=9606 GN=CLDND1 PE=1 SV=1 | Q9NY35 | CLDN1_HUMAN | 29 kDa |
| Claudin-3 OS=Homo sapiens OX=9606 GN=CLDN3 PE=1 SV=1 | O15551 | CLD3_HUMAN | 23 kDa |
| Clusterin OS=Homo sapiens OX=9606 GN=CLU PE=1 SV=1 | P10909 | CLUS_HUMAN | 52 kDa |
| Coagulation factor X OS=Homo sapiens OX=9606 GN=F10 PE=1 SV=2 | P00742 | FA10_HUMAN | 55 kDa |
| Coagulation factor XI OS=Homo sapiens OX=9606 GN=F11 PE=1 SV=1 | P03951 | FA11_HUMAN | 70 kDa |
| Coatomer subunit alpha OS=Homo sapiens OX=9606 GN=COPA PE=1 SV=2 | P53621 | COPA_HUMAN | 138 kDa |
| Coatomer subunit beta OS=Homo sapiens OX=9606 GN=COPB1 PE=1 SV=3 | P53618 | COPB_HUMAN | 107 kDa |
| Coatomer subunit beta' OS=Homo sapiens OX=9606 GN=COPB2 PE=1 SV=2 | P35606 | COPB2_HUMAN | 102 kDa |
| Coatomer subunit epsilon OS=Homo sapiens OX=9606 GN=COPE PE=1 SV=3 | O14579 | COPE_HUMAN | 34 kDa |
| Coatomer subunit gamma-1 OS=Homo sapiens OX=9606 GN=COPG1 PE=1 SV=1 | Q9Y678 | COPG1_HUMAN | 98 kDa |
| Coiled-coil domain-containing protein 69 OS=Homo sapiens OX=9606 GN=CCDC69 PE=1 SV=1 | A6NI79 | CCD69_HUMAN | 35 kDa |
| Collagen alpha-1(IV) chain OS=Homo sapiens OX=9606 GN=COL4A1 PE=1 SV=4 | P02462 | CO4A1_HUMAN | 161 kDa |
| Collagen alpha-1(VI) chain OS=Homo sapiens OX=9606 GN=COL6A1 PE=1 SV=3 | P12109 | CO6A1_HUMAN | 109 kDa |
| Collagen alpha-1(XV) chain OS=Homo sapiens OX=9606 GN=COL15A1 PE=1 SV=2 | P39059 | COFA1_HUMAN | 142 kDa |
| Collagen alpha-1(XVIII) chain OS=Homo sapiens OX=9606 GN=COL18A1 PE=1 SV=5 | P39060 | COIA1_HUMAN | 178 kDa |
| Collagen alpha-2(I) chain OS=Homo sapiens OX=9606 GN=COL1A2 PE=1 SV=7 | P08123 | CO1A2_HUMAN | 129 kDa |
| Collagen alpha-2(IV) chain OS=Homo sapiens OX=9606 GN=COL4A2 PE=1 SV=4 | P08572 | CO4A2_HUMAN | 168 kDa |
| Collagen alpha-2(V) chain OS=Homo sapiens OX=9606 GN=COL5A2 PE=1 SV=3 | P05997 | CO5A2_HUMAN | 145 kDa |
| Collagen alpha-2(VI) chain OS=Homo sapiens OX=9606 GN=COL6A2 PE=1 SV=4 | P12110 | CO6A2_HUMAN | 109 kDa |
| Collagen alpha-3(VI) chain OS=Homo sapiens OX=9606 GN=COL6A3 PE=1 SV=5 | P12111 | CO6A3_HUMAN | 344 kDa |
| Colorectal mutant cancer protein OS=Homo sapiens OX=9606 GN=MCC PE=1 SV=2 | P23508 | CRCM_HUMAN | 93 kDa |
| Complement C1q subcomponent subunit A OS=Homo sapiens OX=9606 GN=C1QA PE=1 SV=2 | P02745 | C1QA_HUMAN | 26 kDa |
| Complement C1q subcomponent subunit B OS=Homo sapiens OX=9606 GN=C1QB PE=1 SV=3 | P02746 | C1QB_HUMAN | 27 kDa |
| Complement C1q subcomponent subunit C OS=Homo sapiens OX=9606 GN=C1QC PE=1 SV=3 | P02747 | C1QC_HUMAN | 26 kDa |
| Complement C1r subcomponent OS=Homo sapiens OX=9606 GN=C1R PE=1 SV=2 | P00736 | C1R_HUMAN | 80 kDa |
| Complement C1s subcomponent OS=Homo sapiens OX=9606 GN=C1S PE=1 SV=1 | P09871 | C1S_HUMAN | 77 kDa |
| Complement C2 OS=Homo sapiens OX=9606 GN=C2 PE=1 SV=2 | P06681 | CO2_HUMAN | 83 kDa |
| Complement C3 OS=Homo sapiens OX=9606 GN=C3 PE=1 SV=2 | P01024 | CO3_HUMAN | 187 kDa |
| Complement C5 OS=Homo sapiens OX=9606 GN=C5 PE=1 SV=4 | P01031 | CO5_HUMAN | 188 kDa |
| Complement component C6 OS=Homo sapiens OX=9606 GN=C6 PE=1 SV=3 | P13671 | CO6_HUMAN | 105 kDa |
| Complement component C7 OS=Homo sapiens OX=9606 GN=C7 PE=1 SV=2 | P10643 | CO7_HUMAN | 94 kDa |
| Complement component C8 alpha chain OS=Homo sapiens OX=9606 GN=C8A PE=1 SV=2 | P07357 | CO8A_HUMAN | 65 kDa |
| Complement component C8 beta chain OS=Homo sapiens OX=9606 GN=C8B PE=1 SV=3 | P07358 | CO8B_HUMAN | 67 kDa |
| Complement component C8 gamma chain OS=Homo sapiens OX=9606 GN=C8G PE=1 SV=3 | P07360 | CO8G_HUMAN | 22 kDa |
| Complement component C9 OS=Homo sapiens OX=9606 GN=C9 PE=1 SV=2 | P02748 | CO9_HUMAN | 63 kDa |
| Complement factor B OS=Homo sapiens OX=9606 GN=CFB PE=1 SV=2 | P00751 | CFAB_HUMAN | 86 kDa |
| Complement factor H OS=Homo sapiens OX=9606 GN=CFH PE=1 SV=4 | P08603 | CFAH_HUMAN | 139 kDa |
| Complement factor H-related protein 1 OS=Homo sapiens OX=9606 GN=CFHR1 PE=1 SV=2 | Q03591 | FHR1_HUMAN | 38 kDa |
| Complement factor I OS=Homo sapiens OX=9606 GN=CFI PE=1 SV=2 | P05156 | CFAI_HUMAN | 66 kDa |
| COP9 signalosome complex subunit 1 OS=Homo sapiens OX=9606 GN=GPS1 PE=1 SV=4 | Q13098 | CSN1_HUMAN | 56 kDa |
| COP9 signalosome complex subunit 4 OS=Homo sapiens OX=9606 GN=COPS4 PE=1 SV=1 | Q9BT78 | CSN4_HUMAN | 46 kDa |
| Copine-2 OS=Homo sapiens OX=9606 GN=CPNE2 PE=1 SV=3 | Q96FN4 | CPNE2_HUMAN | 61 kDa |
| Copine-3 OS=Homo sapiens OX=9606 GN=CPNE3 PE=1 SV=1 | O75131 | CPNE3_HUMAN | 60 kDa |
| Cornulin OS=Homo sapiens OX=9606 GN=CRNN PE=1 SV=1 | Q9UBG3 | CRNN_HUMAN | 54 kDa |
| Coronin-1B OS=Homo sapiens OX=9606 GN=CORO1B PE=1 SV=1 | Q9BR76 | COR1B_HUMAN | 54 kDa |
| Coronin-1C OS=Homo sapiens OX=9606 GN=CORO1C PE=1 SV=1 | Q9ULV4 | COR1C_HUMAN | 53 kDa |
| Coronin-7 OS=Homo sapiens OX=9606 GN=CORO7 PE=1 SV=2 | P57737 | CORO7_HUMAN | 101 kDa |
| Corticosteroid-binding globulin OS=Homo sapiens OX=9606 GN=SERPINA6 PE=1 SV=1 | P08185 | CBG_HUMAN | 45 kDa |
| Creatine kinase B-type OS=Homo sapiens OX=9606 GN=CKB PE=1 SV=1 | P12277 | KCRB_HUMAN | 43 kDa |
| CTP synthase 1 OS=Homo sapiens OX=9606 GN=CTPS1 PE=1 SV=2 | P17812 | PYRG1_HUMAN | 67 kDa |
| Cullin-7 OS=Homo sapiens OX=9606 GN=CUL7 PE=1 SV=2 | Q14999 | CUL7_HUMAN | 191 kDa |
| Cullin-associated NEDD8-dissociated protein 1 OS=Homo sapiens OX=9606 GN=CAND1 PE=1 SV=2 | Q86VP6 | CAND1_HUMAN | 136 kDa |
| Cyclin-dependent kinase 1 OS=Homo sapiens OX=9606 GN=CDK1 PE=1 SV=3 | P06493 | CDK1_HUMAN | 34 kDa |
| CYFIP-related Rac1 interactor A OS=Homo sapiens OX=9606 GN=CYRIA PE=2 SV=1 | Q9H0Q0 | CYRIA_HUMAN | 37 kDa |
| CYFIP-related Rac1 interactor B OS=Homo sapiens OX=9606 GN=CYRIB PE=1 SV=1 | Q9NUQ9 | CYRIB_HUMAN | 37 kDa |
| Cystatin-B OS=Homo sapiens OX=9606 GN=CSTB PE=1 SV=2 | P04080 | CYTB_HUMAN | 11 kDa |
| Cysteine and glycine-rich protein 2 OS=Homo sapiens OX=9606 GN=CSRP2 PE=1 SV=3 | Q16527 | CSRP2_HUMAN | 21 kDa |
| Cysteine-rich C-terminal protein 1 OS=Homo sapiens OX=9606 GN=CRCT1 PE=1 SV=1 | Q9UGL9 | CRCT1_HUMAN | 10 kDa |
| Cytochrome b-c1 complex subunit 2, mitochondrial OS=Homo sapiens OX=9606 GN=UQCRC2 PE=1 SV=3 | P22695 | QCR2_HUMAN | 48 kDa |
| Cytochrome c oxidase subunit 2 OS=Homo sapiens OX=9606 GN=MT-CO2 PE=1 SV=1 | P00403 | COX2_HUMAN | 26 kDa |
| Cytoplasmic aconitate hydratase OS=Homo sapiens OX=9606 GN=ACO1 PE=1 SV=3 | P21399 | ACOHC_HUMAN | 98 kDa |
| Cytoplasmic dynein 1 heavy chain 1 OS=Homo sapiens OX=9606 GN=DYNC1H1 PE=1 SV=5 | Q14204 | DYHC1_HUMAN | 532 kDa |
| Cytoplasmic dynein 1 intermediate chain 2 OS=Homo sapiens OX=9606 GN=DYNC1I2 PE=1 SV=3 | Q13409 | DC1I2_HUMAN | 71 kDa |
| Cytoplasmic dynein 1 light intermediate chain 1 OS=Homo sapiens OX=9606 GN=DYNC1LI1 PE=1 SV=3 | Q9Y6G9 | DC1L1_HUMAN | 57 kDa |
| Cytoplasmic dynein 1 light intermediate chain 2 OS=Homo sapiens OX=9606 GN=DYNC1LI2 PE=1 SV=1 | O43237 | DC1L2_HUMAN | 54 kDa |
| Cytoplasmic FMR1-interacting protein 1 OS=Homo sapiens OX=9606 GN=CYFIP1 PE=1 SV=1 | Q7L576 | CYFP1_HUMAN | 145 kDa |
| Cytoskeleton-associated protein 4 OS=Homo sapiens OX=9606 GN=CKAP4 PE=1 SV=2 | Q07065 | CKAP4_HUMAN | 66 kDa |
| Cytosol aminopeptidase OS=Homo sapiens OX=9606 GN=LAP3 PE=1 SV=3 | P28838 | AMPL_HUMAN | 56 kDa |
| Cytosolic non-specific dipeptidase OS=Homo sapiens OX=9606 GN=CNDP2 PE=1 SV=2 | Q96KP4 | CNDP2_HUMAN | 53 kDa |
| D-3-phosphoglycerate dehydrogenase OS=Homo sapiens OX=9606 GN=PHGDH PE=1 SV=4 | O43175 | SERA_HUMAN | 57 kDa |
| DCC-interacting protein 13-alpha OS=Homo sapiens OX=9606 GN=APPL1 PE=1 SV=1 | Q9UKG1 | DP13A_HUMAN | 80 kDa |
| DCC-interacting protein 13-beta OS=Homo sapiens OX=9606 GN=APPL2 PE=1 SV=3 | Q8NEU8 | DP13B_HUMAN | 74 kDa |
| Dedicator of cytokinesis protein 1 OS=Homo sapiens OX=9606 GN=DOCK1 PE=1 SV=2 | Q14185 | DOCK1_HUMAN | 215 kDa |
| Dedicator of cytokinesis protein 10 OS=Homo sapiens OX=9606 GN=DOCK10 PE=1 SV=3 | Q96BY6 | DOC10_HUMAN | 250 kDa |
| Dedicator of cytokinesis protein 7 OS=Homo sapiens OX=9606 GN=DOCK7 PE=1 SV=4 | Q96N67 | DOCK7_HUMAN | 243 kDa |
| Dedicator of cytokinesis protein 9 OS=Homo sapiens OX=9606 GN=DOCK9 PE=1 SV=2 | Q9BZ29 | DOCK9_HUMAN | 236 kDa |
| Deleted in malignant brain tumors 1 protein OS=Homo sapiens OX=9606 GN=DMBT1 PE=1 SV=2 | Q9UGM3 | DMBT1_HUMAN | 261 kDa |
| Delta(24)-sterol reductase OS=Homo sapiens OX=9606 GN=DHCR24 PE=1 SV=2 | Q15392 | DHC24_HUMAN | 60 kDa |
| Delta(3,5)-Delta(2,4)-dienoyl-CoA isomerase, mitochondrial OS=Homo sapiens OX=9606 GN=ECH1 PE=1 SV=2 | Q13011 | ECH1_HUMAN | 36 kDa |
| Delta-aminolevulinic acid dehydratase OS=Homo sapiens OX=9606 GN=ALAD PE=1 SV=1 | P13716 | HEM2_HUMAN | 36 kDa |
| Deoxycytidylate deaminase OS=Homo sapiens OX=9606 GN=DCTD PE=1 SV=2 | P32321 | DCTD_HUMAN | 20 kDa |
| Deoxyribose-phosphate aldolase OS=Homo sapiens OX=9606 GN=DERA PE=1 SV=2 | Q9Y315 | DEOC_HUMAN | 35 kDa |
| DEP domain-containing mTOR-interacting protein OS=Homo sapiens OX=9606 GN=DEPTOR PE=1 SV=2 | Q8TB45 | DPTOR_HUMAN | 46 kDa |
| Dermcidin OS=Homo sapiens OX=9606 GN=DCD PE=1 SV=2 | P81605 | DCD_HUMAN | 11 kDa |
| Desmocollin-1 OS=Homo sapiens OX=9606 GN=DSC1 PE=1 SV=2 | Q08554 | DSC1_HUMAN | 100 kDa |
| Desmocollin-2 OS=Homo sapiens OX=9606 GN=DSC2 PE=1 SV=1 | Q02487 | DSC2_HUMAN | 100 kDa |
| Desmoglein-1 OS=Homo sapiens OX=9606 GN=DSG1 PE=1 SV=2 | Q02413 | DSG1_HUMAN | 114 kDa |
| Desmoglein-3 OS=Homo sapiens OX=9606 GN=DSG3 PE=1 SV=2 | P32926 | DSG3_HUMAN | 108 kDa |
| Desmoplakin OS=Homo sapiens OX=9606 GN=DSP PE=1 SV=3 | P15924 | DESP_HUMAN | 332 kDa |
| Destrin OS=Homo sapiens OX=9606 GN=DSTN PE=1 SV=3 | P60981 | DEST_HUMAN | 19 kDa |
| Dihydrolipoyllysine-residue succinyltransferase component of 2-oxoglutarate dehydrogenase complex, mitochondrial OS=Homo sapiens OX=9606 GN=DLST PE=1 SV=4 | P36957 | ODO2_HUMAN | 49 kDa |
| Dihydropteridine reductase OS=Homo sapiens OX=9606 GN=QDPR PE=1 SV=2 | P09417 | DHPR_HUMAN | 26 kDa |
| Dihydropyrimidinase-related protein 3 OS=Homo sapiens OX=9606 GN=DPYSL3 PE=1 SV=1 | Q14195 | DPYL3_HUMAN | 62 kDa |
| Dipeptidyl peptidase 3 OS=Homo sapiens OX=9606 GN=DPP3 PE=1 SV=2 | Q9NY33 | DPP3_HUMAN | 83 kDa |
| Dipeptidyl peptidase 4 OS=Homo sapiens OX=9606 GN=DPP4 PE=1 SV=2 | P27487 | DPP4_HUMAN | 88 kDa |
| Diphosphomevalonate decarboxylase OS=Homo sapiens OX=9606 GN=MVD PE=1 SV=1 | P53602 | MVD1_HUMAN | 43 kDa |
| Disco-interacting protein 2 homolog B OS=Homo sapiens OX=9606 GN=DIP2B PE=1 SV=3 | Q9P265 | DIP2B_HUMAN | 171 kDa |
| Disheveled-associated activator of morphogenesis 1 OS=Homo sapiens OX=9606 GN=DAAM1 PE=1 SV=2 | Q9Y4D1 | DAAM1_HUMAN | 123 kDa |
| Disintegrin and metalloproteinase domain-containing protein 10 OS=Homo sapiens OX=9606 GN=ADAM10 PE=1 SV=1 | O14672 | ADA10_HUMAN | 84 kDa |
| Disintegrin and metalloproteinase domain-containing protein 17 OS=Homo sapiens OX=9606 GN=ADAM17 PE=1 SV=1 | P78536 | ADA17_HUMAN | 93 kDa |
| Disintegrin and metalloproteinase domain-containing protein 9 OS=Homo sapiens OX=9606 GN=ADAM9 PE=1 SV=1 | Q13443 | ADAM9_HUMAN | 91 kDa |
| DNA repair protein complementing XP-C cells OS=Homo sapiens OX=9606 GN=XPC PE=1 SV=4 | Q01831 | XPC_HUMAN | 106 kDa |
| DnaJ homolog subfamily A member 1 OS=Homo sapiens OX=9606 GN=DNAJA1 PE=1 SV=2 | P31689 | DNJA1_HUMAN | 45 kDa |
| DnaJ homolog subfamily C member 13 OS=Homo sapiens OX=9606 GN=DNAJC13 PE=1 SV=5 | O75165 | DJC13_HUMAN | 254 kDa |
| DnaJ homolog subfamily C member 5 OS=Homo sapiens OX=9606 GN=DNAJC5 PE=1 SV=1 | Q9H3Z4 | DNJC5_HUMAN | 22 kDa |
| Dolichyl-diphosphooligosaccharide--protein glycosyltransferase 48 kDa subunit OS=Homo sapiens OX=9606 GN=DDOST PE=1 SV=4 | P39656 | OST48_HUMAN | 51 kDa |
| Dolichyl-diphosphooligosaccharide--protein glycosyltransferase subunit 1 OS=Homo sapiens OX=9606 GN=RPN1 PE=1 SV=1 | P04843 | RPN1_HUMAN | 69 kDa |
| Dolichyl-diphosphooligosaccharide--protein glycosyltransferase subunit 2 OS=Homo sapiens OX=9606 GN=RPN2 PE=1 SV=3 | P04844 | RPN2_HUMAN | 69 kDa |
| Drebrin OS=Homo sapiens OX=9606 GN=DBN1 PE=1 SV=4 | Q16643 | DREB_HUMAN | 71 kDa |
| Drebrin-like protein OS=Homo sapiens OX=9606 GN=DBNL PE=1 SV=1 | Q9UJU6 | DBNL_HUMAN | 48 kDa |
| Dynactin subunit 1 OS=Homo sapiens OX=9606 GN=DCTN1 PE=1 SV=3 | Q14203 | DCTN1_HUMAN | 142 kDa |
| Dynactin subunit 2 OS=Homo sapiens OX=9606 GN=DCTN2 PE=1 SV=4 | Q13561 | DCTN2_HUMAN | 44 kDa |
| Dynactin subunit 4 OS=Homo sapiens OX=9606 GN=DCTN4 PE=1 SV=1 | Q9UJW0 | DCTN4_HUMAN | 52 kDa |
| Dynamin-2 OS=Homo sapiens OX=9606 GN=DNM2 PE=1 SV=2 | P50570 | DYN2_HUMAN | 98 kDa |
| Dynein light chain 1, cytoplasmic OS=Homo sapiens OX=9606 GN=DYNLL1 PE=1 SV=1 | P63167 | DYL1_HUMAN | 10 kDa |
| Dynein light chain Tctex-type 1 OS=Homo sapiens OX=9606 GN=DYNLT1 PE=1 SV=1 | P63172 | DYLT1_HUMAN | 12 kDa |
| Dystroglycan 1 OS=Homo sapiens OX=9606 GN=DAG1 PE=1 SV=2 | Q14118 | DAG1_HUMAN | 97 kDa |
| Dystrophin OS=Homo sapiens OX=9606 GN=DMD PE=1 SV=3 | P11532 | DMD_HUMAN | 427 kDa |
| E3 ubiquitin-protein ligase HUWE1 OS=Homo sapiens OX=9606 GN=HUWE1 PE=1 SV=3 | Q7Z6Z7 | HUWE1_HUMAN | 482 kDa |
| E3 ubiquitin-protein ligase MGRN1 OS=Homo sapiens OX=9606 GN=MGRN1 PE=1 SV=2 | O60291 | MGRN1_HUMAN | 61 kDa |
| E3 ubiquitin-protein ligase RBX1 OS=Homo sapiens OX=9606 GN=RBX1 PE=1 SV=1 | P62877 | RBX1_HUMAN | 12 kDa |
| E3 ubiquitin-protein ligase RNF213 OS=Homo sapiens OX=9606 GN=RNF213 PE=1 SV=3 | Q63HN8 | RN213_HUMAN | 591 kDa |
| E3 ubiquitin-protein ligase TRIM4 OS=Homo sapiens OX=9606 GN=TRIM4 PE=1 SV=2 | Q9C037 | TRIM4_HUMAN | 57 kDa |
| E3 ubiquitin-protein ligase UBR4 OS=Homo sapiens OX=9606 GN=UBR4 PE=1 SV=1 | Q5T4S7 | UBR4_HUMAN | 574 kDa |
| EARP and GARP complex-interacting protein 1 OS=Homo sapiens OX=9606 GN=EIPR1 PE=1 SV=2 | Q53HC9 | EIPR1_HUMAN | 44 kDa |
| Echinoderm microtubule-associated protein-like 2 OS=Homo sapiens OX=9606 GN=EML2 PE=1 SV=1 | O95834 | EMAL2_HUMAN | 71 kDa |
| EGF-like repeat and discoidin I-like domain-containing protein 3 OS=Homo sapiens OX=9606 GN=EDIL3 PE=1 SV=1 | O43854 | EDIL3_HUMAN | 54 kDa |
| EH domain-containing protein 1 OS=Homo sapiens OX=9606 GN=EHD1 PE=1 SV=2 | Q9H4M9 | EHD1_HUMAN | 61 kDa |
| EH domain-containing protein 2 OS=Homo sapiens OX=9606 GN=EHD2 PE=1 SV=2 | Q9NZN4 | EHD2_HUMAN | 61 kDa |
| EH domain-containing protein 4 OS=Homo sapiens OX=9606 GN=EHD4 PE=1 SV=1 | Q9H223 | EHD4_HUMAN | 61 kDa |
| eIF-2-alpha kinase activator GCN1 OS=Homo sapiens OX=9606 GN=GCN1 PE=1 SV=7 | Q92616 | GCN1_HUMAN | 293 kDa |
| eIF5-mimic protein 2 OS=Homo sapiens OX=9606 GN=BZW1 PE=1 SV=1 | Q7L1Q6 | 5MP2_HUMAN | 48 kDa |
| Elongation factor 1-alpha 1 OS=Homo sapiens OX=9606 GN=EEF1A1 PE=1 SV=1 | P68104 | EF1A1_HUMAN (+1) | 50 kDa |
| Elongation factor 1-beta OS=Homo sapiens OX=9606 GN=EEF1B2 PE=1 SV=3 | P24534 | EF1B_HUMAN | 25 kDa |
| Elongation factor 1-delta OS=Homo sapiens OX=9606 GN=EEF1D PE=1 SV=5 | P29692 | EF1D_HUMAN | 31 kDa |
| Elongation factor 1-gamma OS=Homo sapiens OX=9606 GN=EEF1G PE=1 SV=3 | P26641 | EF1G_HUMAN | 50 kDa |
| Elongation factor 2 OS=Homo sapiens OX=9606 GN=EEF2 PE=1 SV=4 | P13639 | EF2_HUMAN | 95 kDa |
| Elongation factor Tu, mitochondrial OS=Homo sapiens OX=9606 GN=TUFM PE=1 SV=2 | P49411 | EFTU_HUMAN | 50 kDa |
| Elongin-C OS=Homo sapiens OX=9606 GN=ELOC PE=1 SV=1 | Q15369 | ELOC_HUMAN | 12 kDa |
| EMILIN-1 OS=Homo sapiens OX=9606 GN=EMILIN1 PE=1 SV=3 | Q9Y6C2 | EMIL1_HUMAN | 107 kDa |
| Endophilin-A2 OS=Homo sapiens OX=9606 GN=SH3GL1 PE=1 SV=1 | Q99961 | SH3G1_HUMAN | 41 kDa |
| Endoplasmic reticulum aminopeptidase 1 OS=Homo sapiens OX=9606 GN=ERAP1 PE=1 SV=3 | Q9NZ08 | ERAP1_HUMAN | 107 kDa |
| Endoplasmic reticulum chaperone BiP OS=Homo sapiens OX=9606 GN=HSPA5 PE=1 SV=2 | P11021 | BIP_HUMAN | 72 kDa |
| Endoplasmic reticulum metallopeptidase 1 OS=Homo sapiens OX=9606 GN=ERMP1 PE=1 SV=2 | Q7Z2K6 | ERMP1_HUMAN | 100 kDa |
| Endoplasmic reticulum resident protein 29 OS=Homo sapiens OX=9606 GN=ERP29 PE=1 SV=4 | P30040 | ERP29_HUMAN | 29 kDa |
| Endoplasmic reticulum resident protein 44 OS=Homo sapiens OX=9606 GN=ERP44 PE=1 SV=1 | Q9BS26 | ERP44_HUMAN | 47 kDa |
| Endoplasmin OS=Homo sapiens OX=9606 GN=HSP90B1 PE=1 SV=1 | P14625 | ENPL_HUMAN | 92 kDa |
| Endothelin-converting enzyme 1 OS=Homo sapiens OX=9606 GN=ECE1 PE=1 SV=2 | P42892 | ECE1_HUMAN | 87 kDa |
| Engulfment and cell motility protein 1 OS=Homo sapiens OX=9606 GN=ELMO1 PE=1 SV=2 | Q92556 | ELMO1_HUMAN | 84 kDa |
| Envoplakin OS=Homo sapiens OX=9606 GN=EVPL PE=1 SV=3 | Q92817 | EVPL_HUMAN | 232 kDa |
| Ephrin type-B receptor 4 OS=Homo sapiens OX=9606 GN=EPHB4 PE=1 SV=2 | P54760 | EPHB4_HUMAN | 108 kDa |
| Ephrin-B1 OS=Homo sapiens OX=9606 GN=EFNB1 PE=1 SV=1 | P98172 | EFNB1_HUMAN | 38 kDa |
| Epidermal growth factor receptor kinase substrate 8 OS=Homo sapiens OX=9606 GN=EPS8 PE=1 SV=1 | Q12929 | EPS8_HUMAN | 92 kDa |
| Epidermal growth factor receptor kinase substrate 8-like protein 1 OS=Homo sapiens OX=9606 GN=EPS8L1 PE=1 SV=3 | Q8TE68 | ES8L1_HUMAN | 80 kDa |
| Epidermal growth factor receptor OS=Homo sapiens OX=9606 GN=EGFR PE=1 SV=2 | P00533 | EGFR_HUMAN | 134 kDa |
| Epoxide hydrolase 1 OS=Homo sapiens OX=9606 GN=EPHX1 PE=1 SV=1 | P07099 | HYEP_HUMAN | 53 kDa |
| Epsin-3 OS=Homo sapiens OX=9606 GN=EPN3 PE=2 SV=1 | Q9H201 | EPN3_HUMAN (+1) | 68 kDa |
| Equilibrative nucleobase transporter 1 OS=Homo sapiens OX=9606 GN=SLC43A3 PE=1 SV=2 | Q8NBI5 | S43A3_HUMAN | 55 kDa |
| Equilibrative nucleoside transporter 1 OS=Homo sapiens OX=9606 GN=SLC29A1 PE=1 SV=3 | Q99808 | S29A1_HUMAN | 50 kDa |
| ERO1-like protein alpha OS=Homo sapiens OX=9606 GN=ERO1A PE=1 SV=2 | Q96HE7 | ERO1A_HUMAN | 54 kDa |
| Ethanolamine-phosphate cytidylyltransferase OS=Homo sapiens OX=9606 GN=PCYT2 PE=1 SV=1 | Q99447 | PCY2_HUMAN | 44 kDa |
| Eukaryotic peptide chain release factor GTP-binding subunit ERF3A OS=Homo sapiens OX=9606 GN=GSPT1 PE=1 SV=1 | P15170 | ERF3A_HUMAN | 56 kDa |
| Eukaryotic peptide chain release factor subunit 1 OS=Homo sapiens OX=9606 GN=ETF1 PE=1 SV=3 | P62495 | ERF1_HUMAN | 49 kDa |
| Eukaryotic translation initiation factor 2 subunit 1 OS=Homo sapiens OX=9606 GN=EIF2S1 PE=1 SV=3 | P05198 | IF2A_HUMAN | 36 kDa |
| Eukaryotic translation initiation factor 2 subunit 3 OS=Homo sapiens OX=9606 GN=EIF2S3 PE=1 SV=3 | P41091 | IF2G_HUMAN | 51 kDa |
| Eukaryotic translation initiation factor 3 subunit A OS=Homo sapiens OX=9606 GN=EIF3A PE=1 SV=1 | Q14152 | EIF3A_HUMAN | 167 kDa |
| Eukaryotic translation initiation factor 3 subunit B OS=Homo sapiens OX=9606 GN=EIF3B PE=1 SV=3 | P55884 | EIF3B_HUMAN | 92 kDa |
| Eukaryotic translation initiation factor 3 subunit C-like protein OS=Homo sapiens OX=9606 GN=EIF3CL PE=1 SV=1 | B5ME19 | EIFCL_HUMAN (+1) | 105 kDa |
| Eukaryotic translation initiation factor 3 subunit D OS=Homo sapiens OX=9606 GN=EIF3D PE=1 SV=1 | O15371 | EIF3D_HUMAN | 64 kDa |
| Eukaryotic translation initiation factor 3 subunit E OS=Homo sapiens OX=9606 GN=EIF3E PE=1 SV=1 | P60228 | EIF3E_HUMAN | 52 kDa |
| Eukaryotic translation initiation factor 3 subunit F OS=Homo sapiens OX=9606 GN=EIF3F PE=1 SV=1 | O00303 | EIF3F_HUMAN | 38 kDa |
| Eukaryotic translation initiation factor 3 subunit I OS=Homo sapiens OX=9606 GN=EIF3I PE=1 SV=1 | Q13347 | EIF3I_HUMAN | 37 kDa |
| Eukaryotic translation initiation factor 3 subunit K OS=Homo sapiens OX=9606 GN=EIF3K PE=1 SV=1 | Q9UBQ5 | EIF3K_HUMAN | 25 kDa |
| Eukaryotic translation initiation factor 3 subunit L OS=Homo sapiens OX=9606 GN=EIF3L PE=1 SV=1 | Q9Y262 | EIF3L_HUMAN | 67 kDa |
| Eukaryotic translation initiation factor 5 OS=Homo sapiens OX=9606 GN=EIF5 PE=1 SV=2 | P55010 | IF5_HUMAN | 49 kDa |
| Eukaryotic translation initiation factor 5A-1 OS=Homo sapiens OX=9606 GN=EIF5A PE=1 SV=2 | P63241 | IF5A1_HUMAN (+1) | 17 kDa |
| Eukaryotic translation initiation factor 5B OS=Homo sapiens OX=9606 GN=EIF5B PE=1 SV=4 | O60841 | IF2P_HUMAN | 139 kDa |
| Eukaryotic translation initiation factor 6 OS=Homo sapiens OX=9606 GN=EIF6 PE=1 SV=1 | P56537 | IF6_HUMAN | 27 kDa |
| Exocyst complex component 1 OS=Homo sapiens OX=9606 GN=EXOC1 PE=1 SV=4 | Q9NV70 | EXOC1_HUMAN | 102 kDa |
| Exocyst complex component 2 OS=Homo sapiens OX=9606 GN=EXOC2 PE=1 SV=1 | Q96KP1 | EXOC2_HUMAN | 104 kDa |
| Exocyst complex component 4 OS=Homo sapiens OX=9606 GN=EXOC4 PE=1 SV=1 | Q96A65 | EXOC4_HUMAN | 111 kDa |
| Exportin-1 OS=Homo sapiens OX=9606 GN=XPO1 PE=1 SV=1 | O14980 | XPO1_HUMAN | 123 kDa |
| Extracellular matrix organizing protein FRAS1 OS=Homo sapiens OX=9606 GN=FRAS1 PE=1 SV=2 | Q86XX4 | FRAS1_HUMAN | 443 kDa |
| Ezrin OS=Homo sapiens OX=9606 GN=EZR PE=1 SV=4 | P15311 | EZRI_HUMAN | 69 kDa |
| F-actin-capping protein subunit alpha-1 OS=Homo sapiens OX=9606 GN=CAPZA1 PE=1 SV=3 | P52907 | CAZA1_HUMAN | 33 kDa |
| F-actin-capping protein subunit alpha-2 OS=Homo sapiens OX=9606 GN=CAPZA2 PE=1 SV=3 | P47755 | CAZA2_HUMAN | 33 kDa |
| F-actin-capping protein subunit beta OS=Homo sapiens OX=9606 GN=CAPZB PE=1 SV=5 | P47756 | CAPZB_HUMAN | 31 kDa |
| F-actin-uncapping protein LRRC16A OS=Homo sapiens OX=9606 GN=CARMIL1 PE=1 SV=1 | Q5VZK9 | CARL1_HUMAN | 152 kDa |
| Farnesyl pyrophosphate synthase OS=Homo sapiens OX=9606 GN=FDPS PE=1 SV=4 | P14324 | FPPS_HUMAN | 48 kDa |
| Fascin OS=Homo sapiens OX=9606 GN=FSCN1 PE=1 SV=3 | Q16658 | FSCN1_HUMAN | 55 kDa |
| Fatty acid synthase OS=Homo sapiens OX=9606 GN=FASN PE=1 SV=3 | P49327 | FAS_HUMAN | 273 kDa |
| Fatty acid-binding protein 5 OS=Homo sapiens OX=9606 GN=FABP5 PE=1 SV=3 | Q01469 | FABP5_HUMAN | 15 kDa |
| F-box only protein 40 OS=Homo sapiens OX=9606 GN=FBXO40 PE=2 SV=2 | Q9UH90 | FBX40_HUMAN | 80 kDa |
| F-box only protein 50 OS=Homo sapiens OX=9606 GN=NCCRP1 PE=1 SV=1 | Q6ZVX7 | FBX50_HUMAN | 31 kDa |
| F-box/WD repeat-containing protein 8 OS=Homo sapiens OX=9606 GN=FBXW8 PE=1 SV=2 | Q8N3Y1 | FBXW8_HUMAN | 67 kDa |
| FERM, ARHGEF and pleckstrin domain-containing protein 1 OS=Homo sapiens OX=9606 GN=FARP1 PE=1 SV=1 | Q9Y4F1 | FARP1_HUMAN | 119 kDa |
| FERM, ARHGEF and pleckstrin domain-containing protein 2 OS=Homo sapiens OX=9606 GN=FARP2 PE=1 SV=3 | O94887 | FARP2_HUMAN | 120 kDa |
| Fermitin family homolog 2 OS=Homo sapiens OX=9606 GN=FERMT2 PE=1 SV=1 | Q96AC1 | FERM2_HUMAN | 78 kDa |
| Ferritin heavy chain OS=Homo sapiens OX=9606 GN=FTH1 PE=1 SV=2 | P02794 | FRIH_HUMAN | 21 kDa |
| Ferritin light chain OS=Homo sapiens OX=9606 GN=FTL PE=1 SV=2 | P02792 | FRIL_HUMAN | 20 kDa |
| FHF complex subunit HOOK interacting protein 1B OS=Homo sapiens OX=9606 GN=FHIP1B PE=1 SV=3 | Q8N612 | FHI1B_HUMAN | 106 kDa |
| Fibrinogen alpha chain OS=Homo sapiens OX=9606 GN=FGA PE=1 SV=2 | P02671 | FIBA_HUMAN | 95 kDa |
| Fibrinogen beta chain OS=Homo sapiens OX=9606 GN=FGB PE=1 SV=2 | P02675 | FIBB_HUMAN | 56 kDa |
| Fibrinogen gamma chain OS=Homo sapiens OX=9606 GN=FGG PE=1 SV=3 | P02679 | FIBG_HUMAN | 52 kDa |
| Fibronectin OS=Homo sapiens OX=9606 GN=FN1 PE=1 SV=5 | P02751 | FINC_HUMAN | 272 kDa |
| Fibulin-1 OS=Homo sapiens OX=9606 GN=FBLN1 PE=1 SV=4 | P23142 | FBLN1_HUMAN | 77 kDa |
| Ficolin-2 OS=Homo sapiens OX=9606 GN=FCN2 PE=1 SV=2 | Q15485 | FCN2_HUMAN | 34 kDa |
| Ficolin-3 OS=Homo sapiens OX=9606 GN=FCN3 PE=1 SV=2 | O75636 | FCN3_HUMAN | 33 kDa |
| Filaggrin OS=Homo sapiens OX=9606 GN=FLG PE=1 SV=3 | P20930 | FILA_HUMAN | 435 kDa |
| Filaggrin-2 OS=Homo sapiens OX=9606 GN=FLG2 PE=1 SV=1 | Q5D862 | FILA2_HUMAN | 248 kDa |
| Filamin-A OS=Homo sapiens OX=9606 GN=FLNA PE=1 SV=4 | P21333 | FLNA_HUMAN | 281 kDa |
| Filamin-B OS=Homo sapiens OX=9606 GN=FLNB PE=1 SV=2 | O75369 | FLNB_HUMAN | 278 kDa |
| Flavin reductase (NADPH) OS=Homo sapiens OX=9606 GN=BLVRB PE=1 SV=3 | P30043 | BLVRB_HUMAN | 22 kDa |
| Flotillin-1 OS=Homo sapiens OX=9606 GN=FLOT1 PE=1 SV=3 | O75955 | FLOT1_HUMAN | 47 kDa |
| Flotillin-2 OS=Homo sapiens OX=9606 GN=FLOT2 PE=1 SV=2 | Q14254 | FLOT2_HUMAN | 47 kDa |
| Follistatin-related protein 4 OS=Homo sapiens OX=9606 GN=FSTL4 PE=2 SV=3 | Q6MZW2 | FSTL4_HUMAN | 93 kDa |
| Four and a half LIM domains protein 1 OS=Homo sapiens OX=9606 GN=FHL1 PE=1 SV=4 | Q13642 | FHL1_HUMAN | 36 kDa |
| Four and a half LIM domains protein 2 OS=Homo sapiens OX=9606 GN=FHL2 PE=1 SV=3 | Q14192 | FHL2_HUMAN | 32 kDa |
| Fructose-1,6-bisphosphatase 1 OS=Homo sapiens OX=9606 GN=FBP1 PE=1 SV=5 | P09467 | F16P1_HUMAN | 37 kDa |
| Fructose-bisphosphate aldolase A OS=Homo sapiens OX=9606 GN=ALDOA PE=1 SV=2 | P04075 | ALDOA_HUMAN | 39 kDa |
| Fructose-bisphosphate aldolase C OS=Homo sapiens OX=9606 GN=ALDOC PE=1 SV=2 | P09972 | ALDOC_HUMAN | 39 kDa |
| Galectin-3-binding protein OS=Homo sapiens OX=9606 GN=LGALS3BP PE=1 SV=1 | Q08380 | LG3BP_HUMAN | 65 kDa |
| Galectin-7 OS=Homo sapiens OX=9606 GN=LGALS7B PE=1 SV=2 | P47929 | LEG7_HUMAN | 15 kDa |
| Galectin-8 OS=Homo sapiens OX=9606 GN=LGALS8 PE=1 SV=4 | O00214 | LEG8_HUMAN | 36 kDa |
| Gamma-soluble NSF attachment protein OS=Homo sapiens OX=9606 GN=NAPG PE=1 SV=1 | Q99747 | SNAG_HUMAN | 35 kDa |
| Gap junction alpha-1 protein OS=Homo sapiens OX=9606 GN=GJA1 PE=1 SV=2 | P17302 | CXA1_HUMAN | 43 kDa |
| Gastrin/cholecystokinin type B receptor OS=Homo sapiens OX=9606 GN=CCKBR PE=1 SV=1 | P32239 | GASR_HUMAN | 48 kDa |
| Gelsolin OS=Homo sapiens OX=9606 GN=GSN PE=1 SV=1 | P06396 | GELS_HUMAN | 86 kDa |
| General vesicular transport factor p115 OS=Homo sapiens OX=9606 GN=USO1 PE=1 SV=2 | O60763 | USO1_HUMAN | 108 kDa |
| Glia maturation factor beta OS=Homo sapiens OX=9606 GN=GMFB PE=1 SV=2 | P60983 | GMFB_HUMAN | 17 kDa |
| Glia-derived nexin OS=Homo sapiens OX=9606 GN=SERPINE2 PE=1 SV=1 | P07093 | GDN_HUMAN | 44 kDa |
| Glucose-6-phosphate 1-dehydrogenase OS=Homo sapiens OX=9606 GN=G6PD PE=1 SV=4 | P11413 | G6PD_HUMAN | 59 kDa |
| Glucose-6-phosphate isomerase OS=Homo sapiens OX=9606 GN=GPI PE=1 SV=4 | P06744 | G6PI_HUMAN | 63 kDa |
| Glucosidase 2 subunit beta OS=Homo sapiens OX=9606 GN=PRKCSH PE=1 SV=2 | P14314 | GLU2B_HUMAN | 59 kDa |
| Glutamate dehydrogenase 1, mitochondrial OS=Bos taurus OX=9913 GN=GLUD1 PE=1 SV=2 | P00366 | DHE3_BOVIN (+2) | 62 kDa |
| Glutamate--cysteine ligase catalytic subunit OS=Homo sapiens OX=9606 GN=GCLC PE=1 SV=2 | P48506 | GSH1_HUMAN | 73 kDa |
| Glutamine synthetase OS=Homo sapiens OX=9606 GN=GLUL PE=1 SV=4 | P15104 | GLNA_HUMAN | 42 kDa |
| Glutamine--fructose-6-phosphate aminotransferase [isomerizing] 1 OS=Homo sapiens OX=9606 GN=GFPT1 PE=1 SV=3 | Q06210 | GFPT1_HUMAN | 79 kDa |
| Glutaredoxin-3 OS=Homo sapiens OX=9606 GN=GLRX3 PE=1 SV=2 | O76003 | GLRX3_HUMAN | 37 kDa |
| Glutathione peroxidase 1 OS=Homo sapiens OX=9606 GN=GPX1 PE=1 SV=4 | P07203 | GPX1_HUMAN | 22 kDa |
| Glutathione peroxidase 3 OS=Homo sapiens OX=9606 GN=GPX3 PE=1 SV=2 | P22352 | GPX3_HUMAN | 26 kDa |
| Glutathione reductase, mitochondrial OS=Homo sapiens OX=9606 GN=GSR PE=1 SV=2 | P00390 | GSHR_HUMAN | 56 kDa |
| Glutathione S-transferase A1 OS=Homo sapiens OX=9606 GN=GSTA1 PE=1 SV=3 | P08263 | GSTA1_HUMAN | 26 kDa |
| Glutathione S-transferase A4 OS=Homo sapiens OX=9606 GN=GSTA4 PE=1 SV=1 | O15217 | GSTA4_HUMAN | 26 kDa |
| Glutathione S-transferase Mu 2 OS=Homo sapiens OX=9606 GN=GSTM2 PE=1 SV=2 | P28161 | GSTM2_HUMAN (+1) | 26 kDa |
| Glutathione S-transferase omega-1 OS=Homo sapiens OX=9606 GN=GSTO1 PE=1 SV=2 | P78417 | GSTO1_HUMAN | 28 kDa |
| Glutathione S-transferase P OS=Homo sapiens OX=9606 GN=GSTP1 PE=1 SV=2 | P09211 | GSTP1_HUMAN | 23 kDa |
| Glyceraldehyde-3-phosphate dehydrogenase OS=Homo sapiens OX=9606 GN=GAPDH PE=1 SV=3 | P04406 | G3P_HUMAN | 36 kDa |
| Glycerol-3-phosphate phosphatase OS=Homo sapiens OX=9606 GN=PGP PE=1 SV=1 | A6NDG6 | PGP_HUMAN | 34 kDa |
| Glycine--tRNA ligase OS=Homo sapiens OX=9606 GN=GARS1 PE=1 SV=3 | P41250 | GARS_HUMAN | 83 kDa |
| Glycogen phosphorylase, brain form OS=Homo sapiens OX=9606 GN=PYGB PE=1 SV=5 | P11216 | PYGB_HUMAN | 97 kDa |
| Glycogenin-1 OS=Homo sapiens OX=9606 GN=GYG1 PE=1 SV=4 | P46976 | GLYG_HUMAN | 39 kDa |
| Glyoxylate reductase/hydroxypyruvate reductase OS=Homo sapiens OX=9606 GN=GRHPR PE=1 SV=1 | Q9UBQ7 | GRHPR_HUMAN | 36 kDa |
| Glypican-6 OS=Homo sapiens OX=9606 GN=GPC6 PE=1 SV=1 | Q9Y625 | GPC6_HUMAN | 63 kDa |
| Golgin subfamily A member 7 OS=Homo sapiens OX=9606 GN=GOLGA7 PE=1 SV=2 | Q7Z5G4 | GOGA7_HUMAN | 16 kDa |
| Growth factor receptor-bound protein 14 OS=Homo sapiens OX=9606 GN=GRB14 PE=1 SV=2 | Q14449 | GRB14_HUMAN | 61 kDa |
| GTPase NRas OS=Homo sapiens OX=9606 GN=NRAS PE=1 SV=1 | P01111 | RASN_HUMAN | 21 kDa |
| GTP-binding nuclear protein Ran OS=Homo sapiens OX=9606 GN=RAN PE=1 SV=3 | P62826 | RAN_HUMAN | 24 kDa |
| GTP-binding protein 1 OS=Homo sapiens OX=9606 GN=GTPBP1 PE=1 SV=3 | O00178 | GTPB1_HUMAN | 72 kDa |
| GTP-binding protein Rheb OS=Homo sapiens OX=9606 GN=RHEB PE=1 SV=1 | Q15382 | RHEB_HUMAN | 20 kDa |
| GTP-binding protein SAR1a OS=Homo sapiens OX=9606 GN=SAR1A PE=1 SV=1 | Q9NR31 | SAR1A_HUMAN (+1) | 22 kDa |
| Guanine nucleotide-binding protein G(i) subunit alpha-2 OS=Homo sapiens OX=9606 GN=GNAI2 PE=1 SV=3 | P04899 | GNAI2_HUMAN | 40 kDa |
| Guanine nucleotide-binding protein G(i) subunit alpha-3 OS=Homo sapiens OX=9606 GN=GNAI3 PE=1 SV=3 | P08754 | GNAI3_HUMAN | 41 kDa |
| Guanine nucleotide-binding protein G(I)/G(S)/G(O) subunit gamma-12 OS=Homo sapiens OX=9606 GN=GNG12 PE=1 SV=3 | Q9UBI6 | GBG12_HUMAN | 8 kDa |
| Guanine nucleotide-binding protein G(I)/G(S)/G(O) subunit gamma-5 OS=Homo sapiens OX=9606 GN=GNG5 PE=1 SV=3 | P63218 | GBG5_HUMAN | 7 kDa |
| Guanine nucleotide-binding protein G(q) subunit alpha OS=Homo sapiens OX=9606 GN=GNAQ PE=1 SV=4 | P50148 | GNAQ_HUMAN | 42 kDa |
| Guanine nucleotide-binding protein G(s) subunit alpha isoforms short OS=Homo sapiens OX=9606 GN=GNAS PE=1 SV=1 | P63092 | GNAS2_HUMAN (+1) | 46 kDa |
| Guanine nucleotide-binding protein G(t) subunit alpha-3 OS=Homo sapiens OX=9606 GN=GNAT3 PE=2 SV=2 | A8MTJ3 | GNAT3_HUMAN | 40 kDa |
| Guanine nucleotide-binding protein subunit alpha-11 OS=Homo sapiens OX=9606 GN=GNA11 PE=1 SV=2 | P29992 | GNA11_HUMAN | 42 kDa |
| Guanine nucleotide-binding protein subunit alpha-13 OS=Homo sapiens OX=9606 GN=GNA13 PE=1 SV=2 | Q14344 | GNA13_HUMAN | 44 kDa |
| Guanine nucleotide-binding protein subunit alpha-14 OS=Homo sapiens OX=9606 GN=GNA14 PE=1 SV=1 | O95837 | GNA14_HUMAN | 42 kDa |
| Heat shock 70 kDa protein 1A OS=Homo sapiens OX=9606 GN=HSPA1A PE=1 SV=1 | P0DMV8 | HS71A_HUMAN (+1) | 70 kDa |
| Heat shock 70 kDa protein 4 OS=Homo sapiens OX=9606 GN=HSPA4 PE=1 SV=4 | P34932 | HSP74_HUMAN | 94 kDa |
| Heat shock cognate 71 kDa protein OS=Homo sapiens OX=9606 GN=HSPA8 PE=1 SV=1 | P11142 | HSP7C_HUMAN | 71 kDa |
| Heat shock protein 105 kDa OS=Homo sapiens OX=9606 GN=HSPH1 PE=1 SV=1 | Q92598 | HS105_HUMAN | 97 kDa |
| Heat shock protein beta-1 OS=Homo sapiens OX=9606 GN=HSPB1 PE=1 SV=2 | P04792 | HSPB1_HUMAN | 23 kDa |
| Heat shock protein HSP 90-alpha OS=Homo sapiens OX=9606 GN=HSP90AA1 PE=1 SV=5 | P07900 | HS90A_HUMAN | 85 kDa |
| Heat shock protein HSP 90-beta OS=Homo sapiens OX=9606 GN=HSP90AB1 PE=1 SV=4 | P08238 | HS90B_HUMAN | 83 kDa |
| Helicase MOV-10 OS=Homo sapiens OX=9606 GN=MOV10 PE=1 SV=2 | Q9HCE1 | MOV10_HUMAN | 114 kDa |
| Heme-binding protein 1 OS=Homo sapiens OX=9606 GN=HEBP1 PE=1 SV=1 | Q9NRV9 | HEBP1_HUMAN | 21 kDa |
| Hemoglobin subunit alpha OS=Homo sapiens OX=9606 GN=HBA2 PE=1 SV=2 | P69905 | HBA_HUMAN | 15 kDa |
| Hemoglobin subunit beta OS=Homo sapiens OX=9606 GN=HBB PE=1 SV=2 | P68871 | HBB_HUMAN | 16 kDa |
| Hemopexin OS=Homo sapiens OX=9606 GN=HPX PE=1 SV=2 | P02790 | HEMO_HUMAN | 52 kDa |
| Heparan sulfate glucosamine 3-O-sulfotransferase 1 OS=Homo sapiens OX=9606 GN=HS3ST1 PE=1 SV=1 | O14792 | HS3S1_HUMAN | 36 kDa |
| Heparin cofactor 2 OS=Homo sapiens OX=9606 GN=SERPIND1 PE=1 SV=3 | P05546 | HEP2_HUMAN | 57 kDa |
| Heterogeneous nuclear ribonucleoprotein A1 OS=Homo sapiens OX=9606 GN=HNRNPA1 PE=1 SV=5 | P09651 | ROA1_HUMAN | 39 kDa |
| Heterogeneous nuclear ribonucleoprotein D0 OS=Homo sapiens OX=9606 GN=HNRNPD PE=1 SV=1 | Q14103 | HNRPD_HUMAN | 38 kDa |
| Heterogeneous nuclear ribonucleoprotein H OS=Homo sapiens OX=9606 GN=HNRNPH1 PE=1 SV=4 | P31943 | HNRH1_HUMAN (+1) | 49 kDa |
| Heterogeneous nuclear ribonucleoprotein K OS=Homo sapiens OX=9606 GN=HNRNPK PE=1 SV=1 | P61978 | HNRPK_HUMAN | 51 kDa |
| Heterogeneous nuclear ribonucleoprotein U OS=Homo sapiens OX=9606 GN=HNRNPU PE=1 SV=6 | Q00839 | HNRPU_HUMAN | 91 kDa |
| Heterogeneous nuclear ribonucleoproteins C1/C2 OS=Homo sapiens OX=9606 GN=HNRNPC PE=1 SV=4 | P07910 | HNRPC_HUMAN | 34 kDa |
| High affinity cationic amino acid transporter 1 OS=Homo sapiens OX=9606 GN=SLC7A1 PE=1 SV=1 | P30825 | CTR1_HUMAN | 68 kDa |
| Histidine-rich glycoprotein OS=Homo sapiens OX=9606 GN=HRG PE=1 SV=1 | P04196 | HRG_HUMAN | 60 kDa |
| Histone deacetylase 6 OS=Homo sapiens OX=9606 GN=HDAC6 PE=1 SV=2 | Q9UBN7 | HDAC6_HUMAN | 131 kDa |
| Histone H2A type 1-B/E OS=Homo sapiens OX=9606 GN=H2AC8 PE=1 SV=2 | P04908 | H2A1B_HUMAN (+14) | 14 kDa |
| Histone H2B type 1-K OS=Homo sapiens OX=9606 GN=H2BC12 PE=1 SV=3 | O60814 | H2B1K_HUMAN (+13) | 14 kDa |
| Histone H3.1 OS=Homo sapiens OX=9606 GN=H3C12 PE=1 SV=2 | P68431 | H31_HUMAN (+3) | 15 kDa |
| Histone H4 OS=Homo sapiens OX=9606 GN=H4C16 PE=1 SV=2 | P62805 | H4_HUMAN | 11 kDa |
| HLA class I histocompatibility antigen, A alpha chain OS=Homo sapiens OX=9606 GN=HLA-A PE=1 SV=2 | P04439 | HLAA_HUMAN | 41 kDa |
| HLA class I histocompatibility antigen, B alpha chain OS=Homo sapiens OX=9606 GN=HLA-B PE=1 SV=3 | P01889 | HLAB_HUMAN | 40 kDa |
| HLA class I histocompatibility antigen, B alpha chain OS=Homo sapiens OX=9606 GN=HLA-B PE=1 SV=3 | P01889 | HLAB_HUMAN | 40 kDa |
| HLA class I histocompatibility antigen, C alpha chain OS=Homo sapiens OX=9606 GN=HLA-C PE=1 SV=3 | P10321 | HLAC_HUMAN | 41 kDa |
| Hornerin OS=Homo sapiens OX=9606 GN=HRNR PE=1 SV=2 | Q86YZ3 | HORN_HUMAN | 282 kDa |
| Huntingtin-interacting protein 1 OS=Homo sapiens OX=9606 GN=HIP1 PE=1 SV=5 | O00291 | HIP1_HUMAN | 116 kDa |
| Hydroxymethylglutaryl-CoA synthase, cytoplasmic OS=Homo sapiens OX=9606 GN=HMGCS1 PE=1 SV=2 | Q01581 | HMCS1_HUMAN | 57 kDa |
| Hypoxanthine-guanine phosphoribosyltransferase OS=Homo sapiens OX=9606 GN=HPRT1 PE=1 SV=2 | P00492 | HPRT_HUMAN | 25 kDa |
| Hypoxia up-regulated protein 1 OS=Homo sapiens OX=9606 GN=HYOU1 PE=1 SV=1 | Q9Y4L1 | HYOU1_HUMAN | 111 kDa |
| Immunity-related GTPase family Q protein OS=Homo sapiens OX=9606 GN=IRGQ PE=1 SV=1 | Q8WZA9 | IRGQ_HUMAN | 63 kDa |
| Immunoglobulin heavy constant delta OS=Homo sapiens OX=9606 GN=IGHD PE=1 SV=3 | P01880 | IGHD_HUMAN | 42 kDa |
| Immunoglobulin heavy constant mu OS=Homo sapiens OX=9606 GN=IGHM PE=1 SV=4 | P01871 | IGHM_HUMAN | 49 kDa |
| Immunoglobulin heavy variable 1-2 OS=Homo sapiens OX=9606 GN=IGHV1-2 PE=1 SV=2 | P23083 | HV102_HUMAN | 13 kDa |
| Immunoglobulin heavy variable 3-23 OS=Homo sapiens OX=9606 GN=IGHV3-23 PE=1 SV=2 | P01764 | HV323_HUMAN (+6) | 13 kDa |
| Immunoglobulin heavy variable 3-7 OS=Homo sapiens OX=9606 GN=IGHV3-7 PE=1 SV=2 | P01780 | HV307_HUMAN | 13 kDa |
| Immunoglobulin heavy variable 4-28 OS=Homo sapiens OX=9606 GN=IGHV4-28 PE=3 SV=1 | A0A0C4DH34 | HV428_HUMAN | 13 kDa |
| Immunoglobulin heavy variable 4-61 OS=Homo sapiens OX=9606 GN=IGHV4-61 PE=1 SV=1 | A0A0C4DH41 | HV461_HUMAN (+6) | 13 kDa |
| Immunoglobulin heavy variable 5-51 OS=Homo sapiens OX=9606 GN=IGHV5-51 PE=3 SV=1 | A0A0C4DH38 | HV551_HUMAN (+1) | 13 kDa |
| Immunoglobulin heavy variable 6-1 OS=Homo sapiens OX=9606 GN=IGHV6-1 PE=3 SV=1 | A0A0B4J1U7 | HV601_HUMAN | 13 kDa |
| Immunoglobulin J chain OS=Homo sapiens OX=9606 GN=JCHAIN PE=1 SV=4 | P01591 | IGJ_HUMAN | 18 kDa |
| Immunoglobulin kappa constant OS=Homo sapiens OX=9606 GN=IGKC PE=1 SV=2 | P01834 | IGKC_HUMAN | 12 kDa |
| Immunoglobulin kappa variable 1-6 OS=Homo sapiens OX=9606 GN=IGKV1-6 PE=3 SV=1 | A0A0C4DH72 | KV106_HUMAN | 13 kDa |
| Immunoglobulin kappa variable 1-8 OS=Homo sapiens OX=9606 GN=IGKV1-8 PE=3 SV=1 | A0A0C4DH67 | KV108_HUMAN | 13 kDa |
| Immunoglobulin kappa variable 1D-13 OS=Homo sapiens OX=9606 GN=IGKV1D-13 PE=3 SV=1 | A0A0B4J2D9 | KVD13_HUMAN (+2) | 13 kDa |
| Immunoglobulin kappa variable 1D-33 OS=Homo sapiens OX=9606 GN=IGKV1D-33 PE=1 SV=2 | P01593 | KVD33_HUMAN (+1) | 13 kDa |
| Immunoglobulin kappa variable 2-28 OS=Homo sapiens OX=9606 GN=IGKV2-28 PE=3 SV=1 | A0A075B6P5 | KV228_HUMAN (+3) | 13 kDa |
| Immunoglobulin kappa variable 3-20 OS=Homo sapiens OX=9606 GN=IGKV3-20 PE=1 SV=2 | P01619 | KV320_HUMAN | 13 kDa |
| Immunoglobulin kappa variable 3D-11 OS=Homo sapiens OX=9606 GN=IGKV3D-11 PE=3 SV=6 | A0A0A0MRZ8 | KVD11_HUMAN (+1) | 13 kDa |
| Immunoglobulin kappa variable 4-1 OS=Homo sapiens OX=9606 GN=IGKV4-1 PE=1 SV=1 | P06312 | KV401_HUMAN | 13 kDa |
| Immunoglobulin kappa variable 6-21 OS=Homo sapiens OX=9606 GN=IGKV6-21 PE=3 SV=1 | A0A0C4DH24 | KV621_HUMAN | 12 kDa |
| Immunoglobulin lambda variable 1-47 OS=Homo sapiens OX=9606 GN=IGLV1-47 PE=1 SV=2 | P01700 | LV147_HUMAN | 12 kDa |
| Immunoglobulin lambda variable 2-11 OS=Homo sapiens OX=9606 GN=IGLV2-11 PE=1 SV=2 | P01706 | LV211_HUMAN | 13 kDa |
| Immunoglobulin lambda variable 3-19 OS=Homo sapiens OX=9606 GN=IGLV3-19 PE=1 SV=2 | P01714 | LV319_HUMAN | 12 kDa |
| Immunoglobulin superfamily member 8 OS=Homo sapiens OX=9606 GN=IGSF8 PE=1 SV=1 | Q969P0 | IGSF8_HUMAN | 65 kDa |
| Importin subunit alpha-1 OS=Homo sapiens OX=9606 GN=KPNA2 PE=1 SV=1 | P52292 | IMA1_HUMAN | 58 kDa |
| Importin subunit alpha-3 OS=Homo sapiens OX=9606 GN=KPNA4 PE=1 SV=1 | O00629 | IMA3_HUMAN | 58 kDa |
| Importin subunit alpha-7 OS=Homo sapiens OX=9606 GN=KPNA6 PE=1 SV=1 | O60684 | IMA7_HUMAN | 60 kDa |
| Importin subunit beta-1 OS=Homo sapiens OX=9606 GN=KPNB1 PE=1 SV=2 | Q14974 | IMB1_HUMAN | 97 kDa |
| Importin-5 OS=Homo sapiens OX=9606 GN=IPO5 PE=1 SV=4 | O00410 | IPO5_HUMAN | 124 kDa |
| Importin-7 OS=Homo sapiens OX=9606 GN=IPO7 PE=1 SV=1 | O95373 | IPO7_HUMAN | 120 kDa |
| Inactive dipeptidyl peptidase 10 OS=Homo sapiens OX=9606 GN=DPP10 PE=1 SV=2 | Q8N608 | DPP10_HUMAN | 91 kDa |
| Inactive rhomboid protein 2 OS=Homo sapiens OX=9606 GN=RHBDF2 PE=1 SV=2 | Q6PJF5 | RHDF2_HUMAN | 97 kDa |
| Inactive tyrosine-protein kinase 7 OS=Homo sapiens OX=9606 GN=PTK7 PE=1 SV=2 | Q13308 | PTK7_HUMAN | 118 kDa |
| Influenza virus NS1A-binding protein OS=Homo sapiens OX=9606 GN=IVNS1ABP PE=1 SV=3 | Q9Y6Y0 | NS1BP_HUMAN | 72 kDa |
| Inhibin alpha chain OS=Homo sapiens OX=9606 GN=INHA PE=1 SV=1 | P05111 | INHA_HUMAN | 40 kDa |
| Inorganic pyrophosphatase OS=Homo sapiens OX=9606 GN=PPA1 PE=1 SV=2 | Q15181 | IPYR_HUMAN | 33 kDa |
| Inosine-5'-monophosphate dehydrogenase 2 OS=Homo sapiens OX=9606 GN=IMPDH2 PE=1 SV=2 | P12268 | IMDH2_HUMAN | 56 kDa |
| Insulin receptor OS=Homo sapiens OX=9606 GN=INSR PE=1 SV=4 | P06213 | INSR_HUMAN | 156 kDa |
| Insulin-like growth factor 1 receptor OS=Homo sapiens OX=9606 GN=IGF1R PE=1 SV=1 | P08069 | IGF1R_HUMAN | 155 kDa |
| Insulin-like growth factor-binding protein 5 OS=Homo sapiens OX=9606 GN=IGFBP5 PE=1 SV=1 | P24593 | IBP5_HUMAN | 31 kDa |
| Insulin-like growth factor-binding protein complex acid labile subunit OS=Homo sapiens OX=9606 GN=IGFALS PE=1 SV=1 | P35858 | ALS_HUMAN | 66 kDa |
| Integrin alpha-2 OS=Homo sapiens OX=9606 GN=ITGA2 PE=1 SV=2 | P17301 | ITA2_HUMAN | 129 kDa |
| Integrin alpha-6 OS=Homo sapiens OX=9606 GN=ITGA6 PE=1 SV=5 | P23229 | ITA6_HUMAN | 127 kDa |
| Integrin alpha-7 OS=Homo sapiens OX=9606 GN=ITGA7 PE=1 SV=3 | Q13683 | ITA7_HUMAN | 129 kDa |
| Integrin alpha-9 OS=Homo sapiens OX=9606 GN=ITGA9 PE=1 SV=2 | Q13797 | ITA9_HUMAN | 114 kDa |
| Integrin alpha-V OS=Homo sapiens OX=9606 GN=ITGAV PE=1 SV=2 | P06756 | ITAV_HUMAN | 116 kDa |
| Integrin beta-1 OS=Homo sapiens OX=9606 GN=ITGB1 PE=1 SV=2 | P05556 | ITB1_HUMAN | 88 kDa |
| Integrin-linked protein kinase OS=Homo sapiens OX=9606 GN=ILK PE=1 SV=2 | Q13418 | ILK_HUMAN | 51 kDa |
| Inter-alpha-trypsin inhibitor heavy chain H1 OS=Homo sapiens OX=9606 GN=ITIH1 PE=1 SV=3 | P19827 | ITIH1_HUMAN | 101 kDa |
| Inter-alpha-trypsin inhibitor heavy chain H2 OS=Homo sapiens OX=9606 GN=ITIH2 PE=1 SV=2 | P19823 | ITIH2_HUMAN | 106 kDa |
| Inter-alpha-trypsin inhibitor heavy chain H3 OS=Homo sapiens OX=9606 GN=ITIH3 PE=1 SV=2 | Q06033 | ITIH3_HUMAN | 100 kDa |
| Inter-alpha-trypsin inhibitor heavy chain H4 OS=Homo sapiens OX=9606 GN=ITIH4 PE=1 SV=4 | Q14624 | ITIH4_HUMAN | 103 kDa |
| Interleukin-6 receptor subunit beta OS=Homo sapiens OX=9606 GN=IL6ST PE=1 SV=2 | P40189 | IL6RB_HUMAN | 104 kDa |
| Isochorismatase domain-containing protein 1 OS=Homo sapiens OX=9606 GN=ISOC1 PE=1 SV=3 | Q96CN7 | ISOC1_HUMAN | 32 kDa |
| Isocitrate dehydrogenase [NADP] cytoplasmic OS=Homo sapiens OX=9606 GN=IDH1 PE=1 SV=2 | O75874 | IDHC_HUMAN | 47 kDa |
| Isoleucine--tRNA ligase, cytoplasmic OS=Homo sapiens OX=9606 GN=IARS1 PE=1 SV=2 | P41252 | SYIC_HUMAN | 145 kDa |
| Isopentenyl-diphosphate Delta-isomerase 1 OS=Homo sapiens OX=9606 GN=IDI1 PE=1 SV=2 | Q13907 | IDI1_HUMAN | 26 kDa |
| IST1 homolog OS=Homo sapiens OX=9606 GN=IST1 PE=1 SV=1 | P53990 | IST1_HUMAN | 40 kDa |
| Junction plakoglobin OS=Homo sapiens OX=9606 GN=JUP PE=1 SV=3 | P14923 | PLAK_HUMAN | 82 kDa |
| Junctional adhesion molecule A OS=Homo sapiens OX=9606 GN=F11R PE=1 SV=1 | Q9Y624 | JAM1_HUMAN | 33 kDa |
| Kallikrein-10 OS=Homo sapiens OX=9606 GN=KLK10 PE=1 SV=3 | O43240 | KLK10_HUMAN | 30 kDa |
| Keratin, type I cytoskeletal 13 OS=Homo sapiens OX=9606 GN=KRT13 PE=1 SV=4 | P13646 | K1C13_HUMAN | 50 kDa |
| Keratin, type I cytoskeletal 9 OS=Homo sapiens OX=9606 GN=KRT9 PE=1 SV=3 | P35527 | K1C9_HUMAN | 62 kDa |
| Keratin, type II cytoskeletal 4 OS=Homo sapiens OX=9606 GN=KRT4 PE=1 SV=5 | P19013 | K2C4_HUMAN | 56 kDa |
| Keratin, type II cytoskeletal 78 OS=Homo sapiens OX=9606 GN=KRT78 PE=1 SV=2 | Q8N1N4 | K2C78_HUMAN | 57 kDa |
| Keratin, type II cytoskeletal 80 OS=Homo sapiens OX=9606 GN=KRT80 PE=1 SV=2 | Q6KB66 | K2C80_HUMAN | 51 kDa |
| Keratin, type II microfibrillar (Fragment) OS=Ovis aries OX=9940 PE=1 SV=1 | P02539 | K2M1_SHEEP | 13 kDa |
| Keratinocyte proline-rich protein OS=Homo sapiens OX=9606 GN=KPRP PE=1 SV=1 | Q5T749 | KPRP_HUMAN | 64 kDa |
| KIF-binding protein OS=Homo sapiens OX=9606 GN=KIFBP PE=1 SV=1 | Q96EK5 | KBP_HUMAN | 72 kDa |
| Kinase D-interacting substrate of 220 kDa OS=Homo sapiens OX=9606 GN=KIDINS220 PE=1 SV=3 | Q9ULH0 | KDIS_HUMAN | 197 kDa |
| Kinesin light chain 4 OS=Homo sapiens OX=9606 GN=KLC4 PE=1 SV=3 | Q9NSK0 | KLC4_HUMAN | 69 kDa |
| Kinesin-1 heavy chain OS=Homo sapiens OX=9606 GN=KIF5B PE=1 SV=1 | P33176 | KINH_HUMAN | 110 kDa |
| Kinesin-like protein KIF23 OS=Homo sapiens OX=9606 GN=KIF23 PE=1 SV=3 | Q02241 | KIF23_HUMAN | 110 kDa |
| Kininogen-1 OS=Homo sapiens OX=9606 GN=KNG1 PE=1 SV=2 | P01042 | KNG1_HUMAN | 72 kDa |
| Kynurenine--oxoglutarate transaminase 3 OS=Homo sapiens OX=9606 GN=KYAT3 PE=1 SV=1 | Q6YP21 | KAT3_HUMAN | 51 kDa |
| Lactadherin OS=Homo sapiens OX=9606 GN=MFGE8 PE=1 SV=3 | Q08431 | MFGM_HUMAN | 43 kDa |
| Laminin subunit alpha-1 OS=Homo sapiens OX=9606 GN=LAMA1 PE=1 SV=2 | P25391 | LAMA1_HUMAN | 337 kDa |
| Laminin subunit alpha-3 OS=Homo sapiens OX=9606 GN=LAMA3 PE=1 SV=3 | Q16787 | LAMA3_HUMAN | 367 kDa |
| Laminin subunit beta-2 OS=Homo sapiens OX=9606 GN=LAMB2 PE=1 SV=2 | P55268 | LAMB2_HUMAN | 196 kDa |
| Laminin subunit gamma-1 OS=Homo sapiens OX=9606 GN=LAMC1 PE=1 SV=3 | P11047 | LAMC1_HUMAN | 178 kDa |
| Lanosterol 14-alpha demethylase OS=Homo sapiens OX=9606 GN=CYP51A1 PE=1 SV=4 | Q16850 | CP51A_HUMAN | 57 kDa |
| Lanosterol synthase OS=Homo sapiens OX=9606 GN=LSS PE=1 SV=1 | P48449 | LSS_HUMAN | 83 kDa |
| Large neutral amino acids transporter small subunit 1 OS=Homo sapiens OX=9606 GN=SLC7A5 PE=1 SV=2 | Q01650 | LAT1_HUMAN | 55 kDa |
| Large neutral amino acids transporter small subunit 2 OS=Homo sapiens OX=9606 GN=SLC7A8 PE=1 SV=1 | Q9UHI5 | LAT2_HUMAN | 58 kDa |
| Lethal(2) giant larvae protein homolog 1 OS=Homo sapiens OX=9606 GN=LLGL1 PE=1 SV=3 | Q15334 | L2GL1_HUMAN | 115 kDa |
| Leucine carboxyl methyltransferase 1 OS=Homo sapiens OX=9606 GN=LCMT1 PE=1 SV=2 | Q9UIC8 | LCMT1_HUMAN | 38 kDa |
| Leucine-rich alpha-2-glycoprotein OS=Homo sapiens OX=9606 GN=LRG1 PE=1 SV=2 | P02750 | A2GL_HUMAN | 38 kDa |
| Leucine-rich repeat and calponin homology domain-containing protein 2 OS=Homo sapiens OX=9606 GN=LRCH2 PE=1 SV=2 | Q5VUJ6 | LRCH2_HUMAN | 85 kDa |
| Leucine-rich repeat-containing protein 47 OS=Homo sapiens OX=9606 GN=LRRC47 PE=1 SV=1 | Q8N1G4 | LRC47_HUMAN | 63 kDa |
| Leucine--tRNA ligase, cytoplasmic OS=Homo sapiens OX=9606 GN=LARS1 PE=1 SV=2 | Q9P2J5 | SYLC_HUMAN | 134 kDa |
| Leukocyte elastase inhibitor OS=Homo sapiens OX=9606 GN=SERPINB1 PE=1 SV=1 | P30740 | ILEU_HUMAN | 43 kDa |
| LHFPL tetraspan subfamily member 2 protein OS=Homo sapiens OX=9606 GN=LHFPL2 PE=1 SV=2 | Q6ZUX7 | LHPL2_HUMAN | 24 kDa |
| Lipocalin-1 OS=Homo sapiens OX=9606 GN=LCN1 PE=1 SV=1 | P31025 | LCN1_HUMAN (+1) | 19 kDa |
| Lipolysis-stimulated lipoprotein receptor OS=Homo sapiens OX=9606 GN=LSR PE=1 SV=4 | Q86X29 | LSR_HUMAN | 71 kDa |
| Lipoma-preferred partner OS=Homo sapiens OX=9606 GN=LPP PE=1 SV=1 | Q93052 | LPP_HUMAN | 66 kDa |
| Lipopolysaccharide-binding protein OS=Homo sapiens OX=9606 GN=LBP PE=1 SV=3 | P18428 | LBP_HUMAN | 53 kDa |
| Lipoprotein lipase OS=Homo sapiens OX=9606 GN=LPL PE=1 SV=1 | P06858 | LIPL_HUMAN | 53 kDa |
| Liprin-alpha-1 OS=Homo sapiens OX=9606 GN=PPFIA1 PE=1 SV=1 | Q13136 | LIPA1_HUMAN | 136 kDa |
| L-lactate dehydrogenase A chain OS=Homo sapiens OX=9606 GN=LDHA PE=1 SV=2 | P00338 | LDHA_HUMAN | 37 kDa |
| L-lactate dehydrogenase B chain OS=Homo sapiens OX=9606 GN=LDHB PE=1 SV=2 | P07195 | LDHB_HUMAN | 37 kDa |
| Long-chain-fatty-acid--CoA ligase 4 OS=Homo sapiens OX=9606 GN=ACSL4 PE=1 SV=2 | O60488 | ACSL4_HUMAN | 79 kDa |
| Low-density lipoprotein receptor OS=Homo sapiens OX=9606 GN=LDLR PE=1 SV=1 | P01130 | LDLR_HUMAN | 95 kDa |
| Low-density lipoprotein receptor-related protein 8 OS=Homo sapiens OX=9606 GN=LRP8 PE=1 SV=4 | Q14114 | LRP8_HUMAN | 106 kDa |
| Lutropin-choriogonadotropic hormone receptor OS=Homo sapiens OX=9606 GN=LHCGR PE=1 SV=4 | P22888 | LSHR_HUMAN | 79 kDa |
| Lymphocyte function-associated antigen 3 OS=Homo sapiens OX=9606 GN=CD58 PE=1 SV=1 | P19256 | LFA3_HUMAN | 28 kDa |
| Lysine--tRNA ligase OS=Homo sapiens OX=9606 GN=KARS1 PE=1 SV=3 | Q15046 | SYK_HUMAN | 68 kDa |
| Lysosome membrane protein 2 OS=Homo sapiens OX=9606 GN=SCARB2 PE=1 SV=2 | Q14108 | SCRB2_HUMAN | 54 kDa |
| Lysosome-associated membrane glycoprotein 1 OS=Homo sapiens OX=9606 GN=LAMP1 PE=1 SV=3 | P11279 | LAMP1_HUMAN | 45 kDa |
| Lysosome-associated membrane glycoprotein 2 OS=Homo sapiens OX=9606 GN=LAMP2 PE=1 SV=2 | P13473 | LAMP2_HUMAN | 45 kDa |
| Macrophage migration inhibitory factor OS=Homo sapiens OX=9606 GN=MIF PE=1 SV=4 | P14174 | MIF_HUMAN | 12 kDa |
| Macrophage-capping protein OS=Homo sapiens OX=9606 GN=CAPG PE=1 SV=2 | P40121 | CAPG_HUMAN | 38 kDa |
| Major vault protein OS=Homo sapiens OX=9606 GN=MVP PE=1 SV=4 | Q14764 | MVP_HUMAN | 99 kDa |
| Malate dehydrogenase, cytoplasmic OS=Homo sapiens OX=9606 GN=MDH1 PE=1 SV=4 | P40925 | MDHC_HUMAN | 36 kDa |
| Malectin OS=Homo sapiens OX=9606 GN=MLEC PE=1 SV=1 | Q14165 | MLEC_HUMAN | 32 kDa |
| Mannan-binding lectin serine protease 1 OS=Homo sapiens OX=9606 GN=MASP1 PE=1 SV=3 | P48740 | MASP1_HUMAN | 79 kDa |
| Mannan-binding lectin serine protease 2 OS=Homo sapiens OX=9606 GN=MASP2 PE=1 SV=4 | O00187 | MASP2_HUMAN | 76 kDa |
| Mannose-1-phosphate guanyltransferase alpha OS=Homo sapiens OX=9606 GN=GMPPA PE=1 SV=1 | Q96IJ6 | GMPPA_HUMAN | 46 kDa |
| Mannose-1-phosphate guanyltransferase beta OS=Homo sapiens OX=9606 GN=GMPPB PE=1 SV=2 | Q9Y5P6 | GMPPB_HUMAN | 40 kDa |
| MAP/microtubule affinity-regulating kinase 3 OS=Homo sapiens OX=9606 GN=MARK3 PE=1 SV=5 | P27448 | MARK3_HUMAN | 84 kDa |
| MARCKS-related protein OS=Homo sapiens OX=9606 GN=MARCKSL1 PE=1 SV=2 | P49006 | MRP_HUMAN | 20 kDa |
| Matrix Gla protein OS=Homo sapiens OX=9606 GN=MGP PE=1 SV=2 | P08493 | MGP_HUMAN | 12 kDa |
| Matrix metalloproteinase-14 OS=Homo sapiens OX=9606 GN=MMP14 PE=1 SV=3 | P50281 | MMP14_HUMAN | 66 kDa |
| Melanoma-associated antigen D2 OS=Homo sapiens OX=9606 GN=MAGED2 PE=1 SV=2 | Q9UNF1 | MAGD2_HUMAN | 65 kDa |
| Membrane cofactor protein OS=Homo sapiens OX=9606 GN=CD46 PE=1 SV=3 | P15529 | MCP_HUMAN | 44 kDa |
| Membrane primary amine oxidase OS=Homo sapiens OX=9606 GN=AOC3 PE=1 SV=3 | Q16853 | AOC3_HUMAN | 85 kDa |
| Membrane-associated progesterone receptor component 1 OS=Homo sapiens OX=9606 GN=PGRMC1 PE=1 SV=3 | O00264 | PGRC1_HUMAN | 22 kDa |
| Membrane-associated progesterone receptor component 2 OS=Homo sapiens OX=9606 GN=PGRMC2 PE=1 SV=1 | O15173 | PGRC2_HUMAN | 24 kDa |
| Metal cation symporter ZIP14 OS=Homo sapiens OX=9606 GN=SLC39A14 PE=1 SV=3 | Q15043 | S39AE_HUMAN | 54 kDa |
| Metal transporter CNNM3 OS=Homo sapiens OX=9606 GN=CNNM3 PE=1 SV=1 | Q8NE01 | CNNM3_HUMAN | 76 kDa |
| Metalloreductase STEAP1 OS=Homo sapiens OX=9606 GN=STEAP1 PE=1 SV=1 | Q9UHE8 | STEA1_HUMAN | 40 kDa |
| Methionine adenosyltransferase 2 subunit beta OS=Homo sapiens OX=9606 GN=MAT2B PE=1 SV=1 | Q9NZL9 | MAT2B_HUMAN | 38 kDa |
| Methionine--tRNA ligase, cytoplasmic OS=Homo sapiens OX=9606 GN=MARS1 PE=1 SV=2 | P56192 | SYMC_HUMAN | 101 kDa |
| Methylsterol monooxygenase 1 OS=Homo sapiens OX=9606 GN=MSMO1 PE=1 SV=1 | Q15800 | MSMO1_HUMAN | 35 kDa |
| Mevalonate kinase OS=Homo sapiens OX=9606 GN=MVK PE=1 SV=1 | Q03426 | KIME_HUMAN | 42 kDa |
| MICOS complex subunit MIC60 OS=Homo sapiens OX=9606 GN=IMMT PE=1 SV=1 | Q16891 | MIC60_HUMAN | 84 kDa |
| Microtubule-actin cross-linking factor 1, isoforms 1/2/3/4/5 OS=Homo sapiens OX=9606 GN=MACF1 PE=1 SV=4 | Q9UPN3 | MACF1_HUMAN | 838 kDa |
| Midkine OS=Homo sapiens OX=9606 GN=MDK PE=1 SV=1 | P21741 | MK_HUMAN | 16 kDa |
| Mitochondrial 2-oxoglutarate/malate carrier protein OS=Homo sapiens OX=9606 GN=SLC25A11 PE=1 SV=3 | Q02978 | M2OM_HUMAN | 34 kDa |
| Mitogen-activated protein kinase 1 OS=Homo sapiens OX=9606 GN=MAPK1 PE=1 SV=3 | P28482 | MK01_HUMAN | 41 kDa |
| Mitogen-activated protein kinase 3 OS=Homo sapiens OX=9606 GN=MAPK3 PE=1 SV=4 | P27361 | MK03_HUMAN | 43 kDa |
| MOB kinase activator 1B OS=Homo sapiens OX=9606 GN=MOB1B PE=1 SV=3 | Q7L9L4 | MOB1B_HUMAN (+1) | 25 kDa |
| Moesin OS=Homo sapiens OX=9606 GN=MSN PE=1 SV=3 | P26038 | MOES_HUMAN | 68 kDa |
| Monocarboxylate transporter 1 OS=Homo sapiens OX=9606 GN=SLC16A1 PE=1 SV=3 | P53985 | MOT1_HUMAN | 54 kDa |
| Monocarboxylate transporter 4 OS=Homo sapiens OX=9606 GN=SLC16A3 PE=1 SV=1 | O15427 | MOT4_HUMAN | 49 kDa |
| Mothers against decapentaplegic homolog 9 OS=Homo sapiens OX=9606 GN=SMAD9 PE=1 SV=1 | O15198 | SMAD9_HUMAN (+1) | 52 kDa |
| MTOR-associated protein MEAK7 OS=Homo sapiens OX=9606 GN=MEAK7 PE=1 SV=2 | Q6P9B6 | MEAK7_HUMAN | 51 kDa |
| Mucin-19 OS=Homo sapiens OX=9606 GN=MUC19 PE=1 SV=3 | Q7Z5P9 | MUC19_HUMAN | 805 kDa |
| Multidrug and toxin extrusion protein 1 OS=Homo sapiens OX=9606 GN=SLC47A1 PE=1 SV=1 | Q96FL8 | S47A1_HUMAN | 62 kDa |
| Multidrug resistance-associated protein 1 OS=Homo sapiens OX=9606 GN=ABCC1 PE=1 SV=3 | P33527 | MRP1_HUMAN | 172 kDa |
| Multivesicular body subunit 12A OS=Homo sapiens OX=9606 GN=MVB12A PE=1 SV=1 | Q96EY5 | MB12A_HUMAN | 29 kDa |
| Myelin protein zero-like protein 1 OS=Homo sapiens OX=9606 GN=MPZL1 PE=1 SV=1 | O95297 | MPZL1_HUMAN | 29 kDa |
| Myeloid-derived growth factor OS=Homo sapiens OX=9606 GN=MYDGF PE=1 SV=1 | Q969H8 | MYDGF_HUMAN | 19 kDa |
| Myosin light chain 1/3, skeletal muscle isoform OS=Homo sapiens OX=9606 GN=MYL1 PE=1 SV=3 | P05976 | MYL1_HUMAN (+1) | 21 kDa |
| Myosin light polypeptide 6 OS=Homo sapiens OX=9606 GN=MYL6 PE=1 SV=2 | P60660 | MYL6_HUMAN | 17 kDa |
| Myosin regulatory light chain 12B OS=Homo sapiens OX=9606 GN=MYL12B PE=1 SV=2 | O14950 | ML12B_HUMAN (+1) | 20 kDa |
| Myosin-10 OS=Homo sapiens OX=9606 GN=MYH10 PE=1 SV=3 | P35580 | MYH10_HUMAN | 229 kDa |
| Myosin-9 OS=Homo sapiens OX=9606 GN=MYH9 PE=1 SV=4 | P35579 | MYH9_HUMAN | 227 kDa |
| Myotrophin OS=Homo sapiens OX=9606 GN=MTPN PE=1 SV=2 | P58546 | MTPN_HUMAN | 13 kDa |
| Myristoylated alanine-rich C-kinase substrate OS=Homo sapiens OX=9606 GN=MARCKS PE=1 SV=4 | P29966 | MARCS_HUMAN | 32 kDa |
| N(G),N(G)-dimethylarginine dimethylaminohydrolase 1 OS=Homo sapiens OX=9606 GN=DDAH1 PE=1 SV=3 | O94760 | DDAH1_HUMAN | 31 kDa |
| N(G),N(G)-dimethylarginine dimethylaminohydrolase 2 OS=Homo sapiens OX=9606 GN=DDAH2 PE=1 SV=1 | O95865 | DDAH2_HUMAN | 30 kDa |
| Na(+)/H(+) exchange regulatory cofactor NHE-RF1 OS=Homo sapiens OX=9606 GN=SLC9A3R1 PE=1 SV=4 | O14745 | NHRF1_HUMAN | 39 kDa |
| NADH dehydrogenase [ubiquinone] iron-sulfur protein 3, mitochondrial OS=Homo sapiens OX=9606 GN=NDUFS3 PE=1 SV=1 | O75489 | NDUS3_HUMAN | 30 kDa |
| NADH-cytochrome b5 reductase 3 OS=Homo sapiens OX=9606 GN=CYB5R3 PE=1 SV=3 | P00387 | NB5R3_HUMAN | 34 kDa |
| NADH-ubiquinone oxidoreductase 75 kDa subunit, mitochondrial OS=Homo sapiens OX=9606 GN=NDUFS1 PE=1 SV=3 | P28331 | NDUS1_HUMAN | 79 kDa |
| NADPH:adrenodoxin oxidoreductase, mitochondrial OS=Homo sapiens OX=9606 GN=FDXR PE=1 SV=3 | P22570 | ADRO_HUMAN | 54 kDa |
| NADPH--cytochrome P450 reductase OS=Homo sapiens OX=9606 GN=POR PE=1 SV=2 | P16435 | NCPR_HUMAN | 77 kDa |
| Nascent polypeptide-associated complex subunit alpha, muscle-specific form OS=Homo sapiens OX=9606 GN=NACA PE=1 SV=1 | E9PAV3 | NACAM_HUMAN (+1) | 205 kDa |
| Nck-associated protein 1 OS=Homo sapiens OX=9606 GN=NCKAP1 PE=1 SV=1 | Q9Y2A7 | NCKP1_HUMAN | 129 kDa |
| Nectin-2 OS=Homo sapiens OX=9606 GN=NECTIN2 PE=1 SV=1 | Q92692 | NECT2_HUMAN | 58 kDa |
| NEDD8-conjugating enzyme Ubc12 OS=Homo sapiens OX=9606 GN=UBE2M PE=1 SV=1 | P61081 | UBC12_HUMAN | 21 kDa |
| Neurabin-1 OS=Homo sapiens OX=9606 GN=PPP1R9A PE=1 SV=2 | Q9ULJ8 | NEB1_HUMAN | 123 kDa |
| Neurogenic locus notch homolog protein 2 OS=Homo sapiens OX=9606 GN=NOTCH2 PE=1 SV=3 | Q04721 | NOTC2_HUMAN | 265 kDa |
| Neuropilin-1 OS=Homo sapiens OX=9606 GN=NRP1 PE=1 SV=3 | O14786 | NRP1_HUMAN | 103 kDa |
| Neuroplastin OS=Homo sapiens OX=9606 GN=NPTN PE=1 SV=2 | Q9Y639 | NPTN_HUMAN | 44 kDa |
| Neutral alpha-glucosidase AB OS=Homo sapiens OX=9606 GN=GANAB PE=1 SV=3 | Q14697 | GANAB_HUMAN | 107 kDa |
| Neutral amino acid transporter A OS=Homo sapiens OX=9606 GN=SLC1A4 PE=1 SV=1 | P43007 | SATT_HUMAN | 56 kDa |
| Neutral amino acid transporter B(0) OS=Homo sapiens OX=9606 GN=SLC1A5 PE=1 SV=2 | Q15758 | AAAT_HUMAN | 57 kDa |
| Neutrophil gelatinase-associated lipocalin OS=Homo sapiens OX=9606 GN=LCN2 PE=1 SV=2 | P80188 | NGAL_HUMAN | 23 kDa |
| Nicastrin OS=Homo sapiens OX=9606 GN=NCSTN PE=1 SV=2 | Q92542 | NICA_HUMAN | 78 kDa |
| Nicotinamide phosphoribosyltransferase OS=Homo sapiens OX=9606 GN=NAMPT PE=1 SV=1 | P43490 | NAMPT_HUMAN | 56 kDa |
| Nidogen-1 OS=Homo sapiens OX=9606 GN=NID1 PE=1 SV=3 | P14543 | NID1_HUMAN | 136 kDa |
| Nidogen-2 OS=Homo sapiens OX=9606 GN=NID2 PE=1 SV=3 | Q14112 | NID2_HUMAN | 151 kDa |
| Nodal modulator 3 OS=Homo sapiens OX=9606 GN=NOMO3 PE=1 SV=2 | P69849 | NOMO3_HUMAN (+2) | 134 kDa |
| Non-histone chromosomal protein HMG-17 OS=Homo sapiens OX=9606 GN=HMGN2 PE=1 SV=3 | P05204 | HMGN2_HUMAN | 9 kDa |
| NPC intracellular cholesterol transporter 1 OS=Homo sapiens OX=9606 GN=NPC1 PE=1 SV=2 | O15118 | NPC1_HUMAN | 142 kDa |
| Nuclear migration protein nudC OS=Homo sapiens OX=9606 GN=NUDC PE=1 SV=1 | Q9Y266 | NUDC_HUMAN | 38 kDa |
| Nucleolin OS=Homo sapiens OX=9606 GN=NCL PE=1 SV=3 | P19338 | NUCL_HUMAN | 77 kDa |
| Nucleophosmin OS=Homo sapiens OX=9606 GN=NPM1 PE=1 SV=2 | P06748 | NPM_HUMAN | 33 kDa |
| Nucleoside diphosphate kinase B OS=Homo sapiens OX=9606 GN=NME2 PE=1 SV=1 | P22392 | NDKB_HUMAN | 17 kDa |
| Nucleosome assembly protein 1-like 1 OS=Homo sapiens OX=9606 GN=NAP1L1 PE=1 SV=1 | P55209 | NP1L1_HUMAN | 45 kDa |
| Nucleosome assembly protein 1-like 4 OS=Homo sapiens OX=9606 GN=NAP1L4 PE=1 SV=1 | Q99733 | NP1L4_HUMAN | 43 kDa |
| Obg-like ATPase 1 OS=Homo sapiens OX=9606 GN=OLA1 PE=1 SV=2 | Q9NTK5 | OLA1_HUMAN | 45 kDa |
| Obscurin-like protein 1 OS=Homo sapiens OX=9606 GN=OBSL1 PE=1 SV=4 | O75147 | OBSL1_HUMAN | 207 kDa |
| Olfactomedin-4 OS=Homo sapiens OX=9606 GN=OLFM4 PE=1 SV=1 | Q6UX06 | OLFM4_HUMAN | 57 kDa |
| OTU domain-containing protein 7B OS=Homo sapiens OX=9606 GN=OTUD7B PE=1 SV=1 | Q6GQQ9 | OTU7B_HUMAN | 93 kDa |
| Parkinson disease protein 7 OS=Homo sapiens OX=9606 GN=PARK7 PE=1 SV=2 | Q99497 | PARK7_HUMAN | 20 kDa |
| PDZ domain-containing protein GIPC1 OS=Homo sapiens OX=9606 GN=GIPC1 PE=1 SV=2 | O14908 | GIPC1_HUMAN | 36 kDa |
| Peptidyl-glycine alpha-amidating monooxygenase OS=Homo sapiens OX=9606 GN=PAM PE=1 SV=2 | P19021 | AMD_HUMAN | 108 kDa |
| Peptidyl-prolyl cis-trans isomerase A OS=Homo sapiens OX=9606 GN=PPIA PE=1 SV=2 | P62937 | PPIA_HUMAN | 18 kDa |
| Peptidyl-prolyl cis-trans isomerase B OS=Homo sapiens OX=9606 GN=PPIB PE=1 SV=2 | P23284 | PPIB_HUMAN | 24 kDa |
| Peptidyl-prolyl cis-trans isomerase FKBP1A OS=Homo sapiens OX=9606 GN=FKBP1A PE=1 SV=2 | P62942 | FKB1A_HUMAN | 12 kDa |
| Peptidyl-prolyl cis-trans isomerase FKBP4 OS=Homo sapiens OX=9606 GN=FKBP4 PE=1 SV=3 | Q02790 | FKBP4_HUMAN | 52 kDa |
| Perilipin-3 OS=Homo sapiens OX=9606 GN=PLIN3 PE=1 SV=3 | O60664 | PLIN3_HUMAN | 47 kDa |
| Peripheral plasma membrane protein CASK OS=Homo sapiens OX=9606 GN=CASK PE=1 SV=3 | O14936 | CSKP_HUMAN | 105 kDa |
| Periplakin OS=Homo sapiens OX=9606 GN=PPL PE=1 SV=4 | O60437 | PEPL_HUMAN | 205 kDa |
| Peroxidasin homolog OS=Homo sapiens OX=9606 GN=PXDN PE=1 SV=2 | Q92626 | PXDN_HUMAN | 165 kDa |
| Peroxiredoxin-2 OS=Homo sapiens OX=9606 GN=PRDX2 PE=1 SV=5 | P32119 | PRDX2_HUMAN | 22 kDa |
| Peroxiredoxin-5, mitochondrial OS=Homo sapiens OX=9606 GN=PRDX5 PE=1 SV=4 | P30044 | PRDX5_HUMAN | 22 kDa |
| Peroxiredoxin-6 OS=Homo sapiens OX=9606 GN=PRDX6 PE=1 SV=3 | P30041 | PRDX6_HUMAN | 25 kDa |
| Peroxisomal multifunctional enzyme type 2 OS=Homo sapiens OX=9606 GN=HSD17B4 PE=1 SV=3 | P51659 | DHB4_HUMAN | 80 kDa |
| PH and SEC7 domain-containing protein 3 OS=Homo sapiens OX=9606 GN=PSD3 PE=1 SV=2 | Q9NYI0 | PSD3_HUMAN | 116 kDa |
| Phosphate carrier protein, mitochondrial OS=Homo sapiens OX=9606 GN=SLC25A3 PE=1 SV=2 | Q00325 | MPCP_HUMAN | 40 kDa |
| Phosphate-regulating neutral endopeptidase PHEX OS=Homo sapiens OX=9606 GN=PHEX PE=1 SV=1 | P78562 | PHEX_HUMAN | 86 kDa |
| Phosphatidylethanolamine-binding protein 1 OS=Homo sapiens OX=9606 GN=PEBP1 PE=1 SV=3 | P30086 | PEBP1_HUMAN | 21 kDa |
| Phosphatidylinositol 4-kinase alpha OS=Homo sapiens OX=9606 GN=PI4KA PE=1 SV=4 | P42356 | PI4KA_HUMAN | 237 kDa |
| Phosphatidylinositol 5-phosphate 4-kinase type-2 alpha OS=Homo sapiens OX=9606 GN=PIP4K2A PE=1 SV=2 | P48426 | PI42A_HUMAN | 46 kDa |
| Phosphatidylinositol 5-phosphate 4-kinase type-2 beta OS=Homo sapiens OX=9606 GN=PIP4K2B PE=1 SV=1 | P78356 | PI42B_HUMAN | 47 kDa |
| Phosphatidylinositol transfer protein beta isoform OS=Homo sapiens OX=9606 GN=PITPNB PE=1 SV=2 | P48739 | PIPNB_HUMAN | 32 kDa |
| Phosphatidylinositol-glycan-specific phospholipase D OS=Homo sapiens OX=9606 GN=GPLD1 PE=1 SV=3 | P80108 | PHLD_HUMAN | 92 kDa |
| Phosphoacetylglucosamine mutase OS=Homo sapiens OX=9606 GN=PGM3 PE=1 SV=1 | O95394 | AGM1_HUMAN | 60 kDa |
| Phosphoglucomutase-1 OS=Homo sapiens OX=9606 GN=PGM1 PE=1 SV=3 | P36871 | PGM1_HUMAN | 61 kDa |
| Phosphoglycerate kinase 1 OS=Homo sapiens OX=9606 GN=PGK1 PE=1 SV=3 | P00558 | PGK1_HUMAN | 45 kDa |
| Phosphoglycerate mutase 1 OS=Homo sapiens OX=9606 GN=PGAM1 PE=1 SV=2 | P18669 | PGAM1_HUMAN | 29 kDa |
| Phospholipid-transporting ATPase IG OS=Homo sapiens OX=9606 GN=ATP11C PE=1 SV=3 | Q8NB49 | AT11C_HUMAN | 129 kDa |
| Phospholipid-transporting ATPase IH OS=Homo sapiens OX=9606 GN=ATP11A PE=1 SV=3 | P98196 | AT11A_HUMAN | 130 kDa |
| Phosphomevalonate kinase OS=Homo sapiens OX=9606 GN=PMVK PE=1 SV=3 | Q15126 | PMVK_HUMAN | 22 kDa |
| Phosphoribosylformylglycinamidine synthase OS=Homo sapiens OX=9606 GN=PFAS PE=1 SV=4 | O15067 | PUR4_HUMAN | 145 kDa |
| Pigment epithelium-derived factor OS=Homo sapiens OX=9606 GN=SERPINF1 PE=1 SV=4 | P36955 | PEDF_HUMAN | 46 kDa |
| Plakophilin-1 OS=Homo sapiens OX=9606 GN=PKP1 PE=1 SV=2 | Q13835 | PKP1_HUMAN | 83 kDa |
| Plasma kallikrein OS=Homo sapiens OX=9606 GN=KLKB1 PE=1 SV=1 | P03952 | KLKB1_HUMAN | 71 kDa |
| Plasma membrane calcium-transporting ATPase 4 OS=Homo sapiens OX=9606 GN=ATP2B4 PE=1 SV=2 | P23634 | AT2B4_HUMAN | 138 kDa |
| Plasma protease C1 inhibitor OS=Homo sapiens OX=9606 GN=SERPING1 PE=1 SV=2 | P05155 | IC1_HUMAN | 55 kDa |
| Plasminogen activator inhibitor 1 RNA-binding protein OS=Homo sapiens OX=9606 GN=SERBP1 PE=1 SV=2 | Q8NC51 | PAIRB_HUMAN | 45 kDa |
| Plasminogen OS=Homo sapiens OX=9606 GN=PLG PE=1 SV=2 | P00747 | PLMN_HUMAN | 91 kDa |
| Plastin-1 OS=Homo sapiens OX=9606 GN=PLS1 PE=1 SV=2 | Q14651 | PLSI_HUMAN | 70 kDa |
| Plastin-3 OS=Homo sapiens OX=9606 GN=PLS3 PE=1 SV=4 | P13797 | PLST_HUMAN | 71 kDa |
| Platelet-activating factor acetylhydrolase IB subunit alpha1 OS=Homo sapiens OX=9606 GN=PAFAH1B3 PE=1 SV=1 | Q15102 | PA1B3_HUMAN | 26 kDa |
| Platelet-activating factor acetylhydrolase IB subunit alpha2 OS=Homo sapiens OX=9606 GN=PAFAH1B2 PE=1 SV=1 | P68402 | PA1B2_HUMAN | 26 kDa |
| Platelet-activating factor acetylhydrolase IB subunit beta OS=Homo sapiens OX=9606 GN=PAFAH1B1 PE=1 SV=2 | P43034 | LIS1_HUMAN | 47 kDa |
| Plectin OS=Homo sapiens OX=9606 GN=PLEC PE=1 SV=3 | Q15149 | PLEC_HUMAN | 532 kDa |
| Plexin domain-containing protein 2 OS=Homo sapiens OX=9606 GN=PLXDC2 PE=1 SV=1 | Q6UX71 | PXDC2_HUMAN | 60 kDa |
| Plexin-A1 OS=Homo sapiens OX=9606 GN=PLXNA1 PE=1 SV=3 | Q9UIW2 | PLXA1_HUMAN | 211 kDa |
| Plexin-B2 OS=Homo sapiens OX=9606 GN=PLXNB2 PE=1 SV=3 | O15031 | PLXB2_HUMAN | 205 kDa |
| Poly(rC)-binding protein 1 OS=Homo sapiens OX=9606 GN=PCBP1 PE=1 SV=2 | Q15365 | PCBP1_HUMAN | 37 kDa |
| Poly(rC)-binding protein 2 OS=Homo sapiens OX=9606 GN=PCBP2 PE=1 SV=1 | Q15366 | PCBP2_HUMAN | 39 kDa |
| Polyadenylate-binding protein 1 OS=Homo sapiens OX=9606 GN=PABPC1 PE=1 SV=2 | P11940 | PABP1_HUMAN | 71 kDa |
| Polyubiquitin-B OS=Homo sapiens OX=9606 GN=UBB PE=1 SV=1 | P0CG47 | UBB_HUMAN (+3) | 26 kDa |
| Polyunsaturated fatty acid lipoxygenase ALOX12 OS=Homo sapiens OX=9606 GN=ALOX12 PE=1 SV=4 | P18054 | LOX12_HUMAN | 76 kDa |
| PRAME family member 4 OS=Homo sapiens OX=9606 GN=PRAMEF4 PE=2 SV=5 | O60810 | PRAM4_HUMAN | 55 kDa |
| Prefoldin subunit 2 OS=Homo sapiens OX=9606 GN=PFDN2 PE=1 SV=1 | Q9UHV9 | PFD2_HUMAN | 17 kDa |
| Pre-mRNA-processing-splicing factor 8 OS=Homo sapiens OX=9606 GN=PRPF8 PE=1 SV=2 | Q6P2Q9 | PRP8_HUMAN | 274 kDa |
| Presenilin-1 OS=Homo sapiens OX=9606 GN=PSEN1 PE=1 SV=1 | P49768 | PSN1_HUMAN | 53 kDa |
| Probable non-functional immunoglobulin kappa variable 2D-24 OS=Homo sapiens OX=9606 GN=IGKV2D-24 PE=1 SV=1 | A0A075B6R9 | KVD24_HUMAN (+1) | 13 kDa |
| Probable non-functional immunoglobulin kappa variable 3-7 OS=Homo sapiens OX=9606 GN=IGKV3-7 PE=1 SV=1 | A0A075B6H7 | KV37_HUMAN | 13 kDa |
| Probable ubiquitin carboxyl-terminal hydrolase FAF-X OS=Homo sapiens OX=9606 GN=USP9X PE=1 SV=4 | Q93008 | USP9X_HUMAN | 290 kDa |
| Profilin-1 OS=Homo sapiens OX=9606 GN=PFN1 PE=1 SV=2 | P07737 | PROF1_HUMAN | 15 kDa |
| Profilin-2 OS=Homo sapiens OX=9606 GN=PFN2 PE=1 SV=3 | P35080 | PROF2_HUMAN | 15 kDa |
| Programmed cell death 6-interacting protein OS=Homo sapiens OX=9606 GN=PDCD6IP PE=1 SV=1 | Q8WUM4 | PDC6I_HUMAN | 96 kDa |
| Programmed cell death protein 10 OS=Homo sapiens OX=9606 GN=PDCD10 PE=1 SV=1 | Q9BUL8 | PDC10_HUMAN | 25 kDa |
| Proliferating cell nuclear antigen OS=Homo sapiens OX=9606 GN=PCNA PE=1 SV=1 | P12004 | PCNA_HUMAN | 29 kDa |
| Proliferation-associated protein 2G4 OS=Homo sapiens OX=9606 GN=PA2G4 PE=1 SV=3 | Q9UQ80 | PA2G4_HUMAN | 44 kDa |
| Proline-rich transmembrane protein 1 OS=Homo sapiens OX=9606 GN=PRRT1 PE=1 SV=2 | Q99946 | PRRT1_HUMAN | 31 kDa |
| Properdin OS=Homo sapiens OX=9606 GN=CFP PE=1 SV=2 | P27918 | PROP_HUMAN | 51 kDa |
| Prostaglandin E synthase 3 OS=Homo sapiens OX=9606 GN=PTGES3 PE=1 SV=1 | Q15185 | TEBP_HUMAN | 19 kDa |
| Prostaglandin F2 receptor negative regulator OS=Homo sapiens OX=9606 GN=PTGFRN PE=1 SV=2 | Q9P2B2 | FPRP_HUMAN | 99 kDa |
| Prostatic acid phosphatase OS=Homo sapiens OX=9606 GN=ACP3 PE=1 SV=3 | P15309 | PPAP_HUMAN | 45 kDa |
| Proteasomal ubiquitin receptor ADRM1 OS=Homo sapiens OX=9606 GN=ADRM1 PE=1 SV=2 | Q16186 | ADRM1_HUMAN | 42 kDa |
| Proteasome activator complex subunit 1 OS=Homo sapiens OX=9606 GN=PSME1 PE=1 SV=1 | Q06323 | PSME1_HUMAN | 29 kDa |
| Proteasome activator complex subunit 2 OS=Homo sapiens OX=9606 GN=PSME2 PE=1 SV=4 | Q9UL46 | PSME2_HUMAN | 27 kDa |
| Proteasome adapter and scaffold protein ECM29 OS=Homo sapiens OX=9606 GN=ECPAS PE=1 SV=2 | Q5VYK3 | ECM29_HUMAN | 204 kDa |
| Proteasome subunit alpha type-1 OS=Homo sapiens OX=9606 GN=PSMA1 PE=1 SV=1 | P25786 | PSA1_HUMAN | 30 kDa |
| Proteasome subunit alpha type-2 OS=Homo sapiens OX=9606 GN=PSMA2 PE=1 SV=2 | P25787 | PSA2_HUMAN | 26 kDa |
| Proteasome subunit alpha type-3 OS=Homo sapiens OX=9606 GN=PSMA3 PE=1 SV=2 | P25788 | PSA3_HUMAN | 28 kDa |
| Proteasome subunit alpha type-5 OS=Homo sapiens OX=9606 GN=PSMA5 PE=1 SV=3 | P28066 | PSA5_HUMAN | 26 kDa |
| Proteasome subunit alpha type-6 OS=Homo sapiens OX=9606 GN=PSMA6 PE=1 SV=1 | P60900 | PSA6_HUMAN | 27 kDa |
| Proteasome subunit alpha type-7 OS=Homo sapiens OX=9606 GN=PSMA7 PE=1 SV=1 | O14818 | PSA7_HUMAN (+1) | 28 kDa |
| Proteasome subunit beta type-1 OS=Homo sapiens OX=9606 GN=PSMB1 PE=1 SV=2 | P20618 | PSB1_HUMAN | 26 kDa |
| Proteasome subunit beta type-4 OS=Homo sapiens OX=9606 GN=PSMB4 PE=1 SV=4 | P28070 | PSB4_HUMAN | 29 kDa |
| Proteasome subunit beta type-5 OS=Homo sapiens OX=9606 GN=PSMB5 PE=1 SV=3 | P28074 | PSB5_HUMAN | 28 kDa |
| Proteasome subunit beta type-6 OS=Homo sapiens OX=9606 GN=PSMB6 PE=1 SV=4 | P28072 | PSB6_HUMAN | 25 kDa |
| Proteasome subunit beta type-8 OS=Homo sapiens OX=9606 GN=PSMB8 PE=1 SV=3 | P28062 | PSB8_HUMAN | 30 kDa |
| Proteasome subunit beta type-9 OS=Homo sapiens OX=9606 GN=PSMB9 PE=1 SV=2 | P28065 | PSB9_HUMAN | 23 kDa |
| Protein AMBP OS=Homo sapiens OX=9606 GN=AMBP PE=1 SV=1 | P02760 | AMBP_HUMAN | 39 kDa |
| Protein arginine N-methyltransferase 5 OS=Homo sapiens OX=9606 GN=PRMT5 PE=1 SV=4 | O14744 | ANM5_HUMAN | 73 kDa |
| Protein canopy homolog 2 OS=Homo sapiens OX=9606 GN=CNPY2 PE=1 SV=1 | Q9Y2B0 | CNPY2_HUMAN | 21 kDa |
| Protein disulfide-isomerase A3 OS=Homo sapiens OX=9606 GN=PDIA3 PE=1 SV=4 | P30101 | PDIA3_HUMAN | 57 kDa |
| Protein disulfide-isomerase A4 OS=Homo sapiens OX=9606 GN=PDIA4 PE=1 SV=2 | P13667 | PDIA4_HUMAN | 73 kDa |
| Protein disulfide-isomerase A6 OS=Homo sapiens OX=9606 GN=PDIA6 PE=1 SV=1 | Q15084 | PDIA6_HUMAN | 48 kDa |
| Protein disulfide-isomerase OS=Homo sapiens OX=9606 GN=P4HB PE=1 SV=3 | P07237 | PDIA1_HUMAN | 57 kDa |
| Protein EFR3 homolog A OS=Homo sapiens OX=9606 GN=EFR3A PE=1 SV=2 | Q14156 | EFR3A_HUMAN | 93 kDa |
| Protein enabled homolog OS=Homo sapiens OX=9606 GN=ENAH PE=1 SV=2 | Q8N8S7 | ENAH_HUMAN | 67 kDa |
| Protein FAM234A OS=Homo sapiens OX=9606 GN=FAM234A PE=1 SV=1 | Q9H0X4 | F234A_HUMAN | 60 kDa |
| Protein FAM3C OS=Homo sapiens OX=9606 GN=FAM3C PE=1 SV=1 | Q92520 | FAM3C_HUMAN | 25 kDa |
| Protein FAM98A OS=Homo sapiens OX=9606 GN=FAM98A PE=1 SV=2 | Q8NCA5 | FA98A_HUMAN | 55 kDa |
| Protein GOLM2 OS=Homo sapiens OX=9606 GN=GOLM2 PE=1 SV=2 | Q6P4E1 | GOLM2_HUMAN | 49 kDa |
| Protein Hook homolog 3 OS=Homo sapiens OX=9606 GN=HOOK3 PE=1 SV=2 | Q86VS8 | HOOK3_HUMAN | 83 kDa |
| Protein kinase C alpha type OS=Homo sapiens OX=9606 GN=PRKCA PE=1 SV=4 | P17252 | KPCA_HUMAN | 77 kDa |
| Protein lifeguard 3 OS=Homo sapiens OX=9606 GN=TMBIM1 PE=1 SV=2 | Q969X1 | LFG3_HUMAN | 35 kDa |
| Protein lin-7 homolog C OS=Homo sapiens OX=9606 GN=LIN7C PE=1 SV=1 | Q9NUP9 | LIN7C_HUMAN | 22 kDa |
| Protein NDRG1 OS=Homo sapiens OX=9606 GN=NDRG1 PE=1 SV=1 | Q92597 | NDRG1_HUMAN | 43 kDa |
| Protein NDRG2 OS=Homo sapiens OX=9606 GN=NDRG2 PE=1 SV=2 | Q9UN36 | NDRG2_HUMAN | 41 kDa |
| Protein NDRG3 OS=Homo sapiens OX=9606 GN=NDRG3 PE=1 SV=2 | Q9UGV2 | NDRG3_HUMAN | 41 kDa |
| Protein Niban 2 OS=Homo sapiens OX=9606 GN=NIBAN2 PE=1 SV=3 | Q96TA1 | NIBA2_HUMAN | 84 kDa |
| Protein PALS2 OS=Homo sapiens OX=9606 GN=PALS2 PE=1 SV=2 | Q9NZW5 | PALS2_HUMAN | 61 kDa |
| Protein phosphatase 1 regulatory subunit 12C OS=Homo sapiens OX=9606 GN=PPP1R12C PE=1 SV=1 | Q9BZL4 | PP12C_HUMAN | 85 kDa |
| Protein phosphatase 3 catalytic subunit alpha OS=Homo sapiens OX=9606 GN=PPP3CA PE=1 SV=1 | Q08209 | PP2BA_HUMAN | 59 kDa |
| Protein phosphatase methylesterase 1 OS=Homo sapiens OX=9606 GN=PPME1 PE=1 SV=3 | Q9Y570 | PPME1_HUMAN | 42 kDa |
| Protein RCC2 OS=Homo sapiens OX=9606 GN=RCC2 PE=1 SV=2 | Q9P258 | RCC2_HUMAN | 56 kDa |
| Protein S100-A11 OS=Homo sapiens OX=9606 GN=S100A11 PE=1 SV=2 | P31949 | S10AB_HUMAN | 12 kDa |
| Protein S100-A13 OS=Homo sapiens OX=9606 GN=S100A13 PE=1 SV=1 | Q99584 | S10AD_HUMAN | 11 kDa |
| Protein S100-A16 OS=Homo sapiens OX=9606 GN=S100A16 PE=1 SV=1 | Q96FQ6 | S10AG_HUMAN | 12 kDa |
| Protein S100-A8 OS=Homo sapiens OX=9606 GN=S100A8 PE=1 SV=1 | P05109 | S10A8_HUMAN | 11 kDa |
| Protein S100-A9 OS=Homo sapiens OX=9606 GN=S100A9 PE=1 SV=1 | P06702 | S10A9_HUMAN | 13 kDa |
| Protein scribble homolog OS=Homo sapiens OX=9606 GN=SCRIB PE=1 SV=5 | Q14160 | SCRIB_HUMAN | 175 kDa |
| Protein SEC13 homolog OS=Homo sapiens OX=9606 GN=SEC13 PE=1 SV=3 | P55735 | SEC13_HUMAN | 36 kDa |
| Protein SGT1 homolog OS=Homo sapiens OX=9606 GN=SUGT1 PE=1 SV=3 | Q9Y2Z0 | SGT1_HUMAN | 41 kDa |
| Protein sidekick-1 OS=Homo sapiens OX=9606 GN=SDK1 PE=2 SV=3 | Q7Z5N4 | SDK1_HUMAN | 242 kDa |
| Protein TFG OS=Homo sapiens OX=9606 GN=TFG PE=1 SV=2 | Q92734 | TFG_HUMAN | 43 kDa |
| Protein transport protein Sec23A OS=Homo sapiens OX=9606 GN=SEC23A PE=1 SV=2 | Q15436 | SC23A_HUMAN | 86 kDa |
| Protein tweety homolog 3 OS=Homo sapiens OX=9606 GN=TTYH3 PE=1 SV=3 | Q9C0H2 | TTYH3_HUMAN | 58 kDa |
| Protein tyrosine phosphatase type IVA 2 OS=Homo sapiens OX=9606 GN=PTP4A2 PE=1 SV=1 | Q12974 | TP4A2_HUMAN (+1) | 19 kDa |
| Protein unc-13 homolog D OS=Homo sapiens OX=9606 GN=UNC13D PE=1 SV=1 | Q70J99 | UN13D_HUMAN | 123 kDa |
| Protein unc-45 homolog A OS=Homo sapiens OX=9606 GN=UNC45A PE=1 SV=1 | Q9H3U1 | UN45A_HUMAN | 103 kDa |
| Protein XRP2 OS=Homo sapiens OX=9606 GN=RP2 PE=1 SV=4 | O75695 | XRP2_HUMAN | 40 kDa |
| Protein-glutamine gamma-glutamyltransferase E OS=Homo sapiens OX=9606 GN=TGM3 PE=1 SV=4 | Q08188 | TGM3_HUMAN | 77 kDa |
| Protein-glutamine gamma-glutamyltransferase K OS=Homo sapiens OX=9606 GN=TGM1 PE=1 SV=4 | P22735 | TGM1_HUMAN | 90 kDa |
| Proteolipid protein 2 OS=Homo sapiens OX=9606 GN=PLP2 PE=1 SV=1 | Q04941 | PLP2_HUMAN | 17 kDa |
| Prothrombin OS=Homo sapiens OX=9606 GN=F2 PE=1 SV=2 | P00734 | THRB_HUMAN | 70 kDa |
| Prothymosin alpha OS=Homo sapiens OX=9606 GN=PTMA PE=1 SV=2 | P06454 | PTMA_HUMAN | 12 kDa |
| Protocadherin Fat 1 OS=Homo sapiens OX=9606 GN=FAT1 PE=1 SV=2 | Q14517 | FAT1_HUMAN | 506 kDa |
| Puromycin-sensitive aminopeptidase OS=Homo sapiens OX=9606 GN=NPEPPS PE=1 SV=2 | P55786 | PSA_HUMAN | 103 kDa |
| Pyruvate kinase PKM OS=Homo sapiens OX=9606 GN=PKM PE=1 SV=4 | P14618 | KPYM_HUMAN | 58 kDa |
| Quinone oxidoreductase OS=Homo sapiens OX=9606 GN=CRYZ PE=1 SV=1 | Q08257 | QOR_HUMAN | 35 kDa |
| Rab GDP dissociation inhibitor alpha OS=Homo sapiens OX=9606 GN=GDI1 PE=1 SV=2 | P31150 | GDIA_HUMAN | 51 kDa |
| Rab GDP dissociation inhibitor beta OS=Homo sapiens OX=9606 GN=GDI2 PE=1 SV=2 | P50395 | GDIB_HUMAN | 51 kDa |
| Rab11 family-interacting protein 5 OS=Homo sapiens OX=9606 GN=RAB11FIP5 PE=1 SV=1 | Q9BXF6 | RFIP5_HUMAN | 70 kDa |
| Rabankyrin-5 OS=Homo sapiens OX=9606 GN=ANKFY1 PE=1 SV=2 | Q9P2R3 | ANFY1_HUMAN | 128 kDa |
| Rac GTPase-activating protein 1 OS=Homo sapiens OX=9606 GN=RACGAP1 PE=1 SV=1 | Q9H0H5 | RGAP1_HUMAN | 71 kDa |
| RAC-alpha serine/threonine-protein kinase OS=Homo sapiens OX=9606 GN=AKT1 PE=1 SV=2 | P31749 | AKT1_HUMAN | 56 kDa |
| Radixin OS=Homo sapiens OX=9606 GN=RDX PE=1 SV=1 | P35241 | RADI_HUMAN | 69 kDa |
| Ran GTPase-activating protein 1 OS=Homo sapiens OX=9606 GN=RANGAP1 PE=1 SV=1 | P46060 | RAGP1_HUMAN | 64 kDa |
| Ran-specific GTPase-activating protein OS=Homo sapiens OX=9606 GN=RANBP1 PE=1 SV=1 | P43487 | RANG_HUMAN | 23 kDa |
| Rap1 GTPase-GDP dissociation stimulator 1 OS=Homo sapiens OX=9606 GN=RAP1GDS1 PE=1 SV=3 | P52306 | GDS1_HUMAN | 66 kDa |
| Rapamycin-insensitive companion of mTOR OS=Homo sapiens OX=9606 GN=RICTOR PE=1 SV=1 | Q6R327 | RICTR_HUMAN | 192 kDa |
| Ras GTPase-activating protein nGAP OS=Homo sapiens OX=9606 GN=RASAL2 PE=1 SV=2 | Q9UJF2 | NGAP_HUMAN | 129 kDa |
| Ras GTPase-activating-like protein IQGAP1 OS=Homo sapiens OX=9606 GN=IQGAP1 PE=1 SV=1 | P46940 | IQGA1_HUMAN | 189 kDa |
| Ras-related C3 botulinum toxin substrate 1 OS=Homo sapiens OX=9606 GN=RAC1 PE=1 SV=1 | P63000 | RAC1_HUMAN | 21 kDa |
| Ras-related protein Rab-11A OS=Homo sapiens OX=9606 GN=RAB11A PE=1 SV=3 | P62491 | RB11A_HUMAN (+1) | 24 kDa |
| Ras-related protein Rab-12 OS=Homo sapiens OX=9606 GN=RAB12 PE=1 SV=3 | Q6IQ22 | RAB12_HUMAN | 27 kDa |
| Ras-related protein Rab-13 OS=Homo sapiens OX=9606 GN=RAB13 PE=1 SV=1 | P51153 | RAB13_HUMAN | 23 kDa |
| Ras-related protein Rab-14 OS=Homo sapiens OX=9606 GN=RAB14 PE=1 SV=4 | P61106 | RAB14_HUMAN | 24 kDa |
| Ras-related protein Rab-18 OS=Homo sapiens OX=9606 GN=RAB18 PE=1 SV=1 | Q9NP72 | RAB18_HUMAN | 23 kDa |
| Ras-related protein Rab-21 OS=Homo sapiens OX=9606 GN=RAB21 PE=1 SV=3 | Q9UL25 | RAB21_HUMAN | 24 kDa |
| Ras-related protein Rab-22A OS=Homo sapiens OX=9606 GN=RAB22A PE=1 SV=2 | Q9UL26 | RB22A_HUMAN | 22 kDa |
| Ras-related protein Rab-2A OS=Homo sapiens OX=9606 GN=RAB2A PE=1 SV=1 | P61019 | RAB2A_HUMAN | 24 kDa |
| Ras-related protein Rab-34 OS=Homo sapiens OX=9606 GN=RAB34 PE=1 SV=1 | Q9BZG1 | RAB34_HUMAN | 29 kDa |
| Ras-related protein Rab-35 OS=Homo sapiens OX=9606 GN=RAB35 PE=1 SV=1 | Q15286 | RAB35_HUMAN | 23 kDa |
| Ras-related protein Rab-5A OS=Homo sapiens OX=9606 GN=RAB5A PE=1 SV=2 | P20339 | RAB5A_HUMAN | 24 kDa |
| Ras-related protein Rab-5B OS=Homo sapiens OX=9606 GN=RAB5B PE=1 SV=1 | P61020 | RAB5B_HUMAN | 24 kDa |
| Ras-related protein Rab-5C OS=Homo sapiens OX=9606 GN=RAB5C PE=1 SV=2 | P51148 | RAB5C_HUMAN | 23 kDa |
| Ras-related protein Rab-6A OS=Homo sapiens OX=9606 GN=RAB6A PE=1 SV=3 | P20340 | RAB6A_HUMAN | 24 kDa |
| Ras-related protein Rab-6C OS=Homo sapiens OX=9606 GN=RAB6C PE=1 SV=2 | Q9H0N0 | RAB6C_HUMAN | 28 kDa |
| Ras-related protein Rab-7a OS=Homo sapiens OX=9606 GN=RAB7A PE=1 SV=1 | P51149 | RAB7A_HUMAN | 23 kDa |
| Ras-related protein Rap-1b OS=Homo sapiens OX=9606 GN=RAP1B PE=1 SV=1 | P61224 | RAP1B_HUMAN | 21 kDa |
| Ras-related protein R-Ras2 OS=Homo sapiens OX=9606 GN=RRAS2 PE=1 SV=1 | P62070 | RRAS2_HUMAN | 23 kDa |
| Receptor of activated protein C kinase 1 OS=Homo sapiens OX=9606 GN=RACK1 PE=1 SV=3 | P63244 | RACK1_HUMAN | 35 kDa |
| Receptor-type tyrosine-protein phosphatase alpha OS=Homo sapiens OX=9606 GN=PTPRA PE=1 SV=3 | P18433 | PTPRA_HUMAN | 91 kDa |
| Receptor-type tyrosine-protein phosphatase gamma OS=Homo sapiens OX=9606 GN=PTPRG PE=1 SV=4 | P23470 | PTPRG_HUMAN | 162 kDa |
| Reticulocalbin-2 OS=Homo sapiens OX=9606 GN=RCN2 PE=1 SV=1 | Q14257 | RCN2_HUMAN | 37 kDa |
| Reticulon-4 OS=Homo sapiens OX=9606 GN=RTN4 PE=1 SV=2 | Q9NQC3 | RTN4_HUMAN | 130 kDa |
| Retinol-binding protein 1 OS=Homo sapiens OX=9606 GN=RBP1 PE=1 SV=2 | P09455 | RET1_HUMAN | 16 kDa |
| Rho GDP-dissociation inhibitor 1 OS=Homo sapiens OX=9606 GN=ARHGDIA PE=1 SV=3 | P52565 | GDIR1_HUMAN | 23 kDa |
| Rho GTPase-activating protein 1 OS=Homo sapiens OX=9606 GN=ARHGAP1 PE=1 SV=1 | Q07960 | RHG01_HUMAN | 50 kDa |
| Rho GTPase-activating protein 17 OS=Homo sapiens OX=9606 GN=ARHGAP17 PE=1 SV=1 | Q68EM7 | RHG17_HUMAN | 95 kDa |
| Rho guanine nucleotide exchange factor 1 OS=Homo sapiens OX=9606 GN=ARHGEF1 PE=1 SV=2 | Q92888 | ARHG1_HUMAN | 102 kDa |
| Rho-associated protein kinase 1 OS=Homo sapiens OX=9606 GN=ROCK1 PE=1 SV=1 | Q13464 | ROCK1_HUMAN | 158 kDa |
| Rho-associated protein kinase 2 OS=Homo sapiens OX=9606 GN=ROCK2 PE=1 SV=4 | O75116 | ROCK2_HUMAN | 161 kDa |
| Rho-related GTP-binding protein RhoG OS=Homo sapiens OX=9606 GN=RHOG PE=1 SV=1 | P84095 | RHOG_HUMAN | 21 kDa |
| Ribonuclease inhibitor OS=Homo sapiens OX=9606 GN=RNH1 PE=1 SV=2 | P13489 | RINI_HUMAN | 50 kDa |
| Ribose-phosphate pyrophosphokinase 3 OS=Homo sapiens OX=9606 GN=PRPS1L1 PE=1 SV=2 | P21108 | PRPS3_HUMAN | 35 kDa |
| RNA transcription, translation and transport factor protein OS=Homo sapiens OX=9606 GN=RTRAF PE=1 SV=1 | Q9Y224 | RTRAF_HUMAN | 28 kDa |
| RuvB-like 1 OS=Homo sapiens OX=9606 GN=RUVBL1 PE=1 SV=1 | Q9Y265 | RUVB1_HUMAN | 50 kDa |
| RuvB-like 2 OS=Homo sapiens OX=9606 GN=RUVBL2 PE=1 SV=3 | Q9Y230 | RUVB2_HUMAN | 51 kDa |
| S-adenosylhomocysteine hydrolase-like protein 1 OS=Homo sapiens OX=9606 GN=AHCYL1 PE=1 SV=2 | O43865 | SAHH2_HUMAN | 59 kDa |
| S-adenosylmethionine synthase isoform type-2 OS=Homo sapiens OX=9606 GN=MAT2A PE=1 SV=1 | P31153 | METK2_HUMAN | 44 kDa |
| Sarcoplasmic/endoplasmic reticulum calcium ATPase 2 OS=Homo sapiens OX=9606 GN=ATP2A2 PE=1 SV=1 | P16615 | AT2A2_HUMAN | 115 kDa |
| Scavenger receptor class B member 1 OS=Homo sapiens OX=9606 GN=SCARB1 PE=1 SV=1 | Q8WTV0 | SCRB1_HUMAN | 61 kDa |
| Secreted phosphoprotein 24 OS=Homo sapiens OX=9606 GN=SPP2 PE=1 SV=1 | Q13103 | SPP24_HUMAN | 24 kDa |
| Secretory carrier-associated membrane protein 2 OS=Homo sapiens OX=9606 GN=SCAMP2 PE=1 SV=2 | O15127 | SCAM2_HUMAN | 37 kDa |
| Secretory carrier-associated membrane protein 3 OS=Homo sapiens OX=9606 GN=SCAMP3 PE=1 SV=3 | O14828 | SCAM3_HUMAN | 38 kDa |
| Sedoheptulokinase OS=Homo sapiens OX=9606 GN=SHPK PE=1 SV=4 | Q9UHJ6 | SHPK_HUMAN | 52 kDa |
| Septin-10 OS=Homo sapiens OX=9606 GN=SEPTIN10 PE=1 SV=2 | Q9P0V9 | SEP10_HUMAN | 53 kDa |
| Septin-2 OS=Homo sapiens OX=9606 GN=SEPTIN2 PE=1 SV=1 | Q15019 | SEPT2_HUMAN | 41 kDa |
| Septin-7 OS=Homo sapiens OX=9606 GN=SEPTIN7 PE=1 SV=2 | Q16181 | SEPT7_HUMAN | 51 kDa |
| Septin-8 OS=Homo sapiens OX=9606 GN=SEPTIN8 PE=1 SV=4 | Q92599 | SEPT8_HUMAN | 56 kDa |
| Septin-9 OS=Homo sapiens OX=9606 GN=SEPTIN9 PE=1 SV=2 | Q9UHD8 | SEPT9_HUMAN | 65 kDa |
| Sequestosome-1 OS=Homo sapiens OX=9606 GN=SQSTM1 PE=1 SV=1 | Q13501 | SQSTM_HUMAN | 48 kDa |
| Serine incorporator 1 OS=Homo sapiens OX=9606 GN=SERINC1 PE=1 SV=1 | Q9NRX5 | SERC1_HUMAN | 50 kDa |
| Serine incorporator 5 OS=Homo sapiens OX=9606 GN=SERINC5 PE=2 SV=1 | Q86VE9 | SERC5_HUMAN | 47 kDa |
| Serine protease 23 OS=Homo sapiens OX=9606 GN=PRSS23 PE=1 SV=1 | O95084 | PRS23_HUMAN | 43 kDa |
| Serine protease HTRA1 OS=Homo sapiens OX=9606 GN=HTRA1 PE=1 SV=1 | Q92743 | HTRA1_HUMAN | 51 kDa |
| Serine protease HTRA3 OS=Homo sapiens OX=9606 GN=HTRA3 PE=1 SV=2 | P83110 | HTRA3_HUMAN | 49 kDa |
| Serine/threonine-protein kinase 25 OS=Homo sapiens OX=9606 GN=STK25 PE=1 SV=1 | O00506 | STK25_HUMAN (+2) | 48 kDa |
| Serine/threonine-protein kinase 3 OS=Homo sapiens OX=9606 GN=STK3 PE=1 SV=2 | Q13188 | STK3_HUMAN | 56 kDa |
| Serine/threonine-protein kinase MARK2 OS=Homo sapiens OX=9606 GN=MARK2 PE=1 SV=2 | Q7KZI7 | MARK2_HUMAN | 88 kDa |
| Serine/threonine-protein kinase MRCK beta OS=Homo sapiens OX=9606 GN=CDC42BPB PE=1 SV=2 | Q9Y5S2 | MRCKB_HUMAN | 194 kDa |
| Serine/threonine-protein kinase Nek9 OS=Homo sapiens OX=9606 GN=NEK9 PE=1 SV=2 | Q8TD19 | NEK9_HUMAN | 107 kDa |
| Serine/threonine-protein kinase OSR1 OS=Homo sapiens OX=9606 GN=OXSR1 PE=1 SV=1 | O95747 | OXSR1_HUMAN | 58 kDa |
| Serine/threonine-protein phosphatase 2A 55 kDa regulatory subunit B alpha isoform OS=Homo sapiens OX=9606 GN=PPP2R2A PE=1 SV=1 | P63151 | 2ABA_HUMAN | 52 kDa |
| Serine/threonine-protein phosphatase 2A 65 kDa regulatory subunit A alpha isoform OS=Homo sapiens OX=9606 GN=PPP2R1A PE=1 SV=4 | P30153 | 2AAA_HUMAN | 65 kDa |
| Serine/threonine-protein phosphatase 2A catalytic subunit beta isoform OS=Homo sapiens OX=9606 GN=PPP2CB PE=1 SV=1 | P62714 | PP2AB_HUMAN (+1) | 36 kDa |
| Serine/threonine-protein phosphatase 6 catalytic subunit OS=Homo sapiens OX=9606 GN=PPP6C PE=1 SV=1 | O00743 | PPP6_HUMAN | 35 kDa |
| Serine/threonine-protein phosphatase CPPED1 OS=Homo sapiens OX=9606 GN=CPPED1 PE=1 SV=3 | Q9BRF8 | CPPED_HUMAN | 36 kDa |
| Serine/threonine-protein phosphatase PP1-alpha catalytic subunit OS=Homo sapiens OX=9606 GN=PPP1CA PE=1 SV=1 | P62136 | PP1A_HUMAN | 38 kDa |
| Serine/threonine-protein phosphatase PP1-beta catalytic subunit OS=Homo sapiens OX=9606 GN=PPP1CB PE=1 SV=3 | P62140 | PP1B_HUMAN | 37 kDa |
| Serine/threonine-protein phosphatase PP1-gamma catalytic subunit OS=Homo sapiens OX=9606 GN=PPP1CC PE=1 SV=1 | P36873 | PP1G_HUMAN | 37 kDa |
| Serine-threonine kinase receptor-associated protein OS=Homo sapiens OX=9606 GN=STRAP PE=1 SV=1 | Q9Y3F4 | STRAP_HUMAN | 38 kDa |
| Serine--tRNA ligase, cytoplasmic OS=Homo sapiens OX=9606 GN=SARS1 PE=1 SV=3 | P49591 | SYSC_HUMAN | 59 kDa |
| Serpin B3 OS=Homo sapiens OX=9606 GN=SERPINB3 PE=1 SV=2 | P29508 | SPB3_HUMAN | 45 kDa |
| Serpin B6 OS=Homo sapiens OX=9606 GN=SERPINB6 PE=1 SV=3 | P35237 | SPB6_HUMAN | 43 kDa |
| Serpin H1 OS=Homo sapiens OX=9606 GN=SERPINH1 PE=1 SV=2 | P50454 | SERPH_HUMAN | 46 kDa |
| Serum amyloid A-1 protein OS=Homo sapiens OX=9606 GN=SAA1 PE=1 SV=1 | P0DJI8 | SAA1_HUMAN | 14 kDa |
| Serum amyloid P-component OS=Homo sapiens OX=9606 GN=APCS PE=1 SV=2 | P02743 | SAMP_HUMAN | 25 kDa |
| Serum paraoxonase/arylesterase 1 OS=Homo sapiens OX=9606 GN=PON1 PE=1 SV=3 | P27169 | PON1_HUMAN | 40 kDa |
| Serum paraoxonase/arylesterase 2 OS=Homo sapiens OX=9606 GN=PON2 PE=1 SV=4 | Q15165 | PON2_HUMAN | 39 kDa |
| S-formylglutathione hydrolase OS=Homo sapiens OX=9606 GN=ESD PE=1 SV=2 | P10768 | ESTD_HUMAN | 31 kDa |
| SH3 domain-binding glutamic acid-rich-like protein 3 OS=Homo sapiens OX=9606 GN=SH3BGRL3 PE=1 SV=1 | Q9H299 | SH3L3_HUMAN | 10 kDa |
| Short transient receptor potential channel 3 OS=Homo sapiens OX=9606 GN=TRPC3 PE=1 SV=4 | Q13507 | TRPC3_HUMAN | 106 kDa |
| Sialic acid synthase OS=Homo sapiens OX=9606 GN=NANS PE=1 SV=2 | Q9NR45 | SIAS_HUMAN | 40 kDa |
| SID1 transmembrane family member 2 OS=Homo sapiens OX=9606 GN=SIDT2 PE=1 SV=2 | Q8NBJ9 | SIDT2_HUMAN | 94 kDa |
| Sideroflexin-3 OS=Homo sapiens OX=9606 GN=SFXN3 PE=1 SV=3 | Q9BWM7 | SFXN3_HUMAN | 36 kDa |
| Skin-specific protein 32 OS=Homo sapiens OX=9606 GN=XP32 PE=1 SV=1 | Q5T750 | XP32_HUMAN | 26 kDa |
| SLIT and NTRK-like protein 4 OS=Homo sapiens OX=9606 GN=SLITRK4 PE=1 SV=1 | Q8IW52 | SLIK4_HUMAN | 94 kDa |
| Small glutamine-rich tetratricopeptide repeat-containing protein alpha OS=Homo sapiens OX=9606 GN=SGTA PE=1 SV=1 | O43765 | SGTA_HUMAN | 34 kDa |
| Small integral membrane protein 1 OS=Homo sapiens OX=9606 GN=SMIM1 PE=1 SV=1 | B2RUZ4 | SMIM1_HUMAN | 9 kDa |
| Sodium- and chloride-dependent taurine transporter OS=Homo sapiens OX=9606 GN=SLC6A6 PE=1 SV=2 | P31641 | SC6A6_HUMAN | 70 kDa |
| Sodium/potassium-transporting ATPase subunit alpha-1 OS=Homo sapiens OX=9606 GN=ATP1A1 PE=1 SV=1 | P05023 | AT1A1_HUMAN | 113 kDa |
| Sodium/potassium-transporting ATPase subunit beta-1 OS=Homo sapiens OX=9606 GN=ATP1B1 PE=1 SV=1 | P05026 | AT1B1_HUMAN | 35 kDa |
| Sodium/potassium-transporting ATPase subunit beta-3 OS=Homo sapiens OX=9606 GN=ATP1B3 PE=1 SV=1 | P54709 | AT1B3_HUMAN | 32 kDa |
| Sodium-coupled neutral amino acid symporter 2 OS=Homo sapiens OX=9606 GN=SLC38A2 PE=1 SV=2 | Q96QD8 | S38A2_HUMAN | 56 kDa |
| Sodium-dependent multivitamin transporter OS=Homo sapiens OX=9606 GN=SLC5A6 PE=1 SV=2 | Q9Y289 | SC5A6_HUMAN | 69 kDa |
| Solute carrier family 12 member 7 OS=Homo sapiens OX=9606 GN=SLC12A7 PE=1 SV=3 | Q9Y666 | S12A7_HUMAN | 119 kDa |
| Solute carrier family 2, facilitated glucose transporter member 1 OS=Homo sapiens OX=9606 GN=SLC2A1 PE=1 SV=2 | P11166 | GTR1_HUMAN | 54 kDa |
| Solute carrier family 2, facilitated glucose transporter member 3 OS=Homo sapiens OX=9606 GN=SLC2A3 PE=1 SV=1 | P11169 | GTR3_HUMAN (+1) | 54 kDa |
| Solute carrier family 23 member 2 OS=Homo sapiens OX=9606 GN=SLC23A2 PE=1 SV=1 | Q9UGH3 | S23A2_HUMAN | 70 kDa |
| Solute carrier family 26 member 6 OS=Homo sapiens OX=9606 GN=SLC26A6 PE=1 SV=1 | Q9BXS9 | S26A6_HUMAN | 83 kDa |
| Solute carrier family 35 member F6 OS=Homo sapiens OX=9606 GN=SLC35F6 PE=1 SV=1 | Q8N357 | S35F6_HUMAN | 40 kDa |
| Solute carrier organic anion transporter family member 3A1 OS=Homo sapiens OX=9606 GN=SLCO3A1 PE=1 SV=3 | Q9UIG8 | SO3A1_HUMAN | 77 kDa |
| Sorbitol dehydrogenase OS=Homo sapiens OX=9606 GN=SORD PE=1 SV=4 | Q00796 | DHSO_HUMAN | 38 kDa |
| Sorting nexin-1 OS=Homo sapiens OX=9606 GN=SNX1 PE=1 SV=3 | Q13596 | SNX1_HUMAN | 59 kDa |
| Sorting nexin-3 OS=Homo sapiens OX=9606 GN=SNX3 PE=1 SV=3 | O60493 | SNX3_HUMAN | 19 kDa |
| Sorting nexin-5 OS=Homo sapiens OX=9606 GN=SNX5 PE=1 SV=1 | Q9Y5X3 | SNX5_HUMAN | 47 kDa |
| Sorting nexin-6 OS=Homo sapiens OX=9606 GN=SNX6 PE=1 SV=1 | Q9UNH7 | SNX6_HUMAN | 47 kDa |
| SPARC OS=Homo sapiens OX=9606 GN=SPARC PE=1 SV=1 | P09486 | SPRC_HUMAN | 35 kDa |
| Spartin OS=Homo sapiens OX=9606 GN=SPART PE=1 SV=1 | Q8N0X7 | SPART_HUMAN | 73 kDa |
| Spectrin alpha chain, non-erythrocytic 1 OS=Homo sapiens OX=9606 GN=SPTAN1 PE=1 SV=3 | Q13813 | SPTN1_HUMAN | 285 kDa |
| Spectrin beta chain, non-erythrocytic 1 OS=Homo sapiens OX=9606 GN=SPTBN1 PE=1 SV=2 | Q01082 | SPTB2_HUMAN | 275 kDa |
| Spermidine synthase OS=Homo sapiens OX=9606 GN=SRM PE=1 SV=1 | P19623 | SPEE_HUMAN | 34 kDa |
| Spermine synthase OS=Homo sapiens OX=9606 GN=SMS PE=1 SV=2 | P52788 | SPSY_HUMAN | 41 kDa |
| S-phase kinase-associated protein 1 OS=Homo sapiens OX=9606 GN=SKP1 PE=1 SV=2 | P63208 | SKP1_HUMAN | 19 kDa |
| Spliceosome RNA helicase DDX39B OS=Homo sapiens OX=9606 GN=DDX39B PE=1 SV=1 | Q13838 | DX39B_HUMAN | 49 kDa |
| Squalene synthase OS=Homo sapiens OX=9606 GN=FDFT1 PE=1 SV=1 | P37268 | FDFT_HUMAN | 48 kDa |
| Staphylococcal nuclease domain-containing protein 1 OS=Homo sapiens OX=9606 GN=SND1 PE=1 SV=1 | Q7KZF4 | SND1_HUMAN | 102 kDa |
| Stomatin OS=Homo sapiens OX=9606 GN=STOM PE=1 SV=3 | P27105 | STOM_HUMAN | 32 kDa |
| Stress-70 protein, mitochondrial OS=Homo sapiens OX=9606 GN=HSPA9 PE=1 SV=2 | P38646 | GRP75_HUMAN | 74 kDa |
| Stress-induced-phosphoprotein 1 OS=Homo sapiens OX=9606 GN=STIP1 PE=1 SV=1 | P31948 | STIP1_HUMAN | 63 kDa |
| Striatin OS=Homo sapiens OX=9606 GN=STRN PE=1 SV=4 | O43815 | STRN_HUMAN | 86 kDa |
| Striatin-3 OS=Homo sapiens OX=9606 GN=STRN3 PE=1 SV=3 | Q13033 | STRN3_HUMAN | 87 kDa |
| Sulfhydryl oxidase 1 OS=Homo sapiens OX=9606 GN=QSOX1 PE=1 SV=3 | O00391 | QSOX1_HUMAN | 83 kDa |
| Superoxide dismutase [Mn], mitochondrial OS=Homo sapiens OX=9606 GN=SOD2 PE=1 SV=3 | P04179 | SODM_HUMAN | 25 kDa |
| Suprabasin OS=Homo sapiens OX=9606 GN=SBSN PE=1 SV=2 | Q6UWP8 | SBSN_HUMAN | 61 kDa |
| Sushi repeat-containing protein SRPX OS=Homo sapiens OX=9606 GN=SRPX PE=1 SV=1 | P78539 | SRPX_HUMAN | 52 kDa |
| Synaptic vesicle membrane protein VAT-1 homolog OS=Homo sapiens OX=9606 GN=VAT1 PE=1 SV=2 | Q99536 | VAT1_HUMAN | 42 kDa |
| Synaptophysin-like protein 1 OS=Homo sapiens OX=9606 GN=SYPL1 PE=1 SV=1 | Q16563 | SYPL1_HUMAN | 29 kDa |
| Synaptosomal-associated protein 23 OS=Homo sapiens OX=9606 GN=SNAP23 PE=1 SV=1 | O00161 | SNP23_HUMAN | 23 kDa |
| Syndecan-4 OS=Homo sapiens OX=9606 GN=SDC4 PE=1 SV=2 | P31431 | SDC4_HUMAN | 22 kDa |
| Syntaxin-12 OS=Homo sapiens OX=9606 GN=STX12 PE=1 SV=1 | Q86Y82 | STX12_HUMAN | 32 kDa |
| Syntaxin-3 OS=Homo sapiens OX=9606 GN=STX3 PE=1 SV=3 | Q13277 | STX3_HUMAN | 33 kDa |
| Syntaxin-4 OS=Homo sapiens OX=9606 GN=STX4 PE=1 SV=2 | Q12846 | STX4_HUMAN | 34 kDa |
| Syntaxin-7 OS=Homo sapiens OX=9606 GN=STX7 PE=1 SV=4 | O15400 | STX7_HUMAN | 30 kDa |
| Syntaxin-binding protein 2 OS=Homo sapiens OX=9606 GN=STXBP2 PE=1 SV=2 | Q15833 | STXB2_HUMAN | 66 kDa |
| Syntaxin-binding protein 3 OS=Homo sapiens OX=9606 GN=STXBP3 PE=1 SV=2 | O00186 | STXB3_HUMAN | 68 kDa |
| Syntenin-1 OS=Homo sapiens OX=9606 GN=SDCBP PE=1 SV=1 | O00560 | SDCB1_HUMAN | 32 kDa |
| Talin-1 OS=Homo sapiens OX=9606 GN=TLN1 PE=1 SV=3 | Q9Y490 | TLN1_HUMAN | 270 kDa |
| Target of rapamycin complex subunit LST8 OS=Homo sapiens OX=9606 GN=MLST8 PE=1 SV=1 | Q9BVC4 | LST8_HUMAN | 36 kDa |
| Tax1-binding protein 3 OS=Homo sapiens OX=9606 GN=TAX1BP3 PE=1 SV=2 | O14907 | TX1B3_HUMAN | 14 kDa |
| TBC1 domain family member 10A OS=Homo sapiens OX=9606 GN=TBC1D10A PE=1 SV=1 | Q9BXI6 | TB10A_HUMAN | 57 kDa |
| TBC1 domain family member 10B OS=Homo sapiens OX=9606 GN=TBC1D10B PE=1 SV=3 | Q4KMP7 | TB10B_HUMAN | 87 kDa |
| TBC1 domain family member 4 OS=Homo sapiens OX=9606 GN=TBC1D4 PE=1 SV=2 | O60343 | TBCD4_HUMAN | 147 kDa |
| T-complex protein 1 subunit alpha OS=Homo sapiens OX=9606 GN=TCP1 PE=1 SV=1 | P17987 | TCPA_HUMAN | 60 kDa |
| T-complex protein 1 subunit beta OS=Homo sapiens OX=9606 GN=CCT2 PE=1 SV=4 | P78371 | TCPB_HUMAN | 57 kDa |
| T-complex protein 1 subunit delta OS=Homo sapiens OX=9606 GN=CCT4 PE=1 SV=4 | P50991 | TCPD_HUMAN | 58 kDa |
| T-complex protein 1 subunit epsilon OS=Homo sapiens OX=9606 GN=CCT5 PE=1 SV=1 | P48643 | TCPE_HUMAN | 60 kDa |
| T-complex protein 1 subunit eta OS=Homo sapiens OX=9606 GN=CCT7 PE=1 SV=2 | Q99832 | TCPH_HUMAN | 59 kDa |
| T-complex protein 1 subunit gamma OS=Homo sapiens OX=9606 GN=CCT3 PE=1 SV=4 | P49368 | TCPG_HUMAN | 61 kDa |
| T-complex protein 1 subunit theta OS=Homo sapiens OX=9606 GN=CCT8 PE=1 SV=4 | P50990 | TCPQ_HUMAN | 60 kDa |
| T-complex protein 1 subunit zeta OS=Homo sapiens OX=9606 GN=CCT6A PE=1 SV=3 | P40227 | TCPZ_HUMAN | 58 kDa |
| Tenascin-X OS=Homo sapiens OX=9606 GN=TNXB PE=1 SV=5 | P22105 | TENX_HUMAN | 458 kDa |
| Testin OS=Homo sapiens OX=9606 GN=TES PE=1 SV=1 | Q9UGI8 | TES_HUMAN | 48 kDa |
| Tetranectin OS=Homo sapiens OX=9606 GN=CLEC3B PE=1 SV=3 | P05452 | TETN_HUMAN | 23 kDa |
| Tetraspanin-7 OS=Homo sapiens OX=9606 GN=TSPAN7 PE=1 SV=2 | P41732 | TSN7_HUMAN | 28 kDa |
| Thioredoxin domain-containing protein 17 OS=Homo sapiens OX=9606 GN=TXNDC17 PE=1 SV=1 | Q9BRA2 | TXD17_HUMAN | 14 kDa |
| Thioredoxin-like protein 1 OS=Homo sapiens OX=9606 GN=TXNL1 PE=1 SV=3 | O43396 | TXNL1_HUMAN | 32 kDa |
| Thioredoxin-related transmembrane protein 1 OS=Homo sapiens OX=9606 GN=TMX1 PE=1 SV=1 | Q9H3N1 | TMX1_HUMAN | 32 kDa |
| Threonine--tRNA ligase 1, cytoplasmic OS=Homo sapiens OX=9606 GN=TARS1 PE=1 SV=3 | P26639 | SYTC_HUMAN | 83 kDa |
| Thrombospondin-4 OS=Homo sapiens OX=9606 GN=THBS4 PE=1 SV=2 | P35443 | TSP4_HUMAN | 106 kDa |
| Thymosin beta-10 OS=Homo sapiens OX=9606 GN=TMSB10 PE=1 SV=2 | P63313 | TYB10_HUMAN | 5 kDa |
| Thyrotropin-releasing hormone-degrading ectoenzyme OS=Homo sapiens OX=9606 GN=TRHDE PE=2 SV=1 | Q9UKU6 | TRHDE_HUMAN | 117 kDa |
| Thyroxine-binding globulin OS=Homo sapiens OX=9606 GN=SERPINA7 PE=1 SV=2 | P05543 | THBG_HUMAN | 46 kDa |
| Tight junction protein ZO-1 OS=Homo sapiens OX=9606 GN=TJP1 PE=1 SV=3 | Q07157 | ZO1_HUMAN | 195 kDa |
| Toll-interacting protein OS=Homo sapiens OX=9606 GN=TOLLIP PE=1 SV=1 | Q9H0E2 | TOLIP_HUMAN | 30 kDa |
| Trafficking protein particle complex subunit 3 OS=Homo sapiens OX=9606 GN=TRAPPC3 PE=1 SV=1 | O43617 | TPPC3_HUMAN | 20 kDa |
| Transaldolase OS=Homo sapiens OX=9606 GN=TALDO1 PE=1 SV=2 | P37837 | TALDO_HUMAN | 38 kDa |
| Transcription intermediary factor 1-beta OS=Homo sapiens OX=9606 GN=TRIM28 PE=1 SV=5 | Q13263 | TIF1B_HUMAN | 89 kDa |
| Transferrin receptor protein 1 OS=Homo sapiens OX=9606 GN=TFRC PE=1 SV=2 | P02786 | TFR1_HUMAN | 85 kDa |
| Transforming growth factor beta receptor type 3 OS=Homo sapiens OX=9606 GN=TGFBR3 PE=1 SV=3 | Q03167 | TGBR3_HUMAN | 94 kDa |
| Transforming growth factor-beta-induced protein ig-h3 OS=Homo sapiens OX=9606 GN=TGFBI PE=1 SV=1 | Q15582 | BGH3_HUMAN | 75 kDa |
| Transforming protein RhoA OS=Homo sapiens OX=9606 GN=RHOA PE=1 SV=1 | P61586 | RHOA_HUMAN | 22 kDa |
| Transgelin-2 OS=Homo sapiens OX=9606 GN=TAGLN2 PE=1 SV=3 | P37802 | TAGL2_HUMAN | 22 kDa |
| Trans-Golgi network integral membrane protein 2 OS=Homo sapiens OX=9606 GN=TGOLN2 PE=1 SV=4 | O43493 | TGON2_HUMAN | 46 kDa |
| Transitional endoplasmic reticulum ATPase OS=Homo sapiens OX=9606 GN=VCP PE=1 SV=4 | P55072 | TERA_HUMAN | 89 kDa |
| Transketolase OS=Homo sapiens OX=9606 GN=TKT PE=1 SV=3 | P29401 | TKT_HUMAN | 68 kDa |
| Translationally-controlled tumor protein OS=Homo sapiens OX=9606 GN=TPT1 PE=1 SV=1 | P13693 | TCTP_HUMAN | 20 kDa |
| Translin OS=Homo sapiens OX=9606 GN=TSN PE=1 SV=1 | Q15631 | TSN_HUMAN | 26 kDa |
| Translocon-associated protein subunit alpha OS=Homo sapiens OX=9606 GN=SSR1 PE=1 SV=3 | P43307 | SSRA_HUMAN | 32 kDa |
| Translocon-associated protein subunit delta OS=Homo sapiens OX=9606 GN=SSR4 PE=1 SV=1 | P51571 | SSRD_HUMAN | 19 kDa |
| Transmembrane 4 L6 family member 1 OS=Homo sapiens OX=9606 GN=TM4SF1 PE=1 SV=1 | P30408 | T4S1_HUMAN | 22 kDa |
| Transmembrane 9 superfamily member 2 OS=Homo sapiens OX=9606 GN=TM9SF2 PE=1 SV=1 | Q99805 | TM9S2_HUMAN | 76 kDa |
| Transmembrane emp24 domain-containing protein 10 OS=Homo sapiens OX=9606 GN=TMED10 PE=1 SV=2 | P49755 | TMEDA_HUMAN | 25 kDa |
| Transmembrane protein 109 OS=Homo sapiens OX=9606 GN=TMEM109 PE=1 SV=1 | Q9BVC6 | TM109_HUMAN | 26 kDa |
| Transmembrane protein 200A OS=Homo sapiens OX=9606 GN=TMEM200A PE=1 SV=1 | Q86VY9 | T200A_HUMAN | 54 kDa |
| Transmembrane protein 59 OS=Homo sapiens OX=9606 GN=TMEM59 PE=1 SV=1 | Q9BXS4 | TMM59_HUMAN | 36 kDa |
| Transportin-1 OS=Homo sapiens OX=9606 GN=TNPO1 PE=1 SV=2 | Q92973 | TNPO1_HUMAN | 102 kDa |
| Transthyretin OS=Homo sapiens OX=9606 GN=TTR PE=1 SV=1 | P02766 | TTHY_HUMAN | 16 kDa |
| Trifunctional purine biosynthetic protein adenosine-3 OS=Homo sapiens OX=9606 GN=GART PE=1 SV=1 | P22102 | PUR2_HUMAN | 108 kDa |
| TRIO and F-actin-binding protein OS=Homo sapiens OX=9606 GN=TRIOBP PE=1 SV=3 | Q9H2D6 | TARA_HUMAN | 261 kDa |
| Triokinase/FMN cyclase OS=Homo sapiens OX=9606 GN=TKFC PE=1 SV=2 | Q3LXA3 | TKFC_HUMAN | 59 kDa |
| Triosephosphate isomerase OS=Homo sapiens OX=9606 GN=TPI1 PE=1 SV=4 | P60174 | TPIS_HUMAN | 27 kDa |
| Tripeptidyl-peptidase 2 OS=Homo sapiens OX=9606 GN=TPP2 PE=1 SV=4 | P29144 | TPP2_HUMAN | 138 kDa |
| Tropomodulin-3 OS=Homo sapiens OX=9606 GN=TMOD3 PE=1 SV=1 | Q9NYL9 | TMOD3_HUMAN | 40 kDa |
| TRPM8 channel-associated factor 1 OS=Homo sapiens OX=9606 GN=TCAF1 PE=1 SV=3 | Q9Y4C2 | TCAF1_HUMAN | 102 kDa |
| Trypsin OS=Sus scrofa OX=9823 PE=1 SV=1 | P00761 | TRYP_PIG | 24 kDa |
| Tsukushi OS=Homo sapiens OX=9606 GN=TSKU PE=1 SV=3 | Q8WUA8 | TSK_HUMAN | 38 kDa |
| Tuberin OS=Homo sapiens OX=9606 GN=TSC2 PE=1 SV=2 | P49815 | TSC2_HUMAN | 201 kDa |
| Tubulin-folding cofactor B OS=Homo sapiens OX=9606 GN=TBCB PE=1 SV=2 | Q99426 | TBCB_HUMAN | 27 kDa |
| Tubulin-specific chaperone A OS=Homo sapiens OX=9606 GN=TBCA PE=1 SV=3 | O75347 | TBCA_HUMAN | 13 kDa |
| Tubulin--tyrosine ligase-like protein 12 OS=Homo sapiens OX=9606 GN=TTLL12 PE=1 SV=2 | Q14166 | TTL12_HUMAN | 74 kDa |
| Tumor susceptibility gene 101 protein OS=Homo sapiens OX=9606 GN=TSG101 PE=1 SV=2 | Q99816 | TS101_HUMAN | 44 kDa |
| Twinfilin-1 OS=Homo sapiens OX=9606 GN=TWF1 PE=1 SV=3 | Q12792 | TWF1_HUMAN | 40 kDa |
| Tyrosine--tRNA ligase, cytoplasmic OS=Homo sapiens OX=9606 GN=YARS1 PE=1 SV=4 | P54577 | SYYC_HUMAN | 59 kDa |
| Ubiquitin carboxyl-terminal hydrolase 14 OS=Homo sapiens OX=9606 GN=USP14 PE=1 SV=3 | P54578 | UBP14_HUMAN | 56 kDa |
| Ubiquitin carboxyl-terminal hydrolase 15 OS=Homo sapiens OX=9606 GN=USP15 PE=1 SV=3 | Q9Y4E8 | UBP15_HUMAN | 112 kDa |
| Ubiquitin carboxyl-terminal hydrolase 5 OS=Homo sapiens OX=9606 GN=USP5 PE=1 SV=2 | P45974 | UBP5_HUMAN | 96 kDa |
| Ubiquitin carboxyl-terminal hydrolase isozyme L1 OS=Homo sapiens OX=9606 GN=UCHL1 PE=1 SV=2 | P09936 | UCHL1_HUMAN | 25 kDa |
| Ubiquitin thioesterase OTUB1 OS=Homo sapiens OX=9606 GN=OTUB1 PE=1 SV=2 | Q96FW1 | OTUB1_HUMAN | 31 kDa |
| Ubiquitin-conjugating enzyme E2 L3 OS=Homo sapiens OX=9606 GN=UBE2L3 PE=1 SV=1 | P68036 | UB2L3_HUMAN | 18 kDa |
| Ubiquitin-conjugating enzyme E2 N OS=Homo sapiens OX=9606 GN=UBE2N PE=1 SV=1 | P61088 | UBE2N_HUMAN (+1) | 17 kDa |
| Ubiquitin-like modifier-activating enzyme 1 OS=Homo sapiens OX=9606 GN=UBA1 PE=1 SV=3 | P22314 | UBA1_HUMAN | 118 kDa |
| Ubiquitin-like modifier-activating enzyme 6 OS=Homo sapiens OX=9606 GN=UBA6 PE=1 SV=1 | A0AVT1 | UBA6_HUMAN | 118 kDa |
| UDP-glucose 6-dehydrogenase OS=Homo sapiens OX=9606 GN=UGDH PE=1 SV=1 | O60701 | UGDH_HUMAN | 55 kDa |
| UDP-N-acetylglucosamine transporter OS=Homo sapiens OX=9606 GN=SLC35A3 PE=1 SV=1 | Q9Y2D2 | S35A3_HUMAN | 36 kDa |
| UMP-CMP kinase OS=Homo sapiens OX=9606 GN=CMPK1 PE=1 SV=3 | P30085 | KCY_HUMAN | 22 kDa |
| Uncharacterized protein CXorf38 OS=Homo sapiens OX=9606 GN=CXorf38 PE=1 SV=1 | Q8TB03 | CX038_HUMAN | 37 kDa |
| Unconventional myosin-Ib OS=Homo sapiens OX=9606 GN=MYO1B PE=1 SV=3 | O43795 | MYO1B_HUMAN | 132 kDa |
| Unconventional myosin-Ic OS=Homo sapiens OX=9606 GN=MYO1C PE=1 SV=4 | O00159 | MYO1C_HUMAN | 122 kDa |
| Unconventional myosin-Vb OS=Homo sapiens OX=9606 GN=MYO5B PE=1 SV=3 | Q9ULV0 | MYO5B_HUMAN | 214 kDa |
| Unconventional myosin-VI OS=Homo sapiens OX=9606 GN=MYO6 PE=1 SV=4 | Q9UM54 | MYO6_HUMAN | 150 kDa |
| Unconventional myosin-XVIIIa OS=Homo sapiens OX=9606 GN=MYO18A PE=1 SV=3 | Q92614 | MY18A_HUMAN | 233 kDa |
| Uridine 5'-monophosphate synthase OS=Homo sapiens OX=9606 GN=UMPS PE=1 SV=1 | P11172 | UMPS_HUMAN | 52 kDa |
| UTP--glucose-1-phosphate uridylyltransferase OS=Homo sapiens OX=9606 GN=UGP2 PE=1 SV=5 | Q16851 | UGPA_HUMAN | 57 kDa |
| UV excision repair protein RAD23 homolog B OS=Homo sapiens OX=9606 GN=RAD23B PE=1 SV=1 | P54727 | RD23B_HUMAN | 43 kDa |
| Vacuolar protein sorting-associated protein 26A OS=Homo sapiens OX=9606 GN=VPS26A PE=1 SV=2 | O75436 | VP26A_HUMAN | 38 kDa |
| Vacuolar protein sorting-associated protein 28 homolog OS=Homo sapiens OX=9606 GN=VPS28 PE=1 SV=1 | Q9UK41 | VPS28_HUMAN | 25 kDa |
| Vacuolar protein sorting-associated protein 29 OS=Homo sapiens OX=9606 GN=VPS29 PE=1 SV=1 | Q9UBQ0 | VPS29_HUMAN | 21 kDa |
| Vacuolar protein sorting-associated protein 35 OS=Homo sapiens OX=9606 GN=VPS35 PE=1 SV=2 | Q96QK1 | VPS35_HUMAN | 92 kDa |
| Vacuolar protein sorting-associated protein 37B OS=Homo sapiens OX=9606 GN=VPS37B PE=1 SV=1 | Q9H9H4 | VP37B_HUMAN | 31 kDa |
| Vacuolar protein sorting-associated protein 4B OS=Homo sapiens OX=9606 GN=VPS4B PE=1 SV=2 | O75351 | VPS4B_HUMAN | 49 kDa |
| Valine--tRNA ligase OS=Homo sapiens OX=9606 GN=VARS1 PE=1 SV=4 | P26640 | SYVC_HUMAN | 140 kDa |
| Vascular cell adhesion protein 1 OS=Homo sapiens OX=9606 GN=VCAM1 PE=1 SV=1 | P19320 | VCAM1_HUMAN | 81 kDa |
| Vasodilator-stimulated phosphoprotein OS=Homo sapiens OX=9606 GN=VASP PE=1 SV=3 | P50552 | VASP_HUMAN | 40 kDa |
| Versican core protein OS=Homo sapiens OX=9606 GN=VCAN PE=1 SV=3 | P13611 | CSPG2_HUMAN | 373 kDa |
| Vesicle-associated membrane protein-associated protein A OS=Homo sapiens OX=9606 GN=VAPA PE=1 SV=3 | Q9P0L0 | VAPA_HUMAN | 28 kDa |
| Vesicle-fusing ATPase OS=Homo sapiens OX=9606 GN=NSF PE=1 SV=3 | P46459 | NSF_HUMAN | 83 kDa |
| Vesicle-trafficking protein SEC22b OS=Homo sapiens OX=9606 GN=SEC22B PE=1 SV=5 | O75396 | SC22B_HUMAN | 25 kDa |
| Vigilin OS=Homo sapiens OX=9606 GN=HDLBP PE=1 SV=2 | Q00341 | VIGLN_HUMAN | 141 kDa |
| Vimentin OS=Homo sapiens OX=9606 GN=VIM PE=1 SV=4 | P08670 | VIME_HUMAN | 54 kDa |
| Vinculin OS=Homo sapiens OX=9606 GN=VCL PE=1 SV=4 | P18206 | VINC_HUMAN | 124 kDa |
| Vitamin D-binding protein OS=Homo sapiens OX=9606 GN=GC PE=1 SV=2 | P02774 | VTDB_HUMAN | 53 kDa |
| Vitamin K-dependent protein S OS=Homo sapiens OX=9606 GN=PROS1 PE=1 SV=1 | P07225 | PROS_HUMAN | 75 kDa |
| Vitronectin OS=Homo sapiens OX=9606 GN=VTN PE=1 SV=1 | P04004 | VTNC_HUMAN | 54 kDa |
| Voltage-dependent anion-selective channel protein 1 OS=Homo sapiens OX=9606 GN=VDAC1 PE=1 SV=2 | P21796 | VDAC1_HUMAN | 31 kDa |
| Voltage-dependent anion-selective channel protein 2 OS=Homo sapiens OX=9606 GN=VDAC2 PE=1 SV=2 | P45880 | VDAC2_HUMAN | 32 kDa |
| Voltage-dependent calcium channel subunit alpha-2/delta-1 OS=Homo sapiens OX=9606 GN=CACNA2D1 PE=1 SV=3 | P54289 | CA2D1_HUMAN | 125 kDa |
| Volume-regulated anion channel subunit LRRC8A OS=Homo sapiens OX=9606 GN=LRRC8A PE=1 SV=1 | Q8IWT6 | LRC8A_HUMAN | 94 kDa |
| V-type proton ATPase 116 kDa subunit a 1 OS=Homo sapiens OX=9606 GN=ATP6V0A1 PE=1 SV=3 | Q93050 | VPP1_HUMAN | 96 kDa |
| V-type proton ATPase 16 kDa proteolipid subunit c OS=Homo sapiens OX=9606 GN=ATP6V0C PE=1 SV=1 | P27449 | VATL_HUMAN | 16 kDa |
| V-type proton ATPase catalytic subunit A OS=Homo sapiens OX=9606 GN=ATP6V1A PE=1 SV=2 | P38606 | VATA_HUMAN | 68 kDa |
| V-type proton ATPase subunit B, brain isoform OS=Homo sapiens OX=9606 GN=ATP6V1B2 PE=1 SV=3 | P21281 | VATB2_HUMAN | 57 kDa |
| V-type proton ATPase subunit E 1 OS=Homo sapiens OX=9606 GN=ATP6V1E1 PE=1 SV=1 | P36543 | VATE1_HUMAN | 26 kDa |
| V-type proton ATPase subunit G 1 OS=Homo sapiens OX=9606 GN=ATP6V1G1 PE=1 SV=3 | O75348 | VATG1_HUMAN (+1) | 14 kDa |
| V-type proton ATPase subunit H OS=Homo sapiens OX=9606 GN=ATP6V1H PE=1 SV=1 | Q9UI12 | VATH_HUMAN | 56 kDa |
| WD repeat-containing protein 1 OS=Homo sapiens OX=9606 GN=WDR1 PE=1 SV=4 | O75083 | WDR1_HUMAN | 66 kDa |
| WW domain-binding protein 2 OS=Homo sapiens OX=9606 GN=WBP2 PE=1 SV=1 | Q969T9 | WBP2_HUMAN | 28 kDa |
| Xaa-Pro aminopeptidase 1 OS=Homo sapiens OX=9606 GN=XPNPEP1 PE=1 SV=3 | Q9NQW7 | XPP1_HUMAN | 70 kDa |
| Zinc finger CCCH-type antiviral protein 1 OS=Homo sapiens OX=9606 GN=ZC3HAV1 PE=1 SV=3 | Q7Z2W4 | ZCCHV_HUMAN | 101 kDa |
| Zinc transporter ZIP10 OS=Homo sapiens OX=9606 GN=SLC39A10 PE=1 SV=2 | Q9ULF5 | S39AA_HUMAN | 94 kDa |
| Zinc-alpha-2-glycoprotein OS=Homo sapiens OX=9606 GN=AZGP1 PE=1 SV=2 | P25311 | ZA2G_HUMAN | 34 kDa |
| Zymogen granule protein 16 homolog B OS=Homo sapiens OX=9606 GN=ZG16B PE=1 SV=3 | Q96DA0 | ZG16B_HUMAN | 23 kDa |

| **Supplementary Table 2: Identified proteins (1340) in ffEVs samples coming from individual follicles containing MII, GVBD and GV oocytes (n=21)** | | | | | |
| --- | --- | --- | --- | --- | --- |
| **Name** | **protein_Id** | **UniProt_ID** | **UniProt_name** | **Gene_ID** | **description** |
| IGLV8-61 | sp\|A0A075B6I0\|LV861_HUMAN | A0A075B6I0 | LV861 | 28774 | Immunoglobulin lambda variable 8-61 |
| IGKV2D-24 | sp\|A0A075B6R9\|KVD24_HUMAN | A0A075B6R9 | KVD24 | 28885 | Probable non-functional immunoglobulin kappa variable 2D-24 |
| IGKV3D-11 | sp\|A0A0A0MRZ8\|KVD11_HUMAN | A0A0A0MRZ8 | KVD11 | 28876 | Immunoglobulin kappa variable 3D-11 |
| IGHV3-49 | sp\|A0A0A0MS15\|HV349_HUMAN | A0A0A0MS15 | HV349 | 28423 | Immunoglobulin heavy variable 3-49 |
| IGHV3-15 | sp\|A0A0B4J1V0\|HV315_HUMAN | A0A0B4J1V0 | HV315 | 28448 | Immunoglobulin heavy variable 3-15 |
| IGHV3-72 | sp\|A0A0B4J1Y9\|HV372_HUMAN | A0A0B4J1Y9 | HV372 | 28410 | Immunoglobulin heavy variable 3-72 |
| IGKV1D-13 | sp\|A0A0B4J2D9\|KVD13_HUMAN | A0A0B4J2D9 | KVD13 | 28902 | Immunoglobulin kappa variable 1D-13 |
| IGHV4-28 | sp\|A0A0C4DH34\|HV428_HUMAN | A0A0C4DH34 | HV428 | 28400 | Immunoglobulin heavy variable 4-28 |
| IGHV5-51 | sp\|A0A0C4DH38\|HV551_HUMAN | A0A0C4DH38 | HV551 | 28388 | Immunoglobulin heavy variable 5-51 |
| ESYT2 | sp\|A0FGR8\|ESYT2_HUMAN | A0FGR8 | ESYT2 | 57488 | Extended synaptotagmin-2 |
| A2ML1 | sp\|A8K2U0\|A2ML1_HUMAN | A8K2U0 | A2ML1 | 144568 | Alpha-2-macroglobulin-like protein 1 |
| MYO1C | sp\|O00159\|MYO1C_HUMAN | O00159 | MYO1C | 4641 | Unconventional myosin-Ic |
| SNAP23 | sp\|O00161\|SNP23_HUMAN | O00161 | SNP23 | 8773 | Synaptosomal-associated protein 23 |
| STXBP3 | sp\|O00186\|STXB3_HUMAN | O00186 | STXB3 | 6814 | Syntaxin-binding protein 3 |
| MASP2 | sp\|O00187\|MASP2_HUMAN | O00187 | MASP2 | 10747 | Mannan-binding lectin serine protease 2 |
| PSMD11 | sp\|O00231\|PSD11_HUMAN | O00231 | PSD11 | 5717 | 26S proteasome non-ATPase regulatory subunit 11 |
| PGRMC1 | sp\|O00264\|PGRC1_HUMAN | O00264 | PGRC1 | 10857 | Membrane-associated progesterone receptor component 1 |
| CLIC1 | sp\|O00299\|CLIC1_HUMAN | O00299 | CLIC1 | 1192 | Chloride intracellular channel protein 1 |
| MATN2 | sp\|O00339\|MATN2_HUMAN | O00339 | MATN2 | 4147 | Matrilin-2 |
| QSOX1 | sp\|O00391\|QSOX1_HUMAN | O00391 | QSOX1 | 5768 | Sulfhydryl oxidase 1 |
| PSMD14 | sp\|O00487\|PSDE_HUMAN | O00487 | PSDE | 10213 | 26S proteasome non-ATPase regulatory subunit 14 |
| SDCBP | sp\|O00560\|SDCB1_HUMAN | O00560 | SDCB1 | 6386 | Syntenin-1 |
| SCD | sp\|O00767\|SCD_HUMAN | O00767 | SCD | 6319 | Stearoyl-CoA desaturase |
| COPE | sp\|O14579\|COPE_HUMAN | O14579 | COPE | 11316 | Coatomer subunit epsilon |
| ADAM10 | sp\|O14672\|ADA10_HUMAN | O14672 | ADA10 | 102 | Disintegrin and metalloproteinase domain-containing protein 10 |
| CDIPT | sp\|O14735\|CDIPT_HUMAN | O14735 | CDIPT | 10423 | CDP-diacylglycerol--inositol 3-phosphatidyltransferase |
| SLC9A3R1 | sp\|O14745\|NHRF1_HUMAN | O14745 | NHRF1 | 9368 | Na(+)/H(+) exchange regulatory cofactor NHE-RF1 |
| APOL1 | sp\|O14791\|APOL1_HUMAN | O14791 | APOL1 | 8542 | Apolipoprotein L1 |
| HS3ST1 | sp\|O14792\|HS3S1_HUMAN | O14792 | HS3S1 | 9957 | Heparan sulfate glucosamine 3-O-sulfotransferase 1 |
| SCAMP3 | sp\|O14828\|SCAM3_HUMAN | O14828 | SCAM3 | 10067 | Secretory carrier-associated membrane protein 3 |
| CASK | sp\|O14936\|CSKP_HUMAN | O14936 | CSKP | 8573 | Peripheral plasma membrane protein CASK |
| MYL12B | sp\|O14950\|ML12B_HUMAN | O14950 | ML12B | 103910; 10627 | Myosin regulatory light chain 12B |
| PLXNB2 | sp\|O15031\|PLXB2_HUMAN | O15031 | PLXB2 | 23654 | Plexin-B2 |
| NPC1 | sp\|O15118\|NPC1_HUMAN | O15118 | NPC1 | 4864 | NPC intracellular cholesterol transporter 1 |
| ANGPT2 | sp\|O15123\|ANGP2_HUMAN | O15123 | ANGP2 | 285 | Angiopoietin-2 |
| SCAMP1 | sp\|O15126\|SCAM1_HUMAN | O15126 | SCAM1 | 9522 | Secretory carrier-associated membrane protein 1 |
| SCAMP2 | sp\|O15127\|SCAM2_HUMAN | O15127 | SCAM2 | 10066 | Secretory carrier-associated membrane protein 2 |
| ARPC2 | sp\|O15144\|ARPC2_HUMAN | O15144 | ARPC2 | 10109 | Actin-related protein 2/3 complex subunit 2 |
| ARPC3 | sp\|O15145\|ARPC3_HUMAN | O15145 | ARPC3 | 10094 | Actin-related protein 2/3 complex subunit 3 |
| PGRMC2 | sp\|O15173\|PGRC2_HUMAN | O15173 | PGRC2 | 10424 | Membrane-associated progesterone receptor component 2 |
| VGF | sp\|O15240\|VGF_HUMAN | O15240 | VGF | 7425 | Neurosecretory protein VGF |
| RER1 | sp\|O15258\|RER1_HUMAN | O15258 | RER1 | 11079 | Protein RER1 |
| SURF4 | sp\|O15260\|SURF4_HUMAN | O15260 | SURF4 | 6836 | Surfeit locus protein 4 |
| SPTLC2 | sp\|O15270\|SPTC2_HUMAN | O15270 | SPTC2 | 9517 | Serine palmitoyltransferase 2 |
| SLC16A3 | sp\|O15427\|MOT4_HUMAN | O15427 | MOT4 | 9123 | Monocarboxylate transporter 4 |
| ABCC4 | sp\|O15439\|MRP4_HUMAN | O15439 | MRP4 | 10257 | ATP-binding cassette sub-family C member 4 |
| TAPBP | sp\|O15533\|TPSN_HUMAN | O15533 | TPSN | 6892 | Tapasin |
| CLDN3 | sp\|O15551\|CLD3_HUMAN | O15551 | CLD3 | 1365 | Claudin-3 |
| DYNC1LI2 | sp\|O43237\|DC1L2_HUMAN | O43237 | DC1L2 | 1783 | Cytoplasmic dynein 1 light intermediate chain 2 |
| KLK10 | sp\|O43240\|KLK10_HUMAN | O43240 | KLK10 | 5655 | Kallikrein-10 |
| PSMD3 | sp\|O43242\|PSMD3_HUMAN | O43242 | PSMD3 | 5709 | 26S proteasome non-ATPase regulatory subunit 3 |
| TXNL1 | sp\|O43396\|TXNL1_HUMAN | O43396 | TXNL1 | 9352 | Thioredoxin-like protein 1 |
| TSPAN6 | sp\|O43657\|TSN6_HUMAN | O43657 | TSN6 | 7105 | Tetraspanin-6 |
| CALU | sp\|O43852\|CALU_HUMAN | O43852 | CALU | 813 | Calumenin |
| EDIL3 | sp\|O43854\|EDIL3_HUMAN | O43854 | EDIL3 | 10085 | EGF-like repeat and discoidin I-like domain-containing protein 3 |
| AHCYL1 | sp\|O43865\|SAHH2_HUMAN | O43865 | SAHH2 | 10768 | S-adenosylhomocysteine hydrolase-like protein 1 |
| MGRN1 | sp\|O60291\|MGRN1_HUMAN | O60291 | MGRN1 | 23295 | E3 ubiquitin-protein ligase MGRN1 |
| PPL | sp\|O60437\|PEPL_HUMAN | O60437 | PEPL | 5493 | Periplakin |
| ACSL4 | sp\|O60488\|ACSL4_HUMAN | O60488 | ACSL4 | 2182 | Long-chain-fatty-acid--CoA ligase 4 |
| PLOD3 | sp\|O60568\|PLOD3_HUMAN | O60568 | PLOD3 | 8985 | Multifunctional procollagen lysine hydroxylase and glycosyltransferase LH3 |
| PLIN3 | sp\|O60664\|PLIN3_HUMAN | O60664 | PLIN3 | 10226 | Perilipin-3 |
| UGDH | sp\|O60701\|UGDH_HUMAN | O60701 | UGDH | 7358 | UDP-glucose 6-dehydrogenase |
| CTNND1 | sp\|O60716\|CTND1_HUMAN | O60716 | CTND1 | 1500 | Catenin delta-1 |
| DPM1 | sp\|O60762\|DPM1_HUMAN | O60762 | DPM1 | 8813 | Dolichol-phosphate mannosyltransferase subunit 1 |
| WDR1 | sp\|O75083\|WDR1_HUMAN | O75083 | WDR1 | 9948 | WD repeat-containing protein 1 |
| GGCT | sp\|O75223\|GGCT_HUMAN | O75223 | GGCT | 79017 | Gamma-glutamylcyclotransferase |
| ALOX12B | sp\|O75342\|LX12B_HUMAN | O75342 | LX12B | 242 | Arachidonate 12-lipoxygenase, 12R-type |
| MPDU1 | sp\|O75352\|MPU1_HUMAN | O75352 | MPU1 | 9526 | Mannose-P-dolichol utilization defect 1 protein |
| MACROH2A1 | sp\|O75367\|H2AY_HUMAN | O75367 | H2AY | 9555 | Core histone macro-H2A.1 |
| SEC22B | sp\|O75396\|SC22B_HUMAN | O75396 | SC22B | 9554 | Vesicle-trafficking protein SEC22b |
| ERN1 | sp\|O75460\|ERN1_HUMAN | O75460 | ERN1 | 2081 | Serine/threonine-protein kinase/endoribonuclease IRE1 |
| BANF1 | sp\|O75531\|BAF_HUMAN | O75531 | BAF | 8815 | Barrier-to-autointegration factor |
| FCN3 | sp\|O75636\|FCN3_HUMAN | O75636 | FCN3 | 8547 | Ficolin-3 |
| RP2 | sp\|O75695\|XRP2_HUMAN | O75695 | XRP2 | 6102 | Protein XRP2 |
| ATP6AP2 | sp\|O75787\|RENR_HUMAN | O75787 | RENR | 10159 | Renin receptor |
| ZMPSTE24 | sp\|O75844\|FACE1_HUMAN | O75844 | FACE1 | 10269 | CAAX prenyl protease 1 homolog |
| IDH1 | sp\|O75874\|IDHC_HUMAN | O75874 | IDHC | 3417 | Isocitrate dehydrogenase [NADP] cytoplasmic |
| ATRN | sp\|O75882\|ATRN_HUMAN | O75882 | ATRN | 8455 | Attractin |
| ARL6IP5 | sp\|O75915\|PRAF3_HUMAN | O75915 | PRAF3 | 10550 | PRA1 family protein 3 |
| FLOT1 | sp\|O75955\|FLOT1_HUMAN | O75955 | FLOT1 | 10211 | Flotillin-1 |
| ERLIN2 | sp\|O94905\|ERLN2_HUMAN | O94905 | ERLN2 | 11160 | Erlin-2 |
| ENDOD1 | sp\|O94919\|ENDD1_HUMAN | O94919 | ENDD1 | 23052 | Endonuclease domain-containing 1 protein |
| PRSS23 | sp\|O95084\|PRS23_HUMAN | O95084 | PRS23 | 11098 | Serine protease 23 |
| RTN3 | sp\|O95197\|RTN3_HUMAN | O95197 | RTN3 | 10313 | Reticulon-3 |
| VAPB | sp\|O95292\|VAPB_HUMAN | O95292 | VAPB | 9217 | Vesicle-associated membrane protein-associated protein B/C |
| MPZL1 | sp\|O95297\|MPZL1_HUMAN | O95297 | MPZL1 | 9019 | Myelin protein zero-like protein 1 |
| FKBP9 | sp\|O95302\|FKBP9_HUMAN | O95302 | FKBP9 | 11328 | Peptidyl-prolyl cis-trans isomerase FKBP9 |
| APOM | sp\|O95445\|APOM_HUMAN | O95445 | APOM | 55937 | Apolipoprotein M |
| H6PD | sp\|O95479\|G6PE_HUMAN | O95479 | G6PE | 9563 | GDH/6PGL endoplasmic bifunctional protein |
| CLIC3 | sp\|O95833\|CLIC3_HUMAN | O95833 | CLIC3 | 9022 | Chloride intracellular channel protein 3 |
| DDAH2 | sp\|O95865\|DDAH2_HUMAN | O95865 | DDAH2 | 23564 | N(G),N(G)-dimethylarginine dimethylaminohydrolase 2 |
| TXNDC12 | sp\|O95881\|TXD12_HUMAN | O95881 | TXD12 | 51060 | Thioredoxin domain-containing protein 12 |
| CLPTM1 | sp\|O96005\|CLPT1_HUMAN | O96005 | CLPT1 | 1209 | Cleft lip and palate transmembrane protein 1 |
| CYB5A | sp\|P00167\|CYB5_HUMAN | P00167 | CYB5 | 1528 | Cytochrome b5 |
| CYB5R3 | sp\|P00387\|NB5R3_HUMAN | P00387 | NB5R3 | 1727 | NADH-cytochrome b5 reductase 3 |
| GSR | sp\|P00390\|GSHR_HUMAN | P00390 | GSHR | 2936 | Glutathione reductase, mitochondrial |
| MT-CO2 | sp\|P00403\|COX2_HUMAN | P00403 | COX2 | 4513 | Cytochrome c oxidase subunit 2 |
| F13A1 | sp\|P00488\|F13A_HUMAN | P00488 | F13A | 2162 | Coagulation factor XIII A chain |
| C1R | sp\|P00736\|C1R_HUMAN | P00736 | C1R | 715 | Complement C1r subcomponent |
| F9 | sp\|P00740\|FA9_HUMAN | P00740 | FA9 | 2158 | Coagulation factor IX |
| F10 | sp\|P00742\|FA10_HUMAN | P00742 | FA10 | 2159 | Coagulation factor X |
| CFD | sp\|P00746\|CFAD_HUMAN | P00746 | CFAD | 1675 | Complement factor D |
| F12 | sp\|P00748\|FA12_HUMAN | P00748 | FA12 | 2161 | Coagulation factor XII |
| SERPINA1 | sp\|P01009\|A1AT_HUMAN | P01009 | A1AT | 5265 | Alpha-1-antitrypsin |
| SERPINA3 | sp\|P01011\|AACT_HUMAN | P01011 | AACT | 12 | Alpha-1-antichymotrypsin |
| AGT | sp\|P01019\|ANGT_HUMAN | P01019 | ANGT | 183 | Angiotensinogen |
| TIMP1 | sp\|P01033\|TIMP1_HUMAN | P01033 | TIMP1 | 7076 | Metalloproteinase inhibitor 1 |
| CST3 | sp\|P01034\|CYTC_HUMAN | P01034 | CYTC | 1471 | Cystatin-C |
| CST4 | sp\|P01036\|CYTS_HUMAN | P01036 | CYTS | 1472 | Cystatin-S |
| CST1 | sp\|P01037\|CYTN_HUMAN | P01037 | CYTN | 1469 | Cystatin-SN |
| CSTA | sp\|P01040\|CYTA_HUMAN | P01040 | CYTA | 1475 | Cystatin-A |
| LDLR | sp\|P01130\|LDLR_HUMAN | P01130 | LDLR | 3949 | Low-density lipoprotein receptor |
| IGF2 | sp\|P01344\|IGF2_HUMAN | P01344 | IGF2 | 3481 | Insulin-like growth factor II |
| JCHAIN | sp\|P01591\|IGJ_HUMAN | P01591 | IGJ | 3512 | Immunoglobulin J chain |
| IGKV1D-33 | sp\|P01593\|KVD33_HUMAN | P01593 | KVD33 | 28896 | Immunoglobulin kappa variable 1D-33 |
| IGKV1-5 | sp\|P01602\|KV105_HUMAN | P01602 | KV105 | 28299 | Immunoglobulin kappa variable 1-5 |
| IGKV3-20 | sp\|P01619\|KV320_HUMAN | P01619 | KV320 | 28912 | Immunoglobulin kappa variable 3-20 |
| IGLV1-51 | sp\|P01701\|LV151_HUMAN | P01701 | LV151 | 28820 | Immunoglobulin lambda variable 1-51 |
| IGLV2-14 | sp\|P01704\|LV214_HUMAN | P01704 | LV214 | 28815 | Immunoglobulin lambda variable 2-14 |
| IGLV3-19 | sp\|P01714\|LV319_HUMAN | P01714 | LV319 | 28797 | Immunoglobulin lambda variable 3-19 |
| PIGR | sp\|P01833\|PIGR_HUMAN | P01833 | PIGR | 5284 | Polymeric immunoglobulin receptor |
| IGKC | sp\|P01834\|IGKC_HUMAN | P01834 | IGKC | 3514 | Immunoglobulin kappa constant |
| IGHG1 | sp\|P01857\|IGHG1_HUMAN | P01857 | IGHG1 | 3500 | Immunoglobulin heavy constant gamma 1 |
| IGHG2 | sp\|P01859\|IGHG2_HUMAN | P01859 | IGHG2 | 3501 | Immunoglobulin heavy constant gamma 2 |
| IGHG3 | sp\|P01860\|IGHG3_HUMAN | P01860 | IGHG3 | 3502 | Immunoglobulin heavy constant gamma 3 |
| IGHG4 | sp\|P01861\|IGHG4_HUMAN | P01861 | IGHG4 | 3503 | Immunoglobulin heavy constant gamma 4 |
| IGHA1 | sp\|P01876\|IGHA1_HUMAN | P01876 | IGHA1 | 3493 | Immunoglobulin heavy constant alpha 1 |
| IGHA2 | sp\|P01877\|IGHA2_HUMAN | P01877 | IGHA2 | 3494 | Immunoglobulin heavy constant alpha 2 |
| IGHD | sp\|P01880\|IGHD_HUMAN | P01880 | IGHD | 3495 | Immunoglobulin heavy constant delta |
| HLA-B | sp\|P01889\|HLAB_HUMAN | P01889 | HLAB | 3106 | HLA class I histocompatibility antigen, B alpha chain |
| APOA2 | sp\|P02652\|APOA2_HUMAN | P02652 | APOA2 | 336 | Apolipoprotein A-II |
| APOC1 | sp\|P02654\|APOC1_HUMAN | P02654 | APOC1 | 341 | Apolipoprotein C-I |
| APOC3 | sp\|P02656\|APOC3_HUMAN | P02656 | APOC3 | 345 | Apolipoprotein C-III |
| FGB | sp\|P02675\|FIBB_HUMAN | P02675 | FIBB | 2244 | Fibrinogen beta chain |
| FGG | sp\|P02679\|FIBG_HUMAN | P02679 | FIBG | 2266 | Fibrinogen gamma chain |
| SLC4A1 | sp\|P02730\|B3AT_HUMAN | P02730 | B3AT | 6521 | Band 3 anion transport protein |
| APCS | sp\|P02743\|SAMP_HUMAN | P02743 | SAMP | 325 | Serum amyloid P-component |
| C1QA | sp\|P02745\|C1QA_HUMAN | P02745 | C1QA | 712 | Complement C1q subcomponent subunit A |
| C1QB | sp\|P02746\|C1QB_HUMAN | P02746 | C1QB | 713 | Complement C1q subcomponent subunit B |
| C1QC | sp\|P02747\|C1QC_HUMAN | P02747 | C1QC | 714 | Complement C1q subcomponent subunit C |
| LRG1 | sp\|P02750\|A2GL_HUMAN | P02750 | A2GL | 116844 | Leucine-rich alpha-2-glycoprotein |
| RBP4 | sp\|P02753\|RET4_HUMAN | P02753 | RET4 | 5950 | Retinol-binding protein 4 |
| AMBP | sp\|P02760\|AMBP_HUMAN | P02760 | AMBP | 259 | Protein AMBP |
| ORM1 | sp\|P02763\|A1AG1_HUMAN | P02763 | A1AG1 | 5004 | Alpha-1-acid glycoprotein 1 |
| TTR | sp\|P02766\|TTHY_HUMAN | P02766 | TTHY | 7276 | Transthyretin |
| TFRC | sp\|P02786\|TFR1_HUMAN | P02786 | TFR1 | 7037 | Transferrin receptor protein 1 |
| FTL | sp\|P02792\|FRIL_HUMAN | P02792 | FRIL | 2512 | Ferritin light chain |
| FTH1 | sp\|P02794\|FRIH_HUMAN | P02794 | FRIH | 2495 | Ferritin heavy chain |
| SMR3B | sp\|P02814\|SMR3B_HUMAN | P02814 | SMR3B | 10879 | Submaxillary gland androgen-regulated protein 3B |
| ANG | sp\|P03950\|ANGI_HUMAN | P03950 | ANGI | 283 | Angiogenin |
| F11 | sp\|P03951\|FA11_HUMAN | P03951 | FA11 | 2160 | Coagulation factor XI |
| KLKB1 | sp\|P03952\|KLKB1_HUMAN | P03952 | KLKB1 | 3818 | Plasma kallikrein |
| HMGCR | sp\|P04035\|HMDH_HUMAN | P04035 | HMDH | 3156 | 3-hydroxy-3-methylglutaryl-coenzyme A reductase |
| CAT | sp\|P04040\|CATA_HUMAN | P04040 | CATA | 847 | Catalase |
| PROC | sp\|P04070\|PROC_HUMAN | P04070 | PROC | 5624 | Vitamin K-dependent protein C |
| CSTB | sp\|P04080\|CYTB_HUMAN | P04080 | CYTB | 1476 | Cystatin-B |
| ANXA1 | sp\|P04083\|ANXA1_HUMAN | P04083 | ANXA1 | 301 | Annexin A1 |
| APOB | sp\|P04114\|APOB_HUMAN | P04114 | APOB | 338 | Apolipoprotein B-100 |
| HRG | sp\|P04196\|HRG_HUMAN | P04196 | HRG | 3273 | Histidine-rich glycoprotein |
| A1BG | sp\|P04217\|A1BG_HUMAN | P04217 | A1BG | 1 | Alpha-1B-glycoprotein |
| VWF | sp\|P04275\|VWF_HUMAN | P04275 | VWF | 7450 | von Willebrand factor |
| CAPNS1 | sp\|P04632\|CPNS1_HUMAN | P04632 | CPNS1 | 826 | Calpain small subunit 1 |
| HSPB1 | sp\|P04792\|HSPB1_HUMAN | P04792 | HSPB1 | 3315 | Heat shock protein beta-1 |
| RPN1 | sp\|P04843\|RPN1_HUMAN | P04843 | RPN1 | 6184 | Dolichyl-diphosphooligosaccharide--protein glycosyltransferase subunit 1 |
| RPN2 | sp\|P04844\|RPN2_HUMAN | P04844 | RPN2 | 6185 | Dolichyl-diphosphooligosaccharide--protein glycosyltransferase subunit 2 |
| ATP1B1 | sp\|P05026\|AT1B1_HUMAN | P05026 | AT1B1 | 481 | Sodium/potassium-transporting ATPase subunit beta-1 |
| CHGB | sp\|P05060\|SCG1_HUMAN | P05060 | SCG1 | 1114 | Secretogranin-1 |
| ARG1 | sp\|P05089\|ARGI1_HUMAN | P05089 | ARGI1 | 383 | Arginase-1 |
| CYP17A1 | sp\|P05093\|CP17A_HUMAN | P05093 | CP17A | 1586 | Steroid 17-alpha-hydroxylase/17,20 lyase |
| CYP11A1 | sp\|P05108\|CP11A_HUMAN | P05108 | CP11A | 1583 | Cholesterol side-chain cleavage enzyme, mitochondrial |
| S100A8 | sp\|P05109\|S10A8_HUMAN | P05109 | S10A8 | 6279 | Protein S100-A8 |
| INHA | sp\|P05111\|INHA_HUMAN | P05111 | INHA | 3623 | Inhibin alpha chain |
| SERPINA5 | sp\|P05154\|IPSP_HUMAN | P05154 | IPSP | 5104 | Plasma serine protease inhibitor |
| SERPING1 | sp\|P05155\|IC1_HUMAN | P05155 | IC1 | 710 | Plasma protease C1 inhibitor |
| F13B | sp\|P05160\|F13B_HUMAN | P05160 | F13B | 2165 | Coagulation factor XIII B chain |
| RPLP2 | sp\|P05387\|RLA2_HUMAN | P05387 | RLA2 | 6181 | 60S acidic ribosomal protein P2 |
| CLEC3B | sp\|P05452\|TETN_HUMAN | P05452 | TETN | 7123 | Tetranectin |
| COL5A2 | sp\|P05997\|CO5A2_HUMAN | P05997 | CO5A2 | 1290 | Collagen alpha-2(V) chain |
| BCHE | sp\|P06276\|CHLE_HUMAN | P06276 | CHLE | 590 | Cholinesterase |
| IGKV4-1 | sp\|P06312\|KV401_HUMAN | P06312 | KV401 | 28908 | Immunoglobulin kappa variable 4-1 |
| ATP5F1B | sp\|P06576\|ATPB_HUMAN | P06576 | ATPB | 506 | ATP synthase subunit beta, mitochondrial |
| C2 | sp\|P06681\|CO2_HUMAN | P06681 | CO2 | 717 | Complement C2 |
| S100A9 | sp\|P06702\|S10A9_HUMAN | P06702 | S10A9 | 6280 | Protein S100-A9 |
| APOA4 | sp\|P06727\|APOA4_HUMAN | P06727 | APOA4 | 337 | Apolipoprotein A-IV |
| ITGAV | sp\|P06756\|ITAV_HUMAN | P06756 | ITAV | 3685 | Integrin alpha-V |
| SERPINE2 | sp\|P07093\|GDN_HUMAN | P07093 | GDN | 5270 | Glia-derived nexin |
| EPHX1 | sp\|P07099\|HYEP_HUMAN | P07099 | HYEP | 2052 | Epoxide hydrolase 1 |
| P4HB | sp\|P07237\|PDIA1_HUMAN | P07237 | PDIA1 | 5034 | Protein disulfide-isomerase |
| CTSD | sp\|P07339\|CATD_HUMAN | P07339 | CATD | 1509 | Cathepsin D |
| C8A | sp\|P07357\|CO8A_HUMAN | P07357 | CO8A | 731 | Complement component C8 alpha chain |
| C8B | sp\|P07358\|CO8B_HUMAN | P07358 | CO8B | 732 | Complement component C8 beta chain |
| C8G | sp\|P07360\|CO8G_HUMAN | P07360 | CO8G | 733 | Complement component C8 gamma chain |
| CAPN1 | sp\|P07384\|CAN1_HUMAN | P07384 | CAN1 | 823 | Calpain-1 catalytic subunit |
| IVL | sp\|P07476\|INVO_HUMAN | P07476 | INVO | 3713 | Involucrin |
| PSAP | sp\|P07602\|SAP_HUMAN | P07602 | SAP | 5660 | Prosaposin |
| APRT | sp\|P07741\|APT_HUMAN | P07741 | APT | 353 | Adenine phosphoribosyltransferase |
| EPRS1 | sp\|P07814\|SYEP_HUMAN | P07814 | SYEP | 2058 | Bifunctional glutamate/proline--tRNA ligase |
| CTSB | sp\|P07858\|CATB_HUMAN | P07858 | CATB | 1508 | Cathepsin B |
| COL1A2 | sp\|P08123\|CO1A2_HUMAN | P08123 | CO1A2 | 1278 | Collagen alpha-2(I) chain |
| CD55 | sp\|P08174\|DAF_HUMAN | P08174 | DAF | 1604 | Complement decay-accelerating factor |
| SERPINA6 | sp\|P08185\|CBG_HUMAN | P08185 | CBG | 866 | Corticosteroid-binding globulin |
| SLC3A2 | sp\|P08195\|4F2_HUMAN | P08195 | 4F2 | 6520 | 4F2 cell-surface antigen heavy chain |
| SOD3 | sp\|P08294\|SODE_HUMAN | P08294 | SODE | 6649 | Extracellular superoxide dismutase [Cu-Zn] |
| INHBA | sp\|P08476\|INHBA_HUMAN | P08476 | INHBA | 3624 | Inhibin beta A chain |
| MGP | sp\|P08493\|MGP_HUMAN | P08493 | MGP | 4256 | Matrix Gla protein |
| ITGA2B | sp\|P08514\|ITA2B_HUMAN | P08514 | ITA2B | 3674 | Integrin alpha-IIb |
| CD14 | sp\|P08571\|CD14_HUMAN | P08571 | CD14 | 929 | Monocyte differentiation antigen CD14 |
| COL4A2 | sp\|P08572\|CO4A2_HUMAN | P08572 | CO4A2 | 1284 | Collagen alpha-2(IV) chain |
| ITGA5 | sp\|P08648\|ITA5_HUMAN | P08648 | ITA5 | 3678 | Integrin alpha-5 |
| ANXA5 | sp\|P08758\|ANXA5_HUMAN | P08758 | ANXA5 | 308 | Annexin A5 |
| RPSA | sp\|P08865\|RSSA_HUMAN | P08865 | RSSA | 3921 | 40S ribosomal protein SA |
| CD63 | sp\|P08962\|CD63_HUMAN | P08962 | CD63 | 967 | CD63 antigen |
| DBH | sp\|P09172\|DOPO_HUMAN | P09172 | DOPO | 1621 | Dopamine beta-hydroxylase |
| LGALS1 | sp\|P09382\|LEG1_HUMAN | P09382 | LEG1 | 3956 | Galectin-1 |
| RBP1 | sp\|P09455\|RET1_HUMAN | P09455 | RET1 | 5947 | Retinol-binding protein 1 |
| SPARC | sp\|P09486\|SPRC_HUMAN | P09486 | SPRC | 6678 | SPARC |
| CNP | sp\|P09543\|CN37_HUMAN | P09543 | CN37 | 1267 | 2',3'-cyclic-nucleotide 3'-phosphodiesterase |
| HMOX1 | sp\|P09601\|HMOX1_HUMAN | P09601 | HMOX1 | 3162 | Heme oxygenase 1 |
| UCHL1 | sp\|P09936\|UCHL1_HUMAN | P09936 | UCHL1 | 7345 | Ubiquitin carboxyl-terminal hydrolase isozyme L1 |
| UBB | sp\|P0CG47\|UBB_HUMAN | P0CG47 | UBB | 7314 | Polyubiquitin-B |
| SAA1 | sp\|P0DJI8\|SAA1_HUMAN | P0DJI8 | SAA1 | 6288 | Serum amyloid A-1 protein |
| SAA2 | sp\|P0DJI9\|SAA2_HUMAN | P0DJI9 | SAA2 | 6289 | Serum amyloid A-2 protein |
| CALM1 | sp\|P0DP23\|CALM1_HUMAN | P0DP23 | CALM1 | 801; 805; 808 | Calmodulin-1 |
| TXN | sp\|P10599\|THIO_HUMAN | P10599 | THIO | 7295 | Thioredoxin |
| MGST1 | sp\|P10620\|MGST1_HUMAN | P10620 | MGST1 | 4257 | Microsomal glutathione S-transferase 1 |
| PRKAR1A | sp\|P10644\|KAP0_HUMAN | P10644 | KAP0 | 5573 | cAMP-dependent protein kinase type I-alpha regulatory subunit |
| TFPI | sp\|P10646\|TFPI1_HUMAN | P10646 | TFPI1 | 7035 | Tissue factor pathway inhibitor |
| HSPD1 | sp\|P10809\|CH60_HUMAN | P10809 | CH60 | 3329 | 60 kDa heat shock protein, mitochondrial |
| LAMC1 | sp\|P11047\|LAMC1_HUMAN | P11047 | LAMC1 | 3915 | Laminin subunit gamma-1 |
| SLC2A1 | sp\|P11166\|GTR1_HUMAN | P11166 | GTR1 | 6513 | Solute carrier family 2, facilitated glucose transporter member 1 |
| MBL2 | sp\|P11226\|MBL2_HUMAN | P11226 | MBL2 | 4153 | Mannose-binding protein C |
| RALA | sp\|P11233\|RALA_HUMAN | P11233 | RALA | 5898 | Ras-related protein Ral-A |
| RALB | sp\|P11234\|RALB_HUMAN | P11234 | RALB | 5899 | Ras-related protein Ral-B |
| CYP19A1 | sp\|P11511\|CP19A_HUMAN | P11511 | CP19A | 1588 | Aromatase |
| MTHFD1 | sp\|P11586\|C1TC_HUMAN | P11586 | C1TC | 4522 | C-1-tetrahydrofolate synthase, cytoplasmic |
| CETP | sp\|P11597\|CETP_HUMAN | P11597 | CETP | 1071 | Cholesteryl ester transfer protein |
| COL6A1 | sp\|P12109\|CO6A1_HUMAN | P12109 | CO6A1 | 1291 | Collagen alpha-1(VI) chain |
| COL6A2 | sp\|P12110\|CO6A2_HUMAN | P12110 | CO6A2 | 1292 | Collagen alpha-2(VI) chain |
| COL6A3 | sp\|P12111\|CO6A3_HUMAN | P12111 | CO6A3 | 1293 | Collagen alpha-3(VI) chain |
| IMPDH2 | sp\|P12268\|IMDH2_HUMAN | P12268 | IMDH2 | 3615 | Inosine-5'-monophosphate dehydrogenase 2 |
| PIP | sp\|P12273\|PIP_HUMAN | P12273 | PIP | 5304 | Prolactin-inducible protein |
| CKB | sp\|P12277\|KCRB_HUMAN | P12277 | KCRB | 1152 | Creatine kinase B-type |
| BMP3 | sp\|P12645\|BMP3_HUMAN | P12645 | BMP3 | 651 | Bone morphogenetic protein 3 |
| ACE | sp\|P12821\|ACE_HUMAN | P12821 | ACE | 1636 | Angiotensin-converting enzyme |
| CDH1 | sp\|P12830\|CADH1_HUMAN | P12830 | CADH1 | 999 | Cadherin-1 |
| COX4I1 | sp\|P13073\|COX41_HUMAN | P13073 | COX41 | 1327 | Cytochrome c oxidase subunit 4 isoform 1, mitochondrial |
| RNH1 | sp\|P13489\|RINI_HUMAN | P13489 | RINI | 6050 | Ribonuclease inhibitor |
| VCAN | sp\|P13611\|CSPG2_HUMAN | P13611 | CSPG2 | 1462 | Versican core protein |
| PDIA4 | sp\|P13667\|PDIA4_HUMAN | P13667 | PDIA4 | 9601 | Protein disulfide-isomerase A4 |
| C6 | sp\|P13671\|CO6_HUMAN | P13671 | CO6 | 729 | Complement component C6 |
| PLS3 | sp\|P13797\|PLST_HUMAN | P13797 | PLST | 5358 | Plastin-3 |
| CD59 | sp\|P13987\|CD59_HUMAN | P13987 | CD59 | 966 | CD59 glycoprotein |
| HSD17B1 | sp\|P14061\|DHB1_HUMAN | P14061 | DHB1 | 3292 | 17-beta-hydroxysteroid dehydrogenase type 1 |
| CD99 | sp\|P14209\|CD99_HUMAN | P14209 | CD99 | 4267 | CD99 antigen |
| PRKCSH | sp\|P14314\|GLU2B_HUMAN | P14314 | GLU2B | 5589 | Glucosidase 2 subunit beta |
| FDPS | sp\|P14324\|FPPS_HUMAN | P14324 | FPPS | 2224 | Farnesyl pyrophosphate synthase |
| CPM | sp\|P14384\|CBPM_HUMAN | P14384 | CBPM | 1368 | Carboxypeptidase M |
| NID1 | sp\|P14543\|NID1_HUMAN | P14543 | NID1 | 4811 | Nidogen-1 |
| IDE | sp\|P14735\|IDE_HUMAN | P14735 | IDE | 3416 | Insulin-degrading enzyme |
| DARS1 | sp\|P14868\|SYDC_HUMAN | P14868 | SYDC | 1615 | Aspartate--tRNA ligase, cytoplasmic |
| GLUL | sp\|P15104\|GLNA_HUMAN | P15104 | GLNA | 2752 | Glutamine synthetase |
| AKR1B1 | sp\|P15121\|ALDR_HUMAN | P15121 | ALDR | 231 | Aldo-keto reductase family 1 member B1 |
| PVR | sp\|P15151\|PVR_HUMAN | P15151 | PVR | 5817 | Poliovirus receptor |
| ACP3 | sp\|P15309\|PPAP_HUMAN | P15309 | PPAP | 55 | Prostatic acid phosphatase |
| AREG | sp\|P15514\|AREG_HUMAN | P15514 | AREG | 374 | Amphiregulin |
| CD46 | sp\|P15529\|MCP_HUMAN | P15529 | MCP | 4179 | Membrane cofactor protein |
| RPS2 | sp\|P15880\|RS2_HUMAN | P15880 | RS2 | 6187 | 40S ribosomal protein S2 |
| DSP | sp\|P15924\|DESP_HUMAN | P15924 | DESP | 1832 | Desmoplakin |
| MUC1 | sp\|P15941\|MUC1_HUMAN | P15941 | MUC1 | 4582 | Mucin-1 |
| ACAN | sp\|P16112\|PGCA_HUMAN | P16112 | PGCA | 176 | Aggrecan core protein |
| POR | sp\|P16435\|NCPR_HUMAN | P16435 | NCPR | 5447 | NADPH--cytochrome P450 reductase |
| GOT1 | sp\|P17174\|AATC_HUMAN | P17174 | AATC | 2805 | Aspartate aminotransferase, cytoplasmic |
| ITGA2 | sp\|P17301\|ITA2_HUMAN | P17301 | ITA2 | 3673 | Integrin alpha-2 |
| GJA1 | sp\|P17302\|CXA1_HUMAN | P17302 | CXA1 | 2697 | Gap junction alpha-1 protein |
| PFKL | sp\|P17858\|PFKAL_HUMAN | P17858 | PFKAL | 5211 | ATP-dependent 6-phosphofructokinase, liver type |
| IGFBP3 | sp\|P17936\|IBP3_HUMAN | P17936 | IBP3 | 3486 | Insulin-like growth factor-binding protein 3 |
| PSMC3 | sp\|P17980\|PRS6A_HUMAN | P17980 | PRS6A | 5702 | 26S proteasome regulatory subunit 6A |
| TCP1 | sp\|P17987\|TCPA_HUMAN | P17987 | TCPA | 6950 | T-complex protein 1 subunit alpha |
| IGFBP2 | sp\|P18065\|IBP2_HUMAN | P18065 | IBP2 | 3485 | Insulin-like growth factor-binding protein 2 |
| ARF4 | sp\|P18085\|ARF4_HUMAN | P18085 | ARF4 | 378 | ADP-ribosylation factor 4 |
| RPL7 | sp\|P18124\|RL7_HUMAN | P18124 | RL7 | 6129 | 60S ribosomal protein L7 |
| VCL | sp\|P18206\|VINC_HUMAN | P18206 | VINC | 7414 | Vinculin |
| LBP | sp\|P18428\|LBP_HUMAN | P18428 | LBP | 3929 | Lipopolysaccharide-binding protein |
| PAM | sp\|P19021\|AMD_HUMAN | P19021 | AMD | 5066 | Peptidyl-glycine alpha-amidating monooxygenase |
| CDH2 | sp\|P19022\|CADH2_HUMAN | P19022 | CADH2 | 1000 | Cadherin-2 |
| VCAM1 | sp\|P19320\|VCAM1_HUMAN | P19320 | VCAM1 | 7412 | Vascular cell adhesion protein 1 |
| ORM2 | sp\|P19652\|A1AG2_HUMAN | P19652 | A1AG2 | 5005 | Alpha-1-acid glycoprotein 2 |
| FST | sp\|P19883\|FST_HUMAN | P19883 | FST | 10468 | Follistatin |
| PI3 | sp\|P19957\|ELAF_HUMAN | P19957 | ELAF | 5266 | Elafin |
| ANXA7 | sp\|P20073\|ANXA7_HUMAN | P20073 | ANXA7 | 310 | Annexin A7 |
| PSMB1 | sp\|P20618\|PSB1_HUMAN | P20618 | PSB1 | 5689 | Proteasome subunit beta type-1 |
| M6PR | sp\|P20645\|MPRD_HUMAN | P20645 | MPRD | 4074 | Cation-dependent mannose-6-phosphate receptor |
| C4BPB | sp\|P20851\|C4BPB_HUMAN | P20851 | C4BPB | 725 | C4b-binding protein beta chain |
| FLG | sp\|P20930\|FILA_HUMAN | P20930 | FILA | 2312 | Filaggrin |
| ATP6V1B2 | sp\|P21281\|VATB2_HUMAN | P21281 | VATB2 | 526 | V-type proton ATPase subunit B, brain isoform |
| MDK | sp\|P21741\|MK_HUMAN | P21741 | MK | 4192 | Midkine |
| VDAC1 | sp\|P21796\|VDAC1_HUMAN | P21796 | VDAC1 | 7416 | Voltage-dependent anion-selective channel protein 1 |
| CD9 | sp\|P21926\|CD9_HUMAN | P21926 | CD9 | 928 | CD9 antigen |
| COMT | sp\|P21964\|COMT_HUMAN | P21964 | COMT | 1312 | Catechol O-methyltransferase |
| PAICS | sp\|P22234\|PUR6_HUMAN | P22234 | PUR6 | 10606 | Multifunctional protein ADE2 |
| UBA1 | sp\|P22314\|UBA1_HUMAN | P22314 | UBA1 | 7317 | Ubiquitin-like modifier-activating enzyme 1 |
| GPX3 | sp\|P22352\|GPX3_HUMAN | P22352 | GPX3 | 2878 | Glutathione peroxidase 3 |
| IGFBP4 | sp\|P22692\|IBP4_HUMAN | P22692 | IBP4 | 3487 | Insulin-like growth factor-binding protein 4 |
| UQCRC2 | sp\|P22695\|QCR2_HUMAN | P22695 | QCR2 | 7385 | Cytochrome b-c1 complex subunit 2, mitochondrial |
| TGM1 | sp\|P22735\|TGM1_HUMAN | P22735 | TGM1 | 7051 | Protein-glutamine gamma-glutamyltransferase K |
| AADAC | sp\|P22760\|AAAD_HUMAN | P22760 | AAAD | 13 | Arylacetamide deacetylase |
| CPN2 | sp\|P22792\|CPN2_HUMAN | P22792 | CPN2 | 1370 | Carboxypeptidase N subunit 2 |
| IGHV1-2 | sp\|P23083\|HV102_HUMAN | P23083 | HV102 | 28474 | Immunoglobulin heavy variable 1-2 |
| ITGA6 | sp\|P23229\|ITA6_HUMAN | P23229 | ITA6 | 3655 | Integrin alpha-6 |
| CA6 | sp\|P23280\|CAH6_HUMAN | P23280 | CAH6 | 765 | Carbonic anhydrase 6 |
| RPS3 | sp\|P23396\|RS3_HUMAN | P23396 | RS3 | 6188 | 40S ribosomal protein S3 |
| PTPRG | sp\|P23470\|PTPRG_HUMAN | P23470 | PTPRG | 5793 | Receptor-type tyrosine-protein phosphatase gamma |
| MCC | sp\|P23508\|CRCM_HUMAN | P23508 | CRCM | 4163 | Colorectal mutant cancer protein |
| AHCY | sp\|P23526\|SAHH_HUMAN | P23526 | SAHH | 191 | Adenosylhomocysteinase |
| CFL1 | sp\|P23528\|COF1_HUMAN | P23528 | COF1 | 1072 | Cofilin-1 |
| ATP5PB | sp\|P24539\|AT5F1_HUMAN | P24539 | AT5F1 | 515 | ATP synthase F(0) complex subunit B1, mitochondrial |
| AZGP1 | sp\|P25311\|ZA2G_HUMAN | P25311 | ZA2G | 563 | Zinc-alpha-2-glycoprotein |
| LAMA1 | sp\|P25391\|LAMA1_HUMAN | P25391 | LAMA1 | 284217 | Laminin subunit alpha-1 |
| PSMA1 | sp\|P25786\|PSA1_HUMAN | P25786 | PSA1 | 5682 | Proteasome subunit alpha type-1 |
| PSMA3 | sp\|P25788\|PSA3_HUMAN | P25788 | PSA3 | 5684 | Proteasome subunit alpha type-3 |
| PTX3 | sp\|P26022\|PTX3_HUMAN | P26022 | PTX3 | 5806 | Pentraxin-related protein PTX3 |
| RPL13 | sp\|P26373\|RL13_HUMAN | P26373 | RL13 | 6137 | 60S ribosomal protein L13 |
| TARS1 | sp\|P26639\|SYTC_HUMAN | P26639 | SYTC | 6897 | Threonine--tRNA ligase 1, cytoplasmic |
| EEF1G | sp\|P26641\|EF1G_HUMAN | P26641 | EF1G | 1937 | Elongation factor 1-gamma |
| FKBP2 | sp\|P26885\|FKBP2_HUMAN | P26885 | FKBP2 | 2286 | Peptidyl-prolyl cis-trans isomerase FKBP2 |
| YWHAQ | sp\|P27348\|1433T_HUMAN | P27348 | 1433T | 10971 | 14-3-3 protein theta |
| CALML3 | sp\|P27482\|CALL3_HUMAN | P27482 | CALL3 | 810 | Calmodulin-like protein 3 |
| CALR | sp\|P27797\|CALR_HUMAN | P27797 | CALR | 811 | Calreticulin |
| CFP | sp\|P27918\|PROP_HUMAN | P27918 | PROP | 5199 | Properdin |
| PSMA5 | sp\|P28066\|PSA5_HUMAN | P28066 | PSA5 | 5686 | Proteasome subunit alpha type-5 |
| PSMB4 | sp\|P28070\|PSB4_HUMAN | P28070 | PSB4 | 5692 | Proteasome subunit beta type-4 |
| PSMB5 | sp\|P28074\|PSB5_HUMAN | P28074 | PSB5 | 5693 | Proteasome subunit beta type-5 |
| ABCD3 | sp\|P28288\|ABCD3_HUMAN | P28288 | ABCD3 | 5825 | ATP-binding cassette sub-family D member 3 |
| LAP3 | sp\|P28838\|AMPL_HUMAN | P28838 | AMPL | 51056 | Cytosol aminopeptidase |
| HSD11B1 | sp\|P28845\|DHI1_HUMAN | P28845 | DHI1 | 3290 | 11-beta-hydroxysteroid dehydrogenase 1 |
| CRABP2 | sp\|P29373\|RABP2_HUMAN | P29373 | RABP2 | 1382 | Cellular retinoic acid-binding protein 2 |
| TKT | sp\|P29401\|TKT_HUMAN | P29401 | TKT | 7086 | Transketolase |
| SERPINB3 | sp\|P29508\|SPB3_HUMAN | P29508 | SPB3 | 6317 | Serpin B3 |
| SERPINA4 | sp\|P29622\|KAIN_HUMAN | P29622 | KAIN | 5267 | Kallistatin |
| EEF1D | sp\|P29692\|EF1D_HUMAN | P29692 | EF1D | 1936 | Elongation factor 1-delta |
| MARCKS | sp\|P29966\|MARCS_HUMAN | P29966 | MARCS | 4082 | Myristoylated alanine-rich C-kinase substrate |
| ERP29 | sp\|P30040\|ERP29_HUMAN | P30040 | ERP29 | 10961 | Endoplasmic reticulum resident protein 29 |
| PRDX6 | sp\|P30041\|PRDX6_HUMAN | P30041 | PRDX6 | 9588 | Peroxiredoxin-6 |
| PRDX5 | sp\|P30044\|PRDX5_HUMAN | P30044 | PRDX5 | 25824 | Peroxiredoxin-5, mitochondrial |
| CMPK1 | sp\|P30085\|KCY_HUMAN | P30085 | KCY | 51727 | UMP-CMP kinase |
| PEBP1 | sp\|P30086\|PEBP1_HUMAN | P30086 | PEBP1 | 5037 | Phosphatidylethanolamine-binding protein 1 |
| PDIA3 | sp\|P30101\|PDIA3_HUMAN | P30101 | PDIA3 | 2923 | Protein disulfide-isomerase A3 |
| PPP2R1A | sp\|P30153\|2AAA_HUMAN | P30153 | 2AAA | 5518 | Serine/threonine-protein phosphatase 2A 65 kDa regulatory subunit A alpha isoform |
| SERPINB1 | sp\|P30740\|ILEU_HUMAN | P30740 | ILEU | 1992 | Leukocyte elastase inhibitor |
| NTS | sp\|P30990\|NEUT_HUMAN | P30990 | NEUT | 4922 | Neurotensin/neuromedin N |
| GDI1 | sp\|P31150\|GDIA_HUMAN | P31150 | GDIA | 2664 | Rab GDP dissociation inhibitor alpha |
| PRKAR2B | sp\|P31323\|KAP3_HUMAN | P31323 | KAP3 | 5577 | cAMP-dependent protein kinase type II-beta regulatory subunit |
| UQCRC1 | sp\|P31930\|QCR1_HUMAN | P31930 | QCR1 | 7384 | Cytochrome b-c1 complex subunit 1, mitochondrial |
| ATIC | sp\|P31939\|PUR9_HUMAN | P31939 | PUR9 | 471 | Bifunctional purine biosynthesis protein ATIC |
| CASP14 | sp\|P31944\|CASPE_HUMAN | P31944 | CASPE | 23581 | Caspase-14 |
| YWHAB | sp\|P31946\|1433B_HUMAN | P31946 | 1433B | 7529 | 14-3-3 protein beta/alpha |
| SFN | sp\|P31947\|1433S_HUMAN | P31947 | 1433S | 2810 | 14-3-3 protein sigma |
| STIP1 | sp\|P31948\|STIP1_HUMAN | P31948 | STIP1 | 10963 | Stress-induced-phosphoprotein 1 |
| S100A11 | sp\|P31949\|S10AB_HUMAN | P31949 | S10AB | 6282 | Protein S100-A11 |
| PRDX2 | sp\|P32119\|PRDX2_HUMAN | P32119 | PRDX2 | 7001 | Peroxiredoxin-2 |
| CCKBR | sp\|P32239\|GASR_HUMAN | P32239 | GASR | 887 | Gastrin/cholecystokinin type B receptor |
| DSG3 | sp\|P32926\|DSG3_HUMAN | P32926 | DSG3 | 1830 | Desmoglein-3 |
| MAN1A1 | sp\|P33908\|MA1A1_HUMAN | P33908 | MA1A1 | 4121 | Mannosyl-oligosaccharide 1,2-alpha-mannosidase IA |
| RNASE4 | sp\|P34096\|RNAS4_HUMAN | P34096 | RNAS4 | 6038 | Ribonuclease 4 |
| SDC2 | sp\|P34741\|SDC2_HUMAN | P34741 | SDC2 | 6383 | Syndecan-2 |
| CTNNB1 | sp\|P35222\|CTNB1_HUMAN | P35222 | CTNB1 | 1499 | Catenin beta-1 |
| PHB1 | sp\|P35232\|PHB1_HUMAN | P35232 | PHB1 | 5245 | Prohibitin 1 |
| SAA4 | sp\|P35542\|SAA4_HUMAN | P35542 | SAA4 | 6291 | Serum amyloid A-4 protein |
| FBN1 | sp\|P35555\|FBN1_HUMAN | P35555 | FBN1 | 2200 | Fibrillin-1 |
| MYH9 | sp\|P35579\|MYH9_HUMAN | P35579 | MYH9 | 4627 | Myosin-9 |
| MYH10 | sp\|P35580\|MYH10_HUMAN | P35580 | MYH10 | 4628 | Myosin-10 |
| COPB2 | sp\|P35606\|COPB2_HUMAN | P35606 | COPB2 | 9276 | Coatomer subunit beta' |
| BSG | sp\|P35613\|BASI_HUMAN | P35613 | BASI | 682 | Basigin |
| IGFALS | sp\|P35858\|ALS_HUMAN | P35858 | ALS | 3483 | Insulin-like growth factor-binding protein complex acid labile subunit |
| PSMC2 | sp\|P35998\|PRS7_HUMAN | P35998 | PRS7 | 5701 | 26S proteasome regulatory subunit 7 |
| RPL4 | sp\|P36578\|RL4_HUMAN | P36578 | RL4 | 6124 | 60S ribosomal protein L4 |
| PGM1 | sp\|P36871\|PGM1_HUMAN | P36871 | PGM1 | 5236 | Phosphoglucomutase-1 |
| SERPINB5 | sp\|P36952\|SPB5_HUMAN | P36952 | SPB5 | 5268 | Serpin B5 |
| FDFT1 | sp\|P37268\|FDFT_HUMAN | P37268 | FDFT | 2222 | Squalene synthase |
| TAGLN2 | sp\|P37802\|TAGL2_HUMAN | P37802 | TAGL2 | 8407 | Transgelin-2 |
| ATP6V1A | sp\|P38606\|VATA_HUMAN | P38606 | VATA | 523 | V-type proton ATPase catalytic subunit A |
| HSPA9 | sp\|P38646\|GRP75_HUMAN | P38646 | GRP75 | 3313 | Stress-70 protein, mitochondrial |
| RPL3 | sp\|P39023\|RL3_HUMAN | P39023 | RL3 | 6122 | 60S ribosomal protein L3 |
| COL15A1 | sp\|P39059\|COFA1_HUMAN | P39059 | COFA1 | 1306 | Collagen alpha-1(XV) chain |
| COL18A1 | sp\|P39060\|COIA1_HUMAN | P39060 | COIA1 | 80781 | Collagen alpha-1(XVIII) chain |
| DDOST | sp\|P39656\|OST48_HUMAN | P39656 | OST48 | 1650 | Dolichyl-diphosphooligosaccharide--protein glycosyltransferase 48 kDa subunit |
| IL6ST | sp\|P40189\|IL6RB_HUMAN | P40189 | IL6RB | 3572 | Interleukin-6 receptor subunit beta |
| MDH1 | sp\|P40925\|MDHC_HUMAN | P40925 | MDHC | 4190 | Malate dehydrogenase, cytoplasmic |
| HAL | sp\|P42357\|HUTH_HUMAN | P42357 | HUTH | 3034 | Histidine ammonia-lyase |
| ECE1 | sp\|P42892\|ECE1_HUMAN | P42892 | ECE1 | 1889 | Endothelin-converting enzyme 1 |
| SLC1A4 | sp\|P43007\|SATT_HUMAN | P43007 | SATT | 6509 | Neutral amino acid transporter A |
| PAFAH1B1 | sp\|P43034\|LIS1_HUMAN | P43034 | LIS1 | 5048 | Platelet-activating factor acetylhydrolase IB subunit beta |
| BTD | sp\|P43251\|BTD_HUMAN | P43251 | BTD | 686 | Biotinidase |
| SSR1 | sp\|P43307\|SSRA_HUMAN | P43307 | SSRA | 6745 | Translocon-associated protein subunit alpha |
| NAMPT | sp\|P43490\|NAMPT_HUMAN | P43490 | NAMPT | 10135 | Nicotinamide phosphoribosyltransferase |
| VDAC2 | sp\|P45880\|VDAC2_HUMAN | P45880 | VDAC2 | 7417 | Voltage-dependent anion-selective channel protein 2 |
| USP5 | sp\|P45974\|UBP5_HUMAN | P45974 | UBP5 | 8078 | Ubiquitin carboxyl-terminal hydrolase 5 |
| RPL27A | sp\|P46776\|RL27A_HUMAN | P46776 | RL27A | 6157 | 60S ribosomal protein L27a |
| RPS9 | sp\|P46781\|RS9_HUMAN | P46781 | RS9 | 6203 | 40S ribosomal protein S9 |
| RPS5 | sp\|P46782\|RS5_HUMAN | P46782 | RS5 | 6193 | 40S ribosomal protein S5 |
| STT3A | sp\|P46977\|STT3A_HUMAN | P46977 | STT3A | 3703 | Dolichyl-diphosphooligosaccharide--protein glycosyltransferase subunit STT3A |
| CAPZA2 | sp\|P47755\|CAZA2_HUMAN | P47755 | CAZA2 | 830 | F-actin-capping protein subunit alpha-2 |
| CAPZB | sp\|P47756\|CAPZB_HUMAN | P47756 | CAPZB | 832 | F-actin-capping protein subunit beta |
| LGALS7B; LGALS7 | sp\|P47929\|LEG7_HUMAN | P47929 | LEG7 | 653499; 3963 | Galectin-7 |
| TFPI2 | sp\|P48307\|TFPI2_HUMAN | P48307 | TFPI2 | 7980 | Tissue factor pathway inhibitor 2 |
| LSS | sp\|P48449\|LSS_HUMAN | P48449 | LSS | 4047 | Lanosterol synthase |
| GCLC | sp\|P48506\|GSH1_HUMAN | P48506 | GSH1 | 2729 | Glutamate--cysteine ligase catalytic subunit |
| SERPINB4 | sp\|P48594\|SPB4_HUMAN | P48594 | SPB4 | 6318 | Serpin B4 |
| CCT5 | sp\|P48643\|TCPE_HUMAN | P48643 | TCPE | 22948 | T-complex protein 1 subunit epsilon |
| MASP1 | sp\|P48740\|MASP1_HUMAN | P48740 | MASP1 | 5648 | Mannan-binding lectin serine protease 1 |
| MARCKSL1 | sp\|P49006\|MRP_HUMAN | P49006 | MRP | 65108 | MARCKS-related protein |
| ALDH9A1 | sp\|P49189\|AL9A1_HUMAN | P49189 | AL9A1 | 223 | 4-trimethylaminobutyraldehyde dehydrogenase |
| LMAN1 | sp\|P49257\|LMAN1_HUMAN | P49257 | LMAN1 | 3998 | Protein ERGIC-53 |
| FASN | sp\|P49327\|FAS_HUMAN | P49327 | FAS | 2194 | Fatty acid synthase |
| CCT3 | sp\|P49368\|TCPG_HUMAN | P49368 | TCPG | 7203 | T-complex protein 1 subunit gamma |
| ALDH7A1 | sp\|P49419\|AL7A1_HUMAN | P49419 | AL7A1 | 501 | Alpha-aminoadipic semialdehyde dehydrogenase |
| AARS1 | sp\|P49588\|SYAC_HUMAN | P49588 | SYAC | 16 | Alanine--tRNA ligase, cytoplasmic |
| TMED10 | sp\|P49755\|TMEDA_HUMAN | P49755 | TMEDA | 10972 | Transmembrane emp24 domain-containing protein 10 |
| SELENOP | sp\|P49908\|SEPP1_HUMAN | P49908 | SEPP1 | 6414 | Selenoprotein P |
| CAMP | sp\|P49913\|CAMP_HUMAN | P49913 | CAMP | 820 | Cathelicidin antimicrobial peptide |
| GDI2 | sp\|P50395\|GDIB_HUMAN | P50395 | GDIB | 2665 | Rab GDP dissociation inhibitor beta |
| EMD | sp\|P50402\|EMD_HUMAN | P50402 | EMD | 2010 | Emerin |
| CPT1A | sp\|P50416\|CPT1A_HUMAN | P50416 | CPT1A | 1374 | Carnitine O-palmitoyltransferase 1, liver isoform |
| SERPINH1 | sp\|P50454\|SERPH_HUMAN | P50454 | SERPH | 871 | Serpin H1 |
| CCT8 | sp\|P50990\|TCPQ_HUMAN | P50990 | TCPQ | 10694 | T-complex protein 1 subunit theta |
| CCT4 | sp\|P50991\|TCPD_HUMAN | P50991 | TCPD | 10575 | T-complex protein 1 subunit delta |
| RAB5C | sp\|P51148\|RAB5C_HUMAN | P51148 | RAB5C | 5878 | Ras-related protein Rab-5C |
| RAB7A | sp\|P51149\|RAB7A_HUMAN | P51149 | RAB7A | 7879 | Ras-related protein Rab-7a |
| SSR4 | sp\|P51571\|SSRD_HUMAN | P51571 | SSRD | 6748 | Translocon-associated protein subunit delta |
| BCAP31 | sp\|P51572\|BAP31_HUMAN | P51572 | BAP31 | 10134 | B-cell receptor-associated protein 31 |
| HSD17B4 | sp\|P51659\|DHB4_HUMAN | P51659 | DHB4 | 3295 | Peroxisomal multifunctional enzyme type 2 |
| HNRNPM | sp\|P52272\|HNRPM_HUMAN | P52272 | HNRPM | 4670 | Heterogeneous nuclear ribonucleoprotein M |
| ARHGDIA | sp\|P52565\|GDIR1_HUMAN | P52565 | GDIR1 | 396 | Rho GDP-dissociation inhibitor 1 |
| STC1 | sp\|P52823\|STC1_HUMAN | P52823 | STC1 | 6781 | Stanniocalcin-1 |
| CAPZA1 | sp\|P52907\|CAZA1_HUMAN | P52907 | CAZA1 | 829 | F-actin-capping protein subunit alpha-1 |
| ACLY | sp\|P53396\|ACLY_HUMAN | P53396 | ACLY | 47 | ATP-citrate synthase |
| MVD | sp\|P53602\|MVD1_HUMAN | P53602 | MVD1 | 4597 | Diphosphomevalonate decarboxylase |
| COPA | sp\|P53621\|COPA_HUMAN | P53621 | COPA | 1314 | Coatomer subunit alpha |
| SLC16A1 | sp\|P53985\|MOT1_HUMAN | P53985 | MOT1 | 6566 | Monocarboxylate transporter 1 |
| CACNA2D1 | sp\|P54289\|CA2D1_HUMAN | P54289 | CA2D1 | 781 | Voltage-dependent calcium channel subunit alpha-2/delta-1 |
| ATP1B3 | sp\|P54709\|AT1B3_HUMAN | P54709 | AT1B3 | 483 | Sodium/potassium-transporting ATPase subunit beta-3 |
| SLC12A2 | sp\|P55011\|S12A2_HUMAN | P55011 | S12A2 | 6558 | Solute carrier family 12 member 2 |
| APOC4 | sp\|P55056\|APOC4_HUMAN | P55056 | APOC4 | 346 | Apolipoprotein C-IV |
| PLTP | sp\|P55058\|PLTP_HUMAN | P55058 | PLTP | 5360 | Phospholipid transfer protein |
| TMBIM6 | sp\|P55061\|BI1_HUMAN | P55061 | BI1 | 7009 | Bax inhibitor 1 |
| MTTP | sp\|P55157\|MTP_HUMAN | P55157 | MTP | 4547 | Microsomal triglyceride transfer protein large subunit |
| NAP1L1 | sp\|P55209\|NP1L1_HUMAN | P55209 | NP1L1 | 4673 | Nucleosome assembly protein 1-like 1 |
| LAMB2 | sp\|P55268\|LAMB2_HUMAN | P55268 | LAMB2 | 3913 | Laminin subunit beta-2 |
| MARS1 | sp\|P56192\|SYMC_HUMAN | P56192 | SYMC | 4141 | Methionine--tRNA ligase, cytoplasmic |
| TMEM33 | sp\|P57088\|TMM33_HUMAN | P57088 | TMM33 | 55161 | Transmembrane protein 33 |
| ARPC4 | sp\|P59998\|ARPC4_HUMAN | P59998 | ARPC4 | 10093 | Actin-related protein 2/3 complex subunit 4 |
| CD81 | sp\|P60033\|CD81_HUMAN | P60033 | CD81 | 975 | CD81 antigen |
| TPI1 | sp\|P60174\|TPIS_HUMAN | P60174 | TPIS | 7167 | Triosephosphate isomerase |
| PLP1 | sp\|P60201\|MYPR_HUMAN | P60201 | MYPR | 5354 | Myelin proteolipid protein |
| SEC61B | sp\|P60468\|SC61B_HUMAN | P60468 | SC61B | 10952 | Protein transport protein Sec61 subunit beta |
| PSMA6 | sp\|P60900\|PSA6_HUMAN | P60900 | PSA6 | 5687 | Proteasome subunit alpha type-6 |
| DSTN | sp\|P60981\|DEST_HUMAN | P60981 | DEST | 11034 | Destrin |
| SPCS3 | sp\|P61009\|SPCS3_HUMAN | P61009 | SPCS3 | 60559 | Signal peptidase complex subunit 3 |
| RAB2A | sp\|P61019\|RAB2A_HUMAN | P61019 | RAB2A | 5862 | Ras-related protein Rab-2A |
| RAB5B | sp\|P61020\|RAB5B_HUMAN | P61020 | RAB5B | 5869 | Ras-related protein Rab-5B |
| UBE2N | sp\|P61088\|UBE2N_HUMAN | P61088 | UBE2N | 7334 | Ubiquitin-conjugating enzyme E2 N |
| ACTR2 | sp\|P61160\|ARP2_HUMAN | P61160 | ARP2 | 10097 | Actin-related protein 2 |
| ACTR1A | sp\|P61163\|ACTZ_HUMAN | P61163 | ACTZ | 10121 | Alpha-centractin |
| ARF3 | sp\|P61204\|ARF3_HUMAN | P61204 | ARF3 | 377 | ADP-ribosylation factor 3 |
| RPS3A | sp\|P61247\|RS3A_HUMAN | P61247 | RS3A | 6189 | 40S ribosomal protein S3a |
| RPL15 | sp\|P61313\|RL15_HUMAN | P61313 | RL15 | 6138 | 60S ribosomal protein L15 |
| RHOA | sp\|P61586\|RHOA_HUMAN | P61586 | RHOA | 387 | Transforming protein RhoA |
| HSPE1 | sp\|P61604\|CH10_HUMAN | P61604 | CH10 | 3336 | 10 kDa heat shock protein, mitochondrial |
| B2M | sp\|P61769\|B2MG_HUMAN | P61769 | B2MG | 567 | Beta-2-microglobulin |
| DAD1 | sp\|P61803\|DAD1_HUMAN | P61803 | DAD1 | 1603 | Dolichyl-diphosphooligosaccharide--protein glycosyltransferase subunit DAD1 |
| HNRNPK | sp\|P61978\|HNRPK_HUMAN | P61978 | HNRPK | 3190 | Heterogeneous nuclear ribonucleoprotein K |
| YWHAG | sp\|P61981\|1433G_HUMAN | P61981 | 1433G | 7532 | 14-3-3 protein gamma |
| PSMC5 | sp\|P62195\|PRS8_HUMAN | P62195 | PRS8 | 5705 | 26S proteasome regulatory subunit 8 |
| RPS8 | sp\|P62241\|RS8_HUMAN | P62241 | RS8 | 6202 | 40S ribosomal protein S8 |
| RPS15A | sp\|P62244\|RS15A_HUMAN | P62244 | RS15A | 6210 | 40S ribosomal protein S15a |
| RPS16 | sp\|P62249\|RS16_HUMAN | P62249 | RS16 | 6217 | 40S ribosomal protein S16 |
| YWHAE | sp\|P62258\|1433E_HUMAN | P62258 | 1433E | 7531 | 14-3-3 protein epsilon |
| RPS18 | sp\|P62269\|RS18_HUMAN | P62269 | RS18 | 6222 | 40S ribosomal protein S18 |
| RPS11 | sp\|P62280\|RS11_HUMAN | P62280 | RS11 | 6205 | 40S ribosomal protein S11 |
| TMSB4X | sp\|P62328\|TYB4_HUMAN | P62328 | TYB4 | 7114 | Thymosin beta-4 |
| PSMC6 | sp\|P62333\|PRS10_HUMAN | P62333 | PRS10 | 5706 | 26S proteasome regulatory subunit 10B |
| RPL7A | sp\|P62424\|RL7A_HUMAN | P62424 | RL7A | 6130 | 60S ribosomal protein L7a |
| RAB11A | sp\|P62491\|RB11A_HUMAN | P62491 | RB11A | 8766 | Ras-related protein Rab-11A |
| PPP2CB | sp\|P62714\|PP2AB_HUMAN | P62714 | PP2AB | 5516 | Serine/threonine-protein phosphatase 2A catalytic subunit beta isoform |
| RHOB | sp\|P62745\|RHOB_HUMAN | P62745 | RHOB | 388 | Rho-related GTP-binding protein RhoB |
| RPS6 | sp\|P62753\|RS6_HUMAN | P62753 | RS6 | 6194 | 40S ribosomal protein S6 |
| H4C2; H4-16; H4C11; H4C14; H4C9; H4C5; H4C12; H4C8; H4C13; H4C1; H4C3; H4C4; H4C6; H4C15 | sp\|P62805\|H4_HUMAN | P62805 | H4 | 8361; 8362; 8363; 8359; 8364; 8368; 121504; 8365; 8366; 554313; 8294; 8370; 8360; 8367 | Histone H4 |
| RAN | sp\|P62826\|RAN_HUMAN | P62826 | RAN | 5901 | GTP-binding nuclear protein Ran |
| RPL11 | sp\|P62913\|RL11_HUMAN | P62913 | RL11 | 6135 | 60S ribosomal protein L11 |
| RPL8 | sp\|P62917\|RL8_HUMAN | P62917 | RL8 | 6132 | 60S ribosomal protein L8 |
| YWHAZ | sp\|P63104\|1433Z_HUMAN | P63104 | 1433Z | 7534 | 14-3-3 protein zeta/delta |
| RACK1 | sp\|P63244\|RACK1_HUMAN | P63244 | RACK1 | 10399 | Receptor of activated protein C kinase 1 |
| YBX1 | sp\|P67809\|YBOX1_HUMAN | P67809 | YBOX1 | 4904 | Y-box-binding protein 1 |
| CCT2 | sp\|P78371\|TCPB_HUMAN | P78371 | TCPB | 10576 | T-complex protein 1 subunit beta |
| GSTO1 | sp\|P78417\|GSTO1_HUMAN | P78417 | GSTO1 | 9446 | Glutathione S-transferase omega-1 |
| PRKDC | sp\|P78527\|PRKDC_HUMAN | P78527 | PRKDC | 5591 | DNA-dependent protein kinase catalytic subunit |
| ADAM17 | sp\|P78536\|ADA17_HUMAN | P78536 | ADA17 | 6868 | Disintegrin and metalloproteinase domain-containing protein 17 |
| SRPX | sp\|P78539\|SRPX_HUMAN | P78539 | SRPX | 8406 | Sushi repeat-containing protein SRPX |
| PHEX | sp\|P78562\|PHEX_HUMAN | P78562 | PHEX | 5251 | Phosphate-regulating neutral endopeptidase PHEX |
| GPLD1 | sp\|P80108\|PHLD_HUMAN | P80108 | PHLD | 2822 | Phosphatidylinositol-glycan-specific phospholipase D |
| LCN2 | sp\|P80188\|NGAL_HUMAN | P80188 | NGAL | 3934 | Neutrophil gelatinase-associated lipocalin |
| NUCB2 | sp\|P80303\|NUCB2_HUMAN | P80303 | NUCB2 | 4925 | Nucleobindin-2 |
| HSD11B2 | sp\|P80365\|DHI2_HUMAN | P80365 | DHI2 | 3291 | 11-beta-hydroxysteroid dehydrogenase type 2 |
| BASP1 | sp\|P80723\|BASP1_HUMAN | P80723 | BASP1 | 10409 | Brain acid soluble protein 1 |
| DCD | sp\|P81605\|DCD_HUMAN | P81605 | DCD | 117159 | Dermcidin |
| RHOG | sp\|P84095\|RHOG_HUMAN | P84095 | RHOG | 391 | Rho-related GTP-binding protein RhoG |
| SRSF3 | sp\|P84103\|SRSF3_HUMAN | P84103 | SRSF3 | 6428 | Serine/arginine-rich splicing factor 3 |
| MXRA7 | sp\|P84157\|MXRA7_HUMAN | P84157 | MXRA7 | 439921 | Matrix-remodeling-associated protein 7 |
| ATP11A | sp\|P98196\|AT11A_HUMAN | P98196 | AT11A | 23250 | Phospholipid-transporting ATPase IH |
| SLC25A3 | sp\|Q00325\|MPCP_HUMAN | Q00325 | MPCP | 5250 | Phosphate carrier protein, mitochondrial |
| REEP5 | sp\|Q00765\|REEP5_HUMAN | Q00765 | REEP5 | 7905 | Receptor expression-enhancing protein 5 |
| HNRNPU | sp\|Q00839\|HNRPU_HUMAN | Q00839 | HNRPU | 3192 | Heterogeneous nuclear ribonucleoprotein U |
| CAP1 | sp\|Q01518\|CAP1_HUMAN | Q01518 | CAP1 | 10487 | Adenylyl cyclase-associated protein 1 |
| PFKP | sp\|Q01813\|PFKAP_HUMAN | Q01813 | PFKAP | 5214 | ATP-dependent 6-phosphofructokinase, platelet type |
| DSG1 | sp\|Q02413\|DSG1_HUMAN | Q02413 | DSG1 | 1828 | Desmoglein-1 |
| DSC2 | sp\|Q02487\|DSC2_HUMAN | Q02487 | DSC2 | 1824 | Desmocollin-2 |
| PLOD1 | sp\|Q02809\|PLOD1_HUMAN | Q02809 | PLOD1 | 5351 | Procollagen-lysine,2-oxoglutarate 5-dioxygenase 1 |
| NUCB1 | sp\|Q02818\|NUCB1_HUMAN | Q02818 | NUCB1 | 4924 | Nucleobindin-1 |
| RPL6 | sp\|Q02878\|RL6_HUMAN | Q02878 | RL6 | 6128 | 60S ribosomal protein L6 |
| TGFBR3 | sp\|Q03167\|TGBR3_HUMAN | Q03167 | TGBR3 | 7049 | Transforming growth factor beta receptor type 3 |
| TAP1 | sp\|Q03518\|TAP1_HUMAN | Q03518 | TAP1 | 6890 | Antigen peptide transporter 1 |
| TAP2 | sp\|Q03519\|TAP2_HUMAN | Q03519 | TAP2 | 6891 | Antigen peptide transporter 2 |
| YWHAH | sp\|Q04917\|1433F_HUMAN | Q04917 | 1433F | 7533 | 14-3-3 protein eta |
| ITIH3 | sp\|Q06033\|ITIH3_HUMAN | Q06033 | ITIH3 | 3699 | Inter-alpha-trypsin inhibitor heavy chain H3 |
| KDSR | sp\|Q06136\|KDSR_HUMAN | Q06136 | KDSR | 2531 | 3-ketodihydrosphingosine reductase |
| GFPT1 | sp\|Q06210\|GFPT1_HUMAN | Q06210 | GFPT1 | 2673 | Glutamine--fructose-6-phosphate aminotransferase [isomerizing] 1 |
| PSME1 | sp\|Q06323\|PSME1_HUMAN | Q06323 | PSME1 | 5720 | Proteasome activator complex subunit 1 |
| PRDX1 | sp\|Q06830\|PRDX1_HUMAN | Q06830 | PRDX1 | 5052 | Peroxiredoxin-1 |
| RPL18 | sp\|Q07020\|RL18_HUMAN | Q07020 | RL18 | 6141 | 60S ribosomal protein L18 |
| TGM3 | sp\|Q08188\|TGM3_HUMAN | Q08188 | TGM3 | 7053 | Protein-glutamine gamma-glutamyltransferase E |
| LGALS3BP | sp\|Q08380\|LG3BP_HUMAN | Q08380 | LG3BP | 3959 | Galectin-3-binding protein |
| MFGE8 | sp\|Q08431\|MFGM_HUMAN | Q08431 | MFGM | 4240 | Lactadherin |
| DSC1 | sp\|Q08554\|DSC1_HUMAN | Q08554 | DSC1 | 1823 | Desmocollin-1 |
| BST2 | sp\|Q10589\|BST2_HUMAN | Q10589 | BST2 | 684 | Bone marrow stromal antigen 2 |
| ST3GAL4 | sp\|Q11206\|SIA4C_HUMAN | Q11206 | SIA4C | 6484 | CMP-N-acetylneuraminate-beta-galactosamide-alpha-2,3-sialyltransferase 4 |
| STX4 | sp\|Q12846\|STX4_HUMAN | Q12846 | STX4 | 6810 | Syntaxin-4 |
| OVGP1 | sp\|Q12889\|OVGP1_HUMAN | Q12889 | OVGP1 | 5016 | Oviduct-specific glycoprotein |
| LMAN2 | sp\|Q12907\|LMAN2_HUMAN | Q12907 | LMAN2 | 10960 | Vesicular integral-membrane protein VIP36 |
| ECH1 | sp\|Q13011\|ECH1_HUMAN | Q13011 | ECH1 | 1891 | Delta(3,5)-Delta(2,4)-dienoyl-CoA isomerase, mitochondrial |
| SPP2 | sp\|Q13103\|SPP24_HUMAN | Q13103 | SPP24 | 6694 | Secreted phosphoprotein 24 |
| PRDX4 | sp\|Q13162\|PRDX4_HUMAN | Q13162 | PRDX4 | 10549 | Peroxiredoxin-4 |
| PSMD2 | sp\|Q13200\|PSMD2_HUMAN | Q13200 | PSMD2 | 5708 | 26S proteasome non-ATPase regulatory subunit 2 |
| PAPPA | sp\|Q13219\|PAPP1_HUMAN | Q13219 | PAPP1 | 5069 | Pappalysin-1 |
| DYNC1I2 | sp\|Q13409\|DC1I2_HUMAN | Q13409 | DC1I2 | 1781 | Cytoplasmic dynein 1 intermediate chain 2 |
| TMED1 | sp\|Q13445\|TMED1_HUMAN | Q13445 | TMED1 | 11018 | Transmembrane emp24 domain-containing protein 1 |
| DCTN2 | sp\|Q13561\|DCTN2_HUMAN | Q13561 | DCTN2 | 10540 | Dynactin subunit 2 |
| STIM1 | sp\|Q13586\|STIM1_HUMAN | Q13586 | STIM1 | 6786 | Stromal interaction molecule 1 |
| SNX1 | sp\|Q13596\|SNX1_HUMAN | Q13596 | SNX1 | 6642 | Sorting nexin-1 |
| ITGA7 | sp\|Q13683\|ITA7_HUMAN | Q13683 | ITA7 | 3679 | Integrin alpha-7 |
| MOGS | sp\|Q13724\|MOGS_HUMAN | Q13724 | MOGS | 7841 | Mannosyl-oligosaccharide glucosidase |
| APOF | sp\|Q13790\|APOF_HUMAN | Q13790 | APOF | 319 | Apolipoprotein F |
| SPTAN1 | sp\|Q13813\|SPTN1_HUMAN | Q13813 | SPTN1 | 6709 | Spectrin alpha chain, non-erythrocytic 1 |
| BLMH | sp\|Q13867\|BLMH_HUMAN | Q13867 | BLMH | 642 | Bleomycin hydrolase |
| BMPR2 | sp\|Q13873\|BMPR2_HUMAN | Q13873 | BMPR2 | 659 | Bone morphogenetic protein receptor type-2 |
| CEACAM7 | sp\|Q14002\|CEAM7_HUMAN | Q14002 | CEAM7 | 1087 | Carcinoembryonic antigen-related cell adhesion molecule 7 |
| SCARB2 | sp\|Q14108\|SCRB2_HUMAN | Q14108 | SCRB2 | 950 | Lysosome membrane protein 2 |
| NID2 | sp\|Q14112\|NID2_HUMAN | Q14112 | NID2 | 22795 | Nidogen-2 |
| LRP8 | sp\|Q14114\|LRP8_HUMAN | Q14114 | LRP8 | 7804 | Low-density lipoprotein receptor-related protein 8 |
| SCRIB | sp\|Q14160\|SCRIB_HUMAN | Q14160 | SCRIB | 23513 | Protein scribble homolog |
| MLEC | sp\|Q14165\|MLEC_HUMAN | Q14165 | MLEC | 9761 | Malectin |
| DCTN1 | sp\|Q14203\|DCTN1_HUMAN | Q14203 | DCTN1 | 1639 | Dynactin subunit 1 |
| DYNC1H1 | sp\|Q14204\|DYHC1_HUMAN | Q14204 | DYHC1 | 1778 | Cytoplasmic dynein 1 heavy chain 1 |
| FLOT2 | sp\|Q14254\|FLOT2_HUMAN | Q14254 | FLOT2 | 2319 | Flotillin-2 |
| HABP2 | sp\|Q14520\|HABP2_HUMAN | Q14520 | HABP2 | 3026 | Hyaluronan-binding protein 2 |
| SQLE | sp\|Q14534\|ERG1_HUMAN | Q14534 | ERG1 | 6713 | Squalene monooxygenase |
| PDIA5 | sp\|Q14554\|PDIA5_HUMAN | Q14554 | PDIA5 | 10954 | Protein disulfide-isomerase A5 |
| DSC3 | sp\|Q14574\|DSC3_HUMAN | Q14574 | DSC3 | 1825 | Desmocollin-3 |
| PLS1 | sp\|Q14651\|PLSI_HUMAN | Q14651 | PLSI | 5357 | Plastin-1 |
| GANAB | sp\|Q14697\|GANAB_HUMAN | Q14697 | GANAB | 23193 | Neutral alpha-glucosidase AB |
| MVP | sp\|Q14764\|MVP_HUMAN | Q14764 | MVP | 9961 | Major vault protein |
| GOLGB1 | sp\|Q14789\|GOGB1_HUMAN | Q14789 | GOGB1 | 2804 | Golgin subfamily B member 1 |
| KPNB1 | sp\|Q14974\|IMB1_HUMAN | Q14974 | IMB1 | 3837 | Importin subunit beta-1 |
| CLCA4 | sp\|Q14CN2\|CLCA4_HUMAN | Q14CN2 | CLCA4 | 22802 | Calcium-activated chloride channel regulator 4 |
| PSMD6 | sp\|Q15008\|PSMD6_HUMAN | Q15008 | PSMD6 | 9861 | 26S proteasome non-ATPase regulatory subunit 6 |
| SLC39A14 | sp\|Q15043\|S39AE_HUMAN | Q15043 | S39AE | 23516 | Metal cation symporter ZIP14 |
| PDIA6 | sp\|Q15084\|PDIA6_HUMAN | Q15084 | PDIA6 | 10130 | Protein disulfide-isomerase A6 |
| PCOLCE | sp\|Q15113\|PCOC1_HUMAN | Q15113 | PCOC1 | 5118 | Procollagen C-endopeptidase enhancer 1 |
| EBP | sp\|Q15125\|EBP_HUMAN | Q15125 | EBP | 10682 | 3-beta-hydroxysteroid-Delta(8),Delta(7)-isomerase |
| PON2 | sp\|Q15165\|PON2_HUMAN | Q15165 | PON2 | 5445 | Serum paraoxonase/arylesterase 2 |
| RCN1 | sp\|Q15293\|RCN1_HUMAN | Q15293 | RCN1 | 5954 | Reticulocalbin-1 |
| TMED2 | sp\|Q15363\|TMED2_HUMAN | Q15363 | TMED2 | 10959 | Transmembrane emp24 domain-containing protein 2 |
| DHCR24 | sp\|Q15392\|DHC24_HUMAN | Q15392 | DHC24 | 1718 | Delta(24)-sterol reductase |
| FCN2 | sp\|Q15485\|FCN2_HUMAN | Q15485 | FCN2 | 2220 | Ficolin-2 |
| CDSN | sp\|Q15517\|CDSN_HUMAN | Q15517 | CDSN | 1041 | Corneodesmosin |
| TGFBI | sp\|Q15582\|BGH3_HUMAN | Q15582 | BGH3 | 7045 | Transforming growth factor-beta-induced protein ig-h3 |
| NSDHL | sp\|Q15738\|NSDHL_HUMAN | Q15738 | NSDHL | 50814 | Sterol-4-alpha-carboxylate 3-dehydrogenase, decarboxylating |
| SLC1A5 | sp\|Q15758\|AAAT_HUMAN | Q15758 | AAAT | 6510 | Neutral amino acid transporter B(0) |
| MSMO1 | sp\|Q15800\|MSMO1_HUMAN | Q15800 | MSMO1 | 6307 | Methylsterol monooxygenase 1 |
| STXBP2 | sp\|Q15833\|STXB2_HUMAN | Q15833 | STXB2 | 6813 | Syntaxin-binding protein 2 |
| ADIPOQ | sp\|Q15848\|ADIPO_HUMAN | Q15848 | ADIPO | 9370 | Adiponectin |
| ECM1 | sp\|Q16610\|ECM1_HUMAN | Q16610 | ECM1 | 1893 | Extracellular matrix protein 1 |
| FSCN1 | sp\|Q16658\|FSCN1_HUMAN | Q16658 | FSCN1 | 6624 | Fascin |
| LAMA3 | sp\|Q16787\|LAMA3_HUMAN | Q16787 | LAMA3 | 3909 | Laminin subunit alpha-3 |
| CYP51A1 | sp\|Q16850\|CP51A_HUMAN | Q16850 | CP51A | 1595 | Lanosterol 14-alpha demethylase |
| UGP2 | sp\|Q16851\|UGPA_HUMAN | Q16851 | UGPA | 7360 | UTP--glucose-1-phosphate uridylyltransferase |
| P3H1 | sp\|Q32P28\|P3H1_HUMAN | Q32P28 | P3H1 | 64175 | Prolyl 3-hydroxylase 1 |
| ORMDL2 | sp\|Q53FV1\|ORML2_HUMAN | Q53FV1 | ORML2 | 29095 | ORM1-like protein 2 |
| HSD17B12 | sp\|Q53GQ0\|DHB12_HUMAN | Q53GQ0 | DHB12 | 51144 | Very-long-chain 3-oxoacyl-CoA reductase |
| FLG2 | sp\|Q5D862\|FILA2_HUMAN | Q5D862 | FILA2 | 388698 | Filaggrin-2 |
| MIA3 | sp\|Q5JRA6\|TGO1_HUMAN | Q5JRA6 | TGO1 | 375056 | Transport and Golgi organization protein 1 homolog |
| FREM2 | sp\|Q5SZK8\|FREM2_HUMAN | Q5SZK8 | FREM2 | 341640 | FRAS1-related extracellular matrix protein 2 |
| KPRP | sp\|Q5T749\|KPRP_HUMAN | Q5T749 | KPRP | 448834 | Keratinocyte proline-rich protein |
| XP32 | sp\|Q5T750\|XP32_HUMAN | Q5T750 | XP32 | 1E+08 | Skin-specific protein 32 |
| ACBD5 | sp\|Q5T8D3\|ACBD5_HUMAN | Q5T8D3 | ACBD5 | 91452 | Acyl-CoA-binding domain-containing protein 5 |
| CD276 | sp\|Q5ZPR3\|CD276_HUMAN | Q5ZPR3 | CD276 | 80381 | CD276 antigen |
| MBLAC2 | sp\|Q68D91\|MBLC2_HUMAN | Q68D91 | MBLC2 | 153364 | Acyl-coenzyme A thioesterase MBLAC2 |
| ATL3 | sp\|Q6DD88\|ATLA3_HUMAN | Q6DD88 | ATLA3 | 25923 | Atlastin-3 |
| DMKN | sp\|Q6E0U4\|DMKN_HUMAN | Q6E0U4 | DMKN | 93099 | Dermokine |
| GOLM2 | sp\|Q6P4E1\|GOLM2_HUMAN | Q6P4E1 | GOLM2 | 113201 | Protein GOLM2 |
| NCEH1 | sp\|Q6PIU2\|NCEH1_HUMAN | Q6PIU2 | NCEH1 | 57552 | Neutral cholesterol ester hydrolase 1 |
| APOA5 | sp\|Q6Q788\|APOA5_HUMAN | Q6Q788 | APOA5 | 116519 | Apolipoprotein A-V |
| AGTRAP | sp\|Q6RW13\|ATRAP_HUMAN | Q6RW13 | ATRAP | 57085 | Type-1 angiotensin II receptor-associated protein |
| TMEM205 | sp\|Q6UW68\|TM205_HUMAN | Q6UW68 | TM205 | 374882 | Transmembrane protein 205 |
| SBSN | sp\|Q6UWP8\|SBSN_HUMAN | Q6UWP8 | SBSN | 374897 | Suprabasin |
| OLFM4 | sp\|Q6UX06\|OLFM4_HUMAN | Q6UX06 | OLFM4 | 10562 | Olfactomedin-4 |
| PI16 | sp\|Q6UXB8\|PI16_HUMAN | Q6UXB8 | PI16 | 221476 | Peptidase inhibitor 16 |
| GBP6 | sp\|Q6ZN66\|GBP6_HUMAN | Q6ZN66 | GBP6 | 163351 | Guanylate-binding protein 6 |
| TMTC3 | sp\|Q6ZXV5\|TMTC3_HUMAN | Q6ZXV5 | TMTC3 | 160418 | Protein O-mannosyl-transferase TMTC3 |
| SND1 | sp\|Q7KZF4\|SND1_HUMAN | Q7KZF4 | SND1 | 27044 | Staphylococcal nuclease domain-containing protein 1 |
| CYFIP1 | sp\|Q7L576\|CYFP1_HUMAN | Q7L576 | CYFP1 | 23191 | Cytoplasmic FMR1-interacting protein 1 |
| ERMP1 | sp\|Q7Z2K6\|ERMP1_HUMAN | Q7Z2K6 | ERMP1 | 79956 | Endoplasmic reticulum metallopeptidase 1 |
| TMED4 | sp\|Q7Z7H5\|TMED4_HUMAN | Q7Z7H5 | TMED4 | 222068 | Transmembrane emp24 domain-containing protein 4 |
| SCD5 | sp\|Q86SK9\|SCD5_HUMAN | Q86SK9 | SCD5 | 79966 | Stearoyl-CoA desaturase 5 |
| TMPRSS11B | sp\|Q86T26\|TM11B_HUMAN | Q86T26 | TM11B | 132724 | Transmembrane protease serine 11B |
| KTN1 | sp\|Q86UP2\|KTN1_HUMAN | Q86UP2 | KTN1 | 3895 | Kinectin |
| LSR | sp\|Q86X29\|LSR_HUMAN | Q86X29 | LSR | 51599 | Lipolysis-stimulated lipoprotein receptor |
| FRAS1 | sp\|Q86XX4\|FRAS1_HUMAN | Q86XX4 | FRAS1 | 80144 | Extracellular matrix organizing protein FRAS1 |
| STX12 | sp\|Q86Y82\|STX12_HUMAN | Q86Y82 | STX12 | 23673 | Syntaxin-12 |
| HRNR | sp\|Q86YZ3\|HORN_HUMAN | Q86YZ3 | HORN | 388697 | Hornerin |
| PLD3 | sp\|Q8IV08\|PLD3_HUMAN | Q8IV08 | PLD3 | 23646 | 5'-3' exonuclease PLD3 |
| LRRC8A | sp\|Q8IWT6\|LRC8A_HUMAN | Q8IWT6 | LRC8A | 56262 | Volume-regulated anion channel subunit LRRC8A |
| SULF2 | sp\|Q8IWU5\|SULF2_HUMAN | Q8IWU5 | SULF2 | 55959 | Extracellular sulfatase Sulf-2 |
| CHPF | sp\|Q8IZ52\|CHSS2_HUMAN | Q8IZ52 | CHSS2 | 79586 | Chondroitin sulfate synthase 2 |
| ALDH16A1 | sp\|Q8IZ83\|A16A1_HUMAN | Q8IZ83 | A16A1 | 126133 | Aldehyde dehydrogenase family 16 member A1 |
| SFRP1 | sp\|Q8N474\|SFRP1_HUMAN | Q8N474 | SFRP1 | 6422 | Secreted frizzled-related protein 1 |
| CISD2 | sp\|Q8N5K1\|CISD2_HUMAN | Q8N5K1 | CISD2 | 493856 | CDGSH iron-sulfur domain-containing protein 2 |
| EMC1 | sp\|Q8N766\|EMC1_HUMAN | Q8N766 | EMC1 | 23065 | ER membrane protein complex subunit 1 |
| UBAC2 | sp\|Q8NBM4\|UBAC2_HUMAN | Q8NBM4 | UBAC2 | 337867 | Ubiquitin-associated domain-containing protein 2 |
| TXNDC5 | sp\|Q8NBS9\|TXND5_HUMAN | Q8NBS9 | TXND5 | 81567 | Thioredoxin domain-containing protein 5 |
| DIPK2A | sp\|Q8NDZ4\|DIK2A_HUMAN | Q8NDZ4 | DIK2A | 205428 | Divergent protein kinase domain 2A |
| APPL2 | sp\|Q8NEU8\|DP13B_HUMAN | Q8NEU8 | DP13B | 55198 | DCC-interacting protein 13-beta |
| SLC30A7 | sp\|Q8NEW0\|ZNT7_HUMAN | Q8NEW0 | ZNT7 | 148867 | Zinc transporter 7 |
| TOR1AIP2 | sp\|Q8NFQ8\|TOIP2_HUMAN | Q8NFQ8 | TOIP2 | 163590 | Torsin-1A-interacting protein 2 |
| GOLGA5 | sp\|Q8TBA6\|GOGA5_HUMAN | Q8TBA6 | GOGA5 | 9950 | Golgin subfamily A member 5 |
| RDH11 | sp\|Q8TC12\|RDH11_HUMAN | Q8TC12 | RDH11 | 51109 | Retinol dehydrogenase 11 |
| STT3B | sp\|Q8TCJ2\|STT3B_HUMAN | Q8TCJ2 | STT3B | 201595 | Dolichyl-diphosphooligosaccharide--protein glycosyltransferase subunit STT3B |
| HM13 | sp\|Q8TCT9\|HM13_HUMAN | Q8TCT9 | HM13 | 81502 | Minor histocompatibility antigen H13 |
| BPIFB1 | sp\|Q8TDL5\|BPIB1_HUMAN | Q8TDL5 | BPIB1 | 92747 | BPI fold-containing family B member 1 |
| LRRC15 | sp\|Q8TF66\|LRC15_HUMAN | Q8TF66 | LRC15 | 131578 | Leucine-rich repeat-containing protein 15 |
| SCARB1 | sp\|Q8WTV0\|SCRB1_HUMAN | Q8WTV0 | SCRB1 | 949 | Scavenger receptor class B member 1 |
| TSKU | sp\|Q8WUA8\|TSK_HUMAN | Q8WUA8 | TSK | 25987 | Tsukushi |
| PDCD6IP | sp\|Q8WUM4\|PDC6I_HUMAN | Q8WUM4 | PDC6I | 10015 | Programmed cell death 6-interacting protein |
| POF1B | sp\|Q8WVV4\|POF1B_HUMAN | Q8WVV4 | POF1B | 79983 | Protein POF1B |
| FAR1 | sp\|Q8WVX9\|FACR1_HUMAN | Q8WVX9 | FACR1 | 84188 | Fatty acyl-CoA reductase 1 |
| SLC44A1 | sp\|Q8WWI5\|CTL1_HUMAN | Q8WWI5 | CTL1 | 23446 | Choline transporter-like protein 1 |
| NCSTN | sp\|Q92542\|NICA_HUMAN | Q92542 | NICA | 23385 | Nicastrin |
| TM9SF4 | sp\|Q92544\|TM9S4_HUMAN | Q92544 | TM9S4 | 9777 | Transmembrane 9 superfamily member 4 |
| NDRG1 | sp\|Q92597\|NDRG1_HUMAN | Q92597 | NDRG1 | 10397 | Protein NDRG1 |
| PXDN | sp\|Q92626\|PXDN_HUMAN | Q92626 | PXDN | 7837 | Peroxidasin homolog |
| NECTIN2 | sp\|Q92692\|NECT2_HUMAN | Q92692 | NECT2 | 5819 | Nectin-2 |
| GLG1 | sp\|Q92896\|GSLG1_HUMAN | Q92896 | GSLG1 | 2734 | Golgi apparatus protein 1 |
| PRG4 | sp\|Q92954\|PRG4_HUMAN | Q92954 | PRG4 | 10216 | Proteoglycan 4 |
| ATP6V0A1 | sp\|Q93050\|VPP1_HUMAN | Q93050 | VPP1 | 535 | V-type proton ATPase 116 kDa subunit a 1 |
| MYDGF | sp\|Q969H8\|MYDGF_HUMAN | Q969H8 | MYDGF | 56005 | Myeloid-derived growth factor |
| IGSF8 | sp\|Q969P0\|IGSF8_HUMAN | Q969P0 | IGSF8 | 93185 | Immunoglobulin superfamily member 8 |
| NCLN | sp\|Q969V3\|NCLN_HUMAN | Q969V3 | NCLN | 56926 | Nicalin |
| ERGIC1 | sp\|Q969X5\|ERGI1_HUMAN | Q969X5 | ERGI1 | 57222 | Endoplasmic reticulum-Golgi intermediate compartment protein 1 |
| PBXIP1 | sp\|Q96AQ6\|PBIP1_HUMAN | Q96AQ6 | PBIP1 | 57326 | Pre-B-cell leukemia transcription factor-interacting protein 1 |
| FKBP10 | sp\|Q96AY3\|FKB10_HUMAN | Q96AY3 | FKB10 | 60681 | Peptidyl-prolyl cis-trans isomerase FKBP10 |
| FAF2 | sp\|Q96CS3\|FAF2_HUMAN | Q96CS3 | FAF2 | 23197 | FAS-associated factor 2 |
| HVCN1 | sp\|Q96D96\|HVCN1_HUMAN | Q96D96 | HVCN1 | 84329 | Voltage-gated hydrogen channel 1 |
| ZG16B | sp\|Q96DA0\|ZG16B_HUMAN | Q96DA0 | ZG16B | 124220 | Zymogen granule protein 16 homolog B |
| ERLEC1 | sp\|Q96DZ1\|ERLEC_HUMAN | Q96DZ1 | ERLEC | 27248 | Endoplasmic reticulum lectin 1 |
| SLC47A1 | sp\|Q96FL8\|S47A1_HUMAN | Q96FL8 | S47A1 | 55244 | Multidrug and toxin extrusion protein 1 |
| S100A16 | sp\|Q96FQ6\|S10AG_HUMAN | Q96FQ6 | S10AG | 140576 | Protein S100-A16 |
| ERO1A | sp\|Q96HE7\|ERO1A_HUMAN | Q96HE7 | ERO1A | 30001 | ERO1-like protein alpha |
| REEP6 | sp\|Q96HR9\|REEP6_HUMAN | Q96HR9 | REEP6 | 92840 | Receptor expression-enhancing protein 6 |
| RNF170 | sp\|Q96K19\|RN170_HUMAN | Q96K19 | RN170 | 81790 | E3 ubiquitin-protein ligase RNF170 |
| SLC35E1 | sp\|Q96K37\|S35E1_HUMAN | Q96K37 | S35E1 | 79939 | Solute carrier family 35 member E1 |
| CNDP1 | sp\|Q96KN2\|CNDP1_HUMAN | Q96KN2 | CNDP1 | 84735 | Beta-Ala-His dipeptidase |
| CNDP2 | sp\|Q96KP4\|CNDP2_HUMAN | Q96KP4 | CNDP2 | 55748 | Cytosolic non-specific dipeptidase |
| MBOAT7 | sp\|Q96N66\|MBOA7_HUMAN | Q96N66 | MBOA7 | 79143 | Lysophospholipid acyltransferase 7 |
| CLIC6 | sp\|Q96NY7\|CLIC6_HUMAN | Q96NY7 | CLIC6 | 54102 | Chloride intracellular channel protein 6 |
| VPS35 | sp\|Q96QK1\|VPS35_HUMAN | Q96QK1 | VPS35 | 55737 | Vacuolar protein sorting-associated protein 35 |
| PANX1 | sp\|Q96RD7\|PANX1_HUMAN | Q96RD7 | PANX1 | 24145 | Pannexin-1 |
| PIGS | sp\|Q96S52\|PIGS_HUMAN | Q96S52 | PIGS | 94005 | GPI transamidase component PIG-S |
| CLCC1 | sp\|Q96S66\|CLCC1_HUMAN | Q96S66 | CLCC1 | 23155 | Chloride channel CLIC-like protein 1 |
| NIBAN2 | sp\|Q96TA1\|NIBA2_HUMAN | Q96TA1 | NIBA2 | 64855 | Protein Niban 2 |
| PARK7 | sp\|Q99497\|PARK7_HUMAN | Q99497 | PARK7 | 11315 | Parkinson disease protein 7 |
| VAT1 | sp\|Q99536\|VAT1_HUMAN | Q99536 | VAT1 | 10493 | Synaptic vesicle membrane protein VAT-1 homolog |
| S100A13 | sp\|Q99584\|S10AD_HUMAN | Q99584 | S10AD | 6284 | Protein S100-A13 |
| PHB2 | sp\|Q99623\|PHB2_HUMAN | Q99623 | PHB2 | 11331 | Prohibitin-2 |
| CHP1 | sp\|Q99653\|CHP1_HUMAN | Q99653 | CHP1 | 11261 | Calcineurin B homologous protein 1 |
| TM9SF2 | sp\|Q99805\|TM9S2_HUMAN | Q99805 | TM9S2 | 9375 | Transmembrane 9 superfamily member 2 |
| SLC29A1 | sp\|Q99808\|S29A1_HUMAN | Q99808 | S29A1 | 2030 | Equilibrative nucleoside transporter 1 |
| TSG101 | sp\|Q99816\|TS101_HUMAN | Q99816 | TS101 | 7251 | Tumor susceptibility gene 101 protein |
| CCT7 | sp\|Q99832\|TCPH_HUMAN | Q99832 | TCPH | 10574 | T-complex protein 1 subunit eta |
| PRRT1 | sp\|Q99946\|PRRT1_HUMAN | Q99946 | PRRT1 | 80863 | Proline-rich transmembrane protein 1 |
| CORO1B | sp\|Q9BR76\|COR1B_HUMAN | Q9BR76 | COR1B | 57175 | Coronin-1B |
| SDF4 | sp\|Q9BRK5\|CAB45_HUMAN | Q9BRK5 | CAB45 | 51150 | 45 kDa calcium-binding protein |
| PRXL2A | sp\|Q9BRX8\|PXL2A_HUMAN | Q9BRX8 | PXL2A | 84293 | Peroxiredoxin-like 2A |
| ERP44 | sp\|Q9BS26\|ERP44_HUMAN | Q9BS26 | ERP44 | 23071 | Endoplasmic reticulum resident protein 44 |
| CNPY3 | sp\|Q9BT09\|CNPY3_HUMAN | Q9BT09 | CNPY3 | 10695 | Protein canopy homolog 3 |
| TMEM43 | sp\|Q9BTV4\|TMM43_HUMAN | Q9BTV4 | TMM43 | 79188 | Transmembrane protein 43 |
| LMF2 | sp\|Q9BU23\|LMF2_HUMAN | Q9BU23 | LMF2 | 91289 | Lipase maturation factor 2 |
| TMED9 | sp\|Q9BVK6\|TMED9_HUMAN | Q9BVK6 | TMED9 | 54732 | Transmembrane emp24 domain-containing protein 9 |
| ACAT2 | sp\|Q9BWD1\|THIC_HUMAN | Q9BWD1 | THIC | 39 | Acetyl-CoA acetyltransferase, cytosolic |
| SFXN3 | sp\|Q9BWM7\|SFXN3_HUMAN | Q9BWM7 | SFXN3 | 81855 | Sideroflexin-3 |
| CFHR5 | sp\|Q9BXR6\|FHR5_HUMAN | Q9BXR6 | FHR5 | 81494 | Complement factor H-related protein 5 |
| ANGPTL4 | sp\|Q9BY76\|ANGL4_HUMAN | Q9BY76 | ANGL4 | 51129 | Angiopoietin-related protein 4 |
| TTYH3 | sp\|Q9C0H2\|TTYH3_HUMAN | Q9C0H2 | TTYH3 | 80727 | Protein tweety homolog 3 |
| TOLLIP | sp\|Q9H0E2\|TOLIP_HUMAN | Q9H0E2 | TOLIP | 54472 | Toll-interacting protein |
| MAGT1 | sp\|Q9H0U3\|MAGT1_HUMAN | Q9H0U3 | MAGT1 | 84061 | Magnesium transporter protein 1 |
| FAM234A | sp\|Q9H0X4\|F234A_HUMAN | Q9H0X4 | F234A | 83986 | Protein FAM234A |
| TMX4 | sp\|Q9H1E5\|TMX4_HUMAN | Q9H1E5 | TMX4 | 56255 | Thioredoxin-related transmembrane protein 4 |
| EHD4 | sp\|Q9H223\|EHD4_HUMAN | Q9H223 | EHD4 | 30844 | EH domain-containing protein 4 |
| TMEM245 | sp\|Q9H330\|TM245_HUMAN | Q9H330 | TM245 | 23731 | Transmembrane protein 245 |
| TMX1 | sp\|Q9H3N1\|TMX1_HUMAN | Q9H3N1 | TMX1 | 81542 | Thioredoxin-related transmembrane protein 1 |
| CHMP4B | sp\|Q9H444\|CHM4B_HUMAN | Q9H444 | CHM4B | 128866 | Charged multivesicular body protein 4b |
| VPS37B | sp\|Q9H9H4\|VP37B_HUMAN | Q9H9H4 | VP37B | 79720 | Vacuolar protein sorting-associated protein 37B |
| SPON1 | sp\|Q9HCB6\|SPON1_HUMAN | Q9HCB6 | SPON1 | 10418 | Spondin-1 |
| APMAP | sp\|Q9HDC9\|APMAP_HUMAN | Q9HDC9 | APMAP | 57136 | Adipocyte plasma membrane-associated protein |
| RAB18 | sp\|Q9NP72\|RAB18_HUMAN | Q9NP72 | RAB18 | 22931 | Ras-related protein Rab-18 |
| RTN4 | sp\|Q9NQC3\|RTN4_HUMAN | Q9NQC3 | RTN4 | 57142 | Reticulon-4 |
| SERINC1 | sp\|Q9NRX5\|SERC1_HUMAN | Q9NRX5 | SERC1 | 57515 | Serine incorporator 1 |
| SACM1L | sp\|Q9NTJ5\|SAC1_HUMAN | Q9NTJ5 | SAC1 | 22908 | Phosphatidylinositol-3-phosphatase SAC1 |
| RAB20 | sp\|Q9NX57\|RAB20_HUMAN | Q9NX57 | RAB20 | 55647 | Ras-related protein Rab-20 |
| BPNT2 | sp\|Q9NX62\|IMPA3_HUMAN | Q9NX62 | IMPA3 | 54928 | Golgi-resident adenosine 3',5'-bisphosphate 3'-phosphatase |
| ELOVL5 | sp\|Q9NYP7\|ELOV5_HUMAN | Q9NYP7 | ELOV5 | 60481 | Elongation of very long chain fatty acids protein 5 |
| TECR | sp\|Q9NZ01\|TECR_HUMAN | Q9NZ01 | TECR | 9524 | Very-long-chain enoyl-CoA reductase |
| ERAP1 | sp\|Q9NZ08\|ERAP1_HUMAN | Q9NZ08 | ERAP1 | 51752 | Endoplasmic reticulum aminopeptidase 1 |
| C1RL | sp\|Q9NZP8\|C1RL_HUMAN | Q9NZP8 | C1RL | 51279 | Complement C1r subcomponent-like protein |
| CALML5 | sp\|Q9NZT1\|CALL5_HUMAN | Q9NZT1 | CALL5 | 51806 | Calmodulin-like protein 5 |
| HACD3 | sp\|Q9P035\|HACD3_HUMAN | Q9P035 | HACD3 | 51495 | Very-long-chain (3R)-3-hydroxyacyl-CoA dehydratase 3 |
| PTGFRN | sp\|Q9P2B2\|FPRP_HUMAN | Q9P2B2 | FPRP | 5738 | Prostaglandin F2 receptor negative regulator |
| SPRR3 | sp\|Q9UBC9\|SPRR3_HUMAN | Q9UBC9 | SPRR3 | 6707 | Small proline-rich protein 3 |
| CRNN | sp\|Q9UBG3\|CRNN_HUMAN | Q9UBG3 | CRNN | 49860 | Cornulin |
| GNG12 | sp\|Q9UBI6\|GBG12_HUMAN | Q9UBI6 | GBG12 | 55970 | Guanine nucleotide-binding protein G(I)/G(S)/G(O) subunit gamma-12 |
| DHCR7 | sp\|Q9UBM7\|DHCR7_HUMAN | Q9UBM7 | DHCR7 | 1717 | 7-dehydrocholesterol reductase |
| DNAJB9 | sp\|Q9UBS3\|DNJB9_HUMAN | Q9UBS3 | DNJB9 | 4189 | DnaJ homolog subfamily B member 9 |
| SEL1L | sp\|Q9UBV2\|SE1L1_HUMAN | Q9UBV2 | SE1L1 | 6400 | Protein sel-1 homolog 1 |
| FETUB | sp\|Q9UGM5\|FETUB_HUMAN | Q9UGM5 | FETUB | 26998 | Fetuin-B |
| SEPTIN9 | sp\|Q9UHD8\|SEPT9_HUMAN | Q9UHD8 | 40057 | 10801 | Septin-9 |
| PCYOX1 | sp\|Q9UHG3\|PCYOX_HUMAN | Q9UHG3 | PCYOX | 51449 | Prenylcysteine oxidase 1 |
| SERPINB13 | sp\|Q9UIV8\|SPB13_HUMAN | Q9UIV8 | SPB13 | 5275 | Serpin B13 |
| DBNL | sp\|Q9UJU6\|DBNL_HUMAN | Q9UJU6 | DBNL | 28988 | Drebrin-like protein |
| SERPINA10 | sp\|Q9UK55\|ZPI_HUMAN | Q9UK55 | ZPI | 51156 | Protein Z-dependent protease inhibitor |
| RAB21 | sp\|Q9UL25\|RAB21_HUMAN | Q9UL25 | RAB21 | 23011 | Ras-related protein Rab-21 |
| PSME2 | sp\|Q9UL46\|PSME2_HUMAN | Q9UL46 | PSME2 | 5721 | Proteasome activator complex subunit 2 |
| TMCO1 | sp\|Q9UM00\|TMCO1_HUMAN | Q9UM00 | TMCO1 | 54499 | Calcium load-activated calcium channel |
| NDRG2 | sp\|Q9UN36\|NDRG2_HUMAN | Q9UN36 | NDRG2 | 57447 | Protein NDRG2 |
| SNX6 | sp\|Q9UNH7\|SNX6_HUMAN | Q9UNH7 | SNX6 | 58533 | Sorting nexin-6 |
| PSMD13 | sp\|Q9UNM6\|PSD13_HUMAN | Q9UNM6 | PSD13 | 5719 | 26S proteasome non-ATPase regulatory subunit 13 |
| RUVBL2 | sp\|Q9Y230\|RUVB2_HUMAN | Q9Y230 | RUVB2 | 10856 | RuvB-like 2 |
| RUVBL1 | sp\|Q9Y265\|RUVB1_HUMAN | Q9Y265 | RUVB1 | 8607 | RuvB-like 1 |
| NUDC | sp\|Q9Y266\|NUDC_HUMAN | Q9Y266 | NUDC | 10726 | Nuclear migration protein nudC |
| NCKAP1 | sp\|Q9Y2A7\|NCKP1_HUMAN | Q9Y2A7 | NCKP1 | 10787 | Nck-associated protein 1 |
| CNPY2 | sp\|Q9Y2B0\|CNPY2_HUMAN | Q9Y2B0 | CNPY2 | 10330 | Protein canopy homolog 2 |
| TMED5 | sp\|Q9Y3A6\|TMED5_HUMAN | Q9Y3A6 | TMED5 | 50999 | Transmembrane emp24 domain-containing protein 5 |
| TMED7 | sp\|Q9Y3B3\|TMED7_HUMAN | Q9Y3B3 | TMED7 | 51014; 100302736 | Transmembrane emp24 domain-containing protein 7 |
| TLN1 | sp\|Q9Y490\|TLN1_HUMAN | Q9Y490 | TLN1 | 7094 | Talin-1 |
| HYOU1 | sp\|Q9Y4L1\|HYOU1_HUMAN | Q9Y4L1 | HYOU1 | 10525 | Hypoxia up-regulated protein 1 |
| TBL2 | sp\|Q9Y4P3\|TBL2_HUMAN | Q9Y4P3 | TBL2 | 26608 | Transducin beta-like protein 2 |
| SRPRB | sp\|Q9Y5M8\|SRPRB_HUMAN | Q9Y5M8 | SRPRB | 58477 | Signal recognition particle receptor subunit beta |
| F11R | sp\|Q9Y624\|JAM1_HUMAN | Q9Y624 | JAM1 | 50848 | Junctional adhesion molecule A |
| NPTN | sp\|Q9Y639\|NPTN_HUMAN | Q9Y639 | NPTN | 27020 | Neuroplastin |
| ADGRG1 | sp\|Q9Y653\|AGRG1_HUMAN | Q9Y653 | AGRG1 | 9289 | Adhesion G-protein coupled receptor G1 |
| ALG5 | sp\|Q9Y673\|ALG5_HUMAN | Q9Y673 | ALG5 | 29880 | Dolichyl-phosphate beta-glucosyltransferase |
| CLIC4 | sp\|Q9Y696\|CLIC4_HUMAN | Q9Y696 | CLIC4 | 25932 | Chloride intracellular channel protein 4 |
| EMILIN1 | sp\|Q9Y6C2\|EMIL1_HUMAN | Q9Y6C2 | EMIL1 | 11117 | EMILIN-1 |
| DYNC1LI1 | sp\|Q9Y6G9\|DC1L1_HUMAN | Q9Y6G9 | DC1L1 | 51143 | Cytoplasmic dynein 1 light intermediate chain 1 |
| TEX264 | sp\|Q9Y6I9\|TX264_HUMAN | Q9Y6I9 | TX264 | 51368 | Testis-expressed protein 264 |
| SLC30A1 | sp\|Q9Y6M5\|ZNT1_HUMAN | Q9Y6M5 | ZNT1 | 7779 | Zinc transporter 1 |
| IGKV3D-15 | sp\|A0A087WSY6\|KVD15_HUMAN | A0A087WSY6 | KVD15 | 28875 | Immunoglobulin kappa variable 3D-15 |
| IGKV3-7 | sp\|A0A075B6H7\|KV37_HUMAN | A0A075B6H7 | KV37 | 28915 | Probable non-functional immunoglobulin kappa variable 3-7 |
| IGKV6D-21 | sp\|A0A0A0MT36\|KVD21_HUMAN | A0A0A0MT36 | KVD21 | 28870 | Immunoglobulin kappa variable 6D-21 |
| IGKV6-21 | sp\|A0A0C4DH24\|KV621_HUMAN | A0A0C4DH24 | KV621 | 28906 | Immunoglobulin kappa variable 6-21 |
| IGHV1-69D | sp\|A0A0B4J2H0\|HV69D_HUMAN | A0A0B4J2H0 | HV69D | 1.03E+08 | Immunoglobulin heavy variable 1-69D |
| IGHV1-3 | sp\|A0A0C4DH29\|HV103_HUMAN | A0A0C4DH29 | HV103 | 28473 | Immunoglobulin heavy variable 1-3 |
| IGHV1-46 | sp\|P01743\|HV146_HUMAN | P01743 | HV146 | 28465 | Immunoglobulin heavy variable 1-46 |
| IGKV1-8 | sp\|A0A0C4DH67\|KV108_HUMAN | A0A0C4DH67 | KV108 | 28942 | Immunoglobulin kappa variable 1-8 |
| IGKV1-27 | sp\|A0A075B6S5\|KV127_HUMAN | A0A075B6S5 | KV127 | 28935 | Immunoglobulin kappa variable 1-27 |
| LEFTY2 | sp\|O00292\|LFTY2_HUMAN | O00292 | LFTY2 | 7044 | Left-right determination factor 2 |
| DDX3X | sp\|O00571\|DDX3X_HUMAN | O00571 | DDX3X | 1654 | ATP-dependent RNA helicase DDX3X |
| PSMA7 | sp\|O14818\|PSA7_HUMAN | O14818 | PSA7 | 5688 | Proteasome subunit alpha type-7 |
| PHGDH | sp\|O43175\|SERA_HUMAN | O43175 | SERA | 26227 | D-3-phosphoglycerate dehydrogenase |
| EPB41L2 | sp\|O43491\|E41L2_HUMAN | O43491 | E41L2 | 2037 | Band 4.1-like protein 2 |
| ACTN4 | sp\|O43707\|ACTN4_HUMAN | O43707 | ACTN4 | 81 | Alpha-actinin-4 |
| MYO1B | sp\|O43795\|MYO1B_HUMAN | O43795 | MYO1B | 4430 | Unconventional myosin-Ib |
| CD5L | sp\|O43866\|CD5L_HUMAN | O43866 | CD5L | 922 | CD5 antigen-like |
| AKR1B10 | sp\|O60218\|AK1BA_HUMAN | O60218 | AK1BA | 57016 | Aldo-keto reductase family 1 member B10 |
| H2BC12 | sp\|O60814\|H2B1K_HUMAN | O60814 | H2B1K | 85236 | Histone H2B type 1-K |
| ADAMTS4 | sp\|O75173\|ATS4_HUMAN | O75173 | ATS4 | 9507 | A disintegrin and metalloproteinase with thrombospondin motifs 4 |
| VPS4B | sp\|O75351\|VPS4B_HUMAN | O75351 | VPS4B | 9525 | Vacuolar protein sorting-associated protein 4B |
| FLNB | sp\|O75369\|FLNB_HUMAN | O75369 | FLNB | 2317 | Filamin-B |
| FLNA | sp\|P21333\|FLNA_HUMAN | P21333 | FLNA | 2316 | Filamin-A |
| LDHA | sp\|P00338\|LDHA_HUMAN | P00338 | LDHA | 3939 | L-lactate dehydrogenase A chain |
| LDHB | sp\|P07195\|LDHB_HUMAN | P07195 | LDHB | 3945 | L-lactate dehydrogenase B chain |
| CP | sp\|P00450\|CERU_HUMAN | P00450 | CERU | 1356 | Ceruloplasmin |
| EGFR | sp\|P00533\|EGFR_HUMAN | P00533 | EGFR | 1956 | Epidermal growth factor receptor |
| PGK1 | sp\|P00558\|PGK1_HUMAN | P00558 | PGK1 | 5230 | Phosphoglycerate kinase 1 |
| F2 | sp\|P00734\|THRB_HUMAN | P00734 | THRB | 2147 | Prothrombin |
| HP | sp\|P00738\|HPT_HUMAN | P00738 | HPT | 3240 | Haptoglobin |
| HPR | sp\|P00739\|HPTR_HUMAN | P00739 | HPTR | 3250 | Haptoglobin-related protein |
| PLG | sp\|P00747\|PLMN_HUMAN | P00747 | PLMN | 5340 | Plasminogen |
| LPA | sp\|P08519\|APOA_HUMAN | P08519 | APOA | 4018 | Apolipoprotein(a) |
| CFB | sp\|P00751\|CFAB_HUMAN | P00751 | CFAB | 629 | Complement factor B |
| SERPINC1 | sp\|P01008\|ANT3_HUMAN | P01008 | ANT3 | 462 | Antithrombin-III |
| A2M | sp\|P01023\|A2MG_HUMAN | P01023 | A2MG | 2 | Alpha-2-macroglobulin |
| PZP | sp\|P20742\|PZP_HUMAN | P20742 | PZP | 5858 | Pregnancy zone protein |
| C3 | sp\|P01024\|CO3_HUMAN | P01024 | CO3 | 718 | Complement C3 |
| C5 | sp\|P01031\|CO5_HUMAN | P01031 | CO5 | 727 | Complement C5 |
| KNG1 | sp\|P01042\|KNG1_HUMAN | P01042 | KNG1 | 3827 | Kininogen-1 |
| IGKV1-39 | sp\|P01597\|KV139_HUMAN | P01597 | KV139 | 28930 | Immunoglobulin kappa variable 1-39 |
| IGKV1-17 | sp\|P01599\|KV117_HUMAN | P01599 | KV117 | 28937 | Immunoglobulin kappa variable 1-17 |
| IGKV1D-16 | sp\|P01601\|KVD16_HUMAN | P01601 | KVD16 | 28901 | Immunoglobulin kappa variable 1D-16 |
| IGKV1-6 | sp\|A0A0C4DH72\|KV106_HUMAN | A0A0C4DH72 | KV106 | 28943 | Immunoglobulin kappa variable 1-6 |
| IGLV2-11 | sp\|P01706\|LV211_HUMAN | P01706 | LV211 | 28816 | Immunoglobulin lambda variable 2-11 |
| IGLV1-40 | sp\|P01703\|LV140_HUMAN | P01703 | LV140 | 28825 | Immunoglobulin lambda variable 1-40 |
| IGLV2-18 | sp\|A0A075B6J9\|LV218_HUMAN | A0A075B6J9 | LV218 | 28814 | Immunoglobulin lambda variable 2-18 |
| IGLV2-8 | sp\|P01709\|LV208_HUMAN | P01709 | LV208 | 28817 | Immunoglobulin lambda variable 2-8 |
| IGLV1-47 | sp\|P01700\|LV147_HUMAN | P01700 | LV147 | 28822 | Immunoglobulin lambda variable 1-47 |
| IGLV1-44 | sp\|P01699\|LV144_HUMAN | P01699 | LV144 | 28823 | Immunoglobulin lambda variable 1-44 |
| IGHV3-7 | sp\|P01780\|HV307_HUMAN | P01780 | HV307 | 28452 | Immunoglobulin heavy variable 3-7 |
| IGHV3-43D | sp\|P0DP04\|HV43D_HUMAN | P0DP04 | HV43D | 1.03E+08 | Immunoglobulin heavy variable 3-43D |
| IGHV3-23 | sp\|P01764\|HV323_HUMAN | P01764 | HV323 | 28442 | Immunoglobulin heavy variable 3-23 |
| IGHV3-74 | sp\|A0A0B4J1X5\|HV374_HUMAN | A0A0B4J1X5 | HV374 | 28408 | Immunoglobulin heavy variable 3-74 |
| IGHV3-43 | sp\|A0A0B4J1X8\|HV343_HUMAN | A0A0B4J1X8 | HV343 | 28426 | Immunoglobulin heavy variable 3-43 |
| IGHV3-13 | sp\|P01766\|HV313_HUMAN | P01766 | HV313 | 28449 | Immunoglobulin heavy variable 3-13 |
| IGHM | sp\|P01871\|IGHM_HUMAN | P01871 | IGHM | 3507 | Immunoglobulin heavy constant mu |
| HLA-DRB1 | sp\|P01911\|DRB1_HUMAN | P01911 | DRB1 | 3123 | HLA class II histocompatibility antigen, DRB1 beta chain |
| COL1A1 | sp\|P02452\|CO1A1_HUMAN | P02452 | CO1A1 | 1277 | Collagen alpha-1(I) chain |
| COL4A1 | sp\|P02462\|CO4A1_HUMAN | P02462 | CO4A1 | 1282 | Collagen alpha-1(IV) chain |
| LMNA | sp\|P02545\|LMNA_HUMAN | P02545 | LMNA | 4000 | Prelamin-A/C |
| APOA1 | sp\|P02647\|APOA1_HUMAN | P02647 | APOA1 | 335 | Apolipoprotein A-I |
| APOE | sp\|P02649\|APOE_HUMAN | P02649 | APOE | 348 | Apolipoprotein E |
| FGA | sp\|P02671\|FIBA_HUMAN | P02671 | FIBA | 2243 | Fibrinogen alpha chain |
| C9 | sp\|P02748\|CO9_HUMAN | P02748 | CO9 | 735 | Complement component C9 |
| APOH | sp\|P02749\|APOH_HUMAN | P02749 | APOH | 350 | Beta-2-glycoprotein 1 |
| FN1 | sp\|P02751\|FINC_HUMAN | P02751 | FINC | 2335 | Fibronectin |
| AHSG | sp\|P02765\|FETUA_HUMAN | P02765 | FETUA | 197 | Alpha-2-HS-glycoprotein |
| ALB | sp\|P02768\|ALBU_HUMAN | P02768 | ALBU | 213 | Albumin |
| GC | sp\|P02774\|VTDB_HUMAN | P02774 | VTDB | 2638 | Vitamin D-binding protein |
| HPX | sp\|P02790\|HEMO_HUMAN | P02790 | HEMO | 3263 | Hemopexin |
| C4BPA | sp\|P04003\|C4BPA_HUMAN | P04003 | C4BPA | 722 | C4b-binding protein alpha chain |
| VTN | sp\|P04004\|VTNC_HUMAN | P04004 | VTNC | 7448 | Vitronectin |
| ALDOA | sp\|P04075\|ALDOA_HUMAN | P04075 | ALDOA | 226 | Fructose-bisphosphate aldolase A |
| ALDOC | sp\|P09972\|ALDOC_HUMAN | P09972 | ALDOC | 230 | Fructose-bisphosphate aldolase C |
| SHBG | sp\|P04278\|SHBG_HUMAN | P04278 | SHBG | 6462 | Sex hormone-binding globulin |
| GAPDH | sp\|P04406\|G3P_HUMAN | P04406 | G3P | 2597 | Glyceraldehyde-3-phosphate dehydrogenase |
| HLA-A | sp\|P04439\|HLAA_HUMAN | P04439 | HLAA | 3105 | HLA class I histocompatibility antigen, A alpha chain |
| HLA-C | sp\|P10321\|HLAC_HUMAN | P10321 | HLAC | 3107 | HLA class I histocompatibility antigen, C alpha chain |
| GNAI2 | sp\|P04899\|GNAI2_HUMAN | P04899 | GNAI2 | 2771 | Guanine nucleotide-binding protein G(i) subunit alpha-2 |
| GNA13 | sp\|Q14344\|GNA13_HUMAN | Q14344 | GNA13 | 10672 | Guanine nucleotide-binding protein subunit alpha-13 |
| GNAS | sp\|P63092\|GNAS2_HUMAN | P63092 | GNAS2 | 2778 | Guanine nucleotide-binding protein G(s) subunit alpha isoforms short |
| GNAI3 | sp\|P08754\|GNAI3_HUMAN | P08754 | GNAI3 | 2773 | Guanine nucleotide-binding protein G(i) subunit alpha-3 |
| GNAI1 | sp\|P63096\|GNAI1_HUMAN | P63096 | GNAI1 | 2770 | Guanine nucleotide-binding protein G(i) subunit alpha-1 |
| ATP1A1 | sp\|P05023\|AT1A1_HUMAN | P05023 | AT1A1 | 476 | Sodium/potassium-transporting ATPase subunit alpha-1 |
| APOD | sp\|P05090\|APOD_HUMAN | P05090 | APOD | 347 | Apolipoprotein D |
| SLC25A5 | sp\|P05141\|ADT2_HUMAN | P05141 | ADT2 | 292 | ADP/ATP translocase 2 |
| CFI | sp\|P05156\|CFAI_HUMAN | P05156 | CFAI | 3426 | Complement factor I |
| RPLP0 | sp\|P05388\|RLA0_HUMAN | P05388 | RLA0 | 6175 | 60S acidic ribosomal protein P0 |
| SERPINA7 | sp\|P05543\|THBG_HUMAN | P05543 | THBG | 6906 | Thyroxine-binding globulin |
| SERPIND1 | sp\|P05546\|HEP2_HUMAN | P05546 | HEP2 | 3053 | Heparin cofactor 2 |
| ITGB1 | sp\|P05556\|ITB1_HUMAN | P05556 | ITB1 | 3688 | Integrin beta-1 |
| IGKV2-30 | sp\|P06310\|KV230_HUMAN | P06310 | KV230 | 28919 | Immunoglobulin kappa variable 2-30 |
| IGKV2-28 | sp\|A0A075B6P5\|KV228_HUMAN | A0A075B6P5 | KV228 | 28921 | Immunoglobulin kappa variable 2-28 |
| IGKV2D-29 | sp\|A0A075B6S2\|KVD29_HUMAN | A0A075B6S2 | KVD29 | 28882 | Immunoglobulin kappa variable 2D-29 |
| GSN | sp\|P06396\|GELS_HUMAN | P06396 | GELS | 2934 | Gelsolin |
| CEACAM5 | sp\|P06731\|CEAM5_HUMAN | P06731 | CEAM5 | 1048 | Carcinoembryonic antigen-related cell adhesion molecule 5 |
| CEACAM1 | sp\|P13688\|CEAM1_HUMAN | P13688 | CEAM1 | 634 | Carcinoembryonic antigen-related cell adhesion molecule 1 |
| ENO1 | sp\|P06733\|ENOA_HUMAN | P06733 | ENOA | 2023 | Alpha-enolase |
| GPI | sp\|P06744\|G6PI_HUMAN | P06744 | G6PI | 2821 | Glucose-6-phosphate isomerase |
| LPL | sp\|P06858\|LIPL_HUMAN | P06858 | LIPL | 4023 | Lipoprotein lipase |
| PROS1 | sp\|P07225\|PROS_HUMAN | P07225 | PROS | 5627 | Vitamin K-dependent protein S |
| ANXA2 | sp\|P07355\|ANXA2_HUMAN | P07355 | ANXA2 | 302 | Annexin A2 |
| TUBB | sp\|P07437\|TBB5_HUMAN | P07437 | TBB5 | 203068 | Tubulin beta chain |
| TUBB4B | sp\|P68371\|TBB4B_HUMAN | P68371 | TBB4B | 10383 | Tubulin beta-4B chain |
| TUBB4A | sp\|P04350\|TBB4A_HUMAN | P04350 | TBB4A | 10382 | Tubulin beta-4A chain |
| PFN1 | sp\|P07737\|PROF1_HUMAN | P07737 | PROF1 | 5216 | Profilin-1 |
| IGF1R | sp\|P08069\|IGF1R_HUMAN | P08069 | IGF1R | 3480 | Insulin-like growth factor 1 receptor |
| INSR | sp\|P06213\|INSR_HUMAN | P06213 | INSR | 3643 | Insulin receptor |
| ANXA6 | sp\|P08133\|ANXA6_HUMAN | P08133 | ANXA6 | 309 | Annexin A6 |
| ANXA11 | sp\|P50995\|ANX11_HUMAN | P50995 | ANX11 | 311 | Annexin A11 |
| ANXA4 | sp\|P09525\|ANXA4_HUMAN | P09525 | ANXA4 | 307 | Annexin A4 |
| HSP90AB1 | sp\|P08238\|HS90B_HUMAN | P08238 | HS90B | 3326 | Heat shock protein HSP 90-beta |
| HSP90AA1 | sp\|P07900\|HS90A_HUMAN | P07900 | HS90A | 3320 | Heat shock protein HSP 90-alpha |
| GSTA1 | sp\|P08263\|GSTA1_HUMAN | P08263 | GSTA1 | 2938 | Glutathione S-transferase A1 |
| CFH | sp\|P08603\|CFAH_HUMAN | P08603 | CFAH | 3075 | Complement factor H |
| SERPINF2 | sp\|P08697\|A2AP_HUMAN | P08697 | A2AP | 5345 | Alpha-2-antiplasmin |
| GSTP1 | sp\|P09211\|GSTP1_HUMAN | P09211 | GSTP1 | 2950 | Glutathione S-transferase P |
| C1S | sp\|P09871\|C1S_HUMAN | P09871 | C1S | 716 | Complement C1s subcomponent |
| C4B_2; C4B | sp\|P0C0L5\|CO4B_HUMAN | P0C0L5 | CO4B | 100293534; 721 | Complement C4-B |
| C4A | sp\|P0C0L4\|CO4A_HUMAN | P0C0L4 | CO4A | 721; 720 | Complement C4-A |
| H2AC11; H2AC13; H2AC16; H2AC17; H2AC15 | sp\|P0C0S8\|H2A1_HUMAN | P0C0S8 | H2A1 | 8336; 8330; 8969; 8329; 8332 | Histone H2A type 1 |
| H2AZ1 | sp\|P0C0S5\|H2AZ_HUMAN | P0C0S5 | H2AZ | 3015 | Histone H2A.Z |
| IGLC3 | sp\|P0DOY3\|IGLC3_HUMAN | P0DOY3 | IGLC3 | 3539 | Immunoglobulin lambda constant 3 |
| IGLL5 | sp\|B9A064\|IGLL5_HUMAN | B9A064 | IGLL5 | 1E+08 | Immunoglobulin lambda-like polypeptide 5 |
| IGHV4-38-2 | sp\|P0DP08\|HVD82_HUMAN | P0DP08 | HVD82 | 1.03E+08 | Immunoglobulin heavy variable 4-38-2 |
| IGHV4-4 | sp\|A0A075B6R2\|HV404_HUMAN | A0A075B6R2 | HV404 | 1.03E+08 | Immunoglobulin heavy variable 4-4 |
| AMY1B | sp\|P0DTE7\|AMY1B_HUMAN | P0DTE7 | AMY1B | 276; 277; 278 | Alpha-amylase 1B |
| H1-4 | sp\|P10412\|H14_HUMAN | P10412 | H14 | 3008 | Histone H1.4 |
| C7 | sp\|P10643\|CO7_HUMAN | P10643 | CO7 | 730 | Complement component C7 |
| CLU | sp\|P10909\|CLUS_HUMAN | P10909 | CLUS | 1191 | Clusterin |
| HSPA5 | sp\|P11021\|BIP_HUMAN | P11021 | BIP | 3309 | Endoplasmic reticulum chaperone BiP |
| HSPA8 | sp\|P11142\|HSP7C_HUMAN | P11142 | HSP7C | 3312 | Heat shock cognate 71 kDa protein |
| HSPA1A | sp\|P0DMV8\|HS71A_HUMAN | P0DMV8 | HS71A | 3304; 3303 | Heat shock 70 kDa protein 1A |
| IGF2R | sp\|P11717\|MPRI_HUMAN | P11717 | MPRI | 3482 | Cation-independent mannose-6-phosphate receptor |
| PABPC1 | sp\|P11940\|PABP1_HUMAN | P11940 | PABP1 | 26986 | Polyadenylate-binding protein 1 |
| F5 | sp\|P12259\|FA5_HUMAN | P12259 | FA5 | 2153 | Coagulation factor V |
| YES1 | sp\|P07947\|YES_HUMAN | P07947 | YES | 7525 | Tyrosine-protein kinase Yes |
| EEF2 | sp\|P13639\|EF2_HUMAN | P13639 | EF2 | 1938 | Elongation factor 2 |
| TPT1 | sp\|P13693\|TCTP_HUMAN | P13693 | TCTP | 7178 | Translationally-controlled tumor protein |
| SELL | sp\|P14151\|LYAM1_HUMAN | P14151 | LYAM1 | 6402 | L-selectin |
| PKM | sp\|P14618\|KPYM_HUMAN | P14618 | KPYM | 5315 | Pyruvate kinase PKM |
| HSP90B1 | sp\|P14625\|ENPL_HUMAN | P14625 | ENPL | 7184 | Endoplasmin |
| JUP | sp\|P14923\|PLAK_HUMAN | P14923 | PLAK | 3728 | Junction plakoglobin |
| CPN1 | sp\|P15169\|CBPN_HUMAN | P15169 | CBPN | 1369 | Carboxypeptidase N catalytic chain |
| B4GALT1 | sp\|P15291\|B4GT1_HUMAN | P15291 | B4GT1 | 2683 | Beta-1,4-galactosyltransferase 1 |
| CBR1 | sp\|P16152\|CBR1_HUMAN | P16152 | CBR1 | 873 | Carbonyl reductase [NADPH] 1 |
| ATP2A2 | sp\|P16615\|AT2A2_HUMAN | P16615 | AT2A2 | 488 | Sarcoplasmic/endoplasmic reticulum calcium ATPase 2 |
| PGAM1 | sp\|P18669\|PGAM1_HUMAN | P18669 | PGAM1 | 5223 | Phosphoglycerate mutase 1 |
| ITIH2 | sp\|P19823\|ITIH2_HUMAN | P19823 | ITIH2 | 3698 | Inter-alpha-trypsin inhibitor heavy chain H2 |
| ITIH1 | sp\|P19827\|ITIH1_HUMAN | P19827 | ITIH1 | 3697 | Inter-alpha-trypsin inhibitor heavy chain H1 |
| TNXB | sp\|P22105\|TENX_HUMAN | P22105 | TENX | 7148 | Tenascin-X |
| NME2 | sp\|P22392\|NDKB_HUMAN | P22392 | NDKB | 4831; 654364 | Nucleoside diphosphate kinase B |
| NME1 | sp\|P15531\|NDKA_HUMAN | P15531 | NDKA | 4830 | Nucleoside diphosphate kinase A |
| SPRR2E | sp\|P22531\|SPR2E_HUMAN | P22531 | SPR2E | 6704 | Small proline-rich protein 2E |
| FDXR | sp\|P22570\|ADRO_HUMAN | P22570 | ADRO | 2232 | NADPH:adrenodoxin oxidoreductase, mitochondrial |
| FBLN1 | sp\|P23142\|FBLN1_HUMAN | P23142 | FBLN1 | 2192 | Fibulin-1 |
| PPIB | sp\|P23284\|PPIB_HUMAN | P23284 | PPIB | 5479 | Peptidyl-prolyl cis-trans isomerase B |
| ATP2B4 | sp\|P23634\|AT2B4_HUMAN | P23634 | AT2B4 | 493 | Plasma membrane calcium-transporting ATPase 4 |
| ATP2B1 | sp\|P20020\|AT2B1_HUMAN | P20020 | AT2B1 | 490 | Plasma membrane calcium-transporting ATPase 1 |
| ATP5F1A | sp\|P25705\|ATPA_HUMAN | P25705 | ATPA | 498 | ATP synthase subunit alpha, mitochondrial |
| HSD3B2 | sp\|P26439\|3BHS2_HUMAN | P26439 | 3BHS2 | 3284 | 3 beta-hydroxysteroid dehydrogenase/Delta 5--4-isomerase type 2 |
| MST1 | sp\|P26927\|HGFL_HUMAN | P26927 | HGFL | 4485 | Hepatocyte growth factor-like protein |
| STOM | sp\|P27105\|STOM_HUMAN | P27105 | STOM | 2040 | Stomatin |
| PON1 | sp\|P27169\|PON1_HUMAN | P27169 | PON1 | 5444 | Serum paraoxonase/arylesterase 1 |
| DPP4 | sp\|P27487\|DPP4_HUMAN | P27487 | DPP4 | 1803 | Dipeptidyl peptidase 4 |
| RPL10 | sp\|P27635\|RL10_HUMAN | P27635 | RL10 | 6134 | 60S ribosomal protein L10 |
| CANX | sp\|P27824\|CALX_HUMAN | P27824 | CALX | 821 | Calnexin |
| LCN1 | sp\|P31025\|LCN1_HUMAN | P31025 | LCN1 | 3933 | Lipocalin-1 |
| S100A7 | sp\|P31151\|S10A7_HUMAN | P31151 | S10A7 | 6278 | Protein S100-A7 |
| HNRNPH1 | sp\|P31943\|HNRH1_HUMAN | P31943 | HNRH1 | 3187 | Heterogeneous nuclear ribonucleoprotein H |
| KIF5B | sp\|P33176\|KINH_HUMAN | P33176 | KINH | 3799 | Kinesin-1 heavy chain |
| CTNNA1 | sp\|P35221\|CTNA1_HUMAN | P35221 | CTNA1 | 1495 | Catenin alpha-1 |
| RDX | sp\|P35241\|RADI_HUMAN | P35241 | RADI | 5962 | Radixin |
| EZR | sp\|P15311\|EZRI_HUMAN | P15311 | EZRI | 7430 | Ezrin |
| MSN | sp\|P26038\|MOES_HUMAN | P26038 | MOES | 4478 | Moesin |
| THBS4 | sp\|P35443\|TSP4_HUMAN | P35443 | TSP4 | 7060 | Thrombospondin-4 |
| KRT9 | sp\|P35527\|K1C9_HUMAN | P35527 | K1C9 | 3857 | Keratin, type I cytoskeletal 9 |
| KRT10 | sp\|P13645\|K1C10_HUMAN | P13645 | K1C10 | 3858 | Keratin, type I cytoskeletal 10 |
| KRT14 | sp\|P02533\|K1C14_HUMAN | P02533 | K1C14 | 3861 | Keratin, type I cytoskeletal 14 |
| KRT16 | sp\|P08779\|K1C16_HUMAN | P08779 | K1C16 | 3868 | Keratin, type I cytoskeletal 16 |
| KRT13 | sp\|P13646\|K1C13_HUMAN | P13646 | K1C13 | 3860 | Keratin, type I cytoskeletal 13 |
| KRT17 | sp\|Q04695\|K1C17_HUMAN | Q04695 | K1C17 | 3872 | Keratin, type I cytoskeletal 17 |
| KRT31 | sp\|Q15323\|K1H1_HUMAN | Q15323 | K1H1 | 3881 | Keratin, type I cuticular Ha1 |
| KRT33B | sp\|Q14525\|KT33B_HUMAN | Q14525 | KT33B | 3884 | Keratin, type I cuticular Ha3-II |
| KRT19 | sp\|P08727\|K1C19_HUMAN | P08727 | K1C19 | 3880 | Keratin, type I cytoskeletal 19 |
| KRT34 | sp\|O76011\|KRT34_HUMAN | O76011 | KRT34 | 100653049; 3885 | Keratin, type I cuticular Ha4 |
| KRT36 | sp\|O76013\|KRT36_HUMAN | O76013 | KRT36 | 8689 | Keratin, type I cuticular Ha6 |
| KRT32 | sp\|Q14532\|K1H2_HUMAN | Q14532 | K1H2 | 3882 | Keratin, type I cuticular Ha2 |
| KRT23 | sp\|Q9C075\|K1C23_HUMAN | Q9C075 | K1C23 | 25984 | Keratin, type I cytoskeletal 23 |
| KRT2 | sp\|P35908\|K22E_HUMAN | P35908 | K22E | 3849 | Keratin, type II cytoskeletal 2 epidermal |
| KRT6C | sp\|P48668\|K2C6C_HUMAN | P48668 | K2C6C | 286887 | Keratin, type II cytoskeletal 6C |
| KRT76 | sp\|Q01546\|K22O_HUMAN | Q01546 | K22O | 51350 | Keratin, type II cytoskeletal 2 oral |
| KRT1 | sp\|P04264\|K2C1_HUMAN | P04264 | K2C1 | 3848 | Keratin, type II cytoskeletal 1 |
| KRT5 | sp\|P13647\|K2C5_HUMAN | P13647 | K2C5 | 3852 | Keratin, type II cytoskeletal 5 |
| KRT6A | sp\|P02538\|K2C6A_HUMAN | P02538 | K2C6A | 3853 | Keratin, type II cytoskeletal 6A |
| KRT6B | sp\|P04259\|K2C6B_HUMAN | P04259 | K2C6B | 3854 | Keratin, type II cytoskeletal 6B |
| KRT4 | sp\|P19013\|K2C4_HUMAN | P19013 | K2C4 | 3851 | Keratin, type II cytoskeletal 4 |
| KRT3 | sp\|P12035\|K2C3_HUMAN | P12035 | K2C3 | 3850 | Keratin, type II cytoskeletal 3 |
| KRT84 | sp\|Q9NSB2\|KRT84_HUMAN | Q9NSB2 | KRT84 | 3890 | Keratin, type II cuticular Hb4 |
| KRT85 | sp\|P78386\|KRT85_HUMAN | P78386 | KRT85 | 3891 | Keratin, type II cuticular Hb5 |
| KRT86 | sp\|O43790\|KRT86_HUMAN | O43790 | KRT86 | 3892 | Keratin, type II cuticular Hb6 |
| KRT83 | sp\|P78385\|KRT83_HUMAN | P78385 | KRT83 | 3889 | Keratin, type II cuticular Hb3 |
| KRT77 | sp\|Q7Z794\|K2C1B_HUMAN | Q7Z794 | K2C1B | 374454 | Keratin, type II cytoskeletal 1b |
| KRT78 | sp\|Q8N1N4\|K2C78_HUMAN | Q8N1N4 | K2C78 | 196374 | Keratin, type II cytoskeletal 78 |
| KRT79 | sp\|Q5XKE5\|K2C79_HUMAN | Q5XKE5 | K2C79 | 338785 | Keratin, type II cytoskeletal 79 |
| KRT80 | sp\|Q6KB66\|K2C80_HUMAN | Q6KB66 | K2C80 | 144501 | Keratin, type II cytoskeletal 80 |
| ARL3 | sp\|P36405\|ARL3_HUMAN | P36405 | ARL3 | 403 | ADP-ribosylation factor-like protein 3 |
| SERPINF1 | sp\|P36955\|PEDF_HUMAN | P36955 | PEDF | 5176 | Pigment epithelium-derived factor |
| CCT6A | sp\|P40227\|TCPZ_HUMAN | P40227 | TCPZ | 908 | T-complex protein 1 subunit zeta |
| RPL13A | sp\|P40429\|RL13A_HUMAN | P40429 | RL13A | 23521 | 60S ribosomal protein L13a |
| AFM | sp\|P43652\|AFAM_HUMAN | P43652 | AFAM | 173 | Afamin |
| IQGAP1 | sp\|P46940\|IQGA1_HUMAN | P46940 | IQGA1 | 8826 | Ras GTPase-activating-like protein IQGAP1 |
| NPTX2 | sp\|P47972\|NPTX2_HUMAN | P47972 | NPTX2 | 4885 | Neuronal pentraxin-2 |
| TUFM | sp\|P49411\|EFTU_HUMAN | P49411 | EFTU | 7284 | Elongation factor Tu, mitochondrial |
| GNAQ | sp\|P50148\|GNAQ_HUMAN | P50148 | GNAQ | 2776 | Guanine nucleotide-binding protein G(q) subunit alpha |
| GNA11 | sp\|P29992\|GNA11_HUMAN | P29992 | GNA11 | 2767 | Guanine nucleotide-binding protein subunit alpha-11 |
| ST13 | sp\|P50502\|F10A1_HUMAN | P50502 | F10A1 | 6767 | Hsc70-interacting protein |
| DNM2 | sp\|P50570\|DYN2_HUMAN | P50570 | DYN2 | 1785 | Dynamin-2 |
| ALDH3A2 | sp\|P51648\|AL3A2_HUMAN | P51648 | AL3A2 | 224 | Aldehyde dehydrogenase family 3 member A2 |
| LUM | sp\|P51884\|LUM_HUMAN | P51884 | LUM | 4060 | Lumican |
| PGD | sp\|P52209\|6PGD_HUMAN | P52209 | 6PGD | 5226 | 6-phosphogluconate dehydrogenase, decarboxylating |
| NAPA | sp\|P54920\|SNAA_HUMAN | P54920 | SNAA | 8775 | Alpha-soluble NSF attachment protein |
| VCP | sp\|P55072\|TERA_HUMAN | P55072 | TERA | 7415 | Transitional endoplasmic reticulum ATPase |
| NPEPPS | sp\|P55786\|PSA_HUMAN | P55786 | PSA | 9520 | Puromycin-sensitive aminopeptidase |
| MYL6 | sp\|P60660\|MYL6_HUMAN | P60660 | MYL6 | 4637 | Myosin light polypeptide 6 |
| EIF4A1 | sp\|P60842\|IF4A1_HUMAN | P60842 | IF4A1 | 1973 | Eukaryotic initiation factor 4A-I |
| EIF4A2 | sp\|Q14240\|IF4A2_HUMAN | Q14240 | IF4A2 | 1974 | Eukaryotic initiation factor 4A-II |
| CDC42 | sp\|P60953\|CDC42_HUMAN | P60953 | CDC42 | 998 | Cell division control protein 42 homolog |
| RAB14 | sp\|P61106\|RAB14_HUMAN | P61106 | RAB14 | 51552 | Ras-related protein Rab-14 |
| RAB1B | sp\|Q9H0U4\|RAB1B_HUMAN | Q9H0U4 | RAB1B | 81876 | Ras-related protein Rab-1B |
| RAB6A | sp\|P20340\|RAB6A_HUMAN | P20340 | RAB6A | 5870 | Ras-related protein Rab-6A |
| RAB1A | sp\|P62820\|RAB1A_HUMAN | P62820 | RAB1A | 5861 | Ras-related protein Rab-1A |
| RAB8A | sp\|P61006\|RAB8A_HUMAN | P61006 | RAB8A | 4218 | Ras-related protein Rab-8A |
| RAB35 | sp\|Q15286\|RAB35_HUMAN | Q15286 | RAB35 | 11021 | Ras-related protein Rab-35 |
| RAB34 | sp\|Q9BZG1\|RAB34_HUMAN | Q9BZG1 | RAB34 | 83871 | Ras-related protein Rab-34 |
| RAB4B | sp\|P61018\|RAB4B_HUMAN | P61018 | RAB4B | 53916 | Ras-related protein Rab-4B |
| RAB43 | sp\|Q86YS6\|RAB43_HUMAN | Q86YS6 | RAB43 | 339122 | Ras-related protein Rab-43 |
| RAB10 | sp\|P61026\|RAB10_HUMAN | P61026 | RAB10 | 10890 | Ras-related protein Rab-10 |
| RAB13 | sp\|P51153\|RAB13_HUMAN | P51153 | RAB13 | 5872 | Ras-related protein Rab-13 |
| ACTR3 | sp\|P61158\|ARP3_HUMAN | P61158 | ARP3 | 10096 | Actin-related protein 3 |
| RAP1B | sp\|P61224\|RAP1B_HUMAN | P61224 | RAP1B | 5908 | Ras-related protein Rap-1b |
| RAP1A | sp\|P62834\|RAP1A_HUMAN | P62834 | RAP1A | 5906 | Ras-related protein Rap-1A |
| NRAS | sp\|P01111\|RASN_HUMAN | P01111 | RASN | 4893 | GTPase NRas |
| RAP2B | sp\|P61225\|RAP2B_HUMAN | P61225 | RAP2B | 5912 | Ras-related protein Rap-2b |
| SEC61A1 | sp\|P61619\|S61A1_HUMAN | P61619 | S61A1 | 29927 | Protein transport protein Sec61 subunit alpha isoform 1 |
| LYZ | sp\|P61626\|LYSC_HUMAN | P61626 | LYSC | 4069 | Lysozyme C |
| RRAS2 | sp\|P62070\|RRAS2_HUMAN | P62070 | RRAS2 | 22800 | Ras-related protein R-Ras2 |
| ARF6 | sp\|P62330\|ARF6_HUMAN | P62330 | ARF6 | 382 | ADP-ribosylation factor 6 |
| RPS4X | sp\|P62701\|RS4X_HUMAN | P62701 | RS4X | 6191 | 40S ribosomal protein S4, X isoform |
| GNB1 | sp\|P62873\|GBB1_HUMAN | P62873 | GBB1 | 2782 | Guanine nucleotide-binding protein G(I)/G(S)/G(T) subunit beta-1 |
| GNB2 | sp\|P62879\|GBB2_HUMAN | P62879 | GBB2 | 2783 | Guanine nucleotide-binding protein G(I)/G(S)/G(T) subunit beta-2 |
| PPIA | sp\|P62937\|PPIA_HUMAN | P62937 | PPIA | 5478 | Peptidyl-prolyl cis-trans isomerase A |
| RAC1 | sp\|P63000\|RAC1_HUMAN | P63000 | RAC1 | 5879 | Ras-related C3 botulinum toxin substrate 1 |
| VAMP2 | sp\|P63027\|VAMP2_HUMAN | P63027 | VAMP2 | 6844 | Vesicle-associated membrane protein 2 |
| VAMP3 | sp\|Q15836\|VAMP3_HUMAN | Q15836 | VAMP3 | 9341 | Vesicle-associated membrane protein 3 |
| DYNLL1 | sp\|P63167\|DYL1_HUMAN | P63167 | DYL1 | 8655 | Dynein light chain 1, cytoplasmic |
| EIF5A | sp\|P63241\|IF5A1_HUMAN | P63241 | IF5A1 | 1984 | Eukaryotic translation initiation factor 5A-1 |
| ACTG1 | sp\|P63261\|ACTG_HUMAN | P63261 | ACTG | 71 | Actin, cytoplasmic 2 |
| ACTB | sp\|P60709\|ACTB_HUMAN | P60709 | ACTB | 60 | Actin, cytoplasmic 1 |
| ACTC1 | sp\|P68032\|ACTC_HUMAN | P68032 | ACTC | 70 | Actin, alpha cardiac muscle 1 |
| ACTBL2 | sp\|Q562R1\|ACTBL_HUMAN | Q562R1 | ACTBL | 345651 | Beta-actin-like protein 2 |
| SEC11A | sp\|P67812\|SC11A_HUMAN | P67812 | SC11A | 23478 | Signal peptidase complex catalytic subunit SEC11A |
| TPM4 | sp\|P67936\|TPM4_HUMAN | P67936 | TPM4 | 7171 | Tropomyosin alpha-4 chain |
| TPM3 | sp\|P06753\|TPM3_HUMAN | P06753 | TPM3 | 7170 | Tropomyosin alpha-3 chain |
| EEF1A1 | sp\|P68104\|EF1A1_HUMAN | P68104 | EF1A1 | 1915 | Elongation factor 1-alpha 1 |
| TUBA1B | sp\|P68363\|TBA1B_HUMAN | P68363 | TBA1B | 10376 | Tubulin alpha-1B chain |
| H3C3; H3C7; H3C8; H3C11; H3C4; H3C12; H3C1; H3C2; H3C6; H3C10 | sp\|P68431\|H31_HUMAN | P68431 | H31 | 8350; 8353; 8354; 8357; 8356; 8351; 8355; 8968; 8352; 8358 | Histone H3.1 |
| HBB | sp\|P68871\|HBB_HUMAN | P68871 | HBB | 3043 | Hemoglobin subunit beta |
| NOMO3 | sp\|P69849\|NOMO3_HUMAN | P69849 | NOMO3 | 408050 | Nodal modulator 3 |
| HBA1; HBA2 | sp\|P69905\|HBA_HUMAN | P69905 | HBA | 3039; 3040 | Hemoglobin subunit alpha |
| IGLV3-21 | sp\|P80748\|LV321_HUMAN | P80748 | LV321 | 28796 | Immunoglobulin lambda variable 3-21 |
| IGLV3-10 | sp\|A0A075B6K4\|LV310_HUMAN | A0A075B6K4 | LV310 | 28803 | Immunoglobulin lambda variable 3-10 |
| IGLV3-27 | sp\|P01718\|LV327_HUMAN | P01718 | LV327 | 28791 | Immunoglobulin lambda variable 3-27 |
| HTRA3 | sp\|P83110\|HTRA3_HUMAN | P83110 | HTRA3 | 94031 | Serine protease HTRA3 |
| HTRA1 | sp\|Q92743\|HTRA1_HUMAN | Q92743 | HTRA1 | 5654 | Serine protease HTRA1 |
| HSPG2 | sp\|P98160\|PGBM_HUMAN | P98160 | PGBM | 3339 | Basement membrane-specific heparan sulfate proteoglycan core protein |
| EFNB1 | sp\|P98172\|EFNB1_HUMAN | P98172 | EFNB1 | 1947 | Ephrin-B1 |
| CLTC | sp\|Q00610\|CLH1_HUMAN | Q00610 | CLH1 | 1213 | Clathrin heavy chain 1 |
| SPTBN1 | sp\|Q01082\|SPTB2_HUMAN | Q01082 | SPTB2 | 6711 | Spectrin beta chain, non-erythrocytic 1 |
| FABP5 | sp\|Q01469\|FABP5_HUMAN | Q01469 | FABP5 | 2171 | Fatty acid-binding protein 5 |
| HMGCS1 | sp\|Q01581\|HMCS1_HUMAN | Q01581 | HMCS1 | 3157 | Hydroxymethylglutaryl-CoA synthase, cytoplasmic |
| CFHR1 | sp\|Q03591\|FHR1_HUMAN | Q03591 | FHR1 | 3078 | Complement factor H-related protein 1 |
| CFHR2 | sp\|P36980\|FHR2_HUMAN | P36980 | FHR2 | 3080 | Complement factor H-related protein 2 |
| CKAP4 | sp\|Q07065\|CKAP4_HUMAN | Q07065 | CKAP4 | 10970 | Cytoskeleton-associated protein 4 |
| CD47 | sp\|Q08722\|CD47_HUMAN | Q08722 | CD47 | 961 | Leukocyte surface antigen CD47 |
| AHNAK | sp\|Q09666\|AHNK_HUMAN | Q09666 | AHNK | 79026 | Neuroblast differentiation-associated protein AHNAK |
| MYH7 | sp\|P12883\|MYH7_HUMAN | P12883 | MYH7 | 4625 | Myosin-7 |
| GALNT2 | sp\|Q10471\|GALT2_HUMAN | Q10471 | GALT2 | 2590 | Polypeptide N-acetylgalactosaminyltransferase 2 |
| ITGA9 | sp\|Q13797\|ITA9_HUMAN | Q13797 | ITA9 | 3680 | Integrin alpha-9 |
| PKP1 | sp\|Q13835\|PKP1_HUMAN | Q13835 | PKP1 | 5317 | Plakophilin-1 |
| DPYSL3 | sp\|Q14195\|DPYL3_HUMAN | Q14195 | DPYL3 | 1809 | Dihydropyrimidinase-related protein 3 |
| FAT1 | sp\|Q14517\|FAT1_HUMAN | Q14517 | FAT1 | 2195 | Protocadherin Fat 1 |
| ITIH4 | sp\|Q14624\|ITIH4_HUMAN | Q14624 | ITIH4 | 3700 | Inter-alpha-trypsin inhibitor heavy chain H4 |
| SPCS2 | sp\|Q15005\|SPCS2_HUMAN | Q15005 | SPCS2 | 9789 | Signal peptidase complex subunit 2 |
| POSTN | sp\|Q15063\|POSTN_HUMAN | Q15063 | POSTN | 10631 | Periostin |
| PPA1 | sp\|Q15181\|IPYR_HUMAN | Q15181 | IPYR | 5464 | Inorganic pyrophosphatase |
| PCBP1 | sp\|Q15365\|PCBP1_HUMAN | Q15365 | PCBP1 | 5093 | Poly(rC)-binding protein 1 |
| PCBP2 | sp\|Q15366\|PCBP2_HUMAN | Q15366 | PCBP2 | 5094 | Poly(rC)-binding protein 2 |
| AOC3 | sp\|Q16853\|AOC3_HUMAN | Q16853 | AOC3 | 8639 | Membrane primary amine oxidase |
| CHSY3 | sp\|Q70JA7\|CHSS3_HUMAN | Q70JA7 | CHSS3 | 337876 | Chondroitin sulfate synthase 3 |
| BTN2A1 | sp\|Q7KYR7\|BT2A1_HUMAN | Q7KYR7 | BT2A1 | 11120 | Butyrophilin subfamily 2 member A1 |
| HUWE1 | sp\|Q7Z6Z7\|HUWE1_HUMAN | Q7Z6Z7 | HUWE1 | 10075 | E3 ubiquitin-protein ligase HUWE1 |
| SCUBE1 | sp\|Q8IWY4\|SCUB1_HUMAN | Q8IWY4 | SCUB1 | 80274 | Signal peptide, CUB and EGF-like domain-containing protein 1 |
| DPP10 | sp\|Q8N608\|DPP10_HUMAN | Q8N608 | DPP10 | 57628 | Inactive dipeptidyl peptidase 10 |
| ENAH | sp\|Q8N8S7\|ENAH_HUMAN | Q8N8S7 | ENAH | 55740 | Protein enabled homolog |
| EVPL | sp\|Q92817\|EVPL_HUMAN | Q92817 | EVPL | 2125 | Envoplakin |
| CPNE2 | sp\|Q96FN4\|CPNE2_HUMAN | Q96FN4 | CPNE2 | 221184 | Copine-2 |
| CPNE3 | sp\|O75131\|CPNE3_HUMAN | O75131 | CPNE3 | 8895 | Copine-3 |
| CPB2 | sp\|Q96IY4\|CBPB2_HUMAN | Q96IY4 | CBPB2 | 1361 | Carboxypeptidase B2 |
| SERPINB12 | sp\|Q96P63\|SPB12_HUMAN | Q96P63 | SPB12 | 89777 | Serpin B12 |
| PGLYRP2 | sp\|Q96PD5\|PGRP2_HUMAN | Q96PD5 | PGRP2 | 114770 | N-acetylmuramoyl-L-alanine amidase |
| HAPLN3 | sp\|Q96S86\|HPLN3_HUMAN | Q96S86 | HPLN3 | 145864 | Hyaluronan and proteoglycan link protein 3 |
| HAPLN1 | sp\|P10915\|HPLN1_HUMAN | P10915 | HPLN1 | 1404 | Hyaluronan and proteoglycan link protein 1 |
| SH3GL1 | sp\|Q99961\|SH3G1_HUMAN | Q99961 | SH3G1 | 6455 | Endophilin-A2 |
| ESYT1 | sp\|Q9BSJ8\|ESYT1_HUMAN | Q9BSJ8 | ESYT1 | 23344 | Extended synaptotagmin-1 |
| CRISPLD2 | sp\|Q9H0B8\|CRLD2_HUMAN | Q9H0B8 | CRLD2 | 83716 | Cysteine-rich secretory protein LCCL domain-containing 2 |
| EHD1 | sp\|Q9H4M9\|EHD1_HUMAN | Q9H4M9 | EHD1 | 10938 | EH domain-containing protein 1 |
| ANTXR1 | sp\|Q9H6X2\|ANTR1_HUMAN | Q9H6X2 | ANTR1 | 84168 | Anthrax toxin receptor 1 |
| LIN7C | sp\|Q9NUP9\|LIN7C_HUMAN | Q9NUP9 | LIN7C | 55327 | Protein lin-7 homolog C |
| TMEM30A | sp\|Q9NV96\|CC50A_HUMAN | Q9NV96 | CC50A | 55754 | Cell cycle control protein 50A |
| SEPTIN11 | sp\|Q9NVA2\|SEP11_HUMAN | Q9NVA2 | 40787 | 55752 | Septin-11 |
| UGGT1 | sp\|Q9NYU2\|UGGG1_HUMAN | Q9NYU2 | UGGG1 | 56886 | UDP-glucose:glycoprotein glucosyltransferase 1 |
| RRBP1 | sp\|Q9P2E9\|RRBP1_HUMAN | Q9P2E9 | RRBP1 | 6238 | Ribosome-binding protein 1 |
| PLXNA1 | sp\|Q9UIW2\|PLXA1_HUMAN | Q9UIW2 | PLXA1 | 5361 | Plexin-A1 |
| GPC6 | sp\|Q9Y625\|GPC6_HUMAN | Q9Y625 | GPC6 | 10082 | Glypican-6 |
| PRSS1 | sp\|P07477\|TRY1_HUMAN | P07477 | TRY1 | 5644 | Serine protease 1 |
| LTF | sp\|P02788\|TRFL_HUMAN | P02788 | TRFL | 4057 | Lactotransferrin |
| IGLV9-49 | sp\|A0A0B4J1Y8\|LV949_HUMAN | A0A0B4J1Y8 | LV949 | 28773 | Immunoglobulin lambda variable 9-49 |
| STX7 | sp\|O15400\|STX7_HUMAN | O15400 | STX7 | 8417 | Syntaxin-7 |
| DNAJA2 | sp\|O60884\|DNJA2_HUMAN | O60884 | DNJA2 | 10294 | DnaJ homolog subfamily A member 2 |
| DDAH1 | sp\|O94760\|DDAH1_HUMAN | O94760 | DDAH1 | 23576 | N(G),N(G)-dimethylarginine dimethylaminohydrolase 1 |
| ABCA1 | sp\|O95477\|ABCA1_HUMAN | O95477 | ABCA1 | 19 | Phospholipid-transporting ATPase ABCA1 |
| THBS1 | sp\|P07996\|TSP1_HUMAN | P07996 | TSP1 | 7057 | Thrombospondin-1 |
| CLTA | sp\|P09496\|CLCA_HUMAN | P09496 | CLCA | 1211 | Clathrin light chain A |
| ALAD | sp\|P13716\|HEM2_HUMAN | P13716 | HEM2 | 210 | Delta-aminolevulinic acid dehydratase |
| ANXA8 | sp\|P13928\|ANXA8_HUMAN | P13928 | ANXA8 | 728113; 653145 | Annexin A8 |
| RAB5A | sp\|P20339\|RAB5A_HUMAN | P20339 | RAB5A | 5868 | Ras-related protein Rab-5A |
| GAL | sp\|P22466\|GALA_HUMAN | P22466 | GALA | 51083 | Galanin peptides |
| RPS12 | sp\|P25398\|RS12_HUMAN | P25398 | RS12 | 6206 | 40S ribosomal protein S12 |
| PSMB8 | sp\|P28062\|PSB8_HUMAN | P28062 | PSB8 | 5696 | Proteasome subunit beta type-8 |
| RANBP1 | sp\|P43487\|RANG_HUMAN | P43487 | RANG | 5902 | Ran-specific GTPase-activating protein |
| PSMC4 | sp\|P43686\|PRS6B_HUMAN | P43686 | PRS6B | 5704 | 26S proteasome regulatory subunit 6B |
| RPL21 | sp\|P46778\|RL21_HUMAN | P46778 | RL21 | 6144 | 60S ribosomal protein L21 |
| JAM2 | sp\|P57087\|JAM2_HUMAN | P57087 | JAM2 | 58494 | Junctional adhesion molecule B |
| EPPK1 | sp\|P58107\|EPIPL_HUMAN | P58107 | EPIPL | 83481 | Epiplakin |
| RPS20 | sp\|P60866\|RS20_HUMAN | P60866 | RS20 | 6224 | 40S ribosomal protein S20 |
| S100A10 | sp\|P60903\|S10AA_HUMAN | P60903 | S10AA | 6281 | Protein S100-A10 |
| RPS14 | sp\|P62263\|RS14_HUMAN | P62263 | RS14 | 6208 | 40S ribosomal protein S14 |
| RPL23A | sp\|P62750\|RL23A_HUMAN | P62750 | RL23A | 6147 | 60S ribosomal protein L23a |
| RPS25 | sp\|P62851\|RS25_HUMAN | P62851 | RS25 | 6230 | 40S ribosomal protein S25 |
| FKBP1A | sp\|P62942\|FKB1A_HUMAN | P62942 | FKB1A | 2280 | Peptidyl-prolyl cis-trans isomerase FKBP1A |
| RPL19 | sp\|P84098\|RL19_HUMAN | P84098 | RL19 | 6143 | 60S ribosomal protein L19 |
| VLDLR | sp\|P98155\|VLDLR_HUMAN | P98155 | VLDLR | 7436 | Very low-density lipoprotein receptor |
| SORD | sp\|Q00796\|DHSO_HUMAN | Q00796 | DHSO | 6652 | Sorbitol dehydrogenase |
| SLC7A5 | sp\|Q01650\|LAT1_HUMAN | Q01650 | LAT1 | 8140 | Large neutral amino acids transporter small subunit 1 |
| TWF1 | sp\|Q12792\|TWF1_HUMAN | Q12792 | TWF1 | 5756 | Twinfilin-1 |
| ALCAM | sp\|Q13740\|CD166_HUMAN | Q13740 | CD166 | 214 | CD166 antigen |
| FKBP8 | sp\|Q14318\|FKBP8_HUMAN | Q14318 | FKBP8 | 23770 | Peptidyl-prolyl cis-trans isomerase FKBP8 |
| GASK1B | sp\|Q6UWH4\|GAK1B_HUMAN | Q6UWH4 | GAK1B | 51313 | Golgi-associated kinase 1B |
| HS2ST1 | sp\|Q7LGA3\|HS2ST_HUMAN | Q7LGA3 | HS2ST | 9653 | Heparan sulfate 2-O-sulfotransferase 1 |
| USP9X | sp\|Q93008\|USP9X_HUMAN | Q93008 | USP9X | 8239 | Probable ubiquitin carboxyl-terminal hydrolase FAF-X |
| ISOC1 | sp\|Q96CN7\|ISOC1_HUMAN | Q96CN7 | ISOC1 | 51015 | Isochorismatase domain-containing protein 1 |
| CYFIP2 | sp\|Q96F07\|CYFP2_HUMAN | Q96F07 | CYFP2 | 26999 | Cytoplasmic FMR1-interacting protein 2 |
| TRIOBP | sp\|Q9H2D6\|TARA_HUMAN | Q9H2D6 | TARA | 11078 | TRIO and F-actin-binding protein |
| GLIPR2 | sp\|Q9H4G4\|GAPR1_HUMAN | Q9H4G4 | GAPR1 | 152007 | Golgi-associated plant pathogenesis-related protein 1 |
| SLC38A10 | sp\|Q9HBR0\|S38AA_HUMAN | Q9HBR0 | S38AA | 124565 | Putative sodium-coupled neutral amino acid transporter 10 |
| S100A14 | sp\|Q9HCY8\|S10AE_HUMAN | Q9HCY8 | S10AE | 57402 | Protein S100-A14 |
| ATXN10 | sp\|Q9UBB4\|ATX10_HUMAN | Q9UBB4 | ATX10 | 25814 | Ataxin-10 |
| GRHPR | sp\|Q9UBQ7\|GRHPR_HUMAN | Q9UBQ7 | GRHPR | 9380 | Glyoxylate reductase/hydroxypyruvate reductase |
| DMBT1 | sp\|Q9UGM3\|DMBT1_HUMAN | Q9UGM3 | DMBT1 | 1755 | Deleted in malignant brain tumors 1 protein |
| SLC7A8 | sp\|Q9UHI5\|LAT2_HUMAN | Q9UHI5 | LAT2 | 23428 | Large neutral amino acids transporter small subunit 2 |
| ERG28 | sp\|Q9UKR5\|ERG28_HUMAN | Q9UKR5 | ERG28 | 11161 | Ergosterol biosynthetic protein 28 homolog |
| MACF1 | sp\|Q9UPN3\|MACF1_HUMAN | Q9UPN3 | MACF1 | 23499 | Microtubule-actin cross-linking factor 1, isoforms 1/2/3/5 |
| HNRNPA1L3 | sp\|A0A2R8Y4L2\|RA1L3_HUMAN | A0A2R8Y4L2 | RA1L3 | 642659 | Heterogeneous nuclear ribonucleoprotein A1-like 3 |
| AP3B1 | sp\|O00203\|AP3B1_HUMAN | O00203 | AP3B1 | 8546 | AP-3 complex subunit beta-1 |
| LCP1 | sp\|P13796\|PLSL_HUMAN | P13796 | PLSL | 3936 | Plastin-2 |
| ARPC5 | sp\|O15511\|ARPC5_HUMAN | O15511 | ARPC5 | 10092 | Actin-related protein 2/3 complex subunit 5 |
| AKR7A2 | sp\|O43488\|ARK72_HUMAN | O43488 | ARK72 | 8574 | Aflatoxin B1 aldehyde reductase member 2 |
| SLC4A2 | sp\|P04920\|B3A2_HUMAN | P04920 | B3A2 | 6522 | Anion exchange protein 2 |
| MPO | sp\|P05164\|PERM_HUMAN | P05164 | PERM | 4353 | Myeloperoxidase |
| SCG5 | sp\|P05408\|7B2_HUMAN | P05408 | 7B2 | 6447 | Neuroendocrine protein 7B2 |
| IGFBP5 | sp\|P24593\|IBP5_HUMAN | P24593 | IBP5 | 3488 | Insulin-like growth factor-binding protein 5 |
| PSMA4 | sp\|P25789\|PSA4_HUMAN | P25789 | PSA4 | 5685 | Proteasome subunit alpha type-4 |
| SRI | sp\|P30626\|SORCN_HUMAN | P30626 | SORCN | 6717 | Sorcin |
| PAFAH1B2 | sp\|P68402\|PA1B2_HUMAN | P68402 | PA1B2 | 5049 | Platelet-activating factor acetylhydrolase IB subunit alpha2 |
| HSDL2 | sp\|Q6YN16\|HSDL2_HUMAN | Q6YN16 | HSDL2 | 84263 | Hydroxysteroid dehydrogenase-like protein 2 |
| GOLGA7 | sp\|Q7Z5G4\|GOGA7_HUMAN | Q7Z5G4 | GOGA7 | 51125 | Golgin subfamily A member 7 |
| SDK1 | sp\|Q7Z5N4\|SDK1_HUMAN | Q7Z5N4 | SDK1 | 221935 | Protein sidekick-1 |
| PSEN1 | sp\|P49768\|PSN1_HUMAN | P49768 | PSN1 | 5663 | Presenilin-1 |
| TM7SF2 | sp\|O76062\|ERG24_HUMAN | O76062 | ERG24 | 7108 | Delta(14)-sterol reductase TM7SF2 |
| RPL14 | sp\|P50914\|RL14_HUMAN | P50914 | RL14 | 9045 | 60S ribosomal protein L14 |
| PSMD4 | sp\|P55036\|PSMD4_HUMAN | P55036 | PSMD4 | 5710 | 26S proteasome non-ATPase regulatory subunit 4 |
| HADHB | sp\|P55084\|ECHB_HUMAN | P55084 | ECHB | 3032 | Trifunctional enzyme subunit beta, mitochondrial |
| TSPAN14 | sp\|Q8NG11\|TSN14_HUMAN | Q8NG11 | TSN14 | 81619 | Tetraspanin-14 |
| OTUB1 | sp\|Q96FW1\|OTUB1_HUMAN | Q96FW1 | OTUB1 | 55611 | Ubiquitin thioesterase OTUB1 |
| VPS28 | sp\|Q9UK41\|VPS28_HUMAN | Q9UK41 | VPS28 | 51160 | Vacuolar protein sorting-associated protein 28 homolog |
| RAB22A | sp\|Q9UL26\|RB22A_HUMAN | Q9UL26 | RB22A | 57403 | Ras-related protein Rab-22A |
| MINPP1 | sp\|Q9UNW1\|MINP1_HUMAN | Q9UNW1 | MINP1 | 9562 | Multiple inositol polyphosphate phosphatase 1 |
| TSPAN9 | sp\|O75954\|TSN9_HUMAN | O75954 | TSN9 | 10867 | Tetraspanin-9 |
| EIF2S1 | sp\|P05198\|IF2A_HUMAN | P05198 | IF2A | 1965 | Eukaryotic translation initiation factor 2 subunit 1 |
| RPS19 | sp\|P39019\|RS19_HUMAN | P39019 | RS19 | 6223 | 40S ribosomal protein S19 |
| NSF | sp\|P46459\|NSF_HUMAN | P46459 | NSF | 4905 | Vesicle-fusing ATPase |
| SYPL1 | sp\|Q16563\|SYPL1_HUMAN | Q16563 | SYPL1 | 6856 | Synaptophysin-like protein 1 |
| NCCRP1 | sp\|Q6ZVX7\|FBX50_HUMAN | Q6ZVX7 | FBX50 | 342897 | F-box only protein 50 |
| LACRT | sp\|Q9GZZ8\|LACRT_HUMAN | Q9GZZ8 | LACRT | 90070 | Extracellular glycoprotein lacritin |
| SRSF2 | sp\|Q01130\|SRSF2_HUMAN | Q01130 | SRSF2 | 6427 | Serine/arginine-rich splicing factor 2 |
| ADCY9 | sp\|O60503\|ADCY9_HUMAN | O60503 | ADCY9 | 115 | Adenylate cyclase type 9 |
| VPS26A | sp\|O75436\|VP26A_HUMAN | O75436 | VP26A | 9559 | Vacuolar protein sorting-associated protein 26A |
| LAMP1 | sp\|P11279\|LAMP1_HUMAN | P11279 | LAMP1 | 3916 | Lysosome-associated membrane glycoprotein 1 |
| PSMC1 | sp\|P62191\|PRS4_HUMAN | P62191 | PRS4 | 5700 | 26S proteasome regulatory subunit 4 |
| FERMT2 | sp\|Q96AC1\|FERM2_HUMAN | Q96AC1 | FERM2 | 10979 | Fermitin family homolog 2 |
| SMOC2 | sp\|Q9H3U7\|SMOC2_HUMAN | Q9H3U7 | SMOC2 | 64094 | SPARC-related modular calcium-binding protein 2 |
| PALS2 | sp\|Q9NZW5\|PALS2_HUMAN | Q9NZW5 | PALS2 | 51678 | Protein PALS2 |
| SNX3 | sp\|O60493\|SNX3_HUMAN | O60493 | SNX3 | 8724 | Sorting nexin-3 |
| BLVRB | sp\|P30043\|BLVRB_HUMAN | P30043 | BLVRB | 645 | Flavin reductase (NADPH) |
| AQP5 | sp\|P55064\|AQP5_HUMAN | P55064 | AQP5 | 362 | Aquaporin-5 |
| EFHD2 | sp\|Q96C19\|EFHD2_HUMAN | Q96C19 | EFHD2 | 79180 | EF-hand domain-containing protein D2 |
| PITPNB | sp\|P48739\|PIPNB_HUMAN | P48739 | PIPNB | 23760 | Phosphatidylinositol transfer protein beta isoform |
| RPS23 | sp\|P62266\|RS23_HUMAN | P62266 | RS23 | 6228 | 40S ribosomal protein S23 |
| RPL10A | sp\|P62906\|RL10A_HUMAN | P62906 | RL10A | 4736 | 60S ribosomal protein L10a |
| NEU2 | sp\|Q9Y3R4\|NEUR2_HUMAN | Q9Y3R4 | NEUR2 | 4759 | Sialidase-2 |
| SERPINB6 | sp\|P35237\|SPB6_HUMAN | P35237 | SPB6 | 5269 | Serpin B6 |
| IST1 | sp\|P53990\|IST1_HUMAN | P53990 | IST1 | 9798 | IST1 homolog |
| RPS24 | sp\|P62847\|RS24_HUMAN | P62847 | RS24 | 6229 | 40S ribosomal protein S24 |
| NOTCH2 | sp\|Q04721\|NOTC2_HUMAN | Q04721 | NOTC2 | 4853 | Neurogenic locus notch homolog protein 2 |
| PTGES | sp\|O14684\|PTGES_HUMAN | O14684 | PTGES | 9536 | Prostaglandin E synthase |
| ARPC1B | sp\|O15143\|ARC1B_HUMAN | O15143 | ARC1B | 10095 | Actin-related protein 2/3 complex subunit 1B |
| LCAT | sp\|P04180\|LCAT_HUMAN | P04180 | LCAT | 3931 | Phosphatidylcholine-sterol acyltransferase |
| MGST2 | sp\|Q99735\|MGST2_HUMAN | Q99735 | MGST2 | 4258 | Microsomal glutathione S-transferase 2 |
| MAGED2 | sp\|Q9UNF1\|MAGD2_HUMAN | Q9UNF1 | MAGD2 | 10916 | Melanoma-associated antigen D2 |
| IGHV2-26 | sp\|A0A0B4J1V2\|HV226_HUMAN | A0A0B4J1V2 | HV226 | 28455 | Immunoglobulin heavy variable 2-26 |
| FDX1 | sp\|P10109\|ADX_HUMAN | P10109 | ADX | 2230 | Adrenodoxin, mitochondrial |
| ADH1B | sp\|P00325\|ADH1B_HUMAN | P00325 | ADH1B | 125 | All-trans-retinol dehydrogenase [NAD(+)] ADH1B |
| MIF | sp\|P14174\|MIF_HUMAN | P14174 | MIF | 4282 | Macrophage migration inhibitory factor |
| EIF6 | sp\|P56537\|IF6_HUMAN | P56537 | IF6 | 3692 | Eukaryotic translation initiation factor 6 |
| FKBP4 | sp\|Q02790\|FKBP4_HUMAN | Q02790 | FKBP4 | 2288 | Peptidyl-prolyl cis-trans isomerase FKBP4 |
| MGAT2 | sp\|Q10469\|MGAT2_HUMAN | Q10469 | MGAT2 | 4247 | Alpha-1,6-mannosyl-glycoprotein 2-beta-N-acetylglucosaminyltransferase |
| SQSTM1 | sp\|Q13501\|SQSTM_HUMAN | Q13501 | SQSTM | 8878 | Sequestosome-1 |
| HEBP1 | sp\|Q9NRV9\|HEBP1_HUMAN | Q9NRV9 | HEBP1 | 50865 | Heme-binding protein 1 |
| RPL17 | sp\|P18621\|RL17_HUMAN | P18621 | RL17 | 6139 | 60S ribosomal protein L17 |
| PRR4 | sp\|Q16378\|PROL4_HUMAN | Q16378 | PROL4 | 11272 | Proline-rich protein 4 |
| IGHV3-64D | sp\|A0A0J9YX35\|HV64D_HUMAN | A0A0J9YX35 | HV64D | 1.03E+08 | Immunoglobulin heavy variable 3-64D |
| CXCL12 | sp\|P48061\|SDF1_HUMAN | P48061 | SDF1 | 6387 | Stromal cell-derived factor 1 |
| SEMA3C | sp\|Q99985\|SEM3C_HUMAN | Q99985 | SEM3C | 10512 | Semaphorin-3C |
| MACROH2A2 | sp\|Q9P0M6\|H2AW_HUMAN | Q9P0M6 | H2AW | 55506 | Core histone macro-H2A.2 |
| SPRR1A | sp\|P35321\|SPR1A_HUMAN | P35321 | SPR1A | 6698 | Cornifin-A |
| EXTL2 | sp\|Q9UBQ6\|EXTL2_HUMAN | Q9UBQ6 | EXTL2 | 2135 | Exostosin-like 2 |
| AP2A2 | sp\|O94973\|AP2A2_HUMAN | O94973 | AP2A2 | 161 | AP-2 complex subunit alpha-2 |
| YARS1 | sp\|P54577\|SYYC_HUMAN | P54577 | SYYC | 8565 | Tyrosine--tRNA ligase, cytoplasmic |
| GOLM1 | sp\|Q8NBJ4\|GOLM1_HUMAN | Q8NBJ4 | GOLM1 | 51280 | Golgi membrane protein 1 |
| ARPC1A | sp\|Q92747\|ARC1A_HUMAN | Q92747 | ARC1A | 10552 | Actin-related protein 2/3 complex subunit 1A |
| MAT2A | sp\|P31153\|METK2_HUMAN | P31153 | METK2 | 4144 | S-adenosylmethionine synthase isoform type-2 |
| GPC1 | sp\|P35052\|GPC1_HUMAN | P35052 | GPC1 | 2817 | Glypican-1 |
| TXNDC17 | sp\|Q9BRA2\|TXD17_HUMAN | Q9BRA2 | TXD17 | 84817 | Thioredoxin domain-containing protein 17 |
| DNAJA1 | sp\|P31689\|DNJA1_HUMAN | P31689 | DNJA1 | 3301 | DnaJ homolog subfamily A member 1 |
| IGHV2-70D | sp\|A0A0C4DH43\|HV70D_HUMAN | A0A0C4DH43 | HV70D | 1.03E+08 | Immunoglobulin heavy variable 2-70D |
| APOC2 | sp\|P02655\|APOC2_HUMAN | P02655 | APOC2 | 344 | Apolipoprotein C-II |
| CRISP3 | sp\|P54108\|CRIS3_HUMAN | P54108 | CRIS3 | 10321 | Cysteine-rich secretory protein 3 |
| SLC9A1 | sp\|P19634\|SL9A1_HUMAN | P19634 | SL9A1 | 6548 | Sodium/hydrogen exchanger 1 |
| SH3BGRL3 | sp\|Q9H299\|SH3L3_HUMAN | Q9H299 | SH3L3 | 83442 | SH3 domain-binding glutamic acid-rich-like protein 3 |
| SCGB1D2 | sp\|O95969\|SG1D2_HUMAN | O95969 | SG1D2 | 10647 | Secretoglobin family 1D member 2 |
| TXNRD1 | sp\|Q16881\|TRXR1_HUMAN | Q16881 | TRXR1 | 7296 | Thioredoxin reductase 1, cytoplasmic |
| SAR1A | sp\|Q9NR31\|SAR1A_HUMAN | Q9NR31 | SAR1A | 56681 | GTP-binding protein SAR1a |
| CYB5R1 | sp\|Q9UHQ9\|NB5R1_HUMAN | Q9UHQ9 | NB5R1 | 51706 | NADH-cytochrome b5 reductase 1 |
| MYOF | sp\|Q9NZM1\|MYOF_HUMAN | Q9NZM1 | MYOF | 26509 | Myoferlin |
| GART | sp\|P22102\|PUR2_HUMAN | P22102 | PUR2 | 2618 | Trifunctional purine biosynthetic protein adenosine-3 |
| PSMB9 | sp\|P28065\|PSB9_HUMAN | P28065 | PSB9 | 5698 | Proteasome subunit beta type-9 |
| LASP1 | sp\|Q14847\|LASP1_HUMAN | Q14847 | LASP1 | 3927 | LIM and SH3 domain protein 1 |
| RHOC | sp\|P08134\|RHOC_HUMAN | P08134 | RHOC | 389 | Rho-related GTP-binding protein RhoC |
| PSMB6 | sp\|P28072\|PSB6_HUMAN | P28072 | PSB6 | 5694 | Proteasome subunit beta type-6 |
| IGHV6-1 | sp\|A0A0B4J1U7\|HV601_HUMAN | A0A0B4J1U7 | HV601 | 28385 | Immunoglobulin heavy variable 6-1 |
| GSTA4 | sp\|O15217\|GSTA4_HUMAN | O15217 | GSTA4 | 2941 | Glutathione S-transferase A4 |
| S1PR3 | sp\|Q99500\|S1PR3_HUMAN | Q99500 | S1PR3 | 1903 | Sphingosine 1-phosphate receptor 3 |
| IGLV7-46 | sp\|A0A075B6I9\|LV746_HUMAN | A0A075B6I9 | LV746 | 28775 | Immunoglobulin lambda variable 7-46 |
| UBE2V1 | sp\|Q13404\|UB2V1_HUMAN | Q13404 | UB2V1 | 387522; 7335 | Ubiquitin-conjugating enzyme E2 variant 1 |
| IGHV3-38-3 | sp\|P0DTE1\|HV383_HUMAN | P0DTE1 | HV383 | 28404 | Probable non-functional immunoglobulin heavy variable 3-38-3 |
| IGHV1-18 | sp\|A0A0C4DH31\|HV118_HUMAN | A0A0C4DH31 | HV118 | 28468 | Immunoglobulin heavy variable 1-18 |
| MUC16 | sp\|Q8WXI7\|MUC16_HUMAN | Q8WXI7 | MUC16 | 94025 | Mucin-16 |
| RNASE7 | sp\|Q9H1E1\|RNAS7_HUMAN | Q9H1E1 | RNAS7 | 84659 | Ribonuclease 7 |
| PTP4A1 | sp\|Q93096\|TP4A1_HUMAN | Q93096 | TP4A1 | 7803 | Protein tyrosine phosphatase type IVA 1 |
| IGHV3-73 | sp\|A0A0B4J1V6\|HV373_HUMAN | A0A0B4J1V6 | HV373 | 28409 | Immunoglobulin heavy variable 3-73 |
| ETF1 | sp\|P62495\|ERF1_HUMAN | P62495 | ERF1 | 2107 | Eukaryotic peptide chain release factor subunit 1 |
| GSPT2 | sp\|Q8IYD1\|ERF3B_HUMAN | Q8IYD1 | ERF3B | 23708 | Eukaryotic peptide chain release factor GTP-binding subunit ERF3B |
| IGKV3D-20 | sp\|A0A0C4DH25\|KVD20_HUMAN | A0A0C4DH25 | KVD20 | 28874 | Immunoglobulin kappa variable 3D-20 |
| IGLV4-69 | sp\|A0A075B6H9\|LV469_HUMAN | A0A075B6H9 | LV469 | 28784 | Immunoglobulin lambda variable 4-69 |
| IGHV1-24 | sp\|A0A0C4DH33\|HV124_HUMAN | A0A0C4DH33 | HV124 | 28467 | Immunoglobulin heavy variable 1-24 |
| IGKV1-16 | sp\|P04430\|KV116_HUMAN | P04430 | KV116 | 28938 | Immunoglobulin kappa variable 1-16 |
| IGLL1 | sp\|P15814\|IGLL1_HUMAN | P15814 | IGLL1 | 3543 | Immunoglobulin lambda-like polypeptide 1 |
| MAPK3 | sp\|P27361\|MK03_HUMAN | P27361 | MK03 | 5595 | Mitogen-activated protein kinase 3 |
| PFN2 | sp\|P35080\|PROF2_HUMAN | P35080 | PROF2 | 5217 | Profilin-2 |
| SEC13 | sp\|P55735\|SEC13_HUMAN | P55735 | SEC13 | 6396 | Protein SEC13 homolog |
| FCGRT | sp\|P55899\|FCGRN_HUMAN | P55899 | FCGRN | 2217 | IgG receptor FcRn large subunit p51 |
| FAM3A | sp\|P98173\|FAM3A_HUMAN | P98173 | FAM3A | 60343 | Protein FAM3A |
| RPL18A | sp\|Q02543\|RL18A_HUMAN | Q02543 | RL18A | 6142 | 60S ribosomal protein L18a |
| SLC25A11 | sp\|Q02978\|M2OM_HUMAN | Q02978 | M2OM | 8402 | Mitochondrial 2-oxoglutarate/malate carrier protein |
| TKFC | sp\|Q3LXA3\|TKFC_HUMAN | Q3LXA3 | TKFC | 26007 | Triokinase/FMN cyclase |
| LHFPL2 | sp\|Q6ZUX7\|LHPL2_HUMAN | Q6ZUX7 | LHPL2 | 10184 | LHFPL tetraspan subfamily member 2 protein |
| WDTC1 | sp\|Q8N5D0\|WDTC1_HUMAN | Q8N5D0 | WDTC1 | 23038 | WD and tetratricopeptide repeats protein 1 |
| ATAD1 | sp\|Q8NBU5\|ATAD1_HUMAN | Q8NBU5 | ATAD1 | 84896 | Outer mitochondrial transmembrane helix translocase |
| MGARP | sp\|Q8TDB4\|HUMMR_HUMAN | Q8TDB4 | HUMMR | 84709 | Protein MGARP |
| TFG | sp\|Q92734\|TFG_HUMAN | Q92734 | TFG | 10342 | Protein TFG |
| TMEM230 | sp\|Q96A57\|TM230_HUMAN | Q96A57 | TM230 | 29058 | Transmembrane protein 230 |
| COLEC11 | sp\|Q9BWP8\|COL11_HUMAN | Q9BWP8 | COL11 | 78989 | Collectin-11 |
| TBC1D10A | sp\|Q9BXI6\|TB10A_HUMAN | Q9BXI6 | TB10A | 83874 | TBC1 domain family member 10A |
| SPNS1 | sp\|Q9H2V7\|SPNS1_HUMAN | Q9H2V7 | SPNS1 | 83985 | Protein spinster homolog 1 |
| LOXL2 | sp\|Q9Y4K0\|LOXL2_HUMAN | Q9Y4K0 | LOXL2 | 4017 | Lysyl oxidase homolog 2 |
| CDC42BPB | sp\|Q9Y5S2\|MRCKB_HUMAN | Q9Y5S2 | MRCKB | 9578 | Serine/threonine-protein kinase MRCK beta |
| TMEM109 | sp\|Q9BVC6\|TM109_HUMAN | Q9BVC6 | TM109 | 79073 | Transmembrane protein 109 |
| TINAGL1 | sp\|Q9GZM7\|TINAL_HUMAN | Q9GZM7 | TINAL | 64129 | Tubulointerstitial nephritis antigen-like |
| PGLS | sp\|O95336\|6PGL_HUMAN | O95336 | 6PGL | 25796 | 6-phosphogluconolactonase |
| CRP | sp\|P02741\|CRP_HUMAN | P02741 | CRP | 1401 | C-reactive protein |
| PFKM | sp\|P08237\|PFKAM_HUMAN | P08237 | PFKAM | 5213 | ATP-dependent 6-phosphofructokinase, muscle type |
| ARPC5L | sp\|Q9BPX5\|ARP5L_HUMAN | Q9BPX5 | ARP5L | 81873 | Actin-related protein 2/3 complex subunit 5-like protein |
| IL36G | sp\|Q9NZH8\|IL36G_HUMAN | Q9NZH8 | IL36G | 56300 | Interleukin-36 gamma |
| SPRR1B | sp\|P22528\|SPR1B_HUMAN | P22528 | SPR1B | 6699 | Cornifin-B |
| SPINT2 | sp\|O43291\|SPIT2_HUMAN | O43291 | SPIT2 | 10653 | Kunitz-type protease inhibitor 2 |
| VARS1 | sp\|P26640\|SYVC_HUMAN | P26640 | SYVC | 7407 | Valine--tRNA ligase |
| CUL7 | sp\|Q14999\|CUL7_HUMAN | Q14999 | CUL7 | 9820 | Cullin-7 |
| ABI1 | sp\|Q8IZP0\|ABI1_HUMAN | Q8IZP0 | ABI1 | 10006 | Abl interactor 1 |
| PRKAR2A | sp\|P13861\|KAP2_HUMAN | P13861 | KAP2 | 5576 | cAMP-dependent protein kinase type II-alpha regulatory subunit |
| RHBDF2 | sp\|Q6PJF5\|RHDF2_HUMAN | Q6PJF5 | RHDF2 | 79651 | Inactive rhomboid protein 2 |
| ALOX12 | sp\|P18054\|LOX12_HUMAN | P18054 | LOX12 | 239 | Polyunsaturated fatty acid lipoxygenase ALOX12 |
| HNRNPD | sp\|Q14103\|HNRPD_HUMAN | Q14103 | HNRPD | 3184 | Heterogeneous nuclear ribonucleoprotein D0 |
| EPS8L1 | sp\|Q8TE68\|ES8L1_HUMAN | Q8TE68 | ES8L1 | 54869 | Epidermal growth factor receptor kinase substrate 8-like protein 1 |
| HNRNPH3 | sp\|P31942\|HNRH3_HUMAN | P31942 | HNRH3 | 3189 | Heterogeneous nuclear ribonucleoprotein H3 |
| CPA4 | sp\|Q9UI42\|CBPA4_HUMAN | Q9UI42 | CBPA4 | 51200 | Carboxypeptidase A4 |
| PITPNA | sp\|Q00169\|PIPNA_HUMAN | Q00169 | PIPNA | 5306 | Phosphatidylinositol transfer protein alpha isoform |
| CNNM2 | sp\|Q9H8M5\|CNNM2_HUMAN | Q9H8M5 | CNNM2 | 54805 | Metal transporter CNNM2 |
| APEH | sp\|P13798\|ACPH_HUMAN | P13798 | ACPH | 327 | Acylamino-acid-releasing enzyme |
| CSGALNACT1 | sp\|Q8TDX6\|CGAT1_HUMAN | Q8TDX6 | CGAT1 | 55790 | Chondroitin sulfate N-acetylgalactosaminyltransferase 1 |
| IGLV5-39 | sp\|A0A0G2JS06\|LV539_HUMAN | A0A0G2JS06 | LV539 | 28782 | Immunoglobulin lambda variable 5-39 |
| DNAJB1 | sp\|P25685\|DNJB1_HUMAN | P25685 | DNJB1 | 3337 | DnaJ homolog subfamily B member 1 |
| GALC | sp\|P54803\|GALC_HUMAN | P54803 | GALC | 2581 | Galactocerebrosidase |
| SEC61G | sp\|P60059\|SC61G_HUMAN | P60059 | SC61G | 23480 | Protein transport protein Sec61 subunit gamma |
| GVINP1 | sp\|Q7Z2Y8\|GVIN1_HUMAN | Q7Z2Y8 | GVIN1 | 387751 | Interferon-induced very large GTPase 1 |
| RAB23 | sp\|Q9ULC3\|RAB23_HUMAN | Q9ULC3 | RAB23 | 51715 | Ras-related protein Rab-23 |
| NACA | sp\|E9PAV3\|NACAM_HUMAN | E9PAV3 | NACAM | 4666 | Nascent polypeptide-associated complex subunit alpha, muscle-specific form |
| PAPSS2 | sp\|O95340\|PAPS2_HUMAN | O95340 | PAPS2 | 9060 | Bifunctional 3'-phosphoadenosine 5'-phosphosulfate synthase 2 |
| DCAF8L2 | sp\|P0C7V8\|DC8L2_HUMAN | P0C7V8 | DC8L2 | 347442 | DDB1- and CUL4-associated factor 8-like protein 2 |
| KLC1 | sp\|Q07866\|KLC1_HUMAN | Q07866 | KLC1 | 3831 | Kinesin light chain 1 |
| USP14 | sp\|P54578\|UBP14_HUMAN | P54578 | UBP14 | 9097 | Ubiquitin carboxyl-terminal hydrolase 14 |
| COASY | sp\|Q13057\|COASY_HUMAN | Q13057 | COASY | 80347 | Bifunctional coenzyme A synthase |
| IGHV3-64 | sp\|A0A075B6Q5\|HV364_HUMAN | A0A075B6Q5 | HV364 | 28414 | Immunoglobulin heavy variable 3-64 |
| MSH5 | sp\|O43196\|MSH5_HUMAN | O43196 | MSH5 | 4439 | MutS protein homolog 5 |
| PYGB | sp\|P11216\|PYGB_HUMAN | P11216 | PYGB | 5834 | Glycogen phosphorylase, brain form |
| AKR1A1 | sp\|P14550\|AK1A1_HUMAN | P14550 | AK1A1 | 10327 | Aldo-keto reductase family 1 member A1 |
| IGHV3-38 | sp\|A0A0C4DH36\|HV338_HUMAN | A0A0C4DH36 | HV338 | 28429 | Probable non-functional immunoglobulin heavy variable 3-38 |
| IGLV2-23 | sp\|P01705\|LV223_HUMAN | P01705 | LV223 | 28813 | Immunoglobulin lambda variable 2-23 |
| CCDC8 | sp\|Q9H0W5\|CCDC8_HUMAN | Q9H0W5 | CCDC8 | 83987 | Coiled-coil domain-containing protein 8 |
| PODXL | sp\|O00592\|PODXL_HUMAN | O00592 | PODXL | 5420 | Podocalyxin |
| PLD1 | sp\|Q13393\|PLD1_HUMAN | Q13393 | PLD1 | 5337 | Phospholipase D1 |
| RPL35 | sp\|P42766\|RL35_HUMAN | P42766 | RL35 | 11224 | 60S ribosomal protein L35 |
| CDC5L | sp\|Q99459\|CDC5L_HUMAN | Q99459 | CDC5L | 988 | Cell division cycle 5-like protein |
| BANK1 | sp\|Q8NDB2\|BANK1_HUMAN | Q8NDB2 | BANK1 | 55024 | B-cell scaffold protein with ankyrin repeats |
| PMVK | sp\|Q15126\|PMVK_HUMAN | Q15126 | PMVK | 10654 | Phosphomevalonate kinase |
| DNAJC5 | sp\|Q9H3Z4\|DNJC5_HUMAN | Q9H3Z4 | DNJC5 | 80331 | DnaJ homolog subfamily C member 5 |
| ABCB8 | sp\|Q9NUT2\|MITOS_HUMAN | Q9NUT2 | MITOS | 11194 | Mitochondrial potassium channel ATP-binding subunit |
| ITGA11 | sp\|Q9UKX5\|ITA11_HUMAN | Q9UKX5 | ITA11 | 22801 | Integrin alpha-11 |
| DNAH7 | sp\|Q8WXX0\|DYH7_HUMAN | Q8WXX0 | DYH7 | 56171 | Dynein axonemal heavy chain 7 |
| ATP6V1G1 | sp\|O75348\|VATG1_HUMAN | O75348 | VATG1 | 9550 | V-type proton ATPase subunit G 1 |
| JPT1 | sp\|Q9UK76\|JUPI1_HUMAN | Q9UK76 | JUPI1 | 51155 | Jupiter microtubule associated homolog 1 |
| IGFBP7 | sp\|Q16270\|IBP7_HUMAN | Q16270 | IBP7 | 3490 | Insulin-like growth factor-binding protein 7 |
| C18orf63 | sp\|Q68DL7\|CR063_HUMAN | Q68DL7 | CR063 | 644041 | Uncharacterized protein C18orf63 |
| PLPP3 | sp\|O14495\|PLPP3_HUMAN | O14495 | PLPP3 | 8613 | Phospholipid phosphatase 3 |
| CD44 | sp\|P16070\|CD44_HUMAN | P16070 | CD44 | 960 | CD44 antigen |
| BROX | sp\|Q5VW32\|BROX_HUMAN | Q5VW32 | BROX | 148362 | BRO1 domain-containing protein BROX |

| **Supplementary Table 3: Filtered proteins in ffEVs (790) used in further statistical analysis** | |
| --- | --- |
| **Protein_Gen Symbol** | **description** |
| C4B_2 | Complement C4-B |
| H2AC11 | Histone H2A type 1 |
| H3C3 | Histone H3.1 |
| H4C2 | Histone H4 |
| HBA1 | Hemoglobin subunit alpha |
| LGALS7B | Galectin-7 |
| A1BG | Alpha-1B-glycoprotein |
| A2M | Alpha-2-macroglobulin |
| AADAC | Arylacetamide deacetylase |
| ACAN | Aggrecan core protein |
| ACAT2 | Acetyl-CoA acetyltransferase, cytosolic |
| ACLY | ATP-citrate synthase |
| ACP3 | Prostatic acid phosphatase |
| ACSL4 | Long-chain-fatty-acid--CoA ligase 4 |
| ACTB | Actin, cytoplasmic 1 |
| ACTBL2 | Beta-actin-like protein 2 |
| ACTC1 | Actin, alpha cardiac muscle 1 |
| ACTG1 | Actin, cytoplasmic 2 |
| ACTN4 | Alpha-actinin-4 |
| ACTR1A | Alpha-centractin |
| ACTR2 | Actin-related protein 2 |
| ACTR3 | Actin-related protein 3 |
| ADCY9 | Adenylate cyclase type 9 |
| ADGRG1 | Adhesion G-protein coupled receptor G1 |
| ADIPOQ | Adiponectin |
| AFM | Afamin |
| AGT | Angiotensinogen |
| AHCY | Adenosylhomocysteinase |
| AHSG | Alpha-2-HS-glycoprotein |
| AKR1B1 | Aldo-keto reductase family 1 member B1 |
| ALB | Albumin |
| ALCAM | CD166 antigen |
| ALDH3A2 | Aldehyde dehydrogenase family 3 member A2 |
| ALDOA | Fructose-bisphosphate aldolase A |
| ALDOC | Fructose-bisphosphate aldolase C |
| AMBP | Protein AMBP |
| AMY1B | Alpha-amylase 1B |
| ANG | Angiogenin |
| ANGPTL4 | Angiopoietin-related protein 4 |
| ANXA1 | Annexin A1 |
| ANXA11 | Annexin A11 |
| ANXA2 | Annexin A2 |
| ANXA5 | Annexin A5 |
| ANXA6 | Annexin A6 |
| ANXA7 | Annexin A7 |
| AOC3 | Membrane primary amine oxidase |
| APCS | Serum amyloid P-component |
| APMAP | Adipocyte plasma membrane-associated protein |
| APOA1 | Apolipoprotein A-I |
| APOA2 | Apolipoprotein A-II |
| APOA4 | Apolipoprotein A-IV |
| APOB | Apolipoprotein B-100 |
| APOC1 | Apolipoprotein C-I |
| APOC3 | Apolipoprotein C-III |
| APOC4 | Apolipoprotein C-IV |
| APOD | Apolipoprotein D |
| APOE | Apolipoprotein E |
| APOF | Apolipoprotein F |
| APOH | Beta-2-glycoprotein 1 |
| APOL1 | Apolipoprotein L1 |
| APOM | Apolipoprotein M |
| APRT | Adenine phosphoribosyltransferase |
| ARF3 | ADP-ribosylation factor 3 |
| ARF4 | ADP-ribosylation factor 4 |
| ARF6 | ADP-ribosylation factor 6 |
| ARG1 | Arginase-1 |
| ARHGDIA | Rho GDP-dissociation inhibitor 1 |
| ARL3 | ADP-ribosylation factor-like protein 3 |
| ARL6IP5 | PRA1 family protein 3 |
| ARPC3 | Actin-related protein 2/3 complex subunit 3 |
| ARPC4 | Actin-related protein 2/3 complex subunit 4 |
| ARPC5 | Actin-related protein 2/3 complex subunit 5 |
| ATP1A1 | Sodium/potassium-transporting ATPase subunit alpha-1 |
| ATP1B1 | Sodium/potassium-transporting ATPase subunit beta-1 |
| ATP1B3 | Sodium/potassium-transporting ATPase subunit beta-3 |
| ATP2A2 | Sarcoplasmic/endoplasmic reticulum calcium ATPase 2 |
| ATP2B1 | Plasma membrane calcium-transporting ATPase 1 |
| ATP2B4 | Plasma membrane calcium-transporting ATPase 4 |
| ATP5F1A | ATP synthase subunit alpha, mitochondrial |
| ATP5F1B | ATP synthase subunit beta, mitochondrial |
| ATP6V0A1 | V-type proton ATPase 116 kDa subunit a 1 |
| ATP6V1B2 | V-type proton ATPase subunit B, brain isoform |
| ATRN | Attractin |
| AZGP1 | Zinc-alpha-2-glycoprotein |
| B2M | Beta-2-microglobulin |
| BASP1 | Brain acid soluble protein 1 |
| BCAP31 | B-cell receptor-associated protein 31 |
| BCHE | Cholinesterase |
| BLVRB | Flavin reductase (NADPH) |
| BMPR2 | Bone morphogenetic protein receptor type-2 |
| BSG | Basigin |
| BST2 | Bone marrow stromal antigen 2 |
| BTD | Biotinidase |
| C18orf63 | Uncharacterized protein C18orf63 |
| C1QA | Complement C1q subcomponent subunit A |
| C1QB | Complement C1q subcomponent subunit B |
| C1QC | Complement C1q subcomponent subunit C |
| C1R | Complement C1r subcomponent |
| C1RL | Complement C1r subcomponent-like protein |
| C1S | Complement C1s subcomponent |
| C2 | Complement C2 |
| C3 | Complement C3 |
| C4A | Complement C4-A |
| C4BPA | C4b-binding protein alpha chain |
| C4BPB | C4b-binding protein beta chain |
| C5 | Complement C5 |
| C6 | Complement component C6 |
| C7 | Complement component C7 |
| C8A | Complement component C8 alpha chain |
| C8B | Complement component C8 beta chain |
| C8G | Complement component C8 gamma chain |
| C9 | Complement component C9 |
| CALM1 | Calmodulin-1 |
| CALML3 | Calmodulin-like protein 3 |
| CALML5 | Calmodulin-like protein 5 |
| CALR | Calreticulin |
| CALU | Calumenin |
| CAMP | Cathelicidin antimicrobial peptide |
| CANX | Calnexin |
| CAPN1 | Calpain-1 catalytic subunit |
| CAPZA1 | F-actin-capping protein subunit alpha-1 |
| CAPZB | F-actin-capping protein subunit beta |
| CASK | Peripheral plasma membrane protein CASK |
| CASP14 | Caspase-14 |
| CAT | Catalase |
| CBR1 | Carbonyl reductase [NADPH] 1 |
| CCT2 | T-complex protein 1 subunit beta |
| CCT3 | T-complex protein 1 subunit gamma |
| CCT4 | T-complex protein 1 subunit delta |
| CCT5 | T-complex protein 1 subunit epsilon |
| CCT6A | T-complex protein 1 subunit zeta |
| CCT7 | T-complex protein 1 subunit eta |
| CCT8 | T-complex protein 1 subunit theta |
| CD14 | Monocyte differentiation antigen CD14 |
| CD276 | CD276 antigen |
| CD46 | Membrane cofactor protein |
| CD47 | Leukocyte surface antigen CD47 |
| CD55 | Complement decay-accelerating factor |
| CD59 | CD59 glycoprotein |
| CD5L | CD5 antigen-like |
| CD63 | CD63 antigen |
| CD81 | CD81 antigen |
| CD9 | CD9 antigen |
| CD99 | CD99 antigen |
| CDC42 | Cell division control protein 42 homolog |
| CDC5L | Cell division cycle 5-like protein |
| CDH1 | Cadherin-1 |
| CDH2 | Cadherin-2 |
| CDSN | Corneodesmosin |
| CETP | Cholesteryl ester transfer protein |
| CFB | Complement factor B |
| CFD | Complement factor D |
| CFH | Complement factor H |
| CFHR1 | Complement factor H-related protein 1 |
| CFHR2 | Complement factor H-related protein 2 |
| CFHR5 | Complement factor H-related protein 5 |
| CFI | Complement factor I |
| CFL1 | Cofilin-1 |
| CFP | Properdin |
| CHMP4B | Charged multivesicular body protein 4b |
| CHP1 | Calcineurin B homologous protein 1 |
| CKAP4 | Cytoskeleton-associated protein 4 |
| CKB | Creatine kinase B-type |
| CLEC3B | Tetranectin |
| CLIC1 | Chloride intracellular channel protein 1 |
| CLTA | Clathrin light chain A |
| CLTC | Clathrin heavy chain 1 |
| CLU | Clusterin |
| CNDP2 | Cytosolic non-specific dipeptidase |
| CNP | 2',3'-cyclic-nucleotide 3'-phosphodiesterase |
| CNPY2 | Protein canopy homolog 2 |
| COL15A1 | Collagen alpha-1(XV) chain |
| COL18A1 | Collagen alpha-1(XVIII) chain |
| COL1A1 | Collagen alpha-1(I) chain |
| COL4A1 | Collagen alpha-1(IV) chain |
| COL4A2 | Collagen alpha-2(IV) chain |
| COL5A2 | Collagen alpha-2(V) chain |
| COL6A1 | Collagen alpha-1(VI) chain |
| COL6A2 | Collagen alpha-2(VI) chain |
| COL6A3 | Collagen alpha-3(VI) chain |
| COMT | Catechol O-methyltransferase |
| COPA | Coatomer subunit alpha |
| COX4I1 | Cytochrome c oxidase subunit 4 isoform 1, mitochondrial |
| CP | Ceruloplasmin |
| CPB2 | Carboxypeptidase B2 |
| CPM | Carboxypeptidase M |
| CPN1 | Carboxypeptidase N catalytic chain |
| CPN2 | Carboxypeptidase N subunit 2 |
| CPNE2 | Copine-2 |
| CPNE3 | Copine-3 |
| CRISPLD2 | Cysteine-rich secretory protein LCCL domain-containing 2 |
| CRNN | Cornulin |
| CRP | C-reactive protein |
| CSGALNACT1 | Chondroitin sulfate N-acetylgalactosaminyltransferase 1 |
| CST3 | Cystatin-C |
| CSTA | Cystatin-A |
| CTNNA1 | Catenin alpha-1 |
| CTNNB1 | Catenin beta-1 |
| CTNND1 | Catenin delta-1 |
| CTSD | Cathepsin D |
| CXCL12 | Stromal cell-derived factor 1 |
| CYB5R3 | NADH-cytochrome b5 reductase 3 |
| CYFIP1 | Cytoplasmic FMR1-interacting protein 1 |
| CYP11A1 | Cholesterol side-chain cleavage enzyme, mitochondrial |
| CYP17A1 | Steroid 17-alpha-hydroxylase/17,20 lyase |
| CYP19A1 | Aromatase |
| CYP51A1 | Lanosterol 14-alpha demethylase |
| DAD1 | Dolichyl-diphosphooligosaccharide--protein glycosyltransferase subunit DAD1 |
| DARS1 | Aspartate--tRNA ligase, cytoplasmic |
| DCAF8L2 | DDB1- and CUL4-associated factor 8-like protein 2 |
| DCD | Dermcidin |
| DCTN1 | Dynactin subunit 1 |
| DDAH1 | N(G),N(G)-dimethylarginine dimethylaminohydrolase 1 |
| DDOST | Dolichyl-diphosphooligosaccharide--protein glycosyltransferase 48 kDa subunit |
| DHCR24 | Delta(24)-sterol reductase |
| DHCR7 | 7-dehydrocholesterol reductase |
| DIPK2A | Divergent protein kinase domain 2A |
| DMKN | Dermokine |
| DNAH7 | Dynein axonemal heavy chain 7 |
| DNM2 | Dynamin-2 |
| DPP4 | Dipeptidyl peptidase 4 |
| DPYSL3 | Dihydropyrimidinase-related protein 3 |
| DSC1 | Desmocollin-1 |
| DSC2 | Desmocollin-2 |
| DSC3 | Desmocollin-3 |
| DSG1 | Desmoglein-1 |
| DSP | Desmoplakin |
| DSTN | Destrin |
| DYNC1H1 | Cytoplasmic dynein 1 heavy chain 1 |
| DYNC1I2 | Cytoplasmic dynein 1 intermediate chain 2 |
| DYNC1LI2 | Cytoplasmic dynein 1 light intermediate chain 2 |
| ECE1 | Endothelin-converting enzyme 1 |
| ECH1 | Delta(3,5)-Delta(2,4)-dienoyl-CoA isomerase, mitochondrial |
| ECM1 | Extracellular matrix protein 1 |
| EEF1A1 | Elongation factor 1-alpha 1 |
| EEF1G | Elongation factor 1-gamma |
| EEF2 | Elongation factor 2 |
| EFNB1 | Ephrin-B1 |
| EGFR | Epidermal growth factor receptor |
| EHD1 | EH domain-containing protein 1 |
| EHD4 | EH domain-containing protein 4 |
| EIF2S1 | Eukaryotic translation initiation factor 2 subunit 1 |
| EIF4A1 | Eukaryotic initiation factor 4A-I |
| EIF5A | Eukaryotic translation initiation factor 5A-1 |
| EIF6 | Eukaryotic translation initiation factor 6 |
| EMILIN1 | EMILIN-1 |
| ENO1 | Alpha-enolase |
| EPB41L2 | Band 4.1-like protein 2 |
| EPHX1 | Epoxide hydrolase 1 |
| ERAP1 | Endoplasmic reticulum aminopeptidase 1 |
| ERGIC1 | Endoplasmic reticulum-Golgi intermediate compartment protein 1 |
| ERLIN2 | Erlin-2 |
| ERMP1 | Endoplasmic reticulum metallopeptidase 1 |
| ERP29 | Endoplasmic reticulum resident protein 29 |
| ERP44 | Endoplasmic reticulum resident protein 44 |
| EVPL | Envoplakin |
| EZR | Ezrin |
| F10 | Coagulation factor X |
| F11 | Coagulation factor XI |
| F11R | Junctional adhesion molecule A |
| F12 | Coagulation factor XII |
| F13B | Coagulation factor XIII B chain |
| F2 | Prothrombin |
| F5 | Coagulation factor V |
| F9 | Coagulation factor IX |
| FABP5 | Fatty acid-binding protein 5 |
| FAM234A | Protein FAM234A |
| FASN | Fatty acid synthase |
| FBLN1 | Fibulin-1 |
| FCN2 | Ficolin-2 |
| FCN3 | Ficolin-3 |
| FDFT1 | Squalene synthase |
| FDPS | Farnesyl pyrophosphate synthase |
| FETUB | Fetuin-B |
| FGA | Fibrinogen alpha chain |
| FGB | Fibrinogen beta chain |
| FGG | Fibrinogen gamma chain |
| FKBP1A | Peptidyl-prolyl cis-trans isomerase FKBP1A |
| FLG2 | Filaggrin-2 |
| FLNA | Filamin-A |
| FLNB | Filamin-B |
| FLOT1 | Flotillin-1 |
| FLOT2 | Flotillin-2 |
| FN1 | Fibronectin |
| FREM2 | FRAS1-related extracellular matrix protein 2 |
| FSCN1 | Fascin |
| FST | Follistatin |
| FTH1 | Ferritin heavy chain |
| GAL | Galanin peptides |
| GANAB | Neutral alpha-glucosidase AB |
| GAPDH | Glyceraldehyde-3-phosphate dehydrogenase |
| GC | Vitamin D-binding protein |
| GCLC | Glutamate--cysteine ligase catalytic subunit |
| GDI2 | Rab GDP dissociation inhibitor beta |
| GJA1 | Gap junction alpha-1 protein |
| GLG1 | Golgi apparatus protein 1 |
| GNA11 | Guanine nucleotide-binding protein subunit alpha-11 |
| GNA13 | Guanine nucleotide-binding protein subunit alpha-13 |
| GNAI1 | Guanine nucleotide-binding protein G(i) subunit alpha-1 |
| GNAI2 | Guanine nucleotide-binding protein G(i) subunit alpha-2 |
| GNAI3 | Guanine nucleotide-binding protein G(i) subunit alpha-3 |
| GNAQ | Guanine nucleotide-binding protein G(q) subunit alpha |
| GNAS | Guanine nucleotide-binding protein G(s) subunit alpha isoforms short |
| GNB1 | Guanine nucleotide-binding protein G(I)/G(S)/G(T) subunit beta-1 |
| GNB2 | Guanine nucleotide-binding protein G(I)/G(S)/G(T) subunit beta-2 |
| GOLM2 | Protein GOLM2 |
| GOT1 | Aspartate aminotransferase, cytoplasmic |
| GPC6 | Glypican-6 |
| GPI | Glucose-6-phosphate isomerase |
| GPLD1 | Phosphatidylinositol-glycan-specific phospholipase D |
| GPX3 | Glutathione peroxidase 3 |
| GRHPR | Glyoxylate reductase/hydroxypyruvate reductase |
| GSN | Gelsolin |
| GSTA1 | Glutathione S-transferase A1 |
| GSTP1 | Glutathione S-transferase P |
| GVINP1 | Interferon-induced very large GTPase 1 |
| H1-4 | Histone H1.4 |
| H2BC12 | Histone H2B type 1-K |
| **HABP2** | Hyaluronan-binding protein 2 |
| HAL | Histidine ammonia-lyase |
| HBB | Hemoglobin subunit beta |
| HLA-A | HLA class I histocompatibility antigen, A alpha chain |
| HLA-B | HLA class I histocompatibility antigen, B alpha chain |
| HLA-C | HLA class I histocompatibility antigen, C alpha chain |
| HMGCS1 | Hydroxymethylglutaryl-CoA synthase, cytoplasmic |
| HNRNPK | Heterogeneous nuclear ribonucleoprotein K |
| HP | Haptoglobin |
| HPR | Haptoglobin-related protein |
| HPX | Hemopexin |
| HRG | Histidine-rich glycoprotein |
| HRNR | Hornerin |
| HS3ST1 | Heparan sulfate glucosamine 3-O-sulfotransferase 1 |
| HSD17B1 | 17-beta-hydroxysteroid dehydrogenase type 1 |
| HSD17B12 | Very-long-chain 3-oxoacyl-CoA reductase |
| HSD3B2 | 3 beta-hydroxysteroid dehydrogenase/Delta 5--4-isomerase type 2 |
| HSP90AA1 | Heat shock protein HSP 90-alpha |
| HSP90AB1 | Heat shock protein HSP 90-beta |
| HSP90B1 | Endoplasmin |
| HSPA1A | Heat shock 70 kDa protein 1A |
| HSPA5 | Endoplasmic reticulum chaperone BiP |
| HSPA8 | Heat shock cognate 71 kDa protein |
| HSPB1 | Heat shock protein beta-1 |
| HSPD1 | 60 kDa heat shock protein, mitochondrial |
| HSPG2 | Basement membrane-specific heparan sulfate proteoglycan core protein |
| HTRA1 | Serine protease HTRA1 |
| HTRA3 | Serine protease HTRA3 |
| HUWE1 | E3 ubiquitin-protein ligase HUWE1 |
| HVCN1 | Voltage-gated hydrogen channel 1 |
| HYOU1 | Hypoxia up-regulated protein 1 |
| IDH1 | Isocitrate dehydrogenase [NADP] cytoplasmic |
| IGF1R | Insulin-like growth factor 1 receptor |
| IGF2 | Insulin-like growth factor II |
| IGF2R | Cation-independent mannose-6-phosphate receptor |
| IGFALS | Insulin-like growth factor-binding protein complex acid labile subunit |
| IGFBP2 | Insulin-like growth factor-binding protein 2 |
| IGFBP3 | Insulin-like growth factor-binding protein 3 |
| IGFBP4 | Insulin-like growth factor-binding protein 4 |
| IGFBP7 | Insulin-like growth factor-binding protein 7 |
| IGHA1 | Immunoglobulin heavy constant alpha 1 |
| IGHA2 | Immunoglobulin heavy constant alpha 2 |
| IGHG1 | Immunoglobulin heavy constant gamma 1 |
| IGHG2 | Immunoglobulin heavy constant gamma 2 |
| IGHG3 | Immunoglobulin heavy constant gamma 3 |
| IGHG4 | Immunoglobulin heavy constant gamma 4 |
| IGHM | Immunoglobulin heavy constant mu |
| IGHV1-18 | Immunoglobulin heavy variable 1-18 |
| IGHV1-2 | Immunoglobulin heavy variable 1-2 |
| IGHV1-3 | Immunoglobulin heavy variable 1-3 |
| IGHV1-46 | Immunoglobulin heavy variable 1-46 |
| IGHV2-26 | Immunoglobulin heavy variable 2-26 |
| IGHV2-70D | Immunoglobulin heavy variable 2-70D |
| IGHV3-15 | Immunoglobulin heavy variable 3-15 |
| IGHV3-23 | Immunoglobulin heavy variable 3-23 |
| IGHV3-38 | Probable non-functional immunoglobulin heavy variable 3-38 |
| IGHV3-43D | Immunoglobulin heavy variable 3-43D |
| IGHV3-49 | Immunoglobulin heavy variable 3-49 |
| IGHV3-64D | Immunoglobulin heavy variable 3-64D |
| IGHV3-7 | Immunoglobulin heavy variable 3-7 |
| IGHV3-72 | Immunoglobulin heavy variable 3-72 |
| IGHV3-74 | Immunoglobulin heavy variable 3-74 |
| IGHV4-28 | Immunoglobulin heavy variable 4-28 |
| IGHV4-38-2 | Immunoglobulin heavy variable 4-38-2 |
| IGHV5-51 | Immunoglobulin heavy variable 5-51 |
| IGHV6-1 | Immunoglobulin heavy variable 6-1 |
| IGKC | Immunoglobulin kappa constant |
| IGKV1-16 | Immunoglobulin kappa variable 1-16 |
| IGKV1-17 | Immunoglobulin kappa variable 1-17 |
| IGKV1-39 | Immunoglobulin kappa variable 1-39 |
| IGKV1-5 | Immunoglobulin kappa variable 1-5 |
| IGKV1-6 | Immunoglobulin kappa variable 1-6 |
| IGKV1-8 | Immunoglobulin kappa variable 1-8 |
| IGKV1D-13 | Immunoglobulin kappa variable 1D-13 |
| IGKV1D-16 | Immunoglobulin kappa variable 1D-16 |
| IGKV1D-33 | Immunoglobulin kappa variable 1D-33 |
| IGKV2-28 | Immunoglobulin kappa variable 2-28 |
| IGKV2-30 | Immunoglobulin kappa variable 2-30 |
| IGKV2D-24 | Probable non-functional immunoglobulin kappa variable 2D-24 |
| IGKV3-20 | Immunoglobulin kappa variable 3-20 |
| IGKV3-7 | Probable non-functional immunoglobulin kappa variable 3-7 |
| IGKV3D-11 | Immunoglobulin kappa variable 3D-11 |
| IGKV3D-15 | Immunoglobulin kappa variable 3D-15 |
| IGKV3D-20 | Immunoglobulin kappa variable 3D-20 |
| IGKV4-1 | Immunoglobulin kappa variable 4-1 |
| IGKV6-21 | Immunoglobulin kappa variable 6-21 |
| IGLC3 | Immunoglobulin lambda constant 3 |
| IGLL1 | Immunoglobulin lambda-like polypeptide 1 |
| IGLL5 | Immunoglobulin lambda-like polypeptide 5 |
| IGLV1-40 | Immunoglobulin lambda variable 1-40 |
| IGLV1-44 | Immunoglobulin lambda variable 1-44 |
| IGLV1-47 | Immunoglobulin lambda variable 1-47 |
| IGLV1-51 | Immunoglobulin lambda variable 1-51 |
| IGLV2-11 | Immunoglobulin lambda variable 2-11 |
| IGLV2-14 | Immunoglobulin lambda variable 2-14 |
| IGLV2-18 | Immunoglobulin lambda variable 2-18 |
| IGLV2-23 | Immunoglobulin lambda variable 2-23 |
| IGLV2-8 | Immunoglobulin lambda variable 2-8 |
| IGLV3-10 | Immunoglobulin lambda variable 3-10 |
| IGLV3-21 | Immunoglobulin lambda variable 3-21 |
| IGLV3-27 | Immunoglobulin lambda variable 3-27 |
| IGLV4-69 | Immunoglobulin lambda variable 4-69 |
| IGLV7-46 | Immunoglobulin lambda variable 7-46 |
| IGLV8-61 | Immunoglobulin lambda variable 8-61 |
| IGSF8 | Immunoglobulin superfamily member 8 |
| IL6ST | Interleukin-6 receptor subunit beta |
| INHA | Inhibin alpha chain |
| IQGAP1 | Ras GTPase-activating-like protein IQGAP1 |
| ITGA2 | Integrin alpha-2 |
| ITGA5 | Integrin alpha-5 |
| ITGA6 | Integrin alpha-6 |
| ITGA7 | Integrin alpha-7 |
| ITGA9 | Integrin alpha-9 |
| ITGAV | Integrin alpha-V |
| ITGB1 | Integrin beta-1 |
| ITIH1 | Inter-alpha-trypsin inhibitor heavy chain H1 |
| ITIH2 | Inter-alpha-trypsin inhibitor heavy chain H2 |
| ITIH3 | Inter-alpha-trypsin inhibitor heavy chain H3 |
| ITIH4 | Inter-alpha-trypsin inhibitor heavy chain H4 |
| JAM2 | Junctional adhesion molecule B |
| JCHAIN | Immunoglobulin J chain |
| JUP | Junction plakoglobin |
| KLC1 | Kinesin light chain 1 |
| KLKB1 | Plasma kallikrein |
| KNG1 | Kininogen-1 |
| KPNB1 | Importin subunit beta-1 |
| KPRP | Keratinocyte proline-rich protein |
| KRT1 | Keratin, type II cytoskeletal 1 |
| KRT10 | Keratin, type I cytoskeletal 10 |
| KRT13 | Keratin, type I cytoskeletal 13 |
| KRT14 | Keratin, type I cytoskeletal 14 |
| KRT16 | Keratin, type I cytoskeletal 16 |
| KRT17 | Keratin, type I cytoskeletal 17 |
| KRT2 | Keratin, type II cytoskeletal 2 epidermal |
| KRT33B | Keratin, type I cuticular Ha3-II |
| KRT4 | Keratin, type II cytoskeletal 4 |
| KRT5 | Keratin, type II cytoskeletal 5 |
| KRT6A | Keratin, type II cytoskeletal 6A |
| KRT6B | Keratin, type II cytoskeletal 6B |
| KRT77 | Keratin, type II cytoskeletal 1b |
| KRT78 | Keratin, type II cytoskeletal 78 |
| KRT80 | Keratin, type II cytoskeletal 80 |
| KRT85 | Keratin, type II cuticular Hb5 |
| KRT9 | Keratin, type I cytoskeletal 9 |
| LAMA1 | Laminin subunit alpha-1 |
| LAMB2 | Laminin subunit beta-2 |
| LAMC1 | Laminin subunit gamma-1 |
| LAMP1 | Lysosome-associated membrane glycoprotein 1 |
| LBP | Lipopolysaccharide-binding protein |
| LCAT | Phosphatidylcholine-sterol acyltransferase |
| LCN1 | Lipocalin-1 |
| LDHA | L-lactate dehydrogenase A chain |
| LDHB | L-lactate dehydrogenase B chain |
| LDLR | Low-density lipoprotein receptor |
| LGALS3BP | Galectin-3-binding protein |
| LHFPL2 | LHFPL tetraspan subfamily member 2 protein |
| LIN7C | Protein lin-7 homolog C |
| LMAN1 | Protein ERGIC-53 |
| LMAN2 | Vesicular integral-membrane protein VIP36 |
| LOXL2 | Lysyl oxidase homolog 2 |
| LPA | Apolipoprotein(a) |
| LPL | Lipoprotein lipase |
| LRG1 | Leucine-rich alpha-2-glycoprotein |
| LRP8 | Low-density lipoprotein receptor-related protein 8 |
| LSR | Lipolysis-stimulated lipoprotein receptor |
| LSS | Lanosterol synthase |
| LUM | Lumican |
| LYZ | Lysozyme C |
| M6PR | Cation-dependent mannose-6-phosphate receptor |
| MARCKS | Myristoylated alanine-rich C-kinase substrate |
| MASP1 | Mannan-binding lectin serine protease 1 |
| MASP2 | Mannan-binding lectin serine protease 2 |
| MATN2 | Matrilin-2 |
| MBL2 | Mannose-binding protein C |
| MCC | Colorectal mutant cancer protein |
| MDH1 | Malate dehydrogenase, cytoplasmic |
| MDK | Midkine |
| MFGE8 | Lactadherin |
| MGP | Matrix Gla protein |
| MPO | Myeloperoxidase |
| MPZL1 | Myelin protein zero-like protein 1 |
| MSMO1 | Methylsterol monooxygenase 1 |
| MSN | Moesin |
| MT-CO2 | Cytochrome c oxidase subunit 2 |
| MTHFD1 | C-1-tetrahydrofolate synthase, cytoplasmic |
| MVP | Major vault protein |
| MYDGF | Myeloid-derived growth factor |
| MYH10 | Myosin-10 |
| MYH9 | Myosin-9 |
| MYL12B | Myosin regulatory light chain 12B |
| MYL6 | Myosin light polypeptide 6 |
| MYO1B | Unconventional myosin-Ib |
| NAPA | Alpha-soluble NSF attachment protein |
| NCSTN | Nicastrin |
| NDRG1 | Protein NDRG1 |
| NID1 | Nidogen-1 |
| NME2 | Nucleoside diphosphate kinase B |
| NPTN | Neuroplastin |
| NPTX2 | Neuronal pentraxin-2 |
| NRAS | GTPase NRas |
| NSDHL | Sterol-4-alpha-carboxylate 3-dehydrogenase, decarboxylating |
| NTS | Neurotensin/neuromedin N |
| ORM1 | Alpha-1-acid glycoprotein 1 |
| ORM2 | Alpha-1-acid glycoprotein 2 |
| P3H1 | Prolyl 3-hydroxylase 1 |
| P4HB | Protein disulfide-isomerase |
| PABPC1 | Polyadenylate-binding protein 1 |
| PAICS | Multifunctional protein ADE2 |
| PAM | Peptidyl-glycine alpha-amidating monooxygenase |
| PAPPA | Pappalysin-1 |
| PARK7 | Parkinson disease protein 7 |
| PBXIP1 | Pre-B-cell leukemia transcription factor-interacting protein 1 |
| PCBP1 | Poly(rC)-binding protein 1 |
| PDCD6IP | Programmed cell death 6-interacting protein |
| PDIA3 | Protein disulfide-isomerase A3 |
| PDIA4 | Protein disulfide-isomerase A4 |
| PDIA6 | Protein disulfide-isomerase A6 |
| PEBP1 | Phosphatidylethanolamine-binding protein 1 |
| PFKL | ATP-dependent 6-phosphofructokinase, liver type |
| PFN1 | Profilin-1 |
| PGAM1 | Phosphoglycerate mutase 1 |
| PGD | 6-phosphogluconate dehydrogenase, decarboxylating |
| PGK1 | Phosphoglycerate kinase 1 |
| PGLYRP2 | N-acetylmuramoyl-L-alanine amidase |
| PGM1 | Phosphoglucomutase-1 |
| PGRMC1 | Membrane-associated progesterone receptor component 1 |
| PGRMC2 | Membrane-associated progesterone receptor component 2 |
| PHEX | Phosphate-regulating neutral endopeptidase PHEX |
| PITPNB | Phosphatidylinositol transfer protein beta isoform |
| PKM | Pyruvate kinase PKM |
| PKP1 | Plakophilin-1 |
| PLD3 | 5'-3' exonuclease PLD3 |
| PLG | Plasminogen |
| PLIN3 | Perilipin-3 |
| PLP1 | Myelin proteolipid protein |
| PLS1 | Plastin-1 |
| PLS3 | Plastin-3 |
| PLTP | Phospholipid transfer protein |
| POF1B | Protein POF1B |
| PON1 | Serum paraoxonase/arylesterase 1 |
| POR | NADPH--cytochrome P450 reductase |
| PPA1 | Inorganic pyrophosphatase |
| PPIA | Peptidyl-prolyl cis-trans isomerase A |
| PPIB | Peptidyl-prolyl cis-trans isomerase B |
| PPP2CB | Serine/threonine-protein phosphatase 2A catalytic subunit beta isoform |
| PPP2R1A | Serine/threonine-protein phosphatase 2A 65 kDa regulatory subunit A alpha isoform |
| PRDX1 | Peroxiredoxin-1 |
| PRDX2 | Peroxiredoxin-2 |
| PRDX4 | Peroxiredoxin-4 |
| PRDX5 | Peroxiredoxin-5, mitochondrial |
| PRDX6 | Peroxiredoxin-6 |
| PRKAR2B | cAMP-dependent protein kinase type II-beta regulatory subunit |
| PRKCSH | Glucosidase 2 subunit beta |
| PRKDC | DNA-dependent protein kinase catalytic subunit |
| PROC | Vitamin K-dependent protein C |
| PROS1 | Vitamin K-dependent protein S |
| PRSS1 | Serine protease 1 |
| PRSS23 | Serine protease 23 |
| PSAP | Prosaposin |
| PSMA1 | Proteasome subunit alpha type-1 |
| PSMA3 | Proteasome subunit alpha type-3 |
| PSMA5 | Proteasome subunit alpha type-5 |
| PSMA6 | Proteasome subunit alpha type-6 |
| PSMA7 | Proteasome subunit alpha type-7 |
| PSMB6 | Proteasome subunit beta type-6 |
| PSMB8 | Proteasome subunit beta type-8 |
| PSMC1 | 26S proteasome regulatory subunit 4 |
| PSMC2 | 26S proteasome regulatory subunit 7 |
| PSMD11 | 26S proteasome non-ATPase regulatory subunit 11 |
| PSMD2 | 26S proteasome non-ATPase regulatory subunit 2 |
| PSMD3 | 26S proteasome non-ATPase regulatory subunit 3 |
| PSMD6 | 26S proteasome non-ATPase regulatory subunit 6 |
| PSME1 | Proteasome activator complex subunit 1 |
| PTGFRN | Prostaglandin F2 receptor negative regulator |
| PTPRG | Receptor-type tyrosine-protein phosphatase gamma |
| PTX3 | Pentraxin-related protein PTX3 |
| PXDN | Peroxidasin homolog |
| PZP | Pregnancy zone protein |
| QSOX1 | Sulfhydryl oxidase 1 |
| RAB10 | Ras-related protein Rab-10 |
| RAB11A | Ras-related protein Rab-11A |
| RAB14 | Ras-related protein Rab-14 |
| RAB18 | Ras-related protein Rab-18 |
| RAB1B | Ras-related protein Rab-1B |
| RAB21 | Ras-related protein Rab-21 |
| RAB22A | Ras-related protein Rab-22A |
| RAB2A | Ras-related protein Rab-2A |
| RAB5B | Ras-related protein Rab-5B |
| RAB5C | Ras-related protein Rab-5C |
| RAB6A | Ras-related protein Rab-6A |
| RAB7A | Ras-related protein Rab-7a |
| RAC1 | Ras-related C3 botulinum toxin substrate 1 |
| RALA | Ras-related protein Ral-A |
| RAN | GTP-binding nuclear protein Ran |
| RANBP1 | Ran-specific GTPase-activating protein |
| RAP1B | Ras-related protein Rap-1b |
| RAP2B | Ras-related protein Rap-2b |
| RBP1 | Retinol-binding protein 1 |
| RBP4 | Retinol-binding protein 4 |
| RCN1 | Reticulocalbin-1 |
| RDX | Radixin |
| REEP6 | Receptor expression-enhancing protein 6 |
| RHBDF2 | Inactive rhomboid protein 2 |
| RHOA | Transforming protein RhoA |
| RHOB | Rho-related GTP-binding protein RhoB |
| RHOG | Rho-related GTP-binding protein RhoG |
| RNF170 | E3 ubiquitin-protein ligase RNF170 |
| RNH1 | Ribonuclease inhibitor |
| RPL11 | 60S ribosomal protein L11 |
| RPL13A | 60S ribosomal protein L13a |
| RPL14 | 60S ribosomal protein L14 |
| RPL15 | 60S ribosomal protein L15 |
| RPL18 | 60S ribosomal protein L18 |
| RPL35 | 60S ribosomal protein L35 |
| RPL4 | 60S ribosomal protein L4 |
| RPL6 | 60S ribosomal protein L6 |
| RPL7 | 60S ribosomal protein L7 |
| RPL7A | 60S ribosomal protein L7a |
| RPL8 | 60S ribosomal protein L8 |
| RPLP2 | 60S acidic ribosomal protein P2 |
| RPN1 | Dolichyl-diphosphooligosaccharide--protein glycosyltransferase subunit 1 |
| RPN2 | Dolichyl-diphosphooligosaccharide--protein glycosyltransferase subunit 2 |
| RPS15A | 40S ribosomal protein S15a |
| RPS2 | 40S ribosomal protein S2 |
| RPS25 | 40S ribosomal protein S25 |
| RPS3 | 40S ribosomal protein S3 |
| RPS6 | 40S ribosomal protein S6 |
| RPS8 | 40S ribosomal protein S8 |
| RPS9 | 40S ribosomal protein S9 |
| RPSA | 40S ribosomal protein SA |
| RRAS2 | Ras-related protein R-Ras2 |
| RRBP1 | Ribosome-binding protein 1 |
| RTN4 | Reticulon-4 |
| S100A10 | Protein S100-A10 |
| S100A13 | Protein S100-A13 |
| S100A16 | Protein S100-A16 |
| S100A7 | Protein S100-A7 |
| S100A8 | Protein S100-A8 |
| S100A9 | Protein S100-A9 |
| SAA1 | Serum amyloid A-1 protein |
| SAA2 | Serum amyloid A-2 protein |
| SAA4 | Serum amyloid A-4 protein |
| SAR1A | GTP-binding protein SAR1a |
| SBSN | Suprabasin |
| SCAMP3 | Secretory carrier-associated membrane protein 3 |
| SCARB1 | Scavenger receptor class B member 1 |
| SCARB2 | Lysosome membrane protein 2 |
| SCD | Stearoyl-CoA desaturase |
| SCUBE1 | Signal peptide, CUB and EGF-like domain-containing protein 1 |
| SDCBP | Syntenin-1 |
| SEC11A | Signal peptidase complex catalytic subunit SEC11A |
| SEC22B | Vesicle-trafficking protein SEC22b |
| SEL1L | Protein sel-1 homolog 1 |
| SELENOP | Selenoprotein P |
| SEPTIN11 | Septin-11 |
| SEPTIN9 | Septin-9 |
| SERINC1 | Serine incorporator 1 |
| SERPINA1 | Alpha-1-antitrypsin |
| SERPINA10 | Protein Z-dependent protease inhibitor |
| SERPINA3 | Alpha-1-antichymotrypsin |
| SERPINA4 | Kallistatin |
| SERPINA5 | Plasma serine protease inhibitor |
| SERPINA6 | Corticosteroid-binding globulin |
| SERPINA7 | Thyroxine-binding globulin |
| SERPINB12 | Serpin B12 |
| SERPINB3 | Serpin B3 |
| SERPINC1 | Antithrombin-III |
| SERPIND1 | Heparin cofactor 2 |
| SERPINE2 | Glia-derived nexin |
| SERPINF1 | Pigment epithelium-derived factor |
| SERPINF2 | Alpha-2-antiplasmin |
| SERPING1 | Plasma protease C1 inhibitor |
| SERPINH1 | Serpin H1 |
| SFN | 14-3-3 protein sigma |
| SFRP1 | Secreted frizzled-related protein 1 |
| SHBG | Sex hormone-binding globulin |
| SLC12A2 | Solute carrier family 12 member 2 |
| SLC16A1 | Monocarboxylate transporter 1 |
| SLC16A3 | Monocarboxylate transporter 4 |
| SLC1A4 | Neutral amino acid transporter A |
| SLC25A3 | Phosphate carrier protein, mitochondrial |
| SLC25A5 | ADP/ATP translocase 2 |
| SLC29A1 | Equilibrative nucleoside transporter 1 |
| SLC2A1 | Solute carrier family 2, facilitated glucose transporter member 1 |
| SLC39A14 | Metal cation symporter ZIP14 |
| SLC3A2 | 4F2 cell-surface antigen heavy chain |
| SLC47A1 | Multidrug and toxin extrusion protein 1 |
| SLC4A1 | Band 3 anion transport protein |
| SLC9A3R1 | Na(+)/H(+) exchange regulatory cofactor NHE-RF1 |
| SNAP23 | Synaptosomal-associated protein 23 |
| SNX1 | Sorting nexin-1 |
| SNX3 | Sorting nexin-3 |
| SNX6 | Sorting nexin-6 |
| SPARC | SPARC |
| SPON1 | Spondin-1 |
| SPP2 | Secreted phosphoprotein 24 |
| SPTAN1 | Spectrin alpha chain, non-erythrocytic 1 |
| SPTBN1 | Spectrin beta chain, non-erythrocytic 1 |
| SRPRB | Signal recognition particle receptor subunit beta |
| SRPX | Sushi repeat-containing protein SRPX |
| SSR1 | Translocon-associated protein subunit alpha |
| ST13 | Hsc70-interacting protein |
| STC1 | Stanniocalcin-1 |
| STIM1 | Stromal interaction molecule 1 |
| STOM | Stomatin |
| STX12 | Syntaxin-12 |
| STX4 | Syntaxin-4 |
| STX7 | Syntaxin-7 |
| SULF2 | Extracellular sulfatase Sulf-2 |
| SURF4 | Surfeit locus protein 4 |
| SYPL1 | Synaptophysin-like protein 1 |
| TAGLN2 | Transgelin-2 |
| TCP1 | T-complex protein 1 subunit alpha |
| TFPI | Tissue factor pathway inhibitor |
| TFRC | Transferrin receptor protein 1 |
| TGFBR3 | Transforming growth factor beta receptor type 3 |
| TGM3 | Protein-glutamine gamma-glutamyltransferase E |
| THBS4 | Thrombospondin-4 |
| TIMP1 | Metalloproteinase inhibitor 1 |
| TKT | Transketolase |
| TLN1 | Talin-1 |
| TM9SF2 | Transmembrane 9 superfamily member 2 |
| TMED10 | Transmembrane emp24 domain-containing protein 10 |
| TMED7 | Transmembrane emp24 domain-containing protein 7 |
| TMED9 | Transmembrane emp24 domain-containing protein 9 |
| TMEM109 | Transmembrane protein 109 |
| TMEM30A | Cell cycle control protein 50A |
| TMX1 | Thioredoxin-related transmembrane protein 1 |
| TNXB | Tenascin-X |
| TOLLIP | Toll-interacting protein |
| TPI1 | Triosephosphate isomerase |
| TPM3 | Tropomyosin alpha-3 chain |
| TPM4 | Tropomyosin alpha-4 chain |
| TPT1 | Translationally-controlled tumor protein |
| TRIOBP | TRIO and F-actin-binding protein |
| TSG101 | Tumor susceptibility gene 101 protein |
| TSKU | Tsukushi |
| TTR | Transthyretin |
| TTYH3 | Protein tweety homolog 3 |
| TUBA1B | Tubulin alpha-1B chain |
| TUBB | Tubulin beta chain |
| TUBB4A | Tubulin beta-4A chain |
| TUBB4B | Tubulin beta-4B chain |
| TXN | Thioredoxin |
| TXNDC5 | Thioredoxin domain-containing protein 5 |
| UBA1 | Ubiquitin-like modifier-activating enzyme 1 |
| UBB | Polyubiquitin-B |
| UBE2N | Ubiquitin-conjugating enzyme E2 N |
| UCHL1 | Ubiquitin carboxyl-terminal hydrolase isozyme L1 |
| UGGT1 | UDP-glucose:glycoprotein glucosyltransferase 1 |
| UQCRC2 | Cytochrome b-c1 complex subunit 2, mitochondrial |
| VAMP2 | Vesicle-associated membrane protein 2 |
| VAT1 | Synaptic vesicle membrane protein VAT-1 homolog |
| VCAM1 | Vascular cell adhesion protein 1 |
| VCAN | Versican core protein |
| VCL | Vinculin |
| VCP | Transitional endoplasmic reticulum ATPase |
| VPS28 | Vacuolar protein sorting-associated protein 28 homolog |
| VPS35 | Vacuolar protein sorting-associated protein 35 |
| VPS4B | Vacuolar protein sorting-associated protein 4B |
| VTN | Vitronectin |
| VWF | von Willebrand factor |
| WDR1 | WD repeat-containing protein 1 |
| XP32 | Skin-specific protein 32 |
| YES1 | Tyrosine-protein kinase Yes |
| YWHAB | 14-3-3 protein beta/alpha |
| YWHAE | 14-3-3 protein epsilon |
| YWHAG | 14-3-3 protein gamma |
| YWHAH | 14-3-3 protein eta |
| YWHAQ | 14-3-3 protein theta |
| YWHAZ | 14-3-3 protein zeta/delta |

| **Supplementary Table 4. Differentially abundant proteins (DAPs) in ffEVs (FDR<0.1%)** | | | |
| --- | --- | --- | --- |
| **MII vs GV** | |  |  |
| **Name** | **description** | **Fold change** | **P-value adj.** |
| F12 | Coagulation factor XII | 3.121700777 | 0.073926072 |
| MYDGF | Myeloid-derived growth factor | -2.68995734 | 0.097232583 |
| VWF | von Willebrand factor | -3.015133072 | 0.091929566 |
| CYP17A1 | Steroid 17-alpha-hydroxylase/17,20 lyase | -3.13473712 | 0.027009615 |
| SCD | Stearoyl-CoA desaturase | -3.369200349 | 0.062439065 |
| PRDX4 | Peroxiredoxin-4 | -3.409518609 | 0.027714191 |
| APOB | Apolipoprotein B-100 | -3.485107612 | 0.020898306 |
| ERGIC1 | Endoplasmic reticulum-Golgi intermediate compartment protein 1 | -3.599362105 | 0.062692531 |
| CYP51A1 | Lanosterol 14-alpha demethylase | -3.608985604 | 0.084773948 |
| CALU | Calumenin | -4.102417627 | 0.002575207 |
| PGRMC2 | Membrane-associated progesterone receptor component 2 | -4.387098336 | 0.002575207 |
| PRKCSH | Glucosidase 2 subunit beta | -4.430722567 | 0.006449441 |
| MSMO1 | Methylsterol monooxygenase 1 | -4.477163984 | 0.002575207 |
| HSD3B2 | 3 beta-hydroxysteroid dehydrogenase/Delta 5--4-isomerase type 2 | -4.755870202 | 0.019330173 |
| **MII vs GVBD** | |  |  |
| **Name** | **description** | **Fold change** | **P-value adj.** |
| IGKV1-39 | Immunoglobulin kappa variable 1-39 | 4.73165011 | 3.95988E-06 |
| FREM2 | FRAS1-related extracellular matrix protein 2 | 1.842861235 | 0.054361295 |
| C1QC | Complement C1q subcomponent subunit C | 1.311549088 | 0.047800816 |
| IGF2 | Insulin-like growth factor II | -2.252175843 | 0.049186212 |
| GAL | Galanin peptides | -2.368586916 | 0.000882037 |
| SAA2 | Serum amyloid A-2 protein | -2.396195437 | 0.05173607 |
| MDK | Midkine | -2.870416511 | 0.040394716 |
| **GV vs GVBD** | |  |  |
| **Name** | **description** | **Fold change** | **P-value adj.** |
| CALU | Calumenin | -4.260981673 | 0.099632858 |

| **Supplementary Table 5. Differential abundant proteins (DAPs) in ffEVs (FDR<0.2%)** | | | |
| --- | --- | --- | --- |
| **MII vs GV** |  |  |  |
| **Name** | **description** | **Fold change** | **P-value adj.** |
| FABP5 | Fatty acid-binding protein 5 | 3.641296456 | 0.105326334 |
| S100A9 | Protein S100-A9 | 3.290961272 | 0.147764654 |
| F12 | Coagulation factor XII | 3.121700777 | 0.073926072 |
| SERPINA10 | Protein Z-dependent protease inhibitor | 2.793656563 | 0.118836347 |
| IGKV1-8 | Immunoglobulin kappa variable 1-8 | 2.402710166 | 0.165639147 |
| IGKV1-5 | Immunoglobulin kappa variable 1-5 | 2.24854069 | 0.153172596 |
| CFD | Complement factor D | 2.136555971 | 0.168792836 |
| RAB18 | Ras-related protein Rab-18 | -1.611549507 | 0.198469635 |
| ECH1 | Delta(3,5)-Delta(2,4)-dienoyl-CoA isomerase, mitochondrial | -1.875199707 | 0.141778066 |
| F11R | Junctional adhesion molecule A | -1.876556582 | 0.180301655 |
| ATP2B1 | Plasma membrane calcium-transporting ATPase 1 | -2.068812991 | 0.120478233 |
| RPL6 | 60S ribosomal protein L6 | -2.099349113 | 0.146956594 |
| SEC11A | Signal peptidase complex catalytic subunit SEC11A | -2.182691769 | 0.138404237 |
| P3H1 | Prolyl 3-hydroxylase 1 | -2.200328716 | 0.133174787 |
| MVP | Major vault protein | -2.202801132 | 0.190062416 |
| ERP44 | Endoplasmic reticulum resident protein 44 | -2.246920191 | 0.18663538 |
| SLC2A1 | Solute carrier family 2, facilitated glucose transporter member 1 | -2.251850534 | 0.120947538 |
| ITGA5 | Integrin alpha-5 | -2.314255795 | 0.133086277 |
| TMX1 | Thioredoxin-related transmembrane protein 1 | -2.333840732 | 0.132154815 |
| IGHV3-38 | Probable non-functional immunoglobulin heavy variable 3-38 | -2.350373713 | 0.146597095 |
| GANAB | Neutral alpha-glucosidase AB | -2.394008136 | 0.116661941 |
| HSP90B1 | Endoplasmin | -2.396920654 | 0.101739111 |
| IGF2R | Cation-independent mannose-6-phosphate receptor | -2.473327476 | 0.125946284 |
| HSPA5 | Endoplasmic reticulum chaperone BiP | -2.474034484 | 0.139147021 |
| B2M | Beta-2-microglobulin | -2.511701597 | 0.149497522 |
| APMAP | Adipocyte plasma membrane-associated protein | -2.550365072 | 0.194515635 |
| AADAC | Arylacetamide deacetylase | -2.604798539 | 0.139450221 |
| TMED9 | Transmembrane emp24 domain-containing protein 9 | -2.605644759 | 0.12192245 |
| LMAN2 | Vesicular integral-membrane protein VIP36 | -2.625464613 | 0.130393475 |
| SSR1 | Translocon-associated protein subunit alpha | -2.639777251 | 0.123910364 |
| CD63 | CD63 antigen | -2.65680533 | 0.184438106 |
| COMT | Catechol O-methyltransferase | -2.676787689 | 0.125244739 |
| MYDGF | Myeloid-derived growth factor | -2.68995734 | 0.097232583 |
| UGGT1 | UDP-glucose:glycoprotein glucosyltransferase 1 | -2.781366408 | 0.121657644 |
| CYB5R3 | NADH-cytochrome b5 reductase 3 | -2.858459408 | 0.158350964 |
| SURF4 | Surfeit locus protein 4 | -2.905914054 | 0.12214002 |
| DAD1 | Dolichyl-diphosphooligosaccharide--protein glycosyltransferase subunit DAD1 | -2.916140922 | 0.110673141 |
| ERLIN2 | Erlin-2 | -2.924649841 | 0.101830335 |
| RCN1 | Reticulocalbin-1 | -2.935432529 | 0.129306483 |
| PDIA6 | Protein disulfide-isomerase A6 | -2.969064644 | 0.125845851 |
| ERP29 | Endoplasmic reticulum resident protein 29 | -3.006817078 | 0.129569603 |
| VWF | von Willebrand factor | -3.015133072 | 0.091929566 |
| PDIA3 | Protein disulfide-isomerase A3 | -3.026125209 | 0.119001906 |
| FDFT1 | Squalene synthase | -3.028972887 | 0.134447734 |
| ALDH3A2 | Aldehyde dehydrogenase family 3 member A2 | -3.037112287 | 0.131506416 |
| PDIA4 | Protein disulfide-isomerase A4 | -3.084667136 | 0.138649427 |
| LMAN1 | Protein ERGIC-53 | -3.089644327 | 0.132035522 |
| CYP17A1 | Steroid 17-alpha-hydroxylase/17,20 lyase | -3.13473712 | 0.027009615 |
| SEC22B | Vesicle-trafficking protein SEC22b | -3.156951155 | 0.138152467 |
| CANX | Calnexin | -3.19995337 | 0.126967027 |
| ERMP1 | Endoplasmic reticulum metallopeptidase 1 | -3.250890987 | 0.126392252 |
| SCD | Stearoyl-CoA desaturase | -3.369200349 | 0.062439065 |
| PRDX4 | Peroxiredoxin-4 | -3.409518609 | 0.027714191 |
| APOB | Apolipoprotein B-100 | -3.485107612 | 0.020898306 |
| ERGIC1 | Endoplasmic reticulum-Golgi intermediate compartment protein 1 | -3.599362105 | 0.062692531 |
| CYP51A1 | Lanosterol 14-alpha demethylase | -3.608985604 | 0.084773948 |
| CALU | Calumenin | -4.102417627 | 0.002575207 |
| PGRMC2 | Membrane-associated progesterone receptor component 2 | -4.387098336 | 0.002575207 |
| PRKCSH | Glucosidase 2 subunit beta | -4.430722567 | 0.006449441 |
| MSMO1 | Methylsterol monooxygenase 1 | -4.477163984 | 0.002575207 |
| HSD3B2 | 3 beta-hydroxysteroid dehydrogenase/Delta 5--4-isomerase type 2 | -4.755870202 | 0.019330173 |
| **MII vs GVBD** | |  |  |
| **Name** | **description** | **Fold change** | **P-value adj.** |
| IGKV1-39 | Immunoglobulin kappa variable 1-39 | 4.73165011 | 3.95988E-06 |
| FREM2 | FRAS1-related extracellular matrix protein 2 | 1.842861235 | 0.054361295 |
| C1QC | Complement C1q subcomponent subunit C | 1.311549088 | 0.047800816 |
| C1QA | Complement C1q subcomponent subunit A | 1.224130803 | 0.161617634 |
| ANG | Angiogenin | -2.021442728 | 0.123540987 |
| SERPINA7 | Thyroxine-binding globulin | -2.115793888 | 0.156411186 |
| IGF2 | Insulin-like growth factor II | -2.252175843 | 0.049186212 |
| GAL | Galanin peptides | -2.368586916 | 0.000882037 |
| SAA2 | Serum amyloid A-2 protein | -2.396195437 | 0.05173607 |
| AMY1B | Alpha-amylase 1B | -2.442765998 | 0.153352673 |
| KRT33B | Keratin, type I cuticular Ha3-II | -2.730655057 | 0.187128792 |
| SFRP1 | Secreted frizzled-related protein 1 | -2.793826433 | 0.121377411 |
| MGP | Matrix Gla protein | -2.817840958 | 0.139185784 |
| KRT85 | Keratin, type II cuticular Hb5 | -2.832062121 | 0.129411219 |
| MDK | Midkine | -2.870416511 | 0.040394716 |
| **GV vs GVBD** | |  |  |
| **Name** | **description** | **Fold change** | **P-value adj.** |
| MYDGF | Myeloid-derived growth factor | -3.221258745 | 0.170268305 |
| VWF | von Willebrand factor | -3.47387619 | 0.190163151 |
| PRDX4 | Peroxiredoxin-4 | -3.643961372 | 0.125152841 |
| TMED9 | Transmembrane emp24 domain-containing protein 9 | -3.695513133 | 0.117203187 |
| APOB | Apolipoprotein B-100 | -3.755814341 | 0.116507802 |
| RCN1 | Reticulocalbin-1 | -3.886012824 | 0.177613267 |
| MSMO1 | Methylsterol monooxygenase 1 | -3.908766068 | 0.122990843 |
| ERGIC1 | Endoplasmic reticulum-Golgi intermediate compartment protein 1 | -3.983569153 | 0.16915912 |
| PRKCSH | Glucosidase 2 subunit beta | -4.243521518 | 0.11803634 |
| CALU | Calumenin | -4.260981673 | 0.099632858 |
| PGRMC2 | Membrane-associated progesterone receptor component 2 | -4.287213473 | 0.110921496 |
| PDIA6 | Protein disulfide-isomerase A6 | -4.502365264 | 0.112097412 |
| HSD3B2 | 3 beta-hydroxysteroid dehydrogenase/Delta 5--4-isomerase type 2 | -4.983982318 | 0.117905077 |
| PDIA4 | Protein disulfide-isomerase A4 | -5.014119596 | 0.111408551 |

| **Supplementary Table 6. Functional enrichment analysis with g:Profiler of DAPs (FDR<0.2%) specific to MII ffEVs** | | | | |
| --- | --- | --- | --- | --- |
| **GO.ID** | **Description** | **p.Val** | **FDR** | **Genes** |
| GO:0016860 | intramolecular oxidoreductase activity | 5.62E-07 | 5.62E-07 | ECH1,ERP44,TMX1,PDIA6,PDIA3,PDIA4,HSD3B2 |
| GO:0016864 | intramolecular oxidoreductase activity, transposing S-S bonds | 4.0545E-06 | 4.0545E-06 | ERP44,TMX1,PDIA6,PDIA3,PDIA4 |
| GO:0003756 | protein disulfide isomerase activity | 4.0545E-06 | 4.0545E-06 | ERP44,TMX1,PDIA6,PDIA3,PDIA4 |
| GO:0016491 | oxidoreductase activity | 0.00166047 | 0.00166047 | P3H1,TMX1,CYB5R3,PDIA6,PDIA3,ALDH3A2,PDIA4,CYP17A1,SCD,PRDX4,CYP51A1,MSMO1,HSD3B2 |
| GO:0016853 | isomerase activity | 0.00201623 | 0.00201623 | ECH1,ERP44,TMX1,PDIA6,PDIA3,PDIA4,HSD3B2 |
| GO:0015035 | protein-disulfide reductase activity | 0.00395551 | 0.00395551 | TMX1,PDIA6,PDIA3,PDIA4 |
| GO:0015036 | disulfide oxidoreductase activity | 0.00781978 | 0.00781978 | TMX1,PDIA6,PDIA3,PDIA4 |
| GO:0016667 | oxidoreductase activity, acting on a sulfur group of donors | 0.02884466 | 0.02884466 | TMX1,PDIA6,PDIA3,PDIA4 |
| GO:0051082 | unfolded protein binding | 0.03746717 | 0.03746717 | HSP90B1,HSPA5,UGGT1,LMAN1,CANX |
| GO:0006457 | protein folding | 1.45E-09 | 1.45E-09 | P3H1,ERP44,HSP90B1,HSPA5,B2M,UGGT1,PDIA6,ERP29,PDIA3,PDIA4,LMAN1,CANX,PRDX4 |
| GO:0051604 | protein maturation | 1.2173E-06 | 1.2173E-06 | F12,SEC11A,P3H1,ERP44,HSP90B1,HSPA5,B2M,UGGT1,PDIA6,ERP29,PDIA3,PDIA4,LMAN1,CANX,PRDX4 |
| GO:1902653 | secondary alcohol biosynthetic process | 0.00007275 | 0.00007275 | CYB5R3,ERLIN2,FDFT1,APOB,CYP51A1,MSMO1 |
| GO:0006695 | cholesterol biosynthetic process | 0.00007275 | 0.00007275 | CYB5R3,ERLIN2,FDFT1,APOB,CYP51A1,MSMO1 |
| GO:0034975 | protein folding in endoplasmic reticulum | 0.00009288 | 0.00009288 | HSP90B1,HSPA5,PDIA3,CANX |
| GO:0016126 | sterol biosynthetic process | 0.00015335 | 0.00015335 | CYB5R3,ERLIN2,FDFT1,APOB,CYP51A1,MSMO1 |
| GO:0034976 | response to endoplasmic reticulum stress | 0.00016909 | 0.00016909 | ERP44,TMX1,HSP90B1,HSPA5,PDIA6,ERP29,PDIA3,PDIA4,ERMP1 |
| GO:0006694 | steroid biosynthetic process | 0.00046986 | 0.00046986 | CYB5R3,ERLIN2,FDFT1,CYP17A1,APOB,CYP51A1,MSMO1,HSD3B2 |
| GO:0015031 | protein transport | 0.0007684 | 0.0007684 | RAB18,P3H1,MVP,HSP90B1,HSPA5,TMED9,LMAN2,CD63,SURF4,ERP29,PDIA4,LMAN1,SEC22B,CANX,APOB,CYP51A1,ANG,GAL,SFRP1 |
| GO:0071705 | nitrogen compound transport | 0.00393295 | 0.00393295 | RAB18,P3H1,MVP,HSP90B1,HSPA5,TMED9,LMAN2,CD63,COMT,SURF4,ERP29,PDIA4,LMAN1,SEC22B,CANX,APOB,CYP51A1,PGRMC2,ANG,GAL,SFRP1 |
| GO:0008202 | steroid metabolic process | 0.00443889 | 0.00443889 | CYB5R3,ERLIN2,FDFT1,CYP17A1,APOB,CYP51A1,MSMO1,HSD3B2,GAL |
| GO:0071702 | organic substance transport | 0.00852642 | 0.00852642 | FABP5,RAB18,P3H1,MVP,SLC2A1,HSP90B1,HSPA5,TMED9,LMAN2,CD63,COMT,SURF4,ERP29,PDIA4,LMAN1,SEC22B,CANX,APOB,CYP51A1,PGRMC2,ANG,GAL,SFRP1 |
| GO:0046165 | alcohol biosynthetic process | 0.01684895 | 0.01684895 | CYB5R3,ERLIN2,FDFT1,APOB,CYP51A1,MSMO1 |
| GO:0045184 | establishment of protein localization | 0.01710627 | 0.01710627 | RAB18,P3H1,MVP,HSP90B1,HSPA5,TMED9,LMAN2,SSR1,CD63,SURF4,ERP29,PDIA4,LMAN1,SEC22B,CANX,APOB,CYP51A1,ANG,GAL,SFRP1 |
| GO:0008203 | cholesterol metabolic process | 0.01833062 | 0.01833062 | CYB5R3,ERLIN2,FDFT1,APOB,CYP51A1,MSMO1 |
| GO:1902652 | secondary alcohol metabolic process | 0.028486 | 0.028486 | CYB5R3,ERLIN2,FDFT1,APOB,CYP51A1,MSMO1 |
| GO:0007596 | blood coagulation | 0.03276612 | 0.03276612 | F12,SERPINA10,F11R,TMX1,VWF,PDIA3,LMAN1 |
| GO:0016125 | sterol metabolic process | 0.03314598 | 0.03314598 | CYB5R3,ERLIN2,FDFT1,APOB,CYP51A1,MSMO1 |
| GO:0050817 | coagulation | 0.0376792 | 0.0376792 | F12,SERPINA10,F11R,TMX1,VWF,PDIA3,LMAN1 |
| GO:0007599 | hemostasis | 0.03980764 | 0.03980764 | F12,SERPINA10,F11R,TMX1,VWF,PDIA3,LMAN1 |
| GO:0006629 | lipid metabolic process | 0.0477296 | 0.0477296 | FABP5,ECH1,AADAC,COMT,CYB5R3,ERLIN2,FDFT1,ALDH3A2,CYP17A1,SCD,APOB,CYP51A1,MSMO1,HSD3B2,ANG,GAL |
| GO:0005783 | endoplasmic reticulum | 2.24E-26 | 2.24E-26 | F12,SERPINA10,RAB18,RPL6,SEC11A,P3H1,ERP44,ITGA5,TMX1,GANAB,HSP90B1,HSPA5,B2M,AADAC,TMED9,LMAN2,SSR1,MYDGF,UGGT1,CYB5R3,SURF4,DAD1,ERLIN2,RCN1,PDIA6,ERP29,VWF,PDIA3,FDFT1,ALDH3A2,PDIA4,LMAN1,CYP17A1,SEC22B,CANX,ERMP1,SCD,PRDX4,APOB,ERGIC1,CYP51A1,CALU,PGRMC2,PRKCSH,MSMO1,HSD3B2 |
| GO:0012505 | endomembrane system | 4.14E-20 | 4.14E-20 | FABP5,S100A9,F12,SERPINA10,CFD,RAB18,ATP2B1,RPL6,SEC11A,P3H1,MVP,ERP44,SLC2A1,ITGA5,TMX1,GANAB,HSP90B1,IGF2R,HSPA5,B2M,AADAC,TMED9,LMAN2,SSR1,CD63,MYDGF,UGGT1,CYB5R3,SURF4,DAD1,ERLIN2,RCN1,PDIA6,ERP29,VWF,PDIA3,FDFT1,ALDH3A2,PDIA4,LMAN1,CYP17A1,SEC22B,CANX,ERMP1,SCD,PRDX4,APOB,ERGIC1,CYP51A1,CALU,PGRMC2,PRKCSH,MSMO1,HSD3B2,IGF2,GAL |
| GO:0005789 | endoplasmic reticulum membrane | 1.18E-17 | 1.18E-17 | RAB18,SEC11A,ERP44,TMX1,HSP90B1,HSPA5,B2M,AADAC,TMED9,LMAN2,SSR1,CYB5R3,SURF4,DAD1,ERLIN2,PDIA6,PDIA3,FDFT1,ALDH3A2,LMAN1,CYP17A1,SEC22B,CANX,ERMP1,SCD,APOB,ERGIC1,CYP51A1,CALU,MSMO1,HSD3B2 |
| GO:0098827 | endoplasmic reticulum subcompartment | 1.37E-17 | 1.37E-17 | RAB18,SEC11A,ERP44,TMX1,HSP90B1,HSPA5,B2M,AADAC,TMED9,LMAN2,SSR1,CYB5R3,SURF4,DAD1,ERLIN2,PDIA6,PDIA3,FDFT1,ALDH3A2,LMAN1,CYP17A1,SEC22B,CANX,ERMP1,SCD,APOB,ERGIC1,CYP51A1,CALU,MSMO1,HSD3B2 |
| GO:0042175 | nuclear outer membrane-endoplasmic reticulum membrane network | 2.06E-17 | 2.06E-17 | RAB18,SEC11A,ERP44,TMX1,HSP90B1,HSPA5,B2M,AADAC,TMED9,LMAN2,SSR1,CYB5R3,SURF4,DAD1,ERLIN2,PDIA6,PDIA3,FDFT1,ALDH3A2,LMAN1,CYP17A1,SEC22B,CANX,ERMP1,SCD,APOB,ERGIC1,CYP51A1,CALU,MSMO1,HSD3B2 |
| GO:0031982 | vesicle | 6.87E-17 | 6.87E-17 | FABP5,S100A9,F12,SERPINA10,IGKV1-5,CFD,RAB18,ECH1,F11R,ATP2B1,P3H1,MVP,ERP44,SLC2A1,ITGA5,GANAB,HSP90B1,IGF2R,HSPA5,B2M,TMED9,LMAN2,CD63,COMT,UGGT1,CYB5R3,SURF4,PDIA6,ERP29,VWF,PDIA3,PDIA4,LMAN1,SEC22B,CANX,PRDX4,APOB,ERGIC1,CALU,IGKV1-39,FREM2,ANG,SERPINA7,IGF2,GAL,SAA2,KRT33B,SFRP1,MGP |
| GO:0005576 | extracellular region | 7.56E-17 | 7.56E-17 | FABP5,S100A9,F12,SERPINA10,IGKV1-8,IGKV1-5,CFD,ECH1,F11R,ATP2B1,P3H1,MVP,ERP44,SLC2A1,IGHV3-38,GANAB,HSP90B1,IGF2R,HSPA5,B2M,TMED9,LMAN2,CD63,COMT,MYDGF,UGGT1,CYB5R3,PDIA6,VWF,PDIA3,PDIA4,LMAN1,CANX,PRDX4,APOB,CALU,IGKV1-39,FREM2,C1QC,C1QA,ANG,SERPINA7,IGF2,GAL,SAA2,KRT33B,SFRP1,MGP,KRT85,MDK |
| GO:0005615 | extracellular space | 1.19E-16 | 1.19E-16 | FABP5,S100A9,F12,SERPINA10,IGKV1-8,IGKV1-5,CFD,ECH1,F11R,ATP2B1,P3H1,MVP,ERP44,SLC2A1,GANAB,HSP90B1,IGF2R,HSPA5,B2M,TMED9,LMAN2,CD63,COMT,MYDGF,UGGT1,PDIA6,VWF,PDIA3,PDIA4,LMAN1,CANX,PRDX4,APOB,IGKV1-39,FREM2,C1QC,ANG,SERPINA7,IGF2,GAL,SAA2,KRT33B,SFRP1,MGP,KRT85 |
| GO:0070062 | extracellular exosome | 3.55E-16 | 3.55E-16 | FABP5,S100A9,F12,SERPINA10,IGKV1-5,CFD,ECH1,F11R,ATP2B1,P3H1,MVP,ERP44,SLC2A1,GANAB,HSP90B1,IGF2R,HSPA5,B2M,TMED9,LMAN2,CD63,COMT,UGGT1,PDIA6,VWF,PDIA3,LMAN1,CANX,PRDX4,APOB,IGKV1-39,FREM2,SERPINA7,SAA2,KRT33B,SFRP1,MGP |
| GO:1903561 | extracellular vesicle | 5.17E-16 | 5.17E-16 | FABP5,S100A9,F12,SERPINA10,IGKV1-5,CFD,ECH1,F11R,ATP2B1,P3H1,MVP,ERP44,SLC2A1,GANAB,HSP90B1,IGF2R,HSPA5,B2M,TMED9,LMAN2,CD63,COMT,UGGT1,PDIA6,VWF,PDIA3,LMAN1,CANX,PRDX4,APOB,IGKV1-39,FREM2,SERPINA7,SAA2,KRT33B,SFRP1,MGP |
| GO:0065010 | extracellular membrane-bounded organelle | 5.26E-16 | 5.26E-16 | FABP5,S100A9,F12,SERPINA10,IGKV1-5,CFD,ECH1,F11R,ATP2B1,P3H1,MVP,ERP44,SLC2A1,GANAB,HSP90B1,IGF2R,HSPA5,B2M,TMED9,LMAN2,CD63,COMT,UGGT1,PDIA6,VWF,PDIA3,LMAN1,CANX,PRDX4,APOB,IGKV1-39,FREM2,SERPINA7,SAA2,KRT33B,SFRP1,MGP |
| GO:0043230 | extracellular organelle | 5.26E-16 | 5.26E-16 | FABP5,S100A9,F12,SERPINA10,IGKV1-5,CFD,ECH1,F11R,ATP2B1,P3H1,MVP,ERP44,SLC2A1,GANAB,HSP90B1,IGF2R,HSPA5,B2M,TMED9,LMAN2,CD63,COMT,UGGT1,PDIA6,VWF,PDIA3,LMAN1,CANX,PRDX4,APOB,IGKV1-39,FREM2,SERPINA7,SAA2,KRT33B,SFRP1,MGP |
| GO:0031984 | organelle subcompartment | 1.19E-15 | 1.19E-15 | RAB18,SEC11A,ERP44,TMX1,HSP90B1,IGF2R,HSPA5,B2M,AADAC,TMED9,LMAN2,SSR1,CYB5R3,SURF4,DAD1,ERLIN2,PDIA6,PDIA3,FDFT1,ALDH3A2,LMAN1,CYP17A1,SEC22B,CANX,ERMP1,SCD,APOB,ERGIC1,CYP51A1,CALU,MSMO1,HSD3B2 |
| GO:0005788 | endoplasmic reticulum lumen | 4.39E-15 | 4.39E-15 | SERPINA10,P3H1,ERP44,GANAB,HSP90B1,HSPA5,B2M,MYDGF,UGGT1,RCN1,PDIA6,ERP29,PDIA3,PDIA4,CANX,APOB,CALU,PRKCSH |
| GO:0042470 | melanosome | 8.27E-13 | 8.27E-13 | SLC2A1,GANAB,HSP90B1,HSPA5,CD63,PDIA6,ERP29,PDIA3,PDIA4,SEC22B,CANX,CALU |
| GO:0048770 | pigment granule | 8.27E-13 | 8.27E-13 | SLC2A1,GANAB,HSP90B1,HSPA5,CD63,PDIA6,ERP29,PDIA3,PDIA4,SEC22B,CANX,CALU |
| GO:0031410 | cytoplasmic vesicle | 4.23E-11 | 4.23E-11 | FABP5,S100A9,CFD,RAB18,ATP2B1,MVP,ERP44,SLC2A1,ITGA5,GANAB,HSP90B1,IGF2R,HSPA5,B2M,TMED9,LMAN2,CD63,CYB5R3,SURF4,PDIA6,ERP29,VWF,PDIA3,PDIA4,LMAN1,SEC22B,CANX,PRDX4,APOB,ERGIC1,CALU,ANG,IGF2,GAL |
| GO:0097708 | intracellular vesicle | 4.48E-11 | 4.48E-11 | FABP5,S100A9,CFD,RAB18,ATP2B1,MVP,ERP44,SLC2A1,ITGA5,GANAB,HSP90B1,IGF2R,HSPA5,B2M,TMED9,LMAN2,CD63,CYB5R3,SURF4,PDIA6,ERP29,VWF,PDIA3,PDIA4,LMAN1,SEC22B,CANX,PRDX4,APOB,ERGIC1,CALU,ANG,IGF2,GAL |
| GO:0005793 | endoplasmic reticulum-Golgi intermediate compartment | 3.28E-10 | 3.28E-10 | ERP44,HSPA5,TMED9,LMAN2,MYDGF,UGGT1,SURF4,PDIA6,LMAN1,SEC22B,ERGIC1 |
| GO:0140534 | endoplasmic reticulum protein-containing complex | 1.28E-07 | 1.28E-07 | SEC11A,GANAB,HSP90B1,HSPA5,B2M,DAD1,PDIA6,PDIA3,PRKCSH |
| GO:0031090 | organelle membrane | 1.72E-07 | 1.72E-07 | FABP5,RAB18,ATP2B1,SEC11A,ERP44,SLC2A1,TMX1,HSP90B1,IGF2R,HSPA5,B2M,AADAC,TMED9,LMAN2,SSR1,CD63,CYB5R3,SURF4,DAD1,ERLIN2,PDIA6,PDIA3,FDFT1,ALDH3A2,LMAN1,CYP17A1,SEC22B,CANX,ERMP1,SCD,APOB,ERGIC1,CYP51A1,CALU,MSMO1,HSD3B2 |
| GO:0062023 | collagen-containing extracellular matrix | 5.19E-07 | 5.19E-07 | S100A9,F12,P3H1,HSP90B1,VWF,LMAN1,FREM2,C1QC,C1QA,ANG,SFRP1,MGP,MDK |
| GO:0031012 | extracellular matrix | 0.000012341 | 0.000012341 | S100A9,F12,P3H1,HSP90B1,VWF,LMAN1,FREM2,C1QC,C1QA,ANG,SFRP1,MGP,MDK |
| GO:0030312 | external encapsulating structure | 0.000012603 | 0.000012603 | S100A9,F12,P3H1,HSP90B1,VWF,LMAN1,FREM2,C1QC,C1QA,ANG,SFRP1,MGP,MDK |
| GO:0099503 | secretory vesicle | 0.000023506 | 0.000023506 | FABP5,S100A9,CFD,RAB18,ATP2B1,MVP,ERP44,IGF2R,B2M,TMED9,CD63,CYB5R3,SURF4,VWF,PRDX4,IGF2,GAL |
| GO:0034774 | secretory granule lumen | 0.000037387 | 0.000037387 | FABP5,S100A9,CFD,MVP,ERP44,B2M,CYB5R3,VWF,PRDX4,IGF2 |
| GO:0060205 | cytoplasmic vesicle lumen | 0.000040768 | 0.000040768 | FABP5,S100A9,CFD,MVP,ERP44,B2M,CYB5R3,VWF,PRDX4,IGF2 |
| GO:0031983 | vesicle lumen | 0.000041953 | 0.000041953 | FABP5,S100A9,CFD,MVP,ERP44,B2M,CYB5R3,VWF,PRDX4,IGF2 |
| GO:0030141 | secretory granule | 0.000080783 | 0.000080783 | FABP5,S100A9,CFD,RAB18,MVP,ERP44,IGF2R,B2M,CD63,CYB5R3,SURF4,VWF,PRDX4,IGF2,GAL |
| GO:0033116 | endoplasmic reticulum-Golgi intermediate compartment membrane | 0.000097303 | 0.000097303 | TMED9,LMAN2,SURF4,LMAN1,SEC22B,ERGIC1 |
| GO:0016020 | membrane | 0.00012495 | 0.00012495 | FABP5,S100A9,F12,IGKV1-8,IGKV1-5,RAB18,ECH1,F11R,ATP2B1,RPL6,SEC11A,P3H1,MVP,ERP44,SLC2A1,ITGA5,TMX1,IGHV3-38,GANAB,HSP90B1,IGF2R,HSPA5,B2M,APMAP,AADAC,TMED9,LMAN2,SSR1,CD63,COMT,CYB5R3,SURF4,DAD1,ERLIN2,PDIA6,ERP29,PDIA3,FDFT1,ALDH3A2,LMAN1,CYP17A1,SEC22B,CANX,ERMP1,SCD,APOB,ERGIC1,CYP51A1,CALU,PGRMC2,MSMO1,HSD3B2,IGKV1-39,FREM2,SFRP1 |
| GO:0030134 | COPII-coated ER to Golgi transport vesicle | 0.00017053 | 0.00017053 | B2M,TMED9,LMAN2,LMAN1,SEC22B,ERGIC1 |
| GO:0005737 | cytoplasm | 0.00024408 | 0.00024408 | FABP5,S100A9,F12,SERPINA10,CFD,RAB18,ECH1,ATP2B1,RPL6,SEC11A,P3H1,MVP,ERP44,SLC2A1,ITGA5,TMX1,GANAB,HSP90B1,IGF2R,HSPA5,B2M,AADAC,TMED9,LMAN2,SSR1,CD63,COMT,MYDGF,UGGT1,CYB5R3,SURF4,DAD1,ERLIN2,RCN1,PDIA6,ERP29,VWF,PDIA3,FDFT1,ALDH3A2,PDIA4,LMAN1,CYP17A1,SEC22B,CANX,ERMP1,SCD,PRDX4,APOB,ERGIC1,CYP51A1,CALU,PGRMC2,PRKCSH,MSMO1,HSD3B2,ANG,IGF2,GAL,KRT33B,SFRP1,KRT85 |
| GO:0030135 | coated vesicle | 0.00032434 | 0.00032434 | IGF2R,B2M,TMED9,LMAN2,VWF,LMAN1,SEC22B,APOB,ERGIC1 |
| GO:0005790 | smooth endoplasmic reticulum | 0.00120448 | 0.00120448 | HSP90B1,ERP29,APOB,HSD3B2 |
| GO:0034663 | endoplasmic reticulum chaperone complex | 0.00178723 | 0.00178723 | HSP90B1,HSPA5,PDIA6 |
| GO:0005602 | complement component C1 complex | 0.00334324 | 0.00334324 | C1QC,C1QA |
| GO:0017177 | glucosidase II complex | 0.00334324 | 0.00334324 | GANAB,PRKCSH |
| GO:0062167 | complement component C1q complex | 0.00334324 | 0.00334324 | C1QC,C1QA |
| GO:0043227 | membrane-bounded organelle | 0.00933206 | 0.00933206 | FABP5,S100A9,F12,SERPINA10,IGKV1-5,CFD,RAB18,ECH1,F11R,ATP2B1,RPL6,SEC11A,P3H1,MVP,ERP44,SLC2A1,ITGA5,TMX1,GANAB,HSP90B1,IGF2R,HSPA5,B2M,AADAC,TMED9,LMAN2,SSR1,CD63,COMT,MYDGF,UGGT1,CYB5R3,SURF4,DAD1,ERLIN2,RCN1,PDIA6,ERP29,VWF,PDIA3,FDFT1,ALDH3A2,PDIA4,LMAN1,CYP17A1,SEC22B,CANX,ERMP1,SCD,PRDX4,APOB,ERGIC1,CYP51A1,CALU,PGRMC2,PRKCSH,MSMO1,HSD3B2,IGKV1-39,FREM2,ANG,SERPINA7,IGF2,GAL,SAA2,KRT33B,SFRP1,MGP |
| GO:0009986 | cell surface | 0.01444964 | 0.01444964 | ERP44,ITGA5,IGF2R,HSPA5,B2M,APMAP,LMAN2,CD63,ERP29,PDIA3,PDIA4,SFRP1 |
| GO:0030133 | transport vesicle | 0.03158853 | 0.03158853 | ATP2B1,IGF2R,B2M,TMED9,SURF4,ERP29,LMAN1,SEC22B |
| KEGG:04141 | Protein processing in endoplasmic reticulum | 3.31E-11 | 3.31E-11 | GANAB,HSP90B1,HSPA5,LMAN2,SSR1,UGGT1,DAD1,PDIA6,ERP29,PDIA3,PDIA4,LMAN1,CANX,PRKCSH |
| KEGG:04918 | Thyroid hormone synthesis | 0.00580012 | 0.00580012 | HSP90B1,HSPA5,PDIA4,CANX,SERPINA7 |
| KEGG:04610 | Complement and coagulation cascades | 0.01114795 | 0.01114795 | F12,CFD,VWF,C1QC,C1QA |
| KEGG:00100 | Steroid biosynthesis | 0.01618871 | 0.01618871 | FDFT1,CYP51A1,MSMO1 |
| REAC:R-HSA-901042 | Calnexin/calreticulin cycle | 0.00014062 | 0.00014062 | GANAB,UGGT1,PDIA3,CANX,PRKCSH |
| REAC:R-HSA-381426 | Regulation of Insulin-like Growth Factor (IGF) transport and uptake by Insulin-like Growth Factor Binding Proteins (IGFBPs) | 0.00022242 | 0.00022242 | SERPINA10,HSP90B1,RCN1,PDIA6,APOB,CALU,PRKCSH,IGF2 |
| REAC:R-HSA-532668 | N-glycan trimming in the ER and Calnexin/Calreticulin cycle | 0.00066666 | 0.00066666 | GANAB,UGGT1,PDIA3,CANX,PRKCSH |
| REAC:R-HSA-9683686 | Maturation of spike protein | 0.00077273 | 0.00077273 | GANAB,CANX,PRKCSH |
| REAC:R-HSA-8957275 | Post-translational protein phosphorylation | 0.00109535 | 0.00109535 | SERPINA10,HSP90B1,RCN1,PDIA6,APOB,CALU,PRKCSH |
| REAC:R-HSA-168249 | Innate Immune System | 0.00376363 | 0.00376363 | FABP5,S100A9,IGKV1-5,CFD,RAB18,MVP,ERP44,HSP90B1,IGF2R,B2M,CD63,CYB5R3,SURF4,PRDX4,APOB,PRKCSH,IGKV1-39,C1QC,C1QA |
| REAC:R-HSA-446203 | Asparagine N-linked glycosylation | 0.00380542 | 0.00380542 | GANAB,TMED9,LMAN2,UGGT1,DAD1,PDIA3,LMAN1,SEC22B,CANX,PRKCSH |
| REAC:R-HSA-6798695 | Neutrophil degranulation | 0.00643254 | 0.00643254 | FABP5,S100A9,CFD,RAB18,MVP,ERP44,IGF2R,B2M,CD63,CYB5R3,SURF4,PRDX4 |
| REAC:R-HSA-983170 | Antigen Presentation: Folding, assembly and peptide loading of class I MHC | 0.00918672 | 0.00918672 | HSPA5,B2M,PDIA3,CANX |
| REAC:R-HSA-9694548 | Maturation of spike protein | 0.02467183 | 0.02467183 | GANAB,DAD1,CANX,PRKCSH |
| REAC:R-HSA-166663 | Initial triggering of complement | 0.03160171 | 0.03160171 | IGKV1-5,CFD,IGKV1-39,C1QC,C1QA |
| REAC:R-HSA-114608 | Platelet degranulation | 0.03444771 | 0.03444771 | CFD,HSPA5,CD63,VWF,CALU,IGF2 |
| REAC:R-HSA-76005 | Response to elevated platelet cytosolic Ca2+ | 0.04281893 | 0.04281893 | CFD,HSPA5,CD63,VWF,CALU,IGF2 |
| REAC:R-HSA-109582 | Hemostasis | 0.04331972 | 0.04331972 | F12,IGKV1-5,CFD,F11R,ATP2B1,ITGA5,HSPA5,CD63,VWF,APOB,CALU,IGKV1-39,IGF2 |
| WP:WP558 | Complement and coagulation cascades | 0.00021643 | 0.00021643 | F12,CFD,VWF,LMAN1,C1QC,C1QA |
| WP:WP5193 | Cholesterol synthesis disorders | 0.00059745 | 0.00059745 | FDFT1,CYP51A1,MSMO1,HSD3B2 |
| WP:WP5333 | Enterocyte cholesterol metabolism | 0.01057984 | 0.01057984 | FDFT1,APOB,CYP51A1,MSMO1 |
| WP:WP197 | Cholesterol biosynthesis pathway | 0.01519397 | 0.01519397 | FDFT1,CYP51A1,MSMO1 |
| WP:WP5329 | Cholesterol biosynthesis pathway in hepatocytes | 0.02145843 | 0.02145843 | FDFT1,SCD,CYP51A1,MSMO1 |
| WP:WP5300 | TROP2 regulatory signaling | 0.02349525 | 0.02349525 | ITGA5,IGF2R,IGF2,MDK |
| WP:WP4718 | Cholesterol metabolism with Bloch and Kandutsch Russell pathways | 0.03044503 | 0.03044503 | FDFT1,SCD,CYP51A1,MSMO1 |
| WP:WP545 | Complement activation | 0.04985443 | 0.04985443 | CFD,C1QC,C1QA |

| **Supplementary Table 7. Identified proteins (4589) in single MII oocytes (n=10)** | | |
| --- | --- | --- |
| **protein_Id** | **description** | **nrPeptides** |
| A0A0B4J2B5 | Immunoglobulin heavy variable 3/OR16-9 (non-functional) (Fragment) OS=Homo sapiens OX=9606 GN=IGHV3OR16-9 PE=1 SV=1 | 3 |
| A8K2U0 | Alpha-2-macroglobulin-like protein 1 OS=Homo sapiens OX=9606 GN=A2ML1 PE=1 SV=3 | 22 |
| O00425 | Insulin-like growth factor 2 mRNA-binding protein 3 OS=Homo sapiens OX=9606 GN=IGF2BP3 PE=1 SV=2 | 28 |
| O14578 | Citron Rho-interacting kinase OS=Homo sapiens OX=9606 GN=CIT PE=1 SV=2 | 54 |
| O14732 | Inositol monophosphatase 2 OS=Homo sapiens OX=9606 GN=IMPA2 PE=1 SV=1 | 4 |
| O14818 | Proteasome subunit alpha type-7 OS=Homo sapiens OX=9606 GN=PSMA7 PE=1 SV=1 | 13 |
| O43548 | Protein-glutamine gamma-glutamyltransferase 5 OS=Homo sapiens OX=9606 GN=TGM5 PE=1 SV=4 | 9 |
| O43707 | Alpha-actinin-4 OS=Homo sapiens OX=9606 GN=ACTN4 PE=1 SV=2 | 55 |
| O43790 | Keratin, type II cuticular Hb6 OS=Homo sapiens OX=9606 GN=KRT86 PE=1 SV=1 | 27 |
| O60313 | Dynamin-like 120 kDa protein, mitochondrial OS=Homo sapiens OX=9606 GN=OPA1 PE=1 SV=3 | 60 |
| O60814;P57053;P58876;P62807;Q5QNW6;Q93079;Q99877;Q99879 | Histone H2B type 1-K OS=Homo sapiens OX=9606 GN=H2BC12 PE=1 SV=3 | 7 |
| O60911 | Cathepsin L2 OS=Homo sapiens OX=9606 GN=CTSV PE=1 SV=2 | 2 |
| O75131 | Copine-3 OS=Homo sapiens OX=9606 GN=CPNE3 PE=1 SV=1 | 10 |
| O75223 | Gamma-glutamylcyclotransferase OS=Homo sapiens OX=9606 GN=GGCT PE=1 SV=1 | 10 |
| O75342 | Arachidonate 12-lipoxygenase, 12R-type OS=Homo sapiens OX=9606 GN=ALOX12B PE=1 SV=1 | 15 |
| O75369 | Filamin-B OS=Homo sapiens OX=9606 GN=FLNB PE=1 SV=2 | 103 |
| O75390 | Citrate synthase, mitochondrial OS=Homo sapiens OX=9606 GN=CS PE=1 SV=2 | 19 |
| O75436 | Vacuolar protein sorting-associated protein 26A OS=Homo sapiens OX=9606 GN=VPS26A PE=1 SV=2 | 16 |
| O75556 | Mammaglobin-B OS=Homo sapiens OX=9606 GN=SCGB2A1 PE=1 SV=1 | 3 |
| O75608 | Acyl-protein thioesterase 1 OS=Homo sapiens OX=9606 GN=LYPLA1 PE=1 SV=1 | 8 |
| O75964 | ATP synthase subunit g, mitochondrial OS=Homo sapiens OX=9606 GN=ATP5MG PE=1 SV=3 | 7 |
| O76011 | Keratin, type I cuticular Ha4 OS=Homo sapiens OX=9606 GN=KRT34 PE=1 SV=2 | 11 |
| O95613 | Pericentrin OS=Homo sapiens OX=9606 GN=PCNT PE=1 SV=4 | 1 |
| O95721 | Synaptosomal-associated protein 29 OS=Homo sapiens OX=9606 GN=SNAP29 PE=1 SV=1 | 6 |
| O95757 | Heat shock 70 kDa protein 4L OS=Homo sapiens OX=9606 GN=HSPA4L PE=1 SV=3 | 40 |
| O95848 | Uridine diphosphate glucose pyrophosphatase NUDT14 OS=Homo sapiens OX=9606 GN=NUDT14 PE=1 SV=2 | 7 |
| P00338 | L-lactate dehydrogenase A chain OS=Homo sapiens OX=9606 GN=LDHA PE=1 SV=2 | 14 |
| P00441 | Superoxide dismutase [Cu-Zn] OS=Homo sapiens OX=9606 GN=SOD1 PE=1 SV=2 | 9 |
| P00491 | Purine nucleoside phosphorylase OS=Homo sapiens OX=9606 GN=PNP PE=1 SV=2 | 10 |
| P00505 | Aspartate aminotransferase, mitochondrial OS=Homo sapiens OX=9606 GN=GOT2 PE=1 SV=3 | 28 |
| P00558 | Phosphoglycerate kinase 1 OS=Homo sapiens OX=9606 GN=PGK1 PE=1 SV=3 | 35 |
| P01040 | Cystatin-A OS=Homo sapiens OX=9606 GN=CSTA PE=1 SV=1 | 12 |
| P01591 | Immunoglobulin J chain OS=Homo sapiens OX=9606 GN=JCHAIN PE=1 SV=4 | 2 |
| P01833 | Polymeric immunoglobulin receptor OS=Homo sapiens OX=9606 GN=PIGR PE=1 SV=4 | 7 |
| P01834 | Immunoglobulin kappa constant OS=Homo sapiens OX=9606 GN=IGKC PE=1 SV=2 | 7 |
| P01857 | Immunoglobulin heavy constant gamma 1 OS=Homo sapiens OX=9606 GN=IGHG1 PE=1 SV=1 | 11 |
| P01876 | Immunoglobulin heavy constant alpha 1 OS=Homo sapiens OX=9606 GN=IGHA1 PE=1 SV=2 | 12 |
| P02452 | Collagen alpha-1(I) chain OS=Homo sapiens OX=9606 GN=COL1A1 PE=1 SV=6 | 2 |
| P02533 | Keratin, type I cytoskeletal 14 OS=Homo sapiens OX=9606 GN=KRT14 PE=1 SV=4 | 54 |
| P02538 | Keratin, type II cytoskeletal 6A OS=Homo sapiens OX=9606 GN=KRT6A PE=1 SV=3 | 42 |
| P02545 | Prelamin-A/C OS=Homo sapiens OX=9606 GN=LMNA PE=1 SV=1 | 37 |
| P02647 | Apolipoprotein A-I OS=Homo sapiens OX=9606 GN=APOA1 PE=1 SV=1 | 20 |
| P02768 | Albumin OS=Homo sapiens OX=9606 GN=ALB PE=1 SV=2 | 83 |
| P02788 | Lactotransferrin OS=Homo sapiens OX=9606 GN=LTF PE=1 SV=6 | 24 |
| P02792 | Ferritin light chain OS=Homo sapiens OX=9606 GN=FTL PE=1 SV=2 | 12 |
| P04004 | Vitronectin OS=Homo sapiens OX=9606 GN=VTN PE=1 SV=1 | 12 |
| P04040 | Catalase OS=Homo sapiens OX=9606 GN=CAT PE=1 SV=3 | 23 |
| P04062 | Lysosomal acid glucosylceramidase OS=Homo sapiens OX=9606 GN=GBA1 PE=1 SV=3 | 4 |
| P04075 | Fructose-bisphosphate aldolase A OS=Homo sapiens OX=9606 GN=ALDOA PE=1 SV=2 | 26 |
| P04080 | Cystatin-B OS=Homo sapiens OX=9606 GN=CSTB PE=1 SV=2 | 6 |
| P04083 | Annexin A1 OS=Homo sapiens OX=9606 GN=ANXA1 PE=1 SV=2 | 8 |
| P04114 | Apolipoprotein B-100 OS=Homo sapiens OX=9606 GN=APOB PE=1 SV=2 | 128 |
| P04179 | Superoxide dismutase [Mn], mitochondrial OS=Homo sapiens OX=9606 GN=SOD2 PE=1 SV=3 | 12 |
| P04181 | Ornithine aminotransferase, mitochondrial OS=Homo sapiens OX=9606 GN=OAT PE=1 SV=1 | 29 |
| P04259 | Keratin, type II cytoskeletal 6B OS=Homo sapiens OX=9606 GN=KRT6B PE=1 SV=5 | 10 |
| P04264 | Keratin, type II cytoskeletal 1 OS=Homo sapiens OX=9606 GN=KRT1 PE=1 SV=6 | 63 |
| P04406 | Glyceraldehyde-3-phosphate dehydrogenase OS=Homo sapiens OX=9606 GN=GAPDH PE=1 SV=3 | 21 |
| P04792 | Heat shock protein beta-1 OS=Homo sapiens OX=9606 GN=HSPB1 PE=1 SV=2 | 22 |
| P04908;P0C0S8;P20671;Q16777;Q6FI13;Q7L7L0;Q93077;Q96KK5;Q99878;Q9BTM1 | Histone H2A type 1-B/E OS=Homo sapiens OX=9606 GN=H2AC8 PE=1 SV=2 | 5 |
| P05089 | Arginase-1 OS=Homo sapiens OX=9606 GN=ARG1 PE=1 SV=2 | 21 |
| P05090 | Apolipoprotein D OS=Homo sapiens OX=9606 GN=APOD PE=1 SV=1 | 7 |
| P05109 | Protein S100-A8 OS=Homo sapiens OX=9606 GN=S100A8 PE=1 SV=1 | 12 |
| P05120 | Plasminogen activator inhibitor 2 OS=Homo sapiens OX=9606 GN=SERPINB2 PE=1 SV=2 | 9 |
| P05141 | ADP/ATP translocase 2 OS=Homo sapiens OX=9606 GN=SLC25A5 PE=1 SV=7 | 25 |
| P05386 | 60S acidic ribosomal protein P1 OS=Homo sapiens OX=9606 GN=RPLP1 PE=1 SV=1 | 1 |
| P05387 | 60S acidic ribosomal protein P2 OS=Homo sapiens OX=9606 GN=RPLP2 PE=1 SV=1 | 4 |
| P06396 | Gelsolin OS=Homo sapiens OX=9606 GN=GSN PE=1 SV=1 | 33 |
| P06576 | ATP synthase subunit beta, mitochondrial OS=Homo sapiens OX=9606 GN=ATP5F1B PE=1 SV=3 | 27 |
| P06702 | Protein S100-A9 OS=Homo sapiens OX=9606 GN=S100A9 PE=1 SV=1 | 9 |
| P06703 | Protein S100-A6 OS=Homo sapiens OX=9606 GN=S100A6 PE=1 SV=1 | 1 |
| P06733 | Alpha-enolase OS=Homo sapiens OX=9606 GN=ENO1 PE=1 SV=2 | 31 |
| P06748 | Nucleophosmin OS=Homo sapiens OX=9606 GN=NPM1 PE=1 SV=2 | 9 |
| P06753 | Tropomyosin alpha-3 chain OS=Homo sapiens OX=9606 GN=TPM3 PE=1 SV=2 | 12 |
| P06865 | Beta-hexosaminidase subunit alpha OS=Homo sapiens OX=9606 GN=HEXA PE=1 SV=2 | 6 |
| P07195 | L-lactate dehydrogenase B chain OS=Homo sapiens OX=9606 GN=LDHB PE=1 SV=2 | 7 |
| P07237 | Protein disulfide-isomerase OS=Homo sapiens OX=9606 GN=P4HB PE=1 SV=3 | 36 |
| P07339 | Cathepsin D OS=Homo sapiens OX=9606 GN=CTSD PE=1 SV=1 | 16 |
| P07355 | Annexin A2 OS=Homo sapiens OX=9606 GN=ANXA2 PE=1 SV=2 | 28 |
| P07384 | Calpain-1 catalytic subunit OS=Homo sapiens OX=9606 GN=CAPN1 PE=1 SV=1 | 15 |
| P07437 | Tubulin beta chain OS=Homo sapiens OX=9606 GN=TUBB PE=1 SV=2 | 10 |
| P07476 | Involucrin OS=Homo sapiens OX=9606 GN=IVL PE=1 SV=2 | 4 |
| P07686 | Beta-hexosaminidase subunit beta OS=Homo sapiens OX=9606 GN=HEXB PE=1 SV=4 | 16 |
| P07737 | Profilin-1 OS=Homo sapiens OX=9606 GN=PFN1 PE=1 SV=2 | 15 |
| P07900 | Heat shock protein HSP 90-alpha OS=Homo sapiens OX=9606 GN=HSP90AA1 PE=1 SV=5 | 59 |
| P07910 | Heterogeneous nuclear ribonucleoproteins C1/C2 OS=Homo sapiens OX=9606 GN=HNRNPC PE=1 SV=4 | 3 |
| P07954 | Fumarate hydratase, mitochondrial OS=Homo sapiens OX=9606 GN=FH PE=1 SV=3 | 24 |
| P08123 | Collagen alpha-2(I) chain OS=Homo sapiens OX=9606 GN=COL1A2 PE=1 SV=7 | 1 |
| P08670 | Vimentin OS=Homo sapiens OX=9606 GN=VIM PE=1 SV=4 | 6 |
| P08727 | Keratin, type I cytoskeletal 19 OS=Homo sapiens OX=9606 GN=KRT19 PE=1 SV=4 | 5 |
| P08779 | Keratin, type I cytoskeletal 16 OS=Homo sapiens OX=9606 GN=KRT16 PE=1 SV=4 | 34 |
| P08842 | Steryl-sulfatase OS=Homo sapiens OX=9606 GN=STS PE=1 SV=2 | 2 |
| P08865 | 40S ribosomal protein SA OS=Homo sapiens OX=9606 GN=RPSA PE=1 SV=4 | 9 |
| P09211 | Glutathione S-transferase P OS=Homo sapiens OX=9606 GN=GSTP1 PE=1 SV=2 | 14 |
| P09525 | Annexin A4 OS=Homo sapiens OX=9606 GN=ANXA4 PE=1 SV=4 | 11 |
| P09543 | 2',3'-cyclic-nucleotide 3'-phosphodiesterase OS=Homo sapiens OX=9606 GN=CNP PE=1 SV=2 | 12 |
| P09668 | Pro-cathepsin H OS=Homo sapiens OX=9606 GN=CTSH PE=1 SV=4 | 2 |
| P09936 | Ubiquitin carboxyl-terminal hydrolase isozyme L1 OS=Homo sapiens OX=9606 GN=UCHL1 PE=1 SV=2 | 21 |
| P0DMV8;P0DMV9 | Heat shock 70 kDa protein 1A OS=Homo sapiens OX=9606 GN=HSPA1A PE=1 SV=1 | 38 |
| P0DOY2;P0DOY3 | Immunoglobulin lambda constant 2 OS=Homo sapiens OX=9606 GN=IGLC2 PE=1 SV=1 | 4 |
| P0DP23;P0DP24;P0DP25 | Calmodulin-1 OS=Homo sapiens OX=9606 GN=CALM1 PE=1 SV=1 | 6 |
| P10412;P16402;P16403 | Histone H1.4 OS=Homo sapiens OX=9606 GN=H1-4 PE=1 SV=2 | 2 |
| P10599 | Thioredoxin OS=Homo sapiens OX=9606 GN=TXN PE=1 SV=3 | 9 |
| P10619 | Lysosomal protective protein OS=Homo sapiens OX=9606 GN=CTSA PE=1 SV=2 | 5 |
| P10809 | 60 kDa heat shock protein, mitochondrial OS=Homo sapiens OX=9606 GN=HSPD1 PE=1 SV=2 | 50 |
| P10909 | Clusterin OS=Homo sapiens OX=9606 GN=CLU PE=1 SV=1 | 11 |
| P11021 | Endoplasmic reticulum chaperone BiP OS=Homo sapiens OX=9606 GN=HSPA5 PE=1 SV=2 | 54 |
| P11142 | Heat shock cognate 71 kDa protein OS=Homo sapiens OX=9606 GN=HSPA8 PE=1 SV=1 | 45 |
| P11279 | Lysosome-associated membrane glycoprotein 1 OS=Homo sapiens OX=9606 GN=LAMP1 PE=1 SV=3 | 6 |
| P11413 | Glucose-6-phosphate 1-dehydrogenase OS=Homo sapiens OX=9606 GN=G6PD PE=1 SV=4 | 46 |
| P11586 | C-1-tetrahydrofolate synthase, cytoplasmic OS=Homo sapiens OX=9606 GN=MTHFD1 PE=1 SV=4 | 36 |
| P12035 | Keratin, type II cytoskeletal 3 OS=Homo sapiens OX=9606 GN=KRT3 PE=1 SV=3 | 7 |
| P12273 | Prolactin-inducible protein OS=Homo sapiens OX=9606 GN=PIP PE=1 SV=1 | 6 |
| P13639 | Elongation factor 2 OS=Homo sapiens OX=9606 GN=EEF2 PE=1 SV=4 | 48 |
| P13645 | Keratin, type I cytoskeletal 10 OS=Homo sapiens OX=9606 GN=KRT10 PE=1 SV=6 | 49 |
| P13646 | Keratin, type I cytoskeletal 13 OS=Homo sapiens OX=9606 GN=KRT13 PE=1 SV=4 | 22 |
| P13647 | Keratin, type II cytoskeletal 5 OS=Homo sapiens OX=9606 GN=KRT5 PE=1 SV=3 | 62 |
| P13667 | Protein disulfide-isomerase A4 OS=Homo sapiens OX=9606 GN=PDIA4 PE=1 SV=2 | 53 |
| P13798 | Acylamino-acid-releasing enzyme OS=Homo sapiens OX=9606 GN=APEH PE=1 SV=4 | 25 |
| P13804 | Electron transfer flavoprotein subunit alpha, mitochondrial OS=Homo sapiens OX=9606 GN=ETFA PE=1 SV=1 | 17 |
| P14550 | Aldo-keto reductase family 1 member A1 OS=Homo sapiens OX=9606 GN=AKR1A1 PE=1 SV=3 | 13 |
| P14618 | Pyruvate kinase PKM OS=Homo sapiens OX=9606 GN=PKM PE=1 SV=4 | 39 |
| P14625 | Endoplasmin OS=Homo sapiens OX=9606 GN=HSP90B1 PE=1 SV=1 | 63 |
| P14735 | Insulin-degrading enzyme OS=Homo sapiens OX=9606 GN=IDE PE=1 SV=4 | 46 |
| P14923 | Junction plakoglobin OS=Homo sapiens OX=9606 GN=JUP PE=1 SV=3 | 42 |
| P15121 | Aldo-keto reductase family 1 member B1 OS=Homo sapiens OX=9606 GN=AKR1B1 PE=1 SV=3 | 22 |
| P15311 | Ezrin OS=Homo sapiens OX=9606 GN=EZR PE=1 SV=4 | 50 |
| P15531 | Nucleoside diphosphate kinase A OS=Homo sapiens OX=9606 GN=NME1 PE=1 SV=1 | 10 |
| P15924 | Desmoplakin OS=Homo sapiens OX=9606 GN=DSP PE=1 SV=3 | 241 |
| P16278 | Beta-galactosidase OS=Homo sapiens OX=9606 GN=GLB1 PE=1 SV=2 | 13 |
| P17174 | Aspartate aminotransferase, cytoplasmic OS=Homo sapiens OX=9606 GN=GOT1 PE=1 SV=3 | 19 |
| P17900 | Ganglioside GM2 activator OS=Homo sapiens OX=9606 GN=GM2A PE=1 SV=4 | 6 |
| P17987 | T-complex protein 1 subunit alpha OS=Homo sapiens OX=9606 GN=TCP1 PE=1 SV=1 | 31 |
| P18206 | Vinculin OS=Homo sapiens OX=9606 GN=VCL PE=1 SV=4 | 44 |
| P18583 | Protein SON OS=Homo sapiens OX=9606 GN=SON PE=1 SV=4 | 1 |
| P19013 | Keratin, type II cytoskeletal 4 OS=Homo sapiens OX=9606 GN=KRT4 PE=1 SV=5 | 23 |
| P20073 | Annexin A7 OS=Homo sapiens OX=9606 GN=ANXA7 PE=1 SV=3 | 26 |
| P20618 | Proteasome subunit beta type-1 OS=Homo sapiens OX=9606 GN=PSMB1 PE=1 SV=2 | 7 |
| P20930 | Filaggrin OS=Homo sapiens OX=9606 GN=FLG PE=1 SV=3 | 18 |
| P21333 | Filamin-A OS=Homo sapiens OX=9606 GN=FLNA PE=1 SV=4 | 93 |
| P21754 | Zona pellucida sperm-binding protein 3 OS=Homo sapiens OX=9606 GN=ZP3 PE=1 SV=2 | 21 |
| P22061 | Protein-L-isoaspartate(D-aspartate) O-methyltransferase OS=Homo sapiens OX=9606 GN=PCMT1 PE=1 SV=4 | 15 |
| P22234 | Bifunctional phosphoribosylaminoimidazole carboxylase/phosphoribosylaminoimidazole succinocarboxamide synthetase OS=Homo sapiens OX=9606 GN=PAICS PE=1 SV=3 | 15 |
| P22528 | Cornifin-B OS=Homo sapiens OX=9606 GN=SPRR1B PE=1 SV=2 | 5 |
| P22735 | Protein-glutamine gamma-glutamyltransferase K OS=Homo sapiens OX=9606 GN=TGM1 PE=1 SV=4 | 31 |
| P23284 | Peptidyl-prolyl cis-trans isomerase B OS=Homo sapiens OX=9606 GN=PPIB PE=1 SV=2 | 15 |
| P23490 | Loricrin OS=Homo sapiens OX=9606 GN=LORICRIN PE=1 SV=2 | 4 |
| P23526 | Adenosylhomocysteinase OS=Homo sapiens OX=9606 GN=AHCY PE=1 SV=4 | 28 |
| P23528 | Cofilin-1 OS=Homo sapiens OX=9606 GN=CFL1 PE=1 SV=3 | 16 |
| P24752 | Acetyl-CoA acetyltransferase, mitochondrial OS=Homo sapiens OX=9606 GN=ACAT1 PE=1 SV=1 | 28 |
| P25311 | Zinc-alpha-2-glycoprotein OS=Homo sapiens OX=9606 GN=AZGP1 PE=1 SV=2 | 18 |
| P25705 | ATP synthase subunit alpha, mitochondrial OS=Homo sapiens OX=9606 GN=ATP5F1A PE=1 SV=1 | 34 |
| P25786 | Proteasome subunit alpha type-1 OS=Homo sapiens OX=9606 GN=PSMA1 PE=1 SV=1 | 17 |
| P25787 | Proteasome subunit alpha type-2 OS=Homo sapiens OX=9606 GN=PSMA2 PE=1 SV=2 | 12 |
| P25788 | Proteasome subunit alpha type-3 OS=Homo sapiens OX=9606 GN=PSMA3 PE=1 SV=2 | 7 |
| P25789 | Proteasome subunit alpha type-4 OS=Homo sapiens OX=9606 GN=PSMA4 PE=1 SV=1 | 10 |
| P25815 | Protein S100-P OS=Homo sapiens OX=9606 GN=S100P PE=1 SV=2 | 2 |
| P26641 | Elongation factor 1-gamma OS=Homo sapiens OX=9606 GN=EEF1G PE=1 SV=3 | 19 |
| P26885 | Peptidyl-prolyl cis-trans isomerase FKBP2 OS=Homo sapiens OX=9606 GN=FKBP2 PE=1 SV=2 | 9 |
| P27482 | Calmodulin-like protein 3 OS=Homo sapiens OX=9606 GN=CALML3 PE=1 SV=2 | 6 |
| P27797 | Calreticulin OS=Homo sapiens OX=9606 GN=CALR PE=1 SV=1 | 17 |
| P27816 | Microtubule-associated protein 4 OS=Homo sapiens OX=9606 GN=MAP4 PE=1 SV=3 | 34 |
| P27824 | Calnexin OS=Homo sapiens OX=9606 GN=CANX PE=1 SV=2 | 20 |
| P28066 | Proteasome subunit alpha type-5 OS=Homo sapiens OX=9606 GN=PSMA5 PE=1 SV=3 | 9 |
| P28070 | Proteasome subunit beta type-4 OS=Homo sapiens OX=9606 GN=PSMB4 PE=1 SV=4 | 7 |
| P28072 | Proteasome subunit beta type-6 OS=Homo sapiens OX=9606 GN=PSMB6 PE=1 SV=4 | 7 |
| P28074 | Proteasome subunit beta type-5 OS=Homo sapiens OX=9606 GN=PSMB5 PE=1 SV=3 | 10 |
| P29401 | Transketolase OS=Homo sapiens OX=9606 GN=TKT PE=1 SV=3 | 23 |
| P29508 | Serpin B3 OS=Homo sapiens OX=9606 GN=SERPINB3 PE=1 SV=2 | 26 |
| P29692 | Elongation factor 1-delta OS=Homo sapiens OX=9606 GN=EEF1D PE=1 SV=5 | 15 |
| P30041 | Peroxiredoxin-6 OS=Homo sapiens OX=9606 GN=PRDX6 PE=1 SV=3 | 24 |
| P30050 | 60S ribosomal protein L12 OS=Homo sapiens OX=9606 GN=RPL12 PE=1 SV=1 | 5 |
| P30086 | Phosphatidylethanolamine-binding protein 1 OS=Homo sapiens OX=9606 GN=PEBP1 PE=1 SV=3 | 12 |
| P30101 | Protein disulfide-isomerase A3 OS=Homo sapiens OX=9606 GN=PDIA3 PE=1 SV=4 | 45 |
| P31025 | Lipocalin-1 OS=Homo sapiens OX=9606 GN=LCN1 PE=1 SV=1 | 3 |
| P31151 | Protein S100-A7 OS=Homo sapiens OX=9606 GN=S100A7 PE=1 SV=4 | 11 |
| P31943 | Heterogeneous nuclear ribonucleoprotein H OS=Homo sapiens OX=9606 GN=HNRNPH1 PE=1 SV=4 | 7 |
| P31944 | Caspase-14 OS=Homo sapiens OX=9606 GN=CASP14 PE=1 SV=2 | 24 |
| P31947 | 14-3-3 protein sigma OS=Homo sapiens OX=9606 GN=SFN PE=1 SV=1 | 6 |
| P31948 | Stress-induced-phosphoprotein 1 OS=Homo sapiens OX=9606 GN=STIP1 PE=1 SV=1 | 38 |
| P31949 | Protein S100-A11 OS=Homo sapiens OX=9606 GN=S100A11 PE=1 SV=2 | 3 |
| P32119 | Peroxiredoxin-2 OS=Homo sapiens OX=9606 GN=PRDX2 PE=1 SV=5 | 16 |
| P35232 | Prohibitin 1 OS=Homo sapiens OX=9606 GN=PHB1 PE=1 SV=1 | 21 |
| P35268 | 60S ribosomal protein L22 OS=Homo sapiens OX=9606 GN=RPL22 PE=1 SV=2 | 3 |
| P35270 | Sepiapterin reductase OS=Homo sapiens OX=9606 GN=SPR PE=1 SV=1 | 16 |
| P35527 | Keratin, type I cytoskeletal 9 OS=Homo sapiens OX=9606 GN=KRT9 PE=1 SV=3 | 49 |
| P35579 | Myosin-9 OS=Homo sapiens OX=9606 GN=MYH9 PE=1 SV=4 | 109 |
| P35908 | Keratin, type II cytoskeletal 2 epidermal OS=Homo sapiens OX=9606 GN=KRT2 PE=1 SV=2 | 52 |
| P36542 | ATP synthase subunit gamma, mitochondrial OS=Homo sapiens OX=9606 GN=ATP5F1C PE=1 SV=1 | 11 |
| P36957 | Dihydrolipoyllysine-residue succinyltransferase component of 2-oxoglutarate dehydrogenase complex, mitochondrial OS=Homo sapiens OX=9606 GN=DLST PE=1 SV=4 | 23 |
| P37802 | Transgelin-2 OS=Homo sapiens OX=9606 GN=TAGLN2 PE=1 SV=3 | 13 |
| P37837 | Transaldolase OS=Homo sapiens OX=9606 GN=TALDO1 PE=1 SV=2 | 15 |
| P38646 | Stress-70 protein, mitochondrial OS=Homo sapiens OX=9606 GN=HSPA9 PE=1 SV=2 | 60 |
| P40925 | Malate dehydrogenase, cytoplasmic OS=Homo sapiens OX=9606 GN=MDH1 PE=1 SV=4 | 17 |
| P40926 | Malate dehydrogenase, mitochondrial OS=Homo sapiens OX=9606 GN=MDH2 PE=1 SV=3 | 19 |
| P40939 | Trifunctional enzyme subunit alpha, mitochondrial OS=Homo sapiens OX=9606 GN=HADHA PE=1 SV=2 | 32 |
| P42166 | Lamina-associated polypeptide 2, isoform alpha OS=Homo sapiens OX=9606 GN=TMPO PE=1 SV=2 | 5 |
| P42357 | Histidine ammonia-lyase OS=Homo sapiens OX=9606 GN=HAL PE=1 SV=1 | 15 |
| P42765 | 3-ketoacyl-CoA thiolase, mitochondrial OS=Homo sapiens OX=9606 GN=ACAA2 PE=1 SV=2 | 21 |
| P43307 | Translocon-associated protein subunit alpha OS=Homo sapiens OX=9606 GN=SSR1 PE=1 SV=3 | 4 |
| P45880 | Voltage-dependent anion-selective channel protein 2 OS=Homo sapiens OX=9606 GN=VDAC2 PE=1 SV=2 | 16 |
| P46459 | Vesicle-fusing ATPase OS=Homo sapiens OX=9606 GN=NSF PE=1 SV=3 | 55 |
| P47929 | Galectin-7 OS=Homo sapiens OX=9606 GN=LGALS7B PE=1 SV=2 | 11 |
| P48594 | Serpin B4 OS=Homo sapiens OX=9606 GN=SERPINB4 PE=1 SV=2 | 2 |
| P48681 | Nestin OS=Homo sapiens OX=9606 GN=NES PE=1 SV=2 | 1 |
| P49189 | 4-trimethylaminobutyraldehyde dehydrogenase OS=Homo sapiens OX=9606 GN=ALDH9A1 PE=1 SV=3 | 22 |
| P49411 | Elongation factor Tu, mitochondrial OS=Homo sapiens OX=9606 GN=TUFM PE=1 SV=3 | 30 |
| P49589 | Cysteine--tRNA ligase, cytoplasmic OS=Homo sapiens OX=9606 GN=CARS1 PE=1 SV=3 | 16 |
| P49720 | Proteasome subunit beta type-3 OS=Homo sapiens OX=9606 GN=PSMB3 PE=1 SV=2 | 6 |
| P49748 | Very long-chain specific acyl-CoA dehydrogenase, mitochondrial OS=Homo sapiens OX=9606 GN=ACADVL PE=1 SV=1 | 37 |
| P49862 | Kallikrein-7 OS=Homo sapiens OX=9606 GN=KLK7 PE=1 SV=1 | 7 |
| P50395 | Rab GDP dissociation inhibitor beta OS=Homo sapiens OX=9606 GN=GDI2 PE=1 SV=2 | 37 |
| P50452 | Serpin B8 OS=Homo sapiens OX=9606 GN=SERPINB8 PE=1 SV=2 | 8 |
| P50990 | T-complex protein 1 subunit theta OS=Homo sapiens OX=9606 GN=CCT8 PE=1 SV=4 | 45 |
| P51149 | Ras-related protein Rab-7a OS=Homo sapiens OX=9606 GN=RAB7A PE=1 SV=1 | 13 |
| P51571 | Translocon-associated protein subunit delta OS=Homo sapiens OX=9606 GN=SSR4 PE=1 SV=1 | 7 |
| P52209 | 6-phosphogluconate dehydrogenase, decarboxylating OS=Homo sapiens OX=9606 GN=PGD PE=1 SV=3 | 26 |
| P52907 | F-actin-capping protein subunit alpha-1 OS=Homo sapiens OX=9606 GN=CAPZA1 PE=1 SV=3 | 5 |
| P55072 | Transitional endoplasmic reticulum ATPase OS=Homo sapiens OX=9606 GN=VCP PE=1 SV=4 | 50 |
| P56537 | Eukaryotic translation initiation factor 6 OS=Homo sapiens OX=9606 GN=EIF6 PE=1 SV=1 | 3 |
| P58107 | Epiplakin OS=Homo sapiens OX=9606 GN=EPPK1 PE=1 SV=3 | 30 |
| P59047 | NACHT, LRR and PYD domains-containing protein 5 OS=Homo sapiens OX=9606 GN=NLRP5 PE=1 SV=2 | 71 |
| P59665;P59666 | Neutrophil defensin 1 OS=Homo sapiens OX=9606 GN=DEFA1B PE=1 SV=1 | 2 |
| P60174 | Triosephosphate isomerase OS=Homo sapiens OX=9606 GN=TPI1 PE=1 SV=4 | 18 |
| P60660 | Myosin light polypeptide 6 OS=Homo sapiens OX=9606 GN=MYL6 PE=1 SV=2 | 9 |
| P60842 | Eukaryotic initiation factor 4A-I OS=Homo sapiens OX=9606 GN=EIF4A1 PE=1 SV=1 | 22 |
| P60852 | Zona pellucida sperm-binding protein 1 OS=Homo sapiens OX=9606 GN=ZP1 PE=1 SV=1 | 24 |
| P60900 | Proteasome subunit alpha type-6 OS=Homo sapiens OX=9606 GN=PSMA6 PE=1 SV=1 | 14 |
| P61019 | Ras-related protein Rab-2A OS=Homo sapiens OX=9606 GN=RAB2A PE=1 SV=1 | 16 |
| P61106 | Ras-related protein Rab-14 OS=Homo sapiens OX=9606 GN=RAB14 PE=1 SV=4 | 7 |
| P61604 | 10 kDa heat shock protein, mitochondrial OS=Homo sapiens OX=9606 GN=HSPE1 PE=1 SV=2 | 15 |
| P61626 | Lysozyme C OS=Homo sapiens OX=9606 GN=LYZ PE=1 SV=1 | 10 |
| P61978 | Heterogeneous nuclear ribonucleoprotein K OS=Homo sapiens OX=9606 GN=HNRNPK PE=1 SV=1 | 9 |
| P62249 | 40S ribosomal protein S16 OS=Homo sapiens OX=9606 GN=RPS16 PE=1 SV=2 | 4 |
| P62258 | 14-3-3 protein epsilon OS=Homo sapiens OX=9606 GN=YWHAE PE=1 SV=1 | 23 |
| P62318 | Small nuclear ribonucleoprotein Sm D3 OS=Homo sapiens OX=9606 GN=SNRPD3 PE=1 SV=1 | 3 |
| P62753 | 40S ribosomal protein S6 OS=Homo sapiens OX=9606 GN=RPS6 PE=1 SV=1 | 10 |
| P62805 | Histone H4 OS=Homo sapiens OX=9606 GN=H4C16 PE=1 SV=2 | 11 |
| P62820 | Ras-related protein Rab-1A OS=Homo sapiens OX=9606 GN=RAB1A PE=1 SV=3 | 17 |
| P62857 | 40S ribosomal protein S28 OS=Homo sapiens OX=9606 GN=RPS28 PE=1 SV=1 | 4 |
| P62899 | 60S ribosomal protein L31 OS=Homo sapiens OX=9606 GN=RPL31 PE=1 SV=1 | 4 |
| P62937 | Peptidyl-prolyl cis-trans isomerase A OS=Homo sapiens OX=9606 GN=PPIA PE=1 SV=2 | 12 |
| P62979 | Ubiquitin-40S ribosomal protein S27a OS=Homo sapiens OX=9606 GN=RPS27A PE=1 SV=2 | 11 |
| P63261 | Actin, cytoplasmic 2 OS=Homo sapiens OX=9606 GN=ACTG1 PE=1 SV=1 | 33 |
| P68104;Q5VTE0 | Elongation factor 1-alpha 1 OS=Homo sapiens OX=9606 GN=EEF1A1 PE=1 SV=1 | 12 |
| P68133 | Actin, alpha skeletal muscle OS=Homo sapiens OX=9606 GN=ACTA1 PE=1 SV=1 | 3 |
| P68366 | Tubulin alpha-4A chain OS=Homo sapiens OX=9606 GN=TUBA4A PE=1 SV=1 | 27 |
| P68431;P84243;Q71DI3 | Histone H3.1 OS=Homo sapiens OX=9606 GN=H3C12 PE=1 SV=2 | 7 |
| P68871 | Hemoglobin subunit beta OS=Homo sapiens OX=9606 GN=HBB PE=1 SV=2 | 13 |
| P69905 | Hemoglobin subunit alpha OS=Homo sapiens OX=9606 GN=HBA2 PE=1 SV=2 | 5 |
| P78386 | Keratin, type II cuticular Hb5 OS=Homo sapiens OX=9606 GN=KRT85 PE=1 SV=1 | 16 |
| P78417 | Glutathione S-transferase omega-1 OS=Homo sapiens OX=9606 GN=GSTO1 PE=1 SV=2 | 6 |
| P78540 | Arginase-2, mitochondrial OS=Homo sapiens OX=9606 GN=ARG2 PE=1 SV=1 | 24 |
| P81605 | Dermcidin OS=Homo sapiens OX=9606 GN=DCD PE=1 SV=2 | 6 |
| Q00325 | Phosphate carrier protein, mitochondrial OS=Homo sapiens OX=9606 GN=SLC25A3 PE=1 SV=2 | 17 |
| Q01469 | Fatty acid-binding protein 5 OS=Homo sapiens OX=9606 GN=FABP5 PE=1 SV=3 | 12 |
| Q01518 | Adenylyl cyclase-associated protein 1 OS=Homo sapiens OX=9606 GN=CAP1 PE=1 SV=5 | 27 |
| Q02413 | Desmoglein-1 OS=Homo sapiens OX=9606 GN=DSG1 PE=1 SV=2 | 36 |
| Q03001 | Dystonin OS=Homo sapiens OX=9606 GN=DST PE=1 SV=4 | 1 |
| Q04695 | Keratin, type I cytoskeletal 17 OS=Homo sapiens OX=9606 GN=KRT17 PE=1 SV=2 | 30 |
| Q05639 | Elongation factor 1-alpha 2 OS=Homo sapiens OX=9606 GN=EEF1A2 PE=1 SV=1 | 16 |
| Q05996 | Zona pellucida sperm-binding protein 2 OS=Homo sapiens OX=9606 GN=ZP2 PE=1 SV=1 | 41 |
| Q06830 | Peroxiredoxin-1 OS=Homo sapiens OX=9606 GN=PRDX1 PE=1 SV=1 | 18 |
| Q08188 | Protein-glutamine gamma-glutamyltransferase E OS=Homo sapiens OX=9606 GN=TGM3 PE=1 SV=4 | 34 |
| Q08257 | Quinone oxidoreductase OS=Homo sapiens OX=9606 GN=CRYZ PE=1 SV=1 | 23 |
| Q08554 | Desmocollin-1 OS=Homo sapiens OX=9606 GN=DSC1 PE=1 SV=2 | 31 |
| Q09666 | Neuroblast differentiation-associated protein AHNAK OS=Homo sapiens OX=9606 GN=AHNAK PE=1 SV=2 | 17 |
| Q11206 | CMP-N-acetylneuraminate-beta-galactosamide-alpha-2,3-sialyltransferase 4 OS=Homo sapiens OX=9606 GN=ST3GAL4 PE=1 SV=1 | 4 |
| Q12836 | Zona pellucida sperm-binding protein 4 OS=Homo sapiens OX=9606 GN=ZP4 PE=1 SV=1 | 22 |
| Q12931 | Heat shock protein 75 kDa, mitochondrial OS=Homo sapiens OX=9606 GN=TRAP1 PE=1 SV=3 | 44 |
| Q13296 | Mammaglobin-A OS=Homo sapiens OX=9606 GN=SCGB2A2 PE=1 SV=1 | 1 |
| Q13404 | Ubiquitin-conjugating enzyme E2 variant 1 OS=Homo sapiens OX=9606 GN=UBE2V1 PE=1 SV=2 | 2 |
| Q13510 | Acid ceramidase OS=Homo sapiens OX=9606 GN=ASAH1 PE=1 SV=5 | 10 |
| Q13698 | Voltage-dependent L-type calcium channel subunit alpha-1S OS=Homo sapiens OX=9606 GN=CACNA1S PE=1 SV=4 | 1 |
| Q13835 | Plakophilin-1 OS=Homo sapiens OX=9606 GN=PKP1 PE=1 SV=2 | 34 |
| Q13867 | Bleomycin hydrolase OS=Homo sapiens OX=9606 GN=BLMH PE=1 SV=1 | 18 |
| Q14166 | Tubulin--tyrosine ligase-like protein 12 OS=Homo sapiens OX=9606 GN=TTLL12 PE=1 SV=2 | 33 |
| Q14204 | Cytoplasmic dynein 1 heavy chain 1 OS=Homo sapiens OX=9606 GN=DYNC1H1 PE=1 SV=5 | 169 |
| Q14525 | Keratin, type I cuticular Ha3-II OS=Homo sapiens OX=9606 GN=KRT33B PE=1 SV=3 | 4 |
| Q14574 | Desmocollin-3 OS=Homo sapiens OX=9606 GN=DSC3 PE=1 SV=3 | 15 |
| Q14764 | Major vault protein OS=Homo sapiens OX=9606 GN=MVP PE=1 SV=4 | 82 |
| Q14894 | Ketimine reductase mu-crystallin OS=Homo sapiens OX=9606 GN=CRYM PE=1 SV=1 | 13 |
| Q14974 | Importin subunit beta-1 OS=Homo sapiens OX=9606 GN=KPNB1 PE=1 SV=2 | 23 |
| Q14CN4 | Keratin, type II cytoskeletal 72 OS=Homo sapiens OX=9606 GN=KRT72 PE=1 SV=2 | 5 |
| Q15149 | Plectin OS=Homo sapiens OX=9606 GN=PLEC PE=1 SV=3 | 38 |
| Q15181 | Inorganic pyrophosphatase OS=Homo sapiens OX=9606 GN=PPA1 PE=1 SV=2 | 16 |
| Q15323 | Keratin, type I cuticular Ha1 OS=Homo sapiens OX=9606 GN=KRT31 PE=1 SV=3 | 25 |
| Q15436 | Protein transport protein Sec23A OS=Homo sapiens OX=9606 GN=SEC23A PE=1 SV=2 | 23 |
| Q15517 | Corneodesmosin OS=Homo sapiens OX=9606 GN=CDSN PE=1 SV=3 | 4 |
| Q15819 | Ubiquitin-conjugating enzyme E2 variant 2 OS=Homo sapiens OX=9606 GN=UBE2V2 PE=1 SV=4 | 10 |
| Q15828 | Cystatin-M OS=Homo sapiens OX=9606 GN=CST6 PE=1 SV=1 | 4 |
| Q16610 | Extracellular matrix protein 1 OS=Homo sapiens OX=9606 GN=ECM1 PE=1 SV=2 | 15 |
| Q16881 | Thioredoxin reductase 1, cytoplasmic OS=Homo sapiens OX=9606 GN=TXNRD1 PE=1 SV=3 | 29 |
| Q3ZCM7 | Tubulin beta-8 chain OS=Homo sapiens OX=9606 GN=TUBB8 PE=1 SV=2 | 30 |
| Q3ZCW2 | Galectin-related protein OS=Homo sapiens OX=9606 GN=LGALSL PE=1 SV=2 | 2 |
| Q4AC99 | Probable inactive 1-aminocyclopropane-1-carboxylate synthase-like protein 2 OS=Homo sapiens OX=9606 GN=ACCSL PE=2 SV=3 | 29 |
| Q53RT3 | Retroviral-like aspartic protease 1 OS=Homo sapiens OX=9606 GN=ASPRV1 PE=1 SV=1 | 2 |
| Q587J8 | KH domain-containing protein 3 OS=Homo sapiens OX=9606 GN=KHDC3L PE=1 SV=1 | 17 |
| Q5D862 | Filaggrin-2 OS=Homo sapiens OX=9606 GN=FLG2 PE=1 SV=1 | 25 |
| Q5T2N8 | ATPase family AAA domain-containing protein 3C OS=Homo sapiens OX=9606 GN=ATAD3C PE=2 SV=2 | 18 |
| Q5T749 | Keratinocyte proline-rich protein OS=Homo sapiens OX=9606 GN=KPRP PE=1 SV=1 | 21 |
| Q5T750 | Skin-specific protein 32 OS=Homo sapiens OX=9606 GN=XP32 PE=1 SV=1 | 8 |
| Q5T9A4 | ATPase family AAA domain-containing protein 3B OS=Homo sapiens OX=9606 GN=ATAD3B PE=1 SV=1 | 2 |
| Q5TC12 | ATP synthase mitochondrial F1 complex assembly factor 1 OS=Homo sapiens OX=9606 GN=ATPAF1 PE=1 SV=1 | 10 |
| Q5VVQ6 | Ubiquitin thioesterase OTU1 OS=Homo sapiens OX=9606 GN=YOD1 PE=1 SV=1 | 8 |
| Q5XKE5 | Keratin, type II cytoskeletal 79 OS=Homo sapiens OX=9606 GN=KRT79 PE=1 SV=2 | 10 |
| Q69YQ0 | Cytospin-A OS=Homo sapiens OX=9606 GN=SPECC1L PE=1 SV=3 | 13 |
| Q6E0U4 | Dermokine OS=Homo sapiens OX=9606 GN=DMKN PE=1 SV=3 | 4 |
| Q6HA08 | Astacin-like metalloendopeptidase OS=Homo sapiens OX=9606 GN=ASTL PE=1 SV=4 | 25 |
| Q6KB66 | Keratin, type II cytoskeletal 80 OS=Homo sapiens OX=9606 GN=KRT80 PE=1 SV=2 | 30 |
| Q6TGC4 | Protein-arginine deiminase type-6 OS=Homo sapiens OX=9606 GN=PADI6 PE=1 SV=3 | 50 |
| Q6UWP8 | Suprabasin OS=Homo sapiens OX=9606 GN=SBSN PE=1 SV=2 | 20 |
| Q6ZMN8 | Cyclin-I2 OS=Homo sapiens OX=9606 GN=CCNI2 PE=2 SV=1 | 18 |
| Q6ZMR3 | L-lactate dehydrogenase A-like 6A OS=Homo sapiens OX=9606 GN=LDHAL6A PE=1 SV=1 | 22 |
| Q6ZVX7 | F-box only protein 50 OS=Homo sapiens OX=9606 GN=NCCRP1 PE=1 SV=1 | 7 |
| Q7L5L3 | Lysophospholipase D GDPD3 OS=Homo sapiens OX=9606 GN=GDPD3 PE=1 SV=3 | 7 |
| Q7RTR0 | NACHT, LRR and PYD domains-containing protein 9 OS=Homo sapiens OX=9606 GN=NLRP9 PE=1 SV=1 | 46 |
| Q7Z3Y8 | Keratin, type I cytoskeletal 27 OS=Homo sapiens OX=9606 GN=KRT27 PE=1 SV=2 | 1 |
| Q7Z406 | Myosin-14 OS=Homo sapiens OX=9606 GN=MYH14 PE=1 SV=2 | 1 |
| Q7Z794 | Keratin, type II cytoskeletal 1b OS=Homo sapiens OX=9606 GN=KRT77 PE=1 SV=3 | 31 |
| Q86W25 | NACHT, LRR and PYD domains-containing protein 13 OS=Homo sapiens OX=9606 GN=NLRP13 PE=2 SV=2 | 54 |
| Q86YZ3 | Hornerin OS=Homo sapiens OX=9606 GN=HRNR PE=1 SV=2 | 27 |
| Q8IW75 | Serpin A12 OS=Homo sapiens OX=9606 GN=SERPINA12 PE=1 SV=1 | 13 |
| Q8N1N4 | Keratin, type II cytoskeletal 78 OS=Homo sapiens OX=9606 GN=KRT78 PE=1 SV=2 | 29 |
| Q8NEX9 | Short-chain dehydrogenase/reductase family 9C member 7 OS=Homo sapiens OX=9606 GN=SDR9C7 PE=1 SV=1 | 7 |
| Q8TEH3 | DENN domain-containing protein 1A OS=Homo sapiens OX=9606 GN=DENND1A PE=1 SV=2 | 1 |
| Q8TEK3 | Histone-lysine N-methyltransferase, H3 lysine-79 specific OS=Homo sapiens OX=9606 GN=DOT1L PE=1 SV=3 | 2 |
| Q8WVV4 | Protein POF1B OS=Homo sapiens OX=9606 GN=POF1B PE=1 SV=3 | 27 |
| Q8WX94 | NACHT, LRR and PYD domains-containing protein 7 OS=Homo sapiens OX=9606 GN=NLRP7 PE=1 SV=1 | 64 |
| Q92820 | Gamma-glutamyl hydrolase OS=Homo sapiens OX=9606 GN=GGH PE=1 SV=2 | 12 |
| Q96DA0 | Zymogen granule protein 16 homolog B OS=Homo sapiens OX=9606 GN=ZG16B PE=1 SV=3 | 3 |
| Q96DR8 | Mucin-like protein 1 OS=Homo sapiens OX=9606 GN=MUCL1 PE=1 SV=1 | 2 |
| Q96P63 | Serpin B12 OS=Homo sapiens OX=9606 GN=SERPINB12 PE=1 SV=1 | 21 |
| Q96QA5 | Gasdermin-A OS=Homo sapiens OX=9606 GN=GSDMA PE=1 SV=4 | 15 |
| Q96T88 | E3 ubiquitin-protein ligase UHRF1 OS=Homo sapiens OX=9606 GN=UHRF1 PE=1 SV=1 | 50 |
| Q96TC7 | Regulator of microtubule dynamics protein 3 OS=Homo sapiens OX=9606 GN=RMDN3 PE=1 SV=2 | 22 |
| Q99456 | Keratin, type I cytoskeletal 12 OS=Homo sapiens OX=9606 GN=KRT12 PE=1 SV=1 | 2 |
| Q99497 | Parkinson disease protein 7 OS=Homo sapiens OX=9606 GN=PARK7 PE=1 SV=2 | 15 |
| Q99536 | Synaptic vesicle membrane protein VAT-1 homolog OS=Homo sapiens OX=9606 GN=VAT1 PE=1 SV=2 | 13 |
| Q99735 | Microsomal glutathione S-transferase 2 OS=Homo sapiens OX=9606 GN=MGST2 PE=1 SV=1 | 2 |
| Q99832 | T-complex protein 1 subunit eta OS=Homo sapiens OX=9606 GN=CCT7 PE=1 SV=2 | 40 |
| Q9BQE3 | Tubulin alpha-1C chain OS=Homo sapiens OX=9606 GN=TUBA1C PE=1 SV=1 | 9 |
| Q9BYE4 | Small proline-rich protein 2G OS=Homo sapiens OX=9606 GN=SPRR2G PE=3 SV=1 | 4 |
| Q9C075 | Keratin, type I cytoskeletal 23 OS=Homo sapiens OX=9606 GN=KRT23 PE=1 SV=2 | 20 |
| Q9C0H9 | SRC kinase signaling inhibitor 1 OS=Homo sapiens OX=9606 GN=SRCIN1 PE=1 SV=4 | 1 |
| Q9GZZ8 | Extracellular glycoprotein lacritin OS=Homo sapiens OX=9606 GN=LACRT PE=1 SV=1 | 3 |
| Q9H0U9 | Testis-specific Y-encoded-like protein 1 OS=Homo sapiens OX=9606 GN=TSPYL1 PE=1 SV=3 | 1 |
| Q9H0W9 | Ester hydrolase C11orf54 OS=Homo sapiens OX=9606 GN=C11orf54 PE=1 SV=1 | 7 |
| Q9H1E1 | Ribonuclease 7 OS=Homo sapiens OX=9606 GN=RNASE7 PE=1 SV=2 | 5 |
| Q9H329 | Band 4.1-like protein 4B OS=Homo sapiens OX=9606 GN=EPB41L4B PE=2 SV=2 | 1 |
| Q9H444 | Charged multivesicular body protein 4b OS=Homo sapiens OX=9606 GN=CHMP4B PE=1 SV=1 | 8 |
| Q9H568 | Actin-like protein 8 OS=Homo sapiens OX=9606 GN=ACTL8 PE=1 SV=1 | 19 |
| Q9H808 | Transducin-like enhancer protein 6 OS=Homo sapiens OX=9606 GN=TLE6 PE=1 SV=2 | 33 |
| Q9HCY8 | Protein S100-A14 OS=Homo sapiens OX=9606 GN=S100A14 PE=1 SV=1 | 5 |
| Q9NRW1 | Ras-related protein Rab-6B OS=Homo sapiens OX=9606 GN=RAB6B PE=1 SV=1 | 16 |
| Q9NSB2 | Keratin, type II cuticular Hb4 OS=Homo sapiens OX=9606 GN=KRT84 PE=2 SV=2 | 24 |
| Q9NSB4 | Keratin, type II cuticular Hb2 OS=Homo sapiens OX=9606 GN=KRT82 PE=1 SV=3 | 2 |
| Q9NVI7 | ATPase family AAA domain-containing protein 3A OS=Homo sapiens OX=9606 GN=ATAD3A PE=1 SV=2 | 44 |
| Q9NX02 | NACHT, LRR and PYD domains-containing protein 2 OS=Homo sapiens OX=9606 GN=NLRP2 PE=1 SV=1 | 82 |
| Q9NZH6 | Interleukin-37 OS=Homo sapiens OX=9606 GN=IL37 PE=1 SV=1 | 4 |
| Q9NZH8 | Interleukin-36 gamma OS=Homo sapiens OX=9606 GN=IL36G PE=1 SV=1 | 8 |
| Q9NZT1 | Calmodulin-like protein 5 OS=Homo sapiens OX=9606 GN=CALML5 PE=1 SV=2 | 11 |
| Q9P0U3 | Sentrin-specific protease 1 OS=Homo sapiens OX=9606 GN=SENP1 PE=1 SV=2 | 1 |
| Q9UBQ7 | Glyoxylate reductase/hydroxypyruvate reductase OS=Homo sapiens OX=9606 GN=GRHPR PE=1 SV=1 | 19 |
| Q9UBS4 | DnaJ homolog subfamily B member 11 OS=Homo sapiens OX=9606 GN=DNAJB11 PE=1 SV=1 | 14 |
| Q9UHQ9 | NADH-cytochrome b5 reductase 1 OS=Homo sapiens OX=9606 GN=CYB5R1 PE=1 SV=1 | 1 |
| Q9UI42 | Carboxypeptidase A4 OS=Homo sapiens OX=9606 GN=CPA4 PE=1 SV=2 | 13 |
| Q9UKK9 | ADP-sugar pyrophosphatase OS=Homo sapiens OX=9606 GN=NUDT5 PE=1 SV=1 | 13 |
| Q9UQL6 | Histone deacetylase 5 OS=Homo sapiens OX=9606 GN=HDAC5 PE=1 SV=2 | 1 |
| Q9Y266 | Nuclear migration protein nudC OS=Homo sapiens OX=9606 GN=NUDC PE=1 SV=1 | 20 |
| Q9Y2T3 | Guanine deaminase OS=Homo sapiens OX=9606 GN=GDA PE=1 SV=1 | 16 |
| Q9Y2T7 | Y-box-binding protein 2 OS=Homo sapiens OX=9606 GN=YBX2 PE=1 SV=2 | 21 |
| Q9Y3R4 | Sialidase-2 OS=Homo sapiens OX=9606 GN=NEU2 PE=1 SV=2 | 13 |
| Q9Y487 | V-type proton ATPase 116 kDa subunit a 2 OS=Homo sapiens OX=9606 GN=ATP6V0A2 PE=1 SV=2 | 1 |
| Q9Y4W6 | AFG3-like protein 2 OS=Homo sapiens OX=9606 GN=AFG3L2 PE=1 SV=2 | 32 |
| Y-FGCZCont00028;Y-FGCZCont00058;Y-FGCZCont00341 | zz_FGCZCont0027_P13646_K1C13_HUMAN blastpHomologue_0.0 | 3 |
| Y-FGCZCont00046 | zz_FGCZCont0045_P13645_K1C10_HUMAN blastpHomologue_0.0 | 1 |
| Y-FGCZCont00057 | zz_FGCZCont0056_O77727_K1C15_SHEEP blastpHomologue_0.0 | 1 |
| Y-FGCZCont00078 | zz_FGCZCont0077_P04264_K2C1_HUMAN blastpHomologue_0.0 | 3 |
| Y-FGCZCont00113 | zz_FGCZCont0112_P02769_ALBU_BOVIN blastpHomologue_0.0 | 26 |
| Y-FGCZCont00118;Y-FGCZCont00378 | zz_FGCZCont0117_P02668_CASK_BOVIN blastpHomologue_6.0e-139 | 6 |
| Y-FGCZCont00124;Y-FGCZCont00375 | zz_FGCZCont0123_P02662_CASA1_BOVIN blastpHomologue_1.0e-158 | 7 |
| Y-FGCZCont00154;Y-FGCZCont00377 | zz_FGCZCont0153_P02666_CASB_BOVIN blastpHomologue_3.0e-157 | 3 |
| Y-FGCZCont00180 | zz_FGCZCont0179_P00761_TRYP_PIG blastpHomologue_4.0e-171 | 8 |
| Y-FGCZCont00192 | zz_FGCZCont0191_P15636_API_ACHLY blastpHomologue_0.0 | 12 |
| Y-FGCZCont00231;Y-FGCZCont00376 | zz_FGCZCont0230_P02663_CASA2_BOVIN blastpHomologue_2.0e-164 | 4 |
| Y-FGCZCont00256 | zz_FGCZCont0255_P12763_FETUA_BOVIN blastpHomologue_0.0 | 5 |
| Y-FGCZCont00257 | zz_FGCZCont0256_P15252_REF_HEVBR blastpHomologue_1.0e-96 | 10 |
| Y-FGCZCont00261 | zz_FGCZCont0260_FGCZCont0260_FGCZCont0260 iRT protein with AAAA spacer | 11 |
| Y-FGCZCont00287 | A3EZ79 TREMBL:A3EZ79;Q32W65;Q6UXC7 Tax_Id=9606 Gene_Symbol=DMKN Dermokine beta-1 | 1 |
| Y-FGCZCont00319 | P34955 SWISS-PROT:P34955 (Bos taurus) Alpha-1-antiproteinase precursor | 5 |
| Y-FGCZCont00347 | P01966 SWISS-PROT:P01966 (Bos taurus) Hemoglobin subunit alpha | 2 |
| Y-FGCZCont00380 | P02754 SWISS-PROT:P02754 Beta-lactoglobulin - Bos taurus (Bovine). | 7 |
| Y-FGCZCont00404 | Q3TTY5 SWISS-PROT:Q3TTY5 Tax_Id=10090 Gene_Symbol=Krt2 Keratin, type II cytoskeletal 2 epidermal | 1 |
| Y-FGCZCont00407 | Q61782 TREMBL:Q61782 Tax_Id=10090 Gene_Symbol=Krt14 Type I epidermal keratin (Fragment) | 1 |
| Y-FGCZCont00488 | sp_LYSC_CHICK_ | 12 |
| Y-FGCZCont00493 | sp_SRPP_HEVBR_ | 4 |
| A0A024R1R8;Q9Y2S6 | Translation machinery-associated protein 7B OS=Homo sapiens OX=9606 GN=TMA7B PE=3 SV=1 | 2 |
| A0A075B6H7;A0A0C4DH55;P01624 | Probable non-functional immunoglobulin kappa variable 3-7 OS=Homo sapiens OX=9606 GN=IGKV3-7 PE=1 SV=1 | 1 |
| A0A075B6I0 | Immunoglobulin lambda variable 8-61 OS=Homo sapiens OX=9606 GN=IGLV8-61 PE=3 SV=7 | 1 |
| A0A075B6P5;P01615 | Immunoglobulin kappa variable 2-28 OS=Homo sapiens OX=9606 GN=IGKV2-28 PE=3 SV=1 | 3 |
| A0A075B6R9;A0A0C4DH68 | Probable non-functional immunoglobulin kappa variable 2D-24 OS=Homo sapiens OX=9606 GN=IGKV2D-24 PE=1 SV=1 | 1 |
| A0A087WSY6 | Immunoglobulin kappa variable 3D-15 OS=Homo sapiens OX=9606 GN=IGKV3D-15 PE=3 SV=6 | 1 |
| A0A087X1C5;P10635 | Putative cytochrome P450 2D7 OS=Homo sapiens OX=9606 GN=CYP2D7 PE=5 SV=1 | 1 |
| A0A0A0MRZ8;P04433 | Immunoglobulin kappa variable 3D-11 OS=Homo sapiens OX=9606 GN=IGKV3D-11 PE=3 SV=6 | 1 |
| A0A0A0MS15 | Immunoglobulin heavy variable 3-49 OS=Homo sapiens OX=9606 GN=IGHV3-49 PE=3 SV=1 | 2 |
| A0A0B4J1U7 | Immunoglobulin heavy variable 6-1 OS=Homo sapiens OX=9606 GN=IGHV6-1 PE=3 SV=1 | 1 |
| A0A0B4J1V0;A0A0B4J1V6;A0A0B4J1Y9 | Immunoglobulin heavy variable 3-15 OS=Homo sapiens OX=9606 GN=IGHV3-15 PE=3 SV=1 | 1 |
| A0A0B4J2D5;P0DPI2 | Putative glutamine amidotransferase-like class 1 domain-containing protein 3B, mitochondrial OS=Homo sapiens OX=9606 GN=GATD3B PE=5 SV=1 | 12 |
| A0A0B4J2F0 | Protein PIGBOS1 OS=Homo sapiens OX=9606 GN=PIGBOS1 PE=1 SV=1 | 1 |
| A0A0C4DH25 | Immunoglobulin kappa variable 3D-20 OS=Homo sapiens OX=9606 GN=IGKV3D-20 PE=3 SV=1 | 2 |
| A0A0C4DH41;P01824;P01825;P06331;P0DP06;P0DP08 | Immunoglobulin heavy variable 4-61 OS=Homo sapiens OX=9606 GN=IGHV4-61 PE=1 SV=1 | 2 |
| A0A0G2JKW9 | Uncharacterized protein OS=Homo sapiens OX=9606 PE=4 SV=1 | 1 |
| A0A1B0GUC4 | Myocilin opposite strand protein OS=Homo sapiens OX=9606 GN=MYOCOS PE=4 SV=2 | 4 |
| A0A1B0GVG4 | Coiled-coil domain-containing protein 194 OS=Homo sapiens OX=9606 GN=CCDC194 PE=3 SV=1 | 1 |
| A0A1W2PR48 | Transducin-like enhancer protein 7 OS=Homo sapiens OX=9606 GN=TLE7 PE=3 SV=1 | 13 |
| A0A2R8YFL7 | Oocyte-secreted protein 4A OS=Homo sapiens OX=9606 GN=OOSP4A PE=3 SV=1 | 4 |
| A0A8V8TLL3 | Uncharacterized protein OS=Homo sapiens OX=9606 PE=4 SV=1 | 1 |
| A0AVT1 | Ubiquitin-like modifier-activating enzyme 6 OS=Homo sapiens OX=9606 GN=UBA6 PE=1 SV=1 | 50 |
| A0FGR8 | Extended synaptotagmin-2 OS=Homo sapiens OX=9606 GN=ESYT2 PE=1 SV=1 | 2 |
| A0JP26 | POTE ankyrin domain family member B3 OS=Homo sapiens OX=9606 GN=POTEB3 PE=1 SV=2 | 1 |
| A0MZ66 | Shootin-1 OS=Homo sapiens OX=9606 GN=SHTN1 PE=1 SV=4 | 22 |
| A1A4S6 | Rho GTPase-activating protein 10 OS=Homo sapiens OX=9606 GN=ARHGAP10 PE=1 SV=1 | 1 |
| A2RRP1 | NBAS subunit of NRZ tethering complex OS=Homo sapiens OX=9606 GN=NBAS PE=1 SV=2 | 12 |
| A2RUC4 | tRNA wybutosine-synthesizing protein 5 OS=Homo sapiens OX=9606 GN=TYW5 PE=1 SV=1 | 5 |
| A3KMH1 | von Willebrand factor A domain-containing protein 8 OS=Homo sapiens OX=9606 GN=VWA8 PE=1 SV=2 | 24 |
| A4D126 | D-ribitol-5-phosphate cytidylyltransferase OS=Homo sapiens OX=9606 GN=CRPPA PE=1 SV=2 | 5 |
| A4D1E9 | GTP-binding protein 10 OS=Homo sapiens OX=9606 GN=GTPBP10 PE=1 SV=1 | 19 |
| A4D1P6 | WD repeat-containing protein 91 OS=Homo sapiens OX=9606 GN=WDR91 PE=1 SV=2 | 2 |
| A5D8V6 | Vacuolar protein sorting-associated protein 37C OS=Homo sapiens OX=9606 GN=VPS37C PE=1 SV=2 | 2 |
| A5PLN9 | Trafficking protein particle complex subunit 13 OS=Homo sapiens OX=9606 GN=TRAPPC13 PE=1 SV=2 | 5 |
| A5YKK6 | CCR4-NOT transcription complex subunit 1 OS=Homo sapiens OX=9606 GN=CNOT1 PE=1 SV=2 | 9 |
| A6NC42 | Developmental pluripotency-associated 5 protein OS=Homo sapiens OX=9606 GN=DPPA5 PE=2 SV=1 | 5 |
| A6NCE7;Q9GZQ8 | Microtubule-associated proteins 1A/1B light chain 3 beta 2 OS=Homo sapiens OX=9606 GN=MAP1LC3B2 PE=1 SV=1 | 3 |
| A6NCF5 | Kelch-like protein 33 OS=Homo sapiens OX=9606 GN=KLHL33 PE=4 SV=2 | 1 |
| A6ND01 | Sperm-egg fusion protein Juno OS=Homo sapiens OX=9606 GN=IZUMO1R PE=1 SV=3 | 6 |
| A6NDB9 | Paralemmin-3 OS=Homo sapiens OX=9606 GN=PALM3 PE=1 SV=2 | 6 |
| A6NDG6 | Glycerol-3-phosphate phosphatase OS=Homo sapiens OX=9606 GN=PGP PE=1 SV=1 | 4 |
| A6NDX5 | Putative zinc finger protein 840 OS=Homo sapiens OX=9606 GN=ZNF840P PE=5 SV=5 | 29 |
| A6NDY0 | Embryonic polyadenylate-binding protein 2 OS=Homo sapiens OX=9606 GN=PABPN1L PE=2 SV=1 | 4 |
| A6NED2 | RCC1 domain-containing protein 1 OS=Homo sapiens OX=9606 GN=RCCD1 PE=1 SV=1 | 3 |
| A6NGQ2 | Oocyte-expressed protein homolog OS=Homo sapiens OX=9606 GN=OOEP PE=1 SV=3 | 10 |
| A6NK58 | Putative lipoyltransferase 2, mitochondrial OS=Homo sapiens OX=9606 GN=LIPT2 PE=1 SV=1 | 3 |
| A6NKN8 | Purkinje cell protein 4-like protein 1 OS=Homo sapiens OX=9606 GN=PCP4L1 PE=3 SV=3 | 2 |
| A6NKT7;Q7Z3J3 | RanBP2-like and GRIP domain-containing protein 3 OS=Homo sapiens OX=9606 GN=RGPD3 PE=3 SV=2 | 1 |
| A6NM11;A6NMS7 | Leucine-rich repeat-containing protein 37A2 OS=Homo sapiens OX=9606 GN=LRRC37A2 PE=2 SV=2 | 1 |
| A6NMX2 | Eukaryotic translation initiation factor 4E type 1B OS=Homo sapiens OX=9606 GN=EIF4E1B PE=1 SV=3 | 13 |
| A6NMZ7 | Collagen alpha-6(VI) chain OS=Homo sapiens OX=9606 GN=COL6A6 PE=1 SV=2 | 10 |
| A6NN14 | Zinc finger protein 729 OS=Homo sapiens OX=9606 GN=ZNF729 PE=2 SV=4 | 1 |
| A6NN73;Q08AF8;Q0D2H9 | Golgin subfamily A member 8C OS=Homo sapiens OX=9606 GN=GOLGA8CP PE=3 SV=2 | 1 |
| A6NNL5 | Uncharacterized protein C15orf61 OS=Homo sapiens OX=9606 GN=C15orf61 PE=2 SV=2 | 4 |
| A6NNZ2 | Tubulin beta 8B OS=Homo sapiens OX=9606 GN=TUBB8B PE=1 SV=1 | 5 |
| A6NP61 | Protein ZAR1-like OS=Homo sapiens OX=9606 GN=ZAR1L PE=2 SV=2 | 7 |
| A8K8P3 | Protein SFI1 homolog OS=Homo sapiens OX=9606 GN=SFI1 PE=1 SV=2 | 1 |
| A8MT69 | Centromere protein X OS=Homo sapiens OX=9606 GN=CENPX PE=1 SV=1 | 1 |
| A8MVS5 | Protein HIDE1 OS=Homo sapiens OX=9606 GN=HIDE1 PE=1 SV=2 | 1 |
| A8MWD9;P62308 | Putative small nuclear ribonucleoprotein G-like protein 15 OS=Homo sapiens OX=9606 GN=SNRPGP15 PE=5 SV=2 | 2 |
| A8MWK0 | Putative fatty acid desaturase 2-like protein FADS2B OS=Homo sapiens OX=9606 GN=FADS2B PE=5 SV=2 | 1 |
| A8MWY0 | Endosome/lysosome-associated apoptosis and autophagy regulator family member 2 OS=Homo sapiens OX=9606 GN=ELAPOR2 PE=1 SV=2 | 10 |
| A9QM74 | Importin subunit alpha-8 OS=Homo sapiens OX=9606 GN=KPNA7 PE=1 SV=1 | 13 |
| B2RPK0;P09429 | Putative high mobility group protein B1-like 1 OS=Homo sapiens OX=9606 GN=HMGB1P1 PE=5 SV=1 | 3 |
| B3KNS4 | Endogenous retrovirus group K3 member 1 OS=Homo sapiens OX=9606 GN=ERVK3-1 PE=1 SV=1 | 1 |
| B7ZAQ6;P0CG08 | Golgi pH regulator A OS=Homo sapiens OX=9606 GN=GPR89A PE=1 SV=2 | 1 |
| B9A064 | Immunoglobulin lambda-like polypeptide 5 OS=Homo sapiens OX=9606 GN=IGLL5 PE=2 SV=2 | 2 |
| C4AMC7;Q6VEQ5 | Putative WAS protein family homolog 3 OS=Homo sapiens OX=9606 GN=WASH3P PE=1 SV=2 | 5 |
| C9J7I0 | UBAP1-MVB12-associated (UMA)-domain containing protein 1 OS=Homo sapiens OX=9606 GN=UMAD1 PE=1 SV=2 | 2 |
| C9JE40 | Protein PAT1 homolog 2 OS=Homo sapiens OX=9606 GN=PATL2 PE=1 SV=1 | 20 |
| C9JLW8 | Mapk-regulated corepressor-interacting protein 1 OS=Homo sapiens OX=9606 GN=MCRIP1 PE=1 SV=1 | 1 |
| C9JRZ8 | Aldo-keto reductase family 1 member B15 OS=Homo sapiens OX=9606 GN=AKR1B15 PE=1 SV=2 | 1 |
| E9PAV3 | Nascent polypeptide-associated complex subunit alpha, muscle-specific form OS=Homo sapiens OX=9606 GN=NACA PE=1 SV=1 | 6 |
| O00116 | Alkyldihydroxyacetonephosphate synthase, peroxisomal OS=Homo sapiens OX=9606 GN=AGPS PE=1 SV=1 | 10 |
| O00142 | Thymidine kinase 2, mitochondrial OS=Homo sapiens OX=9606 GN=TK2 PE=1 SV=4 | 2 |
| O00151 | PDZ and LIM domain protein 1 OS=Homo sapiens OX=9606 GN=PDLIM1 PE=1 SV=4 | 9 |
| O00154 | Cytosolic acyl coenzyme A thioester hydrolase OS=Homo sapiens OX=9606 GN=ACOT7 PE=1 SV=3 | 13 |
| O00159 | Unconventional myosin-Ic OS=Homo sapiens OX=9606 GN=MYO1C PE=1 SV=4 | 1 |
| O00161 | Synaptosomal-associated protein 23 OS=Homo sapiens OX=9606 GN=SNAP23 PE=1 SV=1 | 5 |
| O00170 | AH receptor-interacting protein OS=Homo sapiens OX=9606 GN=AIP PE=1 SV=3 | 13 |
| O00186 | Syntaxin-binding protein 3 OS=Homo sapiens OX=9606 GN=STXBP3 PE=1 SV=2 | 15 |
| O00193 | Small acidic protein OS=Homo sapiens OX=9606 GN=SMAP PE=1 SV=1 | 3 |
| O00203 | AP-3 complex subunit beta-1 OS=Homo sapiens OX=9606 GN=AP3B1 PE=1 SV=3 | 18 |
| O00214 | Galectin-8 OS=Homo sapiens OX=9606 GN=LGALS8 PE=1 SV=4 | 7 |
| O00217 | NADH dehydrogenase [ubiquinone] iron-sulfur protein 8, mitochondrial OS=Homo sapiens OX=9606 GN=NDUFS8 PE=1 SV=1 | 4 |
| O00231 | 26S proteasome non-ATPase regulatory subunit 11 OS=Homo sapiens OX=9606 GN=PSMD11 PE=1 SV=3 | 23 |
| O00232 | 26S proteasome non-ATPase regulatory subunit 12 OS=Homo sapiens OX=9606 GN=PSMD12 PE=1 SV=3 | 24 |
| O00233 | 26S proteasome non-ATPase regulatory subunit 9 OS=Homo sapiens OX=9606 GN=PSMD9 PE=1 SV=3 | 7 |
| O00244 | Copper transport protein ATOX1 OS=Homo sapiens OX=9606 GN=ATOX1 PE=1 SV=1 | 5 |
| O00273 | DNA fragmentation factor subunit alpha OS=Homo sapiens OX=9606 GN=DFFA PE=1 SV=1 | 10 |
| O00291 | Huntingtin-interacting protein 1 OS=Homo sapiens OX=9606 GN=HIP1 PE=1 SV=5 | 5 |
| O00303 | Eukaryotic translation initiation factor 3 subunit F OS=Homo sapiens OX=9606 GN=EIF3F PE=1 SV=1 | 7 |
| O00330 | Pyruvate dehydrogenase protein X component, mitochondrial OS=Homo sapiens OX=9606 GN=PDHX PE=1 SV=3 | 17 |
| O00339 | Matrilin-2 OS=Homo sapiens OX=9606 GN=MATN2 PE=1 SV=4 | 5 |
| O00391 | Sulfhydryl oxidase 1 OS=Homo sapiens OX=9606 GN=QSOX1 PE=1 SV=3 | 2 |
| O00399 | Dynactin subunit 6 OS=Homo sapiens OX=9606 GN=DCTN6 PE=1 SV=1 | 2 |
| O00401 | Actin nucleation-promoting factor WASL OS=Homo sapiens OX=9606 GN=WASL PE=1 SV=2 | 4 |
| O00410 | Importin-5 OS=Homo sapiens OX=9606 GN=IPO5 PE=1 SV=4 | 21 |
| O00423 | Echinoderm microtubule-associated protein-like 1 OS=Homo sapiens OX=9606 GN=EML1 PE=1 SV=3 | 3 |
| O00429 | Dynamin-1-like protein OS=Homo sapiens OX=9606 GN=DNM1L PE=1 SV=2 | 41 |
| O00442 | RNA 3'-terminal phosphate cyclase OS=Homo sapiens OX=9606 GN=RTCA PE=1 SV=1 | 11 |
| O00471 | Exocyst complex component 5 OS=Homo sapiens OX=9606 GN=EXOC5 PE=1 SV=1 | 1 |
| O00483 | Cytochrome c oxidase subunit NDUFA4 OS=Homo sapiens OX=9606 GN=NDUFA4 PE=1 SV=1 | 2 |
| O00487 | 26S proteasome non-ATPase regulatory subunit 14 OS=Homo sapiens OX=9606 GN=PSMD14 PE=1 SV=1 | 9 |
| O00515 | Ladinin-1 OS=Homo sapiens OX=9606 GN=LAD1 PE=1 SV=2 | 28 |
| O00522 | Krev interaction trapped protein 1 OS=Homo sapiens OX=9606 GN=KRIT1 PE=1 SV=2 | 6 |
| O00534 | von Willebrand factor A domain-containing protein 5A OS=Homo sapiens OX=9606 GN=VWA5A PE=1 SV=2 | 26 |
| O00560 | Syntenin-1 OS=Homo sapiens OX=9606 GN=SDCBP PE=1 SV=1 | 4 |
| O00571 | ATP-dependent RNA helicase DDX3X OS=Homo sapiens OX=9606 GN=DDX3X PE=1 SV=3 | 25 |
| O00584 | Ribonuclease T2 OS=Homo sapiens OX=9606 GN=RNASET2 PE=1 SV=2 | 4 |
| O00592 | Podocalyxin OS=Homo sapiens OX=9606 GN=PODXL PE=1 SV=2 | 8 |
| O00629 | Importin subunit alpha-3 OS=Homo sapiens OX=9606 GN=KPNA4 PE=1 SV=1 | 3 |
| O00743 | Serine/threonine-protein phosphatase 6 catalytic subunit OS=Homo sapiens OX=9606 GN=PPP6C PE=1 SV=1 | 5 |
| O00754 | Lysosomal alpha-mannosidase OS=Homo sapiens OX=9606 GN=MAN2B1 PE=1 SV=3 | 1 |
| O00764 | Pyridoxal kinase OS=Homo sapiens OX=9606 GN=PDXK PE=1 SV=1 | 15 |
| O14493;O15551;O95484;P56747 | Claudin-4 OS=Homo sapiens OX=9606 GN=CLDN4 PE=1 SV=1 | 1 |
| O14521 | Succinate dehydrogenase [ubiquinone] cytochrome b small subunit, mitochondrial OS=Homo sapiens OX=9606 GN=SDHD PE=1 SV=1 | 1 |
| O14530 | Thioredoxin domain-containing protein 9 OS=Homo sapiens OX=9606 GN=TXNDC9 PE=1 SV=2 | 3 |
| O14548 | Cytochrome c oxidase subunit 7A-related protein, mitochondrial OS=Homo sapiens OX=9606 GN=COX7A2L PE=1 SV=2 | 3 |
| O14558 | Heat shock protein beta-6 OS=Homo sapiens OX=9606 GN=HSPB6 PE=1 SV=2 | 1 |
| O14561 | Acyl carrier protein, mitochondrial OS=Homo sapiens OX=9606 GN=NDUFAB1 PE=1 SV=3 | 4 |
| O14562 | Ubiquitin domain-containing protein UBFD1 OS=Homo sapiens OX=9606 GN=UBFD1 PE=1 SV=2 | 6 |
| O14579 | Coatomer subunit epsilon OS=Homo sapiens OX=9606 GN=COPE PE=1 SV=3 | 7 |
| O14617 | AP-3 complex subunit delta-1 OS=Homo sapiens OX=9606 GN=AP3D1 PE=1 SV=1 | 9 |
| O14618 | Copper chaperone for superoxide dismutase OS=Homo sapiens OX=9606 GN=CCS PE=1 SV=1 | 7 |
| O14662 | Syntaxin-16 OS=Homo sapiens OX=9606 GN=STX16 PE=1 SV=3 | 4 |
| O14672 | Disintegrin and metalloproteinase domain-containing protein 10 OS=Homo sapiens OX=9606 GN=ADAM10 PE=1 SV=1 | 3 |
| O14681 | Etoposide-induced protein 2.4 homolog OS=Homo sapiens OX=9606 GN=EI24 PE=1 SV=4 | 1 |
| O14713 | Integrin beta-1-binding protein 1 OS=Homo sapiens OX=9606 GN=ITGB1BP1 PE=1 SV=1 | 2 |
| O14734 | Acyl-coenzyme A thioesterase 8 OS=Homo sapiens OX=9606 GN=ACOT8 PE=1 SV=1 | 4 |
| O14735 | CDP-diacylglycerol--inositol 3-phosphatidyltransferase OS=Homo sapiens OX=9606 GN=CDIPT PE=1 SV=1 | 1 |
| O14737 | Programmed cell death protein 5 OS=Homo sapiens OX=9606 GN=PDCD5 PE=1 SV=3 | 11 |
| O14744 | Protein arginine N-methyltransferase 5 OS=Homo sapiens OX=9606 GN=PRMT5 PE=1 SV=4 | 7 |
| O14745 | Na(+)/H(+) exchange regulatory cofactor NHE-RF1 OS=Homo sapiens OX=9606 GN=NHERF1 PE=1 SV=4 | 16 |
| O14757 | Serine/threonine-protein kinase Chk1 OS=Homo sapiens OX=9606 GN=CHEK1 PE=1 SV=2 | 7 |
| O14771 | Zinc finger protein 213 OS=Homo sapiens OX=9606 GN=ZNF213 PE=1 SV=2 | 1 |
| O14773 | Tripeptidyl-peptidase 1 OS=Homo sapiens OX=9606 GN=TPP1 PE=1 SV=2 | 2 |
| O14777 | Kinetochore protein NDC80 homolog OS=Homo sapiens OX=9606 GN=NDC80 PE=1 SV=1 | 1 |
| O14782 | Kinesin-like protein KIF3C OS=Homo sapiens OX=9606 GN=KIF3C PE=1 SV=4 | 1 |
| O14787 | Transportin-2 OS=Homo sapiens OX=9606 GN=TNPO2 PE=1 SV=3 | 1 |
| O14791 | Apolipoprotein L1 OS=Homo sapiens OX=9606 GN=APOL1 PE=1 SV=5 | 1 |
| O14817 | Tetraspanin-4 OS=Homo sapiens OX=9606 GN=TSPAN4 PE=1 SV=1 | 1 |
| O14828 | Secretory carrier-associated membrane protein 3 OS=Homo sapiens OX=9606 GN=SCAMP3 PE=1 SV=3 | 2 |
| O14832 | Phytanoyl-CoA dioxygenase, peroxisomal OS=Homo sapiens OX=9606 GN=PHYH PE=1 SV=1 | 1 |
| O14862 | Interferon-inducible protein AIM2 OS=Homo sapiens OX=9606 GN=AIM2 PE=1 SV=1 | 1 |
| O14874 | [3-methyl-2-oxobutanoate dehydrogenase [lipoamide]] kinase, mitochondrial OS=Homo sapiens OX=9606 GN=BCKDK PE=1 SV=2 | 4 |
| O14896 | Interferon regulatory factor 6 OS=Homo sapiens OX=9606 GN=IRF6 PE=1 SV=1 | 9 |
| O14904 | Protein Wnt-9a OS=Homo sapiens OX=9606 GN=WNT9A PE=1 SV=2 | 1 |
| O14908 | PDZ domain-containing protein GIPC1 OS=Homo sapiens OX=9606 GN=GIPC1 PE=1 SV=2 | 7 |
| O14925;Q5SRD1 | Mitochondrial import inner membrane translocase subunit Tim23 OS=Homo sapiens OX=9606 GN=TIMM23 PE=1 SV=1 | 1 |
| O14929 | Histone acetyltransferase type B catalytic subunit OS=Homo sapiens OX=9606 GN=HAT1 PE=1 SV=2 | 2 |
| O14949 | Cytochrome b-c1 complex subunit 8 OS=Homo sapiens OX=9606 GN=UQCRQ PE=1 SV=4 | 5 |
| O14950;P19105 | Myosin regulatory light chain 12B OS=Homo sapiens OX=9606 GN=MYL12B PE=1 SV=2 | 7 |
| O14964 | Hepatocyte growth factor-regulated tyrosine kinase substrate OS=Homo sapiens OX=9606 GN=HGS PE=1 SV=1 | 9 |
| O14965 | Aurora kinase A OS=Homo sapiens OX=9606 GN=AURKA PE=1 SV=3 | 24 |
| O14972 | Vacuolar protein sorting-associated protein 26C OS=Homo sapiens OX=9606 GN=VPS26C PE=1 SV=1 | 2 |
| O14974 | Protein phosphatase 1 regulatory subunit 12A OS=Homo sapiens OX=9606 GN=PPP1R12A PE=1 SV=1 | 11 |
| O14975 | Long-chain fatty acid transport protein 2 OS=Homo sapiens OX=9606 GN=SLC27A2 PE=1 SV=2 | 1 |
| O14976 | Cyclin-G-associated kinase OS=Homo sapiens OX=9606 GN=GAK PE=1 SV=2 | 4 |
| O14979 | Heterogeneous nuclear ribonucleoprotein D-like OS=Homo sapiens OX=9606 GN=HNRNPDL PE=1 SV=3 | 2 |
| O14980 | Exportin-1 OS=Homo sapiens OX=9606 GN=XPO1 PE=1 SV=1 | 20 |
| O15020 | Spectrin beta chain, non-erythrocytic 2 OS=Homo sapiens OX=9606 GN=SPTBN2 PE=1 SV=3 | 1 |
| O15027 | Protein transport protein Sec16A OS=Homo sapiens OX=9606 GN=SEC16A PE=1 SV=4 | 9 |
| O15067 | Phosphoribosylformylglycinamidine synthase OS=Homo sapiens OX=9606 GN=PFAS PE=1 SV=4 | 13 |
| O15078 | Centrosomal protein of 290 kDa OS=Homo sapiens OX=9606 GN=CEP290 PE=1 SV=2 | 1 |
| O15116 | U6 snRNA-associated Sm-like protein LSm1 OS=Homo sapiens OX=9606 GN=LSM1 PE=1 SV=1 | 6 |
| O15118 | NPC intracellular cholesterol transporter 1 OS=Homo sapiens OX=9606 GN=NPC1 PE=1 SV=2 | 4 |
| O15120 | 1-acyl-sn-glycerol-3-phosphate acyltransferase beta OS=Homo sapiens OX=9606 GN=AGPAT2 PE=1 SV=1 | 2 |
| O15121 | Sphingolipid delta(4)-desaturase DES1 OS=Homo sapiens OX=9606 GN=DEGS1 PE=1 SV=1 | 1 |
| O15126 | Secretory carrier-associated membrane protein 1 OS=Homo sapiens OX=9606 GN=SCAMP1 PE=1 SV=2 | 3 |
| O15127 | Secretory carrier-associated membrane protein 2 OS=Homo sapiens OX=9606 GN=SCAMP2 PE=1 SV=2 | 4 |
| O15143 | Actin-related protein 2/3 complex subunit 1B OS=Homo sapiens OX=9606 GN=ARPC1B PE=1 SV=3 | 13 |
| O15144 | Actin-related protein 2/3 complex subunit 2 OS=Homo sapiens OX=9606 GN=ARPC2 PE=1 SV=1 | 13 |
| O15145 | Actin-related protein 2/3 complex subunit 3 OS=Homo sapiens OX=9606 GN=ARPC3 PE=1 SV=3 | 10 |
| O15155 | BET1 homolog OS=Homo sapiens OX=9606 GN=BET1 PE=1 SV=1 | 1 |
| O15160 | DNA-directed RNA polymerases I and III subunit RPAC1 OS=Homo sapiens OX=9606 GN=POLR1C PE=1 SV=1 | 3 |
| O15173 | Membrane-associated progesterone receptor component 2 OS=Homo sapiens OX=9606 GN=PGRMC2 PE=1 SV=1 | 5 |
| O15182 | Centrin-3 OS=Homo sapiens OX=9606 GN=CETN3 PE=1 SV=2 | 2 |
| O15212 | Prefoldin subunit 6 OS=Homo sapiens OX=9606 GN=PFDN6 PE=1 SV=1 | 5 |
| O15217 | Glutathione S-transferase A4 OS=Homo sapiens OX=9606 GN=GSTA4 PE=1 SV=1 | 1 |
| O15228 | Dihydroxyacetone phosphate acyltransferase OS=Homo sapiens OX=9606 GN=GNPAT PE=1 SV=1 | 1 |
| O15254 | Peroxisomal acyl-coenzyme A oxidase 3 OS=Homo sapiens OX=9606 GN=ACOX3 PE=1 SV=2 | 5 |
| O15258 | Protein RER1 OS=Homo sapiens OX=9606 GN=RER1 PE=1 SV=1 | 4 |
| O15260 | Surfeit locus protein 4 OS=Homo sapiens OX=9606 GN=SURF4 PE=1 SV=3 | 1 |
| O15347 | High mobility group protein B3 OS=Homo sapiens OX=9606 GN=HMGB3 PE=1 SV=4 | 6 |
| O15355 | Protein phosphatase 1G OS=Homo sapiens OX=9606 GN=PPM1G PE=1 SV=1 | 2 |
| O15371 | Eukaryotic translation initiation factor 3 subunit D OS=Homo sapiens OX=9606 GN=EIF3D PE=1 SV=1 | 8 |
| O15372 | Eukaryotic translation initiation factor 3 subunit H OS=Homo sapiens OX=9606 GN=EIF3H PE=1 SV=1 | 8 |
| O15382 | Branched-chain-amino-acid aminotransferase, mitochondrial OS=Homo sapiens OX=9606 GN=BCAT2 PE=1 SV=2 | 19 |
| O15397 | Importin-8 OS=Homo sapiens OX=9606 GN=IPO8 PE=1 SV=2 | 8 |
| O15400 | Syntaxin-7 OS=Homo sapiens OX=9606 GN=STX7 PE=1 SV=4 | 4 |
| O15440 | ATP-binding cassette sub-family C member 5 OS=Homo sapiens OX=9606 GN=ABCC5 PE=1 SV=2 | 2 |
| O15498 | Synaptobrevin homolog YKT6 OS=Homo sapiens OX=9606 GN=YKT6 PE=1 SV=1 | 5 |
| O15511 | Actin-related protein 2/3 complex subunit 5 OS=Homo sapiens OX=9606 GN=ARPC5 PE=1 SV=3 | 3 |
| O15537 | Retinoschisin OS=Homo sapiens OX=9606 GN=RS1 PE=1 SV=2 | 1 |
| O43143 | ATP-dependent RNA helicase DHX15 OS=Homo sapiens OX=9606 GN=DHX15 PE=1 SV=2 | 4 |
| O43169 | Cytochrome b5 type B OS=Homo sapiens OX=9606 GN=CYB5B PE=1 SV=3 | 3 |
| O43175 | D-3-phosphoglycerate dehydrogenase OS=Homo sapiens OX=9606 GN=PHGDH PE=1 SV=4 | 5 |
| O43181 | NADH dehydrogenase [ubiquinone] iron-sulfur protein 4, mitochondrial OS=Homo sapiens OX=9606 GN=NDUFS4 PE=1 SV=1 | 3 |
| O43194 | G-protein coupled receptor 39 OS=Homo sapiens OX=9606 GN=GPR39 PE=1 SV=1 | 2 |
| O43240 | Kallikrein-10 OS=Homo sapiens OX=9606 GN=KLK10 PE=1 SV=3 | 4 |
| O43242 | 26S proteasome non-ATPase regulatory subunit 3 OS=Homo sapiens OX=9606 GN=PSMD3 PE=1 SV=2 | 23 |
| O43264 | Centromere/kinetochore protein zw10 homolog OS=Homo sapiens OX=9606 GN=ZW10 PE=1 SV=3 | 8 |
| O43292 | Glycosylphosphatidylinositol anchor attachment 1 protein OS=Homo sapiens OX=9606 GN=GPAA1 PE=1 SV=3 | 5 |
| O43324 | Eukaryotic translation elongation factor 1 epsilon-1 OS=Homo sapiens OX=9606 GN=EEF1E1 PE=1 SV=1 | 2 |
| O43325 | LYR motif-containing protein 1 OS=Homo sapiens OX=9606 GN=LYRM1 PE=2 SV=1 | 1 |
| O43396 | Thioredoxin-like protein 1 OS=Homo sapiens OX=9606 GN=TXNL1 PE=1 SV=3 | 12 |
| O43399 | Tumor protein D54 OS=Homo sapiens OX=9606 GN=TPD52L2 PE=1 SV=2 | 13 |
| O43402 | ER membrane protein complex subunit 8 OS=Homo sapiens OX=9606 GN=EMC8 PE=1 SV=1 | 2 |
| O43405 | Cochlin OS=Homo sapiens OX=9606 GN=COCH PE=1 SV=1 | 25 |
| O43414 | ERI1 exoribonuclease 3 OS=Homo sapiens OX=9606 GN=ERI3 PE=1 SV=2 | 5 |
| O43426 | Synaptojanin-1 OS=Homo sapiens OX=9606 GN=SYNJ1 PE=1 SV=2 | 9 |
| O43447 | Peptidyl-prolyl cis-trans isomerase H OS=Homo sapiens OX=9606 GN=PPIH PE=1 SV=1 | 2 |
| O43488 | Aflatoxin B1 aldehyde reductase member 2 OS=Homo sapiens OX=9606 GN=AKR7A2 PE=1 SV=3 | 14 |
| O43491 | Band 4.1-like protein 2 OS=Homo sapiens OX=9606 GN=EPB41L2 PE=1 SV=1 | 35 |
| O43493 | Trans-Golgi network integral membrane protein 2 OS=Homo sapiens OX=9606 GN=TGOLN2 PE=1 SV=4 | 1 |
| O43592 | Exportin-T OS=Homo sapiens OX=9606 GN=XPOT PE=1 SV=2 | 3 |
| O43615 | Mitochondrial import inner membrane translocase subunit TIM44 OS=Homo sapiens OX=9606 GN=TIMM44 PE=1 SV=2 | 18 |
| O43617 | Trafficking protein particle complex subunit 3 OS=Homo sapiens OX=9606 GN=TRAPPC3 PE=1 SV=1 | 4 |
| O43633 | Charged multivesicular body protein 2a OS=Homo sapiens OX=9606 GN=CHMP2A PE=1 SV=1 | 1 |
| O43639 | Cytoplasmic protein NCK2 OS=Homo sapiens OX=9606 GN=NCK2 PE=1 SV=2 | 17 |
| O43670 | BUB3-interacting and GLEBS motif-containing protein ZNF207 OS=Homo sapiens OX=9606 GN=ZNF207 PE=1 SV=1 | 1 |
| O43674 | NADH dehydrogenase [ubiquinone] 1 beta subcomplex subunit 5, mitochondrial OS=Homo sapiens OX=9606 GN=NDUFB5 PE=1 SV=1 | 3 |
| O43676 | NADH dehydrogenase [ubiquinone] 1 beta subcomplex subunit 3 OS=Homo sapiens OX=9606 GN=NDUFB3 PE=1 SV=3 | 3 |
| O43678 | NADH dehydrogenase [ubiquinone] 1 alpha subcomplex subunit 2 OS=Homo sapiens OX=9606 GN=NDUFA2 PE=1 SV=3 | 3 |
| O43681 | ATPase GET3 OS=Homo sapiens OX=9606 GN=GET3 PE=1 SV=2 | 10 |
| O43684 | Mitotic checkpoint protein BUB3 OS=Homo sapiens OX=9606 GN=BUB3 PE=1 SV=1 | 12 |
| O43708 | Maleylacetoacetate isomerase OS=Homo sapiens OX=9606 GN=GSTZ1 PE=1 SV=3 | 3 |
| O43709 | Probable 18S rRNA (guanine-N(7))-methyltransferase OS=Homo sapiens OX=9606 GN=BUD23 PE=1 SV=2 | 1 |
| O43719 | HIV Tat-specific factor 1 OS=Homo sapiens OX=9606 GN=HTATSF1 PE=1 SV=1 | 6 |
| O43747 | AP-1 complex subunit gamma-1 OS=Homo sapiens OX=9606 GN=AP1G1 PE=1 SV=5 | 10 |
| O43752 | Syntaxin-6 OS=Homo sapiens OX=9606 GN=STX6 PE=1 SV=1 | 7 |
| O43765 | Small glutamine-rich tetratricopeptide repeat-containing protein alpha OS=Homo sapiens OX=9606 GN=SGTA PE=1 SV=1 | 4 |
| O43768 | Alpha-endosulfine OS=Homo sapiens OX=9606 GN=ENSA PE=1 SV=1 | 5 |
| O43772 | Mitochondrial carnitine/acylcarnitine carrier protein OS=Homo sapiens OX=9606 GN=SLC25A20 PE=1 SV=1 | 3 |
| O43776 | Asparagine--tRNA ligase, cytoplasmic OS=Homo sapiens OX=9606 GN=NARS1 PE=1 SV=1 | 15 |
| O43795 | Unconventional myosin-Ib OS=Homo sapiens OX=9606 GN=MYO1B PE=1 SV=3 | 26 |
| O43805 | Microtubule nucleation factor SSNA1 OS=Homo sapiens OX=9606 GN=SSNA1 PE=1 SV=2 | 1 |
| O43809 | Cleavage and polyadenylation specificity factor subunit 5 OS=Homo sapiens OX=9606 GN=NUDT21 PE=1 SV=1 | 3 |
| O43813 | Glutathione S-transferase LANCL1 OS=Homo sapiens OX=9606 GN=LANCL1 PE=1 SV=1 | 15 |
| O43822 | Cilia- and flagella-associated protein 410 OS=Homo sapiens OX=9606 GN=CFAP410 PE=1 SV=1 | 1 |
| O43837 | Isocitrate dehydrogenase [NAD] subunit beta, mitochondrial OS=Homo sapiens OX=9606 GN=IDH3B PE=1 SV=2 | 21 |
| O43847 | Nardilysin OS=Homo sapiens OX=9606 GN=NRDC PE=1 SV=3 | 4 |
| O43852 | Calumenin OS=Homo sapiens OX=9606 GN=CALU PE=1 SV=2 | 18 |
| O43854 | EGF-like repeat and discoidin I-like domain-containing protein 3 OS=Homo sapiens OX=9606 GN=EDIL3 PE=1 SV=1 | 10 |
| O43920 | NADH dehydrogenase [ubiquinone] iron-sulfur protein 5 OS=Homo sapiens OX=9606 GN=NDUFS5 PE=1 SV=3 | 4 |
| O43924 | Retinal rod rhodopsin-sensitive cGMP 3',5'-cyclic phosphodiesterase subunit delta OS=Homo sapiens OX=9606 GN=PDE6D PE=1 SV=1 | 2 |
| O60218 | Aldo-keto reductase family 1 member B10 OS=Homo sapiens OX=9606 GN=AKR1B10 PE=1 SV=2 | 13 |
| O60220 | Mitochondrial import inner membrane translocase subunit Tim8 A OS=Homo sapiens OX=9606 GN=TIMM8A PE=1 SV=1 | 6 |
| O60229 | Kalirin OS=Homo sapiens OX=9606 GN=KALRN PE=1 SV=3 | 12 |
| O60232 | Protein ZNRD2 OS=Homo sapiens OX=9606 GN=ZNRD2 PE=1 SV=1 | 1 |
| O60256 | Phosphoribosyl pyrophosphate synthase-associated protein 2 OS=Homo sapiens OX=9606 GN=PRPSAP2 PE=1 SV=1 | 13 |
| O60282 | Kinesin heavy chain isoform 5C OS=Homo sapiens OX=9606 GN=KIF5C PE=1 SV=1 | 7 |
| O60331 | Phosphatidylinositol 4-phosphate 5-kinase type-1 gamma OS=Homo sapiens OX=9606 GN=PIP5K1C PE=1 SV=2 | 3 |
| O60333 | Kinesin-like protein KIF1B OS=Homo sapiens OX=9606 GN=KIF1B PE=1 SV=5 | 3 |
| O60341 | Lysine-specific histone demethylase 1A OS=Homo sapiens OX=9606 GN=KDM1A PE=1 SV=2 | 2 |
| O60383 | Growth/differentiation factor 9 OS=Homo sapiens OX=9606 GN=GDF9 PE=1 SV=1 | 25 |
| O60391 | Glutamate receptor ionotropic, NMDA 3B OS=Homo sapiens OX=9606 GN=GRIN3B PE=2 SV=2 | 1 |
| O60437 | Periplakin OS=Homo sapiens OX=9606 GN=PPL PE=1 SV=4 | 11 |
| O60443 | Gasdermin-E OS=Homo sapiens OX=9606 GN=GSDME PE=1 SV=2 | 17 |
| O60488 | Long-chain-fatty-acid--CoA ligase 4 OS=Homo sapiens OX=9606 GN=ACSL4 PE=1 SV=2 | 7 |
| O60493 | Sorting nexin-3 OS=Homo sapiens OX=9606 GN=SNX3 PE=1 SV=3 | 3 |
| O60502 | Protein O-GlcNAcase OS=Homo sapiens OX=9606 GN=OGA PE=1 SV=2 | 3 |
| O60506 | Heterogeneous nuclear ribonucleoprotein Q OS=Homo sapiens OX=9606 GN=SYNCRIP PE=1 SV=2 | 19 |
| O60512 | Beta-1,4-galactosyltransferase 3 OS=Homo sapiens OX=9606 GN=B4GALT3 PE=1 SV=2 | 1 |
| O60513 | Beta-1,4-galactosyltransferase 4 OS=Homo sapiens OX=9606 GN=B4GALT4 PE=1 SV=1 | 3 |
| O60522 | Tudor domain-containing protein 6 OS=Homo sapiens OX=9606 GN=TDRD6 PE=2 SV=2 | 15 |
| O60524 | Ribosome quality control complex subunit NEMF OS=Homo sapiens OX=9606 GN=NEMF PE=1 SV=4 | 2 |
| O60547 | GDP-mannose 4,6 dehydratase OS=Homo sapiens OX=9606 GN=GMDS PE=1 SV=1 | 15 |
| O60566 | Mitotic checkpoint serine/threonine-protein kinase BUB1 beta OS=Homo sapiens OX=9606 GN=BUB1B PE=1 SV=3 | 35 |
| O60610 | Protein diaphanous homolog 1 OS=Homo sapiens OX=9606 GN=DIAPH1 PE=1 SV=2 | 37 |
| O60641 | Clathrin coat assembly protein AP180 OS=Homo sapiens OX=9606 GN=SNAP91 PE=1 SV=2 | 3 |
| O60645 | Exocyst complex component 3 OS=Homo sapiens OX=9606 GN=EXOC3 PE=1 SV=3 | 5 |
| O60664 | Perilipin-3 OS=Homo sapiens OX=9606 GN=PLIN3 PE=1 SV=3 | 24 |
| O60684 | Importin subunit alpha-7 OS=Homo sapiens OX=9606 GN=KPNA6 PE=1 SV=1 | 16 |
| O60701 | UDP-glucose 6-dehydrogenase OS=Homo sapiens OX=9606 GN=UGDH PE=1 SV=1 | 26 |
| O60716 | Catenin delta-1 OS=Homo sapiens OX=9606 GN=CTNND1 PE=1 SV=1 | 16 |
| O60739 | Eukaryotic translation initiation factor 1b OS=Homo sapiens OX=9606 GN=EIF1B PE=1 SV=2 | 5 |
| O60749 | Sorting nexin-2 OS=Homo sapiens OX=9606 GN=SNX2 PE=1 SV=2 | 21 |
| O60762 | Dolichol-phosphate mannosyltransferase subunit 1 OS=Homo sapiens OX=9606 GN=DPM1 PE=1 SV=1 | 2 |
| O60763 | General vesicular transport factor p115 OS=Homo sapiens OX=9606 GN=USO1 PE=1 SV=2 | 21 |
| O60784 | Target of Myb1 membrane trafficking protein OS=Homo sapiens OX=9606 GN=TOM1 PE=1 SV=2 | 3 |
| O60825 | 6-phosphofructo-2-kinase/fructose-2,6-bisphosphatase 2 OS=Homo sapiens OX=9606 GN=PFKFB2 PE=1 SV=2 | 20 |
| O60826 | Coiled-coil domain-containing protein 22 OS=Homo sapiens OX=9606 GN=CCDC22 PE=1 SV=1 | 14 |
| O60830 | Mitochondrial import inner membrane translocase subunit Tim17-B OS=Homo sapiens OX=9606 GN=TIMM17B PE=1 SV=1 | 1 |
| O60831 | PRA1 family protein 2 OS=Homo sapiens OX=9606 GN=PRAF2 PE=1 SV=1 | 2 |
| O60841 | Eukaryotic translation initiation factor 5B OS=Homo sapiens OX=9606 GN=EIF5B PE=1 SV=4 | 6 |
| O60861 | Growth arrest-specific protein 7 OS=Homo sapiens OX=9606 GN=GAS7 PE=1 SV=3 | 5 |
| O60869 | Endothelial differentiation-related factor 1 OS=Homo sapiens OX=9606 GN=EDF1 PE=1 SV=1 | 4 |
| O60879 | Protein diaphanous homolog 2 OS=Homo sapiens OX=9606 GN=DIAPH2 PE=1 SV=1 | 10 |
| O60884 | DnaJ homolog subfamily A member 2 OS=Homo sapiens OX=9606 GN=DNAJA2 PE=1 SV=1 | 16 |
| O60888 | Protein CutA OS=Homo sapiens OX=9606 GN=CUTA PE=1 SV=2 | 3 |
| O60925 | Prefoldin subunit 1 OS=Homo sapiens OX=9606 GN=PFDN1 PE=1 SV=2 | 6 |
| O75027 | Iron-sulfur clusters transporter ABCB7, mitochondrial OS=Homo sapiens OX=9606 GN=ABCB7 PE=1 SV=2 | 6 |
| O75054 | Immunoglobulin superfamily member 3 OS=Homo sapiens OX=9606 GN=IGSF3 PE=2 SV=3 | 2 |
| O75083 | WD repeat-containing protein 1 OS=Homo sapiens OX=9606 GN=WDR1 PE=1 SV=4 | 32 |
| O75146 | Huntingtin-interacting protein 1-related protein OS=Homo sapiens OX=9606 GN=HIP1R PE=1 SV=2 | 3 |
| O75165 | DnaJ homolog subfamily C member 13 OS=Homo sapiens OX=9606 GN=DNAJC13 PE=1 SV=5 | 90 |
| O75175 | CCR4-NOT transcription complex subunit 3 OS=Homo sapiens OX=9606 GN=CNOT3 PE=1 SV=1 | 3 |
| O75190 | DnaJ homolog subfamily B member 6 OS=Homo sapiens OX=9606 GN=DNAJB6 PE=1 SV=2 | 1 |
| O75208 | Ubiquinone biosynthesis protein COQ9, mitochondrial OS=Homo sapiens OX=9606 GN=COQ9 PE=1 SV=1 | 3 |
| O75251 | NADH dehydrogenase [ubiquinone] iron-sulfur protein 7, mitochondrial OS=Homo sapiens OX=9606 GN=NDUFS7 PE=1 SV=3 | 3 |
| O75306 | NADH dehydrogenase [ubiquinone] iron-sulfur protein 2, mitochondrial OS=Homo sapiens OX=9606 GN=NDUFS2 PE=1 SV=2 | 10 |
| O75312 | Zinc finger protein ZPR1 OS=Homo sapiens OX=9606 GN=ZPR1 PE=1 SV=1 | 3 |
| O75323 | Protein NipSnap homolog 2 OS=Homo sapiens OX=9606 GN=NIPSNAP2 PE=1 SV=1 | 8 |
| O75340 | Programmed cell death protein 6 OS=Homo sapiens OX=9606 GN=PDCD6 PE=1 SV=1 | 6 |
| O75347 | Tubulin-specific chaperone A OS=Homo sapiens OX=9606 GN=TBCA PE=1 SV=3 | 7 |
| O75348 | V-type proton ATPase subunit G 1 OS=Homo sapiens OX=9606 GN=ATP6V1G1 PE=1 SV=3 | 3 |
| O75351 | Vacuolar protein sorting-associated protein 4B OS=Homo sapiens OX=9606 GN=VPS4B PE=1 SV=2 | 7 |
| O75352 | Mannose-P-dolichol utilization defect 1 protein OS=Homo sapiens OX=9606 GN=MPDU1 PE=1 SV=2 | 3 |
| O75380 | NADH dehydrogenase [ubiquinone] iron-sulfur protein 6, mitochondrial OS=Homo sapiens OX=9606 GN=NDUFS6 PE=1 SV=1 | 3 |
| O75381 | Peroxisomal membrane protein PEX14 OS=Homo sapiens OX=9606 GN=PEX14 PE=1 SV=1 | 3 |
| O75382 | Tripartite motif-containing protein 3 OS=Homo sapiens OX=9606 GN=TRIM3 PE=1 SV=2 | 3 |
| O75386 | Tubby-related protein 3 OS=Homo sapiens OX=9606 GN=TULP3 PE=1 SV=2 | 8 |
| O75396 | Vesicle-trafficking protein SEC22b OS=Homo sapiens OX=9606 GN=SEC22B PE=1 SV=5 | 13 |
| O75431 | Metaxin-2 OS=Homo sapiens OX=9606 GN=MTX2 PE=1 SV=1 | 7 |
| O75439 | Mitochondrial-processing peptidase subunit beta OS=Homo sapiens OX=9606 GN=PMPCB PE=1 SV=2 | 11 |
| O75477 | Erlin-1 OS=Homo sapiens OX=9606 GN=ERLIN1 PE=1 SV=2 | 7 |
| O75489 | NADH dehydrogenase [ubiquinone] iron-sulfur protein 3, mitochondrial OS=Homo sapiens OX=9606 GN=NDUFS3 PE=1 SV=1 | 10 |
| O75521 | Enoyl-CoA delta isomerase 2 OS=Homo sapiens OX=9606 GN=ECI2 PE=1 SV=4 | 14 |
| O75531 | Barrier-to-autointegration factor OS=Homo sapiens OX=9606 GN=BANF1 PE=1 SV=1 | 7 |
| O75534 | Cold shock domain-containing protein E1 OS=Homo sapiens OX=9606 GN=CSDE1 PE=1 SV=2 | 14 |
| O75553 | Disabled homolog 1 OS=Homo sapiens OX=9606 GN=DAB1 PE=1 SV=3 | 1 |
| O75569 | Interferon-inducible double-stranded RNA-dependent protein kinase activator A OS=Homo sapiens OX=9606 GN=PRKRA PE=1 SV=1 | 5 |
| O75575 | DNA-directed RNA polymerase III subunit RPC9 OS=Homo sapiens OX=9606 GN=CRCP PE=1 SV=1 | 1 |
| O75592 | E3 ubiquitin-protein ligase MYCBP2 OS=Homo sapiens OX=9606 GN=MYCBP2 PE=1 SV=4 | 1 |
| O75600 | 2-amino-3-ketobutyrate coenzyme A ligase, mitochondrial OS=Homo sapiens OX=9606 GN=GCAT PE=1 SV=1 | 16 |
| O75607 | Nucleoplasmin-3 OS=Homo sapiens OX=9606 GN=NPM3 PE=1 SV=3 | 2 |
| O75616 | GTPase Era, mitochondrial OS=Homo sapiens OX=9606 GN=ERAL1 PE=1 SV=2 | 2 |
| O75635 | Serpin B7 OS=Homo sapiens OX=9606 GN=SERPINB7 PE=1 SV=1 | 12 |
| O75643 | U5 small nuclear ribonucleoprotein 200 kDa helicase OS=Homo sapiens OX=9606 GN=SNRNP200 PE=1 SV=2 | 3 |
| O75663 | TIP41-like protein OS=Homo sapiens OX=9606 GN=TIPRL PE=1 SV=2 | 10 |
| O75674 | TOM1-like protein 1 OS=Homo sapiens OX=9606 GN=TOM1L1 PE=1 SV=2 | 6 |
| O75688 | Protein phosphatase 1B OS=Homo sapiens OX=9606 GN=PPM1B PE=1 SV=1 | 11 |
| O75717 | WD repeat and HMG-box DNA-binding protein 1 OS=Homo sapiens OX=9606 GN=WDHD1 PE=1 SV=1 | 1 |
| O75746 | Electrogenic aspartate/glutamate antiporter SLC25A12, mitochondrial OS=Homo sapiens OX=9606 GN=SLC25A12 PE=1 SV=2 | 21 |
| O75787 | Renin receptor OS=Homo sapiens OX=9606 GN=ATP6AP2 PE=1 SV=2 | 6 |
| O75821 | Eukaryotic translation initiation factor 3 subunit G OS=Homo sapiens OX=9606 GN=EIF3G PE=1 SV=2 | 4 |
| O75822 | Eukaryotic translation initiation factor 3 subunit J OS=Homo sapiens OX=9606 GN=EIF3J PE=1 SV=2 | 7 |
| O75832 | 26S proteasome non-ATPase regulatory subunit 10 OS=Homo sapiens OX=9606 GN=PSMD10 PE=1 SV=1 | 11 |
| O75874 | Isocitrate dehydrogenase [NADP] cytoplasmic OS=Homo sapiens OX=9606 GN=IDH1 PE=1 SV=2 | 21 |
| O75880 | Protein SCO1 homolog, mitochondrial OS=Homo sapiens OX=9606 GN=SCO1 PE=1 SV=1 | 1 |
| O75882 | Attractin OS=Homo sapiens OX=9606 GN=ATRN PE=1 SV=2 | 5 |
| O75884 | Serine hydrolase RBBP9 OS=Homo sapiens OX=9606 GN=RBBP9 PE=1 SV=2 | 7 |
| O75886 | Signal transducing adapter molecule 2 OS=Homo sapiens OX=9606 GN=STAM2 PE=1 SV=1 | 11 |
| O75896 | Tumor suppressor candidate 2 OS=Homo sapiens OX=9606 GN=TUSC2 PE=1 SV=3 | 2 |
| O75935 | Dynactin subunit 3 OS=Homo sapiens OX=9606 GN=DCTN3 PE=1 SV=1 | 5 |
| O75947 | ATP synthase subunit d, mitochondrial OS=Homo sapiens OX=9606 GN=ATP5PD PE=1 SV=3 | 10 |
| O75952 | Calcium-binding tyrosine phosphorylation-regulated protein OS=Homo sapiens OX=9606 GN=CABYR PE=1 SV=2 | 5 |
| O75954 | Tetraspanin-9 OS=Homo sapiens OX=9606 GN=TSPAN9 PE=1 SV=1 | 2 |
| O75955 | Flotillin-1 OS=Homo sapiens OX=9606 GN=FLOT1 PE=1 SV=3 | 4 |
| O76003 | Glutaredoxin-3 OS=Homo sapiens OX=9606 GN=GLRX3 PE=1 SV=2 | 9 |
| O76024 | Wolframin OS=Homo sapiens OX=9606 GN=WFS1 PE=1 SV=2 | 16 |
| O76031 | ATP-dependent Clp protease ATP-binding subunit clpX-like, mitochondrial OS=Homo sapiens OX=9606 GN=CLPX PE=1 SV=2 | 5 |
| O76071 | Probable cytosolic iron-sulfur protein assembly protein CIAO1 OS=Homo sapiens OX=9606 GN=CIAO1 PE=1 SV=1 | 10 |
| O76094 | Signal recognition particle subunit SRP72 OS=Homo sapiens OX=9606 GN=SRP72 PE=1 SV=3 | 6 |
| O76095 | Protein JTB OS=Homo sapiens OX=9606 GN=JTB PE=1 SV=1 | 1 |
| O94760 | N(G),N(G)-dimethylarginine dimethylaminohydrolase 1 OS=Homo sapiens OX=9606 GN=DDAH1 PE=1 SV=3 | 17 |
| O94763 | Unconventional prefoldin RPB5 interactor 1 OS=Homo sapiens OX=9606 GN=URI1 PE=1 SV=3 | 1 |
| O94766 | Galactosylgalactosylxylosylprotein 3-beta-glucuronosyltransferase 3 OS=Homo sapiens OX=9606 GN=B3GAT3 PE=1 SV=2 | 1 |
| O94788 | Retinal dehydrogenase 2 OS=Homo sapiens OX=9606 GN=ALDH1A2 PE=1 SV=3 | 16 |
| O94826 | Mitochondrial import receptor subunit TOM70 OS=Homo sapiens OX=9606 GN=TOMM70 PE=1 SV=1 | 23 |
| O94832 | Unconventional myosin-Id OS=Homo sapiens OX=9606 GN=MYO1D PE=1 SV=2 | 11 |
| O94874 | E3 UFM1-protein ligase 1 OS=Homo sapiens OX=9606 GN=UFL1 PE=1 SV=2 | 8 |
| O94889 | Kelch-like protein 18 OS=Homo sapiens OX=9606 GN=KLHL18 PE=1 SV=3 | 3 |
| O94901 | SUN domain-containing protein 1 OS=Homo sapiens OX=9606 GN=SUN1 PE=1 SV=4 | 14 |
| O94903 | Pyridoxal phosphate homeostasis protein OS=Homo sapiens OX=9606 GN=PLPBP PE=1 SV=1 | 3 |
| O94905 | Erlin-2 OS=Homo sapiens OX=9606 GN=ERLIN2 PE=1 SV=1 | 9 |
| O94913 | Pre-mRNA cleavage complex 2 protein Pcf11 OS=Homo sapiens OX=9606 GN=PCF11 PE=1 SV=3 | 1 |
| O94919 | Endonuclease domain-containing 1 protein OS=Homo sapiens OX=9606 GN=ENDOD1 PE=1 SV=2 | 9 |
| O94925 | Glutaminase kidney isoform, mitochondrial OS=Homo sapiens OX=9606 GN=GLS PE=1 SV=1 | 6 |
| O94927 | HAUS augmin-like complex subunit 5 OS=Homo sapiens OX=9606 GN=HAUS5 PE=1 SV=2 | 1 |
| O94967 | WD repeat-containing protein 47 OS=Homo sapiens OX=9606 GN=WDR47 PE=1 SV=1 | 4 |
| O94973 | AP-2 complex subunit alpha-2 OS=Homo sapiens OX=9606 GN=AP2A2 PE=1 SV=2 | 4 |
| O94979 | Protein transport protein Sec31A OS=Homo sapiens OX=9606 GN=SEC31A PE=1 SV=3 | 6 |
| O94988 | Protein FAM13A OS=Homo sapiens OX=9606 GN=FAM13A PE=1 SV=2 | 19 |
| O95081 | Arf-GAP domain and FG repeat-containing protein 2 OS=Homo sapiens OX=9606 GN=AGFG2 PE=1 SV=2 | 5 |
| O95084 | Serine protease 23 OS=Homo sapiens OX=9606 GN=PRSS23 PE=1 SV=1 | 1 |
| O95140 | Mitofusin-2 OS=Homo sapiens OX=9606 GN=MFN2 PE=1 SV=3 | 5 |
| O95147 | Dual specificity protein phosphatase 14 OS=Homo sapiens OX=9606 GN=DUSP14 PE=1 SV=1 | 3 |
| O95153 | Peripheral-type benzodiazepine receptor-associated protein 1 OS=Homo sapiens OX=9606 GN=TSPOAP1 PE=1 SV=2 | 1 |
| O95159 | Zinc finger protein-like 1 OS=Homo sapiens OX=9606 GN=ZFPL1 PE=1 SV=2 | 1 |
| O95168 | NADH dehydrogenase [ubiquinone] 1 beta subcomplex subunit 4 OS=Homo sapiens OX=9606 GN=NDUFB4 PE=1 SV=3 | 2 |
| O95169 | NADH dehydrogenase [ubiquinone] 1 beta subcomplex subunit 8, mitochondrial OS=Homo sapiens OX=9606 GN=NDUFB8 PE=1 SV=1 | 3 |
| O95178 | NADH dehydrogenase [ubiquinone] 1 beta subcomplex subunit 2, mitochondrial OS=Homo sapiens OX=9606 GN=NDUFB2 PE=1 SV=1 | 1 |
| O95182 | NADH dehydrogenase [ubiquinone] 1 alpha subcomplex subunit 7 OS=Homo sapiens OX=9606 GN=NDUFA7 PE=1 SV=3 | 5 |
| O95197 | Reticulon-3 OS=Homo sapiens OX=9606 GN=RTN3 PE=1 SV=2 | 4 |
| O95199 | RCC1 and BTB domain-containing protein 2 OS=Homo sapiens OX=9606 GN=RCBTB2 PE=1 SV=1 | 1 |
| O95202 | Mitochondrial proton/calcium exchanger protein OS=Homo sapiens OX=9606 GN=LETM1 PE=1 SV=1 | 17 |
| O95219 | Sorting nexin-4 OS=Homo sapiens OX=9606 GN=SNX4 PE=1 SV=1 | 19 |
| O95229 | ZW10 interactor OS=Homo sapiens OX=9606 GN=ZWINT PE=1 SV=2 | 1 |
| O95239 | Chromosome-associated kinesin KIF4A OS=Homo sapiens OX=9606 GN=KIF4A PE=1 SV=3 | 10 |
| O95248 | Myotubularin-related protein 5 OS=Homo sapiens OX=9606 GN=SBF1 PE=1 SV=4 | 2 |
| O95274 | Ly6/PLAUR domain-containing protein 3 OS=Homo sapiens OX=9606 GN=LYPD3 PE=1 SV=2 | 1 |
| O95292 | Vesicle-associated membrane protein-associated protein B/C OS=Homo sapiens OX=9606 GN=VAPB PE=1 SV=3 | 13 |
| O95294 | RasGAP-activating-like protein 1 OS=Homo sapiens OX=9606 GN=RASAL1 PE=1 SV=3 | 3 |
| O95295 | SNARE-associated protein Snapin OS=Homo sapiens OX=9606 GN=SNAPIN PE=1 SV=1 | 1 |
| O95297 | Myelin protein zero-like protein 1 OS=Homo sapiens OX=9606 GN=MPZL1 PE=1 SV=1 | 1 |
| O95298 | NADH dehydrogenase [ubiquinone] 1 subunit C2 OS=Homo sapiens OX=9606 GN=NDUFC2 PE=1 SV=1 | 3 |
| O95299 | NADH dehydrogenase [ubiquinone] 1 alpha subcomplex subunit 10, mitochondrial OS=Homo sapiens OX=9606 GN=NDUFA10 PE=1 SV=1 | 7 |
| O95336 | 6-phosphogluconolactonase OS=Homo sapiens OX=9606 GN=PGLS PE=1 SV=2 | 12 |
| O95347 | Structural maintenance of chromosomes protein 2 OS=Homo sapiens OX=9606 GN=SMC2 PE=1 SV=2 | 4 |
| O95352 | Ubiquitin-like modifier-activating enzyme ATG7 OS=Homo sapiens OX=9606 GN=ATG7 PE=1 SV=1 | 9 |
| O95363 | Phenylalanine--tRNA ligase, mitochondrial OS=Homo sapiens OX=9606 GN=FARS2 PE=1 SV=1 | 3 |
| O95372 | Acyl-protein thioesterase 2 OS=Homo sapiens OX=9606 GN=LYPLA2 PE=1 SV=1 | 5 |
| O95373 | Importin-7 OS=Homo sapiens OX=9606 GN=IPO7 PE=1 SV=1 | 9 |
| O95379 | Tumor necrosis factor alpha-induced protein 8 OS=Homo sapiens OX=9606 GN=TNFAIP8 PE=1 SV=1 | 2 |
| O95394 | Phosphoacetylglucosamine mutase OS=Homo sapiens OX=9606 GN=PGM3 PE=1 SV=1 | 13 |
| O95433 | Activator of 90 kDa heat shock protein ATPase homolog 1 OS=Homo sapiens OX=9606 GN=AHSA1 PE=1 SV=1 | 22 |
| O95445 | Apolipoprotein M OS=Homo sapiens OX=9606 GN=APOM PE=1 SV=2 | 1 |
| O95456 | Proteasome assembly chaperone 1 OS=Homo sapiens OX=9606 GN=PSMG1 PE=1 SV=1 | 2 |
| O95470 | Sphingosine-1-phosphate lyase 1 OS=Homo sapiens OX=9606 GN=SGPL1 PE=1 SV=3 | 4 |
| O95487 | Protein transport protein Sec24B OS=Homo sapiens OX=9606 GN=SEC24B PE=1 SV=2 | 1 |
| O95571 | Persulfide dioxygenase ETHE1, mitochondrial OS=Homo sapiens OX=9606 GN=ETHE1 PE=1 SV=2 | 11 |
| O95573 | Fatty acid CoA ligase Acsl3 OS=Homo sapiens OX=9606 GN=ACSL3 PE=1 SV=3 | 20 |
| O95630 | STAM-binding protein OS=Homo sapiens OX=9606 GN=STAMBP PE=1 SV=1 | 12 |
| O95671 | Probable bifunctional dTTP/UTP pyrophosphatase/methyltransferase protein OS=Homo sapiens OX=9606 GN=ASMTL PE=1 SV=3 | 25 |
| O95674 | Phosphatidate cytidylyltransferase 2 OS=Homo sapiens OX=9606 GN=CDS2 PE=1 SV=1 | 2 |
| O95678 | Keratin, type II cytoskeletal 75 OS=Homo sapiens OX=9606 GN=KRT75 PE=1 SV=2 | 1 |
| O95684 | Centrosomal protein 43 OS=Homo sapiens OX=9606 GN=CEP43 PE=1 SV=1 | 2 |
| O95716 | Ras-related protein Rab-3D OS=Homo sapiens OX=9606 GN=RAB3D PE=1 SV=1 | 7 |
| O95747 | Serine/threonine-protein kinase OSR1 OS=Homo sapiens OX=9606 GN=OXSR1 PE=1 SV=1 | 10 |
| O95749 | Geranylgeranyl pyrophosphate synthase OS=Homo sapiens OX=9606 GN=GGPS1 PE=1 SV=1 | 3 |
| O95777 | U6 snRNA-associated Sm-like protein LSm8 OS=Homo sapiens OX=9606 GN=LSM8 PE=1 SV=3 | 3 |
| O95782 | AP-2 complex subunit alpha-1 OS=Homo sapiens OX=9606 GN=AP2A1 PE=1 SV=3 | 11 |
| O95786 | Antiviral innate immune response receptor RIG-I OS=Homo sapiens OX=9606 GN=RIGI PE=1 SV=2 | 8 |
| O95793 | Double-stranded RNA-binding protein Staufen homolog 1 OS=Homo sapiens OX=9606 GN=STAU1 PE=1 SV=2 | 10 |
| O95816 | BAG family molecular chaperone regulator 2 OS=Homo sapiens OX=9606 GN=BAG2 PE=1 SV=1 | 2 |
| O95822 | Malonyl-CoA decarboxylase, mitochondrial OS=Homo sapiens OX=9606 GN=MLYCD PE=1 SV=3 | 10 |
| O95825 | Quinone oxidoreductase-like protein 1 OS=Homo sapiens OX=9606 GN=CRYZL1 PE=1 SV=2 | 2 |
| O95831 | Apoptosis-inducing factor 1, mitochondrial OS=Homo sapiens OX=9606 GN=AIFM1 PE=1 SV=1 | 26 |
| O95861 | 3'(2'),5'-bisphosphate nucleotidase 1 OS=Homo sapiens OX=9606 GN=BPNT1 PE=1 SV=1 | 5 |
| O95865 | N(G),N(G)-dimethylarginine dimethylaminohydrolase 2 OS=Homo sapiens OX=9606 GN=DDAH2 PE=1 SV=1 | 7 |
| O95867 | Lymphocyte antigen 6 complex locus protein G6c OS=Homo sapiens OX=9606 GN=LY6G6C PE=1 SV=1 | 2 |
| O95870 | Phosphatidylserine lipase ABHD16A OS=Homo sapiens OX=9606 GN=ABHD16A PE=1 SV=3 | 9 |
| O95881 | Thioredoxin domain-containing protein 12 OS=Homo sapiens OX=9606 GN=TXNDC12 PE=1 SV=1 | 5 |
| O95922 | Polyglutamylase complex subunit TTLL1 OS=Homo sapiens OX=9606 GN=TTLL1 PE=1 SV=1 | 1 |
| O95969 | Secretoglobin family 1D member 2 OS=Homo sapiens OX=9606 GN=SCGB1D2 PE=2 SV=1 | 3 |
| O95972 | Bone morphogenetic protein 15 OS=Homo sapiens OX=9606 GN=BMP15 PE=1 SV=2 | 15 |
| O95988 | T-cell leukemia/lymphoma protein 1B OS=Homo sapiens OX=9606 GN=TCL1B PE=1 SV=1 | 4 |
| O95989 | Diphosphoinositol polyphosphate phosphohydrolase 1 OS=Homo sapiens OX=9606 GN=NUDT3 PE=1 SV=1 | 3 |
| O95997 | Securin OS=Homo sapiens OX=9606 GN=PTTG1 PE=1 SV=1 | 2 |
| O96000 | NADH dehydrogenase [ubiquinone] 1 beta subcomplex subunit 10 OS=Homo sapiens OX=9606 GN=NDUFB10 PE=1 SV=3 | 5 |
| O96005 | Putative lipid scramblase CLPTM1 OS=Homo sapiens OX=9606 GN=CLPTM1 PE=1 SV=1 | 2 |
| O96007 | Molybdopterin synthase catalytic subunit OS=Homo sapiens OX=9606 GN=MOCS2 PE=1 SV=1 | 3 |
| O96008 | Mitochondrial import receptor subunit TOM40 homolog OS=Homo sapiens OX=9606 GN=TOMM40 PE=1 SV=1 | 6 |
| O96011 | Peroxisomal membrane protein 11B OS=Homo sapiens OX=9606 GN=PEX11B PE=1 SV=1 | 1 |
| O96019 | Actin-like protein 6A OS=Homo sapiens OX=9606 GN=ACTL6A PE=1 SV=1 | 4 |
| P00167 | Cytochrome b5 OS=Homo sapiens OX=9606 GN=CYB5A PE=1 SV=2 | 4 |
| P00367 | Glutamate dehydrogenase 1, mitochondrial OS=Homo sapiens OX=9606 GN=GLUD1 PE=1 SV=2 | 17 |
| P00387 | NADH-cytochrome b5 reductase 3 OS=Homo sapiens OX=9606 GN=CYB5R3 PE=1 SV=3 | 15 |
| P00390 | Glutathione reductase, mitochondrial OS=Homo sapiens OX=9606 GN=GSR PE=1 SV=2 | 17 |
| P00403 | Cytochrome c oxidase subunit 2 OS=Homo sapiens OX=9606 GN=MT-CO2 PE=1 SV=1 | 4 |
| P00450 | Ceruloplasmin OS=Homo sapiens OX=9606 GN=CP PE=1 SV=1 | 20 |
| P00488 | Coagulation factor XIII A chain OS=Homo sapiens OX=9606 GN=F13A1 PE=1 SV=5 | 8 |
| P00492 | Hypoxanthine-guanine phosphoribosyltransferase OS=Homo sapiens OX=9606 GN=HPRT1 PE=1 SV=2 | 11 |
| P00568 | Adenylate kinase isoenzyme 1 OS=Homo sapiens OX=9606 GN=AK1 PE=1 SV=3 | 12 |
| P00734 | Prothrombin OS=Homo sapiens OX=9606 GN=F2 PE=1 SV=2 | 9 |
| P00738 | Haptoglobin OS=Homo sapiens OX=9606 GN=HP PE=1 SV=1 | 30 |
| P00742 | Coagulation factor X OS=Homo sapiens OX=9606 GN=F10 PE=1 SV=2 | 1 |
| P00747 | Plasminogen OS=Homo sapiens OX=9606 GN=PLG PE=1 SV=2 | 6 |
| P00751 | Complement factor B OS=Homo sapiens OX=9606 GN=CFB PE=1 SV=2 | 8 |
| P00813 | Adenosine deaminase OS=Homo sapiens OX=9606 GN=ADA PE=1 SV=3 | 5 |
| P00966 | Argininosuccinate synthase OS=Homo sapiens OX=9606 GN=ASS1 PE=1 SV=2 | 1 |
| P01008 | Antithrombin-III OS=Homo sapiens OX=9606 GN=SERPINC1 PE=1 SV=1 | 14 |
| P01009 | Alpha-1-antitrypsin OS=Homo sapiens OX=9606 GN=SERPINA1 PE=1 SV=3 | 20 |
| P01011 | Alpha-1-antichymotrypsin OS=Homo sapiens OX=9606 GN=SERPINA3 PE=1 SV=2 | 9 |
| P01019 | Angiotensinogen OS=Homo sapiens OX=9606 GN=AGT PE=1 SV=3 | 6 |
| P01023 | Alpha-2-macroglobulin OS=Homo sapiens OX=9606 GN=A2M PE=1 SV=3 | 16 |
| P01024 | Complement C3 OS=Homo sapiens OX=9606 GN=C3 PE=1 SV=2 | 78 |
| P01031 | Complement C5 OS=Homo sapiens OX=9606 GN=C5 PE=1 SV=4 | 73 |
| P01037 | Cystatin-SN OS=Homo sapiens OX=9606 GN=CST1 PE=1 SV=3 | 2 |
| P01042 | Kininogen-1 OS=Homo sapiens OX=9606 GN=KNG1 PE=1 SV=2 | 15 |
| P01111 | GTPase NRas OS=Homo sapiens OX=9606 GN=NRAS PE=1 SV=1 | 6 |
| P01112 | GTPase HRas OS=Homo sapiens OX=9606 GN=HRAS PE=1 SV=1 | 1 |
| P01178 | Oxytocin-neurophysin 1 OS=Homo sapiens OX=9606 GN=OXT PE=1 SV=1 | 6 |
| P01619 | Immunoglobulin kappa variable 3-20 OS=Homo sapiens OX=9606 GN=IGKV3-20 PE=1 SV=2 | 1 |
| P01732 | T-cell surface glycoprotein CD8 alpha chain OS=Homo sapiens OX=9606 GN=CD8A PE=1 SV=1 | 1 |
| P01764;P01768;P0DP03 | Immunoglobulin heavy variable 3-23 OS=Homo sapiens OX=9606 GN=IGHV3-23 PE=1 SV=2 | 2 |
| P01780 | Immunoglobulin heavy variable 3-7 OS=Homo sapiens OX=9606 GN=IGHV3-7 PE=1 SV=2 | 2 |
| P01859 | Immunoglobulin heavy constant gamma 2 OS=Homo sapiens OX=9606 GN=IGHG2 PE=1 SV=2 | 7 |
| P01860 | Immunoglobulin heavy constant gamma 3 OS=Homo sapiens OX=9606 GN=IGHG3 PE=1 SV=2 | 4 |
| P01861 | Immunoglobulin heavy constant gamma 4 OS=Homo sapiens OX=9606 GN=IGHG4 PE=1 SV=1 | 2 |
| P01871 | Immunoglobulin heavy constant mu OS=Homo sapiens OX=9606 GN=IGHM PE=1 SV=4 | 11 |
| P01877 | Immunoglobulin heavy constant alpha 2 OS=Homo sapiens OX=9606 GN=IGHA2 PE=1 SV=4 | 1 |
| P01889 | HLA class I histocompatibility antigen, B alpha chain OS=Homo sapiens OX=9606 GN=HLA-B PE=1 SV=3 | 3 |
| P02042 | Hemoglobin subunit delta OS=Homo sapiens OX=9606 GN=HBD PE=1 SV=2 | 2 |
| P02100 | Hemoglobin subunit epsilon OS=Homo sapiens OX=9606 GN=HBE1 PE=1 SV=2 | 2 |
| P02462 | Collagen alpha-1(IV) chain OS=Homo sapiens OX=9606 GN=COL4A1 PE=1 SV=4 | 5 |
| P02649 | Apolipoprotein E OS=Homo sapiens OX=9606 GN=APOE PE=1 SV=1 | 14 |
| P02652 | Apolipoprotein A-II OS=Homo sapiens OX=9606 GN=APOA2 PE=1 SV=1 | 7 |
| P02654 | Apolipoprotein C-I OS=Homo sapiens OX=9606 GN=APOC1 PE=1 SV=1 | 3 |
| P02655 | Apolipoprotein C-II OS=Homo sapiens OX=9606 GN=APOC2 PE=1 SV=1 | 4 |
| P02656 | Apolipoprotein C-III OS=Homo sapiens OX=9606 GN=APOC3 PE=1 SV=1 | 3 |
| P02671 | Fibrinogen alpha chain OS=Homo sapiens OX=9606 GN=FGA PE=1 SV=2 | 18 |
| P02675 | Fibrinogen beta chain OS=Homo sapiens OX=9606 GN=FGB PE=1 SV=2 | 17 |
| P02679 | Fibrinogen gamma chain OS=Homo sapiens OX=9606 GN=FGG PE=1 SV=3 | 15 |
| P02743 | Serum amyloid P-component OS=Homo sapiens OX=9606 GN=APCS PE=1 SV=2 | 3 |
| P02747 | Complement C1q subcomponent subunit C OS=Homo sapiens OX=9606 GN=C1QC PE=1 SV=3 | 1 |
| P02748 | Complement component C9 OS=Homo sapiens OX=9606 GN=C9 PE=1 SV=2 | 5 |
| P02749 | Beta-2-glycoprotein 1 OS=Homo sapiens OX=9606 GN=APOH PE=1 SV=3 | 14 |
| P02750 | Leucine-rich alpha-2-glycoprotein OS=Homo sapiens OX=9606 GN=LRG1 PE=1 SV=2 | 9 |
| P02751 | Fibronectin OS=Homo sapiens OX=9606 GN=FN1 PE=1 SV=5 | 57 |
| P02753 | Retinol-binding protein 4 OS=Homo sapiens OX=9606 GN=RBP4 PE=1 SV=3 | 4 |
| P02760 | Protein AMBP OS=Homo sapiens OX=9606 GN=AMBP PE=1 SV=1 | 11 |
| P02763 | Alpha-1-acid glycoprotein 1 OS=Homo sapiens OX=9606 GN=ORM1 PE=1 SV=2 | 13 |
| P02765 | Alpha-2-HS-glycoprotein OS=Homo sapiens OX=9606 GN=AHSG PE=1 SV=2 | 9 |
| P02766 | Transthyretin OS=Homo sapiens OX=9606 GN=TTR PE=1 SV=1 | 12 |
| P02774 | Vitamin D-binding protein OS=Homo sapiens OX=9606 GN=GC PE=1 SV=2 | 20 |
| P02786 | Transferrin receptor protein 1 OS=Homo sapiens OX=9606 GN=TFRC PE=1 SV=2 | 9 |
| P02787 | Serotransferrin OS=Homo sapiens OX=9606 GN=TF PE=1 SV=4 | 47 |
| P02790 | Hemopexin OS=Homo sapiens OX=9606 GN=HPX PE=1 SV=2 | 27 |
| P02794 | Ferritin heavy chain OS=Homo sapiens OX=9606 GN=FTH1 PE=1 SV=2 | 11 |
| P03973 | Antileukoproteinase OS=Homo sapiens OX=9606 GN=SLPI PE=1 SV=2 | 1 |
| P04066 | Tissue alpha-L-fucosidase OS=Homo sapiens OX=9606 GN=FUCA1 PE=1 SV=4 | 7 |
| P04180 | Phosphatidylcholine-sterol acyltransferase OS=Homo sapiens OX=9606 GN=LCAT PE=1 SV=1 | 3 |
| P04196 | Histidine-rich glycoprotein OS=Homo sapiens OX=9606 GN=HRG PE=1 SV=1 | 5 |
| P04217 | Alpha-1B-glycoprotein OS=Homo sapiens OX=9606 GN=A1BG PE=1 SV=4 | 22 |
| P04278 | Sex hormone-binding globulin OS=Homo sapiens OX=9606 GN=SHBG PE=1 SV=2 | 2 |
| P04424 | Argininosuccinate lyase OS=Homo sapiens OX=9606 GN=ASL PE=1 SV=4 | 9 |
| P04439 | HLA class I histocompatibility antigen, A alpha chain OS=Homo sapiens OX=9606 GN=HLA-A PE=1 SV=2 | 3 |
| P04632 | Calpain small subunit 1 OS=Homo sapiens OX=9606 GN=CAPNS1 PE=1 SV=1 | 8 |
| P04843 | Dolichyl-diphosphooligosaccharide--protein glycosyltransferase subunit 1 OS=Homo sapiens OX=9606 GN=RPN1 PE=1 SV=1 | 30 |
| P04844 | Dolichyl-diphosphooligosaccharide--protein glycosyltransferase subunit 2 OS=Homo sapiens OX=9606 GN=RPN2 PE=1 SV=3 | 11 |
| P04899 | Guanine nucleotide-binding protein G(i) subunit alpha-2 OS=Homo sapiens OX=9606 GN=GNAI2 PE=1 SV=3 | 3 |
| P05023 | Sodium/potassium-transporting ATPase subunit alpha-1 OS=Homo sapiens OX=9606 GN=ATP1A1 PE=1 SV=1 | 35 |
| P05026 | Sodium/potassium-transporting ATPase subunit beta-1 OS=Homo sapiens OX=9606 GN=ATP1B1 PE=1 SV=1 | 8 |
| P05067 | Amyloid-beta precursor protein OS=Homo sapiens OX=9606 GN=APP PE=1 SV=3 | 5 |
| P05091 | Aldehyde dehydrogenase, mitochondrial OS=Homo sapiens OX=9606 GN=ALDH2 PE=1 SV=2 | 4 |
| P05155 | Plasma protease C1 inhibitor OS=Homo sapiens OX=9606 GN=SERPING1 PE=1 SV=2 | 16 |
| P05165 | Propionyl-CoA carboxylase alpha chain, mitochondrial OS=Homo sapiens OX=9606 GN=PCCA PE=1 SV=4 | 2 |
| P05166 | Propionyl-CoA carboxylase beta chain, mitochondrial OS=Homo sapiens OX=9606 GN=PCCB PE=1 SV=3 | 4 |
| P05198 | Eukaryotic translation initiation factor 2 subunit 1 OS=Homo sapiens OX=9606 GN=EIF2S1 PE=1 SV=3 | 10 |
| P05362 | Intercellular adhesion molecule 1 OS=Homo sapiens OX=9606 GN=ICAM1 PE=1 SV=2 | 16 |
| P05388 | 60S acidic ribosomal protein P0 OS=Homo sapiens OX=9606 GN=RPLP0 PE=1 SV=1 | 11 |
| P05452 | Tetranectin OS=Homo sapiens OX=9606 GN=CLEC3B PE=1 SV=3 | 1 |
| P05455 | Lupus La protein OS=Homo sapiens OX=9606 GN=SSB PE=1 SV=2 | 9 |
| P05543 | Thyroxine-binding globulin OS=Homo sapiens OX=9606 GN=SERPINA7 PE=1 SV=2 | 10 |
| P05546 | Heparin cofactor 2 OS=Homo sapiens OX=9606 GN=SERPIND1 PE=1 SV=3 | 3 |
| P05556 | Integrin beta-1 OS=Homo sapiens OX=9606 GN=ITGB1 PE=1 SV=2 | 21 |
| P05976;P08590 | Myosin light chain 1/3, skeletal muscle isoform OS=Homo sapiens OX=9606 GN=MYL1 PE=1 SV=3 | 1 |
| P06132 | Uroporphyrinogen decarboxylase OS=Homo sapiens OX=9606 GN=UROD PE=1 SV=2 | 4 |
| P06213 | Insulin receptor OS=Homo sapiens OX=9606 GN=INSR PE=1 SV=4 | 1 |
| P06280 | Alpha-galactosidase A OS=Homo sapiens OX=9606 GN=GLA PE=1 SV=1 | 4 |
| P06312 | Immunoglobulin kappa variable 4-1 OS=Homo sapiens OX=9606 GN=IGKV4-1 PE=1 SV=1 | 1 |
| P06454 | Prothymosin alpha OS=Homo sapiens OX=9606 GN=PTMA PE=1 SV=2 | 2 |
| P06493 | Cyclin-dependent kinase 1 OS=Homo sapiens OX=9606 GN=CDK1 PE=1 SV=3 | 11 |
| P06727 | Apolipoprotein A-IV OS=Homo sapiens OX=9606 GN=APOA4 PE=1 SV=4 | 8 |
| P06730 | Eukaryotic translation initiation factor 4E OS=Homo sapiens OX=9606 GN=EIF4E PE=1 SV=2 | 8 |
| P06732 | Creatine kinase M-type OS=Homo sapiens OX=9606 GN=CKM PE=1 SV=2 | 3 |
| P06744 | Glucose-6-phosphate isomerase OS=Homo sapiens OX=9606 GN=GPI PE=1 SV=4 | 25 |
| P06746 | DNA polymerase beta OS=Homo sapiens OX=9606 GN=POLB PE=1 SV=3 | 8 |
| P06756 | Integrin alpha-V OS=Homo sapiens OX=9606 GN=ITGAV PE=1 SV=2 | 22 |
| P07093 | Glia-derived nexin OS=Homo sapiens OX=9606 GN=SERPINE2 PE=1 SV=1 | 14 |
| P07099 | Epoxide hydrolase 1 OS=Homo sapiens OX=9606 GN=EPHX1 PE=1 SV=1 | 20 |
| P07108 | Acyl-CoA-binding protein OS=Homo sapiens OX=9606 GN=DBI PE=1 SV=2 | 5 |
| P07197 | Neurofilament medium polypeptide OS=Homo sapiens OX=9606 GN=NEFM PE=1 SV=3 | 1 |
| P07203 | Glutathione peroxidase 1 OS=Homo sapiens OX=9606 GN=GPX1 PE=1 SV=4 | 14 |
| P07205 | Phosphoglycerate kinase 2 OS=Homo sapiens OX=9606 GN=PGK2 PE=1 SV=3 | 1 |
| P07357 | Complement component C8 alpha chain OS=Homo sapiens OX=9606 GN=C8A PE=1 SV=2 | 1 |
| P07358 | Complement component C8 beta chain OS=Homo sapiens OX=9606 GN=C8B PE=1 SV=3 | 3 |
| P07478 | Trypsin-2 OS=Homo sapiens OX=9606 GN=PRSS2 PE=1 SV=1 | 2 |
| P07602 | Prosaposin OS=Homo sapiens OX=9606 GN=PSAP PE=1 SV=2 | 12 |
| P07738 | Bisphosphoglycerate mutase OS=Homo sapiens OX=9606 GN=BPGM PE=1 SV=2 | 24 |
| P07741 | Adenine phosphoribosyltransferase OS=Homo sapiens OX=9606 GN=APRT PE=1 SV=2 | 6 |
| P07814 | Bifunctional glutamate/proline--tRNA ligase OS=Homo sapiens OX=9606 GN=EPRS1 PE=1 SV=5 | 40 |
| P07858 | Cathepsin B OS=Homo sapiens OX=9606 GN=CTSB PE=1 SV=3 | 12 |
| P07902 | Galactose-1-phosphate uridylyltransferase OS=Homo sapiens OX=9606 GN=GALT PE=1 SV=3 | 7 |
| P07919 | Cytochrome b-c1 complex subunit 6, mitochondrial OS=Homo sapiens OX=9606 GN=UQCRH PE=1 SV=2 | 3 |
| P07942 | Laminin subunit beta-1 OS=Homo sapiens OX=9606 GN=LAMB1 PE=1 SV=2 | 34 |
| P07947 | Tyrosine-protein kinase Yes OS=Homo sapiens OX=9606 GN=YES1 PE=1 SV=3 | 6 |
| P07951 | Tropomyosin beta chain OS=Homo sapiens OX=9606 GN=TPM2 PE=1 SV=1 | 2 |
| P07996 | Thrombospondin-1 OS=Homo sapiens OX=9606 GN=THBS1 PE=1 SV=2 | 10 |
| P08069 | Insulin-like growth factor 1 receptor OS=Homo sapiens OX=9606 GN=IGF1R PE=1 SV=1 | 4 |
| P08133 | Annexin A6 OS=Homo sapiens OX=9606 GN=ANXA6 PE=1 SV=3 | 42 |
| P08134 | Rho-related GTP-binding protein RhoC OS=Homo sapiens OX=9606 GN=RHOC PE=1 SV=1 | 2 |
| P08174 | Complement decay-accelerating factor OS=Homo sapiens OX=9606 GN=CD55 PE=1 SV=4 | 4 |
| P08185 | Corticosteroid-binding globulin OS=Homo sapiens OX=9606 GN=SERPINA6 PE=1 SV=1 | 1 |
| P08195 | 4F2 cell-surface antigen heavy chain OS=Homo sapiens OX=9606 GN=SLC3A2 PE=1 SV=3 | 17 |
| P08236 | Beta-glucuronidase OS=Homo sapiens OX=9606 GN=GUSB PE=1 SV=2 | 1 |
| P08238 | Heat shock protein HSP 90-beta OS=Homo sapiens OX=9606 GN=HSP90AB1 PE=1 SV=4 | 29 |
| P08240 | Signal recognition particle receptor subunit alpha OS=Homo sapiens OX=9606 GN=SRPRA PE=1 SV=2 | 6 |
| P08246 | Neutrophil elastase OS=Homo sapiens OX=9606 GN=ELANE PE=1 SV=1 | 1 |
| P08263 | Glutathione S-transferase A1 OS=Homo sapiens OX=9606 GN=GSTA1 PE=1 SV=3 | 5 |
| P08311 | Cathepsin G OS=Homo sapiens OX=9606 GN=CTSG PE=1 SV=2 | 1 |
| P08559 | Pyruvate dehydrogenase E1 component subunit alpha, somatic form, mitochondrial OS=Homo sapiens OX=9606 GN=PDHA1 PE=1 SV=3 | 26 |
| P08571 | Monocyte differentiation antigen CD14 OS=Homo sapiens OX=9606 GN=CD14 PE=1 SV=2 | 3 |
| P08572 | Collagen alpha-2(IV) chain OS=Homo sapiens OX=9606 GN=COL4A2 PE=1 SV=4 | 5 |
| P08574 | Cytochrome c1, heme protein, mitochondrial OS=Homo sapiens OX=9606 GN=CYC1 PE=1 SV=3 | 7 |
| P08603 | Complement factor H OS=Homo sapiens OX=9606 GN=CFH PE=1 SV=4 | 13 |
| P08621 | U1 small nuclear ribonucleoprotein 70 kDa OS=Homo sapiens OX=9606 GN=SNRNP70 PE=1 SV=2 | 4 |
| P08697 | Alpha-2-antiplasmin OS=Homo sapiens OX=9606 GN=SERPINF2 PE=1 SV=3 | 4 |
| P08708 | 40S ribosomal protein S17 OS=Homo sapiens OX=9606 GN=RPS17 PE=1 SV=2 | 8 |
| P08754 | Guanine nucleotide-binding protein G(i) subunit alpha-3 OS=Homo sapiens OX=9606 GN=GNAI3 PE=1 SV=3 | 12 |
| P08758 | Annexin A5 OS=Homo sapiens OX=9606 GN=ANXA5 PE=1 SV=2 | 17 |
| P08833 | Insulin-like growth factor-binding protein 1 OS=Homo sapiens OX=9606 GN=IGFBP1 PE=1 SV=1 | 6 |
| P08962 | CD63 antigen OS=Homo sapiens OX=9606 GN=CD63 PE=1 SV=2 | 2 |
| P09110 | 3-ketoacyl-CoA thiolase, peroxisomal OS=Homo sapiens OX=9606 GN=ACAA1 PE=1 SV=2 | 20 |
| P09132 | Signal recognition particle 19 kDa protein OS=Homo sapiens OX=9606 GN=SRP19 PE=1 SV=3 | 2 |
| P09327 | Villin-1 OS=Homo sapiens OX=9606 GN=VIL1 PE=1 SV=4 | 5 |
| P09382 | Galectin-1 OS=Homo sapiens OX=9606 GN=LGALS1 PE=1 SV=2 | 11 |
| P09417 | Dihydropteridine reductase OS=Homo sapiens OX=9606 GN=QDPR PE=1 SV=2 | 12 |
| P09455 | Retinol-binding protein 1 OS=Homo sapiens OX=9606 GN=RBP1 PE=1 SV=2 | 8 |
| P09493 | Tropomyosin alpha-1 chain OS=Homo sapiens OX=9606 GN=TPM1 PE=1 SV=2 | 6 |
| P09496 | Clathrin light chain A OS=Homo sapiens OX=9606 GN=CLTA PE=1 SV=1 | 8 |
| P09497 | Clathrin light chain B OS=Homo sapiens OX=9606 GN=CLTB PE=1 SV=1 | 9 |
| P09601 | Heme oxygenase 1 OS=Homo sapiens OX=9606 GN=HMOX1 PE=1 SV=1 | 2 |
| P09622 | Dihydrolipoyl dehydrogenase, mitochondrial OS=Homo sapiens OX=9606 GN=DLD PE=1 SV=2 | 26 |
| P09651 | Heterogeneous nuclear ribonucleoprotein A1 OS=Homo sapiens OX=9606 GN=HNRNPA1 PE=1 SV=5 | 12 |
| P09669 | Cytochrome c oxidase subunit 6C OS=Homo sapiens OX=9606 GN=COX6C PE=1 SV=2 | 4 |
| P09769 | Tyrosine-protein kinase Fgr OS=Homo sapiens OX=9606 GN=FGR PE=1 SV=2 | 1 |
| P09874 | Poly [ADP-ribose] polymerase 1 OS=Homo sapiens OX=9606 GN=PARP1 PE=1 SV=4 | 15 |
| P09960 | Leukotriene A-4 hydrolase OS=Homo sapiens OX=9606 GN=LTA4H PE=1 SV=2 | 22 |
| P09972 | Fructose-bisphosphate aldolase C OS=Homo sapiens OX=9606 GN=ALDOC PE=1 SV=2 | 3 |
| P0C024 | Peroxisomal coenzyme A diphosphatase NUDT7 OS=Homo sapiens OX=9606 GN=NUDT7 PE=1 SV=1 | 2 |
| P0C0L4 | Complement C4-A OS=Homo sapiens OX=9606 GN=C4A PE=1 SV=2 | 30 |
| P0C0L5 | Complement C4-B OS=Homo sapiens OX=9606 GN=C4B_2 PE=1 SV=2 | 1 |
| P0C1S8 | Wee1-like protein kinase 2 OS=Homo sapiens OX=9606 GN=WEE2 PE=1 SV=2 | 26 |
| P0C860 | Putative male-specific lethal-3 protein-like 2 OS=Homo sapiens OX=9606 GN=MSL3P1 PE=5 SV=1 | 1 |
| P0C870 | Bifunctional peptidase and (3S)-lysyl hydroxylase JMJD7 OS=Homo sapiens OX=9606 GN=JMJD7 PE=1 SV=1 | 11 |
| P0DI81;P0DI82 | Trafficking protein particle complex subunit 2 OS=Homo sapiens OX=9606 GN=TRAPPC2 PE=1 SV=1 | 1 |
| P0DJD7;P0DJD8;P0DJD9 | Pepsin A-4 OS=Homo sapiens OX=9606 GN=PGA4 PE=1 SV=1 | 1 |
| P0DME0;Q01105 | Protein SETSIP OS=Homo sapiens OX=9606 GN=SETSIP PE=1 SV=1 | 6 |
| P0DO97 | Coiled-coil domain-containing protein 192 OS=Homo sapiens OX=9606 GN=CCDC192 PE=4 SV=1 | 1 |
| P0DP57 | Secreted Ly-6/uPAR domain-containing protein 2 OS=Homo sapiens OX=9606 GN=SLURP2 PE=1 SV=1 | 3 |
| P0DTE7;P0DTE8;P0DUB6 | Alpha-amylase 1B OS=Homo sapiens OX=9606 GN=AMY1B PE=1 SV=1 | 1 |
| P0DTL6 | Zinc finger TRAF-type-containing protein 1 OS=Homo sapiens OX=9606 GN=ZFTRAF1 PE=2 SV=1 | 1 |
| P0DW85 | Histone H2A.N OS=Homo sapiens OX=9606 GN=H2BN1 PE=2 SV=1 | 3 |
| P10109 | Adrenodoxin, mitochondrial OS=Homo sapiens OX=9606 GN=FDX1 PE=1 SV=1 | 4 |
| P10155 | RNA-binding protein RO60 OS=Homo sapiens OX=9606 GN=RO60 PE=1 SV=2 | 1 |
| P10253 | Lysosomal alpha-glucosidase OS=Homo sapiens OX=9606 GN=GAA PE=1 SV=4 | 13 |
| P10321 | HLA class I histocompatibility antigen, C alpha chain OS=Homo sapiens OX=9606 GN=HLA-C PE=1 SV=3 | 1 |
| P10515 | Dihydrolipoyllysine-residue acetyltransferase component of pyruvate dehydrogenase complex, mitochondrial OS=Homo sapiens OX=9606 GN=DLAT PE=1 SV=3 | 17 |
| P10606 | Cytochrome c oxidase subunit 5B, mitochondrial OS=Homo sapiens OX=9606 GN=COX5B PE=1 SV=2 | 6 |
| P10643 | Complement component C7 OS=Homo sapiens OX=9606 GN=C7 PE=1 SV=2 | 1 |
| P10644 | cAMP-dependent protein kinase type I-alpha regulatory subunit OS=Homo sapiens OX=9606 GN=PRKAR1A PE=1 SV=1 | 15 |
| P10645 | Chromogranin-A OS=Homo sapiens OX=9606 GN=CHGA PE=1 SV=7 | 10 |
| P10721 | Mast/stem cell growth factor receptor Kit OS=Homo sapiens OX=9606 GN=KIT PE=1 SV=1 | 3 |
| P10768 | S-formylglutathione hydrolase OS=Homo sapiens OX=9606 GN=ESD PE=1 SV=2 | 10 |
| P10915 | Hyaluronan and proteoglycan link protein 1 OS=Homo sapiens OX=9606 GN=HAPLN1 PE=2 SV=2 | 2 |
| P11047 | Laminin subunit gamma-1 OS=Homo sapiens OX=9606 GN=LAMC1 PE=1 SV=3 | 23 |
| P11117 | Lysosomal acid phosphatase OS=Homo sapiens OX=9606 GN=ACP2 PE=1 SV=3 | 1 |
| P11172 | Uridine 5'-monophosphate synthase OS=Homo sapiens OX=9606 GN=UMPS PE=1 SV=1 | 20 |
| P11177 | Pyruvate dehydrogenase E1 component subunit beta, mitochondrial OS=Homo sapiens OX=9606 GN=PDHB PE=1 SV=3 | 13 |
| P11182 | Lipoamide acyltransferase component of branched-chain alpha-keto acid dehydrogenase complex, mitochondrial OS=Homo sapiens OX=9606 GN=DBT PE=1 SV=3 | 7 |
| P11216 | Glycogen phosphorylase, brain form OS=Homo sapiens OX=9606 GN=PYGB PE=1 SV=5 | 24 |
| P11217 | Glycogen phosphorylase, muscle form OS=Homo sapiens OX=9606 GN=PYGM PE=1 SV=6 | 1 |
| P11233 | Ras-related protein Ral-A OS=Homo sapiens OX=9606 GN=RALA PE=1 SV=1 | 5 |
| P11234 | Ras-related protein Ral-B OS=Homo sapiens OX=9606 GN=RALB PE=1 SV=1 | 12 |
| P11274;Q12979 | Breakpoint cluster region protein OS=Homo sapiens OX=9606 GN=BCR PE=1 SV=2 | 1 |
| P11310 | Medium-chain specific acyl-CoA dehydrogenase, mitochondrial OS=Homo sapiens OX=9606 GN=ACADM PE=1 SV=1 | 26 |
| P11362;P21802;P22607 | Fibroblast growth factor receptor 1 OS=Homo sapiens OX=9606 GN=FGFR1 PE=1 SV=3 | 1 |
| P11441 | Ubiquitin-like protein 4A OS=Homo sapiens OX=9606 GN=UBL4A PE=1 SV=1 | 3 |
| P11464;Q00888;Q00889;Q13046 | Pregnancy-specific beta-1-glycoprotein 1 OS=Homo sapiens OX=9606 GN=PSG1 PE=1 SV=1 | 1 |
| P11498 | Pyruvate carboxylase, mitochondrial OS=Homo sapiens OX=9606 GN=PC PE=1 SV=2 | 34 |
| P11678 | Eosinophil peroxidase OS=Homo sapiens OX=9606 GN=EPX PE=1 SV=2 | 1 |
| P11717 | Cation-independent mannose-6-phosphate receptor OS=Homo sapiens OX=9606 GN=IGF2R PE=1 SV=3 | 5 |
| P11766 | Alcohol dehydrogenase class-3 OS=Homo sapiens OX=9606 GN=ADH5 PE=1 SV=4 | 12 |
| P11908 | Ribose-phosphate pyrophosphokinase 2 OS=Homo sapiens OX=9606 GN=PRPS2 PE=1 SV=2 | 2 |
| P11940 | Polyadenylate-binding protein 1 OS=Homo sapiens OX=9606 GN=PABPC1 PE=1 SV=2 | 2 |
| P12004 | Proliferating cell nuclear antigen OS=Homo sapiens OX=9606 GN=PCNA PE=1 SV=1 | 19 |
| P12074 | Cytochrome c oxidase subunit 6A1, mitochondrial OS=Homo sapiens OX=9606 GN=COX6A1 PE=1 SV=4 | 1 |
| P12081 | Histidine--tRNA ligase, cytoplasmic OS=Homo sapiens OX=9606 GN=HARS1 PE=1 SV=2 | 24 |
| P12110 | Collagen alpha-2(VI) chain OS=Homo sapiens OX=9606 GN=COL6A2 PE=1 SV=4 | 4 |
| P12235 | ADP/ATP translocase 1 OS=Homo sapiens OX=9606 GN=SLC25A4 PE=1 SV=4 | 10 |
| P12236 | ADP/ATP translocase 3 OS=Homo sapiens OX=9606 GN=SLC25A6 PE=1 SV=4 | 2 |
| P12268 | Inosine-5'-monophosphate dehydrogenase 2 OS=Homo sapiens OX=9606 GN=IMPDH2 PE=1 SV=2 | 27 |
| P12270 | Nucleoprotein TPR OS=Homo sapiens OX=9606 GN=TPR PE=1 SV=3 | 19 |
| P12277 | Creatine kinase B-type OS=Homo sapiens OX=9606 GN=CKB PE=1 SV=1 | 21 |
| P12694 | 2-oxoisovalerate dehydrogenase subunit alpha, mitochondrial OS=Homo sapiens OX=9606 GN=BCKDHA PE=1 SV=2 | 7 |
| P12814 | Alpha-actinin-1 OS=Homo sapiens OX=9606 GN=ACTN1 PE=1 SV=2 | 19 |
| P12821 | Angiotensin-converting enzyme OS=Homo sapiens OX=9606 GN=ACE PE=1 SV=1 | 3 |
| P12830 | Cadherin-1 OS=Homo sapiens OX=9606 GN=CDH1 PE=1 SV=3 | 2 |
| P12883 | Myosin-7 OS=Homo sapiens OX=9606 GN=MYH7 PE=1 SV=5 | 1 |
| P12955 | Xaa-Pro dipeptidase OS=Homo sapiens OX=9606 GN=PEPD PE=1 SV=3 | 3 |
| P12956 | X-ray repair cross-complementing protein 6 OS=Homo sapiens OX=9606 GN=XRCC6 PE=1 SV=2 | 17 |
| P13010 | X-ray repair cross-complementing protein 5 OS=Homo sapiens OX=9606 GN=XRCC5 PE=1 SV=3 | 7 |
| P13073 | Cytochrome c oxidase subunit 4 isoform 1, mitochondrial OS=Homo sapiens OX=9606 GN=COX4I1 PE=1 SV=1 | 10 |
| P13164;Q01628;Q01629 | Interferon-induced transmembrane protein 1 OS=Homo sapiens OX=9606 GN=IFITM1 PE=1 SV=3 | 1 |
| P13196 | 5-aminolevulinate synthase, non-specific, mitochondrial OS=Homo sapiens OX=9606 GN=ALAS1 PE=1 SV=2 | 1 |
| P13284 | Gamma-interferon-inducible lysosomal thiol reductase OS=Homo sapiens OX=9606 GN=IFI30 PE=1 SV=3 | 1 |
| P13473 | Lysosome-associated membrane glycoprotein 2 OS=Homo sapiens OX=9606 GN=LAMP2 PE=1 SV=2 | 7 |
| P13489 | Ribonuclease inhibitor OS=Homo sapiens OX=9606 GN=RNH1 PE=1 SV=2 | 21 |
| P13611 | Versican core protein OS=Homo sapiens OX=9606 GN=VCAN PE=1 SV=3 | 11 |
| P13612 | Integrin alpha-4 OS=Homo sapiens OX=9606 GN=ITGA4 PE=1 SV=3 | 1 |
| P13671 | Complement component C6 OS=Homo sapiens OX=9606 GN=C6 PE=1 SV=3 | 2 |
| P13674 | Prolyl 4-hydroxylase subunit alpha-1 OS=Homo sapiens OX=9606 GN=P4HA1 PE=1 SV=2 | 6 |
| P13686 | Tartrate-resistant acid phosphatase type 5 OS=Homo sapiens OX=9606 GN=ACP5 PE=1 SV=3 | 1 |
| P13693 | Translationally-controlled tumor protein OS=Homo sapiens OX=9606 GN=TPT1 PE=1 SV=1 | 8 |
| P13716 | Delta-aminolevulinic acid dehydratase OS=Homo sapiens OX=9606 GN=ALAD PE=1 SV=1 | 3 |
| P13797 | Plastin-3 OS=Homo sapiens OX=9606 GN=PLS3 PE=1 SV=4 | 8 |
| P13861 | cAMP-dependent protein kinase type II-alpha regulatory subunit OS=Homo sapiens OX=9606 GN=PRKAR2A PE=1 SV=2 | 19 |
| P13928;Q5VT79 | Annexin A8 OS=Homo sapiens OX=9606 GN=ANXA8 PE=1 SV=3 | 3 |
| P13929 | Beta-enolase OS=Homo sapiens OX=9606 GN=ENO3 PE=1 SV=5 | 2 |
| P13987 | CD59 glycoprotein OS=Homo sapiens OX=9606 GN=CD59 PE=1 SV=1 | 6 |
| P14174 | Macrophage migration inhibitory factor OS=Homo sapiens OX=9606 GN=MIF PE=1 SV=4 | 2 |
| P14209 | CD99 antigen OS=Homo sapiens OX=9606 GN=CD99 PE=1 SV=1 | 2 |
| P14314 | Glucosidase 2 subunit beta OS=Homo sapiens OX=9606 GN=PRKCSH PE=1 SV=2 | 18 |
| P14317 | Hematopoietic lineage cell-specific protein OS=Homo sapiens OX=9606 GN=HCLS1 PE=1 SV=3 | 1 |
| P14324 | Farnesyl pyrophosphate synthase OS=Homo sapiens OX=9606 GN=FDPS PE=1 SV=4 | 6 |
| P14384 | Carboxypeptidase M OS=Homo sapiens OX=9606 GN=CPM PE=1 SV=2 | 6 |
| P14406 | Cytochrome c oxidase subunit 7A2, mitochondrial OS=Homo sapiens OX=9606 GN=COX7A2 PE=1 SV=1 | 2 |
| P14616 | Insulin receptor-related protein OS=Homo sapiens OX=9606 GN=INSRR PE=1 SV=2 | 1 |
| P14635 | G2/mitotic-specific cyclin-B1 OS=Homo sapiens OX=9606 GN=CCNB1 PE=1 SV=1 | 5 |
| P14649 | Myosin light chain 6B OS=Homo sapiens OX=9606 GN=MYL6B PE=1 SV=1 | 7 |
| P14678;P63162 | Small nuclear ribonucleoprotein-associated proteins B and B' OS=Homo sapiens OX=9606 GN=SNRPB PE=1 SV=2 | 2 |
| P14854 | Cytochrome c oxidase subunit 6B1 OS=Homo sapiens OX=9606 GN=COX6B1 PE=1 SV=2 | 3 |
| P14866 | Heterogeneous nuclear ribonucleoprotein L OS=Homo sapiens OX=9606 GN=HNRNPL PE=1 SV=2 | 7 |
| P14868 | Aspartate--tRNA ligase, cytoplasmic OS=Homo sapiens OX=9606 GN=DARS1 PE=1 SV=2 | 23 |
| P14927 | Cytochrome b-c1 complex subunit 7 OS=Homo sapiens OX=9606 GN=UQCRB PE=1 SV=2 | 8 |
| P15056 | Serine/threonine-protein kinase B-raf OS=Homo sapiens OX=9606 GN=BRAF PE=1 SV=4 | 1 |
| P15153 | Ras-related C3 botulinum toxin substrate 2 OS=Homo sapiens OX=9606 GN=RAC2 PE=1 SV=1 | 2 |
| P15170 | Eukaryotic peptide chain release factor GTP-binding subunit ERF3A OS=Homo sapiens OX=9606 GN=GSPT1 PE=1 SV=1 | 22 |
| P15289 | Arylsulfatase A OS=Homo sapiens OX=9606 GN=ARSA PE=1 SV=3 | 2 |
| P15309 | Prostatic acid phosphatase OS=Homo sapiens OX=9606 GN=ACP3 PE=1 SV=3 | 5 |
| P15328 | Folate receptor alpha OS=Homo sapiens OX=9606 GN=FOLR1 PE=1 SV=3 | 7 |
| P15374 | Ubiquitin carboxyl-terminal hydrolase isozyme L3 OS=Homo sapiens OX=9606 GN=UCHL3 PE=1 SV=1 | 14 |
| P15586 | N-acetylglucosamine-6-sulfatase OS=Homo sapiens OX=9606 GN=GNS PE=1 SV=3 | 15 |
| P15880 | 40S ribosomal protein S2 OS=Homo sapiens OX=9606 GN=RPS2 PE=1 SV=2 | 10 |
| P15927 | Replication protein A 32 kDa subunit OS=Homo sapiens OX=9606 GN=RPA2 PE=1 SV=1 | 7 |
| P16035 | Metalloproteinase inhibitor 2 OS=Homo sapiens OX=9606 GN=TIMP2 PE=1 SV=2 | 5 |
| P16070 | CD44 antigen OS=Homo sapiens OX=9606 GN=CD44 PE=1 SV=3 | 2 |
| P16083 | Ribosyldihydronicotinamide dehydrogenase [quinone] OS=Homo sapiens OX=9606 GN=NQO2 PE=1 SV=5 | 3 |
| P16112 | Aggrecan core protein OS=Homo sapiens OX=9606 GN=ACAN PE=1 SV=3 | 4 |
| P16152 | Carbonyl reductase [NADPH] 1 OS=Homo sapiens OX=9606 GN=CBR1 PE=1 SV=3 | 4 |
| P16219 | Short-chain specific acyl-CoA dehydrogenase, mitochondrial OS=Homo sapiens OX=9606 GN=ACADS PE=1 SV=1 | 12 |
| P16284 | Platelet endothelial cell adhesion molecule OS=Homo sapiens OX=9606 GN=PECAM1 PE=1 SV=2 | 3 |
| P16298 | Serine/threonine-protein phosphatase 2B catalytic subunit beta isoform OS=Homo sapiens OX=9606 GN=PPP3CB PE=1 SV=2 | 2 |
| P16333 | Cytoplasmic protein NCK1 OS=Homo sapiens OX=9606 GN=NCK1 PE=1 SV=1 | 6 |
| P16422 | Epithelial cell adhesion molecule OS=Homo sapiens OX=9606 GN=EPCAM PE=1 SV=2 | 1 |
| P16435 | NADPH--cytochrome P450 reductase OS=Homo sapiens OX=9606 GN=POR PE=1 SV=2 | 24 |
| P16444 | Dipeptidase 1 OS=Homo sapiens OX=9606 GN=DPEP1 PE=1 SV=3 | 12 |
| P16455 | Methylated-DNA--protein-cysteine methyltransferase OS=Homo sapiens OX=9606 GN=MGMT PE=1 SV=1 | 6 |
| P16615 | Sarcoplasmic/endoplasmic reticulum calcium ATPase 2 OS=Homo sapiens OX=9606 GN=ATP2A2 PE=1 SV=1 | 13 |
| P16671 | Platelet glycoprotein 4 OS=Homo sapiens OX=9606 GN=CD36 PE=1 SV=2 | 3 |
| P16870 | Carboxypeptidase E OS=Homo sapiens OX=9606 GN=CPE PE=1 SV=1 | 1 |
| P16885 | 1-phosphatidylinositol 4,5-bisphosphate phosphodiesterase gamma-2 OS=Homo sapiens OX=9606 GN=PLCG2 PE=1 SV=4 | 29 |
| P16949 | Stathmin OS=Homo sapiens OX=9606 GN=STMN1 PE=1 SV=3 | 8 |
| P16989 | Y-box-binding protein 3 OS=Homo sapiens OX=9606 GN=YBX3 PE=1 SV=4 | 3 |
| P17066 | Heat shock 70 kDa protein 6 OS=Homo sapiens OX=9606 GN=HSPA6 PE=1 SV=2 | 1 |
| P17152 | Transmembrane protein 11, mitochondrial OS=Homo sapiens OX=9606 GN=TMEM11 PE=1 SV=1 | 6 |
| P17301 | Integrin alpha-2 OS=Homo sapiens OX=9606 GN=ITGA2 PE=1 SV=2 | 22 |
| P17302 | Gap junction alpha-1 protein OS=Homo sapiens OX=9606 GN=GJA1 PE=1 SV=2 | 7 |
| P17405 | Sphingomyelin phosphodiesterase OS=Homo sapiens OX=9606 GN=SMPD1 PE=1 SV=5 | 1 |
| P17568 | NADH dehydrogenase [ubiquinone] 1 beta subcomplex subunit 7 OS=Homo sapiens OX=9606 GN=NDUFB7 PE=1 SV=4 | 3 |
| P17655 | Calpain-2 catalytic subunit OS=Homo sapiens OX=9606 GN=CAPN2 PE=1 SV=6 | 20 |
| P17707 | S-adenosylmethionine decarboxylase proenzyme OS=Homo sapiens OX=9606 GN=AMD1 PE=1 SV=2 | 1 |
| P17812 | CTP synthase 1 OS=Homo sapiens OX=9606 GN=CTPS1 PE=1 SV=2 | 10 |
| P17844 | Probable ATP-dependent RNA helicase DDX5 OS=Homo sapiens OX=9606 GN=DDX5 PE=1 SV=1 | 1 |
| P17858 | ATP-dependent 6-phosphofructokinase, liver type OS=Homo sapiens OX=9606 GN=PFKL PE=1 SV=6 | 25 |
| P17931 | Galectin-3 OS=Homo sapiens OX=9606 GN=LGALS3 PE=1 SV=5 | 5 |
| P17936 | Insulin-like growth factor-binding protein 3 OS=Homo sapiens OX=9606 GN=IGFBP3 PE=1 SV=2 | 2 |
| P17980 | 26S proteasome regulatory subunit 6A OS=Homo sapiens OX=9606 GN=PSMC3 PE=1 SV=3 | 28 |
| P18031 | Tyrosine-protein phosphatase non-receptor type 1 OS=Homo sapiens OX=9606 GN=PTPN1 PE=1 SV=1 | 15 |
| P18065 | Insulin-like growth factor-binding protein 2 OS=Homo sapiens OX=9606 GN=IGFBP2 PE=1 SV=2 | 10 |
| P18077 | 60S ribosomal protein L35a OS=Homo sapiens OX=9606 GN=RPL35A PE=1 SV=2 | 3 |
| P18084 | Integrin beta-5 OS=Homo sapiens OX=9606 GN=ITGB5 PE=1 SV=1 | 5 |
| P18085 | ADP-ribosylation factor 4 OS=Homo sapiens OX=9606 GN=ARF4 PE=1 SV=3 | 2 |
| P18124 | 60S ribosomal protein L7 OS=Homo sapiens OX=9606 GN=RPL7 PE=1 SV=1 | 9 |
| P18510 | Interleukin-1 receptor antagonist protein OS=Homo sapiens OX=9606 GN=IL1RN PE=1 SV=1 | 3 |
| P18564 | Integrin beta-6 OS=Homo sapiens OX=9606 GN=ITGB6 PE=1 SV=2 | 1 |
| P18615 | Negative elongation factor E OS=Homo sapiens OX=9606 GN=NELFE PE=1 SV=3 | 1 |
| P18621 | 60S ribosomal protein L17 OS=Homo sapiens OX=9606 GN=RPL17 PE=1 SV=3 | 6 |
| P18669 | Phosphoglycerate mutase 1 OS=Homo sapiens OX=9606 GN=PGAM1 PE=1 SV=2 | 13 |
| P18859 | ATP synthase-coupling factor 6, mitochondrial OS=Homo sapiens OX=9606 GN=ATP5PF PE=1 SV=1 | 3 |
| P18887 | DNA repair protein XRCC1 OS=Homo sapiens OX=9606 GN=XRCC1 PE=1 SV=3 | 7 |
| P19012 | Keratin, type I cytoskeletal 15 OS=Homo sapiens OX=9606 GN=KRT15 PE=1 SV=3 | 6 |
| P19022 | Cadherin-2 OS=Homo sapiens OX=9606 GN=CDH2 PE=1 SV=4 | 5 |
| P19087 | Guanine nucleotide-binding protein G(t) subunit alpha-2 OS=Homo sapiens OX=9606 GN=GNAT2 PE=1 SV=4 | 3 |
| P19338 | Nucleolin OS=Homo sapiens OX=9606 GN=NCL PE=1 SV=3 | 12 |
| P19367 | Hexokinase-1 OS=Homo sapiens OX=9606 GN=HK1 PE=1 SV=3 | 4 |
| P19387 | DNA-directed RNA polymerase II subunit RPB3 OS=Homo sapiens OX=9606 GN=POLR2C PE=1 SV=2 | 3 |
| P19388 | DNA-directed RNA polymerases I, II, and III subunit RPABC1 OS=Homo sapiens OX=9606 GN=POLR2E PE=1 SV=4 | 1 |
| P19404 | NADH dehydrogenase [ubiquinone] flavoprotein 2, mitochondrial OS=Homo sapiens OX=9606 GN=NDUFV2 PE=1 SV=2 | 7 |
| P19419 | ETS domain-containing protein Elk-1 OS=Homo sapiens OX=9606 GN=ELK1 PE=1 SV=2 | 1 |
| P19623 | Spermidine synthase OS=Homo sapiens OX=9606 GN=SRM PE=1 SV=1 | 14 |
| P19652 | Alpha-1-acid glycoprotein 2 OS=Homo sapiens OX=9606 GN=ORM2 PE=1 SV=2 | 8 |
| P19823 | Inter-alpha-trypsin inhibitor heavy chain H2 OS=Homo sapiens OX=9606 GN=ITIH2 PE=1 SV=2 | 20 |
| P19827 | Inter-alpha-trypsin inhibitor heavy chain H1 OS=Homo sapiens OX=9606 GN=ITIH1 PE=1 SV=3 | 20 |
| P19957 | Elafin OS=Homo sapiens OX=9606 GN=PI3 PE=1 SV=3 | 1 |
| P19971 | Thymidine phosphorylase OS=Homo sapiens OX=9606 GN=TYMP PE=1 SV=2 | 6 |
| P20020 | Plasma membrane calcium-transporting ATPase 1 OS=Homo sapiens OX=9606 GN=ATP2B1 PE=1 SV=4 | 25 |
| P20023 | Complement receptor type 2 OS=Homo sapiens OX=9606 GN=CR2 PE=1 SV=2 | 2 |
| P20042 | Eukaryotic translation initiation factor 2 subunit 2 OS=Homo sapiens OX=9606 GN=EIF2S2 PE=1 SV=2 | 2 |
| P20132 | L-serine dehydratase/L-threonine deaminase OS=Homo sapiens OX=9606 GN=SDS PE=1 SV=2 | 3 |
| P20290 | Transcription factor BTF3 OS=Homo sapiens OX=9606 GN=BTF3 PE=1 SV=1 | 1 |
| P20336 | Ras-related protein Rab-3A OS=Homo sapiens OX=9606 GN=RAB3A PE=1 SV=1 | 2 |
| P20338 | Ras-related protein Rab-4A OS=Homo sapiens OX=9606 GN=RAB4A PE=1 SV=3 | 5 |
| P20339 | Ras-related protein Rab-5A OS=Homo sapiens OX=9606 GN=RAB5A PE=1 SV=2 | 3 |
| P20594 | Atrial natriuretic peptide receptor 2 OS=Homo sapiens OX=9606 GN=NPR2 PE=1 SV=1 | 1 |
| P20645 | Cation-dependent mannose-6-phosphate receptor OS=Homo sapiens OX=9606 GN=M6PR PE=1 SV=1 | 5 |
| P20674 | Cytochrome c oxidase subunit 5A, mitochondrial OS=Homo sapiens OX=9606 GN=COX5A PE=1 SV=2 | 3 |
| P20700 | Lamin-B1 OS=Homo sapiens OX=9606 GN=LMNB1 PE=1 SV=2 | 8 |
| P20810 | Calpastatin OS=Homo sapiens OX=9606 GN=CAST PE=1 SV=4 | 8 |
| P20839 | Inosine-5'-monophosphate dehydrogenase 1 OS=Homo sapiens OX=9606 GN=IMPDH1 PE=1 SV=2 | 3 |
| P20929 | Nebulin OS=Homo sapiens OX=9606 GN=NEB PE=1 SV=5 | 3 |
| P20933 | N(4)-(beta-N-acetylglucosaminyl)-L-asparaginase OS=Homo sapiens OX=9606 GN=AGA PE=1 SV=2 | 1 |
| P20936 | Ras GTPase-activating protein 1 OS=Homo sapiens OX=9606 GN=RASA1 PE=1 SV=1 | 13 |
| P21217;P51993;Q11128 | 3-galactosyl-N-acetylglucosaminide 4-alpha-L-fucosyltransferase FUT3 OS=Homo sapiens OX=9606 GN=FUT3 PE=1 SV=1 | 1 |
| P21246 | Pleiotrophin OS=Homo sapiens OX=9606 GN=PTN PE=1 SV=1 | 1 |
| P21266 | Glutathione S-transferase Mu 3 OS=Homo sapiens OX=9606 GN=GSTM3 PE=1 SV=3 | 25 |
| P21281 | V-type proton ATPase subunit B, brain isoform OS=Homo sapiens OX=9606 GN=ATP6V1B2 PE=1 SV=3 | 26 |
| P21283 | V-type proton ATPase subunit C 1 OS=Homo sapiens OX=9606 GN=ATP6V1C1 PE=1 SV=4 | 8 |
| P21291 | Cysteine and glycine-rich protein 1 OS=Homo sapiens OX=9606 GN=CSRP1 PE=1 SV=3 | 10 |
| P21359 | Neurofibromin OS=Homo sapiens OX=9606 GN=NF1 PE=1 SV=2 | 2 |
| P21399 | Cytoplasmic aconitate hydratase OS=Homo sapiens OX=9606 GN=ACO1 PE=1 SV=3 | 35 |
| P21695 | Glycerol-3-phosphate dehydrogenase [NAD(+)], cytoplasmic OS=Homo sapiens OX=9606 GN=GPD1 PE=1 SV=4 | 24 |
| P21796 | Voltage-dependent anion-selective channel protein 1 OS=Homo sapiens OX=9606 GN=VDAC1 PE=1 SV=2 | 20 |
| P21912 | Succinate dehydrogenase [ubiquinone] iron-sulfur subunit, mitochondrial OS=Homo sapiens OX=9606 GN=SDHB PE=1 SV=3 | 9 |
| P21926 | CD9 antigen OS=Homo sapiens OX=9606 GN=CD9 PE=1 SV=4 | 4 |
| P21953 | 2-oxoisovalerate dehydrogenase subunit beta, mitochondrial OS=Homo sapiens OX=9606 GN=BCKDHB PE=1 SV=2 | 2 |
| P21980 | Protein-glutamine gamma-glutamyltransferase 2 OS=Homo sapiens OX=9606 GN=TGM2 PE=1 SV=2 | 1 |
| P22033 | Methylmalonyl-CoA mutase, mitochondrial OS=Homo sapiens OX=9606 GN=MMUT PE=1 SV=4 | 24 |
| P22059 | Oxysterol-binding protein 1 OS=Homo sapiens OX=9606 GN=OSBP PE=1 SV=1 | 20 |
| P22087 | rRNA 2'-O-methyltransferase fibrillarin OS=Homo sapiens OX=9606 GN=FBL PE=1 SV=2 | 1 |
| P22102 | Trifunctional purine biosynthetic protein adenosine-3 OS=Homo sapiens OX=9606 GN=GART PE=1 SV=1 | 3 |
| P22304 | Iduronate 2-sulfatase OS=Homo sapiens OX=9606 GN=IDS PE=1 SV=1 | 1 |
| P22307 | Sterol carrier protein 2 OS=Homo sapiens OX=9606 GN=SCP2 PE=1 SV=2 | 18 |
| P22314 | Ubiquitin-like modifier-activating enzyme 1 OS=Homo sapiens OX=9606 GN=UBA1 PE=1 SV=3 | 36 |
| P22352 | Glutathione peroxidase 3 OS=Homo sapiens OX=9606 GN=GPX3 PE=1 SV=2 | 1 |
| P22392 | Nucleoside diphosphate kinase B OS=Homo sapiens OX=9606 GN=NME2 PE=1 SV=1 | 7 |
| P22532 | Small proline-rich protein 2D OS=Homo sapiens OX=9606 GN=SPRR2D PE=2 SV=2 | 3 |
| P22570 | NADPH:adrenodoxin oxidoreductase, mitochondrial OS=Homo sapiens OX=9606 GN=FDXR PE=1 SV=3 | 29 |
| P22626 | Heterogeneous nuclear ribonucleoproteins A2/B1 OS=Homo sapiens OX=9606 GN=HNRNPA2B1 PE=1 SV=2 | 13 |
| P22676 | Calretinin OS=Homo sapiens OX=9606 GN=CALB2 PE=2 SV=2 | 6 |
| P22695 | Cytochrome b-c1 complex subunit 2, mitochondrial OS=Homo sapiens OX=9606 GN=UQCRC2 PE=1 SV=3 | 13 |
| P22830 | Ferrochelatase, mitochondrial OS=Homo sapiens OX=9606 GN=FECH PE=1 SV=2 | 13 |
| P22897 | Macrophage mannose receptor 1 OS=Homo sapiens OX=9606 GN=MRC1 PE=1 SV=1 | 3 |
| P23142 | Fibulin-1 OS=Homo sapiens OX=9606 GN=FBLN1 PE=1 SV=4 | 12 |
| P23229 | Integrin alpha-6 OS=Homo sapiens OX=9606 GN=ITGA6 PE=1 SV=5 | 23 |
| P23246 | Splicing factor, proline- and glutamine-rich OS=Homo sapiens OX=9606 GN=SFPQ PE=1 SV=2 | 8 |
| P23258 | Tubulin gamma-1 chain OS=Homo sapiens OX=9606 GN=TUBG1 PE=1 SV=2 | 18 |
| P23381 | Tryptophan--tRNA ligase, cytoplasmic OS=Homo sapiens OX=9606 GN=WARS1 PE=1 SV=2 | 21 |
| P23396 | 40S ribosomal protein S3 OS=Homo sapiens OX=9606 GN=RPS3 PE=1 SV=2 | 12 |
| P23434 | Glycine cleavage system H protein, mitochondrial OS=Homo sapiens OX=9606 GN=GCSH PE=1 SV=2 | 2 |
| P23443 | Ribosomal protein S6 kinase beta-1 OS=Homo sapiens OX=9606 GN=RPS6KB1 PE=1 SV=2 | 2 |
| P23588 | Eukaryotic translation initiation factor 4B OS=Homo sapiens OX=9606 GN=EIF4B PE=1 SV=2 | 6 |
| P23634 | Plasma membrane calcium-transporting ATPase 4 OS=Homo sapiens OX=9606 GN=ATP2B4 PE=1 SV=2 | 6 |
| P23786 | Carnitine O-palmitoyltransferase 2, mitochondrial OS=Homo sapiens OX=9606 GN=CPT2 PE=1 SV=2 | 13 |
| P23919 | Thymidylate kinase OS=Homo sapiens OX=9606 GN=DTYMK PE=1 SV=4 | 12 |
| P23921 | Ribonucleoside-diphosphate reductase large subunit OS=Homo sapiens OX=9606 GN=RRM1 PE=1 SV=1 | 4 |
| P24534 | Elongation factor 1-beta OS=Homo sapiens OX=9606 GN=EEF1B2 PE=1 SV=3 | 7 |
| P24539 | ATP synthase F(0) complex subunit B1, mitochondrial OS=Homo sapiens OX=9606 GN=ATP5PB PE=1 SV=2 | 14 |
| P24592 | Insulin-like growth factor-binding protein 6 OS=Homo sapiens OX=9606 GN=IGFBP6 PE=1 SV=1 | 4 |
| P24666 | Low molecular weight phosphotyrosine protein phosphatase OS=Homo sapiens OX=9606 GN=ACP1 PE=1 SV=3 | 8 |
| P25098;P35626 | Beta-adrenergic receptor kinase 1 OS=Homo sapiens OX=9606 GN=GRK2 PE=1 SV=2 | 1 |
| P25205 | DNA replication licensing factor MCM3 OS=Homo sapiens OX=9606 GN=MCM3 PE=1 SV=3 | 5 |
| P25398 | 40S ribosomal protein S12 OS=Homo sapiens OX=9606 GN=RPS12 PE=1 SV=3 | 7 |
| P25685 | DnaJ homolog subfamily B member 1 OS=Homo sapiens OX=9606 GN=DNAJB1 PE=1 SV=4 | 21 |
| P26006 | Integrin alpha-3 OS=Homo sapiens OX=9606 GN=ITGA3 PE=1 SV=5 | 1 |
| P26196 | Probable ATP-dependent RNA helicase DDX6 OS=Homo sapiens OX=9606 GN=DDX6 PE=1 SV=2 | 26 |
| P26232 | Catenin alpha-2 OS=Homo sapiens OX=9606 GN=CTNNA2 PE=1 SV=5 | 1 |
| P26358 | DNA (cytosine-5)-methyltransferase 1 OS=Homo sapiens OX=9606 GN=DNMT1 PE=1 SV=2 | 70 |
| P26368 | Splicing factor U2AF 65 kDa subunit OS=Homo sapiens OX=9606 GN=U2AF2 PE=1 SV=4 | 3 |
| P26373 | 60S ribosomal protein L13 OS=Homo sapiens OX=9606 GN=RPL13 PE=1 SV=4 | 8 |
| P26440 | Isovaleryl-CoA dehydrogenase, mitochondrial OS=Homo sapiens OX=9606 GN=IVD PE=1 SV=2 | 15 |
| P26583 | High mobility group protein B2 OS=Homo sapiens OX=9606 GN=HMGB2 PE=1 SV=2 | 7 |
| P26599 | Polypyrimidine tract-binding protein 1 OS=Homo sapiens OX=9606 GN=PTBP1 PE=1 SV=2 | 1 |
| P26639 | Threonine--tRNA ligase 1, cytoplasmic OS=Homo sapiens OX=9606 GN=TARS1 PE=1 SV=3 | 11 |
| P26640 | Valine--tRNA ligase OS=Homo sapiens OX=9606 GN=VARS1 PE=1 SV=4 | 16 |
| P27105 | Stomatin OS=Homo sapiens OX=9606 GN=STOM PE=1 SV=3 | 13 |
| P27144 | Adenylate kinase 4, mitochondrial OS=Homo sapiens OX=9606 GN=AK4 PE=1 SV=1 | 15 |
| P27169 | Serum paraoxonase/arylesterase 1 OS=Homo sapiens OX=9606 GN=PON1 PE=1 SV=3 | 3 |
| P27348 | 14-3-3 protein theta OS=Homo sapiens OX=9606 GN=YWHAQ PE=1 SV=1 | 12 |
| P27361 | Mitogen-activated protein kinase 3 OS=Homo sapiens OX=9606 GN=MAPK3 PE=1 SV=4 | 6 |
| P27448;Q7KZI7;Q9P0L2 | MAP/microtubule affinity-regulating kinase 3 OS=Homo sapiens OX=9606 GN=MARK3 PE=1 SV=5 | 1 |
| P27449 | V-type proton ATPase 16 kDa proteolipid subunit c OS=Homo sapiens OX=9606 GN=ATP6V0C PE=1 SV=1 | 1 |
| P27635 | 60S ribosomal protein L10 OS=Homo sapiens OX=9606 GN=RPL10 PE=1 SV=5 | 8 |
| P27694 | Replication protein A 70 kDa DNA-binding subunit OS=Homo sapiens OX=9606 GN=RPA1 PE=1 SV=2 | 18 |
| P27708 | CAD protein OS=Homo sapiens OX=9606 GN=CAD PE=1 SV=3 | 42 |
| P27815 | cAMP-specific 3',5'-cyclic phosphodiesterase 4A OS=Homo sapiens OX=9606 GN=PDE4A PE=1 SV=3 | 1 |
| P27986 | Phosphatidylinositol 3-kinase regulatory subunit alpha OS=Homo sapiens OX=9606 GN=PIK3R1 PE=1 SV=2 | 6 |
| P28161 | Glutathione S-transferase Mu 2 OS=Homo sapiens OX=9606 GN=GSTM2 PE=1 SV=2 | 6 |
| P28288 | ATP-binding cassette sub-family D member 3 OS=Homo sapiens OX=9606 GN=ABCD3 PE=1 SV=1 | 7 |
| P28329 | Choline O-acetyltransferase OS=Homo sapiens OX=9606 GN=CHAT PE=1 SV=4 | 3 |
| P28331 | NADH-ubiquinone oxidoreductase 75 kDa subunit, mitochondrial OS=Homo sapiens OX=9606 GN=NDUFS1 PE=1 SV=3 | 24 |
| P28340 | DNA polymerase delta catalytic subunit OS=Homo sapiens OX=9606 GN=POLD1 PE=1 SV=2 | 2 |
| P28482 | Mitogen-activated protein kinase 1 OS=Homo sapiens OX=9606 GN=MAPK1 PE=1 SV=3 | 16 |
| P28676 | Grancalcin OS=Homo sapiens OX=9606 GN=GCA PE=1 SV=2 | 6 |
| P28799 | Progranulin OS=Homo sapiens OX=9606 GN=GRN PE=1 SV=2 | 8 |
| P28838 | Cytosol aminopeptidase OS=Homo sapiens OX=9606 GN=LAP3 PE=1 SV=3 | 27 |
| P29034 | Protein S100-A2 OS=Homo sapiens OX=9606 GN=S100A2 PE=1 SV=3 | 1 |
| P29083 | General transcription factor IIE subunit 1 OS=Homo sapiens OX=9606 GN=GTF2E1 PE=1 SV=2 | 5 |
| P29144 | Tripeptidyl-peptidase 2 OS=Homo sapiens OX=9606 GN=TPP2 PE=1 SV=4 | 26 |
| P29218 | Inositol monophosphatase 1 OS=Homo sapiens OX=9606 GN=IMPA1 PE=1 SV=1 | 14 |
| P29590 | Protein PML OS=Homo sapiens OX=9606 GN=PML PE=1 SV=3 | 1 |
| P29622 | Kallistatin OS=Homo sapiens OX=9606 GN=SERPINA4 PE=1 SV=3 | 2 |
| P29762 | Cellular retinoic acid-binding protein 1 OS=Homo sapiens OX=9606 GN=CRABP1 PE=1 SV=2 | 9 |
| P29966 | Myristoylated alanine-rich C-kinase substrate OS=Homo sapiens OX=9606 GN=MARCKS PE=1 SV=4 | 5 |
| P29992 | Guanine nucleotide-binding protein subunit alpha-11 OS=Homo sapiens OX=9606 GN=GNA11 PE=1 SV=2 | 2 |
| P30038 | Delta-1-pyrroline-5-carboxylate dehydrogenase, mitochondrial OS=Homo sapiens OX=9606 GN=ALDH4A1 PE=1 SV=3 | 25 |
| P30040 | Endoplasmic reticulum resident protein 29 OS=Homo sapiens OX=9606 GN=ERP29 PE=1 SV=4 | 13 |
| P30044 | Peroxiredoxin-5, mitochondrial OS=Homo sapiens OX=9606 GN=PRDX5 PE=1 SV=4 | 13 |
| P30046 | D-dopachrome decarboxylase OS=Homo sapiens OX=9606 GN=DDT PE=1 SV=3 | 4 |
| P30047 | GTP cyclohydrolase 1 feedback regulatory protein OS=Homo sapiens OX=9606 GN=GCHFR PE=1 SV=3 | 5 |
| P30048 | Thioredoxin-dependent peroxide reductase, mitochondrial OS=Homo sapiens OX=9606 GN=PRDX3 PE=1 SV=3 | 10 |
| P30049 | ATP synthase subunit delta, mitochondrial OS=Homo sapiens OX=9606 GN=ATP5F1D PE=1 SV=2 | 3 |
| P30084 | Enoyl-CoA hydratase, mitochondrial OS=Homo sapiens OX=9606 GN=ECHS1 PE=1 SV=4 | 17 |
| P30085 | UMP-CMP kinase OS=Homo sapiens OX=9606 GN=CMPK1 PE=1 SV=3 | 16 |
| P30153 | Serine/threonine-protein phosphatase 2A 65 kDa regulatory subunit A alpha isoform OS=Homo sapiens OX=9606 GN=PPP2R1A PE=1 SV=4 | 21 |
| P30405 | Peptidyl-prolyl cis-trans isomerase F, mitochondrial OS=Homo sapiens OX=9606 GN=PPIF PE=1 SV=1 | 7 |
| P30419 | Glycylpeptide N-tetradecanoyltransferase 1 OS=Homo sapiens OX=9606 GN=NMT1 PE=1 SV=2 | 4 |
| P30519 | Heme oxygenase 2 OS=Homo sapiens OX=9606 GN=HMOX2 PE=1 SV=2 | 15 |
| P30520 | Adenylosuccinate synthetase isozyme 2 OS=Homo sapiens OX=9606 GN=ADSS2 PE=1 SV=3 | 5 |
| P30530 | Tyrosine-protein kinase receptor UFO OS=Homo sapiens OX=9606 GN=AXL PE=1 SV=4 | 8 |
| P30533 | Alpha-2-macroglobulin receptor-associated protein OS=Homo sapiens OX=9606 GN=LRPAP1 PE=1 SV=1 | 23 |
| P30566 | Adenylosuccinate lyase OS=Homo sapiens OX=9606 GN=ADSL PE=1 SV=2 | 24 |
| P30622 | CAP-Gly domain-containing linker protein 1 OS=Homo sapiens OX=9606 GN=CLIP1 PE=1 SV=2 | 8 |
| P30626 | Sorcin OS=Homo sapiens OX=9606 GN=SRI PE=1 SV=1 | 8 |
| P30837 | Aldehyde dehydrogenase X, mitochondrial OS=Homo sapiens OX=9606 GN=ALDH1B1 PE=1 SV=4 | 3 |
| P31040 | Succinate dehydrogenase [ubiquinone] flavoprotein subunit, mitochondrial OS=Homo sapiens OX=9606 GN=SDHA PE=1 SV=2 | 20 |
| P31150 | Rab GDP dissociation inhibitor alpha OS=Homo sapiens OX=9606 GN=GDI1 PE=1 SV=2 | 23 |
| P31153 | S-adenosylmethionine synthase isoform type-2 OS=Homo sapiens OX=9606 GN=MAT2A PE=1 SV=1 | 8 |
| P31327 | Carbamoyl-phosphate synthase [ammonia], mitochondrial OS=Homo sapiens OX=9606 GN=CPS1 PE=1 SV=2 | 6 |
| P31689 | DnaJ homolog subfamily A member 1 OS=Homo sapiens OX=9606 GN=DNAJA1 PE=1 SV=2 | 16 |
| P31751 | RAC-beta serine/threonine-protein kinase OS=Homo sapiens OX=9606 GN=AKT2 PE=1 SV=2 | 15 |
| P31930 | Cytochrome b-c1 complex subunit 1, mitochondrial OS=Homo sapiens OX=9606 GN=UQCRC1 PE=1 SV=3 | 26 |
| P31937 | 3-hydroxyisobutyrate dehydrogenase, mitochondrial OS=Homo sapiens OX=9606 GN=HIBADH PE=1 SV=2 | 7 |
| P31939 | Bifunctional purine biosynthesis protein ATIC OS=Homo sapiens OX=9606 GN=ATIC PE=1 SV=3 | 33 |
| P31942 | Heterogeneous nuclear ribonucleoprotein H3 OS=Homo sapiens OX=9606 GN=HNRNPH3 PE=1 SV=2 | 5 |
| P31946 | 14-3-3 protein beta/alpha OS=Homo sapiens OX=9606 GN=YWHAB PE=1 SV=3 | 7 |
| P32121 | Beta-arrestin-2 OS=Homo sapiens OX=9606 GN=ARRB2 PE=1 SV=2 | 1 |
| P32320 | Cytidine deaminase OS=Homo sapiens OX=9606 GN=CDA PE=1 SV=2 | 3 |
| P32322 | Pyrroline-5-carboxylate reductase 1, mitochondrial OS=Homo sapiens OX=9606 GN=PYCR1 PE=1 SV=2 | 7 |
| P32929 | Cystathionine gamma-lyase OS=Homo sapiens OX=9606 GN=CTH PE=1 SV=3 | 3 |
| P32969 | 60S ribosomal protein L9 OS=Homo sapiens OX=9606 GN=RPL9P9 PE=1 SV=1 | 4 |
| P33121 | Long-chain-fatty-acid--CoA ligase 1 OS=Homo sapiens OX=9606 GN=ACSL1 PE=1 SV=1 | 14 |
| P33176 | Kinesin-1 heavy chain OS=Homo sapiens OX=9606 GN=KIF5B PE=1 SV=1 | 38 |
| P33240 | Cleavage stimulation factor subunit 2 OS=Homo sapiens OX=9606 GN=CSTF2 PE=1 SV=1 | 11 |
| P33316 | Deoxyuridine 5'-triphosphate nucleotidohydrolase, mitochondrial OS=Homo sapiens OX=9606 GN=DUT PE=1 SV=4 | 6 |
| P33552 | Cyclin-dependent kinases regulatory subunit 2 OS=Homo sapiens OX=9606 GN=CKS2 PE=1 SV=1 | 1 |
| P33764 | Protein S100-A3 OS=Homo sapiens OX=9606 GN=S100A3 PE=1 SV=1 | 1 |
| P33897 | ATP-binding cassette sub-family D member 1 OS=Homo sapiens OX=9606 GN=ABCD1 PE=1 SV=2 | 2 |
| P33981 | Dual specificity protein kinase TTK OS=Homo sapiens OX=9606 GN=TTK PE=1 SV=2 | 3 |
| P34059 | N-acetylgalactosamine-6-sulfatase OS=Homo sapiens OX=9606 GN=GALNS PE=1 SV=1 | 2 |
| P34896 | Serine hydroxymethyltransferase, cytosolic OS=Homo sapiens OX=9606 GN=SHMT1 PE=1 SV=1 | 10 |
| P34897 | Serine hydroxymethyltransferase, mitochondrial OS=Homo sapiens OX=9606 GN=SHMT2 PE=1 SV=3 | 31 |
| P34913 | Bifunctional epoxide hydrolase 2 OS=Homo sapiens OX=9606 GN=EPHX2 PE=1 SV=2 | 17 |
| P34932 | Heat shock 70 kDa protein 4 OS=Homo sapiens OX=9606 GN=HSPA4 PE=1 SV=4 | 62 |
| P34949 | Mannose-6-phosphate isomerase OS=Homo sapiens OX=9606 GN=MPI PE=1 SV=2 | 11 |
| P35080 | Profilin-2 OS=Homo sapiens OX=9606 GN=PFN2 PE=1 SV=3 | 7 |
| P35221 | Catenin alpha-1 OS=Homo sapiens OX=9606 GN=CTNNA1 PE=1 SV=1 | 19 |
| P35222 | Catenin beta-1 OS=Homo sapiens OX=9606 GN=CTNNB1 PE=1 SV=1 | 13 |
| P35237 | Serpin B6 OS=Homo sapiens OX=9606 GN=SERPINB6 PE=1 SV=3 | 6 |
| P35241 | Radixin OS=Homo sapiens OX=9606 GN=RDX PE=1 SV=1 | 15 |
| P35244 | Replication protein A 14 kDa subunit OS=Homo sapiens OX=9606 GN=RPA3 PE=1 SV=1 | 4 |
| P35249 | Replication factor C subunit 4 OS=Homo sapiens OX=9606 GN=RFC4 PE=1 SV=2 | 2 |
| P35443 | Thrombospondin-4 OS=Homo sapiens OX=9606 GN=THBS4 PE=1 SV=2 | 1 |
| P35542 | Serum amyloid A-4 protein OS=Homo sapiens OX=9606 GN=SAA4 PE=1 SV=2 | 2 |
| P35573 | Glycogen debranching enzyme OS=Homo sapiens OX=9606 GN=AGL PE=1 SV=3 | 2 |
| P35580 | Myosin-10 OS=Homo sapiens OX=9606 GN=MYH10 PE=1 SV=3 | 53 |
| P35606 | Coatomer subunit beta' OS=Homo sapiens OX=9606 GN=COPB2 PE=1 SV=2 | 24 |
| P35612 | Beta-adducin OS=Homo sapiens OX=9606 GN=ADD2 PE=1 SV=3 | 4 |
| P35613 | Basigin OS=Homo sapiens OX=9606 GN=BSG PE=1 SV=2 | 7 |
| P35658 | Nuclear pore complex protein Nup214 OS=Homo sapiens OX=9606 GN=NUP214 PE=1 SV=2 | 7 |
| P35749 | Myosin-11 OS=Homo sapiens OX=9606 GN=MYH11 PE=1 SV=3 | 38 |
| P35754 | Glutaredoxin-1 OS=Homo sapiens OX=9606 GN=GLRX PE=1 SV=2 | 6 |
| P35813 | Protein phosphatase 1A OS=Homo sapiens OX=9606 GN=PPM1A PE=1 SV=1 | 3 |
| P35914 | Hydroxymethylglutaryl-CoA lyase, mitochondrial OS=Homo sapiens OX=9606 GN=HMGCL PE=1 SV=2 | 13 |
| P35998 | 26S proteasome regulatory subunit 7 OS=Homo sapiens OX=9606 GN=PSMC2 PE=1 SV=3 | 25 |
| P36222 | Chitinase-3-like protein 1 OS=Homo sapiens OX=9606 GN=CHI3L1 PE=1 SV=2 | 2 |
| P36404 | ADP-ribosylation factor-like protein 2 OS=Homo sapiens OX=9606 GN=ARL2 PE=1 SV=4 | 13 |
| P36405 | ADP-ribosylation factor-like protein 3 OS=Homo sapiens OX=9606 GN=ARL3 PE=1 SV=2 | 12 |
| P36507 | Dual specificity mitogen-activated protein kinase kinase 2 OS=Homo sapiens OX=9606 GN=MAP2K2 PE=1 SV=1 | 10 |
| P36543 | V-type proton ATPase subunit E 1 OS=Homo sapiens OX=9606 GN=ATP6V1E1 PE=1 SV=1 | 11 |
| P36551 | Oxygen-dependent coproporphyrinogen-III oxidase, mitochondrial OS=Homo sapiens OX=9606 GN=CPOX PE=1 SV=3 | 28 |
| P36578 | 60S ribosomal protein L4 OS=Homo sapiens OX=9606 GN=RPL4 PE=1 SV=5 | 12 |
| P36639 | Oxidized purine nucleoside triphosphate hydrolase OS=Homo sapiens OX=9606 GN=NUDT1 PE=1 SV=4 | 2 |
| P36776 | Lon protease homolog, mitochondrial OS=Homo sapiens OX=9606 GN=LONP1 PE=1 SV=2 | 26 |
| P36871 | Phosphoglucomutase-1 OS=Homo sapiens OX=9606 GN=PGM1 PE=1 SV=3 | 21 |
| P36873 | Serine/threonine-protein phosphatase PP1-gamma catalytic subunit OS=Homo sapiens OX=9606 GN=PPP1CC PE=1 SV=1 | 1 |
| P36915 | Guanine nucleotide-binding protein-like 1 OS=Homo sapiens OX=9606 GN=GNL1 PE=1 SV=2 | 6 |
| P36952 | Serpin B5 OS=Homo sapiens OX=9606 GN=SERPINB5 PE=1 SV=2 | 10 |
| P36954 | DNA-directed RNA polymerase II subunit RPB9 OS=Homo sapiens OX=9606 GN=POLR2I PE=1 SV=1 | 1 |
| P36955 | Pigment epithelium-derived factor OS=Homo sapiens OX=9606 GN=SERPINF1 PE=1 SV=4 | 12 |
| P36969 | Phospholipid hydroperoxide glutathione peroxidase OS=Homo sapiens OX=9606 GN=GPX4 PE=1 SV=3 | 6 |
| P37108 | Signal recognition particle 14 kDa protein OS=Homo sapiens OX=9606 GN=SRP14 PE=1 SV=2 | 2 |
| P37198 | Nuclear pore glycoprotein p62 OS=Homo sapiens OX=9606 GN=NUP62 PE=1 SV=3 | 3 |
| P37235;P61601;P84074 | Hippocalcin-like protein 1 OS=Homo sapiens OX=9606 GN=HPCAL1 PE=1 SV=3 | 1 |
| P38117 | Electron transfer flavoprotein subunit beta OS=Homo sapiens OX=9606 GN=ETFB PE=1 SV=3 | 21 |
| P38159;Q96E39 | RNA-binding motif protein, X chromosome OS=Homo sapiens OX=9606 GN=RBMX PE=1 SV=3 | 1 |
| P38606 | V-type proton ATPase catalytic subunit A OS=Homo sapiens OX=9606 GN=ATP6V1A PE=1 SV=2 | 24 |
| P39019 | 40S ribosomal protein S19 OS=Homo sapiens OX=9606 GN=RPS19 PE=1 SV=2 | 9 |
| P39023 | 60S ribosomal protein L3 OS=Homo sapiens OX=9606 GN=RPL3 PE=1 SV=2 | 10 |
| P39210 | Protein Mpv17 OS=Homo sapiens OX=9606 GN=MPV17 PE=1 SV=1 | 1 |
| P39656 | Dolichyl-diphosphooligosaccharide--protein glycosyltransferase 48 kDa subunit OS=Homo sapiens OX=9606 GN=DDOST PE=1 SV=4 | 8 |
| P39687 | Acidic leucine-rich nuclear phosphoprotein 32 family member A OS=Homo sapiens OX=9606 GN=ANP32A PE=1 SV=1 | 5 |
| P39748 | Flap endonuclease 1 OS=Homo sapiens OX=9606 GN=FEN1 PE=1 SV=1 | 6 |
| P40121 | Macrophage-capping protein OS=Homo sapiens OX=9606 GN=CAPG PE=1 SV=2 | 2 |
| P40227 | T-complex protein 1 subunit zeta OS=Homo sapiens OX=9606 GN=CCT6A PE=1 SV=3 | 29 |
| P40429 | 60S ribosomal protein L13a OS=Homo sapiens OX=9606 GN=RPL13A PE=1 SV=2 | 5 |
| P40616 | ADP-ribosylation factor-like protein 1 OS=Homo sapiens OX=9606 GN=ARL1 PE=1 SV=1 | 5 |
| P40818 | Ubiquitin carboxyl-terminal hydrolase 8 OS=Homo sapiens OX=9606 GN=USP8 PE=1 SV=1 | 3 |
| P40855 | Peroxisomal biogenesis factor 19 OS=Homo sapiens OX=9606 GN=PEX19 PE=1 SV=1 | 1 |
| P41091 | Eukaryotic translation initiation factor 2 subunit 3 OS=Homo sapiens OX=9606 GN=EIF2S3 PE=1 SV=3 | 5 |
| P41208 | Centrin-2 OS=Homo sapiens OX=9606 GN=CETN2 PE=1 SV=1 | 3 |
| P41222 | Prostaglandin-H2 D-isomerase OS=Homo sapiens OX=9606 GN=PTGDS PE=1 SV=1 | 6 |
| P41227 | N-alpha-acetyltransferase 10 OS=Homo sapiens OX=9606 GN=NAA10 PE=1 SV=1 | 4 |
| P41236 | Protein phosphatase inhibitor 2 OS=Homo sapiens OX=9606 GN=PPP1R2 PE=1 SV=2 | 5 |
| P41250 | Glycine--tRNA ligase OS=Homo sapiens OX=9606 GN=GARS1 PE=1 SV=3 | 26 |
| P41252 | Isoleucine--tRNA ligase, cytoplasmic OS=Homo sapiens OX=9606 GN=IARS1 PE=1 SV=2 | 12 |
| P41567 | Eukaryotic translation initiation factor 1 OS=Homo sapiens OX=9606 GN=EIF1 PE=1 SV=1 | 1 |
| P41732 | Tetraspanin-7 OS=Homo sapiens OX=9606 GN=TSPAN7 PE=1 SV=2 | 2 |
| P42025 | Beta-centractin OS=Homo sapiens OX=9606 GN=ACTR1B PE=1 SV=1 | 16 |
| P42126 | Enoyl-CoA delta isomerase 1, mitochondrial OS=Homo sapiens OX=9606 GN=ECI1 PE=1 SV=1 | 8 |
| P42224 | Signal transducer and activator of transcription 1-alpha/beta OS=Homo sapiens OX=9606 GN=STAT1 PE=1 SV=2 | 10 |
| P42336 | Phosphatidylinositol 4,5-bisphosphate 3-kinase catalytic subunit alpha isoform OS=Homo sapiens OX=9606 GN=PIK3CA PE=1 SV=2 | 2 |
| P42338 | Phosphatidylinositol 4,5-bisphosphate 3-kinase catalytic subunit beta isoform OS=Homo sapiens OX=9606 GN=PIK3CB PE=1 SV=1 | 2 |
| P42345 | Serine/threonine-protein kinase mTOR OS=Homo sapiens OX=9606 GN=MTOR PE=1 SV=1 | 15 |
| P42566 | Epidermal growth factor receptor substrate 15 OS=Homo sapiens OX=9606 GN=EPS15 PE=1 SV=2 | 3 |
| P42574 | Caspase-3 OS=Homo sapiens OX=9606 GN=CASP3 PE=1 SV=2 | 4 |
| P42677 | 40S ribosomal protein S27 OS=Homo sapiens OX=9606 GN=RPS27 PE=1 SV=3 | 1 |
| P42704 | Leucine-rich PPR motif-containing protein, mitochondrial OS=Homo sapiens OX=9606 GN=LRPPRC PE=1 SV=3 | 63 |
| P42766 | 60S ribosomal protein L35 OS=Homo sapiens OX=9606 GN=RPL35 PE=1 SV=2 | 2 |
| P42785 | Lysosomal Pro-X carboxypeptidase OS=Homo sapiens OX=9606 GN=PRCP PE=1 SV=1 | 3 |
| P42892 | Endothelin-converting enzyme 1 OS=Homo sapiens OX=9606 GN=ECE1 PE=1 SV=2 | 9 |
| P43007 | Neutral amino acid transporter A OS=Homo sapiens OX=9606 GN=SLC1A4 PE=1 SV=1 | 6 |
| P43034 | Platelet-activating factor acetylhydrolase IB subunit beta OS=Homo sapiens OX=9606 GN=PAFAH1B1 PE=1 SV=2 | 23 |
| P43121 | Cell surface glycoprotein MUC18 OS=Homo sapiens OX=9606 GN=MCAM PE=1 SV=2 | 10 |
| P43155 | Carnitine O-acetyltransferase OS=Homo sapiens OX=9606 GN=CRAT PE=1 SV=5 | 7 |
| P43243 | Matrin-3 OS=Homo sapiens OX=9606 GN=MATR3 PE=1 SV=2 | 1 |
| P43251 | Biotinidase OS=Homo sapiens OX=9606 GN=BTD PE=1 SV=2 | 3 |
| P43304 | Glycerol-3-phosphate dehydrogenase, mitochondrial OS=Homo sapiens OX=9606 GN=GPD2 PE=1 SV=3 | 22 |
| P43487 | Ran-specific GTPase-activating protein OS=Homo sapiens OX=9606 GN=RANBP1 PE=1 SV=1 | 3 |
| P43490 | Nicotinamide phosphoribosyltransferase OS=Homo sapiens OX=9606 GN=NAMPT PE=1 SV=1 | 5 |
| P43652 | Afamin OS=Homo sapiens OX=9606 GN=AFM PE=1 SV=1 | 31 |
| P43686 | 26S proteasome regulatory subunit 6B OS=Homo sapiens OX=9606 GN=PSMC4 PE=1 SV=2 | 14 |
| P43897 | Elongation factor Ts, mitochondrial OS=Homo sapiens OX=9606 GN=TSFM PE=1 SV=2 | 14 |
| P45877 | Peptidyl-prolyl cis-trans isomerase C OS=Homo sapiens OX=9606 GN=PPIC PE=1 SV=1 | 2 |
| P45974 | Ubiquitin carboxyl-terminal hydrolase 5 OS=Homo sapiens OX=9606 GN=USP5 PE=1 SV=2 | 45 |
| P45985 | Dual specificity mitogen-activated protein kinase kinase 4 OS=Homo sapiens OX=9606 GN=MAP2K4 PE=1 SV=1 | 4 |
| P46019 | Phosphorylase b kinase regulatory subunit alpha, liver isoform OS=Homo sapiens OX=9606 GN=PHKA2 PE=1 SV=1 | 1 |
| P46060 | Ran GTPase-activating protein 1 OS=Homo sapiens OX=9606 GN=RANGAP1 PE=1 SV=1 | 6 |
| P46108 | Adapter molecule crk OS=Homo sapiens OX=9606 GN=CRK PE=1 SV=2 | 14 |
| P46109 | Crk-like protein OS=Homo sapiens OX=9606 GN=CRKL PE=1 SV=1 | 12 |
| P46199 | Translation initiation factor IF-2, mitochondrial OS=Homo sapiens OX=9606 GN=MTIF2 PE=1 SV=2 | 1 |
| P46379 | Large proline-rich protein BAG6 OS=Homo sapiens OX=9606 GN=BAG6 PE=1 SV=2 | 12 |
| P46736 | Lys-63-specific deubiquitinase BRCC36 OS=Homo sapiens OX=9606 GN=BRCC3 PE=1 SV=2 | 1 |
| P46776 | 60S ribosomal protein L27a OS=Homo sapiens OX=9606 GN=RPL27A PE=1 SV=2 | 5 |
| P46777 | 60S ribosomal protein L5 OS=Homo sapiens OX=9606 GN=RPL5 PE=1 SV=3 | 10 |
| P46778 | 60S ribosomal protein L21 OS=Homo sapiens OX=9606 GN=RPL21 PE=1 SV=2 | 4 |
| P46779 | 60S ribosomal protein L28 OS=Homo sapiens OX=9606 GN=RPL28 PE=1 SV=3 | 4 |
| P46781 | 40S ribosomal protein S9 OS=Homo sapiens OX=9606 GN=RPS9 PE=1 SV=3 | 9 |
| P46782 | 40S ribosomal protein S5 OS=Homo sapiens OX=9606 GN=RPS5 PE=1 SV=4 | 8 |
| P46783 | 40S ribosomal protein S10 OS=Homo sapiens OX=9606 GN=RPS10 PE=1 SV=1 | 5 |
| P46926 | Glucosamine-6-phosphate isomerase 1 OS=Homo sapiens OX=9606 GN=GNPDA1 PE=1 SV=1 | 10 |
| P46937 | Transcriptional coactivator YAP1 OS=Homo sapiens OX=9606 GN=YAP1 PE=1 SV=2 | 1 |
| P46940 | Ras GTPase-activating-like protein IQGAP1 OS=Homo sapiens OX=9606 GN=IQGAP1 PE=1 SV=1 | 28 |
| P46976 | Glycogenin-1 OS=Homo sapiens OX=9606 GN=GYG1 PE=1 SV=4 | 15 |
| P46977 | Dolichyl-diphosphooligosaccharide--protein glycosyltransferase subunit STT3A OS=Homo sapiens OX=9606 GN=STT3A PE=1 SV=2 | 7 |
| P47224 | Guanine nucleotide exchange factor MSS4 OS=Homo sapiens OX=9606 GN=RABIF PE=1 SV=2 | 3 |
| P47755 | F-actin-capping protein subunit alpha-2 OS=Homo sapiens OX=9606 GN=CAPZA2 PE=1 SV=3 | 11 |
| P47756 | F-actin-capping protein subunit beta OS=Homo sapiens OX=9606 GN=CAPZB PE=1 SV=5 | 18 |
| P47813 | Eukaryotic translation initiation factor 1A, X-chromosomal OS=Homo sapiens OX=9606 GN=EIF1AX PE=1 SV=2 | 7 |
| P47897 | Glutamine--tRNA ligase OS=Homo sapiens OX=9606 GN=QARS1 PE=1 SV=1 | 16 |
| P47914 | 60S ribosomal protein L29 OS=Homo sapiens OX=9606 GN=RPL29 PE=1 SV=2 | 3 |
| P47985 | Cytochrome b-c1 complex subunit Rieske, mitochondrial OS=Homo sapiens OX=9606 GN=UQCRFS1 PE=1 SV=2 | 11 |
| P48047 | ATP synthase subunit O, mitochondrial OS=Homo sapiens OX=9606 GN=ATP5PO PE=1 SV=1 | 11 |
| P48147 | Prolyl endopeptidase OS=Homo sapiens OX=9606 GN=PREP PE=1 SV=2 | 15 |
| P48163 | NADP-dependent malic enzyme OS=Homo sapiens OX=9606 GN=ME1 PE=1 SV=1 | 5 |
| P48426 | Phosphatidylinositol 5-phosphate 4-kinase type-2 alpha OS=Homo sapiens OX=9606 GN=PIP4K2A PE=1 SV=2 | 2 |
| P48444 | Coatomer subunit delta OS=Homo sapiens OX=9606 GN=ARCN1 PE=1 SV=1 | 34 |
| P48449 | Lanosterol synthase OS=Homo sapiens OX=9606 GN=LSS PE=1 SV=1 | 2 |
| P48506 | Glutamate--cysteine ligase catalytic subunit OS=Homo sapiens OX=9606 GN=GCLC PE=1 SV=2 | 12 |
| P48507 | Glutamate--cysteine ligase regulatory subunit OS=Homo sapiens OX=9606 GN=GCLM PE=1 SV=1 | 4 |
| P48556 | 26S proteasome non-ATPase regulatory subunit 8 OS=Homo sapiens OX=9606 GN=PSMD8 PE=1 SV=2 | 10 |
| P48637 | Glutathione synthetase OS=Homo sapiens OX=9606 GN=GSS PE=1 SV=1 | 26 |
| P48643 | T-complex protein 1 subunit epsilon OS=Homo sapiens OX=9606 GN=CCT5 PE=1 SV=1 | 38 |
| P48729 | Casein kinase I isoform alpha OS=Homo sapiens OX=9606 GN=CSNK1A1 PE=1 SV=2 | 2 |
| P48735 | Isocitrate dehydrogenase [NADP], mitochondrial OS=Homo sapiens OX=9606 GN=IDH2 PE=1 SV=2 | 18 |
| P48739 | Phosphatidylinositol transfer protein beta isoform OS=Homo sapiens OX=9606 GN=PITPNB PE=1 SV=2 | 2 |
| P49006 | MARCKS-related protein OS=Homo sapiens OX=9606 GN=MARCKSL1 PE=1 SV=2 | 2 |
| P49069 | Guided entry of tail-anchored proteins factor CAMLG OS=Homo sapiens OX=9606 GN=CAMLG PE=1 SV=1 | 4 |
| P49207 | 60S ribosomal protein L34 OS=Homo sapiens OX=9606 GN=RPL34 PE=1 SV=3 | 2 |
| P49247 | Ribose-5-phosphate isomerase OS=Homo sapiens OX=9606 GN=RPIA PE=1 SV=3 | 5 |
| P49257 | Protein ERGIC-53 OS=Homo sapiens OX=9606 GN=LMAN1 PE=1 SV=2 | 5 |
| P49321 | Nuclear autoantigenic sperm protein OS=Homo sapiens OX=9606 GN=NASP PE=1 SV=2 | 34 |
| P49327 | Fatty acid synthase OS=Homo sapiens OX=9606 GN=FASN PE=1 SV=3 | 49 |
| P49354 | Protein farnesyltransferase/geranylgeranyltransferase type-1 subunit alpha OS=Homo sapiens OX=9606 GN=FNTA PE=1 SV=1 | 6 |
| P49368 | T-complex protein 1 subunit gamma OS=Homo sapiens OX=9606 GN=CCT3 PE=1 SV=4 | 34 |
| P49406 | 39S ribosomal protein L19, mitochondrial OS=Homo sapiens OX=9606 GN=MRPL19 PE=1 SV=2 | 2 |
| P49419 | Alpha-aminoadipic semialdehyde dehydrogenase OS=Homo sapiens OX=9606 GN=ALDH7A1 PE=1 SV=5 | 31 |
| P49427 | Ubiquitin-conjugating enzyme E2 R1 OS=Homo sapiens OX=9606 GN=CDC34 PE=1 SV=2 | 3 |
| P49441 | Inositol polyphosphate 1-phosphatase OS=Homo sapiens OX=9606 GN=INPP1 PE=1 SV=1 | 14 |
| P49458 | Signal recognition particle 9 kDa protein OS=Homo sapiens OX=9606 GN=SRP9 PE=1 SV=2 | 4 |
| P49459 | Ubiquitin-conjugating enzyme E2 A OS=Homo sapiens OX=9606 GN=UBE2A PE=1 SV=2 | 1 |
| P49588 | Alanine--tRNA ligase, cytoplasmic OS=Homo sapiens OX=9606 GN=AARS1 PE=1 SV=2 | 26 |
| P49591 | Serine--tRNA ligase, cytoplasmic OS=Homo sapiens OX=9606 GN=SARS1 PE=1 SV=3 | 20 |
| P49593 | Protein phosphatase 1F OS=Homo sapiens OX=9606 GN=PPM1F PE=1 SV=3 | 22 |
| P49721 | Proteasome subunit beta type-2 OS=Homo sapiens OX=9606 GN=PSMB2 PE=1 SV=1 | 5 |
| P49736 | DNA replication licensing factor MCM2 OS=Homo sapiens OX=9606 GN=MCM2 PE=1 SV=4 | 18 |
| P49753;Q86TX2 | Acyl-coenzyme A thioesterase 2, mitochondrial OS=Homo sapiens OX=9606 GN=ACOT2 PE=1 SV=6 | 10 |
| P49755 | Transmembrane emp24 domain-containing protein 10 OS=Homo sapiens OX=9606 GN=TMED10 PE=1 SV=2 | 8 |
| P49757 | Protein numb homolog OS=Homo sapiens OX=9606 GN=NUMB PE=1 SV=2 | 6 |
| P49773 | Adenosine 5'-monophosphoramidase HINT1 OS=Homo sapiens OX=9606 GN=HINT1 PE=1 SV=2 | 11 |
| P49789 | Bis(5'-adenosyl)-triphosphatase OS=Homo sapiens OX=9606 GN=FHIT PE=1 SV=3 | 11 |
| P49792 | E3 SUMO-protein ligase RanBP2 OS=Homo sapiens OX=9606 GN=RANBP2 PE=1 SV=2 | 10 |
| P49821 | NADH dehydrogenase [ubiquinone] flavoprotein 1, mitochondrial OS=Homo sapiens OX=9606 GN=NDUFV1 PE=1 SV=4 | 12 |
| P49840 | Glycogen synthase kinase-3 alpha OS=Homo sapiens OX=9606 GN=GSK3A PE=1 SV=2 | 2 |
| P49902 | Cytosolic purine 5'-nucleotidase OS=Homo sapiens OX=9606 GN=NT5C2 PE=1 SV=1 | 1 |
| P49903 | Selenide, water dikinase 1 OS=Homo sapiens OX=9606 GN=SEPHS1 PE=1 SV=2 | 7 |
| P49915 | GMP synthase [glutamine-hydrolyzing] OS=Homo sapiens OX=9606 GN=GMPS PE=1 SV=1 | 2 |
| P49916 | DNA ligase 3 OS=Homo sapiens OX=9606 GN=LIG3 PE=1 SV=2 | 3 |
| P49959 | Double-strand break repair protein MRE11 OS=Homo sapiens OX=9606 GN=MRE11 PE=1 SV=3 | 4 |
| P50053 | Ketohexokinase OS=Homo sapiens OX=9606 GN=KHK PE=1 SV=2 | 12 |
| P50213 | Isocitrate dehydrogenase [NAD] subunit alpha, mitochondrial OS=Homo sapiens OX=9606 GN=IDH3A PE=1 SV=1 | 17 |
| P50336 | Protoporphyrinogen oxidase OS=Homo sapiens OX=9606 GN=PPOX PE=1 SV=1 | 13 |
| P50402 | Emerin OS=Homo sapiens OX=9606 GN=EMD PE=1 SV=1 | 2 |
| P50502 | Hsc70-interacting protein OS=Homo sapiens OX=9606 GN=ST13 PE=1 SV=2 | 13 |
| P50539 | Max-interacting protein 1 OS=Homo sapiens OX=9606 GN=MXI1 PE=1 SV=2 | 1 |
| P50542 | Peroxisomal targeting signal 1 receptor OS=Homo sapiens OX=9606 GN=PEX5 PE=1 SV=3 | 2 |
| P50552 | Vasodilator-stimulated phosphoprotein OS=Homo sapiens OX=9606 GN=VASP PE=1 SV=3 | 3 |
| P50570 | Dynamin-2 OS=Homo sapiens OX=9606 GN=DNM2 PE=1 SV=2 | 16 |
| P50579 | Methionine aminopeptidase 2 OS=Homo sapiens OX=9606 GN=METAP2 PE=1 SV=1 | 5 |
| P50583 | Bis(5'-nucleosyl)-tetraphosphatase [asymmetrical] OS=Homo sapiens OX=9606 GN=NUDT2 PE=1 SV=3 | 5 |
| P50747 | Biotin--protein ligase OS=Homo sapiens OX=9606 GN=HLCS PE=1 SV=1 | 2 |
| P50897 | Palmitoyl-protein thioesterase 1 OS=Homo sapiens OX=9606 GN=PPT1 PE=1 SV=1 | 3 |
| P50914 | 60S ribosomal protein L14 OS=Homo sapiens OX=9606 GN=RPL14 PE=1 SV=4 | 5 |
| P50991 | T-complex protein 1 subunit delta OS=Homo sapiens OX=9606 GN=CCT4 PE=1 SV=4 | 32 |
| P50995 | Annexin A11 OS=Homo sapiens OX=9606 GN=ANXA11 PE=1 SV=1 | 21 |
| P51114 | RNA-binding protein FXR1 OS=Homo sapiens OX=9606 GN=FXR1 PE=1 SV=3 | 15 |
| P51116 | RNA-binding protein FXR2 OS=Homo sapiens OX=9606 GN=FXR2 PE=1 SV=2 | 6 |
| P51148 | Ras-related protein Rab-5C OS=Homo sapiens OX=9606 GN=RAB5C PE=1 SV=2 | 3 |
| P51159 | Ras-related protein Rab-27A OS=Homo sapiens OX=9606 GN=RAB27A PE=1 SV=3 | 7 |
| P51398 | 28S ribosomal protein S29, mitochondrial OS=Homo sapiens OX=9606 GN=DAP3 PE=1 SV=1 | 4 |
| P51452 | Dual specificity protein phosphatase 3 OS=Homo sapiens OX=9606 GN=DUSP3 PE=1 SV=1 | 5 |
| P51531 | Probable global transcription activator SNF2L2 OS=Homo sapiens OX=9606 GN=SMARCA2 PE=1 SV=2 | 5 |
| P51532 | Transcription activator BRG1 OS=Homo sapiens OX=9606 GN=SMARCA4 PE=1 SV=2 | 1 |
| P51553 | Isocitrate dehydrogenase [NAD] subunit gamma, mitochondrial OS=Homo sapiens OX=9606 GN=IDH3G PE=1 SV=1 | 9 |
| P51570 | Galactokinase OS=Homo sapiens OX=9606 GN=GALK1 PE=1 SV=1 | 18 |
| P51572 | B-cell receptor-associated protein 31 OS=Homo sapiens OX=9606 GN=BCAP31 PE=1 SV=3 | 14 |
| P51648 | Aldehyde dehydrogenase family 3 member A2 OS=Homo sapiens OX=9606 GN=ALDH3A2 PE=1 SV=1 | 8 |
| P51649 | Succinate-semialdehyde dehydrogenase, mitochondrial OS=Homo sapiens OX=9606 GN=ALDH5A1 PE=1 SV=2 | 2 |
| P51659 | Peroxisomal multifunctional enzyme type 2 OS=Homo sapiens OX=9606 GN=HSD17B4 PE=1 SV=3 | 36 |
| P51665 | 26S proteasome non-ATPase regulatory subunit 7 OS=Homo sapiens OX=9606 GN=PSMD7 PE=1 SV=2 | 10 |
| P51688 | N-sulphoglucosamine sulphohydrolase OS=Homo sapiens OX=9606 GN=SGSH PE=1 SV=1 | 2 |
| P51795 | H(+)/Cl(-) exchange transporter 5 OS=Homo sapiens OX=9606 GN=CLCN5 PE=1 SV=2 | 1 |
| P51809 | Vesicle-associated membrane protein 7 OS=Homo sapiens OX=9606 GN=VAMP7 PE=1 SV=3 | 8 |
| P51812 | Ribosomal protein S6 kinase alpha-3 OS=Homo sapiens OX=9606 GN=RPS6KA3 PE=1 SV=1 | 19 |
| P51858 | Hepatoma-derived growth factor OS=Homo sapiens OX=9606 GN=HDGF PE=1 SV=1 | 4 |
| P51884 | Lumican OS=Homo sapiens OX=9606 GN=LUM PE=1 SV=2 | 2 |
| P51970 | NADH dehydrogenase [ubiquinone] 1 alpha subcomplex subunit 8 OS=Homo sapiens OX=9606 GN=NDUFA8 PE=1 SV=3 | 4 |
| P51991 | Heterogeneous nuclear ribonucleoprotein A3 OS=Homo sapiens OX=9606 GN=HNRNPA3 PE=1 SV=2 | 8 |
| P52272 | Heterogeneous nuclear ribonucleoprotein M OS=Homo sapiens OX=9606 GN=HNRNPM PE=1 SV=3 | 14 |
| P52292 | Importin subunit alpha-1 OS=Homo sapiens OX=9606 GN=KPNA2 PE=1 SV=1 | 8 |
| P52306 | Rap1 GTPase-GDP dissociation stimulator 1 OS=Homo sapiens OX=9606 GN=RAP1GDS1 PE=1 SV=3 | 7 |
| P52434 | DNA-directed RNA polymerases I, II, and III subunit RPABC3 OS=Homo sapiens OX=9606 GN=POLR2H PE=1 SV=4 | 3 |
| P52564 | Dual specificity mitogen-activated protein kinase kinase 6 OS=Homo sapiens OX=9606 GN=MAP2K6 PE=1 SV=1 | 6 |
| P52565 | Rho GDP-dissociation inhibitor 1 OS=Homo sapiens OX=9606 GN=ARHGDIA PE=1 SV=3 | 12 |
| P52594 | Arf-GAP domain and FG repeat-containing protein 1 OS=Homo sapiens OX=9606 GN=AGFG1 PE=1 SV=2 | 6 |
| P52597 | Heterogeneous nuclear ribonucleoprotein F OS=Homo sapiens OX=9606 GN=HNRNPF PE=1 SV=3 | 11 |
| P52758 | 2-iminobutanoate/2-iminopropanoate deaminase OS=Homo sapiens OX=9606 GN=RIDA PE=1 SV=1 | 1 |
| P52788 | Spermine synthase OS=Homo sapiens OX=9606 GN=SMS PE=1 SV=2 | 26 |
| P52815 | 39S ribosomal protein L12, mitochondrial OS=Homo sapiens OX=9606 GN=MRPL12 PE=1 SV=2 | 2 |
| P52888 | Thimet oligopeptidase OS=Homo sapiens OX=9606 GN=THOP1 PE=1 SV=2 | 14 |
| P52948 | Nuclear pore complex protein Nup98-Nup96 OS=Homo sapiens OX=9606 GN=NUP98 PE=1 SV=4 | 8 |
| P53007 | Tricarboxylate transport protein, mitochondrial OS=Homo sapiens OX=9606 GN=SLC25A1 PE=1 SV=2 | 12 |
| P53041 | Serine/threonine-protein phosphatase 5 OS=Homo sapiens OX=9606 GN=PPP5C PE=1 SV=1 | 17 |
| P53350 | Serine/threonine-protein kinase PLK1 OS=Homo sapiens OX=9606 GN=PLK1 PE=1 SV=1 | 5 |
| P53365 | Arfaptin-2 OS=Homo sapiens OX=9606 GN=ARFIP2 PE=1 SV=1 | 5 |
| P53367 | Arfaptin-1 OS=Homo sapiens OX=9606 GN=ARFIP1 PE=1 SV=2 | 2 |
| P53396 | ATP-citrate synthase OS=Homo sapiens OX=9606 GN=ACLY PE=1 SV=3 | 24 |
| P53597 | Succinate--CoA ligase [ADP/GDP-forming] subunit alpha, mitochondrial OS=Homo sapiens OX=9606 GN=SUCLG1 PE=1 SV=4 | 12 |
| P53602 | Diphosphomevalonate decarboxylase OS=Homo sapiens OX=9606 GN=MVD PE=1 SV=1 | 12 |
| P53609 | Geranylgeranyl transferase type-1 subunit beta OS=Homo sapiens OX=9606 GN=PGGT1B PE=1 SV=2 | 1 |
| P53611 | Geranylgeranyl transferase type-2 subunit beta OS=Homo sapiens OX=9606 GN=RABGGTB PE=1 SV=2 | 1 |
| P53618 | Coatomer subunit beta OS=Homo sapiens OX=9606 GN=COPB1 PE=1 SV=3 | 28 |
| P53621 | Coatomer subunit alpha OS=Homo sapiens OX=9606 GN=COPA PE=1 SV=2 | 30 |
| P53634 | Dipeptidyl peptidase 1 OS=Homo sapiens OX=9606 GN=CTSC PE=1 SV=2 | 7 |
| P53674 | Beta-crystallin B1 OS=Homo sapiens OX=9606 GN=CRYBB1 PE=1 SV=2 | 1 |
| P53677 | AP-3 complex subunit mu-2 OS=Homo sapiens OX=9606 GN=AP3M2 PE=1 SV=1 | 5 |
| P53680 | AP-2 complex subunit sigma OS=Homo sapiens OX=9606 GN=AP2S1 PE=1 SV=2 | 3 |
| P53701 | Holocytochrome c-type synthase OS=Homo sapiens OX=9606 GN=HCCS PE=1 SV=1 | 2 |
| P53778 | Mitogen-activated protein kinase 12 OS=Homo sapiens OX=9606 GN=MAPK12 PE=1 SV=3 | 2 |
| P53803 | DNA-directed RNA polymerases I, II, and III subunit RPABC4 OS=Homo sapiens OX=9606 GN=POLR2K PE=1 SV=1 | 1 |
| P53816 | Phospholipase A and acyltransferase 3 OS=Homo sapiens OX=9606 GN=PLAAT3 PE=1 SV=2 | 3 |
| P53985 | Monocarboxylate transporter 1 OS=Homo sapiens OX=9606 GN=SLC16A1 PE=1 SV=3 | 2 |
| P53990 | IST1 homolog OS=Homo sapiens OX=9606 GN=IST1 PE=1 SV=1 | 6 |
| P53992 | Protein transport protein Sec24C OS=Homo sapiens OX=9606 GN=SEC24C PE=1 SV=3 | 15 |
| P53999 | Activated RNA polymerase II transcriptional coactivator p15 OS=Homo sapiens OX=9606 GN=SUB1 PE=1 SV=3 | 7 |
| P54098 | DNA polymerase subunit gamma-1 OS=Homo sapiens OX=9606 GN=POLG PE=1 SV=1 | 12 |
| P54136 | Arginine--tRNA ligase, cytoplasmic OS=Homo sapiens OX=9606 GN=RARS1 PE=1 SV=2 | 32 |
| P54252 | Ataxin-3 OS=Homo sapiens OX=9606 GN=ATXN3 PE=1 SV=5 | 1 |
| P54289 | Voltage-dependent calcium channel subunit alpha-2/delta-1 OS=Homo sapiens OX=9606 GN=CACNA2D1 PE=1 SV=3 | 9 |
| P54577 | Tyrosine--tRNA ligase, cytoplasmic OS=Homo sapiens OX=9606 GN=YARS1 PE=1 SV=4 | 10 |
| P54578 | Ubiquitin carboxyl-terminal hydrolase 14 OS=Homo sapiens OX=9606 GN=USP14 PE=1 SV=3 | 25 |
| P54619 | 5'-AMP-activated protein kinase subunit gamma-1 OS=Homo sapiens OX=9606 GN=PRKAG1 PE=1 SV=1 | 2 |
| P54652 | Heat shock-related 70 kDa protein 2 OS=Homo sapiens OX=9606 GN=HSPA2 PE=1 SV=1 | 17 |
| P54709 | Sodium/potassium-transporting ATPase subunit beta-3 OS=Homo sapiens OX=9606 GN=ATP1B3 PE=1 SV=1 | 4 |
| P54727 | UV excision repair protein RAD23 homolog B OS=Homo sapiens OX=9606 GN=RAD23B PE=1 SV=1 | 8 |
| P54819 | Adenylate kinase 2, mitochondrial OS=Homo sapiens OX=9606 GN=AK2 PE=1 SV=2 | 16 |
| P54886 | Delta-1-pyrroline-5-carboxylate synthase OS=Homo sapiens OX=9606 GN=ALDH18A1 PE=1 SV=2 | 10 |
| P54920 | Alpha-soluble NSF attachment protein OS=Homo sapiens OX=9606 GN=NAPA PE=1 SV=3 | 23 |
| P55000 | Secreted Ly-6/uPAR-related protein 1 OS=Homo sapiens OX=9606 GN=SLURP1 PE=1 SV=2 | 2 |
| P55010 | Eukaryotic translation initiation factor 5 OS=Homo sapiens OX=9606 GN=EIF5 PE=1 SV=2 | 16 |
| P55011 | Solute carrier family 12 member 2 OS=Homo sapiens OX=9606 GN=SLC12A2 PE=1 SV=1 | 7 |
| P55036 | 26S proteasome non-ATPase regulatory subunit 4 OS=Homo sapiens OX=9606 GN=PSMD4 PE=1 SV=1 | 13 |
| P55039 | Developmentally-regulated GTP-binding protein 2 OS=Homo sapiens OX=9606 GN=DRG2 PE=1 SV=1 | 2 |
| P55060 | Exportin-2 OS=Homo sapiens OX=9606 GN=CSE1L PE=1 SV=3 | 26 |
| P55084 | Trifunctional enzyme subunit beta, mitochondrial OS=Homo sapiens OX=9606 GN=HADHB PE=1 SV=3 | 26 |
| P55145 | Mesencephalic astrocyte-derived neurotrophic factor OS=Homo sapiens OX=9606 GN=MANF PE=1 SV=3 | 11 |
| P55196 | Afadin OS=Homo sapiens OX=9606 GN=AFDN PE=1 SV=3 | 12 |
| P55209 | Nucleosome assembly protein 1-like 1 OS=Homo sapiens OX=9606 GN=NAP1L1 PE=1 SV=1 | 9 |
| P55212 | Caspase-6 OS=Homo sapiens OX=9606 GN=CASP6 PE=1 SV=2 | 3 |
| P55263 | Adenosine kinase OS=Homo sapiens OX=9606 GN=ADK PE=1 SV=2 | 11 |
| P55327 | Tumor protein D52 OS=Homo sapiens OX=9606 GN=TPD52 PE=1 SV=2 | 6 |
| P55735 | Protein SEC13 homolog OS=Homo sapiens OX=9606 GN=SEC13 PE=1 SV=3 | 5 |
| P55769 | NHP2-like protein 1 OS=Homo sapiens OX=9606 GN=SNU13 PE=1 SV=3 | 2 |
| P55786 | Puromycin-sensitive aminopeptidase OS=Homo sapiens OX=9606 GN=NPEPPS PE=1 SV=2 | 34 |
| P55789 | FAD-linked sulfhydryl oxidase ALR OS=Homo sapiens OX=9606 GN=GFER PE=1 SV=2 | 5 |
| P55795 | Heterogeneous nuclear ribonucleoprotein H2 OS=Homo sapiens OX=9606 GN=HNRNPH2 PE=1 SV=1 | 2 |
| P55809 | Succinyl-CoA:3-ketoacid coenzyme A transferase 1, mitochondrial OS=Homo sapiens OX=9606 GN=OXCT1 PE=1 SV=1 | 16 |
| P55884 | Eukaryotic translation initiation factor 3 subunit B OS=Homo sapiens OX=9606 GN=EIF3B PE=1 SV=3 | 12 |
| P56134 | ATP synthase subunit f, mitochondrial OS=Homo sapiens OX=9606 GN=ATP5MF PE=1 SV=3 | 2 |
| P56181 | NADH dehydrogenase [ubiquinone] flavoprotein 3, mitochondrial OS=Homo sapiens OX=9606 GN=NDUFV3 PE=1 SV=2 | 1 |
| P56192 | Methionine--tRNA ligase, cytoplasmic OS=Homo sapiens OX=9606 GN=MARS1 PE=1 SV=2 | 11 |
| P56199 | Integrin alpha-1 OS=Homo sapiens OX=9606 GN=ITGA1 PE=1 SV=2 | 4 |
| P56277 | Cx9C motif-containing protein 4 OS=Homo sapiens OX=9606 GN=CMC4 PE=1 SV=1 | 2 |
| P56279 | T-cell leukemia/lymphoma protein 1A OS=Homo sapiens OX=9606 GN=TCL1A PE=1 SV=1 | 9 |
| P56378 | ATP synthase subunit ATP5MJ, mitochondrial OS=Homo sapiens OX=9606 GN=ATP5MJ PE=1 SV=1 | 3 |
| P56385 | ATP synthase subunit e, mitochondrial OS=Homo sapiens OX=9606 GN=ATP5ME PE=1 SV=2 | 6 |
| P56556 | NADH dehydrogenase [ubiquinone] 1 alpha subcomplex subunit 6 OS=Homo sapiens OX=9606 GN=NDUFA6 PE=1 SV=4 | 2 |
| P56589 | Peroxisomal biogenesis factor 3 OS=Homo sapiens OX=9606 GN=PEX3 PE=1 SV=1 | 1 |
| P56937 | 3-keto-steroid reductase/17-beta-hydroxysteroid dehydrogenase 7 OS=Homo sapiens OX=9606 GN=HSD17B7 PE=1 SV=1 | 3 |
| P57058 | Hormonally up-regulated neu tumor-associated kinase OS=Homo sapiens OX=9606 GN=HUNK PE=1 SV=1 | 1 |
| P57076 | Cilia- and flagella-associated protein 298 OS=Homo sapiens OX=9606 GN=CFAP298 PE=1 SV=1 | 1 |
| P57105 | Synaptojanin-2-binding protein OS=Homo sapiens OX=9606 GN=SYNJ2BP PE=1 SV=2 | 3 |
| P57678 | Gem-associated protein 4 OS=Homo sapiens OX=9606 GN=GEMIN4 PE=1 SV=2 | 1 |
| P57721 | Poly(rC)-binding protein 3 OS=Homo sapiens OX=9606 GN=PCBP3 PE=1 SV=2 | 1 |
| P57740 | Nuclear pore complex protein Nup107 OS=Homo sapiens OX=9606 GN=NUP107 PE=1 SV=1 | 14 |
| P57764 | Gasdermin-D OS=Homo sapiens OX=9606 GN=GSDMD PE=1 SV=1 | 1 |
| P58546 | Myotrophin OS=Homo sapiens OX=9606 GN=MTPN PE=1 SV=2 | 8 |
| P59045 | NACHT, LRR and PYD domains-containing protein 11 OS=Homo sapiens OX=9606 GN=NLRP11 PE=2 SV=2 | 45 |
| P59190 | Ras-related protein Rab-15 OS=Homo sapiens OX=9606 GN=RAB15 PE=1 SV=1 | 1 |
| P59780 | AP-3 complex subunit sigma-2 OS=Homo sapiens OX=9606 GN=AP3S2 PE=2 SV=1 | 5 |
| P59998 | Actin-related protein 2/3 complex subunit 4 OS=Homo sapiens OX=9606 GN=ARPC4 PE=1 SV=3 | 7 |
| P60033 | CD81 antigen OS=Homo sapiens OX=9606 GN=CD81 PE=1 SV=1 | 2 |
| P60201 | Myelin proteolipid protein OS=Homo sapiens OX=9606 GN=PLP1 PE=1 SV=2 | 3 |
| P60228 | Eukaryotic translation initiation factor 3 subunit E OS=Homo sapiens OX=9606 GN=EIF3E PE=1 SV=1 | 8 |
| P60468 | Protein transport protein Sec61 subunit beta OS=Homo sapiens OX=9606 GN=SEC61B PE=1 SV=2 | 2 |
| P60510 | Serine/threonine-protein phosphatase 4 catalytic subunit OS=Homo sapiens OX=9606 GN=PPP4C PE=1 SV=1 | 3 |
| P60520 | Gamma-aminobutyric acid receptor-associated protein-like 2 OS=Homo sapiens OX=9606 GN=GABARAPL2 PE=1 SV=1 | 7 |
| P60604 | Ubiquitin-conjugating enzyme E2 G2 OS=Homo sapiens OX=9606 GN=UBE2G2 PE=1 SV=1 | 3 |
| P60866 | 40S ribosomal protein S20 OS=Homo sapiens OX=9606 GN=RPS20 PE=1 SV=1 | 6 |
| P60891 | Ribose-phosphate pyrophosphokinase 1 OS=Homo sapiens OX=9606 GN=PRPS1 PE=1 SV=2 | 14 |
| P60903 | Protein S100-A10 OS=Homo sapiens OX=9606 GN=S100A10 PE=1 SV=2 | 1 |
| P60953 | Cell division control protein 42 homolog OS=Homo sapiens OX=9606 GN=CDC42 PE=1 SV=2 | 9 |
| P60981 | Destrin OS=Homo sapiens OX=9606 GN=DSTN PE=1 SV=3 | 12 |
| P60983 | Glia maturation factor beta OS=Homo sapiens OX=9606 GN=GMFB PE=1 SV=2 | 4 |
| P61006 | Ras-related protein Rab-8A OS=Homo sapiens OX=9606 GN=RAB8A PE=1 SV=1 | 10 |
| P61009 | Signal peptidase complex subunit 3 OS=Homo sapiens OX=9606 GN=SPCS3 PE=1 SV=1 | 3 |
| P61011 | Signal recognition particle subunit SRP54 OS=Homo sapiens OX=9606 GN=SRP54 PE=1 SV=1 | 12 |
| P61018 | Ras-related protein Rab-4B OS=Homo sapiens OX=9606 GN=RAB4B PE=1 SV=1 | 10 |
| P61020 | Ras-related protein Rab-5B OS=Homo sapiens OX=9606 GN=RAB5B PE=1 SV=1 | 11 |
| P61024 | Cyclin-dependent kinases regulatory subunit 1 OS=Homo sapiens OX=9606 GN=CKS1B PE=1 SV=1 | 2 |
| P61026 | Ras-related protein Rab-10 OS=Homo sapiens OX=9606 GN=RAB10 PE=1 SV=1 | 7 |
| P61077 | Ubiquitin-conjugating enzyme E2 D3 OS=Homo sapiens OX=9606 GN=UBE2D3 PE=1 SV=1 | 5 |
| P61081 | NEDD8-conjugating enzyme Ubc12 OS=Homo sapiens OX=9606 GN=UBE2M PE=1 SV=1 | 8 |
| P61086 | Ubiquitin-conjugating enzyme E2 K OS=Homo sapiens OX=9606 GN=UBE2K PE=1 SV=3 | 6 |
| P61088 | Ubiquitin-conjugating enzyme E2 N OS=Homo sapiens OX=9606 GN=UBE2N PE=1 SV=1 | 9 |
| P61158 | Actin-related protein 3 OS=Homo sapiens OX=9606 GN=ACTR3 PE=1 SV=3 | 20 |
| P61160 | Actin-related protein 2 OS=Homo sapiens OX=9606 GN=ACTR2 PE=1 SV=1 | 15 |
| P61163 | Alpha-centractin OS=Homo sapiens OX=9606 GN=ACTR1A PE=1 SV=1 | 5 |
| P61165 | Transmembrane protein 258 OS=Homo sapiens OX=9606 GN=TMEM258 PE=1 SV=1 | 1 |
| P61201 | COP9 signalosome complex subunit 2 OS=Homo sapiens OX=9606 GN=COPS2 PE=1 SV=1 | 24 |
| P61204;P84077 | ADP-ribosylation factor 3 OS=Homo sapiens OX=9606 GN=ARF3 PE=1 SV=2 | 3 |
| P61221 | ATP-binding cassette sub-family E member 1 OS=Homo sapiens OX=9606 GN=ABCE1 PE=1 SV=1 | 12 |
| P61224 | Ras-related protein Rap-1b OS=Homo sapiens OX=9606 GN=RAP1B PE=1 SV=1 | 9 |
| P61225 | Ras-related protein Rap-2b OS=Homo sapiens OX=9606 GN=RAP2B PE=1 SV=1 | 4 |
| P61247 | 40S ribosomal protein S3a OS=Homo sapiens OX=9606 GN=RPS3A PE=1 SV=2 | 14 |
| P61254 | 60S ribosomal protein L26 OS=Homo sapiens OX=9606 GN=RPL26 PE=1 SV=1 | 6 |
| P61289 | Proteasome activator complex subunit 3 OS=Homo sapiens OX=9606 GN=PSME3 PE=1 SV=1 | 3 |
| P61313 | 60S ribosomal protein L15 OS=Homo sapiens OX=9606 GN=RPL15 PE=1 SV=2 | 7 |
| P61353 | 60S ribosomal protein L27 OS=Homo sapiens OX=9606 GN=RPL27 PE=1 SV=2 | 4 |
| P61421 | V-type proton ATPase subunit d 1 OS=Homo sapiens OX=9606 GN=ATP6V0D1 PE=1 SV=1 | 9 |
| P61457 | Pterin-4-alpha-carbinolamine dehydratase OS=Homo sapiens OX=9606 GN=PCBD1 PE=1 SV=2 | 1 |
| P61586 | Transforming protein RhoA OS=Homo sapiens OX=9606 GN=RHOA PE=1 SV=1 | 9 |
| P61758 | Prefoldin subunit 3 OS=Homo sapiens OX=9606 GN=VBP1 PE=1 SV=4 | 6 |
| P61769 | Beta-2-microglobulin OS=Homo sapiens OX=9606 GN=B2M PE=1 SV=1 | 1 |
| P61916 | NPC intracellular cholesterol transporter 2 OS=Homo sapiens OX=9606 GN=NPC2 PE=1 SV=1 | 5 |
| P61923 | Coatomer subunit zeta-1 OS=Homo sapiens OX=9606 GN=COPZ1 PE=1 SV=1 | 3 |
| P61956 | Small ubiquitin-related modifier 2 OS=Homo sapiens OX=9606 GN=SUMO2 PE=1 SV=3 | 2 |
| P61960 | Ubiquitin-fold modifier 1 OS=Homo sapiens OX=9606 GN=UFM1 PE=1 SV=1 | 5 |
| P61966 | AP-1 complex subunit sigma-1A OS=Homo sapiens OX=9606 GN=AP1S1 PE=1 SV=1 | 4 |
| P61968 | LIM domain transcription factor LMO4 OS=Homo sapiens OX=9606 GN=LMO4 PE=1 SV=1 | 1 |
| P61970 | Nuclear transport factor 2 OS=Homo sapiens OX=9606 GN=NUTF2 PE=1 SV=1 | 4 |
| P61981 | 14-3-3 protein gamma OS=Homo sapiens OX=9606 GN=YWHAG PE=1 SV=2 | 10 |
| P62072 | Mitochondrial import inner membrane translocase subunit Tim10 OS=Homo sapiens OX=9606 GN=TIMM10 PE=1 SV=1 | 3 |
| P62081 | 40S ribosomal protein S7 OS=Homo sapiens OX=9606 GN=RPS7 PE=1 SV=1 | 8 |
| P62136 | Serine/threonine-protein phosphatase PP1-alpha catalytic subunit OS=Homo sapiens OX=9606 GN=PPP1CA PE=1 SV=1 | 11 |
| P62140 | Serine/threonine-protein phosphatase PP1-beta catalytic subunit OS=Homo sapiens OX=9606 GN=PPP1CB PE=1 SV=3 | 3 |
| P62191 | 26S proteasome regulatory subunit 4 OS=Homo sapiens OX=9606 GN=PSMC1 PE=1 SV=1 | 23 |
| P62195 | 26S proteasome regulatory subunit 8 OS=Homo sapiens OX=9606 GN=PSMC5 PE=1 SV=1 | 20 |
| P62241 | 40S ribosomal protein S8 OS=Homo sapiens OX=9606 GN=RPS8 PE=1 SV=2 | 10 |
| P62244 | 40S ribosomal protein S15a OS=Homo sapiens OX=9606 GN=RPS15A PE=1 SV=2 | 6 |
| P62253 | Ubiquitin-conjugating enzyme E2 G1 OS=Homo sapiens OX=9606 GN=UBE2G1 PE=1 SV=3 | 2 |
| P62256 | Ubiquitin-conjugating enzyme E2 H OS=Homo sapiens OX=9606 GN=UBE2H PE=1 SV=1 | 5 |
| P62263 | 40S ribosomal protein S14 OS=Homo sapiens OX=9606 GN=RPS14 PE=1 SV=3 | 3 |
| P62266 | 40S ribosomal protein S23 OS=Homo sapiens OX=9606 GN=RPS23 PE=1 SV=3 | 3 |
| P62269 | 40S ribosomal protein S18 OS=Homo sapiens OX=9606 GN=RPS18 PE=1 SV=3 | 7 |
| P62273 | 40S ribosomal protein S29 OS=Homo sapiens OX=9606 GN=RPS29 PE=1 SV=2 | 3 |
| P62277 | 40S ribosomal protein S13 OS=Homo sapiens OX=9606 GN=RPS13 PE=1 SV=2 | 6 |
| P62280 | 40S ribosomal protein S11 OS=Homo sapiens OX=9606 GN=RPS11 PE=1 SV=3 | 7 |
| P62304 | Small nuclear ribonucleoprotein E OS=Homo sapiens OX=9606 GN=SNRPE PE=1 SV=1 | 2 |
| P62306 | Small nuclear ribonucleoprotein F OS=Homo sapiens OX=9606 GN=SNRPF PE=1 SV=1 | 1 |
| P62310 | U6 snRNA-associated Sm-like protein LSm3 OS=Homo sapiens OX=9606 GN=LSM3 PE=1 SV=2 | 3 |
| P62312 | U6 snRNA-associated Sm-like protein LSm6 OS=Homo sapiens OX=9606 GN=LSM6 PE=1 SV=1 | 7 |
| P62316 | Small nuclear ribonucleoprotein Sm D2 OS=Homo sapiens OX=9606 GN=SNRPD2 PE=1 SV=1 | 2 |
| P62328 | Thymosin beta-4 OS=Homo sapiens OX=9606 GN=TMSB4X PE=1 SV=2 | 2 |
| P62330 | ADP-ribosylation factor 6 OS=Homo sapiens OX=9606 GN=ARF6 PE=1 SV=2 | 5 |
| P62333 | 26S proteasome regulatory subunit 10B OS=Homo sapiens OX=9606 GN=PSMC6 PE=1 SV=1 | 23 |
| P62424 | 60S ribosomal protein L7a OS=Homo sapiens OX=9606 GN=RPL7A PE=1 SV=2 | 11 |
| P62495 | Eukaryotic peptide chain release factor subunit 1 OS=Homo sapiens OX=9606 GN=ETF1 PE=1 SV=3 | 15 |
| P62633 | CCHC-type zinc finger nucleic acid binding protein OS=Homo sapiens OX=9606 GN=CNBP PE=1 SV=1 | 3 |
| P62701 | 40S ribosomal protein S4, X isoform OS=Homo sapiens OX=9606 GN=RPS4X PE=1 SV=2 | 14 |
| P62745 | Rho-related GTP-binding protein RhoB OS=Homo sapiens OX=9606 GN=RHOB PE=1 SV=1 | 1 |
| P62750 | 60S ribosomal protein L23a OS=Homo sapiens OX=9606 GN=RPL23A PE=1 SV=1 | 9 |
| P62826 | GTP-binding nuclear protein Ran OS=Homo sapiens OX=9606 GN=RAN PE=1 SV=3 | 15 |
| P62829 | 60S ribosomal protein L23 OS=Homo sapiens OX=9606 GN=RPL23 PE=1 SV=1 | 5 |
| P62841 | 40S ribosomal protein S15 OS=Homo sapiens OX=9606 GN=RPS15 PE=1 SV=2 | 1 |
| P62847 | 40S ribosomal protein S24 OS=Homo sapiens OX=9606 GN=RPS24 PE=1 SV=1 | 3 |
| P62851 | 40S ribosomal protein S25 OS=Homo sapiens OX=9606 GN=RPS25 PE=1 SV=1 | 3 |
| P62861 | FAU ubiquitin-like and ribosomal protein S30 OS=Homo sapiens OX=9606 GN=FAU PE=1 SV=2 | 2 |
| P62873 | Guanine nucleotide-binding protein G(I)/G(S)/G(T) subunit beta-1 OS=Homo sapiens OX=9606 GN=GNB1 PE=1 SV=3 | 12 |
| P62877 | E3 ubiquitin-protein ligase RBX1 OS=Homo sapiens OX=9606 GN=RBX1 PE=1 SV=1 | 1 |
| P62879 | Guanine nucleotide-binding protein G(I)/G(S)/G(T) subunit beta-2 OS=Homo sapiens OX=9606 GN=GNB2 PE=1 SV=3 | 3 |
| P62888 | 60S ribosomal protein L30 OS=Homo sapiens OX=9606 GN=RPL30 PE=1 SV=2 | 4 |
| P62891;Q59GN2 | 60S ribosomal protein L39 OS=Homo sapiens OX=9606 GN=RPL39 PE=1 SV=2 | 1 |
| P62906 | 60S ribosomal protein L10a OS=Homo sapiens OX=9606 GN=RPL10A PE=1 SV=2 | 6 |
| P62910 | 60S ribosomal protein L32 OS=Homo sapiens OX=9606 GN=RPL32 PE=1 SV=2 | 5 |
| P62913 | 60S ribosomal protein L11 OS=Homo sapiens OX=9606 GN=RPL11 PE=1 SV=2 | 4 |
| P62917 | 60S ribosomal protein L8 OS=Homo sapiens OX=9606 GN=RPL8 PE=1 SV=2 | 6 |
| P62942 | Peptidyl-prolyl cis-trans isomerase FKBP1A OS=Homo sapiens OX=9606 GN=FKBP1A PE=1 SV=2 | 5 |
| P62993 | Growth factor receptor-bound protein 2 OS=Homo sapiens OX=9606 GN=GRB2 PE=1 SV=1 | 7 |
| P63000 | Ras-related C3 botulinum toxin substrate 1 OS=Homo sapiens OX=9606 GN=RAC1 PE=1 SV=1 | 6 |
| P63010 | AP-2 complex subunit beta OS=Homo sapiens OX=9606 GN=AP2B1 PE=1 SV=1 | 12 |
| P63096 | Guanine nucleotide-binding protein G(i) subunit alpha-1 OS=Homo sapiens OX=9606 GN=GNAI1 PE=1 SV=2 | 3 |
| P63104 | 14-3-3 protein zeta/delta OS=Homo sapiens OX=9606 GN=YWHAZ PE=1 SV=1 | 18 |
| P63151 | Serine/threonine-protein phosphatase 2A 55 kDa regulatory subunit B alpha isoform OS=Homo sapiens OX=9606 GN=PPP2R2A PE=1 SV=1 | 10 |
| P63167 | Dynein light chain 1, cytoplasmic OS=Homo sapiens OX=9606 GN=DYNLL1 PE=1 SV=1 | 8 |
| P63172 | Dynein light chain Tctex-type 1 OS=Homo sapiens OX=9606 GN=DYNLT1 PE=1 SV=1 | 3 |
| P63173 | 60S ribosomal protein L38 OS=Homo sapiens OX=9606 GN=RPL38 PE=1 SV=2 | 2 |
| P63208 | S-phase kinase-associated protein 1 OS=Homo sapiens OX=9606 GN=SKP1 PE=1 SV=2 | 10 |
| P63220 | 40S ribosomal protein S21 OS=Homo sapiens OX=9606 GN=RPS21 PE=1 SV=1 | 5 |
| P63241 | Eukaryotic translation initiation factor 5A-1 OS=Homo sapiens OX=9606 GN=EIF5A PE=1 SV=2 | 8 |
| P63244 | Receptor of activated protein C kinase 1 OS=Homo sapiens OX=9606 GN=RACK1 PE=1 SV=3 | 15 |
| P63279 | SUMO-conjugating enzyme UBC9 OS=Homo sapiens OX=9606 GN=UBE2I PE=1 SV=1 | 7 |
| P67775 | Serine/threonine-protein phosphatase 2A catalytic subunit alpha isoform OS=Homo sapiens OX=9606 GN=PPP2CA PE=1 SV=1 | 9 |
| P67809 | Y-box-binding protein 1 OS=Homo sapiens OX=9606 GN=YBX1 PE=1 SV=3 | 4 |
| P67812 | Signal peptidase complex catalytic subunit SEC11A OS=Homo sapiens OX=9606 GN=SEC11A PE=1 SV=1 | 2 |
| P67870 | Casein kinase II subunit beta OS=Homo sapiens OX=9606 GN=CSNK2B PE=1 SV=1 | 3 |
| P67936 | Tropomyosin alpha-4 chain OS=Homo sapiens OX=9606 GN=TPM4 PE=1 SV=3 | 7 |
| P68036 | Ubiquitin-conjugating enzyme E2 L3 OS=Homo sapiens OX=9606 GN=UBE2L3 PE=1 SV=1 | 8 |
| P68363;Q71U36 | Tubulin alpha-1B chain OS=Homo sapiens OX=9606 GN=TUBA1B PE=1 SV=1 | 1 |
| P68400 | Casein kinase II subunit alpha OS=Homo sapiens OX=9606 GN=CSNK2A1 PE=1 SV=1 | 4 |
| P68402 | Platelet-activating factor acetylhydrolase IB subunit alpha2 OS=Homo sapiens OX=9606 GN=PAFAH1B2 PE=1 SV=1 | 5 |
| P69849;Q15155;Q5JPE7 | BOS complex subunit NOMO3 OS=Homo sapiens OX=9606 GN=NOMO3 PE=1 SV=2 | 28 |
| P78318 | Immunoglobulin-binding protein 1 OS=Homo sapiens OX=9606 GN=IGBP1 PE=1 SV=1 | 6 |
| P78330 | Phosphoserine phosphatase OS=Homo sapiens OX=9606 GN=PSPH PE=1 SV=2 | 1 |
| P78333 | Glypican-5 OS=Homo sapiens OX=9606 GN=GPC5 PE=1 SV=1 | 1 |
| P78344 | Eukaryotic translation initiation factor 4 gamma 2 OS=Homo sapiens OX=9606 GN=EIF4G2 PE=1 SV=1 | 23 |
| P78347 | General transcription factor II-I OS=Homo sapiens OX=9606 GN=GTF2I PE=1 SV=2 | 4 |
| P78371 | T-complex protein 1 subunit beta OS=Homo sapiens OX=9606 GN=CCT2 PE=1 SV=4 | 41 |
| P78406 | mRNA export factor RAE1 OS=Homo sapiens OX=9606 GN=RAE1 PE=1 SV=1 | 10 |
| P78504 | Protein jagged-1 OS=Homo sapiens OX=9606 GN=JAG1 PE=1 SV=3 | 2 |
| P78527 | DNA-dependent protein kinase catalytic subunit OS=Homo sapiens OX=9606 GN=PRKDC PE=1 SV=3 | 9 |
| P78537 | Biogenesis of lysosome-related organelles complex 1 subunit 1 OS=Homo sapiens OX=9606 GN=BLOC1S1 PE=1 SV=2 | 1 |
| P78562 | Phosphate-regulating neutral endopeptidase PHEX OS=Homo sapiens OX=9606 GN=PHEX PE=1 SV=1 | 19 |
| P80188 | Neutrophil gelatinase-associated lipocalin OS=Homo sapiens OX=9606 GN=LCN2 PE=1 SV=2 | 4 |
| P80303 | Nucleobindin-2 OS=Homo sapiens OX=9606 GN=NUCB2 PE=1 SV=3 | 15 |
| P80723 | Brain acid soluble protein 1 OS=Homo sapiens OX=9606 GN=BASP1 PE=1 SV=2 | 8 |
| P80748 | Immunoglobulin lambda variable 3-21 OS=Homo sapiens OX=9606 GN=IGLV3-21 PE=1 SV=2 | 1 |
| P82650 | 28S ribosomal protein S22, mitochondrial OS=Homo sapiens OX=9606 GN=MRPS22 PE=1 SV=1 | 7 |
| P82663 | 28S ribosomal protein S25, mitochondrial OS=Homo sapiens OX=9606 GN=MRPS25 PE=1 SV=1 | 7 |
| P82664 | 28S ribosomal protein S10, mitochondrial OS=Homo sapiens OX=9606 GN=MRPS10 PE=1 SV=2 | 2 |
| P82673 | 28S ribosomal protein S35, mitochondrial OS=Homo sapiens OX=9606 GN=MRPS35 PE=1 SV=1 | 3 |
| P82909 | Alpha-ketoglutarate dehydrogenase component 4 OS=Homo sapiens OX=9606 GN=MRPS36 PE=1 SV=2 | 6 |
| P82912 | 28S ribosomal protein S11, mitochondrial OS=Homo sapiens OX=9606 GN=MRPS11 PE=1 SV=2 | 1 |
| P82930 | 28S ribosomal protein S34, mitochondrial OS=Homo sapiens OX=9606 GN=MRPS34 PE=1 SV=2 | 3 |
| P82932 | 28S ribosomal protein S6, mitochondrial OS=Homo sapiens OX=9606 GN=MRPS6 PE=1 SV=3 | 1 |
| P82979 | SAP domain-containing ribonucleoprotein OS=Homo sapiens OX=9606 GN=SARNP PE=1 SV=3 | 1 |
| P83111 | Serine beta-lactamase-like protein LACTB, mitochondrial OS=Homo sapiens OX=9606 GN=LACTB PE=1 SV=2 | 9 |
| P83436 | Conserved oligomeric Golgi complex subunit 7 OS=Homo sapiens OX=9606 GN=COG7 PE=1 SV=1 | 2 |
| P83731 | 60S ribosomal protein L24 OS=Homo sapiens OX=9606 GN=RPL24 PE=1 SV=1 | 5 |
| P83876 | Thioredoxin-like protein 4A OS=Homo sapiens OX=9606 GN=TXNL4A PE=1 SV=1 | 1 |
| P83881 | 60S ribosomal protein L36a OS=Homo sapiens OX=9606 GN=RPL36A PE=1 SV=2 | 2 |
| P83916 | Chromobox protein homolog 1 OS=Homo sapiens OX=9606 GN=CBX1 PE=1 SV=1 | 5 |
| P84085 | ADP-ribosylation factor 5 OS=Homo sapiens OX=9606 GN=ARF5 PE=1 SV=2 | 9 |
| P84090 | Enhancer of rudimentary homolog OS=Homo sapiens OX=9606 GN=ERH PE=1 SV=1 | 2 |
| P84095 | Rho-related GTP-binding protein RhoG OS=Homo sapiens OX=9606 GN=RHOG PE=1 SV=1 | 6 |
| P84098 | 60S ribosomal protein L19 OS=Homo sapiens OX=9606 GN=RPL19 PE=1 SV=1 | 2 |
| P84103 | Serine/arginine-rich splicing factor 3 OS=Homo sapiens OX=9606 GN=SRSF3 PE=1 SV=1 | 1 |
| P86790;P86791 | Vacuolar fusion protein CCZ1 homolog B OS=Homo sapiens OX=9606 GN=CCZ1B PE=1 SV=1 | 14 |
| P98155 | Very low-density lipoprotein receptor OS=Homo sapiens OX=9606 GN=VLDLR PE=1 SV=1 | 5 |
| P98160 | Basement membrane-specific heparan sulfate proteoglycan core protein OS=Homo sapiens OX=9606 GN=HSPG2 PE=1 SV=4 | 41 |
| P98179 | RNA-binding protein 3 OS=Homo sapiens OX=9606 GN=RBM3 PE=1 SV=1 | 2 |
| P98194 | Calcium-transporting ATPase type 2C member 1 OS=Homo sapiens OX=9606 GN=ATP2C1 PE=1 SV=3 | 10 |
| P99999 | Cytochrome c OS=Homo sapiens OX=9606 GN=CYCS PE=1 SV=2 | 7 |
| Q00059 | Transcription factor A, mitochondrial OS=Homo sapiens OX=9606 GN=TFAM PE=1 SV=1 | 9 |
| Q00169 | Phosphatidylinositol transfer protein alpha isoform OS=Homo sapiens OX=9606 GN=PITPNA PE=1 SV=2 | 20 |
| Q00266 | S-adenosylmethionine synthase isoform type-1 OS=Homo sapiens OX=9606 GN=MAT1A PE=1 SV=2 | 10 |
| Q00341 | Vigilin OS=Homo sapiens OX=9606 GN=HDLBP PE=1 SV=2 | 2 |
| Q00535 | Cyclin-dependent kinase 5 OS=Homo sapiens OX=9606 GN=CDK5 PE=1 SV=3 | 19 |
| Q00577 | Transcriptional activator protein Pur-alpha OS=Homo sapiens OX=9606 GN=PURA PE=1 SV=2 | 2 |
| Q00610 | Clathrin heavy chain 1 OS=Homo sapiens OX=9606 GN=CLTC PE=1 SV=5 | 79 |
| Q00688 | Peptidyl-prolyl cis-trans isomerase FKBP3 OS=Homo sapiens OX=9606 GN=FKBP3 PE=1 SV=1 | 9 |
| Q00765 | Receptor expression-enhancing protein 5 OS=Homo sapiens OX=9606 GN=REEP5 PE=1 SV=3 | 3 |
| Q01082 | Spectrin beta chain, non-erythrocytic 1 OS=Homo sapiens OX=9606 GN=SPTBN1 PE=1 SV=2 | 28 |
| Q01085 | Nucleolysin TIAR OS=Homo sapiens OX=9606 GN=TIAL1 PE=1 SV=1 | 7 |
| Q01130 | Serine/arginine-rich splicing factor 2 OS=Homo sapiens OX=9606 GN=SRSF2 PE=1 SV=4 | 2 |
| Q01433 | AMP deaminase 2 OS=Homo sapiens OX=9606 GN=AMPD2 PE=1 SV=2 | 20 |
| Q01459 | Di-N-acetylchitobiase OS=Homo sapiens OX=9606 GN=CTBS PE=1 SV=1 | 4 |
| Q01546 | Keratin, type II cytoskeletal 2 oral OS=Homo sapiens OX=9606 GN=KRT76 PE=1 SV=2 | 14 |
| Q01804 | OTU domain-containing protein 4 OS=Homo sapiens OX=9606 GN=OTUD4 PE=1 SV=4 | 6 |
| Q01813 | ATP-dependent 6-phosphofructokinase, platelet type OS=Homo sapiens OX=9606 GN=PFKP PE=1 SV=2 | 1 |
| Q01844 | RNA-binding protein EWS OS=Homo sapiens OX=9606 GN=EWSR1 PE=1 SV=1 | 3 |
| Q01955 | Collagen alpha-3(IV) chain OS=Homo sapiens OX=9606 GN=COL4A3 PE=1 SV=3 | 1 |
| Q01968 | Inositol polyphosphate 5-phosphatase OCRL OS=Homo sapiens OX=9606 GN=OCRL PE=1 SV=3 | 7 |
| Q01970 | 1-phosphatidylinositol 4,5-bisphosphate phosphodiesterase beta-3 OS=Homo sapiens OX=9606 GN=PLCB3 PE=1 SV=2 | 2 |
| Q01974 | Tyrosine-protein kinase transmembrane receptor ROR2 OS=Homo sapiens OX=9606 GN=ROR2 PE=1 SV=2 | 1 |
| Q02218 | 2-oxoglutarate dehydrogenase complex component E1 OS=Homo sapiens OX=9606 GN=OGDH PE=1 SV=3 | 54 |
| Q02224 | Centromere-associated protein E OS=Homo sapiens OX=9606 GN=CENPE PE=1 SV=2 | 38 |
| Q02241 | Kinesin-like protein KIF23 OS=Homo sapiens OX=9606 GN=KIF23 PE=1 SV=3 | 9 |
| Q02252 | Methylmalonate-semialdehyde dehydrogenase [acylating], mitochondrial OS=Homo sapiens OX=9606 GN=ALDH6A1 PE=1 SV=2 | 26 |
| Q02338 | D-beta-hydroxybutyrate dehydrogenase, mitochondrial OS=Homo sapiens OX=9606 GN=BDH1 PE=1 SV=3 | 5 |
| Q02410 | Amyloid-beta A4 precursor protein-binding family A member 1 OS=Homo sapiens OX=9606 GN=APBA1 PE=1 SV=3 | 7 |
| Q02543 | 60S ribosomal protein L18a OS=Homo sapiens OX=9606 GN=RPL18A PE=1 SV=2 | 5 |
| Q02750 | Dual specificity mitogen-activated protein kinase kinase 1 OS=Homo sapiens OX=9606 GN=MAP2K1 PE=1 SV=2 | 5 |
| Q02790 | Peptidyl-prolyl cis-trans isomerase FKBP4 OS=Homo sapiens OX=9606 GN=FKBP4 PE=1 SV=3 | 35 |
| Q02818 | Nucleobindin-1 OS=Homo sapiens OX=9606 GN=NUCB1 PE=1 SV=4 | 13 |
| Q02878 | 60S ribosomal protein L6 OS=Homo sapiens OX=9606 GN=RPL6 PE=1 SV=3 | 10 |
| Q02978 | Mitochondrial 2-oxoglutarate/malate carrier protein OS=Homo sapiens OX=9606 GN=SLC25A11 PE=1 SV=3 | 3 |
| Q03252 | Lamin-B2 OS=Homo sapiens OX=9606 GN=LMNB2 PE=1 SV=4 | 7 |
| Q03393 | 6-pyruvoyl tetrahydrobiopterin synthase OS=Homo sapiens OX=9606 GN=PTS PE=1 SV=1 | 3 |
| Q03426 | Mevalonate kinase OS=Homo sapiens OX=9606 GN=MVK PE=1 SV=1 | 12 |
| Q03518 | Antigen peptide transporter 1 OS=Homo sapiens OX=9606 GN=TAP1 PE=1 SV=3 | 1 |
| Q04323 | UBX domain-containing protein 1 OS=Homo sapiens OX=9606 GN=UBXN1 PE=1 SV=2 | 3 |
| Q04446 | 1,4-alpha-glucan-branching enzyme OS=Homo sapiens OX=9606 GN=GBE1 PE=1 SV=3 | 22 |
| Q04637 | Eukaryotic translation initiation factor 4 gamma 1 OS=Homo sapiens OX=9606 GN=EIF4G1 PE=1 SV=4 | 25 |
| Q04721 | Neurogenic locus notch homolog protein 2 OS=Homo sapiens OX=9606 GN=NOTCH2 PE=1 SV=3 | 1 |
| Q04725 | Transducin-like enhancer protein 2 OS=Homo sapiens OX=9606 GN=TLE2 PE=1 SV=2 | 5 |
| Q04760 | Lactoylglutathione lyase OS=Homo sapiens OX=9606 GN=GLO1 PE=1 SV=4 | 12 |
| Q04837 | Single-stranded DNA-binding protein, mitochondrial OS=Homo sapiens OX=9606 GN=SSBP1 PE=1 SV=1 | 10 |
| Q04900 | Sialomucin core protein 24 OS=Homo sapiens OX=9606 GN=CD164 PE=1 SV=2 | 1 |
| Q04917 | 14-3-3 protein eta OS=Homo sapiens OX=9606 GN=YWHAH PE=1 SV=4 | 14 |
| Q05048 | Cleavage stimulation factor subunit 1 OS=Homo sapiens OX=9606 GN=CSTF1 PE=1 SV=1 | 3 |
| Q05397 | Focal adhesion kinase 1 OS=Homo sapiens OX=9606 GN=PTK2 PE=1 SV=2 | 18 |
| Q05707 | Collagen alpha-1(XIV) chain OS=Homo sapiens OX=9606 GN=COL14A1 PE=1 SV=3 | 3 |
| Q05BV3 | Echinoderm microtubule-associated protein-like 5 OS=Homo sapiens OX=9606 GN=EML5 PE=2 SV=3 | 1 |
| Q06033 | Inter-alpha-trypsin inhibitor heavy chain H3 OS=Homo sapiens OX=9606 GN=ITIH3 PE=1 SV=2 | 10 |
| Q06203 | Amidophosphoribosyltransferase OS=Homo sapiens OX=9606 GN=PPAT PE=1 SV=1 | 20 |
| Q06210 | Glutamine--fructose-6-phosphate aminotransferase [isomerizing] 1 OS=Homo sapiens OX=9606 GN=GFPT1 PE=1 SV=3 | 12 |
| Q06265 | Exosome complex component RRP45 OS=Homo sapiens OX=9606 GN=EXOSC9 PE=1 SV=3 | 1 |
| Q06278 | Aldehyde oxidase OS=Homo sapiens OX=9606 GN=AOX1 PE=1 SV=2 | 8 |
| Q06323 | Proteasome activator complex subunit 1 OS=Homo sapiens OX=9606 GN=PSME1 PE=1 SV=1 | 5 |
| Q06418 | Tyrosine-protein kinase receptor TYRO3 OS=Homo sapiens OX=9606 GN=TYRO3 PE=1 SV=1 | 5 |
| Q06787 | Fragile X messenger ribonucleoprotein 1 OS=Homo sapiens OX=9606 GN=FMR1 PE=1 SV=1 | 3 |
| Q07020 | 60S ribosomal protein L18 OS=Homo sapiens OX=9606 GN=RPL18 PE=1 SV=2 | 7 |
| Q07021 | Complement component 1 Q subcomponent-binding protein, mitochondrial OS=Homo sapiens OX=9606 GN=C1QBP PE=1 SV=1 | 6 |
| Q07065 | Cytoskeleton-associated protein 4 OS=Homo sapiens OX=9606 GN=CKAP4 PE=1 SV=2 | 3 |
| Q07157 | Tight junction protein ZO-1 OS=Homo sapiens OX=9606 GN=TJP1 PE=1 SV=3 | 6 |
| Q07666 | KH domain-containing, RNA-binding, signal transduction-associated protein 1 OS=Homo sapiens OX=9606 GN=KHDRBS1 PE=1 SV=1 | 2 |
| Q07812 | Apoptosis regulator BAX OS=Homo sapiens OX=9606 GN=BAX PE=1 SV=1 | 3 |
| Q07866 | Kinesin light chain 1 OS=Homo sapiens OX=9606 GN=KLC1 PE=1 SV=2 | 19 |
| Q07955 | Serine/arginine-rich splicing factor 1 OS=Homo sapiens OX=9606 GN=SRSF1 PE=1 SV=2 | 5 |
| Q07960 | Rho GTPase-activating protein 1 OS=Homo sapiens OX=9606 GN=ARHGAP1 PE=1 SV=1 | 12 |
| Q08043 | Alpha-actinin-3 OS=Homo sapiens OX=9606 GN=ACTN3 PE=1 SV=2 | 9 |
| Q08211 | ATP-dependent RNA helicase A OS=Homo sapiens OX=9606 GN=DHX9 PE=1 SV=4 | 2 |
| Q08345 | Epithelial discoidin domain-containing receptor 1 OS=Homo sapiens OX=9606 GN=DDR1 PE=1 SV=1 | 1 |
| Q08378 | Golgin subfamily A member 3 OS=Homo sapiens OX=9606 GN=GOLGA3 PE=1 SV=2 | 22 |
| Q08379 | Golgin subfamily A member 2 OS=Homo sapiens OX=9606 GN=GOLGA2 PE=1 SV=3 | 4 |
| Q08431 | Lactadherin OS=Homo sapiens OX=9606 GN=MFGE8 PE=1 SV=3 | 25 |
| Q08499 | cAMP-specific 3',5'-cyclic phosphodiesterase 4D OS=Homo sapiens OX=9606 GN=PDE4D PE=1 SV=2 | 1 |
| Q08629 | Testican-1 OS=Homo sapiens OX=9606 GN=SPOCK1 PE=1 SV=1 | 6 |
| Q08722 | Leukocyte surface antigen CD47 OS=Homo sapiens OX=9606 GN=CD47 PE=1 SV=1 | 2 |
| Q08752 | Peptidyl-prolyl cis-trans isomerase D OS=Homo sapiens OX=9606 GN=PPID PE=1 SV=3 | 13 |
| Q08AM6 | Protein VAC14 homolog OS=Homo sapiens OX=9606 GN=VAC14 PE=1 SV=1 | 2 |
| Q09028 | Histone-binding protein RBBP4 OS=Homo sapiens OX=9606 GN=RBBP4 PE=1 SV=3 | 13 |
| Q0VDF9 | Heat shock 70 kDa protein 14 OS=Homo sapiens OX=9606 GN=HSPA14 PE=1 SV=1 | 1 |
| Q0VDG4 | Secernin-3 OS=Homo sapiens OX=9606 GN=SCRN3 PE=1 SV=1 | 3 |
| Q0VGL1 | Ragulator complex protein LAMTOR4 OS=Homo sapiens OX=9606 GN=LAMTOR4 PE=1 SV=1 | 1 |
| Q10471 | Polypeptide N-acetylgalactosaminyltransferase 2 OS=Homo sapiens OX=9606 GN=GALNT2 PE=1 SV=1 | 6 |
| Q10567 | AP-1 complex subunit beta-1 OS=Homo sapiens OX=9606 GN=AP1B1 PE=1 SV=3 | 31 |
| Q10570 | Cleavage and polyadenylation specificity factor subunit 1 OS=Homo sapiens OX=9606 GN=CPSF1 PE=1 SV=2 | 1 |
| Q10589 | Bone marrow stromal antigen 2 OS=Homo sapiens OX=9606 GN=BST2 PE=1 SV=1 | 2 |
| Q10713 | Mitochondrial-processing peptidase subunit alpha OS=Homo sapiens OX=9606 GN=PMPCA PE=1 SV=2 | 13 |
| Q12756 | Kinesin-like protein KIF1A OS=Homo sapiens OX=9606 GN=KIF1A PE=1 SV=2 | 14 |
| Q12765 | Secernin-1 OS=Homo sapiens OX=9606 GN=SCRN1 PE=1 SV=2 | 6 |
| Q12768 | WASH complex subunit 5 OS=Homo sapiens OX=9606 GN=WASHC5 PE=1 SV=1 | 9 |
| Q12792 | Twinfilin-1 OS=Homo sapiens OX=9606 GN=TWF1 PE=1 SV=3 | 5 |
| Q12797 | Aspartyl/asparaginyl beta-hydroxylase OS=Homo sapiens OX=9606 GN=ASPH PE=1 SV=3 | 2 |
| Q12802 | A-kinase anchor protein 13 OS=Homo sapiens OX=9606 GN=AKAP13 PE=1 SV=2 | 2 |
| Q12846 | Syntaxin-4 OS=Homo sapiens OX=9606 GN=STX4 PE=1 SV=2 | 9 |
| Q12849 | G-rich sequence factor 1 OS=Homo sapiens OX=9606 GN=GRSF1 PE=1 SV=3 | 14 |
| Q12864 | Cadherin-17 OS=Homo sapiens OX=9606 GN=CDH17 PE=1 SV=3 | 5 |
| Q12879 | Glutamate receptor ionotropic, NMDA 2A OS=Homo sapiens OX=9606 GN=GRIN2A PE=1 SV=1 | 2 |
| Q12899 | Tripartite motif-containing protein 26 OS=Homo sapiens OX=9606 GN=TRIM26 PE=1 SV=1 | 2 |
| Q12904 | Aminoacyl tRNA synthase complex-interacting multifunctional protein 1 OS=Homo sapiens OX=9606 GN=AIMP1 PE=1 SV=2 | 9 |
| Q12905 | Interleukin enhancer-binding factor 2 OS=Homo sapiens OX=9606 GN=ILF2 PE=1 SV=2 | 4 |
| Q12906 | Interleukin enhancer-binding factor 3 OS=Homo sapiens OX=9606 GN=ILF3 PE=1 SV=3 | 3 |
| Q12907 | Vesicular integral-membrane protein VIP36 OS=Homo sapiens OX=9606 GN=LMAN2 PE=1 SV=1 | 19 |
| Q12926 | ELAV-like protein 2 OS=Homo sapiens OX=9606 GN=ELAVL2 PE=1 SV=2 | 3 |
| Q12933 | TNF receptor-associated factor 2 OS=Homo sapiens OX=9606 GN=TRAF2 PE=1 SV=2 | 12 |
| Q12965 | Unconventional myosin-Ie OS=Homo sapiens OX=9606 GN=MYO1E PE=1 SV=2 | 2 |
| Q12981 | Vesicle transport protein SEC20 OS=Homo sapiens OX=9606 GN=BNIP1 PE=1 SV=3 | 8 |
| Q12986 | Transcriptional repressor NF-X1 OS=Homo sapiens OX=9606 GN=NFX1 PE=1 SV=2 | 2 |
| Q13009 | Rho guanine nucleotide exchange factor TIAM1 OS=Homo sapiens OX=9606 GN=TIAM1 PE=1 SV=2 | 8 |
| Q13011 | Delta(3,5)-Delta(2,4)-dienoyl-CoA isomerase, mitochondrial OS=Homo sapiens OX=9606 GN=ECH1 PE=1 SV=2 | 17 |
| Q13015 | Protein AF1q OS=Homo sapiens OX=9606 GN=MLLT11 PE=1 SV=1 | 3 |
| Q13045 | Protein flightless-1 homolog OS=Homo sapiens OX=9606 GN=FLII PE=1 SV=2 | 1 |
| Q13057 | Bifunctional coenzyme A synthase OS=Homo sapiens OX=9606 GN=COASY PE=1 SV=4 | 7 |
| Q13085 | Acetyl-CoA carboxylase 1 OS=Homo sapiens OX=9606 GN=ACACA PE=1 SV=2 | 6 |
| Q13098 | COP9 signalosome complex subunit 1 OS=Homo sapiens OX=9606 GN=GPS1 PE=1 SV=4 | 16 |
| Q13107 | Ubiquitin carboxyl-terminal hydrolase 4 OS=Homo sapiens OX=9606 GN=USP4 PE=1 SV=3 | 1 |
| Q13126 | S-methyl-5'-thioadenosine phosphorylase OS=Homo sapiens OX=9606 GN=MTAP PE=1 SV=2 | 10 |
| Q13129 | Zinc finger protein Rlf OS=Homo sapiens OX=9606 GN=RLF PE=1 SV=2 | 1 |
| Q13131 | 5'-AMP-activated protein kinase catalytic subunit alpha-1 OS=Homo sapiens OX=9606 GN=PRKAA1 PE=1 SV=4 | 7 |
| Q13144 | Translation initiation factor eIF-2B subunit epsilon OS=Homo sapiens OX=9606 GN=EIF2B5 PE=1 SV=3 | 1 |
| Q13148 | TAR DNA-binding protein 43 OS=Homo sapiens OX=9606 GN=TARDBP PE=1 SV=1 | 5 |
| Q13153 | Serine/threonine-protein kinase PAK 1 OS=Homo sapiens OX=9606 GN=PAK1 PE=1 SV=2 | 9 |
| Q13155 | Aminoacyl tRNA synthase complex-interacting multifunctional protein 2 OS=Homo sapiens OX=9606 GN=AIMP2 PE=1 SV=2 | 4 |
| Q13158 | FAS-associated death domain protein OS=Homo sapiens OX=9606 GN=FADD PE=1 SV=1 | 1 |
| Q13162 | Peroxiredoxin-4 OS=Homo sapiens OX=9606 GN=PRDX4 PE=1 SV=1 | 7 |
| Q13164 | Mitogen-activated protein kinase 7 OS=Homo sapiens OX=9606 GN=MAPK7 PE=1 SV=2 | 4 |
| Q13177 | Serine/threonine-protein kinase PAK 2 OS=Homo sapiens OX=9606 GN=PAK2 PE=1 SV=3 | 27 |
| Q13185 | Chromobox protein homolog 3 OS=Homo sapiens OX=9606 GN=CBX3 PE=1 SV=4 | 5 |
| Q13200 | 26S proteasome non-ATPase regulatory subunit 2 OS=Homo sapiens OX=9606 GN=PSMD2 PE=1 SV=3 | 38 |
| Q13217 | DnaJ homolog subfamily C member 3 OS=Homo sapiens OX=9606 GN=DNAJC3 PE=1 SV=1 | 27 |
| Q13228 | Methanethiol oxidase OS=Homo sapiens OX=9606 GN=SELENBP1 PE=1 SV=2 | 1 |
| Q13237 | cGMP-dependent protein kinase 2 OS=Homo sapiens OX=9606 GN=PRKG2 PE=1 SV=1 | 1 |
| Q13247 | Serine/arginine-rich splicing factor 6 OS=Homo sapiens OX=9606 GN=SRSF6 PE=1 SV=2 | 3 |
| Q13263 | Transcription intermediary factor 1-beta OS=Homo sapiens OX=9606 GN=TRIM28 PE=1 SV=5 | 12 |
| Q13277 | Syntaxin-3 OS=Homo sapiens OX=9606 GN=STX3 PE=1 SV=3 | 14 |
| Q13283 | Ras GTPase-activating protein-binding protein 1 OS=Homo sapiens OX=9606 GN=G3BP1 PE=1 SV=1 | 6 |
| Q13308 | Inactive tyrosine-protein kinase 7 OS=Homo sapiens OX=9606 GN=PTK7 PE=1 SV=2 | 1 |
| Q13310 | Polyadenylate-binding protein 4 OS=Homo sapiens OX=9606 GN=PABPC4 PE=1 SV=1 | 20 |
| Q13347 | Eukaryotic translation initiation factor 3 subunit I OS=Homo sapiens OX=9606 GN=EIF3I PE=1 SV=1 | 13 |
| Q13362 | Serine/threonine-protein phosphatase 2A 56 kDa regulatory subunit gamma isoform OS=Homo sapiens OX=9606 GN=PPP2R5C PE=1 SV=3 | 1 |
| Q13371 | Phosducin-like protein OS=Homo sapiens OX=9606 GN=PDCL PE=1 SV=3 | 5 |
| Q13395 | Probable methyltransferase TARBP1 OS=Homo sapiens OX=9606 GN=TARBP1 PE=1 SV=1 | 3 |
| Q13405 | 39S ribosomal protein L49, mitochondrial OS=Homo sapiens OX=9606 GN=MRPL49 PE=1 SV=1 | 3 |
| Q13409 | Cytoplasmic dynein 1 intermediate chain 2 OS=Homo sapiens OX=9606 GN=DYNC1I2 PE=1 SV=3 | 10 |
| Q13415 | Origin recognition complex subunit 1 OS=Homo sapiens OX=9606 GN=ORC1 PE=1 SV=2 | 1 |
| Q13418 | Integrin-linked protein kinase OS=Homo sapiens OX=9606 GN=ILK PE=1 SV=2 | 2 |
| Q13428 | Treacle protein OS=Homo sapiens OX=9606 GN=TCOF1 PE=1 SV=3 | 7 |
| Q13433 | Zinc transporter ZIP6 OS=Homo sapiens OX=9606 GN=SLC39A6 PE=1 SV=3 | 1 |
| Q13435 | Splicing factor 3B subunit 2 OS=Homo sapiens OX=9606 GN=SF3B2 PE=1 SV=2 | 1 |
| Q13438 | Protein OS-9 OS=Homo sapiens OX=9606 GN=OS9 PE=1 SV=1 | 18 |
| Q13442 | 28 kDa heat- and acid-stable phosphoprotein OS=Homo sapiens OX=9606 GN=PDAP1 PE=1 SV=1 | 7 |
| Q13444 | Disintegrin and metalloproteinase domain-containing protein 15 OS=Homo sapiens OX=9606 GN=ADAM15 PE=1 SV=4 | 1 |
| Q13445 | Transmembrane emp24 domain-containing protein 1 OS=Homo sapiens OX=9606 GN=TMED1 PE=1 SV=1 | 2 |
| Q13490 | Baculoviral IAP repeat-containing protein 2 OS=Homo sapiens OX=9606 GN=BIRC2 PE=1 SV=2 | 13 |
| Q13492 | Phosphatidylinositol-binding clathrin assembly protein OS=Homo sapiens OX=9606 GN=PICALM PE=1 SV=2 | 7 |
| Q13501 | Sequestosome-1 OS=Homo sapiens OX=9606 GN=SQSTM1 PE=1 SV=1 | 1 |
| Q13505 | Metaxin-1 OS=Homo sapiens OX=9606 GN=MTX1 PE=1 SV=3 | 9 |
| Q13526 | Peptidyl-prolyl cis-trans isomerase NIMA-interacting 1 OS=Homo sapiens OX=9606 GN=PIN1 PE=1 SV=1 | 9 |
| Q13541 | Eukaryotic translation initiation factor 4E-binding protein 1 OS=Homo sapiens OX=9606 GN=EIF4EBP1 PE=1 SV=3 | 1 |
| Q13547;Q92769 | Histone deacetylase 1 OS=Homo sapiens OX=9606 GN=HDAC1 PE=1 SV=1 | 1 |
| Q13555 | Calcium/calmodulin-dependent protein kinase type II subunit gamma OS=Homo sapiens OX=9606 GN=CAMK2G PE=1 SV=4 | 2 |
| Q13557 | Calcium/calmodulin-dependent protein kinase type II subunit delta OS=Homo sapiens OX=9606 GN=CAMK2D PE=1 SV=3 | 1 |
| Q13561 | Dynactin subunit 2 OS=Homo sapiens OX=9606 GN=DCTN2 PE=1 SV=4 | 16 |
| Q13564 | NEDD8-activating enzyme E1 regulatory subunit OS=Homo sapiens OX=9606 GN=NAE1 PE=1 SV=1 | 8 |
| Q13572 | Inositol-tetrakisphosphate 1-kinase OS=Homo sapiens OX=9606 GN=ITPK1 PE=1 SV=2 | 1 |
| Q13576 | Ras GTPase-activating-like protein IQGAP2 OS=Homo sapiens OX=9606 GN=IQGAP2 PE=1 SV=4 | 14 |
| Q13596 | Sorting nexin-1 OS=Homo sapiens OX=9606 GN=SNX1 PE=1 SV=3 | 14 |
| Q13610 | Periodic tryptophan protein 1 homolog OS=Homo sapiens OX=9606 GN=PWP1 PE=1 SV=1 | 3 |
| Q13613 | Myotubularin-related protein 1 OS=Homo sapiens OX=9606 GN=MTMR1 PE=1 SV=4 | 4 |
| Q13614 | Myotubularin-related protein 2 OS=Homo sapiens OX=9606 GN=MTMR2 PE=1 SV=4 | 4 |
| Q13615 | Myotubularin-related protein 3 OS=Homo sapiens OX=9606 GN=MTMR3 PE=1 SV=3 | 32 |
| Q13616 | Cullin-1 OS=Homo sapiens OX=9606 GN=CUL1 PE=1 SV=2 | 25 |
| Q13618 | Cullin-3 OS=Homo sapiens OX=9606 GN=CUL3 PE=1 SV=2 | 17 |
| Q13630 | GDP-L-fucose synthase OS=Homo sapiens OX=9606 GN=GFUS PE=1 SV=1 | 18 |
| Q13670 | Putative postmeiotic segregation increased 2-like protein 11 OS=Homo sapiens OX=9606 GN=PMS2P11 PE=5 SV=1 | 1 |
| Q13724 | Mannosyl-oligosaccharide glucosidase OS=Homo sapiens OX=9606 GN=MOGS PE=1 SV=5 | 2 |
| Q13769 | THO complex subunit 5 homolog OS=Homo sapiens OX=9606 GN=THOC5 PE=1 SV=2 | 1 |
| Q13796 | Protein Shroom2 OS=Homo sapiens OX=9606 GN=SHROOM2 PE=1 SV=1 | 4 |
| Q13797 | Integrin alpha-9 OS=Homo sapiens OX=9606 GN=ITGA9 PE=1 SV=2 | 12 |
| Q13813 | Spectrin alpha chain, non-erythrocytic 1 OS=Homo sapiens OX=9606 GN=SPTAN1 PE=1 SV=3 | 26 |
| Q13822 | Ectonucleotide pyrophosphatase/phosphodiesterase family member 2 OS=Homo sapiens OX=9606 GN=ENPP2 PE=1 SV=3 | 1 |
| Q13825 | Methylglutaconyl-CoA hydratase, mitochondrial OS=Homo sapiens OX=9606 GN=AUH PE=1 SV=1 | 2 |
| Q13838 | Spliceosome RNA helicase DDX39B OS=Homo sapiens OX=9606 GN=DDX39B PE=1 SV=1 | 8 |
| Q13885;Q9BVA1 | Tubulin beta-2A chain OS=Homo sapiens OX=9606 GN=TUBB2A PE=1 SV=1 | 1 |
| Q13907 | Isopentenyl-diphosphate Delta-isomerase 1 OS=Homo sapiens OX=9606 GN=IDI1 PE=1 SV=2 | 5 |
| Q14005 | Pro-interleukin-16 OS=Homo sapiens OX=9606 GN=IL16 PE=1 SV=4 | 2 |
| Q14008 | Cytoskeleton-associated protein 5 OS=Homo sapiens OX=9606 GN=CKAP5 PE=1 SV=3 | 64 |
| Q14011 | Cold-inducible RNA-binding protein OS=Homo sapiens OX=9606 GN=CIRBP PE=1 SV=1 | 8 |
| Q14019 | Coactosin-like protein OS=Homo sapiens OX=9606 GN=COTL1 PE=1 SV=3 | 7 |
| Q14028 | Cyclic nucleotide-gated cation channel beta-1 OS=Homo sapiens OX=9606 GN=CNGB1 PE=1 SV=2 | 1 |
| Q14061 | Cytochrome c oxidase copper chaperone OS=Homo sapiens OX=9606 GN=COX17 PE=1 SV=2 | 2 |
| Q14094 | Cyclin-I OS=Homo sapiens OX=9606 GN=CCNI PE=1 SV=1 | 2 |
| Q14103 | Heterogeneous nuclear ribonucleoprotein D0 OS=Homo sapiens OX=9606 GN=HNRNPD PE=1 SV=1 | 6 |
| Q14108 | Lysosome membrane protein 2 OS=Homo sapiens OX=9606 GN=SCARB2 PE=1 SV=2 | 9 |
| Q14118 | Dystroglycan 1 OS=Homo sapiens OX=9606 GN=DAG1 PE=1 SV=2 | 3 |
| Q14140 | SERTA domain-containing protein 2 OS=Homo sapiens OX=9606 GN=SERTAD2 PE=1 SV=1 | 2 |
| Q14147 | Probable ATP-dependent RNA helicase DHX34 OS=Homo sapiens OX=9606 GN=DHX34 PE=1 SV=2 | 1 |
| Q14152 | Eukaryotic translation initiation factor 3 subunit A OS=Homo sapiens OX=9606 GN=EIF3A PE=1 SV=1 | 14 |
| Q14155 | Rho guanine nucleotide exchange factor 7 OS=Homo sapiens OX=9606 GN=ARHGEF7 PE=1 SV=2 | 16 |
| Q14157 | Ubiquitin-associated protein 2-like OS=Homo sapiens OX=9606 GN=UBAP2L PE=1 SV=2 | 6 |
| Q14161 | ARF GTPase-activating protein GIT2 OS=Homo sapiens OX=9606 GN=GIT2 PE=1 SV=2 | 15 |
| Q14165 | Malectin OS=Homo sapiens OX=9606 GN=MLEC PE=1 SV=1 | 12 |
| Q14197 | Peptidyl-tRNA hydrolase ICT1, mitochondrial OS=Homo sapiens OX=9606 GN=MRPL58 PE=1 SV=1 | 5 |
| Q14203 | Dynactin subunit 1 OS=Homo sapiens OX=9606 GN=DCTN1 PE=1 SV=3 | 33 |
| Q14232 | Translation initiation factor eIF-2B subunit alpha OS=Homo sapiens OX=9606 GN=EIF2B1 PE=1 SV=1 | 1 |
| Q14240 | Eukaryotic initiation factor 4A-II OS=Homo sapiens OX=9606 GN=EIF4A2 PE=1 SV=2 | 7 |
| Q14247 | Src substrate cortactin OS=Homo sapiens OX=9606 GN=CTTN PE=1 SV=2 | 17 |
| Q14254 | Flotillin-2 OS=Homo sapiens OX=9606 GN=FLOT2 PE=1 SV=2 | 6 |
| Q14257 | Reticulocalbin-2 OS=Homo sapiens OX=9606 GN=RCN2 PE=1 SV=1 | 15 |
| Q14344 | Guanine nucleotide-binding protein subunit alpha-13 OS=Homo sapiens OX=9606 GN=GNA13 PE=1 SV=2 | 3 |
| Q14376 | UDP-glucose 4-epimerase OS=Homo sapiens OX=9606 GN=GALE PE=1 SV=2 | 1 |
| Q14393 | Growth arrest-specific protein 6 OS=Homo sapiens OX=9606 GN=GAS6 PE=1 SV=3 | 9 |
| Q14444 | Caprin-1 OS=Homo sapiens OX=9606 GN=CAPRIN1 PE=1 SV=2 | 6 |
| Q14451 | Growth factor receptor-bound protein 7 OS=Homo sapiens OX=9606 GN=GRB7 PE=1 SV=2 | 21 |
| Q14508 | WAP four-disulfide core domain protein 2 OS=Homo sapiens OX=9606 GN=WFDC2 PE=1 SV=2 | 3 |
| Q14532 | Keratin, type I cuticular Ha2 OS=Homo sapiens OX=9606 GN=KRT32 PE=1 SV=3 | 2 |
| Q14534 | Squalene monooxygenase OS=Homo sapiens OX=9606 GN=SQLE PE=1 SV=3 | 1 |
| Q14554 | Protein disulfide-isomerase A5 OS=Homo sapiens OX=9606 GN=PDIA5 PE=1 SV=1 | 14 |
| Q14558 | Phosphoribosyl pyrophosphate synthase-associated protein 1 OS=Homo sapiens OX=9606 GN=PRPSAP1 PE=1 SV=2 | 15 |
| Q14566 | DNA replication licensing factor MCM6 OS=Homo sapiens OX=9606 GN=MCM6 PE=1 SV=1 | 5 |
| Q14568 | Heat shock protein HSP 90-alpha A2 OS=Homo sapiens OX=9606 GN=HSP90AA2P PE=1 SV=2 | 1 |
| Q14571 | Inositol 1,4,5-trisphosphate receptor type 2 OS=Homo sapiens OX=9606 GN=ITPR2 PE=1 SV=2 | 1 |
| Q14573 | Inositol 1,4,5-trisphosphate receptor type 3 OS=Homo sapiens OX=9606 GN=ITPR3 PE=1 SV=2 | 1 |
| Q14624 | Inter-alpha-trypsin inhibitor heavy chain H4 OS=Homo sapiens OX=9606 GN=ITIH4 PE=1 SV=4 | 15 |
| Q14643 | Inositol 1,4,5-trisphosphate receptor type 1 OS=Homo sapiens OX=9606 GN=ITPR1 PE=1 SV=3 | 54 |
| Q14651 | Plastin-1 OS=Homo sapiens OX=9606 GN=PLS1 PE=1 SV=2 | 27 |
| Q14669 | E3 ubiquitin-protein ligase TRIP12 OS=Homo sapiens OX=9606 GN=TRIP12 PE=1 SV=1 | 4 |
| Q14677 | Clathrin interactor 1 OS=Homo sapiens OX=9606 GN=CLINT1 PE=1 SV=1 | 14 |
| Q14681 | BTB/POZ domain-containing protein KCTD2 OS=Homo sapiens OX=9606 GN=KCTD2 PE=1 SV=3 | 2 |
| Q14696 | LRP chaperone MESD OS=Homo sapiens OX=9606 GN=MESD PE=1 SV=2 | 10 |
| Q14697 | Neutral alpha-glucosidase AB OS=Homo sapiens OX=9606 GN=GANAB PE=1 SV=3 | 39 |
| Q14738 | Serine/threonine-protein phosphatase 2A 56 kDa regulatory subunit delta isoform OS=Homo sapiens OX=9606 GN=PPP2R5D PE=1 SV=1 | 17 |
| Q14766 | Latent-transforming growth factor beta-binding protein 1 OS=Homo sapiens OX=9606 GN=LTBP1 PE=1 SV=4 | 2 |
| Q14847 | LIM and SH3 domain protein 1 OS=Homo sapiens OX=9606 GN=LASP1 PE=1 SV=2 | 9 |
| Q14914 | Prostaglandin reductase 1 OS=Homo sapiens OX=9606 GN=PTGR1 PE=1 SV=2 | 17 |
| Q14956 | Transmembrane glycoprotein NMB OS=Homo sapiens OX=9606 GN=GPNMB PE=1 SV=2 | 1 |
| Q14997 | Proteasome activator complex subunit 4 OS=Homo sapiens OX=9606 GN=PSME4 PE=1 SV=2 | 4 |
| Q14C86 | GTPase-activating protein and VPS9 domain-containing protein 1 OS=Homo sapiens OX=9606 GN=GAPVD1 PE=1 SV=2 | 3 |
| Q14CX7 | N-alpha-acetyltransferase 25, NatB auxiliary subunit OS=Homo sapiens OX=9606 GN=NAA25 PE=1 SV=1 | 1 |
| Q15003 | Condensin complex subunit 2 OS=Homo sapiens OX=9606 GN=NCAPH PE=1 SV=3 | 3 |
| Q15005 | Signal peptidase complex subunit 2 OS=Homo sapiens OX=9606 GN=SPCS2 PE=1 SV=3 | 3 |
| Q15006 | ER membrane protein complex subunit 2 OS=Homo sapiens OX=9606 GN=EMC2 PE=1 SV=1 | 5 |
| Q15008 | 26S proteasome non-ATPase regulatory subunit 6 OS=Homo sapiens OX=9606 GN=PSMD6 PE=1 SV=1 | 18 |
| Q15018 | BRISC complex subunit Abraxas 2 OS=Homo sapiens OX=9606 GN=ABRAXAS2 PE=1 SV=2 | 4 |
| Q15019 | Septin-2 OS=Homo sapiens OX=9606 GN=SEPTIN2 PE=1 SV=1 | 7 |
| Q15020 | Squamous cell carcinoma antigen recognized by T-cells 3 OS=Homo sapiens OX=9606 GN=SART3 PE=1 SV=1 | 3 |
| Q15029 | 116 kDa U5 small nuclear ribonucleoprotein component OS=Homo sapiens OX=9606 GN=EFTUD2 PE=1 SV=1 | 12 |
| Q15042 | Rab3 GTPase-activating protein catalytic subunit OS=Homo sapiens OX=9606 GN=RAB3GAP1 PE=1 SV=3 | 18 |
| Q15043 | Metal cation symporter ZIP14 OS=Homo sapiens OX=9606 GN=SLC39A14 PE=1 SV=3 | 2 |
| Q15046 | Lysine--tRNA ligase OS=Homo sapiens OX=9606 GN=KARS1 PE=1 SV=3 | 19 |
| Q15051 | IQ calmodulin-binding motif-containing protein 1 OS=Homo sapiens OX=9606 GN=IQCB1 PE=1 SV=1 | 6 |
| Q15052 | Rho guanine nucleotide exchange factor 6 OS=Homo sapiens OX=9606 GN=ARHGEF6 PE=1 SV=2 | 1 |
| Q15056 | Eukaryotic translation initiation factor 4H OS=Homo sapiens OX=9606 GN=EIF4H PE=1 SV=5 | 4 |
| Q15067 | Peroxisomal acyl-coenzyme A oxidase 1 OS=Homo sapiens OX=9606 GN=ACOX1 PE=1 SV=3 | 7 |
| Q15084 | Protein disulfide-isomerase A6 OS=Homo sapiens OX=9606 GN=PDIA6 PE=1 SV=1 | 18 |
| Q15102 | Platelet-activating factor acetylhydrolase IB subunit alpha1 OS=Homo sapiens OX=9606 GN=PAFAH1B3 PE=1 SV=1 | 11 |
| Q15111 | Inactive phospholipase C-like protein 1 OS=Homo sapiens OX=9606 GN=PLCL1 PE=1 SV=3 | 8 |
| Q15120 | [Pyruvate dehydrogenase (acetyl-transferring)] kinase isozyme 3, mitochondrial OS=Homo sapiens OX=9606 GN=PDK3 PE=1 SV=1 | 3 |
| Q15121 | Astrocytic phosphoprotein PEA-15 OS=Homo sapiens OX=9606 GN=PEA15 PE=1 SV=2 | 4 |
| Q15126 | Phosphomevalonate kinase OS=Homo sapiens OX=9606 GN=PMVK PE=1 SV=3 | 6 |
| Q15165 | Serum paraoxonase/arylesterase 2 OS=Homo sapiens OX=9606 GN=PON2 PE=1 SV=4 | 9 |
| Q15166 | Serum paraoxonase/lactonase 3 OS=Homo sapiens OX=9606 GN=PON3 PE=1 SV=3 | 6 |
| Q15185 | Prostaglandin E synthase 3 OS=Homo sapiens OX=9606 GN=PTGES3 PE=1 SV=1 | 10 |
| Q15208 | Serine/threonine-protein kinase 38 OS=Homo sapiens OX=9606 GN=STK38 PE=1 SV=1 | 5 |
| Q15233 | Non-POU domain-containing octamer-binding protein OS=Homo sapiens OX=9606 GN=NONO PE=1 SV=4 | 3 |
| Q15257 | Serine/threonine-protein phosphatase 2A activator OS=Homo sapiens OX=9606 GN=PTPA PE=1 SV=3 | 14 |
| Q15269 | Periodic tryptophan protein 2 homolog OS=Homo sapiens OX=9606 GN=PWP2 PE=1 SV=2 | 1 |
| Q15276 | Rab GTPase-binding effector protein 1 OS=Homo sapiens OX=9606 GN=RABEP1 PE=1 SV=2 | 13 |
| Q15286 | Ras-related protein Rab-35 OS=Homo sapiens OX=9606 GN=RAB35 PE=1 SV=1 | 3 |
| Q15311 | RalA-binding protein 1 OS=Homo sapiens OX=9606 GN=RALBP1 PE=1 SV=3 | 1 |
| Q15349;Q86UE8;Q9UKI8 | Ribosomal protein S6 kinase alpha-2 OS=Homo sapiens OX=9606 GN=RPS6KA2 PE=1 SV=2 | 1 |
| Q15363 | Transmembrane emp24 domain-containing protein 2 OS=Homo sapiens OX=9606 GN=TMED2 PE=1 SV=1 | 4 |
| Q15365 | Poly(rC)-binding protein 1 OS=Homo sapiens OX=9606 GN=PCBP1 PE=1 SV=2 | 15 |
| Q15366 | Poly(rC)-binding protein 2 OS=Homo sapiens OX=9606 GN=PCBP2 PE=1 SV=1 | 8 |
| Q15369 | Elongin-C OS=Homo sapiens OX=9606 GN=ELOC PE=1 SV=1 | 8 |
| Q15370 | Elongin-B OS=Homo sapiens OX=9606 GN=ELOB PE=1 SV=1 | 5 |
| Q15382 | GTP-binding protein Rheb OS=Homo sapiens OX=9606 GN=RHEB PE=1 SV=1 | 4 |
| Q15386 | Ubiquitin-protein ligase E3C OS=Homo sapiens OX=9606 GN=UBE3C PE=1 SV=3 | 2 |
| Q15388 | Mitochondrial import receptor subunit TOM20 homolog OS=Homo sapiens OX=9606 GN=TOMM20 PE=1 SV=1 | 2 |
| Q15392 | Delta(24)-sterol reductase OS=Homo sapiens OX=9606 GN=DHCR24 PE=1 SV=2 | 8 |
| Q15393 | Splicing factor 3B subunit 3 OS=Homo sapiens OX=9606 GN=SF3B3 PE=1 SV=4 | 1 |
| Q15398 | Disks large-associated protein 5 OS=Homo sapiens OX=9606 GN=DLGAP5 PE=1 SV=2 | 35 |
| Q15404 | Ras suppressor protein 1 OS=Homo sapiens OX=9606 GN=RSU1 PE=1 SV=3 | 3 |
| Q15413 | Ryanodine receptor 3 OS=Homo sapiens OX=9606 GN=RYR3 PE=1 SV=3 | 1 |
| Q15417 | Calponin-3 OS=Homo sapiens OX=9606 GN=CNN3 PE=1 SV=1 | 10 |
| Q15418 | Ribosomal protein S6 kinase alpha-1 OS=Homo sapiens OX=9606 GN=RPS6KA1 PE=1 SV=2 | 3 |
| Q15427 | Splicing factor 3B subunit 4 OS=Homo sapiens OX=9606 GN=SF3B4 PE=1 SV=1 | 2 |
| Q15435 | Protein phosphatase 1 regulatory subunit 7 OS=Homo sapiens OX=9606 GN=PPP1R7 PE=1 SV=1 | 18 |
| Q15437 | Protein transport protein Sec23B OS=Homo sapiens OX=9606 GN=SEC23B PE=1 SV=2 | 5 |
| Q15477 | SKI2 subunit of superkiller complex protein OS=Homo sapiens OX=9606 GN=SKIC2 PE=1 SV=3 | 2 |
| Q15526 | Surfeit locus protein 1 OS=Homo sapiens OX=9606 GN=SURF1 PE=1 SV=1 | 4 |
| Q15555 | Microtubule-associated protein RP/EB family member 2 OS=Homo sapiens OX=9606 GN=MAPRE2 PE=1 SV=1 | 1 |
| Q15631 | Translin OS=Homo sapiens OX=9606 GN=TSN PE=1 SV=1 | 5 |
| Q15645 | Pachytene checkpoint protein 2 homolog OS=Homo sapiens OX=9606 GN=TRIP13 PE=1 SV=2 | 8 |
| Q15691 | Microtubule-associated protein RP/EB family member 1 OS=Homo sapiens OX=9606 GN=MAPRE1 PE=1 SV=3 | 14 |
| Q15714 | TSC22 domain family protein 1 OS=Homo sapiens OX=9606 GN=TSC22D1 PE=1 SV=3 | 3 |
| Q15717 | ELAV-like protein 1 OS=Homo sapiens OX=9606 GN=ELAVL1 PE=1 SV=2 | 11 |
| Q15738 | Sterol-4-alpha-carboxylate 3-dehydrogenase, decarboxylating OS=Homo sapiens OX=9606 GN=NSDHL PE=1 SV=2 | 9 |
| Q15758 | Neutral amino acid transporter B(0) OS=Homo sapiens OX=9606 GN=SLC1A5 PE=1 SV=2 | 2 |
| Q15771 | Ras-related protein Rab-30 OS=Homo sapiens OX=9606 GN=RAB30 PE=1 SV=2 | 3 |
| Q15773 | Myeloid leukemia factor 2 OS=Homo sapiens OX=9606 GN=MLF2 PE=1 SV=1 | 5 |
| Q15785 | Mitochondrial import receptor subunit TOM34 OS=Homo sapiens OX=9606 GN=TOMM34 PE=1 SV=2 | 25 |
| Q15797 | Mothers against decapentaplegic homolog 1 OS=Homo sapiens OX=9606 GN=SMAD1 PE=1 SV=1 | 5 |
| Q15811;Q9NZM3 | Intersectin-1 OS=Homo sapiens OX=9606 GN=ITSN1 PE=1 SV=3 | 1 |
| Q15813 | Tubulin-specific chaperone E OS=Homo sapiens OX=9606 GN=TBCE PE=1 SV=1 | 16 |
| Q15814 | Tubulin-specific chaperone C OS=Homo sapiens OX=9606 GN=TBCC PE=1 SV=2 | 13 |
| Q15833 | Syntaxin-binding protein 2 OS=Homo sapiens OX=9606 GN=STXBP2 PE=1 SV=2 | 21 |
| Q15836 | Vesicle-associated membrane protein 3 OS=Homo sapiens OX=9606 GN=VAMP3 PE=1 SV=3 | 3 |
| Q15843 | NEDD8 OS=Homo sapiens OX=9606 GN=NEDD8 PE=1 SV=1 | 4 |
| Q15904 | V-type proton ATPase subunit S1 OS=Homo sapiens OX=9606 GN=ATP6AP1 PE=1 SV=2 | 2 |
| Q15907 | Ras-related protein Rab-11B OS=Homo sapiens OX=9606 GN=RAB11B PE=1 SV=4 | 15 |
| Q16134 | Electron transfer flavoprotein-ubiquinone oxidoreductase, mitochondrial OS=Homo sapiens OX=9606 GN=ETFDH PE=1 SV=2 | 7 |
| Q16181 | Septin-7 OS=Homo sapiens OX=9606 GN=SEPTIN7 PE=1 SV=2 | 10 |
| Q16186 | Proteasomal ubiquitin receptor ADRM1 OS=Homo sapiens OX=9606 GN=ADRM1 PE=1 SV=2 | 5 |
| Q16204 | Coiled-coil domain-containing protein 6 OS=Homo sapiens OX=9606 GN=CCDC6 PE=1 SV=2 | 25 |
| Q16222;Q3KQV9 | UDP-N-acetylhexosamine pyrophosphorylase OS=Homo sapiens OX=9606 GN=UAP1 PE=1 SV=3 | 1 |
| Q16342 | Programmed cell death protein 2 OS=Homo sapiens OX=9606 GN=PDCD2 PE=1 SV=2 | 1 |
| Q16348 | Solute carrier family 15 member 2 OS=Homo sapiens OX=9606 GN=SLC15A2 PE=1 SV=2 | 1 |
| Q16401 | 26S proteasome non-ATPase regulatory subunit 5 OS=Homo sapiens OX=9606 GN=PSMD5 PE=1 SV=3 | 9 |
| Q16527 | Cysteine and glycine-rich protein 2 OS=Homo sapiens OX=9606 GN=CSRP2 PE=1 SV=3 | 1 |
| Q16531 | DNA damage-binding protein 1 OS=Homo sapiens OX=9606 GN=DDB1 PE=1 SV=1 | 36 |
| Q16539 | Mitogen-activated protein kinase 14 OS=Homo sapiens OX=9606 GN=MAPK14 PE=1 SV=3 | 1 |
| Q16540 | 39S ribosomal protein L23, mitochondrial OS=Homo sapiens OX=9606 GN=MRPL23 PE=1 SV=1 | 5 |
| Q16543 | Hsp90 co-chaperone Cdc37 OS=Homo sapiens OX=9606 GN=CDC37 PE=1 SV=1 | 14 |
| Q16555 | Dihydropyrimidinase-related protein 2 OS=Homo sapiens OX=9606 GN=DPYSL2 PE=1 SV=1 | 22 |
| Q16563 | Synaptophysin-like protein 1 OS=Homo sapiens OX=9606 GN=SYPL1 PE=1 SV=1 | 5 |
| Q16576 | Histone-binding protein RBBP7 OS=Homo sapiens OX=9606 GN=RBBP7 PE=1 SV=1 | 4 |
| Q16595 | Frataxin, mitochondrial OS=Homo sapiens OX=9606 GN=FXN PE=1 SV=2 | 7 |
| Q16626 | Male-enhanced antigen 1 OS=Homo sapiens OX=9606 GN=MEA1 PE=1 SV=2 | 2 |
| Q16637 | Survival motor neuron protein OS=Homo sapiens OX=9606 GN=SMN2 PE=1 SV=1 | 3 |
| Q16658 | Fascin OS=Homo sapiens OX=9606 GN=FSCN1 PE=1 SV=3 | 15 |
| Q16698 | 2,4-dienoyl-CoA reductase [(3E)-enoyl-CoA-producing], mitochondrial OS=Homo sapiens OX=9606 GN=DECR1 PE=1 SV=1 | 12 |
| Q16718 | NADH dehydrogenase [ubiquinone] 1 alpha subcomplex subunit 5 OS=Homo sapiens OX=9606 GN=NDUFA5 PE=1 SV=3 | 6 |
| Q16720 | Plasma membrane calcium-transporting ATPase 3 OS=Homo sapiens OX=9606 GN=ATP2B3 PE=1 SV=3 | 2 |
| Q16774 | Guanylate kinase OS=Homo sapiens OX=9606 GN=GUK1 PE=1 SV=2 | 4 |
| Q16775 | Hydroxyacylglutathione hydrolase, mitochondrial OS=Homo sapiens OX=9606 GN=HAGH PE=1 SV=2 | 1 |
| Q16795 | NADH dehydrogenase [ubiquinone] 1 alpha subcomplex subunit 9, mitochondrial OS=Homo sapiens OX=9606 GN=NDUFA9 PE=1 SV=2 | 6 |
| Q16836 | Hydroxyacyl-coenzyme A dehydrogenase, mitochondrial OS=Homo sapiens OX=9606 GN=HADH PE=1 SV=3 | 14 |
| Q16851 | UTP--glucose-1-phosphate uridylyltransferase OS=Homo sapiens OX=9606 GN=UGP2 PE=1 SV=5 | 28 |
| Q16854 | Deoxyguanosine kinase, mitochondrial OS=Homo sapiens OX=9606 GN=DGUOK PE=1 SV=2 | 5 |
| Q16864 | V-type proton ATPase subunit F OS=Homo sapiens OX=9606 GN=ATP6V1F PE=1 SV=2 | 2 |
| Q16891 | MICOS complex subunit MIC60 OS=Homo sapiens OX=9606 GN=IMMT PE=1 SV=1 | 47 |
| Q17R89 | Rho GTPase-activating protein 44 OS=Homo sapiens OX=9606 GN=ARHGAP44 PE=1 SV=1 | 5 |
| Q17RD7 | Synaptotagmin-16 OS=Homo sapiens OX=9606 GN=SYT16 PE=1 SV=2 | 2 |
| Q17RH7 | Tetra-peptide repeat homeobox-like protein OS=Homo sapiens OX=9606 GN=TPRXL PE=1 SV=3 | 3 |
| Q17RN3 | Protein FAM98C OS=Homo sapiens OX=9606 GN=FAM98C PE=1 SV=1 | 1 |
| Q27J81 | Inverted formin-2 OS=Homo sapiens OX=9606 GN=INF2 PE=1 SV=2 | 3 |
| Q2M1P5 | Kinesin-like protein KIF7 OS=Homo sapiens OX=9606 GN=KIF7 PE=1 SV=2 | 1 |
| Q2M2I5 | Keratin, type I cytoskeletal 24 OS=Homo sapiens OX=9606 GN=KRT24 PE=1 SV=1 | 2 |
| Q2M389 | WASH complex subunit 4 OS=Homo sapiens OX=9606 GN=WASHC4 PE=1 SV=2 | 6 |
| Q2NL98 | Vimentin-type intermediate filament-associated coiled-coil protein OS=Homo sapiens OX=9606 GN=VMAC PE=1 SV=1 | 1 |
| Q2Q1W2 | E3 ubiquitin-protein ligase TRIM71 OS=Homo sapiens OX=9606 GN=TRIM71 PE=1 SV=1 | 9 |
| Q2TAA2 | Isoamyl acetate-hydrolyzing esterase 1 homolog OS=Homo sapiens OX=9606 GN=IAH1 PE=1 SV=1 | 3 |
| Q2UY09 | Collagen alpha-1(XXVIII) chain OS=Homo sapiens OX=9606 GN=COL28A1 PE=2 SV=2 | 1 |
| Q32NB8 | CDP-diacylglycerol--glycerol-3-phosphate 3-phosphatidyltransferase, mitochondrial OS=Homo sapiens OX=9606 GN=PGS1 PE=2 SV=1 | 3 |
| Q3KRA6 | UPF0538 protein C2orf76 OS=Homo sapiens OX=9606 GN=C2orf76 PE=1 SV=3 | 3 |
| Q3LXA3 | Triokinase/FMN cyclase OS=Homo sapiens OX=9606 GN=TKFC PE=1 SV=2 | 13 |
| Q3MHD2 | Protein LSM12 OS=Homo sapiens OX=9606 GN=LSM12 PE=1 SV=2 | 1 |
| Q3MIX3 | Uncharacterized aarF domain-containing protein kinase 5 OS=Homo sapiens OX=9606 GN=ADCK5 PE=1 SV=3 | 2 |
| Q3MJ16 | Cytosolic phospholipase A2 epsilon OS=Homo sapiens OX=9606 GN=PLA2G4E PE=1 SV=4 | 5 |
| Q3SXM5 | Inactive hydroxysteroid dehydrogenase-like protein 1 OS=Homo sapiens OX=9606 GN=HSDL1 PE=1 SV=3 | 12 |
| Q3SYB3;Q5VV16;Q6VB84;Q8WXT5 | Forkhead box protein D4-like 6 OS=Homo sapiens OX=9606 GN=FOXD4L6 PE=1 SV=2 | 1 |
| Q3V6T2 | Girdin OS=Homo sapiens OX=9606 GN=CCDC88A PE=1 SV=2 | 2 |
| Q3YEC7 | Rab-like protein 6 OS=Homo sapiens OX=9606 GN=RABL6 PE=1 SV=2 | 5 |
| Q3ZCQ3 | Membrane protein FAM174B OS=Homo sapiens OX=9606 GN=FAM174B PE=2 SV=2 | 1 |
| Q3ZCQ8 | Mitochondrial import inner membrane translocase subunit TIM50 OS=Homo sapiens OX=9606 GN=TIMM50 PE=1 SV=2 | 4 |
| Q49A26 | Cytokine-like nuclear factor N-PAC OS=Homo sapiens OX=9606 GN=GLYR1 PE=1 SV=4 | 1 |
| Q49AH0 | Cerebral dopamine neurotrophic factor OS=Homo sapiens OX=9606 GN=CDNF PE=1 SV=2 | 1 |
| Q4G0F5 | Vacuolar protein sorting-associated protein 26B OS=Homo sapiens OX=9606 GN=VPS26B PE=1 SV=2 | 7 |
| Q4G0N4 | NAD kinase 2, mitochondrial OS=Homo sapiens OX=9606 GN=NADK2 PE=1 SV=2 | 14 |
| Q4G0X4 | BTB/POZ domain-containing protein KCTD21 OS=Homo sapiens OX=9606 GN=KCTD21 PE=1 SV=1 | 2 |
| Q4G176 | Malonate--CoA ligase ACSF3, mitochondrial OS=Homo sapiens OX=9606 GN=ACSF3 PE=1 SV=3 | 5 |
| Q4J6C6 | Prolyl endopeptidase-like OS=Homo sapiens OX=9606 GN=PREPL PE=1 SV=1 | 12 |
| Q4KMP7 | TBC1 domain family member 10B OS=Homo sapiens OX=9606 GN=TBC1D10B PE=1 SV=3 | 3 |
| Q4KWH8 | 1-phosphatidylinositol 4,5-bisphosphate phosphodiesterase eta-1 OS=Homo sapiens OX=9606 GN=PLCH1 PE=1 SV=1 | 1 |
| Q4L180 | Filamin A-interacting protein 1-like OS=Homo sapiens OX=9606 GN=FILIP1L PE=1 SV=2 | 1 |
| Q4V328 | GRIP1-associated protein 1 OS=Homo sapiens OX=9606 GN=GRIPAP1 PE=1 SV=2 | 16 |
| Q4V339;Q5JTY5;Q5RIA9;Q8IUF1;Q9BRT8 | Zinc-regulated GTPase metalloprotein activator 1F OS=Homo sapiens OX=9606 GN=ZNG1F PE=3 SV=1 | 2 |
| Q4VC31 | Protein MIX23 OS=Homo sapiens OX=9606 GN=MIX23 PE=1 SV=1 | 6 |
| Q4VCS5 | Angiomotin OS=Homo sapiens OX=9606 GN=AMOT PE=1 SV=1 | 17 |
| Q4VXU2 | Polyadenylate-binding protein 1-like OS=Homo sapiens OX=9606 GN=PABPC1L PE=2 SV=1 | 23 |
| Q52LJ0 | Protein FAM98B OS=Homo sapiens OX=9606 GN=FAM98B PE=1 SV=2 | 9 |
| Q53EU6 | Glycerol-3-phosphate acyltransferase 3 OS=Homo sapiens OX=9606 GN=GPAT3 PE=1 SV=2 | 1 |
| Q53F39 | Metallophosphoesterase 1 OS=Homo sapiens OX=9606 GN=MPPE1 PE=1 SV=2 | 1 |
| Q53FT3 | Protein Hikeshi OS=Homo sapiens OX=9606 GN=HIKESHI PE=1 SV=2 | 1 |
| Q53GG5 | PDZ and LIM domain protein 3 OS=Homo sapiens OX=9606 GN=PDLIM3 PE=1 SV=1 | 1 |
| Q53GQ0 | Very-long-chain 3-oxoacyl-CoA reductase OS=Homo sapiens OX=9606 GN=HSD17B12 PE=1 SV=2 | 6 |
| Q53H12 | Acylglycerol kinase, mitochondrial OS=Homo sapiens OX=9606 GN=AGK PE=1 SV=2 | 9 |
| Q53H82 | Endoribonuclease LACTB2 OS=Homo sapiens OX=9606 GN=LACTB2 PE=1 SV=2 | 9 |
| Q53H96 | Pyrroline-5-carboxylate reductase 3 OS=Homo sapiens OX=9606 GN=PYCR3 PE=1 SV=3 | 2 |
| Q567U6 | Coiled-coil domain-containing protein 93 OS=Homo sapiens OX=9606 GN=CCDC93 PE=1 SV=2 | 5 |
| Q587J7 | Putative ATP-dependent RNA helicase TDRD12 OS=Homo sapiens OX=9606 GN=TDRD12 PE=2 SV=2 | 2 |
| Q59H18 | Serine/threonine-protein kinase TNNI3K OS=Homo sapiens OX=9606 GN=TNNI3K PE=1 SV=3 | 1 |
| Q5BLP8 | Neuropeptide-like protein C4orf48 OS=Homo sapiens OX=9606 GN=C4orf48 PE=1 SV=3 | 3 |
| Q5EB52 | Mesoderm-specific transcript homolog protein OS=Homo sapiens OX=9606 GN=MEST PE=1 SV=2 | 1 |
| Q5GLZ8 | Probable E3 ubiquitin-protein ligase HERC4 OS=Homo sapiens OX=9606 GN=HERC4 PE=1 SV=1 | 7 |
| Q5H9R7 | Serine/threonine-protein phosphatase 6 regulatory subunit 3 OS=Homo sapiens OX=9606 GN=PPP6R3 PE=1 SV=2 | 3 |
| Q5HYI8 | Rab-like protein 3 OS=Homo sapiens OX=9606 GN=RABL3 PE=1 SV=1 | 3 |
| Q5HYK3 | 2-methoxy-6-polyprenyl-1,4-benzoquinol methylase, mitochondrial OS=Homo sapiens OX=9606 GN=COQ5 PE=1 SV=2 | 8 |
| Q5J8M3 | ER membrane protein complex subunit 4 OS=Homo sapiens OX=9606 GN=EMC4 PE=1 SV=2 | 2 |
| Q5JRA6 | Transport and Golgi organization protein 1 homolog OS=Homo sapiens OX=9606 GN=MIA3 PE=1 SV=1 | 27 |
| Q5JRX3 | Presequence protease, mitochondrial OS=Homo sapiens OX=9606 GN=PITRM1 PE=1 SV=3 | 52 |
| Q5JS37 | NHL repeat-containing protein 3 OS=Homo sapiens OX=9606 GN=NHLRC3 PE=2 SV=1 | 2 |
| Q5JSH3 | WD repeat-containing protein 44 OS=Homo sapiens OX=9606 GN=WDR44 PE=1 SV=1 | 24 |
| Q5JSP0 | FYVE, RhoGEF and PH domain-containing protein 3 OS=Homo sapiens OX=9606 GN=FGD3 PE=1 SV=1 | 1 |
| Q5JTD0 | Tight junction-associated protein 1 OS=Homo sapiens OX=9606 GN=TJAP1 PE=1 SV=1 | 8 |
| Q5JTV8 | Torsin-1A-interacting protein 1 OS=Homo sapiens OX=9606 GN=TOR1AIP1 PE=1 SV=2 | 3 |
| Q5JVS0 | Intracellular hyaluronan-binding protein 4 OS=Homo sapiens OX=9606 GN=HABP4 PE=1 SV=1 | 5 |
| Q5JWF2 | Guanine nucleotide-binding protein G(s) subunit alpha isoforms XLas OS=Homo sapiens OX=9606 GN=GNAS PE=1 SV=2 | 5 |
| Q5JWR5 | Protein dopey-1 OS=Homo sapiens OX=9606 GN=DOP1A PE=2 SV=1 | 4 |
| Q5M775 | Cytospin-B OS=Homo sapiens OX=9606 GN=SPECC1 PE=1 SV=1 | 2 |
| Q5MNZ6 | WD repeat domain phosphoinositide-interacting protein 3 OS=Homo sapiens OX=9606 GN=WDR45B PE=1 SV=2 | 1 |
| Q5NDL2 | EGF domain-specific O-linked N-acetylglucosamine transferase OS=Homo sapiens OX=9606 GN=EOGT PE=1 SV=1 | 1 |
| Q5QJ38 | Trichohyalin-like protein 1 OS=Homo sapiens OX=9606 GN=TCHHL1 PE=2 SV=1 | 1 |
| Q5R372 | Rab GTPase-activating protein 1-like OS=Homo sapiens OX=9606 GN=RABGAP1L PE=1 SV=1 | 8 |
| Q5RHP9 | Glutamate-rich protein 3 OS=Homo sapiens OX=9606 GN=ERICH3 PE=1 SV=1 | 1 |
| Q5RI15 | Cytochrome c oxidase assembly protein COX20, mitochondrial OS=Homo sapiens OX=9606 GN=COX20 PE=1 SV=2 | 4 |
| Q5SRE7 | Phytanoyl-CoA dioxygenase domain-containing protein 1 OS=Homo sapiens OX=9606 GN=PHYHD1 PE=1 SV=2 | 5 |
| Q5ST30 | Valine--tRNA ligase, mitochondrial OS=Homo sapiens OX=9606 GN=VARS2 PE=1 SV=2 | 9 |
| Q5SW96 | Low density lipoprotein receptor adapter protein 1 OS=Homo sapiens OX=9606 GN=LDLRAP1 PE=1 SV=3 | 8 |
| Q5SWX8 | Protein odr-4 homolog OS=Homo sapiens OX=9606 GN=ODR4 PE=1 SV=1 | 6 |
| Q5SXM8 | DNL-type zinc finger protein OS=Homo sapiens OX=9606 GN=DNLZ PE=1 SV=1 | 2 |
| Q5T013 | Putative hydroxypyruvate isomerase OS=Homo sapiens OX=9606 GN=HYI PE=1 SV=2 | 10 |
| Q5T124 | UBX domain-containing protein 11 OS=Homo sapiens OX=9606 GN=UBXN11 PE=1 SV=2 | 16 |
| Q5T160 | Probable arginine--tRNA ligase, mitochondrial OS=Homo sapiens OX=9606 GN=RARS2 PE=1 SV=1 | 14 |
| Q5T1M5 | FK506-binding protein 15 OS=Homo sapiens OX=9606 GN=FKBP15 PE=1 SV=2 | 9 |
| Q5T2W1 | Na(+)/H(+) exchange regulatory cofactor NHE-RF3 OS=Homo sapiens OX=9606 GN=PDZK1 PE=1 SV=2 | 14 |
| Q5T3F8 | CSC1-like protein 2 OS=Homo sapiens OX=9606 GN=TMEM63B PE=1 SV=1 | 4 |
| Q5T440 | Putative transferase CAF17, mitochondrial OS=Homo sapiens OX=9606 GN=IBA57 PE=1 SV=1 | 2 |
| Q5T447 | E3 ubiquitin-protein ligase HECTD3 OS=Homo sapiens OX=9606 GN=HECTD3 PE=1 SV=1 | 1 |
| Q5T4I8 | Putative uncharacterized protein C6orf52 OS=Homo sapiens OX=9606 GN=C6orf52 PE=2 SV=2 | 2 |
| Q5T4S7 | E3 ubiquitin-protein ligase UBR4 OS=Homo sapiens OX=9606 GN=UBR4 PE=1 SV=1 | 13 |
| Q5T5Y3 | Calmodulin-regulated spectrin-associated protein 1 OS=Homo sapiens OX=9606 GN=CAMSAP1 PE=1 SV=2 | 16 |
| Q5T6F2 | Ubiquitin-associated protein 2 OS=Homo sapiens OX=9606 GN=UBAP2 PE=1 SV=1 | 9 |
| Q5T6V5 | Queuosine salvage protein OS=Homo sapiens OX=9606 GN=C9orf64 PE=1 SV=1 | 5 |
| Q5T7P8 | Synaptotagmin-6 OS=Homo sapiens OX=9606 GN=SYT6 PE=1 SV=3 | 1 |
| Q5T8I9 | Small RNA 2'-O-methyltransferase OS=Homo sapiens OX=9606 GN=HENMT1 PE=1 SV=1 | 8 |
| Q5T8P6 | RNA-binding protein 26 OS=Homo sapiens OX=9606 GN=RBM26 PE=1 SV=3 | 4 |
| Q5TAQ9 | DDB1- and CUL4-associated factor 8 OS=Homo sapiens OX=9606 GN=DCAF8 PE=1 SV=1 | 1 |
| Q5TAX3 | Terminal uridylyltransferase 4 OS=Homo sapiens OX=9606 GN=TUT4 PE=1 SV=3 | 2 |
| Q5TCY1 | Tau-tubulin kinase 1 OS=Homo sapiens OX=9606 GN=TTBK1 PE=1 SV=2 | 1 |
| Q5TDH0 | Protein DDI1 homolog 2 OS=Homo sapiens OX=9606 GN=DDI2 PE=1 SV=1 | 4 |
| Q5TFE4 | 5'-nucleotidase domain-containing protein 1 OS=Homo sapiens OX=9606 GN=NT5DC1 PE=1 SV=1 | 3 |
| Q5TGZ0 | MICOS complex subunit MIC10 OS=Homo sapiens OX=9606 GN=MICOS10 PE=1 SV=1 | 1 |
| Q5UCC4 | ER membrane protein complex subunit 10 OS=Homo sapiens OX=9606 GN=EMC10 PE=1 SV=1 | 2 |
| Q5VIR6 | Vacuolar protein sorting-associated protein 53 homolog OS=Homo sapiens OX=9606 GN=VPS53 PE=1 SV=2 | 4 |
| Q5VT25 | Serine/threonine-protein kinase MRCK alpha OS=Homo sapiens OX=9606 GN=CDC42BPA PE=1 SV=1 | 1 |
| Q5VT97 | Rho GTPase-activating protein SYDE2 OS=Homo sapiens OX=9606 GN=SYDE2 PE=1 SV=2 | 1 |
| Q5VTE6 | Protein angel homolog 2 OS=Homo sapiens OX=9606 GN=ANGEL2 PE=1 SV=1 | 3 |
| Q5VTU8 | ATP synthase subunit epsilon-like protein, mitochondrial OS=Homo sapiens OX=9606 GN=ATP5F1EP2 PE=1 SV=1 | 2 |
| Q5VUM1 | Succinate dehydrogenase assembly factor 4, mitochondrial OS=Homo sapiens OX=9606 GN=SDHAF4 PE=1 SV=1 | 2 |
| Q5VV41 | Rho guanine nucleotide exchange factor 16 OS=Homo sapiens OX=9606 GN=ARHGEF16 PE=1 SV=1 | 5 |
| Q5VVH2 | Peptidyl-prolyl cis-trans isomerase FKBP1C OS=Homo sapiens OX=9606 GN=FKBP1C PE=3 SV=1 | 1 |
| Q5VVW2 | GTPase-activating Rap/Ran-GAP domain-like protein 3 OS=Homo sapiens OX=9606 GN=GARNL3 PE=2 SV=2 | 1 |
| Q5VW32 | BRO1 domain-containing protein BROX OS=Homo sapiens OX=9606 GN=BROX PE=1 SV=1 | 1 |
| Q5VW36 | Focadhesin OS=Homo sapiens OX=9606 GN=FOCAD PE=1 SV=2 | 6 |
| Q5VW38 | Protein GPR107 OS=Homo sapiens OX=9606 GN=GPR107 PE=1 SV=1 | 7 |
| Q5VWJ9 | Sorting nexin-30 OS=Homo sapiens OX=9606 GN=SNX30 PE=1 SV=1 | 16 |
| Q5VWZ2 | Lysophospholipase-like protein 1 OS=Homo sapiens OX=9606 GN=LYPLAL1 PE=1 SV=3 | 8 |
| Q5VY09 | Immediate early response gene 5 protein OS=Homo sapiens OX=9606 GN=IER5 PE=1 SV=3 | 1 |
| Q5VYK3 | Proteasome adapter and scaffold protein ECM29 OS=Homo sapiens OX=9606 GN=ECPAS PE=1 SV=2 | 7 |
| Q5VYY1 | Ankyrin repeat domain-containing protein 22 OS=Homo sapiens OX=9606 GN=ANKRD22 PE=2 SV=1 | 1 |
| Q5VZE5 | N-alpha-acetyltransferase 35, NatC auxiliary subunit OS=Homo sapiens OX=9606 GN=NAA35 PE=1 SV=1 | 2 |
| Q5VZK9 | F-actin-uncapping protein LRRC16A OS=Homo sapiens OX=9606 GN=CARMIL1 PE=1 SV=1 | 9 |
| Q5VZM2 | Ras-related GTP-binding protein B OS=Homo sapiens OX=9606 GN=RRAGB PE=1 SV=1 | 6 |
| Q5W111 | SPRY domain-containing protein 7 OS=Homo sapiens OX=9606 GN=SPRYD7 PE=1 SV=2 | 1 |
| Q5XKP0 | MICOS complex subunit MIC13 OS=Homo sapiens OX=9606 GN=MICOS13 PE=1 SV=1 | 2 |
| Q5XUX1 | F-box/WD repeat-containing protein 9 OS=Homo sapiens OX=9606 GN=FBXW9 PE=1 SV=3 | 9 |
| Q641Q2 | WASH complex subunit 2A OS=Homo sapiens OX=9606 GN=WASHC2A PE=1 SV=3 | 7 |
| Q66K14 | TBC1 domain family member 9B OS=Homo sapiens OX=9606 GN=TBC1D9B PE=1 SV=3 | 4 |
| Q66K74 | Microtubule-associated protein 1S OS=Homo sapiens OX=9606 GN=MAP1S PE=1 SV=2 | 27 |
| Q676U5 | Autophagy-related protein 16-1 OS=Homo sapiens OX=9606 GN=ATG16L1 PE=1 SV=2 | 3 |
| Q684P5 | Rap1 GTPase-activating protein 2 OS=Homo sapiens OX=9606 GN=RAP1GAP2 PE=1 SV=2 | 2 |
| Q68EM7 | Rho GTPase-activating protein 17 OS=Homo sapiens OX=9606 GN=ARHGAP17 PE=1 SV=1 | 15 |
| Q6AHZ1 | Zinc finger protein 518A OS=Homo sapiens OX=9606 GN=ZNF518A PE=1 SV=2 | 2 |
| Q6BCY4 | NADH-cytochrome b5 reductase 2 OS=Homo sapiens OX=9606 GN=CYB5R2 PE=1 SV=1 | 1 |
| Q6DD88 | Atlastin-3 OS=Homo sapiens OX=9606 GN=ATL3 PE=1 SV=1 | 19 |
| Q6DKJ4 | Nucleoredoxin OS=Homo sapiens OX=9606 GN=NXN PE=1 SV=2 | 3 |
| Q6DN14 | Multiple C2 and transmembrane domain-containing protein 1 OS=Homo sapiens OX=9606 GN=MCTP1 PE=2 SV=2 | 1 |
| Q6DN90 | IQ motif and SEC7 domain-containing protein 1 OS=Homo sapiens OX=9606 GN=IQSEC1 PE=1 SV=1 | 1 |
| Q6EMK4 | Vasorin OS=Homo sapiens OX=9606 GN=VASN PE=1 SV=1 | 1 |
| Q6FI81 | Anamorsin OS=Homo sapiens OX=9606 GN=CIAPIN1 PE=1 SV=2 | 13 |
| Q6GMV2 | Histone-lysine N-trimethyltransferase SMYD5 OS=Homo sapiens OX=9606 GN=SMYD5 PE=1 SV=2 | 1 |
| Q6GMV3 | Putative peptidyl-tRNA hydrolase PTRHD1 OS=Homo sapiens OX=9606 GN=PTRHD1 PE=1 SV=1 | 8 |
| Q6GQQ9 | OTU domain-containing protein 7B OS=Homo sapiens OX=9606 GN=OTUD7B PE=1 SV=1 | 3 |
| Q6IA69 | Glutamine-dependent NAD(+) synthetase OS=Homo sapiens OX=9606 GN=NADSYN1 PE=1 SV=3 | 8 |
| Q6IAA8 | Ragulator complex protein LAMTOR1 OS=Homo sapiens OX=9606 GN=LAMTOR1 PE=1 SV=2 | 3 |
| Q6IAN0 | Dehydrogenase/reductase SDR family member 7B OS=Homo sapiens OX=9606 GN=DHRS7B PE=1 SV=2 | 5 |
| Q6IMN6 | Caprin-2 OS=Homo sapiens OX=9606 GN=CAPRIN2 PE=1 SV=1 | 30 |
| Q6IN84 | rRNA methyltransferase 1, mitochondrial OS=Homo sapiens OX=9606 GN=MRM1 PE=1 SV=1 | 1 |
| Q6IPR1 | Electron transfer flavoprotein regulatory factor 1 OS=Homo sapiens OX=9606 GN=ETFRF1 PE=1 SV=2 | 2 |
| Q6IQ26 | DENN domain-containing protein 5A OS=Homo sapiens OX=9606 GN=DENND5A PE=1 SV=2 | 2 |
| Q6KCM7 | Calcium-binding mitochondrial carrier protein SCaMC-2 OS=Homo sapiens OX=9606 GN=SLC25A25 PE=1 SV=1 | 10 |
| Q6NUJ1 | Proactivator polypeptide-like 1 OS=Homo sapiens OX=9606 GN=PSAPL1 PE=2 SV=2 | 3 |
| Q6NUQ1 | RAD50-interacting protein 1 OS=Homo sapiens OX=9606 GN=RINT1 PE=1 SV=1 | 5 |
| Q6NVY1 | 3-hydroxyisobutyryl-CoA hydrolase, mitochondrial OS=Homo sapiens OX=9606 GN=HIBCH PE=1 SV=2 | 14 |
| Q6NW29 | RWD domain-containing protein 4 OS=Homo sapiens OX=9606 GN=RWDD4 PE=1 SV=3 | 1 |
| Q6NXE6 | Armadillo repeat-containing protein 6 OS=Homo sapiens OX=9606 GN=ARMC6 PE=1 SV=2 | 4 |
| Q6NXG1 | Epithelial splicing regulatory protein 1 OS=Homo sapiens OX=9606 GN=ESRP1 PE=1 SV=2 | 4 |
| Q6P1J6 | Phospholipase B1, membrane-associated OS=Homo sapiens OX=9606 GN=PLB1 PE=1 SV=3 | 3 |
| Q6P1K1 | Heme transporter HRG1 OS=Homo sapiens OX=9606 GN=SLC48A1 PE=1 SV=1 | 1 |
| Q6P1M0 | Long-chain fatty acid transport protein 4 OS=Homo sapiens OX=9606 GN=SLC27A4 PE=1 SV=1 | 16 |
| Q6P1N9 | Deoxyribonuclease TATDN1 OS=Homo sapiens OX=9606 GN=TATDN1 PE=1 SV=2 | 12 |
| Q6P1S2 | Protein C3orf33 OS=Homo sapiens OX=9606 GN=C3orf33 PE=1 SV=2 | 4 |
| Q6P1X6 | UPF0598 protein C8orf82 OS=Homo sapiens OX=9606 GN=C8orf82 PE=1 SV=2 | 1 |
| Q6P2E9 | Enhancer of mRNA-decapping protein 4 OS=Homo sapiens OX=9606 GN=EDC4 PE=1 SV=1 | 7 |
| Q6P2Q9 | Pre-mRNA-processing-splicing factor 8 OS=Homo sapiens OX=9606 GN=PRPF8 PE=1 SV=2 | 4 |
| Q6P3S1 | DENN domain-containing protein 1B OS=Homo sapiens OX=9606 GN=DENND1B PE=1 SV=2 | 1 |
| Q6P499 | NIPA-like protein 3 OS=Homo sapiens OX=9606 GN=NIPAL3 PE=1 SV=1 | 1 |
| Q6P4A8 | Phospholipase B-like 1 OS=Homo sapiens OX=9606 GN=PLBD1 PE=1 SV=2 | 16 |
| Q6P4F2 | Ferredoxin-2, mitochondrial OS=Homo sapiens OX=9606 GN=FDX2 PE=1 SV=2 | 2 |
| Q6P582 | Mitotic-spindle organizing protein 2A OS=Homo sapiens OX=9606 GN=MZT2A PE=1 SV=2 | 1 |
| Q6P587 | Acylpyruvase FAHD1, mitochondrial OS=Homo sapiens OX=9606 GN=FAHD1 PE=1 SV=2 | 4 |
| Q6P6B1 | Glutamate-rich protein 5 OS=Homo sapiens OX=9606 GN=ERICH5 PE=1 SV=1 | 1 |
| Q6P6C2 | RNA demethylase ALKBH5 OS=Homo sapiens OX=9606 GN=ALKBH5 PE=1 SV=2 | 1 |
| Q6P996 | Pyridoxal-dependent decarboxylase domain-containing protein 1 OS=Homo sapiens OX=9606 GN=PDXDC1 PE=1 SV=2 | 24 |
| Q6P9B6 | MTOR-associated protein MEAK7 OS=Homo sapiens OX=9606 GN=MEAK7 PE=1 SV=2 | 6 |
| Q6PGP7 | SKI3 subunit of superkiller complex protein OS=Homo sapiens OX=9606 GN=SKIC3 PE=1 SV=1 | 4 |
| Q6PHR2 | Serine/threonine-protein kinase ULK3 OS=Homo sapiens OX=9606 GN=ULK3 PE=1 SV=2 | 1 |
| Q6PI48 | Aspartate--tRNA ligase, mitochondrial OS=Homo sapiens OX=9606 GN=DARS2 PE=1 SV=1 | 3 |
| Q6PIU2 | Neutral cholesterol ester hydrolase 1 OS=Homo sapiens OX=9606 GN=NCEH1 PE=1 SV=3 | 13 |
| Q6PKG0 | La-related protein 1 OS=Homo sapiens OX=9606 GN=LARP1 PE=1 SV=2 | 13 |
| Q6PL24 | Protein TMED8 OS=Homo sapiens OX=9606 GN=TMED8 PE=1 SV=1 | 8 |
| Q6PML9 | Proton-coupled zinc antiporter SLC30A9, mitochondrial OS=Homo sapiens OX=9606 GN=SLC30A9 PE=1 SV=1 | 1 |
| Q6QHC5 | Sphingolipid delta(4)-desaturase/C4-monooxygenase DES2 OS=Homo sapiens OX=9606 GN=DEGS2 PE=1 SV=2 | 3 |
| Q6QNY0 | Biogenesis of lysosome-related organelles complex 1 subunit 3 OS=Homo sapiens OX=9606 GN=BLOC1S3 PE=1 SV=1 | 2 |
| Q6R327 | Rapamycin-insensitive companion of mTOR OS=Homo sapiens OX=9606 GN=RICTOR PE=1 SV=1 | 4 |
| Q6RW13 | Type-1 angiotensin II receptor-associated protein OS=Homo sapiens OX=9606 GN=AGTRAP PE=1 SV=1 | 2 |
| Q6UB35 | Monofunctional C1-tetrahydrofolate synthase, mitochondrial OS=Homo sapiens OX=9606 GN=MTHFD1L PE=1 SV=1 | 1 |
| Q6UN15 | Pre-mRNA 3'-end-processing factor FIP1 OS=Homo sapiens OX=9606 GN=FIP1L1 PE=1 SV=1 | 3 |
| Q6UUV7 | CREB-regulated transcription coactivator 3 OS=Homo sapiens OX=9606 GN=CRTC3 PE=1 SV=2 | 1 |
| Q6UUV9 | CREB-regulated transcription coactivator 1 OS=Homo sapiens OX=9606 GN=CRTC1 PE=1 SV=2 | 2 |
| Q6UW02 | Cytochrome P450 20A1 OS=Homo sapiens OX=9606 GN=CYP20A1 PE=1 SV=1 | 2 |
| Q6UW68 | Transmembrane protein 205 OS=Homo sapiens OX=9606 GN=TMEM205 PE=1 SV=1 | 5 |
| Q6UWP2 | Dehydrogenase/reductase SDR family member 11 OS=Homo sapiens OX=9606 GN=DHRS11 PE=1 SV=1 | 10 |
| Q6UWV6;Q9Y6X5 | Ectonucleotide pyrophosphatase/phosphodiesterase family member 7 OS=Homo sapiens OX=9606 GN=ENPP7 PE=1 SV=4 | 1 |
| Q6UXB8 | Peptidase inhibitor 16 OS=Homo sapiens OX=9606 GN=PI16 PE=1 SV=1 | 2 |
| Q6UXH1 | Protein disulfide isomerase CRELD2 OS=Homo sapiens OX=9606 GN=CRELD2 PE=1 SV=1 | 6 |
| Q6UXV4 | MICOS complex subunit MIC27 OS=Homo sapiens OX=9606 GN=APOOL PE=1 SV=1 | 6 |
| Q6VY07 | Phosphofurin acidic cluster sorting protein 1 OS=Homo sapiens OX=9606 GN=PACS1 PE=1 SV=2 | 4 |
| Q6W0C5 | Developmental pluripotency-associated protein 3 OS=Homo sapiens OX=9606 GN=DPPA3 PE=1 SV=1 | 7 |
| Q6WRI0 | Immunoglobulin superfamily member 10 OS=Homo sapiens OX=9606 GN=IGSF10 PE=1 SV=1 | 19 |
| Q6X9E4 | F-box/WD repeat-containing protein 12 OS=Homo sapiens OX=9606 GN=FBXW12 PE=2 SV=2 | 5 |
| Q6XQN6 | Nicotinate phosphoribosyltransferase OS=Homo sapiens OX=9606 GN=NAPRT PE=1 SV=2 | 11 |
| Q6Y288 | Beta-1,3-glucosyltransferase OS=Homo sapiens OX=9606 GN=B3GLCT PE=1 SV=2 | 8 |
| Q6Y7W6 | GRB10-interacting GYF protein 2 OS=Homo sapiens OX=9606 GN=GIGYF2 PE=1 SV=1 | 1 |
| Q6YN16 | Hydroxysteroid dehydrogenase-like protein 2 OS=Homo sapiens OX=9606 GN=HSDL2 PE=1 SV=1 | 7 |
| Q6YP21 | Kynurenine--oxoglutarate transaminase 3 OS=Homo sapiens OX=9606 GN=KYAT3 PE=1 SV=1 | 6 |
| Q6ZMI0 | Protein phosphatase 1 regulatory subunit 21 OS=Homo sapiens OX=9606 GN=PPP1R21 PE=1 SV=1 | 1 |
| Q6ZMV8 | Putative zinc finger protein 730 OS=Homo sapiens OX=9606 GN=ZNF730 PE=2 SV=1 | 1 |
| Q6ZRY4 | RNA-binding protein with multiple splicing 2 OS=Homo sapiens OX=9606 GN=RBPMS2 PE=1 SV=1 | 2 |
| Q6ZSJ8 | Uncharacterized protein C1orf122 OS=Homo sapiens OX=9606 GN=C1orf122 PE=4 SV=2 | 1 |
| Q6ZSS7 | Major facilitator superfamily domain-containing protein 6 OS=Homo sapiens OX=9606 GN=MFSD6 PE=1 SV=2 | 1 |
| Q6ZVM7 | TOM1-like protein 2 OS=Homo sapiens OX=9606 GN=TOM1L2 PE=1 SV=1 | 4 |
| Q709C8 | Intermembrane lipid transfer protein VPS13C OS=Homo sapiens OX=9606 GN=VPS13C PE=1 SV=1 | 44 |
| Q709F0 | Acyl-CoA dehydrogenase family member 11 OS=Homo sapiens OX=9606 GN=ACAD11 PE=1 SV=3 | 19 |
| Q70E73 | Ras-associated and pleckstrin homology domains-containing protein 1 OS=Homo sapiens OX=9606 GN=RAPH1 PE=1 SV=3 | 6 |
| Q70IA6 | MOB kinase activator 2 OS=Homo sapiens OX=9606 GN=MOB2 PE=1 SV=1 | 6 |
| Q70IA8 | MOB kinase activator 3C OS=Homo sapiens OX=9606 GN=MOB3C PE=1 SV=1 | 1 |
| Q70SY1 | Cyclic AMP-responsive element-binding protein 3-like protein 2 OS=Homo sapiens OX=9606 GN=CREB3L2 PE=1 SV=3 | 1 |
| Q712K3 | Ubiquitin-conjugating enzyme E2 R2 OS=Homo sapiens OX=9606 GN=UBE2R2 PE=1 SV=1 | 3 |
| Q71RC2 | La-related protein 4 OS=Homo sapiens OX=9606 GN=LARP4 PE=1 SV=3 | 2 |
| Q71UM5 | 40S ribosomal protein S27-like OS=Homo sapiens OX=9606 GN=RPS27L PE=1 SV=3 | 3 |
| Q7KZF4 | Staphylococcal nuclease domain-containing protein 1 OS=Homo sapiens OX=9606 GN=SND1 PE=1 SV=1 | 22 |
| Q7L099 | Protein RUFY3 OS=Homo sapiens OX=9606 GN=RUFY3 PE=1 SV=1 | 5 |
| Q7L0L9 | Transmembrane protein LOC653160 OS=Homo sapiens OX=9606 PE=2 SV=2 | 1 |
| Q7L0Y3 | tRNA methyltransferase 10 homolog C OS=Homo sapiens OX=9606 GN=TRMT10C PE=1 SV=2 | 13 |
| Q7L1Q6 | eIF5-mimic protein 2 OS=Homo sapiens OX=9606 GN=BZW1 PE=1 SV=1 | 2 |
| Q7L2E3 | ATP-dependent RNA helicase DHX30 OS=Homo sapiens OX=9606 GN=DHX30 PE=1 SV=1 | 15 |
| Q7L2H7 | Eukaryotic translation initiation factor 3 subunit M OS=Homo sapiens OX=9606 GN=EIF3M PE=1 SV=1 | 7 |
| Q7L3B6 | Hsp90 co-chaperone Cdc37-like 1 OS=Homo sapiens OX=9606 GN=CDC37L1 PE=1 SV=1 | 3 |
| Q7L3T8 | Probable proline--tRNA ligase, mitochondrial OS=Homo sapiens OX=9606 GN=PARS2 PE=1 SV=1 | 5 |
| Q7L576 | Cytoplasmic FMR1-interacting protein 1 OS=Homo sapiens OX=9606 GN=CYFIP1 PE=1 SV=1 | 9 |
| Q7L5D6 | Golgi to ER traffic protein 4 homolog OS=Homo sapiens OX=9606 GN=GET4 PE=1 SV=1 | 6 |
| Q7L5N1 | COP9 signalosome complex subunit 6 OS=Homo sapiens OX=9606 GN=COPS6 PE=1 SV=1 | 11 |
| Q7L5Y1 | Mitochondrial enolase superfamily member 1 OS=Homo sapiens OX=9606 GN=ENOSF1 PE=1 SV=1 | 7 |
| Q7LBR1 | Charged multivesicular body protein 1b OS=Homo sapiens OX=9606 GN=CHMP1B PE=1 SV=1 | 2 |
| Q7LG56 | Ribonucleoside-diphosphate reductase subunit M2 B OS=Homo sapiens OX=9606 GN=RRM2B PE=1 SV=1 | 5 |
| Q7RTV0 | PHD finger-like domain-containing protein 5A OS=Homo sapiens OX=9606 GN=PHF5A PE=1 SV=1 | 1 |
| Q7RTY7 | Ovochymase-1 OS=Homo sapiens OX=9606 GN=OVCH1 PE=2 SV=2 | 9 |
| Q7Z2D5 | Phospholipid phosphatase-related protein type 4 OS=Homo sapiens OX=9606 GN=PLPPR4 PE=1 SV=1 | 1 |
| Q7Z2K6 | Endoplasmic reticulum metallopeptidase 1 OS=Homo sapiens OX=9606 GN=ERMP1 PE=1 SV=2 | 16 |
| Q7Z2W9 | 39S ribosomal protein L21, mitochondrial OS=Homo sapiens OX=9606 GN=MRPL21 PE=1 SV=2 | 2 |
| Q7Z2Z2 | Elongation factor-like GTPase 1 OS=Homo sapiens OX=9606 GN=EFL1 PE=1 SV=2 | 18 |
| Q7Z392 | Trafficking protein particle complex subunit 11 OS=Homo sapiens OX=9606 GN=TRAPPC11 PE=1 SV=2 | 4 |
| Q7Z3B4 | Nucleoporin p54 OS=Homo sapiens OX=9606 GN=NUP54 PE=1 SV=2 | 9 |
| Q7Z3E5 | LisH domain-containing protein ARMC9 OS=Homo sapiens OX=9606 GN=ARMC9 PE=1 SV=3 | 10 |
| Q7Z3J2 | VPS35 endosomal protein-sorting factor-like OS=Homo sapiens OX=9606 GN=VPS35L PE=1 SV=2 | 4 |
| Q7Z3U7 | Protein MON2 homolog OS=Homo sapiens OX=9606 GN=MON2 PE=1 SV=3 | 7 |
| Q7Z3Z0 | Keratin, type I cytoskeletal 25 OS=Homo sapiens OX=9606 GN=KRT25 PE=1 SV=1 | 1 |
| Q7Z3Z3 | Piwi-like protein 3 OS=Homo sapiens OX=9606 GN=PIWIL3 PE=2 SV=2 | 20 |
| Q7Z408 | CUB and sushi domain-containing protein 2 OS=Homo sapiens OX=9606 GN=CSMD2 PE=1 SV=2 | 1 |
| Q7Z478 | ATP-dependent RNA helicase DHX29 OS=Homo sapiens OX=9606 GN=DHX29 PE=1 SV=2 | 10 |
| Q7Z4G1 | COMM domain-containing protein 6 OS=Homo sapiens OX=9606 GN=COMMD6 PE=1 SV=1 | 4 |
| Q7Z4H3 | 5'-deoxynucleotidase HDDC2 OS=Homo sapiens OX=9606 GN=HDDC2 PE=1 SV=1 | 10 |
| Q7Z4V5 | Hepatoma-derived growth factor-related protein 2 OS=Homo sapiens OX=9606 GN=HDGFL2 PE=1 SV=1 | 1 |
| Q7Z4W1 | L-xylulose reductase OS=Homo sapiens OX=9606 GN=DCXR PE=1 SV=2 | 15 |
| Q7Z569 | BRCA1-associated protein OS=Homo sapiens OX=9606 GN=BRAP PE=1 SV=2 | 4 |
| Q7Z5G4 | Golgin subfamily A member 7 OS=Homo sapiens OX=9606 GN=GOLGA7 PE=1 SV=2 | 2 |
| Q7Z5N4 | Protein sidekick-1 OS=Homo sapiens OX=9606 GN=SDK1 PE=2 SV=3 | 1 |
| Q7Z6M4 | Transcription termination factor 4, mitochondrial OS=Homo sapiens OX=9606 GN=MTERF4 PE=1 SV=3 | 1 |
| Q7Z6Z7 | E3 ubiquitin-protein ligase HUWE1 OS=Homo sapiens OX=9606 GN=HUWE1 PE=1 SV=3 | 22 |
| Q7Z739 | YTH domain-containing family protein 3 OS=Homo sapiens OX=9606 GN=YTHDF3 PE=1 SV=1 | 3 |
| Q7Z7C7 | Stimulated by retinoic acid gene 8 protein homolog OS=Homo sapiens OX=9606 GN=STRA8 PE=1 SV=1 | 1 |
| Q7Z7F7 | 39S ribosomal protein L55, mitochondrial OS=Homo sapiens OX=9606 GN=MRPL55 PE=1 SV=1 | 2 |
| Q7Z7H5 | Transmembrane emp24 domain-containing protein 4 OS=Homo sapiens OX=9606 GN=TMED4 PE=1 SV=1 | 3 |
| Q7Z7J7 | LHFPL tetraspan subfamily member 4 protein OS=Homo sapiens OX=9606 GN=LHFPL4 PE=1 SV=1 | 1 |
| Q7Z7N9 | Transmembrane protein 179B OS=Homo sapiens OX=9606 GN=TMEM179B PE=1 SV=1 | 1 |
| Q86SE8 | Nucleoplasmin-2 OS=Homo sapiens OX=9606 GN=NPM2 PE=1 SV=1 | 8 |
| Q86SH2 | Zygote arrest protein 1 OS=Homo sapiens OX=9606 GN=ZAR1 PE=1 SV=1 | 11 |
| Q86SQ4 | Adhesion G-protein coupled receptor G6 OS=Homo sapiens OX=9606 GN=ADGRG6 PE=1 SV=3 | 1 |
| Q86SX6 | Glutaredoxin-related protein 5, mitochondrial OS=Homo sapiens OX=9606 GN=GLRX5 PE=1 SV=2 | 3 |
| Q86SZ2 | Trafficking protein particle complex subunit 6B OS=Homo sapiens OX=9606 GN=TRAPPC6B PE=1 SV=1 | 6 |
| Q86T03 | Type 1 phosphatidylinositol 4,5-bisphosphate 4-phosphatase OS=Homo sapiens OX=9606 GN=PIP4P1 PE=1 SV=1 | 1 |
| Q86TI0 | TBC1 domain family member 1 OS=Homo sapiens OX=9606 GN=TBC1D1 PE=1 SV=2 | 3 |
| Q86TM6 | E3 ubiquitin-protein ligase synoviolin OS=Homo sapiens OX=9606 GN=SYVN1 PE=1 SV=2 | 2 |
| Q86TP1 | Exopolyphosphatase PRUNE1 OS=Homo sapiens OX=9606 GN=PRUNE1 PE=1 SV=2 | 5 |
| Q86U28 | Iron-sulfur cluster assembly 2 homolog, mitochondrial OS=Homo sapiens OX=9606 GN=ISCA2 PE=1 SV=2 | 3 |
| Q86U42 | Polyadenylate-binding protein 2 OS=Homo sapiens OX=9606 GN=PABPN1 PE=1 SV=3 | 2 |
| Q86U90 | Threonylcarbamoyl-AMP synthase OS=Homo sapiens OX=9606 GN=YRDC PE=1 SV=1 | 5 |
| Q86UA1 | Pre-mRNA-processing factor 39 OS=Homo sapiens OX=9606 GN=PRPF39 PE=1 SV=3 | 4 |
| Q86UE4 | Protein LYRIC OS=Homo sapiens OX=9606 GN=MTDH PE=1 SV=2 | 3 |
| Q86UL3 | Glycerol-3-phosphate acyltransferase 4 OS=Homo sapiens OX=9606 GN=GPAT4 PE=1 SV=1 | 2 |
| Q86UP2 | Kinectin OS=Homo sapiens OX=9606 GN=KTN1 PE=1 SV=1 | 58 |
| Q86UW7 | Calcium-dependent secretion activator 2 OS=Homo sapiens OX=9606 GN=CADPS2 PE=1 SV=2 | 4 |
| Q86UW9 | Probable E3 ubiquitin-protein ligase DTX2 OS=Homo sapiens OX=9606 GN=DTX2 PE=1 SV=3 | 2 |
| Q86UY8 | 5'-nucleotidase domain-containing protein 3 OS=Homo sapiens OX=9606 GN=NT5DC3 PE=1 SV=1 | 5 |
| Q86V21 | Acetoacetyl-CoA synthetase OS=Homo sapiens OX=9606 GN=AACS PE=1 SV=1 | 5 |
| Q86VE9 | Serine incorporator 5 OS=Homo sapiens OX=9606 GN=SERINC5 PE=2 SV=1 | 1 |
| Q86VN1 | Vacuolar protein-sorting-associated protein 36 OS=Homo sapiens OX=9606 GN=VPS36 PE=1 SV=1 | 8 |
| Q86VP1 | Tax1-binding protein 1 OS=Homo sapiens OX=9606 GN=TAX1BP1 PE=1 SV=2 | 12 |
| Q86VP6 | Cullin-associated NEDD8-dissociated protein 1 OS=Homo sapiens OX=9606 GN=CAND1 PE=1 SV=2 | 40 |
| Q86VQ1 | Glucocorticoid-induced transcript 1 protein OS=Homo sapiens OX=9606 GN=GLCCI1 PE=1 SV=1 | 2 |
| Q86VR2 | Reticulophagy regulator 3 OS=Homo sapiens OX=9606 GN=RETREG3 PE=1 SV=1 | 3 |
| Q86VS8 | Protein Hook homolog 3 OS=Homo sapiens OX=9606 GN=HOOK3 PE=1 SV=2 | 8 |
| Q86VX2 | COMM domain-containing protein 7 OS=Homo sapiens OX=9606 GN=COMMD7 PE=1 SV=2 | 3 |
| Q86W24 | NACHT, LRR and PYD domains-containing protein 14 OS=Homo sapiens OX=9606 GN=NLRP14 PE=1 SV=1 | 35 |
| Q86W56 | Poly(ADP-ribose) glycohydrolase OS=Homo sapiens OX=9606 GN=PARG PE=1 SV=1 | 1 |
| Q86WA6 | Valacyclovir hydrolase OS=Homo sapiens OX=9606 GN=BPHL PE=1 SV=1 | 9 |
| Q86WB0 | Zinc finger C3HC-type protein 1 OS=Homo sapiens OX=9606 GN=ZC3HC1 PE=1 SV=1 | 1 |
| Q86WI1 | Fibrocystin-L OS=Homo sapiens OX=9606 GN=PKHD1L1 PE=2 SV=2 | 1 |
| Q86WJ1 | Chromodomain-helicase-DNA-binding protein 1-like OS=Homo sapiens OX=9606 GN=CHD1L PE=1 SV=3 | 2 |
| Q86WQ0 | Nuclear receptor 2C2-associated protein OS=Homo sapiens OX=9606 GN=NR2C2AP PE=1 SV=1 | 3 |
| Q86WR0 | Coiled-coil domain-containing protein 25 OS=Homo sapiens OX=9606 GN=CCDC25 PE=1 SV=2 | 10 |
| Q86WS3 | Oocyte-secreted protein 2 OS=Homo sapiens OX=9606 GN=OOSP2 PE=1 SV=1 | 3 |
| Q86WU2 | Probable D-lactate dehydrogenase, mitochondrial OS=Homo sapiens OX=9606 GN=LDHD PE=1 SV=1 | 4 |
| Q86X29 | Lipolysis-stimulated lipoprotein receptor OS=Homo sapiens OX=9606 GN=LSR PE=1 SV=4 | 7 |
| Q86X40 | Leucine-rich repeat-containing protein 28 OS=Homo sapiens OX=9606 GN=LRRC28 PE=2 SV=1 | 1 |
| Q86X51 | EZH inhibitory protein OS=Homo sapiens OX=9606 GN=EZHIP PE=1 SV=1 | 7 |
| Q86X55 | Histone-arginine methyltransferase CARM1 OS=Homo sapiens OX=9606 GN=CARM1 PE=1 SV=3 | 2 |
| Q86X76 | Deaminated glutathione amidase OS=Homo sapiens OX=9606 GN=NIT1 PE=1 SV=2 | 9 |
| Q86X83 | COMM domain-containing protein 2 OS=Homo sapiens OX=9606 GN=COMMD2 PE=1 SV=2 | 2 |
| Q86XL3 | Ankyrin repeat and LEM domain-containing protein 2 OS=Homo sapiens OX=9606 GN=ANKLE2 PE=1 SV=4 | 3 |
| Q86XN7 | Proline and serine-rich protein 1 OS=Homo sapiens OX=9606 GN=PROSER1 PE=1 SV=2 | 12 |
| Q86Y82 | Syntaxin-12 OS=Homo sapiens OX=9606 GN=STX12 PE=1 SV=1 | 4 |
| Q86YB8 | ERO1-like protein beta OS=Homo sapiens OX=9606 GN=ERO1B PE=1 SV=2 | 10 |
| Q86YL5 | Testis development-related protein OS=Homo sapiens OX=9606 GN=TDRP PE=1 SV=2 | 4 |
| Q86YS6 | Ras-related protein Rab-43 OS=Homo sapiens OX=9606 GN=RAB43 PE=1 SV=1 | 1 |
| Q86YS7 | C2 domain-containing protein 5 OS=Homo sapiens OX=9606 GN=C2CD5 PE=1 SV=1 | 4 |
| Q8IUC0 | Keratin-associated protein 13-1 OS=Homo sapiens OX=9606 GN=KRTAP13-1 PE=2 SV=2 | 1 |
| Q8IUD2 | ELKS/Rab6-interacting/CAST family member 1 OS=Homo sapiens OX=9606 GN=ERC1 PE=1 SV=1 | 9 |
| Q8IUH4 | Palmitoyltransferase ZDHHC13 OS=Homo sapiens OX=9606 GN=ZDHHC13 PE=1 SV=3 | 5 |
| Q8IUX8 | Epidermal growth factor-like protein 6 OS=Homo sapiens OX=9606 GN=EGFL6 PE=1 SV=1 | 1 |
| Q8IVD9 | NudC domain-containing protein 3 OS=Homo sapiens OX=9606 GN=NUDCD3 PE=1 SV=3 | 12 |
| Q8IVF2 | Protein AHNAK2 OS=Homo sapiens OX=9606 GN=AHNAK2 PE=1 SV=2 | 1 |
| Q8IVF6 | Ankyrin repeat domain-containing protein 18A OS=Homo sapiens OX=9606 GN=ANKRD18A PE=2 SV=3 | 1 |
| Q8IVH4 | Methylmalonic aciduria type A protein, mitochondrial OS=Homo sapiens OX=9606 GN=MMAA PE=1 SV=1 | 8 |
| Q8IVJ1;Q96GZ6;Q96JW4 | Solute carrier family 41 member 1 OS=Homo sapiens OX=9606 GN=SLC41A1 PE=1 SV=2 | 1 |
| Q8IVM0 | Coiled-coil domain-containing protein 50 OS=Homo sapiens OX=9606 GN=CCDC50 PE=1 SV=1 | 4 |
| Q8IVS2 | Malonyl-CoA-acyl carrier protein transacylase, mitochondrial OS=Homo sapiens OX=9606 GN=MCAT PE=1 SV=2 | 6 |
| Q8IVT5 | Kinase suppressor of Ras 1 OS=Homo sapiens OX=9606 GN=KSR1 PE=1 SV=3 | 6 |
| Q8IW35 | Centrosomal protein of 97 kDa OS=Homo sapiens OX=9606 GN=CEP97 PE=1 SV=1 | 2 |
| Q8IW45 | ATP-dependent (S)-NAD(P)H-hydrate dehydratase OS=Homo sapiens OX=9606 GN=NAXD PE=1 SV=1 | 8 |
| Q8IWA4 | Mitofusin-1 OS=Homo sapiens OX=9606 GN=MFN1 PE=1 SV=3 | 18 |
| Q8IWB7 | WD repeat and FYVE domain-containing protein 1 OS=Homo sapiens OX=9606 GN=WDFY1 PE=1 SV=1 | 22 |
| Q8IWB9 | Testis-expressed protein 2 OS=Homo sapiens OX=9606 GN=TEX2 PE=1 SV=2 | 4 |
| Q8IWJ2 | GRIP and coiled-coil domain-containing protein 2 OS=Homo sapiens OX=9606 GN=GCC2 PE=1 SV=4 | 4 |
| Q8IWL3 | Iron-sulfur cluster co-chaperone protein HscB OS=Homo sapiens OX=9606 GN=HSCB PE=1 SV=3 | 3 |
| Q8IWV8 | E3 ubiquitin-protein ligase UBR2 OS=Homo sapiens OX=9606 GN=UBR2 PE=1 SV=1 | 6 |
| Q8IWW6 | Rho GTPase-activating protein 12 OS=Homo sapiens OX=9606 GN=ARHGAP12 PE=1 SV=1 | 5 |
| Q8IWY4 | Signal peptide, CUB and EGF-like domain-containing protein 1 OS=Homo sapiens OX=9606 GN=SCUBE1 PE=1 SV=3 | 1 |
| Q8IX90 | Spindle and kinetochore-associated protein 3 OS=Homo sapiens OX=9606 GN=SKA3 PE=1 SV=2 | 1 |
| Q8IXB1 | DnaJ homolog subfamily C member 10 OS=Homo sapiens OX=9606 GN=DNAJC10 PE=1 SV=2 | 7 |
| Q8IXI1 | Mitochondrial Rho GTPase 2 OS=Homo sapiens OX=9606 GN=RHOT2 PE=1 SV=2 | 4 |
| Q8IXI2 | Mitochondrial Rho GTPase 1 OS=Homo sapiens OX=9606 GN=RHOT1 PE=1 SV=2 | 6 |
| Q8IXM3 | 39S ribosomal protein L41, mitochondrial OS=Homo sapiens OX=9606 GN=MRPL41 PE=1 SV=1 | 5 |
| Q8IXN7 | N-acetylaspartylglutamate synthase A OS=Homo sapiens OX=9606 GN=RIMKLA PE=1 SV=2 | 3 |
| Q8IXQ3 | Uncharacterized protein C9orf40 OS=Homo sapiens OX=9606 GN=C9orf40 PE=1 SV=1 | 2 |
| Q8IXQ5 | Kelch-like protein 7 OS=Homo sapiens OX=9606 GN=KLHL7 PE=1 SV=2 | 6 |
| Q8IXZ2 | Zinc finger CCCH domain-containing protein 3 OS=Homo sapiens OX=9606 GN=ZC3H3 PE=1 SV=3 | 1 |
| Q8IY17 | Patatin-like phospholipase domain-containing protein 6 OS=Homo sapiens OX=9606 GN=PNPLA6 PE=1 SV=3 | 8 |
| Q8IYB1 | Nucleotidyltransferase MB21D2 OS=Homo sapiens OX=9606 GN=MB21D2 PE=1 SV=3 | 10 |
| Q8IYB8 | ATP-dependent RNA helicase SUPV3L1, mitochondrial OS=Homo sapiens OX=9606 GN=SUPV3L1 PE=1 SV=1 | 14 |
| Q8IYD1 | Eukaryotic peptide chain release factor GTP-binding subunit ERF3B OS=Homo sapiens OX=9606 GN=GSPT2 PE=1 SV=2 | 4 |
| Q8IYI6 | Exocyst complex component 8 OS=Homo sapiens OX=9606 GN=EXOC8 PE=1 SV=2 | 3 |
| Q8IYL2 | Probable tRNA (uracil-O(2)-)-methyltransferase OS=Homo sapiens OX=9606 GN=TRMT44 PE=1 SV=2 | 1 |
| Q8IYQ7 | Threonine synthase-like 1 OS=Homo sapiens OX=9606 GN=THNSL1 PE=1 SV=2 | 21 |
| Q8IYS2 | Uncharacterized protein KIAA2013 OS=Homo sapiens OX=9606 GN=KIAA2013 PE=1 SV=1 | 2 |
| Q8IZ07 | Ankyrin repeat domain-containing protein 13A OS=Homo sapiens OX=9606 GN=ANKRD13A PE=1 SV=3 | 7 |
| Q8IZ81 | ELMO domain-containing protein 2 OS=Homo sapiens OX=9606 GN=ELMOD2 PE=1 SV=1 | 5 |
| Q8IZ83 | Aldehyde dehydrogenase family 16 member A1 OS=Homo sapiens OX=9606 GN=ALDH16A1 PE=1 SV=2 | 8 |
| Q8IZA3 | Histone H1.8 OS=Homo sapiens OX=9606 GN=H1-8 PE=2 SV=1 | 7 |
| Q8IZP0 | Abl interactor 1 OS=Homo sapiens OX=9606 GN=ABI1 PE=1 SV=4 | 3 |
| Q8IZU2 | WD repeat-containing protein 17 OS=Homo sapiens OX=9606 GN=WDR17 PE=2 SV=2 | 1 |
| Q8N0U8 | Vitamin K epoxide reductase complex subunit 1-like protein 1 OS=Homo sapiens OX=9606 GN=VKORC1L1 PE=1 SV=2 | 1 |
| Q8N0W3 | L-fucose kinase OS=Homo sapiens OX=9606 GN=FCSK PE=1 SV=2 | 17 |
| Q8N0X4 | Citramalyl-CoA lyase, mitochondrial OS=Homo sapiens OX=9606 GN=CLYBL PE=1 SV=2 | 11 |
| Q8N0Z6 | Tetratricopeptide repeat protein 5 OS=Homo sapiens OX=9606 GN=TTC5 PE=1 SV=2 | 3 |
| Q8N111 | Cell cycle exit and neuronal differentiation protein 1 OS=Homo sapiens OX=9606 GN=CEND1 PE=1 SV=1 | 3 |
| Q8N122 | Regulatory-associated protein of mTOR OS=Homo sapiens OX=9606 GN=RPTOR PE=1 SV=1 | 1 |
| Q8N128 | Protein FAM177A1 OS=Homo sapiens OX=9606 GN=FAM177A1 PE=1 SV=1 | 1 |
| Q8N129 | Protein canopy homolog 4 OS=Homo sapiens OX=9606 GN=CNPY4 PE=1 SV=1 | 2 |
| Q8N131 | Porimin OS=Homo sapiens OX=9606 GN=TMEM123 PE=1 SV=1 | 2 |
| Q8N183 | NADH dehydrogenase [ubiquinone] 1 alpha subcomplex assembly factor 2 OS=Homo sapiens OX=9606 GN=NDUFAF2 PE=1 SV=1 | 8 |
| Q8N1F7 | Nuclear pore complex protein Nup93 OS=Homo sapiens OX=9606 GN=NUP93 PE=1 SV=2 | 16 |
| Q8N1G4 | Leucine-rich repeat-containing protein 47 OS=Homo sapiens OX=9606 GN=LRRC47 PE=1 SV=1 | 22 |
| Q8N1I0 | Dedicator of cytokinesis protein 4 OS=Homo sapiens OX=9606 GN=DOCK4 PE=1 SV=3 | 1 |
| Q8N1Y9 | Putative uncharacterized protein FLJ37218 OS=Homo sapiens OX=9606 PE=5 SV=2 | 1 |
| Q8N2F6 | Armadillo repeat-containing protein 10 OS=Homo sapiens OX=9606 GN=ARMC10 PE=1 SV=1 | 6 |
| Q8N2K0 | Lysophosphatidylserine lipase ABHD12 OS=Homo sapiens OX=9606 GN=ABHD12 PE=1 SV=2 | 14 |
| Q8N2U0 | Transmembrane protein 256 OS=Homo sapiens OX=9606 GN=TMEM256 PE=3 SV=1 | 1 |
| Q8N2Z9 | Centromere protein S OS=Homo sapiens OX=9606 GN=CENPS PE=1 SV=1 | 3 |
| Q8N307 | Mucin-20 OS=Homo sapiens OX=9606 GN=MUC20 PE=1 SV=3 | 3 |
| Q8N392 | Rho GTPase-activating protein 18 OS=Homo sapiens OX=9606 GN=ARHGAP18 PE=1 SV=3 | 6 |
| Q8N427 | Thioredoxin domain-containing protein 3 OS=Homo sapiens OX=9606 GN=NME8 PE=2 SV=2 | 1 |
| Q8N4C7 | Syntaxin-19 OS=Homo sapiens OX=9606 GN=STX19 PE=1 SV=1 | 1 |
| Q8N4C8 | Misshapen-like kinase 1 OS=Homo sapiens OX=9606 GN=MINK1 PE=1 SV=2 | 5 |
| Q8N4H5 | Mitochondrial import receptor subunit TOM5 homolog OS=Homo sapiens OX=9606 GN=TOMM5 PE=1 SV=1 | 1 |
| Q8N4Q1 | Mitochondrial intermembrane space import and assembly protein 40 OS=Homo sapiens OX=9606 GN=CHCHD4 PE=1 SV=1 | 4 |
| Q8N4T8 | 3-oxoacyl-[acyl-carrier-protein] reductase OS=Homo sapiens OX=9606 GN=CBR4 PE=1 SV=3 | 5 |
| Q8N4V1 | ER membrane protein complex subunit 5 OS=Homo sapiens OX=9606 GN=MMGT1 PE=1 SV=1 | 2 |
| Q8N535 | Putative uncharacterized protein encoded by LINC00471 OS=Homo sapiens OX=9606 GN=LINC00471 PE=5 SV=1 | 1 |
| Q8N568 | Serine/threonine-protein kinase DCLK2 OS=Homo sapiens OX=9606 GN=DCLK2 PE=1 SV=4 | 5 |
| Q8N573 | Oxidation resistance protein 1 OS=Homo sapiens OX=9606 GN=OXR1 PE=1 SV=2 | 8 |
| Q8N5G0 | Small integral membrane protein 20 OS=Homo sapiens OX=9606 GN=SMIM20 PE=1 SV=3 | 1 |
| Q8N5G2 | Macoilin OS=Homo sapiens OX=9606 GN=MACO1 PE=1 SV=1 | 3 |
| Q8N5I2 | Arrestin domain-containing protein 1 OS=Homo sapiens OX=9606 GN=ARRDC1 PE=1 SV=1 | 1 |
| Q8N5K1 | CDGSH iron-sulfur domain-containing protein 2 OS=Homo sapiens OX=9606 GN=CISD2 PE=1 SV=1 | 3 |
| Q8N5M4 | Tetratricopeptide repeat protein 9C OS=Homo sapiens OX=9606 GN=TTC9C PE=1 SV=1 | 5 |
| Q8N5N7 | 39S ribosomal protein L50, mitochondrial OS=Homo sapiens OX=9606 GN=MRPL50 PE=1 SV=2 | 2 |
| Q8N5X7 | Eukaryotic translation initiation factor 4E type 3 OS=Homo sapiens OX=9606 GN=EIF4E3 PE=1 SV=4 | 1 |
| Q8N5Y2 | Male-specific lethal 3 homolog OS=Homo sapiens OX=9606 GN=MSL3 PE=1 SV=1 | 1 |
| Q8N5Y8 | Protein mono-ADP-ribosyltransferase PARP16 OS=Homo sapiens OX=9606 GN=PARP16 PE=1 SV=2 | 8 |
| Q8N668 | COMM domain-containing protein 1 OS=Homo sapiens OX=9606 GN=COMMD1 PE=1 SV=1 | 8 |
| Q8N6M9 | AN1-type zinc finger protein 2A OS=Homo sapiens OX=9606 GN=ZFAND2A PE=1 SV=2 | 1 |
| Q8N6T3 | ADP-ribosylation factor GTPase-activating protein 1 OS=Homo sapiens OX=9606 GN=ARFGAP1 PE=1 SV=2 | 4 |
| Q8N755 | Solute carrier family 66 member 3 OS=Homo sapiens OX=9606 GN=SLC66A3 PE=2 SV=1 | 2 |
| Q8N766 | ER membrane protein complex subunit 1 OS=Homo sapiens OX=9606 GN=EMC1 PE=1 SV=1 | 6 |
| Q8N7G0 | POU domain, class 5, transcription factor 2 OS=Homo sapiens OX=9606 GN=POU5F2 PE=1 SV=1 | 1 |
| Q8N7R7 | Cyclin-Y-like protein 1 OS=Homo sapiens OX=9606 GN=CCNYL1 PE=1 SV=2 | 2 |
| Q8N806 | Putative E3 ubiquitin-protein ligase UBR7 OS=Homo sapiens OX=9606 GN=UBR7 PE=1 SV=2 | 5 |
| Q8N841 | Tubulin polyglutamylase TTLL6 OS=Homo sapiens OX=9606 GN=TTLL6 PE=1 SV=2 | 1 |
| Q8N8S7 | Protein enabled homolog OS=Homo sapiens OX=9606 GN=ENAH PE=1 SV=2 | 5 |
| Q8N8Y2 | V-type proton ATPase subunit d 2 OS=Homo sapiens OX=9606 GN=ATP6V0D2 PE=1 SV=1 | 1 |
| Q8N944 | APC membrane recruitment protein 3 OS=Homo sapiens OX=9606 GN=AMER3 PE=1 SV=2 | 1 |
| Q8N967 | Leucine-rich repeat and transmembrane domain-containing protein 2 OS=Homo sapiens OX=9606 GN=LRTM2 PE=2 SV=1 | 1 |
| Q8N983 | 39S ribosomal protein L43, mitochondrial OS=Homo sapiens OX=9606 GN=MRPL43 PE=1 SV=1 | 4 |
| Q8N999 | Uncharacterized protein C12orf29 OS=Homo sapiens OX=9606 GN=C12orf29 PE=1 SV=2 | 2 |
| Q8N9I0 | Synaptotagmin-2 OS=Homo sapiens OX=9606 GN=SYT2 PE=1 SV=2 | 14 |
| Q8N9V3 | WD repeat, SAM and U-box domain-containing protein 1 OS=Homo sapiens OX=9606 GN=WDSUB1 PE=1 SV=4 | 9 |
| Q8N9W4 | Golgin subfamily A member 6-like protein 2 OS=Homo sapiens OX=9606 GN=GOLGA6L2 PE=1 SV=3 | 1 |
| Q8NA58 | Poly(A)-specific ribonuclease PNLDC1 OS=Homo sapiens OX=9606 GN=PNLDC1 PE=1 SV=2 | 8 |
| Q8NA72 | Centrosomal protein POC5 OS=Homo sapiens OX=9606 GN=POC5 PE=1 SV=2 | 1 |
| Q8NAT1 | Protein O-linked-mannose beta-1,4-N-acetylglucosaminyltransferase 2 OS=Homo sapiens OX=9606 GN=POMGNT2 PE=1 SV=1 | 2 |
| Q8NB37 | Glutamine amidotransferase-like class 1 domain-containing protein 1 OS=Homo sapiens OX=9606 GN=GATD1 PE=1 SV=1 | 1 |
| Q8NB78 | Lysine-specific histone demethylase 2 OS=Homo sapiens OX=9606 GN=KDM1B PE=1 SV=3 | 27 |
| Q8NB90 | Ribosome biogenesis protein SPATA5 OS=Homo sapiens OX=9606 GN=SPATA5 PE=1 SV=3 | 3 |
| Q8NBF2 | NHL repeat-containing protein 2 OS=Homo sapiens OX=9606 GN=NHLRC2 PE=1 SV=1 | 4 |
| Q8NBF6 | Late secretory pathway protein AVL9 homolog OS=Homo sapiens OX=9606 GN=AVL9 PE=1 SV=1 | 4 |
| Q8NBJ4 | Golgi membrane protein 1 OS=Homo sapiens OX=9606 GN=GOLM1 PE=1 SV=1 | 6 |
| Q8NBJ5 | Procollagen galactosyltransferase 1 OS=Homo sapiens OX=9606 GN=COLGALT1 PE=1 SV=1 | 9 |
| Q8NBJ7 | Inactive C-alpha-formylglycine-generating enzyme 2 OS=Homo sapiens OX=9606 GN=SUMF2 PE=1 SV=2 | 6 |
| Q8NBJ9 | SID1 transmembrane family member 2 OS=Homo sapiens OX=9606 GN=SIDT2 PE=1 SV=2 | 1 |
| Q8NBM4 | Ubiquitin-associated domain-containing protein 2 OS=Homo sapiens OX=9606 GN=UBAC2 PE=1 SV=1 | 2 |
| Q8NBM8 | Prenylcysteine oxidase-like OS=Homo sapiens OX=9606 GN=PCYOX1L PE=1 SV=2 | 3 |
| Q8NBS9 | Thioredoxin domain-containing protein 5 OS=Homo sapiens OX=9606 GN=TXNDC5 PE=1 SV=2 | 21 |
| Q8NBU5 | Outer mitochondrial transmembrane helix translocase OS=Homo sapiens OX=9606 GN=ATAD1 PE=1 SV=1 | 5 |
| Q8NBX0 | Saccharopine dehydrogenase-like oxidoreductase OS=Homo sapiens OX=9606 GN=SCCPDH PE=1 SV=1 | 14 |
| Q8NBZ7 | UDP-glucuronic acid decarboxylase 1 OS=Homo sapiens OX=9606 GN=UXS1 PE=1 SV=1 | 4 |
| Q8NC51 | Plasminogen activator inhibitor 1 RNA-binding protein OS=Homo sapiens OX=9606 GN=SERBP1 PE=1 SV=2 | 3 |
| Q8NC56 | LEM domain-containing protein 2 OS=Homo sapiens OX=9606 GN=LEMD2 PE=1 SV=1 | 5 |
| Q8NC96 | Adaptin ear-binding coat-associated protein 1 OS=Homo sapiens OX=9606 GN=NECAP1 PE=1 SV=2 | 7 |
| Q8NCA5 | Protein FAM98A OS=Homo sapiens OX=9606 GN=FAM98A PE=1 SV=2 | 2 |
| Q8NCE2 | Myotubularin-related protein 14 OS=Homo sapiens OX=9606 GN=MTMR14 PE=1 SV=2 | 12 |
| Q8NCG7 | Diacylglycerol lipase-beta OS=Homo sapiens OX=9606 GN=DAGLB PE=1 SV=2 | 8 |
| Q8NCN5 | Pyruvate dehydrogenase phosphatase regulatory subunit, mitochondrial OS=Homo sapiens OX=9606 GN=PDPR PE=1 SV=2 | 17 |
| Q8NCW5 | NAD(P)H-hydrate epimerase OS=Homo sapiens OX=9606 GN=NAXE PE=1 SV=2 | 9 |
| Q8ND30 | Liprin-beta-2 OS=Homo sapiens OX=9606 GN=PPFIBP2 PE=1 SV=3 | 3 |
| Q8ND56 | Protein LSM14 homolog A OS=Homo sapiens OX=9606 GN=LSM14A PE=1 SV=3 | 2 |
| Q8NDA8 | Maestro heat-like repeat-containing protein family member 1 OS=Homo sapiens OX=9606 GN=MROH1 PE=2 SV=3 | 1 |
| Q8NDH3 | Probable aminopeptidase NPEPL1 OS=Homo sapiens OX=9606 GN=NPEPL1 PE=1 SV=3 | 1 |
| Q8NE01 | Metal transporter CNNM3 OS=Homo sapiens OX=9606 GN=CNNM3 PE=1 SV=1 | 2 |
| Q8NE63 | Homeodomain-interacting protein kinase 4 OS=Homo sapiens OX=9606 GN=HIPK4 PE=1 SV=1 | 1 |
| Q8NE71 | ATP-binding cassette sub-family F member 1 OS=Homo sapiens OX=9606 GN=ABCF1 PE=1 SV=2 | 7 |
| Q8NE86 | Calcium uniporter protein, mitochondrial OS=Homo sapiens OX=9606 GN=MCU PE=1 SV=1 | 6 |
| Q8NEH6 | Meiosis-specific nuclear structural protein 1 OS=Homo sapiens OX=9606 GN=MNS1 PE=1 SV=2 | 1 |
| Q8NEL9 | Phospholipase DDHD1 OS=Homo sapiens OX=9606 GN=DDHD1 PE=1 SV=2 | 9 |
| Q8NEM2 | SHC SH2 domain-binding protein 1 OS=Homo sapiens OX=9606 GN=SHCBP1 PE=1 SV=3 | 2 |
| Q8NEM8 | Cytosolic carboxypeptidase 3 OS=Homo sapiens OX=9606 GN=AGBL3 PE=2 SV=2 | 1 |
| Q8NEZ2 | Vacuolar protein sorting-associated protein 37A OS=Homo sapiens OX=9606 GN=VPS37A PE=1 SV=1 | 4 |
| Q8NEZ4 | Histone-lysine N-methyltransferase 2C OS=Homo sapiens OX=9606 GN=KMT2C PE=1 SV=3 | 17 |
| Q8NEZ5 | F-box only protein 22 OS=Homo sapiens OX=9606 GN=FBXO22 PE=1 SV=1 | 13 |
| Q8NF37 | Lysophosphatidylcholine acyltransferase 1 OS=Homo sapiens OX=9606 GN=LPCAT1 PE=1 SV=2 | 7 |
| Q8NF50 | Dedicator of cytokinesis protein 8 OS=Homo sapiens OX=9606 GN=DOCK8 PE=1 SV=3 | 13 |
| Q8NF91 | Nesprin-1 OS=Homo sapiens OX=9606 GN=SYNE1 PE=1 SV=4 | 2 |
| Q8NFF5 | FAD synthase OS=Homo sapiens OX=9606 GN=FLAD1 PE=1 SV=1 | 14 |
| Q8NFH3 | Nucleoporin Nup43 OS=Homo sapiens OX=9606 GN=NUP43 PE=1 SV=1 | 3 |
| Q8NFH8 | RalBP1-associated Eps domain-containing protein 2 OS=Homo sapiens OX=9606 GN=REPS2 PE=1 SV=2 | 3 |
| Q8NFU3 | Thiosulfate:glutathione sulfurtransferase OS=Homo sapiens OX=9606 GN=TSTD1 PE=1 SV=3 | 7 |
| Q8NFV4 | Protein ABHD11 OS=Homo sapiens OX=9606 GN=ABHD11 PE=1 SV=1 | 4 |
| Q8NFW9 | Rab effector MyRIP OS=Homo sapiens OX=9606 GN=MYRIP PE=1 SV=2 | 1 |
| Q8NFX7 | Syntaxin-binding protein 6 OS=Homo sapiens OX=9606 GN=STXBP6 PE=1 SV=2 | 4 |
| Q8NG11 | Tetraspanin-14 OS=Homo sapiens OX=9606 GN=TSPAN14 PE=1 SV=1 | 3 |
| Q8NHC8 | Olfactory receptor 2T6 OS=Homo sapiens OX=9606 GN=OR2T6 PE=3 SV=2 | 1 |
| Q8NHG7 | Small VCP/p97-interacting protein OS=Homo sapiens OX=9606 GN=SVIP PE=1 SV=1 | 2 |
| Q8NHH9 | Atlastin-2 OS=Homo sapiens OX=9606 GN=ATL2 PE=1 SV=2 | 8 |
| Q8NHP8 | Putative phospholipase B-like 2 OS=Homo sapiens OX=9606 GN=PLBD2 PE=1 SV=2 | 1 |
| Q8NHV4 | Protein NEDD1 OS=Homo sapiens OX=9606 GN=NEDD1 PE=1 SV=1 | 4 |
| Q8TAA5 | GrpE protein homolog 2, mitochondrial OS=Homo sapiens OX=9606 GN=GRPEL2 PE=1 SV=1 | 4 |
| Q8TAE8 | Growth arrest and DNA damage-inducible proteins-interacting protein 1 OS=Homo sapiens OX=9606 GN=GADD45GIP1 PE=1 SV=1 | 3 |
| Q8TAG9 | Exocyst complex component 6 OS=Homo sapiens OX=9606 GN=EXOC6 PE=1 SV=3 | 2 |
| Q8TAT6 | Nuclear protein localization protein 4 homolog OS=Homo sapiens OX=9606 GN=NPLOC4 PE=1 SV=3 | 4 |
| Q8TB03 | Uncharacterized protein CXorf38 OS=Homo sapiens OX=9606 GN=CXorf38 PE=1 SV=1 | 1 |
| Q8TB36 | Ganglioside-induced differentiation-associated protein 1 OS=Homo sapiens OX=9606 GN=GDAP1 PE=1 SV=3 | 17 |
| Q8TB37 | Iron-sulfur protein NUBPL OS=Homo sapiens OX=9606 GN=NUBPL PE=1 SV=3 | 6 |
| Q8TB61 | Adenosine 3'-phospho 5'-phosphosulfate transporter 1 OS=Homo sapiens OX=9606 GN=SLC35B2 PE=1 SV=1 | 1 |
| Q8TB96 | T-cell immunomodulatory protein OS=Homo sapiens OX=9606 GN=ITFG1 PE=1 SV=1 | 2 |
| Q8TBB5 | Kelch domain-containing protein 4 OS=Homo sapiens OX=9606 GN=KLHDC4 PE=1 SV=1 | 1 |
| Q8TBC4 | NEDD8-activating enzyme E1 catalytic subunit OS=Homo sapiens OX=9606 GN=UBA3 PE=1 SV=2 | 6 |
| Q8TBF2 | Prostamide/prostaglandin F synthase OS=Homo sapiens OX=9606 GN=PRXL2B PE=1 SV=1 | 1 |
| Q8TBX8 | Phosphatidylinositol 5-phosphate 4-kinase type-2 gamma OS=Homo sapiens OX=9606 GN=PIP4K2C PE=1 SV=3 | 17 |
| Q8TC05 | Nuclear protein MDM1 OS=Homo sapiens OX=9606 GN=MDM1 PE=1 SV=2 | 2 |
| Q8TC07 | TBC1 domain family member 15 OS=Homo sapiens OX=9606 GN=TBC1D15 PE=1 SV=2 | 15 |
| Q8TC12 | Retinol dehydrogenase 11 OS=Homo sapiens OX=9606 GN=RDH11 PE=1 SV=2 | 2 |
| Q8TCA0 | Leucine-rich repeat-containing protein 20 OS=Homo sapiens OX=9606 GN=LRRC20 PE=1 SV=1 | 3 |
| Q8TCC3 | 39S ribosomal protein L30, mitochondrial OS=Homo sapiens OX=9606 GN=MRPL30 PE=1 SV=1 | 2 |
| Q8TCD5 | 5'(3')-deoxyribonucleotidase, cytosolic type OS=Homo sapiens OX=9606 GN=NT5C PE=1 SV=2 | 3 |
| Q8TCE6 | DENN domain-containing protein 10 OS=Homo sapiens OX=9606 GN=DENND10 PE=1 SV=1 | 4 |
| Q8TCG1 | Protein CIP2A OS=Homo sapiens OX=9606 GN=CIP2A PE=1 SV=2 | 5 |
| Q8TCJ2 | Dolichyl-diphosphooligosaccharide--protein glycosyltransferase subunit STT3B OS=Homo sapiens OX=9606 GN=STT3B PE=1 SV=1 | 3 |
| Q8TCS8 | Polyribonucleotide nucleotidyltransferase 1, mitochondrial OS=Homo sapiens OX=9606 GN=PNPT1 PE=1 SV=2 | 7 |
| Q8TCT8 | Signal peptide peptidase-like 2A OS=Homo sapiens OX=9606 GN=SPPL2A PE=1 SV=2 | 1 |
| Q8TCX5 | Rhophilin-1 OS=Homo sapiens OX=9606 GN=RHPN1 PE=1 SV=2 | 1 |
| Q8TD08 | Mitogen-activated protein kinase 15 OS=Homo sapiens OX=9606 GN=MAPK15 PE=1 SV=1 | 1 |
| Q8TD30 | Alanine aminotransferase 2 OS=Homo sapiens OX=9606 GN=GPT2 PE=1 SV=1 | 21 |
| Q8TDQ7 | Glucosamine-6-phosphate isomerase 2 OS=Homo sapiens OX=9606 GN=GNPDA2 PE=1 SV=1 | 19 |
| Q8TDZ2 | [F-actin]-monooxygenase MICAL1 OS=Homo sapiens OX=9606 GN=MICAL1 PE=1 SV=2 | 6 |
| Q8TE56 | A disintegrin and metalloproteinase with thrombospondin motifs 17 OS=Homo sapiens OX=9606 GN=ADAMTS17 PE=2 SV=2 | 3 |
| Q8TE85 | Grainyhead-like protein 3 homolog OS=Homo sapiens OX=9606 GN=GRHL3 PE=1 SV=3 | 1 |
| Q8TEM1 | Nuclear pore membrane glycoprotein 210 OS=Homo sapiens OX=9606 GN=NUP210 PE=1 SV=3 | 4 |
| Q8TEP8 | Centrosomal protein of 192 kDa OS=Homo sapiens OX=9606 GN=CEP192 PE=1 SV=3 | 1 |
| Q8TEQ6 | Gem-associated protein 5 OS=Homo sapiens OX=9606 GN=GEMIN5 PE=1 SV=3 | 15 |
| Q8TET4 | Neutral alpha-glucosidase C OS=Homo sapiens OX=9606 GN=GANC PE=2 SV=3 | 1 |
| Q8TF05 | Serine/threonine-protein phosphatase 4 regulatory subunit 1 OS=Homo sapiens OX=9606 GN=PPP4R1 PE=1 SV=1 | 4 |
| Q8TF65 | PDZ domain-containing protein GIPC2 OS=Homo sapiens OX=9606 GN=GIPC2 PE=1 SV=1 | 13 |
| Q8TF74 | WAS/WASL-interacting protein family member 2 OS=Homo sapiens OX=9606 GN=WIPF2 PE=1 SV=1 | 8 |
| Q8WTV0 | Scavenger receptor class B member 1 OS=Homo sapiens OX=9606 GN=SCARB1 PE=1 SV=1 | 5 |
| Q8WU39 | Marginal zone B- and B1-cell-specific protein OS=Homo sapiens OX=9606 GN=MZB1 PE=1 SV=1 | 1 |
| Q8WU79 | Stromal membrane-associated protein 2 OS=Homo sapiens OX=9606 GN=SMAP2 PE=1 SV=1 | 1 |
| Q8WU90 | Zinc finger CCCH domain-containing protein 15 OS=Homo sapiens OX=9606 GN=ZC3H15 PE=1 SV=1 | 7 |
| Q8WUD1 | Ras-related protein Rab-2B OS=Homo sapiens OX=9606 GN=RAB2B PE=1 SV=1 | 5 |
| Q8WUK0 | Phosphatidylglycerophosphatase and protein-tyrosine phosphatase 1 OS=Homo sapiens OX=9606 GN=PTPMT1 PE=1 SV=1 | 3 |
| Q8WUM0 | Nuclear pore complex protein Nup133 OS=Homo sapiens OX=9606 GN=NUP133 PE=1 SV=2 | 30 |
| Q8WUM4 | Programmed cell death 6-interacting protein OS=Homo sapiens OX=9606 GN=PDCD6IP PE=1 SV=1 | 39 |
| Q8WUT4 | Leucine-rich repeat neuronal protein 4 OS=Homo sapiens OX=9606 GN=LRRN4 PE=1 SV=3 | 2 |
| Q8WUX9 | Charged multivesicular body protein 7 OS=Homo sapiens OX=9606 GN=CHMP7 PE=1 SV=1 | 1 |
| Q8WUY1 | Protein THEM6 OS=Homo sapiens OX=9606 GN=THEM6 PE=1 SV=2 | 3 |
| Q8WUY3 | Protein prune homolog 2 OS=Homo sapiens OX=9606 GN=PRUNE2 PE=1 SV=3 | 1 |
| Q8WV22 | Non-structural maintenance of chromosomes element 1 homolog OS=Homo sapiens OX=9606 GN=NSMCE1 PE=1 SV=5 | 1 |
| Q8WV74 | Mitochondrial coenzyme A diphosphatase NUDT8 OS=Homo sapiens OX=9606 GN=NUDT8 PE=1 SV=2 | 1 |
| Q8WVC6 | Dephospho-CoA kinase domain-containing protein OS=Homo sapiens OX=9606 GN=DCAKD PE=1 SV=1 | 2 |
| Q8WVF1 | Protein OSCP1 OS=Homo sapiens OX=9606 GN=OSCP1 PE=1 SV=4 | 9 |
| Q8WVJ2 | NudC domain-containing protein 2 OS=Homo sapiens OX=9606 GN=NUDCD2 PE=1 SV=1 | 6 |
| Q8WVM0 | Dimethyladenosine transferase 1, mitochondrial OS=Homo sapiens OX=9606 GN=TFB1M PE=1 SV=1 | 1 |
| Q8WVM8 | Sec1 family domain-containing protein 1 OS=Homo sapiens OX=9606 GN=SCFD1 PE=1 SV=4 | 27 |
| Q8WVN8 | Ubiquitin-conjugating enzyme E2 Q2 OS=Homo sapiens OX=9606 GN=UBE2Q2 PE=1 SV=1 | 8 |
| Q8WVS4 | Cytoplasmic dynein 2 intermediate chain 1 OS=Homo sapiens OX=9606 GN=DYNC2I1 PE=1 SV=3 | 1 |
| Q8WVT3 | Trafficking protein particle complex subunit 12 OS=Homo sapiens OX=9606 GN=TRAPPC12 PE=1 SV=3 | 5 |
| Q8WVY7 | Ubiquitin-like domain-containing CTD phosphatase 1 OS=Homo sapiens OX=9606 GN=UBLCP1 PE=1 SV=2 | 3 |
| Q8WW22 | DnaJ homolog subfamily A member 4 OS=Homo sapiens OX=9606 GN=DNAJA4 PE=1 SV=1 | 13 |
| Q8WW33 | Gametocyte-specific factor 1 OS=Homo sapiens OX=9606 GN=GTSF1 PE=1 SV=2 | 8 |
| Q8WWC4 | m-AAA protease-interacting protein 1, mitochondrial OS=Homo sapiens OX=9606 GN=MAIP1 PE=1 SV=1 | 2 |
| Q8WWI1 | LIM domain only protein 7 OS=Homo sapiens OX=9606 GN=LMO7 PE=1 SV=3 | 3 |
| Q8WWM7 | Ataxin-2-like protein OS=Homo sapiens OX=9606 GN=ATXN2L PE=1 SV=2 | 5 |
| Q8WWM9 | Cytoglobin OS=Homo sapiens OX=9606 GN=CYGB PE=1 SV=1 | 1 |
| Q8WWV3 | Reticulon-4-interacting protein 1, mitochondrial OS=Homo sapiens OX=9606 GN=RTN4IP1 PE=1 SV=2 | 10 |
| Q8WWY7 | WAP four-disulfide core domain protein 12 OS=Homo sapiens OX=9606 GN=WFDC12 PE=1 SV=1 | 3 |
| Q8WWZ7 | Cholesterol transporter ABCA5 OS=Homo sapiens OX=9606 GN=ABCA5 PE=1 SV=2 | 1 |
| Q8WXB1 | Protein N-lysine methyltransferase METTL21A OS=Homo sapiens OX=9606 GN=METTL21A PE=1 SV=2 | 3 |
| Q8WXG6 | MAP kinase-activating death domain protein OS=Homo sapiens OX=9606 GN=MADD PE=1 SV=2 | 3 |
| Q8WXG8 | Protein S100-Z OS=Homo sapiens OX=9606 GN=S100Z PE=1 SV=4 | 1 |
| Q8WXX5 | DnaJ homolog subfamily C member 9 OS=Homo sapiens OX=9606 GN=DNAJC9 PE=1 SV=1 | 17 |
| Q8WY54 | Protein phosphatase 1E OS=Homo sapiens OX=9606 GN=PPM1E PE=1 SV=3 | 6 |
| Q8WY91 | Peroxynitrite isomerase THAP4 OS=Homo sapiens OX=9606 GN=THAP4 PE=1 SV=2 | 2 |
| Q8WZA0 | Protein LZIC OS=Homo sapiens OX=9606 GN=LZIC PE=1 SV=1 | 10 |
| Q8WZA1 | Protein O-linked-mannose beta-1,2-N-acetylglucosaminyltransferase 1 OS=Homo sapiens OX=9606 GN=POMGNT1 PE=1 SV=2 | 4 |
| Q92485 | Acid sphingomyelinase-like phosphodiesterase 3b OS=Homo sapiens OX=9606 GN=SMPDL3B PE=1 SV=2 | 7 |
| Q92499 | ATP-dependent RNA helicase DDX1 OS=Homo sapiens OX=9606 GN=DDX1 PE=1 SV=2 | 26 |
| Q92506 | (3R)-3-hydroxyacyl-CoA dehydrogenase OS=Homo sapiens OX=9606 GN=HSD17B8 PE=1 SV=2 | 4 |
| Q92520 | Protein FAM3C OS=Homo sapiens OX=9606 GN=FAM3C PE=1 SV=1 | 4 |
| Q92526 | T-complex protein 1 subunit zeta-2 OS=Homo sapiens OX=9606 GN=CCT6B PE=1 SV=5 | 4 |
| Q92529 | SHC-transforming protein 3 OS=Homo sapiens OX=9606 GN=SHC3 PE=1 SV=1 | 1 |
| Q92530 | Proteasome inhibitor PI31 subunit OS=Homo sapiens OX=9606 GN=PSMF1 PE=1 SV=2 | 3 |
| Q92538 | Golgi-specific brefeldin A-resistance guanine nucleotide exchange factor 1 OS=Homo sapiens OX=9606 GN=GBF1 PE=1 SV=2 | 16 |
| Q92539 | Phosphatidate phosphatase LPIN2 OS=Homo sapiens OX=9606 GN=LPIN2 PE=1 SV=1 | 5 |
| Q92542 | Nicastrin OS=Homo sapiens OX=9606 GN=NCSTN PE=1 SV=2 | 5 |
| Q92544 | Transmembrane 9 superfamily member 4 OS=Homo sapiens OX=9606 GN=TM9SF4 PE=1 SV=2 | 3 |
| Q92552 | 28S ribosomal protein S27, mitochondrial OS=Homo sapiens OX=9606 GN=MRPS27 PE=1 SV=3 | 2 |
| Q92558 | Actin-binding protein WASF1 OS=Homo sapiens OX=9606 GN=WASF1 PE=1 SV=1 | 1 |
| Q92569 | Phosphatidylinositol 3-kinase regulatory subunit gamma OS=Homo sapiens OX=9606 GN=PIK3R3 PE=1 SV=2 | 1 |
| Q92572 | AP-3 complex subunit sigma-1 OS=Homo sapiens OX=9606 GN=AP3S1 PE=1 SV=1 | 2 |
| Q92575 | UBX domain-containing protein 4 OS=Homo sapiens OX=9606 GN=UBXN4 PE=1 SV=2 | 1 |
| Q92581 | Sodium/hydrogen exchanger 6 OS=Homo sapiens OX=9606 GN=SLC9A6 PE=1 SV=3 | 5 |
| Q92597 | Protein NDRG1 OS=Homo sapiens OX=9606 GN=NDRG1 PE=1 SV=1 | 6 |
| Q92598 | Heat shock protein 105 kDa OS=Homo sapiens OX=9606 GN=HSPH1 PE=1 SV=1 | 34 |
| Q92599 | Septin-8 OS=Homo sapiens OX=9606 GN=SEPTIN8 PE=1 SV=4 | 5 |
| Q92614 | Unconventional myosin-XVIIIa OS=Homo sapiens OX=9606 GN=MYO18A PE=1 SV=3 | 9 |
| Q92616 | eIF-2-alpha kinase activator GCN1 OS=Homo sapiens OX=9606 GN=GCN1 PE=1 SV=7 | 17 |
| Q92621 | Nuclear pore complex protein Nup205 OS=Homo sapiens OX=9606 GN=NUP205 PE=1 SV=3 | 5 |
| Q92643 | GPI-anchor transamidase OS=Homo sapiens OX=9606 GN=PIGK PE=1 SV=2 | 4 |
| Q92665 | 28S ribosomal protein S31, mitochondrial OS=Homo sapiens OX=9606 GN=MRPS31 PE=1 SV=3 | 3 |
| Q92667 | A-kinase anchor protein 1, mitochondrial OS=Homo sapiens OX=9606 GN=AKAP1 PE=1 SV=1 | 20 |
| Q92688 | Acidic leucine-rich nuclear phosphoprotein 32 family member B OS=Homo sapiens OX=9606 GN=ANP32B PE=1 SV=1 | 4 |
| Q92696 | Geranylgeranyl transferase type-2 subunit alpha OS=Homo sapiens OX=9606 GN=RABGGTA PE=1 SV=2 | 10 |
| Q92734 | Protein TFG OS=Homo sapiens OX=9606 GN=TFG PE=1 SV=2 | 8 |
| Q92743 | Serine protease HTRA1 OS=Homo sapiens OX=9606 GN=HTRA1 PE=1 SV=1 | 7 |
| Q92747 | Actin-related protein 2/3 complex subunit 1A OS=Homo sapiens OX=9606 GN=ARPC1A PE=1 SV=2 | 1 |
| Q92783 | Signal transducing adapter molecule 1 OS=Homo sapiens OX=9606 GN=STAM PE=1 SV=3 | 3 |
| Q92796 | Disks large homolog 3 OS=Homo sapiens OX=9606 GN=DLG3 PE=1 SV=2 | 4 |
| Q92804 | TATA-binding protein-associated factor 2N OS=Homo sapiens OX=9606 GN=TAF15 PE=1 SV=1 | 4 |
| Q92817 | Envoplakin OS=Homo sapiens OX=9606 GN=EVPL PE=1 SV=3 | 2 |
| Q92832 | Protein kinase C-binding protein NELL1 OS=Homo sapiens OX=9606 GN=NELL1 PE=1 SV=4 | 5 |
| Q92839 | Hyaluronan synthase 1 OS=Homo sapiens OX=9606 GN=HAS1 PE=1 SV=2 | 1 |
| Q92841 | Probable ATP-dependent RNA helicase DDX17 OS=Homo sapiens OX=9606 GN=DDX17 PE=1 SV=2 | 4 |
| Q92874 | Deoxyribonuclease-1-like 2 OS=Homo sapiens OX=9606 GN=DNASE1L2 PE=1 SV=1 | 1 |
| Q92879 | CUGBP Elav-like family member 1 OS=Homo sapiens OX=9606 GN=CELF1 PE=1 SV=2 | 4 |
| Q92882 | Osteoclast-stimulating factor 1 OS=Homo sapiens OX=9606 GN=OSTF1 PE=1 SV=2 | 11 |
| Q92890 | Ubiquitin recognition factor in ER-associated degradation protein 1 OS=Homo sapiens OX=9606 GN=UFD1 PE=1 SV=3 | 3 |
| Q92896 | Golgi apparatus protein 1 OS=Homo sapiens OX=9606 GN=GLG1 PE=1 SV=2 | 21 |
| Q92900 | Regulator of nonsense transcripts 1 OS=Homo sapiens OX=9606 GN=UPF1 PE=1 SV=2 | 10 |
| Q92903 | Phosphatidate cytidylyltransferase 1 OS=Homo sapiens OX=9606 GN=CDS1 PE=1 SV=2 | 1 |
| Q92905 | COP9 signalosome complex subunit 5 OS=Homo sapiens OX=9606 GN=COPS5 PE=1 SV=4 | 15 |
| Q92930 | Ras-related protein Rab-8B OS=Homo sapiens OX=9606 GN=RAB8B PE=1 SV=2 | 3 |
| Q92945 | Far upstream element-binding protein 2 OS=Homo sapiens OX=9606 GN=KHSRP PE=1 SV=4 | 6 |
| Q92947 | Glutaryl-CoA dehydrogenase, mitochondrial OS=Homo sapiens OX=9606 GN=GCDH PE=1 SV=1 | 9 |
| Q92973 | Transportin-1 OS=Homo sapiens OX=9606 GN=TNPO1 PE=1 SV=2 | 8 |
| Q92990 | Glomulin OS=Homo sapiens OX=9606 GN=GLMN PE=1 SV=2 | 6 |
| Q93008 | Probable ubiquitin carboxyl-terminal hydrolase FAF-X OS=Homo sapiens OX=9606 GN=USP9X PE=1 SV=4 | 56 |
| Q93009 | Ubiquitin carboxyl-terminal hydrolase 7 OS=Homo sapiens OX=9606 GN=USP7 PE=1 SV=2 | 10 |
| Q93034 | Cullin-5 OS=Homo sapiens OX=9606 GN=CUL5 PE=1 SV=4 | 33 |
| Q93045 | Stathmin-2 OS=Homo sapiens OX=9606 GN=STMN2 PE=1 SV=3 | 1 |
| Q93050 | V-type proton ATPase 116 kDa subunit a 1 OS=Homo sapiens OX=9606 GN=ATP6V0A1 PE=1 SV=3 | 18 |
| Q969H8 | Myeloid-derived growth factor OS=Homo sapiens OX=9606 GN=MYDGF PE=1 SV=1 | 7 |
| Q969J3 | BLOC-1-related complex subunit 5 OS=Homo sapiens OX=9606 GN=BORCS5 PE=1 SV=1 | 1 |
| Q969L2 | Protein MAL2 OS=Homo sapiens OX=9606 GN=MAL2 PE=1 SV=1 | 1 |
| Q969M3 | Protein YIPF5 OS=Homo sapiens OX=9606 GN=YIPF5 PE=1 SV=1 | 2 |
| Q969M7 | NEDD8-conjugating enzyme UBE2F OS=Homo sapiens OX=9606 GN=UBE2F PE=1 SV=1 | 6 |
| Q969N2 | GPI transamidase component PIG-T OS=Homo sapiens OX=9606 GN=PIGT PE=1 SV=1 | 1 |
| Q969P0 | Immunoglobulin superfamily member 8 OS=Homo sapiens OX=9606 GN=IGSF8 PE=1 SV=1 | 2 |
| Q969S9 | Ribosome-releasing factor 2, mitochondrial OS=Homo sapiens OX=9606 GN=GFM2 PE=1 SV=1 | 6 |
| Q969T9 | WW domain-binding protein 2 OS=Homo sapiens OX=9606 GN=WBP2 PE=1 SV=1 | 4 |
| Q969U7 | Proteasome assembly chaperone 2 OS=Homo sapiens OX=9606 GN=PSMG2 PE=1 SV=1 | 3 |
| Q969V3 | BOS complex subunit NCLN OS=Homo sapiens OX=9606 GN=NCLN PE=1 SV=2 | 15 |
| Q969X1 | Protein lifeguard 3 OS=Homo sapiens OX=9606 GN=TMBIM1 PE=1 SV=2 | 1 |
| Q969Y2 | tRNA modification GTPase GTPBP3, mitochondrial OS=Homo sapiens OX=9606 GN=GTPBP3 PE=1 SV=2 | 2 |
| Q969Z0 | FAST kinase domain-containing protein 4 OS=Homo sapiens OX=9606 GN=TBRG4 PE=1 SV=1 | 6 |
| Q96A35 | 39S ribosomal protein L24, mitochondrial OS=Homo sapiens OX=9606 GN=MRPL24 PE=1 SV=1 | 4 |
| Q96A49 | Synapse-associated protein 1 OS=Homo sapiens OX=9606 GN=SYAP1 PE=1 SV=1 | 13 |
| Q96A57 | Transmembrane protein 230 OS=Homo sapiens OX=9606 GN=TMEM230 PE=1 SV=1 | 2 |
| Q96A65 | Exocyst complex component 4 OS=Homo sapiens OX=9606 GN=EXOC4 PE=1 SV=1 | 8 |
| Q96AA3 | Protein RFT1 homolog OS=Homo sapiens OX=9606 GN=RFT1 PE=1 SV=1 | 1 |
| Q96AB3 | Isochorismatase domain-containing protein 2 OS=Homo sapiens OX=9606 GN=ISOC2 PE=1 SV=1 | 6 |
| Q96AB6 | Protein N-terminal asparagine amidohydrolase OS=Homo sapiens OX=9606 GN=NTAN1 PE=1 SV=3 | 16 |
| Q96AE4 | Far upstream element-binding protein 1 OS=Homo sapiens OX=9606 GN=FUBP1 PE=1 SV=3 | 14 |
| Q96AE7 | Tetratricopeptide repeat protein 17 OS=Homo sapiens OX=9606 GN=TTC17 PE=1 SV=1 | 13 |
| Q96AG3 | Mitochondrial outer membrane protein SLC25A46 OS=Homo sapiens OX=9606 GN=SLC25A46 PE=1 SV=1 | 2 |
| Q96AG4 | Leucine-rich repeat-containing protein 59 OS=Homo sapiens OX=9606 GN=LRRC59 PE=1 SV=1 | 10 |
| Q96AJ9 | Vesicle transport through interaction with t-SNAREs homolog 1A OS=Homo sapiens OX=9606 GN=VTI1A PE=1 SV=2 | 2 |
| Q96AQ6 | Pre-B-cell leukemia transcription factor-interacting protein 1 OS=Homo sapiens OX=9606 GN=PBXIP1 PE=1 SV=1 | 13 |
| Q96AQ8 | Mitochondrial calcium uniporter regulator 1 OS=Homo sapiens OX=9606 GN=MCUR1 PE=1 SV=1 | 8 |
| Q96AT9 | Ribulose-phosphate 3-epimerase OS=Homo sapiens OX=9606 GN=RPE PE=1 SV=1 | 1 |
| Q96AX1 | Vacuolar protein sorting-associated protein 33A OS=Homo sapiens OX=9606 GN=VPS33A PE=1 SV=1 | 3 |
| Q96B36 | Proline-rich AKT1 substrate 1 OS=Homo sapiens OX=9606 GN=AKT1S1 PE=1 SV=1 | 1 |
| Q96B49 | Mitochondrial import receptor subunit TOM6 homolog OS=Homo sapiens OX=9606 GN=TOMM6 PE=1 SV=1 | 1 |
| Q96B97 | SH3 domain-containing kinase-binding protein 1 OS=Homo sapiens OX=9606 GN=SH3KBP1 PE=1 SV=2 | 19 |
| Q96BJ3 | Axin interactor, dorsalization-associated protein OS=Homo sapiens OX=9606 GN=AIDA PE=1 SV=1 | 4 |
| Q96BN8 | Ubiquitin thioesterase otulin OS=Homo sapiens OX=9606 GN=OTULIN PE=1 SV=3 | 10 |
| Q96BS2 | Calcineurin B homologous protein 3 OS=Homo sapiens OX=9606 GN=TESC PE=1 SV=3 | 3 |
| Q96BW9 | Phosphatidate cytidylyltransferase, mitochondrial OS=Homo sapiens OX=9606 GN=TAMM41 PE=1 SV=2 | 4 |
| Q96BY7 | Autophagy-related protein 2 homolog B OS=Homo sapiens OX=9606 GN=ATG2B PE=1 SV=5 | 2 |
| Q96C10 | ATP-dependent RNA helicase DHX58 OS=Homo sapiens OX=9606 GN=DHX58 PE=1 SV=1 | 12 |
| Q96C11 | FGGY carbohydrate kinase domain-containing protein OS=Homo sapiens OX=9606 GN=FGGY PE=1 SV=2 | 3 |
| Q96C19 | EF-hand domain-containing protein D2 OS=Homo sapiens OX=9606 GN=EFHD2 PE=1 SV=1 | 4 |
| Q96C36 | Pyrroline-5-carboxylate reductase 2 OS=Homo sapiens OX=9606 GN=PYCR2 PE=1 SV=1 | 9 |
| Q96CD2 | Phosphopantothenoylcysteine decarboxylase OS=Homo sapiens OX=9606 GN=PPCDC PE=1 SV=2 | 2 |
| Q96CM3 | Pseudouridylate synthase RPUSD4, mitochondrial OS=Homo sapiens OX=9606 GN=RPUSD4 PE=1 SV=1 | 1 |
| Q96CN7 | Isochorismatase domain-containing protein 1 OS=Homo sapiens OX=9606 GN=ISOC1 PE=1 SV=3 | 8 |
| Q96CP2 | FLYWCH family member 2 OS=Homo sapiens OX=9606 GN=FLYWCH2 PE=1 SV=1 | 3 |
| Q96CS3 | FAS-associated factor 2 OS=Homo sapiens OX=9606 GN=FAF2 PE=1 SV=2 | 5 |
| Q96CT7 | Coiled-coil domain-containing protein 124 OS=Homo sapiens OX=9606 GN=CCDC124 PE=1 SV=1 | 11 |
| Q96CU9 | FAD-dependent oxidoreductase domain-containing protein 1 OS=Homo sapiens OX=9606 GN=FOXRED1 PE=1 SV=2 | 2 |
| Q96CW1 | AP-2 complex subunit mu OS=Homo sapiens OX=9606 GN=AP2M1 PE=1 SV=2 | 10 |
| Q96CW5 | Gamma-tubulin complex component 3 OS=Homo sapiens OX=9606 GN=TUBGCP3 PE=1 SV=2 | 8 |
| Q96D96 | Voltage-gated hydrogen channel 1 OS=Homo sapiens OX=9606 GN=HVCN1 PE=1 SV=1 | 6 |
| Q96DA6 | Mitochondrial import inner membrane translocase subunit TIM14 OS=Homo sapiens OX=9606 GN=DNAJC19 PE=1 SV=3 | 5 |
| Q96DB5 | Regulator of microtubule dynamics protein 1 OS=Homo sapiens OX=9606 GN=RMDN1 PE=1 SV=1 | 8 |
| Q96DC8 | Enoyl-CoA hydratase domain-containing protein 3, mitochondrial OS=Homo sapiens OX=9606 GN=ECHDC3 PE=1 SV=2 | 4 |
| Q96DN5 | TBC1 domain family member 31 OS=Homo sapiens OX=9606 GN=TBC1D31 PE=1 SV=2 | 1 |
| Q96DP5 | Methionyl-tRNA formyltransferase, mitochondrial OS=Homo sapiens OX=9606 GN=MTFMT PE=1 SV=2 | 2 |
| Q96DT6 | Cysteine protease ATG4C OS=Homo sapiens OX=9606 GN=ATG4C PE=1 SV=1 | 6 |
| Q96DV4 | 39S ribosomal protein L38, mitochondrial OS=Homo sapiens OX=9606 GN=MRPL38 PE=1 SV=2 | 4 |
| Q96DZ1 | Endoplasmic reticulum lectin 1 OS=Homo sapiens OX=9606 GN=ERLEC1 PE=1 SV=1 | 22 |
| Q96E11 | Ribosome-recycling factor, mitochondrial OS=Homo sapiens OX=9606 GN=MRRF PE=1 SV=1 | 1 |
| Q96E29 | Transcription termination factor 3, mitochondrial OS=Homo sapiens OX=9606 GN=MTERF3 PE=1 SV=2 | 7 |
| Q96EA4 | Protein Spindly OS=Homo sapiens OX=9606 GN=SPDL1 PE=1 SV=2 | 23 |
| Q96EC8 | Protein YIPF6 OS=Homo sapiens OX=9606 GN=YIPF6 PE=1 SV=2 | 1 |
| Q96EE3 | Nucleoporin SEH1 OS=Homo sapiens OX=9606 GN=SEH1L PE=1 SV=3 | 3 |
| Q96EK5 | KIF-binding protein OS=Homo sapiens OX=9606 GN=KIFBP PE=1 SV=1 | 14 |
| Q96EK6 | Glucosamine 6-phosphate N-acetyltransferase OS=Homo sapiens OX=9606 GN=GNPNAT1 PE=1 SV=1 | 3 |
| Q96EK7 | Constitutive coactivator of peroxisome proliferator-activated receptor gamma OS=Homo sapiens OX=9606 GN=FAM120B PE=1 SV=1 | 2 |
| Q96EP5 | DAZ-associated protein 1 OS=Homo sapiens OX=9606 GN=DAZAP1 PE=1 SV=1 | 2 |
| Q96EY1 | DnaJ homolog subfamily A member 3, mitochondrial OS=Homo sapiens OX=9606 GN=DNAJA3 PE=1 SV=2 | 13 |
| Q96EY5 | Multivesicular body subunit 12A OS=Homo sapiens OX=9606 GN=MVB12A PE=1 SV=1 | 6 |
| Q96EY7 | Pentatricopeptide repeat domain-containing protein 3, mitochondrial OS=Homo sapiens OX=9606 GN=PTCD3 PE=1 SV=3 | 3 |
| Q96F63 | Coiled-coil domain-containing protein 97 OS=Homo sapiens OX=9606 GN=CCDC97 PE=1 SV=1 | 2 |
| Q96F85 | CB1 cannabinoid receptor-interacting protein 1 OS=Homo sapiens OX=9606 GN=CNRIP1 PE=1 SV=1 | 4 |
| Q96FF7 | Uncharacterized protein MISP3 OS=Homo sapiens OX=9606 GN=MISP3 PE=2 SV=4 | 2 |
| Q96FJ2 | Dynein light chain 2, cytoplasmic OS=Homo sapiens OX=9606 GN=DYNLL2 PE=1 SV=1 | 3 |
| Q96FN4 | Copine-2 OS=Homo sapiens OX=9606 GN=CPNE2 PE=1 SV=3 | 1 |
| Q96FN9 | D-aminoacyl-tRNA deacylase 2 OS=Homo sapiens OX=9606 GN=DTD2 PE=1 SV=1 | 1 |
| Q96FQ6 | Protein S100-A16 OS=Homo sapiens OX=9606 GN=S100A16 PE=1 SV=1 | 3 |
| Q96FV2 | Secernin-2 OS=Homo sapiens OX=9606 GN=SCRN2 PE=1 SV=3 | 7 |
| Q96FW1 | Ubiquitin thioesterase OTUB1 OS=Homo sapiens OX=9606 GN=OTUB1 PE=1 SV=2 | 11 |
| Q96FX7 | tRNA (adenine(58)-N(1))-methyltransferase catalytic subunit TRMT61A OS=Homo sapiens OX=9606 GN=TRMT61A PE=1 SV=1 | 1 |
| Q96FZ2 | Abasic site processing protein HMCES OS=Homo sapiens OX=9606 GN=HMCES PE=1 SV=1 | 7 |
| Q96FZ7 | Charged multivesicular body protein 6 OS=Homo sapiens OX=9606 GN=CHMP6 PE=1 SV=3 | 3 |
| Q96G03 | Phosphopentomutase OS=Homo sapiens OX=9606 GN=PGM2 PE=1 SV=4 | 13 |
| Q96G42 | Kelch domain-containing protein 7B OS=Homo sapiens OX=9606 GN=KLHDC7B PE=1 SV=2 | 1 |
| Q96G97 | Seipin OS=Homo sapiens OX=9606 GN=BSCL2 PE=1 SV=3 | 4 |
| Q96GA7 | Serine dehydratase-like OS=Homo sapiens OX=9606 GN=SDSL PE=1 SV=1 | 11 |
| Q96GC5 | 39S ribosomal protein L48, mitochondrial OS=Homo sapiens OX=9606 GN=MRPL48 PE=1 SV=2 | 1 |
| Q96GD0 | Chronophin OS=Homo sapiens OX=9606 GN=PDXP PE=1 SV=2 | 8 |
| Q96GG9 | DCN1-like protein 1 OS=Homo sapiens OX=9606 GN=DCUN1D1 PE=1 SV=1 | 7 |
| Q96GK7 | Fumarylacetoacetate hydrolase domain-containing protein 2A OS=Homo sapiens OX=9606 GN=FAHD2A PE=1 SV=1 | 13 |
| Q96GQ5 | RUS family member 1 OS=Homo sapiens OX=9606 GN=RUSF1 PE=1 SV=2 | 4 |
| Q96H20 | Vacuolar-sorting protein SNF8 OS=Homo sapiens OX=9606 GN=SNF8 PE=1 SV=1 | 3 |
| Q96HD1 | Protein disulfide isomerase CRELD1 OS=Homo sapiens OX=9606 GN=CRELD1 PE=1 SV=3 | 3 |
| Q96HN2 | Adenosylhomocysteinase 3 OS=Homo sapiens OX=9606 GN=AHCYL2 PE=1 SV=1 | 3 |
| Q96HP4 | Oxidoreductase NAD-binding domain-containing protein 1 OS=Homo sapiens OX=9606 GN=OXNAD1 PE=1 SV=1 | 7 |
| Q96HY6 | DDRGK domain-containing protein 1 OS=Homo sapiens OX=9606 GN=DDRGK1 PE=1 SV=2 | 1 |
| Q96I59 | Probable asparagine--tRNA ligase, mitochondrial OS=Homo sapiens OX=9606 GN=NARS2 PE=1 SV=3 | 2 |
| Q96I99 | Succinate--CoA ligase [GDP-forming] subunit beta, mitochondrial OS=Homo sapiens OX=9606 GN=SUCLG2 PE=1 SV=2 | 5 |
| Q96IJ6 | Mannose-1-phosphate guanyltransferase alpha OS=Homo sapiens OX=9606 GN=GMPPA PE=1 SV=1 | 9 |
| Q96IU4 | Putative protein-lysine deacylase ABHD14B OS=Homo sapiens OX=9606 GN=ABHD14B PE=1 SV=1 | 7 |
| Q96IV0 | Peptide-N(4)-(N-acetyl-beta-glucosaminyl)asparagine amidase OS=Homo sapiens OX=9606 GN=NGLY1 PE=1 SV=1 | 3 |
| Q96IX5 | ATP synthase membrane subunit K, mitochondrial OS=Homo sapiens OX=9606 GN=ATP5MK PE=1 SV=1 | 3 |
| Q96J01 | THO complex subunit 3 OS=Homo sapiens OX=9606 GN=THOC3 PE=1 SV=1 | 5 |
| Q96J94 | Piwi-like protein 1 OS=Homo sapiens OX=9606 GN=PIWIL1 PE=1 SV=1 | 7 |
| Q96JB2 | Conserved oligomeric Golgi complex subunit 3 OS=Homo sapiens OX=9606 GN=COG3 PE=1 SV=3 | 3 |
| Q96JB5 | CDK5 regulatory subunit-associated protein 3 OS=Homo sapiens OX=9606 GN=CDK5RAP3 PE=1 SV=2 | 5 |
| Q96JD6 | 1,5-anhydro-D-fructose reductase OS=Homo sapiens OX=9606 GN=AKR1E2 PE=1 SV=2 | 1 |
| Q96JH7 | Deubiquitinating protein VCPIP1 OS=Homo sapiens OX=9606 GN=VCPIP1 PE=1 SV=2 | 15 |
| Q96JI7 | Spatacsin OS=Homo sapiens OX=9606 GN=SPG11 PE=1 SV=3 | 1 |
| Q96JJ3 | Engulfment and cell motility protein 2 OS=Homo sapiens OX=9606 GN=ELMO2 PE=1 SV=2 | 6 |
| Q96JJ7 | Protein disulfide-isomerase TMX3 OS=Homo sapiens OX=9606 GN=TMX3 PE=1 SV=2 | 4 |
| Q96JQ0 | Protocadherin-16 OS=Homo sapiens OX=9606 GN=DCHS1 PE=1 SV=1 | 2 |
| Q96JY0 | Protein maelstrom homolog OS=Homo sapiens OX=9606 GN=MAEL PE=1 SV=1 | 12 |
| Q96JY6 | PDZ and LIM domain protein 2 OS=Homo sapiens OX=9606 GN=PDLIM2 PE=1 SV=1 | 6 |
| Q96K17 | Transcription factor BTF3 homolog 4 OS=Homo sapiens OX=9606 GN=BTF3L4 PE=1 SV=1 | 5 |
| Q96K19 | E3 ubiquitin-protein ligase RNF170 OS=Homo sapiens OX=9606 GN=RNF170 PE=1 SV=2 | 3 |
| Q96K21 | Abscission/NoCut checkpoint regulator OS=Homo sapiens OX=9606 GN=ZFYVE19 PE=1 SV=3 | 3 |
| Q96K76 | Ubiquitin carboxyl-terminal hydrolase 47 OS=Homo sapiens OX=9606 GN=USP47 PE=1 SV=3 | 7 |
| Q96KA5 | Lipid scramblase CLPTM1L OS=Homo sapiens OX=9606 GN=CLPTM1L PE=1 SV=1 | 5 |
| Q96KC8 | DnaJ homolog subfamily C member 1 OS=Homo sapiens OX=9606 GN=DNAJC1 PE=1 SV=1 | 3 |
| Q96KG9 | N-terminal kinase-like protein OS=Homo sapiens OX=9606 GN=SCYL1 PE=1 SV=1 | 14 |
| Q96KN1 | Protein LRATD2 OS=Homo sapiens OX=9606 GN=LRATD2 PE=1 SV=1 | 1 |
| Q96KP1 | Exocyst complex component 2 OS=Homo sapiens OX=9606 GN=EXOC2 PE=1 SV=1 | 5 |
| Q96KP4 | Cytosolic non-specific dipeptidase OS=Homo sapiens OX=9606 GN=CNDP2 PE=1 SV=2 | 29 |
| Q96KR7 | Phosphatase and actin regulator 3 OS=Homo sapiens OX=9606 GN=PHACTR3 PE=1 SV=1 | 7 |
| Q96L46 | Calpain small subunit 2 OS=Homo sapiens OX=9606 GN=CAPNS2 PE=1 SV=2 | 3 |
| Q96L92 | Sorting nexin-27 OS=Homo sapiens OX=9606 GN=SNX27 PE=1 SV=2 | 7 |
| Q96L93 | Kinesin-like protein KIF16B OS=Homo sapiens OX=9606 GN=KIF16B PE=1 SV=2 | 1 |
| Q96LB3 | Intraflagellar transport protein 74 homolog OS=Homo sapiens OX=9606 GN=IFT74 PE=1 SV=1 | 3 |
| Q96LD8 | Sentrin-specific protease 8 OS=Homo sapiens OX=9606 GN=SENP8 PE=1 SV=1 | 5 |
| Q96LJ7 | Dehydrogenase/reductase SDR family member 1 OS=Homo sapiens OX=9606 GN=DHRS1 PE=1 SV=1 | 2 |
| Q96LM1 | Putative uncharacterized protein encoded by LINC00615 OS=Homo sapiens OX=9606 GN=LINC00615 PE=5 SV=1 | 1 |
| Q96M27 | Protein PRRC1 OS=Homo sapiens OX=9606 GN=PRRC1 PE=1 SV=1 | 4 |
| Q96M93 | Adenosine deaminase domain-containing protein 1 OS=Homo sapiens OX=9606 GN=ADAD1 PE=2 SV=1 | 5 |
| Q96MN2 | NACHT, LRR and PYD domains-containing protein 4 OS=Homo sapiens OX=9606 GN=NLRP4 PE=1 SV=3 | 38 |
| Q96MP5 | Zinc finger SWIM domain-containing protein 3 OS=Homo sapiens OX=9606 GN=ZSWIM3 PE=1 SV=2 | 10 |
| Q96MW1 | Coiled-coil domain-containing protein 43 OS=Homo sapiens OX=9606 GN=CCDC43 PE=1 SV=2 | 1 |
| Q96N06 | Spermatogenesis-associated protein 33 OS=Homo sapiens OX=9606 GN=SPATA33 PE=1 SV=1 | 2 |
| Q96N66 | Lysophospholipid acyltransferase 7 OS=Homo sapiens OX=9606 GN=MBOAT7 PE=1 SV=2 | 4 |
| Q96NA8 | t-SNARE domain-containing protein 1 OS=Homo sapiens OX=9606 GN=TSNARE1 PE=1 SV=2 | 1 |
| Q96NT0 | Coiled-coil domain-containing protein 115 OS=Homo sapiens OX=9606 GN=CCDC115 PE=1 SV=1 | 2 |
| Q96NU0 | Contactin-associated protein-like 3B OS=Homo sapiens OX=9606 GN=CNTNAP3B PE=2 SV=3 | 1 |
| Q96P70 | Importin-9 OS=Homo sapiens OX=9606 GN=IPO9 PE=1 SV=3 | 3 |
| Q96PC5 | Melanoma inhibitory activity protein 2 OS=Homo sapiens OX=9606 GN=MIA2 PE=1 SV=4 | 26 |
| Q96PD5 | N-acetylmuramoyl-L-alanine amidase OS=Homo sapiens OX=9606 GN=PGLYRP2 PE=1 SV=1 | 14 |
| Q96PE5 | Opalin OS=Homo sapiens OX=9606 GN=OPALIN PE=2 SV=1 | 3 |
| Q96PF1 | Protein-glutamine gamma-glutamyltransferase Z OS=Homo sapiens OX=9606 GN=TGM7 PE=1 SV=1 | 2 |
| Q96PU4 | E3 ubiquitin-protein ligase UHRF2 OS=Homo sapiens OX=9606 GN=UHRF2 PE=1 SV=1 | 21 |
| Q96PU5 | E3 ubiquitin-protein ligase NEDD4-like OS=Homo sapiens OX=9606 GN=NEDD4L PE=1 SV=2 | 10 |
| Q96PY5 | Formin-like protein 2 OS=Homo sapiens OX=9606 GN=FMNL2 PE=1 SV=3 | 2 |
| Q96PY6 | Serine/threonine-protein kinase Nek1 OS=Homo sapiens OX=9606 GN=NEK1 PE=1 SV=2 | 10 |
| Q96Q05 | Trafficking protein particle complex subunit 9 OS=Homo sapiens OX=9606 GN=TRAPPC9 PE=1 SV=2 | 1 |
| Q96Q06 | Perilipin-4 OS=Homo sapiens OX=9606 GN=PLIN4 PE=1 SV=3 | 1 |
| Q96Q11 | CCA tRNA nucleotidyltransferase 1, mitochondrial OS=Homo sapiens OX=9606 GN=TRNT1 PE=1 SV=2 | 5 |
| Q96Q42 | Alsin OS=Homo sapiens OX=9606 GN=ALS2 PE=1 SV=2 | 11 |
| Q96Q83 | Alpha-ketoglutarate-dependent dioxygenase alkB homolog 3 OS=Homo sapiens OX=9606 GN=ALKBH3 PE=1 SV=1 | 5 |
| Q96QG7 | Myotubularin-related protein 9 OS=Homo sapiens OX=9606 GN=MTMR9 PE=1 SV=1 | 2 |
| Q96QK1 | Vacuolar protein sorting-associated protein 35 OS=Homo sapiens OX=9606 GN=VPS35 PE=1 SV=2 | 22 |
| Q96QZ7 | Membrane-associated guanylate kinase, WW and PDZ domain-containing protein 1 OS=Homo sapiens OX=9606 GN=MAGI1 PE=1 SV=3 | 1 |
| Q96RD7 | Pannexin-1 OS=Homo sapiens OX=9606 GN=PANX1 PE=1 SV=4 | 2 |
| Q96RF0 | Sorting nexin-18 OS=Homo sapiens OX=9606 GN=SNX18 PE=1 SV=2 | 1 |
| Q96RK0 | Protein capicua homolog OS=Homo sapiens OX=9606 GN=CIC PE=1 SV=2 | 1 |
| Q96RL7 | Intermembrane lipid transfer protein VPS13A OS=Homo sapiens OX=9606 GN=VPS13A PE=1 SV=2 | 6 |
| Q96RM1 | Small proline-rich protein 2F OS=Homo sapiens OX=9606 GN=SPRR2F PE=3 SV=1 | 3 |
| Q96RN5 | Mediator of RNA polymerase II transcription subunit 15 OS=Homo sapiens OX=9606 GN=MED15 PE=1 SV=2 | 1 |
| Q96RP9 | Elongation factor G, mitochondrial OS=Homo sapiens OX=9606 GN=GFM1 PE=1 SV=2 | 31 |
| Q96RS6 | NudC domain-containing protein 1 OS=Homo sapiens OX=9606 GN=NUDCD1 PE=1 SV=2 | 5 |
| Q96RT7 | Gamma-tubulin complex component 6 OS=Homo sapiens OX=9606 GN=TUBGCP6 PE=1 SV=3 | 1 |
| Q96S19 | Methyltransferase-like 26 OS=Homo sapiens OX=9606 GN=METTL26 PE=1 SV=2 | 5 |
| Q96S52 | GPI transamidase component PIG-S OS=Homo sapiens OX=9606 GN=PIGS PE=1 SV=3 | 8 |
| Q96SI9 | Spermatid perinuclear RNA-binding protein OS=Homo sapiens OX=9606 GN=STRBP PE=1 SV=1 | 2 |
| Q96ST3 | Paired amphipathic helix protein Sin3a OS=Homo sapiens OX=9606 GN=SIN3A PE=1 SV=2 | 16 |
| Q96SU4 | Oxysterol-binding protein-related protein 9 OS=Homo sapiens OX=9606 GN=OSBPL9 PE=1 SV=2 | 7 |
| Q96T17 | MAP7 domain-containing protein 2 OS=Homo sapiens OX=9606 GN=MAP7D2 PE=1 SV=2 | 19 |
| Q96T91 | Glycoprotein hormone alpha-2 OS=Homo sapiens OX=9606 GN=GPHA2 PE=1 SV=1 | 1 |
| Q99417 | c-Myc-binding protein OS=Homo sapiens OX=9606 GN=MYCBP PE=1 SV=3 | 7 |
| Q99426 | Tubulin-folding cofactor B OS=Homo sapiens OX=9606 GN=TBCB PE=1 SV=2 | 15 |
| Q99436 | Proteasome subunit beta type-7 OS=Homo sapiens OX=9606 GN=PSMB7 PE=1 SV=1 | 5 |
| Q99442 | Translocation protein SEC62 OS=Homo sapiens OX=9606 GN=SEC62 PE=1 SV=1 | 2 |
| Q99447 | Ethanolamine-phosphate cytidylyltransferase OS=Homo sapiens OX=9606 GN=PCYT2 PE=1 SV=1 | 7 |
| Q99460 | 26S proteasome non-ATPase regulatory subunit 1 OS=Homo sapiens OX=9606 GN=PSMD1 PE=1 SV=2 | 31 |
| Q99471 | Prefoldin subunit 5 OS=Homo sapiens OX=9606 GN=PFDN5 PE=1 SV=2 | 6 |
| Q99487 | Platelet-activating factor acetylhydrolase 2, cytoplasmic OS=Homo sapiens OX=9606 GN=PAFAH2 PE=1 SV=1 | 2 |
| Q99501 | GAS2-like protein 1 OS=Homo sapiens OX=9606 GN=GAS2L1 PE=1 SV=2 | 1 |
| Q99523 | Sortilin OS=Homo sapiens OX=9606 GN=SORT1 PE=1 SV=3 | 5 |
| Q99543 | DnaJ homolog subfamily C member 2 OS=Homo sapiens OX=9606 GN=DNAJC2 PE=1 SV=4 | 1 |
| Q99567 | Nuclear pore complex protein Nup88 OS=Homo sapiens OX=9606 GN=NUP88 PE=1 SV=2 | 2 |
| Q99574 | Neuroserpin OS=Homo sapiens OX=9606 GN=SERPINI1 PE=1 SV=1 | 2 |
| Q99598 | Translin-associated protein X OS=Homo sapiens OX=9606 GN=TSNAX PE=1 SV=1 | 13 |
| Q99611 | Selenide, water dikinase 2 OS=Homo sapiens OX=9606 GN=SEPHS2 PE=1 SV=3 | 10 |
| Q99613 | Eukaryotic translation initiation factor 3 subunit C OS=Homo sapiens OX=9606 GN=EIF3C PE=1 SV=1 | 12 |
| Q99614 | Tetratricopeptide repeat protein 1 OS=Homo sapiens OX=9606 GN=TTC1 PE=1 SV=1 | 6 |
| Q99615 | DnaJ homolog subfamily C member 7 OS=Homo sapiens OX=9606 GN=DNAJC7 PE=1 SV=2 | 9 |
| Q99623 | Prohibitin-2 OS=Homo sapiens OX=9606 GN=PHB2 PE=1 SV=2 | 15 |
| Q99627 | COP9 signalosome complex subunit 8 OS=Homo sapiens OX=9606 GN=COPS8 PE=1 SV=1 | 4 |
| Q99653 | Calcineurin B homologous protein 1 OS=Homo sapiens OX=9606 GN=CHP1 PE=1 SV=3 | 5 |
| Q99661 | Kinesin-like protein KIF2C OS=Homo sapiens OX=9606 GN=KIF2C PE=1 SV=2 | 19 |
| Q99674 | Cell growth regulator with EF hand domain protein 1 OS=Homo sapiens OX=9606 GN=CGREF1 PE=1 SV=3 | 2 |
| Q99714 | 3-hydroxyacyl-CoA dehydrogenase type-2 OS=Homo sapiens OX=9606 GN=HSD17B10 PE=1 SV=3 | 14 |
| Q99726 | Probable proton-coupled zinc antiporter SLC30A3 OS=Homo sapiens OX=9606 GN=SLC30A3 PE=1 SV=2 | 1 |
| Q99729 | Heterogeneous nuclear ribonucleoprotein A/B OS=Homo sapiens OX=9606 GN=HNRNPAB PE=1 SV=2 | 6 |
| Q99733 | Nucleosome assembly protein 1-like 4 OS=Homo sapiens OX=9606 GN=NAP1L4 PE=1 SV=1 | 12 |
| Q99747 | Gamma-soluble NSF attachment protein OS=Homo sapiens OX=9606 GN=NAPG PE=1 SV=1 | 17 |
| Q99757 | Thioredoxin, mitochondrial OS=Homo sapiens OX=9606 GN=TXN2 PE=1 SV=2 | 3 |
| Q99766 | ATP synthase subunit s, mitochondrial OS=Homo sapiens OX=9606 GN=DMAC2L PE=1 SV=3 | 7 |
| Q99797 | Mitochondrial intermediate peptidase OS=Homo sapiens OX=9606 GN=MIPEP PE=1 SV=2 | 2 |
| Q99798 | Aconitate hydratase, mitochondrial OS=Homo sapiens OX=9606 GN=ACO2 PE=1 SV=2 | 35 |
| Q99805 | Transmembrane 9 superfamily member 2 OS=Homo sapiens OX=9606 GN=TM9SF2 PE=1 SV=1 | 11 |
| Q99807 | 5-demethoxyubiquinone hydroxylase, mitochondrial OS=Homo sapiens OX=9606 GN=COQ7 PE=1 SV=3 | 4 |
| Q99808 | Equilibrative nucleoside transporter 1 OS=Homo sapiens OX=9606 GN=SLC29A1 PE=1 SV=3 | 3 |
| Q99816 | Tumor susceptibility gene 101 protein OS=Homo sapiens OX=9606 GN=TSG101 PE=1 SV=2 | 10 |
| Q99829 | Copine-1 OS=Homo sapiens OX=9606 GN=CPNE1 PE=1 SV=1 | 10 |
| Q99961 | Endophilin-A2 OS=Homo sapiens OX=9606 GN=SH3GL1 PE=1 SV=1 | 2 |
| Q99962 | Endophilin-A1 OS=Homo sapiens OX=9606 GN=SH3GL2 PE=1 SV=1 | 10 |
| Q99969 | Retinoic acid receptor responder protein 2 OS=Homo sapiens OX=9606 GN=RARRES2 PE=1 SV=1 | 6 |
| Q99973 | Telomerase protein component 1 OS=Homo sapiens OX=9606 GN=TEP1 PE=1 SV=2 | 1 |
| Q99986 | Serine/threonine-protein kinase VRK1 OS=Homo sapiens OX=9606 GN=VRK1 PE=1 SV=1 | 3 |
| Q9BPU6 | Dihydropyrimidinase-related protein 5 OS=Homo sapiens OX=9606 GN=DPYSL5 PE=1 SV=1 | 18 |
| Q9BPW8 | Protein NipSnap homolog 1 OS=Homo sapiens OX=9606 GN=NIPSNAP1 PE=1 SV=1 | 13 |
| Q9BPX5 | Actin-related protein 2/3 complex subunit 5-like protein OS=Homo sapiens OX=9606 GN=ARPC5L PE=1 SV=1 | 6 |
| Q9BPX6 | Calcium uptake protein 1, mitochondrial OS=Homo sapiens OX=9606 GN=MICU1 PE=1 SV=1 | 1 |
| Q9BPX7 | UPF0415 protein C7orf25 OS=Homo sapiens OX=9606 GN=C7orf25 PE=1 SV=1 | 2 |
| Q9BQ50 | Three prime repair exonuclease 2 OS=Homo sapiens OX=9606 GN=TREX2 PE=1 SV=2 | 2 |
| Q9BQ52 | Zinc phosphodiesterase ELAC protein 2 OS=Homo sapiens OX=9606 GN=ELAC2 PE=1 SV=2 | 3 |
| Q9BQ61 | Telomerase RNA component interacting RNase OS=Homo sapiens OX=9606 GN=TRIR PE=1 SV=1 | 1 |
| Q9BQ69 | ADP-ribose glycohydrolase MACROD1 OS=Homo sapiens OX=9606 GN=MACROD1 PE=1 SV=2 | 1 |
| Q9BQ95 | Evolutionarily conserved signaling intermediate in Toll pathway, mitochondrial OS=Homo sapiens OX=9606 GN=ECSIT PE=1 SV=1 | 5 |
| Q9BQA1 | Methylosome protein 50 OS=Homo sapiens OX=9606 GN=WDR77 PE=1 SV=1 | 7 |
| Q9BQA9 | Cytochrome b-245 chaperone 1 OS=Homo sapiens OX=9606 GN=CYBC1 PE=1 SV=1 | 1 |
| Q9BQB6 | Vitamin K epoxide reductase complex subunit 1 OS=Homo sapiens OX=9606 GN=VKORC1 PE=1 SV=1 | 1 |
| Q9BQP7 | Mitochondrial genome maintenance exonuclease 1 OS=Homo sapiens OX=9606 GN=MGME1 PE=1 SV=1 | 13 |
| Q9BR39 | Junctophilin-2 OS=Homo sapiens OX=9606 GN=JPH2 PE=1 SV=2 | 1 |
| Q9BR76 | Coronin-1B OS=Homo sapiens OX=9606 GN=CORO1B PE=1 SV=1 | 2 |
| Q9BRA0 | N-alpha-acetyltransferase 38, NatC auxiliary subunit OS=Homo sapiens OX=9606 GN=NAA38 PE=1 SV=1 | 1 |
| Q9BRA2 | Thioredoxin domain-containing protein 17 OS=Homo sapiens OX=9606 GN=TXNDC17 PE=1 SV=1 | 11 |
| Q9BRF8 | Serine/threonine-protein phosphatase CPPED1 OS=Homo sapiens OX=9606 GN=CPPED1 PE=1 SV=3 | 3 |
| Q9BRG1 | Vacuolar protein-sorting-associated protein 25 OS=Homo sapiens OX=9606 GN=VPS25 PE=1 SV=1 | 5 |
| Q9BRJ6 | Uncharacterized protein C7orf50 OS=Homo sapiens OX=9606 GN=C7orf50 PE=1 SV=1 | 1 |
| Q9BRK0 | Receptor expression-enhancing protein 2 OS=Homo sapiens OX=9606 GN=REEP2 PE=1 SV=2 | 6 |
| Q9BRK5 | 45 kDa calcium-binding protein OS=Homo sapiens OX=9606 GN=SDF4 PE=1 SV=1 | 4 |
| Q9BRP8 | Partner of Y14 and mago OS=Homo sapiens OX=9606 GN=PYM1 PE=1 SV=1 | 3 |
| Q9BRQ6 | MICOS complex subunit MIC25 OS=Homo sapiens OX=9606 GN=CHCHD6 PE=1 SV=1 | 9 |
| Q9BRR6 | ADP-dependent glucokinase OS=Homo sapiens OX=9606 GN=ADPGK PE=1 SV=1 | 3 |
| Q9BRT3 | Migration and invasion enhancer 1 OS=Homo sapiens OX=9606 GN=MIEN1 PE=1 SV=1 | 2 |
| Q9BRX2 | Protein pelota homolog OS=Homo sapiens OX=9606 GN=PELO PE=1 SV=2 | 8 |
| Q9BRX8 | Peroxiredoxin-like 2A OS=Homo sapiens OX=9606 GN=PRXL2A PE=1 SV=3 | 10 |
| Q9BS26 | Endoplasmic reticulum resident protein 44 OS=Homo sapiens OX=9606 GN=ERP44 PE=1 SV=1 | 14 |
| Q9BSD7 | Cancer-related nucleoside-triphosphatase OS=Homo sapiens OX=9606 GN=NTPCR PE=1 SV=1 | 6 |
| Q9BSF4 | Mitochondrial import inner membrane translocase subunit Tim29 OS=Homo sapiens OX=9606 GN=TIMM29 PE=1 SV=2 | 1 |
| Q9BSG0 | Protease-associated domain-containing protein 1 OS=Homo sapiens OX=9606 GN=PRADC1 PE=1 SV=1 | 2 |
| Q9BSH4 | Translational activator of cytochrome c oxidase 1 OS=Homo sapiens OX=9606 GN=TACO1 PE=1 SV=1 | 3 |
| Q9BSH5 | Haloacid dehalogenase-like hydrolase domain-containing protein 3 OS=Homo sapiens OX=9606 GN=HDHD3 PE=1 SV=1 | 6 |
| Q9BSJ2 | Gamma-tubulin complex component 2 OS=Homo sapiens OX=9606 GN=TUBGCP2 PE=1 SV=2 | 12 |
| Q9BSJ8 | Extended synaptotagmin-1 OS=Homo sapiens OX=9606 GN=ESYT1 PE=1 SV=1 | 1 |
| Q9BSL1 | Ubiquitin-associated domain-containing protein 1 OS=Homo sapiens OX=9606 GN=UBAC1 PE=1 SV=1 | 1 |
| Q9BSQ5 | Cerebral cavernous malformations 2 protein OS=Homo sapiens OX=9606 GN=CCM2 PE=1 SV=1 | 5 |
| Q9BSU1 | Phagosome assembly factor 1 OS=Homo sapiens OX=9606 GN=PHAF1 PE=1 SV=1 | 1 |
| Q9BT09 | Protein canopy homolog 3 OS=Homo sapiens OX=9606 GN=CNPY3 PE=1 SV=1 | 8 |
| Q9BT22 | Chitobiosyldiphosphodolichol beta-mannosyltransferase OS=Homo sapiens OX=9606 GN=ALG1 PE=1 SV=2 | 2 |
| Q9BT25 | HAUS augmin-like complex subunit 8 OS=Homo sapiens OX=9606 GN=HAUS8 PE=1 SV=3 | 1 |
| Q9BT73 | Proteasome assembly chaperone 3 OS=Homo sapiens OX=9606 GN=PSMG3 PE=1 SV=1 | 4 |
| Q9BT78 | COP9 signalosome complex subunit 4 OS=Homo sapiens OX=9606 GN=COPS4 PE=1 SV=1 | 30 |
| Q9BTD8 | RNA-binding protein 42 OS=Homo sapiens OX=9606 GN=RBM42 PE=1 SV=1 | 1 |
| Q9BTE3 | Mini-chromosome maintenance complex-binding protein OS=Homo sapiens OX=9606 GN=MCMBP PE=1 SV=2 | 2 |
| Q9BTE7 | DCN1-like protein 5 OS=Homo sapiens OX=9606 GN=DCUN1D5 PE=1 SV=1 | 4 |
| Q9BTT0 | Acidic leucine-rich nuclear phosphoprotein 32 family member E OS=Homo sapiens OX=9606 GN=ANP32E PE=1 SV=1 | 1 |
| Q9BTV4 | Transmembrane protein 43 OS=Homo sapiens OX=9606 GN=TMEM43 PE=1 SV=1 | 15 |
| Q9BTV5 | Fibronectin type III and SPRY domain-containing protein 1 OS=Homo sapiens OX=9606 GN=FSD1 PE=1 SV=1 | 7 |
| Q9BTW9 | Tubulin-specific chaperone D OS=Homo sapiens OX=9606 GN=TBCD PE=1 SV=2 | 42 |
| Q9BTX1 | Nucleoporin NDC1 OS=Homo sapiens OX=9606 GN=NDC1 PE=1 SV=2 | 3 |
| Q9BU23 | Lipase maturation factor 2 OS=Homo sapiens OX=9606 GN=LMF2 PE=1 SV=2 | 5 |
| Q9BU61 | NADH dehydrogenase [ubiquinone] 1 alpha subcomplex assembly factor 3 OS=Homo sapiens OX=9606 GN=NDUFAF3 PE=1 SV=1 | 5 |
| Q9BUH6 | Protein PAXX OS=Homo sapiens OX=9606 GN=PAXX PE=1 SV=2 | 3 |
| Q9BUK0 | Coiled-coil-helix-coiled-coil-helix domain-containing protein 7 OS=Homo sapiens OX=9606 GN=CHCHD7 PE=1 SV=1 | 1 |
| Q9BUL8 | Programmed cell death protein 10 OS=Homo sapiens OX=9606 GN=PDCD10 PE=1 SV=1 | 6 |
| Q9BUN8 | Derlin-1 OS=Homo sapiens OX=9606 GN=DERL1 PE=1 SV=1 | 2 |
| Q9BUP3 | Oxidoreductase HTATIP2 OS=Homo sapiens OX=9606 GN=HTATIP2 PE=1 SV=2 | 3 |
| Q9BUR5 | MICOS complex subunit MIC26 OS=Homo sapiens OX=9606 GN=APOO PE=1 SV=1 | 6 |
| Q9BUT1 | Dehydrogenase/reductase SDR family member 6 OS=Homo sapiens OX=9606 GN=BDH2 PE=1 SV=2 | 9 |
| Q9BV19 | Uncharacterized protein C1orf50 OS=Homo sapiens OX=9606 GN=C1orf50 PE=1 SV=2 | 7 |
| Q9BV38 | WD repeat-containing protein 18 OS=Homo sapiens OX=9606 GN=WDR18 PE=1 SV=2 | 5 |
| Q9BV40 | Vesicle-associated membrane protein 8 OS=Homo sapiens OX=9606 GN=VAMP8 PE=1 SV=1 | 5 |
| Q9BV44 | tRNA (guanine(6)-N2)-methyltransferase THUMP3 OS=Homo sapiens OX=9606 GN=THUMPD3 PE=1 SV=1 | 1 |
| Q9BV57 | Acireductone dioxygenase OS=Homo sapiens OX=9606 GN=ADI1 PE=1 SV=1 | 3 |
| Q9BV73 | Centrosome-associated protein CEP250 OS=Homo sapiens OX=9606 GN=CEP250 PE=1 SV=2 | 33 |
| Q9BV79 | Enoyl-[acyl-carrier-protein] reductase, mitochondrial OS=Homo sapiens OX=9606 GN=MECR PE=1 SV=2 | 7 |
| Q9BVC4 | Target of rapamycin complex subunit LST8 OS=Homo sapiens OX=9606 GN=MLST8 PE=1 SV=1 | 6 |
| Q9BVC6 | Transmembrane protein 109 OS=Homo sapiens OX=9606 GN=TMEM109 PE=1 SV=1 | 2 |
| Q9BVG3 | E3 ubiquitin-protein ligase TRIM62 OS=Homo sapiens OX=9606 GN=TRIM62 PE=1 SV=1 | 1 |
| Q9BVK6 | Transmembrane emp24 domain-containing protein 9 OS=Homo sapiens OX=9606 GN=TMED9 PE=1 SV=2 | 6 |
| Q9BVL2 | Nucleoporin p58/p45 OS=Homo sapiens OX=9606 GN=NUP58 PE=1 SV=1 | 3 |
| Q9BVL4 | Protein adenylyltransferase SelO, mitochondrial OS=Homo sapiens OX=9606 GN=SELENOO PE=1 SV=3 | 1 |
| Q9BVM2 | Protein DPCD OS=Homo sapiens OX=9606 GN=DPCD PE=1 SV=2 | 5 |
| Q9BVM4 | Gamma-glutamylaminecyclotransferase OS=Homo sapiens OX=9606 GN=GGACT PE=1 SV=2 | 2 |
| Q9BW91 | ADP-ribose pyrophosphatase, mitochondrial OS=Homo sapiens OX=9606 GN=NUDT9 PE=1 SV=1 | 3 |
| Q9BWD1 | Acetyl-CoA acetyltransferase, cytosolic OS=Homo sapiens OX=9606 GN=ACAT2 PE=1 SV=2 | 16 |
| Q9BWF3 | RNA-binding protein 4 OS=Homo sapiens OX=9606 GN=RBM4 PE=1 SV=1 | 2 |
| Q9BWS9 | Chitinase domain-containing protein 1 OS=Homo sapiens OX=9606 GN=CHID1 PE=1 SV=1 | 8 |
| Q9BX40 | Protein LSM14 homolog B OS=Homo sapiens OX=9606 GN=LSM14B PE=1 SV=1 | 14 |
| Q9BX46;Q9H0Z9 | RNA-binding protein 24 OS=Homo sapiens OX=9606 GN=RBM24 PE=1 SV=1 | 1 |
| Q9BX66 | Sorbin and SH3 domain-containing protein 1 OS=Homo sapiens OX=9606 GN=SORBS1 PE=1 SV=3 | 1 |
| Q9BX67 | Junctional adhesion molecule C OS=Homo sapiens OX=9606 GN=JAM3 PE=1 SV=1 | 2 |
| Q9BX68 | Adenosine 5'-monophosphoramidase HINT2 OS=Homo sapiens OX=9606 GN=HINT2 PE=1 SV=1 | 6 |
| Q9BXB5 | Oxysterol-binding protein-related protein 10 OS=Homo sapiens OX=9606 GN=OSBPL10 PE=1 SV=2 | 5 |
| Q9BXF6 | Rab11 family-interacting protein 5 OS=Homo sapiens OX=9606 GN=RAB11FIP5 PE=1 SV=1 | 8 |
| Q9BXK5 | Bcl-2-like protein 13 OS=Homo sapiens OX=9606 GN=BCL2L13 PE=1 SV=1 | 5 |
| Q9BXS5 | AP-1 complex subunit mu-1 OS=Homo sapiens OX=9606 GN=AP1M1 PE=1 SV=3 | 12 |
| Q9BXU1 | Serine/threonine-protein kinase 31 OS=Homo sapiens OX=9606 GN=STK31 PE=2 SV=2 | 32 |
| Q9BXW7 | Haloacid dehalogenase-like hydrolase domain-containing 5 OS=Homo sapiens OX=9606 GN=HDHD5 PE=1 SV=1 | 4 |
| Q9BY44 | Eukaryotic translation initiation factor 2A OS=Homo sapiens OX=9606 GN=EIF2A PE=1 SV=3 | 17 |
| Q9BY67 | Cell adhesion molecule 1 OS=Homo sapiens OX=9606 GN=CADM1 PE=1 SV=2 | 2 |
| Q9BYD5 | Cornifelin OS=Homo sapiens OX=9606 GN=CNFN PE=1 SV=2 | 1 |
| Q9BYJ1 | Hydroperoxide isomerase ALOXE3 OS=Homo sapiens OX=9606 GN=ALOXE3 PE=1 SV=1 | 5 |
| Q9BYN0 | Sulfiredoxin-1 OS=Homo sapiens OX=9606 GN=SRXN1 PE=1 SV=2 | 1 |
| Q9BYN8 | 28S ribosomal protein S26, mitochondrial OS=Homo sapiens OX=9606 GN=MRPS26 PE=1 SV=1 | 2 |
| Q9BYR8 | Keratin-associated protein 3-1 OS=Homo sapiens OX=9606 GN=KRTAP3-1 PE=1 SV=1 | 1 |
| Q9BYT8 | Neurolysin, mitochondrial OS=Homo sapiens OX=9606 GN=NLN PE=1 SV=1 | 16 |
| Q9BYX2 | TBC1 domain family member 2A OS=Homo sapiens OX=9606 GN=TBC1D2 PE=1 SV=3 | 3 |
| Q9BZ23 | Pantothenate kinase 2, mitochondrial OS=Homo sapiens OX=9606 GN=PANK2 PE=1 SV=3 | 5 |
| Q9BZ29 | Dedicator of cytokinesis protein 9 OS=Homo sapiens OX=9606 GN=DOCK9 PE=1 SV=2 | 1 |
| Q9BZB8 | Cytoplasmic polyadenylation element-binding protein 1 OS=Homo sapiens OX=9606 GN=CPEB1 PE=1 SV=1 | 6 |
| Q9BZE1 | 39S ribosomal protein L37, mitochondrial OS=Homo sapiens OX=9606 GN=MRPL37 PE=1 SV=2 | 1 |
| Q9BZE9 | Tether containing UBX domain for GLUT4 OS=Homo sapiens OX=9606 GN=ASPSCR1 PE=1 SV=1 | 14 |
| Q9BZF1 | Oxysterol-binding protein-related protein 8 OS=Homo sapiens OX=9606 GN=OSBPL8 PE=1 SV=3 | 12 |
| Q9BZI7 | Regulator of nonsense transcripts 3B OS=Homo sapiens OX=9606 GN=UPF3B PE=1 SV=1 | 2 |
| Q9BZZ5 | Apoptosis inhibitor 5 OS=Homo sapiens OX=9606 GN=API5 PE=1 SV=3 | 10 |
| Q9C002 | Normal mucosa of esophagus-specific gene 1 protein OS=Homo sapiens OX=9606 GN=NMES1 PE=1 SV=1 | 5 |
| Q9C0C9 | (E3-independent) E2 ubiquitin-conjugating enzyme OS=Homo sapiens OX=9606 GN=UBE2O PE=1 SV=3 | 17 |
| Q9C0E4 | Glutamate receptor-interacting protein 2 OS=Homo sapiens OX=9606 GN=GRIP2 PE=1 SV=4 | 1 |
| Q9C0E8 | Endoplasmic reticulum junction formation protein lunapark OS=Homo sapiens OX=9606 GN=LNPK PE=1 SV=2 | 11 |
| Q9C0I1 | Myotubularin-related protein 12 OS=Homo sapiens OX=9606 GN=MTMR12 PE=1 SV=2 | 5 |
| Q9GZM7 | Tubulointerstitial nephritis antigen-like OS=Homo sapiens OX=9606 GN=TINAGL1 PE=1 SV=1 | 3 |
| Q9GZN8 | UPF0687 protein C20orf27 OS=Homo sapiens OX=9606 GN=C20orf27 PE=1 SV=3 | 1 |
| Q9GZP4 | PITH domain-containing protein 1 OS=Homo sapiens OX=9606 GN=PITHD1 PE=1 SV=1 | 15 |
| Q9GZQ3 | COMM domain-containing protein 5 OS=Homo sapiens OX=9606 GN=COMMD5 PE=1 SV=1 | 2 |
| Q9GZS3 | SKI8 subunit of superkiller complex protein OS=Homo sapiens OX=9606 GN=SKIC8 PE=1 SV=1 | 9 |
| Q9GZT3 | SRA stem-loop-interacting RNA-binding protein, mitochondrial OS=Homo sapiens OX=9606 GN=SLIRP PE=1 SV=1 | 8 |
| Q9GZT6 | Coiled-coil domain-containing protein 90B, mitochondrial OS=Homo sapiens OX=9606 GN=CCDC90B PE=1 SV=2 | 5 |
| Q9GZT8 | NIF3-like protein 1 OS=Homo sapiens OX=9606 GN=NIF3L1 PE=1 SV=2 | 8 |
| Q9GZY4 | Cytochrome c oxidase assembly factor 1 homolog OS=Homo sapiens OX=9606 GN=COA1 PE=1 SV=1 | 4 |
| Q9GZY8 | Mitochondrial fission factor OS=Homo sapiens OX=9606 GN=MFF PE=1 SV=1 | 6 |
| Q9GZZ1 | N-alpha-acetyltransferase 50 OS=Homo sapiens OX=9606 GN=NAA50 PE=1 SV=1 | 9 |
| Q9GZZ9 | Ubiquitin-like modifier-activating enzyme 5 OS=Homo sapiens OX=9606 GN=UBA5 PE=1 SV=1 | 11 |
| Q9H008 | Phospholysine phosphohistidine inorganic pyrophosphate phosphatase OS=Homo sapiens OX=9606 GN=LHPP PE=1 SV=2 | 8 |
| Q9H019 | Mitochondrial fission regulator 1-like OS=Homo sapiens OX=9606 GN=MTFR1L PE=1 SV=2 | 1 |
| Q9H061 | Transmembrane protein 126A OS=Homo sapiens OX=9606 GN=TMEM126A PE=1 SV=1 | 8 |
| Q9H074 | Polyadenylate-binding protein-interacting protein 1 OS=Homo sapiens OX=9606 GN=PAIP1 PE=1 SV=1 | 7 |
| Q9H078 | Caseinolytic peptidase B protein homolog OS=Homo sapiens OX=9606 GN=CLPB PE=1 SV=1 | 6 |
| Q9H098 | Protein FAM107B OS=Homo sapiens OX=9606 GN=FAM107B PE=1 SV=1 | 2 |
| Q9H0A8 | COMM domain-containing protein 4 OS=Homo sapiens OX=9606 GN=COMMD4 PE=1 SV=1 | 8 |
| Q9H0B3 | IQ domain-containing protein N OS=Homo sapiens OX=9606 GN=IQCN PE=1 SV=1 | 1 |
| Q9H0B6 | Kinesin light chain 2 OS=Homo sapiens OX=9606 GN=KLC2 PE=1 SV=1 | 3 |
| Q9H0C2 | ADP/ATP translocase 4 OS=Homo sapiens OX=9606 GN=SLC25A31 PE=1 SV=1 | 3 |
| Q9H0C8 | Integrin-linked kinase-associated serine/threonine phosphatase 2C OS=Homo sapiens OX=9606 GN=ILKAP PE=1 SV=1 | 5 |
| Q9H0D6 | 5'-3' exoribonuclease 2 OS=Homo sapiens OX=9606 GN=XRN2 PE=1 SV=1 | 3 |
| Q9H0E2 | Toll-interacting protein OS=Homo sapiens OX=9606 GN=TOLLIP PE=1 SV=1 | 7 |
| Q9H0H3 | Kelch-like protein 25 OS=Homo sapiens OX=9606 GN=KLHL25 PE=1 SV=1 | 4 |
| Q9H0H5 | Rac GTPase-activating protein 1 OS=Homo sapiens OX=9606 GN=RACGAP1 PE=1 SV=1 | 4 |
| Q9H0J9 | Protein mono-ADP-ribosyltransferase PARP12 OS=Homo sapiens OX=9606 GN=PARP12 PE=1 SV=1 | 2 |
| Q9H0N5 | Pterin-4-alpha-carbinolamine dehydratase 2 OS=Homo sapiens OX=9606 GN=PCBD2 PE=1 SV=4 | 3 |
| Q9H0R4 | Haloacid dehalogenase-like hydrolase domain-containing protein 2 OS=Homo sapiens OX=9606 GN=HDHD2 PE=1 SV=1 | 3 |
| Q9H0U3 | Magnesium transporter protein 1 OS=Homo sapiens OX=9606 GN=MAGT1 PE=1 SV=1 | 4 |
| Q9H0U4 | Ras-related protein Rab-1B OS=Homo sapiens OX=9606 GN=RAB1B PE=1 SV=1 | 5 |
| Q9H0U6 | 39S ribosomal protein L18, mitochondrial OS=Homo sapiens OX=9606 GN=MRPL18 PE=1 SV=1 | 3 |
| Q9H0V1 | Transmembrane protein 168 OS=Homo sapiens OX=9606 GN=TMEM168 PE=1 SV=2 | 2 |
| Q9H0V9 | VIP36-like protein OS=Homo sapiens OX=9606 GN=LMAN2L PE=1 SV=1 | 6 |
| Q9H0W8 | Nonsense-mediated mRNA decay factor SMG9 OS=Homo sapiens OX=9606 GN=SMG9 PE=1 SV=1 | 1 |
| Q9H115 | Beta-soluble NSF attachment protein OS=Homo sapiens OX=9606 GN=NAPB PE=1 SV=2 | 12 |
| Q9H116 | GDNF-inducible zinc finger protein 1 OS=Homo sapiens OX=9606 GN=GZF1 PE=1 SV=1 | 1 |
| Q9H173 | Nucleotide exchange factor SIL1 OS=Homo sapiens OX=9606 GN=SIL1 PE=1 SV=1 | 10 |
| Q9H1B7 | Probable E3 ubiquitin-protein ligase IRF2BPL OS=Homo sapiens OX=9606 GN=IRF2BPL PE=1 SV=1 | 2 |
| Q9H1E3 | Nuclear ubiquitous casein and cyclin-dependent kinase substrate 1 OS=Homo sapiens OX=9606 GN=NUCKS1 PE=1 SV=1 | 1 |
| Q9H1E5 | Thioredoxin-related transmembrane protein 4 OS=Homo sapiens OX=9606 GN=TMX4 PE=1 SV=1 | 3 |
| Q9H1K1 | Iron-sulfur cluster assembly enzyme ISCU OS=Homo sapiens OX=9606 GN=ISCU PE=1 SV=2 | 9 |
| Q9H223 | EH domain-containing protein 4 OS=Homo sapiens OX=9606 GN=EHD4 PE=1 SV=1 | 5 |
| Q9H267 | Vacuolar protein sorting-associated protein 33B OS=Homo sapiens OX=9606 GN=VPS33B PE=1 SV=2 | 5 |
| Q9H269 | Vacuolar protein sorting-associated protein 16 homolog OS=Homo sapiens OX=9606 GN=VPS16 PE=1 SV=2 | 8 |
| Q9H270 | Vacuolar protein sorting-associated protein 11 homolog OS=Homo sapiens OX=9606 GN=VPS11 PE=1 SV=1 | 3 |
| Q9H2C0 | Gigaxonin OS=Homo sapiens OX=9606 GN=GAN PE=1 SV=1 | 3 |
| Q9H2D6 | TRIO and F-actin-binding protein OS=Homo sapiens OX=9606 GN=TRIOBP PE=1 SV=3 | 3 |
| Q9H2G2 | STE20-like serine/threonine-protein kinase OS=Homo sapiens OX=9606 GN=SLK PE=1 SV=1 | 4 |
| Q9H2H8 | Peptidyl-prolyl cis-trans isomerase-like 3 OS=Homo sapiens OX=9606 GN=PPIL3 PE=1 SV=1 | 2 |
| Q9H2J4 | Phosducin-like protein 3 OS=Homo sapiens OX=9606 GN=PDCL3 PE=1 SV=1 | 13 |
| Q9H2K0 | Translation initiation factor IF-3, mitochondrial OS=Homo sapiens OX=9606 GN=MTIF3 PE=1 SV=4 | 2 |
| Q9H2M9 | Rab3 GTPase-activating protein non-catalytic subunit OS=Homo sapiens OX=9606 GN=RAB3GAP2 PE=1 SV=1 | 18 |
| Q9H2U1 | ATP-dependent DNA/RNA helicase DHX36 OS=Homo sapiens OX=9606 GN=DHX36 PE=1 SV=2 | 10 |
| Q9H2U2 | Inorganic pyrophosphatase 2, mitochondrial OS=Homo sapiens OX=9606 GN=PPA2 PE=1 SV=2 | 12 |
| Q9H2V7 | Protein spinster homolog 1 OS=Homo sapiens OX=9606 GN=SPNS1 PE=1 SV=1 | 2 |
| Q9H2W6 | 39S ribosomal protein L46, mitochondrial OS=Homo sapiens OX=9606 GN=MRPL46 PE=1 SV=1 | 5 |
| Q9H300 | Presenilins-associated rhomboid-like protein, mitochondrial OS=Homo sapiens OX=9606 GN=PARL PE=1 SV=2 | 8 |
| Q9H313 | Protein tweety homolog 1 OS=Homo sapiens OX=9606 GN=TTYH1 PE=1 SV=1 | 3 |
| Q9H330 | Transmembrane protein 245 OS=Homo sapiens OX=9606 GN=TMEM245 PE=1 SV=3 | 5 |
| Q9H336 | Cysteine-rich secretory protein LCCL domain-containing 1 OS=Homo sapiens OX=9606 GN=CRISPLD1 PE=2 SV=1 | 11 |
| Q9H3G5 | Probable serine carboxypeptidase CPVL OS=Homo sapiens OX=9606 GN=CPVL PE=1 SV=2 | 7 |
| Q9H3K2 | Growth hormone-inducible transmembrane protein OS=Homo sapiens OX=9606 GN=GHITM PE=1 SV=2 | 1 |
| Q9H3K6 | BolA-like protein 2 OS=Homo sapiens OX=9606 GN=BOLA2B PE=1 SV=1 | 4 |
| Q9H3N1 | Thioredoxin-related transmembrane protein 1 OS=Homo sapiens OX=9606 GN=TMX1 PE=1 SV=1 | 7 |
| Q9H3P7 | Golgi resident protein GCP60 OS=Homo sapiens OX=9606 GN=ACBD3 PE=1 SV=4 | 7 |
| Q9H3S7 | Tyrosine-protein phosphatase non-receptor type 23 OS=Homo sapiens OX=9606 GN=PTPN23 PE=1 SV=1 | 10 |
| Q9H3Z4 | DnaJ homolog subfamily C member 5 OS=Homo sapiens OX=9606 GN=DNAJC5 PE=1 SV=1 | 1 |
| Q9H446 | RWD domain-containing protein 1 OS=Homo sapiens OX=9606 GN=RWDD1 PE=1 SV=1 | 2 |
| Q9H469 | F-box/LRR-repeat protein 15 OS=Homo sapiens OX=9606 GN=FBXL15 PE=1 SV=2 | 2 |
| Q9H477 | Ribokinase OS=Homo sapiens OX=9606 GN=RBKS PE=1 SV=1 | 7 |
| Q9H479 | Fructosamine-3-kinase OS=Homo sapiens OX=9606 GN=FN3K PE=1 SV=1 | 3 |
| Q9H488 | GDP-fucose protein O-fucosyltransferase 1 OS=Homo sapiens OX=9606 GN=POFUT1 PE=1 SV=1 | 7 |
| Q9H4A3 | Serine/threonine-protein kinase WNK1 OS=Homo sapiens OX=9606 GN=WNK1 PE=1 SV=2 | 2 |
| Q9H4A4 | Aminopeptidase B OS=Homo sapiens OX=9606 GN=RNPEP PE=1 SV=2 | 12 |
| Q9H4A5 | Golgi phosphoprotein 3-like OS=Homo sapiens OX=9606 GN=GOLPH3L PE=1 SV=1 | 11 |
| Q9H4H8 | Protein FAM83D OS=Homo sapiens OX=9606 GN=FAM83D PE=1 SV=3 | 1 |
| Q9H4I9 | Essential MCU regulator, mitochondrial OS=Homo sapiens OX=9606 GN=SMDT1 PE=1 SV=1 | 1 |
| Q9H4M9 | EH domain-containing protein 1 OS=Homo sapiens OX=9606 GN=EHD1 PE=1 SV=2 | 15 |
| Q9H5Q4 | Dimethyladenosine transferase 2, mitochondrial OS=Homo sapiens OX=9606 GN=TFB2M PE=1 SV=1 | 1 |
| Q9H5X1 | Cytosolic iron-sulfur assembly component 2A OS=Homo sapiens OX=9606 GN=CIAO2A PE=1 SV=1 | 1 |
| Q9H6E4 | Coiled-coil domain-containing protein 134 OS=Homo sapiens OX=9606 GN=CCDC134 PE=1 SV=1 | 2 |
| Q9H6H4 | Receptor expression-enhancing protein 4 OS=Homo sapiens OX=9606 GN=REEP4 PE=1 SV=1 | 2 |
| Q9H6K1 | Protein ILRUN OS=Homo sapiens OX=9606 GN=ILRUN PE=1 SV=2 | 1 |
| Q9H6K4 | Optic atrophy 3 protein OS=Homo sapiens OX=9606 GN=OPA3 PE=1 SV=1 | 2 |
| Q9H6R7 | WD repeat and coiled-coil-containing protein OS=Homo sapiens OX=9606 GN=WDCP PE=1 SV=1 | 7 |
| Q9H6T3 | RNA polymerase II-associated protein 3 OS=Homo sapiens OX=9606 GN=RPAP3 PE=1 SV=2 | 10 |
| Q9H6U6 | BCAS3 microtubule associated cell migration factor OS=Homo sapiens OX=9606 GN=BCAS3 PE=1 SV=3 | 4 |
| Q9H6W3 | Ribosomal oxygenase 1 OS=Homo sapiens OX=9606 GN=RIOX1 PE=1 SV=2 | 1 |
| Q9H6Z4 | Ran-binding protein 3 OS=Homo sapiens OX=9606 GN=RANBP3 PE=1 SV=1 | 6 |
| Q9H773 | dCTP pyrophosphatase 1 OS=Homo sapiens OX=9606 GN=DCTPP1 PE=1 SV=1 | 3 |
| Q9H7C9 | Mth938 domain-containing protein OS=Homo sapiens OX=9606 GN=AAMDC PE=1 SV=1 | 4 |
| Q9H7L9 | Sin3 histone deacetylase corepressor complex component SDS3 OS=Homo sapiens OX=9606 GN=SUDS3 PE=1 SV=2 | 5 |
| Q9H7M9 | V-type immunoglobulin domain-containing suppressor of T-cell activation OS=Homo sapiens OX=9606 GN=VSIR PE=1 SV=3 | 1 |
| Q9H7Z7 | Prostaglandin E synthase 2 OS=Homo sapiens OX=9606 GN=PTGES2 PE=1 SV=1 | 3 |
| Q9H832 | Ubiquitin-conjugating enzyme E2 Z OS=Homo sapiens OX=9606 GN=UBE2Z PE=1 SV=2 | 13 |
| Q9H845 | Complex I assembly factor ACAD9, mitochondrial OS=Homo sapiens OX=9606 GN=ACAD9 PE=1 SV=1 | 15 |
| Q9H8M7 | Ubiquitin carboxyl-terminal hydrolase MINDY-3 OS=Homo sapiens OX=9606 GN=MINDY3 PE=1 SV=1 | 8 |
| Q9H8T0 | AKT-interacting protein OS=Homo sapiens OX=9606 GN=AKTIP PE=1 SV=1 | 2 |
| Q9H8Y8 | Golgi reassembly-stacking protein 2 OS=Homo sapiens OX=9606 GN=GORASP2 PE=1 SV=3 | 4 |
| Q9H910 | Jupiter microtubule associated homolog 2 OS=Homo sapiens OX=9606 GN=JPT2 PE=1 SV=1 | 8 |
| Q9H936 | Mitochondrial glutamate carrier 1 OS=Homo sapiens OX=9606 GN=SLC25A22 PE=1 SV=1 | 6 |
| Q9H944 | Mediator of RNA polymerase II transcription subunit 20 OS=Homo sapiens OX=9606 GN=MED20 PE=1 SV=1 | 2 |
| Q9H974 | Queuine tRNA-ribosyltransferase accessory subunit 2 OS=Homo sapiens OX=9606 GN=QTRT2 PE=1 SV=1 | 1 |
| Q9H977 | WD repeat-containing protein 54 OS=Homo sapiens OX=9606 GN=WDR54 PE=1 SV=1 | 5 |
| Q9H9A5 | CCR4-NOT transcription complex subunit 10 OS=Homo sapiens OX=9606 GN=CNOT10 PE=1 SV=1 | 2 |
| Q9H9A6 | Leucine-rich repeat-containing protein 40 OS=Homo sapiens OX=9606 GN=LRRC40 PE=1 SV=1 | 32 |
| Q9H9B4 | Sideroflexin-1 OS=Homo sapiens OX=9606 GN=SFXN1 PE=1 SV=4 | 16 |
| Q9H9C1 | Spermatogenesis-defective protein 39 homolog OS=Homo sapiens OX=9606 GN=VIPAS39 PE=1 SV=1 | 3 |
| Q9H9H4 | Vacuolar protein sorting-associated protein 37B OS=Homo sapiens OX=9606 GN=VPS37B PE=1 SV=1 | 2 |
| Q9H9J2 | 39S ribosomal protein L44, mitochondrial OS=Homo sapiens OX=9606 GN=MRPL44 PE=1 SV=1 | 9 |
| Q9H9P8 | L-2-hydroxyglutarate dehydrogenase, mitochondrial OS=Homo sapiens OX=9606 GN=L2HGDH PE=1 SV=3 | 11 |
| Q9H9Q2 | COP9 signalosome complex subunit 7b OS=Homo sapiens OX=9606 GN=COPS7B PE=1 SV=1 | 8 |
| Q9H9Z2 | Protein lin-28 homolog A OS=Homo sapiens OX=9606 GN=LIN28A PE=1 SV=1 | 2 |
| Q9HA64 | Ketosamine-3-kinase OS=Homo sapiens OX=9606 GN=FN3KRP PE=1 SV=2 | 9 |
| Q9HA65 | TBC1 domain family member 17 OS=Homo sapiens OX=9606 GN=TBC1D17 PE=1 SV=3 | 1 |
| Q9HA77 | Probable cysteine--tRNA ligase, mitochondrial OS=Homo sapiens OX=9606 GN=CARS2 PE=1 SV=1 | 3 |
| Q9HAB8 | Phosphopantothenate--cysteine ligase OS=Homo sapiens OX=9606 GN=PPCS PE=1 SV=2 | 13 |
| Q9HAD4 | WD repeat-containing protein 41 OS=Homo sapiens OX=9606 GN=WDR41 PE=1 SV=3 | 1 |
| Q9HAS3 | Solute carrier family 28 member 3 OS=Homo sapiens OX=9606 GN=SLC28A3 PE=1 SV=1 | 1 |
| Q9HAU0 | Pleckstrin homology domain-containing family A member 5 OS=Homo sapiens OX=9606 GN=PLEKHA5 PE=1 SV=1 | 1 |
| Q9HAU5 | Regulator of nonsense transcripts 2 OS=Homo sapiens OX=9606 GN=UPF2 PE=1 SV=1 | 3 |
| Q9HAV0 | Guanine nucleotide-binding protein subunit beta-4 OS=Homo sapiens OX=9606 GN=GNB4 PE=1 SV=3 | 5 |
| Q9HAV7 | GrpE protein homolog 1, mitochondrial OS=Homo sapiens OX=9606 GN=GRPEL1 PE=1 SV=2 | 11 |
| Q9HB07 | MYG1 exonuclease OS=Homo sapiens OX=9606 GN=MYG1 PE=1 SV=3 | 2 |
| Q9HB20 | Pleckstrin homology domain-containing family A member 3 OS=Homo sapiens OX=9606 GN=PLEKHA3 PE=1 SV=2 | 1 |
| Q9HB40 | Retinoid-inducible serine carboxypeptidase OS=Homo sapiens OX=9606 GN=SCPEP1 PE=1 SV=1 | 2 |
| Q9HB66 | Alternative protein MKKS OS=Homo sapiens OX=9606 GN=MKKS PE=1 SV=1 | 1 |
| Q9HB71 | Calcyclin-binding protein OS=Homo sapiens OX=9606 GN=CACYBP PE=1 SV=2 | 14 |
| Q9HB90 | Ras-related GTP-binding protein C OS=Homo sapiens OX=9606 GN=RRAGC PE=1 SV=1 | 12 |
| Q9HBE4 | Interleukin-21 OS=Homo sapiens OX=9606 GN=IL21 PE=1 SV=3 | 1 |
| Q9HBH1 | Peptide deformylase, mitochondrial OS=Homo sapiens OX=9606 GN=PDF PE=1 SV=1 | 7 |
| Q9HBH5 | Retinol dehydrogenase 14 OS=Homo sapiens OX=9606 GN=RDH14 PE=1 SV=1 | 3 |
| Q9HC21 | Mitochondrial thiamine pyrophosphate carrier OS=Homo sapiens OX=9606 GN=SLC25A19 PE=1 SV=1 | 4 |
| Q9HC35 | Echinoderm microtubule-associated protein-like 4 OS=Homo sapiens OX=9606 GN=EML4 PE=1 SV=3 | 18 |
| Q9HC36 | rRNA methyltransferase 3, mitochondrial OS=Homo sapiens OX=9606 GN=MRM3 PE=1 SV=2 | 4 |
| Q9HC38 | Glyoxalase domain-containing protein 4 OS=Homo sapiens OX=9606 GN=GLOD4 PE=1 SV=1 | 14 |
| Q9HC77 | Centromere protein J OS=Homo sapiens OX=9606 GN=CENPJ PE=1 SV=2 | 1 |
| Q9HC84 | Mucin-5B OS=Homo sapiens OX=9606 GN=MUC5B PE=1 SV=3 | 1 |
| Q9HCC0 | Methylcrotonoyl-CoA carboxylase beta chain, mitochondrial OS=Homo sapiens OX=9606 GN=MCCC2 PE=1 SV=1 | 18 |
| Q9HCD6 | Protein TANC2 OS=Homo sapiens OX=9606 GN=TANC2 PE=1 SV=3 | 1 |
| Q9HCF6 | Transient receptor potential cation channel subfamily M member 3 OS=Homo sapiens OX=9606 GN=TRPM3 PE=2 SV=4 | 1 |
| Q9HCI5 | Melanoma-associated antigen E1 OS=Homo sapiens OX=9606 GN=MAGEE1 PE=1 SV=2 | 1 |
| Q9HCN8 | Stromal cell-derived factor 2-like protein 1 OS=Homo sapiens OX=9606 GN=SDF2L1 PE=1 SV=2 | 8 |
| Q9HCU4 | Cadherin EGF LAG seven-pass G-type receptor 2 OS=Homo sapiens OX=9606 GN=CELSR2 PE=1 SV=1 | 1 |
| Q9HCU5 | Prolactin regulatory element-binding protein OS=Homo sapiens OX=9606 GN=PREB PE=1 SV=2 | 9 |
| Q9HD20 | Endoplasmic reticulum transmembrane helix translocase OS=Homo sapiens OX=9606 GN=ATP13A1 PE=1 SV=2 | 5 |
| Q9HD26 | Golgi-associated PDZ and coiled-coil motif-containing protein OS=Homo sapiens OX=9606 GN=GOPC PE=1 SV=1 | 7 |
| Q9HD33 | 39S ribosomal protein L47, mitochondrial OS=Homo sapiens OX=9606 GN=MRPL47 PE=1 SV=2 | 7 |
| Q9HD34 | LYR motif-containing protein 4 OS=Homo sapiens OX=9606 GN=LYRM4 PE=1 SV=1 | 3 |
| Q9HD42 | Charged multivesicular body protein 1a OS=Homo sapiens OX=9606 GN=CHMP1A PE=1 SV=1 | 3 |
| Q9HD45 | Transmembrane 9 superfamily member 3 OS=Homo sapiens OX=9606 GN=TM9SF3 PE=1 SV=2 | 2 |
| Q9HD67 | Unconventional myosin-X OS=Homo sapiens OX=9606 GN=MYO10 PE=1 SV=3 | 25 |
| Q9HDC9 | Adipocyte plasma membrane-associated protein OS=Homo sapiens OX=9606 GN=APMAP PE=1 SV=2 | 12 |
| Q9NNW7 | Thioredoxin reductase 2, mitochondrial OS=Homo sapiens OX=9606 GN=TXNRD2 PE=1 SV=3 | 1 |
| Q9NP58 | ATP-binding cassette sub-family B member 6 OS=Homo sapiens OX=9606 GN=ABCB6 PE=1 SV=1 | 1 |
| Q9NP61 | ADP-ribosylation factor GTPase-activating protein 3 OS=Homo sapiens OX=9606 GN=ARFGAP3 PE=1 SV=1 | 8 |
| Q9NP72 | Ras-related protein Rab-18 OS=Homo sapiens OX=9606 GN=RAB18 PE=1 SV=1 | 11 |
| Q9NP73 | Putative bifunctional UDP-N-acetylglucosamine transferase and deubiquitinase ALG13 OS=Homo sapiens OX=9606 GN=ALG13 PE=1 SV=2 | 6 |
| Q9NP79 | Vacuolar protein sorting-associated protein VTA1 homolog OS=Homo sapiens OX=9606 GN=VTA1 PE=1 SV=1 | 8 |
| Q9NP81 | Serine--tRNA ligase, mitochondrial OS=Homo sapiens OX=9606 GN=SARS2 PE=1 SV=1 | 25 |
| Q9NP97 | Dynein light chain roadblock-type 1 OS=Homo sapiens OX=9606 GN=DYNLRB1 PE=1 SV=3 | 5 |
| Q9NPA0 | ER membrane protein complex subunit 7 OS=Homo sapiens OX=9606 GN=EMC7 PE=1 SV=1 | 3 |
| Q9NPB8 | Glycerophosphocholine phosphodiesterase GPCPD1 OS=Homo sapiens OX=9606 GN=GPCPD1 PE=1 SV=2 | 3 |
| Q9NPD3 | Exosome complex component RRP41 OS=Homo sapiens OX=9606 GN=EXOSC4 PE=1 SV=3 | 1 |
| Q9NPE2 | Neugrin OS=Homo sapiens OX=9606 GN=NGRN PE=1 SV=2 | 8 |
| Q9NPH2 | Inositol-3-phosphate synthase 1 OS=Homo sapiens OX=9606 GN=ISYNA1 PE=1 SV=1 | 8 |
| Q9NPJ3 | Acyl-coenzyme A thioesterase 13 OS=Homo sapiens OX=9606 GN=ACOT13 PE=1 SV=1 | 4 |
| Q9NPQ8 | Synembryn-A OS=Homo sapiens OX=9606 GN=RIC8A PE=1 SV=3 | 2 |
| Q9NQ50 | 39S ribosomal protein L40, mitochondrial OS=Homo sapiens OX=9606 GN=MRPL40 PE=1 SV=1 | 3 |
| Q9NQ86 | E3 ubiquitin-protein ligase TRIM36 OS=Homo sapiens OX=9606 GN=TRIM36 PE=1 SV=2 | 6 |
| Q9NQ88 | Fructose-2,6-bisphosphatase TIGAR OS=Homo sapiens OX=9606 GN=TIGAR PE=1 SV=1 | 2 |
| Q9NQ92 | Coordinator of PRMT5 and differentiation stimulator OS=Homo sapiens OX=9606 GN=COPRS PE=1 SV=3 | 1 |
| Q9NQC3 | Reticulon-4 OS=Homo sapiens OX=9606 GN=RTN4 PE=1 SV=2 | 15 |
| Q9NQH7 | Xaa-Pro aminopeptidase 3 OS=Homo sapiens OX=9606 GN=XPNPEP3 PE=1 SV=1 | 1 |
| Q9NQI0 | Probable ATP-dependent RNA helicase DDX4 OS=Homo sapiens OX=9606 GN=DDX4 PE=1 SV=2 | 28 |
| Q9NQP4 | Prefoldin subunit 4 OS=Homo sapiens OX=9606 GN=PFDN4 PE=1 SV=1 | 4 |
| Q9NQR4 | Omega-amidase NIT2 OS=Homo sapiens OX=9606 GN=NIT2 PE=1 SV=1 | 17 |
| Q9NQS1 | Cell death regulator Aven OS=Homo sapiens OX=9606 GN=AVEN PE=1 SV=1 | 9 |
| Q9NQT8 | Kinesin-like protein KIF13B OS=Homo sapiens OX=9606 GN=KIF13B PE=1 SV=2 | 72 |
| Q9NQW6 | Anillin OS=Homo sapiens OX=9606 GN=ANLN PE=1 SV=2 | 16 |
| Q9NQW7 | Xaa-Pro aminopeptidase 1 OS=Homo sapiens OX=9606 GN=XPNPEP1 PE=1 SV=3 | 10 |
| Q9NQX3 | Gephyrin OS=Homo sapiens OX=9606 GN=GPHN PE=1 SV=1 | 2 |
| Q9NR09 | Baculoviral IAP repeat-containing protein 6 OS=Homo sapiens OX=9606 GN=BIRC6 PE=1 SV=3 | 3 |
| Q9NR28 | Diablo IAP-binding mitochondrial protein OS=Homo sapiens OX=9606 GN=DIABLO PE=1 SV=1 | 11 |
| Q9NR31 | GTP-binding protein SAR1a OS=Homo sapiens OX=9606 GN=SAR1A PE=1 SV=1 | 9 |
| Q9NR45 | Sialic acid synthase OS=Homo sapiens OX=9606 GN=NANS PE=1 SV=2 | 13 |
| Q9NR46 | Endophilin-B2 OS=Homo sapiens OX=9606 GN=SH3GLB2 PE=1 SV=1 | 14 |
| Q9NR50 | Translation initiation factor eIF-2B subunit gamma OS=Homo sapiens OX=9606 GN=EIF2B3 PE=1 SV=1 | 8 |
| Q9NR96 | Toll-like receptor 9 OS=Homo sapiens OX=9606 GN=TLR9 PE=1 SV=2 | 1 |
| Q9NR99 | Matrix-remodeling-associated protein 5 OS=Homo sapiens OX=9606 GN=MXRA5 PE=1 SV=3 | 1 |
| Q9NRA8 | Eukaryotic translation initiation factor 4E transporter OS=Homo sapiens OX=9606 GN=EIF4ENIF1 PE=1 SV=2 | 20 |
| Q9NRG9 | Aladin OS=Homo sapiens OX=9606 GN=AAAS PE=1 SV=1 | 12 |
| Q9NRH3 | Tubulin gamma-2 chain OS=Homo sapiens OX=9606 GN=TUBG2 PE=2 SV=1 | 2 |
| Q9NRP0 | Oligosaccharyltransferase complex subunit OSTC OS=Homo sapiens OX=9606 GN=OSTC PE=1 SV=1 | 1 |
| Q9NRP2 | COX assembly mitochondrial protein 2 homolog OS=Homo sapiens OX=9606 GN=CMC2 PE=1 SV=1 | 1 |
| Q9NRR5 | Ubiquilin-4 OS=Homo sapiens OX=9606 GN=UBQLN4 PE=1 SV=2 | 3 |
| Q9NRW4 | Dual specificity protein phosphatase 22 OS=Homo sapiens OX=9606 GN=DUSP22 PE=1 SV=1 | 1 |
| Q9NRW7 | Vacuolar protein sorting-associated protein 45 OS=Homo sapiens OX=9606 GN=VPS45 PE=1 SV=1 | 7 |
| Q9NRX2 | 39S ribosomal protein L17, mitochondrial OS=Homo sapiens OX=9606 GN=MRPL17 PE=1 SV=1 | 2 |
| Q9NRX4 | 14 kDa phosphohistidine phosphatase OS=Homo sapiens OX=9606 GN=PHPT1 PE=1 SV=1 | 7 |
| Q9NRY5 | Protein FAM114A2 OS=Homo sapiens OX=9606 GN=FAM114A2 PE=1 SV=4 | 20 |
| Q9NRZ7 | 1-acyl-sn-glycerol-3-phosphate acyltransferase gamma OS=Homo sapiens OX=9606 GN=AGPAT3 PE=1 SV=1 | 2 |
| Q9NS69 | Mitochondrial import receptor subunit TOM22 homolog OS=Homo sapiens OX=9606 GN=TOMM22 PE=1 SV=3 | 4 |
| Q9NS87 | Kinesin-like protein KIF15 OS=Homo sapiens OX=9606 GN=KIF15 PE=1 SV=1 | 2 |
| Q9NSA3 | Beta-catenin-interacting protein 1 OS=Homo sapiens OX=9606 GN=CTNNBIP1 PE=1 SV=1 | 1 |
| Q9NSD9 | Phenylalanine--tRNA ligase beta subunit OS=Homo sapiens OX=9606 GN=FARSB PE=1 SV=3 | 21 |
| Q9NSE4 | Isoleucine--tRNA ligase, mitochondrial OS=Homo sapiens OX=9606 GN=IARS2 PE=1 SV=2 | 34 |
| Q9NSV4 | Protein diaphanous homolog 3 OS=Homo sapiens OX=9606 GN=DIAPH3 PE=1 SV=4 | 3 |
| Q9NSY1 | BMP-2-inducible protein kinase OS=Homo sapiens OX=9606 GN=BMP2K PE=1 SV=2 | 5 |
| Q9NSY2 | StAR-related lipid transfer protein 5 OS=Homo sapiens OX=9606 GN=STARD5 PE=1 SV=2 | 1 |
| Q9NT62 | Ubiquitin-like-conjugating enzyme ATG3 OS=Homo sapiens OX=9606 GN=ATG3 PE=1 SV=1 | 4 |
| Q9NTJ3 | Structural maintenance of chromosomes protein 4 OS=Homo sapiens OX=9606 GN=SMC4 PE=1 SV=2 | 3 |
| Q9NTJ4 | Alpha-mannosidase 2C1 OS=Homo sapiens OX=9606 GN=MAN2C1 PE=1 SV=1 | 3 |
| Q9NTJ5 | Phosphatidylinositol-3-phosphatase SAC1 OS=Homo sapiens OX=9606 GN=SACM1L PE=1 SV=2 | 24 |
| Q9NTK5 | Obg-like ATPase 1 OS=Homo sapiens OX=9606 GN=OLA1 PE=1 SV=2 | 9 |
| Q9NTX5 | Ethylmalonyl-CoA decarboxylase OS=Homo sapiens OX=9606 GN=ECHDC1 PE=1 SV=2 | 10 |
| Q9NTZ6 | RNA-binding protein 12 OS=Homo sapiens OX=9606 GN=RBM12 PE=1 SV=1 | 5 |
| Q9NUJ1 | Palmitoyl-protein thioesterase ABHD10, mitochondrial OS=Homo sapiens OX=9606 GN=ABHD10 PE=1 SV=1 | 16 |
| Q9NUL5 | Shiftless antiviral inhibitor of ribosomal frameshifting protein OS=Homo sapiens OX=9606 GN=SHFL PE=1 SV=2 | 4 |
| Q9NUL7 | Probable ATP-dependent RNA helicase DDX28 OS=Homo sapiens OX=9606 GN=DDX28 PE=1 SV=2 | 6 |
| Q9NUP9 | Protein lin-7 homolog C OS=Homo sapiens OX=9606 GN=LIN7C PE=1 SV=1 | 3 |
| Q9NUQ3 | Gamma-taxilin OS=Homo sapiens OX=9606 GN=TXLNG PE=1 SV=2 | 2 |
| Q9NUQ7 | Ufm1-specific protease 2 OS=Homo sapiens OX=9606 GN=UFSP2 PE=1 SV=3 | 7 |
| Q9NUQ8 | ATP-binding cassette sub-family F member 3 OS=Homo sapiens OX=9606 GN=ABCF3 PE=1 SV=2 | 6 |
| Q9NUQ9 | CYFIP-related Rac1 interactor B OS=Homo sapiens OX=9606 GN=CYRIB PE=1 SV=1 | 13 |
| Q9NUT2 | Mitochondrial potassium channel ATP-binding subunit OS=Homo sapiens OX=9606 GN=ABCB8 PE=1 SV=3 | 6 |
| Q9NUU7 | ATP-dependent RNA helicase DDX19A OS=Homo sapiens OX=9606 GN=DDX19A PE=1 SV=1 | 5 |
| Q9NV70 | Exocyst complex component 1 OS=Homo sapiens OX=9606 GN=EXOC1 PE=1 SV=4 | 6 |
| Q9NV96 | Cell cycle control protein 50A OS=Homo sapiens OX=9606 GN=TMEM30A PE=1 SV=1 | 1 |
| Q9NVA2 | Septin-11 OS=Homo sapiens OX=9606 GN=SEPTIN11 PE=1 SV=3 | 18 |
| Q9NVH1 | DnaJ homolog subfamily C member 11 OS=Homo sapiens OX=9606 GN=DNAJC11 PE=1 SV=2 | 13 |
| Q9NVJ2 | ADP-ribosylation factor-like protein 8B OS=Homo sapiens OX=9606 GN=ARL8B PE=1 SV=1 | 5 |
| Q9NVM4 | Protein arginine N-methyltransferase 7 OS=Homo sapiens OX=9606 GN=PRMT7 PE=1 SV=1 | 4 |
| Q9NVR5 | Protein kintoun OS=Homo sapiens OX=9606 GN=DNAAF2 PE=1 SV=2 | 1 |
| Q9NVS9 | Pyridoxine-5'-phosphate oxidase OS=Homo sapiens OX=9606 GN=PNPO PE=1 SV=1 | 1 |
| Q9NVT9 | Armadillo repeat-containing protein 1 OS=Homo sapiens OX=9606 GN=ARMC1 PE=1 SV=1 | 5 |
| Q9NVZ3 | Adaptin ear-binding coat-associated protein 2 OS=Homo sapiens OX=9606 GN=NECAP2 PE=1 SV=1 | 2 |
| Q9NW15 | Anoctamin-10 OS=Homo sapiens OX=9606 GN=ANO10 PE=1 SV=2 | 7 |
| Q9NW68 | BSD domain-containing protein 1 OS=Homo sapiens OX=9606 GN=BSDC1 PE=1 SV=1 | 5 |
| Q9NWM8 | Peptidyl-prolyl cis-trans isomerase FKBP14 OS=Homo sapiens OX=9606 GN=FKBP14 PE=1 SV=1 | 1 |
| Q9NWS0 | PIH1 domain-containing protein 1 OS=Homo sapiens OX=9606 GN=PIH1D1 PE=1 SV=1 | 4 |
| Q9NWS8 | Required for meiotic nuclear division protein 1 homolog OS=Homo sapiens OX=9606 GN=RMND1 PE=1 SV=2 | 7 |
| Q9NWT8 | Aurora kinase A-interacting protein OS=Homo sapiens OX=9606 GN=AURKAIP1 PE=1 SV=1 | 1 |
| Q9NWU1 | 3-oxoacyl-[acyl-carrier-protein] synthase, mitochondrial OS=Homo sapiens OX=9606 GN=OXSM PE=1 SV=1 | 4 |
| Q9NWU5 | 39S ribosomal protein L22, mitochondrial OS=Homo sapiens OX=9606 GN=MRPL22 PE=1 SV=1 | 12 |
| Q9NWV4 | CXXC motif containing zinc binding protein OS=Homo sapiens OX=9606 GN=CZIB PE=1 SV=1 | 7 |
| Q9NWV8 | BRISC and BRCA1-A complex member 1 OS=Homo sapiens OX=9606 GN=BABAM1 PE=1 SV=1 | 2 |
| Q9NWY4 | Histone PARylation factor 1 OS=Homo sapiens OX=9606 GN=HPF1 PE=1 SV=2 | 11 |
| Q9NWZ8 | Gem-associated protein 8 OS=Homo sapiens OX=9606 GN=GEMIN8 PE=1 SV=1 | 1 |
| Q9NX00 | Transmembrane protein 160 OS=Homo sapiens OX=9606 GN=TMEM160 PE=1 SV=1 | 1 |
| Q9NX08 | COMM domain-containing protein 8 OS=Homo sapiens OX=9606 GN=COMMD8 PE=1 SV=1 | 4 |
| Q9NX14 | NADH dehydrogenase [ubiquinone] 1 beta subcomplex subunit 11, mitochondrial OS=Homo sapiens OX=9606 GN=NDUFB11 PE=1 SV=1 | 2 |
| Q9NX18 | Succinate dehydrogenase assembly factor 2, mitochondrial OS=Homo sapiens OX=9606 GN=SDHAF2 PE=1 SV=1 | 2 |
| Q9NX40 | OCIA domain-containing protein 1 OS=Homo sapiens OX=9606 GN=OCIAD1 PE=1 SV=1 | 4 |
| Q9NX46 | ADP-ribosylhydrolase ARH3 OS=Homo sapiens OX=9606 GN=ADPRS PE=1 SV=1 | 8 |
| Q9NX55 | Huntingtin-interacting protein K OS=Homo sapiens OX=9606 GN=HYPK PE=1 SV=3 | 3 |
| Q9NX57 | Ras-related protein Rab-20 OS=Homo sapiens OX=9606 GN=RAB20 PE=1 SV=1 | 1 |
| Q9NX63 | MICOS complex subunit MIC19 OS=Homo sapiens OX=9606 GN=CHCHD3 PE=1 SV=1 | 12 |
| Q9NXA8 | NAD-dependent protein deacylase sirtuin-5, mitochondrial OS=Homo sapiens OX=9606 GN=SIRT5 PE=1 SV=2 | 5 |
| Q9NXC5 | GATOR complex protein MIOS OS=Homo sapiens OX=9606 GN=MIOS PE=1 SV=2 | 2 |
| Q9NXG2 | THUMP domain-containing protein 1 OS=Homo sapiens OX=9606 GN=THUMPD1 PE=1 SV=2 | 2 |
| Q9NXJ5 | Pyroglutamyl-peptidase 1 OS=Homo sapiens OX=9606 GN=PGPEP1 PE=1 SV=1 | 4 |
| Q9NXN4 | Ganglioside-induced differentiation-associated protein 2 OS=Homo sapiens OX=9606 GN=GDAP2 PE=1 SV=1 | 1 |
| Q9NXZ2 | Probable ATP-dependent RNA helicase DDX43 OS=Homo sapiens OX=9606 GN=DDX43 PE=1 SV=2 | 7 |
| Q9NY33 | Dipeptidyl peptidase 3 OS=Homo sapiens OX=9606 GN=DPP3 PE=1 SV=2 | 28 |
| Q9NY64 | Solute carrier family 2, facilitated glucose transporter member 8 OS=Homo sapiens OX=9606 GN=SLC2A8 PE=1 SV=3 | 1 |
| Q9NY97 | N-acetyllactosaminide beta-1,3-N-acetylglucosaminyltransferase 2 OS=Homo sapiens OX=9606 GN=B3GNT2 PE=1 SV=2 | 9 |
| Q9NYC9 | Dynein axonemal heavy chain 9 OS=Homo sapiens OX=9606 GN=DNAH9 PE=1 SV=3 | 1 |
| Q9NYG2 | Palmitoyltransferase ZDHHC3 OS=Homo sapiens OX=9606 GN=ZDHHC3 PE=1 SV=2 | 2 |
| Q9NYK5 | 39S ribosomal protein L39, mitochondrial OS=Homo sapiens OX=9606 GN=MRPL39 PE=1 SV=3 | 9 |
| Q9NYL9 | Tropomodulin-3 OS=Homo sapiens OX=9606 GN=TMOD3 PE=1 SV=1 | 3 |
| Q9NYR9 | NF-kappa-B inhibitor-interacting Ras-like protein 2 OS=Homo sapiens OX=9606 GN=NKIRAS2 PE=1 SV=1 | 1 |
| Q9NYU2 | UDP-glucose:glycoprotein glucosyltransferase 1 OS=Homo sapiens OX=9606 GN=UGGT1 PE=1 SV=3 | 48 |
| Q9NYY8 | FAST kinase domain-containing protein 2, mitochondrial OS=Homo sapiens OX=9606 GN=FASTKD2 PE=1 SV=1 | 5 |
| Q9NYZ4 | Sialic acid-binding Ig-like lectin 8 OS=Homo sapiens OX=9606 GN=SIGLEC8 PE=1 SV=2 | 1 |
| Q9NZ01 | Very-long-chain enoyl-CoA reductase OS=Homo sapiens OX=9606 GN=TECR PE=1 SV=1 | 8 |
| Q9NZ32 | Actin-related protein 10 OS=Homo sapiens OX=9606 GN=ACTR10 PE=1 SV=1 | 4 |
| Q9NZ43 | Vesicle transport protein USE1 OS=Homo sapiens OX=9606 GN=USE1 PE=1 SV=2 | 6 |
| Q9NZ45 | CDGSH iron-sulfur domain-containing protein 1 OS=Homo sapiens OX=9606 GN=CISD1 PE=1 SV=1 | 5 |
| Q9NZ52 | ADP-ribosylation factor-binding protein GGA3 OS=Homo sapiens OX=9606 GN=GGA3 PE=1 SV=1 | 1 |
| Q9NZD8 | Maspardin OS=Homo sapiens OX=9606 GN=SPG21 PE=1 SV=1 | 3 |
| Q9NZJ6 | Ubiquinone biosynthesis O-methyltransferase, mitochondrial OS=Homo sapiens OX=9606 GN=COQ3 PE=1 SV=3 | 5 |
| Q9NZJ7 | Mitochondrial carrier homolog 1 OS=Homo sapiens OX=9606 GN=MTCH1 PE=1 SV=1 | 5 |
| Q9NZK5 | Adenosine deaminase 2 OS=Homo sapiens OX=9606 GN=ADA2 PE=1 SV=2 | 1 |
| Q9NZL4 | Hsp70-binding protein 1 OS=Homo sapiens OX=9606 GN=HSPBP1 PE=1 SV=2 | 7 |
| Q9NZL9 | Methionine adenosyltransferase 2 subunit beta OS=Homo sapiens OX=9606 GN=MAT2B PE=1 SV=1 | 9 |
| Q9NZN3 | EH domain-containing protein 3 OS=Homo sapiens OX=9606 GN=EHD3 PE=1 SV=2 | 2 |
| Q9NZP8 | Complement C1r subcomponent-like protein OS=Homo sapiens OX=9606 GN=C1RL PE=1 SV=2 | 5 |
| Q9NZR2 | Low-density lipoprotein receptor-related protein 1B OS=Homo sapiens OX=9606 GN=LRP1B PE=1 SV=2 | 2 |
| Q9NZT2 | Opioid growth factor receptor OS=Homo sapiens OX=9606 GN=OGFR PE=1 SV=3 | 4 |
| Q9NZW5 | Protein PALS2 OS=Homo sapiens OX=9606 GN=PALS2 PE=1 SV=2 | 1 |
| Q9NZZ3 | Charged multivesicular body protein 5 OS=Homo sapiens OX=9606 GN=CHMP5 PE=1 SV=1 | 1 |
| Q9P000 | COMM domain-containing protein 9 OS=Homo sapiens OX=9606 GN=COMMD9 PE=1 SV=2 | 4 |
| Q9P015 | 39S ribosomal protein L15, mitochondrial OS=Homo sapiens OX=9606 GN=MRPL15 PE=1 SV=1 | 3 |
| Q9P016 | Thymocyte nuclear protein 1 OS=Homo sapiens OX=9606 GN=THYN1 PE=1 SV=1 | 1 |
| Q9P032 | NADH dehydrogenase [ubiquinone] 1 alpha subcomplex assembly factor 4 OS=Homo sapiens OX=9606 GN=NDUFAF4 PE=1 SV=1 | 6 |
| Q9P0I2 | ER membrane protein complex subunit 3 OS=Homo sapiens OX=9606 GN=EMC3 PE=1 SV=3 | 4 |
| Q9P0J0 | NADH dehydrogenase [ubiquinone] 1 alpha subcomplex subunit 13 OS=Homo sapiens OX=9606 GN=NDUFA13 PE=1 SV=3 | 5 |
| Q9P0J1 | [Pyruvate dehydrogenase [acetyl-transferring]]-phosphatase 1, mitochondrial OS=Homo sapiens OX=9606 GN=PDP1 PE=1 SV=3 | 11 |
| Q9P0K7 | Ankycorbin OS=Homo sapiens OX=9606 GN=RAI14 PE=1 SV=2 | 2 |
| Q9P0L0 | Vesicle-associated membrane protein-associated protein A OS=Homo sapiens OX=9606 GN=VAPA PE=1 SV=3 | 9 |
| Q9P0M2 | A-kinase anchor protein 7 isoform gamma OS=Homo sapiens OX=9606 GN=AKAP7 PE=1 SV=2 | 2 |
| Q9P0M9 | 39S ribosomal protein L27, mitochondrial OS=Homo sapiens OX=9606 GN=MRPL27 PE=1 SV=1 | 4 |
| Q9P0N9 | TBC1 domain family member 7 OS=Homo sapiens OX=9606 GN=TBC1D7 PE=1 SV=1 | 1 |
| Q9P0P8 | Mitochondrial transcription rescue factor 1 OS=Homo sapiens OX=9606 GN=MTRES1 PE=1 SV=1 | 3 |
| Q9P0R6 | GSK3B-interacting protein OS=Homo sapiens OX=9606 GN=GSKIP PE=1 SV=2 | 3 |
| Q9P0T7 | Proton-transporting V-type ATPase complex assembly regulator TMEM9 OS=Homo sapiens OX=9606 GN=TMEM9 PE=1 SV=1 | 2 |
| Q9P0V9 | Septin-10 OS=Homo sapiens OX=9606 GN=SEPTIN10 PE=1 SV=2 | 4 |
| Q9P1Y5 | Calmodulin-regulated spectrin-associated protein 3 OS=Homo sapiens OX=9606 GN=CAMSAP3 PE=1 SV=2 | 5 |
| Q9P1Z2 | Calcium-binding and coiled-coil domain-containing protein 1 OS=Homo sapiens OX=9606 GN=CALCOCO1 PE=1 SV=2 | 9 |
| Q9P219 | Protein Daple OS=Homo sapiens OX=9606 GN=CCDC88C PE=1 SV=3 | 1 |
| Q9P258 | Protein RCC2 OS=Homo sapiens OX=9606 GN=RCC2 PE=1 SV=2 | 11 |
| Q9P265 | Disco-interacting protein 2 homolog B OS=Homo sapiens OX=9606 GN=DIP2B PE=1 SV=3 | 7 |
| Q9P287 | BRCA2 and CDKN1A-interacting protein OS=Homo sapiens OX=9606 GN=BCCIP PE=1 SV=1 | 2 |
| Q9P2B2 | Prostaglandin F2 receptor negative regulator OS=Homo sapiens OX=9606 GN=PTGFRN PE=1 SV=2 | 26 |
| Q9P2D1 | Chromodomain-helicase-DNA-binding protein 7 OS=Homo sapiens OX=9606 GN=CHD7 PE=1 SV=3 | 1 |
| Q9P2E9 | Ribosome-binding protein 1 OS=Homo sapiens OX=9606 GN=RRBP1 PE=1 SV=5 | 26 |
| Q9P2F5 | Storkhead-box protein 2 OS=Homo sapiens OX=9606 GN=STOX2 PE=2 SV=2 | 1 |
| Q9P2H5 | Ubiquitin carboxyl-terminal hydrolase 35 OS=Homo sapiens OX=9606 GN=USP35 PE=1 SV=3 | 1 |
| Q9P2J5 | Leucine--tRNA ligase, cytoplasmic OS=Homo sapiens OX=9606 GN=LARS1 PE=1 SV=2 | 21 |
| Q9P2K8 | eIF-2-alpha kinase GCN2 OS=Homo sapiens OX=9606 GN=EIF2AK4 PE=1 SV=3 | 1 |
| Q9P2M7 | Cingulin OS=Homo sapiens OX=9606 GN=CGN PE=1 SV=3 | 13 |
| Q9P2N4 | A disintegrin and metalloproteinase with thrombospondin motifs 9 OS=Homo sapiens OX=9606 GN=ADAMTS9 PE=1 SV=4 | 1 |
| Q9P2R3 | Rabankyrin-5 OS=Homo sapiens OX=9606 GN=ANKFY1 PE=1 SV=2 | 4 |
| Q9P2R7 | Succinate--CoA ligase [ADP-forming] subunit beta, mitochondrial OS=Homo sapiens OX=9606 GN=SUCLA2 PE=1 SV=3 | 21 |
| Q9P2T1 | GMP reductase 2 OS=Homo sapiens OX=9606 GN=GMPR2 PE=1 SV=1 | 3 |
| Q9P2W9 | Syntaxin-18 OS=Homo sapiens OX=9606 GN=STX18 PE=1 SV=1 | 4 |
| Q9P2X3 | Protein IMPACT OS=Homo sapiens OX=9606 GN=IMPACT PE=1 SV=2 | 4 |
| Q9UBC2 | Epidermal growth factor receptor substrate 15-like 1 OS=Homo sapiens OX=9606 GN=EPS15L1 PE=1 SV=1 | 14 |
| Q9UBD0 | Heat shock transcription factor, X-linked OS=Homo sapiens OX=9606 GN=HSFX2 PE=1 SV=1 | 1 |
| Q9UBE0 | SUMO-activating enzyme subunit 1 OS=Homo sapiens OX=9606 GN=SAE1 PE=1 SV=1 | 3 |
| Q9UBF2 | Coatomer subunit gamma-2 OS=Homo sapiens OX=9606 GN=COPG2 PE=1 SV=1 | 8 |
| Q9UBF6 | RING-box protein 2 OS=Homo sapiens OX=9606 GN=RNF7 PE=1 SV=1 | 3 |
| Q9UBI1 | COMM domain-containing protein 3 OS=Homo sapiens OX=9606 GN=COMMD3 PE=1 SV=1 | 3 |
| Q9UBI6 | Guanine nucleotide-binding protein G(I)/G(S)/G(O) subunit gamma-12 OS=Homo sapiens OX=9606 GN=GNG12 PE=1 SV=3 | 4 |
| Q9UBM7 | 7-dehydrocholesterol reductase OS=Homo sapiens OX=9606 GN=DHCR7 PE=1 SV=1 | 1 |
| Q9UBQ0 | Vacuolar protein sorting-associated protein 29 OS=Homo sapiens OX=9606 GN=VPS29 PE=1 SV=1 | 10 |
| Q9UBQ5 | Eukaryotic translation initiation factor 3 subunit K OS=Homo sapiens OX=9606 GN=EIF3K PE=1 SV=1 | 5 |
| Q9UBS3 | DnaJ homolog subfamily B member 9 OS=Homo sapiens OX=9606 GN=DNAJB9 PE=1 SV=1 | 9 |
| Q9UBS5 | Gamma-aminobutyric acid type B receptor subunit 1 OS=Homo sapiens OX=9606 GN=GABBR1 PE=1 SV=1 | 1 |
| Q9UBS8 | E3 ubiquitin-protein ligase RNF14 OS=Homo sapiens OX=9606 GN=RNF14 PE=1 SV=1 | 4 |
| Q9UBU6 | Protein FAM8A1 OS=Homo sapiens OX=9606 GN=FAM8A1 PE=1 SV=1 | 1 |
| Q9UBV2 | Protein sel-1 homolog 1 OS=Homo sapiens OX=9606 GN=SEL1L PE=1 SV=3 | 21 |
| Q9UBV8 | Peflin OS=Homo sapiens OX=9606 GN=PEF1 PE=1 SV=1 | 3 |
| Q9UBX3 | Mitochondrial dicarboxylate carrier OS=Homo sapiens OX=9606 GN=SLC25A10 PE=1 SV=2 | 2 |
| Q9UBX5 | Fibulin-5 OS=Homo sapiens OX=9606 GN=FBLN5 PE=1 SV=1 | 4 |
| Q9UBX7 | Kallikrein-11 OS=Homo sapiens OX=9606 GN=KLK11 PE=1 SV=2 | 1 |
| Q9UDW1 | Cytochrome b-c1 complex subunit 9 OS=Homo sapiens OX=9606 GN=UQCR10 PE=1 SV=3 | 2 |
| Q9UDY2 | Tight junction protein ZO-2 OS=Homo sapiens OX=9606 GN=TJP2 PE=1 SV=2 | 8 |
| Q9UDY4 | DnaJ homolog subfamily B member 4 OS=Homo sapiens OX=9606 GN=DNAJB4 PE=1 SV=1 | 3 |
| Q9UEW8 | STE20/SPS1-related proline-alanine-rich protein kinase OS=Homo sapiens OX=9606 GN=STK39 PE=1 SV=3 | 6 |
| Q9UF11 | Pleckstrin homology domain-containing family B member 1 OS=Homo sapiens OX=9606 GN=PLEKHB1 PE=1 SV=1 | 1 |
| Q9UFC0 | Leucine-rich repeat and WD repeat-containing protein 1 OS=Homo sapiens OX=9606 GN=LRWD1 PE=1 SV=2 | 1 |
| Q9UFF9 | CCR4-NOT transcription complex subunit 8 OS=Homo sapiens OX=9606 GN=CNOT8 PE=1 SV=1 | 2 |
| Q9UFG5 | UPF0449 protein C19orf25 OS=Homo sapiens OX=9606 GN=C19orf25 PE=1 SV=2 | 2 |
| Q9UFN0 | Protein NipSnap homolog 3A OS=Homo sapiens OX=9606 GN=NIPSNAP3A PE=1 SV=2 | 20 |
| Q9UG56 | Phosphatidylserine decarboxylase proenzyme, mitochondrial OS=Homo sapiens OX=9606 GN=PISD PE=1 SV=4 | 3 |
| Q9UG63 | ATP-binding cassette sub-family F member 2 OS=Homo sapiens OX=9606 GN=ABCF2 PE=1 SV=2 | 2 |
| Q9UGC7 | Peptide chain release factor 1-like, mitochondrial OS=Homo sapiens OX=9606 GN=MTRF1L PE=1 SV=1 | 2 |
| Q9UGI8 | Testin OS=Homo sapiens OX=9606 GN=TES PE=1 SV=1 | 5 |
| Q9UGJ1 | Gamma-tubulin complex component 4 OS=Homo sapiens OX=9606 GN=TUBGCP4 PE=1 SV=1 | 4 |
| Q9UGM3 | Deleted in malignant brain tumors 1 protein OS=Homo sapiens OX=9606 GN=DMBT1 PE=1 SV=2 | 2 |
| Q9UGM5 | Fetuin-B OS=Homo sapiens OX=9606 GN=FETUB PE=1 SV=2 | 5 |
| Q9UGM6 | Tryptophan--tRNA ligase, mitochondrial OS=Homo sapiens OX=9606 GN=WARS2 PE=1 SV=1 | 2 |
| Q9UGP8 | Translocation protein SEC63 homolog OS=Homo sapiens OX=9606 GN=SEC63 PE=1 SV=2 | 15 |
| Q9UGV2 | Protein NDRG3 OS=Homo sapiens OX=9606 GN=NDRG3 PE=1 SV=2 | 3 |
| Q9UH62 | Armadillo repeat-containing X-linked protein 3 OS=Homo sapiens OX=9606 GN=ARMCX3 PE=1 SV=1 | 1 |
| Q9UH65 | Switch-associated protein 70 OS=Homo sapiens OX=9606 GN=SWAP70 PE=1 SV=1 | 9 |
| Q9UHA4 | Ragulator complex protein LAMTOR3 OS=Homo sapiens OX=9606 GN=LAMTOR3 PE=1 SV=1 | 1 |
| Q9UHB6 | LIM domain and actin-binding protein 1 OS=Homo sapiens OX=9606 GN=LIMA1 PE=1 SV=1 | 34 |
| Q9UHB9 | Signal recognition particle subunit SRP68 OS=Homo sapiens OX=9606 GN=SRP68 PE=1 SV=2 | 7 |
| Q9UHD1 | Cysteine and histidine-rich domain-containing protein 1 OS=Homo sapiens OX=9606 GN=CHORDC1 PE=1 SV=2 | 10 |
| Q9UHD8 | Septin-9 OS=Homo sapiens OX=9606 GN=SEPTIN9 PE=1 SV=2 | 24 |
| Q9UHF5 | Interleukin-17B OS=Homo sapiens OX=9606 GN=IL17B PE=2 SV=1 | 8 |
| Q9UHG3 | Prenylcysteine oxidase 1 OS=Homo sapiens OX=9606 GN=PCYOX1 PE=1 SV=3 | 15 |
| Q9UHK6 | Alpha-methylacyl-CoA racemase OS=Homo sapiens OX=9606 GN=AMACR PE=1 SV=2 | 13 |
| Q9UHN1 | DNA polymerase subunit gamma-2, mitochondrial OS=Homo sapiens OX=9606 GN=POLG2 PE=1 SV=1 | 6 |
| Q9UHN6 | Cell surface hyaluronidase OS=Homo sapiens OX=9606 GN=CEMIP2 PE=1 SV=1 | 2 |
| Q9UHQ4 | B-cell receptor-associated protein 29 OS=Homo sapiens OX=9606 GN=BCAP29 PE=1 SV=2 | 3 |
| Q9UHR4 | Brain-specific angiogenesis inhibitor 1-associated protein 2-like protein 1 OS=Homo sapiens OX=9606 GN=BAIAP2L1 PE=1 SV=2 | 10 |
| Q9UHV9 | Prefoldin subunit 2 OS=Homo sapiens OX=9606 GN=PFDN2 PE=1 SV=1 | 6 |
| Q9UHY7 | Enolase-phosphatase E1 OS=Homo sapiens OX=9606 GN=ENOPH1 PE=1 SV=1 | 4 |
| Q9UI09 | NADH dehydrogenase [ubiquinone] 1 alpha subcomplex subunit 12 OS=Homo sapiens OX=9606 GN=NDUFA12 PE=1 SV=1 | 2 |
| Q9UI12 | V-type proton ATPase subunit H OS=Homo sapiens OX=9606 GN=ATP6V1H PE=1 SV=1 | 14 |
| Q9UI14 | Prenylated Rab acceptor protein 1 OS=Homo sapiens OX=9606 GN=RABAC1 PE=1 SV=1 | 1 |
| Q9UI17 | Dimethylglycine dehydrogenase, mitochondrial OS=Homo sapiens OX=9606 GN=DMGDH PE=1 SV=2 | 2 |
| Q9UI30 | Multifunctional methyltransferase subunit TRM112-like protein OS=Homo sapiens OX=9606 GN=TRMT112 PE=1 SV=1 | 6 |
| Q9UI43 | rRNA methyltransferase 2, mitochondrial OS=Homo sapiens OX=9606 GN=MRM2 PE=1 SV=1 | 2 |
| Q9UIA9 | Exportin-7 OS=Homo sapiens OX=9606 GN=XPO7 PE=1 SV=3 | 6 |
| Q9UIC8 | Leucine carboxyl methyltransferase 1 OS=Homo sapiens OX=9606 GN=LCMT1 PE=1 SV=2 | 18 |
| Q9UII2 | ATPase inhibitor, mitochondrial OS=Homo sapiens OX=9606 GN=ATP5IF1 PE=1 SV=1 | 6 |
| Q9UIJ7 | GTP:AMP phosphotransferase AK3, mitochondrial OS=Homo sapiens OX=9606 GN=AK3 PE=1 SV=4 | 12 |
| Q9UIQ6 | Leucyl-cystinyl aminopeptidase OS=Homo sapiens OX=9606 GN=LNPEP PE=1 SV=3 | 8 |
| Q9UIV1 | CCR4-NOT transcription complex subunit 7 OS=Homo sapiens OX=9606 GN=CNOT7 PE=1 SV=3 | 1 |
| Q9UIV8 | Serpin B13 OS=Homo sapiens OX=9606 GN=SERPINB13 PE=1 SV=2 | 10 |
| Q9UJ55 | MAGE-like protein 2 OS=Homo sapiens OX=9606 GN=MAGEL2 PE=1 SV=2 | 1 |
| Q9UJ70 | N-acetyl-D-glucosamine kinase OS=Homo sapiens OX=9606 GN=NAGK PE=1 SV=4 | 9 |
| Q9UJ72 | Annexin A10 OS=Homo sapiens OX=9606 GN=ANXA10 PE=1 SV=3 | 1 |
| Q9UJ83 | 2-hydroxyacyl-CoA lyase 1 OS=Homo sapiens OX=9606 GN=HACL1 PE=1 SV=2 | 7 |
| Q9UJA5 | tRNA (adenine(58)-N(1))-methyltransferase non-catalytic subunit TRM6 OS=Homo sapiens OX=9606 GN=TRMT6 PE=1 SV=1 | 1 |
| Q9UJC3 | Protein Hook homolog 1 OS=Homo sapiens OX=9606 GN=HOOK1 PE=1 SV=2 | 19 |
| Q9UJC5 | SH3 domain-binding glutamic acid-rich-like protein 2 OS=Homo sapiens OX=9606 GN=SH3BGRL2 PE=1 SV=2 | 2 |
| Q9UJS0 | Electrogenic aspartate/glutamate antiporter SLC25A13, mitochondrial OS=Homo sapiens OX=9606 GN=SLC25A13 PE=1 SV=2 | 13 |
| Q9UJU6 | Drebrin-like protein OS=Homo sapiens OX=9606 GN=DBNL PE=1 SV=1 | 3 |
| Q9UJW0 | Dynactin subunit 4 OS=Homo sapiens OX=9606 GN=DCTN4 PE=1 SV=1 | 8 |
| Q9UJX2 | Cell division cycle protein 23 homolog OS=Homo sapiens OX=9606 GN=CDC23 PE=1 SV=3 | 1 |
| Q9UJX3 | Anaphase-promoting complex subunit 7 OS=Homo sapiens OX=9606 GN=ANAPC7 PE=1 SV=5 | 1 |
| Q9UJZ1 | Stomatin-like protein 2, mitochondrial OS=Homo sapiens OX=9606 GN=STOML2 PE=1 SV=1 | 17 |
| Q9UK22 | F-box only protein 2 OS=Homo sapiens OX=9606 GN=FBXO2 PE=1 SV=2 | 10 |
| Q9UK41 | Vacuolar protein sorting-associated protein 28 homolog OS=Homo sapiens OX=9606 GN=VPS28 PE=1 SV=1 | 6 |
| Q9UK45 | U6 snRNA-associated Sm-like protein LSm7 OS=Homo sapiens OX=9606 GN=LSM7 PE=1 SV=1 | 5 |
| Q9UK61 | Protein TASOR OS=Homo sapiens OX=9606 GN=TASOR PE=1 SV=3 | 1 |
| Q9UK76 | Jupiter microtubule associated homolog 1 OS=Homo sapiens OX=9606 GN=JPT1 PE=1 SV=3 | 2 |
| Q9UKA1 | F-box/LRR-repeat protein 5 OS=Homo sapiens OX=9606 GN=FBXL5 PE=1 SV=2 | 1 |
| Q9UKA9 | Polypyrimidine tract-binding protein 2 OS=Homo sapiens OX=9606 GN=PTBP2 PE=1 SV=1 | 8 |
| Q9UKF6 | Cleavage and polyadenylation specificity factor subunit 3 OS=Homo sapiens OX=9606 GN=CPSF3 PE=1 SV=1 | 7 |
| Q9UKG1 | DCC-interacting protein 13-alpha OS=Homo sapiens OX=9606 GN=APPL1 PE=1 SV=1 | 19 |
| Q9UKK3 | Protein mono-ADP-ribosyltransferase PARP4 OS=Homo sapiens OX=9606 GN=PARP4 PE=1 SV=3 | 2 |
| Q9UKL6 | Phosphatidylcholine transfer protein OS=Homo sapiens OX=9606 GN=PCTP PE=1 SV=1 | 1 |
| Q9UKN7 | Unconventional myosin-XV OS=Homo sapiens OX=9606 GN=MYO15A PE=1 SV=2 | 1 |
| Q9UKU7 | Isobutyryl-CoA dehydrogenase, mitochondrial OS=Homo sapiens OX=9606 GN=ACAD8 PE=1 SV=1 | 14 |
| Q9UKX2 | Myosin-2 OS=Homo sapiens OX=9606 GN=MYH2 PE=1 SV=1 | 2 |
| Q9UKX3 | Myosin-13 OS=Homo sapiens OX=9606 GN=MYH13 PE=2 SV=2 | 2 |
| Q9UKY7 | Protein CDV3 homolog OS=Homo sapiens OX=9606 GN=CDV3 PE=1 SV=1 | 7 |
| Q9UKZ1 | CCR4-NOT transcription complex subunit 11 OS=Homo sapiens OX=9606 GN=CNOT11 PE=1 SV=1 | 4 |
| Q9UL25 | Ras-related protein Rab-21 OS=Homo sapiens OX=9606 GN=RAB21 PE=1 SV=3 | 7 |
| Q9UL26 | Ras-related protein Rab-22A OS=Homo sapiens OX=9606 GN=RAB22A PE=1 SV=2 | 10 |
| Q9UL33 | Trafficking protein particle complex subunit 2-like protein OS=Homo sapiens OX=9606 GN=TRAPPC2L PE=1 SV=1 | 4 |
| Q9UL54 | Serine/threonine-protein kinase TAO2 OS=Homo sapiens OX=9606 GN=TAOK2 PE=1 SV=2 | 3 |
| Q9ULA0 | Aspartyl aminopeptidase OS=Homo sapiens OX=9606 GN=DNPEP PE=1 SV=2 | 10 |
| Q9ULC4 | Malignant T-cell-amplified sequence 1 OS=Homo sapiens OX=9606 GN=MCTS1 PE=1 SV=1 | 5 |
| Q9ULD0 | 2-oxoglutarate dehydrogenase-like, mitochondrial OS=Homo sapiens OX=9606 GN=OGDHL PE=1 SV=3 | 19 |
| Q9ULE4 | Protein FAM184B OS=Homo sapiens OX=9606 GN=FAM184B PE=2 SV=3 | 2 |
| Q9ULF5 | Zinc transporter ZIP10 OS=Homo sapiens OX=9606 GN=SLC39A10 PE=1 SV=2 | 1 |
| Q9ULJ8 | Neurabin-1 OS=Homo sapiens OX=9606 GN=PPP1R9A PE=1 SV=2 | 1 |
| Q9ULP0 | Protein NDRG4 OS=Homo sapiens OX=9606 GN=NDRG4 PE=1 SV=2 | 2 |
| Q9ULR3 | Protein phosphatase 1H OS=Homo sapiens OX=9606 GN=PPM1H PE=1 SV=2 | 6 |
| Q9ULR5 | Polyadenylate-binding protein-interacting protein 2B OS=Homo sapiens OX=9606 GN=PAIP2B PE=1 SV=2 | 3 |
| Q9ULT8 | E3 ubiquitin-protein ligase HECTD1 OS=Homo sapiens OX=9606 GN=HECTD1 PE=1 SV=4 | 8 |
| Q9ULV0 | Unconventional myosin-Vb OS=Homo sapiens OX=9606 GN=MYO5B PE=1 SV=3 | 12 |
| Q9ULV4 | Coronin-1C OS=Homo sapiens OX=9606 GN=CORO1C PE=1 SV=1 | 5 |
| Q9UM00 | Calcium load-activated calcium channel OS=Homo sapiens OX=9606 GN=TMCO1 PE=1 SV=2 | 2 |
| Q9UM44 | HERV-H LTR-associating protein 2 OS=Homo sapiens OX=9606 GN=HHLA2 PE=1 SV=1 | 1 |
| Q9UM54 | Unconventional myosin-VI OS=Homo sapiens OX=9606 GN=MYO6 PE=1 SV=4 | 35 |
| Q9UMS0 | NFU1 iron-sulfur cluster scaffold homolog, mitochondrial OS=Homo sapiens OX=9606 GN=NFU1 PE=1 SV=2 | 6 |
| Q9UMX0 | Ubiquilin-1 OS=Homo sapiens OX=9606 GN=UBQLN1 PE=1 SV=2 | 6 |
| Q9UMX5 | Neudesin OS=Homo sapiens OX=9606 GN=NENF PE=1 SV=1 | 9 |
| Q9UMY4 | Sorting nexin-12 OS=Homo sapiens OX=9606 GN=SNX12 PE=1 SV=4 | 13 |
| Q9UMZ2 | Synergin gamma OS=Homo sapiens OX=9606 GN=SYNRG PE=1 SV=2 | 1 |
| Q9UN37 | Vacuolar protein sorting-associated protein 4A OS=Homo sapiens OX=9606 GN=VPS4A PE=1 SV=1 | 15 |
| Q9UN86 | Ras GTPase-activating protein-binding protein 2 OS=Homo sapiens OX=9606 GN=G3BP2 PE=1 SV=2 | 12 |
| Q9UNE7 | E3 ubiquitin-protein ligase CHIP OS=Homo sapiens OX=9606 GN=STUB1 PE=1 SV=2 | 16 |
| Q9UNF0 | Protein kinase C and casein kinase substrate in neurons protein 2 OS=Homo sapiens OX=9606 GN=PACSIN2 PE=1 SV=2 | 22 |
| Q9UNF1 | Melanoma-associated antigen D2 OS=Homo sapiens OX=9606 GN=MAGED2 PE=1 SV=2 | 2 |
| Q9UNH7 | Sorting nexin-6 OS=Homo sapiens OX=9606 GN=SNX6 PE=1 SV=1 | 1 |
| Q9UNK0 | Syntaxin-8 OS=Homo sapiens OX=9606 GN=STX8 PE=1 SV=2 | 1 |
| Q9UNL2 | Translocon-associated protein subunit gamma OS=Homo sapiens OX=9606 GN=SSR3 PE=1 SV=1 | 1 |
| Q9UNM6 | 26S proteasome non-ATPase regulatory subunit 13 OS=Homo sapiens OX=9606 GN=PSMD13 PE=1 SV=2 | 18 |
| Q9UNN4 | TFIIA-alpha and beta-like factor OS=Homo sapiens OX=9606 GN=GTF2A1L PE=1 SV=2 | 2 |
| Q9UNN5 | FAS-associated factor 1 OS=Homo sapiens OX=9606 GN=FAF1 PE=1 SV=2 | 4 |
| Q9UNS2 | COP9 signalosome complex subunit 3 OS=Homo sapiens OX=9606 GN=COPS3 PE=1 SV=3 | 17 |
| Q9UNX3 | 60S ribosomal protein L26-like 1 OS=Homo sapiens OX=9606 GN=RPL26L1 PE=1 SV=1 | 1 |
| Q9UNZ2 | NSFL1 cofactor p47 OS=Homo sapiens OX=9606 GN=NSFL1C PE=1 SV=2 | 16 |
| Q9UPN3 | Microtubule-actin cross-linking factor 1, isoforms 1/2/3/4/5 OS=Homo sapiens OX=9606 GN=MACF1 PE=1 SV=4 | 19 |
| Q9UPN7 | Serine/threonine-protein phosphatase 6 regulatory subunit 1 OS=Homo sapiens OX=9606 GN=PPP6R1 PE=1 SV=5 | 2 |
| Q9UPQ9 | Trinucleotide repeat-containing gene 6B protein OS=Homo sapiens OX=9606 GN=TNRC6B PE=1 SV=4 | 6 |
| Q9UPT5 | Exocyst complex component 7 OS=Homo sapiens OX=9606 GN=EXOC7 PE=1 SV=3 | 10 |
| Q9UPY8 | Microtubule-associated protein RP/EB family member 3 OS=Homo sapiens OX=9606 GN=MAPRE3 PE=1 SV=1 | 8 |
| Q9UQ13 | Leucine-rich repeat protein SHOC-2 OS=Homo sapiens OX=9606 GN=SHOC2 PE=1 SV=2 | 13 |
| Q9UQ35 | Serine/arginine repetitive matrix protein 2 OS=Homo sapiens OX=9606 GN=SRRM2 PE=1 SV=2 | 3 |
| Q9UQ80 | Proliferation-associated protein 2G4 OS=Homo sapiens OX=9606 GN=PA2G4 PE=1 SV=3 | 14 |
| Q9UQ90 | Paraplegin OS=Homo sapiens OX=9606 GN=SPG7 PE=1 SV=2 | 3 |
| Q9UQB8 | Brain-specific angiogenesis inhibitor 1-associated protein 2 OS=Homo sapiens OX=9606 GN=BAIAP2 PE=1 SV=1 | 9 |
| Q9UQC2 | GRB2-associated-binding protein 2 OS=Homo sapiens OX=9606 GN=GAB2 PE=1 SV=1 | 1 |
| Q9UQN3 | Charged multivesicular body protein 2b OS=Homo sapiens OX=9606 GN=CHMP2B PE=1 SV=1 | 4 |
| Q9Y217 | Myotubularin-related protein 6 OS=Homo sapiens OX=9606 GN=MTMR6 PE=1 SV=3 | 3 |
| Q9Y224 | RNA transcription, translation and transport factor protein OS=Homo sapiens OX=9606 GN=RTRAF PE=1 SV=1 | 7 |
| Q9Y230 | RuvB-like 2 OS=Homo sapiens OX=9606 GN=RUVBL2 PE=1 SV=3 | 29 |
| Q9Y237 | Peptidyl-prolyl cis-trans isomerase NIMA-interacting 4 OS=Homo sapiens OX=9606 GN=PIN4 PE=1 SV=1 | 4 |
| Q9Y248 | DNA replication complex GINS protein PSF2 OS=Homo sapiens OX=9606 GN=GINS2 PE=1 SV=1 | 1 |
| Q9Y262 | Eukaryotic translation initiation factor 3 subunit L OS=Homo sapiens OX=9606 GN=EIF3L PE=1 SV=1 | 11 |
| Q9Y263 | Phospholipase A-2-activating protein OS=Homo sapiens OX=9606 GN=PLAA PE=1 SV=2 | 22 |
| Q9Y265 | RuvB-like 1 OS=Homo sapiens OX=9606 GN=RUVBL1 PE=1 SV=1 | 14 |
| Q9Y276 | Mitochondrial chaperone BCS1 OS=Homo sapiens OX=9606 GN=BCS1L PE=1 SV=1 | 3 |
| Q9Y277 | Voltage-dependent anion-selective channel protein 3 OS=Homo sapiens OX=9606 GN=VDAC3 PE=1 SV=1 | 13 |
| Q9Y281 | Cofilin-2 OS=Homo sapiens OX=9606 GN=CFL2 PE=1 SV=1 | 8 |
| Q9Y282 | Endoplasmic reticulum-Golgi intermediate compartment protein 3 OS=Homo sapiens OX=9606 GN=ERGIC3 PE=1 SV=1 | 5 |
| Q9Y285 | Phenylalanine--tRNA ligase alpha subunit OS=Homo sapiens OX=9606 GN=FARSA PE=1 SV=3 | 10 |
| Q9Y294 | Histone chaperone ASF1A OS=Homo sapiens OX=9606 GN=ASF1A PE=1 SV=1 | 3 |
| Q9Y295 | Developmentally-regulated GTP-binding protein 1 OS=Homo sapiens OX=9606 GN=DRG1 PE=1 SV=1 | 6 |
| Q9Y296 | Trafficking protein particle complex subunit 4 OS=Homo sapiens OX=9606 GN=TRAPPC4 PE=1 SV=1 | 6 |
| Q9Y2A7 | Nck-associated protein 1 OS=Homo sapiens OX=9606 GN=NCKAP1 PE=1 SV=1 | 2 |
| Q9Y2B0 | Protein canopy homolog 2 OS=Homo sapiens OX=9606 GN=CNPY2 PE=1 SV=1 | 3 |
| Q9Y2H1 | Serine/threonine-protein kinase 38-like OS=Homo sapiens OX=9606 GN=STK38L PE=1 SV=3 | 8 |
| Q9Y2I8 | WD repeat-containing protein 37 OS=Homo sapiens OX=9606 GN=WDR37 PE=1 SV=2 | 2 |
| Q9Y2J0 | Rabphilin-3A OS=Homo sapiens OX=9606 GN=RPH3A PE=1 SV=1 | 7 |
| Q9Y2J4 | Angiomotin-like protein 2 OS=Homo sapiens OX=9606 GN=AMOTL2 PE=1 SV=3 | 1 |
| Q9Y2L5 | Trafficking protein particle complex subunit 8 OS=Homo sapiens OX=9606 GN=TRAPPC8 PE=1 SV=2 | 7 |
| Q9Y2L9 | Leucine-rich repeat and calponin homology domain-containing protein 1 OS=Homo sapiens OX=9606 GN=LRCH1 PE=1 SV=3 | 5 |
| Q9Y2P4 | Long-chain fatty acid transport protein 6 OS=Homo sapiens OX=9606 GN=SLC27A6 PE=2 SV=1 | 4 |
| Q9Y2Q3 | Glutathione S-transferase kappa 1 OS=Homo sapiens OX=9606 GN=GSTK1 PE=1 SV=3 | 10 |
| Q9Y2Q5 | Ragulator complex protein LAMTOR2 OS=Homo sapiens OX=9606 GN=LAMTOR2 PE=1 SV=1 | 4 |
| Q9Y2Q9 | 28S ribosomal protein S28, mitochondrial OS=Homo sapiens OX=9606 GN=MRPS28 PE=1 SV=1 | 3 |
| Q9Y2R0 | Cytochrome c oxidase assembly factor 3 homolog, mitochondrial OS=Homo sapiens OX=9606 GN=COA3 PE=1 SV=1 | 3 |
| Q9Y2R5 | 28S ribosomal protein S17, mitochondrial OS=Homo sapiens OX=9606 GN=MRPS17 PE=1 SV=1 | 2 |
| Q9Y2R9 | 28S ribosomal protein S7, mitochondrial OS=Homo sapiens OX=9606 GN=MRPS7 PE=1 SV=2 | 4 |
| Q9Y2S7 | Polymerase delta-interacting protein 2 OS=Homo sapiens OX=9606 GN=POLDIP2 PE=1 SV=1 | 11 |
| Q9Y2T2 | AP-3 complex subunit mu-1 OS=Homo sapiens OX=9606 GN=AP3M1 PE=1 SV=1 | 9 |
| Q9Y2V2 | Calcium-regulated heat-stable protein 1 OS=Homo sapiens OX=9606 GN=CARHSP1 PE=1 SV=2 | 4 |
| Q9Y2W6 | Tudor and KH domain-containing protein OS=Homo sapiens OX=9606 GN=TDRKH PE=1 SV=2 | 20 |
| Q9Y2X7 | ARF GTPase-activating protein GIT1 OS=Homo sapiens OX=9606 GN=GIT1 PE=1 SV=2 | 1 |
| Q9Y2Z0 | Protein SGT1 homolog OS=Homo sapiens OX=9606 GN=SUGT1 PE=1 SV=3 | 28 |
| Q9Y2Z2 | Protein MTO1 homolog, mitochondrial OS=Homo sapiens OX=9606 GN=MTO1 PE=1 SV=2 | 2 |
| Q9Y2Z4 | Tyrosine--tRNA ligase, mitochondrial OS=Homo sapiens OX=9606 GN=YARS2 PE=1 SV=2 | 17 |
| Q9Y2Z9 | Ubiquinone biosynthesis monooxygenase COQ6, mitochondrial OS=Homo sapiens OX=9606 GN=COQ6 PE=1 SV=2 | 6 |
| Q9Y303 | N-acetylglucosamine-6-phosphate deacetylase OS=Homo sapiens OX=9606 GN=AMDHD2 PE=1 SV=2 | 4 |
| Q9Y305 | Acyl-coenzyme A thioesterase 9, mitochondrial OS=Homo sapiens OX=9606 GN=ACOT9 PE=1 SV=2 | 30 |
| Q9Y312 | Protein AAR2 homolog OS=Homo sapiens OX=9606 GN=AAR2 PE=1 SV=2 | 1 |
| Q9Y315 | Deoxyribose-phosphate aldolase OS=Homo sapiens OX=9606 GN=DERA PE=1 SV=2 | 1 |
| Q9Y316 | Protein MEMO1 OS=Homo sapiens OX=9606 GN=MEMO1 PE=1 SV=1 | 2 |
| Q9Y333 | U6 snRNA-associated Sm-like protein LSm2 OS=Homo sapiens OX=9606 GN=LSM2 PE=1 SV=1 | 8 |
| Q9Y337 | Kallikrein-5 OS=Homo sapiens OX=9606 GN=KLK5 PE=1 SV=3 | 3 |
| Q9Y371 | Endophilin-B1 OS=Homo sapiens OX=9606 GN=SH3GLB1 PE=1 SV=1 | 13 |
| Q9Y375 | Complex I intermediate-associated protein 30, mitochondrial OS=Homo sapiens OX=9606 GN=NDUFAF1 PE=1 SV=2 | 6 |
| Q9Y376 | Calcium-binding protein 39 OS=Homo sapiens OX=9606 GN=CAB39 PE=1 SV=1 | 13 |
| Q9Y383 | Putative RNA-binding protein Luc7-like 2 OS=Homo sapiens OX=9606 GN=LUC7L2 PE=1 SV=2 | 3 |
| Q9Y3A6 | Transmembrane emp24 domain-containing protein 5 OS=Homo sapiens OX=9606 GN=TMED5 PE=1 SV=1 | 3 |
| Q9Y3B3 | Transmembrane emp24 domain-containing protein 7 OS=Homo sapiens OX=9606 GN=TMED7 PE=1 SV=2 | 4 |
| Q9Y3B4 | Splicing factor 3B subunit 6 OS=Homo sapiens OX=9606 GN=SF3B6 PE=1 SV=1 | 2 |
| Q9Y3B8 | Oligoribonuclease, mitochondrial OS=Homo sapiens OX=9606 GN=REXO2 PE=1 SV=3 | 11 |
| Q9Y3C0 | WASH complex subunit 3 OS=Homo sapiens OX=9606 GN=WASHC3 PE=1 SV=1 | 2 |
| Q9Y3C6 | Peptidyl-prolyl cis-trans isomerase-like 1 OS=Homo sapiens OX=9606 GN=PPIL1 PE=1 SV=1 | 4 |
| Q9Y3C8 | Ubiquitin-fold modifier-conjugating enzyme 1 OS=Homo sapiens OX=9606 GN=UFC1 PE=1 SV=3 | 12 |
| Q9Y3D0 | Cytosolic iron-sulfur assembly component 2B OS=Homo sapiens OX=9606 GN=CIAO2B PE=1 SV=1 | 3 |
| Q9Y3D3 | 28S ribosomal protein S16, mitochondrial OS=Homo sapiens OX=9606 GN=MRPS16 PE=1 SV=1 | 2 |
| Q9Y3D6 | Mitochondrial fission 1 protein OS=Homo sapiens OX=9606 GN=FIS1 PE=1 SV=2 | 4 |
| Q9Y3D7 | Mitochondrial import inner membrane translocase subunit TIM16 OS=Homo sapiens OX=9606 GN=PAM16 PE=1 SV=2 | 5 |
| Q9Y3D9 | 28S ribosomal protein S23, mitochondrial OS=Homo sapiens OX=9606 GN=MRPS23 PE=1 SV=2 | 2 |
| Q9Y3E7 | Charged multivesicular body protein 3 OS=Homo sapiens OX=9606 GN=CHMP3 PE=1 SV=3 | 5 |
| Q9Y3F4 | Serine-threonine kinase receptor-associated protein OS=Homo sapiens OX=9606 GN=STRAP PE=1 SV=1 | 15 |
| Q9Y3I0 | RNA-splicing ligase RtcB homolog OS=Homo sapiens OX=9606 GN=RTCB PE=1 SV=1 | 24 |
| Q9Y3I1 | F-box only protein 7 OS=Homo sapiens OX=9606 GN=FBXO7 PE=1 SV=1 | 1 |
| Q9Y3P9 | Rab GTPase-activating protein 1 OS=Homo sapiens OX=9606 GN=RABGAP1 PE=1 SV=3 | 17 |
| Q9Y3Q3 | Transmembrane emp24 domain-containing protein 3 OS=Homo sapiens OX=9606 GN=TMED3 PE=1 SV=1 | 3 |
| Q9Y3U8 | 60S ribosomal protein L36 OS=Homo sapiens OX=9606 GN=RPL36 PE=1 SV=3 | 2 |
| Q9Y3Z3 | Deoxynucleoside triphosphate triphosphohydrolase SAMHD1 OS=Homo sapiens OX=9606 GN=SAMHD1 PE=1 SV=2 | 4 |
| Q9Y446 | Plakophilin-3 OS=Homo sapiens OX=9606 GN=PKP3 PE=1 SV=1 | 1 |
| Q9Y450 | HBS1-like protein OS=Homo sapiens OX=9606 GN=HBS1L PE=1 SV=1 | 22 |
| Q9Y478 | 5'-AMP-activated protein kinase subunit beta-1 OS=Homo sapiens OX=9606 GN=PRKAB1 PE=1 SV=4 | 1 |
| Q9Y490 | Talin-1 OS=Homo sapiens OX=9606 GN=TLN1 PE=1 SV=3 | 39 |
| Q9Y496 | Kinesin-like protein KIF3A OS=Homo sapiens OX=9606 GN=KIF3A PE=1 SV=4 | 4 |
| Q9Y4B6 | DDB1- and CUL4-associated factor 1 OS=Homo sapiens OX=9606 GN=DCAF1 PE=1 SV=3 | 1 |
| Q9Y4D1 | Disheveled-associated activator of morphogenesis 1 OS=Homo sapiens OX=9606 GN=DAAM1 PE=1 SV=2 | 1 |
| Q9Y4E8 | Ubiquitin carboxyl-terminal hydrolase 15 OS=Homo sapiens OX=9606 GN=USP15 PE=1 SV=3 | 22 |
| Q9Y4G6 | Talin-2 OS=Homo sapiens OX=9606 GN=TLN2 PE=1 SV=4 | 22 |
| Q9Y4I1 | Unconventional myosin-Va OS=Homo sapiens OX=9606 GN=MYO5A PE=1 SV=2 | 27 |
| Q9Y4K0 | Lysyl oxidase homolog 2 OS=Homo sapiens OX=9606 GN=LOXL2 PE=1 SV=1 | 14 |
| Q9Y4K1 | Beta/gamma crystallin domain-containing protein 1 OS=Homo sapiens OX=9606 GN=CRYBG1 PE=1 SV=3 | 9 |
| Q9Y4L1 | Hypoxia up-regulated protein 1 OS=Homo sapiens OX=9606 GN=HYOU1 PE=1 SV=1 | 44 |
| Q9Y4P1 | Cysteine protease ATG4B OS=Homo sapiens OX=9606 GN=ATG4B PE=1 SV=2 | 2 |
| Q9Y4P8 | WD repeat domain phosphoinositide-interacting protein 2 OS=Homo sapiens OX=9606 GN=WIPI2 PE=1 SV=1 | 4 |
| Q9Y4X5 | E3 ubiquitin-protein ligase ARIH1 OS=Homo sapiens OX=9606 GN=ARIH1 PE=1 SV=2 | 4 |
| Q9Y4Z0 | U6 snRNA-associated Sm-like protein LSm4 OS=Homo sapiens OX=9606 GN=LSM4 PE=1 SV=1 | 8 |
| Q9Y508 | E3 ubiquitin-protein ligase RNF114 OS=Homo sapiens OX=9606 GN=RNF114 PE=1 SV=1 | 8 |
| Q9Y512 | Sorting and assembly machinery component 50 homolog OS=Homo sapiens OX=9606 GN=SAMM50 PE=1 SV=3 | 16 |
| Q9Y530 | ADP-ribose glycohydrolase OARD1 OS=Homo sapiens OX=9606 GN=OARD1 PE=1 SV=2 | 6 |
| Q9Y547 | Intraflagellar transport protein 25 homolog OS=Homo sapiens OX=9606 GN=HSPB11 PE=1 SV=1 | 3 |
| Q9Y570 | Protein phosphatase methylesterase 1 OS=Homo sapiens OX=9606 GN=PPME1 PE=1 SV=3 | 8 |
| Q9Y584 | Mitochondrial import inner membrane translocase subunit Tim22 OS=Homo sapiens OX=9606 GN=TIMM22 PE=1 SV=2 | 1 |
| Q9Y5B9 | FACT complex subunit SPT16 OS=Homo sapiens OX=9606 GN=SUPT16H PE=1 SV=1 | 2 |
| Q9Y5J3 | Hairy/enhancer-of-split related with YRPW motif protein 1 OS=Homo sapiens OX=9606 GN=HEY1 PE=1 SV=1 | 1 |
| Q9Y5J7 | Mitochondrial import inner membrane translocase subunit Tim9 OS=Homo sapiens OX=9606 GN=TIMM9 PE=1 SV=1 | 4 |
| Q9Y5K8 | V-type proton ATPase subunit D OS=Homo sapiens OX=9606 GN=ATP6V1D PE=1 SV=1 | 3 |
| Q9Y5L0 | Transportin-3 OS=Homo sapiens OX=9606 GN=TNPO3 PE=1 SV=3 | 1 |
| Q9Y5L4 | Mitochondrial import inner membrane translocase subunit Tim13 OS=Homo sapiens OX=9606 GN=TIMM13 PE=1 SV=1 | 5 |
| Q9Y5M8 | Signal recognition particle receptor subunit beta OS=Homo sapiens OX=9606 GN=SRPRB PE=1 SV=3 | 9 |
| Q9Y5P4 | Ceramide transfer protein OS=Homo sapiens OX=9606 GN=CERT1 PE=1 SV=1 | 7 |
| Q9Y5P6 | Mannose-1-phosphate guanyltransferase beta OS=Homo sapiens OX=9606 GN=GMPPB PE=1 SV=2 | 7 |
| Q9Y5R8 | Trafficking protein particle complex subunit 1 OS=Homo sapiens OX=9606 GN=TRAPPC1 PE=1 SV=1 | 4 |
| Q9Y5S9 | RNA-binding protein 8A OS=Homo sapiens OX=9606 GN=RBM8A PE=1 SV=1 | 3 |
| Q9Y5T4 | DnaJ homolog subfamily C member 15 OS=Homo sapiens OX=9606 GN=DNAJC15 PE=1 SV=2 | 5 |
| Q9Y5W9 | Sorting nexin-11 OS=Homo sapiens OX=9606 GN=SNX11 PE=1 SV=2 | 4 |
| Q9Y5X1 | Sorting nexin-9 OS=Homo sapiens OX=9606 GN=SNX9 PE=1 SV=1 | 11 |
| Q9Y5X3 | Sorting nexin-5 OS=Homo sapiens OX=9606 GN=SNX5 PE=1 SV=1 | 12 |
| Q9Y5Y2 | Cytosolic Fe-S cluster assembly factor NUBP2 OS=Homo sapiens OX=9606 GN=NUBP2 PE=1 SV=1 | 1 |
| Q9Y5Z4 | Heme-binding protein 2 OS=Homo sapiens OX=9606 GN=HEBP2 PE=1 SV=1 | 4 |
| Q9Y605 | MORF4 family-associated protein 1 OS=Homo sapiens OX=9606 GN=MRFAP1 PE=1 SV=1 | 3 |
| Q9Y606 | Pseudouridylate synthase 1 homolog OS=Homo sapiens OX=9606 GN=PUS1 PE=1 SV=3 | 7 |
| Q9Y608 | Leucine-rich repeat flightless-interacting protein 2 OS=Homo sapiens OX=9606 GN=LRRFIP2 PE=1 SV=1 | 2 |
| Q9Y619 | Mitochondrial ornithine transporter 1 OS=Homo sapiens OX=9606 GN=SLC25A15 PE=1 SV=1 | 5 |
| Q9Y623 | Myosin-4 OS=Homo sapiens OX=9606 GN=MYH4 PE=2 SV=2 | 9 |
| Q9Y639 | Neuroplastin OS=Homo sapiens OX=9606 GN=NPTN PE=1 SV=2 | 4 |
| Q9Y646 | Carboxypeptidase Q OS=Homo sapiens OX=9606 GN=CPQ PE=1 SV=1 | 4 |
| Q9Y653 | Adhesion G-protein coupled receptor G1 OS=Homo sapiens OX=9606 GN=ADGRG1 PE=1 SV=2 | 4 |
| Q9Y678 | Coatomer subunit gamma-1 OS=Homo sapiens OX=9606 GN=COPG1 PE=1 SV=1 | 29 |
| Q9Y679 | Lipid droplet-regulating VLDL assembly factor AUP1 OS=Homo sapiens OX=9606 GN=AUP1 PE=1 SV=2 | 4 |
| Q9Y696 | Chloride intracellular channel protein 4 OS=Homo sapiens OX=9606 GN=CLIC4 PE=1 SV=4 | 13 |
| Q9Y697 | Cysteine desulfurase OS=Homo sapiens OX=9606 GN=NFS1 PE=1 SV=3 | 18 |
| Q9Y6A5 | Transforming acidic coiled-coil-containing protein 3 OS=Homo sapiens OX=9606 GN=TACC3 PE=1 SV=1 | 20 |
| Q9Y6A9 | Signal peptidase complex subunit 1 OS=Homo sapiens OX=9606 GN=SPCS1 PE=1 SV=5 | 1 |
| Q9Y6C9 | Mitochondrial carrier homolog 2 OS=Homo sapiens OX=9606 GN=MTCH2 PE=1 SV=1 | 8 |
| Q9Y6D6 | Brefeldin A-inhibited guanine nucleotide-exchange protein 1 OS=Homo sapiens OX=9606 GN=ARFGEF1 PE=1 SV=2 | 3 |
| Q9Y6E0 | Serine/threonine-protein kinase 24 OS=Homo sapiens OX=9606 GN=STK24 PE=1 SV=1 | 5 |
| Q9Y6F6 | Inositol 1,4,5-triphosphate receptor associated 1 OS=Homo sapiens OX=9606 GN=IRAG1 PE=1 SV=4 | 4 |
| Q9Y6G3 | 39S ribosomal protein L42, mitochondrial OS=Homo sapiens OX=9606 GN=MRPL42 PE=1 SV=1 | 1 |
| Q9Y6G5 | COMM domain-containing protein 10 OS=Homo sapiens OX=9606 GN=COMMD10 PE=1 SV=1 | 4 |
| Q9Y6G9 | Cytoplasmic dynein 1 light intermediate chain 1 OS=Homo sapiens OX=9606 GN=DYNC1LI1 PE=1 SV=3 | 21 |
| Q9Y6H1 | Coiled-coil-helix-coiled-coil-helix domain-containing protein 2 OS=Homo sapiens OX=9606 GN=CHCHD2 PE=1 SV=1 | 4 |
| Q9Y6I3 | Epsin-1 OS=Homo sapiens OX=9606 GN=EPN1 PE=1 SV=2 | 4 |
| Q9Y6I9 | Testis-expressed protein 264 OS=Homo sapiens OX=9606 GN=TEX264 PE=1 SV=1 | 3 |
| Q9Y6K9 | NF-kappa-B essential modulator OS=Homo sapiens OX=9606 GN=IKBKG PE=1 SV=2 | 3 |
| Q9Y6M1 | Insulin-like growth factor 2 mRNA-binding protein 2 OS=Homo sapiens OX=9606 GN=IGF2BP2 PE=1 SV=2 | 13 |
| Q9Y6M5 | Proton-coupled zinc antiporter SLC30A1 OS=Homo sapiens OX=9606 GN=SLC30A1 PE=1 SV=3 | 1 |
| Q9Y6M9 | NADH dehydrogenase [ubiquinone] 1 beta subcomplex subunit 9 OS=Homo sapiens OX=9606 GN=NDUFB9 PE=1 SV=3 | 5 |
| Q9Y6N5 | Sulfide:quinone oxidoreductase, mitochondrial OS=Homo sapiens OX=9606 GN=SQOR PE=1 SV=1 | 22 |
| Q9Y6R0 | Numb-like protein OS=Homo sapiens OX=9606 GN=NUMBL PE=1 SV=1 | 1 |
| Q9Y6Y8 | SEC23-interacting protein OS=Homo sapiens OX=9606 GN=SEC23IP PE=1 SV=1 | 17 |
| Y-FGCZCont00002;Y-FGCZCont00120;Y-FGCZCont00225 | zz_FGCZCont0001_P02534_K1M1_SHEEP blastpHomologue_0.0 | 1 |
| Y-FGCZCont00021 | zz_FGCZCont0020_P02770_ALBU_RAT blastpHomologue_0.0 | 3 |
| Y-FGCZCont00023;Y-FGCZCont00381 | zz_FGCZCont0022_P00711_LALBA_BOVIN blastpHomologue_3.0e-103 | 1 |
| Y-FGCZCont00034;Y-FGCZCont00039;Y-FGCZCont00174 | zz_FGCZCont0033_P04259_K2C6B_HUMAN blastpHomologue_0.0 | 2 |
| Y-FGCZCont00051 | zz_FGCZCont0050_P04264_K2C1_HUMAN blastpHomologue_0.0 | 2 |
| Y-FGCZCont00060;Y-FGCZCont00237 | zz_FGCZCont0059_P02538_K2C6A_HUMAN blastpHomologue_0.0 | 1 |
| Y-FGCZCont00061 | zz_FGCZCont0060_P35527_K1C9_HUMAN blastpHomologue_2.0e-07 | 1 |
| Y-FGCZCont00063 | zz_FGCZCont0062_P12035_K2C3_HUMAN blastpHomologue_0.0 | 2 |
| Y-FGCZCont00100;Y-FGCZCont00220 | zz_FGCZCont0099_P35527_K1C9_HUMAN blastpHomologue_0.0 | 1 |
| Y-FGCZCont00196 | zz_FGCZCont0195_P35527_K1C9_HUMAN blastpHomologue_2.0e-171 | 1 |
| Y-FGCZCont00223 | zz_FGCZCont0222_FGCZCont0222_FGCZCont0222 ambigous_blastpHomologue | 2 |
| Y-FGCZCont00293 | Q3SZ57 SWISS-PROT:Q3SZ57 (Bos taurus) Alpha-fetoprotein precursor | 1 |
| Y-FGCZCont00310 | Q1RMN8 TREMBL:Q1RMN8 (Bos taurus) Similar to Immunoglobulin lambda-like polypeptide 1 | 2 |
| Y-FGCZCont00351 | Q2UVX4 SWISS-PROT:Q2UVX4 (Bos taurus) Complement C3 precursor | 6 |
| Y-FGCZCont00354 | Q3SX09 TREMBL:Q3SX09 (Bos taurus) similar to HBG protein | 2 |
| Y-FGCZCont00359 | P15497 SWISS-PROT:P15497 (Bos taurus) Apolipoprotein A-I precursor | 2 |
| Y-FGCZCont00362 | P02070 SWISS-PROT:P02070 (Bos taurus) Hemoglobin subunit beta | 9 |
| Y-FGCZCont00373;Y-FGCZCont00383 | Q0IIK2 TREMBL:Q0IIK2 (Bos taurus) Transferrin | 5 |
| Y-FGCZCont00384 | P19001 SWISS-PROT:P19001 Tax_Id=10090 Gene_Symbol=Krt19 Keratin, type I cytoskeletal 19 | 2 |
| Y-FGCZCont00394 | Q8BGZ7 TREMBL:Q8BGZ7;Q99MH7 Tax_Id=10090 Gene_Symbol=Krt75 10 days neonate skin cDNA, RIKEN full-length enriched library, clone:4732475I03 product:CYTOKERATIN homolog | 2 |
| Y-FGCZCont00401 | Q6IFZ6 SWISS-PROT:Q6IFZ6 Tax_Id=10090 Gene_Symbol=Krt77 Keratin, type II cytoskeletal 1b | 2 |
| Y-FGCZCont00402 | Q6IFX2 SWISS-PROT:Q6IFX2 Tax_Id=10090 Gene_Symbol=Krt42 Keratin, type I cytoskeletal 42 | 1 |
| Y-FGCZCont00457 | ENSEMBL:ENSBTAP00000038253 (Bos taurus) 63 kDa protein | 1 |
| Y-FGCZCont00463 | ENSEMBL:ENSBTAP00000024462 (Bos taurus) 47 kDa protein | 3 |
| Y-FGCZCont00483 | sp_GAG_SCVLA_ | 1 |
| Y-FGCZCont00491 | sp_BGAL_ECOLI_ | 1 |
| A0A1W2PQ73 | ETS domain-containing transcription factor ERF-like OS=Homo sapiens OX=9606 GN=ERFL PE=3 SV=1 | 1 |
| A0AUZ9 | KAT8 regulatory NSL complex subunit 1-like protein OS=Homo sapiens OX=9606 GN=KANSL1L PE=1 SV=2 | 1 |
| A2RUB1 | Meiosis-specific coiled-coil domain-containing protein MEIOC OS=Homo sapiens OX=9606 GN=MEIOC PE=2 SV=3 | 1 |
| A6NE01 | Protein FAM186A OS=Homo sapiens OX=9606 GN=FAM186A PE=2 SV=3 | 1 |
| C9J202 | Putative glycosyltransferase ALG1L2 OS=Homo sapiens OX=9606 GN=ALG1L2 PE=3 SV=1 | 1 |
| O00194 | Ras-related protein Rab-27B OS=Homo sapiens OX=9606 GN=RAB27B PE=1 SV=4 | 2 |
| O00299 | Chloride intracellular channel protein 1 OS=Homo sapiens OX=9606 GN=CLIC1 PE=1 SV=4 | 4 |
| O00762 | Ubiquitin-conjugating enzyme E2 C OS=Homo sapiens OX=9606 GN=UBE2C PE=1 SV=1 | 5 |
| O14497 | AT-rich interactive domain-containing protein 1A OS=Homo sapiens OX=9606 GN=ARID1A PE=1 SV=3 | 1 |
| O14525 | Astrotactin-1 OS=Homo sapiens OX=9606 GN=ASTN1 PE=1 SV=3 | 1 |
| O14545 | TRAF-type zinc finger domain-containing protein 1 OS=Homo sapiens OX=9606 GN=TRAFD1 PE=1 SV=1 | 14 |
| O14653 | Golgi SNAP receptor complex member 2 OS=Homo sapiens OX=9606 GN=GOSR2 PE=1 SV=2 | 1 |
| O14764 | Gamma-aminobutyric acid receptor subunit delta OS=Homo sapiens OX=9606 GN=GABRD PE=1 SV=2 | 1 |
| O14776 | Transcription elongation regulator 1 OS=Homo sapiens OX=9606 GN=TCERG1 PE=1 SV=2 | 1 |
| O14867 | Transcription regulator protein BACH1 OS=Homo sapiens OX=9606 GN=BACH1 PE=1 SV=2 | 4 |
| O15198;P84022;Q15796 | Mothers against decapentaplegic homolog 9 OS=Homo sapiens OX=9606 GN=SMAD9 PE=1 SV=1 | 1 |
| O15234 | Protein CASC3 OS=Homo sapiens OX=9606 GN=CASC3 PE=1 SV=2 | 3 |
| O15439 | ATP-binding cassette sub-family C member 4 OS=Homo sapiens OX=9606 GN=ABCC4 PE=1 SV=3 | 2 |
| O15530 | 3-phosphoinositide-dependent protein kinase 1 OS=Homo sapiens OX=9606 GN=PDPK1 PE=1 SV=1 | 2 |
| O43291 | Kunitz-type protease inhibitor 2 OS=Homo sapiens OX=9606 GN=SPINT2 PE=1 SV=2 | 2 |
| O43301 | Heat shock 70 kDa protein 12A OS=Homo sapiens OX=9606 GN=HSPA12A PE=1 SV=2 | 2 |
| O43314 | Inositol hexakisphosphate and diphosphoinositol-pentakisphosphate kinase 2 OS=Homo sapiens OX=9606 GN=PPIP5K2 PE=1 SV=3 | 3 |
| O43464 | Serine protease HTRA2, mitochondrial OS=Homo sapiens OX=9606 GN=HTRA2 PE=1 SV=2 | 1 |
| O43586 | Proline-serine-threonine phosphatase-interacting protein 1 OS=Homo sapiens OX=9606 GN=PSTPIP1 PE=1 SV=1 | 1 |
| O43663 | Protein regulator of cytokinesis 1 OS=Homo sapiens OX=9606 GN=PRC1 PE=1 SV=2 | 8 |
| O43716 | Glutamyl-tRNA(Gln) amidotransferase subunit C, mitochondrial OS=Homo sapiens OX=9606 GN=GATC PE=1 SV=1 | 2 |
| O43766 | Lipoyl synthase, mitochondrial OS=Homo sapiens OX=9606 GN=LIAS PE=1 SV=3 | 1 |
| O43897 | Tolloid-like protein 1 OS=Homo sapiens OX=9606 GN=TLL1 PE=1 SV=1 | 1 |
| O60231 | Pre-mRNA-splicing factor ATP-dependent RNA helicase DHX16 OS=Homo sapiens OX=9606 GN=DHX16 PE=1 SV=2 | 1 |
| O60244 | Mediator of RNA polymerase II transcription subunit 14 OS=Homo sapiens OX=9606 GN=MED14 PE=1 SV=2 | 1 |
| O60264 | SWI/SNF-related matrix-associated actin-dependent regulator of chromatin subfamily A member 5 OS=Homo sapiens OX=9606 GN=SMARCA5 PE=1 SV=1 | 4 |
| O60271 | C-Jun-amino-terminal kinase-interacting protein 4 OS=Homo sapiens OX=9606 GN=SPAG9 PE=1 SV=4 | 2 |
| O60613 | Selenoprotein F OS=Homo sapiens OX=9606 GN=SELENOF PE=1 SV=4 | 1 |
| O60783 | 28S ribosomal protein S14, mitochondrial OS=Homo sapiens OX=9606 GN=MRPS14 PE=1 SV=1 | 1 |
| O60927 | E3 ubiquitin-protein ligase PPP1R11 OS=Homo sapiens OX=9606 GN=PPP1R11 PE=1 SV=1 | 1 |
| O75044;P0DJJ0;P0DMP2 | SLIT-ROBO Rho GTPase-activating protein 2 OS=Homo sapiens OX=9606 GN=SRGAP2 PE=1 SV=3 | 1 |
| O75122;Q7Z460 | CLIP-associating protein 2 OS=Homo sapiens OX=9606 GN=CLASP2 PE=1 SV=3 | 1 |
| O75446 | Histone deacetylase complex subunit SAP30 OS=Homo sapiens OX=9606 GN=SAP30 PE=1 SV=1 | 2 |
| O75449 | Katanin p60 ATPase-containing subunit A1 OS=Homo sapiens OX=9606 GN=KATNA1 PE=1 SV=1 | 2 |
| O75478 | Transcriptional adapter 2-alpha OS=Homo sapiens OX=9606 GN=TADA2A PE=1 SV=3 | 1 |
| O75496 | Geminin OS=Homo sapiens OX=9606 GN=GMNN PE=1 SV=1 | 1 |
| O75506 | Heat shock factor-binding protein 1 OS=Homo sapiens OX=9606 GN=HSBP1 PE=1 SV=1 | 3 |
| O75794 | Cell division cycle protein 123 homolog OS=Homo sapiens OX=9606 GN=CDC123 PE=1 SV=1 | 1 |
| O75891 | Cytosolic 10-formyltetrahydrofolate dehydrogenase OS=Homo sapiens OX=9606 GN=ALDH1L1 PE=1 SV=2 | 1 |
| O75937 | DnaJ homolog subfamily C member 8 OS=Homo sapiens OX=9606 GN=DNAJC8 PE=1 SV=2 | 1 |
| O75976 | Carboxypeptidase D OS=Homo sapiens OX=9606 GN=CPD PE=1 SV=2 | 1 |
| O76013 | Keratin, type I cuticular Ha6 OS=Homo sapiens OX=9606 GN=KRT36 PE=1 SV=1 | 5 |
| O94830 | Phospholipase DDHD2 OS=Homo sapiens OX=9606 GN=DDHD2 PE=1 SV=2 | 2 |
| O94851 | [F-actin]-monooxygenase MICAL2 OS=Homo sapiens OX=9606 GN=MICAL2 PE=1 SV=2 | 1 |
| O95049 | Tight junction protein ZO-3 OS=Homo sapiens OX=9606 GN=TJP3 PE=1 SV=3 | 1 |
| O95235 | Kinesin-like protein KIF20A OS=Homo sapiens OX=9606 GN=KIF20A PE=1 SV=1 | 3 |
| O95259 | Potassium voltage-gated channel subfamily H member 1 OS=Homo sapiens OX=9606 GN=KCNH1 PE=1 SV=1 | 1 |
| O95359 | Transforming acidic coiled-coil-containing protein 2 OS=Homo sapiens OX=9606 GN=TACC2 PE=1 SV=3 | 1 |
| O95486 | Protein transport protein Sec24A OS=Homo sapiens OX=9606 GN=SEC24A PE=1 SV=2 | 2 |
| O95563 | Mitochondrial pyruvate carrier 2 OS=Homo sapiens OX=9606 GN=MPC2 PE=1 SV=1 | 1 |
| O95833 | Chloride intracellular channel protein 3 OS=Homo sapiens OX=9606 GN=CLIC3 PE=1 SV=2 | 2 |
| P00395 | Cytochrome c oxidase subunit 1 OS=Homo sapiens OX=9606 GN=MT-CO1 PE=1 SV=1 | 1 |
| P00540 | Proto-oncogene serine/threonine-protein kinase mos OS=Homo sapiens OX=9606 GN=MOS PE=1 SV=1 | 1 |
| P00846 | ATP synthase subunit a OS=Homo sapiens OX=9606 GN=MT-ATP6 PE=1 SV=1 | 1 |
| P01100 | Protein c-Fos OS=Homo sapiens OX=9606 GN=FOS PE=1 SV=1 | 2 |
| P02795;P13640;P80297 | Metallothionein-2 OS=Homo sapiens OX=9606 GN=MT2A PE=1 SV=1 | 1 |
| P04350 | Tubulin beta-4A chain OS=Homo sapiens OX=9606 GN=TUBB4A PE=1 SV=2 | 1 |
| P06737 | Glycogen phosphorylase, liver form OS=Homo sapiens OX=9606 GN=PYGL PE=1 SV=4 | 1 |
| P07711 | Procathepsin L OS=Homo sapiens OX=9606 GN=CTSL PE=1 SV=2 | 2 |
| P08237 | ATP-dependent 6-phosphofructokinase, muscle type OS=Homo sapiens OX=9606 GN=PFKM PE=1 SV=2 | 15 |
| P08493 | Matrix Gla protein OS=Homo sapiens OX=9606 GN=MGP PE=1 SV=2 | 1 |
| P09661 | U2 small nuclear ribonucleoprotein A' OS=Homo sapiens OX=9606 GN=SNRPA1 PE=1 SV=2 | 1 |
| P09958 | Furin OS=Homo sapiens OX=9606 GN=FURIN PE=1 SV=2 | 1 |
| P0C221 | Coiled-coil domain-containing protein 175 OS=Homo sapiens OX=9606 GN=CCDC175 PE=4 SV=2 | 1 |
| P0DMM9;P0DMN0;P50225;P50226 | Sulfotransferase 1A3 OS=Homo sapiens OX=9606 GN=SULT1A3 PE=1 SV=1 | 1 |
| P0DP04 | Immunoglobulin heavy variable 3-43D OS=Homo sapiens OX=9606 GN=IGHV3-43D PE=3 SV=1 | 1 |
| P11166 | Solute carrier family 2, facilitated glucose transporter member 1 OS=Homo sapiens OX=9606 GN=SLC2A1 PE=1 SV=2 | 3 |
| P12532 | Creatine kinase U-type, mitochondrial OS=Homo sapiens OX=9606 GN=CKMT1B PE=1 SV=1 | 2 |
| P13051 | Uracil-DNA glycosylase OS=Homo sapiens OX=9606 GN=UNG PE=1 SV=2 | 1 |
| P13533 | Myosin-6 OS=Homo sapiens OX=9606 GN=MYH6 PE=1 SV=5 | 1 |
| P13726 | Tissue factor OS=Homo sapiens OX=9606 GN=F3 PE=1 SV=1 | 3 |
| P13807 | Glycogen [starch] synthase, muscle OS=Homo sapiens OX=9606 GN=GYS1 PE=1 SV=2 | 1 |
| P15085 | Carboxypeptidase A1 OS=Homo sapiens OX=9606 GN=CPA1 PE=1 SV=2 | 1 |
| P15954 | Cytochrome c oxidase subunit 7C, mitochondrial OS=Homo sapiens OX=9606 GN=COX7C PE=1 SV=1 | 1 |
| P16104;Q96QV6 | Histone H2AX OS=Homo sapiens OX=9606 GN=H2AX PE=1 SV=2 | 1 |
| P18754 | Regulator of chromosome condensation OS=Homo sapiens OX=9606 GN=RCC1 PE=1 SV=1 | 1 |
| P18858 | DNA ligase 1 OS=Homo sapiens OX=9606 GN=LIG1 PE=1 SV=1 | 2 |
| P19320 | Vascular cell adhesion protein 1 OS=Homo sapiens OX=9606 GN=VCAM1 PE=1 SV=1 | 1 |
| P22413 | Ectonucleotide pyrophosphatase/phosphodiesterase family member 1 OS=Homo sapiens OX=9606 GN=ENPP1 PE=1 SV=2 | 1 |
| P23352 | Anosmin-1 OS=Homo sapiens OX=9606 GN=ANOS1 PE=1 SV=3 | 4 |
| P23368 | NAD-dependent malic enzyme, mitochondrial OS=Homo sapiens OX=9606 GN=ME2 PE=1 SV=1 | 3 |
| P24928 | DNA-directed RNA polymerase II subunit RPB1 OS=Homo sapiens OX=9606 GN=POLR2A PE=1 SV=2 | 1 |
| P25440 | Bromodomain-containing protein 2 OS=Homo sapiens OX=9606 GN=BRD2 PE=1 SV=2 | 2 |
| P27338 | Amine oxidase [flavin-containing] B OS=Homo sapiens OX=9606 GN=MAOB PE=1 SV=3 | 2 |
| P27695 | DNA-(apurinic or apyrimidinic site) endonuclease OS=Homo sapiens OX=9606 GN=APEX1 PE=1 SV=2 | 1 |
| P27918 | Properdin OS=Homo sapiens OX=9606 GN=CFP PE=1 SV=2 | 1 |
| P28749 | Retinoblastoma-like protein 1 OS=Homo sapiens OX=9606 GN=RBL1 PE=1 SV=3 | 1 |
| P29353 | SHC-transforming protein 1 OS=Homo sapiens OX=9606 GN=SHC1 PE=1 SV=4 | 1 |
| P30203 | T-cell differentiation antigen CD6 OS=Homo sapiens OX=9606 GN=CD6 PE=1 SV=3 | 1 |
| P30793 | GTP cyclohydrolase 1 OS=Homo sapiens OX=9606 GN=GCH1 PE=1 SV=1 | 4 |
| P31350 | Ribonucleoside-diphosphate reductase subunit M2 OS=Homo sapiens OX=9606 GN=RRM2 PE=1 SV=1 | 2 |
| P31629 | Transcription factor HIVEP2 OS=Homo sapiens OX=9606 GN=HIVEP2 PE=1 SV=2 | 1 |
| P32243 | Homeobox protein OTX2 OS=Homo sapiens OX=9606 GN=OTX2 PE=1 SV=1 | 1 |
| P33527 | Multidrug resistance-associated protein 1 OS=Homo sapiens OX=9606 GN=ABCC1 PE=1 SV=3 | 2 |
| P33991 | DNA replication licensing factor MCM4 OS=Homo sapiens OX=9606 GN=MCM4 PE=1 SV=5 | 3 |
| P33992 | DNA replication licensing factor MCM5 OS=Homo sapiens OX=9606 GN=MCM5 PE=1 SV=5 | 3 |
| P33993 | DNA replication licensing factor MCM7 OS=Homo sapiens OX=9606 GN=MCM7 PE=1 SV=4 | 3 |
| P35030 | Trypsin-3 OS=Homo sapiens OX=9606 GN=PRSS3 PE=1 SV=2 | 1 |
| P35052 | Glypican-1 OS=Homo sapiens OX=9606 GN=GPC1 PE=1 SV=2 | 1 |
| P35321 | Cornifin-A OS=Homo sapiens OX=9606 GN=SPRR1A PE=1 SV=2 | 1 |
| P37268 | Squalene synthase OS=Homo sapiens OX=9606 GN=FDFT1 PE=1 SV=1 | 9 |
| P38571 | Lysosomal acid lipase/cholesteryl ester hydrolase OS=Homo sapiens OX=9606 GN=LIPA PE=1 SV=2 | 1 |
| P41220 | Regulator of G-protein signaling 2 OS=Homo sapiens OX=9606 GN=RGS2 PE=1 SV=1 | 5 |
| P41247 | Patatin-like phospholipase domain-containing protein 4 OS=Homo sapiens OX=9606 GN=PNPLA4 PE=1 SV=3 | 2 |
| P42684 | Tyrosine-protein kinase ABL2 OS=Homo sapiens OX=9606 GN=ABL2 PE=1 SV=1 | 1 |
| P43246 | DNA mismatch repair protein Msh2 OS=Homo sapiens OX=9606 GN=MSH2 PE=1 SV=1 | 2 |
| P47972 | Neuronal pentraxin-2 OS=Homo sapiens OX=9606 GN=NPTX2 PE=1 SV=2 | 1 |
| P48509 | CD151 antigen OS=Homo sapiens OX=9606 GN=CD151 PE=1 SV=3 | 2 |
| P48539 | Calmodulin regulator protein PCP4 OS=Homo sapiens OX=9606 GN=PCP4 PE=1 SV=3 | 1 |
| P48668 | Keratin, type II cytoskeletal 6C OS=Homo sapiens OX=9606 GN=KRT6C PE=1 SV=3 | 1 |
| P48723 | Heat shock 70 kDa protein 13 OS=Homo sapiens OX=9606 GN=HSPA13 PE=1 SV=1 | 1 |
| P49005 | DNA polymerase delta subunit 2 OS=Homo sapiens OX=9606 GN=POLD2 PE=1 SV=1 | 1 |
| P49790 | Nuclear pore complex protein Nup153 OS=Homo sapiens OX=9606 GN=NUP153 PE=1 SV=2 | 1 |
| P50416 | Carnitine O-palmitoyltransferase 1, liver isoform OS=Homo sapiens OX=9606 GN=CPT1A PE=1 SV=2 | 1 |
| P50453 | Serpin B9 OS=Homo sapiens OX=9606 GN=SERPINB9 PE=1 SV=1 | 1 |
| P50613 | Cyclin-dependent kinase 7 OS=Homo sapiens OX=9606 GN=CDK7 PE=1 SV=1 | 6 |
| P50851 | Lipopolysaccharide-responsive and beige-like anchor protein OS=Homo sapiens OX=9606 GN=LRBA PE=1 SV=4 | 1 |
| P51608 | Methyl-CpG-binding protein 2 OS=Homo sapiens OX=9606 GN=MECP2 PE=1 SV=1 | 1 |
| P51808 | Dynein light chain Tctex-type 3 OS=Homo sapiens OX=9606 GN=DYNLT3 PE=1 SV=1 | 1 |
| P52294 | Importin subunit alpha-5 OS=Homo sapiens OX=9606 GN=KPNA1 PE=1 SV=3 | 1 |
| P52732 | Kinesin-like protein KIF11 OS=Homo sapiens OX=9606 GN=KIF11 PE=1 SV=2 | 2 |
| P53675 | Clathrin heavy chain 2 OS=Homo sapiens OX=9606 GN=CLTCL1 PE=1 SV=2 | 1 |
| P54132 | RecQ-like DNA helicase BLM OS=Homo sapiens OX=9606 GN=BLM PE=1 SV=1 | 1 |
| P55061 | Bax inhibitor 1 OS=Homo sapiens OX=9606 GN=TMBIM6 PE=1 SV=2 | 1 |
| P55064 | Aquaporin-5 OS=Homo sapiens OX=9606 GN=AQP5 PE=1 SV=1 | 1 |
| P55197 | Protein AF-10 OS=Homo sapiens OX=9606 GN=MLLT10 PE=1 SV=2 | 1 |
| P55210 | Caspase-7 OS=Homo sapiens OX=9606 GN=CASP7 PE=1 SV=1 | 3 |
| P56749 | Claudin-12 OS=Homo sapiens OX=9606 GN=CLDN12 PE=1 SV=1 | 1 |
| P61326;Q96A72 | Protein mago nashi homolog OS=Homo sapiens OX=9606 GN=MAGOH PE=1 SV=1 | 2 |
| P61513 | 60S ribosomal protein L37a OS=Homo sapiens OX=9606 GN=RPL37A PE=1 SV=2 | 2 |
| P61803 | Dolichyl-diphosphooligosaccharide--protein glycosyltransferase subunit DAD1 OS=Homo sapiens OX=9606 GN=DAD1 PE=1 SV=3 | 1 |
| P62314 | Small nuclear ribonucleoprotein Sm D1 OS=Homo sapiens OX=9606 GN=SNRPD1 PE=1 SV=1 | 1 |
| P62995 | Transformer-2 protein homolog beta OS=Homo sapiens OX=9606 GN=TRA2B PE=1 SV=1 | 6 |
| P63165 | Small ubiquitin-related modifier 1 OS=Homo sapiens OX=9606 GN=SUMO1 PE=1 SV=1 | 3 |
| P80404 | 4-aminobutyrate aminotransferase, mitochondrial OS=Homo sapiens OX=9606 GN=ABAT PE=1 SV=3 | 2 |
| P82675 | 28S ribosomal protein S5, mitochondrial OS=Homo sapiens OX=9606 GN=MRPS5 PE=1 SV=2 | 1 |
| P98095 | Fibulin-2 OS=Homo sapiens OX=9606 GN=FBLN2 PE=1 SV=2 | 1 |
| P98172 | Ephrin-B1 OS=Homo sapiens OX=9606 GN=EFNB1 PE=1 SV=1 | 1 |
| Q00403 | Transcription initiation factor IIB OS=Homo sapiens OX=9606 GN=GTF2B PE=1 SV=1 | 5 |
| Q01081 | Splicing factor U2AF 35 kDa subunit OS=Homo sapiens OX=9606 GN=U2AF1 PE=1 SV=3 | 1 |
| Q01432 | AMP deaminase 3 OS=Homo sapiens OX=9606 GN=AMPD3 PE=1 SV=1 | 3 |
| Q01814 | Plasma membrane calcium-transporting ATPase 2 OS=Homo sapiens OX=9606 GN=ATP2B2 PE=1 SV=2 | 1 |
| Q02487 | Desmocollin-2 OS=Homo sapiens OX=9606 GN=DSC2 PE=1 SV=1 | 2 |
| Q04941 | Proteolipid protein 2 OS=Homo sapiens OX=9606 GN=PLP2 PE=1 SV=1 | 2 |
| Q05193 | Dynamin-1 OS=Homo sapiens OX=9606 GN=DNM1 PE=1 SV=2 | 1 |
| Q06124 | Tyrosine-protein phosphatase non-receptor type 11 OS=Homo sapiens OX=9606 GN=PTPN11 PE=1 SV=3 | 1 |
| Q07817 | Bcl-2-like protein 1 OS=Homo sapiens OX=9606 GN=BCL2L1 PE=1 SV=1 | 3 |
| Q08945 | FACT complex subunit SSRP1 OS=Homo sapiens OX=9606 GN=SSRP1 PE=1 SV=1 | 2 |
| Q12815 | Tastin OS=Homo sapiens OX=9606 GN=TROAP PE=1 SV=3 | 2 |
| Q12834 | Cell division cycle protein 20 homolog OS=Homo sapiens OX=9606 GN=CDC20 PE=1 SV=2 | 3 |
| Q12874 | Splicing factor 3A subunit 3 OS=Homo sapiens OX=9606 GN=SF3A3 PE=1 SV=1 | 2 |
| Q12923 | Tyrosine-protein phosphatase non-receptor type 13 OS=Homo sapiens OX=9606 GN=PTPN13 PE=1 SV=2 | 1 |
| Q12959 | Disks large homolog 1 OS=Homo sapiens OX=9606 GN=DLG1 PE=1 SV=2 | 1 |
| Q13112 | Chromatin assembly factor 1 subunit B OS=Homo sapiens OX=9606 GN=CHAF1B PE=1 SV=1 | 1 |
| Q13136 | Liprin-alpha-1 OS=Homo sapiens OX=9606 GN=PPFIA1 PE=1 SV=1 | 1 |
| Q13137 | Calcium-binding and coiled-coil domain-containing protein 2 OS=Homo sapiens OX=9606 GN=CALCOCO2 PE=1 SV=1 | 2 |
| Q13151 | Heterogeneous nuclear ribonucleoprotein A0 OS=Homo sapiens OX=9606 GN=HNRNPA0 PE=1 SV=1 | 2 |
| Q13190 | Syntaxin-5 OS=Homo sapiens OX=9606 GN=STX5 PE=1 SV=2 | 1 |
| Q13257 | Mitotic spindle assembly checkpoint protein MAD2A OS=Homo sapiens OX=9606 GN=MAD2L1 PE=1 SV=1 | 7 |
| Q13309 | S-phase kinase-associated protein 2 OS=Homo sapiens OX=9606 GN=SKP2 PE=1 SV=2 | 5 |
| Q13423 | NAD(P) transhydrogenase, mitochondrial OS=Homo sapiens OX=9606 GN=NNT PE=1 SV=3 | 1 |
| Q13491 | Neuronal membrane glycoprotein M6-b OS=Homo sapiens OX=9606 GN=GPM6B PE=1 SV=2 | 1 |
| Q13620 | Cullin-4B OS=Homo sapiens OX=9606 GN=CUL4B PE=1 SV=4 | 4 |
| Q13951 | Core-binding factor subunit beta OS=Homo sapiens OX=9606 GN=CBFB PE=1 SV=2 | 1 |
| Q14134 | Tripartite motif-containing protein 29 OS=Homo sapiens OX=9606 GN=TRIM29 PE=1 SV=2 | 2 |
| Q14195 | Dihydropyrimidinase-related protein 3 OS=Homo sapiens OX=9606 GN=DPYSL3 PE=1 SV=1 | 4 |
| Q14315 | Filamin-C OS=Homo sapiens OX=9606 GN=FLNC PE=1 SV=3 | 1 |
| Q14493 | Histone RNA hairpin-binding protein OS=Homo sapiens OX=9606 GN=SLBP PE=1 SV=1 | 6 |
| Q14674 | Separin OS=Homo sapiens OX=9606 GN=ESPL1 PE=1 SV=3 | 3 |
| Q14728 | Major facilitator superfamily domain-containing protein 10 OS=Homo sapiens OX=9606 GN=MFSD10 PE=1 SV=1 | 3 |
| Q14746 | Conserved oligomeric Golgi complex subunit 2 OS=Homo sapiens OX=9606 GN=COG2 PE=1 SV=1 | 2 |
| Q14980 | Nuclear mitotic apparatus protein 1 OS=Homo sapiens OX=9606 GN=NUMA1 PE=1 SV=2 | 11 |
| Q15007 | Pre-mRNA-splicing regulator WTAP OS=Homo sapiens OX=9606 GN=WTAP PE=1 SV=2 | 2 |
| Q15024 | Exosome complex component RRP42 OS=Homo sapiens OX=9606 GN=EXOSC7 PE=1 SV=3 | 1 |
| Q15041 | ADP-ribosylation factor-like protein 6-interacting protein 1 OS=Homo sapiens OX=9606 GN=ARL6IP1 PE=1 SV=2 | 1 |
| Q15075 | Early endosome antigen 1 OS=Homo sapiens OX=9606 GN=EEA1 PE=1 SV=2 | 1 |
| Q15746 | Myosin light chain kinase, smooth muscle OS=Homo sapiens OX=9606 GN=MYLK PE=1 SV=4 | 1 |
| Q16099 | Glutamate receptor ionotropic, kainate 4 OS=Homo sapiens OX=9606 GN=GRIK4 PE=2 SV=2 | 2 |
| Q16143 | Beta-synuclein OS=Homo sapiens OX=9606 GN=SNCB PE=1 SV=1 | 2 |
| Q16281 | Cyclic nucleotide-gated cation channel alpha-3 OS=Homo sapiens OX=9606 GN=CNGA3 PE=1 SV=2 | 1 |
| Q16629 | Serine/arginine-rich splicing factor 7 OS=Homo sapiens OX=9606 GN=SRSF7 PE=1 SV=1 | 2 |
| Q16763 | Ubiquitin-conjugating enzyme E2 S OS=Homo sapiens OX=9606 GN=UBE2S PE=1 SV=2 | 1 |
| Q16773 | Kynurenine--oxoglutarate transaminase 1 OS=Homo sapiens OX=9606 GN=KYAT1 PE=1 SV=1 | 1 |
| Q2T9J0 | Peroxisomal leader peptide-processing protease OS=Homo sapiens OX=9606 GN=TYSND1 PE=1 SV=3 | 2 |
| Q32MZ4 | Leucine-rich repeat flightless-interacting protein 1 OS=Homo sapiens OX=9606 GN=LRRFIP1 PE=1 SV=2 | 1 |
| Q3SXM0 | DDB1- and CUL4-associated factor 4-like protein 1 OS=Homo sapiens OX=9606 GN=DCAF4L1 PE=2 SV=1 | 2 |
| Q4ADV7 | Guanine nucleotide exchange factor subunit RIC1 OS=Homo sapiens OX=9606 GN=RIC1 PE=1 SV=2 | 1 |
| Q4G0J3 | La-related protein 7 OS=Homo sapiens OX=9606 GN=LARP7 PE=1 SV=1 | 3 |
| Q4G0P3 | Hydrocephalus-inducing protein homolog OS=Homo sapiens OX=9606 GN=HYDIN PE=1 SV=3 | 1 |
| Q53H80 | Akirin-2 OS=Homo sapiens OX=9606 GN=AKIRIN2 PE=1 SV=2 | 5 |
| Q53HC9 | EARP and GARP complex-interacting protein 1 OS=Homo sapiens OX=9606 GN=EIPR1 PE=1 SV=2 | 2 |
| Q53SF7 | Cordon-bleu protein-like 1 OS=Homo sapiens OX=9606 GN=COBLL1 PE=1 SV=3 | 2 |
| Q53T59 | HCLS1-binding protein 3 OS=Homo sapiens OX=9606 GN=HS1BP3 PE=1 SV=1 | 1 |
| Q56NI9 | N-acetyltransferase ESCO2 OS=Homo sapiens OX=9606 GN=ESCO2 PE=1 SV=1 | 2 |
| Q5BJF2 | Sigma intracellular receptor 2 OS=Homo sapiens OX=9606 GN=TMEM97 PE=1 SV=1 | 1 |
| Q5BJH2 | Transmembrane protein 128 OS=Homo sapiens OX=9606 GN=TMEM128 PE=1 SV=2 | 1 |
| Q5DT21 | Serine protease inhibitor Kazal-type 9 OS=Homo sapiens OX=9606 GN=SPINK9 PE=1 SV=1 | 1 |
| Q5EBM0 | UMP-CMP kinase 2, mitochondrial OS=Homo sapiens OX=9606 GN=CMPK2 PE=1 SV=3 | 3 |
| Q5JTZ9 | Alanine--tRNA ligase, mitochondrial OS=Homo sapiens OX=9606 GN=AARS2 PE=1 SV=1 | 1 |
| Q5T1H1 | Protein eyes shut homolog OS=Homo sapiens OX=9606 GN=EYS PE=1 SV=5 | 1 |
| Q5T1V6 | Probable ATP-dependent RNA helicase DDX59 OS=Homo sapiens OX=9606 GN=DDX59 PE=1 SV=1 | 3 |
| Q5T2D3 | OTU domain-containing protein 3 OS=Homo sapiens OX=9606 GN=OTUD3 PE=1 SV=1 | 1 |
| Q5T7W7 | Thiosulfate sulfurtransferase/rhodanese-like domain-containing protein 2 OS=Homo sapiens OX=9606 GN=TSTD2 PE=1 SV=1 | 1 |
| Q5TCS8 | Adenylate kinase 9 OS=Homo sapiens OX=9606 GN=AK9 PE=1 SV=2 | 1 |
| Q5TF58 | Intermediate filament family orphan 2 OS=Homo sapiens OX=9606 GN=IFFO2 PE=2 SV=3 | 1 |
| Q5TZA2 | Rootletin OS=Homo sapiens OX=9606 GN=CROCC PE=1 SV=2 | 1 |
| Q5UIP0 | Telomere-associated protein RIF1 OS=Homo sapiens OX=9606 GN=RIF1 PE=1 SV=2 | 1 |
| Q5VSL9 | Striatin-interacting protein 1 OS=Homo sapiens OX=9606 GN=STRIP1 PE=1 SV=1 | 2 |
| Q5VST9 | Obscurin OS=Homo sapiens OX=9606 GN=OBSCN PE=1 SV=3 | 2 |
| Q5VTR2 | E3 ubiquitin-protein ligase BRE1A OS=Homo sapiens OX=9606 GN=RNF20 PE=1 SV=2 | 1 |
| Q63HM2 | Pecanex-like protein 4 OS=Homo sapiens OX=9606 GN=PCNX4 PE=1 SV=4 | 1 |
| Q68CJ6 | Nuclear GTPase SLIP-GC OS=Homo sapiens OX=9606 GN=NUGGC PE=2 SV=3 | 1 |
| Q68DD2 | Cytosolic phospholipase A2 zeta OS=Homo sapiens OX=9606 GN=PLA2G4F PE=1 SV=3 | 1 |
| Q6DKK2 | Tetratricopeptide repeat protein 19, mitochondrial OS=Homo sapiens OX=9606 GN=TTC19 PE=1 SV=4 | 1 |
| Q6GMR7 | Fatty-acid amide hydrolase 2 OS=Homo sapiens OX=9606 GN=FAAH2 PE=1 SV=1 | 1 |
| Q6ICL3 | Transport and Golgi organization protein 2 homolog OS=Homo sapiens OX=9606 GN=TANGO2 PE=1 SV=1 | 1 |
| Q6IQ49 | Splicing regulator SDE2 OS=Homo sapiens OX=9606 GN=SDE2 PE=1 SV=1 | 3 |
| Q6JQN1 | Acyl-CoA dehydrogenase family member 10 OS=Homo sapiens OX=9606 GN=ACAD10 PE=1 SV=1 | 1 |
| Q6NUK1 | Calcium-binding mitochondrial carrier protein SCaMC-1 OS=Homo sapiens OX=9606 GN=SLC25A24 PE=1 SV=2 | 1 |
| Q6P2I3 | Fumarylacetoacetate hydrolase domain-containing protein 2B OS=Homo sapiens OX=9606 GN=FAHD2B PE=1 SV=1 | 1 |
| Q6PCB7 | Long-chain fatty acid transport protein 1 OS=Homo sapiens OX=9606 GN=SLC27A1 PE=1 SV=1 | 1 |
| Q6Q795 | Putative viral protein-binding protein C1 OS=Homo sapiens OX=9606 PE=5 SV=1 | 1 |
| Q6U841 | Sodium-driven chloride bicarbonate exchanger OS=Homo sapiens OX=9606 GN=SLC4A10 PE=1 SV=1 | 1 |
| Q6UVJ0 | Spindle assembly abnormal protein 6 homolog OS=Homo sapiens OX=9606 GN=SASS6 PE=1 SV=1 | 7 |
| Q6UWN5 | Ly6/PLAUR domain-containing protein 5 OS=Homo sapiens OX=9606 GN=LYPD5 PE=1 SV=2 | 1 |
| Q6UWY5 | Olfactomedin-like protein 1 OS=Homo sapiens OX=9606 GN=OLFML1 PE=1 SV=2 | 1 |
| Q6UX71 | Plexin domain-containing protein 2 OS=Homo sapiens OX=9606 GN=PLXDC2 PE=1 SV=1 | 1 |
| Q6V1X1 | Dipeptidyl peptidase 8 OS=Homo sapiens OX=9606 GN=DPP8 PE=1 SV=1 | 1 |
| Q6ZMG9 | Ceramide synthase 6 OS=Homo sapiens OX=9606 GN=CERS6 PE=1 SV=1 | 1 |
| Q6ZMW3 | Echinoderm microtubule-associated protein-like 6 OS=Homo sapiens OX=9606 GN=EML6 PE=2 SV=2 | 1 |
| Q70JA7 | Chondroitin sulfate synthase 3 OS=Homo sapiens OX=9606 GN=CHSY3 PE=2 SV=3 | 1 |
| Q71SY5 | Mediator of RNA polymerase II transcription subunit 25 OS=Homo sapiens OX=9606 GN=MED25 PE=1 SV=2 | 1 |
| Q7L592 | Protein arginine methyltransferase NDUFAF7, mitochondrial OS=Homo sapiens OX=9606 GN=NDUFAF7 PE=1 SV=1 | 7 |
| Q7L9L4;Q9H8S9 | MOB kinase activator 1B OS=Homo sapiens OX=9606 GN=MOB1B PE=1 SV=3 | 1 |
| Q7Z3K6 | Mesoderm induction early response protein 3 OS=Homo sapiens OX=9606 GN=MIER3 PE=1 SV=2 | 1 |
| Q7Z449 | Cytochrome P450 2U1 OS=Homo sapiens OX=9606 GN=CYP2U1 PE=1 SV=1 | 1 |
| Q7Z4G4 | tRNA (guanine(10)-N2)-methyltransferase homolog OS=Homo sapiens OX=9606 GN=TRMT11 PE=1 SV=1 | 1 |
| Q7Z5K2 | Wings apart-like protein homolog OS=Homo sapiens OX=9606 GN=WAPL PE=1 SV=1 | 3 |
| Q7Z6B0 | Coiled-coil domain-containing protein 91 OS=Homo sapiens OX=9606 GN=CCDC91 PE=1 SV=2 | 1 |
| Q7Z7A1 | Centriolin OS=Homo sapiens OX=9606 GN=CNTRL PE=1 SV=2 | 1 |
| Q86T24 | Transcriptional regulator Kaiso OS=Homo sapiens OX=9606 GN=ZBTB33 PE=1 SV=2 | 1 |
| Q86TM3 | Probable ATP-dependent RNA helicase DDX53 OS=Homo sapiens OX=9606 GN=DDX53 PE=1 SV=3 | 1 |
| Q86TS9 | 39S ribosomal protein L52, mitochondrial OS=Homo sapiens OX=9606 GN=MRPL52 PE=1 SV=2 | 1 |
| Q86UX6 | Serine/threonine-protein kinase 32C OS=Homo sapiens OX=9606 GN=STK32C PE=1 SV=1 | 1 |
| Q86VP3 | Phosphofurin acidic cluster sorting protein 2 OS=Homo sapiens OX=9606 GN=PACS2 PE=1 SV=3 | 1 |
| Q86VV8 | Rotatin OS=Homo sapiens OX=9606 GN=RTTN PE=1 SV=3 | 1 |
| Q86W28 | NACHT, LRR and PYD domains-containing protein 8 OS=Homo sapiens OX=9606 GN=NLRP8 PE=2 SV=2 | 1 |
| Q86WA8 | Lon protease homolog 2, peroxisomal OS=Homo sapiens OX=9606 GN=LONP2 PE=1 SV=1 | 1 |
| Q86WT1 | Tetratricopeptide repeat protein 30A OS=Homo sapiens OX=9606 GN=TTC30A PE=2 SV=3 | 1 |
| Q86XP0 | Cytosolic phospholipase A2 delta OS=Homo sapiens OX=9606 GN=PLA2G4D PE=1 SV=2 | 1 |
| Q86YT6 | E3 ubiquitin-protein ligase MIB1 OS=Homo sapiens OX=9606 GN=MIB1 PE=1 SV=1 | 1 |
| Q8IUF8 | Ribosomal oxygenase 2 OS=Homo sapiens OX=9606 GN=RIOX2 PE=1 SV=1 | 7 |
| Q8IUZ5 | 5-phosphohydroxy-L-lysine phospho-lyase OS=Homo sapiens OX=9606 GN=PHYKPL PE=1 SV=1 | 1 |
| Q8IW41 | MAP kinase-activated protein kinase 5 OS=Homo sapiens OX=9606 GN=MAPKAPK5 PE=1 SV=2 | 1 |
| Q8IW92 | Beta-galactosidase-1-like protein 2 OS=Homo sapiens OX=9606 GN=GLB1L2 PE=1 SV=1 | 1 |
| Q8IWA0 | WD repeat-containing protein 75 OS=Homo sapiens OX=9606 GN=WDR75 PE=1 SV=1 | 1 |
| Q8IWI9 | MAX gene-associated protein OS=Homo sapiens OX=9606 GN=MGA PE=1 SV=4 | 2 |
| Q8IWR0 | Zinc finger CCCH domain-containing protein 7A OS=Homo sapiens OX=9606 GN=ZC3H7A PE=1 SV=1 | 1 |
| Q8IWV7 | E3 ubiquitin-protein ligase UBR1 OS=Homo sapiens OX=9606 GN=UBR1 PE=1 SV=1 | 2 |
| Q8IWX8 | Calcium homeostasis endoplasmic reticulum protein OS=Homo sapiens OX=9606 GN=CHERP PE=1 SV=3 | 4 |
| Q8IXM2 | Chromatin complexes subunit BAP18 OS=Homo sapiens OX=9606 GN=BAP18 PE=1 SV=1 | 3 |
| Q8IXT5 | RNA-binding protein 12B OS=Homo sapiens OX=9606 GN=RBM12B PE=1 SV=2 | 1 |
| Q8IY18 | Structural maintenance of chromosomes protein 5 OS=Homo sapiens OX=9606 GN=SMC5 PE=1 SV=2 | 1 |
| Q8IY47 | Kelch repeat and BTB domain-containing protein 2 OS=Homo sapiens OX=9606 GN=KBTBD2 PE=1 SV=2 | 1 |
| Q8IYF3 | Testis-expressed protein 11 OS=Homo sapiens OX=9606 GN=TEX11 PE=1 SV=3 | 2 |
| Q8N136 | Dynein assembly factor with WDR repeat domains 1 OS=Homo sapiens OX=9606 GN=DAW1 PE=1 SV=1 | 5 |
| Q8N1B4 | Vacuolar protein sorting-associated protein 52 homolog OS=Homo sapiens OX=9606 GN=VPS52 PE=1 SV=1 | 2 |
| Q8N1G2 | Cap-specific mRNA (nucleoside-2'-O-)-methyltransferase 1 OS=Homo sapiens OX=9606 GN=CMTR1 PE=1 SV=1 | 12 |
| Q8N257 | Histone H2B type 3-B OS=Homo sapiens OX=9606 GN=H2BC26 PE=1 SV=3 | 1 |
| Q8N2G8 | GH3 domain-containing protein OS=Homo sapiens OX=9606 GN=GHDC PE=1 SV=2 | 1 |
| Q8N335 | Glycerol-3-phosphate dehydrogenase 1-like protein OS=Homo sapiens OX=9606 GN=GPD1L PE=1 SV=1 | 1 |
| Q8N442 | Translation factor GUF1, mitochondrial OS=Homo sapiens OX=9606 GN=GUF1 PE=1 SV=1 | 1 |
| Q8N6M0 | Deubiquitinase OTUD6B OS=Homo sapiens OX=9606 GN=OTUD6B PE=1 SV=1 | 2 |
| Q8N987 | N-terminal EF-hand calcium-binding protein 1 OS=Homo sapiens OX=9606 GN=NECAB1 PE=1 SV=1 | 1 |
| Q8N9B5 | Junction-mediating and -regulatory protein OS=Homo sapiens OX=9606 GN=JMY PE=1 SV=2 | 1 |
| Q8NBI6 | Xyloside xylosyltransferase 1 OS=Homo sapiens OX=9606 GN=XXYLT1 PE=1 SV=1 | 2 |
| Q8NBN7 | Retinol dehydrogenase 13 OS=Homo sapiens OX=9606 GN=RDH13 PE=1 SV=2 | 1 |
| Q8NE62 | Choline dehydrogenase, mitochondrial OS=Homo sapiens OX=9606 GN=CHDH PE=1 SV=2 | 1 |
| Q8NEE6 | Dynein regulatory complex subunit 6 OS=Homo sapiens OX=9606 GN=FBXL13 PE=2 SV=3 | 1 |
| Q8NEY8 | Periphilin-1 OS=Homo sapiens OX=9606 GN=PPHLN1 PE=1 SV=2 | 2 |
| Q8NFG4 | Folliculin OS=Homo sapiens OX=9606 GN=FLCN PE=1 SV=1 | 2 |
| Q8NFH4 | Nucleoporin Nup37 OS=Homo sapiens OX=9606 GN=NUP37 PE=1 SV=1 | 7 |
| Q8NFH5 | Nucleoporin NUP35 OS=Homo sapiens OX=9606 GN=NUP35 PE=1 SV=1 | 2 |
| Q8NFP9 | Neurobeachin OS=Homo sapiens OX=9606 GN=NBEA PE=1 SV=3 | 1 |
| Q8NG68 | Tubulin--tyrosine ligase OS=Homo sapiens OX=9606 GN=TTL PE=1 SV=2 | 1 |
| Q8NHP6 | Motile sperm domain-containing protein 2 OS=Homo sapiens OX=9606 GN=MOSPD2 PE=1 SV=1 | 2 |
| Q8NHU6 | Tudor domain-containing protein 7 OS=Homo sapiens OX=9606 GN=TDRD7 PE=1 SV=2 | 5 |
| Q8NI22 | Multiple coagulation factor deficiency protein 2 OS=Homo sapiens OX=9606 GN=MCFD2 PE=1 SV=1 | 1 |
| Q8TAL5 | Uncharacterized protein C9orf43 OS=Homo sapiens OX=9606 GN=C9orf43 PE=1 SV=1 | 1 |
| Q8TAP8 | Protein phosphatase 1 regulatory subunit 35 OS=Homo sapiens OX=9606 GN=PPP1R35 PE=1 SV=1 | 3 |
| Q8TB52 | F-box only protein 30 OS=Homo sapiens OX=9606 GN=FBXO30 PE=1 SV=3 | 6 |
| Q8TBY0 | Probable RNA-binding protein 46 OS=Homo sapiens OX=9606 GN=RBM46 PE=1 SV=1 | 2 |
| Q8TCG5 | Carnitine O-palmitoyltransferase 1, brain isoform OS=Homo sapiens OX=9606 GN=CPT1C PE=1 SV=1 | 1 |
| Q8WTR2 | Dual specificity protein phosphatase 19 OS=Homo sapiens OX=9606 GN=DUSP19 PE=1 SV=1 | 1 |
| Q8WU76 | Sec1 family domain-containing protein 2 OS=Homo sapiens OX=9606 GN=SCFD2 PE=1 SV=2 | 1 |
| Q8WUA7 | TBC1 domain family member 22A OS=Homo sapiens OX=9606 GN=TBC1D22A PE=1 SV=2 | 1 |
| Q8WUR7 | UPF0235 protein C15orf40 OS=Homo sapiens OX=9606 GN=C15orf40 PE=1 SV=2 | 1 |
| Q8WUY8 | Probable N-acetyltransferase 14 OS=Homo sapiens OX=9606 GN=NAT14 PE=1 SV=1 | 2 |
| Q8WW12 | PEST proteolytic signal-containing nuclear protein OS=Homo sapiens OX=9606 GN=PCNP PE=1 SV=2 | 2 |
| Q8WW59 | SPRY domain-containing protein 4 OS=Homo sapiens OX=9606 GN=SPRYD4 PE=1 SV=2 | 2 |
| Q8WXF1 | Paraspeckle component 1 OS=Homo sapiens OX=9606 GN=PSPC1 PE=1 SV=1 | 6 |
| Q8WXI7 | Mucin-16 OS=Homo sapiens OX=9606 GN=MUC16 PE=1 SV=3 | 2 |
| Q8WYJ6 | Septin-1 OS=Homo sapiens OX=9606 GN=SEPTIN1 PE=1 SV=3 | 1 |
| Q8WYN0 | Cysteine protease ATG4A OS=Homo sapiens OX=9606 GN=ATG4A PE=1 SV=1 | 2 |
| Q8WYP5 | Protein ELYS OS=Homo sapiens OX=9606 GN=AHCTF1 PE=1 SV=3 | 2 |
| Q8WZA9 | Immunity-related GTPase family Q protein OS=Homo sapiens OX=9606 GN=IRGQ PE=1 SV=1 | 1 |
| Q92523 | Carnitine O-palmitoyltransferase 1, muscle isoform OS=Homo sapiens OX=9606 GN=CPT1B PE=1 SV=2 | 1 |
| Q92574 | Hamartin OS=Homo sapiens OX=9606 GN=TSC1 PE=1 SV=2 | 1 |
| Q92620 | Pre-mRNA-splicing factor ATP-dependent RNA helicase PRP16 OS=Homo sapiens OX=9606 GN=DHX38 PE=1 SV=2 | 1 |
| Q92622 | Run domain Beclin-1-interacting and cysteine-rich domain-containing protein OS=Homo sapiens OX=9606 GN=RUBCN PE=1 SV=4 | 1 |
| Q92729 | Receptor-type tyrosine-protein phosphatase U OS=Homo sapiens OX=9606 GN=PTPRU PE=1 SV=2 | 1 |
| Q92904 | Deleted in azoospermia-like OS=Homo sapiens OX=9606 GN=DAZL PE=1 SV=1 | 3 |
| Q92934 | Bcl2-associated agonist of cell death OS=Homo sapiens OX=9606 GN=BAD PE=1 SV=3 | 1 |
| Q92968 | Peroxisomal membrane protein PEX13 OS=Homo sapiens OX=9606 GN=PEX13 PE=1 SV=2 | 2 |
| Q969E8 | Pre-rRNA-processing protein TSR2 homolog OS=Homo sapiens OX=9606 GN=TSR2 PE=1 SV=1 | 1 |
| Q969I3 | Glycine N-acyltransferase-like protein 1 OS=Homo sapiens OX=9606 GN=GLYATL1 PE=1 SV=1 | 2 |
| Q969K3 | E3 ubiquitin-protein ligase RNF34 OS=Homo sapiens OX=9606 GN=RNF34 PE=1 SV=1 | 5 |
| Q969T3 | Sorting nexin-21 OS=Homo sapiens OX=9606 GN=SNX21 PE=1 SV=1 | 1 |
| Q96A26 | Protein FAM162A OS=Homo sapiens OX=9606 GN=FAM162A PE=1 SV=2 | 1 |
| Q96A33 | PAT complex subunit CCDC47 OS=Homo sapiens OX=9606 GN=CCDC47 PE=1 SV=1 | 1 |
| Q96AN5 | Transmembrane protein 143 OS=Homo sapiens OX=9606 GN=TMEM143 PE=1 SV=1 | 2 |
| Q96B02 | Ubiquitin-conjugating enzyme E2 W OS=Homo sapiens OX=9606 GN=UBE2W PE=1 SV=1 | 1 |
| Q96B86 | Repulsive guidance molecule A OS=Homo sapiens OX=9606 GN=RGMA PE=1 SV=4 | 1 |
| Q96BP3 | Peptidylprolyl isomerase domain and WD repeat-containing protein 1 OS=Homo sapiens OX=9606 GN=PPWD1 PE=1 SV=1 | 1 |
| Q96CG3 | TRAF-interacting protein with FHA domain-containing protein A OS=Homo sapiens OX=9606 GN=TIFA PE=1 SV=1 | 3 |
| Q96CS7 | Pleckstrin homology domain-containing family B member 2 OS=Homo sapiens OX=9606 GN=PLEKHB2 PE=1 SV=1 | 1 |
| Q96E09 | PPP2R1A-PPP2R2A-interacting phosphatase regulator 1 OS=Homo sapiens OX=9606 GN=PABIR1 PE=1 SV=1 | 1 |
| Q96EL3 | 39S ribosomal protein L53, mitochondrial OS=Homo sapiens OX=9606 GN=MRPL53 PE=1 SV=1 | 1 |
| Q96ES7 | SAGA-associated factor 29 OS=Homo sapiens OX=9606 GN=SGF29 PE=1 SV=1 | 1 |
| Q96EW2 | HSPB1-associated protein 1 OS=Homo sapiens OX=9606 GN=HSPBAP1 PE=1 SV=1 | 1 |
| Q96EY8 | Corrinoid adenosyltransferase MMAB OS=Homo sapiens OX=9606 GN=MMAB PE=1 SV=1 | 3 |
| Q96G23 | Ceramide synthase 2 OS=Homo sapiens OX=9606 GN=CERS2 PE=1 SV=1 | 1 |
| Q96GW9 | Methionine--tRNA ligase, mitochondrial OS=Homo sapiens OX=9606 GN=MARS2 PE=1 SV=2 | 1 |
| Q96GX5 | Serine/threonine-protein kinase greatwall OS=Homo sapiens OX=9606 GN=MASTL PE=1 SV=1 | 3 |
| Q96HE7 | ERO1-like protein alpha OS=Homo sapiens OX=9606 GN=ERO1A PE=1 SV=2 | 2 |
| Q96HY7 | 2-oxoadipate dehydrogenase complex component E1 OS=Homo sapiens OX=9606 GN=DHTKD1 PE=1 SV=2 | 1 |
| Q96I24 | Far upstream element-binding protein 3 OS=Homo sapiens OX=9606 GN=FUBP3 PE=1 SV=2 | 1 |
| Q96I25 | Splicing factor 45 OS=Homo sapiens OX=9606 GN=RBM17 PE=1 SV=1 | 1 |
| Q96IK1 | Biorientation of chromosomes in cell division protein 1 OS=Homo sapiens OX=9606 GN=BOD1 PE=1 SV=2 | 10 |
| Q96IW7 | Vesicle-trafficking protein SEC22a OS=Homo sapiens OX=9606 GN=SEC22A PE=1 SV=1 | 1 |
| Q96J02 | E3 ubiquitin-protein ligase Itchy homolog OS=Homo sapiens OX=9606 GN=ITCH PE=1 SV=2 | 1 |
| Q96J88 | Epithelial-stromal interaction protein 1 OS=Homo sapiens OX=9606 GN=EPSTI1 PE=1 SV=2 | 8 |
| Q96JG6 | Syndetin OS=Homo sapiens OX=9606 GN=VPS50 PE=1 SV=3 | 2 |
| Q96L91 | E1A-binding protein p400 OS=Homo sapiens OX=9606 GN=EP400 PE=1 SV=4 | 1 |
| Q96L96 | Alpha-protein kinase 3 OS=Homo sapiens OX=9606 GN=ALPK3 PE=1 SV=2 | 2 |
| Q96M83 | Coiled-coil domain-containing protein 7 OS=Homo sapiens OX=9606 GN=CCDC7 PE=1 SV=3 | 3 |
| Q96MA6 | Adenylate kinase 8 OS=Homo sapiens OX=9606 GN=AK8 PE=1 SV=1 | 1 |
| Q96MI6 | Protein phosphatase 1M OS=Homo sapiens OX=9606 GN=PPM1M PE=2 SV=2 | 1 |
| Q96N23 | Cilia- and flagella-associated protein 54 OS=Homo sapiens OX=9606 GN=CFAP54 PE=2 SV=3 | 1 |
| Q96NB2 | Sideroflexin-2 OS=Homo sapiens OX=9606 GN=SFXN2 PE=1 SV=2 | 2 |
| Q96NZ9 | Proline-rich acidic protein 1 OS=Homo sapiens OX=9606 GN=PRAP1 PE=1 SV=2 | 2 |
| Q96PZ0 | Pseudouridylate synthase 7 homolog OS=Homo sapiens OX=9606 GN=PUS7 PE=1 SV=2 | 1 |
| Q96Q89 | Kinesin-like protein KIF20B OS=Homo sapiens OX=9606 GN=KIF20B PE=1 SV=3 | 4 |
| Q96QD5 | DEP domain-containing protein 7 OS=Homo sapiens OX=9606 GN=DEPDC7 PE=2 SV=1 | 10 |
| Q96RQ1 | Endoplasmic reticulum-Golgi intermediate compartment protein 2 OS=Homo sapiens OX=9606 GN=ERGIC2 PE=1 SV=2 | 2 |
| Q96S97 | Myeloid-associated differentiation marker OS=Homo sapiens OX=9606 GN=MYADM PE=1 SV=2 | 1 |
| Q96SQ9 | Cytochrome P450 2S1 OS=Homo sapiens OX=9606 GN=CYP2S1 PE=1 SV=2 | 3 |
| Q96SZ5 | 2-aminoethanethiol dioxygenase OS=Homo sapiens OX=9606 GN=ADO PE=1 SV=2 | 2 |
| Q96TA2 | ATP-dependent zinc metalloprotease YME1L1 OS=Homo sapiens OX=9606 GN=YME1L1 PE=1 SV=2 | 2 |
| Q99538 | Legumain OS=Homo sapiens OX=9606 GN=LGMN PE=1 SV=1 | 2 |
| Q99622 | Protein C10 OS=Homo sapiens OX=9606 GN=C12orf57 PE=1 SV=1 | 1 |
| Q99707 | Methionine synthase OS=Homo sapiens OX=9606 GN=MTR PE=1 SV=2 | 2 |
| Q99873 | Protein arginine N-methyltransferase 1 OS=Homo sapiens OX=9606 GN=PRMT1 PE=1 SV=3 | 12 |
| Q9BPZ7 | Target of rapamycin complex 2 subunit MAPKAP1 OS=Homo sapiens OX=9606 GN=MAPKAP1 PE=1 SV=2 | 1 |
| Q9BQI0 | Allograft inflammatory factor 1-like OS=Homo sapiens OX=9606 GN=AIF1L PE=1 SV=1 | 1 |
| Q9BR61 | Acyl-CoA-binding domain-containing protein 6 OS=Homo sapiens OX=9606 GN=ACBD6 PE=1 SV=1 | 1 |
| Q9BRX5 | DNA replication complex GINS protein PSF3 OS=Homo sapiens OX=9606 GN=GINS3 PE=1 SV=1 | 1 |
| Q9BTE1 | Dynactin subunit 5 OS=Homo sapiens OX=9606 GN=DCTN5 PE=1 SV=1 | 1 |
| Q9BTL3 | RNA guanine-N7 methyltransferase activating subunit OS=Homo sapiens OX=9606 GN=RAMAC PE=1 SV=1 | 1 |
| Q9BUE6 | Iron-sulfur cluster assembly 1 homolog, mitochondrial OS=Homo sapiens OX=9606 GN=ISCA1 PE=1 SV=1 | 2 |
| Q9BUQ8 | Probable ATP-dependent RNA helicase DDX23 OS=Homo sapiens OX=9606 GN=DDX23 PE=1 SV=3 | 1 |
| Q9BV10 | Dol-P-Man:Man(7)GlcNAc(2)-PP-Dol alpha-1,6-mannosyltransferase OS=Homo sapiens OX=9606 GN=ALG12 PE=1 SV=1 | 1 |
| Q9BW27 | Nuclear pore complex protein Nup85 OS=Homo sapiens OX=9606 GN=NUP85 PE=1 SV=1 | 1 |
| Q9BW71 | HIRA-interacting protein 3 OS=Homo sapiens OX=9606 GN=HIRIP3 PE=1 SV=3 | 1 |
| Q9BW83 | Intraflagellar transport protein 27 homolog OS=Homo sapiens OX=9606 GN=IFT27 PE=1 SV=1 | 2 |
| Q9BWH2 | FUN14 domain-containing protein 2 OS=Homo sapiens OX=9606 GN=FUNDC2 PE=1 SV=2 | 1 |
| Q9BWM7 | Sideroflexin-3 OS=Homo sapiens OX=9606 GN=SFXN3 PE=1 SV=3 | 4 |
| Q9BWQ6 | Protein YIPF2 OS=Homo sapiens OX=9606 GN=YIPF2 PE=1 SV=1 | 1 |
| Q9BXF3 | Chromatin remodeling regulator CECR2 OS=Homo sapiens OX=9606 GN=CECR2 PE=1 SV=2 | 1 |
| Q9BXI6 | TBC1 domain family member 10A OS=Homo sapiens OX=9606 GN=TBC1D10A PE=1 SV=1 | 1 |
| Q9BXJ9 | N-alpha-acetyltransferase 15, NatA auxiliary subunit OS=Homo sapiens OX=9606 GN=NAA15 PE=1 SV=1 | 2 |
| Q9BXT5 | Testis-expressed protein 15 OS=Homo sapiens OX=9606 GN=TEX15 PE=1 SV=2 | 1 |
| Q9BY32 | Inosine triphosphate pyrophosphatase OS=Homo sapiens OX=9606 GN=ITPA PE=1 SV=2 | 1 |
| Q9BYC5 | Alpha-(1,6)-fucosyltransferase OS=Homo sapiens OX=9606 GN=FUT8 PE=1 SV=2 | 1 |
| Q9BYD1 | 39S ribosomal protein L13, mitochondrial OS=Homo sapiens OX=9606 GN=MRPL13 PE=1 SV=1 | 1 |
| Q9BYG4;Q9BYG5;Q9NPB6 | Partitioning defective 6 homolog gamma OS=Homo sapiens OX=9606 GN=PARD6G PE=1 SV=1 | 1 |
| Q9BZ71 | Membrane-associated phosphatidylinositol transfer protein 3 OS=Homo sapiens OX=9606 GN=PITPNM3 PE=1 SV=2 | 1 |
| Q9C005 | Protein dpy-30 homolog OS=Homo sapiens OX=9606 GN=DPY30 PE=1 SV=1 | 2 |
| Q9C0A1 | Zinc finger homeobox protein 2 OS=Homo sapiens OX=9606 GN=ZFHX2 PE=1 SV=3 | 1 |
| Q9C0C7 | Activating molecule in BECN1-regulated autophagy protein 1 OS=Homo sapiens OX=9606 GN=AMBRA1 PE=1 SV=2 | 1 |
| Q9GZL7 | Ribosome biogenesis protein WDR12 OS=Homo sapiens OX=9606 GN=WDR12 PE=1 SV=2 | 1 |
| Q9GZR5 | Elongation of very long chain fatty acids protein 4 OS=Homo sapiens OX=9606 GN=ELOVL4 PE=1 SV=1 | 1 |
| Q9H000 | E3 ubiquitin-protein ligase makorin-2 OS=Homo sapiens OX=9606 GN=MKRN2 PE=1 SV=2 | 1 |
| Q9H1A4 | Anaphase-promoting complex subunit 1 OS=Homo sapiens OX=9606 GN=ANAPC1 PE=1 SV=1 | 1 |
| Q9H1Y0 | Autophagy protein 5 OS=Homo sapiens OX=9606 GN=ATG5 PE=1 SV=2 | 1 |
| Q9H3M9 | Ataxin-3-like protein OS=Homo sapiens OX=9606 GN=ATXN3L PE=1 SV=2 | 1 |
| Q9H3R5 | Centromere protein H OS=Homo sapiens OX=9606 GN=CENPH PE=1 SV=1 | 1 |
| Q9H4A6 | Golgi phosphoprotein 3 OS=Homo sapiens OX=9606 GN=GOLPH3 PE=1 SV=1 | 1 |
| Q9H4F8 | SPARC-related modular calcium-binding protein 1 OS=Homo sapiens OX=9606 GN=SMOC1 PE=1 SV=1 | 1 |
| Q9H5V9 | STING ER exit protein OS=Homo sapiens OX=9606 GN=STEEP1 PE=1 SV=1 | 1 |
| Q9H6D7 | HAUS augmin-like complex subunit 4 OS=Homo sapiens OX=9606 GN=HAUS4 PE=1 SV=1 | 1 |
| Q9H7B4 | Histone-lysine N-methyltransferase SMYD3 OS=Homo sapiens OX=9606 GN=SMYD3 PE=1 SV=4 | 1 |
| Q9H814 | Phosphorylated adapter RNA export protein OS=Homo sapiens OX=9606 GN=PHAX PE=1 SV=1 | 1 |
| Q9H8V3 | Protein ECT2 OS=Homo sapiens OX=9606 GN=ECT2 PE=1 SV=4 | 4 |
| Q9H8Y5 | Ankyrin repeat and zinc finger domain-containing protein 1 OS=Homo sapiens OX=9606 GN=ANKZF1 PE=1 SV=1 | 1 |
| Q9HBG6 | Intraflagellar transport protein 122 homolog OS=Homo sapiens OX=9606 GN=IFT122 PE=1 SV=2 | 1 |
| Q9HBR0 | Putative sodium-coupled neutral amino acid transporter 10 OS=Homo sapiens OX=9606 GN=SLC38A10 PE=1 SV=2 | 3 |
| Q9HBW9 | Adhesion G protein-coupled receptor L4 OS=Homo sapiens OX=9606 GN=ADGRL4 PE=1 SV=3 | 1 |
| Q9HCD5 | Nuclear receptor coactivator 5 OS=Homo sapiens OX=9606 GN=NCOA5 PE=1 SV=2 | 1 |
| Q9HD36 | Bcl-2-like protein 10 OS=Homo sapiens OX=9606 GN=BCL2L10 PE=1 SV=3 | 3 |
| Q9NPA8 | Transcription and mRNA export factor ENY2 OS=Homo sapiens OX=9606 GN=ENY2 PE=1 SV=1 | 1 |
| Q9NPD8 | Ubiquitin-conjugating enzyme E2 T OS=Homo sapiens OX=9606 GN=UBE2T PE=1 SV=1 | 5 |
| Q9NQ48 | Leucine zipper transcription factor-like protein 1 OS=Homo sapiens OX=9606 GN=LZTFL1 PE=1 SV=1 | 1 |
| Q9NQT4 | Exosome complex component RRP46 OS=Homo sapiens OX=9606 GN=EXOSC5 PE=1 SV=1 | 1 |
| Q9NQT5 | Exosome complex component RRP40 OS=Homo sapiens OX=9606 GN=EXOSC3 PE=1 SV=3 | 2 |
| Q9NRD1 | F-box only protein 6 OS=Homo sapiens OX=9606 GN=FBXO6 PE=1 SV=1 | 2 |
| Q9NRF8 | CTP synthase 2 OS=Homo sapiens OX=9606 GN=CTPS2 PE=1 SV=1 | 1 |
| Q9NRL2 | Bromodomain adjacent to zinc finger domain protein 1A OS=Homo sapiens OX=9606 GN=BAZ1A PE=1 SV=2 | 2 |
| Q9NUW8 | Tyrosyl-DNA phosphodiesterase 1 OS=Homo sapiens OX=9606 GN=TDP1 PE=1 SV=2 | 3 |
| Q9NVG8 | TBC1 domain family member 13 OS=Homo sapiens OX=9606 GN=TBC1D13 PE=1 SV=3 | 2 |
| Q9NWS1 | PCNA-interacting partner OS=Homo sapiens OX=9606 GN=PARPBP PE=1 SV=3 | 1 |
| Q9NXR7 | BRISC and BRCA1-A complex member 2 OS=Homo sapiens OX=9606 GN=BABAM2 PE=1 SV=2 | 1 |
| Q9NXR8 | Inhibitor of growth protein 3 OS=Homo sapiens OX=9606 GN=ING3 PE=1 SV=2 | 2 |
| Q9NXV2 | BTB/POZ domain-containing protein KCTD5 OS=Homo sapiens OX=9606 GN=KCTD5 PE=1 SV=1 | 2 |
| Q9NY30 | Protein BTG4 OS=Homo sapiens OX=9606 GN=BTG4 PE=1 SV=1 | 1 |
| Q9NY35 | Claudin domain-containing protein 1 OS=Homo sapiens OX=9606 GN=CLDND1 PE=1 SV=1 | 1 |
| Q9NY59 | Sphingomyelin phosphodiesterase 3 OS=Homo sapiens OX=9606 GN=SMPD3 PE=1 SV=1 | 2 |
| Q9NYV6 | RNA polymerase I-specific transcription initiation factor RRN3 OS=Homo sapiens OX=9606 GN=RRN3 PE=1 SV=1 | 1 |
| Q9NYZ3 | G2 and S phase-expressed protein 1 OS=Homo sapiens OX=9606 GN=GTSE1 PE=1 SV=3 | 1 |
| Q9P0U1 | Mitochondrial import receptor subunit TOM7 homolog OS=Homo sapiens OX=9606 GN=TOMM7 PE=1 SV=1 | 3 |
| Q9P203 | BTB/POZ domain-containing protein 7 OS=Homo sapiens OX=9606 GN=BTBD7 PE=1 SV=3 | 1 |
| Q9P225 | Dynein axonemal heavy chain 2 OS=Homo sapiens OX=9606 GN=DNAH2 PE=1 SV=3 | 1 |
| Q9P2D0 | Inhibitor of Bruton tyrosine kinase OS=Homo sapiens OX=9606 GN=IBTK PE=1 SV=3 | 1 |
| Q9P2D6 | Protein FAM135A OS=Homo sapiens OX=9606 GN=FAM135A PE=1 SV=2 | 1 |
| Q9P2I0 | Cleavage and polyadenylation specificity factor subunit 2 OS=Homo sapiens OX=9606 GN=CPSF2 PE=1 SV=2 | 1 |
| Q9P2L0 | WD repeat-containing protein 35 OS=Homo sapiens OX=9606 GN=WDR35 PE=1 SV=3 | 1 |
| Q9P2M4 | TBC1 domain family member 14 OS=Homo sapiens OX=9606 GN=TBC1D14 PE=1 SV=3 | 4 |
| Q9UBH0 | Interleukin-36 receptor antagonist protein OS=Homo sapiens OX=9606 GN=IL36RN PE=1 SV=1 | 1 |
| Q9UBU8 | Mortality factor 4-like protein 1 OS=Homo sapiens OX=9606 GN=MORF4L1 PE=1 SV=2 | 2 |
| Q9UDR5 | Alpha-aminoadipic semialdehyde synthase, mitochondrial OS=Homo sapiens OX=9606 GN=AASS PE=1 SV=1 | 1 |
| Q9UHF7 | Zinc finger transcription factor Trps1 OS=Homo sapiens OX=9606 GN=TRPS1 PE=1 SV=2 | 1 |
| Q9UHI6 | Probable ATP-dependent RNA helicase DDX20 OS=Homo sapiens OX=9606 GN=DDX20 PE=1 SV=2 | 1 |
| Q9UHY1 | Nuclear receptor-binding protein OS=Homo sapiens OX=9606 GN=NRBP1 PE=1 SV=1 | 1 |
| Q9UID3 | Vacuolar protein sorting-associated protein 51 homolog OS=Homo sapiens OX=9606 GN=VPS51 PE=1 SV=2 | 1 |
| Q9UIW2 | Plexin-A1 OS=Homo sapiens OX=9606 GN=PLXNA1 PE=1 SV=3 | 1 |
| Q9UK99 | F-box only protein 3 OS=Homo sapiens OX=9606 GN=FBXO3 PE=1 SV=3 | 1 |
| Q9UKL0 | REST corepressor 1 OS=Homo sapiens OX=9606 GN=RCOR1 PE=1 SV=2 | 3 |
| Q9UKX7 | Nuclear pore complex protein Nup50 OS=Homo sapiens OX=9606 GN=NUP50 PE=1 SV=2 | 1 |
| Q9UL15 | BAG family molecular chaperone regulator 5 OS=Homo sapiens OX=9606 GN=BAG5 PE=1 SV=1 | 2 |
| Q9UMR5 | Lysosomal thioesterase PPT2 OS=Homo sapiens OX=9606 GN=PPT2 PE=1 SV=4 | 1 |
| Q9UPN9 | E3 ubiquitin-protein ligase TRIM33 OS=Homo sapiens OX=9606 GN=TRIM33 PE=1 SV=3 | 5 |
| Q9UQ16 | Dynamin-3 OS=Homo sapiens OX=9606 GN=DNM3 PE=1 SV=4 | 1 |
| Q9UQ26 | Regulating synaptic membrane exocytosis protein 2 OS=Homo sapiens OX=9606 GN=RIMS2 PE=1 SV=2 | 1 |
| Q9UQE7 | Structural maintenance of chromosomes protein 3 OS=Homo sapiens OX=9606 GN=SMC3 PE=1 SV=2 | 2 |
| Q9Y2I1 | Nischarin OS=Homo sapiens OX=9606 GN=NISCH PE=1 SV=3 | 2 |
| Q9Y2Q0 | Phospholipid-transporting ATPase IA OS=Homo sapiens OX=9606 GN=ATP8A1 PE=1 SV=1 | 1 |
| Q9Y2V7 | Conserved oligomeric Golgi complex subunit 6 OS=Homo sapiens OX=9606 GN=COG6 PE=1 SV=2 | 1 |
| Q9Y2X9 | Zinc finger protein 281 OS=Homo sapiens OX=9606 GN=ZNF281 PE=1 SV=1 | 1 |
| Q9Y320 | Thioredoxin-related transmembrane protein 2 OS=Homo sapiens OX=9606 GN=TMX2 PE=1 SV=1 | 1 |
| Q9Y343 | Sorting nexin-24 OS=Homo sapiens OX=9606 GN=SNX24 PE=1 SV=1 | 1 |
| Q9Y3A3 | MOB-like protein phocein OS=Homo sapiens OX=9606 GN=MOB4 PE=1 SV=1 | 1 |
| Q9Y3B7 | 39S ribosomal protein L11, mitochondrial OS=Homo sapiens OX=9606 GN=MRPL11 PE=1 SV=1 | 3 |
| Q9Y3B9 | RRP15-like protein OS=Homo sapiens OX=9606 GN=RRP15 PE=1 SV=2 | 1 |
| Q9Y3D2 | Methionine-R-sulfoxide reductase B2, mitochondrial OS=Homo sapiens OX=9606 GN=MSRB2 PE=1 SV=2 | 1 |
| Q9Y3D8 | Adenylate kinase isoenzyme 6 OS=Homo sapiens OX=9606 GN=AK6 PE=1 SV=1 | 1 |
| Q9Y3E0 | Vesicle transport protein GOT1B OS=Homo sapiens OX=9606 GN=GOLT1B PE=1 SV=1 | 1 |
| Q9Y3E5 | Peptidyl-tRNA hydrolase 2, mitochondrial OS=Homo sapiens OX=9606 GN=PTRH2 PE=1 SV=1 | 2 |
| Q9Y3S1 | Serine/threonine-protein kinase WNK2 OS=Homo sapiens OX=9606 GN=WNK2 PE=1 SV=4 | 1 |
| Q9Y4C1 | Lysine-specific demethylase 3A OS=Homo sapiens OX=9606 GN=KDM3A PE=1 SV=4 | 5 |
| Q9Y520 | Protein PRRC2C OS=Homo sapiens OX=9606 GN=PRRC2C PE=1 SV=4 | 1 |
| Q9Y5B8 | Nucleoside diphosphate kinase 7 OS=Homo sapiens OX=9606 GN=NME7 PE=1 SV=1 | 1 |
| Q9Y5J6 | Mitochondrial import inner membrane translocase subunit Tim10 B OS=Homo sapiens OX=9606 GN=TIMM10B PE=1 SV=1 | 1 |
| Q9Y5R5 | Doublesex- and mab-3-related transcription factor 2 OS=Homo sapiens OX=9606 GN=DMRT2 PE=1 SV=2 | 1 |
| Q9Y5U9 | Immediate early response 3-interacting protein 1 OS=Homo sapiens OX=9606 GN=IER3IP1 PE=1 SV=1 | 1 |
| Q9Y5W7 | Sorting nexin-14 OS=Homo sapiens OX=9606 GN=SNX14 PE=1 SV=3 | 1 |
| Q9Y624 | Junctional adhesion molecule A OS=Homo sapiens OX=9606 GN=F11R PE=1 SV=1 | 1 |
| Q9Y625 | Glypican-6 OS=Homo sapiens OX=9606 GN=GPC6 PE=1 SV=1 | 1 |
| Q9Y676 | 28S ribosomal protein S18b, mitochondrial OS=Homo sapiens OX=9606 GN=MRPS18B PE=1 SV=1 | 1 |
| Q9Y6M7 | Sodium bicarbonate cotransporter 3 OS=Homo sapiens OX=9606 GN=SLC4A7 PE=1 SV=2 | 4 |
| Q9Y6R4 | Mitogen-activated protein kinase kinase kinase 4 OS=Homo sapiens OX=9606 GN=MAP3K4 PE=1 SV=2 | 1 |
| Y-FGCZCont00324 | Q5XQN5 SWISS-PROT:Q5XQN5 (Bos taurus) Keratin, type II cytoskeletal 5 | 2 |
| A0A1B0GTI1 | Coiled-coil domain-containing protein 201 OS=Homo sapiens OX=9606 GN=CCDC201 PE=4 SV=1 | 1 |
| A8MYA2 | Uncharacterized protein CXorf49 OS=Homo sapiens OX=9606 GN=CXorf49B PE=2 SV=3 | 3 |
| O14531 | Dihydropyrimidinase-related protein 4 OS=Homo sapiens OX=9606 GN=DPYSL4 PE=1 SV=2 | 1 |
| O15031 | Plexin-B2 OS=Homo sapiens OX=9606 GN=PLXNB2 PE=1 SV=3 | 1 |
| O15084 | Serine/threonine-protein phosphatase 6 regulatory ankyrin repeat subunit A OS=Homo sapiens OX=9606 GN=ANKRD28 PE=1 SV=5 | 1 |
| O15232 | Matrilin-3 OS=Homo sapiens OX=9606 GN=MATN3 PE=1 SV=2 | 2 |
| O43323 | Desert hedgehog protein OS=Homo sapiens OX=9606 GN=DHH PE=1 SV=1 | 1 |
| O43598 | 2'-deoxynucleoside 5'-phosphate N-hydrolase 1 OS=Homo sapiens OX=9606 GN=DNPH1 PE=1 SV=1 | 2 |
| O43657 | Tetraspanin-6 OS=Homo sapiens OX=9606 GN=TSPAN6 PE=1 SV=1 | 1 |
| O75064 | DENN domain-containing protein 4B OS=Homo sapiens OX=9606 GN=DENND4B PE=1 SV=4 | 1 |
| O75533 | Splicing factor 3B subunit 1 OS=Homo sapiens OX=9606 GN=SF3B1 PE=1 SV=3 | 1 |
| O95670 | V-type proton ATPase subunit G 2 OS=Homo sapiens OX=9606 GN=ATP6V1G2 PE=1 SV=1 | 1 |
| P00739 | Haptoglobin-related protein OS=Homo sapiens OX=9606 GN=HPR PE=2 SV=2 | 1 |
| P00915 | Carbonic anhydrase 1 OS=Homo sapiens OX=9606 GN=CA1 PE=1 SV=2 | 2 |
| P06681 | Complement C2 OS=Homo sapiens OX=9606 GN=C2 PE=1 SV=2 | 1 |
| P08397 | Porphobilinogen deaminase OS=Homo sapiens OX=9606 GN=HMBS PE=1 SV=2 | 1 |
| P0DKL9 | ARL14 effector protein-like OS=Homo sapiens OX=9606 GN=ARL14EPL PE=4 SV=1 | 1 |
| P10242 | Transcriptional activator Myb OS=Homo sapiens OX=9606 GN=MYB PE=1 SV=2 | 1 |
| P13591 | Neural cell adhesion molecule 1 OS=Homo sapiens OX=9606 GN=NCAM1 PE=1 SV=3 | 1 |
| P14151 | L-selectin OS=Homo sapiens OX=9606 GN=SELL PE=1 SV=2 | 2 |
| P15407 | Fos-related antigen 1 OS=Homo sapiens OX=9606 GN=FOSL1 PE=1 SV=1 | 1 |
| P19525 | Interferon-induced, double-stranded RNA-activated protein kinase OS=Homo sapiens OX=9606 GN=EIF2AK2 PE=1 SV=2 | 1 |
| P19784 | Casein kinase II subunit alpha' OS=Homo sapiens OX=9606 GN=CSNK2A2 PE=1 SV=1 | 1 |
| P22223 | Cadherin-3 OS=Homo sapiens OX=9606 GN=CDH3 PE=1 SV=2 | 1 |
| P22748 | Carbonic anhydrase 4 OS=Homo sapiens OX=9606 GN=CA4 PE=1 SV=2 | 2 |
| P23469 | Receptor-type tyrosine-protein phosphatase epsilon OS=Homo sapiens OX=9606 GN=PTPRE PE=1 SV=1 | 1 |
| P27540 | Aryl hydrocarbon receptor nuclear translocator OS=Homo sapiens OX=9606 GN=ARNT PE=1 SV=1 | 2 |
| P29474 | Nitric oxide synthase, endothelial OS=Homo sapiens OX=9606 GN=NOS3 PE=1 SV=4 | 1 |
| P35611 | Alpha-adducin OS=Homo sapiens OX=9606 GN=ADD1 PE=1 SV=2 | 4 |
| P35711;P35712;Q9UN79 | Transcription factor SOX-5 OS=Homo sapiens OX=9606 GN=SOX5 PE=1 SV=3 | 1 |
| P39060 | Collagen alpha-1(XVIII) chain OS=Homo sapiens OX=9606 GN=COL18A1 PE=1 SV=5 | 1 |
| P42858 | Huntingtin OS=Homo sapiens OX=9606 GN=HTT PE=1 SV=2 | 1 |
| P49643 | DNA primase large subunit OS=Homo sapiens OX=9606 GN=PRIM2 PE=1 SV=2 | 1 |
| P49754 | Vacuolar protein sorting-associated protein 41 homolog OS=Homo sapiens OX=9606 GN=VPS41 PE=1 SV=3 | 1 |
| P51692 | Signal transducer and activator of transcription 5B OS=Homo sapiens OX=9606 GN=STAT5B PE=1 SV=2 | 1 |
| P51946 | Cyclin-H OS=Homo sapiens OX=9606 GN=CCNH PE=1 SV=1 | 1 |
| P51956 | Serine/threonine-protein kinase Nek3 OS=Homo sapiens OX=9606 GN=NEK3 PE=1 SV=2 | 1 |
| P53004 | Biliverdin reductase A OS=Homo sapiens OX=9606 GN=BLVRA PE=1 SV=2 | 1 |
| P53420 | Collagen alpha-4(IV) chain OS=Homo sapiens OX=9606 GN=COL4A4 PE=1 SV=3 | 1 |
| P54802 | Alpha-N-acetylglucosaminidase OS=Homo sapiens OX=9606 GN=NAGLU PE=1 SV=2 | 1 |
| P56962 | Syntaxin-17 OS=Homo sapiens OX=9606 GN=STX17 PE=1 SV=2 | 1 |
| P57735 | Ras-related protein Rab-25 OS=Homo sapiens OX=9606 GN=RAB25 PE=1 SV=2 | 1 |
| P81877 | Single-stranded DNA-binding protein 2 OS=Homo sapiens OX=9606 GN=SSBP2 PE=1 SV=2 | 1 |
| P82921 | 28S ribosomal protein S21, mitochondrial OS=Homo sapiens OX=9606 GN=MRPS21 PE=1 SV=3 | 1 |
| Q00839 | Heterogeneous nuclear ribonucleoprotein U OS=Homo sapiens OX=9606 GN=HNRNPU PE=1 SV=6 | 2 |
| Q05655 | Protein kinase C delta type OS=Homo sapiens OX=9606 GN=PRKCD PE=1 SV=2 | 2 |
| Q12912 | Inositol 1,4,5-triphosphate receptor associated 2 OS=Homo sapiens OX=9606 GN=IRAG2 PE=1 SV=3 | 3 |
| Q13740 | CD166 antigen OS=Homo sapiens OX=9606 GN=ALCAM PE=1 SV=2 | 1 |
| Q13873 | Bone morphogenetic protein receptor type-2 OS=Homo sapiens OX=9606 GN=BMPR2 PE=1 SV=2 | 1 |
| Q15058 | Kinesin-like protein KIF14 OS=Homo sapiens OX=9606 GN=KIF14 PE=1 SV=1 | 1 |
| Q15154 | Pericentriolar material 1 protein OS=Homo sapiens OX=9606 GN=PCM1 PE=1 SV=5 | 1 |
| Q15287 | RNA-binding protein with serine-rich domain 1 OS=Homo sapiens OX=9606 GN=RNPS1 PE=1 SV=1 | 1 |
| Q15293 | Reticulocalbin-1 OS=Homo sapiens OX=9606 GN=RCN1 PE=1 SV=1 | 2 |
| Q15750 | TGF-beta-activated kinase 1 and MAP3K7-binding protein 1 OS=Homo sapiens OX=9606 GN=TAB1 PE=1 SV=1 | 2 |
| Q16352 | Alpha-internexin OS=Homo sapiens OX=9606 GN=INA PE=1 SV=2 | 1 |
| Q2TAY7 | WD40 repeat-containing protein SMU1 OS=Homo sapiens OX=9606 GN=SMU1 PE=1 SV=2 | 1 |
| Q3KQU3 | MAP7 domain-containing protein 1 OS=Homo sapiens OX=9606 GN=MAP7D1 PE=1 SV=1 | 1 |
| Q3LI77 | Keratin-associated protein 13-4 OS=Homo sapiens OX=9606 GN=KRTAP13-4 PE=1 SV=1 | 2 |
| Q49AR2 | UPF0489 protein C5orf22 OS=Homo sapiens OX=9606 GN=C5orf22 PE=1 SV=2 | 2 |
| Q52LG2 | Keratin-associated protein 13-2 OS=Homo sapiens OX=9606 GN=KRTAP13-2 PE=1 SV=1 | 3 |
| Q58FG0 | Putative heat shock protein HSP 90-alpha A5 OS=Homo sapiens OX=9606 GN=HSP90AA5P PE=2 SV=1 | 1 |
| Q5JXA9 | Signal-regulatory protein beta-2 OS=Homo sapiens OX=9606 GN=SIRPB2 PE=2 SV=1 | 1 |
| Q5SZK8 | FRAS1-related extracellular matrix protein 2 OS=Homo sapiens OX=9606 GN=FREM2 PE=1 SV=3 | 1 |
| Q5T848 | Probable G-protein coupled receptor 158 OS=Homo sapiens OX=9606 GN=GPR158 PE=1 SV=1 | 1 |
| Q5T9L3 | Protein wntless homolog OS=Homo sapiens OX=9606 GN=WLS PE=1 SV=2 | 1 |
| Q5TB30;Q8WUY9 | DEP domain-containing protein 1A OS=Homo sapiens OX=9606 GN=DEPDC1 PE=1 SV=2 | 1 |
| Q6BDS2 | Bridge-like lipid transfer protein family member 3A OS=Homo sapiens OX=9606 GN=BLTP3A PE=1 SV=1 | 1 |
| Q6DT37 | Serine/threonine-protein kinase MRCK gamma OS=Homo sapiens OX=9606 GN=CDC42BPG PE=1 SV=2 | 1 |
| Q6NZ67 | Mitotic-spindle organizing protein 2B OS=Homo sapiens OX=9606 GN=MZT2B PE=1 SV=1 | 1 |
| Q6P1L8 | 39S ribosomal protein L14, mitochondrial OS=Homo sapiens OX=9606 GN=MRPL14 PE=1 SV=1 | 1 |
| Q6P4A7 | Sideroflexin-4 OS=Homo sapiens OX=9606 GN=SFXN4 PE=1 SV=1 | 1 |
| Q6PEV8 | Protein FAM199X OS=Homo sapiens OX=9606 GN=FAM199X PE=1 SV=1 | 1 |
| Q6ZMZ0 | E3 ubiquitin-protein ligase RNF19B OS=Homo sapiens OX=9606 GN=RNF19B PE=1 SV=2 | 1 |
| Q6ZN66 | Guanylate-binding protein 6 OS=Homo sapiens OX=9606 GN=GBP6 PE=2 SV=1 | 1 |
| Q6ZVH6 | Putative uncharacterized protein FLJ42569 OS=Homo sapiens OX=9606 PE=2 SV=1 | 1 |
| Q702N8 | Xin actin-binding repeat-containing protein 1 OS=Homo sapiens OX=9606 GN=XIRP1 PE=1 SV=1 | 1 |
| Q7Z3T8 | Zinc finger FYVE domain-containing protein 16 OS=Homo sapiens OX=9606 GN=ZFYVE16 PE=1 SV=3 | 1 |
| Q7Z4R8 | UPF0669 protein C6orf120 OS=Homo sapiens OX=9606 GN=C6orf120 PE=1 SV=1 | 1 |
| Q86SG5 | Protein S100-A7A OS=Homo sapiens OX=9606 GN=S100A7A PE=1 SV=3 | 3 |
| Q86VI3 | Ras GTPase-activating-like protein IQGAP3 OS=Homo sapiens OX=9606 GN=IQGAP3 PE=1 SV=2 | 1 |
| Q8IW52 | SLIT and NTRK-like protein 4 OS=Homo sapiens OX=9606 GN=SLITRK4 PE=1 SV=1 | 1 |
| Q8IY63 | Angiomotin-like protein 1 OS=Homo sapiens OX=9606 GN=AMOTL1 PE=1 SV=1 | 1 |
| Q8IY67 | Ribonucleoprotein PTB-binding 1 OS=Homo sapiens OX=9606 GN=RAVER1 PE=1 SV=1 | 1 |
| Q8IY85 | EF-hand calcium-binding domain-containing protein 13 OS=Homo sapiens OX=9606 GN=EFCAB13 PE=2 SV=2 | 1 |
| Q8IZH2 | 5'-3' exoribonuclease 1 OS=Homo sapiens OX=9606 GN=XRN1 PE=1 SV=1 | 1 |
| Q8IZI9 | Interferon lambda-3 OS=Homo sapiens OX=9606 GN=IFNL3 PE=1 SV=2 | 1 |
| Q8IZR5 | CKLF-like MARVEL transmembrane domain-containing protein 4 OS=Homo sapiens OX=9606 GN=CMTM4 PE=1 SV=1 | 1 |
| Q8IZU8 | Dermatan-sulfate epimerase-like protein OS=Homo sapiens OX=9606 GN=DSEL PE=2 SV=2 | 1 |
| Q8N143 | B-cell CLL/lymphoma 6 member B protein OS=Homo sapiens OX=9606 GN=BCL6B PE=2 SV=2 | 1 |
| Q8N5P1 | Zinc finger CCCH domain-containing protein 8 OS=Homo sapiens OX=9606 GN=ZC3H8 PE=1 SV=2 | 1 |
| Q8N8R5 | Mitochondrial protein C2orf69 OS=Homo sapiens OX=9606 GN=C2orf69 PE=1 SV=1 | 1 |
| Q8NA31 | Telomere repeats-binding bouquet formation protein 1 OS=Homo sapiens OX=9606 GN=TERB1 PE=1 SV=3 | 1 |
| Q8NE18 | Putative methyltransferase NSUN7 OS=Homo sapiens OX=9606 GN=NSUN7 PE=2 SV=4 | 1 |
| Q8TA86 | Retinitis pigmentosa 9 protein OS=Homo sapiens OX=9606 GN=RP9 PE=1 SV=2 | 1 |
| Q8TD19 | Serine/threonine-protein kinase Nek9 OS=Homo sapiens OX=9606 GN=NEK9 PE=1 SV=2 | 1 |
| Q8TDD5 | Mucolipin-3 OS=Homo sapiens OX=9606 GN=MCOLN3 PE=1 SV=1 | 1 |
| Q8TF66 | Leucine-rich repeat-containing protein 15 OS=Homo sapiens OX=9606 GN=LRRC15 PE=2 SV=2 | 2 |
| Q8WUS8 | Short-chain dehydrogenase/reductase family 42E member 1 OS=Homo sapiens OX=9606 GN=SDR42E1 PE=2 SV=2 | 1 |
| Q8WX92 | Negative elongation factor B OS=Homo sapiens OX=9606 GN=NELFB PE=1 SV=1 | 1 |
| Q8WY22 | BRI3-binding protein OS=Homo sapiens OX=9606 GN=BRI3BP PE=1 SV=1 | 1 |
| Q92537 | Sushi domain-containing protein 6 OS=Homo sapiens OX=9606 GN=SUSD6 PE=1 SV=1 | 1 |
| Q92692 | Nectin-2 OS=Homo sapiens OX=9606 GN=NECTIN2 PE=1 SV=1 | 1 |
| Q92797 | Symplekin OS=Homo sapiens OX=9606 GN=SYMPK PE=1 SV=2 | 1 |
| Q92917 | G-patch domain and KOW motifs-containing protein OS=Homo sapiens OX=9606 GN=GPKOW PE=1 SV=2 | 1 |
| Q96B45 | BLOC-1-related complex subunit 7 OS=Homo sapiens OX=9606 GN=BORCS7 PE=3 SV=2 | 2 |
| Q96B54 | Zinc finger protein 428 OS=Homo sapiens OX=9606 GN=ZNF428 PE=1 SV=2 | 1 |
| Q96EP1 | E3 ubiquitin-protein ligase CHFR OS=Homo sapiens OX=9606 GN=CHFR PE=1 SV=2 | 1 |
| Q96J42 | Thioredoxin domain-containing protein 15 OS=Homo sapiens OX=9606 GN=TXNDC15 PE=1 SV=1 | 1 |
| Q96MP8 | BTB/POZ domain-containing protein KCTD7 OS=Homo sapiens OX=9606 GN=KCTD7 PE=1 SV=1 | 1 |
| Q96P44 | Collagen alpha-1(XXI) chain OS=Homo sapiens OX=9606 GN=COL21A1 PE=2 SV=1 | 1 |
| Q99715 | Collagen alpha-1(XII) chain OS=Homo sapiens OX=9606 GN=COL12A1 PE=1 SV=2 | 1 |
| Q99755 | Phosphatidylinositol 4-phosphate 5-kinase type-1 alpha OS=Homo sapiens OX=9606 GN=PIP5K1A PE=1 SV=1 | 1 |
| Q99946 | Proline-rich transmembrane protein 1 OS=Homo sapiens OX=9606 GN=PRRT1 PE=1 SV=2 | 1 |
| Q99959 | Plakophilin-2 OS=Homo sapiens OX=9606 GN=PKP2 PE=1 SV=2 | 1 |
| Q9BQ67 | Glutamate-rich WD repeat-containing protein 1 OS=Homo sapiens OX=9606 GN=GRWD1 PE=1 SV=1 | 1 |
| Q9BTC0 | Death-inducer obliterator 1 OS=Homo sapiens OX=9606 GN=DIDO1 PE=1 SV=5 | 1 |
| Q9BTE6 | Alanyl-tRNA editing protein Aarsd1 OS=Homo sapiens OX=9606 GN=AARSD1 PE=1 SV=2 | 1 |
| Q9BTY7 | Protein HGH1 homolog OS=Homo sapiens OX=9606 GN=HGH1 PE=1 SV=1 | 1 |
| Q9BUB7 | Transmembrane protein 70, mitochondrial OS=Homo sapiens OX=9606 GN=TMEM70 PE=1 SV=2 | 1 |
| Q9BV86 | N-terminal Xaa-Pro-Lys N-methyltransferase 1 OS=Homo sapiens OX=9606 GN=NTMT1 PE=1 SV=3 | 1 |
| Q9BXM0 | Periaxin OS=Homo sapiens OX=9606 GN=PRX PE=1 SV=2 | 1 |
| Q9BXX0 | EMILIN-2 OS=Homo sapiens OX=9606 GN=EMILIN2 PE=1 SV=3 | 1 |
| Q9BYZ6 | Rho-related BTB domain-containing protein 2 OS=Homo sapiens OX=9606 GN=RHOBTB2 PE=1 SV=2 | 1 |
| Q9BZE4 | GTP-binding protein 4 OS=Homo sapiens OX=9606 GN=GTPBP4 PE=1 SV=3 | 1 |
| Q9C0G0 | Zinc finger protein 407 OS=Homo sapiens OX=9606 GN=ZNF407 PE=1 SV=2 | 1 |
| Q9GZM8;Q9NXR1 | Nuclear distribution protein nudE-like 1 OS=Homo sapiens OX=9606 GN=NDEL1 PE=1 SV=1 | 1 |
| Q9H082 | Ras-related protein Rab-33B OS=Homo sapiens OX=9606 GN=RAB33B PE=1 SV=1 | 1 |
| Q9H3A6 | PRO2179 OS=Homo sapiens OX=9606 PE=2 SV=1 | 1 |
| Q9H4G4 | Golgi-associated plant pathogenesis-related protein 1 OS=Homo sapiens OX=9606 GN=GLIPR2 PE=1 SV=3 | 1 |
| Q9H4K7 | Mitochondrial ribosome-associated GTPase 2 OS=Homo sapiens OX=9606 GN=MTG2 PE=1 SV=1 | 1 |
| Q9H6N6 | Putative uncharacterized protein MYH16 OS=Homo sapiens OX=9606 GN=MYH16 PE=1 SV=2 | 1 |
| Q9HAY6 | Beta,beta-carotene 15,15'-dioxygenase OS=Homo sapiens OX=9606 GN=BCO1 PE=1 SV=1 | 1 |
| Q9HBL0 | Tensin-1 OS=Homo sapiens OX=9606 GN=TNS1 PE=1 SV=3 | 1 |
| Q9HCQ5 | Polypeptide N-acetylgalactosaminyltransferase 9 OS=Homo sapiens OX=9606 GN=GALNT9 PE=1 SV=3 | 1 |
| Q9HCX3 | Zinc finger protein 304 OS=Homo sapiens OX=9606 GN=ZNF304 PE=1 SV=2 | 1 |
| Q9NPC3 | E3 ubiquitin-protein ligase CCNB1IP1 OS=Homo sapiens OX=9606 GN=CCNB1IP1 PE=1 SV=1 | 1 |
| Q9NPH3 | Interleukin-1 receptor accessory protein OS=Homo sapiens OX=9606 GN=IL1RAP PE=1 SV=2 | 1 |
| Q9NQC8 | Intraflagellar transport protein 46 homolog OS=Homo sapiens OX=9606 GN=IFT46 PE=1 SV=1 | 1 |
| Q9NQX4 | Unconventional myosin-Vc OS=Homo sapiens OX=9606 GN=MYO5C PE=1 SV=2 | 1 |
| Q9NRN7 | L-aminoadipate-semialdehyde dehydrogenase-phosphopantetheinyl transferase OS=Homo sapiens OX=9606 GN=AASDHPPT PE=1 SV=2 | 2 |
| Q9NT68 | Teneurin-2 OS=Homo sapiens OX=9606 GN=TENM2 PE=1 SV=3 | 1 |
| Q9NW82 | WD repeat-containing protein 70 OS=Homo sapiens OX=9606 GN=WDR70 PE=1 SV=1 | 1 |
| Q9NWX6 | Probable tRNA(His) guanylyltransferase OS=Homo sapiens OX=9606 GN=THG1L PE=1 SV=2 | 1 |
| Q9NYB0 | Telomeric repeat-binding factor 2-interacting protein 1 OS=Homo sapiens OX=9606 GN=TERF2IP PE=1 SV=1 | 1 |
| Q9NYP9 | Protein Mis18-alpha OS=Homo sapiens OX=9606 GN=MIS18A PE=1 SV=1 | 2 |
| Q9P035 | Very-long-chain (3R)-3-hydroxyacyl-CoA dehydratase 3 OS=Homo sapiens OX=9606 GN=HACD3 PE=1 SV=2 | 1 |
| Q9P260 | RAB11-binding protein RELCH OS=Homo sapiens OX=9606 GN=RELCH PE=1 SV=2 | 1 |
| Q9UBR1 | Beta-ureidopropionase OS=Homo sapiens OX=9606 GN=UPB1 PE=1 SV=1 | 2 |
| Q9UDX4 | SEC14-like protein 3 OS=Homo sapiens OX=9606 GN=SEC14L3 PE=1 SV=1 | 1 |
| Q9ULK4 | Mediator of RNA polymerase II transcription subunit 23 OS=Homo sapiens OX=9606 GN=MED23 PE=1 SV=2 | 1 |
| Q9UM13 | Anaphase-promoting complex subunit 10 OS=Homo sapiens OX=9606 GN=ANAPC10 PE=1 SV=1 | 1 |
| Q9UMS4 | Pre-mRNA-processing factor 19 OS=Homo sapiens OX=9606 GN=PRPF19 PE=1 SV=1 | 2 |
| Q9UNP9 | Peptidyl-prolyl cis-trans isomerase E OS=Homo sapiens OX=9606 GN=PPIE PE=1 SV=1 | 1 |
| Q9Y4E1 | WASH complex subunit 2C OS=Homo sapiens OX=9606 GN=WASHC2C PE=1 SV=4 | 1 |
| Q9Y5J9 | Mitochondrial import inner membrane translocase subunit Tim8 B OS=Homo sapiens OX=9606 GN=TIMM8B PE=1 SV=1 | 1 |
| Q9Y666 | Solute carrier family 12 member 7 OS=Homo sapiens OX=9606 GN=SLC12A7 PE=1 SV=3 | 1 |
| Y-FGCZCont00399 | A2VCT4 TREMBL:A2VCT4;Q0VDS4;Q6IFX4 Tax_Id=10090 Gene_Symbol=Krt39 keratin 39 | 1 |
| A0A0C4DGP1 | Salivary gland specific protein SAGSIN1 OS=Homo sapiens OX=9606 GN=SAGSIN1 PE=4 SV=1 | 1 |
| A6NGU5;P19440;P36268 | Putative glutathione hydrolase 3 proenzyme OS=Homo sapiens OX=9606 GN=GGT3P PE=5 SV=2 | 1 |
| A6NNA2 | Serine/arginine repetitive matrix protein 3 OS=Homo sapiens OX=9606 GN=SRRM3 PE=2 SV=4 | 1 |
| O15056 | Synaptojanin-2 OS=Homo sapiens OX=9606 GN=SYNJ2 PE=1 SV=3 | 1 |
| O15231 | Zinc finger protein 185 OS=Homo sapiens OX=9606 GN=ZNF185 PE=1 SV=3 | 1 |
| O15265 | Ataxin-7 OS=Homo sapiens OX=9606 GN=ATXN7 PE=1 SV=1 | 1 |
| O15304 | Apoptosis regulatory protein Siva OS=Homo sapiens OX=9606 GN=SIVA1 PE=1 SV=2 | 1 |
| O15427 | Monocarboxylate transporter 4 OS=Homo sapiens OX=9606 GN=SLC16A3 PE=1 SV=1 | 2 |
| O15514 | DNA-directed RNA polymerase II subunit RPB4 OS=Homo sapiens OX=9606 GN=POLR2D PE=1 SV=1 | 1 |
| O43252 | Bifunctional 3'-phosphoadenosine 5'-phosphosulfate synthase 1 OS=Homo sapiens OX=9606 GN=PAPSS1 PE=1 SV=2 | 1 |
| O43390 | Heterogeneous nuclear ribonucleoprotein R OS=Homo sapiens OX=9606 GN=HNRNPR PE=1 SV=1 | 1 |
| O43422 | 52 kDa repressor of the inhibitor of the protein kinase OS=Homo sapiens OX=9606 GN=THAP12 PE=1 SV=2 | 1 |
| O75339 | Cartilage intermediate layer protein 1 OS=Homo sapiens OX=9606 GN=CILP PE=1 SV=4 | 1 |
| O75419 | Cell division control protein 45 homolog OS=Homo sapiens OX=9606 GN=CDC45 PE=1 SV=1 | 1 |
| O75445 | Usherin OS=Homo sapiens OX=9606 GN=USH2A PE=1 SV=3 | 3 |
| O75467;Q6AW86 | Zinc finger protein 324A OS=Homo sapiens OX=9606 GN=ZNF324 PE=2 SV=1 | 1 |
| O75475 | PC4 and SFRS1-interacting protein OS=Homo sapiens OX=9606 GN=PSIP1 PE=1 SV=1 | 1 |
| O75604 | Ubiquitin carboxyl-terminal hydrolase 2 OS=Homo sapiens OX=9606 GN=USP2 PE=1 SV=2 | 1 |
| O75695 | Protein XRP2 OS=Homo sapiens OX=9606 GN=RP2 PE=1 SV=4 | 1 |
| O94993 | Transcription factor SOX-30 OS=Homo sapiens OX=9606 GN=SOX30 PE=1 SV=1 | 1 |
| O95376 | E3 ubiquitin-protein ligase ARIH2 OS=Homo sapiens OX=9606 GN=ARIH2 PE=1 SV=1 | 1 |
| O95619 | YEATS domain-containing protein 4 OS=Homo sapiens OX=9606 GN=YEATS4 PE=1 SV=1 | 1 |
| P01903 | HLA class II histocompatibility antigen, DR alpha chain OS=Homo sapiens OX=9606 GN=HLA-DRA PE=1 SV=2 | 1 |
| P09012 | U1 small nuclear ribonucleoprotein A OS=Homo sapiens OX=9606 GN=SNRPA PE=1 SV=3 | 1 |
| P0CL83 | Putative STAG3-like protein 1 OS=Homo sapiens OX=9606 GN=STAG3L1 PE=5 SV=1 | 1 |
| P10746 | Uroporphyrinogen-III synthase OS=Homo sapiens OX=9606 GN=UROS PE=1 SV=1 | 1 |
| P12111 | Collagen alpha-3(VI) chain OS=Homo sapiens OX=9606 GN=COL6A3 PE=1 SV=5 | 2 |
| P31152 | Mitogen-activated protein kinase 4 OS=Homo sapiens OX=9606 GN=MAPK4 PE=1 SV=2 | 1 |
| P42356 | Phosphatidylinositol 4-kinase alpha OS=Homo sapiens OX=9606 GN=PI4KA PE=1 SV=4 | 1 |
| P42695 | Condensin-2 complex subunit D3 OS=Homo sapiens OX=9606 GN=NCAPD3 PE=1 SV=2 | 1 |
| P49585;Q9Y5K3 | Choline-phosphate cytidylyltransferase A OS=Homo sapiens OX=9606 GN=PCYT1A PE=1 SV=2 | 1 |
| P52701 | DNA mismatch repair protein Msh6 OS=Homo sapiens OX=9606 GN=MSH6 PE=1 SV=2 | 1 |
| P62875 | DNA-directed RNA polymerases I, II, and III subunit RPABC5 OS=Homo sapiens OX=9606 GN=POLR2L PE=1 SV=1 | 1 |
| P68543 | UBX domain-containing protein 2A OS=Homo sapiens OX=9606 GN=UBXN2A PE=1 SV=1 | 1 |
| P98073 | Enteropeptidase OS=Homo sapiens OX=9606 GN=TMPRSS15 PE=1 SV=3 | 1 |
| Q04656 | Copper-transporting ATPase 1 OS=Homo sapiens OX=9606 GN=ATP7A PE=1 SV=4 | 1 |
| Q13114 | TNF receptor-associated factor 3 OS=Homo sapiens OX=9606 GN=TRAF3 PE=1 SV=2 | 1 |
| Q13683 | Integrin alpha-7 OS=Homo sapiens OX=9606 GN=ITGA7 PE=1 SV=3 | 2 |
| Q13868 | Exosome complex component RRP4 OS=Homo sapiens OX=9606 GN=EXOSC2 PE=1 SV=2 | 1 |
| Q14003 | Potassium voltage-gated channel subfamily C member 3 OS=Homo sapiens OX=9606 GN=KCNC3 PE=1 SV=3 | 1 |
| Q15080 | Neutrophil cytosol factor 4 OS=Homo sapiens OX=9606 GN=NCF4 PE=1 SV=2 | 1 |
| Q15697 | Zinc finger protein 174 OS=Homo sapiens OX=9606 GN=ZNF174 PE=1 SV=1 | 1 |
| Q19T08 | Endothelial cell-specific chemotaxis regulator OS=Homo sapiens OX=9606 GN=ECSCR PE=1 SV=1 | 1 |
| Q3MJ40 | Putative coiled-coil domain-containing protein 144B OS=Homo sapiens OX=9606 GN=CCDC144BP PE=5 SV=1 | 2 |
| Q5R3I4 | Tetratricopeptide repeat protein 38 OS=Homo sapiens OX=9606 GN=TTC38 PE=1 SV=1 | 2 |
| Q5U5X0 | Complex III assembly factor LYRM7 OS=Homo sapiens OX=9606 GN=LYRM7 PE=1 SV=1 | 1 |
| Q5VU65 | Nuclear pore membrane glycoprotein 210-like OS=Homo sapiens OX=9606 GN=NUP210L PE=2 SV=1 | 1 |
| Q6NUM9 | All-trans-retinol 13,14-reductase OS=Homo sapiens OX=9606 GN=RETSAT PE=1 SV=2 | 2 |
| Q6T310 | Ras-like protein family member 11A OS=Homo sapiens OX=9606 GN=RASL11A PE=1 SV=1 | 1 |
| Q6UVK1 | Chondroitin sulfate proteoglycan 4 OS=Homo sapiens OX=9606 GN=CSPG4 PE=1 SV=2 | 1 |
| Q6UXS9 | Inactive caspase-12 OS=Homo sapiens OX=9606 GN=CASP12 PE=2 SV=3 | 1 |
| Q86VY4 | Testis-specific Y-encoded-like protein 5 OS=Homo sapiens OX=9606 GN=TSPYL5 PE=1 SV=2 | 1 |
| Q8IX12 | Cell division cycle and apoptosis regulator protein 1 OS=Homo sapiens OX=9606 GN=CCAR1 PE=1 SV=2 | 1 |
| Q8IZD9 | Dedicator of cytokinesis protein 3 OS=Homo sapiens OX=9606 GN=DOCK3 PE=1 SV=1 | 1 |
| Q8N3C0 | Activating signal cointegrator 1 complex subunit 3 OS=Homo sapiens OX=9606 GN=ASCC3 PE=1 SV=3 | 2 |
| Q8N3K9 | Cardiomyopathy-associated protein 5 OS=Homo sapiens OX=9606 GN=CMYA5 PE=1 SV=3 | 1 |
| Q8N6H7 | ADP-ribosylation factor GTPase-activating protein 2 OS=Homo sapiens OX=9606 GN=ARFGAP2 PE=1 SV=1 | 1 |
| Q8N9N8 | Probable RNA-binding protein EIF1AD OS=Homo sapiens OX=9606 GN=EIF1AD PE=1 SV=1 | 1 |
| Q8NBP0 | Tetratricopeptide repeat protein 13 OS=Homo sapiens OX=9606 GN=TTC13 PE=2 SV=3 | 1 |
| Q8NEU8 | DCC-interacting protein 13-beta OS=Homo sapiens OX=9606 GN=APPL2 PE=1 SV=3 | 1 |
| Q8TBZ3 | WD repeat-containing protein 20 OS=Homo sapiens OX=9606 GN=WDR20 PE=1 SV=2 | 1 |
| Q8WTU0 | Protein DDI1 homolog 1 OS=Homo sapiens OX=9606 GN=DDI1 PE=1 SV=1 | 1 |
| Q8WXD5 | Gem-associated protein 6 OS=Homo sapiens OX=9606 GN=GEMIN6 PE=1 SV=1 | 1 |
| Q8WXI2 | Connector enhancer of kinase suppressor of ras 2 OS=Homo sapiens OX=9606 GN=CNKSR2 PE=1 SV=1 | 1 |
| Q8WZ42 | Titin OS=Homo sapiens OX=9606 GN=TTN PE=1 SV=4 | 4 |
| Q92508 | Piezo-type mechanosensitive ion channel component 1 OS=Homo sapiens OX=9606 GN=PIEZO1 PE=1 SV=4 | 1 |
| Q96BY6 | Dedicator of cytokinesis protein 10 OS=Homo sapiens OX=9606 GN=DOCK10 PE=1 SV=3 | 1 |
| Q96DY7 | Mdm2-binding protein OS=Homo sapiens OX=9606 GN=MTBP PE=1 SV=1 | 1 |
| Q96ED9 | Protein Hook homolog 2 OS=Homo sapiens OX=9606 GN=HOOK2 PE=1 SV=3 | 1 |
| Q96GX9 | Methylthioribulose-1-phosphate dehydratase OS=Homo sapiens OX=9606 GN=APIP PE=1 SV=1 | 1 |
| Q96HJ9 | Protein FMC1 homolog OS=Homo sapiens OX=9606 GN=FMC1 PE=1 SV=2 | 1 |
| Q96Q80 | Derlin-3 OS=Homo sapiens OX=9606 GN=DERL3 PE=1 SV=2 | 1 |
| Q96S82 | Ubiquitin-like protein 7 OS=Homo sapiens OX=9606 GN=UBL7 PE=1 SV=2 | 1 |
| Q9BQG1 | Synaptotagmin-3 OS=Homo sapiens OX=9606 GN=SYT3 PE=1 SV=1 | 1 |
| Q9BSE4 | Homocysteine-responsive endoplasmic reticulum-resident ubiquitin-like domain member 2 protein OS=Homo sapiens OX=9606 GN=HERPUD2 PE=1 SV=2 | 1 |
| Q9BVQ7 | Ribosome biogenesis protein SPATA5L1 OS=Homo sapiens OX=9606 GN=SPATA5L1 PE=1 SV=2 | 1 |
| Q9BW62 | Katanin p60 ATPase-containing subunit A-like 1 OS=Homo sapiens OX=9606 GN=KATNAL1 PE=1 SV=1 | 1 |
| Q9BYD2 | 39S ribosomal protein L9, mitochondrial OS=Homo sapiens OX=9606 GN=MRPL9 PE=1 SV=2 | 1 |
| Q9BZJ0 | Crooked neck-like protein 1 OS=Homo sapiens OX=9606 GN=CRNKL1 PE=1 SV=4 | 1 |
| Q9H0J4 | Glutamine-rich protein 2 OS=Homo sapiens OX=9606 GN=QRICH2 PE=1 SV=1 | 2 |
| Q9H813 | Proton-activated chloride channel OS=Homo sapiens OX=9606 GN=PACC1 PE=1 SV=1 | 1 |
| Q9NQV6 | PR domain zinc finger protein 10 OS=Homo sapiens OX=9606 GN=PRDM10 PE=1 SV=3 | 1 |
| Q9NUJ7 | PI-PLC X domain-containing protein 1 OS=Homo sapiens OX=9606 GN=PLCXD1 PE=2 SV=1 | 1 |
| Q9NYF8 | Bcl-2-associated transcription factor 1 OS=Homo sapiens OX=9606 GN=BCLAF1 PE=1 SV=2 | 1 |
| Q9P218 | Collagen alpha-1(XX) chain OS=Homo sapiens OX=9606 GN=COL20A1 PE=1 SV=4 | 1 |
| Q9P2N6 | KAT8 regulatory NSL complex subunit 3 OS=Homo sapiens OX=9606 GN=KANSL3 PE=1 SV=2 | 1 |
| Q9UBC5 | Unconventional myosin-Ia OS=Homo sapiens OX=9606 GN=MYO1A PE=1 SV=1 | 1 |
| Q9UK17 | Potassium voltage-gated channel subfamily D member 3 OS=Homo sapiens OX=9606 GN=KCND3 PE=1 SV=3 | 1 |
| Q9ULL0 | Acrosomal protein KIAA1210 OS=Homo sapiens OX=9606 GN=KIAA1210 PE=2 SV=3 | 1 |
| Q9Y6K0 | Choline/ethanolaminephosphotransferase 1 OS=Homo sapiens OX=9606 GN=CEPT1 PE=1 SV=1 | 1 |
| O00257 | E3 SUMO-protein ligase CBX4 OS=Homo sapiens OX=9606 GN=CBX4 PE=1 SV=3 | 1 |
| O15294 | UDP-N-acetylglucosamine--peptide N-acetylglucosaminyltransferase 110 kDa subunit OS=Homo sapiens OX=9606 GN=OGT PE=1 SV=3 | 1 |
| O75691 | Small subunit processome component 20 homolog OS=Homo sapiens OX=9606 GN=UTP20 PE=1 SV=3 | 1 |
| O94880 | PHD finger protein 14 OS=Homo sapiens OX=9606 GN=PHF14 PE=1 SV=3 | 1 |
| P19801 | Amiloride-sensitive amine oxidase [copper-containing] OS=Homo sapiens OX=9606 GN=AOC1 PE=1 SV=4 | 1 |
| P49023 | Paxillin OS=Homo sapiens OX=9606 GN=PXN PE=1 SV=3 | 1 |
| P53384 | Cytosolic Fe-S cluster assembly factor NUBP1 OS=Homo sapiens OX=9606 GN=NUBP1 PE=1 SV=2 | 1 |
| Q13029 | PR domain zinc finger protein 2 OS=Homo sapiens OX=9606 GN=PRDM2 PE=1 SV=3 | 1 |
| Q15124 | Phosphoglucomutase-like protein 5 OS=Homo sapiens OX=9606 GN=PGM5 PE=1 SV=2 | 1 |
| Q3B820 | Protein FAM161A OS=Homo sapiens OX=9606 GN=FAM161A PE=1 SV=2 | 1 |
| Q587I9 | Vesicle transport protein SFT2C OS=Homo sapiens OX=9606 GN=SFT2D3 PE=2 SV=1 | 1 |
| Q6IEE7 | Transmembrane protein 132E OS=Homo sapiens OX=9606 GN=TMEM132E PE=1 SV=2 | 1 |
| Q86UK7 | E3 ubiquitin-protein ligase ZNF598 OS=Homo sapiens OX=9606 GN=ZNF598 PE=1 SV=1 | 1 |
| Q8IVB4 | Sodium/hydrogen exchanger 9 OS=Homo sapiens OX=9606 GN=SLC9A9 PE=1 SV=1 | 1 |
| Q8WXE9 | Stonin-2 OS=Homo sapiens OX=9606 GN=STON2 PE=1 SV=1 | 1 |
| Q92772 | Cyclin-dependent kinase-like 2 OS=Homo sapiens OX=9606 GN=CDKL2 PE=1 SV=1 | 1 |
| Q96BN6 | Primary cilium assembly protein FAM149B1 OS=Homo sapiens OX=9606 GN=FAM149B1 PE=1 SV=2 | 1 |
| Q96NH3 | Protein broad-minded OS=Homo sapiens OX=9606 GN=TBC1D32 PE=1 SV=4 | 1 |
| Q9BUF5 | Tubulin beta-6 chain OS=Homo sapiens OX=9606 GN=TUBB6 PE=1 SV=1 | 1 |
| Q9BUH8 | Brain-enriched guanylate kinase-associated protein OS=Homo sapiens OX=9606 GN=BEGAIN PE=1 SV=1 | 1 |
| Q9C0E2 | Exportin-4 OS=Homo sapiens OX=9606 GN=XPO4 PE=1 SV=2 | 1 |
| Q9GZV1 | Ankyrin repeat domain-containing protein 2 OS=Homo sapiens OX=9606 GN=ANKRD2 PE=1 SV=3 | 1 |
| Q9H2U9 | Disintegrin and metalloproteinase domain-containing protein 7 OS=Homo sapiens OX=9606 GN=ADAM7 PE=1 SV=3 | 1 |
| Q9HAT2 | Sialate O-acetylesterase OS=Homo sapiens OX=9606 GN=SIAE PE=1 SV=1 | 1 |
| Q9HB19 | Pleckstrin homology domain-containing family A member 2 OS=Homo sapiens OX=9606 GN=PLEKHA2 PE=1 SV=2 | 1 |
| Q9NTM9 | Copper homeostasis protein cutC homolog OS=Homo sapiens OX=9606 GN=CUTC PE=1 SV=1 | 1 |
| Q9UEY8 | Gamma-adducin OS=Homo sapiens OX=9606 GN=ADD3 PE=1 SV=1 | 1 |
| Q9UHG0 | Doublecortin domain-containing protein 2 OS=Homo sapiens OX=9606 GN=DCDC2 PE=1 SV=2 | 1 |
| Q9ULJ1 | Protein BCAP OS=Homo sapiens OX=9606 GN=ODF2L PE=2 SV=2 | 1 |
| Y-FGCZCont00499 | sp_P19909_SPG2_STRSG Immunoglobulin G-binding protein G OS=Streptococcus sp. group G OX=1320 GN=spg PE=1 SV=1 | 2 |
| A0A494C086 | Putative speedy protein E21 OS=Homo sapiens OX=9606 GN=SPDYE21 PE=5 SV=1 | 1 |
| O00411 | DNA-directed RNA polymerase, mitochondrial OS=Homo sapiens OX=9606 GN=POLRMT PE=1 SV=2 | 1 |
| O15075 | Serine/threonine-protein kinase DCLK1 OS=Homo sapiens OX=9606 GN=DCLK1 PE=1 SV=2 | 1 |
| O43930 | Putative serine/threonine-protein kinase PRKY OS=Homo sapiens OX=9606 GN=PRKY PE=5 SV=1 | 1 |
| O60287 | Nucleolar pre-ribosomal-associated protein 1 OS=Homo sapiens OX=9606 GN=URB1 PE=1 SV=4 | 1 |
| O75400 | Pre-mRNA-processing factor 40 homolog A OS=Homo sapiens OX=9606 GN=PRPF40A PE=1 SV=2 | 1 |
| O75648 | Mitochondrial tRNA-specific 2-thiouridylase 1 OS=Homo sapiens OX=9606 GN=TRMU PE=1 SV=2 | 1 |
| O75694 | Nuclear pore complex protein Nup155 OS=Homo sapiens OX=9606 GN=NUP155 PE=1 SV=1 | 1 |
| O95171 | Sciellin OS=Homo sapiens OX=9606 GN=SCEL PE=1 SV=2 | 1 |
| P05783 | Keratin, type I cytoskeletal 18 OS=Homo sapiens OX=9606 GN=KRT18 PE=1 SV=2 | 1 |
| P08648 | Integrin alpha-5 OS=Homo sapiens OX=9606 GN=ITGA5 PE=1 SV=2 | 1 |
| P11388 | DNA topoisomerase 2-alpha OS=Homo sapiens OX=9606 GN=TOP2A PE=1 SV=3 | 1 |
| P13521 | Secretogranin-2 OS=Homo sapiens OX=9606 GN=SCG2 PE=1 SV=2 | 1 |
| P26038 | Moesin OS=Homo sapiens OX=9606 GN=MSN PE=1 SV=3 | 1 |
| P30305 | M-phase inducer phosphatase 2 OS=Homo sapiens OX=9606 GN=CDC25B PE=1 SV=2 | 1 |
| Q13454 | Tumor suppressor candidate 3 OS=Homo sapiens OX=9606 GN=TUSC3 PE=1 SV=1 | 1 |
| Q5VWP2 | Terminal nucleotidyltransferase 5C OS=Homo sapiens OX=9606 GN=TENT5C PE=1 SV=1 | 1 |
| Q6L8Q7 | 2',5'-phosphodiesterase 12 OS=Homo sapiens OX=9606 GN=PDE12 PE=1 SV=2 | 1 |
| Q6STE5 | SWI/SNF-related matrix-associated actin-dependent regulator of chromatin subfamily D member 3 OS=Homo sapiens OX=9606 GN=SMARCD3 PE=1 SV=1 | 1 |
| Q86Y79 | Probable peptidyl-tRNA hydrolase OS=Homo sapiens OX=9606 GN=PTRH1 PE=1 SV=1 | 1 |
| Q8NEE8 | Tetratricopeptide repeat protein 16 OS=Homo sapiens OX=9606 GN=TTC16 PE=2 SV=2 | 1 |
| Q96NW4 | Ankyrin repeat domain-containing protein 27 OS=Homo sapiens OX=9606 GN=ANKRD27 PE=1 SV=2 | 1 |
| Q96QF0 | Rab-3A-interacting protein OS=Homo sapiens OX=9606 GN=RAB3IP PE=1 SV=1 | 1 |
| Q9Y617 | Phosphoserine aminotransferase OS=Homo sapiens OX=9606 GN=PSAT1 PE=1 SV=2 | 1 |
| Q9Y6V7 | Probable ATP-dependent RNA helicase DDX49 OS=Homo sapiens OX=9606 GN=DDX49 PE=1 SV=1 | 1 |
| Y-FGCZCont00365 | Q3T052 TREMBL:Q3T052;Q5EA67 (Bos taurus) Inter-alpha (Globulin) inhibitor H4 | 1 |
| Y-FGCZCont00470 | ENSEMBL:ENSBTAP00000018574 (Bos taurus) 55 kDa protein | 1 |
| O94986 | Centrosomal protein of 152 kDa OS=Homo sapiens OX=9606 GN=CEP152 PE=1 SV=4 | 1 |
| P28062 | Proteasome subunit beta type-8 OS=Homo sapiens OX=9606 GN=PSMB8 PE=1 SV=3 | 1 |
| P45973 | Chromobox protein homolog 5 OS=Homo sapiens OX=9606 GN=CBX5 PE=1 SV=1 | 1 |
| P51610 | Host cell factor 1 OS=Homo sapiens OX=9606 GN=HCFC1 PE=1 SV=2 | 1 |
| Q13402 | Unconventional myosin-VIIa OS=Homo sapiens OX=9606 GN=MYO7A PE=1 SV=2 | 1 |
| Q86V48 | Leucine zipper protein 1 OS=Homo sapiens OX=9606 GN=LUZP1 PE=1 SV=2 | 1 |
| Q8N0V3 | Putative ribosome-binding factor A, mitochondrial OS=Homo sapiens OX=9606 GN=RBFA PE=1 SV=3 | 1 |
| Q9BZA7;Q9BZA8 | Protocadherin-11 X-linked OS=Homo sapiens OX=9606 GN=PCDH11X PE=2 SV=1 | 1 |
| Q9H5N1 | Rab GTPase-binding effector protein 2 OS=Homo sapiens OX=9606 GN=RABEP2 PE=1 SV=2 | 1 |
| Q9UBG3 | Cornulin OS=Homo sapiens OX=9606 GN=CRNN PE=1 SV=1 | 2 |
| Q9Y399 | 28S ribosomal protein S2, mitochondrial OS=Homo sapiens OX=9606 GN=MRPS2 PE=1 SV=1 | 1 |
| Q9Y5C1 | Angiopoietin-related protein 3 OS=Homo sapiens OX=9606 GN=ANGPTL3 PE=1 SV=1 | 1 |
| Q9Y5G7 | Protocadherin gamma-A6 OS=Homo sapiens OX=9606 GN=PCDHGA6 PE=2 SV=1 | 1 |
| O14746 | Telomerase reverse transcriptase OS=Homo sapiens OX=9606 GN=TERT PE=1 SV=1 | 1 |
| O77932 | Decapping and exoribonuclease protein OS=Homo sapiens OX=9606 GN=DXO PE=1 SV=2 | 1 |
| P06239 | Tyrosine-protein kinase Lck OS=Homo sapiens OX=9606 GN=LCK PE=1 SV=6 | 1 |
| P30039 | Phenazine biosynthesis-like domain-containing protein OS=Homo sapiens OX=9606 GN=PBLD PE=1 SV=2 | 1 |
| P46013 | Proliferation marker protein Ki-67 OS=Homo sapiens OX=9606 GN=MKI67 PE=1 SV=2 | 1 |
| P49796 | Regulator of G-protein signaling 3 OS=Homo sapiens OX=9606 GN=RGS3 PE=1 SV=2 | 1 |
| Q13573 | SNW domain-containing protein 1 OS=Homo sapiens OX=9606 GN=SNW1 PE=1 SV=1 | 1 |
| Q14210 | Lymphocyte antigen 6D OS=Homo sapiens OX=9606 GN=LY6D PE=1 SV=1 | 1 |
| Q15262 | Receptor-type tyrosine-protein phosphatase kappa OS=Homo sapiens OX=9606 GN=PTPRK PE=1 SV=2 | 1 |
| Q15345 | Leucine-rich repeat-containing protein 41 OS=Homo sapiens OX=9606 GN=LRRC41 PE=1 SV=3 | 1 |
| Q3B8N5 | Prospero homeobox protein 2 OS=Homo sapiens OX=9606 GN=PROX2 PE=2 SV=3 | 1 |
| Q3KNT7 | Putative NOL1/NOP2/Sun domain family member 5B OS=Homo sapiens OX=9606 GN=NSUN5P1 PE=5 SV=1 | 1 |
| Q53F19 | Nuclear cap-binding protein subunit 3 OS=Homo sapiens OX=9606 GN=NCBP3 PE=1 SV=2 | 1 |
| Q5TBK1 | NEDD4-binding protein 2-like 1 OS=Homo sapiens OX=9606 GN=N4BP2L1 PE=2 SV=1 | 1 |
| Q6QNY1 | Biogenesis of lysosome-related organelles complex 1 subunit 2 OS=Homo sapiens OX=9606 GN=BLOC1S2 PE=1 SV=1 | 1 |
| Q86TV6 | Tetratricopeptide repeat protein 7B OS=Homo sapiens OX=9606 GN=TTC7B PE=1 SV=3 | 1 |
| Q8WVX9 | Fatty acyl-CoA reductase 1 OS=Homo sapiens OX=9606 GN=FAR1 PE=1 SV=1 | 1 |
| Q8WYR4 | Radial spoke head 1 homolog OS=Homo sapiens OX=9606 GN=RSPH1 PE=1 SV=1 | 1 |
| Q96PK6 | RNA-binding protein 14 OS=Homo sapiens OX=9606 GN=RBM14 PE=1 SV=2 | 1 |
| Q9P0B6 | Coiled-coil domain-containing protein 167 OS=Homo sapiens OX=9606 GN=CCDC167 PE=1 SV=2 | 1 |
| Q9P2X0 | Dolichol-phosphate mannosyltransferase subunit 3 OS=Homo sapiens OX=9606 GN=DPM3 PE=1 SV=2 | 1 |
| Q9ULB1 | Neurexin-1 OS=Homo sapiens OX=9606 GN=NRXN1 PE=1 SV=1 | 1 |
| Q9UQR0 | Sex comb on midleg-like protein 2 OS=Homo sapiens OX=9606 GN=SCML2 PE=1 SV=1 | 1 |
| Q9Y3R5 | Protein dopey-2 OS=Homo sapiens OX=9606 GN=DOP1B PE=1 SV=5 | 1 |
| P14060;P26439 | 3 beta-hydroxysteroid dehydrogenase/Delta 5--4-isomerase type 1 OS=Homo sapiens OX=9606 GN=HSD3B1 PE=1 SV=2 | 1 |
| P78559 | Microtubule-associated protein 1A OS=Homo sapiens OX=9606 GN=MAP1A PE=1 SV=6 | 1 |
| Q15485 | Ficolin-2 OS=Homo sapiens OX=9606 GN=FCN2 PE=1 SV=2 | 1 |
| Q7RTS7 | Keratin, type II cytoskeletal 74 OS=Homo sapiens OX=9606 GN=KRT74 PE=1 SV=2 | 1 |
| Q8NFU7 | Methylcytosine dioxygenase TET1 OS=Homo sapiens OX=9606 GN=TET1 PE=1 SV=2 | 1 |
| Q96DT5 | Dynein axonemal heavy chain 11 OS=Homo sapiens OX=9606 GN=DNAH11 PE=1 SV=4 | 1 |
| Q96MZ0 | Ganglioside-induced differentiation-associated protein 1-like 1 OS=Homo sapiens OX=9606 GN=GDAP1L1 PE=2 SV=2 | 1 |
| Q9Y223 | Bifunctional UDP-N-acetylglucosamine 2-epimerase/N-acetylmannosamine kinase OS=Homo sapiens OX=9606 GN=GNE PE=1 SV=1 | 1 |
| P05060 | Secretogranin-1 OS=Homo sapiens OX=9606 GN=CHGB PE=1 SV=2 | 1 |
| Q07954 | Prolow-density lipoprotein receptor-related protein 1 OS=Homo sapiens OX=9606 GN=LRP1 PE=1 SV=2 | 1 |
| Q16799 | Reticulon-1 OS=Homo sapiens OX=9606 GN=RTN1 PE=1 SV=1 | 1 |
| Q4VNC0 | Probable cation-transporting ATPase 13A5 OS=Homo sapiens OX=9606 GN=ATP13A5 PE=2 SV=1 | 1 |
| Q7L8J4 | SH3 domain-binding protein 5-like OS=Homo sapiens OX=9606 GN=SH3BP5L PE=1 SV=1 | 1 |
| Q7Z589 | BRCA2-interacting transcriptional repressor EMSY OS=Homo sapiens OX=9606 GN=EMSY PE=1 SV=2 | 1 |
| Q8IUC4 | Rhophilin-2 OS=Homo sapiens OX=9606 GN=RHPN2 PE=1 SV=1 | 1 |
| Q99467 | CD180 antigen OS=Homo sapiens OX=9606 GN=CD180 PE=1 SV=2 | 1 |
| Q99683 | Mitogen-activated protein kinase kinase kinase 5 OS=Homo sapiens OX=9606 GN=MAP3K5 PE=1 SV=1 | 1 |
| Q9BYF1 | Angiotensin-converting enzyme 2 OS=Homo sapiens OX=9606 GN=ACE2 PE=1 SV=2 | 1 |
| Q9Y6D9 | Mitotic spindle assembly checkpoint protein MAD1 OS=Homo sapiens OX=9606 GN=MAD1L1 PE=1 SV=2 | 1 |
| A0A183 | Late cornified envelope protein 6A OS=Homo sapiens OX=9606 GN=LCE6A PE=3 SV=1 | 1 |
| A6NKK0 | Olfactory receptor 5H1 OS=Homo sapiens OX=9606 GN=OR5H1 PE=1 SV=1 | 1 |
| O00267 | Transcription elongation factor SPT5 OS=Homo sapiens OX=9606 GN=SUPT5H PE=1 SV=1 | 1 |
| O95834 | Echinoderm microtubule-associated protein-like 2 OS=Homo sapiens OX=9606 GN=EML2 PE=1 SV=1 | 3 |
| P30838 | Aldehyde dehydrogenase, dimeric NADP-preferring OS=Homo sapiens OX=9606 GN=ALDH3A1 PE=1 SV=3 | 1 |
| Q13591 | Semaphorin-5A OS=Homo sapiens OX=9606 GN=SEMA5A PE=1 SV=3 | 1 |
| Q5VWQ0 | Lysine-specific demethylase 9 OS=Homo sapiens OX=9606 GN=RSBN1 PE=1 SV=2 | 1 |
| Q68BL8 | Olfactomedin-like protein 2B OS=Homo sapiens OX=9606 GN=OLFML2B PE=2 SV=2 | 1 |
| Q6ZMP0 | Thrombospondin type-1 domain-containing protein 4 OS=Homo sapiens OX=9606 GN=THSD4 PE=1 SV=2 | 1 |
| Q6ZNF0 | Acid phosphatase type 7 OS=Homo sapiens OX=9606 GN=ACP7 PE=2 SV=2 | 1 |
| Q8TEU8 | WAP, Kazal, immunoglobulin, Kunitz and NTR domain-containing protein 2 OS=Homo sapiens OX=9606 GN=WFIKKN2 PE=1 SV=1 | 1 |
| Q9BYG8 | Gasdermin-C OS=Homo sapiens OX=9606 GN=GSDMC PE=1 SV=3 | 1 |
| Q9H094 | Neuroblastoma breakpoint family member 3 OS=Homo sapiens OX=9606 GN=NBPF3 PE=2 SV=1 | 1 |
| Q9H4Y5 | Glutathione S-transferase omega-2 OS=Homo sapiens OX=9606 GN=GSTO2 PE=1 SV=1 | 1 |
| Q9H6B9 | Epoxide hydrolase 3 OS=Homo sapiens OX=9606 GN=EPHX3 PE=1 SV=1 | 1 |
| Q9HA90 | EF-hand and coiled-coil domain-containing protein 1 OS=Homo sapiens OX=9606 GN=EFCC1 PE=2 SV=2 | 1 |
| Q9NSK0 | Kinesin light chain 4 OS=Homo sapiens OX=9606 GN=KLC4 PE=1 SV=3 | 1 |

| **Supplementary Table 8: Filtered proteins in MII oocytes (3884) used in further statistical analysis** | |  |
| --- | --- | --- |
| **Name** | **description** | **nrPeptides** |
| ERFL | ETS domain-containing transcription factor ERF-like OS=Homo sapiens OX=9606 GN=ERFL PE=3 SV=1 | 1 |
| LINC00615 | Putative uncharacterized protein encoded by LINC00615 OS=Homo sapiens OX=9606 GN=LINC00615 PE=5 SV=1 | 1 |
| MED23 | Mediator of RNA polymerase II transcription subunit 23 OS=Homo sapiens OX=9606 GN=MED23 PE=1 SV=2 | 1 |
| CDC42BPG | Serine/threonine-protein kinase MRCK gamma OS=Homo sapiens OX=9606 GN=CDC42BPG PE=1 SV=2 | 1 |
| PIP5K1A | Phosphatidylinositol 4-phosphate 5-kinase type-1 alpha OS=Homo sapiens OX=9606 GN=PIP5K1A PE=1 SV=1 | 1 |
| BRI3BP | BRI3-binding protein OS=Homo sapiens OX=9606 GN=BRI3BP PE=1 SV=1 | 1 |
| HGH1 | Protein HGH1 homolog OS=Homo sapiens OX=9606 GN=HGH1 PE=1 SV=1 | 1 |
| PPIE | Peptidyl-prolyl cis-trans isomerase E OS=Homo sapiens OX=9606 GN=PPIE PE=1 SV=1 | 1 |
| HPX | Hemopexin OS=Homo sapiens OX=9606 GN=HPX PE=1 SV=2 | 27 |
| TMEM179B | Transmembrane protein 179B OS=Homo sapiens OX=9606 GN=TMEM179B PE=1 SV=1 | 1 |
| HP | Haptoglobin OS=Homo sapiens OX=9606 GN=HP PE=1 SV=1 | 30 |
| UTP20 | Small subunit processome component 20 homolog OS=Homo sapiens OX=9606 GN=UTP20 PE=1 SV=3 | 1 |
| RYR3 | Ryanodine receptor 3 OS=Homo sapiens OX=9606 GN=RYR3 PE=1 SV=3 | 1 |
| DOCK9 | Dedicator of cytokinesis protein 9 OS=Homo sapiens OX=9606 GN=DOCK9 PE=1 SV=2 | 1 |
| AHSG | Alpha-2-HS-glycoprotein OS=Homo sapiens OX=9606 GN=AHSG PE=1 SV=2 | 9 |
| SMARCA4 | Transcription activator BRG1 OS=Homo sapiens OX=9606 GN=SMARCA4 PE=1 SV=2 | 1 |
| -.25 | Zinc finger TRAF-type-containing protein 1 OS=Homo sapiens OX=9606 GN=ZFTRAF1 PE=2 SV=1 | 1 |
| PRIM2 | DNA primase large subunit OS=Homo sapiens OX=9606 GN=PRIM2 PE=1 SV=2 | 1 |
| FBXL5 | F-box/LRR-repeat protein 5 OS=Homo sapiens OX=9606 GN=FBXL5 PE=1 SV=2 | 1 |
| PIEZO1 | Piezo-type mechanosensitive ion channel component 1 OS=Homo sapiens OX=9606 GN=PIEZO1 PE=1 SV=4 | 1 |
| SIRPB2 | Signal-regulatory protein beta-2 OS=Homo sapiens OX=9606 GN=SIRPB2 PE=2 SV=1 | 1 |
| ATP6V1G2 | V-type proton ATPase subunit G 2 OS=Homo sapiens OX=9606 GN=ATP6V1G2 PE=1 SV=1 | 1 |
| ALB | Albumin OS=Homo sapiens OX=9606 GN=ALB PE=1 SV=2 | 83 |
| PCNX4 | Pecanex-like protein 4 OS=Homo sapiens OX=9606 GN=PCNX4 PE=1 SV=4 | 1 |
| TMEM143 | Transmembrane protein 143 OS=Homo sapiens OX=9606 GN=TMEM143 PE=1 SV=1 | 2 |
| LRG1 | Leucine-rich alpha-2-glycoprotein OS=Homo sapiens OX=9606 GN=LRG1 PE=1 SV=2 | 9 |
| GALE | UDP-glucose 4-epimerase OS=Homo sapiens OX=9606 GN=GALE PE=1 SV=2 | 1 |
| PKP2 | Plakophilin-2 OS=Homo sapiens OX=9606 GN=PKP2 PE=1 SV=2 | 1 |
| TMEM97 | Sigma intracellular receptor 2 OS=Homo sapiens OX=9606 GN=TMEM97 PE=1 SV=1 | 1 |
| AFM | Afamin OS=Homo sapiens OX=9606 GN=AFM PE=1 SV=1 | 31 |
| SLC27A1 | Long-chain fatty acid transport protein 1 OS=Homo sapiens OX=9606 GN=SLC27A1 PE=1 SV=1 | 1 |
| TXNRD2 | Thioredoxin reductase 2, mitochondrial OS=Homo sapiens OX=9606 GN=TXNRD2 PE=1 SV=3 | 1 |
| CA4 | Carbonic anhydrase 4 OS=Homo sapiens OX=9606 GN=CA4 PE=1 SV=2 | 2 |
| NECTIN2 | Nectin-2 OS=Homo sapiens OX=9606 GN=NECTIN2 PE=1 SV=1 | 1 |
| NEK9 | Serine/threonine-protein kinase Nek9 OS=Homo sapiens OX=9606 GN=NEK9 PE=1 SV=2 | 1 |
| DCHS1 | Protocadherin-16 OS=Homo sapiens OX=9606 GN=DCHS1 PE=1 SV=1 | 2 |
| A1BG | Alpha-1B-glycoprotein OS=Homo sapiens OX=9606 GN=A1BG PE=1 SV=4 | 22 |
| PRRT1 | Proline-rich transmembrane protein 1 OS=Homo sapiens OX=9606 GN=PRRT1 PE=1 SV=2 | 1 |
| BCO1 | Beta,beta-carotene 15,15'-dioxygenase OS=Homo sapiens OX=9606 GN=BCO1 PE=1 SV=1 | 1 |
| MRPS16 | 28S ribosomal protein S16, mitochondrial OS=Homo sapiens OX=9606 GN=MRPS16 PE=1 SV=1 | 2 |
| CNGA3 | Cyclic nucleotide-gated cation channel alpha-3 OS=Homo sapiens OX=9606 GN=CNGA3 PE=1 SV=2 | 1 |
| PGLYRP2 | N-acetylmuramoyl-L-alanine amidase OS=Homo sapiens OX=9606 GN=PGLYRP2 PE=1 SV=1 | 14 |
| APOH | Beta-2-glycoprotein 1 OS=Homo sapiens OX=9606 GN=APOH PE=1 SV=3 | 14 |
| MYH6 | Myosin-6 OS=Homo sapiens OX=9606 GN=MYH6 PE=1 SV=5 | 1 |
| TERB1 | Telomere repeats-binding bouquet formation protein 1 OS=Homo sapiens OX=9606 GN=TERB1 PE=1 SV=3 | 1 |
| MZT2B | Mitotic-spindle organizing protein 2B OS=Homo sapiens OX=9606 GN=MZT2B PE=1 SV=1 | 1 |
| LONP2 | Lon protease homolog 2, peroxisomal OS=Homo sapiens OX=9606 GN=LONP2 PE=1 SV=1 | 1 |
| SUSD6 | Sushi domain-containing protein 6 OS=Homo sapiens OX=9606 GN=SUSD6 PE=1 SV=1 | 1 |
| ANKRD18A | Ankyrin repeat domain-containing protein 18A OS=Homo sapiens OX=9606 GN=ANKRD18A PE=2 SV=3 | 1 |
| ARL6IP1 | ADP-ribosylation factor-like protein 6-interacting protein 1 OS=Homo sapiens OX=9606 GN=ARL6IP1 PE=1 SV=2 | 1 |
| KYAT1 | Kynurenine--oxoglutarate transaminase 1 OS=Homo sapiens OX=9606 GN=KYAT1 PE=1 SV=1 | 1 |
| WDHD1 | WD repeat and HMG-box DNA-binding protein 1 OS=Homo sapiens OX=9606 GN=WDHD1 PE=1 SV=1 | 1 |
| SERPING1 | Plasma protease C1 inhibitor OS=Homo sapiens OX=9606 GN=SERPING1 PE=1 SV=2 | 16 |
| IFNL3 | Interferon lambda-3 OS=Homo sapiens OX=9606 GN=IFNL3 PE=1 SV=2 | 1 |
| MYO5C | Unconventional myosin-Vc OS=Homo sapiens OX=9606 GN=MYO5C PE=1 SV=2 | 1 |
| EP400 | E1A-binding protein p400 OS=Homo sapiens OX=9606 GN=EP400 PE=1 SV=4 | 1 |
| PLEKHB2 | Pleckstrin homology domain-containing family B member 2 OS=Homo sapiens OX=9606 GN=PLEKHB2 PE=1 SV=1 | 1 |
| GLB1L2 | Beta-galactosidase-1-like protein 2 OS=Homo sapiens OX=9606 GN=GLB1L2 PE=1 SV=1 | 1 |
| RAB15 | Ras-related protein Rab-15 OS=Homo sapiens OX=9606 GN=RAB15 PE=1 SV=1 | 1 |
| MOS | Proto-oncogene serine/threonine-protein kinase mos OS=Homo sapiens OX=9606 GN=MOS PE=1 SV=1 | 1 |
| ZWINT | ZW10 interactor OS=Homo sapiens OX=9606 GN=ZWINT PE=1 SV=2 | 1 |
| COX6A1 | Cytochrome c oxidase subunit 6A1, mitochondrial OS=Homo sapiens OX=9606 GN=COX6A1 PE=1 SV=4 | 1 |
| H2BU1 | Histone H2B type 3-B OS=Homo sapiens OX=9606 GN=H2BC26 PE=1 SV=3 | 1 |
| GPC5 | Glypican-5 OS=Homo sapiens OX=9606 GN=GPC5 PE=1 SV=1 | 1 |
| DENND4B | DENN domain-containing protein 4B OS=Homo sapiens OX=9606 GN=DENND4B PE=1 SV=4 | 1 |
| FOXD4L6 | Forkhead box protein D4-like 6 OS=Homo sapiens OX=9606 GN=FOXD4L6 PE=1 SV=2 | 1 |
| DEPDC1 | DEP domain-containing protein 1A OS=Homo sapiens OX=9606 GN=DEPDC1 PE=1 SV=2 | 1 |
| ZNF518A | Zinc finger protein 518A OS=Homo sapiens OX=9606 GN=ZNF518A PE=1 SV=2 | 2 |
| TNS1 | Tensin-1 OS=Homo sapiens OX=9606 GN=TNS1 PE=1 SV=3 | 1 |
| TINAGL1 | Tubulointerstitial nephritis antigen-like OS=Homo sapiens OX=9606 GN=TINAGL1 PE=1 SV=1 | 3 |
| ZFHX2 | Zinc finger homeobox protein 2 OS=Homo sapiens OX=9606 GN=ZFHX2 PE=1 SV=3 | 1 |
| CMTM4 | CKLF-like MARVEL transmembrane domain-containing protein 4 OS=Homo sapiens OX=9606 GN=CMTM4 PE=1 SV=1 | 1 |
| PEX13 | Peroxisomal membrane protein PEX13 OS=Homo sapiens OX=9606 GN=PEX13 PE=1 SV=2 | 2 |
| BEGAIN | Brain-enriched guanylate kinase-associated protein OS=Homo sapiens OX=9606 GN=BEGAIN PE=1 SV=1 | 1 |
| ARIH2 | E3 ubiquitin-protein ligase ARIH2 OS=Homo sapiens OX=9606 GN=ARIH2 PE=1 SV=1 | 1 |
| PTPRU | Receptor-type tyrosine-protein phosphatase U OS=Homo sapiens OX=9606 GN=PTPRU PE=1 SV=2 | 1 |
| TTR | Transthyretin OS=Homo sapiens OX=9606 GN=TTR PE=1 SV=1 | 12 |
| IFT46 | Intraflagellar transport protein 46 homolog OS=Homo sapiens OX=9606 GN=IFT46 PE=1 SV=1 | 1 |
| HBB | Hemoglobin subunit beta OS=Homo sapiens OX=9606 GN=HBB PE=1 SV=2 | 13 |
| OTUD6B | Deubiquitinase OTUD6B OS=Homo sapiens OX=9606 GN=OTUD6B PE=1 SV=1 | 2 |
| TMEM258 | Transmembrane protein 258 OS=Homo sapiens OX=9606 GN=TMEM258 PE=1 SV=1 | 1 |
| GAS2L1 | GAS2-like protein 1 OS=Homo sapiens OX=9606 GN=GAS2L1 PE=1 SV=2 | 1 |
| ORM2 | Alpha-1-acid glycoprotein 2 OS=Homo sapiens OX=9606 GN=ORM2 PE=1 SV=2 | 8 |
| RIC1 | Guanine nucleotide exchange factor subunit RIC1 OS=Homo sapiens OX=9606 GN=RIC1 PE=1 SV=2 | 1 |
| THYN1 | Thymocyte nuclear protein 1 OS=Homo sapiens OX=9606 GN=THYN1 PE=1 SV=1 | 1 |
| FAM199X | Protein FAM199X OS=Homo sapiens OX=9606 GN=FAM199X PE=1 SV=1 | 1 |
| TSPAN6 | Tetraspanin-6 OS=Homo sapiens OX=9606 GN=TSPAN6 PE=1 SV=1 | 1 |
| PAFAH2 | Platelet-activating factor acetylhydrolase 2, cytoplasmic OS=Homo sapiens OX=9606 GN=PAFAH2 PE=1 SV=1 | 2 |
| TRPM3 | Transient receptor potential cation channel subfamily M member 3 OS=Homo sapiens OX=9606 GN=TRPM3 PE=2 SV=4 | 1 |
| AHNAK2 | Protein AHNAK2 OS=Homo sapiens OX=9606 GN=AHNAK2 PE=1 SV=2 | 1 |
| SRP19 | Signal recognition particle 19 kDa protein OS=Homo sapiens OX=9606 GN=SRP19 PE=1 SV=3 | 2 |
| TTC38 | Tetratricopeptide repeat protein 38 OS=Homo sapiens OX=9606 GN=TTC38 PE=1 SV=1 | 2 |
| AKR1E2 | 1,5-anhydro-D-fructose reductase OS=Homo sapiens OX=9606 GN=AKR1E2 PE=1 SV=2 | 1 |
| SFXN4 | Sideroflexin-4 OS=Homo sapiens OX=9606 GN=SFXN4 PE=1 SV=1 | 1 |
| MRPL21 | 39S ribosomal protein L21, mitochondrial OS=Homo sapiens OX=9606 GN=MRPL21 PE=1 SV=2 | 2 |
| ISCA1 | Iron-sulfur cluster assembly 1 homolog, mitochondrial OS=Homo sapiens OX=9606 GN=ISCA1 PE=1 SV=1 | 2 |
| BAG5 | BAG family molecular chaperone regulator 5 OS=Homo sapiens OX=9606 GN=BAG5 PE=1 SV=1 | 2 |
| UHRF1BP1 | Bridge-like lipid transfer protein family member 3A OS=Homo sapiens OX=9606 GN=BLTP3A PE=1 SV=1 | 1 |
| MMAB | Corrinoid adenosyltransferase MMAB OS=Homo sapiens OX=9606 GN=MMAB PE=1 SV=1 | 3 |
| PARPBP | PCNA-interacting partner OS=Homo sapiens OX=9606 GN=PARPBP PE=1 SV=3 | 1 |
| CRNKL1 | Crooked neck-like protein 1 OS=Homo sapiens OX=9606 GN=CRNKL1 PE=1 SV=4 | 1 |
| ORM1 | Alpha-1-acid glycoprotein 1 OS=Homo sapiens OX=9606 GN=ORM1 PE=1 SV=2 | 13 |
| FAM162A | Protein FAM162A OS=Homo sapiens OX=9606 GN=FAM162A PE=1 SV=2 | 1 |
| MYL1 | Myosin light chain 1/3, skeletal muscle isoform OS=Homo sapiens OX=9606 GN=MYL1 PE=1 SV=3 | 1 |
| HSBP1 | Heat shock factor-binding protein 1 OS=Homo sapiens OX=9606 GN=HSBP1 PE=1 SV=1 | 3 |
| MECP2 | Methyl-CpG-binding protein 2 OS=Homo sapiens OX=9606 GN=MECP2 PE=1 SV=1 | 1 |
| VPS37C | Vacuolar protein sorting-associated protein 37C OS=Homo sapiens OX=9606 GN=VPS37C PE=1 SV=2 | 2 |
| RDH13 | Retinol dehydrogenase 13 OS=Homo sapiens OX=9606 GN=RDH13 PE=1 SV=2 | 1 |
| HACD3 | Very-long-chain (3R)-3-hydroxyacyl-CoA dehydratase 3 OS=Homo sapiens OX=9606 GN=HACD3 PE=1 SV=2 | 1 |
| SFI1 | Protein SFI1 homolog OS=Homo sapiens OX=9606 GN=SFI1 PE=1 SV=2 | 1 |
| NTMT1 | N-terminal Xaa-Pro-Lys N-methyltransferase 1 OS=Homo sapiens OX=9606 GN=NTMT1 PE=1 SV=3 | 1 |
| HPCAL1 | Hippocalcin-like protein 1 OS=Homo sapiens OX=9606 GN=HPCAL1 PE=1 SV=3 | 1 |
| IGKV3-7 | Probable non-functional immunoglobulin kappa variable 3-7 OS=Homo sapiens OX=9606 GN=IGKV3-7 PE=1 SV=1 | 1 |
| RNF34 | E3 ubiquitin-protein ligase RNF34 OS=Homo sapiens OX=9606 GN=RNF34 PE=1 SV=1 | 5 |
| BCL2L10 | Bcl-2-like protein 10 OS=Homo sapiens OX=9606 GN=BCL2L10 PE=1 SV=3 | 3 |
| UBE2A | Ubiquitin-conjugating enzyme E2 A OS=Homo sapiens OX=9606 GN=UBE2A PE=1 SV=2 | 1 |
| CHDH | Choline dehydrogenase, mitochondrial OS=Homo sapiens OX=9606 GN=CHDH PE=1 SV=2 | 1 |
| ANKRD28 | Serine/threonine-protein phosphatase 6 regulatory ankyrin repeat subunit A OS=Homo sapiens OX=9606 GN=ANKRD28 PE=1 SV=5 | 1 |
| MFSD10 | Major facilitator superfamily domain-containing protein 10 OS=Homo sapiens OX=9606 GN=MFSD10 PE=1 SV=1 | 3 |
| CEND1 | Cell cycle exit and neuronal differentiation protein 1 OS=Homo sapiens OX=9606 GN=CEND1 PE=1 SV=1 | 3 |
| STEEP1 | STING ER exit protein OS=Homo sapiens OX=9606 GN=STEEP1 PE=1 SV=1 | 1 |
| MGA | MAX gene-associated protein OS=Homo sapiens OX=9606 GN=MGA PE=1 SV=4 | 2 |
| KBTBD2 | Kelch repeat and BTB domain-containing protein 2 OS=Homo sapiens OX=9606 GN=KBTBD2 PE=1 SV=2 | 1 |
| U2AF1 | Splicing factor U2AF 35 kDa subunit OS=Homo sapiens OX=9606 GN=U2AF1 PE=1 SV=3 | 1 |
| HSPA13 | Heat shock 70 kDa protein 13 OS=Homo sapiens OX=9606 GN=HSPA13 PE=1 SV=1 | 1 |
| DMAC2L | ATP synthase subunit s, mitochondrial OS=Homo sapiens OX=9606 GN=DMAC2L PE=1 SV=3 | 7 |
| PRAP1 | Proline-rich acidic protein 1 OS=Homo sapiens OX=9606 GN=PRAP1 PE=1 SV=2 | 2 |
| GARNL3 | GTPase-activating Rap/Ran-GAP domain-like protein 3 OS=Homo sapiens OX=9606 GN=GARNL3 PE=2 SV=2 | 1 |
| SF3A3 | Splicing factor 3A subunit 3 OS=Homo sapiens OX=9606 GN=SF3A3 PE=1 SV=1 | 2 |
| PPWD1 | Peptidylprolyl isomerase domain and WD repeat-containing protein 1 OS=Homo sapiens OX=9606 GN=PPWD1 PE=1 SV=1 | 1 |
| EIF2B1 | Translation initiation factor eIF-2B subunit alpha OS=Homo sapiens OX=9606 GN=EIF2B1 PE=1 SV=1 | 1 |
| VPS35L | VPS35 endosomal protein-sorting factor-like OS=Homo sapiens OX=9606 GN=VPS35L PE=1 SV=2 | 4 |
| BABAM2 | BRISC and BRCA1-A complex member 2 OS=Homo sapiens OX=9606 GN=BABAM2 PE=1 SV=2 | 1 |
| SELENOF | Selenoprotein F OS=Homo sapiens OX=9606 GN=SELENOF PE=1 SV=4 | 1 |
| DDAH1 | N(G),N(G)-dimethylarginine dimethylaminohydrolase 1 OS=Homo sapiens OX=9606 GN=DDAH1 PE=1 SV=3 | 17 |
| SDR42E1 | Short-chain dehydrogenase/reductase family 42E member 1 OS=Homo sapiens OX=9606 GN=SDR42E1 PE=2 SV=2 | 1 |
| NAGLU | Alpha-N-acetylglucosaminidase OS=Homo sapiens OX=9606 GN=NAGLU PE=1 SV=2 | 1 |
| OOSP4A | Oocyte-secreted protein 4A OS=Homo sapiens OX=9606 GN=OOSP4A PE=3 SV=1 | 4 |
| PPP1CC | Serine/threonine-protein phosphatase PP1-gamma catalytic subunit OS=Homo sapiens OX=9606 GN=PPP1CC PE=1 SV=1 | 1 |
| KDM3A | Lysine-specific demethylase 3A OS=Homo sapiens OX=9606 GN=KDM3A PE=1 SV=4 | 5 |
| MED14 | Mediator of RNA polymerase II transcription subunit 14 OS=Homo sapiens OX=9606 GN=MED14 PE=1 SV=2 | 1 |
| TASOR | Protein TASOR OS=Homo sapiens OX=9606 GN=TASOR PE=1 SV=3 | 1 |
| SMYD3 | Histone-lysine N-methyltransferase SMYD3 OS=Homo sapiens OX=9606 GN=SMYD3 PE=1 SV=4 | 1 |
| CEPT1 | Choline/ethanolaminephosphotransferase 1 OS=Homo sapiens OX=9606 GN=CEPT1 PE=1 SV=1 | 1 |
| PKHD1L1 | Fibrocystin-L OS=Homo sapiens OX=9606 GN=PKHD1L1 PE=2 SV=2 | 1 |
| -.27 | Transmembrane protein LOC653160 OS=Homo sapiens OX=9606 PE=2 SV=2 | 1 |
| MAGEL2 | MAGE-like protein 2 OS=Homo sapiens OX=9606 GN=MAGEL2 PE=1 SV=2 | 1 |
| RGS2 | Regulator of G-protein signaling 2 OS=Homo sapiens OX=9606 GN=RGS2 PE=1 SV=1 | 5 |
| MADD | MAP kinase-activating death domain protein OS=Homo sapiens OX=9606 GN=MADD PE=1 SV=2 | 3 |
| MACROD1 | ADP-ribose glycohydrolase MACROD1 OS=Homo sapiens OX=9606 GN=MACROD1 PE=1 SV=2 | 1 |
| SMAP2 | Stromal membrane-associated protein 2 OS=Homo sapiens OX=9606 GN=SMAP2 PE=1 SV=1 | 1 |
| MKRN2 | E3 ubiquitin-protein ligase makorin-2 OS=Homo sapiens OX=9606 GN=MKRN2 PE=1 SV=2 | 1 |
| TMX2 | Thioredoxin-related transmembrane protein 2 OS=Homo sapiens OX=9606 GN=TMX2 PE=1 SV=1 | 1 |
| AMBP | Protein AMBP OS=Homo sapiens OX=9606 GN=AMBP PE=1 SV=1 | 11 |
| C7orf50 | Uncharacterized protein C7orf50 OS=Homo sapiens OX=9606 GN=C7orf50 PE=1 SV=1 | 1 |
| FAM174B | Membrane protein FAM174B OS=Homo sapiens OX=9606 GN=FAM174B PE=2 SV=2 | 1 |
| GCA | Grancalcin OS=Homo sapiens OX=9606 GN=GCA PE=1 SV=2 | 6 |
| C2orf69 | Mitochondrial protein C2orf69 OS=Homo sapiens OX=9606 GN=C2orf69 PE=1 SV=1 | 1 |
| CCZ1B | Vacuolar fusion protein CCZ1 homolog B OS=Homo sapiens OX=9606 GN=CCZ1B PE=1 SV=1 | 14 |
| OLFML1 | Olfactomedin-like protein 1 OS=Homo sapiens OX=9606 GN=OLFML1 PE=1 SV=2 | 1 |
| SMPD3 | Sphingomyelin phosphodiesterase 3 OS=Homo sapiens OX=9606 GN=SMPD3 PE=1 SV=1 | 2 |
| CASC3 | Protein CASC3 OS=Homo sapiens OX=9606 GN=CASC3 PE=1 SV=2 | 3 |
| SFXN2 | Sideroflexin-2 OS=Homo sapiens OX=9606 GN=SFXN2 PE=1 SV=2 | 2 |
| WEE2 | Wee1-like protein kinase 2 OS=Homo sapiens OX=9606 GN=WEE2 PE=1 SV=2 | 26 |
| PRKG2 | cGMP-dependent protein kinase 2 OS=Homo sapiens OX=9606 GN=PRKG2 PE=1 SV=1 | 1 |
| GTPBP4 | GTP-binding protein 4 OS=Homo sapiens OX=9606 GN=GTPBP4 PE=1 SV=3 | 1 |
| PHF5A | PHD finger-like domain-containing protein 5A OS=Homo sapiens OX=9606 GN=PHF5A PE=1 SV=1 | 1 |
| VPS53 | Vacuolar protein sorting-associated protein 53 homolog OS=Homo sapiens OX=9606 GN=VPS53 PE=1 SV=2 | 4 |
| PCCA | Propionyl-CoA carboxylase alpha chain, mitochondrial OS=Homo sapiens OX=9606 GN=PCCA PE=1 SV=4 | 2 |
| KIF11 | Kinesin-like protein KIF11 OS=Homo sapiens OX=9606 GN=KIF11 PE=1 SV=2 | 2 |
| OSBPL9 | Oxysterol-binding protein-related protein 9 OS=Homo sapiens OX=9606 GN=OSBPL9 PE=1 SV=2 | 7 |
| RAB3A | Ras-related protein Rab-3A OS=Homo sapiens OX=9606 GN=RAB3A PE=1 SV=1 | 2 |
| MRPL42 | 39S ribosomal protein L42, mitochondrial OS=Homo sapiens OX=9606 GN=MRPL42 PE=1 SV=1 | 1 |
| FKBP14 | Peptidyl-prolyl cis-trans isomerase FKBP14 OS=Homo sapiens OX=9606 GN=FKBP14 PE=1 SV=1 | 1 |
| COL18A1 | Collagen alpha-1(XVIII) chain OS=Homo sapiens OX=9606 GN=COL18A1 PE=1 SV=5 | 1 |
| HSP90AA2P | Heat shock protein HSP 90-alpha A2 OS=Homo sapiens OX=9606 GN=HSP90AA2P PE=1 SV=2 | 1 |
| C1RL | Complement C1r subcomponent-like protein OS=Homo sapiens OX=9606 GN=C1RL PE=1 SV=2 | 5 |
| PRC1 | Protein regulator of cytokinesis 1 OS=Homo sapiens OX=9606 GN=PRC1 PE=1 SV=2 | 8 |
| MNS1 | Meiosis-specific nuclear structural protein 1 OS=Homo sapiens OX=9606 GN=MNS1 PE=1 SV=2 | 1 |
| ADAMTS9 | A disintegrin and metalloproteinase with thrombospondin motifs 9 OS=Homo sapiens OX=9606 GN=ADAMTS9 PE=1 SV=4 | 1 |
| ABCC1 | Multidrug resistance-associated protein 1 OS=Homo sapiens OX=9606 GN=ABCC1 PE=1 SV=3 | 2 |
| GSTP1 | Glutathione S-transferase P OS=Homo sapiens OX=9606 GN=GSTP1 PE=1 SV=2 | 14 |
| TOMM7 | Mitochondrial import receptor subunit TOM7 homolog OS=Homo sapiens OX=9606 GN=TOMM7 PE=1 SV=1 | 3 |
| ESCO2 | N-acetyltransferase ESCO2 OS=Homo sapiens OX=9606 GN=ESCO2 PE=1 SV=1 | 2 |
| GSK3A | Glycogen synthase kinase-3 alpha OS=Homo sapiens OX=9606 GN=GSK3A PE=1 SV=2 | 2 |
| CCDC7 | Coiled-coil domain-containing protein 7 OS=Homo sapiens OX=9606 GN=CCDC7 PE=1 SV=3 | 3 |
| SERPINA7 | Thyroxine-binding globulin OS=Homo sapiens OX=9606 GN=SERPINA7 PE=1 SV=2 | 10 |
| RPS6KB1 | Ribosomal protein S6 kinase beta-1 OS=Homo sapiens OX=9606 GN=RPS6KB1 PE=1 SV=2 | 2 |
| RS1 | Retinoschisin OS=Homo sapiens OX=9606 GN=RS1 PE=1 SV=2 | 1 |
| RGPD3 | RanBP2-like and GRIP domain-containing protein 3 OS=Homo sapiens OX=9606 GN=RGPD3 PE=3 SV=2 | 1 |
| LGMN | Legumain OS=Homo sapiens OX=9606 GN=LGMN PE=1 SV=1 | 2 |
| SEC22A | Vesicle-trafficking protein SEC22a OS=Homo sapiens OX=9606 GN=SEC22A PE=1 SV=1 | 1 |
| LRRC37A2 | Leucine-rich repeat-containing protein 37A2 OS=Homo sapiens OX=9606 GN=LRRC37A2 PE=2 SV=2 | 1 |
| HIP1 | Huntingtin-interacting protein 1 OS=Homo sapiens OX=9606 GN=HIP1 PE=1 SV=5 | 5 |
| TIMM17B | Mitochondrial import inner membrane translocase subunit Tim17-B OS=Homo sapiens OX=9606 GN=TIMM17B PE=1 SV=1 | 1 |
| DIAPH3 | Protein diaphanous homolog 3 OS=Homo sapiens OX=9606 GN=DIAPH3 PE=1 SV=4 | 3 |
| DAD1 | Dolichyl-diphosphooligosaccharide--protein glycosyltransferase subunit DAD1 OS=Homo sapiens OX=9606 GN=DAD1 PE=1 SV=3 | 1 |
| SMAD9 | Mothers against decapentaplegic homolog 9 OS=Homo sapiens OX=9606 GN=SMAD9 PE=1 SV=1 | 1 |
| CYP2S1 | Cytochrome P450 2S1 OS=Homo sapiens OX=9606 GN=CYP2S1 PE=1 SV=2 | 3 |
| MAN2C1 | Alpha-mannosidase 2C1 OS=Homo sapiens OX=9606 GN=MAN2C1 PE=1 SV=1 | 3 |
| CDK7 | Cyclin-dependent kinase 7 OS=Homo sapiens OX=9606 GN=CDK7 PE=1 SV=1 | 6 |
| BCKDHB | 2-oxoisovalerate dehydrogenase subunit beta, mitochondrial OS=Homo sapiens OX=9606 GN=BCKDHB PE=1 SV=2 | 2 |
| ATP5MF | ATP synthase subunit f, mitochondrial OS=Homo sapiens OX=9606 GN=ATP5MF PE=1 SV=3 | 2 |
| CMPK2 | UMP-CMP kinase 2, mitochondrial OS=Homo sapiens OX=9606 GN=CMPK2 PE=1 SV=3 | 3 |
| RCN2 | Reticulocalbin-2 OS=Homo sapiens OX=9606 GN=RCN2 PE=1 SV=1 | 15 |
| C6orf120 | UPF0669 protein C6orf120 OS=Homo sapiens OX=9606 GN=C6orf120 PE=1 SV=1 | 1 |
| FTL | Ferritin light chain OS=Homo sapiens OX=9606 GN=FTL PE=1 SV=2 | 12 |
| GPAT4 | Glycerol-3-phosphate acyltransferase 4 OS=Homo sapiens OX=9606 GN=GPAT4 PE=1 SV=1 | 2 |
| IRAG2 | Inositol 1,4,5-triphosphate receptor associated 2 OS=Homo sapiens OX=9606 GN=IRAG2 PE=1 SV=3 | 3 |
| SNX14 | Sorting nexin-14 OS=Homo sapiens OX=9606 GN=SNX14 PE=1 SV=3 | 1 |
| STX17 | Syntaxin-17 OS=Homo sapiens OX=9606 GN=STX17 PE=1 SV=2 | 1 |
| BAP18 | Chromatin complexes subunit BAP18 OS=Homo sapiens OX=9606 GN=BAP18 PE=1 SV=1 | 3 |
| RRN3 | RNA polymerase I-specific transcription initiation factor RRN3 OS=Homo sapiens OX=9606 GN=RRN3 PE=1 SV=1 | 1 |
| COL4A1 | Collagen alpha-1(IV) chain OS=Homo sapiens OX=9606 GN=COL4A1 PE=1 SV=4 | 5 |
| ATG2B | Autophagy-related protein 2 homolog B OS=Homo sapiens OX=9606 GN=ATG2B PE=1 SV=5 | 2 |
| ILRUN | Protein ILRUN OS=Homo sapiens OX=9606 GN=ILRUN PE=1 SV=2 | 1 |
| CENPS | Centromere protein S OS=Homo sapiens OX=9606 GN=CENPS PE=1 SV=1 | 3 |
| WDR20 | WD repeat-containing protein 20 OS=Homo sapiens OX=9606 GN=WDR20 PE=1 SV=2 | 1 |
| YIPF2 | Protein YIPF2 OS=Homo sapiens OX=9606 GN=YIPF2 PE=1 SV=1 | 1 |
| DDHD2 | Phospholipase DDHD2 OS=Homo sapiens OX=9606 GN=DDHD2 PE=1 SV=2 | 2 |
| DNMT1 | DNA (cytosine-5)-methyltransferase 1 OS=Homo sapiens OX=9606 GN=DNMT1 PE=1 SV=2 | 70 |
| ING3 | Inhibitor of growth protein 3 OS=Homo sapiens OX=9606 GN=ING3 PE=1 SV=2 | 2 |
| ATP5PF | ATP synthase-coupling factor 6, mitochondrial OS=Homo sapiens OX=9606 GN=ATP5PF PE=1 SV=1 | 3 |
| SNX24 | Sorting nexin-24 OS=Homo sapiens OX=9606 GN=SNX24 PE=1 SV=1 | 1 |
| MAPRE2 | Microtubule-associated protein RP/EB family member 2 OS=Homo sapiens OX=9606 GN=MAPRE2 PE=1 SV=1 | 1 |
| SLC29A1 | Equilibrative nucleoside transporter 1 OS=Homo sapiens OX=9606 GN=SLC29A1 PE=1 SV=3 | 3 |
| DYNC2I1 | Cytoplasmic dynein 2 intermediate chain 1 OS=Homo sapiens OX=9606 GN=DYNC2I1 PE=1 SV=3 | 1 |
| EXOC5 | Exocyst complex component 5 OS=Homo sapiens OX=9606 GN=EXOC5 PE=1 SV=1 | 1 |
| ADO | 2-aminoethanethiol dioxygenase OS=Homo sapiens OX=9606 GN=ADO PE=1 SV=2 | 2 |
| KLHL33 | Kelch-like protein 33 OS=Homo sapiens OX=9606 GN=KLHL33 PE=4 SV=2 | 1 |
| APOC1 | Apolipoprotein C-I OS=Homo sapiens OX=9606 GN=APOC1 PE=1 SV=1 | 3 |
| CCNI | Cyclin-I OS=Homo sapiens OX=9606 GN=CCNI PE=1 SV=1 | 2 |
| VASN | Vasorin OS=Homo sapiens OX=9606 GN=VASN PE=1 SV=1 | 1 |
| DNPH1 | 2'-deoxynucleoside 5'-phosphate N-hydrolase 1 OS=Homo sapiens OX=9606 GN=DNPH1 PE=1 SV=1 | 2 |
| AKIRIN2 | Akirin-2 OS=Homo sapiens OX=9606 GN=AKIRIN2 PE=1 SV=2 | 5 |
| PNPLA4 | Patatin-like phospholipase domain-containing protein 4 OS=Homo sapiens OX=9606 GN=PNPLA4 PE=1 SV=3 | 2 |
| PCP4 | Calmodulin regulator protein PCP4 OS=Homo sapiens OX=9606 GN=PCP4 PE=1 SV=3 | 1 |
| C12orf57 | Protein C10 OS=Homo sapiens OX=9606 GN=C12orf57 PE=1 SV=1 | 1 |
| DERL1 | Derlin-1 OS=Homo sapiens OX=9606 GN=DERL1 PE=1 SV=1 | 2 |
| SERTAD2 | SERTA domain-containing protein 2 OS=Homo sapiens OX=9606 GN=SERTAD2 PE=1 SV=1 | 2 |
| ACTL8 | Actin-like protein 8 OS=Homo sapiens OX=9606 GN=ACTL8 PE=1 SV=1 | 19 |
| UBE2C | Ubiquitin-conjugating enzyme E2 C OS=Homo sapiens OX=9606 GN=UBE2C PE=1 SV=1 | 5 |
| TFB2M | Dimethyladenosine transferase 2, mitochondrial OS=Homo sapiens OX=9606 GN=TFB2M PE=1 SV=1 | 1 |
| SEC24A | Protein transport protein Sec24A OS=Homo sapiens OX=9606 GN=SEC24A PE=1 SV=2 | 2 |
| CHMP2A | Charged multivesicular body protein 2a OS=Homo sapiens OX=9606 GN=CHMP2A PE=1 SV=1 | 1 |
| DAW1 | Dynein assembly factor with WDR repeat domains 1 OS=Homo sapiens OX=9606 GN=DAW1 PE=1 SV=1 | 5 |
| MOGS | Mannosyl-oligosaccharide glucosidase OS=Homo sapiens OX=9606 GN=MOGS PE=1 SV=5 | 2 |
| APOA2 | Apolipoprotein A-II OS=Homo sapiens OX=9606 GN=APOA2 PE=1 SV=1 | 7 |
| SUCLG2 | Succinate--CoA ligase [GDP-forming] subunit beta, mitochondrial OS=Homo sapiens OX=9606 GN=SUCLG2 PE=1 SV=2 | 5 |
| LIPA | Lysosomal acid lipase/cholesteryl ester hydrolase OS=Homo sapiens OX=9606 GN=LIPA PE=1 SV=2 | 1 |
| HIP1R | Huntingtin-interacting protein 1-related protein OS=Homo sapiens OX=9606 GN=HIP1R PE=1 SV=2 | 3 |
| EXOSC3 | Exosome complex component RRP40 OS=Homo sapiens OX=9606 GN=EXOSC3 PE=1 SV=3 | 2 |
| MRM2 | rRNA methyltransferase 2, mitochondrial OS=Homo sapiens OX=9606 GN=MRM2 PE=1 SV=1 | 2 |
| AGPAT3 | 1-acyl-sn-glycerol-3-phosphate acyltransferase gamma OS=Homo sapiens OX=9606 GN=AGPAT3 PE=1 SV=1 | 2 |
| MRPS27 | 28S ribosomal protein S27, mitochondrial OS=Homo sapiens OX=9606 GN=MRPS27 PE=1 SV=3 | 2 |
| VPS37B | Vacuolar protein sorting-associated protein 37B OS=Homo sapiens OX=9606 GN=VPS37B PE=1 SV=1 | 2 |
| GNA11 | Guanine nucleotide-binding protein subunit alpha-11 OS=Homo sapiens OX=9606 GN=GNA11 PE=1 SV=2 | 2 |
| TIMM29 | Mitochondrial import inner membrane translocase subunit Tim29 OS=Homo sapiens OX=9606 GN=TIMM29 PE=1 SV=2 | 1 |
| RTTN | Rotatin OS=Homo sapiens OX=9606 GN=RTTN PE=1 SV=3 | 1 |
| SDHAF4 | Succinate dehydrogenase assembly factor 4, mitochondrial OS=Homo sapiens OX=9606 GN=SDHAF4 PE=1 SV=1 | 2 |
| SPECC1 | Cytospin-B OS=Homo sapiens OX=9606 GN=SPECC1 PE=1 SV=1 | 2 |
| YEATS4 | YEATS domain-containing protein 4 OS=Homo sapiens OX=9606 GN=YEATS4 PE=1 SV=1 | 1 |
| NDC1 | Nucleoporin NDC1 OS=Homo sapiens OX=9606 GN=NDC1 PE=1 SV=2 | 3 |
| ATG4B | Cysteine protease ATG4B OS=Homo sapiens OX=9606 GN=ATG4B PE=1 SV=2 | 2 |
| THUMPD3 | tRNA (guanine(6)-N2)-methyltransferase THUMP3 OS=Homo sapiens OX=9606 GN=THUMPD3 PE=1 SV=1 | 1 |
| CKS1B | Cyclin-dependent kinases regulatory subunit 1 OS=Homo sapiens OX=9606 GN=CKS1B PE=1 SV=1 | 2 |
| CETN3 | Centrin-3 OS=Homo sapiens OX=9606 GN=CETN3 PE=1 SV=2 | 2 |
| STRIP1 | Striatin-interacting protein 1 OS=Homo sapiens OX=9606 GN=STRIP1 PE=1 SV=1 | 2 |
| C15orf61 | Uncharacterized protein C15orf61 OS=Homo sapiens OX=9606 GN=C15orf61 PE=2 SV=2 | 4 |
| GINS2 | DNA replication complex GINS protein PSF2 OS=Homo sapiens OX=9606 GN=GINS2 PE=1 SV=1 | 1 |
| PPOX | Protoporphyrinogen oxidase OS=Homo sapiens OX=9606 GN=PPOX PE=1 SV=1 | 13 |
| C2orf76 | UPF0538 protein C2orf76 OS=Homo sapiens OX=9606 GN=C2orf76 PE=1 SV=3 | 3 |
| PACS2 | Phosphofurin acidic cluster sorting protein 2 OS=Homo sapiens OX=9606 GN=PACS2 PE=1 SV=3 | 1 |
| TAB1 | TGF-beta-activated kinase 1 and MAP3K7-binding protein 1 OS=Homo sapiens OX=9606 GN=TAB1 PE=1 SV=1 | 2 |
| DARS2 | Aspartate--tRNA ligase, mitochondrial OS=Homo sapiens OX=9606 GN=DARS2 PE=1 SV=1 | 3 |
| BCL2L1 | Bcl-2-like protein 1 OS=Homo sapiens OX=9606 GN=BCL2L1 PE=1 SV=1 | 3 |
| CNNM3 | Metal transporter CNNM3 OS=Homo sapiens OX=9606 GN=CNNM3 PE=1 SV=1 | 2 |
| TRAPPC2 | Trafficking protein particle complex subunit 2 OS=Homo sapiens OX=9606 GN=TRAPPC2 PE=1 SV=1 | 1 |
| MICAL2 | [F-actin]-monooxygenase MICAL2 OS=Homo sapiens OX=9606 GN=MICAL2 PE=1 SV=2 | 1 |
| SLC2A1 | Solute carrier family 2, facilitated glucose transporter member 1 OS=Homo sapiens OX=9606 GN=SLC2A1 PE=1 SV=2 | 3 |
| GMNN | Geminin OS=Homo sapiens OX=9606 GN=GMNN PE=1 SV=1 | 1 |
| AIDA | Axin interactor, dorsalization-associated protein OS=Homo sapiens OX=9606 GN=AIDA PE=1 SV=1 | 4 |
| ACCSL | Probable inactive 1-aminocyclopropane-1-carboxylate synthase-like protein 2 OS=Homo sapiens OX=9606 GN=ACCSL PE=2 SV=3 | 29 |
| HTRA1 | Serine protease HTRA1 OS=Homo sapiens OX=9606 GN=HTRA1 PE=1 SV=1 | 7 |
| LRRC20 | Leucine-rich repeat-containing protein 20 OS=Homo sapiens OX=9606 GN=LRRC20 PE=1 SV=1 | 3 |
| OTUD3 | OTU domain-containing protein 3 OS=Homo sapiens OX=9606 GN=OTUD3 PE=1 SV=1 | 1 |
| GGPS1 | Geranylgeranyl pyrophosphate synthase OS=Homo sapiens OX=9606 GN=GGPS1 PE=1 SV=1 | 3 |
| COL14A1 | Collagen alpha-1(XIV) chain OS=Homo sapiens OX=9606 GN=COL14A1 PE=1 SV=3 | 3 |
| CDNF | Cerebral dopamine neurotrophic factor OS=Homo sapiens OX=9606 GN=CDNF PE=1 SV=2 | 1 |
| ADGRL4 | Adhesion G protein-coupled receptor L4 OS=Homo sapiens OX=9606 GN=ADGRL4 PE=1 SV=3 | 1 |
| ALDH1B1 | Aldehyde dehydrogenase X, mitochondrial OS=Homo sapiens OX=9606 GN=ALDH1B1 PE=1 SV=4 | 3 |
| ITIH4 | Inter-alpha-trypsin inhibitor heavy chain H4 OS=Homo sapiens OX=9606 GN=ITIH4 PE=1 SV=4 | 15 |
| SYDE2 | Rho GTPase-activating protein SYDE2 OS=Homo sapiens OX=9606 GN=SYDE2 PE=1 SV=2 | 1 |
| TUT4 | Terminal uridylyltransferase 4 OS=Homo sapiens OX=9606 GN=TUT4 PE=1 SV=3 | 2 |
| HYDIN | Hydrocephalus-inducing protein homolog OS=Homo sapiens OX=9606 GN=HYDIN PE=1 SV=3 | 1 |
| CPQ | Carboxypeptidase Q OS=Homo sapiens OX=9606 GN=CPQ PE=1 SV=1 | 4 |
| QTRT2 | Queuine tRNA-ribosyltransferase accessory subunit 2 OS=Homo sapiens OX=9606 GN=QTRT2 PE=1 SV=1 | 1 |
| CYCS | Cytochrome c OS=Homo sapiens OX=9606 GN=CYCS PE=1 SV=2 | 7 |
| WDR37 | WD repeat-containing protein 37 OS=Homo sapiens OX=9606 GN=WDR37 PE=1 SV=2 | 2 |
| DCAF4L1 | DDB1- and CUL4-associated factor 4-like protein 1 OS=Homo sapiens OX=9606 GN=DCAF4L1 PE=2 SV=1 | 2 |
| OSTC | Oligosaccharyltransferase complex subunit OSTC OS=Homo sapiens OX=9606 GN=OSTC PE=1 SV=1 | 1 |
| TRAFD1 | TRAF-type zinc finger domain-containing protein 1 OS=Homo sapiens OX=9606 GN=TRAFD1 PE=1 SV=1 | 14 |
| CEP290 | Centrosomal protein of 290 kDa OS=Homo sapiens OX=9606 GN=CEP290 PE=1 SV=2 | 1 |
| HLCS | Biotin--protein ligase OS=Homo sapiens OX=9606 GN=HLCS PE=1 SV=1 | 2 |
| VPS52 | Vacuolar protein sorting-associated protein 52 homolog OS=Homo sapiens OX=9606 GN=VPS52 PE=1 SV=1 | 2 |
| CERS2 | Ceramide synthase 2 OS=Homo sapiens OX=9606 GN=CERS2 PE=1 SV=1 | 1 |
| SURF4 | Surfeit locus protein 4 OS=Homo sapiens OX=9606 GN=SURF4 PE=1 SV=3 | 1 |
| SLC41A1 | Solute carrier family 41 member 1 OS=Homo sapiens OX=9606 GN=SLC41A1 PE=1 SV=2 | 1 |
| DERA | Deoxyribose-phosphate aldolase OS=Homo sapiens OX=9606 GN=DERA PE=1 SV=2 | 1 |
| YME1L1 | ATP-dependent zinc metalloprotease YME1L1 OS=Homo sapiens OX=9606 GN=YME1L1 PE=1 SV=2 | 2 |
| AMPD3 | AMP deaminase 3 OS=Homo sapiens OX=9606 GN=AMPD3 PE=1 SV=1 | 3 |
| RBP4 | Retinol-binding protein 4 OS=Homo sapiens OX=9606 GN=RBP4 PE=1 SV=3 | 4 |
| SPATA5 | Ribosome biogenesis protein SPATA5 OS=Homo sapiens OX=9606 GN=SPATA5 PE=1 SV=3 | 3 |
| SCRN3 | Secernin-3 OS=Homo sapiens OX=9606 GN=SCRN3 PE=1 SV=1 | 3 |
| CPVL | Probable serine carboxypeptidase CPVL OS=Homo sapiens OX=9606 GN=CPVL PE=1 SV=2 | 7 |
| CENPX | Centromere protein X OS=Homo sapiens OX=9606 GN=CENPX PE=1 SV=1 | 1 |
| UNG | Uracil-DNA glycosylase OS=Homo sapiens OX=9606 GN=UNG PE=1 SV=2 | 1 |
| CALCOCO2 | Calcium-binding and coiled-coil domain-containing protein 2 OS=Homo sapiens OX=9606 GN=CALCOCO2 PE=1 SV=1 | 2 |
| GRIK4 | Glutamate receptor ionotropic, kainate 4 OS=Homo sapiens OX=9606 GN=GRIK4 PE=2 SV=2 | 2 |
| DPY30 | Protein dpy-30 homolog OS=Homo sapiens OX=9606 GN=DPY30 PE=1 SV=1 | 2 |
| SLC28A3 | Solute carrier family 28 member 3 OS=Homo sapiens OX=9606 GN=SLC28A3 PE=1 SV=1 | 1 |
| SNX6 | Sorting nexin-6 OS=Homo sapiens OX=9606 GN=SNX6 PE=1 SV=1 | 1 |
| CD164 | Sialomucin core protein 24 OS=Homo sapiens OX=9606 GN=CD164 PE=1 SV=2 | 1 |
| VPS50 | Syndetin OS=Homo sapiens OX=9606 GN=VPS50 PE=1 SV=3 | 2 |
| NDUFAF7 | Protein arginine methyltransferase NDUFAF7, mitochondrial OS=Homo sapiens OX=9606 GN=NDUFAF7 PE=1 SV=1 | 7 |
| LRRN4 | Leucine-rich repeat neuronal protein 4 OS=Homo sapiens OX=9606 GN=LRRN4 PE=1 SV=3 | 2 |
| HAT1 | Histone acetyltransferase type B catalytic subunit OS=Homo sapiens OX=9606 GN=HAT1 PE=1 SV=2 | 2 |
| MOB4 | MOB-like protein phocein OS=Homo sapiens OX=9606 GN=MOB4 PE=1 SV=1 | 1 |
| SMIM20 | Small integral membrane protein 20 OS=Homo sapiens OX=9606 GN=SMIM20 PE=1 SV=3 | 1 |
| EMC8 | ER membrane protein complex subunit 8 OS=Homo sapiens OX=9606 GN=EMC8 PE=1 SV=1 | 2 |
| PPP1R35 | Protein phosphatase 1 regulatory subunit 35 OS=Homo sapiens OX=9606 GN=PPP1R35 PE=1 SV=1 | 3 |
| CDC20 | Cell division cycle protein 20 homolog OS=Homo sapiens OX=9606 GN=CDC20 PE=1 SV=2 | 3 |
| SYNJ2BP | Synaptojanin-2-binding protein OS=Homo sapiens OX=9606 GN=SYNJ2BP PE=1 SV=2 | 3 |
| F13A1 | Coagulation factor XIII A chain OS=Homo sapiens OX=9606 GN=F13A1 PE=1 SV=5 | 8 |
| TSPOAP1 | Peripheral-type benzodiazepine receptor-associated protein 1 OS=Homo sapiens OX=9606 GN=TSPOAP1 PE=1 SV=2 | 1 |
| PLIN4 | Perilipin-4 OS=Homo sapiens OX=9606 GN=PLIN4 PE=1 SV=3 | 1 |
| SIL1 | Nucleotide exchange factor SIL1 OS=Homo sapiens OX=9606 GN=SIL1 PE=1 SV=1 | 10 |
| NUP37 | Nucleoporin Nup37 OS=Homo sapiens OX=9606 GN=NUP37 PE=1 SV=1 | 7 |
| SNCB | Beta-synuclein OS=Homo sapiens OX=9606 GN=SNCB PE=1 SV=1 | 2 |
| PHPT1 | 14 kDa phosphohistidine phosphatase OS=Homo sapiens OX=9606 GN=PHPT1 PE=1 SV=1 | 7 |
| EDIL3 | EGF-like repeat and discoidin I-like domain-containing protein 3 OS=Homo sapiens OX=9606 GN=EDIL3 PE=1 SV=1 | 10 |
| SMYD5 | Histone-lysine N-trimethyltransferase SMYD5 OS=Homo sapiens OX=9606 GN=SMYD5 PE=1 SV=2 | 1 |
| EZHIP | EZH inhibitory protein OS=Homo sapiens OX=9606 GN=EZHIP PE=1 SV=1 | 7 |
| NUMA1 | Nuclear mitotic apparatus protein 1 OS=Homo sapiens OX=9606 GN=NUMA1 PE=1 SV=2 | 11 |
| SYMPK | Symplekin OS=Homo sapiens OX=9606 GN=SYMPK PE=1 SV=2 | 1 |
| RPL26L1 | 60S ribosomal protein L26-like 1 OS=Homo sapiens OX=9606 GN=RPL26L1 PE=1 SV=1 | 1 |
| CTSL | Procathepsin L OS=Homo sapiens OX=9606 GN=CTSL PE=1 SV=2 | 2 |
| RIF1 | Telomere-associated protein RIF1 OS=Homo sapiens OX=9606 GN=RIF1 PE=1 SV=2 | 1 |
| BDH1 | D-beta-hydroxybutyrate dehydrogenase, mitochondrial OS=Homo sapiens OX=9606 GN=BDH1 PE=1 SV=3 | 5 |
| PEX11B | Peroxisomal membrane protein 11B OS=Homo sapiens OX=9606 GN=PEX11B PE=1 SV=1 | 1 |
| EPSTI1 | Epithelial-stromal interaction protein 1 OS=Homo sapiens OX=9606 GN=EPSTI1 PE=1 SV=2 | 8 |
| TSPAN9 | Tetraspanin-9 OS=Homo sapiens OX=9606 GN=TSPAN9 PE=1 SV=1 | 2 |
| LRRFIP2 | Leucine-rich repeat flightless-interacting protein 2 OS=Homo sapiens OX=9606 GN=LRRFIP2 PE=1 SV=1 | 2 |
| BOD1 | Biorientation of chromosomes in cell division protein 1 OS=Homo sapiens OX=9606 GN=BOD1 PE=1 SV=2 | 10 |
| WNK1 | Serine/threonine-protein kinase WNK1 OS=Homo sapiens OX=9606 GN=WNK1 PE=1 SV=2 | 2 |
| POLD2 | DNA polymerase delta subunit 2 OS=Homo sapiens OX=9606 GN=POLD2 PE=1 SV=1 | 1 |
| NLRP14 | NACHT, LRR and PYD domains-containing protein 14 OS=Homo sapiens OX=9606 GN=NLRP14 PE=1 SV=1 | 35 |
| PIP5K1C | Phosphatidylinositol 4-phosphate 5-kinase type-1 gamma OS=Homo sapiens OX=9606 GN=PIP5K1C PE=1 SV=2 | 3 |
| MPPE1 | Metallophosphoesterase 1 OS=Homo sapiens OX=9606 GN=MPPE1 PE=1 SV=2 | 1 |
| TRMT44 | Probable tRNA (uracil-O(2)-)-methyltransferase OS=Homo sapiens OX=9606 GN=TRMT44 PE=1 SV=2 | 1 |
| TSTD2 | Thiosulfate sulfurtransferase/rhodanese-like domain-containing protein 2 OS=Homo sapiens OX=9606 GN=TSTD2 PE=1 SV=1 | 1 |
| NUP35 | Nucleoporin NUP35 OS=Homo sapiens OX=9606 GN=NUP35 PE=1 SV=1 | 2 |
| CSRP1 | Cysteine and glycine-rich protein 1 OS=Homo sapiens OX=9606 GN=CSRP1 PE=1 SV=3 | 10 |
| COG3 | Conserved oligomeric Golgi complex subunit 3 OS=Homo sapiens OX=9606 GN=COG3 PE=1 SV=3 | 3 |
| ATP6V1F | V-type proton ATPase subunit F OS=Homo sapiens OX=9606 GN=ATP6V1F PE=1 SV=2 | 2 |
| TRIM3 | Tripartite motif-containing protein 3 OS=Homo sapiens OX=9606 GN=TRIM3 PE=1 SV=2 | 3 |
| LMF2 | Lipase maturation factor 2 OS=Homo sapiens OX=9606 GN=LMF2 PE=1 SV=2 | 5 |
| GPS1 | COP9 signalosome complex subunit 1 OS=Homo sapiens OX=9606 GN=GPS1 PE=1 SV=4 | 16 |
| BTD | Biotinidase OS=Homo sapiens OX=9606 GN=BTD PE=1 SV=2 | 3 |
| RAB8B | Ras-related protein Rab-8B OS=Homo sapiens OX=9606 GN=RAB8B PE=1 SV=2 | 3 |
| RAC2 | Ras-related C3 botulinum toxin substrate 2 OS=Homo sapiens OX=9606 GN=RAC2 PE=1 SV=1 | 2 |
| TMEM30A | Cell cycle control protein 50A OS=Homo sapiens OX=9606 GN=TMEM30A PE=1 SV=1 | 1 |
| RFC4 | Replication factor C subunit 4 OS=Homo sapiens OX=9606 GN=RFC4 PE=1 SV=2 | 2 |
| NRBP1 | Nuclear receptor-binding protein OS=Homo sapiens OX=9606 GN=NRBP1 PE=1 SV=1 | 1 |
| TANGO2 | Transport and Golgi organization protein 2 homolog OS=Homo sapiens OX=9606 GN=TANGO2 PE=1 SV=1 | 1 |
| CNOT8 | CCR4-NOT transcription complex subunit 8 OS=Homo sapiens OX=9606 GN=CNOT8 PE=1 SV=1 | 2 |
| RPL39 | 60S ribosomal protein L39 OS=Homo sapiens OX=9606 GN=RPL39 PE=1 SV=2 | 1 |
| IGFBP1 | Insulin-like growth factor-binding protein 1 OS=Homo sapiens OX=9606 GN=IGFBP1 PE=1 SV=1 | 6 |
| TRAPPC3 | Trafficking protein particle complex subunit 3 OS=Homo sapiens OX=9606 GN=TRAPPC3 PE=1 SV=1 | 4 |
| AK6 | Adenylate kinase isoenzyme 6 OS=Homo sapiens OX=9606 GN=AK6 PE=1 SV=1 | 1 |
| TSR2 | Pre-rRNA-processing protein TSR2 homolog OS=Homo sapiens OX=9606 GN=TSR2 PE=1 SV=1 | 1 |
| FUNDC2 | FUN14 domain-containing protein 2 OS=Homo sapiens OX=9606 GN=FUNDC2 PE=1 SV=2 | 1 |
| PMS2P11 | Putative postmeiotic segregation increased 2-like protein 11 OS=Homo sapiens OX=9606 GN=PMS2P11 PE=5 SV=1 | 1 |
| LGALS8 | Galectin-8 OS=Homo sapiens OX=9606 GN=LGALS8 PE=1 SV=4 | 7 |
| DAGLB | Diacylglycerol lipase-beta OS=Homo sapiens OX=9606 GN=DAGLB PE=1 SV=2 | 8 |
| GET3 | ATPase GET3 OS=Homo sapiens OX=9606 GN=GET3 PE=1 SV=2 | 10 |
| PSME3 | Proteasome activator complex subunit 3 OS=Homo sapiens OX=9606 GN=PSME3 PE=1 SV=1 | 3 |
| MKKS | Alternative protein MKKS OS=Homo sapiens OX=9606 GN=MKKS PE=1 SV=1 | 1 |
| APOC3 | Apolipoprotein C-III OS=Homo sapiens OX=9606 GN=APOC3 PE=1 SV=1 | 3 |
| PPM1G | Protein phosphatase 1G OS=Homo sapiens OX=9606 GN=PPM1G PE=1 SV=1 | 2 |
| B4GALT4 | Beta-1,4-galactosyltransferase 4 OS=Homo sapiens OX=9606 GN=B4GALT4 PE=1 SV=1 | 3 |
| MUC16 | Mucin-16 OS=Homo sapiens OX=9606 GN=MUC16 PE=1 SV=3 | 2 |
| PRKCD | Protein kinase C delta type OS=Homo sapiens OX=9606 GN=PRKCD PE=1 SV=2 | 2 |
| WDR70 | WD repeat-containing protein 70 OS=Homo sapiens OX=9606 GN=WDR70 PE=1 SV=1 | 1 |
| RCN1 | Reticulocalbin-1 OS=Homo sapiens OX=9606 GN=RCN1 PE=1 SV=1 | 2 |
| SF3B4 | Splicing factor 3B subunit 4 OS=Homo sapiens OX=9606 GN=SF3B4 PE=1 SV=1 | 2 |
| DNAJB1 | DnaJ homolog subfamily B member 1 OS=Homo sapiens OX=9606 GN=DNAJB1 PE=1 SV=4 | 21 |
| XPNPEP3 | Xaa-Pro aminopeptidase 3 OS=Homo sapiens OX=9606 GN=XPNPEP3 PE=1 SV=1 | 1 |
| ERGIC2 | Endoplasmic reticulum-Golgi intermediate compartment protein 2 OS=Homo sapiens OX=9606 GN=ERGIC2 PE=1 SV=2 | 2 |
| SLC25A31 | ADP/ATP translocase 4 OS=Homo sapiens OX=9606 GN=SLC25A31 PE=1 SV=1 | 3 |
| TMSB4X | Thymosin beta-4 OS=Homo sapiens OX=9606 GN=TMSB4X PE=1 SV=2 | 2 |
| TPD52 | Tumor protein D52 OS=Homo sapiens OX=9606 GN=TPD52 PE=1 SV=2 | 6 |
| TBC1D13 | TBC1 domain family member 13 OS=Homo sapiens OX=9606 GN=TBC1D13 PE=1 SV=3 | 2 |
| RHOB | Rho-related GTP-binding protein RhoB OS=Homo sapiens OX=9606 GN=RHOB PE=1 SV=1 | 1 |
| FBLN5 | Fibulin-5 OS=Homo sapiens OX=9606 GN=FBLN5 PE=1 SV=1 | 4 |
| NAT14 | Probable N-acetyltransferase 14 OS=Homo sapiens OX=9606 GN=NAT14 PE=1 SV=1 | 2 |
| COL4A2 | Collagen alpha-2(IV) chain OS=Homo sapiens OX=9606 GN=COL4A2 PE=1 SV=4 | 5 |
| UBE2F | NEDD8-conjugating enzyme UBE2F OS=Homo sapiens OX=9606 GN=UBE2F PE=1 SV=1 | 6 |
| RBM42 | RNA-binding protein 42 OS=Homo sapiens OX=9606 GN=RBM42 PE=1 SV=1 | 1 |
| MAD2L1 | Mitotic spindle assembly checkpoint protein MAD2A OS=Homo sapiens OX=9606 GN=MAD2L1 PE=1 SV=1 | 7 |
| VMAC | Vimentin-type intermediate filament-associated coiled-coil protein OS=Homo sapiens OX=9606 GN=VMAC PE=1 SV=1 | 1 |
| TTC17 | Tetratricopeptide repeat protein 17 OS=Homo sapiens OX=9606 GN=TTC17 PE=1 SV=1 | 13 |
| GRB7 | Growth factor receptor-bound protein 7 OS=Homo sapiens OX=9606 GN=GRB7 PE=1 SV=2 | 21 |
| PGS1 | CDP-diacylglycerol--glycerol-3-phosphate 3-phosphatidyltransferase, mitochondrial OS=Homo sapiens OX=9606 GN=PGS1 PE=2 SV=1 | 3 |
| RCC1 | Regulator of chromosome condensation OS=Homo sapiens OX=9606 GN=RCC1 PE=1 SV=1 | 1 |
| SEC13 | Protein SEC13 homolog OS=Homo sapiens OX=9606 GN=SEC13 PE=1 SV=3 | 5 |
| NLRP4 | NACHT, LRR and PYD domains-containing protein 4 OS=Homo sapiens OX=9606 GN=NLRP4 PE=1 SV=3 | 38 |
| TBC1D2 | TBC1 domain family member 2A OS=Homo sapiens OX=9606 GN=TBC1D2 PE=1 SV=3 | 3 |
| SAP30 | Histone deacetylase complex subunit SAP30 OS=Homo sapiens OX=9606 GN=SAP30 PE=1 SV=1 | 2 |
| FAHD2B | Fumarylacetoacetate hydrolase domain-containing protein 2B OS=Homo sapiens OX=9606 GN=FAHD2B PE=1 SV=1 | 1 |
| HNRNPA0 | Heterogeneous nuclear ribonucleoprotein A0 OS=Homo sapiens OX=9606 GN=HNRNPA0 PE=1 SV=1 | 2 |
| MOB1B | MOB kinase activator 1B OS=Homo sapiens OX=9606 GN=MOB1B PE=1 SV=3 | 1 |
| IPO8 | Importin-8 OS=Homo sapiens OX=9606 GN=IPO8 PE=1 SV=2 | 8 |
| MSRB2 | Methionine-R-sulfoxide reductase B2, mitochondrial OS=Homo sapiens OX=9606 GN=MSRB2 PE=1 SV=2 | 1 |
| TEX2 | Testis-expressed protein 2 OS=Homo sapiens OX=9606 GN=TEX2 PE=1 SV=2 | 4 |
| PGK2 | Phosphoglycerate kinase 2 OS=Homo sapiens OX=9606 GN=PGK2 PE=1 SV=3 | 1 |
| GRK2 | Beta-adrenergic receptor kinase 1 OS=Homo sapiens OX=9606 GN=GRK2 PE=1 SV=2 | 1 |
| TUBA1B | Tubulin alpha-1B chain OS=Homo sapiens OX=9606 GN=TUBA1B PE=1 SV=1 | 1 |
| ASF1A | Histone chaperone ASF1A OS=Homo sapiens OX=9606 GN=ASF1A PE=1 SV=1 | 3 |
| AGT | Angiotensinogen OS=Homo sapiens OX=9606 GN=AGT PE=1 SV=3 | 6 |
| GJA1 | Gap junction alpha-1 protein OS=Homo sapiens OX=9606 GN=GJA1 PE=1 SV=2 | 7 |
| TUBA1C | Tubulin alpha-1C chain OS=Homo sapiens OX=9606 GN=TUBA1C PE=1 SV=1 | 9 |
| GHDC | GH3 domain-containing protein OS=Homo sapiens OX=9606 GN=GHDC PE=1 SV=2 | 1 |
| HNRNPDL | Heterogeneous nuclear ribonucleoprotein D-like OS=Homo sapiens OX=9606 GN=HNRNPDL PE=1 SV=3 | 2 |
| AKR1B10 | Aldo-keto reductase family 1 member B10 OS=Homo sapiens OX=9606 GN=AKR1B10 PE=1 SV=2 | 13 |
| DCTPP1 | dCTP pyrophosphatase 1 OS=Homo sapiens OX=9606 GN=DCTPP1 PE=1 SV=1 | 3 |
| MYADM | Myeloid-associated differentiation marker OS=Homo sapiens OX=9606 GN=MYADM PE=1 SV=2 | 1 |
| RAP1GDS1 | Rap1 GTPase-GDP dissociation stimulator 1 OS=Homo sapiens OX=9606 GN=RAP1GDS1 PE=1 SV=3 | 7 |
| KCTD21 | BTB/POZ domain-containing protein KCTD21 OS=Homo sapiens OX=9606 GN=KCTD21 PE=1 SV=1 | 2 |
| ADPGK | ADP-dependent glucokinase OS=Homo sapiens OX=9606 GN=ADPGK PE=1 SV=1 | 3 |
| DPPA5 | Developmental pluripotency-associated 5 protein OS=Homo sapiens OX=9606 GN=DPPA5 PE=2 SV=1 | 5 |
| FAM107B | Protein FAM107B OS=Homo sapiens OX=9606 GN=FAM107B PE=1 SV=1 | 2 |
| TRMT11 | tRNA (guanine(10)-N2)-methyltransferase homolog OS=Homo sapiens OX=9606 GN=TRMT11 PE=1 SV=1 | 1 |
| GTSF1 | Gametocyte-specific factor 1 OS=Homo sapiens OX=9606 GN=GTSF1 PE=1 SV=2 | 8 |
| TRAPPC9 | Trafficking protein particle complex subunit 9 OS=Homo sapiens OX=9606 GN=TRAPPC9 PE=1 SV=2 | 1 |
| CTNND1 | Catenin delta-1 OS=Homo sapiens OX=9606 GN=CTNND1 PE=1 SV=1 | 16 |
| TLE2 | Transducin-like enhancer protein 2 OS=Homo sapiens OX=9606 GN=TLE2 PE=1 SV=2 | 5 |
| FOXRED1 | FAD-dependent oxidoreductase domain-containing protein 1 OS=Homo sapiens OX=9606 GN=FOXRED1 PE=1 SV=2 | 2 |
| AHCTF1 | Protein ELYS OS=Homo sapiens OX=9606 GN=AHCTF1 PE=1 SV=3 | 2 |
| NT5C2 | Cytosolic purine 5'-nucleotidase OS=Homo sapiens OX=9606 GN=NT5C2 PE=1 SV=1 | 1 |
| MARS2 | Methionine--tRNA ligase, mitochondrial OS=Homo sapiens OX=9606 GN=MARS2 PE=1 SV=2 | 1 |
| PCF11 | Pre-mRNA cleavage complex 2 protein Pcf11 OS=Homo sapiens OX=9606 GN=PCF11 PE=1 SV=3 | 1 |
| DENND10 | DENN domain-containing protein 10 OS=Homo sapiens OX=9606 GN=DENND10 PE=1 SV=1 | 4 |
| DOCK8 | Dedicator of cytokinesis protein 8 OS=Homo sapiens OX=9606 GN=DOCK8 PE=1 SV=3 | 13 |
| LIMA1 | LIM domain and actin-binding protein 1 OS=Homo sapiens OX=9606 GN=LIMA1 PE=1 SV=1 | 34 |
| TUSC2 | Tumor suppressor candidate 2 OS=Homo sapiens OX=9606 GN=TUSC2 PE=1 SV=3 | 2 |
| MRPL15 | 39S ribosomal protein L15, mitochondrial OS=Homo sapiens OX=9606 GN=MRPL15 PE=1 SV=1 | 3 |
| KSR1 | Kinase suppressor of Ras 1 OS=Homo sapiens OX=9606 GN=KSR1 PE=1 SV=3 | 6 |
| HSPB11 | Intraflagellar transport protein 25 homolog OS=Homo sapiens OX=9606 GN=HSPB11 PE=1 SV=1 | 3 |
| VPS33A | Vacuolar protein sorting-associated protein 33A OS=Homo sapiens OX=9606 GN=VPS33A PE=1 SV=1 | 3 |
| LRPAP1 | Alpha-2-macroglobulin receptor-associated protein OS=Homo sapiens OX=9606 GN=LRPAP1 PE=1 SV=1 | 23 |
| PON3 | Serum paraoxonase/lactonase 3 OS=Homo sapiens OX=9606 GN=PON3 PE=1 SV=3 | 6 |
| MFN2 | Mitofusin-2 OS=Homo sapiens OX=9606 GN=MFN2 PE=1 SV=3 | 5 |
| COPG2 | Coatomer subunit gamma-2 OS=Homo sapiens OX=9606 GN=COPG2 PE=1 SV=1 | 8 |
| RTRAF | RNA transcription, translation and transport factor protein OS=Homo sapiens OX=9606 GN=RTRAF PE=1 SV=1 | 7 |
| CUL4B | Cullin-4B OS=Homo sapiens OX=9606 GN=CUL4B PE=1 SV=4 | 4 |
| SRSF7 | Serine/arginine-rich splicing factor 7 OS=Homo sapiens OX=9606 GN=SRSF7 PE=1 SV=1 | 2 |
| SMC3 | Structural maintenance of chromosomes protein 3 OS=Homo sapiens OX=9606 GN=SMC3 PE=1 SV=2 | 2 |
| TTC37 | SKI3 subunit of superkiller complex protein OS=Homo sapiens OX=9606 GN=SKIC3 PE=1 SV=1 | 4 |
| MAT2B | Methionine adenosyltransferase 2 subunit beta OS=Homo sapiens OX=9606 GN=MAT2B PE=1 SV=1 | 9 |
| IQGAP2 | Ras GTPase-activating-like protein IQGAP2 OS=Homo sapiens OX=9606 GN=IQGAP2 PE=1 SV=4 | 14 |
| APOC2 | Apolipoprotein C-II OS=Homo sapiens OX=9606 GN=APOC2 PE=1 SV=1 | 4 |
| PPP4C | Serine/threonine-protein phosphatase 4 catalytic subunit OS=Homo sapiens OX=9606 GN=PPP4C PE=1 SV=1 | 3 |
| PROSER1 | Proline and serine-rich protein 1 OS=Homo sapiens OX=9606 GN=PROSER1 PE=1 SV=2 | 12 |
| GBE1 | 1,4-alpha-glucan-branching enzyme OS=Homo sapiens OX=9606 GN=GBE1 PE=1 SV=3 | 22 |
| ABAT | 4-aminobutyrate aminotransferase, mitochondrial OS=Homo sapiens OX=9606 GN=ABAT PE=1 SV=3 | 2 |
| RPL37A | 60S ribosomal protein L37a OS=Homo sapiens OX=9606 GN=RPL37A PE=1 SV=2 | 2 |
| NUBP2 | Cytosolic Fe-S cluster assembly factor NUBP2 OS=Homo sapiens OX=9606 GN=NUBP2 PE=1 SV=1 | 1 |
| SMARCA5 | SWI/SNF-related matrix-associated actin-dependent regulator of chromatin subfamily A member 5 OS=Homo sapiens OX=9606 GN=SMARCA5 PE=1 SV=1 | 4 |
| CNPY2 | Protein canopy homolog 2 OS=Homo sapiens OX=9606 GN=CNPY2 PE=1 SV=1 | 3 |
| NUBPL | Iron-sulfur protein NUBPL OS=Homo sapiens OX=9606 GN=NUBPL PE=1 SV=3 | 6 |
| TSPAN7 | Tetraspanin-7 OS=Homo sapiens OX=9606 GN=TSPAN7 PE=1 SV=2 | 2 |
| NLRP11 | NACHT, LRR and PYD domains-containing protein 11 OS=Homo sapiens OX=9606 GN=NLRP11 PE=2 SV=2 | 45 |
| LARP7 | La-related protein 7 OS=Homo sapiens OX=9606 GN=LARP7 PE=1 SV=1 | 3 |
| FBL | rRNA 2'-O-methyltransferase fibrillarin OS=Homo sapiens OX=9606 GN=FBL PE=1 SV=2 | 1 |
| THEM6 | Protein THEM6 OS=Homo sapiens OX=9606 GN=THEM6 PE=1 SV=2 | 3 |
| FOCAD | Focadhesin OS=Homo sapiens OX=9606 GN=FOCAD PE=1 SV=2 | 6 |
| CFAP410 | Cilia- and flagella-associated protein 410 OS=Homo sapiens OX=9606 GN=CFAP410 PE=1 SV=1 | 1 |
| MMGT1 | ER membrane protein complex subunit 5 OS=Homo sapiens OX=9606 GN=MMGT1 PE=1 SV=1 | 2 |
| GPAA1 | Glycosylphosphatidylinositol anchor attachment 1 protein OS=Homo sapiens OX=9606 GN=GPAA1 PE=1 SV=3 | 5 |
| ACTR10 | Actin-related protein 10 OS=Homo sapiens OX=9606 GN=ACTR10 PE=1 SV=1 | 4 |
| SLC1A4 | Neutral amino acid transporter A OS=Homo sapiens OX=9606 GN=SLC1A4 PE=1 SV=1 | 6 |
| ALG1 | Chitobiosyldiphosphodolichol beta-mannosyltransferase OS=Homo sapiens OX=9606 GN=ALG1 PE=1 SV=2 | 2 |
| RABGAP1 | Rab GTPase-activating protein 1 OS=Homo sapiens OX=9606 GN=RABGAP1 PE=1 SV=3 | 17 |
| H2AC4_H2AC8 | Histone H2A type 1-B/E OS=Homo sapiens OX=9606 GN=H2AC8 PE=1 SV=2 | 5 |
| FTH1 | Ferritin heavy chain OS=Homo sapiens OX=9606 GN=FTH1 PE=1 SV=2 | 11 |
| NDC80 | Kinetochore protein NDC80 homolog OS=Homo sapiens OX=9606 GN=NDC80 PE=1 SV=1 | 1 |
| DHX58 | ATP-dependent RNA helicase DHX58 OS=Homo sapiens OX=9606 GN=DHX58 PE=1 SV=1 | 12 |
| ALDH18A1 | Delta-1-pyrroline-5-carboxylate synthase OS=Homo sapiens OX=9606 GN=ALDH18A1 PE=1 SV=2 | 10 |
| INSRR | Insulin receptor-related protein OS=Homo sapiens OX=9606 GN=INSRR PE=1 SV=2 | 1 |
| TRIOBP | TRIO and F-actin-binding protein OS=Homo sapiens OX=9606 GN=TRIOBP PE=1 SV=3 | 3 |
| BSDC1 | BSD domain-containing protein 1 OS=Homo sapiens OX=9606 GN=BSDC1 PE=1 SV=1 | 5 |
| GPM6B | Neuronal membrane glycoprotein M6-b OS=Homo sapiens OX=9606 GN=GPM6B PE=1 SV=2 | 1 |
| CMTR1 | Cap-specific mRNA (nucleoside-2'-O-)-methyltransferase 1 OS=Homo sapiens OX=9606 GN=CMTR1 PE=1 SV=1 | 12 |
| TMEM63B | CSC1-like protein 2 OS=Homo sapiens OX=9606 GN=TMEM63B PE=1 SV=1 | 4 |
| RPL38 | 60S ribosomal protein L38 OS=Homo sapiens OX=9606 GN=RPL38 PE=1 SV=2 | 2 |
| ANP32E | Acidic leucine-rich nuclear phosphoprotein 32 family member E OS=Homo sapiens OX=9606 GN=ANP32E PE=1 SV=1 | 1 |
| NARS2 | Probable asparagine--tRNA ligase, mitochondrial OS=Homo sapiens OX=9606 GN=NARS2 PE=1 SV=3 | 2 |
| SNRPA | U1 small nuclear ribonucleoprotein A OS=Homo sapiens OX=9606 GN=SNRPA PE=1 SV=3 | 1 |
| ATP13A1 | Endoplasmic reticulum transmembrane helix translocase OS=Homo sapiens OX=9606 GN=ATP13A1 PE=1 SV=2 | 5 |
| TIMM10B | Mitochondrial import inner membrane translocase subunit Tim10 B OS=Homo sapiens OX=9606 GN=TIMM10B PE=1 SV=1 | 1 |
| CTBS | Di-N-acetylchitobiase OS=Homo sapiens OX=9606 GN=CTBS PE=1 SV=1 | 4 |
| TBCD | Tubulin-specific chaperone D OS=Homo sapiens OX=9606 GN=TBCD PE=1 SV=2 | 42 |
| CHCHD7 | Coiled-coil-helix-coiled-coil-helix domain-containing protein 7 OS=Homo sapiens OX=9606 GN=CHCHD7 PE=1 SV=1 | 1 |
| HVCN1 | Voltage-gated hydrogen channel 1 OS=Homo sapiens OX=9606 GN=HVCN1 PE=1 SV=1 | 6 |
| TBCC | Tubulin-specific chaperone C OS=Homo sapiens OX=9606 GN=TBCC PE=1 SV=2 | 13 |
| FOS | Protein c-Fos OS=Homo sapiens OX=9606 GN=FOS PE=1 SV=1 | 2 |
| COMMD8 | COMM domain-containing protein 8 OS=Homo sapiens OX=9606 GN=COMMD8 PE=1 SV=1 | 4 |
| RHOA | Transforming protein RhoA OS=Homo sapiens OX=9606 GN=RHOA PE=1 SV=1 | 9 |
| OPA3 | Optic atrophy 3 protein OS=Homo sapiens OX=9606 GN=OPA3 PE=1 SV=1 | 2 |
| MAPKAP1 | Target of rapamycin complex 2 subunit MAPKAP1 OS=Homo sapiens OX=9606 GN=MAPKAP1 PE=1 SV=2 | 1 |
| RAB2B | Ras-related protein Rab-2B OS=Homo sapiens OX=9606 GN=RAB2B PE=1 SV=1 | 5 |
| WASL | Actin nucleation-promoting factor WASL OS=Homo sapiens OX=9606 GN=WASL PE=1 SV=2 | 4 |
| TMBIM6 | Bax inhibitor 1 OS=Homo sapiens OX=9606 GN=TMBIM6 PE=1 SV=2 | 1 |
| MIS18A | Protein Mis18-alpha OS=Homo sapiens OX=9606 GN=MIS18A PE=1 SV=1 | 2 |
| M6PR | Cation-dependent mannose-6-phosphate receptor OS=Homo sapiens OX=9606 GN=M6PR PE=1 SV=1 | 5 |
| DHX16 | Pre-mRNA-splicing factor ATP-dependent RNA helicase DHX16 OS=Homo sapiens OX=9606 GN=DHX16 PE=1 SV=2 | 1 |
| TUBB8B | Tubulin beta 8B OS=Homo sapiens OX=9606 GN=TUBB8B PE=1 SV=1 | 5 |
| SPRYD4 | SPRY domain-containing protein 4 OS=Homo sapiens OX=9606 GN=SPRYD4 PE=1 SV=2 | 2 |
| GLS | Glutaminase kidney isoform, mitochondrial OS=Homo sapiens OX=9606 GN=GLS PE=1 SV=1 | 6 |
| SDE2 | Splicing regulator SDE2 OS=Homo sapiens OX=9606 GN=SDE2 PE=1 SV=1 | 3 |
| SNAP91 | Clathrin coat assembly protein AP180 OS=Homo sapiens OX=9606 GN=SNAP91 PE=1 SV=2 | 3 |
| CTSB | Cathepsin B OS=Homo sapiens OX=9606 GN=CTSB PE=1 SV=3 | 12 |
| ELAPOR2 | Endosome/lysosome-associated apoptosis and autophagy regulator family member 2 OS=Homo sapiens OX=9606 GN=ELAPOR2 PE=1 SV=2 | 10 |
| SCAMP1 | Secretory carrier-associated membrane protein 1 OS=Homo sapiens OX=9606 GN=SCAMP1 PE=1 SV=2 | 3 |
| DNAJC1 | DnaJ homolog subfamily C member 1 OS=Homo sapiens OX=9606 GN=DNAJC1 PE=1 SV=1 | 3 |
| WASHC5 | WASH complex subunit 5 OS=Homo sapiens OX=9606 GN=WASHC5 PE=1 SV=1 | 9 |
| MOSPD2 | Motile sperm domain-containing protein 2 OS=Homo sapiens OX=9606 GN=MOSPD2 PE=1 SV=1 | 2 |
| ACSF3 | Malonate--CoA ligase ACSF3, mitochondrial OS=Homo sapiens OX=9606 GN=ACSF3 PE=1 SV=3 | 5 |
| DPPA3 | Developmental pluripotency-associated protein 3 OS=Homo sapiens OX=9606 GN=DPPA3 PE=1 SV=1 | 7 |
| VPS37A | Vacuolar protein sorting-associated protein 37A OS=Homo sapiens OX=9606 GN=VPS37A PE=1 SV=1 | 4 |
| PPIL3 | Peptidyl-prolyl cis-trans isomerase-like 3 OS=Homo sapiens OX=9606 GN=PPIL3 PE=1 SV=1 | 2 |
| ITSN1 | Intersectin-1 OS=Homo sapiens OX=9606 GN=ITSN1 PE=1 SV=3 | 1 |
| MROH1 | Maestro heat-like repeat-containing protein family member 1 OS=Homo sapiens OX=9606 GN=MROH1 PE=2 SV=3 | 1 |
| GTF2B | Transcription initiation factor IIB OS=Homo sapiens OX=9606 GN=GTF2B PE=1 SV=1 | 5 |
| IER3IP1 | Immediate early response 3-interacting protein 1 OS=Homo sapiens OX=9606 GN=IER3IP1 PE=1 SV=1 | 1 |
| PLEKHA3 | Pleckstrin homology domain-containing family A member 3 OS=Homo sapiens OX=9606 GN=PLEKHA3 PE=1 SV=2 | 1 |
| LSM14A | Protein LSM14 homolog A OS=Homo sapiens OX=9606 GN=LSM14A PE=1 SV=3 | 2 |
| HSD17B7 | 3-keto-steroid reductase/17-beta-hydroxysteroid dehydrogenase 7 OS=Homo sapiens OX=9606 GN=HSD17B7 PE=1 SV=1 | 3 |
| SLC25A46 | Mitochondrial outer membrane protein SLC25A46 OS=Homo sapiens OX=9606 GN=SLC25A46 PE=1 SV=1 | 2 |
| TK2 | Thymidine kinase 2, mitochondrial OS=Homo sapiens OX=9606 GN=TK2 PE=1 SV=4 | 2 |
| RHOC | Rho-related GTP-binding protein RhoC OS=Homo sapiens OX=9606 GN=RHOC PE=1 SV=1 | 2 |
| LSM8 | U6 snRNA-associated Sm-like protein LSm8 OS=Homo sapiens OX=9606 GN=LSM8 PE=1 SV=3 | 3 |
| LTBP1 | Latent-transforming growth factor beta-binding protein 1 OS=Homo sapiens OX=9606 GN=LTBP1 PE=1 SV=4 | 2 |
| NPM3 | Nucleoplasmin-3 OS=Homo sapiens OX=9606 GN=NPM3 PE=1 SV=3 | 2 |
| COMMD3 | COMM domain-containing protein 3 OS=Homo sapiens OX=9606 GN=COMMD3 PE=1 SV=1 | 3 |
| TRIM33 | E3 ubiquitin-protein ligase TRIM33 OS=Homo sapiens OX=9606 GN=TRIM33 PE=1 SV=3 | 5 |
| VAC14 | Protein VAC14 homolog OS=Homo sapiens OX=9606 GN=VAC14 PE=1 SV=1 | 2 |
| ATG4A | Cysteine protease ATG4A OS=Homo sapiens OX=9606 GN=ATG4A PE=1 SV=1 | 2 |
| PCM1 | Pericentriolar material 1 protein OS=Homo sapiens OX=9606 GN=PCM1 PE=1 SV=5 | 1 |
| SNRPD2 | Small nuclear ribonucleoprotein Sm D2 OS=Homo sapiens OX=9606 GN=SNRPD2 PE=1 SV=1 | 2 |
| IZUMO1R | Sperm-egg fusion protein Juno OS=Homo sapiens OX=9606 GN=IZUMO1R PE=1 SV=3 | 6 |
| HNRNPU | Heterogeneous nuclear ribonucleoprotein U OS=Homo sapiens OX=9606 GN=HNRNPU PE=1 SV=6 | 2 |
| AMOT | Angiomotin OS=Homo sapiens OX=9606 GN=AMOT PE=1 SV=1 | 17 |
| CREB3L2 | Cyclic AMP-responsive element-binding protein 3-like protein 2 OS=Homo sapiens OX=9606 GN=CREB3L2 PE=1 SV=3 | 1 |
| IL16 | Pro-interleukin-16 OS=Homo sapiens OX=9606 GN=IL16 PE=1 SV=4 | 2 |
| IKBKG | NF-kappa-B essential modulator OS=Homo sapiens OX=9606 GN=IKBKG PE=1 SV=2 | 3 |
| BTF3L4 | Transcription factor BTF3 homolog 4 OS=Homo sapiens OX=9606 GN=BTF3L4 PE=1 SV=1 | 5 |
| C9orf43 | Uncharacterized protein C9orf43 OS=Homo sapiens OX=9606 GN=C9orf43 PE=1 SV=1 | 1 |
| IGBP1 | Immunoglobulin-binding protein 1 OS=Homo sapiens OX=9606 GN=IGBP1 PE=1 SV=1 | 6 |
| TECR | Very-long-chain enoyl-CoA reductase OS=Homo sapiens OX=9606 GN=TECR PE=1 SV=1 | 8 |
| MRPL53 | 39S ribosomal protein L53, mitochondrial OS=Homo sapiens OX=9606 GN=MRPL53 PE=1 SV=1 | 1 |
| ARMC6 | Armadillo repeat-containing protein 6 OS=Homo sapiens OX=9606 GN=ARMC6 PE=1 SV=2 | 4 |
| HSPA12A | Heat shock 70 kDa protein 12A OS=Homo sapiens OX=9606 GN=HSPA12A PE=1 SV=2 | 2 |
| PRMT7 | Protein arginine N-methyltransferase 7 OS=Homo sapiens OX=9606 GN=PRMT7 PE=1 SV=1 | 4 |
| PRADC1 | Protease-associated domain-containing protein 1 OS=Homo sapiens OX=9606 GN=PRADC1 PE=1 SV=1 | 2 |
| SYVN1 | E3 ubiquitin-protein ligase synoviolin OS=Homo sapiens OX=9606 GN=SYVN1 PE=1 SV=2 | 2 |
| GNL1 | Guanine nucleotide-binding protein-like 1 OS=Homo sapiens OX=9606 GN=GNL1 PE=1 SV=2 | 6 |
| BCS1L | Mitochondrial chaperone BCS1 OS=Homo sapiens OX=9606 GN=BCS1L PE=1 SV=1 | 3 |
| ALDH1A2 | Retinal dehydrogenase 2 OS=Homo sapiens OX=9606 GN=ALDH1A2 PE=1 SV=3 | 16 |
| SPCS2 | Signal peptidase complex subunit 2 OS=Homo sapiens OX=9606 GN=SPCS2 PE=1 SV=3 | 3 |
| VRK1 | Serine/threonine-protein kinase VRK1 OS=Homo sapiens OX=9606 GN=VRK1 PE=1 SV=1 | 3 |
| ANLN | Anillin OS=Homo sapiens OX=9606 GN=ANLN PE=1 SV=2 | 16 |
| PPP1R11 | E3 ubiquitin-protein ligase PPP1R11 OS=Homo sapiens OX=9606 GN=PPP1R11 PE=1 SV=1 | 1 |
| PIP4P1 | Type 1 phosphatidylinositol 4,5-bisphosphate 4-phosphatase OS=Homo sapiens OX=9606 GN=PIP4P1 PE=1 SV=1 | 1 |
| HNRNPA1 | Heterogeneous nuclear ribonucleoprotein A1 OS=Homo sapiens OX=9606 GN=HNRNPA1 PE=1 SV=5 | 12 |
| NUP133 | Nuclear pore complex protein Nup133 OS=Homo sapiens OX=9606 GN=NUP133 PE=1 SV=2 | 30 |
| MON2 | Protein MON2 homolog OS=Homo sapiens OX=9606 GN=MON2 PE=1 SV=3 | 7 |
| TBCE | Tubulin-specific chaperone E OS=Homo sapiens OX=9606 GN=TBCE PE=1 SV=1 | 16 |
| MZT2A | Mitotic-spindle organizing protein 2A OS=Homo sapiens OX=9606 GN=MZT2A PE=1 SV=2 | 1 |
| ZP1 | Zona pellucida sperm-binding protein 1 OS=Homo sapiens OX=9606 GN=ZP1 PE=1 SV=1 | 24 |
| CLYBL | Citramalyl-CoA lyase, mitochondrial OS=Homo sapiens OX=9606 GN=CLYBL PE=1 SV=2 | 11 |
| VPS26C | Vacuolar protein sorting-associated protein 26C OS=Homo sapiens OX=9606 GN=VPS26C PE=1 SV=1 | 2 |
| SH3BGRL2 | SH3 domain-binding glutamic acid-rich-like protein 2 OS=Homo sapiens OX=9606 GN=SH3BGRL2 PE=1 SV=2 | 2 |
| MAIP1 | m-AAA protease-interacting protein 1, mitochondrial OS=Homo sapiens OX=9606 GN=MAIP1 PE=1 SV=1 | 2 |
| DCTN4 | Dynactin subunit 4 OS=Homo sapiens OX=9606 GN=DCTN4 PE=1 SV=1 | 8 |
| CMC4 | Cx9C motif-containing protein 4 OS=Homo sapiens OX=9606 GN=CMC4 PE=1 SV=1 | 2 |
| LYRM1 | LYR motif-containing protein 1 OS=Homo sapiens OX=9606 GN=LYRM1 PE=2 SV=1 | 1 |
| HSD17B8 | (3R)-3-hydroxyacyl-CoA dehydrogenase OS=Homo sapiens OX=9606 GN=HSD17B8 PE=1 SV=2 | 4 |
| SLC4A7 | Sodium bicarbonate cotransporter 3 OS=Homo sapiens OX=9606 GN=SLC4A7 PE=1 SV=2 | 4 |
| FLAD1 | FAD synthase OS=Homo sapiens OX=9606 GN=FLAD1 PE=1 SV=1 | 14 |
| MTMR2 | Myotubularin-related protein 2 OS=Homo sapiens OX=9606 GN=MTMR2 PE=1 SV=4 | 4 |
| ITGB6 | Integrin beta-6 OS=Homo sapiens OX=9606 GN=ITGB6 PE=1 SV=2 | 1 |
| NME7 | Nucleoside diphosphate kinase 7 OS=Homo sapiens OX=9606 GN=NME7 PE=1 SV=1 | 1 |
| RARS2 | Probable arginine--tRNA ligase, mitochondrial OS=Homo sapiens OX=9606 GN=RARS2 PE=1 SV=1 | 14 |
| STT3B | Dolichyl-diphosphooligosaccharide--protein glycosyltransferase subunit STT3B OS=Homo sapiens OX=9606 GN=STT3B PE=1 SV=1 | 3 |
| PARP4 | Protein mono-ADP-ribosyltransferase PARP4 OS=Homo sapiens OX=9606 GN=PARP4 PE=1 SV=3 | 2 |
| RPS17 | 40S ribosomal protein S17 OS=Homo sapiens OX=9606 GN=RPS17 PE=1 SV=2 | 8 |
| GDAP1 | Ganglioside-induced differentiation-associated protein 1 OS=Homo sapiens OX=9606 GN=GDAP1 PE=1 SV=3 | 17 |
| TMED8 | Protein TMED8 OS=Homo sapiens OX=9606 GN=TMED8 PE=1 SV=1 | 8 |
| TROAP | Tastin OS=Homo sapiens OX=9606 GN=TROAP PE=1 SV=3 | 2 |
| SHMT1 | Serine hydroxymethyltransferase, cytosolic OS=Homo sapiens OX=9606 GN=SHMT1 PE=1 SV=1 | 10 |
| B3GNT2 | N-acetyllactosaminide beta-1,3-N-acetylglucosaminyltransferase 2 OS=Homo sapiens OX=9606 GN=B3GNT2 PE=1 SV=2 | 9 |
| UBE2G2 | Ubiquitin-conjugating enzyme E2 G2 OS=Homo sapiens OX=9606 GN=UBE2G2 PE=1 SV=1 | 3 |
| NUP107 | Nuclear pore complex protein Nup107 OS=Homo sapiens OX=9606 GN=NUP107 PE=1 SV=1 | 14 |
| ERP44 | Endoplasmic reticulum resident protein 44 OS=Homo sapiens OX=9606 GN=ERP44 PE=1 SV=1 | 14 |
| ADAMTS17 | A disintegrin and metalloproteinase with thrombospondin motifs 17 OS=Homo sapiens OX=9606 GN=ADAMTS17 PE=2 SV=2 | 3 |
| GOLGA2 | Golgin subfamily A member 2 OS=Homo sapiens OX=9606 GN=GOLGA2 PE=1 SV=3 | 4 |
| OTUD7B | OTU domain-containing protein 7B OS=Homo sapiens OX=9606 GN=OTUD7B PE=1 SV=1 | 3 |
| BACH1 | Transcription regulator protein BACH1 OS=Homo sapiens OX=9606 GN=BACH1 PE=1 SV=2 | 4 |
| UBE2Q2 | Ubiquitin-conjugating enzyme E2 Q2 OS=Homo sapiens OX=9606 GN=UBE2Q2 PE=1 SV=1 | 8 |
| SUMO1 | Small ubiquitin-related modifier 1 OS=Homo sapiens OX=9606 GN=SUMO1 PE=1 SV=1 | 3 |
| ATRN | Attractin OS=Homo sapiens OX=9606 GN=ATRN PE=1 SV=2 | 5 |
| PDZK1 | Na(+)/H(+) exchange regulatory cofactor NHE-RF3 OS=Homo sapiens OX=9606 GN=PDZK1 PE=1 SV=2 | 14 |
| TOMM5 | Mitochondrial import receptor subunit TOM5 homolog OS=Homo sapiens OX=9606 GN=TOMM5 PE=1 SV=1 | 1 |
| ZNF840P | Putative zinc finger protein 840 OS=Homo sapiens OX=9606 GN=ZNF840P PE=5 SV=5 | 29 |
| LRWD1 | Leucine-rich repeat and WD repeat-containing protein 1 OS=Homo sapiens OX=9606 GN=LRWD1 PE=1 SV=2 | 1 |
| COL6A6 | Collagen alpha-6(VI) chain OS=Homo sapiens OX=9606 GN=COL6A6 PE=1 SV=2 | 10 |
| DNAJA4 | DnaJ homolog subfamily A member 4 OS=Homo sapiens OX=9606 GN=DNAJA4 PE=1 SV=1 | 13 |
| FETUB | Fetuin-B OS=Homo sapiens OX=9606 GN=FETUB PE=1 SV=2 | 5 |
| MIOS | GATOR complex protein MIOS OS=Homo sapiens OX=9606 GN=MIOS PE=1 SV=2 | 2 |
| SHOC2 | Leucine-rich repeat protein SHOC-2 OS=Homo sapiens OX=9606 GN=SHOC2 PE=1 SV=2 | 13 |
| TMEM230 | Transmembrane protein 230 OS=Homo sapiens OX=9606 GN=TMEM230 PE=1 SV=1 | 2 |
| ACADVL | Very long-chain specific acyl-CoA dehydrogenase, mitochondrial OS=Homo sapiens OX=9606 GN=ACADVL PE=1 SV=1 | 37 |
| GLMN | Glomulin OS=Homo sapiens OX=9606 GN=GLMN PE=1 SV=2 | 6 |
| CSNK2B | Casein kinase II subunit beta OS=Homo sapiens OX=9606 GN=CSNK2B PE=1 SV=1 | 3 |
| GOSR2 | Golgi SNAP receptor complex member 2 OS=Homo sapiens OX=9606 GN=GOSR2 PE=1 SV=2 | 1 |
| ITFG1 | T-cell immunomodulatory protein OS=Homo sapiens OX=9606 GN=ITFG1 PE=1 SV=1 | 2 |
| ESPL1 | Separin OS=Homo sapiens OX=9606 GN=ESPL1 PE=1 SV=3 | 3 |
| OOSP2 | Oocyte-secreted protein 2 OS=Homo sapiens OX=9606 GN=OOSP2 PE=1 SV=1 | 3 |
| MYG1 | MYG1 exonuclease OS=Homo sapiens OX=9606 GN=MYG1 PE=1 SV=3 | 2 |
| DPCD | Protein DPCD OS=Homo sapiens OX=9606 GN=DPCD PE=1 SV=2 | 5 |
| HDAC1 | Histone deacetylase 1 OS=Homo sapiens OX=9606 GN=HDAC1 PE=1 SV=1 | 1 |
| LASP1 | LIM and SH3 domain protein 1 OS=Homo sapiens OX=9606 GN=LASP1 PE=1 SV=2 | 9 |
| MAP1S | Microtubule-associated protein 1S OS=Homo sapiens OX=9606 GN=MAP1S PE=1 SV=2 | 27 |
| NUMB | Protein numb homolog OS=Homo sapiens OX=9606 GN=NUMB PE=1 SV=2 | 6 |
| AMDHD2 | N-acetylglucosamine-6-phosphate deacetylase OS=Homo sapiens OX=9606 GN=AMDHD2 PE=1 SV=2 | 4 |
| BMP15 | Bone morphogenetic protein 15 OS=Homo sapiens OX=9606 GN=BMP15 PE=1 SV=2 | 15 |
| SNX18 | Sorting nexin-18 OS=Homo sapiens OX=9606 GN=SNX18 PE=1 SV=2 | 1 |
| PPHLN1 | Periphilin-1 OS=Homo sapiens OX=9606 GN=PPHLN1 PE=1 SV=2 | 2 |
| SLC9A6 | Sodium/hydrogen exchanger 6 OS=Homo sapiens OX=9606 GN=SLC9A6 PE=1 SV=3 | 5 |
| DECR1 | 2,4-dienoyl-CoA reductase [(3E)-enoyl-CoA-producing], mitochondrial OS=Homo sapiens OX=9606 GN=DECR1 PE=1 SV=1 | 12 |
| NQO2 | Ribosyldihydronicotinamide dehydrogenase [quinone] OS=Homo sapiens OX=9606 GN=NQO2 PE=1 SV=5 | 3 |
| PPP1R12A | Protein phosphatase 1 regulatory subunit 12A OS=Homo sapiens OX=9606 GN=PPP1R12A PE=1 SV=1 | 11 |
| LANCL1 | Glutathione S-transferase LANCL1 OS=Homo sapiens OX=9606 GN=LANCL1 PE=1 SV=1 | 15 |
| HIBCH | 3-hydroxyisobutyryl-CoA hydrolase, mitochondrial OS=Homo sapiens OX=9606 GN=HIBCH PE=1 SV=2 | 14 |
| TACC3 | Transforming acidic coiled-coil-containing protein 3 OS=Homo sapiens OX=9606 GN=TACC3 PE=1 SV=1 | 20 |
| KIF15 | Kinesin-like protein KIF15 OS=Homo sapiens OX=9606 GN=KIF15 PE=1 SV=1 | 2 |
| BAZ1A | Bromodomain adjacent to zinc finger domain protein 1A OS=Homo sapiens OX=9606 GN=BAZ1A PE=1 SV=2 | 2 |
| SSNA1 | Microtubule nucleation factor SSNA1 OS=Homo sapiens OX=9606 GN=SSNA1 PE=1 SV=2 | 1 |
| SPAG9 | C-Jun-amino-terminal kinase-interacting protein 4 OS=Homo sapiens OX=9606 GN=SPAG9 PE=1 SV=4 | 2 |
| MRPL43 | 39S ribosomal protein L43, mitochondrial OS=Homo sapiens OX=9606 GN=MRPL43 PE=1 SV=1 | 4 |
| DDX59 | Probable ATP-dependent RNA helicase DDX59 OS=Homo sapiens OX=9606 GN=DDX59 PE=1 SV=1 | 3 |
| UBR2 | E3 ubiquitin-protein ligase UBR2 OS=Homo sapiens OX=9606 GN=UBR2 PE=1 SV=1 | 6 |
| C19orf25 | UPF0449 protein C19orf25 OS=Homo sapiens OX=9606 GN=C19orf25 PE=1 SV=2 | 2 |
| TRIM62 | E3 ubiquitin-protein ligase TRIM62 OS=Homo sapiens OX=9606 GN=TRIM62 PE=1 SV=1 | 1 |
| RABIF | Guanine nucleotide exchange factor MSS4 OS=Homo sapiens OX=9606 GN=RABIF PE=1 SV=2 | 3 |
| GOLT1B | Vesicle transport protein GOT1B OS=Homo sapiens OX=9606 GN=GOLT1B PE=1 SV=1 | 1 |
| ARF3 | ADP-ribosylation factor 3 OS=Homo sapiens OX=9606 GN=ARF3 PE=1 SV=2 | 3 |
| EXOC7 | Exocyst complex component 7 OS=Homo sapiens OX=9606 GN=EXOC7 PE=1 SV=3 | 10 |
| NIPSNAP2 | Protein NipSnap homolog 2 OS=Homo sapiens OX=9606 GN=NIPSNAP2 PE=1 SV=1 | 8 |
| MTAP | S-methyl-5'-thioadenosine phosphorylase OS=Homo sapiens OX=9606 GN=MTAP PE=1 SV=2 | 10 |
| TRPS1 | Zinc finger transcription factor Trps1 OS=Homo sapiens OX=9606 GN=TRPS1 PE=1 SV=2 | 1 |
| DDT | D-dopachrome decarboxylase OS=Homo sapiens OX=9606 GN=DDT PE=1 SV=3 | 4 |
| TIMM23 | Mitochondrial import inner membrane translocase subunit Tim23 OS=Homo sapiens OX=9606 GN=TIMM23 PE=1 SV=1 | 1 |
| LRCH1 | Leucine-rich repeat and calponin homology domain-containing protein 1 OS=Homo sapiens OX=9606 GN=LRCH1 PE=1 SV=3 | 5 |
| RNF114 | E3 ubiquitin-protein ligase RNF114 OS=Homo sapiens OX=9606 GN=RNF114 PE=1 SV=1 | 8 |
| ARRDC1 | Arrestin domain-containing protein 1 OS=Homo sapiens OX=9606 GN=ARRDC1 PE=1 SV=1 | 1 |
| CAST | Calpastatin OS=Homo sapiens OX=9606 GN=CAST PE=1 SV=4 | 8 |
| FDPS | Farnesyl pyrophosphate synthase OS=Homo sapiens OX=9606 GN=FDPS PE=1 SV=4 | 6 |
| GCH1 | GTP cyclohydrolase 1 OS=Homo sapiens OX=9606 GN=GCH1 PE=1 SV=1 | 4 |
| BIRC2 | Baculoviral IAP repeat-containing protein 2 OS=Homo sapiens OX=9606 GN=BIRC2 PE=1 SV=2 | 13 |
| PAIP2B | Polyadenylate-binding protein-interacting protein 2B OS=Homo sapiens OX=9606 GN=PAIP2B PE=1 SV=2 | 3 |
| RER1 | Protein RER1 OS=Homo sapiens OX=9606 GN=RER1 PE=1 SV=1 | 4 |
| RPL11 | 60S ribosomal protein L11 OS=Homo sapiens OX=9606 GN=RPL11 PE=1 SV=2 | 4 |
| PFN2 | Profilin-2 OS=Homo sapiens OX=9606 GN=PFN2 PE=1 SV=3 | 7 |
| ARHGAP44 | Rho GTPase-activating protein 44 OS=Homo sapiens OX=9606 GN=ARHGAP44 PE=1 SV=1 | 5 |
| NDRG4 | Protein NDRG4 OS=Homo sapiens OX=9606 GN=NDRG4 PE=1 SV=2 | 2 |
| PLPBP | Pyridoxal phosphate homeostasis protein OS=Homo sapiens OX=9606 GN=PLPBP PE=1 SV=1 | 3 |
| CAD | CAD protein OS=Homo sapiens OX=9606 GN=CAD PE=1 SV=3 | 42 |
| MANF | Mesencephalic astrocyte-derived neurotrophic factor OS=Homo sapiens OX=9606 GN=MANF PE=1 SV=3 | 11 |
| DNAJC10 | DnaJ homolog subfamily C member 10 OS=Homo sapiens OX=9606 GN=DNAJC10 PE=1 SV=2 | 7 |
| MED20 | Mediator of RNA polymerase II transcription subunit 20 OS=Homo sapiens OX=9606 GN=MED20 PE=1 SV=1 | 2 |
| STMN2 | Stathmin-2 OS=Homo sapiens OX=9606 GN=STMN2 PE=1 SV=3 | 1 |
| XRCC5 | X-ray repair cross-complementing protein 5 OS=Homo sapiens OX=9606 GN=XRCC5 PE=1 SV=3 | 7 |
| SF3B1 | Splicing factor 3B subunit 1 OS=Homo sapiens OX=9606 GN=SF3B1 PE=1 SV=3 | 1 |
| COPG1 | Coatomer subunit gamma-1 OS=Homo sapiens OX=9606 GN=COPG1 PE=1 SV=1 | 29 |
| GSTM2 | Glutathione S-transferase Mu 2 OS=Homo sapiens OX=9606 GN=GSTM2 PE=1 SV=2 | 6 |
| AIMP2 | Aminoacyl tRNA synthase complex-interacting multifunctional protein 2 OS=Homo sapiens OX=9606 GN=AIMP2 PE=1 SV=2 | 4 |
| VKORC1L1 | Vitamin K epoxide reductase complex subunit 1-like protein 1 OS=Homo sapiens OX=9606 GN=VKORC1L1 PE=1 SV=2 | 1 |
| ATG5 | Autophagy protein 5 OS=Homo sapiens OX=9606 GN=ATG5 PE=1 SV=2 | 1 |
| TSG101 | Tumor susceptibility gene 101 protein OS=Homo sapiens OX=9606 GN=TSG101 PE=1 SV=2 | 10 |
| C5orf22 | UPF0489 protein C5orf22 OS=Homo sapiens OX=9606 GN=C5orf22 PE=1 SV=2 | 2 |
| ELAC2 | Zinc phosphodiesterase ELAC protein 2 OS=Homo sapiens OX=9606 GN=ELAC2 PE=1 SV=2 | 3 |
| ATP5MJ | ATP synthase subunit ATP5MJ, mitochondrial OS=Homo sapiens OX=9606 GN=ATP5MJ PE=1 SV=1 | 3 |
| CD9 | CD9 antigen OS=Homo sapiens OX=9606 GN=CD9 PE=1 SV=4 | 4 |
| GUK1 | Guanylate kinase OS=Homo sapiens OX=9606 GN=GUK1 PE=1 SV=2 | 4 |
| NPM1 | Nucleophosmin OS=Homo sapiens OX=9606 GN=NPM1 PE=1 SV=2 | 9 |
| C12orf29 | Uncharacterized protein C12orf29 OS=Homo sapiens OX=9606 GN=C12orf29 PE=1 SV=2 | 2 |
| MVB12A | Multivesicular body subunit 12A OS=Homo sapiens OX=9606 GN=MVB12A PE=1 SV=1 | 6 |
| VAMP3 | Vesicle-associated membrane protein 3 OS=Homo sapiens OX=9606 GN=VAMP3 PE=1 SV=3 | 3 |
| GTPBP10 | GTP-binding protein 10 OS=Homo sapiens OX=9606 GN=GTPBP10 PE=1 SV=1 | 19 |
| ZNF213 | Zinc finger protein 213 OS=Homo sapiens OX=9606 GN=ZNF213 PE=1 SV=2 | 1 |
| SPCS1 | Signal peptidase complex subunit 1 OS=Homo sapiens OX=9606 GN=SPCS1 PE=1 SV=5 | 1 |
| PTTG1 | Securin OS=Homo sapiens OX=9606 GN=PTTG1 PE=1 SV=1 | 2 |
| AVL9 | Late secretory pathway protein AVL9 homolog OS=Homo sapiens OX=9606 GN=AVL9 PE=1 SV=1 | 4 |
| ALDH5A1 | Succinate-semialdehyde dehydrogenase, mitochondrial OS=Homo sapiens OX=9606 GN=ALDH5A1 PE=1 SV=2 | 2 |
| ABHD16A | Phosphatidylserine lipase ABHD16A OS=Homo sapiens OX=9606 GN=ABHD16A PE=1 SV=3 | 9 |
| CKB | Creatine kinase B-type OS=Homo sapiens OX=9606 GN=CKB PE=1 SV=1 | 21 |
| KIF20B | Kinesin-like protein KIF20B OS=Homo sapiens OX=9606 GN=KIF20B PE=1 SV=3 | 4 |
| DCTN5 | Dynactin subunit 5 OS=Homo sapiens OX=9606 GN=DCTN5 PE=1 SV=1 | 1 |
| GATC | Glutamyl-tRNA(Gln) amidotransferase subunit C, mitochondrial OS=Homo sapiens OX=9606 GN=GATC PE=1 SV=1 | 2 |
| EXOC6 | Exocyst complex component 6 OS=Homo sapiens OX=9606 GN=EXOC6 PE=1 SV=3 | 2 |
| PRUNE2 | Protein prune homolog 2 OS=Homo sapiens OX=9606 GN=PRUNE2 PE=1 SV=3 | 1 |
| COPB1 | Coatomer subunit beta OS=Homo sapiens OX=9606 GN=COPB1 PE=1 SV=3 | 28 |
| MAGOH | Protein mago nashi homolog OS=Homo sapiens OX=9606 GN=MAGOH PE=1 SV=1 | 2 |
| DEPDC7 | DEP domain-containing protein 7 OS=Homo sapiens OX=9606 GN=DEPDC7 PE=2 SV=1 | 10 |
| BMP2K | BMP-2-inducible protein kinase OS=Homo sapiens OX=9606 GN=BMP2K PE=1 SV=2 | 5 |
| RLF | Zinc finger protein Rlf OS=Homo sapiens OX=9606 GN=RLF PE=1 SV=2 | 1 |
| PDCD2 | Programmed cell death protein 2 OS=Homo sapiens OX=9606 GN=PDCD2 PE=1 SV=2 | 1 |
| ARIH1 | E3 ubiquitin-protein ligase ARIH1 OS=Homo sapiens OX=9606 GN=ARIH1 PE=1 SV=2 | 4 |
| MCM2 | DNA replication licensing factor MCM2 OS=Homo sapiens OX=9606 GN=MCM2 PE=1 SV=4 | 18 |
| ECT2 | Protein ECT2 OS=Homo sapiens OX=9606 GN=ECT2 PE=1 SV=4 | 4 |
| UBA1 | Ubiquitin-like modifier-activating enzyme 1 OS=Homo sapiens OX=9606 GN=UBA1 PE=1 SV=3 | 36 |
| CASP3 | Caspase-3 OS=Homo sapiens OX=9606 GN=CASP3 PE=1 SV=2 | 4 |
| RABGAP1L | Rab GTPase-activating protein 1-like OS=Homo sapiens OX=9606 GN=RABGAP1L PE=1 SV=1 | 8 |
| KNG1 | Kininogen-1 OS=Homo sapiens OX=9606 GN=KNG1 PE=1 SV=2 | 15 |
| HEXA | Beta-hexosaminidase subunit alpha OS=Homo sapiens OX=9606 GN=HEXA PE=1 SV=2 | 6 |
| LRRC28 | Leucine-rich repeat-containing protein 28 OS=Homo sapiens OX=9606 GN=LRRC28 PE=2 SV=1 | 1 |
| TXNRD1 | Thioredoxin reductase 1, cytoplasmic OS=Homo sapiens OX=9606 GN=TXNRD1 PE=1 SV=3 | 29 |
| GALK1 | Galactokinase OS=Homo sapiens OX=9606 GN=GALK1 PE=1 SV=1 | 18 |
| PPT1 | Palmitoyl-protein thioesterase 1 OS=Homo sapiens OX=9606 GN=PPT1 PE=1 SV=1 | 3 |
| SART3 | Squamous cell carcinoma antigen recognized by T-cells 3 OS=Homo sapiens OX=9606 GN=SART3 PE=1 SV=1 | 3 |
| CDC42 | Cell division control protein 42 homolog OS=Homo sapiens OX=9606 GN=CDC42 PE=1 SV=2 | 9 |
| CLASP2 | CLIP-associating protein 2 OS=Homo sapiens OX=9606 GN=CLASP2 PE=1 SV=3 | 1 |
| MTFMT | Methionyl-tRNA formyltransferase, mitochondrial OS=Homo sapiens OX=9606 GN=MTFMT PE=1 SV=2 | 2 |
| MXI1 | Max-interacting protein 1 OS=Homo sapiens OX=9606 GN=MXI1 PE=1 SV=2 | 1 |
| DAZL | Deleted in azoospermia-like OS=Homo sapiens OX=9606 GN=DAZL PE=1 SV=1 | 3 |
| MGST2 | Microsomal glutathione S-transferase 2 OS=Homo sapiens OX=9606 GN=MGST2 PE=1 SV=1 | 2 |
| STX5 | Syntaxin-5 OS=Homo sapiens OX=9606 GN=STX5 PE=1 SV=2 | 1 |
| HSD17B12 | Very-long-chain 3-oxoacyl-CoA reductase OS=Homo sapiens OX=9606 GN=HSD17B12 PE=1 SV=2 | 6 |
| THNSL1 | Threonine synthase-like 1 OS=Homo sapiens OX=9606 GN=THNSL1 PE=1 SV=2 | 21 |
| ADAM10 | Disintegrin and metalloproteinase domain-containing protein 10 OS=Homo sapiens OX=9606 GN=ADAM10 PE=1 SV=1 | 3 |
| CISD1 | CDGSH iron-sulfur domain-containing protein 1 OS=Homo sapiens OX=9606 GN=CISD1 PE=1 SV=1 | 5 |
| EFL1 | Elongation factor-like GTPase 1 OS=Homo sapiens OX=9606 GN=EFL1 PE=1 SV=2 | 18 |
| CSNK2A1 | Casein kinase II subunit alpha OS=Homo sapiens OX=9606 GN=CSNK2A1 PE=1 SV=1 | 4 |
| ZC3H8 | Zinc finger CCCH domain-containing protein 8 OS=Homo sapiens OX=9606 GN=ZC3H8 PE=1 SV=2 | 1 |
| TARBP1 | Probable methyltransferase TARBP1 OS=Homo sapiens OX=9606 GN=TARBP1 PE=1 SV=1 | 3 |
| PREB | Prolactin regulatory element-binding protein OS=Homo sapiens OX=9606 GN=PREB PE=1 SV=2 | 9 |
| PABIR1 | PPP2R1A-PPP2R2A-interacting phosphatase regulator 1 OS=Homo sapiens OX=9606 GN=PABIR1 PE=1 SV=1 | 1 |
| ACOX3 | Peroxisomal acyl-coenzyme A oxidase 3 OS=Homo sapiens OX=9606 GN=ACOX3 PE=1 SV=2 | 5 |
| ARHGEF6 | Rho guanine nucleotide exchange factor 6 OS=Homo sapiens OX=9606 GN=ARHGEF6 PE=1 SV=2 | 1 |
| IL17B | Interleukin-17B OS=Homo sapiens OX=9606 GN=IL17B PE=2 SV=1 | 8 |
| BST2 | Bone marrow stromal antigen 2 OS=Homo sapiens OX=9606 GN=BST2 PE=1 SV=1 | 2 |
| KPNA2 | Importin subunit alpha-1 OS=Homo sapiens OX=9606 GN=KPNA2 PE=1 SV=1 | 8 |
| NDUFA12 | NADH dehydrogenase [ubiquinone] 1 alpha subcomplex subunit 12 OS=Homo sapiens OX=9606 GN=NDUFA12 PE=1 SV=1 | 2 |
| LACTB2 | Endoribonuclease LACTB2 OS=Homo sapiens OX=9606 GN=LACTB2 PE=1 SV=2 | 9 |
| NISCH | Nischarin OS=Homo sapiens OX=9606 GN=NISCH PE=1 SV=3 | 2 |
| NLRP2 | NACHT, LRR and PYD domains-containing protein 2 OS=Homo sapiens OX=9606 GN=NLRP2 PE=1 SV=1 | 82 |
| TXNDC5 | Thioredoxin domain-containing protein 5 OS=Homo sapiens OX=9606 GN=TXNDC5 PE=1 SV=2 | 21 |
| ECI2 | Enoyl-CoA delta isomerase 2 OS=Homo sapiens OX=9606 GN=ECI2 PE=1 SV=4 | 14 |
| DPYSL3 | Dihydropyrimidinase-related protein 3 OS=Homo sapiens OX=9606 GN=DPYSL3 PE=1 SV=1 | 4 |
| PLCL1 | Inactive phospholipase C-like protein 1 OS=Homo sapiens OX=9606 GN=PLCL1 PE=1 SV=3 | 8 |
| STX12 | Syntaxin-12 OS=Homo sapiens OX=9606 GN=STX12 PE=1 SV=1 | 4 |
| GSTA1 | Glutathione S-transferase A1 OS=Homo sapiens OX=9606 GN=GSTA1 PE=1 SV=3 | 5 |
| NDUFS3 | NADH dehydrogenase [ubiquinone] iron-sulfur protein 3, mitochondrial OS=Homo sapiens OX=9606 GN=NDUFS3 PE=1 SV=1 | 10 |
| MTMR1 | Myotubularin-related protein 1 OS=Homo sapiens OX=9606 GN=MTMR1 PE=1 SV=4 | 4 |
| ABHD11 | Protein ABHD11 OS=Homo sapiens OX=9606 GN=ABHD11 PE=1 SV=1 | 4 |
| MAP2K6 | Dual specificity mitogen-activated protein kinase kinase 6 OS=Homo sapiens OX=9606 GN=MAP2K6 PE=1 SV=1 | 6 |
| DNAJB9 | DnaJ homolog subfamily B member 9 OS=Homo sapiens OX=9606 GN=DNAJB9 PE=1 SV=1 | 9 |
| C1orf50 | Uncharacterized protein C1orf50 OS=Homo sapiens OX=9606 GN=C1orf50 PE=1 SV=2 | 7 |
| TAOK2 | Serine/threonine-protein kinase TAO2 OS=Homo sapiens OX=9606 GN=TAOK2 PE=1 SV=2 | 3 |
| MAP2K4 | Dual specificity mitogen-activated protein kinase kinase 4 OS=Homo sapiens OX=9606 GN=MAP2K4 PE=1 SV=1 | 4 |
| AGFG2 | Arf-GAP domain and FG repeat-containing protein 2 OS=Homo sapiens OX=9606 GN=AGFG2 PE=1 SV=2 | 5 |
| CHP1 | Calcineurin B homologous protein 1 OS=Homo sapiens OX=9606 GN=CHP1 PE=1 SV=3 | 5 |
| PAFAH1B2 | Platelet-activating factor acetylhydrolase IB subunit alpha2 OS=Homo sapiens OX=9606 GN=PAFAH1B2 PE=1 SV=1 | 5 |
| STX7 | Syntaxin-7 OS=Homo sapiens OX=9606 GN=STX7 PE=1 SV=4 | 4 |
| PYM1 | Partner of Y14 and mago OS=Homo sapiens OX=9606 GN=PYM1 PE=1 SV=1 | 3 |
| ARFGEF1 | Brefeldin A-inhibited guanine nucleotide-exchange protein 1 OS=Homo sapiens OX=9606 GN=ARFGEF1 PE=1 SV=2 | 3 |
| MAP2K1 | Dual specificity mitogen-activated protein kinase kinase 1 OS=Homo sapiens OX=9606 GN=MAP2K1 PE=1 SV=2 | 5 |
| PCNP | PEST proteolytic signal-containing nuclear protein OS=Homo sapiens OX=9606 GN=PCNP PE=1 SV=2 | 2 |
| ATP2B4 | Plasma membrane calcium-transporting ATPase 4 OS=Homo sapiens OX=9606 GN=ATP2B4 PE=1 SV=2 | 6 |
| CLTC | Clathrin heavy chain 1 OS=Homo sapiens OX=9606 GN=CLTC PE=1 SV=5 | 79 |
| HSD17B10 | 3-hydroxyacyl-CoA dehydrogenase type-2 OS=Homo sapiens OX=9606 GN=HSD17B10 PE=1 SV=3 | 14 |
| UBLCP1 | Ubiquitin-like domain-containing CTD phosphatase 1 OS=Homo sapiens OX=9606 GN=UBLCP1 PE=1 SV=2 | 3 |
| EOGT | EGF domain-specific O-linked N-acetylglucosamine transferase OS=Homo sapiens OX=9606 GN=EOGT PE=1 SV=1 | 1 |
| H1-4 | Histone H1.4 OS=Homo sapiens OX=9606 GN=H1-4 PE=1 SV=2 | 2 |
| SELENOO | Protein adenylyltransferase SelO, mitochondrial OS=Homo sapiens OX=9606 GN=SELENOO PE=1 SV=3 | 1 |
| LCMT1 | Leucine carboxyl methyltransferase 1 OS=Homo sapiens OX=9606 GN=LCMT1 PE=1 SV=2 | 18 |
| ABHD10 | Palmitoyl-protein thioesterase ABHD10, mitochondrial OS=Homo sapiens OX=9606 GN=ABHD10 PE=1 SV=1 | 16 |
| CPOX | Oxygen-dependent coproporphyrinogen-III oxidase, mitochondrial OS=Homo sapiens OX=9606 GN=CPOX PE=1 SV=3 | 28 |
| IDH3G | Isocitrate dehydrogenase [NAD] subunit gamma, mitochondrial OS=Homo sapiens OX=9606 GN=IDH3G PE=1 SV=1 | 9 |
| IGFBP6 | Insulin-like growth factor-binding protein 6 OS=Homo sapiens OX=9606 GN=IGFBP6 PE=1 SV=1 | 4 |
| TXN2 | Thioredoxin, mitochondrial OS=Homo sapiens OX=9606 GN=TXN2 PE=1 SV=2 | 3 |
| POMGNT2 | Protein O-linked-mannose beta-1,4-N-acetylglucosaminyltransferase 2 OS=Homo sapiens OX=9606 GN=POMGNT2 PE=1 SV=1 | 2 |
| DCTN2 | Dynactin subunit 2 OS=Homo sapiens OX=9606 GN=DCTN2 PE=1 SV=4 | 16 |
| OTUD4 | OTU domain-containing protein 4 OS=Homo sapiens OX=9606 GN=OTUD4 PE=1 SV=4 | 6 |
| MFN1 | Mitofusin-1 OS=Homo sapiens OX=9606 GN=MFN1 PE=1 SV=3 | 18 |
| TTYH1 | Protein tweety homolog 1 OS=Homo sapiens OX=9606 GN=TTYH1 PE=1 SV=1 | 3 |
| API5 | Apoptosis inhibitor 5 OS=Homo sapiens OX=9606 GN=API5 PE=1 SV=3 | 10 |
| LMAN2 | Vesicular integral-membrane protein VIP36 OS=Homo sapiens OX=9606 GN=LMAN2 PE=1 SV=1 | 19 |
| PBXIP1 | Pre-B-cell leukemia transcription factor-interacting protein 1 OS=Homo sapiens OX=9606 GN=PBXIP1 PE=1 SV=1 | 13 |
| MTMR6 | Myotubularin-related protein 6 OS=Homo sapiens OX=9606 GN=MTMR6 PE=1 SV=3 | 3 |
| NLRP13 | NACHT, LRR and PYD domains-containing protein 13 OS=Homo sapiens OX=9606 GN=NLRP13 PE=2 SV=2 | 54 |
| SHFL | Shiftless antiviral inhibitor of ribosomal frameshifting protein OS=Homo sapiens OX=9606 GN=SHFL PE=1 SV=2 | 4 |
| TDRP | Testis development-related protein OS=Homo sapiens OX=9606 GN=TDRP PE=1 SV=2 | 4 |
| DYNLT1 | Dynein light chain Tctex-type 1 OS=Homo sapiens OX=9606 GN=DYNLT1 PE=1 SV=1 | 3 |
| PIN1 | Peptidyl-prolyl cis-trans isomerase NIMA-interacting 1 OS=Homo sapiens OX=9606 GN=PIN1 PE=1 SV=1 | 9 |
| MAGT1 | Magnesium transporter protein 1 OS=Homo sapiens OX=9606 GN=MAGT1 PE=1 SV=1 | 4 |
| WFDC2 | WAP four-disulfide core domain protein 2 OS=Homo sapiens OX=9606 GN=WFDC2 PE=1 SV=2 | 3 |
| SLC25A22 | Mitochondrial glutamate carrier 1 OS=Homo sapiens OX=9606 GN=SLC25A22 PE=1 SV=1 | 6 |
| NUP153 | Nuclear pore complex protein Nup153 OS=Homo sapiens OX=9606 GN=NUP153 PE=1 SV=2 | 1 |
| VAMP7 | Vesicle-associated membrane protein 7 OS=Homo sapiens OX=9606 GN=VAMP7 PE=1 SV=3 | 8 |
| H1-8 | Histone H1.8 OS=Homo sapiens OX=9606 GN=H1-8 PE=2 SV=1 | 7 |
| CUL5 | Cullin-5 OS=Homo sapiens OX=9606 GN=CUL5 PE=1 SV=4 | 33 |
| CARMIL1 | F-actin-uncapping protein LRRC16A OS=Homo sapiens OX=9606 GN=CARMIL1 PE=1 SV=1 | 9 |
| CNPY3 | Protein canopy homolog 3 OS=Homo sapiens OX=9606 GN=CNPY3 PE=1 SV=1 | 8 |
| PCNA | Proliferating cell nuclear antigen OS=Homo sapiens OX=9606 GN=PCNA PE=1 SV=1 | 19 |
| SGPL1 | Sphingosine-1-phosphate lyase 1 OS=Homo sapiens OX=9606 GN=SGPL1 PE=1 SV=3 | 4 |
| CLDN12 | Claudin-12 OS=Homo sapiens OX=9606 GN=CLDN12 PE=1 SV=1 | 1 |
| PHKA2 | Phosphorylase b kinase regulatory subunit alpha, liver isoform OS=Homo sapiens OX=9606 GN=PHKA2 PE=1 SV=1 | 1 |
| PPIP5K2 | Inositol hexakisphosphate and diphosphoinositol-pentakisphosphate kinase 2 OS=Homo sapiens OX=9606 GN=PPIP5K2 PE=1 SV=3 | 3 |
| PRPS2 | Ribose-phosphate pyrophosphokinase 2 OS=Homo sapiens OX=9606 GN=PRPS2 PE=1 SV=2 | 2 |
| RABAC1 | Prenylated Rab acceptor protein 1 OS=Homo sapiens OX=9606 GN=RABAC1 PE=1 SV=1 | 1 |
| DDX28 | Probable ATP-dependent RNA helicase DDX28 OS=Homo sapiens OX=9606 GN=DDX28 PE=1 SV=2 | 6 |
| HUNK | Hormonally up-regulated neu tumor-associated kinase OS=Homo sapiens OX=9606 GN=HUNK PE=1 SV=1 | 1 |
| IGSF10 | Immunoglobulin superfamily member 10 OS=Homo sapiens OX=9606 GN=IGSF10 PE=1 SV=1 | 19 |
| ASPH | Aspartyl/asparaginyl beta-hydroxylase OS=Homo sapiens OX=9606 GN=ASPH PE=1 SV=3 | 2 |
| ERO1B | ERO1-like protein beta OS=Homo sapiens OX=9606 GN=ERO1B PE=1 SV=2 | 10 |
| ELANE | Neutrophil elastase OS=Homo sapiens OX=9606 GN=ELANE PE=1 SV=1 | 1 |
| SIN3A | Paired amphipathic helix protein Sin3a OS=Homo sapiens OX=9606 GN=SIN3A PE=1 SV=2 | 16 |
| **HABP4** | Intracellular hyaluronan-binding protein 4 OS=Homo sapiens OX=9606 GN=HABP4 PE=1 SV=1 | 5 |
| PDF | Peptide deformylase, mitochondrial OS=Homo sapiens OX=9606 GN=PDF PE=1 SV=1 | 7 |
| MRPS23 | 28S ribosomal protein S23, mitochondrial OS=Homo sapiens OX=9606 GN=MRPS23 PE=1 SV=2 | 2 |
| ERAL1 | GTPase Era, mitochondrial OS=Homo sapiens OX=9606 GN=ERAL1 PE=1 SV=2 | 2 |
| SEC23IP | SEC23-interacting protein OS=Homo sapiens OX=9606 GN=SEC23IP PE=1 SV=1 | 17 |
| PRDX6 | Peroxiredoxin-6 OS=Homo sapiens OX=9606 GN=PRDX6 PE=1 SV=3 | 24 |
| PRDX4 | Peroxiredoxin-4 OS=Homo sapiens OX=9606 GN=PRDX4 PE=1 SV=1 | 7 |
| TJP1 | Tight junction protein ZO-1 OS=Homo sapiens OX=9606 GN=TJP1 PE=1 SV=3 | 6 |
| SACM1L | Phosphatidylinositol-3-phosphatase SAC1 OS=Homo sapiens OX=9606 GN=SACM1L PE=1 SV=2 | 24 |
| CHD1L | Chromodomain-helicase-DNA-binding protein 1-like OS=Homo sapiens OX=9606 GN=CHD1L PE=1 SV=3 | 2 |
| UBR4 | E3 ubiquitin-protein ligase UBR4 OS=Homo sapiens OX=9606 GN=UBR4 PE=1 SV=1 | 13 |
| WDFY1 | WD repeat and FYVE domain-containing protein 1 OS=Homo sapiens OX=9606 GN=WDFY1 PE=1 SV=1 | 22 |
| HIRIP3 | HIRA-interacting protein 3 OS=Homo sapiens OX=9606 GN=HIRIP3 PE=1 SV=3 | 1 |
| GTF2A1L | TFIIA-alpha and beta-like factor OS=Homo sapiens OX=9606 GN=GTF2A1L PE=1 SV=2 | 2 |
| NEMF | Ribosome quality control complex subunit NEMF OS=Homo sapiens OX=9606 GN=NEMF PE=1 SV=4 | 2 |
| CHCHD4 | Mitochondrial intermembrane space import and assembly protein 40 OS=Homo sapiens OX=9606 GN=CHCHD4 PE=1 SV=1 | 4 |
| COPS8 | COP9 signalosome complex subunit 8 OS=Homo sapiens OX=9606 GN=COPS8 PE=1 SV=1 | 4 |
| SPOCK1 | Testican-1 OS=Homo sapiens OX=9606 GN=SPOCK1 PE=1 SV=1 | 6 |
| S100A7A | Protein S100-A7A OS=Homo sapiens OX=9606 GN=S100A7A PE=1 SV=3 | 3 |
| DYNC1H1 | Cytoplasmic dynein 1 heavy chain 1 OS=Homo sapiens OX=9606 GN=DYNC1H1 PE=1 SV=5 | 169 |
| PRPSAP1 | Phosphoribosyl pyrophosphate synthase-associated protein 1 OS=Homo sapiens OX=9606 GN=PRPSAP1 PE=1 SV=2 | 15 |
| SMC4 | Structural maintenance of chromosomes protein 4 OS=Homo sapiens OX=9606 GN=SMC4 PE=1 SV=2 | 3 |
| MRE11 | Double-strand break repair protein MRE11 OS=Homo sapiens OX=9606 GN=MRE11 PE=1 SV=3 | 4 |
| RPL18A | 60S ribosomal protein L18a OS=Homo sapiens OX=9606 GN=RPL18A PE=1 SV=2 | 5 |
| BCAT2 | Branched-chain-amino-acid aminotransferase, mitochondrial OS=Homo sapiens OX=9606 GN=BCAT2 PE=1 SV=2 | 19 |
| FAM177A1 | Protein FAM177A1 OS=Homo sapiens OX=9606 GN=FAM177A1 PE=1 SV=1 | 1 |
| BAX | Apoptosis regulator BAX OS=Homo sapiens OX=9606 GN=BAX PE=1 SV=1 | 3 |
| PABPN1L | Embryonic polyadenylate-binding protein 2 OS=Homo sapiens OX=9606 GN=PABPN1L PE=2 SV=1 | 4 |
| CD59 | CD59 glycoprotein OS=Homo sapiens OX=9606 GN=CD59 PE=1 SV=1 | 6 |
| MPI | Mannose-6-phosphate isomerase OS=Homo sapiens OX=9606 GN=MPI PE=1 SV=2 | 11 |
| TRIM71 | E3 ubiquitin-protein ligase TRIM71 OS=Homo sapiens OX=9606 GN=TRIM71 PE=1 SV=1 | 9 |
| DRG2 | Developmentally-regulated GTP-binding protein 2 OS=Homo sapiens OX=9606 GN=DRG2 PE=1 SV=1 | 2 |
| RIC8A | Synembryn-A OS=Homo sapiens OX=9606 GN=RIC8A PE=1 SV=3 | 2 |
| SPINT2 | Kunitz-type protease inhibitor 2 OS=Homo sapiens OX=9606 GN=SPINT2 PE=1 SV=2 | 2 |
| PRAF2 | PRA1 family protein 2 OS=Homo sapiens OX=9606 GN=PRAF2 PE=1 SV=1 | 2 |
| MBOAT7 | Lysophospholipid acyltransferase 7 OS=Homo sapiens OX=9606 GN=MBOAT7 PE=1 SV=2 | 4 |
| AAAS | Aladin OS=Homo sapiens OX=9606 GN=AAAS PE=1 SV=1 | 12 |
| SNX27 | Sorting nexin-27 OS=Homo sapiens OX=9606 GN=SNX27 PE=1 SV=2 | 7 |
| EPHX1 | Epoxide hydrolase 1 OS=Homo sapiens OX=9606 GN=EPHX1 PE=1 SV=1 | 20 |
| DNAJC19 | Mitochondrial import inner membrane translocase subunit TIM14 OS=Homo sapiens OX=9606 GN=DNAJC19 PE=1 SV=3 | 5 |
| MTMR3 | Myotubularin-related protein 3 OS=Homo sapiens OX=9606 GN=MTMR3 PE=1 SV=3 | 32 |
| ACAD11 | Acyl-CoA dehydrogenase family member 11 OS=Homo sapiens OX=9606 GN=ACAD11 PE=1 SV=3 | 19 |
| SLC25A4 | ADP/ATP translocase 1 OS=Homo sapiens OX=9606 GN=SLC25A4 PE=1 SV=4 | 10 |
| BPHL | Valacyclovir hydrolase OS=Homo sapiens OX=9606 GN=BPHL PE=1 SV=1 | 9 |
| FSD1 | Fibronectin type III and SPRY domain-containing protein 1 OS=Homo sapiens OX=9606 GN=FSD1 PE=1 SV=1 | 7 |
| TPR | Nucleoprotein TPR OS=Homo sapiens OX=9606 GN=TPR PE=1 SV=3 | 19 |
| ABHD12 | Lysophosphatidylserine lipase ABHD12 OS=Homo sapiens OX=9606 GN=ABHD12 PE=1 SV=2 | 14 |
| MFSD6 | Major facilitator superfamily domain-containing protein 6 OS=Homo sapiens OX=9606 GN=MFSD6 PE=1 SV=2 | 1 |
| CSTB | Cystatin-B OS=Homo sapiens OX=9606 GN=CSTB PE=1 SV=2 | 6 |
| SCRN1 | Secernin-1 OS=Homo sapiens OX=9606 GN=SCRN1 PE=1 SV=2 | 6 |
| NLRP7 | NACHT, LRR and PYD domains-containing protein 7 OS=Homo sapiens OX=9606 GN=NLRP7 PE=1 SV=1 | 64 |
| CUL3 | Cullin-3 OS=Homo sapiens OX=9606 GN=CUL3 PE=1 SV=2 | 17 |
| PGM1 | Phosphoglucomutase-1 OS=Homo sapiens OX=9606 GN=PGM1 PE=1 SV=3 | 21 |
| APOO | MICOS complex subunit MIC26 OS=Homo sapiens OX=9606 GN=APOO PE=1 SV=1 | 6 |
| APOD | Apolipoprotein D OS=Homo sapiens OX=9606 GN=APOD PE=1 SV=1 | 7 |
| BCKDHA | 2-oxoisovalerate dehydrogenase subunit alpha, mitochondrial OS=Homo sapiens OX=9606 GN=BCKDHA PE=1 SV=2 | 7 |
| RAB30 | Ras-related protein Rab-30 OS=Homo sapiens OX=9606 GN=RAB30 PE=1 SV=2 | 3 |
| ASL | Argininosuccinate lyase OS=Homo sapiens OX=9606 GN=ASL PE=1 SV=4 | 9 |
| DDX39B | Spliceosome RNA helicase DDX39B OS=Homo sapiens OX=9606 GN=DDX39B PE=1 SV=1 | 8 |
| LSM4 | U6 snRNA-associated Sm-like protein LSm4 OS=Homo sapiens OX=9606 GN=LSM4 PE=1 SV=1 | 8 |
| AP3D1 | AP-3 complex subunit delta-1 OS=Homo sapiens OX=9606 GN=AP3D1 PE=1 SV=1 | 9 |
| PALM3 | Paralemmin-3 OS=Homo sapiens OX=9606 GN=PALM3 PE=1 SV=2 | 6 |
| MRPS31 | 28S ribosomal protein S31, mitochondrial OS=Homo sapiens OX=9606 GN=MRPS31 PE=1 SV=3 | 3 |
| CAPRIN1 | Caprin-1 OS=Homo sapiens OX=9606 GN=CAPRIN1 PE=1 SV=2 | 6 |
| STK38L | Serine/threonine-protein kinase 38-like OS=Homo sapiens OX=9606 GN=STK38L PE=1 SV=3 | 8 |
| USE1 | Vesicle transport protein USE1 OS=Homo sapiens OX=9606 GN=USE1 PE=1 SV=2 | 6 |
| RUFY3 | Protein RUFY3 OS=Homo sapiens OX=9606 GN=RUFY3 PE=1 SV=1 | 5 |
| SOD1 | Superoxide dismutase [Cu-Zn] OS=Homo sapiens OX=9606 GN=SOD1 PE=1 SV=2 | 9 |
| LAMP2 | Lysosome-associated membrane glycoprotein 2 OS=Homo sapiens OX=9606 GN=LAMP2 PE=1 SV=2 | 7 |
| MYH10 | Myosin-10 OS=Homo sapiens OX=9606 GN=MYH10 PE=1 SV=3 | 53 |
| RTN3 | Reticulon-3 OS=Homo sapiens OX=9606 GN=RTN3 PE=1 SV=2 | 4 |
| WFS1 | Wolframin OS=Homo sapiens OX=9606 GN=WFS1 PE=1 SV=2 | 16 |
| TCL1A | T-cell leukemia/lymphoma protein 1A OS=Homo sapiens OX=9606 GN=TCL1A PE=1 SV=1 | 9 |
| IMPDH2 | Inosine-5'-monophosphate dehydrogenase 2 OS=Homo sapiens OX=9606 GN=IMPDH2 PE=1 SV=2 | 27 |
| ADD1 | Alpha-adducin OS=Homo sapiens OX=9606 GN=ADD1 PE=1 SV=2 | 4 |
| EIF4B | Eukaryotic translation initiation factor 4B OS=Homo sapiens OX=9606 GN=EIF4B PE=1 SV=2 | 6 |
| NUCKS1 | Nuclear ubiquitous casein and cyclin-dependent kinase substrate 1 OS=Homo sapiens OX=9606 GN=NUCKS1 PE=1 SV=1 | 1 |
| UFSP2 | Ufm1-specific protease 2 OS=Homo sapiens OX=9606 GN=UFSP2 PE=1 SV=3 | 7 |
| GCAT | 2-amino-3-ketobutyrate coenzyme A ligase, mitochondrial OS=Homo sapiens OX=9606 GN=GCAT PE=1 SV=1 | 16 |
| PMVK | Phosphomevalonate kinase OS=Homo sapiens OX=9606 GN=PMVK PE=1 SV=3 | 6 |
| HSPA1A | Heat shock 70 kDa protein 1A OS=Homo sapiens OX=9606 GN=HSPA1A PE=1 SV=1 | 38 |
| TMED2 | Transmembrane emp24 domain-containing protein 2 OS=Homo sapiens OX=9606 GN=TMED2 PE=1 SV=1 | 4 |
| COMMD4 | COMM domain-containing protein 4 OS=Homo sapiens OX=9606 GN=COMMD4 PE=1 SV=1 | 8 |
| MAN2B1 | Lysosomal alpha-mannosidase OS=Homo sapiens OX=9606 GN=MAN2B1 PE=1 SV=3 | 1 |
| COPS3 | COP9 signalosome complex subunit 3 OS=Homo sapiens OX=9606 GN=COPS3 PE=1 SV=3 | 17 |
| TRIP12 | E3 ubiquitin-protein ligase TRIP12 OS=Homo sapiens OX=9606 GN=TRIP12 PE=1 SV=1 | 4 |
| VCAN | Versican core protein OS=Homo sapiens OX=9606 GN=VCAN PE=1 SV=3 | 11 |
| SGTA | Small glutamine-rich tetratricopeptide repeat-containing protein alpha OS=Homo sapiens OX=9606 GN=SGTA PE=1 SV=1 | 4 |
| CDH3 | Cadherin-3 OS=Homo sapiens OX=9606 GN=CDH3 PE=1 SV=2 | 1 |
| KALRN | Kalirin OS=Homo sapiens OX=9606 GN=KALRN PE=1 SV=3 | 12 |
| STXBP3 | Syntaxin-binding protein 3 OS=Homo sapiens OX=9606 GN=STXBP3 PE=1 SV=2 | 15 |
| NANS | Sialic acid synthase OS=Homo sapiens OX=9606 GN=NANS PE=1 SV=2 | 13 |
| RAB4B | Ras-related protein Rab-4B OS=Homo sapiens OX=9606 GN=RAB4B PE=1 SV=1 | 10 |
| RPS28 | 40S ribosomal protein S28 OS=Homo sapiens OX=9606 GN=RPS28 PE=1 SV=1 | 4 |
| NT5DC3 | 5'-nucleotidase domain-containing protein 3 OS=Homo sapiens OX=9606 GN=NT5DC3 PE=1 SV=1 | 5 |
| C1QBP | Complement component 1 Q subcomponent-binding protein, mitochondrial OS=Homo sapiens OX=9606 GN=C1QBP PE=1 SV=1 | 6 |
| TXNDC9 | Thioredoxin domain-containing protein 9 OS=Homo sapiens OX=9606 GN=TXNDC9 PE=1 SV=2 | 3 |
| CDK1 | Cyclin-dependent kinase 1 OS=Homo sapiens OX=9606 GN=CDK1 PE=1 SV=3 | 11 |
| RUVBL1 | RuvB-like 1 OS=Homo sapiens OX=9606 GN=RUVBL1 PE=1 SV=1 | 14 |
| PDPK1 | 3-phosphoinositide-dependent protein kinase 1 OS=Homo sapiens OX=9606 GN=PDPK1 PE=1 SV=1 | 2 |
| NCEH1 | Neutral cholesterol ester hydrolase 1 OS=Homo sapiens OX=9606 GN=NCEH1 PE=1 SV=3 | 13 |
| UPF3B | Regulator of nonsense transcripts 3B OS=Homo sapiens OX=9606 GN=UPF3B PE=1 SV=1 | 2 |
| PTPN23 | Tyrosine-protein phosphatase non-receptor type 23 OS=Homo sapiens OX=9606 GN=PTPN23 PE=1 SV=1 | 10 |
| RAB3GAP1 | Rab3 GTPase-activating protein catalytic subunit OS=Homo sapiens OX=9606 GN=RAB3GAP1 PE=1 SV=3 | 18 |
| KIF1B | Kinesin-like protein KIF1B OS=Homo sapiens OX=9606 GN=KIF1B PE=1 SV=5 | 3 |
| SUB1 | Activated RNA polymerase II transcriptional coactivator p15 OS=Homo sapiens OX=9606 GN=SUB1 PE=1 SV=3 | 7 |
| TTLL12 | Tubulin--tyrosine ligase-like protein 12 OS=Homo sapiens OX=9606 GN=TTLL12 PE=1 SV=2 | 33 |
| DTYMK | Thymidylate kinase OS=Homo sapiens OX=9606 GN=DTYMK PE=1 SV=4 | 12 |
| ESYT2 | Extended synaptotagmin-2 OS=Homo sapiens OX=9606 GN=ESYT2 PE=1 SV=1 | 2 |
| TBC1D14 | TBC1 domain family member 14 OS=Homo sapiens OX=9606 GN=TBC1D14 PE=1 SV=3 | 4 |
| MEA1 | Male-enhanced antigen 1 OS=Homo sapiens OX=9606 GN=MEA1 PE=1 SV=2 | 2 |
| AGK | Acylglycerol kinase, mitochondrial OS=Homo sapiens OX=9606 GN=AGK PE=1 SV=2 | 9 |
| KIF23 | Kinesin-like protein KIF23 OS=Homo sapiens OX=9606 GN=KIF23 PE=1 SV=3 | 9 |
| ARL2 | ADP-ribosylation factor-like protein 2 OS=Homo sapiens OX=9606 GN=ARL2 PE=1 SV=4 | 13 |
| LDLRAP1 | Low density lipoprotein receptor adapter protein 1 OS=Homo sapiens OX=9606 GN=LDLRAP1 PE=1 SV=3 | 8 |
| VPS13A | Intermembrane lipid transfer protein VPS13A OS=Homo sapiens OX=9606 GN=VPS13A PE=1 SV=2 | 6 |
| PPM1H | Protein phosphatase 1H OS=Homo sapiens OX=9606 GN=PPM1H PE=1 SV=2 | 6 |
| MRPS36 | Alpha-ketoglutarate dehydrogenase component 4 OS=Homo sapiens OX=9606 GN=MRPS36 PE=1 SV=2 | 6 |
| SAE1 | SUMO-activating enzyme subunit 1 OS=Homo sapiens OX=9606 GN=SAE1 PE=1 SV=1 | 3 |
| FKBP1C | Peptidyl-prolyl cis-trans isomerase FKBP1C OS=Homo sapiens OX=9606 GN=FKBP1C PE=3 SV=1 | 1 |
| RAB3D | Ras-related protein Rab-3D OS=Homo sapiens OX=9606 GN=RAB3D PE=1 SV=1 | 7 |
| HSDL2 | Hydroxysteroid dehydrogenase-like protein 2 OS=Homo sapiens OX=9606 GN=HSDL2 PE=1 SV=1 | 7 |
| GCDH | Glutaryl-CoA dehydrogenase, mitochondrial OS=Homo sapiens OX=9606 GN=GCDH PE=1 SV=1 | 9 |
| DDX53 | Probable ATP-dependent RNA helicase DDX53 OS=Homo sapiens OX=9606 GN=DDX53 PE=1 SV=3 | 1 |
| AKT2 | RAC-beta serine/threonine-protein kinase OS=Homo sapiens OX=9606 GN=AKT2 PE=1 SV=2 | 15 |
| SPG21 | Maspardin OS=Homo sapiens OX=9606 GN=SPG21 PE=1 SV=1 | 3 |
| ALDH7A1 | Alpha-aminoadipic semialdehyde dehydrogenase OS=Homo sapiens OX=9606 GN=ALDH7A1 PE=1 SV=5 | 31 |
| SEC24B | Protein transport protein Sec24B OS=Homo sapiens OX=9606 GN=SEC24B PE=1 SV=2 | 1 |
| PTGR1 | Prostaglandin reductase 1 OS=Homo sapiens OX=9606 GN=PTGR1 PE=1 SV=2 | 17 |
| TEX264 | Testis-expressed protein 264 OS=Homo sapiens OX=9606 GN=TEX264 PE=1 SV=1 | 3 |
| NDUFS5 | NADH dehydrogenase [ubiquinone] iron-sulfur protein 5 OS=Homo sapiens OX=9606 GN=NDUFS5 PE=1 SV=3 | 4 |
| UBE2D3 | Ubiquitin-conjugating enzyme E2 D3 OS=Homo sapiens OX=9606 GN=UBE2D3 PE=1 SV=1 | 5 |
| VPS26B | Vacuolar protein sorting-associated protein 26B OS=Homo sapiens OX=9606 GN=VPS26B PE=1 SV=2 | 7 |
| TDP1 | Tyrosyl-DNA phosphodiesterase 1 OS=Homo sapiens OX=9606 GN=TDP1 PE=1 SV=2 | 3 |
| CCDC201 | Coiled-coil domain-containing protein 201 OS=Homo sapiens OX=9606 GN=CCDC201 PE=4 SV=1 | 1 |
| COPA | Coatomer subunit alpha OS=Homo sapiens OX=9606 GN=COPA PE=1 SV=2 | 30 |
| PDXP | Chronophin OS=Homo sapiens OX=9606 GN=PDXP PE=1 SV=2 | 8 |
| NAPG | Gamma-soluble NSF attachment protein OS=Homo sapiens OX=9606 GN=NAPG PE=1 SV=1 | 17 |
| CIAO2B | Cytosolic iron-sulfur assembly component 2B OS=Homo sapiens OX=9606 GN=CIAO2B PE=1 SV=1 | 3 |
| ARMC1 | Armadillo repeat-containing protein 1 OS=Homo sapiens OX=9606 GN=ARMC1 PE=1 SV=1 | 5 |
| BCL2L13 | Bcl-2-like protein 13 OS=Homo sapiens OX=9606 GN=BCL2L13 PE=1 SV=1 | 5 |
| NLRP5 | NACHT, LRR and PYD domains-containing protein 5 OS=Homo sapiens OX=9606 GN=NLRP5 PE=1 SV=2 | 71 |
| ZFAND2A | AN1-type zinc finger protein 2A OS=Homo sapiens OX=9606 GN=ZFAND2A PE=1 SV=2 | 1 |
| SAMM50 | Sorting and assembly machinery component 50 homolog OS=Homo sapiens OX=9606 GN=SAMM50 PE=1 SV=3 | 16 |
| JMJD7 | Bifunctional peptidase and (3S)-lysyl hydroxylase JMJD7 OS=Homo sapiens OX=9606 GN=JMJD7 PE=1 SV=1 | 11 |
| NUDT3 | Diphosphoinositol polyphosphate phosphohydrolase 1 OS=Homo sapiens OX=9606 GN=NUDT3 PE=1 SV=1 | 3 |
| AP3M1 | AP-3 complex subunit mu-1 OS=Homo sapiens OX=9606 GN=AP3M1 PE=1 SV=1 | 9 |
| NT5DC1 | 5'-nucleotidase domain-containing protein 1 OS=Homo sapiens OX=9606 GN=NT5DC1 PE=1 SV=1 | 3 |
| RBM8A | RNA-binding protein 8A OS=Homo sapiens OX=9606 GN=RBM8A PE=1 SV=1 | 3 |
| SLC38A10 | Putative sodium-coupled neutral amino acid transporter 10 OS=Homo sapiens OX=9606 GN=SLC38A10 PE=1 SV=2 | 3 |
| SKP1 | S-phase kinase-associated protein 1 OS=Homo sapiens OX=9606 GN=SKP1 PE=1 SV=2 | 10 |
| CPNE1 | Copine-1 OS=Homo sapiens OX=9606 GN=CPNE1 PE=1 SV=1 | 10 |
| AP1S1 | AP-1 complex subunit sigma-1A OS=Homo sapiens OX=9606 GN=AP1S1 PE=1 SV=1 | 4 |
| DNAJC8 | DnaJ homolog subfamily C member 8 OS=Homo sapiens OX=9606 GN=DNAJC8 PE=1 SV=2 | 1 |
| EPS15 | Epidermal growth factor receptor substrate 15 OS=Homo sapiens OX=9606 GN=EPS15 PE=1 SV=2 | 3 |
| TRAPPC8 | Trafficking protein particle complex subunit 8 OS=Homo sapiens OX=9606 GN=TRAPPC8 PE=1 SV=2 | 7 |
| RPL36A | 60S ribosomal protein L36a OS=Homo sapiens OX=9606 GN=RPL36A PE=1 SV=2 | 2 |
| SLC25A12 | Electrogenic aspartate/glutamate antiporter SLC25A12, mitochondrial OS=Homo sapiens OX=9606 GN=SLC25A12 PE=1 SV=2 | 21 |
| FAM13A | Protein FAM13A OS=Homo sapiens OX=9606 GN=FAM13A PE=1 SV=2 | 19 |
| PACS1 | Phosphofurin acidic cluster sorting protein 1 OS=Homo sapiens OX=9606 GN=PACS1 PE=1 SV=2 | 4 |
| P4HB | Protein disulfide-isomerase OS=Homo sapiens OX=9606 GN=P4HB PE=1 SV=3 | 36 |
| SPG7 | Paraplegin OS=Homo sapiens OX=9606 GN=SPG7 PE=1 SV=2 | 3 |
| CRYZL1 | Quinone oxidoreductase-like protein 1 OS=Homo sapiens OX=9606 GN=CRYZL1 PE=1 SV=2 | 2 |
| STK38 | Serine/threonine-protein kinase 38 OS=Homo sapiens OX=9606 GN=STK38 PE=1 SV=1 | 5 |
| TNPO1 | Transportin-1 OS=Homo sapiens OX=9606 GN=TNPO1 PE=1 SV=2 | 8 |
| RPS25 | 40S ribosomal protein S25 OS=Homo sapiens OX=9606 GN=RPS25 PE=1 SV=1 | 3 |
| SHROOM2 | Protein Shroom2 OS=Homo sapiens OX=9606 GN=SHROOM2 PE=1 SV=1 | 4 |
| GPX1 | Glutathione peroxidase 1 OS=Homo sapiens OX=9606 GN=GPX1 PE=1 SV=4 | 14 |
| RAC1 | Ras-related C3 botulinum toxin substrate 1 OS=Homo sapiens OX=9606 GN=RAC1 PE=1 SV=1 | 6 |
| SDHD | Succinate dehydrogenase [ubiquinone] cytochrome b small subunit, mitochondrial OS=Homo sapiens OX=9606 GN=SDHD PE=1 SV=1 | 1 |
| CHGA | Chromogranin-A OS=Homo sapiens OX=9606 GN=CHGA PE=1 SV=7 | 10 |
| TIPRL | TIP41-like protein OS=Homo sapiens OX=9606 GN=TIPRL PE=1 SV=2 | 10 |
| MOB2 | MOB kinase activator 2 OS=Homo sapiens OX=9606 GN=MOB2 PE=1 SV=1 | 6 |
| NAA15 | N-alpha-acetyltransferase 15, NatA auxiliary subunit OS=Homo sapiens OX=9606 GN=NAA15 PE=1 SV=1 | 2 |
| FMNL2 | Formin-like protein 2 OS=Homo sapiens OX=9606 GN=FMNL2 PE=1 SV=3 | 2 |
| GFER | FAD-linked sulfhydryl oxidase ALR OS=Homo sapiens OX=9606 GN=GFER PE=1 SV=2 | 5 |
| UHRF2 | E3 ubiquitin-protein ligase UHRF2 OS=Homo sapiens OX=9606 GN=UHRF2 PE=1 SV=1 | 21 |
| MLYCD | Malonyl-CoA decarboxylase, mitochondrial OS=Homo sapiens OX=9606 GN=MLYCD PE=1 SV=3 | 10 |
| ALDH6A1 | Methylmalonate-semialdehyde dehydrogenase [acylating], mitochondrial OS=Homo sapiens OX=9606 GN=ALDH6A1 PE=1 SV=2 | 26 |
| FBXO30 | F-box only protein 30 OS=Homo sapiens OX=9606 GN=FBXO30 PE=1 SV=3 | 6 |
| ATP5F1C | ATP synthase subunit gamma, mitochondrial OS=Homo sapiens OX=9606 GN=ATP5F1C PE=1 SV=1 | 11 |
| PAIP1 | Polyadenylate-binding protein-interacting protein 1 OS=Homo sapiens OX=9606 GN=PAIP1 PE=1 SV=1 | 7 |
| ZDHHC3 | Palmitoyltransferase ZDHHC3 OS=Homo sapiens OX=9606 GN=ZDHHC3 PE=1 SV=2 | 2 |
| NSMCE1 | Non-structural maintenance of chromosomes element 1 homolog OS=Homo sapiens OX=9606 GN=NSMCE1 PE=1 SV=5 | 1 |
| PGRMC2 | Membrane-associated progesterone receptor component 2 OS=Homo sapiens OX=9606 GN=PGRMC2 PE=1 SV=1 | 5 |
| RPL14 | 60S ribosomal protein L14 OS=Homo sapiens OX=9606 GN=RPL14 PE=1 SV=4 | 5 |
| FLOT1 | Flotillin-1 OS=Homo sapiens OX=9606 GN=FLOT1 PE=1 SV=3 | 4 |
| NLRP9 | NACHT, LRR and PYD domains-containing protein 9 OS=Homo sapiens OX=9606 GN=NLRP9 PE=1 SV=1 | 46 |
| CHCHD6 | MICOS complex subunit MIC25 OS=Homo sapiens OX=9606 GN=CHCHD6 PE=1 SV=1 | 9 |
| DCAF8 | DDB1- and CUL4-associated factor 8 OS=Homo sapiens OX=9606 GN=DCAF8 PE=1 SV=1 | 1 |
| EI24 | Etoposide-induced protein 2.4 homolog OS=Homo sapiens OX=9606 GN=EI24 PE=1 SV=4 | 1 |
| ECH1 | Delta(3,5)-Delta(2,4)-dienoyl-CoA isomerase, mitochondrial OS=Homo sapiens OX=9606 GN=ECH1 PE=1 SV=2 | 17 |
| CASP6 | Caspase-6 OS=Homo sapiens OX=9606 GN=CASP6 PE=1 SV=2 | 3 |
| FN3K | Fructosamine-3-kinase OS=Homo sapiens OX=9606 GN=FN3K PE=1 SV=1 | 3 |
| UBE2O | (E3-independent) E2 ubiquitin-conjugating enzyme OS=Homo sapiens OX=9606 GN=UBE2O PE=1 SV=3 | 17 |
| HADHB | Trifunctional enzyme subunit beta, mitochondrial OS=Homo sapiens OX=9606 GN=HADHB PE=1 SV=3 | 26 |
| NSDHL | Sterol-4-alpha-carboxylate 3-dehydrogenase, decarboxylating OS=Homo sapiens OX=9606 GN=NSDHL PE=1 SV=2 | 9 |
| PEX14 | Peroxisomal membrane protein PEX14 OS=Homo sapiens OX=9606 GN=PEX14 PE=1 SV=1 | 3 |
| KIF20A | Kinesin-like protein KIF20A OS=Homo sapiens OX=9606 GN=KIF20A PE=1 SV=1 | 3 |
| ISYNA1 | Inositol-3-phosphate synthase 1 OS=Homo sapiens OX=9606 GN=ISYNA1 PE=1 SV=1 | 8 |
| KPNA6 | Importin subunit alpha-7 OS=Homo sapiens OX=9606 GN=KPNA6 PE=1 SV=1 | 16 |
| MARS1 | Methionine--tRNA ligase, cytoplasmic OS=Homo sapiens OX=9606 GN=MARS1 PE=1 SV=2 | 11 |
| BROX | BRO1 domain-containing protein BROX OS=Homo sapiens OX=9606 GN=BROX PE=1 SV=1 | 1 |
| OSBPL10 | Oxysterol-binding protein-related protein 10 OS=Homo sapiens OX=9606 GN=OSBPL10 PE=1 SV=2 | 5 |
| ABRAXAS2 | BRISC complex subunit Abraxas 2 OS=Homo sapiens OX=9606 GN=ABRAXAS2 PE=1 SV=2 | 4 |
| ITGA9 | Integrin alpha-9 OS=Homo sapiens OX=9606 GN=ITGA9 PE=1 SV=2 | 12 |
| MRPL38 | 39S ribosomal protein L38, mitochondrial OS=Homo sapiens OX=9606 GN=MRPL38 PE=1 SV=2 | 4 |
| PHYHD1 | Phytanoyl-CoA dioxygenase domain-containing protein 1 OS=Homo sapiens OX=9606 GN=PHYHD1 PE=1 SV=2 | 5 |
| GRB2 | Growth factor receptor-bound protein 2 OS=Homo sapiens OX=9606 GN=GRB2 PE=1 SV=1 | 7 |
| RHOT2 | Mitochondrial Rho GTPase 2 OS=Homo sapiens OX=9606 GN=RHOT2 PE=1 SV=2 | 4 |
| PHACTR3 | Phosphatase and actin regulator 3 OS=Homo sapiens OX=9606 GN=PHACTR3 PE=1 SV=1 | 7 |
| TAMM41 | Phosphatidate cytidylyltransferase, mitochondrial OS=Homo sapiens OX=9606 GN=TAMM41 PE=1 SV=2 | 4 |
| UBA3 | NEDD8-activating enzyme E1 catalytic subunit OS=Homo sapiens OX=9606 GN=UBA3 PE=1 SV=2 | 6 |
| DNAJA3 | DnaJ homolog subfamily A member 3, mitochondrial OS=Homo sapiens OX=9606 GN=DNAJA3 PE=1 SV=2 | 13 |
| UBE2Z | Ubiquitin-conjugating enzyme E2 Z OS=Homo sapiens OX=9606 GN=UBE2Z PE=1 SV=2 | 13 |
| NUP210 | Nuclear pore membrane glycoprotein 210 OS=Homo sapiens OX=9606 GN=NUP210 PE=1 SV=3 | 4 |
| CNN3 | Calponin-3 OS=Homo sapiens OX=9606 GN=CNN3 PE=1 SV=1 | 10 |
| NDEL1 | Nuclear distribution protein nudE-like 1 OS=Homo sapiens OX=9606 GN=NDEL1 PE=1 SV=1 | 1 |
| GIPC2 | PDZ domain-containing protein GIPC2 OS=Homo sapiens OX=9606 GN=GIPC2 PE=1 SV=1 | 13 |
| CISD2 | CDGSH iron-sulfur domain-containing protein 2 OS=Homo sapiens OX=9606 GN=CISD2 PE=1 SV=1 | 3 |
| TRIM36 | E3 ubiquitin-protein ligase TRIM36 OS=Homo sapiens OX=9606 GN=TRIM36 PE=1 SV=2 | 6 |
| BABAM1 | BRISC and BRCA1-A complex member 1 OS=Homo sapiens OX=9606 GN=BABAM1 PE=1 SV=1 | 2 |
| TMEM160 | Transmembrane protein 160 OS=Homo sapiens OX=9606 GN=TMEM160 PE=1 SV=1 | 1 |
| KLC1 | Kinesin light chain 1 OS=Homo sapiens OX=9606 GN=KLC1 PE=1 SV=2 | 19 |
| GAK | Cyclin-G-associated kinase OS=Homo sapiens OX=9606 GN=GAK PE=1 SV=2 | 4 |
| DCXR | L-xylulose reductase OS=Homo sapiens OX=9606 GN=DCXR PE=1 SV=2 | 15 |
| ACOT2 | Acyl-coenzyme A thioesterase 2, mitochondrial OS=Homo sapiens OX=9606 GN=ACOT2 PE=1 SV=6 | 10 |
| SNX12 | Sorting nexin-12 OS=Homo sapiens OX=9606 GN=SNX12 PE=1 SV=4 | 13 |
| BUB1B | Mitotic checkpoint serine/threonine-protein kinase BUB1 beta OS=Homo sapiens OX=9606 GN=BUB1B PE=1 SV=3 | 35 |
| CALM1 | Calmodulin-1 OS=Homo sapiens OX=9606 GN=CALM1 PE=1 SV=1 | 6 |
| CAND1 | Cullin-associated NEDD8-dissociated protein 1 OS=Homo sapiens OX=9606 GN=CAND1 PE=1 SV=2 | 40 |
| FKBP1A | Peptidyl-prolyl cis-trans isomerase FKBP1A OS=Homo sapiens OX=9606 GN=FKBP1A PE=1 SV=2 | 5 |
| PPP2CA | Serine/threonine-protein phosphatase 2A catalytic subunit alpha isoform OS=Homo sapiens OX=9606 GN=PPP2CA PE=1 SV=1 | 9 |
| SERPINF2 | Alpha-2-antiplasmin OS=Homo sapiens OX=9606 GN=SERPINF2 PE=1 SV=3 | 4 |
| SCAMP3 | Secretory carrier-associated membrane protein 3 OS=Homo sapiens OX=9606 GN=SCAMP3 PE=1 SV=3 | 2 |
| LMAN2L | VIP36-like protein OS=Homo sapiens OX=9606 GN=LMAN2L PE=1 SV=1 | 6 |
| IPO5 | Importin-5 OS=Homo sapiens OX=9606 GN=IPO5 PE=1 SV=4 | 21 |
| LAMC1 | Laminin subunit gamma-1 OS=Homo sapiens OX=9606 GN=LAMC1 PE=1 SV=3 | 23 |
| TANC2 | Protein TANC2 OS=Homo sapiens OX=9606 GN=TANC2 PE=1 SV=3 | 1 |
| EXOC4 | Exocyst complex component 4 OS=Homo sapiens OX=9606 GN=EXOC4 PE=1 SV=1 | 8 |
| UCHL1 | Ubiquitin carboxyl-terminal hydrolase isozyme L1 OS=Homo sapiens OX=9606 GN=UCHL1 PE=1 SV=2 | 21 |
| MPDU1 | Mannose-P-dolichol utilization defect 1 protein OS=Homo sapiens OX=9606 GN=MPDU1 PE=1 SV=2 | 3 |
| OARD1 | ADP-ribose glycohydrolase OARD1 OS=Homo sapiens OX=9606 GN=OARD1 PE=1 SV=2 | 6 |
| SEC61B | Protein transport protein Sec61 subunit beta OS=Homo sapiens OX=9606 GN=SEC61B PE=1 SV=2 | 2 |
| MAT1A | S-adenosylmethionine synthase isoform type-1 OS=Homo sapiens OX=9606 GN=MAT1A PE=1 SV=2 | 10 |
| MT-CO1 | Cytochrome c oxidase subunit 1 OS=Homo sapiens OX=9606 GN=MT-CO1 PE=1 SV=1 | 1 |
| PTMA | Prothymosin alpha OS=Homo sapiens OX=9606 GN=PTMA PE=1 SV=2 | 2 |
| UBE2N | Ubiquitin-conjugating enzyme E2 N OS=Homo sapiens OX=9606 GN=UBE2N PE=1 SV=1 | 9 |
| BANF1 | Barrier-to-autointegration factor OS=Homo sapiens OX=9606 GN=BANF1 PE=1 SV=1 | 7 |
| POR | NADPH--cytochrome P450 reductase OS=Homo sapiens OX=9606 GN=POR PE=1 SV=2 | 24 |
| TDRD7 | Tudor domain-containing protein 7 OS=Homo sapiens OX=9606 GN=TDRD7 PE=1 SV=2 | 5 |
| ANXA6 | Annexin A6 OS=Homo sapiens OX=9606 GN=ANXA6 PE=1 SV=3 | 42 |
| PSME1 | Proteasome activator complex subunit 1 OS=Homo sapiens OX=9606 GN=PSME1 PE=1 SV=1 | 5 |
| DNM1 | Dynamin-1 OS=Homo sapiens OX=9606 GN=DNM1 PE=1 SV=2 | 1 |
| ZFYVE19 | Abscission/NoCut checkpoint regulator OS=Homo sapiens OX=9606 GN=ZFYVE19 PE=1 SV=3 | 3 |
| CYRIB | CYFIP-related Rac1 interactor B OS=Homo sapiens OX=9606 GN=CYRIB PE=1 SV=1 | 13 |
| CCDC93 | Coiled-coil domain-containing protein 93 OS=Homo sapiens OX=9606 GN=CCDC93 PE=1 SV=2 | 5 |
| CANX | Calnexin OS=Homo sapiens OX=9606 GN=CANX PE=1 SV=2 | 20 |
| ARMC9 | LisH domain-containing protein ARMC9 OS=Homo sapiens OX=9606 GN=ARMC9 PE=1 SV=3 | 10 |
| TUBB8 | Tubulin beta-8 chain OS=Homo sapiens OX=9606 GN=TUBB8 PE=1 SV=2 | 30 |
| OPALIN | Opalin OS=Homo sapiens OX=9606 GN=OPALIN PE=2 SV=1 | 3 |
| GSTK1 | Glutathione S-transferase kappa 1 OS=Homo sapiens OX=9606 GN=GSTK1 PE=1 SV=3 | 10 |
| FHIT | Bis(5'-adenosyl)-triphosphatase OS=Homo sapiens OX=9606 GN=FHIT PE=1 SV=3 | 11 |
| TMEM109 | Transmembrane protein 109 OS=Homo sapiens OX=9606 GN=TMEM109 PE=1 SV=1 | 2 |
| ANKFY1 | Rabankyrin-5 OS=Homo sapiens OX=9606 GN=ANKFY1 PE=1 SV=2 | 4 |
| FUCA1 | Tissue alpha-L-fucosidase OS=Homo sapiens OX=9606 GN=FUCA1 PE=1 SV=4 | 7 |
| MCM4 | DNA replication licensing factor MCM4 OS=Homo sapiens OX=9606 GN=MCM4 PE=1 SV=5 | 3 |
| PARP16 | Protein mono-ADP-ribosyltransferase PARP16 OS=Homo sapiens OX=9606 GN=PARP16 PE=1 SV=2 | 8 |
| NIPSNAP1 | Protein NipSnap homolog 1 OS=Homo sapiens OX=9606 GN=NIPSNAP1 PE=1 SV=1 | 13 |
| DFFA | DNA fragmentation factor subunit alpha OS=Homo sapiens OX=9606 GN=DFFA PE=1 SV=1 | 10 |
| UFM1 | Ubiquitin-fold modifier 1 OS=Homo sapiens OX=9606 GN=UFM1 PE=1 SV=1 | 5 |
| STAMBP | STAM-binding protein OS=Homo sapiens OX=9606 GN=STAMBP PE=1 SV=1 | 12 |
| RAB4A | Ras-related protein Rab-4A OS=Homo sapiens OX=9606 GN=RAB4A PE=1 SV=3 | 5 |
| SQOR | Sulfide:quinone oxidoreductase, mitochondrial OS=Homo sapiens OX=9606 GN=SQOR PE=1 SV=1 | 22 |
| BPGM | Bisphosphoglycerate mutase OS=Homo sapiens OX=9606 GN=BPGM PE=1 SV=2 | 24 |
| SRSF1 | Serine/arginine-rich splicing factor 1 OS=Homo sapiens OX=9606 GN=SRSF1 PE=1 SV=2 | 5 |
| LZIC | Protein LZIC OS=Homo sapiens OX=9606 GN=LZIC PE=1 SV=1 | 10 |
| ALG12 | Dol-P-Man:Man(7)GlcNAc(2)-PP-Dol alpha-1,6-mannosyltransferase OS=Homo sapiens OX=9606 GN=ALG12 PE=1 SV=1 | 1 |
| PPP2R1A | Serine/threonine-protein phosphatase 2A 65 kDa regulatory subunit A alpha isoform OS=Homo sapiens OX=9606 GN=PPP2R1A PE=1 SV=4 | 21 |
| KDM1B | Lysine-specific histone demethylase 2 OS=Homo sapiens OX=9606 GN=KDM1B PE=1 SV=3 | 27 |
| ARHGDIA | Rho GDP-dissociation inhibitor 1 OS=Homo sapiens OX=9606 GN=ARHGDIA PE=1 SV=3 | 12 |
| VDAC3 | Voltage-dependent anion-selective channel protein 3 OS=Homo sapiens OX=9606 GN=VDAC3 PE=1 SV=1 | 13 |
| CYC1 | Cytochrome c1, heme protein, mitochondrial OS=Homo sapiens OX=9606 GN=CYC1 PE=1 SV=3 | 7 |
| AUP1 | Lipid droplet-regulating VLDL assembly factor AUP1 OS=Homo sapiens OX=9606 GN=AUP1 PE=1 SV=2 | 4 |
| ATOX1 | Copper transport protein ATOX1 OS=Homo sapiens OX=9606 GN=ATOX1 PE=1 SV=1 | 5 |
| BRD2 | Bromodomain-containing protein 2 OS=Homo sapiens OX=9606 GN=BRD2 PE=1 SV=2 | 2 |
| DIP2B | Disco-interacting protein 2 homolog B OS=Homo sapiens OX=9606 GN=DIP2B PE=1 SV=3 | 7 |
| WASHC3 | WASH complex subunit 3 OS=Homo sapiens OX=9606 GN=WASHC3 PE=1 SV=1 | 2 |
| MB21D2 | Nucleotidyltransferase MB21D2 OS=Homo sapiens OX=9606 GN=MB21D2 PE=1 SV=3 | 10 |
| CLCN5 | H(+)/Cl(-) exchange transporter 5 OS=Homo sapiens OX=9606 GN=CLCN5 PE=1 SV=2 | 1 |
| TMED1 | Transmembrane emp24 domain-containing protein 1 OS=Homo sapiens OX=9606 GN=TMED1 PE=1 SV=1 | 2 |
| ENPP2 | Ectonucleotide pyrophosphatase/phosphodiesterase family member 2 OS=Homo sapiens OX=9606 GN=ENPP2 PE=1 SV=3 | 1 |
| YES1 | Tyrosine-protein kinase Yes OS=Homo sapiens OX=9606 GN=YES1 PE=1 SV=3 | 6 |
| PSMG1 | Proteasome assembly chaperone 1 OS=Homo sapiens OX=9606 GN=PSMG1 PE=1 SV=1 | 2 |
| PLIN3 | Perilipin-3 OS=Homo sapiens OX=9606 GN=PLIN3 PE=1 SV=3 | 24 |
| SHTN1 | Shootin-1 OS=Homo sapiens OX=9606 GN=SHTN1 PE=1 SV=4 | 22 |
| RAB22A | Ras-related protein Rab-22A OS=Homo sapiens OX=9606 GN=RAB22A PE=1 SV=2 | 10 |
| PIK3CA | Phosphatidylinositol 4,5-bisphosphate 3-kinase catalytic subunit alpha isoform OS=Homo sapiens OX=9606 GN=PIK3CA PE=1 SV=2 | 2 |
| TRAP1 | Heat shock protein 75 kDa, mitochondrial OS=Homo sapiens OX=9606 GN=TRAP1 PE=1 SV=3 | 44 |
| TXNDC17 | Thioredoxin domain-containing protein 17 OS=Homo sapiens OX=9606 GN=TXNDC17 PE=1 SV=1 | 11 |
| PSMD5 | 26S proteasome non-ATPase regulatory subunit 5 OS=Homo sapiens OX=9606 GN=PSMD5 PE=1 SV=3 | 9 |
| LARP1 | La-related protein 1 OS=Homo sapiens OX=9606 GN=LARP1 PE=1 SV=2 | 13 |
| ARSA | Arylsulfatase A OS=Homo sapiens OX=9606 GN=ARSA PE=1 SV=3 | 2 |
| UBE2T | Ubiquitin-conjugating enzyme E2 T OS=Homo sapiens OX=9606 GN=UBE2T PE=1 SV=1 | 5 |
| ATP6V1G1 | V-type proton ATPase subunit G 1 OS=Homo sapiens OX=9606 GN=ATP6V1G1 PE=1 SV=3 | 3 |
| WASHC4 | WASH complex subunit 4 OS=Homo sapiens OX=9606 GN=WASHC4 PE=1 SV=2 | 6 |
| RIOX2 | Ribosomal oxygenase 2 OS=Homo sapiens OX=9606 GN=RIOX2 PE=1 SV=1 | 7 |
| MOB3C | MOB kinase activator 3C OS=Homo sapiens OX=9606 GN=MOB3C PE=1 SV=1 | 1 |
| AIP | AH receptor-interacting protein OS=Homo sapiens OX=9606 GN=AIP PE=1 SV=3 | 13 |
| ALPK3 | Alpha-protein kinase 3 OS=Homo sapiens OX=9606 GN=ALPK3 PE=1 SV=2 | 2 |
| EEF1G | Elongation factor 1-gamma OS=Homo sapiens OX=9606 GN=EEF1G PE=1 SV=3 | 19 |
| NPM2 | Nucleoplasmin-2 OS=Homo sapiens OX=9606 GN=NPM2 PE=1 SV=1 | 8 |
| RAB21 | Ras-related protein Rab-21 OS=Homo sapiens OX=9606 GN=RAB21 PE=1 SV=3 | 7 |
| ATP1B1 | Sodium/potassium-transporting ATPase subunit beta-1 OS=Homo sapiens OX=9606 GN=ATP1B1 PE=1 SV=1 | 8 |
| TUBGCP3 | Gamma-tubulin complex component 3 OS=Homo sapiens OX=9606 GN=TUBGCP3 PE=1 SV=2 | 8 |
| UBXN1 | UBX domain-containing protein 1 OS=Homo sapiens OX=9606 GN=UBXN1 PE=1 SV=2 | 3 |
| LNPK | Endoplasmic reticulum junction formation protein lunapark OS=Homo sapiens OX=9606 GN=LNPK PE=1 SV=2 | 11 |
| TTC1 | Tetratricopeptide repeat protein 1 OS=Homo sapiens OX=9606 GN=TTC1 PE=1 SV=1 | 6 |
| PHAX | Phosphorylated adapter RNA export protein OS=Homo sapiens OX=9606 GN=PHAX PE=1 SV=1 | 1 |
| PRPS1 | Ribose-phosphate pyrophosphokinase 1 OS=Homo sapiens OX=9606 GN=PRPS1 PE=1 SV=2 | 14 |
| PPP1CA | Serine/threonine-protein phosphatase PP1-alpha catalytic subunit OS=Homo sapiens OX=9606 GN=PPP1CA PE=1 SV=1 | 11 |
| NDUFA7 | NADH dehydrogenase [ubiquinone] 1 alpha subcomplex subunit 7 OS=Homo sapiens OX=9606 GN=NDUFA7 PE=1 SV=3 | 5 |
| RBBP7 | Histone-binding protein RBBP7 OS=Homo sapiens OX=9606 GN=RBBP7 PE=1 SV=1 | 4 |
| HRG | Histidine-rich glycoprotein OS=Homo sapiens OX=9606 GN=HRG PE=1 SV=1 | 5 |
| GPD1 | Glycerol-3-phosphate dehydrogenase [NAD(+)], cytoplasmic OS=Homo sapiens OX=9606 GN=GPD1 PE=1 SV=4 | 24 |
| PREPL | Prolyl endopeptidase-like OS=Homo sapiens OX=9606 GN=PREPL PE=1 SV=1 | 12 |
| TXNDC12 | Thioredoxin domain-containing protein 12 OS=Homo sapiens OX=9606 GN=TXNDC12 PE=1 SV=1 | 5 |
| GNAI3 | Guanine nucleotide-binding protein G(i) subunit alpha-3 OS=Homo sapiens OX=9606 GN=GNAI3 PE=1 SV=3 | 12 |
| ERMP1 | Endoplasmic reticulum metallopeptidase 1 OS=Homo sapiens OX=9606 GN=ERMP1 PE=1 SV=2 | 16 |
| DNAJC3 | DnaJ homolog subfamily C member 3 OS=Homo sapiens OX=9606 GN=DNAJC3 PE=1 SV=1 | 27 |
| LIN7C | Protein lin-7 homolog C OS=Homo sapiens OX=9606 GN=LIN7C PE=1 SV=1 | 3 |
| EXOC1 | Exocyst complex component 1 OS=Homo sapiens OX=9606 GN=EXOC1 PE=1 SV=4 | 6 |
| HPRT1 | Hypoxanthine-guanine phosphoribosyltransferase OS=Homo sapiens OX=9606 GN=HPRT1 PE=1 SV=2 | 11 |
| NFX1 | Transcriptional repressor NF-X1 OS=Homo sapiens OX=9606 GN=NFX1 PE=1 SV=2 | 2 |
| ATP6V0A1 | V-type proton ATPase 116 kDa subunit a 1 OS=Homo sapiens OX=9606 GN=ATP6V0A1 PE=1 SV=3 | 18 |
| STAM2 | Signal transducing adapter molecule 2 OS=Homo sapiens OX=9606 GN=STAM2 PE=1 SV=1 | 11 |
| IPO7 | Importin-7 OS=Homo sapiens OX=9606 GN=IPO7 PE=1 SV=1 | 9 |
| LIN28A | Protein lin-28 homolog A OS=Homo sapiens OX=9606 GN=LIN28A PE=1 SV=1 | 2 |
| PGLS | 6-phosphogluconolactonase OS=Homo sapiens OX=9606 GN=PGLS PE=1 SV=2 | 12 |
| DHRS7B | Dehydrogenase/reductase SDR family member 7B OS=Homo sapiens OX=9606 GN=DHRS7B PE=1 SV=2 | 5 |
| CHORDC1 | Cysteine and histidine-rich domain-containing protein 1 OS=Homo sapiens OX=9606 GN=CHORDC1 PE=1 SV=2 | 10 |
| DUSP3 | Dual specificity protein phosphatase 3 OS=Homo sapiens OX=9606 GN=DUSP3 PE=1 SV=1 | 5 |
| LIPT2 | Putative lipoyltransferase 2, mitochondrial OS=Homo sapiens OX=9606 GN=LIPT2 PE=1 SV=1 | 3 |
| LAMB1 | Laminin subunit beta-1 OS=Homo sapiens OX=9606 GN=LAMB1 PE=1 SV=2 | 34 |
| PGP | Glycerol-3-phosphate phosphatase OS=Homo sapiens OX=9606 GN=PGP PE=1 SV=1 | 4 |
| HIDE1 | Protein HIDE1 OS=Homo sapiens OX=9606 GN=HIDE1 PE=1 SV=2 | 1 |
| UBR7 | Putative E3 ubiquitin-protein ligase UBR7 OS=Homo sapiens OX=9606 GN=UBR7 PE=1 SV=2 | 5 |
| RPS9 | 40S ribosomal protein S9 OS=Homo sapiens OX=9606 GN=RPS9 PE=1 SV=3 | 9 |
| PLAA | Phospholipase A-2-activating protein OS=Homo sapiens OX=9606 GN=PLAA PE=1 SV=2 | 22 |
| NPTN | Neuroplastin OS=Homo sapiens OX=9606 GN=NPTN PE=1 SV=2 | 4 |
| EIF2S3 | Eukaryotic translation initiation factor 2 subunit 3 OS=Homo sapiens OX=9606 GN=EIF2S3 PE=1 SV=3 | 5 |
| SPECC1L | Cytospin-A OS=Homo sapiens OX=9606 GN=SPECC1L PE=1 SV=3 | 13 |
| PPP1R21 | Protein phosphatase 1 regulatory subunit 21 OS=Homo sapiens OX=9606 GN=PPP1R21 PE=1 SV=1 | 1 |
| DNAJC5 | DnaJ homolog subfamily C member 5 OS=Homo sapiens OX=9606 GN=DNAJC5 PE=1 SV=1 | 1 |
| ERGIC3 | Endoplasmic reticulum-Golgi intermediate compartment protein 3 OS=Homo sapiens OX=9606 GN=ERGIC3 PE=1 SV=1 | 5 |
| MMUT | Methylmalonyl-CoA mutase, mitochondrial OS=Homo sapiens OX=9606 GN=MMUT PE=1 SV=4 | 24 |
| NR2C2AP | Nuclear receptor 2C2-associated protein OS=Homo sapiens OX=9606 GN=NR2C2AP PE=1 SV=1 | 3 |
| BSCL2 | Seipin OS=Homo sapiens OX=9606 GN=BSCL2 PE=1 SV=3 | 4 |
| FXN | Frataxin, mitochondrial OS=Homo sapiens OX=9606 GN=FXN PE=1 SV=2 | 7 |
| IDH3A | Isocitrate dehydrogenase [NAD] subunit alpha, mitochondrial OS=Homo sapiens OX=9606 GN=IDH3A PE=1 SV=1 | 17 |
| RAB6B | Ras-related protein Rab-6B OS=Homo sapiens OX=9606 GN=RAB6B PE=1 SV=1 | 16 |
| CHMP6 | Charged multivesicular body protein 6 OS=Homo sapiens OX=9606 GN=CHMP6 PE=1 SV=3 | 3 |
| FAF1 | FAS-associated factor 1 OS=Homo sapiens OX=9606 GN=FAF1 PE=1 SV=2 | 4 |
| RWDD1 | RWD domain-containing protein 1 OS=Homo sapiens OX=9606 GN=RWDD1 PE=1 SV=1 | 2 |
| NFS1 | Cysteine desulfurase OS=Homo sapiens OX=9606 GN=NFS1 PE=1 SV=3 | 18 |
| TUBA4A | Tubulin alpha-4A chain OS=Homo sapiens OX=9606 GN=TUBA4A PE=1 SV=1 | 27 |
| ATG7 | Ubiquitin-like modifier-activating enzyme ATG7 OS=Homo sapiens OX=9606 GN=ATG7 PE=1 SV=1 | 9 |
| B3GLCT | Beta-1,3-glucosyltransferase OS=Homo sapiens OX=9606 GN=B3GLCT PE=1 SV=2 | 8 |
| RDH11 | Retinol dehydrogenase 11 OS=Homo sapiens OX=9606 GN=RDH11 PE=1 SV=2 | 2 |
| RAB11B | Ras-related protein Rab-11B OS=Homo sapiens OX=9606 GN=RAB11B PE=1 SV=4 | 15 |
| PLS1 | Plastin-1 OS=Homo sapiens OX=9606 GN=PLS1 PE=1 SV=2 | 27 |
| SNX9 | Sorting nexin-9 OS=Homo sapiens OX=9606 GN=SNX9 PE=1 SV=1 | 11 |
| RPS13 | 40S ribosomal protein S13 OS=Homo sapiens OX=9606 GN=RPS13 PE=1 SV=2 | 6 |
| PDIA6 | Protein disulfide-isomerase A6 OS=Homo sapiens OX=9606 GN=PDIA6 PE=1 SV=1 | 18 |
| LPIN2 | Phosphatidate phosphatase LPIN2 OS=Homo sapiens OX=9606 GN=LPIN2 PE=1 SV=1 | 5 |
| IDH3B | Isocitrate dehydrogenase [NAD] subunit beta, mitochondrial OS=Homo sapiens OX=9606 GN=IDH3B PE=1 SV=2 | 21 |
| HIBADH | 3-hydroxyisobutyrate dehydrogenase, mitochondrial OS=Homo sapiens OX=9606 GN=HIBADH PE=1 SV=2 | 7 |
| XPO1 | Exportin-1 OS=Homo sapiens OX=9606 GN=XPO1 PE=1 SV=1 | 20 |
| OGA | Protein O-GlcNAcase OS=Homo sapiens OX=9606 GN=OGA PE=1 SV=2 | 3 |
| PICALM | Phosphatidylinositol-binding clathrin assembly protein OS=Homo sapiens OX=9606 GN=PICALM PE=1 SV=2 | 7 |
| OGDH | 2-oxoglutarate dehydrogenase complex component E1 OS=Homo sapiens OX=9606 GN=OGDH PE=1 SV=3 | 54 |
| TMPO | Lamina-associated polypeptide 2, isoform alpha OS=Homo sapiens OX=9606 GN=TMPO PE=1 SV=2 | 5 |
| SNX4 | Sorting nexin-4 OS=Homo sapiens OX=9606 GN=SNX4 PE=1 SV=1 | 19 |
| DNM1L | Dynamin-1-like protein OS=Homo sapiens OX=9606 GN=DNM1L PE=1 SV=2 | 41 |
| LONP1 | Lon protease homolog, mitochondrial OS=Homo sapiens OX=9606 GN=LONP1 PE=1 SV=2 | 26 |
| COPS4 | COP9 signalosome complex subunit 4 OS=Homo sapiens OX=9606 GN=COPS4 PE=1 SV=1 | 30 |
| FBXL15 | F-box/LRR-repeat protein 15 OS=Homo sapiens OX=9606 GN=FBXL15 PE=1 SV=2 | 2 |
| ALS2 | Alsin OS=Homo sapiens OX=9606 GN=ALS2 PE=1 SV=2 | 11 |
| POLG2 | DNA polymerase subunit gamma-2, mitochondrial OS=Homo sapiens OX=9606 GN=POLG2 PE=1 SV=1 | 6 |
| USP14 | Ubiquitin carboxyl-terminal hydrolase 14 OS=Homo sapiens OX=9606 GN=USP14 PE=1 SV=3 | 25 |
| "SMN1 | Survival motor neuron protein OS=Homo sapiens OX=9606 GN=SMN2 PE=1 SV=1 | 3 |
| ACAA1 | 3-ketoacyl-CoA thiolase, peroxisomal OS=Homo sapiens OX=9606 GN=ACAA1 PE=1 SV=2 | 20 |
| PPM1E | Protein phosphatase 1E OS=Homo sapiens OX=9606 GN=PPM1E PE=1 SV=3 | 6 |
| RBM17 | Splicing factor 45 OS=Homo sapiens OX=9606 GN=RBM17 PE=1 SV=1 | 1 |
| COMMD1 | COMM domain-containing protein 1 OS=Homo sapiens OX=9606 GN=COMMD1 PE=1 SV=1 | 8 |
| YWHAQ | 14-3-3 protein theta OS=Homo sapiens OX=9606 GN=YWHAQ PE=1 SV=1 | 12 |
| MLST8 | Target of rapamycin complex subunit LST8 OS=Homo sapiens OX=9606 GN=MLST8 PE=1 SV=1 | 6 |
| CNTNAP3B | Contactin-associated protein-like 3B OS=Homo sapiens OX=9606 GN=CNTNAP3B PE=2 SV=3 | 1 |
| VPS35 | Vacuolar protein sorting-associated protein 35 OS=Homo sapiens OX=9606 GN=VPS35 PE=1 SV=2 | 22 |
| TBC1D7 | TBC1 domain family member 7 OS=Homo sapiens OX=9606 GN=TBC1D7 PE=1 SV=1 | 1 |
| EEF1B2 | Elongation factor 1-beta OS=Homo sapiens OX=9606 GN=EEF1B2 PE=1 SV=3 | 7 |
| COLGALT1 | Procollagen galactosyltransferase 1 OS=Homo sapiens OX=9606 GN=COLGALT1 PE=1 SV=1 | 9 |
| FKBP15 | FK506-binding protein 15 OS=Homo sapiens OX=9606 GN=FKBP15 PE=1 SV=2 | 9 |
| CTTN | Src substrate cortactin OS=Homo sapiens OX=9606 GN=CTTN PE=1 SV=2 | 17 |
| NDUFAF3 | NADH dehydrogenase [ubiquinone] 1 alpha subcomplex assembly factor 3 OS=Homo sapiens OX=9606 GN=NDUFAF3 PE=1 SV=1 | 5 |
| RIMKLA | N-acetylaspartylglutamate synthase A OS=Homo sapiens OX=9606 GN=RIMKLA PE=1 SV=2 | 3 |
| CDC34 | Ubiquitin-conjugating enzyme E2 R1 OS=Homo sapiens OX=9606 GN=CDC34 PE=1 SV=2 | 3 |
| ARFGAP3 | ADP-ribosylation factor GTPase-activating protein 3 OS=Homo sapiens OX=9606 GN=ARFGAP3 PE=1 SV=1 | 8 |
| RPS27 | 40S ribosomal protein S27 OS=Homo sapiens OX=9606 GN=RPS27 PE=1 SV=3 | 1 |
| HMCES | Abasic site processing protein HMCES OS=Homo sapiens OX=9606 GN=HMCES PE=1 SV=1 | 7 |
| GABARAPL2 | Gamma-aminobutyric acid receptor-associated protein-like 2 OS=Homo sapiens OX=9606 GN=GABARAPL2 PE=1 SV=1 | 7 |
| RRM1 | Ribonucleoside-diphosphate reductase large subunit OS=Homo sapiens OX=9606 GN=RRM1 PE=1 SV=1 | 4 |
| VPS33B | Vacuolar protein sorting-associated protein 33B OS=Homo sapiens OX=9606 GN=VPS33B PE=1 SV=2 | 5 |
| IGSF8 | Immunoglobulin superfamily member 8 OS=Homo sapiens OX=9606 GN=IGSF8 PE=1 SV=1 | 2 |
| PTRHD1 | Putative peptidyl-tRNA hydrolase PTRHD1 OS=Homo sapiens OX=9606 GN=PTRHD1 PE=1 SV=1 | 8 |
| GRIPAP1 | GRIP1-associated protein 1 OS=Homo sapiens OX=9606 GN=GRIPAP1 PE=1 SV=2 | 16 |
| FAM98B | Protein FAM98B OS=Homo sapiens OX=9606 GN=FAM98B PE=1 SV=2 | 9 |
| CLPX | ATP-dependent Clp protease ATP-binding subunit clpX-like, mitochondrial OS=Homo sapiens OX=9606 GN=CLPX PE=1 SV=2 | 5 |
| NOMO3 | BOS complex subunit NOMO3 OS=Homo sapiens OX=9606 GN=NOMO3 PE=1 SV=2 | 28 |
| USP9X | Probable ubiquitin carboxyl-terminal hydrolase FAF-X OS=Homo sapiens OX=9606 GN=USP9X PE=1 SV=4 | 56 |
| KPNA1 | Importin subunit alpha-5 OS=Homo sapiens OX=9606 GN=KPNA1 PE=1 SV=3 | 1 |
| SDHAF2 | Succinate dehydrogenase assembly factor 2, mitochondrial OS=Homo sapiens OX=9606 GN=SDHAF2 PE=1 SV=1 | 2 |
| ACOT7 | Cytosolic acyl coenzyme A thioester hydrolase OS=Homo sapiens OX=9606 GN=ACOT7 PE=1 SV=3 | 13 |
| ATAD3C | ATPase family AAA domain-containing protein 3C OS=Homo sapiens OX=9606 GN=ATAD3C PE=2 SV=2 | 18 |
| PLAAT3 | Phospholipase A and acyltransferase 3 OS=Homo sapiens OX=9606 GN=PLAAT3 PE=1 SV=2 | 3 |
| STT3A | Dolichyl-diphosphooligosaccharide--protein glycosyltransferase subunit STT3A OS=Homo sapiens OX=9606 GN=STT3A PE=1 SV=2 | 7 |
| CALR | Calreticulin OS=Homo sapiens OX=9606 GN=CALR PE=1 SV=1 | 17 |
| PHYKPL | 5-phosphohydroxy-L-lysine phospho-lyase OS=Homo sapiens OX=9606 GN=PHYKPL PE=1 SV=1 | 1 |
| MOCS2 | Molybdopterin synthase catalytic subunit OS=Homo sapiens OX=9606 GN=MOCS2 PE=1 SV=1 | 3 |
| TIFA | TRAF-interacting protein with FHA domain-containing protein A OS=Homo sapiens OX=9606 GN=TIFA PE=1 SV=1 | 3 |
| TMEM245 | Transmembrane protein 245 OS=Homo sapiens OX=9606 GN=TMEM245 PE=1 SV=3 | 5 |
| KIF5B | Kinesin-1 heavy chain OS=Homo sapiens OX=9606 GN=KIF5B PE=1 SV=1 | 38 |
| FARSA | Phenylalanine--tRNA ligase alpha subunit OS=Homo sapiens OX=9606 GN=FARSA PE=1 SV=3 | 10 |
| PPM1A | Protein phosphatase 1A OS=Homo sapiens OX=9606 GN=PPM1A PE=1 SV=1 | 3 |
| SQSTM1 | Sequestosome-1 OS=Homo sapiens OX=9606 GN=SQSTM1 PE=1 SV=1 | 1 |
| MFF | Mitochondrial fission factor OS=Homo sapiens OX=9606 GN=MFF PE=1 SV=1 | 6 |
| VPS4A | Vacuolar protein sorting-associated protein 4A OS=Homo sapiens OX=9606 GN=VPS4A PE=1 SV=1 | 15 |
| CNBP | CCHC-type zinc finger nucleic acid binding protein OS=Homo sapiens OX=9606 GN=CNBP PE=1 SV=1 | 3 |
| MCAT | Malonyl-CoA-acyl carrier protein transacylase, mitochondrial OS=Homo sapiens OX=9606 GN=MCAT PE=1 SV=2 | 6 |
| AKR7A2 | Aflatoxin B1 aldehyde reductase member 2 OS=Homo sapiens OX=9606 GN=AKR7A2 PE=1 SV=3 | 14 |
| PTK2 | Focal adhesion kinase 1 OS=Homo sapiens OX=9606 GN=PTK2 PE=1 SV=2 | 18 |
| CSE1L | Exportin-2 OS=Homo sapiens OX=9606 GN=CSE1L PE=1 SV=3 | 26 |
| AK1 | Adenylate kinase isoenzyme 1 OS=Homo sapiens OX=9606 GN=AK1 PE=1 SV=3 | 12 |
| GANAB | Neutral alpha-glucosidase AB OS=Homo sapiens OX=9606 GN=GANAB PE=1 SV=3 | 39 |
| RAN | GTP-binding nuclear protein Ran OS=Homo sapiens OX=9606 GN=RAN PE=1 SV=3 | 15 |
| GYS1 | Glycogen [starch] synthase, muscle OS=Homo sapiens OX=9606 GN=GYS1 PE=1 SV=2 | 1 |
| GPR39 | G-protein coupled receptor 39 OS=Homo sapiens OX=9606 GN=GPR39 PE=1 SV=1 | 2 |
| EMC2 | ER membrane protein complex subunit 2 OS=Homo sapiens OX=9606 GN=EMC2 PE=1 SV=1 | 5 |
| KIF5C | Kinesin heavy chain isoform 5C OS=Homo sapiens OX=9606 GN=KIF5C PE=1 SV=1 | 7 |
| CYB5R3 | NADH-cytochrome b5 reductase 3 OS=Homo sapiens OX=9606 GN=CYB5R3 PE=1 SV=3 | 15 |
| PLCH1 | 1-phosphatidylinositol 4,5-bisphosphate phosphodiesterase eta-1 OS=Homo sapiens OX=9606 GN=PLCH1 PE=1 SV=1 | 1 |
| DNAJC13 | DnaJ homolog subfamily C member 13 OS=Homo sapiens OX=9606 GN=DNAJC13 PE=1 SV=5 | 90 |
| DBT | Lipoamide acyltransferase component of branched-chain alpha-keto acid dehydrogenase complex, mitochondrial OS=Homo sapiens OX=9606 GN=DBT PE=1 SV=3 | 7 |
| COPS6 | COP9 signalosome complex subunit 6 OS=Homo sapiens OX=9606 GN=COPS6 PE=1 SV=1 | 11 |
| FLOT2 | Flotillin-2 OS=Homo sapiens OX=9606 GN=FLOT2 PE=1 SV=2 | 6 |
| ITGB1 | Integrin beta-1 OS=Homo sapiens OX=9606 GN=ITGB1 PE=1 SV=2 | 21 |
| AHCY | Adenosylhomocysteinase OS=Homo sapiens OX=9606 GN=AHCY PE=1 SV=4 | 28 |
| NAP1L4 | Nucleosome assembly protein 1-like 4 OS=Homo sapiens OX=9606 GN=NAP1L4 PE=1 SV=1 | 12 |
| PLCG2 | 1-phosphatidylinositol 4,5-bisphosphate phosphodiesterase gamma-2 OS=Homo sapiens OX=9606 GN=PLCG2 PE=1 SV=4 | 29 |
| RPL13A | 60S ribosomal protein L13a OS=Homo sapiens OX=9606 GN=RPL13A PE=1 SV=2 | 5 |
| SEC24C | Protein transport protein Sec24C OS=Homo sapiens OX=9606 GN=SEC24C PE=1 SV=3 | 15 |
| CZIB | CXXC motif containing zinc binding protein OS=Homo sapiens OX=9606 GN=CZIB PE=1 SV=1 | 7 |
| TIMM13 | Mitochondrial import inner membrane translocase subunit Tim13 OS=Homo sapiens OX=9606 GN=TIMM13 PE=1 SV=1 | 5 |
| FDFT1 | Squalene synthase OS=Homo sapiens OX=9606 GN=FDFT1 PE=1 SV=1 | 9 |
| UBE2H | Ubiquitin-conjugating enzyme E2 H OS=Homo sapiens OX=9606 GN=UBE2H PE=1 SV=1 | 5 |
| GSPT2 | Eukaryotic peptide chain release factor GTP-binding subunit ERF3B OS=Homo sapiens OX=9606 GN=GSPT2 PE=1 SV=2 | 4 |
| -.26 | Histone H2A.N OS=Homo sapiens OX=9606 GN=H2BN1 PE=2 SV=1 | 3 |
| CCDC22 | Coiled-coil domain-containing protein 22 OS=Homo sapiens OX=9606 GN=CCDC22 PE=1 SV=1 | 14 |
| DYNC1I2 | Cytoplasmic dynein 1 intermediate chain 2 OS=Homo sapiens OX=9606 GN=DYNC1I2 PE=1 SV=3 | 10 |
| RPS2 | 40S ribosomal protein S2 OS=Homo sapiens OX=9606 GN=RPS2 PE=1 SV=2 | 10 |
| PPIB | Peptidyl-prolyl cis-trans isomerase B OS=Homo sapiens OX=9606 GN=PPIB PE=1 SV=2 | 15 |
| NDUFB3 | NADH dehydrogenase [ubiquinone] 1 beta subcomplex subunit 3 OS=Homo sapiens OX=9606 GN=NDUFB3 PE=1 SV=3 | 3 |
| SDF2L1 | Stromal cell-derived factor 2-like protein 1 OS=Homo sapiens OX=9606 GN=SDF2L1 PE=1 SV=2 | 8 |
| ELMOD2 | ELMO domain-containing protein 2 OS=Homo sapiens OX=9606 GN=ELMOD2 PE=1 SV=1 | 5 |
| IGFBP3 | Insulin-like growth factor-binding protein 3 OS=Homo sapiens OX=9606 GN=IGFBP3 PE=1 SV=2 | 2 |
| ATP5PO | ATP synthase subunit O, mitochondrial OS=Homo sapiens OX=9606 GN=ATP5PO PE=1 SV=1 | 11 |
| ARHGEF7 | Rho guanine nucleotide exchange factor 7 OS=Homo sapiens OX=9606 GN=ARHGEF7 PE=1 SV=2 | 16 |
| SH3KBP1 | SH3 domain-containing kinase-binding protein 1 OS=Homo sapiens OX=9606 GN=SH3KBP1 PE=1 SV=2 | 19 |
| DLG3 | Disks large homolog 3 OS=Homo sapiens OX=9606 GN=DLG3 PE=1 SV=2 | 4 |
| RBX1 | E3 ubiquitin-protein ligase RBX1 OS=Homo sapiens OX=9606 GN=RBX1 PE=1 SV=1 | 1 |
| NAPA | Alpha-soluble NSF attachment protein OS=Homo sapiens OX=9606 GN=NAPA PE=1 SV=3 | 23 |
| KIF13B | Kinesin-like protein KIF13B OS=Homo sapiens OX=9606 GN=KIF13B PE=1 SV=2 | 72 |
| RPL36 | 60S ribosomal protein L36 OS=Homo sapiens OX=9606 GN=RPL36 PE=1 SV=3 | 2 |
| HGS | Hepatocyte growth factor-regulated tyrosine kinase substrate OS=Homo sapiens OX=9606 GN=HGS PE=1 SV=1 | 9 |
| SDSL | Serine dehydratase-like OS=Homo sapiens OX=9606 GN=SDSL PE=1 SV=1 | 11 |
| NIF3L1 | NIF3-like protein 1 OS=Homo sapiens OX=9606 GN=NIF3L1 PE=1 SV=2 | 8 |
| VIM | Vimentin OS=Homo sapiens OX=9606 GN=VIM PE=1 SV=4 | 6 |
| PSMF1 | Proteasome inhibitor PI31 subunit OS=Homo sapiens OX=9606 GN=PSMF1 PE=1 SV=2 | 3 |
| GDI1 | Rab GDP dissociation inhibitor alpha OS=Homo sapiens OX=9606 GN=GDI1 PE=1 SV=2 | 23 |
| GPHN | Gephyrin OS=Homo sapiens OX=9606 GN=GPHN PE=1 SV=1 | 2 |
| SOD2 | Superoxide dismutase [Mn], mitochondrial OS=Homo sapiens OX=9606 GN=SOD2 PE=1 SV=3 | 12 |
| SDHB | Succinate dehydrogenase [ubiquinone] iron-sulfur subunit, mitochondrial OS=Homo sapiens OX=9606 GN=SDHB PE=1 SV=3 | 9 |
| TRAPPC11 | Trafficking protein particle complex subunit 11 OS=Homo sapiens OX=9606 GN=TRAPPC11 PE=1 SV=2 | 4 |
| ST13 | Hsc70-interacting protein OS=Homo sapiens OX=9606 GN=ST13 PE=1 SV=2 | 13 |
| MYO5A | Unconventional myosin-Va OS=Homo sapiens OX=9606 GN=MYO5A PE=1 SV=2 | 27 |
| TBC1D22A | TBC1 domain family member 22A OS=Homo sapiens OX=9606 GN=TBC1D22A PE=1 SV=2 | 1 |
| ABCD1 | ATP-binding cassette sub-family D member 1 OS=Homo sapiens OX=9606 GN=ABCD1 PE=1 SV=2 | 2 |
| CNOT3 | CCR4-NOT transcription complex subunit 3 OS=Homo sapiens OX=9606 GN=CNOT3 PE=1 SV=1 | 3 |
| SMPDL3B | Acid sphingomyelinase-like phosphodiesterase 3b OS=Homo sapiens OX=9606 GN=SMPDL3B PE=1 SV=2 | 7 |
| CCDC90B | Coiled-coil domain-containing protein 90B, mitochondrial OS=Homo sapiens OX=9606 GN=CCDC90B PE=1 SV=2 | 5 |
| CLPTM1L | Lipid scramblase CLPTM1L OS=Homo sapiens OX=9606 GN=CLPTM1L PE=1 SV=1 | 5 |
| GEMIN5 | Gem-associated protein 5 OS=Homo sapiens OX=9606 GN=GEMIN5 PE=1 SV=3 | 15 |
| COPS7B | COP9 signalosome complex subunit 7b OS=Homo sapiens OX=9606 GN=COPS7B PE=1 SV=1 | 8 |
| RPA3 | Replication protein A 14 kDa subunit OS=Homo sapiens OX=9606 GN=RPA3 PE=1 SV=1 | 4 |
| ACP1 | Low molecular weight phosphotyrosine protein phosphatase OS=Homo sapiens OX=9606 GN=ACP1 PE=1 SV=3 | 8 |
| CALB2 | Calretinin OS=Homo sapiens OX=9606 GN=CALB2 PE=2 SV=2 | 6 |
| IRAG1 | Inositol 1,4,5-triphosphate receptor associated 1 OS=Homo sapiens OX=9606 GN=IRAG1 PE=1 SV=4 | 4 |
| NTAN1 | Protein N-terminal asparagine amidohydrolase OS=Homo sapiens OX=9606 GN=NTAN1 PE=1 SV=3 | 16 |
| TOMM40 | Mitochondrial import receptor subunit TOM40 homolog OS=Homo sapiens OX=9606 GN=TOMM40 PE=1 SV=1 | 6 |
| CYB5B | Cytochrome b5 type B OS=Homo sapiens OX=9606 GN=CYB5B PE=1 SV=3 | 3 |
| DTX2 | Probable E3 ubiquitin-protein ligase DTX2 OS=Homo sapiens OX=9606 GN=DTX2 PE=1 SV=3 | 2 |
| WDCP | WD repeat and coiled-coil-containing protein OS=Homo sapiens OX=9606 GN=WDCP PE=1 SV=1 | 7 |
| ITGB5 | Integrin beta-5 OS=Homo sapiens OX=9606 GN=ITGB5 PE=1 SV=1 | 5 |
| ECI1 | Enoyl-CoA delta isomerase 1, mitochondrial OS=Homo sapiens OX=9606 GN=ECI1 PE=1 SV=1 | 8 |
| SSB | Lupus La protein OS=Homo sapiens OX=9606 GN=SSB PE=1 SV=2 | 9 |
| PRMT1 | Protein arginine N-methyltransferase 1 OS=Homo sapiens OX=9606 GN=PRMT1 PE=1 SV=3 | 12 |
| RAB2A | Ras-related protein Rab-2A OS=Homo sapiens OX=9606 GN=RAB2A PE=1 SV=1 | 16 |
| PNPLA6 | Patatin-like phospholipase domain-containing protein 6 OS=Homo sapiens OX=9606 GN=PNPLA6 PE=1 SV=3 | 8 |
| NDUFS7 | NADH dehydrogenase [ubiquinone] iron-sulfur protein 7, mitochondrial OS=Homo sapiens OX=9606 GN=NDUFS7 PE=1 SV=3 | 3 |
| FARSB | Phenylalanine--tRNA ligase beta subunit OS=Homo sapiens OX=9606 GN=FARSB PE=1 SV=3 | 21 |
| AMACR | Alpha-methylacyl-CoA racemase OS=Homo sapiens OX=9606 GN=AMACR PE=1 SV=2 | 13 |
| CPPED1 | Serine/threonine-protein phosphatase CPPED1 OS=Homo sapiens OX=9606 GN=CPPED1 PE=1 SV=3 | 3 |
| CFL2 | Cofilin-2 OS=Homo sapiens OX=9606 GN=CFL2 PE=1 SV=1 | 8 |
| TOM1 | Target of Myb1 membrane trafficking protein OS=Homo sapiens OX=9606 GN=TOM1 PE=1 SV=2 | 3 |
| C3orf33 | Protein C3orf33 OS=Homo sapiens OX=9606 GN=C3orf33 PE=1 SV=2 | 4 |
| EIF5A | Eukaryotic translation initiation factor 5A-1 OS=Homo sapiens OX=9606 GN=EIF5A PE=1 SV=2 | 8 |
| SEPTIN9 | Septin-9 OS=Homo sapiens OX=9606 GN=SEPTIN9 PE=1 SV=2 | 24 |
| TBCB | Tubulin-folding cofactor B OS=Homo sapiens OX=9606 GN=TBCB PE=1 SV=2 | 15 |
| LYRM4 | LYR motif-containing protein 4 OS=Homo sapiens OX=9606 GN=LYRM4 PE=1 SV=1 | 3 |
| HSP90B1 | Endoplasmin OS=Homo sapiens OX=9606 GN=HSP90B1 PE=1 SV=1 | 63 |
| POFUT1 | GDP-fucose protein O-fucosyltransferase 1 OS=Homo sapiens OX=9606 GN=POFUT1 PE=1 SV=1 | 7 |
| UBA6 | Ubiquitin-like modifier-activating enzyme 6 OS=Homo sapiens OX=9606 GN=UBA6 PE=1 SV=1 | 50 |
| DIAPH1 | Protein diaphanous homolog 1 OS=Homo sapiens OX=9606 GN=DIAPH1 PE=1 SV=2 | 37 |
| UBE2M | NEDD8-conjugating enzyme Ubc12 OS=Homo sapiens OX=9606 GN=UBE2M PE=1 SV=1 | 8 |
| BIRC6 | Baculoviral IAP repeat-containing protein 6 OS=Homo sapiens OX=9606 GN=BIRC6 PE=1 SV=3 | 3 |
| SPR | Sepiapterin reductase OS=Homo sapiens OX=9606 GN=SPR PE=1 SV=1 | 16 |
| SEC11A | Signal peptidase complex catalytic subunit SEC11A OS=Homo sapiens OX=9606 GN=SEC11A PE=1 SV=1 | 2 |
| IBA57 | Putative transferase CAF17, mitochondrial OS=Homo sapiens OX=9606 GN=IBA57 PE=1 SV=1 | 2 |
| RAB7A | Ras-related protein Rab-7a OS=Homo sapiens OX=9606 GN=RAB7A PE=1 SV=1 | 13 |
| HMOX2 | Heme oxygenase 2 OS=Homo sapiens OX=9606 GN=HMOX2 PE=1 SV=2 | 15 |
| EIF4A2 | Eukaryotic initiation factor 4A-II OS=Homo sapiens OX=9606 GN=EIF4A2 PE=1 SV=2 | 7 |
| AP1B1 | AP-1 complex subunit beta-1 OS=Homo sapiens OX=9606 GN=AP1B1 PE=1 SV=3 | 31 |
| TSFM | Elongation factor Ts, mitochondrial OS=Homo sapiens OX=9606 GN=TSFM PE=1 SV=2 | 14 |
| NDUFA2 | NADH dehydrogenase [ubiquinone] 1 alpha subcomplex subunit 2 OS=Homo sapiens OX=9606 GN=NDUFA2 PE=1 SV=3 | 3 |
| CHMP7 | Charged multivesicular body protein 7 OS=Homo sapiens OX=9606 GN=CHMP7 PE=1 SV=1 | 1 |
| GMFB | Glia maturation factor beta OS=Homo sapiens OX=9606 GN=GMFB PE=1 SV=2 | 4 |
| PPM1F | Protein phosphatase 1F OS=Homo sapiens OX=9606 GN=PPM1F PE=1 SV=3 | 22 |
| CIAO1 | Probable cytosolic iron-sulfur protein assembly protein CIAO1 OS=Homo sapiens OX=9606 GN=CIAO1 PE=1 SV=1 | 10 |
| NAP1L1 | Nucleosome assembly protein 1-like 1 OS=Homo sapiens OX=9606 GN=NAP1L1 PE=1 SV=1 | 9 |
| SRP14 | Signal recognition particle 14 kDa protein OS=Homo sapiens OX=9606 GN=SRP14 PE=1 SV=2 | 2 |
| ADA | Adenosine deaminase OS=Homo sapiens OX=9606 GN=ADA PE=1 SV=3 | 5 |
| SLC30A3 | Probable proton-coupled zinc antiporter SLC30A3 OS=Homo sapiens OX=9606 GN=SLC30A3 PE=1 SV=2 | 1 |
| PCYOX1L | Prenylcysteine oxidase-like OS=Homo sapiens OX=9606 GN=PCYOX1L PE=1 SV=2 | 3 |
| DNPEP | Aspartyl aminopeptidase OS=Homo sapiens OX=9606 GN=DNPEP PE=1 SV=2 | 10 |
| SCAMP2 | Secretory carrier-associated membrane protein 2 OS=Homo sapiens OX=9606 GN=SCAMP2 PE=1 SV=2 | 4 |
| RPL4 | 60S ribosomal protein L4 OS=Homo sapiens OX=9606 GN=RPL4 PE=1 SV=5 | 12 |
| UBAC1 | Ubiquitin-associated domain-containing protein 1 OS=Homo sapiens OX=9606 GN=UBAC1 PE=1 SV=1 | 1 |
| DCTN3 | Dynactin subunit 3 OS=Homo sapiens OX=9606 GN=DCTN3 PE=1 SV=1 | 5 |
| RBBP4 | Histone-binding protein RBBP4 OS=Homo sapiens OX=9606 GN=RBBP4 PE=1 SV=3 | 13 |
| PDIA4 | Protein disulfide-isomerase A4 OS=Homo sapiens OX=9606 GN=PDIA4 PE=1 SV=2 | 53 |
| HINT2 | Adenosine 5'-monophosphoramidase HINT2 OS=Homo sapiens OX=9606 GN=HINT2 PE=1 SV=1 | 6 |
| NGRN | Neugrin OS=Homo sapiens OX=9606 GN=NGRN PE=1 SV=2 | 8 |
| PPP2R2A | Serine/threonine-protein phosphatase 2A 55 kDa regulatory subunit B alpha isoform OS=Homo sapiens OX=9606 GN=PPP2R2A PE=1 SV=1 | 10 |
| UBL4A | Ubiquitin-like protein 4A OS=Homo sapiens OX=9606 GN=UBL4A PE=1 SV=1 | 3 |
| CAPN2 | Calpain-2 catalytic subunit OS=Homo sapiens OX=9606 GN=CAPN2 PE=1 SV=6 | 20 |
| ABCD3 | ATP-binding cassette sub-family D member 3 OS=Homo sapiens OX=9606 GN=ABCD3 PE=1 SV=1 | 7 |
| BAG6 | Large proline-rich protein BAG6 OS=Homo sapiens OX=9606 GN=BAG6 PE=1 SV=2 | 12 |
| TLN2 | Talin-2 OS=Homo sapiens OX=9606 GN=TLN2 PE=1 SV=4 | 22 |
| TDRD6 | Tudor domain-containing protein 6 OS=Homo sapiens OX=9606 GN=TDRD6 PE=2 SV=2 | 15 |
| PITPNA | Phosphatidylinositol transfer protein alpha isoform OS=Homo sapiens OX=9606 GN=PITPNA PE=1 SV=2 | 20 |
| XPO7 | Exportin-7 OS=Homo sapiens OX=9606 GN=XPO7 PE=1 SV=3 | 6 |
| VARS2 | Valine--tRNA ligase, mitochondrial OS=Homo sapiens OX=9606 GN=VARS2 PE=1 SV=2 | 9 |
| GTSE1 | G2 and S phase-expressed protein 1 OS=Homo sapiens OX=9606 GN=GTSE1 PE=1 SV=3 | 1 |
| AHSA1 | Activator of 90 kDa heat shock protein ATPase homolog 1 OS=Homo sapiens OX=9606 GN=AHSA1 PE=1 SV=1 | 22 |
| ATP5F1A | ATP synthase subunit alpha, mitochondrial OS=Homo sapiens OX=9606 GN=ATP5F1A PE=1 SV=1 | 34 |
| ATP2B1 | Plasma membrane calcium-transporting ATPase 1 OS=Homo sapiens OX=9606 GN=ATP2B1 PE=1 SV=4 | 25 |
| CCNYL1 | Cyclin-Y-like protein 1 OS=Homo sapiens OX=9606 GN=CCNYL1 PE=1 SV=2 | 2 |
| CHERP | Calcium homeostasis endoplasmic reticulum protein OS=Homo sapiens OX=9606 GN=CHERP PE=1 SV=3 | 4 |
| RWDD4 | RWD domain-containing protein 4 OS=Homo sapiens OX=9606 GN=RWDD4 PE=1 SV=3 | 1 |
| CDC37 | Hsp90 co-chaperone Cdc37 OS=Homo sapiens OX=9606 GN=CDC37 PE=1 SV=1 | 14 |
| WBP2 | WW domain-binding protein 2 OS=Homo sapiens OX=9606 GN=WBP2 PE=1 SV=1 | 4 |
| STOML2 | Stomatin-like protein 2, mitochondrial OS=Homo sapiens OX=9606 GN=STOML2 PE=1 SV=1 | 17 |
| SDS | L-serine dehydratase/L-threonine deaminase OS=Homo sapiens OX=9606 GN=SDS PE=1 SV=2 | 3 |
| ZW10 | Centromere/kinetochore protein zw10 homolog OS=Homo sapiens OX=9606 GN=ZW10 PE=1 SV=3 | 8 |
| CBR1 | Carbonyl reductase [NADPH] 1 OS=Homo sapiens OX=9606 GN=CBR1 PE=1 SV=3 | 4 |
| GOPC | Golgi-associated PDZ and coiled-coil motif-containing protein OS=Homo sapiens OX=9606 GN=GOPC PE=1 SV=1 | 7 |
| PPA2 | Inorganic pyrophosphatase 2, mitochondrial OS=Homo sapiens OX=9606 GN=PPA2 PE=1 SV=2 | 12 |
| DOP1A | Protein dopey-1 OS=Homo sapiens OX=9606 GN=DOP1A PE=2 SV=1 | 4 |
| FAM114A2 | Protein FAM114A2 OS=Homo sapiens OX=9606 GN=FAM114A2 PE=1 SV=4 | 20 |
| RUSF1 | RUS family member 1 OS=Homo sapiens OX=9606 GN=RUSF1 PE=1 SV=2 | 4 |
| ISOC1 | Isochorismatase domain-containing protein 1 OS=Homo sapiens OX=9606 GN=ISOC1 PE=1 SV=3 | 8 |
| ECHDC1 | Ethylmalonyl-CoA decarboxylase OS=Homo sapiens OX=9606 GN=ECHDC1 PE=1 SV=2 | 10 |
| MESD | LRP chaperone MESD OS=Homo sapiens OX=9606 GN=MESD PE=1 SV=2 | 10 |
| FCSK | L-fucose kinase OS=Homo sapiens OX=9606 GN=FCSK PE=1 SV=2 | 17 |
| PLP1 | Myelin proteolipid protein OS=Homo sapiens OX=9606 GN=PLP1 PE=1 SV=2 | 3 |
| PFKM | ATP-dependent 6-phosphofructokinase, muscle type OS=Homo sapiens OX=9606 GN=PFKM PE=1 SV=2 | 15 |
| MRPL44 | 39S ribosomal protein L44, mitochondrial OS=Homo sapiens OX=9606 GN=MRPL44 PE=1 SV=1 | 9 |
| CSTF2 | Cleavage stimulation factor subunit 2 OS=Homo sapiens OX=9606 GN=CSTF2 PE=1 SV=1 | 11 |
| YARS2 | Tyrosine--tRNA ligase, mitochondrial OS=Homo sapiens OX=9606 GN=YARS2 PE=1 SV=2 | 17 |
| MVD | Diphosphomevalonate decarboxylase OS=Homo sapiens OX=9606 GN=MVD PE=1 SV=1 | 12 |
| GDF9 | Growth/differentiation factor 9 OS=Homo sapiens OX=9606 GN=GDF9 PE=1 SV=1 | 25 |
| DYNLL1 | Dynein light chain 1, cytoplasmic OS=Homo sapiens OX=9606 GN=DYNLL1 PE=1 SV=1 | 8 |
| SDCBP | Syntenin-1 OS=Homo sapiens OX=9606 GN=SDCBP PE=1 SV=1 | 4 |
| HADHA | Trifunctional enzyme subunit alpha, mitochondrial OS=Homo sapiens OX=9606 GN=HADHA PE=1 SV=2 | 32 |
| AP1M1 | AP-1 complex subunit mu-1 OS=Homo sapiens OX=9606 GN=AP1M1 PE=1 SV=3 | 12 |
| "BOLA2B | BolA-like protein 2 OS=Homo sapiens OX=9606 GN=BOLA2B PE=1 SV=1 | 4 |
| PRKAR1A | cAMP-dependent protein kinase type I-alpha regulatory subunit OS=Homo sapiens OX=9606 GN=PRKAR1A PE=1 SV=1 | 15 |
| DLST | Dihydrolipoyllysine-residue succinyltransferase component of 2-oxoglutarate dehydrogenase complex, mitochondrial OS=Homo sapiens OX=9606 GN=DLST PE=1 SV=4 | 23 |
| TLE6 | Transducin-like enhancer protein 6 OS=Homo sapiens OX=9606 GN=TLE6 PE=1 SV=2 | 33 |
| AP3B1 | AP-3 complex subunit beta-1 OS=Homo sapiens OX=9606 GN=AP3B1 PE=1 SV=3 | 18 |
| TKFC | Triokinase/FMN cyclase OS=Homo sapiens OX=9606 GN=TKFC PE=1 SV=2 | 13 |
| CMPK1 | UMP-CMP kinase OS=Homo sapiens OX=9606 GN=CMPK1 PE=1 SV=3 | 16 |
| PTGFRN | Prostaglandin F2 receptor negative regulator OS=Homo sapiens OX=9606 GN=PTGFRN PE=1 SV=2 | 26 |
| VCPIP1 | Deubiquitinating protein VCPIP1 OS=Homo sapiens OX=9606 GN=VCPIP1 PE=1 SV=2 | 15 |
| LSM6 | U6 snRNA-associated Sm-like protein LSm6 OS=Homo sapiens OX=9606 GN=LSM6 PE=1 SV=1 | 7 |
| WASH3P | Putative WAS protein family homolog 3 OS=Homo sapiens OX=9606 GN=WASH3P PE=1 SV=2 | 5 |
| STX3 | Syntaxin-3 OS=Homo sapiens OX=9606 GN=STX3 PE=1 SV=3 | 14 |
| CCDC6 | Coiled-coil domain-containing protein 6 OS=Homo sapiens OX=9606 GN=CCDC6 PE=1 SV=2 | 25 |
| RINT1 | RAD50-interacting protein 1 OS=Homo sapiens OX=9606 GN=RINT1 PE=1 SV=1 | 5 |
| TRMT10C | tRNA methyltransferase 10 homolog C OS=Homo sapiens OX=9606 GN=TRMT10C PE=1 SV=2 | 13 |
| NCK2 | Cytoplasmic protein NCK2 OS=Homo sapiens OX=9606 GN=NCK2 PE=1 SV=2 | 17 |
| CTH | Cystathionine gamma-lyase OS=Homo sapiens OX=9606 GN=CTH PE=1 SV=3 | 3 |
| AKTIP | AKT-interacting protein OS=Homo sapiens OX=9606 GN=AKTIP PE=1 SV=1 | 2 |
| MAPRE1 | Microtubule-associated protein RP/EB family member 1 OS=Homo sapiens OX=9606 GN=MAPRE1 PE=1 SV=3 | 14 |
| AK2 | Adenylate kinase 2, mitochondrial OS=Homo sapiens OX=9606 GN=AK2 PE=1 SV=2 | 16 |
| TTC9C | Tetratricopeptide repeat protein 9C OS=Homo sapiens OX=9606 GN=TTC9C PE=1 SV=1 | 5 |
| ATP5MK | ATP synthase membrane subunit K, mitochondrial OS=Homo sapiens OX=9606 GN=ATP5MK PE=1 SV=1 | 3 |
| SEC23A | Protein transport protein Sec23A OS=Homo sapiens OX=9606 GN=SEC23A PE=1 SV=2 | 23 |
| UBE2I | SUMO-conjugating enzyme UBC9 OS=Homo sapiens OX=9606 GN=UBE2I PE=1 SV=1 | 7 |
| GOT2 | Aspartate aminotransferase, mitochondrial OS=Homo sapiens OX=9606 GN=GOT2 PE=1 SV=3 | 28 |
| CARM1 | Histone-arginine methyltransferase CARM1 OS=Homo sapiens OX=9606 GN=CARM1 PE=1 SV=3 | 2 |
| FKBP2 | Peptidyl-prolyl cis-trans isomerase FKBP2 OS=Homo sapiens OX=9606 GN=FKBP2 PE=1 SV=2 | 9 |
| CETN2 | Centrin-2 OS=Homo sapiens OX=9606 GN=CETN2 PE=1 SV=1 | 3 |
| MVP | Major vault protein OS=Homo sapiens OX=9606 GN=MVP PE=1 SV=4 | 82 |
| TMEM11 | Transmembrane protein 11, mitochondrial OS=Homo sapiens OX=9606 GN=TMEM11 PE=1 SV=1 | 6 |
| ALKBH3 | Alpha-ketoglutarate-dependent dioxygenase alkB homolog 3 OS=Homo sapiens OX=9606 GN=ALKBH3 PE=1 SV=1 | 5 |
| COX20 | Cytochrome c oxidase assembly protein COX20, mitochondrial OS=Homo sapiens OX=9606 GN=COX20 PE=1 SV=2 | 4 |
| AP1G1 | AP-1 complex subunit gamma-1 OS=Homo sapiens OX=9606 GN=AP1G1 PE=1 SV=5 | 10 |
| VPS13C | Intermembrane lipid transfer protein VPS13C OS=Homo sapiens OX=9606 GN=VPS13C PE=1 SV=1 | 44 |
| ARHGAP10 | Rho GTPase-activating protein 10 OS=Homo sapiens OX=9606 GN=ARHGAP10 PE=1 SV=1 | 1 |
| PSMD12 | 26S proteasome non-ATPase regulatory subunit 12 OS=Homo sapiens OX=9606 GN=PSMD12 PE=1 SV=3 | 24 |
| ARPC1B | Actin-related protein 2/3 complex subunit 1B OS=Homo sapiens OX=9606 GN=ARPC1B PE=1 SV=3 | 13 |
| HSPA4L | Heat shock 70 kDa protein 4L OS=Homo sapiens OX=9606 GN=HSPA4L PE=1 SV=3 | 40 |
| USP5 | Ubiquitin carboxyl-terminal hydrolase 5 OS=Homo sapiens OX=9606 GN=USP5 PE=1 SV=2 | 45 |
| HTATSF1 | HIV Tat-specific factor 1 OS=Homo sapiens OX=9606 GN=HTATSF1 PE=1 SV=1 | 6 |
| XPOT | Exportin-T OS=Homo sapiens OX=9606 GN=XPOT PE=1 SV=2 | 3 |
| PGAM1 | Phosphoglycerate mutase 1 OS=Homo sapiens OX=9606 GN=PGAM1 PE=1 SV=2 | 13 |
| UBAP2L | Ubiquitin-associated protein 2-like OS=Homo sapiens OX=9606 GN=UBAP2L PE=1 SV=2 | 6 |
| EFTUD2 | 116 kDa U5 small nuclear ribonucleoprotein component OS=Homo sapiens OX=9606 GN=EFTUD2 PE=1 SV=1 | 12 |
| RAB1B | Ras-related protein Rab-1B OS=Homo sapiens OX=9606 GN=RAB1B PE=1 SV=1 | 5 |
| TSPYL1 | Testis-specific Y-encoded-like protein 1 OS=Homo sapiens OX=9606 GN=TSPYL1 PE=1 SV=3 | 1 |
| PPIA | Peptidyl-prolyl cis-trans isomerase A OS=Homo sapiens OX=9606 GN=PPIA PE=1 SV=2 | 12 |
| WDR47 | WD repeat-containing protein 47 OS=Homo sapiens OX=9606 GN=WDR47 PE=1 SV=1 | 4 |
| RTCA | RNA 3'-terminal phosphate cyclase OS=Homo sapiens OX=9606 GN=RTCA PE=1 SV=1 | 11 |
| UBFD1 | Ubiquitin domain-containing protein UBFD1 OS=Homo sapiens OX=9606 GN=UBFD1 PE=1 SV=2 | 6 |
| SNX30 | Sorting nexin-30 OS=Homo sapiens OX=9606 GN=SNX30 PE=1 SV=1 | 16 |
| SYNJ1 | Synaptojanin-1 OS=Homo sapiens OX=9606 GN=SYNJ1 PE=1 SV=2 | 9 |
| GOLPH3L | Golgi phosphoprotein 3-like OS=Homo sapiens OX=9606 GN=GOLPH3L PE=1 SV=1 | 11 |
| CNOT10 | CCR4-NOT transcription complex subunit 10 OS=Homo sapiens OX=9606 GN=CNOT10 PE=1 SV=1 | 2 |
| PCMT1 | Protein-L-isoaspartate(D-aspartate) O-methyltransferase OS=Homo sapiens OX=9606 GN=PCMT1 PE=1 SV=4 | 15 |
| EIF3A | Eukaryotic translation initiation factor 3 subunit A OS=Homo sapiens OX=9606 GN=EIF3A PE=1 SV=1 | 14 |
| SEC23B | Protein transport protein Sec23B OS=Homo sapiens OX=9606 GN=SEC23B PE=1 SV=2 | 5 |
| GPCPD1 | Glycerophosphocholine phosphodiesterase GPCPD1 OS=Homo sapiens OX=9606 GN=GPCPD1 PE=1 SV=2 | 3 |
| DPYSL5 | Dihydropyrimidinase-related protein 5 OS=Homo sapiens OX=9606 GN=DPYSL5 PE=1 SV=1 | 18 |
| SLC27A4 | Long-chain fatty acid transport protein 4 OS=Homo sapiens OX=9606 GN=SLC27A4 PE=1 SV=1 | 16 |
| WARS1 | Tryptophan--tRNA ligase, cytoplasmic OS=Homo sapiens OX=9606 GN=WARS1 PE=1 SV=2 | 21 |
| CRISPLD1 | Cysteine-rich secretory protein LCCL domain-containing 1 OS=Homo sapiens OX=9606 GN=CRISPLD1 PE=2 SV=1 | 11 |
| IAH1 | Isoamyl acetate-hydrolyzing esterase 1 homolog OS=Homo sapiens OX=9606 GN=IAH1 PE=1 SV=1 | 3 |
| NPC2 | NPC intracellular cholesterol transporter 2 OS=Homo sapiens OX=9606 GN=NPC2 PE=1 SV=1 | 5 |
| COPS2 | COP9 signalosome complex subunit 2 OS=Homo sapiens OX=9606 GN=COPS2 PE=1 SV=1 | 24 |
| DNAJB11 | DnaJ homolog subfamily B member 11 OS=Homo sapiens OX=9606 GN=DNAJB11 PE=1 SV=1 | 14 |
| CLDND1 | Claudin domain-containing protein 1 OS=Homo sapiens OX=9606 GN=CLDND1 PE=1 SV=1 | 1 |
| NFU1 | NFU1 iron-sulfur cluster scaffold homolog, mitochondrial OS=Homo sapiens OX=9606 GN=NFU1 PE=1 SV=2 | 6 |
| GIT2 | ARF GTPase-activating protein GIT2 OS=Homo sapiens OX=9606 GN=GIT2 PE=1 SV=2 | 15 |
| IGSF3 | Immunoglobulin superfamily member 3 OS=Homo sapiens OX=9606 GN=IGSF3 PE=2 SV=3 | 2 |
| NDUFV3 | NADH dehydrogenase [ubiquinone] flavoprotein 3, mitochondrial OS=Homo sapiens OX=9606 GN=NDUFV3 PE=1 SV=2 | 1 |
| H2BC12 | Histone H2B type 1-K OS=Homo sapiens OX=9606 GN=H2BC12 PE=1 SV=3 | 7 |
| ARHGAP1 | Rho GTPase-activating protein 1 OS=Homo sapiens OX=9606 GN=ARHGAP1 PE=1 SV=1 | 12 |
| FAM120B | Constitutive coactivator of peroxisome proliferator-activated receptor gamma OS=Homo sapiens OX=9606 GN=FAM120B PE=1 SV=1 | 2 |
| GMDS | GDP-mannose 4,6 dehydratase OS=Homo sapiens OX=9606 GN=GMDS PE=1 SV=1 | 15 |
| ACO2 | Aconitate hydratase, mitochondrial OS=Homo sapiens OX=9606 GN=ACO2 PE=1 SV=2 | 35 |
| PRPSAP2 | Phosphoribosyl pyrophosphate synthase-associated protein 2 OS=Homo sapiens OX=9606 GN=PRPSAP2 PE=1 SV=1 | 13 |
| GLCCI1 | Glucocorticoid-induced transcript 1 protein OS=Homo sapiens OX=9606 GN=GLCCI1 PE=1 SV=1 | 2 |
| RPS6KA3 | Ribosomal protein S6 kinase alpha-3 OS=Homo sapiens OX=9606 GN=RPS6KA3 PE=1 SV=1 | 19 |
| FH | Fumarate hydratase, mitochondrial OS=Homo sapiens OX=9606 GN=FH PE=1 SV=3 | 24 |
| TUBGCP2 | Gamma-tubulin complex component 2 OS=Homo sapiens OX=9606 GN=TUBGCP2 PE=1 SV=2 | 12 |
| SRXN1 | Sulfiredoxin-1 OS=Homo sapiens OX=9606 GN=SRXN1 PE=1 SV=2 | 1 |
| MYO1E | Unconventional myosin-Ie OS=Homo sapiens OX=9606 GN=MYO1E PE=1 SV=2 | 2 |
| NEDD4L | E3 ubiquitin-protein ligase NEDD4-like OS=Homo sapiens OX=9606 GN=NEDD4L PE=1 SV=2 | 10 |
| PTGES3 | Prostaglandin E synthase 3 OS=Homo sapiens OX=9606 GN=PTGES3 PE=1 SV=1 | 10 |
| NDUFA9 | NADH dehydrogenase [ubiquinone] 1 alpha subcomplex subunit 9, mitochondrial OS=Homo sapiens OX=9606 GN=NDUFA9 PE=1 SV=2 | 6 |
| STX16 | Syntaxin-16 OS=Homo sapiens OX=9606 GN=STX16 PE=1 SV=3 | 4 |
| POLD1 | DNA polymerase delta catalytic subunit OS=Homo sapiens OX=9606 GN=POLD1 PE=1 SV=2 | 2 |
| OTULIN | Ubiquitin thioesterase otulin OS=Homo sapiens OX=9606 GN=OTULIN PE=1 SV=3 | 10 |
| TIMM44 | Mitochondrial import inner membrane translocase subunit TIM44 OS=Homo sapiens OX=9606 GN=TIMM44 PE=1 SV=2 | 18 |
| ELOC | Elongin-C OS=Homo sapiens OX=9606 GN=ELOC PE=1 SV=1 | 8 |
| ENSA | Alpha-endosulfine OS=Homo sapiens OX=9606 GN=ENSA PE=1 SV=1 | 5 |
| APBA1 | Amyloid-beta A4 precursor protein-binding family A member 1 OS=Homo sapiens OX=9606 GN=APBA1 PE=1 SV=3 | 7 |
| TMED9 | Transmembrane emp24 domain-containing protein 9 OS=Homo sapiens OX=9606 GN=TMED9 PE=1 SV=2 | 6 |
| ZP3 | Zona pellucida sperm-binding protein 3 OS=Homo sapiens OX=9606 GN=ZP3 PE=1 SV=2 | 21 |
| CCT6B | T-complex protein 1 subunit zeta-2 OS=Homo sapiens OX=9606 GN=CCT6B PE=1 SV=5 | 4 |
| PGPEP1 | Pyroglutamyl-peptidase 1 OS=Homo sapiens OX=9606 GN=PGPEP1 PE=1 SV=1 | 4 |
| RAB8A | Ras-related protein Rab-8A OS=Homo sapiens OX=9606 GN=RAB8A PE=1 SV=1 | 10 |
| RPL27A | 60S ribosomal protein L27a OS=Homo sapiens OX=9606 GN=RPL27A PE=1 SV=2 | 5 |
| PATL2 | Protein PAT1 homolog 2 OS=Homo sapiens OX=9606 GN=PATL2 PE=1 SV=1 | 20 |
| COQ6 | Ubiquinone biosynthesis monooxygenase COQ6, mitochondrial OS=Homo sapiens OX=9606 GN=COQ6 PE=1 SV=2 | 6 |
| NUP214 | Nuclear pore complex protein Nup214 OS=Homo sapiens OX=9606 GN=NUP214 PE=1 SV=2 | 7 |
| RRAGB | Ras-related GTP-binding protein B OS=Homo sapiens OX=9606 GN=RRAGB PE=1 SV=1 | 6 |
| YBX1 | Y-box-binding protein 1 OS=Homo sapiens OX=9606 GN=YBX1 PE=1 SV=3 | 4 |
| MRPL23 | 39S ribosomal protein L23, mitochondrial OS=Homo sapiens OX=9606 GN=MRPL23 PE=1 SV=1 | 5 |
| KPNB1 | Importin subunit beta-1 OS=Homo sapiens OX=9606 GN=KPNB1 PE=1 SV=2 | 23 |
| PRKAA1 | 5'-AMP-activated protein kinase catalytic subunit alpha-1 OS=Homo sapiens OX=9606 GN=PRKAA1 PE=1 SV=4 | 7 |
| SHMT2 | Serine hydroxymethyltransferase, mitochondrial OS=Homo sapiens OX=9606 GN=SHMT2 PE=1 SV=3 | 31 |
| CHID1 | Chitinase domain-containing protein 1 OS=Homo sapiens OX=9606 GN=CHID1 PE=1 SV=1 | 8 |
| CAPZB | F-actin-capping protein subunit beta OS=Homo sapiens OX=9606 GN=CAPZB PE=1 SV=5 | 18 |
| AGPAT2 | 1-acyl-sn-glycerol-3-phosphate acyltransferase beta OS=Homo sapiens OX=9606 GN=AGPAT2 PE=1 SV=1 | 2 |
| EIF4E1B | Eukaryotic translation initiation factor 4E type 1B OS=Homo sapiens OX=9606 GN=EIF4E1B PE=1 SV=3 | 13 |
| STUB1 | E3 ubiquitin-protein ligase CHIP OS=Homo sapiens OX=9606 GN=STUB1 PE=1 SV=2 | 16 |
| IMMT | MICOS complex subunit MIC60 OS=Homo sapiens OX=9606 GN=IMMT PE=1 SV=1 | 47 |
| VPS36 | Vacuolar protein-sorting-associated protein 36 OS=Homo sapiens OX=9606 GN=VPS36 PE=1 SV=1 | 8 |
| VPS16 | Vacuolar protein sorting-associated protein 16 homolog OS=Homo sapiens OX=9606 GN=VPS16 PE=1 SV=2 | 8 |
| PRKCSH | Glucosidase 2 subunit beta OS=Homo sapiens OX=9606 GN=PRKCSH PE=1 SV=2 | 18 |
| ATP6AP2 | Renin receptor OS=Homo sapiens OX=9606 GN=ATP6AP2 PE=1 SV=2 | 6 |
| PEX5 | Peroxisomal targeting signal 1 receptor OS=Homo sapiens OX=9606 GN=PEX5 PE=1 SV=3 | 2 |
| RAB3GAP2 | Rab3 GTPase-activating protein non-catalytic subunit OS=Homo sapiens OX=9606 GN=RAB3GAP2 PE=1 SV=1 | 18 |
| ARFIP1 | Arfaptin-1 OS=Homo sapiens OX=9606 GN=ARFIP1 PE=1 SV=2 | 2 |
| TFRC | Transferrin receptor protein 1 OS=Homo sapiens OX=9606 GN=TFRC PE=1 SV=2 | 9 |
| ACTR1B | Beta-centractin OS=Homo sapiens OX=9606 GN=ACTR1B PE=1 SV=1 | 16 |
| PDHA1 | Pyruvate dehydrogenase E1 component subunit alpha, somatic form, mitochondrial OS=Homo sapiens OX=9606 GN=PDHA1 PE=1 SV=3 | 26 |
| PPP6R1 | Serine/threonine-protein phosphatase 6 regulatory subunit 1 OS=Homo sapiens OX=9606 GN=PPP6R1 PE=1 SV=5 | 2 |
| QDPR | Dihydropteridine reductase OS=Homo sapiens OX=9606 GN=QDPR PE=1 SV=2 | 12 |
| GCSH | Glycine cleavage system H protein, mitochondrial OS=Homo sapiens OX=9606 GN=GCSH PE=1 SV=2 | 2 |
| CELF1 | CUGBP Elav-like family member 1 OS=Homo sapiens OX=9606 GN=CELF1 PE=1 SV=2 | 4 |
| RTN4 | Reticulon-4 OS=Homo sapiens OX=9606 GN=RTN4 PE=1 SV=2 | 15 |
| PCBP3 | Poly(rC)-binding protein 3 OS=Homo sapiens OX=9606 GN=PCBP3 PE=1 SV=2 | 1 |
| ALDH4A1 | Delta-1-pyrroline-5-carboxylate dehydrogenase, mitochondrial OS=Homo sapiens OX=9606 GN=ALDH4A1 PE=1 SV=3 | 25 |
| RARS1 | Arginine--tRNA ligase, cytoplasmic OS=Homo sapiens OX=9606 GN=RARS1 PE=1 SV=2 | 32 |
| CNOT11 | CCR4-NOT transcription complex subunit 11 OS=Homo sapiens OX=9606 GN=CNOT11 PE=1 SV=1 | 4 |
| SWAP70 | Switch-associated protein 70 OS=Homo sapiens OX=9606 GN=SWAP70 PE=1 SV=1 | 9 |
| ITCH | E3 ubiquitin-protein ligase Itchy homolog OS=Homo sapiens OX=9606 GN=ITCH PE=1 SV=2 | 1 |
| HAUS5 | HAUS augmin-like complex subunit 5 OS=Homo sapiens OX=9606 GN=HAUS5 PE=1 SV=2 | 1 |
| RPS12 | 40S ribosomal protein S12 OS=Homo sapiens OX=9606 GN=RPS12 PE=1 SV=3 | 7 |
| GFUS | GDP-L-fucose synthase OS=Homo sapiens OX=9606 GN=GFUS PE=1 SV=1 | 18 |
| HNRNPH2 | Heterogeneous nuclear ribonucleoprotein H2 OS=Homo sapiens OX=9606 GN=HNRNPH2 PE=1 SV=1 | 2 |
| CIAPIN1 | Anamorsin OS=Homo sapiens OX=9606 GN=CIAPIN1 PE=1 SV=2 | 13 |
| PPP4R1 | Serine/threonine-protein phosphatase 4 regulatory subunit 1 OS=Homo sapiens OX=9606 GN=PPP4R1 PE=1 SV=1 | 4 |
| GRHPR | Glyoxylate reductase/hydroxypyruvate reductase OS=Homo sapiens OX=9606 GN=GRHPR PE=1 SV=1 | 19 |
| PRUNE1 | Exopolyphosphatase PRUNE1 OS=Homo sapiens OX=9606 GN=PRUNE1 PE=1 SV=2 | 5 |
| UBE2W | Ubiquitin-conjugating enzyme E2 W OS=Homo sapiens OX=9606 GN=UBE2W PE=1 SV=1 | 1 |
| TPP2 | Tripeptidyl-peptidase 2 OS=Homo sapiens OX=9606 GN=TPP2 PE=1 SV=4 | 26 |
| GNS | N-acetylglucosamine-6-sulfatase OS=Homo sapiens OX=9606 GN=GNS PE=1 SV=3 | 15 |
| PSMC5 | 26S proteasome regulatory subunit 8 OS=Homo sapiens OX=9606 GN=PSMC5 PE=1 SV=1 | 20 |
| MINK1 | Misshapen-like kinase 1 OS=Homo sapiens OX=9606 GN=MINK1 PE=1 SV=2 | 5 |
| EHD3 | EH domain-containing protein 3 OS=Homo sapiens OX=9606 GN=EHD3 PE=1 SV=2 | 2 |
| CACYBP | Calcyclin-binding protein OS=Homo sapiens OX=9606 GN=CACYBP PE=1 SV=2 | 14 |
| MAEL | Protein maelstrom homolog OS=Homo sapiens OX=9606 GN=MAEL PE=1 SV=1 | 12 |
| PCP4L1 | Purkinje cell protein 4-like protein 1 OS=Homo sapiens OX=9606 GN=PCP4L1 PE=3 SV=3 | 2 |
| BRAP | BRCA1-associated protein OS=Homo sapiens OX=9606 GN=BRAP PE=1 SV=2 | 4 |
| DDB1 | DNA damage-binding protein 1 OS=Homo sapiens OX=9606 GN=DDB1 PE=1 SV=1 | 36 |
| PRXL2A | Peroxiredoxin-like 2A OS=Homo sapiens OX=9606 GN=PRXL2A PE=1 SV=3 | 10 |
| MLEC | Malectin OS=Homo sapiens OX=9606 GN=MLEC PE=1 SV=1 | 12 |
| DNAJA1 | DnaJ homolog subfamily A member 1 OS=Homo sapiens OX=9606 GN=DNAJA1 PE=1 SV=2 | 16 |
| CLTA | Clathrin light chain A OS=Homo sapiens OX=9606 GN=CLTA PE=1 SV=1 | 8 |
| NDUFAB1 | Acyl carrier protein, mitochondrial OS=Homo sapiens OX=9606 GN=NDUFAB1 PE=1 SV=3 | 4 |
| GDI2 | Rab GDP dissociation inhibitor beta OS=Homo sapiens OX=9606 GN=GDI2 PE=1 SV=2 | 37 |
| ITPR1 | Inositol 1,4,5-trisphosphate receptor type 1 OS=Homo sapiens OX=9606 GN=ITPR1 PE=1 SV=3 | 54 |
| SCP2 | Sterol carrier protein 2 OS=Homo sapiens OX=9606 GN=SCP2 PE=1 SV=2 | 18 |
| CAMSAP1 | Calmodulin-regulated spectrin-associated protein 1 OS=Homo sapiens OX=9606 GN=CAMSAP1 PE=1 SV=2 | 16 |
| PRDX3 | Thioredoxin-dependent peroxide reductase, mitochondrial OS=Homo sapiens OX=9606 GN=PRDX3 PE=1 SV=3 | 10 |
| OXSR1 | Serine/threonine-protein kinase OSR1 OS=Homo sapiens OX=9606 GN=OXSR1 PE=1 SV=1 | 10 |
| MRPL19 | 39S ribosomal protein L19, mitochondrial OS=Homo sapiens OX=9606 GN=MRPL19 PE=1 SV=2 | 2 |
| PABPC1L | Polyadenylate-binding protein 1-like OS=Homo sapiens OX=9606 GN=PABPC1L PE=2 SV=1 | 23 |
| HSP90AA1 | Heat shock protein HSP 90-alpha OS=Homo sapiens OX=9606 GN=HSP90AA1 PE=1 SV=5 | 59 |
| FBXW12 | F-box/WD repeat-containing protein 12 OS=Homo sapiens OX=9606 GN=FBXW12 PE=2 SV=2 | 5 |
| OSTF1 | Osteoclast-stimulating factor 1 OS=Homo sapiens OX=9606 GN=OSTF1 PE=1 SV=2 | 11 |
| OLA1 | Obg-like ATPase 1 OS=Homo sapiens OX=9606 GN=OLA1 PE=1 SV=2 | 9 |
| ANXA7 | Annexin A7 OS=Homo sapiens OX=9606 GN=ANXA7 PE=1 SV=3 | 26 |
| SRM | Spermidine synthase OS=Homo sapiens OX=9606 GN=SRM PE=1 SV=1 | 14 |
| PDCD6 | Programmed cell death protein 6 OS=Homo sapiens OX=9606 GN=PDCD6 PE=1 SV=1 | 6 |
| EEF1D | Elongation factor 1-delta OS=Homo sapiens OX=9606 GN=EEF1D PE=1 SV=5 | 15 |
| NSFL1C | NSFL1 cofactor p47 OS=Homo sapiens OX=9606 GN=NSFL1C PE=1 SV=2 | 16 |
| PPME1 | Protein phosphatase methylesterase 1 OS=Homo sapiens OX=9606 GN=PPME1 PE=1 SV=3 | 8 |
| DDI2 | Protein DDI1 homolog 2 OS=Homo sapiens OX=9606 GN=DDI2 PE=1 SV=1 | 4 |
| F3 | Tissue factor OS=Homo sapiens OX=9606 GN=F3 PE=1 SV=1 | 3 |
| CRELD2 | Protein disulfide isomerase CRELD2 OS=Homo sapiens OX=9606 GN=CRELD2 PE=1 SV=1 | 6 |
| TMX1 | Thioredoxin-related transmembrane protein 1 OS=Homo sapiens OX=9606 GN=TMX1 PE=1 SV=1 | 7 |
| NUDCD2 | NudC domain-containing protein 2 OS=Homo sapiens OX=9606 GN=NUDCD2 PE=1 SV=1 | 6 |
| NUDT2 | Bis(5'-nucleosyl)-tetraphosphatase [asymmetrical] OS=Homo sapiens OX=9606 GN=NUDT2 PE=1 SV=3 | 5 |
| PWP1 | Periodic tryptophan protein 1 homolog OS=Homo sapiens OX=9606 GN=PWP1 PE=1 SV=1 | 3 |
| CCT7 | T-complex protein 1 subunit eta OS=Homo sapiens OX=9606 GN=CCT7 PE=1 SV=2 | 40 |
| SUCLA2 | Succinate--CoA ligase [ADP-forming] subunit beta, mitochondrial OS=Homo sapiens OX=9606 GN=SUCLA2 PE=1 SV=3 | 21 |
| COBLL1 | Cordon-bleu protein-like 1 OS=Homo sapiens OX=9606 GN=COBLL1 PE=1 SV=3 | 2 |
| MTMR14 | Myotubularin-related protein 14 OS=Homo sapiens OX=9606 GN=MTMR14 PE=1 SV=2 | 12 |
| RPN2 | Dolichyl-diphosphooligosaccharide--protein glycosyltransferase subunit 2 OS=Homo sapiens OX=9606 GN=RPN2 PE=1 SV=3 | 11 |
| PHB1 | Prohibitin 1 OS=Homo sapiens OX=9606 GN=PHB1 PE=1 SV=1 | 21 |
| CLTB | Clathrin light chain B OS=Homo sapiens OX=9606 GN=CLTB PE=1 SV=1 | 9 |
| NUP88 | Nuclear pore complex protein Nup88 OS=Homo sapiens OX=9606 GN=NUP88 PE=1 SV=2 | 2 |
| PFKL | ATP-dependent 6-phosphofructokinase, liver type OS=Homo sapiens OX=9606 GN=PFKL PE=1 SV=6 | 25 |
| SMARCA2 | Probable global transcription activator SNF2L2 OS=Homo sapiens OX=9606 GN=SMARCA2 PE=1 SV=2 | 5 |
| KTN1 | Kinectin OS=Homo sapiens OX=9606 GN=KTN1 PE=1 SV=1 | 58 |
| GFPT1 | Glutamine--fructose-6-phosphate aminotransferase [isomerizing] 1 OS=Homo sapiens OX=9606 GN=GFPT1 PE=1 SV=3 | 12 |
| YWHAB | 14-3-3 protein beta/alpha OS=Homo sapiens OX=9606 GN=YWHAB PE=1 SV=3 | 7 |
| TYSND1 | Peroxisomal leader peptide-processing protease OS=Homo sapiens OX=9606 GN=TYSND1 PE=1 SV=3 | 2 |
| VPS29 | Vacuolar protein sorting-associated protein 29 OS=Homo sapiens OX=9606 GN=VPS29 PE=1 SV=1 | 10 |
| STXBP6 | Syntaxin-binding protein 6 OS=Homo sapiens OX=9606 GN=STXBP6 PE=1 SV=2 | 4 |
| PSTPIP1 | Proline-serine-threonine phosphatase-interacting protein 1 OS=Homo sapiens OX=9606 GN=PSTPIP1 PE=1 SV=1 | 1 |
| MRPL58 | Peptidyl-tRNA hydrolase ICT1, mitochondrial OS=Homo sapiens OX=9606 GN=MRPL58 PE=1 SV=1 | 5 |
| CRELD1 | Protein disulfide isomerase CRELD1 OS=Homo sapiens OX=9606 GN=CRELD1 PE=1 SV=3 | 3 |
| MRPL12 | 39S ribosomal protein L12, mitochondrial OS=Homo sapiens OX=9606 GN=MRPL12 PE=1 SV=2 | 2 |
| SRSF6 | Serine/arginine-rich splicing factor 6 OS=Homo sapiens OX=9606 GN=SRSF6 PE=1 SV=2 | 3 |
| CNDP2 | Cytosolic non-specific dipeptidase OS=Homo sapiens OX=9606 GN=CNDP2 PE=1 SV=2 | 29 |
| EXOSC4 | Exosome complex component RRP41 OS=Homo sapiens OX=9606 GN=EXOSC4 PE=1 SV=3 | 1 |
| PRKAR2A | cAMP-dependent protein kinase type II-alpha regulatory subunit OS=Homo sapiens OX=9606 GN=PRKAR2A PE=1 SV=2 | 19 |
| GSS | Glutathione synthetase OS=Homo sapiens OX=9606 GN=GSS PE=1 SV=1 | 26 |
| PIGK | GPI-anchor transamidase OS=Homo sapiens OX=9606 GN=PIGK PE=1 SV=2 | 4 |
| DCLK2 | Serine/threonine-protein kinase DCLK2 OS=Homo sapiens OX=9606 GN=DCLK2 PE=1 SV=4 | 5 |
| ARHGAP17 | Rho GTPase-activating protein 17 OS=Homo sapiens OX=9606 GN=ARHGAP17 PE=1 SV=1 | 15 |
| NIPSNAP3A | Protein NipSnap homolog 3A OS=Homo sapiens OX=9606 GN=NIPSNAP3A PE=1 SV=2 | 20 |
| ERLEC1 | Endoplasmic reticulum lectin 1 OS=Homo sapiens OX=9606 GN=ERLEC1 PE=1 SV=1 | 22 |
| GOT1 | Aspartate aminotransferase, cytoplasmic OS=Homo sapiens OX=9606 GN=GOT1 PE=1 SV=3 | 19 |
| UGGT1 | UDP-glucose:glycoprotein glucosyltransferase 1 OS=Homo sapiens OX=9606 GN=UGGT1 PE=1 SV=3 | 48 |
| GLRX3 | Glutaredoxin-3 OS=Homo sapiens OX=9606 GN=GLRX3 PE=1 SV=2 | 9 |
| ACTG1 | Actin, cytoplasmic 2 OS=Homo sapiens OX=9606 GN=ACTG1 PE=1 SV=1 | 33 |
| NUCB2 | Nucleobindin-2 OS=Homo sapiens OX=9606 GN=NUCB2 PE=1 SV=3 | 15 |
| TBCA | Tubulin-specific chaperone A OS=Homo sapiens OX=9606 GN=TBCA PE=1 SV=3 | 7 |
| PEA15 | Astrocytic phosphoprotein PEA-15 OS=Homo sapiens OX=9606 GN=PEA15 PE=1 SV=2 | 4 |
| MRPL40 | 39S ribosomal protein L40, mitochondrial OS=Homo sapiens OX=9606 GN=MRPL40 PE=1 SV=1 | 3 |
| DCUN1D1 | DCN1-like protein 1 OS=Homo sapiens OX=9606 GN=DCUN1D1 PE=1 SV=1 | 7 |
| SCCPDH | Saccharopine dehydrogenase-like oxidoreductase OS=Homo sapiens OX=9606 GN=SCCPDH PE=1 SV=1 | 14 |
| CD55 | Complement decay-accelerating factor OS=Homo sapiens OX=9606 GN=CD55 PE=1 SV=4 | 4 |
| ATP2C1 | Calcium-transporting ATPase type 2C member 1 OS=Homo sapiens OX=9606 GN=ATP2C1 PE=1 SV=3 | 10 |
| LRRC40 | Leucine-rich repeat-containing protein 40 OS=Homo sapiens OX=9606 GN=LRRC40 PE=1 SV=1 | 32 |
| PPM1B | Protein phosphatase 1B OS=Homo sapiens OX=9606 GN=PPM1B PE=1 SV=1 | 11 |
| ARHGAP12 | Rho GTPase-activating protein 12 OS=Homo sapiens OX=9606 GN=ARHGAP12 PE=1 SV=1 | 5 |
| HSPH1 | Heat shock protein 105 kDa OS=Homo sapiens OX=9606 GN=HSPH1 PE=1 SV=1 | 34 |
| AGFG1 | Arf-GAP domain and FG repeat-containing protein 1 OS=Homo sapiens OX=9606 GN=AGFG1 PE=1 SV=2 | 6 |
| PARG | Poly(ADP-ribose) glycohydrolase OS=Homo sapiens OX=9606 GN=PARG PE=1 SV=1 | 1 |
| HNRNPF | Heterogeneous nuclear ribonucleoprotein F OS=Homo sapiens OX=9606 GN=HNRNPF PE=1 SV=3 | 11 |
| RPL3 | 60S ribosomal protein L3 OS=Homo sapiens OX=9606 GN=RPL3 PE=1 SV=2 | 10 |
| PSMD13 | 26S proteasome non-ATPase regulatory subunit 13 OS=Homo sapiens OX=9606 GN=PSMD13 PE=1 SV=2 | 18 |
| OSBP | Oxysterol-binding protein 1 OS=Homo sapiens OX=9606 GN=OSBP PE=1 SV=1 | 20 |
| CPT2 | Carnitine O-palmitoyltransferase 2, mitochondrial OS=Homo sapiens OX=9606 GN=CPT2 PE=1 SV=2 | 13 |
| RPS19 | 40S ribosomal protein S19 OS=Homo sapiens OX=9606 GN=RPS19 PE=1 SV=2 | 9 |
| SNRPE | Small nuclear ribonucleoprotein E OS=Homo sapiens OX=9606 GN=SNRPE PE=1 SV=1 | 2 |
| PSMD11 | 26S proteasome non-ATPase regulatory subunit 11 OS=Homo sapiens OX=9606 GN=PSMD11 PE=1 SV=3 | 23 |
| ANAPC1 | Anaphase-promoting complex subunit 1 OS=Homo sapiens OX=9606 GN=ANAPC1 PE=1 SV=1 | 1 |
| MYO5B | Unconventional myosin-Vb OS=Homo sapiens OX=9606 GN=MYO5B PE=1 SV=3 | 12 |
| NDUFAF4 | NADH dehydrogenase [ubiquinone] 1 alpha subcomplex assembly factor 4 OS=Homo sapiens OX=9606 GN=NDUFAF4 PE=1 SV=1 | 6 |
| PIH1D1 | PIH1 domain-containing protein 1 OS=Homo sapiens OX=9606 GN=PIH1D1 PE=1 SV=1 | 4 |
| FKBP3 | Peptidyl-prolyl cis-trans isomerase FKBP3 OS=Homo sapiens OX=9606 GN=FKBP3 PE=1 SV=1 | 9 |
| SEL1L | Protein sel-1 homolog 1 OS=Homo sapiens OX=9606 GN=SEL1L PE=1 SV=3 | 21 |
| MYH11 | Myosin-11 OS=Homo sapiens OX=9606 GN=MYH11 PE=1 SV=3 | 38 |
| RPS16 | 40S ribosomal protein S16 OS=Homo sapiens OX=9606 GN=RPS16 PE=1 SV=2 | 4 |
| ETFA | Electron transfer flavoprotein subunit alpha, mitochondrial OS=Homo sapiens OX=9606 GN=ETFA PE=1 SV=1 | 17 |
| CTNNB1 | Catenin beta-1 OS=Homo sapiens OX=9606 GN=CTNNB1 PE=1 SV=1 | 13 |
| ITGA3 | Integrin alpha-3 OS=Homo sapiens OX=9606 GN=ITGA3 PE=1 SV=5 | 1 |
| KIF2C | Kinesin-like protein KIF2C OS=Homo sapiens OX=9606 GN=KIF2C PE=1 SV=2 | 19 |
| ATAD1 | Outer mitochondrial transmembrane helix translocase OS=Homo sapiens OX=9606 GN=ATAD1 PE=1 SV=1 | 5 |
| NTPCR | Cancer-related nucleoside-triphosphatase OS=Homo sapiens OX=9606 GN=NTPCR PE=1 SV=1 | 6 |
| GLB1 | Beta-galactosidase OS=Homo sapiens OX=9606 GN=GLB1 PE=1 SV=2 | 13 |
| FBXW9 | F-box/WD repeat-containing protein 9 OS=Homo sapiens OX=9606 GN=FBXW9 PE=1 SV=3 | 9 |
| SRSF2 | Serine/arginine-rich splicing factor 2 OS=Homo sapiens OX=9606 GN=SRSF2 PE=1 SV=4 | 2 |
| PDPR | Pyruvate dehydrogenase phosphatase regulatory subunit, mitochondrial OS=Homo sapiens OX=9606 GN=PDPR PE=1 SV=2 | 17 |
| COMMD10 | COMM domain-containing protein 10 OS=Homo sapiens OX=9606 GN=COMMD10 PE=1 SV=1 | 4 |
| GIPC1 | PDZ domain-containing protein GIPC1 OS=Homo sapiens OX=9606 GN=GIPC1 PE=1 SV=2 | 7 |
| EEF1A1 | Elongation factor 1-alpha 1 OS=Homo sapiens OX=9606 GN=EEF1A1 PE=1 SV=1 | 12 |
| APMAP | Adipocyte plasma membrane-associated protein OS=Homo sapiens OX=9606 GN=APMAP PE=1 SV=2 | 12 |
| RACGAP1 | Rac GTPase-activating protein 1 OS=Homo sapiens OX=9606 GN=RACGAP1 PE=1 SV=1 | 4 |
| TCOF1 | Treacle protein OS=Homo sapiens OX=9606 GN=TCOF1 PE=1 SV=3 | 7 |
| CRK | Adapter molecule crk OS=Homo sapiens OX=9606 GN=CRK PE=1 SV=2 | 14 |
| SARS2 | Serine--tRNA ligase, mitochondrial OS=Homo sapiens OX=9606 GN=SARS2 PE=1 SV=1 | 25 |
| MMAA | Methylmalonic aciduria type A protein, mitochondrial OS=Homo sapiens OX=9606 GN=MMAA PE=1 SV=1 | 8 |
| IVD | Isovaleryl-CoA dehydrogenase, mitochondrial OS=Homo sapiens OX=9606 GN=IVD PE=1 SV=2 | 15 |
| ACE | Angiotensin-converting enzyme OS=Homo sapiens OX=9606 GN=ACE PE=1 SV=1 | 3 |
| COPS5 | COP9 signalosome complex subunit 5 OS=Homo sapiens OX=9606 GN=COPS5 PE=1 SV=4 | 15 |
| HYOU1 | Hypoxia up-regulated protein 1 OS=Homo sapiens OX=9606 GN=HYOU1 PE=1 SV=1 | 44 |
| DLD | Dihydrolipoyl dehydrogenase, mitochondrial OS=Homo sapiens OX=9606 GN=DLD PE=1 SV=2 | 26 |
| ADH5 | Alcohol dehydrogenase class-3 OS=Homo sapiens OX=9606 GN=ADH5 PE=1 SV=4 | 12 |
| EIF5 | Eukaryotic translation initiation factor 5 OS=Homo sapiens OX=9606 GN=EIF5 PE=1 SV=2 | 16 |
| HYI | Putative hydroxypyruvate isomerase OS=Homo sapiens OX=9606 GN=HYI PE=1 SV=2 | 10 |
| HENMT1 | Small RNA 2'-O-methyltransferase OS=Homo sapiens OX=9606 GN=HENMT1 PE=1 SV=1 | 8 |
| CD63 | CD63 antigen OS=Homo sapiens OX=9606 GN=CD63 PE=1 SV=2 | 2 |
| GPHA2 | Glycoprotein hormone alpha-2 OS=Homo sapiens OX=9606 GN=GPHA2 PE=1 SV=1 | 1 |
| UCHL3 | Ubiquitin carboxyl-terminal hydrolase isozyme L3 OS=Homo sapiens OX=9606 GN=UCHL3 PE=1 SV=1 | 14 |
| KIF1A | Kinesin-like protein KIF1A OS=Homo sapiens OX=9606 GN=KIF1A PE=1 SV=2 | 14 |
| ADA2 | Adenosine deaminase 2 OS=Homo sapiens OX=9606 GN=ADA2 PE=1 SV=2 | 1 |
| AKAP1 | A-kinase anchor protein 1, mitochondrial OS=Homo sapiens OX=9606 GN=AKAP1 PE=1 SV=1 | 20 |
| HEXB | Beta-hexosaminidase subunit beta OS=Homo sapiens OX=9606 GN=HEXB PE=1 SV=4 | 16 |
| LAD1 | Ladinin-1 OS=Homo sapiens OX=9606 GN=LAD1 PE=1 SV=2 | 28 |
| KLHL7 | Kelch-like protein 7 OS=Homo sapiens OX=9606 GN=KLHL7 PE=1 SV=2 | 6 |
| SEPTIN2 | Septin-2 OS=Homo sapiens OX=9606 GN=SEPTIN2 PE=1 SV=1 | 7 |
| ATAD3B | ATPase family AAA domain-containing protein 3B OS=Homo sapiens OX=9606 GN=ATAD3B PE=1 SV=1 | 2 |
| PIP4K2C | Phosphatidylinositol 5-phosphate 4-kinase type-2 gamma OS=Homo sapiens OX=9606 GN=PIP4K2C PE=1 SV=3 | 17 |
| MAPRE3 | Microtubule-associated protein RP/EB family member 3 OS=Homo sapiens OX=9606 GN=MAPRE3 PE=1 SV=1 | 8 |
| ABHD14B | Putative protein-lysine deacylase ABHD14B OS=Homo sapiens OX=9606 GN=ABHD14B PE=1 SV=1 | 7 |
| ORC1 | Origin recognition complex subunit 1 OS=Homo sapiens OX=9606 GN=ORC1 PE=1 SV=2 | 1 |
| PPT2 | Lysosomal thioesterase PPT2 OS=Homo sapiens OX=9606 GN=PPT2 PE=1 SV=4 | 1 |
| SERPINA1 | Alpha-1-antitrypsin OS=Homo sapiens OX=9606 GN=SERPINA1 PE=1 SV=3 | 20 |
| PSMD14 | 26S proteasome non-ATPase regulatory subunit 14 OS=Homo sapiens OX=9606 GN=PSMD14 PE=1 SV=1 | 9 |
| GLO1 | Lactoylglutathione lyase OS=Homo sapiens OX=9606 GN=GLO1 PE=1 SV=4 | 12 |
| MISP3 | Uncharacterized protein MISP3 OS=Homo sapiens OX=9606 GN=MISP3 PE=2 SV=4 | 2 |
| SEC62 | Translocation protein SEC62 OS=Homo sapiens OX=9606 GN=SEC62 PE=1 SV=1 | 2 |
| NAMPT | Nicotinamide phosphoribosyltransferase OS=Homo sapiens OX=9606 GN=NAMPT PE=1 SV=1 | 5 |
| HDHD2 | Haloacid dehalogenase-like hydrolase domain-containing protein 2 OS=Homo sapiens OX=9606 GN=HDHD2 PE=1 SV=1 | 3 |
| AK4 | Adenylate kinase 4, mitochondrial OS=Homo sapiens OX=9606 GN=AK4 PE=1 SV=1 | 15 |
| RMDN3 | Regulator of microtubule dynamics protein 3 OS=Homo sapiens OX=9606 GN=RMDN3 PE=1 SV=2 | 22 |
| SIRT5 | NAD-dependent protein deacylase sirtuin-5, mitochondrial OS=Homo sapiens OX=9606 GN=SIRT5 PE=1 SV=2 | 5 |
| TBRG4 | FAST kinase domain-containing protein 4 OS=Homo sapiens OX=9606 GN=TBRG4 PE=1 SV=1 | 6 |
| DDX4 | Probable ATP-dependent RNA helicase DDX4 OS=Homo sapiens OX=9606 GN=DDX4 PE=1 SV=2 | 28 |
| CAB39 | Calcium-binding protein 39 OS=Homo sapiens OX=9606 GN=CAB39 PE=1 SV=1 | 13 |
| GTF2I | General transcription factor II-I OS=Homo sapiens OX=9606 GN=GTF2I PE=1 SV=2 | 4 |
| EPHX2 | Bifunctional epoxide hydrolase 2 OS=Homo sapiens OX=9606 GN=EPHX2 PE=1 SV=2 | 17 |
| ANKRD13A | Ankyrin repeat domain-containing protein 13A OS=Homo sapiens OX=9606 GN=ANKRD13A PE=1 SV=3 | 7 |
| TJAP1 | Tight junction-associated protein 1 OS=Homo sapiens OX=9606 GN=TJAP1 PE=1 SV=1 | 8 |
| AP3M2 | AP-3 complex subunit mu-2 OS=Homo sapiens OX=9606 GN=AP3M2 PE=1 SV=1 | 5 |
| POLR2H | DNA-directed RNA polymerases I, II, and III subunit RPABC3 OS=Homo sapiens OX=9606 GN=POLR2H PE=1 SV=4 | 3 |
| MCM5 | DNA replication licensing factor MCM5 OS=Homo sapiens OX=9606 GN=MCM5 PE=1 SV=5 | 3 |
| RPS3A | 40S ribosomal protein S3a OS=Homo sapiens OX=9606 GN=RPS3A PE=1 SV=2 | 14 |
| MTOR | Serine/threonine-protein kinase mTOR OS=Homo sapiens OX=9606 GN=MTOR PE=1 SV=1 | 15 |
| STARD5 | StAR-related lipid transfer protein 5 OS=Homo sapiens OX=9606 GN=STARD5 PE=1 SV=2 | 1 |
| DBNL | Drebrin-like protein OS=Homo sapiens OX=9606 GN=DBNL PE=1 SV=1 | 3 |
| DDX43 | Probable ATP-dependent RNA helicase DDX43 OS=Homo sapiens OX=9606 GN=DDX43 PE=1 SV=2 | 7 |
| LSR | Lipolysis-stimulated lipoprotein receptor OS=Homo sapiens OX=9606 GN=LSR PE=1 SV=4 | 7 |
| MYL12B | Myosin regulatory light chain 12B OS=Homo sapiens OX=9606 GN=MYL12B PE=1 SV=2 | 7 |
| PDCD6IP | Programmed cell death 6-interacting protein OS=Homo sapiens OX=9606 GN=PDCD6IP PE=1 SV=1 | 39 |
| STK39 | STE20/SPS1-related proline-alanine-rich protein kinase OS=Homo sapiens OX=9606 GN=STK39 PE=1 SV=3 | 6 |
| CKAP5 | Cytoskeleton-associated protein 5 OS=Homo sapiens OX=9606 GN=CKAP5 PE=1 SV=3 | 64 |
| ADSL | Adenylosuccinate lyase OS=Homo sapiens OX=9606 GN=ADSL PE=1 SV=2 | 24 |
| MYO1B | Unconventional myosin-Ib OS=Homo sapiens OX=9606 GN=MYO1B PE=1 SV=3 | 26 |
| EDC4 | Enhancer of mRNA-decapping protein 4 OS=Homo sapiens OX=9606 GN=EDC4 PE=1 SV=1 | 7 |
| MRPS11 | 28S ribosomal protein S11, mitochondrial OS=Homo sapiens OX=9606 GN=MRPS11 PE=1 SV=2 | 1 |
| SEPTIN8 | Septin-8 OS=Homo sapiens OX=9606 GN=SEPTIN8 PE=1 SV=4 | 5 |
| NBAS | NBAS subunit of NRZ tethering complex OS=Homo sapiens OX=9606 GN=NBAS PE=1 SV=2 | 12 |
| PDCD5 | Programmed cell death protein 5 OS=Homo sapiens OX=9606 GN=PDCD5 PE=1 SV=3 | 11 |
| TRAPPC6B | Trafficking protein particle complex subunit 6B OS=Homo sapiens OX=9606 GN=TRAPPC6B PE=1 SV=1 | 6 |
| FBXO2 | F-box only protein 2 OS=Homo sapiens OX=9606 GN=FBXO2 PE=1 SV=2 | 10 |
| ACTN4 | Alpha-actinin-4 OS=Homo sapiens OX=9606 GN=ACTN4 PE=1 SV=2 | 55 |
| EIF2S1 | Eukaryotic translation initiation factor 2 subunit 1 OS=Homo sapiens OX=9606 GN=EIF2S1 PE=1 SV=3 | 10 |
| VAPB | Vesicle-associated membrane protein-associated protein B/C OS=Homo sapiens OX=9606 GN=VAPB PE=1 SV=3 | 13 |
| RUVBL2 | RuvB-like 2 OS=Homo sapiens OX=9606 GN=RUVBL2 PE=1 SV=3 | 29 |
| HNRNPD | Heterogeneous nuclear ribonucleoprotein D0 OS=Homo sapiens OX=9606 GN=HNRNPD PE=1 SV=1 | 6 |
| RAB18 | Ras-related protein Rab-18 OS=Homo sapiens OX=9606 GN=RAB18 PE=1 SV=1 | 11 |
| RPS18 | 40S ribosomal protein S18 OS=Homo sapiens OX=9606 GN=RPS18 PE=1 SV=3 | 7 |
| RPS27A | Ubiquitin-40S ribosomal protein S27a OS=Homo sapiens OX=9606 GN=RPS27A PE=1 SV=2 | 11 |
| HSPA4 | Heat shock 70 kDa protein 4 OS=Homo sapiens OX=9606 GN=HSPA4 PE=1 SV=4 | 62 |
| PMPCA | Mitochondrial-processing peptidase subunit alpha OS=Homo sapiens OX=9606 GN=PMPCA PE=1 SV=2 | 13 |
| ATP5F1B | ATP synthase subunit beta, mitochondrial OS=Homo sapiens OX=9606 GN=ATP5F1B PE=1 SV=3 | 27 |
| APEH | Acylamino-acid-releasing enzyme OS=Homo sapiens OX=9606 GN=APEH PE=1 SV=4 | 25 |
| ALG13 | Putative bifunctional UDP-N-acetylglucosamine transferase and deubiquitinase ALG13 OS=Homo sapiens OX=9606 GN=ALG13 PE=1 SV=2 | 6 |
| PSMD8 | 26S proteasome non-ATPase regulatory subunit 8 OS=Homo sapiens OX=9606 GN=PSMD8 PE=1 SV=2 | 10 |
| HSPA5 | Endoplasmic reticulum chaperone BiP OS=Homo sapiens OX=9606 GN=HSPA5 PE=1 SV=2 | 54 |
| PCBP2 | Poly(rC)-binding protein 2 OS=Homo sapiens OX=9606 GN=PCBP2 PE=1 SV=1 | 8 |
| GORASP2 | Golgi reassembly-stacking protein 2 OS=Homo sapiens OX=9606 GN=GORASP2 PE=1 SV=3 | 4 |
| FLYWCH2 | FLYWCH family member 2 OS=Homo sapiens OX=9606 GN=FLYWCH2 PE=1 SV=1 | 3 |
| DDX1 | ATP-dependent RNA helicase DDX1 OS=Homo sapiens OX=9606 GN=DDX1 PE=1 SV=2 | 26 |
| MAP4 | Microtubule-associated protein 4 OS=Homo sapiens OX=9606 GN=MAP4 PE=1 SV=3 | 34 |
| OXNAD1 | Oxidoreductase NAD-binding domain-containing protein 1 OS=Homo sapiens OX=9606 GN=OXNAD1 PE=1 SV=1 | 7 |
| EPS15L1 | Epidermal growth factor receptor substrate 15-like 1 OS=Homo sapiens OX=9606 GN=EPS15L1 PE=1 SV=1 | 14 |
| PHAF1 | Phagosome assembly factor 1 OS=Homo sapiens OX=9606 GN=PHAF1 PE=1 SV=1 | 1 |
| NAPB | Beta-soluble NSF attachment protein OS=Homo sapiens OX=9606 GN=NAPB PE=1 SV=2 | 12 |
| PSMD1 | 26S proteasome non-ATPase regulatory subunit 1 OS=Homo sapiens OX=9606 GN=PSMD1 PE=1 SV=2 | 31 |
| TMBIM1 | Protein lifeguard 3 OS=Homo sapiens OX=9606 GN=TMBIM1 PE=1 SV=2 | 1 |
| NENF | Neudesin OS=Homo sapiens OX=9606 GN=NENF PE=1 SV=1 | 9 |
| CLPTM1 | Putative lipid scramblase CLPTM1 OS=Homo sapiens OX=9606 GN=CLPTM1 PE=1 SV=1 | 2 |
| SLC25A25 | Calcium-binding mitochondrial carrier protein SCaMC-2 OS=Homo sapiens OX=9606 GN=SLC25A25 PE=1 SV=1 | 10 |
| ACBD6 | Acyl-CoA-binding domain-containing protein 6 OS=Homo sapiens OX=9606 GN=ACBD6 PE=1 SV=1 | 1 |
| AURKA | Aurora kinase A OS=Homo sapiens OX=9606 GN=AURKA PE=1 SV=3 | 24 |
| MGME1 | Mitochondrial genome maintenance exonuclease 1 OS=Homo sapiens OX=9606 GN=MGME1 PE=1 SV=1 | 13 |
| TSTD1 | Thiosulfate:glutathione sulfurtransferase OS=Homo sapiens OX=9606 GN=TSTD1 PE=1 SV=3 | 7 |
| GRPEL2 | GrpE protein homolog 2, mitochondrial OS=Homo sapiens OX=9606 GN=GRPEL2 PE=1 SV=1 | 4 |
| APRT | Adenine phosphoribosyltransferase OS=Homo sapiens OX=9606 GN=APRT PE=1 SV=2 | 6 |
| ANOS1 | Anosmin-1 OS=Homo sapiens OX=9606 GN=ANOS1 PE=1 SV=3 | 4 |
| TRIP13 | Pachytene checkpoint protein 2 homolog OS=Homo sapiens OX=9606 GN=TRIP13 PE=1 SV=2 | 8 |
| ACAN | Aggrecan core protein OS=Homo sapiens OX=9606 GN=ACAN PE=1 SV=3 | 4 |
| MPC2 | Mitochondrial pyruvate carrier 2 OS=Homo sapiens OX=9606 GN=MPC2 PE=1 SV=1 | 1 |
| PFAS | Phosphoribosylformylglycinamidine synthase OS=Homo sapiens OX=9606 GN=PFAS PE=1 SV=4 | 13 |
| PTPA | Serine/threonine-protein phosphatase 2A activator OS=Homo sapiens OX=9606 GN=PTPA PE=1 SV=3 | 14 |
| FN3KRP | Ketosamine-3-kinase OS=Homo sapiens OX=9606 GN=FN3KRP PE=1 SV=2 | 9 |
| NEDD8 | NEDD8 OS=Homo sapiens OX=9606 GN=NEDD8 PE=1 SV=1 | 4 |
| AIFM1 | Apoptosis-inducing factor 1, mitochondrial OS=Homo sapiens OX=9606 GN=AIFM1 PE=1 SV=1 | 26 |
| PSMD4 | 26S proteasome non-ATPase regulatory subunit 4 OS=Homo sapiens OX=9606 GN=PSMD4 PE=1 SV=1 | 13 |
| OPA1 | Dynamin-like 120 kDa protein, mitochondrial OS=Homo sapiens OX=9606 GN=OPA1 PE=1 SV=3 | 60 |
| SEC31A | Protein transport protein Sec31A OS=Homo sapiens OX=9606 GN=SEC31A PE=1 SV=3 | 6 |
| DCAKD | Dephospho-CoA kinase domain-containing protein OS=Homo sapiens OX=9606 GN=DCAKD PE=1 SV=1 | 2 |
| PDIA3 | Protein disulfide-isomerase A3 OS=Homo sapiens OX=9606 GN=PDIA3 PE=1 SV=4 | 45 |
| SRP9 | Signal recognition particle 9 kDa protein OS=Homo sapiens OX=9606 GN=SRP9 PE=1 SV=2 | 4 |
| TSN | Translin OS=Homo sapiens OX=9606 GN=TSN PE=1 SV=1 | 5 |
| COPE | Coatomer subunit epsilon OS=Homo sapiens OX=9606 GN=COPE PE=1 SV=3 | 7 |
| MYO10 | Unconventional myosin-X OS=Homo sapiens OX=9606 GN=MYO10 PE=1 SV=3 | 25 |
| RHEB | GTP-binding protein Rheb OS=Homo sapiens OX=9606 GN=RHEB PE=1 SV=1 | 4 |
| CRAT | Carnitine O-acetyltransferase OS=Homo sapiens OX=9606 GN=CRAT PE=1 SV=5 | 7 |
| KPNA7 | Importin subunit alpha-8 OS=Homo sapiens OX=9606 GN=KPNA7 PE=1 SV=1 | 13 |
| ELOB | Elongin-B OS=Homo sapiens OX=9606 GN=ELOB PE=1 SV=1 | 5 |
| EHD1 | EH domain-containing protein 1 OS=Homo sapiens OX=9606 GN=EHD1 PE=1 SV=2 | 15 |
| BZW1 | eIF5-mimic protein 2 OS=Homo sapiens OX=9606 GN=BZW1 PE=1 SV=1 | 2 |
| WASHC2C | WASH complex subunit 2C OS=Homo sapiens OX=9606 GN=WASHC2C PE=1 SV=4 | 1 |
| ABCB7 | Iron-sulfur clusters transporter ABCB7, mitochondrial OS=Homo sapiens OX=9606 GN=ABCB7 PE=1 SV=2 | 6 |
| BAG2 | BAG family molecular chaperone regulator 2 OS=Homo sapiens OX=9606 GN=BAG2 PE=1 SV=1 | 2 |
| TRAPPC12 | Trafficking protein particle complex subunit 12 OS=Homo sapiens OX=9606 GN=TRAPPC12 PE=1 SV=3 | 5 |
| KHDC3L | KH domain-containing protein 3 OS=Homo sapiens OX=9606 GN=KHDC3L PE=1 SV=1 | 17 |
| GSTZ1 | Maleylacetoacetate isomerase OS=Homo sapiens OX=9606 GN=GSTZ1 PE=1 SV=3 | 3 |
| ATP1A1 | Sodium/potassium-transporting ATPase subunit alpha-1 OS=Homo sapiens OX=9606 GN=ATP1A1 PE=1 SV=1 | 35 |
| RABL6 | Rab-like protein 6 OS=Homo sapiens OX=9606 GN=RABL6 PE=1 SV=2 | 5 |
| HLA-A | HLA class I histocompatibility antigen, A alpha chain OS=Homo sapiens OX=9606 GN=HLA-A PE=1 SV=2 | 3 |
| UHRF1 | E3 ubiquitin-protein ligase UHRF1 OS=Homo sapiens OX=9606 GN=UHRF1 PE=1 SV=1 | 50 |
| SRPRB | Signal recognition particle receptor subunit beta OS=Homo sapiens OX=9606 GN=SRPRB PE=1 SV=3 | 9 |
| ITGA1 | Integrin alpha-1 OS=Homo sapiens OX=9606 GN=ITGA1 PE=1 SV=2 | 4 |
| TPD52L2 | Tumor protein D54 OS=Homo sapiens OX=9606 GN=TPD52L2 PE=1 SV=2 | 13 |
| GPAT3 | Glycerol-3-phosphate acyltransferase 3 OS=Homo sapiens OX=9606 GN=GPAT3 PE=1 SV=2 | 1 |
| TLE7 | Transducin-like enhancer protein 7 OS=Homo sapiens OX=9606 GN=TLE7 PE=3 SV=1 | 13 |
| VDAC2 | Voltage-dependent anion-selective channel protein 2 OS=Homo sapiens OX=9606 GN=VDAC2 PE=1 SV=2 | 16 |
| BLOC1S1 | Biogenesis of lysosome-related organelles complex 1 subunit 1 OS=Homo sapiens OX=9606 GN=BLOC1S1 PE=1 SV=2 | 1 |
| PAFAH1B3 | Platelet-activating factor acetylhydrolase IB subunit alpha1 OS=Homo sapiens OX=9606 GN=PAFAH1B3 PE=1 SV=1 | 11 |
| ARPC5 | Actin-related protein 2/3 complex subunit 5 OS=Homo sapiens OX=9606 GN=ARPC5 PE=1 SV=3 | 3 |
| TXNL1 | Thioredoxin-like protein 1 OS=Homo sapiens OX=9606 GN=TXNL1 PE=1 SV=3 | 12 |
| ETFB | Electron transfer flavoprotein subunit beta OS=Homo sapiens OX=9606 GN=ETFB PE=1 SV=3 | 21 |
| PHB2 | Prohibitin-2 OS=Homo sapiens OX=9606 GN=PHB2 PE=1 SV=2 | 15 |
| NCLN | BOS complex subunit NCLN OS=Homo sapiens OX=9606 GN=NCLN PE=1 SV=2 | 15 |
| CERT1 | Ceramide transfer protein OS=Homo sapiens OX=9606 GN=CERT1 PE=1 SV=1 | 7 |
| WDSUB1 | WD repeat, SAM and U-box domain-containing protein 1 OS=Homo sapiens OX=9606 GN=WDSUB1 PE=1 SV=4 | 9 |
| OXR1 | Oxidation resistance protein 1 OS=Homo sapiens OX=9606 GN=OXR1 PE=1 SV=2 | 8 |
| PSMD2 | 26S proteasome non-ATPase regulatory subunit 2 OS=Homo sapiens OX=9606 GN=PSMD2 PE=1 SV=3 | 38 |
| RRAGC | Ras-related GTP-binding protein C OS=Homo sapiens OX=9606 GN=RRAGC PE=1 SV=1 | 12 |
| RNH1 | Ribonuclease inhibitor OS=Homo sapiens OX=9606 GN=RNH1 PE=1 SV=2 | 21 |
| FIS1 | Mitochondrial fission 1 protein OS=Homo sapiens OX=9606 GN=FIS1 PE=1 SV=2 | 4 |
| NDUFB7 | NADH dehydrogenase [ubiquinone] 1 beta subcomplex subunit 7 OS=Homo sapiens OX=9606 GN=NDUFB7 PE=1 SV=4 | 3 |
| ALDH9A1 | 4-trimethylaminobutyraldehyde dehydrogenase OS=Homo sapiens OX=9606 GN=ALDH9A1 PE=1 SV=3 | 22 |
| PHEX | Phosphate-regulating neutral endopeptidase PHEX OS=Homo sapiens OX=9606 GN=PHEX PE=1 SV=1 | 19 |
| COX5A | Cytochrome c oxidase subunit 5A, mitochondrial OS=Homo sapiens OX=9606 GN=COX5A PE=1 SV=2 | 3 |
| DYNLL2 | Dynein light chain 2, cytoplasmic OS=Homo sapiens OX=9606 GN=DYNLL2 PE=1 SV=1 | 3 |
| GGH | Gamma-glutamyl hydrolase OS=Homo sapiens OX=9606 GN=GGH PE=1 SV=2 | 12 |
| RTN4IP1 | Reticulon-4-interacting protein 1, mitochondrial OS=Homo sapiens OX=9606 GN=RTN4IP1 PE=1 SV=2 | 10 |
| MRPL47 | 39S ribosomal protein L47, mitochondrial OS=Homo sapiens OX=9606 GN=MRPL47 PE=1 SV=2 | 7 |
| KIFBP | KIF-binding protein OS=Homo sapiens OX=9606 GN=KIFBP PE=1 SV=1 | 14 |
| KLC2 | Kinesin light chain 2 OS=Homo sapiens OX=9606 GN=KLC2 PE=1 SV=1 | 3 |
| FDXR | NADPH:adrenodoxin oxidoreductase, mitochondrial OS=Homo sapiens OX=9606 GN=FDXR PE=1 SV=3 | 29 |
| PDHB | Pyruvate dehydrogenase E1 component subunit beta, mitochondrial OS=Homo sapiens OX=9606 GN=PDHB PE=1 SV=3 | 13 |
| UBA5 | Ubiquitin-like modifier-activating enzyme 5 OS=Homo sapiens OX=9606 GN=UBA5 PE=1 SV=1 | 11 |
| DNAJC9 | DnaJ homolog subfamily C member 9 OS=Homo sapiens OX=9606 GN=DNAJC9 PE=1 SV=1 | 17 |
| COMMD6 | COMM domain-containing protein 6 OS=Homo sapiens OX=9606 GN=COMMD6 PE=1 SV=1 | 4 |
| HPF1 | Histone PARylation factor 1 OS=Homo sapiens OX=9606 GN=HPF1 PE=1 SV=2 | 11 |
| CALCOCO1 | Calcium-binding and coiled-coil domain-containing protein 1 OS=Homo sapiens OX=9606 GN=CALCOCO1 PE=1 SV=2 | 9 |
| SUPV3L1 | ATP-dependent RNA helicase SUPV3L1, mitochondrial OS=Homo sapiens OX=9606 GN=SUPV3L1 PE=1 SV=1 | 14 |
| SUGT1 | Protein SGT1 homolog OS=Homo sapiens OX=9606 GN=SUGT1 PE=1 SV=3 | 28 |
| ADRM1 | Proteasomal ubiquitin receptor ADRM1 OS=Homo sapiens OX=9606 GN=ADRM1 PE=1 SV=2 | 5 |
| CUL1 | Cullin-1 OS=Homo sapiens OX=9606 GN=CUL1 PE=1 SV=2 | 25 |
| VBP1 | Prefoldin subunit 3 OS=Homo sapiens OX=9606 GN=VBP1 PE=1 SV=4 | 6 |
| PSMD9 | 26S proteasome non-ATPase regulatory subunit 9 OS=Homo sapiens OX=9606 GN=PSMD9 PE=1 SV=3 | 7 |
| WDR1 | WD repeat-containing protein 1 OS=Homo sapiens OX=9606 GN=WDR1 PE=1 SV=4 | 32 |
| CHCHD3 | MICOS complex subunit MIC19 OS=Homo sapiens OX=9606 GN=CHCHD3 PE=1 SV=1 | 12 |
| SHC1 | SHC-transforming protein 1 OS=Homo sapiens OX=9606 GN=SHC1 PE=1 SV=4 | 1 |
| ASMTL | Probable bifunctional dTTP/UTP pyrophosphatase/methyltransferase protein OS=Homo sapiens OX=9606 GN=ASMTL PE=1 SV=3 | 25 |
| AKR1A1 | Aldo-keto reductase family 1 member A1 OS=Homo sapiens OX=9606 GN=AKR1A1 PE=1 SV=3 | 13 |
| PSAP | Prosaposin OS=Homo sapiens OX=9606 GN=PSAP PE=1 SV=2 | 12 |
| CD36 | Platelet glycoprotein 4 OS=Homo sapiens OX=9606 GN=CD36 PE=1 SV=2 | 3 |
| NXN | Nucleoredoxin OS=Homo sapiens OX=9606 GN=NXN PE=1 SV=2 | 3 |
| RPS29 | 40S ribosomal protein S29 OS=Homo sapiens OX=9606 GN=RPS29 PE=1 SV=2 | 3 |
| OTUB1 | Ubiquitin thioesterase OTUB1 OS=Homo sapiens OX=9606 GN=OTUB1 PE=1 SV=2 | 11 |
| UBE2V2 | Ubiquitin-conjugating enzyme E2 variant 2 OS=Homo sapiens OX=9606 GN=UBE2V2 PE=1 SV=4 | 10 |
| ARFGAP1 | ADP-ribosylation factor GTPase-activating protein 1 OS=Homo sapiens OX=9606 GN=ARFGAP1 PE=1 SV=2 | 4 |
| ATAD3A | ATPase family AAA domain-containing protein 3A OS=Homo sapiens OX=9606 GN=ATAD3A PE=1 SV=2 | 44 |
| COMMD2 | COMM domain-containing protein 2 OS=Homo sapiens OX=9606 GN=COMMD2 PE=1 SV=2 | 2 |
| RAB27B | Ras-related protein Rab-27B OS=Homo sapiens OX=9606 GN=RAB27B PE=1 SV=4 | 2 |
| FECH | Ferrochelatase, mitochondrial OS=Homo sapiens OX=9606 GN=FECH PE=1 SV=2 | 13 |
| STRBP | Spermatid perinuclear RNA-binding protein OS=Homo sapiens OX=9606 GN=STRBP PE=1 SV=1 | 2 |
| SENP8 | Sentrin-specific protease 8 OS=Homo sapiens OX=9606 GN=SENP8 PE=1 SV=1 | 5 |
| NSF | Vesicle-fusing ATPase OS=Homo sapiens OX=9606 GN=NSF PE=1 SV=3 | 55 |
| TIMM10 | Mitochondrial import inner membrane translocase subunit Tim10 OS=Homo sapiens OX=9606 GN=TIMM10 PE=1 SV=1 | 3 |
| ERP29 | Endoplasmic reticulum resident protein 29 OS=Homo sapiens OX=9606 GN=ERP29 PE=1 SV=4 | 13 |
| TUFM | Elongation factor Tu, mitochondrial OS=Homo sapiens OX=9606 GN=TUFM PE=1 SV=3 | 30 |
| SORBS1 | Sorbin and SH3 domain-containing protein 1 OS=Homo sapiens OX=9606 GN=SORBS1 PE=1 SV=3 | 1 |
| SEC63 | Translocation protein SEC63 homolog OS=Homo sapiens OX=9606 GN=SEC63 PE=1 SV=2 | 15 |
| TPM1 | Tropomyosin alpha-1 chain OS=Homo sapiens OX=9606 GN=TPM1 PE=1 SV=2 | 6 |
| ISCU | Iron-sulfur cluster assembly enzyme ISCU OS=Homo sapiens OX=9606 GN=ISCU PE=1 SV=2 | 9 |
| MACF1 | Microtubule-actin cross-linking factor 1, isoforms 1/2/3/4/5 OS=Homo sapiens OX=9606 GN=MACF1 PE=1 SV=4 | 19 |
| ACTR1A | Alpha-centractin OS=Homo sapiens OX=9606 GN=ACTR1A PE=1 SV=1 | 5 |
| UFC1 | Ubiquitin-fold modifier-conjugating enzyme 1 OS=Homo sapiens OX=9606 GN=UFC1 PE=1 SV=3 | 12 |
| AXL | Tyrosine-protein kinase receptor UFO OS=Homo sapiens OX=9606 GN=AXL PE=1 SV=4 | 8 |
| AKT1S1 | Proline-rich AKT1 substrate 1 OS=Homo sapiens OX=9606 GN=AKT1S1 PE=1 SV=1 | 1 |
| RNF7 | RING-box protein 2 OS=Homo sapiens OX=9606 GN=RNF7 PE=1 SV=1 | 3 |
| CCT2 | T-complex protein 1 subunit beta OS=Homo sapiens OX=9606 GN=CCT2 PE=1 SV=4 | 41 |
| RPS14 | 40S ribosomal protein S14 OS=Homo sapiens OX=9606 GN=RPS14 PE=1 SV=3 | 3 |
| JPT2 | Jupiter microtubule associated homolog 2 OS=Homo sapiens OX=9606 GN=JPT2 PE=1 SV=1 | 8 |
| KMT2C | Histone-lysine N-methyltransferase 2C OS=Homo sapiens OX=9606 GN=KMT2C PE=1 SV=3 | 17 |
| ARL8B | ADP-ribosylation factor-like protein 8B OS=Homo sapiens OX=9606 GN=ARL8B PE=1 SV=1 | 5 |
| GRPEL1 | GrpE protein homolog 1, mitochondrial OS=Homo sapiens OX=9606 GN=GRPEL1 PE=1 SV=2 | 11 |
| RPL21 | 60S ribosomal protein L21 OS=Homo sapiens OX=9606 GN=RPL21 PE=1 SV=2 | 4 |
| CIAO2A | Cytosolic iron-sulfur assembly component 2A OS=Homo sapiens OX=9606 GN=CIAO2A PE=1 SV=1 | 1 |
| P4HA1 | Prolyl 4-hydroxylase subunit alpha-1 OS=Homo sapiens OX=9606 GN=P4HA1 PE=1 SV=2 | 6 |
| AMPD2 | AMP deaminase 2 OS=Homo sapiens OX=9606 GN=AMPD2 PE=1 SV=2 | 20 |
| SLC39A14 | Metal cation symporter ZIP14 OS=Homo sapiens OX=9606 GN=SLC39A14 PE=1 SV=3 | 2 |
| UXS1 | UDP-glucuronic acid decarboxylase 1 OS=Homo sapiens OX=9606 GN=UXS1 PE=1 SV=1 | 4 |
| THOP1 | Thimet oligopeptidase OS=Homo sapiens OX=9606 GN=THOP1 PE=1 SV=2 | 14 |
| RPS21 | 40S ribosomal protein S21 OS=Homo sapiens OX=9606 GN=RPS21 PE=1 SV=1 | 5 |
| EPB41L2 | Band 4.1-like protein 2 OS=Homo sapiens OX=9606 GN=EPB41L2 PE=1 SV=1 | 35 |
| PPCS | Phosphopantothenate--cysteine ligase OS=Homo sapiens OX=9606 GN=PPCS PE=1 SV=2 | 13 |
| REPS2 | RalBP1-associated Eps domain-containing protein 2 OS=Homo sapiens OX=9606 GN=REPS2 PE=1 SV=2 | 3 |
| ATP5PD | ATP synthase subunit d, mitochondrial OS=Homo sapiens OX=9606 GN=ATP5PD PE=1 SV=3 | 10 |
| MAP7D2 | MAP7 domain-containing protein 2 OS=Homo sapiens OX=9606 GN=MAP7D2 PE=1 SV=2 | 19 |
| FXR2 | RNA-binding protein FXR2 OS=Homo sapiens OX=9606 GN=FXR2 PE=1 SV=2 | 6 |
| RAB5B | Ras-related protein Rab-5B OS=Homo sapiens OX=9606 GN=RAB5B PE=1 SV=1 | 11 |
| USO1 | General vesicular transport factor p115 OS=Homo sapiens OX=9606 GN=USO1 PE=1 SV=2 | 21 |
| NUDT5 | ADP-sugar pyrophosphatase OS=Homo sapiens OX=9606 GN=NUDT5 PE=1 SV=1 | 13 |
| ZP2 | Zona pellucida sperm-binding protein 2 OS=Homo sapiens OX=9606 GN=ZP2 PE=1 SV=1 | 41 |
| PIGS | GPI transamidase component PIG-S OS=Homo sapiens OX=9606 GN=PIGS PE=1 SV=3 | 8 |
| LETM1 | Mitochondrial proton/calcium exchanger protein OS=Homo sapiens OX=9606 GN=LETM1 PE=1 SV=1 | 17 |
| TFAM | Transcription factor A, mitochondrial OS=Homo sapiens OX=9606 GN=TFAM PE=1 SV=1 | 9 |
| PDXK | Pyridoxal kinase OS=Homo sapiens OX=9606 GN=PDXK PE=1 SV=1 | 15 |
| PACSIN2 | Protein kinase C and casein kinase substrate in neurons protein 2 OS=Homo sapiens OX=9606 GN=PACSIN2 PE=1 SV=2 | 22 |
| TPT1 | Translationally-controlled tumor protein OS=Homo sapiens OX=9606 GN=TPT1 PE=1 SV=1 | 8 |
| NDUFS2 | NADH dehydrogenase [ubiquinone] iron-sulfur protein 2, mitochondrial OS=Homo sapiens OX=9606 GN=NDUFS2 PE=1 SV=2 | 10 |
| UBE2L3 | Ubiquitin-conjugating enzyme E2 L3 OS=Homo sapiens OX=9606 GN=UBE2L3 PE=1 SV=1 | 8 |
| NDUFV2 | NADH dehydrogenase [ubiquinone] flavoprotein 2, mitochondrial OS=Homo sapiens OX=9606 GN=NDUFV2 PE=1 SV=2 | 7 |
| STIP1 | Stress-induced-phosphoprotein 1 OS=Homo sapiens OX=9606 GN=STIP1 PE=1 SV=1 | 38 |
| PPCDC | Phosphopantothenoylcysteine decarboxylase OS=Homo sapiens OX=9606 GN=PPCDC PE=1 SV=2 | 2 |
| ERC1 | ELKS/Rab6-interacting/CAST family member 1 OS=Homo sapiens OX=9606 GN=ERC1 PE=1 SV=1 | 9 |
| PDCL3 | Phosducin-like protein 3 OS=Homo sapiens OX=9606 GN=PDCL3 PE=1 SV=1 | 13 |
| PIGT | GPI transamidase component PIG-T OS=Homo sapiens OX=9606 GN=PIGT PE=1 SV=1 | 1 |
| VAT1 | Synaptic vesicle membrane protein VAT-1 homolog OS=Homo sapiens OX=9606 GN=VAT1 PE=1 SV=2 | 13 |
| SDK1 | Protein sidekick-1 OS=Homo sapiens OX=9606 GN=SDK1 PE=2 SV=3 | 1 |
| ZFYVE16 | Zinc finger FYVE domain-containing protein 16 OS=Homo sapiens OX=9606 GN=ZFYVE16 PE=1 SV=3 | 1 |
| SLC35B2 | Adenosine 3'-phospho 5'-phosphosulfate transporter 1 OS=Homo sapiens OX=9606 GN=SLC35B2 PE=1 SV=1 | 1 |
| GPT2 | Alanine aminotransferase 2 OS=Homo sapiens OX=9606 GN=GPT2 PE=1 SV=1 | 21 |
| CBR4 | 3-oxoacyl-[acyl-carrier-protein] reductase OS=Homo sapiens OX=9606 GN=CBR4 PE=1 SV=3 | 5 |
| HADH | Hydroxyacyl-coenzyme A dehydrogenase, mitochondrial OS=Homo sapiens OX=9606 GN=HADH PE=1 SV=3 | 14 |
| DDX17 | Probable ATP-dependent RNA helicase DDX17 OS=Homo sapiens OX=9606 GN=DDX17 PE=1 SV=2 | 4 |
| HSDL1 | Inactive hydroxysteroid dehydrogenase-like protein 1 OS=Homo sapiens OX=9606 GN=HSDL1 PE=1 SV=3 | 12 |
| IARS2 | Isoleucine--tRNA ligase, mitochondrial OS=Homo sapiens OX=9606 GN=IARS2 PE=1 SV=2 | 34 |
| STMN1 | Stathmin OS=Homo sapiens OX=9606 GN=STMN1 PE=1 SV=3 | 8 |
| RPN1 | Dolichyl-diphosphooligosaccharide--protein glycosyltransferase subunit 1 OS=Homo sapiens OX=9606 GN=RPN1 PE=1 SV=1 | 30 |
| YRDC | Threonylcarbamoyl-AMP synthase OS=Homo sapiens OX=9606 GN=YRDC PE=1 SV=1 | 5 |
| G3BP2 | Ras GTPase-activating protein-binding protein 2 OS=Homo sapiens OX=9606 GN=G3BP2 PE=1 SV=2 | 12 |
| ACTR2 | Actin-related protein 2 OS=Homo sapiens OX=9606 GN=ACTR2 PE=1 SV=1 | 15 |
| TCP1 | T-complex protein 1 subunit alpha OS=Homo sapiens OX=9606 GN=TCP1 PE=1 SV=1 | 31 |
| GTF2E1 | General transcription factor IIE subunit 1 OS=Homo sapiens OX=9606 GN=GTF2E1 PE=1 SV=2 | 5 |
| ACSL3 | Fatty acid CoA ligase Acsl3 OS=Homo sapiens OX=9606 GN=ACSL3 PE=1 SV=3 | 20 |
| UMPS | Uridine 5'-monophosphate synthase OS=Homo sapiens OX=9606 GN=UMPS PE=1 SV=1 | 20 |
| RAB5C | Ras-related protein Rab-5C OS=Homo sapiens OX=9606 GN=RAB5C PE=1 SV=2 | 3 |
| GLRX | Glutaredoxin-1 OS=Homo sapiens OX=9606 GN=GLRX PE=1 SV=2 | 6 |
| RPL7 | 60S ribosomal protein L7 OS=Homo sapiens OX=9606 GN=RPL7 PE=1 SV=1 | 9 |
| ENAH | Protein enabled homolog OS=Homo sapiens OX=9606 GN=ENAH PE=1 SV=2 | 5 |
| CLIP1 | CAP-Gly domain-containing linker protein 1 OS=Homo sapiens OX=9606 GN=CLIP1 PE=1 SV=2 | 8 |
| NUDC | Nuclear migration protein nudC OS=Homo sapiens OX=9606 GN=NUDC PE=1 SV=1 | 20 |
| SASS6 | Spindle assembly abnormal protein 6 homolog OS=Homo sapiens OX=9606 GN=SASS6 PE=1 SV=1 | 7 |
| GSN | Gelsolin OS=Homo sapiens OX=9606 GN=GSN PE=1 SV=1 | 33 |
| GSKIP | GSK3B-interacting protein OS=Homo sapiens OX=9606 GN=GSKIP PE=1 SV=2 | 3 |
| CNOT1 | CCR4-NOT transcription complex subunit 1 OS=Homo sapiens OX=9606 GN=CNOT1 PE=1 SV=2 | 9 |
| FADD | FAS-associated death domain protein OS=Homo sapiens OX=9606 GN=FADD PE=1 SV=1 | 1 |
| MDH1 | Malate dehydrogenase, cytoplasmic OS=Homo sapiens OX=9606 GN=MDH1 PE=1 SV=4 | 17 |
| GALT | Galactose-1-phosphate uridylyltransferase OS=Homo sapiens OX=9606 GN=GALT PE=1 SV=3 | 7 |
| AP3S2 | AP-3 complex subunit sigma-2 OS=Homo sapiens OX=9606 GN=AP3S2 PE=2 SV=1 | 5 |
| ADPRS | ADP-ribosylhydrolase ARH3 OS=Homo sapiens OX=9606 GN=ADPRS PE=1 SV=1 | 8 |
| YBX2 | Y-box-binding protein 2 OS=Homo sapiens OX=9606 GN=YBX2 PE=1 SV=2 | 21 |
| CGREF1 | Cell growth regulator with EF hand domain protein 1 OS=Homo sapiens OX=9606 GN=CGREF1 PE=1 SV=3 | 2 |
| STOM | Stomatin OS=Homo sapiens OX=9606 GN=STOM PE=1 SV=3 | 13 |
| ATP6V0D1 | V-type proton ATPase subunit d 1 OS=Homo sapiens OX=9606 GN=ATP6V0D1 PE=1 SV=1 | 9 |
| GNPNAT1 | Glucosamine 6-phosphate N-acetyltransferase OS=Homo sapiens OX=9606 GN=GNPNAT1 PE=1 SV=1 | 3 |
| CHMP1B | Charged multivesicular body protein 1b OS=Homo sapiens OX=9606 GN=CHMP1B PE=1 SV=1 | 2 |
| MZB1 | Marginal zone B- and B1-cell-specific protein OS=Homo sapiens OX=9606 GN=MZB1 PE=1 SV=1 | 1 |
| PSMB2 | Proteasome subunit beta type-2 OS=Homo sapiens OX=9606 GN=PSMB2 PE=1 SV=1 | 5 |
| PPP1CB | Serine/threonine-protein phosphatase PP1-beta catalytic subunit OS=Homo sapiens OX=9606 GN=PPP1CB PE=1 SV=3 | 3 |
| BCAP31 | B-cell receptor-associated protein 31 OS=Homo sapiens OX=9606 GN=BCAP31 PE=1 SV=3 | 14 |
| IDH2 | Isocitrate dehydrogenase [NADP], mitochondrial OS=Homo sapiens OX=9606 GN=IDH2 PE=1 SV=2 | 18 |
| CENPE | Centromere-associated protein E OS=Homo sapiens OX=9606 GN=CENPE PE=1 SV=2 | 38 |
| GMPPB | Mannose-1-phosphate guanyltransferase beta OS=Homo sapiens OX=9606 GN=GMPPB PE=1 SV=2 | 7 |
| GAA | Lysosomal alpha-glucosidase OS=Homo sapiens OX=9606 GN=GAA PE=1 SV=4 | 13 |
| COX4I1 | Cytochrome c oxidase subunit 4 isoform 1, mitochondrial OS=Homo sapiens OX=9606 GN=COX4I1 PE=1 SV=1 | 10 |
| DHX15 | ATP-dependent RNA helicase DHX15 OS=Homo sapiens OX=9606 GN=DHX15 PE=1 SV=2 | 4 |
| DDI1 | Protein DDI1 homolog 1 OS=Homo sapiens OX=9606 GN=DDI1 PE=1 SV=1 | 1 |
| RAE1 | mRNA export factor RAE1 OS=Homo sapiens OX=9606 GN=RAE1 PE=1 SV=1 | 10 |
| LEMD2 | LEM domain-containing protein 2 OS=Homo sapiens OX=9606 GN=LEMD2 PE=1 SV=1 | 5 |
| TES | Testin OS=Homo sapiens OX=9606 GN=TES PE=1 SV=1 | 5 |
| NUCB1 | Nucleobindin-1 OS=Homo sapiens OX=9606 GN=NUCB1 PE=1 SV=4 | 13 |
| TOM1L1 | TOM1-like protein 1 OS=Homo sapiens OX=9606 GN=TOM1L1 PE=1 SV=2 | 6 |
| GNPDA1 | Glucosamine-6-phosphate isomerase 1 OS=Homo sapiens OX=9606 GN=GNPDA1 PE=1 SV=1 | 10 |
| CCT3 | T-complex protein 1 subunit gamma OS=Homo sapiens OX=9606 GN=CCT3 PE=1 SV=4 | 34 |
| STXBP2 | Syntaxin-binding protein 2 OS=Homo sapiens OX=9606 GN=STXBP2 PE=1 SV=2 | 21 |
| HSP90AB1 | Heat shock protein HSP 90-beta OS=Homo sapiens OX=9606 GN=HSP90AB1 PE=1 SV=4 | 29 |
| PITPNB | Phosphatidylinositol transfer protein beta isoform OS=Homo sapiens OX=9606 GN=PITPNB PE=1 SV=2 | 2 |
| TOMM34 | Mitochondrial import receptor subunit TOM34 OS=Homo sapiens OX=9606 GN=TOMM34 PE=1 SV=2 | 25 |
| IARS1 | Isoleucine--tRNA ligase, cytoplasmic OS=Homo sapiens OX=9606 GN=IARS1 PE=1 SV=2 | 12 |
| GNAI2 | Guanine nucleotide-binding protein G(i) subunit alpha-2 OS=Homo sapiens OX=9606 GN=GNAI2 PE=1 SV=3 | 3 |
| SH3GLB1 | Endophilin-B1 OS=Homo sapiens OX=9606 GN=SH3GLB1 PE=1 SV=1 | 13 |
| NDUFA10 | NADH dehydrogenase [ubiquinone] 1 alpha subcomplex subunit 10, mitochondrial OS=Homo sapiens OX=9606 GN=NDUFA10 PE=1 SV=1 | 7 |
| RDX | Radixin OS=Homo sapiens OX=9606 GN=RDX PE=1 SV=1 | 15 |
| NASP | Nuclear autoantigenic sperm protein OS=Homo sapiens OX=9606 GN=NASP PE=1 SV=2 | 34 |
| SYT2 | Synaptotagmin-2 OS=Homo sapiens OX=9606 GN=SYT2 PE=1 SV=2 | 14 |
| PAK2 | Serine/threonine-protein kinase PAK 2 OS=Homo sapiens OX=9606 GN=PAK2 PE=1 SV=3 | 27 |
| RHOT1 | Mitochondrial Rho GTPase 1 OS=Homo sapiens OX=9606 GN=RHOT1 PE=1 SV=2 | 6 |
| TUBG1 | Tubulin gamma-1 chain OS=Homo sapiens OX=9606 GN=TUBG1 PE=1 SV=2 | 18 |
| BCCIP | BRCA2 and CDKN1A-interacting protein OS=Homo sapiens OX=9606 GN=BCCIP PE=1 SV=1 | 2 |
| DPP3 | Dipeptidyl peptidase 3 OS=Homo sapiens OX=9606 GN=DPP3 PE=1 SV=2 | 28 |
| INF2 | Inverted formin-2 OS=Homo sapiens OX=9606 GN=INF2 PE=1 SV=2 | 3 |
| ITGB1BP1 | Integrin beta-1-binding protein 1 OS=Homo sapiens OX=9606 GN=ITGB1BP1 PE=1 SV=1 | 2 |
| SUCLG1 | Succinate--CoA ligase [ADP/GDP-forming] subunit alpha, mitochondrial OS=Homo sapiens OX=9606 GN=SUCLG1 PE=1 SV=4 | 12 |
| VAPA | Vesicle-associated membrane protein-associated protein A OS=Homo sapiens OX=9606 GN=VAPA PE=1 SV=3 | 9 |
| MTX2 | Metaxin-2 OS=Homo sapiens OX=9606 GN=MTX2 PE=1 SV=1 | 7 |
| LTA4H | Leukotriene A-4 hydrolase OS=Homo sapiens OX=9606 GN=LTA4H PE=1 SV=2 | 22 |
| UFL1 | E3 UFM1-protein ligase 1 OS=Homo sapiens OX=9606 GN=UFL1 PE=1 SV=2 | 8 |
| PRPF8 | Pre-mRNA-processing-splicing factor 8 OS=Homo sapiens OX=9606 GN=PRPF8 PE=1 SV=2 | 4 |
| CRKL | Crk-like protein OS=Homo sapiens OX=9606 GN=CRKL PE=1 SV=1 | 12 |
| DDRGK1 | DDRGK domain-containing protein 1 OS=Homo sapiens OX=9606 GN=DDRGK1 PE=1 SV=2 | 1 |
| EIF2B3 | Translation initiation factor eIF-2B subunit gamma OS=Homo sapiens OX=9606 GN=EIF2B3 PE=1 SV=1 | 8 |
| SLC15A2 | Solute carrier family 15 member 2 OS=Homo sapiens OX=9606 GN=SLC15A2 PE=1 SV=2 | 1 |
| TMEM43 | Transmembrane protein 43 OS=Homo sapiens OX=9606 GN=TMEM43 PE=1 SV=1 | 15 |
| TCL1B | T-cell leukemia/lymphoma protein 1B OS=Homo sapiens OX=9606 GN=TCL1B PE=1 SV=1 | 4 |
| VIPAS39 | Spermatogenesis-defective protein 39 homolog OS=Homo sapiens OX=9606 GN=VIPAS39 PE=1 SV=1 | 3 |
| PEX3 | Peroxisomal biogenesis factor 3 OS=Homo sapiens OX=9606 GN=PEX3 PE=1 SV=1 | 1 |
| SEPHS1 | Selenide, water dikinase 1 OS=Homo sapiens OX=9606 GN=SEPHS1 PE=1 SV=2 | 7 |
| BRCC3 | Lys-63-specific deubiquitinase BRCC36 OS=Homo sapiens OX=9606 GN=BRCC3 PE=1 SV=2 | 1 |
| HNRNPAB | Heterogeneous nuclear ribonucleoprotein A/B OS=Homo sapiens OX=9606 GN=HNRNPAB PE=1 SV=2 | 6 |
| ITGAV | Integrin alpha-V OS=Homo sapiens OX=9606 GN=ITGAV PE=1 SV=2 | 22 |
| NUP93 | Nuclear pore complex protein Nup93 OS=Homo sapiens OX=9606 GN=NUP93 PE=1 SV=2 | 16 |
| PPAT | Amidophosphoribosyltransferase OS=Homo sapiens OX=9606 GN=PPAT PE=1 SV=1 | 20 |
| TRA2B | Transformer-2 protein homolog beta OS=Homo sapiens OX=9606 GN=TRA2B PE=1 SV=1 | 6 |
| PELO | Protein pelota homolog OS=Homo sapiens OX=9606 GN=PELO PE=1 SV=2 | 8 |
| RPL24 | 60S ribosomal protein L24 OS=Homo sapiens OX=9606 GN=RPL24 PE=1 SV=1 | 5 |
| SYAP1 | Synapse-associated protein 1 OS=Homo sapiens OX=9606 GN=SYAP1 PE=1 SV=1 | 13 |
| CNRIP1 | CB1 cannabinoid receptor-interacting protein 1 OS=Homo sapiens OX=9606 GN=CNRIP1 PE=1 SV=1 | 4 |
| RBBP9 | Serine hydrolase RBBP9 OS=Homo sapiens OX=9606 GN=RBBP9 PE=1 SV=2 | 7 |
| APOE | Apolipoprotein E OS=Homo sapiens OX=9606 GN=APOE PE=1 SV=1 | 14 |
| CYFIP1 | Cytoplasmic FMR1-interacting protein 1 OS=Homo sapiens OX=9606 GN=CYFIP1 PE=1 SV=1 | 9 |
| SUN1 | SUN domain-containing protein 1 OS=Homo sapiens OX=9606 GN=SUN1 PE=1 SV=4 | 14 |
| LPCAT1 | Lysophosphatidylcholine acyltransferase 1 OS=Homo sapiens OX=9606 GN=LPCAT1 PE=1 SV=2 | 7 |
| RRP15 | RRP15-like protein OS=Homo sapiens OX=9606 GN=RRP15 PE=1 SV=2 | 1 |
| TOM1L2 | TOM1-like protein 2 OS=Homo sapiens OX=9606 GN=TOM1L2 PE=1 SV=1 | 4 |
| TLN1 | Talin-1 OS=Homo sapiens OX=9606 GN=TLN1 PE=1 SV=3 | 39 |
| RCC2 | Protein RCC2 OS=Homo sapiens OX=9606 GN=RCC2 PE=1 SV=2 | 11 |
| NELFB | Negative elongation factor B OS=Homo sapiens OX=9606 GN=NELFB PE=1 SV=1 | 1 |
| GLUD1 | Glutamate dehydrogenase 1, mitochondrial OS=Homo sapiens OX=9606 GN=GLUD1 PE=1 SV=2 | 17 |
| SCUBE1 | Signal peptide, CUB and EGF-like domain-containing protein 1 OS=Homo sapiens OX=9606 GN=SCUBE1 PE=1 SV=3 | 1 |
| RPS24 | 40S ribosomal protein S24 OS=Homo sapiens OX=9606 GN=RPS24 PE=1 SV=1 | 3 |
| ATPAF1 | ATP synthase mitochondrial F1 complex assembly factor 1 OS=Homo sapiens OX=9606 GN=ATPAF1 PE=1 SV=1 | 10 |
| RBKS | Ribokinase OS=Homo sapiens OX=9606 GN=RBKS PE=1 SV=1 | 7 |
| PSMC6 | 26S proteasome regulatory subunit 10B OS=Homo sapiens OX=9606 GN=PSMC6 PE=1 SV=1 | 23 |
| MDH2 | Malate dehydrogenase, mitochondrial OS=Homo sapiens OX=9606 GN=MDH2 PE=1 SV=3 | 19 |
| C9orf64 | Queuosine salvage protein OS=Homo sapiens OX=9606 GN=C9orf64 PE=1 SV=1 | 5 |
| GOLM1 | Golgi membrane protein 1 OS=Homo sapiens OX=9606 GN=GOLM1 PE=1 SV=1 | 6 |
| TAGLN2 | Transgelin-2 OS=Homo sapiens OX=9606 GN=TAGLN2 PE=1 SV=3 | 13 |
| LACTB | Serine beta-lactamase-like protein LACTB, mitochondrial OS=Homo sapiens OX=9606 GN=LACTB PE=1 SV=2 | 9 |
| PSMD6 | 26S proteasome non-ATPase regulatory subunit 6 OS=Homo sapiens OX=9606 GN=PSMD6 PE=1 SV=1 | 18 |
| HIKESHI | Protein Hikeshi OS=Homo sapiens OX=9606 GN=HIKESHI PE=1 SV=2 | 1 |
| RPS10 | 40S ribosomal protein S10 OS=Homo sapiens OX=9606 GN=RPS10 PE=1 SV=1 | 5 |
| MCM3 | DNA replication licensing factor MCM3 OS=Homo sapiens OX=9606 GN=MCM3 PE=1 SV=3 | 5 |
| NUP205 | Nuclear pore complex protein Nup205 OS=Homo sapiens OX=9606 GN=NUP205 PE=1 SV=3 | 5 |
| CYP2D7 | Putative cytochrome P450 2D7 OS=Homo sapiens OX=9606 GN=CYP2D7 PE=5 SV=1 | 1 |
| GAPVD1 | GTPase-activating protein and VPS9 domain-containing protein 1 OS=Homo sapiens OX=9606 GN=GAPVD1 PE=1 SV=2 | 3 |
| PTCD3 | Pentatricopeptide repeat domain-containing protein 3, mitochondrial OS=Homo sapiens OX=9606 GN=PTCD3 PE=1 SV=3 | 3 |
| RPL27 | 60S ribosomal protein L27 OS=Homo sapiens OX=9606 GN=RPL27 PE=1 SV=2 | 4 |
| SRP72 | Signal recognition particle subunit SRP72 OS=Homo sapiens OX=9606 GN=SRP72 PE=1 SV=3 | 6 |
| MRPL46 | 39S ribosomal protein L46, mitochondrial OS=Homo sapiens OX=9606 GN=MRPL46 PE=1 SV=1 | 5 |
| PFDN2 | Prefoldin subunit 2 OS=Homo sapiens OX=9606 GN=PFDN2 PE=1 SV=1 | 6 |
| EIF4A1 | Eukaryotic initiation factor 4A-I OS=Homo sapiens OX=9606 GN=EIF4A1 PE=1 SV=1 | 22 |
| EIF2A | Eukaryotic translation initiation factor 2A OS=Homo sapiens OX=9606 GN=EIF2A PE=1 SV=3 | 17 |
| PIK3R1 | Phosphatidylinositol 3-kinase regulatory subunit alpha OS=Homo sapiens OX=9606 GN=PIK3R1 PE=1 SV=2 | 6 |
| PIGBOS1 | Protein PIGBOS1 OS=Homo sapiens OX=9606 GN=PIGBOS1 PE=1 SV=1 | 1 |
| RALA | Ras-related protein Ral-A OS=Homo sapiens OX=9606 GN=RALA PE=1 SV=1 | 5 |
| MRPL55 | 39S ribosomal protein L55, mitochondrial OS=Homo sapiens OX=9606 GN=MRPL55 PE=1 SV=1 | 2 |
| HSPA9 | Stress-70 protein, mitochondrial OS=Homo sapiens OX=9606 GN=HSPA9 PE=1 SV=2 | 60 |
| MTHFD1L | Monofunctional C1-tetrahydrofolate synthase, mitochondrial OS=Homo sapiens OX=9606 GN=MTHFD1L PE=1 SV=1 | 1 |
| PDLIM3 | PDZ and LIM domain protein 3 OS=Homo sapiens OX=9606 GN=PDLIM3 PE=1 SV=1 | 1 |
| ALDH3A2 | Aldehyde dehydrogenase family 3 member A2 OS=Homo sapiens OX=9606 GN=ALDH3A2 PE=1 SV=1 | 8 |
| HNRNPC | Heterogeneous nuclear ribonucleoproteins C1/C2 OS=Homo sapiens OX=9606 GN=HNRNPC PE=1 SV=4 | 3 |
| PFN1 | Profilin-1 OS=Homo sapiens OX=9606 GN=PFN1 PE=1 SV=2 | 15 |
| UBE3C | Ubiquitin-protein ligase E3C OS=Homo sapiens OX=9606 GN=UBE3C PE=1 SV=3 | 2 |
| ZNF729 | Zinc finger protein 729 OS=Homo sapiens OX=9606 GN=ZNF729 PE=2 SV=4 | 1 |
| SPNS1 | Protein spinster homolog 1 OS=Homo sapiens OX=9606 GN=SPNS1 PE=1 SV=1 | 2 |
| RAB5A | Ras-related protein Rab-5A OS=Homo sapiens OX=9606 GN=RAB5A PE=1 SV=2 | 3 |
| NUP98 | Nuclear pore complex protein Nup98-Nup96 OS=Homo sapiens OX=9606 GN=NUP98 PE=1 SV=4 | 8 |
| EIF4H | Eukaryotic translation initiation factor 4H OS=Homo sapiens OX=9606 GN=EIF4H PE=1 SV=5 | 4 |
| CCDC97 | Coiled-coil domain-containing protein 97 OS=Homo sapiens OX=9606 GN=CCDC97 PE=1 SV=1 | 2 |
| DCTN1 | Dynactin subunit 1 OS=Homo sapiens OX=9606 GN=DCTN1 PE=1 SV=3 | 33 |
| ATG3 | Ubiquitin-like-conjugating enzyme ATG3 OS=Homo sapiens OX=9606 GN=ATG3 PE=1 SV=1 | 4 |
| DIAPH2 | Protein diaphanous homolog 2 OS=Homo sapiens OX=9606 GN=DIAPH2 PE=1 SV=1 | 10 |
| RPS8 | 40S ribosomal protein S8 OS=Homo sapiens OX=9606 GN=RPS8 PE=1 SV=2 | 10 |
| TOMM70 | Mitochondrial import receptor subunit TOM70 OS=Homo sapiens OX=9606 GN=TOMM70 PE=1 SV=1 | 23 |
| ELAVL1 | ELAV-like protein 1 OS=Homo sapiens OX=9606 GN=ELAVL1 PE=1 SV=2 | 11 |
| PABPC1 | Polyadenylate-binding protein 1 OS=Homo sapiens OX=9606 GN=PABPC1 PE=1 SV=2 | 2 |
| BSG | Basigin OS=Homo sapiens OX=9606 GN=BSG PE=1 SV=2 | 7 |
| ETFDH | Electron transfer flavoprotein-ubiquinone oxidoreductase, mitochondrial OS=Homo sapiens OX=9606 GN=ETFDH PE=1 SV=2 | 7 |
| TBC1D1 | TBC1 domain family member 1 OS=Homo sapiens OX=9606 GN=TBC1D1 PE=1 SV=2 | 3 |
| CAPNS1 | Calpain small subunit 1 OS=Homo sapiens OX=9606 GN=CAPNS1 PE=1 SV=1 | 8 |
| GALNS | N-acetylgalactosamine-6-sulfatase OS=Homo sapiens OX=9606 GN=GALNS PE=1 SV=1 | 2 |
| SH3GL2 | Endophilin-A1 OS=Homo sapiens OX=9606 GN=SH3GL2 PE=1 SV=1 | 10 |
| PGK1 | Phosphoglycerate kinase 1 OS=Homo sapiens OX=9606 GN=PGK1 PE=1 SV=3 | 35 |
| DYNC1LI1 | Cytoplasmic dynein 1 light intermediate chain 1 OS=Homo sapiens OX=9606 GN=DYNC1LI1 PE=1 SV=3 | 21 |
| CHEK1 | Serine/threonine-protein kinase Chk1 OS=Homo sapiens OX=9606 GN=CHEK1 PE=1 SV=2 | 7 |
| CALU | Calumenin OS=Homo sapiens OX=9606 GN=CALU PE=1 SV=2 | 18 |
| DMGDH | Dimethylglycine dehydrogenase, mitochondrial OS=Homo sapiens OX=9606 GN=DMGDH PE=1 SV=2 | 2 |
| DNAJA2 | DnaJ homolog subfamily A member 2 OS=Homo sapiens OX=9606 GN=DNAJA2 PE=1 SV=1 | 16 |
| ACAA2 | 3-ketoacyl-CoA thiolase, mitochondrial OS=Homo sapiens OX=9606 GN=ACAA2 PE=1 SV=2 | 21 |
| HMGB2 | High mobility group protein B2 OS=Homo sapiens OX=9606 GN=HMGB2 PE=1 SV=2 | 7 |
| RTCB | RNA-splicing ligase RtcB homolog OS=Homo sapiens OX=9606 GN=RTCB PE=1 SV=1 | 24 |
| GPI | Glucose-6-phosphate isomerase OS=Homo sapiens OX=9606 GN=GPI PE=1 SV=4 | 25 |
| ETHE1 | Persulfide dioxygenase ETHE1, mitochondrial OS=Homo sapiens OX=9606 GN=ETHE1 PE=1 SV=2 | 11 |
| OS9 | Protein OS-9 OS=Homo sapiens OX=9606 GN=OS9 PE=1 SV=1 | 18 |
| ADAD1 | Adenosine deaminase domain-containing protein 1 OS=Homo sapiens OX=9606 GN=ADAD1 PE=2 SV=1 | 5 |
| GYG1 | Glycogenin-1 OS=Homo sapiens OX=9606 GN=GYG1 PE=1 SV=4 | 15 |
| KPNA4 | Importin subunit alpha-3 OS=Homo sapiens OX=9606 GN=KPNA4 PE=1 SV=1 | 3 |
| HSPA8 | Heat shock cognate 71 kDa protein OS=Homo sapiens OX=9606 GN=HSPA8 PE=1 SV=1 | 45 |
| PSMD10 | 26S proteasome non-ATPase regulatory subunit 10 OS=Homo sapiens OX=9606 GN=PSMD10 PE=1 SV=1 | 11 |
| EXOC3 | Exocyst complex component 3 OS=Homo sapiens OX=9606 GN=EXOC3 PE=1 SV=3 | 5 |
| MCTP1 | Multiple C2 and transmembrane domain-containing protein 1 OS=Homo sapiens OX=9606 GN=MCTP1 PE=2 SV=2 | 1 |
| LRRC59 | Leucine-rich repeat-containing protein 59 OS=Homo sapiens OX=9606 GN=LRRC59 PE=1 SV=1 | 10 |
| GC | Vitamin D-binding protein OS=Homo sapiens OX=9606 GN=GC PE=1 SV=2 | 20 |
| ATP5PB | ATP synthase F(0) complex subunit B1, mitochondrial OS=Homo sapiens OX=9606 GN=ATP5PB PE=1 SV=2 | 14 |
| MEAK7 | MTOR-associated protein MEAK7 OS=Homo sapiens OX=9606 GN=MEAK7 PE=1 SV=2 | 6 |
| AGPS | Alkyldihydroxyacetonephosphate synthase, peroxisomal OS=Homo sapiens OX=9606 GN=AGPS PE=1 SV=1 | 10 |
| CDC37L1 | Hsp90 co-chaperone Cdc37-like 1 OS=Homo sapiens OX=9606 GN=CDC37L1 PE=1 SV=1 | 3 |
| ECE1 | Endothelin-converting enzyme 1 OS=Homo sapiens OX=9606 GN=ECE1 PE=1 SV=2 | 9 |
| ANXA5 | Annexin A5 OS=Homo sapiens OX=9606 GN=ANXA5 PE=1 SV=2 | 17 |
| NCL | Nucleolin OS=Homo sapiens OX=9606 GN=NCL PE=1 SV=3 | 12 |
| WASHC2A | WASH complex subunit 2A OS=Homo sapiens OX=9606 GN=WASHC2A PE=1 SV=3 | 7 |
| YWHAG | 14-3-3 protein gamma OS=Homo sapiens OX=9606 GN=YWHAG PE=1 SV=2 | 10 |
| TOMM22 | Mitochondrial import receptor subunit TOM22 homolog OS=Homo sapiens OX=9606 GN=TOMM22 PE=1 SV=3 | 4 |
| RETREG3 | Reticulophagy regulator 3 OS=Homo sapiens OX=9606 GN=RETREG3 PE=1 SV=1 | 3 |
| POLB | DNA polymerase beta OS=Homo sapiens OX=9606 GN=POLB PE=1 SV=3 | 8 |
| ITGA2 | Integrin alpha-2 OS=Homo sapiens OX=9606 GN=ITGA2 PE=1 SV=2 | 22 |
| MCCC2 | Methylcrotonoyl-CoA carboxylase beta chain, mitochondrial OS=Homo sapiens OX=9606 GN=MCCC2 PE=1 SV=1 | 18 |
| NARS1 | Asparagine--tRNA ligase, cytoplasmic OS=Homo sapiens OX=9606 GN=NARS1 PE=1 SV=1 | 15 |
| TPRXL | Tetra-peptide repeat homeobox-like protein OS=Homo sapiens OX=9606 GN=TPRXL PE=1 SV=3 | 3 |
| PRMT5 | Protein arginine N-methyltransferase 5 OS=Homo sapiens OX=9606 GN=PRMT5 PE=1 SV=4 | 7 |
| UBE2K | Ubiquitin-conjugating enzyme E2 K OS=Homo sapiens OX=9606 GN=UBE2K PE=1 SV=3 | 6 |
| RANBP3 | Ran-binding protein 3 OS=Homo sapiens OX=9606 GN=RANBP3 PE=1 SV=1 | 6 |
| TARS1 | Threonine--tRNA ligase 1, cytoplasmic OS=Homo sapiens OX=9606 GN=TARS1 PE=1 SV=3 | 11 |
| PYGB | Glycogen phosphorylase, brain form OS=Homo sapiens OX=9606 GN=PYGB PE=1 SV=5 | 24 |
| SNX11 | Sorting nexin-11 OS=Homo sapiens OX=9606 GN=SNX11 PE=1 SV=2 | 4 |
| MIEN1 | Migration and invasion enhancer 1 OS=Homo sapiens OX=9606 GN=MIEN1 PE=1 SV=1 | 2 |
| LRRC47 | Leucine-rich repeat-containing protein 47 OS=Homo sapiens OX=9606 GN=LRRC47 PE=1 SV=1 | 22 |
| RPLP0 | 60S acidic ribosomal protein P0 OS=Homo sapiens OX=9606 GN=RPLP0 PE=1 SV=1 | 11 |
| LARS1 | Leucine--tRNA ligase, cytoplasmic OS=Homo sapiens OX=9606 GN=LARS1 PE=1 SV=2 | 21 |
| SNX1 | Sorting nexin-1 OS=Homo sapiens OX=9606 GN=SNX1 PE=1 SV=3 | 14 |
| SMS | Spermine synthase OS=Homo sapiens OX=9606 GN=SMS PE=1 SV=2 | 26 |
| EIF5B | Eukaryotic translation initiation factor 5B OS=Homo sapiens OX=9606 GN=EIF5B PE=1 SV=4 | 6 |
| TMED10 | Transmembrane emp24 domain-containing protein 10 OS=Homo sapiens OX=9606 GN=TMED10 PE=1 SV=2 | 8 |
| RPS6KA1 | Ribosomal protein S6 kinase alpha-1 OS=Homo sapiens OX=9606 GN=RPS6KA1 PE=1 SV=2 | 3 |
| MYL6 | Myosin light polypeptide 6 OS=Homo sapiens OX=9606 GN=MYL6 PE=1 SV=2 | 9 |
| C11orf54 | Ester hydrolase C11orf54 OS=Homo sapiens OX=9606 GN=C11orf54 PE=1 SV=1 | 7 |
| AUH | Methylglutaconyl-CoA hydratase, mitochondrial OS=Homo sapiens OX=9606 GN=AUH PE=1 SV=1 | 2 |
| FIP1L1 | Pre-mRNA 3'-end-processing factor FIP1 OS=Homo sapiens OX=9606 GN=FIP1L1 PE=1 SV=1 | 3 |
| ATP5F1EP2 | ATP synthase subunit epsilon-like protein, mitochondrial OS=Homo sapiens OX=9606 GN=ATP5F1EP2 PE=1 SV=1 | 2 |
| GLA | Alpha-galactosidase A OS=Homo sapiens OX=9606 GN=GLA PE=1 SV=1 | 4 |
| CKM | Creatine kinase M-type OS=Homo sapiens OX=9606 GN=CKM PE=1 SV=2 | 3 |
| EXOC2 | Exocyst complex component 2 OS=Homo sapiens OX=9606 GN=EXOC2 PE=1 SV=1 | 5 |
| EIF3C | Eukaryotic translation initiation factor 3 subunit C OS=Homo sapiens OX=9606 GN=EIF3C PE=1 SV=1 | 12 |
| PPP5C | Serine/threonine-protein phosphatase 5 OS=Homo sapiens OX=9606 GN=PPP5C PE=1 SV=1 | 17 |
| QARS1 | Glutamine--tRNA ligase OS=Homo sapiens OX=9606 GN=QARS1 PE=1 SV=1 | 16 |
| **CD44** | **CD44 antigen OS=Homo sapiens OX=9606 GN=CD44 PE=1 SV=3** | **2** |
| ATXN2L | Ataxin-2-like protein OS=Homo sapiens OX=9606 GN=ATXN2L PE=1 SV=2 | 5 |
| "H4C2 | Histone H4 OS=Homo sapiens OX=9606 GN=H4C16 PE=1 SV=2 | 11 |
| ACADS | Short-chain specific acyl-CoA dehydrogenase, mitochondrial OS=Homo sapiens OX=9606 GN=ACADS PE=1 SV=1 | 12 |
| DDX3X | ATP-dependent RNA helicase DDX3X OS=Homo sapiens OX=9606 GN=DDX3X PE=1 SV=3 | 25 |
| RABEP1 | Rab GTPase-binding effector protein 1 OS=Homo sapiens OX=9606 GN=RABEP1 PE=1 SV=2 | 13 |
| CCT8 | T-complex protein 1 subunit theta OS=Homo sapiens OX=9606 GN=CCT8 PE=1 SV=4 | 45 |
| PIWIL1 | Piwi-like protein 1 OS=Homo sapiens OX=9606 GN=PIWIL1 PE=1 SV=1 | 7 |
| SLC25A3 | Phosphate carrier protein, mitochondrial OS=Homo sapiens OX=9606 GN=SLC25A3 PE=1 SV=2 | 17 |
| CCS | Copper chaperone for superoxide dismutase OS=Homo sapiens OX=9606 GN=CCS PE=1 SV=1 | 7 |
| ATP5MG | ATP synthase subunit g, mitochondrial OS=Homo sapiens OX=9606 GN=ATP5MG PE=1 SV=3 | 7 |
| NUP85 | Nuclear pore complex protein Nup85 OS=Homo sapiens OX=9606 GN=NUP85 PE=1 SV=1 | 1 |
| RAP1B | Ras-related protein Rap-1b OS=Homo sapiens OX=9606 GN=RAP1B PE=1 SV=1 | 9 |
| SNF8 | Vacuolar-sorting protein SNF8 OS=Homo sapiens OX=9606 GN=SNF8 PE=1 SV=1 | 3 |
| GCN1 | eIF-2-alpha kinase activator GCN1 OS=Homo sapiens OX=9606 GN=GCN1 PE=1 SV=7 | 17 |
| PC | Pyruvate carboxylase, mitochondrial OS=Homo sapiens OX=9606 GN=PC PE=1 SV=2 | 34 |
| XPNPEP1 | Xaa-Pro aminopeptidase 1 OS=Homo sapiens OX=9606 GN=XPNPEP1 PE=1 SV=3 | 10 |
| MYCBP | c-Myc-binding protein OS=Homo sapiens OX=9606 GN=MYCBP PE=1 SV=3 | 7 |
| HECTD1 | E3 ubiquitin-protein ligase HECTD1 OS=Homo sapiens OX=9606 GN=HECTD1 PE=1 SV=4 | 8 |
| LRPPRC | Leucine-rich PPR motif-containing protein, mitochondrial OS=Homo sapiens OX=9606 GN=LRPPRC PE=1 SV=3 | 63 |
| ARPC2 | Actin-related protein 2/3 complex subunit 2 OS=Homo sapiens OX=9606 GN=ARPC2 PE=1 SV=1 | 13 |
| UBQLN1 | Ubiquilin-1 OS=Homo sapiens OX=9606 GN=UBQLN1 PE=1 SV=2 | 6 |
| TM9SF2 | Transmembrane 9 superfamily member 2 OS=Homo sapiens OX=9606 GN=TM9SF2 PE=1 SV=1 | 11 |
| PADI6 | Protein-arginine deiminase type-6 OS=Homo sapiens OX=9606 GN=PADI6 PE=1 SV=3 | 50 |
| DLGAP5 | Disks large-associated protein 5 OS=Homo sapiens OX=9606 GN=DLGAP5 PE=1 SV=2 | 35 |
| MYO6 | Unconventional myosin-VI OS=Homo sapiens OX=9606 GN=MYO6 PE=1 SV=4 | 35 |
| NDRG1 | Protein NDRG1 OS=Homo sapiens OX=9606 GN=NDRG1 PE=1 SV=1 | 6 |
| LGALS1 | Galectin-1 OS=Homo sapiens OX=9606 GN=LGALS1 PE=1 SV=2 | 11 |
| SCARB2 | Lysosome membrane protein 2 OS=Homo sapiens OX=9606 GN=SCARB2 PE=1 SV=2 | 9 |
| DYNLRB1 | Dynein light chain roadblock-type 1 OS=Homo sapiens OX=9606 GN=DYNLRB1 PE=1 SV=3 | 5 |
| MTHFD1 | C-1-tetrahydrofolate synthase, cytoplasmic OS=Homo sapiens OX=9606 GN=MTHFD1 PE=1 SV=4 | 36 |
| CHMP4B | Charged multivesicular body protein 4b OS=Homo sapiens OX=9606 GN=CHMP4B PE=1 SV=1 | 8 |
| MIA3 | Transport and Golgi organization protein 1 homolog OS=Homo sapiens OX=9606 GN=MIA3 PE=1 SV=1 | 27 |
| IMPA1 | Inositol monophosphatase 1 OS=Homo sapiens OX=9606 GN=IMPA1 PE=1 SV=1 | 14 |
| LOXL2 | Lysyl oxidase homolog 2 OS=Homo sapiens OX=9606 GN=LOXL2 PE=1 SV=1 | 14 |
| ATP6V1A | V-type proton ATPase catalytic subunit A OS=Homo sapiens OX=9606 GN=ATP6V1A PE=1 SV=2 | 24 |
| PSMD3 | 26S proteasome non-ATPase regulatory subunit 3 OS=Homo sapiens OX=9606 GN=PSMD3 PE=1 SV=2 | 23 |
| ARHGAP18 | Rho GTPase-activating protein 18 OS=Homo sapiens OX=9606 GN=ARHGAP18 PE=1 SV=3 | 6 |
| BNIP1 | Vesicle transport protein SEC20 OS=Homo sapiens OX=9606 GN=BNIP1 PE=1 SV=3 | 8 |
| AFG3L2 | AFG3-like protein 2 OS=Homo sapiens OX=9606 GN=AFG3L2 PE=1 SV=2 | 32 |
| POMGNT1 | Protein O-linked-mannose beta-1,2-N-acetylglucosaminyltransferase 1 OS=Homo sapiens OX=9606 GN=POMGNT1 PE=1 SV=2 | 4 |
| CUTA | Protein CutA OS=Homo sapiens OX=9606 GN=CUTA PE=1 SV=2 | 3 |
| DARS1 | Aspartate--tRNA ligase, cytoplasmic OS=Homo sapiens OX=9606 GN=DARS1 PE=1 SV=2 | 23 |
| ATP5ME | ATP synthase subunit e, mitochondrial OS=Homo sapiens OX=9606 GN=ATP5ME PE=1 SV=2 | 6 |
| DIABLO | Diablo IAP-binding mitochondrial protein OS=Homo sapiens OX=9606 GN=DIABLO PE=1 SV=1 | 11 |
| MAPK1 | Mitogen-activated protein kinase 1 OS=Homo sapiens OX=9606 GN=MAPK1 PE=1 SV=3 | 16 |
| OVCH1 | Ovochymase-1 OS=Homo sapiens OX=9606 GN=OVCH1 PE=2 SV=2 | 9 |
| ENDOD1 | Endonuclease domain-containing 1 protein OS=Homo sapiens OX=9606 GN=ENDOD1 PE=1 SV=2 | 9 |
| UBE2R2 | Ubiquitin-conjugating enzyme E2 R2 OS=Homo sapiens OX=9606 GN=UBE2R2 PE=1 SV=1 | 3 |
| PGD | 6-phosphogluconate dehydrogenase, decarboxylating OS=Homo sapiens OX=9606 GN=PGD PE=1 SV=3 | 26 |
| CCNB1 | G2/mitotic-specific cyclin-B1 OS=Homo sapiens OX=9606 GN=CCNB1 PE=1 SV=1 | 5 |
| BDH2 | Dehydrogenase/reductase SDR family member 6 OS=Homo sapiens OX=9606 GN=BDH2 PE=1 SV=2 | 9 |
| MATN2 | Matrilin-2 OS=Homo sapiens OX=9606 GN=MATN2 PE=1 SV=4 | 5 |
| MCM6 | DNA replication licensing factor MCM6 OS=Homo sapiens OX=9606 GN=MCM6 PE=1 SV=1 | 5 |
| ATG16L1 | Autophagy-related protein 16-1 OS=Homo sapiens OX=9606 GN=ATG16L1 PE=1 SV=2 | 3 |
| PFDN6 | Prefoldin subunit 6 OS=Homo sapiens OX=9606 GN=PFDN6 PE=1 SV=1 | 5 |
| LYPLA1 | Acyl-protein thioesterase 1 OS=Homo sapiens OX=9606 GN=LYPLA1 PE=1 SV=1 | 8 |
| C8orf82 | UPF0598 protein C8orf82 OS=Homo sapiens OX=9606 GN=C8orf82 PE=1 SV=2 | 1 |
| NKIRAS2 | NF-kappa-B inhibitor-interacting Ras-like protein 2 OS=Homo sapiens OX=9606 GN=NKIRAS2 PE=1 SV=1 | 1 |
| METTL26 | Methyltransferase-like 26 OS=Homo sapiens OX=9606 GN=METTL26 PE=1 SV=2 | 5 |
| SLC25A1 | Tricarboxylate transport protein, mitochondrial OS=Homo sapiens OX=9606 GN=SLC25A1 PE=1 SV=2 | 12 |
| ZC3H3 | Zinc finger CCCH domain-containing protein 3 OS=Homo sapiens OX=9606 GN=ZC3H3 PE=1 SV=3 | 1 |
| DHX30 | ATP-dependent RNA helicase DHX30 OS=Homo sapiens OX=9606 GN=DHX30 PE=1 SV=1 | 15 |
| FMR1 | Fragile X messenger ribonucleoprotein 1 OS=Homo sapiens OX=9606 GN=FMR1 PE=1 SV=1 | 3 |
| KIF4A | Chromosome-associated kinesin KIF4A OS=Homo sapiens OX=9606 GN=KIF4A PE=1 SV=3 | 10 |
| PSMG2 | Proteasome assembly chaperone 2 OS=Homo sapiens OX=9606 GN=PSMG2 PE=1 SV=1 | 3 |
| ADK | Adenosine kinase OS=Homo sapiens OX=9606 GN=ADK PE=1 SV=2 | 11 |
| NCK1 | Cytoplasmic protein NCK1 OS=Homo sapiens OX=9606 GN=NCK1 PE=1 SV=1 | 6 |
| ACOT13 | Acyl-coenzyme A thioesterase 13 OS=Homo sapiens OX=9606 GN=ACOT13 PE=1 SV=1 | 4 |
| COQ9 | Ubiquinone biosynthesis protein COQ9, mitochondrial OS=Homo sapiens OX=9606 GN=COQ9 PE=1 SV=1 | 3 |
| ZPR1 | Zinc finger protein ZPR1 OS=Homo sapiens OX=9606 GN=ZPR1 PE=1 SV=1 | 3 |
| OSCP1 | Protein OSCP1 OS=Homo sapiens OX=9606 GN=OSCP1 PE=1 SV=4 | 9 |
| AP2S1 | AP-2 complex subunit sigma OS=Homo sapiens OX=9606 GN=AP2S1 PE=1 SV=2 | 3 |
| KLHDC7B | Kelch domain-containing protein 7B OS=Homo sapiens OX=9606 GN=KLHDC7B PE=1 SV=2 | 1 |
| RBP1 | Retinol-binding protein 1 OS=Homo sapiens OX=9606 GN=RBP1 PE=1 SV=2 | 8 |
| SARS1 | Serine--tRNA ligase, cytoplasmic OS=Homo sapiens OX=9606 GN=SARS1 PE=1 SV=3 | 20 |
| DEGS2 | Sphingolipid delta(4)-desaturase/C4-monooxygenase DES2 OS=Homo sapiens OX=9606 GN=DEGS2 PE=1 SV=2 | 3 |
| CNP | 2',3'-cyclic-nucleotide 3'-phosphodiesterase OS=Homo sapiens OX=9606 GN=CNP PE=1 SV=2 | 12 |
| TMEM9 | Proton-transporting V-type ATPase complex assembly regulator TMEM9 OS=Homo sapiens OX=9606 GN=TMEM9 PE=1 SV=1 | 2 |
| ATP2A2 | Sarcoplasmic/endoplasmic reticulum calcium ATPase 2 OS=Homo sapiens OX=9606 GN=ATP2A2 PE=1 SV=1 | 13 |
| ATL3 | Atlastin-3 OS=Homo sapiens OX=9606 GN=ATL3 PE=1 SV=1 | 19 |
| COCH | Cochlin OS=Homo sapiens OX=9606 GN=COCH PE=1 SV=1 | 25 |
| EZR | Ezrin OS=Homo sapiens OX=9606 GN=EZR PE=1 SV=4 | 50 |
| SCFD1 | Sec1 family domain-containing protein 1 OS=Homo sapiens OX=9606 GN=SCFD1 PE=1 SV=4 | 27 |
| ACTR3 | Actin-related protein 3 OS=Homo sapiens OX=9606 GN=ACTR3 PE=1 SV=3 | 20 |
| NUDCD3 | NudC domain-containing protein 3 OS=Homo sapiens OX=9606 GN=NUDCD3 PE=1 SV=3 | 12 |
| XRN2 | 5'-3' exoribonuclease 2 OS=Homo sapiens OX=9606 GN=XRN2 PE=1 SV=1 | 3 |
| MYO18A | Unconventional myosin-XVIIIa OS=Homo sapiens OX=9606 GN=MYO18A PE=1 SV=3 | 9 |
| ARG2 | Arginase-2, mitochondrial OS=Homo sapiens OX=9606 GN=ARG2 PE=1 SV=1 | 24 |
| NUTF2 | Nuclear transport factor 2 OS=Homo sapiens OX=9606 GN=NUTF2 PE=1 SV=1 | 4 |
| SMU1 | WD40 repeat-containing protein SMU1 OS=Homo sapiens OX=9606 GN=SMU1 PE=1 SV=2 | 1 |
| ANP32B | Acidic leucine-rich nuclear phosphoprotein 32 family member B OS=Homo sapiens OX=9606 GN=ANP32B PE=1 SV=1 | 4 |
| RUBCN | Run domain Beclin-1-interacting and cysteine-rich domain-containing protein OS=Homo sapiens OX=9606 GN=RUBCN PE=1 SV=4 | 1 |
| NADK2 | NAD kinase 2, mitochondrial OS=Homo sapiens OX=9606 GN=NADK2 PE=1 SV=2 | 14 |
| EIF3F | Eukaryotic translation initiation factor 3 subunit F OS=Homo sapiens OX=9606 GN=EIF3F PE=1 SV=1 | 7 |
| PARK7 | Parkinson disease protein 7 OS=Homo sapiens OX=9606 GN=PARK7 PE=1 SV=2 | 15 |
| DHX29 | ATP-dependent RNA helicase DHX29 OS=Homo sapiens OX=9606 GN=DHX29 PE=1 SV=2 | 10 |
| EIF4G2 | Eukaryotic translation initiation factor 4 gamma 2 OS=Homo sapiens OX=9606 GN=EIF4G2 PE=1 SV=1 | 23 |
| ILKAP | Integrin-linked kinase-associated serine/threonine phosphatase 2C OS=Homo sapiens OX=9606 GN=ILKAP PE=1 SV=1 | 5 |
| EPN1 | Epsin-1 OS=Homo sapiens OX=9606 GN=EPN1 PE=1 SV=2 | 4 |
| RPL22 | 60S ribosomal protein L22 OS=Homo sapiens OX=9606 GN=RPL22 PE=1 SV=2 | 3 |
| DDX6 | Probable ATP-dependent RNA helicase DDX6 OS=Homo sapiens OX=9606 GN=DDX6 PE=1 SV=2 | 26 |
| FBXO22 | F-box only protein 22 OS=Homo sapiens OX=9606 GN=FBXO22 PE=1 SV=1 | 13 |
| CCT5 | T-complex protein 1 subunit epsilon OS=Homo sapiens OX=9606 GN=CCT5 PE=1 SV=1 | 38 |
| LSM1 | U6 snRNA-associated Sm-like protein LSm1 OS=Homo sapiens OX=9606 GN=LSM1 PE=1 SV=1 | 6 |
| RPS5 | 40S ribosomal protein S5 OS=Homo sapiens OX=9606 GN=RPS5 PE=1 SV=4 | 8 |
| VPS26A | Vacuolar protein sorting-associated protein 26A OS=Homo sapiens OX=9606 GN=VPS26A PE=1 SV=2 | 16 |
| MAPK7 | Mitogen-activated protein kinase 7 OS=Homo sapiens OX=9606 GN=MAPK7 PE=1 SV=2 | 4 |
| DDX58 | Antiviral innate immune response receptor RIG-I OS=Homo sapiens OX=9606 GN=RIGI PE=1 SV=2 | 8 |
| GAPDH | Glyceraldehyde-3-phosphate dehydrogenase OS=Homo sapiens OX=9606 GN=GAPDH PE=1 SV=3 | 21 |
| CPSF3 | Cleavage and polyadenylation specificity factor subunit 3 OS=Homo sapiens OX=9606 GN=CPSF3 PE=1 SV=1 | 7 |
| CD81 | CD81 antigen OS=Homo sapiens OX=9606 GN=CD81 PE=1 SV=1 | 2 |
| BCKDK | [3-methyl-2-oxobutanoate dehydrogenase [lipoamide]] kinase, mitochondrial OS=Homo sapiens OX=9606 GN=BCKDK PE=1 SV=2 | 4 |
| PEBP1 | Phosphatidylethanolamine-binding protein 1 OS=Homo sapiens OX=9606 GN=PEBP1 PE=1 SV=3 | 12 |
| ACLY | ATP-citrate synthase OS=Homo sapiens OX=9606 GN=ACLY PE=1 SV=3 | 24 |
| KHSRP | Far upstream element-binding protein 2 OS=Homo sapiens OX=9606 GN=KHSRP PE=1 SV=4 | 6 |
| MUC20 | Mucin-20 OS=Homo sapiens OX=9606 GN=MUC20 PE=1 SV=3 | 3 |
| G6PD | Glucose-6-phosphate 1-dehydrogenase OS=Homo sapiens OX=9606 GN=G6PD PE=1 SV=4 | 46 |
| RPL23 | 60S ribosomal protein L23 OS=Homo sapiens OX=9606 GN=RPL23 PE=1 SV=1 | 5 |
| MFGE8 | Lactadherin OS=Homo sapiens OX=9606 GN=MFGE8 PE=1 SV=3 | 25 |
| ADGRG1 | Adhesion G-protein coupled receptor G1 OS=Homo sapiens OX=9606 GN=ADGRG1 PE=1 SV=2 | 4 |
| ATP5F1D | ATP synthase subunit delta, mitochondrial OS=Homo sapiens OX=9606 GN=ATP5F1D PE=1 SV=2 | 3 |
| DUSP19 | Dual specificity protein phosphatase 19 OS=Homo sapiens OX=9606 GN=DUSP19 PE=1 SV=1 | 1 |
| SLC25A11 | Mitochondrial 2-oxoglutarate/malate carrier protein OS=Homo sapiens OX=9606 GN=SLC25A11 PE=1 SV=3 | 3 |
| PPA1 | Inorganic pyrophosphatase OS=Homo sapiens OX=9606 GN=PPA1 PE=1 SV=2 | 16 |
| SERPINF1 | Pigment epithelium-derived factor OS=Homo sapiens OX=9606 GN=SERPINF1 PE=1 SV=4 | 12 |
| ATP6V1H | V-type proton ATPase subunit H OS=Homo sapiens OX=9606 GN=ATP6V1H PE=1 SV=1 | 14 |
| PISD | Phosphatidylserine decarboxylase proenzyme, mitochondrial OS=Homo sapiens OX=9606 GN=PISD PE=1 SV=4 | 3 |
| SLC25A19 | Mitochondrial thiamine pyrophosphate carrier OS=Homo sapiens OX=9606 GN=SLC25A19 PE=1 SV=1 | 4 |
| REEP2 | Receptor expression-enhancing protein 2 OS=Homo sapiens OX=9606 GN=REEP2 PE=1 SV=2 | 6 |
| RPL10A | 60S ribosomal protein L10a OS=Homo sapiens OX=9606 GN=RPL10A PE=1 SV=2 | 6 |
| ACOT8 | Acyl-coenzyme A thioesterase 8 OS=Homo sapiens OX=9606 GN=ACOT8 PE=1 SV=1 | 4 |
| RNASET2 | Ribonuclease T2 OS=Homo sapiens OX=9606 GN=RNASET2 PE=1 SV=2 | 4 |
| KARS1 | Lysine--tRNA ligase OS=Homo sapiens OX=9606 GN=KARS1 PE=1 SV=3 | 19 |
| HINT1 | Adenosine 5'-monophosphoramidase HINT1 OS=Homo sapiens OX=9606 GN=HINT1 PE=1 SV=2 | 11 |
| RPS4X | 40S ribosomal protein S4, X isoform OS=Homo sapiens OX=9606 GN=RPS4X PE=1 SV=2 | 14 |
| NEFM | Neurofilament medium polypeptide OS=Homo sapiens OX=9606 GN=NEFM PE=1 SV=3 | 1 |
| SLC25A6 | ADP/ATP translocase 3 OS=Homo sapiens OX=9606 GN=SLC25A6 PE=1 SV=4 | 2 |
| PPIF | Peptidyl-prolyl cis-trans isomerase F, mitochondrial OS=Homo sapiens OX=9606 GN=PPIF PE=1 SV=1 | 7 |
| REXO2 | Oligoribonuclease, mitochondrial OS=Homo sapiens OX=9606 GN=REXO2 PE=1 SV=3 | 11 |
| KIF3A | Kinesin-like protein KIF3A OS=Homo sapiens OX=9606 GN=KIF3A PE=1 SV=4 | 4 |
| RNF170 | E3 ubiquitin-protein ligase RNF170 OS=Homo sapiens OX=9606 GN=RNF170 PE=1 SV=2 | 3 |
| IQCB1 | IQ calmodulin-binding motif-containing protein 1 OS=Homo sapiens OX=9606 GN=IQCB1 PE=1 SV=1 | 6 |
| COASY | Bifunctional coenzyme A synthase OS=Homo sapiens OX=9606 GN=COASY PE=1 SV=4 | 7 |
| TMED5 | Transmembrane emp24 domain-containing protein 5 OS=Homo sapiens OX=9606 GN=TMED5 PE=1 SV=1 | 3 |
| PCTP | Phosphatidylcholine transfer protein OS=Homo sapiens OX=9606 GN=PCTP PE=1 SV=1 | 1 |
| TRIM28 | Transcription intermediary factor 1-beta OS=Homo sapiens OX=9606 GN=TRIM28 PE=1 SV=5 | 12 |
| TULP3 | Tubby-related protein 3 OS=Homo sapiens OX=9606 GN=TULP3 PE=1 SV=2 | 8 |
| ACTL6A | Actin-like protein 6A OS=Homo sapiens OX=9606 GN=ACTL6A PE=1 SV=1 | 4 |
| "RPL9P8 | 60S ribosomal protein L9 OS=Homo sapiens OX=9606 GN=RPL9P9 PE=1 SV=1 | 4 |
| ASTL | Astacin-like metalloendopeptidase OS=Homo sapiens OX=9606 GN=ASTL PE=1 SV=4 | 25 |
| GLOD4 | Glyoxalase domain-containing protein 4 OS=Homo sapiens OX=9606 GN=GLOD4 PE=1 SV=1 | 14 |
| UBXN11 | UBX domain-containing protein 11 OS=Homo sapiens OX=9606 GN=UBXN11 PE=1 SV=2 | 16 |
| TBC1D9B | TBC1 domain family member 9B OS=Homo sapiens OX=9606 GN=TBC1D9B PE=1 SV=3 | 4 |
| UGP2 | UTP--glucose-1-phosphate uridylyltransferase OS=Homo sapiens OX=9606 GN=UGP2 PE=1 SV=5 | 28 |
| HERC4 | Probable E3 ubiquitin-protein ligase HERC4 OS=Homo sapiens OX=9606 GN=HERC4 PE=1 SV=1 | 7 |
| PUS1 | Pseudouridylate synthase 1 homolog OS=Homo sapiens OX=9606 GN=PUS1 PE=1 SV=3 | 7 |
| IMPDH1 | Inosine-5'-monophosphate dehydrogenase 1 OS=Homo sapiens OX=9606 GN=IMPDH1 PE=1 SV=2 | 3 |
| RALB | Ras-related protein Ral-B OS=Homo sapiens OX=9606 GN=RALB PE=1 SV=1 | 12 |
| PSIP1 | PC4 and SFRS1-interacting protein OS=Homo sapiens OX=9606 GN=PSIP1 PE=1 SV=1 | 1 |
| AURKAIP1 | Aurora kinase A-interacting protein OS=Homo sapiens OX=9606 GN=AURKAIP1 PE=1 SV=1 | 1 |
| CRTC1 | CREB-regulated transcription coactivator 1 OS=Homo sapiens OX=9606 GN=CRTC1 PE=1 SV=2 | 2 |
| MICAL1 | [F-actin]-monooxygenase MICAL1 OS=Homo sapiens OX=9606 GN=MICAL1 PE=1 SV=2 | 6 |
| EIF3J | Eukaryotic translation initiation factor 3 subunit J OS=Homo sapiens OX=9606 GN=EIF3J PE=1 SV=2 | 7 |
| ZP4 | Zona pellucida sperm-binding protein 4 OS=Homo sapiens OX=9606 GN=ZP4 PE=1 SV=1 | 22 |
| AACS | Acetoacetyl-CoA synthetase OS=Homo sapiens OX=9606 GN=AACS PE=1 SV=1 | 5 |
| RPS15A | 40S ribosomal protein S15a OS=Homo sapiens OX=9606 GN=RPS15A PE=1 SV=2 | 6 |
| ITIH1 | Inter-alpha-trypsin inhibitor heavy chain H1 OS=Homo sapiens OX=9606 GN=ITIH1 PE=1 SV=3 | 20 |
| PSPC1 | Paraspeckle component 1 OS=Homo sapiens OX=9606 GN=PSPC1 PE=1 SV=1 | 6 |
| TMEM126A | Transmembrane protein 126A OS=Homo sapiens OX=9606 GN=TMEM126A PE=1 SV=1 | 8 |
| SEPHS2 | Selenide, water dikinase 2 OS=Homo sapiens OX=9606 GN=SEPHS2 PE=1 SV=3 | 10 |
| ARF5 | ADP-ribosylation factor 5 OS=Homo sapiens OX=9606 GN=ARF5 PE=1 SV=2 | 9 |
| PAFAH1B1 | Platelet-activating factor acetylhydrolase IB subunit beta OS=Homo sapiens OX=9606 GN=PAFAH1B1 PE=1 SV=2 | 23 |
| EMC10 | ER membrane protein complex subunit 10 OS=Homo sapiens OX=9606 GN=EMC10 PE=1 SV=1 | 2 |
| LSM14B | Protein LSM14 homolog B OS=Homo sapiens OX=9606 GN=LSM14B PE=1 SV=1 | 14 |
| RPS3 | 40S ribosomal protein S3 OS=Homo sapiens OX=9606 GN=RPS3 PE=1 SV=2 | 12 |
| ANP32A | Acidic leucine-rich nuclear phosphoprotein 32 family member A OS=Homo sapiens OX=9606 GN=ANP32A PE=1 SV=1 | 5 |
| CLPB | Caseinolytic peptidase B protein homolog OS=Homo sapiens OX=9606 GN=CLPB PE=1 SV=1 | 6 |
| RPLP2 | 60S acidic ribosomal protein P2 OS=Homo sapiens OX=9606 GN=RPLP2 PE=1 SV=1 | 4 |
| SLC25A5 | ADP/ATP translocase 2 OS=Homo sapiens OX=9606 GN=SLC25A5 PE=1 SV=7 | 25 |
| NIT2 | Omega-amidase NIT2 OS=Homo sapiens OX=9606 GN=NIT2 PE=1 SV=1 | 17 |
| RPL13 | 60S ribosomal protein L13 OS=Homo sapiens OX=9606 GN=RPL13 PE=1 SV=4 | 8 |
| OCIAD1 | OCIA domain-containing protein 1 OS=Homo sapiens OX=9606 GN=OCIAD1 PE=1 SV=1 | 4 |
| GMPPA | Mannose-1-phosphate guanyltransferase alpha OS=Homo sapiens OX=9606 GN=GMPPA PE=1 SV=1 | 9 |
| PODXL | Podocalyxin OS=Homo sapiens OX=9606 GN=PODXL PE=1 SV=2 | 8 |
| ACAD8 | Isobutyryl-CoA dehydrogenase, mitochondrial OS=Homo sapiens OX=9606 GN=ACAD8 PE=1 SV=1 | 14 |
| MRPL27 | 39S ribosomal protein L27, mitochondrial OS=Homo sapiens OX=9606 GN=MRPL27 PE=1 SV=1 | 4 |
| CHCHD2 | Coiled-coil-helix-coiled-coil-helix domain-containing protein 2 OS=Homo sapiens OX=9606 GN=CHCHD2 PE=1 SV=1 | 4 |
| TARDBP | TAR DNA-binding protein 43 OS=Homo sapiens OX=9606 GN=TARDBP PE=1 SV=1 | 5 |
| PSMC2 | 26S proteasome regulatory subunit 7 OS=Homo sapiens OX=9606 GN=PSMC2 PE=1 SV=3 | 25 |
| HSPD1 | 60 kDa heat shock protein, mitochondrial OS=Homo sapiens OX=9606 GN=HSPD1 PE=1 SV=2 | 50 |
| NDUFAF2 | NADH dehydrogenase [ubiquinone] 1 alpha subcomplex assembly factor 2 OS=Homo sapiens OX=9606 GN=NDUFAF2 PE=1 SV=1 | 8 |
| DHRS1 | Dehydrogenase/reductase SDR family member 1 OS=Homo sapiens OX=9606 GN=DHRS1 PE=1 SV=1 | 2 |
| ACO1 | Cytoplasmic aconitate hydratase OS=Homo sapiens OX=9606 GN=ACO1 PE=1 SV=3 | 35 |
| ATP1B3 | Sodium/potassium-transporting ATPase subunit beta-3 OS=Homo sapiens OX=9606 GN=ATP1B3 PE=1 SV=1 | 4 |
| MRPL24 | 39S ribosomal protein L24, mitochondrial OS=Homo sapiens OX=9606 GN=MRPL24 PE=1 SV=1 | 4 |
| SVIP | Small VCP/p97-interacting protein OS=Homo sapiens OX=9606 GN=SVIP PE=1 SV=1 | 2 |
| CSDE1 | Cold shock domain-containing protein E1 OS=Homo sapiens OX=9606 GN=CSDE1 PE=1 SV=2 | 14 |
| RNPEP | Aminopeptidase B OS=Homo sapiens OX=9606 GN=RNPEP PE=1 SV=2 | 12 |
| SLURP2 | Secreted Ly-6/uPAR domain-containing protein 2 OS=Homo sapiens OX=9606 GN=SLURP2 PE=1 SV=1 | 3 |
| PDLIM1 | PDZ and LIM domain protein 1 OS=Homo sapiens OX=9606 GN=PDLIM1 PE=1 SV=4 | 9 |
| SLC3A2 | 4F2 cell-surface antigen heavy chain OS=Homo sapiens OX=9606 GN=SLC3A2 PE=1 SV=3 | 17 |
| SRP54 | Signal recognition particle subunit SRP54 OS=Homo sapiens OX=9606 GN=SRP54 PE=1 SV=1 | 12 |
| FAHD2A | Fumarylacetoacetate hydrolase domain-containing protein 2A OS=Homo sapiens OX=9606 GN=FAHD2A PE=1 SV=1 | 13 |
| YWHAE | 14-3-3 protein epsilon OS=Homo sapiens OX=9606 GN=YWHAE PE=1 SV=1 | 23 |
| HCCS | Holocytochrome c-type synthase OS=Homo sapiens OX=9606 GN=HCCS PE=1 SV=1 | 2 |
| PPIC | Peptidyl-prolyl cis-trans isomerase C OS=Homo sapiens OX=9606 GN=PPIC PE=1 SV=1 | 2 |
| LAMTOR1 | Ragulator complex protein LAMTOR1 OS=Homo sapiens OX=9606 GN=LAMTOR1 PE=1 SV=2 | 3 |
| DLAT | Dihydrolipoyllysine-residue acetyltransferase component of pyruvate dehydrogenase complex, mitochondrial OS=Homo sapiens OX=9606 GN=DLAT PE=1 SV=3 | 17 |
| ABCC5 | ATP-binding cassette sub-family C member 5 OS=Homo sapiens OX=9606 GN=ABCC5 PE=1 SV=2 | 2 |
| TPI1 | Triosephosphate isomerase OS=Homo sapiens OX=9606 GN=TPI1 PE=1 SV=4 | 18 |
| CLU | Clusterin OS=Homo sapiens OX=9606 GN=CLU PE=1 SV=1 | 11 |
| CCDC91 | Coiled-coil domain-containing protein 91 OS=Homo sapiens OX=9606 GN=CCDC91 PE=1 SV=2 | 1 |
| EMC7 | ER membrane protein complex subunit 7 OS=Homo sapiens OX=9606 GN=EMC7 PE=1 SV=1 | 3 |
| SYNE1 | Nesprin-1 OS=Homo sapiens OX=9606 GN=SYNE1 PE=1 SV=4 | 2 |
| VPS28 | Vacuolar protein sorting-associated protein 28 homolog OS=Homo sapiens OX=9606 GN=VPS28 PE=1 SV=1 | 6 |
| TIMM9 | Mitochondrial import inner membrane translocase subunit Tim9 OS=Homo sapiens OX=9606 GN=TIMM9 PE=1 SV=1 | 4 |
| HNRNPK | Heterogeneous nuclear ribonucleoprotein K OS=Homo sapiens OX=9606 GN=HNRNPK PE=1 SV=1 | 9 |
| SLC25A13 | Electrogenic aspartate/glutamate antiporter SLC25A13, mitochondrial OS=Homo sapiens OX=9606 GN=SLC25A13 PE=1 SV=2 | 13 |
| MINDY3 | Ubiquitin carboxyl-terminal hydrolase MINDY-3 OS=Homo sapiens OX=9606 GN=MINDY3 PE=1 SV=1 | 8 |
| CPS1 | Carbamoyl-phosphate synthase [ammonia], mitochondrial OS=Homo sapiens OX=9606 GN=CPS1 PE=1 SV=2 | 6 |
| SEPTIN11 | Septin-11 OS=Homo sapiens OX=9606 GN=SEPTIN11 PE=1 SV=3 | 18 |
| NDUFA5 | NADH dehydrogenase [ubiquinone] 1 alpha subcomplex subunit 5 OS=Homo sapiens OX=9606 GN=NDUFA5 PE=1 SV=3 | 6 |
| EIF4E | Eukaryotic translation initiation factor 4E OS=Homo sapiens OX=9606 GN=EIF4E PE=1 SV=2 | 8 |
| CS | Citrate synthase, mitochondrial OS=Homo sapiens OX=9606 GN=CS PE=1 SV=2 | 19 |
| AHCYL2 | Adenosylhomocysteinase 3 OS=Homo sapiens OX=9606 GN=AHCYL2 PE=1 SV=1 | 3 |
| ACACA | Acetyl-CoA carboxylase 1 OS=Homo sapiens OX=9606 GN=ACACA PE=1 SV=2 | 6 |
| PSMC3 | 26S proteasome regulatory subunit 6A OS=Homo sapiens OX=9606 GN=PSMC3 PE=1 SV=3 | 28 |
| MTIF3 | Translation initiation factor IF-3, mitochondrial OS=Homo sapiens OX=9606 GN=MTIF3 PE=1 SV=4 | 2 |
| PFDN5 | Prefoldin subunit 5 OS=Homo sapiens OX=9606 GN=PFDN5 PE=1 SV=2 | 6 |
| MRPL17 | 39S ribosomal protein L17, mitochondrial OS=Homo sapiens OX=9606 GN=MRPL17 PE=1 SV=1 | 2 |
| GNG12 | Guanine nucleotide-binding protein G(I)/G(S)/G(O) subunit gamma-12 OS=Homo sapiens OX=9606 GN=GNG12 PE=1 SV=3 | 4 |
| FAAH2 | Fatty-acid amide hydrolase 2 OS=Homo sapiens OX=9606 GN=FAAH2 PE=1 SV=1 | 1 |
| CCT6A | T-complex protein 1 subunit zeta OS=Homo sapiens OX=9606 GN=CCT6A PE=1 SV=3 | 29 |
| RANBP2 | E3 SUMO-protein ligase RanBP2 OS=Homo sapiens OX=9606 GN=RANBP2 PE=1 SV=2 | 10 |
| GALNT2 | Polypeptide N-acetylgalactosaminyltransferase 2 OS=Homo sapiens OX=9606 GN=GALNT2 PE=1 SV=1 | 6 |
| ARF4 | ADP-ribosylation factor 4 OS=Homo sapiens OX=9606 GN=ARF4 PE=1 SV=3 | 2 |
| DDOST | Dolichyl-diphosphooligosaccharide--protein glycosyltransferase 48 kDa subunit OS=Homo sapiens OX=9606 GN=DDOST PE=1 SV=4 | 8 |
| "H3C3 | Histone H3.1 OS=Homo sapiens OX=9606 GN=H3C12 PE=1 SV=2 | 7 |
| BUB3 | Mitotic checkpoint protein BUB3 OS=Homo sapiens OX=9606 GN=BUB3 PE=1 SV=1 | 12 |
| CAMLG | Guided entry of tail-anchored proteins factor CAMLG OS=Homo sapiens OX=9606 GN=CAMLG PE=1 SV=1 | 4 |
| PPIH | Peptidyl-prolyl cis-trans isomerase H OS=Homo sapiens OX=9606 GN=PPIH PE=1 SV=1 | 2 |
| PPP2R5D | Serine/threonine-protein phosphatase 2A 56 kDa regulatory subunit delta isoform OS=Homo sapiens OX=9606 GN=PPP2R5D PE=1 SV=1 | 17 |
| PIWIL3 | Piwi-like protein 3 OS=Homo sapiens OX=9606 GN=PIWIL3 PE=2 SV=2 | 20 |
| VDAC1 | Voltage-dependent anion-selective channel protein 1 OS=Homo sapiens OX=9606 GN=VDAC1 PE=1 SV=2 | 20 |
| OOEP | Oocyte-expressed protein homolog OS=Homo sapiens OX=9606 GN=OOEP PE=1 SV=3 | 10 |
| RPIA | Ribose-5-phosphate isomerase OS=Homo sapiens OX=9606 GN=RPIA PE=1 SV=3 | 5 |
| HBS1L | HBS1-like protein OS=Homo sapiens OX=9606 GN=HBS1L PE=1 SV=1 | 22 |
| NAXE | NAD(P)H-hydrate epimerase OS=Homo sapiens OX=9606 GN=NAXE PE=1 SV=2 | 9 |
| PSMC1 | 26S proteasome regulatory subunit 4 OS=Homo sapiens OX=9606 GN=PSMC1 PE=1 SV=1 | 23 |
| PTPMT1 | Phosphatidylglycerophosphatase and protein-tyrosine phosphatase 1 OS=Homo sapiens OX=9606 GN=PTPMT1 PE=1 SV=1 | 3 |
| SSR1 | Translocon-associated protein subunit alpha OS=Homo sapiens OX=9606 GN=SSR1 PE=1 SV=3 | 4 |
| APP | Amyloid-beta precursor protein OS=Homo sapiens OX=9606 GN=APP PE=1 SV=3 | 5 |
| CCT4 | T-complex protein 1 subunit delta OS=Homo sapiens OX=9606 GN=CCT4 PE=1 SV=4 | 32 |
| ATP6V1C1 | V-type proton ATPase subunit C 1 OS=Homo sapiens OX=9606 GN=ATP6V1C1 PE=1 SV=4 | 8 |
| GUF1 | Translation factor GUF1, mitochondrial OS=Homo sapiens OX=9606 GN=GUF1 PE=1 SV=1 | 1 |
| GPD2 | Glycerol-3-phosphate dehydrogenase, mitochondrial OS=Homo sapiens OX=9606 GN=GPD2 PE=1 SV=3 | 22 |
| FASN | Fatty acid synthase OS=Homo sapiens OX=9606 GN=FASN PE=1 SV=3 | 49 |
| COX17 | Cytochrome c oxidase copper chaperone OS=Homo sapiens OX=9606 GN=COX17 PE=1 SV=2 | 2 |
| EIF1 | Eukaryotic translation initiation factor 1 OS=Homo sapiens OX=9606 GN=EIF1 PE=1 SV=1 | 1 |
| PARS2 | Probable proline--tRNA ligase, mitochondrial OS=Homo sapiens OX=9606 GN=PARS2 PE=1 SV=1 | 5 |
| ECPAS | Proteasome adapter and scaffold protein ECM29 OS=Homo sapiens OX=9606 GN=ECPAS PE=1 SV=2 | 7 |
| HHLA2 | HERV-H LTR-associating protein 2 OS=Homo sapiens OX=9606 GN=HHLA2 PE=1 SV=1 | 1 |
| WARS2 | Tryptophan--tRNA ligase, mitochondrial OS=Homo sapiens OX=9606 GN=WARS2 PE=1 SV=1 | 2 |
| DHX36 | ATP-dependent DNA/RNA helicase DHX36 OS=Homo sapiens OX=9606 GN=DHX36 PE=1 SV=2 | 10 |
| MTMR12 | Myotubularin-related protein 12 OS=Homo sapiens OX=9606 GN=MTMR12 PE=1 SV=2 | 5 |
| PDCL | Phosducin-like protein OS=Homo sapiens OX=9606 GN=PDCL PE=1 SV=3 | 5 |
| GATD3B | Putative glutamine amidotransferase-like class 1 domain-containing protein 3B, mitochondrial OS=Homo sapiens OX=9606 GN=GATD3B PE=5 SV=1 | 12 |
| TPM3 | Tropomyosin alpha-3 chain OS=Homo sapiens OX=9606 GN=TPM3 PE=1 SV=2 | 12 |
| LHPP | Phospholysine phosphohistidine inorganic pyrophosphate phosphatase OS=Homo sapiens OX=9606 GN=LHPP PE=1 SV=2 | 8 |
| TMX4 | Thioredoxin-related transmembrane protein 4 OS=Homo sapiens OX=9606 GN=TMX4 PE=1 SV=1 | 3 |
| GBF1 | Golgi-specific brefeldin A-resistance guanine nucleotide exchange factor 1 OS=Homo sapiens OX=9606 GN=GBF1 PE=1 SV=2 | 16 |
| PKM | Pyruvate kinase PKM OS=Homo sapiens OX=9606 GN=PKM PE=1 SV=4 | 39 |
| DPM1 | Dolichol-phosphate mannosyltransferase subunit 1 OS=Homo sapiens OX=9606 GN=DPM1 PE=1 SV=1 | 2 |
| SNX2 | Sorting nexin-2 OS=Homo sapiens OX=9606 GN=SNX2 PE=1 SV=2 | 21 |
| RAB43 | Ras-related protein Rab-43 OS=Homo sapiens OX=9606 GN=RAB43 PE=1 SV=1 | 1 |
| ACAT1 | Acetyl-CoA acetyltransferase, mitochondrial OS=Homo sapiens OX=9606 GN=ACAT1 PE=1 SV=1 | 28 |
| ENPP7 | Ectonucleotide pyrophosphatase/phosphodiesterase family member 7 OS=Homo sapiens OX=9606 GN=ENPP7 PE=1 SV=4 | 1 |
| AIF1L | Allograft inflammatory factor 1-like OS=Homo sapiens OX=9606 GN=AIF1L PE=1 SV=1 | 1 |
| DAAM1 | Disheveled-associated activator of morphogenesis 1 OS=Homo sapiens OX=9606 GN=DAAM1 PE=1 SV=2 | 1 |
| CCM2 | Cerebral cavernous malformations 2 protein OS=Homo sapiens OX=9606 GN=CCM2 PE=1 SV=1 | 5 |
| GNAI1 | Guanine nucleotide-binding protein G(i) subunit alpha-1 OS=Homo sapiens OX=9606 GN=GNAI1 PE=1 SV=2 | 3 |
| C2CD5 | C2 domain-containing protein 5 OS=Homo sapiens OX=9606 GN=C2CD5 PE=1 SV=1 | 4 |
| JPT1 | Jupiter microtubule associated homolog 1 OS=Homo sapiens OX=9606 GN=JPT1 PE=1 SV=3 | 2 |
| FUBP3 | Far upstream element-binding protein 3 OS=Homo sapiens OX=9606 GN=FUBP3 PE=1 SV=2 | 1 |
| NDUFB9 | NADH dehydrogenase [ubiquinone] 1 beta subcomplex subunit 9 OS=Homo sapiens OX=9606 GN=NDUFB9 PE=1 SV=3 | 5 |
| HDDC2 | 5'-deoxynucleotidase HDDC2 OS=Homo sapiens OX=9606 GN=HDDC2 PE=1 SV=1 | 10 |
| XRCC1 | DNA repair protein XRCC1 OS=Homo sapiens OX=9606 GN=XRCC1 PE=1 SV=3 | 7 |
| RPL18 | 60S ribosomal protein L18 OS=Homo sapiens OX=9606 GN=RPL18 PE=1 SV=2 | 7 |
| USP35 | Ubiquitin carboxyl-terminal hydrolase 35 OS=Homo sapiens OX=9606 GN=USP35 PE=1 SV=3 | 1 |
| SCARB1 | Scavenger receptor class B member 1 OS=Homo sapiens OX=9606 GN=SCARB1 PE=1 SV=1 | 5 |
| TESC | Calcineurin B homologous protein 3 OS=Homo sapiens OX=9606 GN=TESC PE=1 SV=3 | 3 |
| TAP1 | Antigen peptide transporter 1 OS=Homo sapiens OX=9606 GN=TAP1 PE=1 SV=3 | 1 |
| ELK1 | ETS domain-containing protein Elk-1 OS=Homo sapiens OX=9606 GN=ELK1 PE=1 SV=2 | 1 |
| VSIR | V-type immunoglobulin domain-containing suppressor of T-cell activation OS=Homo sapiens OX=9606 GN=VSIR PE=1 SV=3 | 1 |
| EIF2AK4 | eIF-2-alpha kinase GCN2 OS=Homo sapiens OX=9606 GN=EIF2AK4 PE=1 SV=3 | 1 |
| RPA2 | Replication protein A 32 kDa subunit OS=Homo sapiens OX=9606 GN=RPA2 PE=1 SV=1 | 7 |
| MTCH1 | Mitochondrial carrier homolog 1 OS=Homo sapiens OX=9606 GN=MTCH1 PE=1 SV=1 | 5 |
| PSMD7 | 26S proteasome non-ATPase regulatory subunit 7 OS=Homo sapiens OX=9606 GN=PSMD7 PE=1 SV=2 | 10 |
| ERLIN2 | Erlin-2 OS=Homo sapiens OX=9606 GN=ERLIN2 PE=1 SV=1 | 9 |
| CHMP3 | Charged multivesicular body protein 3 OS=Homo sapiens OX=9606 GN=CHMP3 PE=1 SV=3 | 5 |
| CAP1 | Adenylyl cyclase-associated protein 1 OS=Homo sapiens OX=9606 GN=CAP1 PE=1 SV=5 | 27 |
| SPDL1 | Protein Spindly OS=Homo sapiens OX=9606 GN=SPDL1 PE=1 SV=2 | 23 |
| YWHAZ | 14-3-3 protein zeta/delta OS=Homo sapiens OX=9606 GN=YWHAZ PE=1 SV=1 | 18 |
| DAP3 | 28S ribosomal protein S29, mitochondrial OS=Homo sapiens OX=9606 GN=DAP3 PE=1 SV=1 | 4 |
| ATP6V1B2 | V-type proton ATPase subunit B, brain isoform OS=Homo sapiens OX=9606 GN=ATP6V1B2 PE=1 SV=3 | 26 |
| CAMSAP3 | Calmodulin-regulated spectrin-associated protein 3 OS=Homo sapiens OX=9606 GN=CAMSAP3 PE=1 SV=2 | 5 |
| CIT | Citron Rho-interacting kinase OS=Homo sapiens OX=9606 GN=CIT PE=1 SV=2 | 54 |
| NLN | Neurolysin, mitochondrial OS=Homo sapiens OX=9606 GN=NLN PE=1 SV=1 | 16 |
| POLDIP2 | Polymerase delta-interacting protein 2 OS=Homo sapiens OX=9606 GN=POLDIP2 PE=1 SV=1 | 11 |
| AVEN | Cell death regulator Aven OS=Homo sapiens OX=9606 GN=AVEN PE=1 SV=1 | 9 |
| ABCB8 | Mitochondrial potassium channel ATP-binding subunit OS=Homo sapiens OX=9606 GN=ABCB8 PE=1 SV=3 | 6 |
| NPLOC4 | Nuclear protein localization protein 4 homolog OS=Homo sapiens OX=9606 GN=NPLOC4 PE=1 SV=3 | 4 |
| UROD | Uroporphyrinogen decarboxylase OS=Homo sapiens OX=9606 GN=UROD PE=1 SV=2 | 4 |
| TM9SF4 | Transmembrane 9 superfamily member 4 OS=Homo sapiens OX=9606 GN=TM9SF4 PE=1 SV=2 | 3 |
| ACADM | Medium-chain specific acyl-CoA dehydrogenase, mitochondrial OS=Homo sapiens OX=9606 GN=ACADM PE=1 SV=1 | 26 |
| PMPCB | Mitochondrial-processing peptidase subunit beta OS=Homo sapiens OX=9606 GN=PMPCB PE=1 SV=2 | 11 |
| MIF | Macrophage migration inhibitory factor OS=Homo sapiens OX=9606 GN=MIF PE=1 SV=4 | 2 |
| RPL19 | 60S ribosomal protein L19 OS=Homo sapiens OX=9606 GN=RPL19 PE=1 SV=1 | 2 |
| RASA1 | Ras GTPase-activating protein 1 OS=Homo sapiens OX=9606 GN=RASA1 PE=1 SV=1 | 13 |
| DNAJC11 | DnaJ homolog subfamily C member 11 OS=Homo sapiens OX=9606 GN=DNAJC11 PE=1 SV=2 | 13 |
| KRIT1 | Krev interaction trapped protein 1 OS=Homo sapiens OX=9606 GN=KRIT1 PE=1 SV=2 | 6 |
| PPP3CB | Serine/threonine-protein phosphatase 2B catalytic subunit beta isoform OS=Homo sapiens OX=9606 GN=PPP3CB PE=1 SV=2 | 2 |
| RPL12 | 60S ribosomal protein L12 OS=Homo sapiens OX=9606 GN=RPL12 PE=1 SV=1 | 5 |
| CEP250 | Centrosome-associated protein CEP250 OS=Homo sapiens OX=9606 GN=CEP250 PE=1 SV=2 | 33 |
| GSTM3 | Glutathione S-transferase Mu 3 OS=Homo sapiens OX=9606 GN=GSTM3 PE=1 SV=3 | 25 |
| GNAS | Guanine nucleotide-binding protein G(s) subunit alpha isoforms XLas OS=Homo sapiens OX=9606 GN=GNAS PE=1 SV=2 | 5 |
| YWHAH | 14-3-3 protein eta OS=Homo sapiens OX=9606 GN=YWHAH PE=1 SV=4 | 14 |
| PGM3 | Phosphoacetylglucosamine mutase OS=Homo sapiens OX=9606 GN=PGM3 PE=1 SV=1 | 13 |
| EFHD2 | EF-hand domain-containing protein D2 OS=Homo sapiens OX=9606 GN=EFHD2 PE=1 SV=1 | 4 |
| BCR | Breakpoint cluster region protein OS=Homo sapiens OX=9606 GN=BCR PE=1 SV=2 | 1 |
| MTCH2 | Mitochondrial carrier homolog 2 OS=Homo sapiens OX=9606 GN=MTCH2 PE=1 SV=1 | 8 |
| WIPF2 | WAS/WASL-interacting protein family member 2 OS=Homo sapiens OX=9606 GN=WIPF2 PE=1 SV=1 | 8 |
| EDF1 | Endothelial differentiation-related factor 1 OS=Homo sapiens OX=9606 GN=EDF1 PE=1 SV=1 | 4 |
| SND1 | Staphylococcal nuclease domain-containing protein 1 OS=Homo sapiens OX=9606 GN=SND1 PE=1 SV=1 | 22 |
| IQGAP1 | Ras GTPase-activating-like protein IQGAP1 OS=Homo sapiens OX=9606 GN=IQGAP1 PE=1 SV=1 | 28 |
| DPEP1 | Dipeptidase 1 OS=Homo sapiens OX=9606 GN=DPEP1 PE=1 SV=3 | 12 |
| WIPI2 | WD repeat domain phosphoinositide-interacting protein 2 OS=Homo sapiens OX=9606 GN=WIPI2 PE=1 SV=1 | 4 |
| MAT2A | S-adenosylmethionine synthase isoform type-2 OS=Homo sapiens OX=9606 GN=MAT2A PE=1 SV=1 | 8 |
| TRAF2 | TNF receptor-associated factor 2 OS=Homo sapiens OX=9606 GN=TRAF2 PE=1 SV=2 | 12 |
| UBE2S | Ubiquitin-conjugating enzyme E2 S OS=Homo sapiens OX=9606 GN=UBE2S PE=1 SV=2 | 1 |
| SMAD1 | Mothers against decapentaplegic homolog 1 OS=Homo sapiens OX=9606 GN=SMAD1 PE=1 SV=1 | 5 |
| SCYL1 | N-terminal kinase-like protein OS=Homo sapiens OX=9606 GN=SCYL1 PE=1 SV=1 | 14 |
| MYO1D | Unconventional myosin-Id OS=Homo sapiens OX=9606 GN=MYO1D PE=1 SV=2 | 11 |
| EIF4G1 | Eukaryotic translation initiation factor 4 gamma 1 OS=Homo sapiens OX=9606 GN=EIF4G1 PE=1 SV=4 | 25 |
| MTX1 | Metaxin-1 OS=Homo sapiens OX=9606 GN=MTX1 PE=1 SV=3 | 9 |
| CLIC4 | Chloride intracellular channel protein 4 OS=Homo sapiens OX=9606 GN=CLIC4 PE=1 SV=4 | 13 |
| XRCC6 | X-ray repair cross-complementing protein 6 OS=Homo sapiens OX=9606 GN=XRCC6 PE=1 SV=2 | 17 |
| PDHX | Pyruvate dehydrogenase protein X component, mitochondrial OS=Homo sapiens OX=9606 GN=PDHX PE=1 SV=3 | 17 |
| PYCR1 | Pyrroline-5-carboxylate reductase 1, mitochondrial OS=Homo sapiens OX=9606 GN=PYCR1 PE=1 SV=2 | 7 |
| VWA8 | von Willebrand factor A domain-containing protein 8 OS=Homo sapiens OX=9606 GN=VWA8 PE=1 SV=2 | 24 |
| OCRL | Inositol polyphosphate 5-phosphatase OCRL OS=Homo sapiens OX=9606 GN=OCRL PE=1 SV=3 | 7 |
| GLYR1 | Cytokine-like nuclear factor N-PAC OS=Homo sapiens OX=9606 GN=GLYR1 PE=1 SV=4 | 1 |
| RPL17 | 60S ribosomal protein L17 OS=Homo sapiens OX=9606 GN=RPL17 PE=1 SV=3 | 6 |
| HMGB3 | High mobility group protein B3 OS=Homo sapiens OX=9606 GN=HMGB3 PE=1 SV=4 | 6 |
| SERPINE2 | Glia-derived nexin OS=Homo sapiens OX=9606 GN=SERPINE2 PE=1 SV=1 | 14 |
| DGUOK | Deoxyguanosine kinase, mitochondrial OS=Homo sapiens OX=9606 GN=DGUOK PE=1 SV=2 | 5 |
| HUWE1 | E3 ubiquitin-protein ligase HUWE1 OS=Homo sapiens OX=9606 GN=HUWE1 PE=1 SV=3 | 22 |
| SAR1A | GTP-binding protein SAR1a OS=Homo sapiens OX=9606 GN=SAR1A PE=1 SV=1 | 9 |
| C4A | Complement C4-A OS=Homo sapiens OX=9606 GN=C4A PE=1 SV=2 | 30 |
| RDH14 | Retinol dehydrogenase 14 OS=Homo sapiens OX=9606 GN=RDH14 PE=1 SV=1 | 3 |
| TMOD3 | Tropomodulin-3 OS=Homo sapiens OX=9606 GN=TMOD3 PE=1 SV=1 | 3 |
| MARCKS | Myristoylated alanine-rich C-kinase substrate OS=Homo sapiens OX=9606 GN=MARCKS PE=1 SV=4 | 5 |
| REEP4 | Receptor expression-enhancing protein 4 OS=Homo sapiens OX=9606 GN=REEP4 PE=1 SV=1 | 2 |
| PANK2 | Pantothenate kinase 2, mitochondrial OS=Homo sapiens OX=9606 GN=PANK2 PE=1 SV=3 | 5 |
| CARS1 | Cysteine--tRNA ligase, cytoplasmic OS=Homo sapiens OX=9606 GN=CARS1 PE=1 SV=3 | 16 |
| NDUFB10 | NADH dehydrogenase [ubiquinone] 1 beta subcomplex subunit 10 OS=Homo sapiens OX=9606 GN=NDUFB10 PE=1 SV=3 | 5 |
| PPP6C | Serine/threonine-protein phosphatase 6 catalytic subunit OS=Homo sapiens OX=9606 GN=PPP6C PE=1 SV=1 | 5 |
| RPH3A | Rabphilin-3A OS=Homo sapiens OX=9606 GN=RPH3A PE=1 SV=1 | 7 |
| AP2M1 | AP-2 complex subunit mu OS=Homo sapiens OX=9606 GN=AP2M1 PE=1 SV=2 | 10 |
| WAPL | Wings apart-like protein homolog OS=Homo sapiens OX=9606 GN=WAPL PE=1 SV=1 | 3 |
| EIF4ENIF1 | Eukaryotic translation initiation factor 4E transporter OS=Homo sapiens OX=9606 GN=EIF4ENIF1 PE=1 SV=2 | 20 |
| ALDH16A1 | Aldehyde dehydrogenase family 16 member A1 OS=Homo sapiens OX=9606 GN=ALDH16A1 PE=1 SV=2 | 8 |
| TRMT112 | Multifunctional methyltransferase subunit TRM112-like protein OS=Homo sapiens OX=9606 GN=TRMT112 PE=1 SV=1 | 6 |
| VCP | Transitional endoplasmic reticulum ATPase OS=Homo sapiens OX=9606 GN=VCP PE=1 SV=4 | 50 |
| RPSA | 40S ribosomal protein SA OS=Homo sapiens OX=9606 GN=RPSA PE=1 SV=4 | 9 |
| RAP1GAP2 | Rap1 GTPase-activating protein 2 OS=Homo sapiens OX=9606 GN=RAP1GAP2 PE=1 SV=2 | 2 |
| FNTA | Protein farnesyltransferase/geranylgeranyltransferase type-1 subunit alpha OS=Homo sapiens OX=9606 GN=FNTA PE=1 SV=1 | 6 |
| SRPRA | Signal recognition particle receptor subunit alpha OS=Homo sapiens OX=9606 GN=SRPRA PE=1 SV=2 | 6 |
| CXorf38 | Uncharacterized protein CXorf38 OS=Homo sapiens OX=9606 GN=CXorf38 PE=1 SV=1 | 1 |
| NPC1 | NPC intracellular cholesterol transporter 1 OS=Homo sapiens OX=9606 GN=NPC1 PE=1 SV=2 | 4 |
| PPID | Peptidyl-prolyl cis-trans isomerase D OS=Homo sapiens OX=9606 GN=PPID PE=1 SV=3 | 13 |
| CORO1B | Coronin-1B OS=Homo sapiens OX=9606 GN=CORO1B PE=1 SV=1 | 2 |
| CENPJ | Centromere protein J OS=Homo sapiens OX=9606 GN=CENPJ PE=1 SV=2 | 1 |
| RCOR1 | REST corepressor 1 OS=Homo sapiens OX=9606 GN=RCOR1 PE=1 SV=2 | 3 |
| GINS3 | DNA replication complex GINS protein PSF3 OS=Homo sapiens OX=9606 GN=GINS3 PE=1 SV=1 | 1 |
| EIF3G | Eukaryotic translation initiation factor 3 subunit G OS=Homo sapiens OX=9606 GN=EIF3G PE=1 SV=2 | 4 |
| FAF2 | FAS-associated factor 2 OS=Homo sapiens OX=9606 GN=FAF2 PE=1 SV=2 | 5 |
| TRNT1 | CCA tRNA nucleotidyltransferase 1, mitochondrial OS=Homo sapiens OX=9606 GN=TRNT1 PE=1 SV=2 | 5 |
| PCYOX1 | Prenylcysteine oxidase 1 OS=Homo sapiens OX=9606 GN=PCYOX1 PE=1 SV=3 | 15 |
| ISCA2 | Iron-sulfur cluster assembly 2 homolog, mitochondrial OS=Homo sapiens OX=9606 GN=ISCA2 PE=1 SV=2 | 3 |
| MCMBP | Mini-chromosome maintenance complex-binding protein OS=Homo sapiens OX=9606 GN=MCMBP PE=1 SV=2 | 2 |
| NUP58 | Nucleoporin p58/p45 OS=Homo sapiens OX=9606 GN=NUP58 PE=1 SV=1 | 3 |
| RCCD1 | RCC1 domain-containing protein 1 OS=Homo sapiens OX=9606 GN=RCCD1 PE=1 SV=1 | 3 |
| VTA1 | Vacuolar protein sorting-associated protein VTA1 homolog OS=Homo sapiens OX=9606 GN=VTA1 PE=1 SV=1 | 8 |
| CYB5A | Cytochrome b5 OS=Homo sapiens OX=9606 GN=CYB5A PE=1 SV=2 | 4 |
| MEST | Mesoderm-specific transcript homolog protein OS=Homo sapiens OX=9606 GN=MEST PE=1 SV=2 | 1 |
| TSNAX | Translin-associated protein X OS=Homo sapiens OX=9606 GN=TSNAX PE=1 SV=1 | 13 |
| ANKZF1 | Ankyrin repeat and zinc finger domain-containing protein 1 OS=Homo sapiens OX=9606 GN=ANKZF1 PE=1 SV=1 | 1 |
| DDX20 | Probable ATP-dependent RNA helicase DDX20 OS=Homo sapiens OX=9606 GN=DDX20 PE=1 SV=2 | 1 |
| NUDT14 | Uridine diphosphate glucose pyrophosphatase NUDT14 OS=Homo sapiens OX=9606 GN=NUDT14 PE=1 SV=2 | 7 |
| NDUFS8 | NADH dehydrogenase [ubiquinone] iron-sulfur protein 8, mitochondrial OS=Homo sapiens OX=9606 GN=NDUFS8 PE=1 SV=1 | 4 |
| VTI1A | Vesicle transport through interaction with t-SNAREs homolog 1A OS=Homo sapiens OX=9606 GN=VTI1A PE=1 SV=2 | 2 |
| GLG1 | Golgi apparatus protein 1 OS=Homo sapiens OX=9606 GN=GLG1 PE=1 SV=2 | 21 |
| LDHD | Probable D-lactate dehydrogenase, mitochondrial OS=Homo sapiens OX=9606 GN=LDHD PE=1 SV=1 | 4 |
| SDF4 | 45 kDa calcium-binding protein OS=Homo sapiens OX=9606 GN=SDF4 PE=1 SV=1 | 4 |
| THOC3 | THO complex subunit 3 OS=Homo sapiens OX=9606 GN=THOC3 PE=1 SV=1 | 5 |
| CLTCL1 | Clathrin heavy chain 2 OS=Homo sapiens OX=9606 GN=CLTCL1 PE=1 SV=2 | 1 |
| SRP68 | Signal recognition particle subunit SRP68 OS=Homo sapiens OX=9606 GN=SRP68 PE=1 SV=2 | 7 |
| RPS20 | 40S ribosomal protein S20 OS=Homo sapiens OX=9606 GN=RPS20 PE=1 SV=1 | 6 |
| SRI | Sorcin OS=Homo sapiens OX=9606 GN=SRI PE=1 SV=1 | 8 |
| ACAD9 | Complex I assembly factor ACAD9, mitochondrial OS=Homo sapiens OX=9606 GN=ACAD9 PE=1 SV=1 | 15 |
| DDHD1 | Phospholipase DDHD1 OS=Homo sapiens OX=9606 GN=DDHD1 PE=1 SV=2 | 9 |
| CDK5RAP3 | CDK5 regulatory subunit-associated protein 3 OS=Homo sapiens OX=9606 GN=CDK5RAP3 PE=1 SV=2 | 5 |
| RPL28 | 60S ribosomal protein L28 OS=Homo sapiens OX=9606 GN=RPL28 PE=1 SV=3 | 4 |
| RPS11 | 40S ribosomal protein S11 OS=Homo sapiens OX=9606 GN=RPS11 PE=1 SV=3 | 7 |
| PFDN1 | Prefoldin subunit 1 OS=Homo sapiens OX=9606 GN=PFDN1 PE=1 SV=2 | 6 |
| CDV3 | Protein CDV3 homolog OS=Homo sapiens OX=9606 GN=CDV3 PE=1 SV=1 | 7 |
| BAIAP2L1 | Brain-specific angiogenesis inhibitor 1-associated protein 2-like protein 1 OS=Homo sapiens OX=9606 GN=BAIAP2L1 PE=1 SV=2 | 10 |
| RAPH1 | Ras-associated and pleckstrin homology domains-containing protein 1 OS=Homo sapiens OX=9606 GN=RAPH1 PE=1 SV=3 | 6 |
| AGTRAP | Type-1 angiotensin II receptor-associated protein OS=Homo sapiens OX=9606 GN=AGTRAP PE=1 SV=1 | 2 |
| RALBP1 | RalA-binding protein 1 OS=Homo sapiens OX=9606 GN=RALBP1 PE=1 SV=3 | 1 |
| CACNA2D1 | Voltage-dependent calcium channel subunit alpha-2/delta-1 OS=Homo sapiens OX=9606 GN=CACNA2D1 PE=1 SV=3 | 9 |
| EPRS1 | Bifunctional glutamate/proline--tRNA ligase OS=Homo sapiens OX=9606 GN=EPRS1 PE=1 SV=5 | 40 |
| TMED4 | Transmembrane emp24 domain-containing protein 4 OS=Homo sapiens OX=9606 GN=TMED4 PE=1 SV=1 | 3 |
| VIL1 | Villin-1 OS=Homo sapiens OX=9606 GN=VIL1 PE=1 SV=4 | 5 |
| PDAP1 | 28 kDa heat- and acid-stable phosphoprotein OS=Homo sapiens OX=9606 GN=PDAP1 PE=1 SV=1 | 7 |
| ECHS1 | Enoyl-CoA hydratase, mitochondrial OS=Homo sapiens OX=9606 GN=ECHS1 PE=1 SV=4 | 17 |
| EEF1E1 | Eukaryotic translation elongation factor 1 epsilon-1 OS=Homo sapiens OX=9606 GN=EEF1E1 PE=1 SV=1 | 2 |
| RAD23B | UV excision repair protein RAD23 homolog B OS=Homo sapiens OX=9606 GN=RAD23B PE=1 SV=1 | 8 |
| ARRB2 | Beta-arrestin-2 OS=Homo sapiens OX=9606 GN=ARRB2 PE=1 SV=2 | 1 |
| MAP2K2 | Dual specificity mitogen-activated protein kinase kinase 2 OS=Homo sapiens OX=9606 GN=MAP2K2 PE=1 SV=1 | 10 |
| AP2B1 | AP-2 complex subunit beta OS=Homo sapiens OX=9606 GN=AP2B1 PE=1 SV=1 | 12 |
| MICOS10 | MICOS complex subunit MIC10 OS=Homo sapiens OX=9606 GN=MICOS10 PE=1 SV=1 | 1 |
| HNRNPL | Heterogeneous nuclear ribonucleoprotein L OS=Homo sapiens OX=9606 GN=HNRNPL PE=1 SV=2 | 7 |
| CDH2 | Cadherin-2 OS=Homo sapiens OX=9606 GN=CDH2 PE=1 SV=4 | 5 |
| C7orf25 | UPF0415 protein C7orf25 OS=Homo sapiens OX=9606 GN=C7orf25 PE=1 SV=1 | 2 |
| FBXO6 | F-box only protein 6 OS=Homo sapiens OX=9606 GN=FBXO6 PE=1 SV=1 | 2 |
| DUT | Deoxyuridine 5'-triphosphate nucleotidohydrolase, mitochondrial OS=Homo sapiens OX=9606 GN=DUT PE=1 SV=4 | 6 |
| HDGF | Hepatoma-derived growth factor OS=Homo sapiens OX=9606 GN=HDGF PE=1 SV=1 | 4 |
| STX4 | Syntaxin-4 OS=Homo sapiens OX=9606 GN=STX4 PE=1 SV=2 | 9 |
| NELL1 | Protein kinase C-binding protein NELL1 OS=Homo sapiens OX=9606 GN=NELL1 PE=1 SV=4 | 5 |
| LIG1 | DNA ligase 1 OS=Homo sapiens OX=9606 GN=LIG1 PE=1 SV=1 | 2 |
| VPS4B | Vacuolar protein sorting-associated protein 4B OS=Homo sapiens OX=9606 GN=VPS4B PE=1 SV=2 | 7 |
| NPEPPS | Puromycin-sensitive aminopeptidase OS=Homo sapiens OX=9606 GN=NPEPPS PE=1 SV=2 | 34 |
| RPLP1 | 60S acidic ribosomal protein P1 OS=Homo sapiens OX=9606 GN=RPLP1 PE=1 SV=1 | 1 |
| KHK | Ketohexokinase OS=Homo sapiens OX=9606 GN=KHK PE=1 SV=2 | 12 |
| ATIC | Bifunctional purine biosynthesis protein ATIC OS=Homo sapiens OX=9606 GN=ATIC PE=1 SV=3 | 33 |
| MRPS26 | 28S ribosomal protein S26, mitochondrial OS=Homo sapiens OX=9606 GN=MRPS26 PE=1 SV=1 | 2 |
| NHLRC3 | NHL repeat-containing protein 3 OS=Homo sapiens OX=9606 GN=NHLRC3 PE=2 SV=1 | 2 |
| RMND1 | Required for meiotic nuclear division protein 1 homolog OS=Homo sapiens OX=9606 GN=RMND1 PE=1 SV=2 | 7 |
| TMX3 | Protein disulfide-isomerase TMX3 OS=Homo sapiens OX=9606 GN=TMX3 PE=1 SV=2 | 4 |
| ACTA1 | Actin, alpha skeletal muscle OS=Homo sapiens OX=9606 GN=ACTA1 PE=1 SV=1 | 3 |
| NDUFA4 | Cytochrome c oxidase subunit NDUFA4 OS=Homo sapiens OX=9606 GN=NDUFA4 PE=1 SV=1 | 2 |
| MAPK12 | Mitogen-activated protein kinase 12 OS=Homo sapiens OX=9606 GN=MAPK12 PE=1 SV=3 | 2 |
| VPS25 | Vacuolar protein-sorting-associated protein 25 OS=Homo sapiens OX=9606 GN=VPS25 PE=1 SV=1 | 5 |
| RABGGTA | Geranylgeranyl transferase type-2 subunit alpha OS=Homo sapiens OX=9606 GN=RABGGTA PE=1 SV=2 | 10 |
| MVK | Mevalonate kinase OS=Homo sapiens OX=9606 GN=MVK PE=1 SV=1 | 12 |
| PRDX5 | Peroxiredoxin-5, mitochondrial OS=Homo sapiens OX=9606 GN=PRDX5 PE=1 SV=4 | 13 |
| TRAPPC4 | Trafficking protein particle complex subunit 4 OS=Homo sapiens OX=9606 GN=TRAPPC4 PE=1 SV=1 | 6 |
| AIMP1 | Aminoacyl tRNA synthase complex-interacting multifunctional protein 1 OS=Homo sapiens OX=9606 GN=AIMP1 PE=1 SV=2 | 9 |
| MT-ATP6 | ATP synthase subunit a OS=Homo sapiens OX=9606 GN=MT-ATP6 PE=1 SV=1 | 1 |
| OXSM | 3-oxoacyl-[acyl-carrier-protein] synthase, mitochondrial OS=Homo sapiens OX=9606 GN=OXSM PE=1 SV=1 | 4 |
| NAXD | ATP-dependent (S)-NAD(P)H-hydrate dehydratase OS=Homo sapiens OX=9606 GN=NAXD PE=1 SV=1 | 8 |
| SLC25A20 | Mitochondrial carnitine/acylcarnitine carrier protein OS=Homo sapiens OX=9606 GN=SLC25A20 PE=1 SV=1 | 3 |
| TMEM123 | Porimin OS=Homo sapiens OX=9606 GN=TMEM123 PE=1 SV=1 | 2 |
| ERLIN1 | Erlin-1 OS=Homo sapiens OX=9606 GN=ERLIN1 PE=1 SV=2 | 7 |
| GNPAT | Dihydroxyacetone phosphate acyltransferase OS=Homo sapiens OX=9606 GN=GNPAT PE=1 SV=1 | 1 |
| MRPL41 | 39S ribosomal protein L41, mitochondrial OS=Homo sapiens OX=9606 GN=MRPL41 PE=1 SV=1 | 5 |
| PRXL2B | Prostamide/prostaglandin F synthase OS=Homo sapiens OX=9606 GN=PRXL2B PE=1 SV=1 | 1 |
| NUDT9 | ADP-ribose pyrophosphatase, mitochondrial OS=Homo sapiens OX=9606 GN=NUDT9 PE=1 SV=1 | 3 |
| PNPT1 | Polyribonucleotide nucleotidyltransferase 1, mitochondrial OS=Homo sapiens OX=9606 GN=PNPT1 PE=1 SV=2 | 7 |
| MTRES1 | Mitochondrial transcription rescue factor 1 OS=Homo sapiens OX=9606 GN=MTRES1 PE=1 SV=1 | 3 |
| OGDHL | 2-oxoglutarate dehydrogenase-like, mitochondrial OS=Homo sapiens OX=9606 GN=OGDHL PE=1 SV=3 | 19 |
| NDUFV1 | NADH dehydrogenase [ubiquinone] flavoprotein 1, mitochondrial OS=Homo sapiens OX=9606 GN=NDUFV1 PE=1 SV=4 | 12 |
| TATDN1 | Deoxyribonuclease TATDN1 OS=Homo sapiens OX=9606 GN=TATDN1 PE=1 SV=2 | 12 |
| RPS6 | 40S ribosomal protein S6 OS=Homo sapiens OX=9606 GN=RPS6 PE=1 SV=1 | 10 |
| SBF1 | Myotubularin-related protein 5 OS=Homo sapiens OX=9606 GN=SBF1 PE=1 SV=4 | 2 |
| PARP1 | Poly [ADP-ribose] polymerase 1 OS=Homo sapiens OX=9606 GN=PARP1 PE=1 SV=4 | 15 |
| SUMF2 | Inactive C-alpha-formylglycine-generating enzyme 2 OS=Homo sapiens OX=9606 GN=SUMF2 PE=1 SV=2 | 6 |
| COMMD9 | COMM domain-containing protein 9 OS=Homo sapiens OX=9606 GN=COMMD9 PE=1 SV=2 | 4 |
| C9orf40 | Uncharacterized protein C9orf40 OS=Homo sapiens OX=9606 GN=C9orf40 PE=1 SV=1 | 2 |
| SERBP1 | Plasminogen activator inhibitor 1 RNA-binding protein OS=Homo sapiens OX=9606 GN=SERBP1 PE=1 SV=2 | 3 |
| DCTN6 | Dynactin subunit 6 OS=Homo sapiens OX=9606 GN=DCTN6 PE=1 SV=1 | 2 |
| PSMA5 | Proteasome subunit alpha type-5 OS=Homo sapiens OX=9606 GN=PSMA5 PE=1 SV=3 | 9 |
| LYPLAL1 | Lysophospholipase-like protein 1 OS=Homo sapiens OX=9606 GN=LYPLAL1 PE=1 SV=3 | 8 |
| ARPC5L | Actin-related protein 2/3 complex subunit 5-like protein OS=Homo sapiens OX=9606 GN=ARPC5L PE=1 SV=1 | 6 |
| STRAP | Serine-threonine kinase receptor-associated protein OS=Homo sapiens OX=9606 GN=STRAP PE=1 SV=1 | 15 |
| HNRNPH1 | Heterogeneous nuclear ribonucleoprotein H OS=Homo sapiens OX=9606 GN=HNRNPH1 PE=1 SV=4 | 7 |
| WDR54 | WD repeat-containing protein 54 OS=Homo sapiens OX=9606 GN=WDR54 PE=1 SV=1 | 5 |
| AP2A1 | AP-2 complex subunit alpha-1 OS=Homo sapiens OX=9606 GN=AP2A1 PE=1 SV=3 | 11 |
| MIA2 | Melanoma inhibitory activity protein 2 OS=Homo sapiens OX=9606 GN=MIA2 PE=1 SV=4 | 26 |
| FEN1 | Flap endonuclease 1 OS=Homo sapiens OX=9606 GN=FEN1 PE=1 SV=1 | 6 |
| CADM1 | Cell adhesion molecule 1 OS=Homo sapiens OX=9606 GN=CADM1 PE=1 SV=2 | 2 |
| TBC1D15 | TBC1 domain family member 15 OS=Homo sapiens OX=9606 GN=TBC1D15 PE=1 SV=2 | 15 |
| COX6C | Cytochrome c oxidase subunit 6C OS=Homo sapiens OX=9606 GN=COX6C PE=1 SV=2 | 4 |
| SLC9A3R1 | Na(+)/H(+) exchange regulatory cofactor NHE-RF1 OS=Homo sapiens OX=9606 GN=NHERF1 PE=1 SV=4 | 16 |
| PITRM1 | Presequence protease, mitochondrial OS=Homo sapiens OX=9606 GN=PITRM1 PE=1 SV=3 | 52 |
| GNB4 | Guanine nucleotide-binding protein subunit beta-4 OS=Homo sapiens OX=9606 GN=GNB4 PE=1 SV=3 | 5 |
| ERI3 | ERI1 exoribonuclease 3 OS=Homo sapiens OX=9606 GN=ERI3 PE=1 SV=2 | 5 |
| FASTKD2 | FAST kinase domain-containing protein 2, mitochondrial OS=Homo sapiens OX=9606 GN=FASTKD2 PE=1 SV=1 | 5 |
| COPZ1 | Coatomer subunit zeta-1 OS=Homo sapiens OX=9606 GN=COPZ1 PE=1 SV=1 | 3 |
| NDUFA8 | NADH dehydrogenase [ubiquinone] 1 alpha subcomplex subunit 8 OS=Homo sapiens OX=9606 GN=NDUFA8 PE=1 SV=3 | 4 |
| ATG4C | Cysteine protease ATG4C OS=Homo sapiens OX=9606 GN=ATG4C PE=1 SV=1 | 6 |
| UBAC2 | Ubiquitin-associated domain-containing protein 2 OS=Homo sapiens OX=9606 GN=UBAC2 PE=1 SV=1 | 2 |
| HSPA2 | Heat shock-related 70 kDa protein 2 OS=Homo sapiens OX=9606 GN=HSPA2 PE=1 SV=1 | 17 |
| CEP97 | Centrosomal protein of 97 kDa OS=Homo sapiens OX=9606 GN=CEP97 PE=1 SV=1 | 2 |
| MIX23 | Protein MIX23 OS=Homo sapiens OX=9606 GN=MIX23 PE=1 SV=1 | 6 |
| EIF3I | Eukaryotic translation initiation factor 3 subunit I OS=Homo sapiens OX=9606 GN=EIF3I PE=1 SV=1 | 13 |
| ZNRD2 | Protein ZNRD2 OS=Homo sapiens OX=9606 GN=ZNRD2 PE=1 SV=1 | 1 |
| GSDME | Gasdermin-E OS=Homo sapiens OX=9606 GN=GSDME PE=1 SV=2 | 17 |
| VPS11 | Vacuolar protein sorting-associated protein 11 homolog OS=Homo sapiens OX=9606 GN=VPS11 PE=1 SV=1 | 3 |
| SLBP | Histone RNA hairpin-binding protein OS=Homo sapiens OX=9606 GN=SLBP PE=1 SV=1 | 6 |
| RPA1 | Replication protein A 70 kDa DNA-binding subunit OS=Homo sapiens OX=9606 GN=RPA1 PE=1 SV=2 | 18 |
| IFT27 | Intraflagellar transport protein 27 homolog OS=Homo sapiens OX=9606 GN=IFT27 PE=1 SV=1 | 2 |
| EIF1AX | Eukaryotic translation initiation factor 1A, X-chromosomal OS=Homo sapiens OX=9606 GN=EIF1AX PE=1 SV=2 | 7 |
| EMC1 | ER membrane protein complex subunit 1 OS=Homo sapiens OX=9606 GN=EMC1 PE=1 SV=1 | 6 |
| MYH9 | Myosin-9 OS=Homo sapiens OX=9606 GN=MYH9 PE=1 SV=4 | 109 |
| SH3GLB2 | Endophilin-B2 OS=Homo sapiens OX=9606 GN=SH3GLB2 PE=1 SV=1 | 14 |
| COMMD7 | COMM domain-containing protein 7 OS=Homo sapiens OX=9606 GN=COMMD7 PE=1 SV=2 | 3 |
| EML5 | Echinoderm microtubule-associated protein-like 5 OS=Homo sapiens OX=9606 GN=EML5 PE=2 SV=3 | 1 |
| GCLC | Glutamate--cysteine ligase catalytic subunit OS=Homo sapiens OX=9606 GN=GCLC PE=1 SV=2 | 12 |
| B4GALT3 | Beta-1,4-galactosyltransferase 3 OS=Homo sapiens OX=9606 GN=B4GALT3 PE=1 SV=2 | 1 |
| CAPRIN2 | Caprin-2 OS=Homo sapiens OX=9606 GN=CAPRIN2 PE=1 SV=1 | 30 |
| TGM2 | Protein-glutamine gamma-glutamyltransferase 2 OS=Homo sapiens OX=9606 GN=TGM2 PE=1 SV=2 | 1 |
| EIF3D | Eukaryotic translation initiation factor 3 subunit D OS=Homo sapiens OX=9606 GN=EIF3D PE=1 SV=1 | 8 |
| TFG | Protein TFG OS=Homo sapiens OX=9606 GN=TFG PE=1 SV=2 | 8 |
| RPL6 | 60S ribosomal protein L6 OS=Homo sapiens OX=9606 GN=RPL6 PE=1 SV=3 | 10 |
| ADI1 | Acireductone dioxygenase OS=Homo sapiens OX=9606 GN=ADI1 PE=1 SV=1 | 3 |
| DNAJB4 | DnaJ homolog subfamily B member 4 OS=Homo sapiens OX=9606 GN=DNAJB4 PE=1 SV=1 | 3 |
| -.23 | Uncharacterized protein OS=Homo sapiens OX=9606 PE=4 SV=1 | 1 |
| GARS1 | Glycine--tRNA ligase OS=Homo sapiens OX=9606 GN=GARS1 PE=1 SV=3 | 26 |
| RAB10 | Ras-related protein Rab-10 OS=Homo sapiens OX=9606 GN=RAB10 PE=1 SV=1 | 7 |
| CERS6 | Ceramide synthase 6 OS=Homo sapiens OX=9606 GN=CERS6 PE=1 SV=1 | 1 |
| EML4 | Echinoderm microtubule-associated protein-like 4 OS=Homo sapiens OX=9606 GN=EML4 PE=1 SV=3 | 18 |
| CARS2 | Probable cysteine--tRNA ligase, mitochondrial OS=Homo sapiens OX=9606 GN=CARS2 PE=1 SV=1 | 3 |
| MPZL1 | Myelin protein zero-like protein 1 OS=Homo sapiens OX=9606 GN=MPZL1 PE=1 SV=1 | 1 |
| SLC25A10 | Mitochondrial dicarboxylate carrier OS=Homo sapiens OX=9606 GN=SLC25A10 PE=1 SV=2 | 2 |
| JPH2 | Junctophilin-2 OS=Homo sapiens OX=9606 GN=JPH2 PE=1 SV=2 | 1 |
| GPX4 | Phospholipid hydroperoxide glutathione peroxidase OS=Homo sapiens OX=9606 GN=GPX4 PE=1 SV=3 | 6 |
| MYDGF | Myeloid-derived growth factor OS=Homo sapiens OX=9606 GN=MYDGF PE=1 SV=1 | 7 |
| ETFRF1 | Electron transfer flavoprotein regulatory factor 1 OS=Homo sapiens OX=9606 GN=ETFRF1 PE=1 SV=2 | 2 |
| RRBP1 | Ribosome-binding protein 1 OS=Homo sapiens OX=9606 GN=RRBP1 PE=1 SV=5 | 26 |
| MRPS25 | 28S ribosomal protein S25, mitochondrial OS=Homo sapiens OX=9606 GN=MRPS25 PE=1 SV=1 | 7 |
| POLR2A | DNA-directed RNA polymerase II subunit RPB1 OS=Homo sapiens OX=9606 GN=POLR2A PE=1 SV=2 | 1 |
| TIMM8A | Mitochondrial import inner membrane translocase subunit Tim8 A OS=Homo sapiens OX=9606 GN=TIMM8A PE=1 SV=1 | 6 |
| HMGCL | Hydroxymethylglutaryl-CoA lyase, mitochondrial OS=Homo sapiens OX=9606 GN=HMGCL PE=1 SV=2 | 13 |
| YAP1 | Transcriptional coactivator YAP1 OS=Homo sapiens OX=9606 GN=YAP1 PE=1 SV=2 | 1 |
| ALDOC | Fructose-bisphosphate aldolase C OS=Homo sapiens OX=9606 GN=ALDOC PE=1 SV=2 | 3 |
| PABPN1 | Polyadenylate-binding protein 2 OS=Homo sapiens OX=9606 GN=PABPN1 PE=1 SV=3 | 2 |
| UQCRQ | Cytochrome b-c1 complex subunit 8 OS=Homo sapiens OX=9606 GN=UQCRQ PE=1 SV=4 | 5 |
| GOLGA3 | Golgin subfamily A member 3 OS=Homo sapiens OX=9606 GN=GOLGA3 PE=1 SV=2 | 22 |
| GNPDA2 | Glucosamine-6-phosphate isomerase 2 OS=Homo sapiens OX=9606 GN=GNPDA2 PE=1 SV=1 | 19 |
| CIRBP | Cold-inducible RNA-binding protein OS=Homo sapiens OX=9606 GN=CIRBP PE=1 SV=1 | 8 |
| CCDC50 | Coiled-coil domain-containing protein 50 OS=Homo sapiens OX=9606 GN=CCDC50 PE=1 SV=1 | 4 |
| FBLN1 | Fibulin-1 OS=Homo sapiens OX=9606 GN=FBLN1 PE=1 SV=4 | 12 |
| PECAM1 | Platelet endothelial cell adhesion molecule OS=Homo sapiens OX=9606 GN=PECAM1 PE=1 SV=2 | 3 |
| MYO1C | Unconventional myosin-Ic OS=Homo sapiens OX=9606 GN=MYO1C PE=1 SV=4 | 1 |
| NDUFB8 | NADH dehydrogenase [ubiquinone] 1 beta subcomplex subunit 8, mitochondrial OS=Homo sapiens OX=9606 GN=NDUFB8 PE=1 SV=1 | 3 |
| ITGA6 | Integrin alpha-6 OS=Homo sapiens OX=9606 GN=ITGA6 PE=1 SV=5 | 23 |
| ARCN1 | Coatomer subunit delta OS=Homo sapiens OX=9606 GN=ARCN1 PE=1 SV=1 | 34 |
| CCNI2 | Cyclin-I2 OS=Homo sapiens OX=9606 GN=CCNI2 PE=2 SV=1 | 18 |
| APOOL | MICOS complex subunit MIC27 OS=Homo sapiens OX=9606 GN=APOOL PE=1 SV=1 | 6 |
| RPL31 | 60S ribosomal protein L31 OS=Homo sapiens OX=9606 GN=RPL31 PE=1 SV=1 | 4 |
| YKT6 | Synaptobrevin homolog YKT6 OS=Homo sapiens OX=9606 GN=YKT6 PE=1 SV=1 | 5 |
| SYPL1 | Synaptophysin-like protein 1 OS=Homo sapiens OX=9606 GN=SYPL1 PE=1 SV=1 | 5 |
| FUBP1 | Far upstream element-binding protein 1 OS=Homo sapiens OX=9606 GN=FUBP1 PE=1 SV=3 | 14 |
| DNM2 | Dynamin-2 OS=Homo sapiens OX=9606 GN=DNM2 PE=1 SV=2 | 16 |
| FGFR1 | Fibroblast growth factor receptor 1 OS=Homo sapiens OX=9606 GN=FGFR1 PE=1 SV=3 | 1 |
| G3BP1 | Ras GTPase-activating protein-binding protein 1 OS=Homo sapiens OX=9606 GN=G3BP1 PE=1 SV=1 | 6 |
| ERICH5 | Glutamate-rich protein 5 OS=Homo sapiens OX=9606 GN=ERICH5 PE=1 SV=1 | 1 |
| ABI1 | Abl interactor 1 OS=Homo sapiens OX=9606 GN=ABI1 PE=1 SV=4 | 3 |
| ESD | S-formylglutathione hydrolase OS=Homo sapiens OX=9606 GN=ESD PE=1 SV=2 | 10 |
| NAGK | N-acetyl-D-glucosamine kinase OS=Homo sapiens OX=9606 GN=NAGK PE=1 SV=4 | 9 |
| AARS1 | Alanine--tRNA ligase, cytoplasmic OS=Homo sapiens OX=9606 GN=AARS1 PE=1 SV=2 | 26 |
| EMC3 | ER membrane protein complex subunit 3 OS=Homo sapiens OX=9606 GN=EMC3 PE=1 SV=3 | 4 |
| NECAP1 | Adaptin ear-binding coat-associated protein 1 OS=Homo sapiens OX=9606 GN=NECAP1 PE=1 SV=2 | 7 |
| CTSH | Pro-cathepsin H OS=Homo sapiens OX=9606 GN=CTSH PE=1 SV=4 | 2 |
| SNAP29 | Synaptosomal-associated protein 29 OS=Homo sapiens OX=9606 GN=SNAP29 PE=1 SV=1 | 6 |
| CAPZA2 | F-actin-capping protein subunit alpha-2 OS=Homo sapiens OX=9606 GN=CAPZA2 PE=1 SV=3 | 11 |
| RAB27A | Ras-related protein Rab-27A OS=Homo sapiens OX=9606 GN=RAB27A PE=1 SV=3 | 7 |
| SDHA | Succinate dehydrogenase [ubiquinone] flavoprotein subunit, mitochondrial OS=Homo sapiens OX=9606 GN=SDHA PE=1 SV=2 | 20 |
| EIF3E | Eukaryotic translation initiation factor 3 subunit E OS=Homo sapiens OX=9606 GN=EIF3E PE=1 SV=1 | 8 |
| OSBPL8 | Oxysterol-binding protein-related protein 8 OS=Homo sapiens OX=9606 GN=OSBPL8 PE=1 SV=3 | 12 |
| FLNB | Filamin-B OS=Homo sapiens OX=9606 GN=FLNB PE=1 SV=2 | 103 |
| IGF2BP2 | Insulin-like growth factor 2 mRNA-binding protein 2 OS=Homo sapiens OX=9606 GN=IGF2BP2 PE=1 SV=2 | 13 |
| CDC123 | Cell division cycle protein 123 homolog OS=Homo sapiens OX=9606 GN=CDC123 PE=1 SV=1 | 1 |
| ARPC3 | Actin-related protein 2/3 complex subunit 3 OS=Homo sapiens OX=9606 GN=ARPC3 PE=1 SV=3 | 10 |
| NADSYN1 | Glutamine-dependent NAD(+) synthetase OS=Homo sapiens OX=9606 GN=NADSYN1 PE=1 SV=3 | 8 |
| ENOSF1 | Mitochondrial enolase superfamily member 1 OS=Homo sapiens OX=9606 GN=ENOSF1 PE=1 SV=1 | 7 |
| MRPS21 | 28S ribosomal protein S21, mitochondrial OS=Homo sapiens OX=9606 GN=MRPS21 PE=1 SV=3 | 1 |
| DNLZ | DNL-type zinc finger protein OS=Homo sapiens OX=9606 GN=DNLZ PE=1 SV=1 | 2 |
| RPL32 | 60S ribosomal protein L32 OS=Homo sapiens OX=9606 GN=RPL32 PE=1 SV=2 | 5 |
| IGF2R | Cation-independent mannose-6-phosphate receptor OS=Homo sapiens OX=9606 GN=IGF2R PE=1 SV=3 | 5 |
| STX6 | Syntaxin-6 OS=Homo sapiens OX=9606 GN=STX6 PE=1 SV=1 | 7 |
| HOOK3 | Protein Hook homolog 3 OS=Homo sapiens OX=9606 GN=HOOK3 PE=1 SV=2 | 8 |
| ITPR3 | Inositol 1,4,5-trisphosphate receptor type 3 OS=Homo sapiens OX=9606 GN=ITPR3 PE=1 SV=2 | 1 |
| POLR1C | DNA-directed RNA polymerases I and III subunit RPAC1 OS=Homo sapiens OX=9606 GN=POLR1C PE=1 SV=1 | 3 |
| RPS7 | 40S ribosomal protein S7 OS=Homo sapiens OX=9606 GN=RPS7 PE=1 SV=1 | 8 |
| DDX19A | ATP-dependent RNA helicase DDX19A OS=Homo sapiens OX=9606 GN=DDX19A PE=1 SV=1 | 5 |
| FN1 | Fibronectin OS=Homo sapiens OX=9606 GN=FN1 PE=1 SV=5 | 57 |
| COX5B | Cytochrome c oxidase subunit 5B, mitochondrial OS=Homo sapiens OX=9606 GN=COX5B PE=1 SV=2 | 6 |
| IDH1 | Isocitrate dehydrogenase [NADP] cytoplasmic OS=Homo sapiens OX=9606 GN=IDH1 PE=1 SV=2 | 21 |
| FREM2 | FRAS1-related extracellular matrix protein 2 OS=Homo sapiens OX=9606 GN=FREM2 PE=1 SV=3 | 1 |
| NECAP2 | Adaptin ear-binding coat-associated protein 2 OS=Homo sapiens OX=9606 GN=NECAP2 PE=1 SV=1 | 2 |
| CCDC124 | Coiled-coil domain-containing protein 124 OS=Homo sapiens OX=9606 GN=CCDC124 PE=1 SV=1 | 11 |
| HACL1 | 2-hydroxyacyl-CoA lyase 1 OS=Homo sapiens OX=9606 GN=HACL1 PE=1 SV=2 | 7 |
| COPB2 | Coatomer subunit beta' OS=Homo sapiens OX=9606 GN=COPB2 PE=1 SV=2 | 24 |
| HNRNPH3 | Heterogeneous nuclear ribonucleoprotein H3 OS=Homo sapiens OX=9606 GN=HNRNPH3 PE=1 SV=2 | 5 |
| EIF3K | Eukaryotic translation initiation factor 3 subunit K OS=Homo sapiens OX=9606 GN=EIF3K PE=1 SV=1 | 5 |
| ARL3 | ADP-ribosylation factor-like protein 3 OS=Homo sapiens OX=9606 GN=ARL3 PE=1 SV=2 | 12 |
| NMT1 | Glycylpeptide N-tetradecanoyltransferase 1 OS=Homo sapiens OX=9606 GN=NMT1 PE=1 SV=2 | 4 |
| ACTN1 | Alpha-actinin-1 OS=Homo sapiens OX=9606 GN=ACTN1 PE=1 SV=2 | 19 |
| EEF1A2 | Elongation factor 1-alpha 2 OS=Homo sapiens OX=9606 GN=EEF1A2 PE=1 SV=1 | 16 |
| PDXDC1 | Pyridoxal-dependent decarboxylase domain-containing protein 1 OS=Homo sapiens OX=9606 GN=PDXDC1 PE=1 SV=2 | 24 |
| ATP5IF1 | ATPase inhibitor, mitochondrial OS=Homo sapiens OX=9606 GN=ATP5IF1 PE=1 SV=1 | 6 |
| NUDT7 | Peroxisomal coenzyme A diphosphatase NUDT7 OS=Homo sapiens OX=9606 GN=NUDT7 PE=1 SV=1 | 2 |
| RAB1A | Ras-related protein Rab-1A OS=Homo sapiens OX=9606 GN=RAB1A PE=1 SV=3 | 17 |
| CHMP2B | Charged multivesicular body protein 2b OS=Homo sapiens OX=9606 GN=CHMP2B PE=1 SV=1 | 4 |
| DNAJC7 | DnaJ homolog subfamily C member 7 OS=Homo sapiens OX=9606 GN=DNAJC7 PE=1 SV=2 | 9 |
| TIAM1 | Rho guanine nucleotide exchange factor TIAM1 OS=Homo sapiens OX=9606 GN=TIAM1 PE=1 SV=2 | 8 |
| ST3GAL4 | CMP-N-acetylneuraminate-beta-galactosamide-alpha-2,3-sialyltransferase 4 OS=Homo sapiens OX=9606 GN=ST3GAL4 PE=1 SV=1 | 4 |
| MRPL49 | 39S ribosomal protein L49, mitochondrial OS=Homo sapiens OX=9606 GN=MRPL49 PE=1 SV=1 | 3 |
| CSTF1 | Cleavage stimulation factor subunit 1 OS=Homo sapiens OX=9606 GN=CSTF1 PE=1 SV=1 | 3 |
| CDK5 | Cyclin-dependent kinase 5 OS=Homo sapiens OX=9606 GN=CDK5 PE=1 SV=3 | 19 |
| TALDO1 | Transaldolase OS=Homo sapiens OX=9606 GN=TALDO1 PE=1 SV=2 | 15 |
| ACOX1 | Peroxisomal acyl-coenzyme A oxidase 1 OS=Homo sapiens OX=9606 GN=ACOX1 PE=1 SV=3 | 7 |
| OGFR | Opioid growth factor receptor OS=Homo sapiens OX=9606 GN=OGFR PE=1 SV=3 | 4 |
| SEC22B | Vesicle-trafficking protein SEC22b OS=Homo sapiens OX=9606 GN=SEC22B PE=1 SV=5 | 13 |
| RARRES2 | Retinoic acid receptor responder protein 2 OS=Homo sapiens OX=9606 GN=RARRES2 PE=1 SV=1 | 6 |
| VARS1 | Valine--tRNA ligase OS=Homo sapiens OX=9606 GN=VARS1 PE=1 SV=4 | 16 |
| ACAT2 | Acetyl-CoA acetyltransferase, cytosolic OS=Homo sapiens OX=9606 GN=ACAT2 PE=1 SV=2 | 16 |
| MCU | Calcium uniporter protein, mitochondrial OS=Homo sapiens OX=9606 GN=MCU PE=1 SV=1 | 6 |
| SETSIP | Protein SETSIP OS=Homo sapiens OX=9606 GN=SETSIP PE=1 SV=1 | 6 |
| HSPE1 | 10 kDa heat shock protein, mitochondrial OS=Homo sapiens OX=9606 GN=HSPE1 PE=1 SV=2 | 15 |
| PPP1R2 | Protein phosphatase inhibitor 2 OS=Homo sapiens OX=9606 GN=PPP1R2 PE=1 SV=2 | 5 |
| ADAM15 | Disintegrin and metalloproteinase domain-containing protein 15 OS=Homo sapiens OX=9606 GN=ADAM15 PE=1 SV=4 | 1 |
| LAMTOR3 | Ragulator complex protein LAMTOR3 OS=Homo sapiens OX=9606 GN=LAMTOR3 PE=1 SV=1 | 1 |
| THAP4 | Peroxynitrite isomerase THAP4 OS=Homo sapiens OX=9606 GN=THAP4 PE=1 SV=2 | 2 |
| TOR1AIP1 | Torsin-1A-interacting protein 1 OS=Homo sapiens OX=9606 GN=TOR1AIP1 PE=1 SV=2 | 3 |
| PSMG3 | Proteasome assembly chaperone 3 OS=Homo sapiens OX=9606 GN=PSMG3 PE=1 SV=1 | 4 |
| ALDOA | Fructose-bisphosphate aldolase A OS=Homo sapiens OX=9606 GN=ALDOA PE=1 SV=2 | 26 |
| PNPO | Pyridoxine-5'-phosphate oxidase OS=Homo sapiens OX=9606 GN=PNPO PE=1 SV=1 | 1 |
| MORF4L1 | Mortality factor 4-like protein 1 OS=Homo sapiens OX=9606 GN=MORF4L1 PE=1 SV=2 | 2 |
| WDR41 | WD repeat-containing protein 41 OS=Homo sapiens OX=9606 GN=WDR41 PE=1 SV=3 | 1 |
| RPL34 | 60S ribosomal protein L34 OS=Homo sapiens OX=9606 GN=RPL34 PE=1 SV=3 | 2 |
| CFL1 | Cofilin-1 OS=Homo sapiens OX=9606 GN=CFL1 PE=1 SV=3 | 16 |
| RIDA | 2-iminobutanoate/2-iminopropanoate deaminase OS=Homo sapiens OX=9606 GN=RIDA PE=1 SV=1 | 1 |
| NDUFB11 | NADH dehydrogenase [ubiquinone] 1 beta subcomplex subunit 11, mitochondrial OS=Homo sapiens OX=9606 GN=NDUFB11 PE=1 SV=1 | 2 |
| ICAM1 | Intercellular adhesion molecule 1 OS=Homo sapiens OX=9606 GN=ICAM1 PE=1 SV=2 | 16 |
| C4orf48 | Neuropeptide-like protein C4orf48 OS=Homo sapiens OX=9606 GN=C4orf48 PE=1 SV=3 | 3 |
| CLINT1 | Clathrin interactor 1 OS=Homo sapiens OX=9606 GN=CLINT1 PE=1 SV=1 | 14 |
| BCAP29 | B-cell receptor-associated protein 29 OS=Homo sapiens OX=9606 GN=BCAP29 PE=1 SV=2 | 3 |
| ETF1 | Eukaryotic peptide chain release factor subunit 1 OS=Homo sapiens OX=9606 GN=ETF1 PE=1 SV=3 | 15 |
| DCUN1D5 | DCN1-like protein 5 OS=Homo sapiens OX=9606 GN=DCUN1D5 PE=1 SV=1 | 4 |
| APOB | Apolipoprotein B-100 OS=Homo sapiens OX=9606 GN=APOB PE=1 SV=2 | 128 |
| GAS7 | Growth arrest-specific protein 7 OS=Homo sapiens OX=9606 GN=GAS7 PE=1 SV=3 | 5 |
| MRPL50 | 39S ribosomal protein L50, mitochondrial OS=Homo sapiens OX=9606 GN=MRPL50 PE=1 SV=2 | 2 |
| FAM98C | Protein FAM98C OS=Homo sapiens OX=9606 GN=FAM98C PE=1 SV=1 | 1 |
| SH3GL1 | Endophilin-A2 OS=Homo sapiens OX=9606 GN=SH3GL1 PE=1 SV=1 | 2 |
| CDH1 | Cadherin-1 OS=Homo sapiens OX=9606 GN=CDH1 PE=1 SV=3 | 2 |
| GET4 | Golgi to ER traffic protein 4 homolog OS=Homo sapiens OX=9606 GN=GET4 PE=1 SV=1 | 6 |
| FDX2 | Ferredoxin-2, mitochondrial OS=Homo sapiens OX=9606 GN=FDX2 PE=1 SV=2 | 2 |
| TNFAIP8 | Tumor necrosis factor alpha-induced protein 8 OS=Homo sapiens OX=9606 GN=TNFAIP8 PE=1 SV=1 | 2 |
| GCHFR | GTP cyclohydrolase 1 feedback regulatory protein OS=Homo sapiens OX=9606 GN=GCHFR PE=1 SV=3 | 5 |
| SYNCRIP | Heterogeneous nuclear ribonucleoprotein Q OS=Homo sapiens OX=9606 GN=SYNCRIP PE=1 SV=2 | 19 |
| JAG1 | Protein jagged-1 OS=Homo sapiens OX=9606 GN=JAG1 PE=1 SV=3 | 2 |
| NME1 | Nucleoside diphosphate kinase A OS=Homo sapiens OX=9606 GN=NME1 PE=1 SV=1 | 10 |
| PPP1R7 | Protein phosphatase 1 regulatory subunit 7 OS=Homo sapiens OX=9606 GN=PPP1R7 PE=1 SV=1 | 18 |
| FOLR1 | Folate receptor alpha OS=Homo sapiens OX=9606 GN=FOLR1 PE=1 SV=3 | 7 |
| RMDN1 | Regulator of microtubule dynamics protein 1 OS=Homo sapiens OX=9606 GN=RMDN1 PE=1 SV=1 | 8 |
| GSPT1 | Eukaryotic peptide chain release factor GTP-binding subunit ERF3A OS=Homo sapiens OX=9606 GN=GSPT1 PE=1 SV=1 | 22 |
| PTPN1 | Tyrosine-protein phosphatase non-receptor type 1 OS=Homo sapiens OX=9606 GN=PTPN1 PE=1 SV=1 | 15 |
| DAG1 | Dystroglycan 1 OS=Homo sapiens OX=9606 GN=DAG1 PE=1 SV=2 | 3 |
| PLEKHB1 | Pleckstrin homology domain-containing family B member 1 OS=Homo sapiens OX=9606 GN=PLEKHB1 PE=1 SV=1 | 1 |
| MLF2 | Myeloid leukemia factor 2 OS=Homo sapiens OX=9606 GN=MLF2 PE=1 SV=1 | 5 |
| VWA5A | von Willebrand factor A domain-containing protein 5A OS=Homo sapiens OX=9606 GN=VWA5A PE=1 SV=2 | 26 |
| LNPEP | Leucyl-cystinyl aminopeptidase OS=Homo sapiens OX=9606 GN=LNPEP PE=1 SV=3 | 8 |
| GADD45GIP1 | Growth arrest and DNA damage-inducible proteins-interacting protein 1 OS=Homo sapiens OX=9606 GN=GADD45GIP1 PE=1 SV=1 | 3 |
| NDUFS1 | NADH-ubiquinone oxidoreductase 75 kDa subunit, mitochondrial OS=Homo sapiens OX=9606 GN=NDUFS1 PE=1 SV=3 | 24 |
| KLHL18 | Kelch-like protein 18 OS=Homo sapiens OX=9606 GN=KLHL18 PE=1 SV=3 | 3 |
| UBE2V1 | Ubiquitin-conjugating enzyme E2 variant 1 OS=Homo sapiens OX=9606 GN=UBE2V1 PE=1 SV=2 | 2 |
| HSD17B4 | Peroxisomal multifunctional enzyme type 2 OS=Homo sapiens OX=9606 GN=HSD17B4 PE=1 SV=3 | 36 |
| FAM3C | Protein FAM3C OS=Homo sapiens OX=9606 GN=FAM3C PE=1 SV=1 | 4 |
| NAE1 | NEDD8-activating enzyme E1 regulatory subunit OS=Homo sapiens OX=9606 GN=NAE1 PE=1 SV=1 | 8 |
| SURF1 | Surfeit locus protein 1 OS=Homo sapiens OX=9606 GN=SURF1 PE=1 SV=1 | 4 |
| ACSL4 | Long-chain-fatty-acid--CoA ligase 4 OS=Homo sapiens OX=9606 GN=ACSL4 PE=1 SV=2 | 7 |
| UMAD1 | UBAP1-MVB12-associated (UMA)-domain containing protein 1 OS=Homo sapiens OX=9606 GN=UMAD1 PE=1 SV=2 | 2 |
| IGF2BP3 | Insulin-like growth factor 2 mRNA-binding protein 3 OS=Homo sapiens OX=9606 GN=IGF2BP3 PE=1 SV=2 | 28 |
| FARS2 | Phenylalanine--tRNA ligase, mitochondrial OS=Homo sapiens OX=9606 GN=FARS2 PE=1 SV=1 | 3 |
| ACOT9 | Acyl-coenzyme A thioesterase 9, mitochondrial OS=Homo sapiens OX=9606 GN=ACOT9 PE=1 SV=2 | 30 |
| APPL1 | DCC-interacting protein 13-alpha OS=Homo sapiens OX=9606 GN=APPL1 PE=1 SV=1 | 19 |
| NUDT21 | Cleavage and polyadenylation specificity factor subunit 5 OS=Homo sapiens OX=9606 GN=NUDT21 PE=1 SV=1 | 3 |
| EFNB1 | Ephrin-B1 OS=Homo sapiens OX=9606 GN=EFNB1 PE=1 SV=1 | 1 |
| RPS23 | 40S ribosomal protein S23 OS=Homo sapiens OX=9606 GN=RPS23 PE=1 SV=3 | 3 |
| RPS27L | 40S ribosomal protein S27-like OS=Homo sapiens OX=9606 GN=RPS27L PE=1 SV=3 | 3 |
| SUPT16H | FACT complex subunit SPT16 OS=Homo sapiens OX=9606 GN=SUPT16H PE=1 SV=1 | 2 |
| SLC4A10 | Sodium-driven chloride bicarbonate exchanger OS=Homo sapiens OX=9606 GN=SLC4A10 PE=1 SV=1 | 1 |
| DSTN | Destrin OS=Homo sapiens OX=9606 GN=DSTN PE=1 SV=3 | 12 |
| AK3 | GTP:AMP phosphotransferase AK3, mitochondrial OS=Homo sapiens OX=9606 GN=AK3 PE=1 SV=4 | 12 |
| YIPF5 | Protein YIPF5 OS=Homo sapiens OX=9606 GN=YIPF5 PE=1 SV=1 | 2 |
| SPPL2A | Signal peptide peptidase-like 2A OS=Homo sapiens OX=9606 GN=SPPL2A PE=1 SV=2 | 1 |
| NUP54 | Nucleoporin p54 OS=Homo sapiens OX=9606 GN=NUP54 PE=1 SV=2 | 9 |
| SLC25A15 | Mitochondrial ornithine transporter 1 OS=Homo sapiens OX=9606 GN=SLC25A15 PE=1 SV=1 | 5 |
| TIMM50 | Mitochondrial import inner membrane translocase subunit TIM50 OS=Homo sapiens OX=9606 GN=TIMM50 PE=1 SV=2 | 4 |
| TACC2 | Transforming acidic coiled-coil-containing protein 2 OS=Homo sapiens OX=9606 GN=TACC2 PE=1 SV=3 | 1 |
| ABCF3 | ATP-binding cassette sub-family F member 3 OS=Homo sapiens OX=9606 GN=ABCF3 PE=1 SV=2 | 6 |
| EMC4 | ER membrane protein complex subunit 4 OS=Homo sapiens OX=9606 GN=EMC4 PE=1 SV=2 | 2 |
| GSTO1 | Glutathione S-transferase omega-1 OS=Homo sapiens OX=9606 GN=GSTO1 PE=1 SV=2 | 6 |
| PDP1 | [Pyruvate dehydrogenase [acetyl-transferring]]-phosphatase 1, mitochondrial OS=Homo sapiens OX=9606 GN=PDP1 PE=1 SV=3 | 11 |
| NDUFAF1 | Complex I intermediate-associated protein 30, mitochondrial OS=Homo sapiens OX=9606 GN=NDUFAF1 PE=1 SV=2 | 6 |
| TWF1 | Twinfilin-1 OS=Homo sapiens OX=9606 GN=TWF1 PE=1 SV=3 | 5 |
| GLRX5 | Glutaredoxin-related protein 5, mitochondrial OS=Homo sapiens OX=9606 GN=GLRX5 PE=1 SV=2 | 3 |
| COX6B1 | Cytochrome c oxidase subunit 6B1 OS=Homo sapiens OX=9606 GN=COX6B1 PE=1 SV=2 | 3 |
| KYAT3 | Kynurenine--oxoglutarate transaminase 3 OS=Homo sapiens OX=9606 GN=KYAT3 PE=1 SV=1 | 6 |
| L2HGDH | L-2-hydroxyglutarate dehydrogenase, mitochondrial OS=Homo sapiens OX=9606 GN=L2HGDH PE=1 SV=3 | 11 |
| HCLS1 | Hematopoietic lineage cell-specific protein OS=Homo sapiens OX=9606 GN=HCLS1 PE=1 SV=3 | 1 |
| ABCE1 | ATP-binding cassette sub-family E member 1 OS=Homo sapiens OX=9606 GN=ABCE1 PE=1 SV=1 | 12 |
| PYCR2 | Pyrroline-5-carboxylate reductase 2 OS=Homo sapiens OX=9606 GN=PYCR2 PE=1 SV=1 | 9 |
| TRAPPC1 | Trafficking protein particle complex subunit 1 OS=Homo sapiens OX=9606 GN=TRAPPC1 PE=1 SV=1 | 4 |
| HSPBP1 | Hsp70-binding protein 1 OS=Homo sapiens OX=9606 GN=HSPBP1 PE=1 SV=2 | 7 |
| GTPBP3 | tRNA modification GTPase GTPBP3, mitochondrial OS=Homo sapiens OX=9606 GN=GTPBP3 PE=1 SV=2 | 2 |
| PLK1 | Serine/threonine-protein kinase PLK1 OS=Homo sapiens OX=9606 GN=PLK1 PE=1 SV=1 | 5 |
| REEP5 | Receptor expression-enhancing protein 5 OS=Homo sapiens OX=9606 GN=REEP5 PE=1 SV=3 | 3 |
| STAU1 | Double-stranded RNA-binding protein Staufen homolog 1 OS=Homo sapiens OX=9606 GN=STAU1 PE=1 SV=2 | 10 |
| HMGB1P1 | Putative high mobility group protein B1-like 1 OS=Homo sapiens OX=9606 GN=HMGB1P1 PE=5 SV=1 | 3 |
| MRPS17 | 28S ribosomal protein S17, mitochondrial OS=Homo sapiens OX=9606 GN=MRPS17 PE=1 SV=1 | 2 |
| POLG | DNA polymerase subunit gamma-1 OS=Homo sapiens OX=9606 GN=POLG PE=1 SV=1 | 12 |
| ARMC10 | Armadillo repeat-containing protein 10 OS=Homo sapiens OX=9606 GN=ARMC10 PE=1 SV=1 | 6 |
| HAUS8 | HAUS augmin-like complex subunit 8 OS=Homo sapiens OX=9606 GN=HAUS8 PE=1 SV=3 | 1 |
| FKBP4 | Peptidyl-prolyl cis-trans isomerase FKBP4 OS=Homo sapiens OX=9606 GN=FKBP4 PE=1 SV=3 | 35 |
| PIN4 | Peptidyl-prolyl cis-trans isomerase NIMA-interacting 4 OS=Homo sapiens OX=9606 GN=PIN4 PE=1 SV=1 | 4 |
| PABPC4 | Polyadenylate-binding protein 4 OS=Homo sapiens OX=9606 GN=PABPC4 PE=1 SV=1 | 20 |
| NONO | Non-POU domain-containing octamer-binding protein OS=Homo sapiens OX=9606 GN=NONO PE=1 SV=4 | 3 |
| PARL | Presenilins-associated rhomboid-like protein, mitochondrial OS=Homo sapiens OX=9606 GN=PARL PE=1 SV=2 | 8 |
| SLC30A1 | Proton-coupled zinc antiporter SLC30A1 OS=Homo sapiens OX=9606 GN=SLC30A1 PE=1 SV=3 | 1 |
| DNAJC2 | DnaJ homolog subfamily C member 2 OS=Homo sapiens OX=9606 GN=DNAJC2 PE=1 SV=4 | 1 |
| GNA13 | Guanine nucleotide-binding protein subunit alpha-13 OS=Homo sapiens OX=9606 GN=GNA13 PE=1 SV=2 | 3 |
| NEK1 | Serine/threonine-protein kinase Nek1 OS=Homo sapiens OX=9606 GN=NEK1 PE=1 SV=2 | 10 |
| VPS45 | Vacuolar protein sorting-associated protein 45 OS=Homo sapiens OX=9606 GN=VPS45 PE=1 SV=1 | 7 |
| GFM1 | Elongation factor G, mitochondrial OS=Homo sapiens OX=9606 GN=GFM1 PE=1 SV=2 | 31 |
| RBM12 | RNA-binding protein 12 OS=Homo sapiens OX=9606 GN=RBM12 PE=1 SV=1 | 5 |
| ZAR1L | Protein ZAR1-like OS=Homo sapiens OX=9606 GN=ZAR1L PE=2 SV=2 | 7 |
| RPL23A | 60S ribosomal protein L23a OS=Homo sapiens OX=9606 GN=RPL23A PE=1 SV=1 | 9 |
| ATP6V1E1 | V-type proton ATPase subunit E 1 OS=Homo sapiens OX=9606 GN=ATP6V1E1 PE=1 SV=1 | 11 |
| UQCRH | Cytochrome b-c1 complex subunit 6, mitochondrial OS=Homo sapiens OX=9606 GN=UQCRH PE=1 SV=2 | 3 |
| ASPSCR1 | Tether containing UBX domain for GLUT4 OS=Homo sapiens OX=9606 GN=ASPSCR1 PE=1 SV=1 | 14 |
| TACO1 | Translational activator of cytochrome c oxidase 1 OS=Homo sapiens OX=9606 GN=TACO1 PE=1 SV=1 | 3 |
| AKR1B1 | Aldo-keto reductase family 1 member B1 OS=Homo sapiens OX=9606 GN=AKR1B1 PE=1 SV=3 | 22 |
| STX18 | Syntaxin-18 OS=Homo sapiens OX=9606 GN=STX18 PE=1 SV=1 | 4 |
| CRYBG1 | Beta/gamma crystallin domain-containing protein 1 OS=Homo sapiens OX=9606 GN=CRYBG1 PE=1 SV=3 | 9 |
| TUBG2 | Tubulin gamma-2 chain OS=Homo sapiens OX=9606 GN=TUBG2 PE=2 SV=1 | 2 |
| HNRNPA3 | Heterogeneous nuclear ribonucleoprotein A3 OS=Homo sapiens OX=9606 GN=HNRNPA3 PE=1 SV=2 | 8 |
| PGGT1B | Geranylgeranyl transferase type-1 subunit beta OS=Homo sapiens OX=9606 GN=PGGT1B PE=1 SV=2 | 1 |
| HLA-C | HLA class I histocompatibility antigen, C alpha chain OS=Homo sapiens OX=9606 GN=HLA-C PE=1 SV=3 | 1 |
| EIF3L | Eukaryotic translation initiation factor 3 subunit L OS=Homo sapiens OX=9606 GN=EIF3L PE=1 SV=1 | 11 |
| PPL | Periplakin OS=Homo sapiens OX=9606 GN=PPL PE=1 SV=4 | 11 |
| LIAS | Lipoyl synthase, mitochondrial OS=Homo sapiens OX=9606 GN=LIAS PE=1 SV=3 | 1 |
| ME2 | NAD-dependent malic enzyme, mitochondrial OS=Homo sapiens OX=9606 GN=ME2 PE=1 SV=1 | 3 |
| PCCB | Propionyl-CoA carboxylase beta chain, mitochondrial OS=Homo sapiens OX=9606 GN=PCCB PE=1 SV=3 | 4 |
| CFAP298 | Cilia- and flagella-associated protein 298 OS=Homo sapiens OX=9606 GN=CFAP298 PE=1 SV=1 | 1 |
| SLC39A10 | Zinc transporter ZIP10 OS=Homo sapiens OX=9606 GN=SLC39A10 PE=1 SV=2 | 1 |
| DHCR24 | Delta(24)-sterol reductase OS=Homo sapiens OX=9606 GN=DHCR24 PE=1 SV=2 | 8 |
| PFDN4 | Prefoldin subunit 4 OS=Homo sapiens OX=9606 GN=PFDN4 PE=1 SV=1 | 4 |
| SSR3 | Translocon-associated protein subunit gamma OS=Homo sapiens OX=9606 GN=SSR3 PE=1 SV=1 | 1 |
| CAMK2G | Calcium/calmodulin-dependent protein kinase type II subunit gamma OS=Homo sapiens OX=9606 GN=CAMK2G PE=1 SV=4 | 2 |
| KRT84 | Keratin, type II cuticular Hb4 OS=Homo sapiens OX=9606 GN=KRT84 PE=2 SV=2 | 24 |
| PRKAG1 | 5'-AMP-activated protein kinase subunit gamma-1 OS=Homo sapiens OX=9606 GN=PRKAG1 PE=1 SV=1 | 2 |
| NDUFS4 | NADH dehydrogenase [ubiquinone] iron-sulfur protein 4, mitochondrial OS=Homo sapiens OX=9606 GN=NDUFS4 PE=1 SV=1 | 3 |
| INPP1 | Inositol polyphosphate 1-phosphatase OS=Homo sapiens OX=9606 GN=INPP1 PE=1 SV=1 | 14 |
| GRSF1 | G-rich sequence factor 1 OS=Homo sapiens OX=9606 GN=GRSF1 PE=1 SV=3 | 14 |
| MTRF1L | Peptide chain release factor 1-like, mitochondrial OS=Homo sapiens OX=9606 GN=MTRF1L PE=1 SV=1 | 2 |
| IST1 | IST1 homolog OS=Homo sapiens OX=9606 GN=IST1 PE=1 SV=1 | 6 |
| ISOC2 | Isochorismatase domain-containing protein 2 OS=Homo sapiens OX=9606 GN=ISOC2 PE=1 SV=1 | 6 |
| NAA38 | N-alpha-acetyltransferase 38, NatC auxiliary subunit OS=Homo sapiens OX=9606 GN=NAA38 PE=1 SV=1 | 1 |
| ERO1A | ERO1-like protein alpha OS=Homo sapiens OX=9606 GN=ERO1A PE=1 SV=2 | 2 |
| PAICS | Bifunctional phosphoribosylaminoimidazole carboxylase/phosphoribosylaminoimidazole succinocarboxamide synthetase OS=Homo sapiens OX=9606 GN=PAICS PE=1 SV=3 | 15 |
| ZC3H15 | Zinc finger CCCH domain-containing protein 15 OS=Homo sapiens OX=9606 GN=ZC3H15 PE=1 SV=1 | 7 |
| SEPTIN10 | Septin-10 OS=Homo sapiens OX=9606 GN=SEPTIN10 PE=1 SV=2 | 4 |
| NACA | Nascent polypeptide-associated complex subunit alpha, muscle-specific form OS=Homo sapiens OX=9606 GN=NACA PE=1 SV=1 | 6 |
| MYOCOS | Myocilin opposite strand protein OS=Homo sapiens OX=9606 GN=MYOCOS PE=4 SV=2 | 4 |
| SLC16A1 | Monocarboxylate transporter 1 OS=Homo sapiens OX=9606 GN=SLC16A1 PE=1 SV=3 | 2 |
| PPIL1 | Peptidyl-prolyl cis-trans isomerase-like 1 OS=Homo sapiens OX=9606 GN=PPIL1 PE=1 SV=1 | 4 |
| DPYSL2 | Dihydropyrimidinase-related protein 2 OS=Homo sapiens OX=9606 GN=DPYSL2 PE=1 SV=1 | 22 |
| KIAA2013 | Uncharacterized protein KIAA2013 OS=Homo sapiens OX=9606 GN=KIAA2013 PE=1 SV=1 | 2 |
| MRPS35 | 28S ribosomal protein S35, mitochondrial OS=Homo sapiens OX=9606 GN=MRPS35 PE=1 SV=1 | 3 |
| PCYT2 | Ethanolamine-phosphate cytidylyltransferase OS=Homo sapiens OX=9606 GN=PCYT2 PE=1 SV=1 | 7 |
| SPTBN1 | Spectrin beta chain, non-erythrocytic 1 OS=Homo sapiens OX=9606 GN=SPTBN1 PE=1 SV=2 | 28 |
| COL28A1 | Collagen alpha-1(XXVIII) chain OS=Homo sapiens OX=9606 GN=COL28A1 PE=2 SV=2 | 1 |
| HOOK1 | Protein Hook homolog 1 OS=Homo sapiens OX=9606 GN=HOOK1 PE=1 SV=2 | 19 |
| PDE6D | Retinal rod rhodopsin-sensitive cGMP 3',5'-cyclic phosphodiesterase subunit delta OS=Homo sapiens OX=9606 GN=PDE6D PE=1 SV=1 | 2 |
| TYW5 | tRNA wybutosine-synthesizing protein 5 OS=Homo sapiens OX=9606 GN=TYW5 PE=1 SV=1 | 5 |
| LSM2 | U6 snRNA-associated Sm-like protein LSm2 OS=Homo sapiens OX=9606 GN=LSM2 PE=1 SV=1 | 8 |
| RPL7A | 60S ribosomal protein L7a OS=Homo sapiens OX=9606 GN=RPL7A PE=1 SV=2 | 11 |
| THBS1 | Thrombospondin-1 OS=Homo sapiens OX=9606 GN=THBS1 PE=1 SV=2 | 10 |
| GSR | Glutathione reductase, mitochondrial OS=Homo sapiens OX=9606 GN=GSR PE=1 SV=2 | 17 |
| PAK1 | Serine/threonine-protein kinase PAK 1 OS=Homo sapiens OX=9606 GN=PAK1 PE=1 SV=2 | 9 |
| FAM8A1 | Protein FAM8A1 OS=Homo sapiens OX=9606 GN=FAM8A1 PE=1 SV=1 | 1 |
| RPL35A | 60S ribosomal protein L35a OS=Homo sapiens OX=9606 GN=RPL35A PE=1 SV=2 | 3 |
| PLCB3 | 1-phosphatidylinositol 4,5-bisphosphate phosphodiesterase beta-3 OS=Homo sapiens OX=9606 GN=PLCB3 PE=1 SV=2 | 2 |
| PPFIBP2 | Liprin-beta-2 OS=Homo sapiens OX=9606 GN=PPFIBP2 PE=1 SV=3 | 3 |
| ZDHHC13 | Palmitoyltransferase ZDHHC13 OS=Homo sapiens OX=9606 GN=ZDHHC13 PE=1 SV=3 | 5 |
| NAA10 | N-alpha-acetyltransferase 10 OS=Homo sapiens OX=9606 GN=NAA10 PE=1 SV=1 | 4 |
| RPL29 | 60S ribosomal protein L29 OS=Homo sapiens OX=9606 GN=RPL29 PE=1 SV=2 | 3 |
| LSM3 | U6 snRNA-associated Sm-like protein LSm3 OS=Homo sapiens OX=9606 GN=LSM3 PE=1 SV=2 | 3 |
| SSBP1 | Single-stranded DNA-binding protein, mitochondrial OS=Homo sapiens OX=9606 GN=SSBP1 PE=1 SV=1 | 10 |
| WTAP | Pre-mRNA-splicing regulator WTAP OS=Homo sapiens OX=9606 GN=WTAP PE=1 SV=2 | 2 |
| HARS1 | Histidine--tRNA ligase, cytoplasmic OS=Homo sapiens OX=9606 GN=HARS1 PE=1 SV=2 | 24 |
| HDHD3 | Haloacid dehalogenase-like hydrolase domain-containing protein 3 OS=Homo sapiens OX=9606 GN=HDHD3 PE=1 SV=1 | 6 |
| ADSS2 | Adenylosuccinate synthetase isozyme 2 OS=Homo sapiens OX=9606 GN=ADSS2 PE=1 SV=3 | 5 |
| ARF6 | ADP-ribosylation factor 6 OS=Homo sapiens OX=9606 GN=ARF6 PE=1 SV=2 | 5 |
| CABYR | Calcium-binding tyrosine phosphorylation-regulated protein OS=Homo sapiens OX=9606 GN=CABYR PE=1 SV=2 | 5 |
| YARS1 | Tyrosine--tRNA ligase, cytoplasmic OS=Homo sapiens OX=9606 GN=YARS1 PE=1 SV=4 | 10 |
| ACSL1 | Long-chain-fatty-acid--CoA ligase 1 OS=Homo sapiens OX=9606 GN=ACSL1 PE=1 SV=1 | 14 |
| HNRNPA2B1 | Heterogeneous nuclear ribonucleoproteins A2/B1 OS=Homo sapiens OX=9606 GN=HNRNPA2B1 PE=1 SV=2 | 13 |
| SNX5 | Sorting nexin-5 OS=Homo sapiens OX=9606 GN=SNX5 PE=1 SV=1 | 12 |
| GNB1 | Guanine nucleotide-binding protein G(I)/G(S)/G(T) subunit beta-1 OS=Homo sapiens OX=9606 GN=GNB1 PE=1 SV=3 | 12 |
| RAP2B | Ras-related protein Rap-2b OS=Homo sapiens OX=9606 GN=RAP2B PE=1 SV=1 | 4 |
| GHITM | Growth hormone-inducible transmembrane protein OS=Homo sapiens OX=9606 GN=GHITM PE=1 SV=2 | 1 |
| PDIA5 | Protein disulfide-isomerase A5 OS=Homo sapiens OX=9606 GN=PDIA5 PE=1 SV=1 | 14 |
| PSMA6 | Proteasome subunit alpha type-6 OS=Homo sapiens OX=9606 GN=PSMA6 PE=1 SV=1 | 14 |
| CORO1C | Coronin-1C OS=Homo sapiens OX=9606 GN=CORO1C PE=1 SV=1 | 5 |
| BASP1 | Brain acid soluble protein 1 OS=Homo sapiens OX=9606 GN=BASP1 PE=1 SV=2 | 8 |
| USP47 | Ubiquitin carboxyl-terminal hydrolase 47 OS=Homo sapiens OX=9606 GN=USP47 PE=1 SV=3 | 7 |
| NEB | Nebulin OS=Homo sapiens OX=9606 GN=NEB PE=1 SV=5 | 3 |
| MRPL37 | 39S ribosomal protein L37, mitochondrial OS=Homo sapiens OX=9606 GN=MRPL37 PE=1 SV=2 | 1 |
| EIF2B5 | Translation initiation factor eIF-2B subunit epsilon OS=Homo sapiens OX=9606 GN=EIF2B5 PE=1 SV=3 | 1 |
| GART | Trifunctional purine biosynthetic protein adenosine-3 OS=Homo sapiens OX=9606 GN=GART PE=1 SV=1 | 3 |
| AFDN | Afadin OS=Homo sapiens OX=9606 GN=AFDN PE=1 SV=3 | 12 |
| ADD2 | Beta-adducin OS=Homo sapiens OX=9606 GN=ADD2 PE=1 SV=3 | 4 |
| FXR1 | RNA-binding protein FXR1 OS=Homo sapiens OX=9606 GN=FXR1 PE=1 SV=3 | 15 |
| RBPMS2 | RNA-binding protein with multiple splicing 2 OS=Homo sapiens OX=9606 GN=RBPMS2 PE=1 SV=1 | 2 |
| BORCS7 | BLOC-1-related complex subunit 7 OS=Homo sapiens OX=9606 GN=BORCS7 PE=3 SV=2 | 2 |
| MRM3 | rRNA methyltransferase 3, mitochondrial OS=Homo sapiens OX=9606 GN=MRM3 PE=1 SV=2 | 4 |
| SAMHD1 | Deoxynucleoside triphosphate triphosphohydrolase SAMHD1 OS=Homo sapiens OX=9606 GN=SAMHD1 PE=1 SV=2 | 4 |
| TIMP2 | Metalloproteinase inhibitor 2 OS=Homo sapiens OX=9606 GN=TIMP2 PE=1 SV=2 | 5 |
| FAU | FAU ubiquitin-like and ribosomal protein S30 OS=Homo sapiens OX=9606 GN=FAU PE=1 SV=2 | 2 |
| SEPTIN7 | Septin-7 OS=Homo sapiens OX=9606 GN=SEPTIN7 PE=1 SV=2 | 10 |
| AP3S1 | AP-3 complex subunit sigma-1 OS=Homo sapiens OX=9606 GN=AP3S1 PE=1 SV=1 | 2 |
| SUMO2 | Small ubiquitin-related modifier 2 OS=Homo sapiens OX=9606 GN=SUMO2 PE=1 SV=3 | 2 |
| ACBD3 | Golgi resident protein GCP60 OS=Homo sapiens OX=9606 GN=ACBD3 PE=1 SV=4 | 7 |
| BLOC1S3 | Biogenesis of lysosome-related organelles complex 1 subunit 3 OS=Homo sapiens OX=9606 GN=BLOC1S3 PE=1 SV=1 | 2 |
| ECHDC3 | Enoyl-CoA hydratase domain-containing protein 3, mitochondrial OS=Homo sapiens OX=9606 GN=ECHDC3 PE=1 SV=2 | 4 |
| CADPS2 | Calcium-dependent secretion activator 2 OS=Homo sapiens OX=9606 GN=CADPS2 PE=1 SV=2 | 4 |
| RABGGTB | Geranylgeranyl transferase type-2 subunit beta OS=Homo sapiens OX=9606 GN=RABGGTB PE=1 SV=2 | 1 |
| ATL2 | Atlastin-2 OS=Homo sapiens OX=9606 GN=ATL2 PE=1 SV=2 | 8 |
| MYL6B | Myosin light chain 6B OS=Homo sapiens OX=9606 GN=MYL6B PE=1 SV=1 | 7 |
| IDI1 | Isopentenyl-diphosphate Delta-isomerase 1 OS=Homo sapiens OX=9606 GN=IDI1 PE=1 SV=2 | 5 |
| RANBP1 | Ran-specific GTPase-activating protein OS=Homo sapiens OX=9606 GN=RANBP1 PE=1 SV=1 | 3 |
| KIT | Mast/stem cell growth factor receptor Kit OS=Homo sapiens OX=9606 GN=KIT PE=1 SV=1 | 3 |
| DBI | Acyl-CoA-binding protein OS=Homo sapiens OX=9606 GN=DBI PE=1 SV=2 | 5 |
| FAM98A | Protein FAM98A OS=Homo sapiens OX=9606 GN=FAM98A PE=1 SV=2 | 2 |
| RPL26 | 60S ribosomal protein L26 OS=Homo sapiens OX=9606 GN=RPL26 PE=1 SV=1 | 6 |
| BAIAP2 | Brain-specific angiogenesis inhibitor 1-associated protein 2 OS=Homo sapiens OX=9606 GN=BAIAP2 PE=1 SV=1 | 9 |
| ERH | Enhancer of rudimentary homolog OS=Homo sapiens OX=9606 GN=ERH PE=1 SV=1 | 2 |
| SEC16A | Protein transport protein Sec16A OS=Homo sapiens OX=9606 GN=SEC16A PE=1 SV=4 | 9 |
| SNX3 | Sorting nexin-3 OS=Homo sapiens OX=9606 GN=SNX3 PE=1 SV=3 | 3 |
| CBX3 | Chromobox protein homolog 3 OS=Homo sapiens OX=9606 GN=CBX3 PE=1 SV=4 | 5 |
| PDCD10 | Programmed cell death protein 10 OS=Homo sapiens OX=9606 GN=PDCD10 PE=1 SV=1 | 6 |
| EXOC8 | Exocyst complex component 8 OS=Homo sapiens OX=9606 GN=EXOC8 PE=1 SV=2 | 3 |
| NRAS | GTPase NRas OS=Homo sapiens OX=9606 GN=NRAS PE=1 SV=1 | 6 |
| ATP6AP1 | V-type proton ATPase subunit S1 OS=Homo sapiens OX=9606 GN=ATP6AP1 PE=1 SV=2 | 2 |
| CD99 | CD99 antigen OS=Homo sapiens OX=9606 GN=CD99 PE=1 SV=1 | 2 |
| SFXN1 | Sideroflexin-1 OS=Homo sapiens OX=9606 GN=SFXN1 PE=1 SV=4 | 16 |
| SCRN2 | Secernin-2 OS=Homo sapiens OX=9606 GN=SCRN2 PE=1 SV=3 | 7 |
| CGN | Cingulin OS=Homo sapiens OX=9606 GN=CGN PE=1 SV=3 | 13 |
| AASS | Alpha-aminoadipic semialdehyde synthase, mitochondrial OS=Homo sapiens OX=9606 GN=AASS PE=1 SV=1 | 1 |
| TMEM205 | Transmembrane protein 205 OS=Homo sapiens OX=9606 GN=TMEM205 PE=1 SV=1 | 5 |
| CTPS1 | CTP synthase 1 OS=Homo sapiens OX=9606 GN=CTPS1 PE=1 SV=2 | 10 |
| PRPF19 | Pre-mRNA-processing factor 19 OS=Homo sapiens OX=9606 GN=PRPF19 PE=1 SV=1 | 2 |
| CAPZA1 | F-actin-capping protein subunit alpha-1 OS=Homo sapiens OX=9606 GN=CAPZA1 PE=1 SV=3 | 5 |
| SLIRP | SRA stem-loop-interacting RNA-binding protein, mitochondrial OS=Homo sapiens OX=9606 GN=SLIRP PE=1 SV=1 | 8 |
| POLR2I | DNA-directed RNA polymerase II subunit RPB9 OS=Homo sapiens OX=9606 GN=POLR2I PE=1 SV=1 | 1 |
| CHMP1A | Charged multivesicular body protein 1a OS=Homo sapiens OX=9606 GN=CHMP1A PE=1 SV=1 | 3 |
| PEPD | Xaa-Pro dipeptidase OS=Homo sapiens OX=9606 GN=PEPD PE=1 SV=3 | 3 |
| PAM16 | Mitochondrial import inner membrane translocase subunit TIM16 OS=Homo sapiens OX=9606 GN=PAM16 PE=1 SV=2 | 5 |
| RPAP3 | RNA polymerase II-associated protein 3 OS=Homo sapiens OX=9606 GN=RPAP3 PE=1 SV=2 | 10 |
| TAF15 | TATA-binding protein-associated factor 2N OS=Homo sapiens OX=9606 GN=TAF15 PE=1 SV=1 | 4 |
| VTN | Vitronectin OS=Homo sapiens OX=9606 GN=VTN PE=1 SV=1 | 12 |
| HLA-B | HLA class I histocompatibility antigen, B alpha chain OS=Homo sapiens OX=9606 GN=HLA-B PE=1 SV=3 | 3 |
| FGR | Tyrosine-protein kinase Fgr OS=Homo sapiens OX=9606 GN=FGR PE=1 SV=2 | 1 |
| COA3 | Cytochrome c oxidase assembly factor 3 homolog, mitochondrial OS=Homo sapiens OX=9606 GN=COA3 PE=1 SV=1 | 3 |
| COG2 | Conserved oligomeric Golgi complex subunit 2 OS=Homo sapiens OX=9606 GN=COG2 PE=1 SV=1 | 2 |
| MIPEP | Mitochondrial intermediate peptidase OS=Homo sapiens OX=9606 GN=MIPEP PE=1 SV=2 | 2 |
| LAMP1 | Lysosome-associated membrane glycoprotein 1 OS=Homo sapiens OX=9606 GN=LAMP1 PE=1 SV=3 | 6 |
| MICOS13 | MICOS complex subunit MIC13 OS=Homo sapiens OX=9606 GN=MICOS13 PE=1 SV=1 | 2 |
| UBAP2 | Ubiquitin-associated protein 2 OS=Homo sapiens OX=9606 GN=UBAP2 PE=1 SV=1 | 9 |
| TDRD12 | Putative ATP-dependent RNA helicase TDRD12 OS=Homo sapiens OX=9606 GN=TDRD12 PE=2 SV=2 | 2 |
| LAMTOR2 | Ragulator complex protein LAMTOR2 OS=Homo sapiens OX=9606 GN=LAMTOR2 PE=1 SV=1 | 4 |
| KRT79 | Keratin, type II cytoskeletal 79 OS=Homo sapiens OX=9606 GN=KRT79 PE=1 SV=2 | 10 |
| NUDCD1 | NudC domain-containing protein 1 OS=Homo sapiens OX=9606 GN=NUDCD1 PE=1 SV=2 | 5 |
| USP7 | Ubiquitin carboxyl-terminal hydrolase 7 OS=Homo sapiens OX=9606 GN=USP7 PE=1 SV=2 | 10 |
| FDX1 | Adrenodoxin, mitochondrial OS=Homo sapiens OX=9606 GN=FDX1 PE=1 SV=1 | 4 |
| OAT | Ornithine aminotransferase, mitochondrial OS=Homo sapiens OX=9606 GN=OAT PE=1 SV=1 | 29 |
| ANXA11 | Annexin A11 OS=Homo sapiens OX=9606 GN=ANXA11 PE=1 SV=1 | 21 |
| PCBD2 | Pterin-4-alpha-carbinolamine dehydratase 2 OS=Homo sapiens OX=9606 GN=PCBD2 PE=1 SV=4 | 3 |
| PRDX1 | Peroxiredoxin-1 OS=Homo sapiens OX=9606 GN=PRDX1 PE=1 SV=1 | 18 |
| MAPK3 | Mitogen-activated protein kinase 3 OS=Homo sapiens OX=9606 GN=MAPK3 PE=1 SV=4 | 6 |
| POLR2E | DNA-directed RNA polymerases I, II, and III subunit RPABC1 OS=Homo sapiens OX=9606 GN=POLR2E PE=1 SV=4 | 1 |
| WASF1 | Actin-binding protein WASF1 OS=Homo sapiens OX=9606 GN=WASF1 PE=1 SV=1 | 1 |
| HSPBAP1 | HSPB1-associated protein 1 OS=Homo sapiens OX=9606 GN=HSPBAP1 PE=1 SV=1 | 1 |
| HSPG2 | Basement membrane-specific heparan sulfate proteoglycan core protein OS=Homo sapiens OX=9606 GN=HSPG2 PE=1 SV=4 | 41 |
| MCUR1 | Mitochondrial calcium uniporter regulator 1 OS=Homo sapiens OX=9606 GN=MCUR1 PE=1 SV=1 | 8 |
| UQCRC2 | Cytochrome b-c1 complex subunit 2, mitochondrial OS=Homo sapiens OX=9606 GN=UQCRC2 PE=1 SV=3 | 13 |
| SNAP23 | Synaptosomal-associated protein 23 OS=Homo sapiens OX=9606 GN=SNAP23 PE=1 SV=1 | 5 |
| PTS | 6-pyruvoyl tetrahydrobiopterin synthase OS=Homo sapiens OX=9606 GN=PTS PE=1 SV=1 | 3 |
| SLC1A5 | Neutral amino acid transporter B(0) OS=Homo sapiens OX=9606 GN=SLC1A5 PE=1 SV=2 | 2 |
| KDM1A | Lysine-specific histone demethylase 1A OS=Homo sapiens OX=9606 GN=KDM1A PE=1 SV=2 | 2 |
| SHCBP1 | SHC SH2 domain-binding protein 1 OS=Homo sapiens OX=9606 GN=SHCBP1 PE=1 SV=3 | 2 |
| ENOPH1 | Enolase-phosphatase E1 OS=Homo sapiens OX=9606 GN=ENOPH1 PE=1 SV=1 | 4 |
| RPL10 | 60S ribosomal protein L10 OS=Homo sapiens OX=9606 GN=RPL10 PE=1 SV=5 | 8 |
| PON2 | Serum paraoxonase/arylesterase 2 OS=Homo sapiens OX=9606 GN=PON2 PE=1 SV=4 | 9 |
| PRDX2 | Peroxiredoxin-2 OS=Homo sapiens OX=9606 GN=PRDX2 PE=1 SV=5 | 16 |
| MRPL22 | 39S ribosomal protein L22, mitochondrial OS=Homo sapiens OX=9606 GN=MRPL22 PE=1 SV=1 | 12 |
| PIK3CB | Phosphatidylinositol 4,5-bisphosphate 3-kinase catalytic subunit beta isoform OS=Homo sapiens OX=9606 GN=PIK3CB PE=1 SV=1 | 2 |
| RANGAP1 | Ran GTPase-activating protein 1 OS=Homo sapiens OX=9606 GN=RANGAP1 PE=1 SV=1 | 6 |
| COX7A2 | Cytochrome c oxidase subunit 7A2, mitochondrial OS=Homo sapiens OX=9606 GN=COX7A2 PE=1 SV=1 | 2 |
| IGHG3 | Immunoglobulin heavy constant gamma 3 OS=Homo sapiens OX=9606 GN=IGHG3 PE=1 SV=2 | 4 |
| FLNA | Filamin-A OS=Homo sapiens OX=9606 GN=FLNA PE=1 SV=4 | 93 |
| SRRM2 | Serine/arginine repetitive matrix protein 2 OS=Homo sapiens OX=9606 GN=SRRM2 PE=1 SV=2 | 3 |
| RPL15 | 60S ribosomal protein L15 OS=Homo sapiens OX=9606 GN=RPL15 PE=1 SV=2 | 7 |
| PSMC4 | 26S proteasome regulatory subunit 6B OS=Homo sapiens OX=9606 GN=PSMC4 PE=1 SV=2 | 14 |
| NME2 | Nucleoside diphosphate kinase B OS=Homo sapiens OX=9606 GN=NME2 PE=1 SV=1 | 7 |
| ARL1 | ADP-ribosylation factor-like protein 1 OS=Homo sapiens OX=9606 GN=ARL1 PE=1 SV=1 | 5 |
| TYRO3 | Tyrosine-protein kinase receptor TYRO3 OS=Homo sapiens OX=9606 GN=TYRO3 PE=1 SV=1 | 5 |
| SLK | STE20-like serine/threonine-protein kinase OS=Homo sapiens OX=9606 GN=SLK PE=1 SV=1 | 4 |
| COQ7 | 5-demethoxyubiquinone hydroxylase, mitochondrial OS=Homo sapiens OX=9606 GN=COQ7 PE=1 SV=3 | 4 |
| HYPK | Huntingtin-interacting protein K OS=Homo sapiens OX=9606 GN=HYPK PE=1 SV=3 | 3 |
| NDUFA6 | NADH dehydrogenase [ubiquinone] 1 alpha subcomplex subunit 6 OS=Homo sapiens OX=9606 GN=NDUFA6 PE=1 SV=4 | 2 |
| PYGM | Glycogen phosphorylase, muscle form OS=Homo sapiens OX=9606 GN=PYGM PE=1 SV=6 | 1 |
| STK31 | Serine/threonine-protein kinase 31 OS=Homo sapiens OX=9606 GN=STK31 PE=2 SV=2 | 32 |
| ZSWIM3 | Zinc finger SWIM domain-containing protein 3 OS=Homo sapiens OX=9606 GN=ZSWIM3 PE=1 SV=2 | 10 |
| TBC1D10B | TBC1 domain family member 10B OS=Homo sapiens OX=9606 GN=TBC1D10B PE=1 SV=3 | 3 |
| TJP2 | Tight junction protein ZO-2 OS=Homo sapiens OX=9606 GN=TJP2 PE=1 SV=2 | 8 |
| TMEM168 | Transmembrane protein 168 OS=Homo sapiens OX=9606 GN=TMEM168 PE=1 SV=2 | 2 |
| UBQLN4 | Ubiquilin-4 OS=Homo sapiens OX=9606 GN=UBQLN4 PE=1 SV=2 | 3 |
| LDHAL6A | L-lactate dehydrogenase A-like 6A OS=Homo sapiens OX=9606 GN=LDHAL6A PE=1 SV=1 | 22 |
| HEBP2 | Heme-binding protein 2 OS=Homo sapiens OX=9606 GN=HEBP2 PE=1 SV=1 | 4 |
| WDR77 | Methylosome protein 50 OS=Homo sapiens OX=9606 GN=WDR77 PE=1 SV=1 | 7 |
| NUP62 | Nuclear pore glycoprotein p62 OS=Homo sapiens OX=9606 GN=NUP62 PE=1 SV=3 | 3 |
| MRPS22 | 28S ribosomal protein S22, mitochondrial OS=Homo sapiens OX=9606 GN=MRPS22 PE=1 SV=1 | 7 |
| AOX1 | Aldehyde oxidase OS=Homo sapiens OX=9606 GN=AOX1 PE=1 SV=2 | 8 |
| PSMA4 | Proteasome subunit alpha type-4 OS=Homo sapiens OX=9606 GN=PSMA4 PE=1 SV=1 | 10 |
| PAXX | Protein PAXX OS=Homo sapiens OX=9606 GN=PAXX PE=1 SV=2 | 3 |
| ZNF407 | Zinc finger protein 407 OS=Homo sapiens OX=9606 GN=ZNF407 PE=1 SV=2 | 1 |
| F2 | Prothrombin OS=Homo sapiens OX=9606 GN=F2 PE=1 SV=2 | 9 |
| SMDT1 | Essential MCU regulator, mitochondrial OS=Homo sapiens OX=9606 GN=SMDT1 PE=1 SV=1 | 1 |
| PPP6R3 | Serine/threonine-protein phosphatase 6 regulatory subunit 3 OS=Homo sapiens OX=9606 GN=PPP6R3 PE=1 SV=2 | 3 |
| ECSIT | Evolutionarily conserved signaling intermediate in Toll pathway, mitochondrial OS=Homo sapiens OX=9606 GN=ECSIT PE=1 SV=1 | 5 |
| HSCB | Iron-sulfur cluster co-chaperone protein HscB OS=Homo sapiens OX=9606 GN=HSCB PE=1 SV=3 | 3 |
| SLC2A8 | Solute carrier family 2, facilitated glucose transporter member 8 OS=Homo sapiens OX=9606 GN=SLC2A8 PE=1 SV=3 | 1 |
| SNRPGP15 | Putative small nuclear ribonucleoprotein G-like protein 15 OS=Homo sapiens OX=9606 GN=SNRPGP15 PE=5 SV=2 | 2 |
| SLC12A2 | Solute carrier family 12 member 2 OS=Homo sapiens OX=9606 GN=SLC12A2 PE=1 SV=1 | 7 |
| APCS | Serum amyloid P-component OS=Homo sapiens OX=9606 GN=APCS PE=1 SV=2 | 3 |
| RICTOR | Rapamycin-insensitive companion of mTOR OS=Homo sapiens OX=9606 GN=RICTOR PE=1 SV=1 | 4 |
| TSPAN14 | Tetraspanin-14 OS=Homo sapiens OX=9606 GN=TSPAN14 PE=1 SV=1 | 3 |
| MARCKSL1 | MARCKS-related protein OS=Homo sapiens OX=9606 GN=MARCKSL1 PE=1 SV=2 | 2 |
| PSME4 | Proteasome activator complex subunit 4 OS=Homo sapiens OX=9606 GN=PSME4 PE=1 SV=2 | 4 |
| GGACT | Gamma-glutamylaminecyclotransferase OS=Homo sapiens OX=9606 GN=GGACT PE=1 SV=2 | 2 |
| NCSTN | Nicastrin OS=Homo sapiens OX=9606 GN=NCSTN PE=1 SV=2 | 5 |
| SPRYD7 | SPRY domain-containing protein 7 OS=Homo sapiens OX=9606 GN=SPRYD7 PE=1 SV=2 | 1 |
| UGDH | UDP-glucose 6-dehydrogenase OS=Homo sapiens OX=9606 GN=UGDH PE=1 SV=1 | 26 |
| RPTOR | Regulatory-associated protein of mTOR OS=Homo sapiens OX=9606 GN=RPTOR PE=1 SV=1 | 1 |
| DNAJC15 | DnaJ homolog subfamily C member 15 OS=Homo sapiens OX=9606 GN=DNAJC15 PE=1 SV=2 | 5 |
| MTPN | Myotrophin OS=Homo sapiens OX=9606 GN=MTPN PE=1 SV=2 | 8 |
| ABCF2 | ATP-binding cassette sub-family F member 2 OS=Homo sapiens OX=9606 GN=ABCF2 PE=1 SV=2 | 2 |
| TRAPPC13 | Trafficking protein particle complex subunit 13 OS=Homo sapiens OX=9606 GN=TRAPPC13 PE=1 SV=2 | 5 |
| SLC27A2 | Long-chain fatty acid transport protein 2 OS=Homo sapiens OX=9606 GN=SLC27A2 PE=1 SV=2 | 1 |
| MAP1LC3B2 | Microtubule-associated proteins 1A/1B light chain 3 beta 2 OS=Homo sapiens OX=9606 GN=MAP1LC3B2 PE=1 SV=1 | 3 |
| COA1 | Cytochrome c oxidase assembly factor 1 homolog OS=Homo sapiens OX=9606 GN=COA1 PE=1 SV=1 | 4 |
| SNRNP200 | U5 small nuclear ribonucleoprotein 200 kDa helicase OS=Homo sapiens OX=9606 GN=SNRNP200 PE=1 SV=2 | 3 |
| MECR | Enoyl-[acyl-carrier-protein] reductase, mitochondrial OS=Homo sapiens OX=9606 GN=MECR PE=1 SV=2 | 7 |
| CRYZ | Quinone oxidoreductase OS=Homo sapiens OX=9606 GN=CRYZ PE=1 SV=1 | 23 |
| LUC7L2 | Putative RNA-binding protein Luc7-like 2 OS=Homo sapiens OX=9606 GN=LUC7L2 PE=1 SV=2 | 3 |
| IFT74 | Intraflagellar transport protein 74 homolog OS=Homo sapiens OX=9606 GN=IFT74 PE=1 SV=1 | 3 |
| GCC2 | GRIP and coiled-coil domain-containing protein 2 OS=Homo sapiens OX=9606 GN=GCC2 PE=1 SV=4 | 4 |
| TRIM26 | Tripartite motif-containing protein 26 OS=Homo sapiens OX=9606 GN=TRIM26 PE=1 SV=1 | 2 |
| RAB11FIP5 | Rab11 family-interacting protein 5 OS=Homo sapiens OX=9606 GN=RAB11FIP5 PE=1 SV=1 | 8 |
| DDX5 | Probable ATP-dependent RNA helicase DDX5 OS=Homo sapiens OX=9606 GN=DDX5 PE=1 SV=1 | 1 |
| GEMIN8 | Gem-associated protein 8 OS=Homo sapiens OX=9606 GN=GEMIN8 PE=1 SV=1 | 1 |
| GMPS | GMP synthase [glutamine-hydrolyzing] OS=Homo sapiens OX=9606 GN=GMPS PE=1 SV=1 | 2 |
| STK32C | Serine/threonine-protein kinase 32C OS=Homo sapiens OX=9606 GN=STK32C PE=1 SV=1 | 1 |
| TOLLIP | Toll-interacting protein OS=Homo sapiens OX=9606 GN=TOLLIP PE=1 SV=1 | 7 |
| AP2A2 | AP-2 complex subunit alpha-2 OS=Homo sapiens OX=9606 GN=AP2A2 PE=1 SV=2 | 4 |
| SPCS3 | Signal peptidase complex subunit 3 OS=Homo sapiens OX=9606 GN=SPCS3 PE=1 SV=1 | 3 |
| PRRC1 | Protein PRRC1 OS=Homo sapiens OX=9606 GN=PRRC1 PE=1 SV=1 | 4 |
| NDUFS6 | NADH dehydrogenase [ubiquinone] iron-sulfur protein 6, mitochondrial OS=Homo sapiens OX=9606 GN=NDUFS6 PE=1 SV=1 | 3 |
| DHRS11 | Dehydrogenase/reductase SDR family member 11 OS=Homo sapiens OX=9606 GN=DHRS11 PE=1 SV=1 | 10 |
| PITHD1 | PITH domain-containing protein 1 OS=Homo sapiens OX=9606 GN=PITHD1 PE=1 SV=1 | 15 |
| ODR4 | Protein odr-4 homolog OS=Homo sapiens OX=9606 GN=ODR4 PE=1 SV=1 | 6 |
| COMMD5 | COMM domain-containing protein 5 OS=Homo sapiens OX=9606 GN=COMMD5 PE=1 SV=1 | 2 |
| UQCRB | Cytochrome b-c1 complex subunit 7 OS=Homo sapiens OX=9606 GN=UQCRB PE=1 SV=2 | 8 |
| RHOG | Rho-related GTP-binding protein RhoG OS=Homo sapiens OX=9606 GN=RHOG PE=1 SV=1 | 6 |
| SARNP | SAP domain-containing ribonucleoprotein OS=Homo sapiens OX=9606 GN=SARNP PE=1 SV=3 | 1 |
| RFT1 | Protein RFT1 homolog OS=Homo sapiens OX=9606 GN=RFT1 PE=1 SV=1 | 1 |
| UQCRC1 | Cytochrome b-c1 complex subunit 1, mitochondrial OS=Homo sapiens OX=9606 GN=UQCRC1 PE=1 SV=3 | 26 |
| BET1 | BET1 homolog OS=Homo sapiens OX=9606 GN=BET1 PE=1 SV=1 | 1 |
| MRPL30 | 39S ribosomal protein L30, mitochondrial OS=Homo sapiens OX=9606 GN=MRPL30 PE=1 SV=1 | 2 |
| UBE2G1 | Ubiquitin-conjugating enzyme E2 G1 OS=Homo sapiens OX=9606 GN=UBE2G1 PE=1 SV=3 | 2 |
| NGLY1 | Peptide-N(4)-(N-acetyl-beta-glucosaminyl)asparagine amidase OS=Homo sapiens OX=9606 GN=NGLY1 PE=1 SV=1 | 3 |
| TPM4 | Tropomyosin alpha-4 chain OS=Homo sapiens OX=9606 GN=TPM4 PE=1 SV=3 | 7 |
| LIG3 | DNA ligase 3 OS=Homo sapiens OX=9606 GN=LIG3 PE=1 SV=2 | 3 |
| EEF2 | Elongation factor 2 OS=Homo sapiens OX=9606 GN=EEF2 PE=1 SV=4 | 48 |
| IMPACT | Protein IMPACT OS=Homo sapiens OX=9606 GN=IMPACT PE=1 SV=2 | 4 |
| PCBP1 | Poly(rC)-binding protein 1 OS=Homo sapiens OX=9606 GN=PCBP1 PE=1 SV=2 | 15 |
| SUDS3 | Sin3 histone deacetylase corepressor complex component SDS3 OS=Homo sapiens OX=9606 GN=SUDS3 PE=1 SV=2 | 5 |
| CHAF1B | Chromatin assembly factor 1 subunit B OS=Homo sapiens OX=9606 GN=CHAF1B PE=1 SV=1 | 1 |
| NDUFA13 | NADH dehydrogenase [ubiquinone] 1 alpha subcomplex subunit 13 OS=Homo sapiens OX=9606 GN=NDUFA13 PE=1 SV=3 | 5 |
| GCLM | Glutamate--cysteine ligase regulatory subunit OS=Homo sapiens OX=9606 GN=GCLM PE=1 SV=1 | 4 |
| OXT | Oxytocin-neurophysin 1 OS=Homo sapiens OX=9606 GN=OXT PE=1 SV=1 | 6 |
| LSM7 | U6 snRNA-associated Sm-like protein LSm7 OS=Homo sapiens OX=9606 GN=LSM7 PE=1 SV=1 | 5 |
| TTK | Dual specificity protein kinase TTK OS=Homo sapiens OX=9606 GN=TTK PE=1 SV=2 | 3 |
| KATNA1 | Katanin p60 ATPase-containing subunit A1 OS=Homo sapiens OX=9606 GN=KATNA1 PE=1 SV=1 | 2 |
| TUBGCP4 | Gamma-tubulin complex component 4 OS=Homo sapiens OX=9606 GN=TUBGCP4 PE=1 SV=1 | 4 |
| HSPA6 | Heat shock 70 kDa protein 6 OS=Homo sapiens OX=9606 GN=HSPA6 PE=1 SV=2 | 1 |
| NUDT8 | Mitochondrial coenzyme A diphosphatase NUDT8 OS=Homo sapiens OX=9606 GN=NUDT8 PE=1 SV=2 | 1 |
| PSMB1 | Proteasome subunit beta type-1 OS=Homo sapiens OX=9606 GN=PSMB1 PE=1 SV=2 | 7 |
| EIF2S2 | Eukaryotic translation initiation factor 2 subunit 2 OS=Homo sapiens OX=9606 GN=EIF2S2 PE=1 SV=2 | 2 |
| RPL5 | 60S ribosomal protein L5 OS=Homo sapiens OX=9606 GN=RPL5 PE=1 SV=3 | 10 |
| PSMA2 | Proteasome subunit alpha type-2 OS=Homo sapiens OX=9606 GN=PSMA2 PE=1 SV=2 | 12 |
| PURA | Transcriptional activator protein Pur-alpha OS=Homo sapiens OX=9606 GN=PURA PE=1 SV=2 | 2 |
| USP8 | Ubiquitin carboxyl-terminal hydrolase 8 OS=Homo sapiens OX=9606 GN=USP8 PE=1 SV=1 | 3 |
| TADA2A | Transcriptional adapter 2-alpha OS=Homo sapiens OX=9606 GN=TADA2A PE=1 SV=3 | 1 |
| NF1 | Neurofibromin OS=Homo sapiens OX=9606 GN=NF1 PE=1 SV=2 | 2 |
| SSRP1 | FACT complex subunit SSRP1 OS=Homo sapiens OX=9606 GN=SSRP1 PE=1 SV=1 | 2 |
| UBXN4 | UBX domain-containing protein 4 OS=Homo sapiens OX=9606 GN=UBXN4 PE=1 SV=2 | 1 |
| EIF3H | Eukaryotic translation initiation factor 3 subunit H OS=Homo sapiens OX=9606 GN=EIF3H PE=1 SV=1 | 8 |
| EIF3B | Eukaryotic translation initiation factor 3 subunit B OS=Homo sapiens OX=9606 GN=EIF3B PE=1 SV=3 | 12 |
| CCDC25 | Coiled-coil domain-containing protein 25 OS=Homo sapiens OX=9606 GN=CCDC25 PE=1 SV=2 | 10 |
| LMNB1 | Lamin-B1 OS=Homo sapiens OX=9606 GN=LMNB1 PE=1 SV=2 | 8 |
| EIF1B | Eukaryotic translation initiation factor 1b OS=Homo sapiens OX=9606 GN=EIF1B PE=1 SV=2 | 5 |
| BORCS5 | BLOC-1-related complex subunit 5 OS=Homo sapiens OX=9606 GN=BORCS5 PE=1 SV=1 | 1 |
| ILF3 | Interleukin enhancer-binding factor 3 OS=Homo sapiens OX=9606 GN=ILF3 PE=1 SV=3 | 3 |
| SRSF3 | Serine/arginine-rich splicing factor 3 OS=Homo sapiens OX=9606 GN=SRSF3 PE=1 SV=1 | 1 |
| CMC2 | COX assembly mitochondrial protein 2 homolog OS=Homo sapiens OX=9606 GN=CMC2 PE=1 SV=1 | 1 |
| SSR4 | Translocon-associated protein subunit delta OS=Homo sapiens OX=9606 GN=SSR4 PE=1 SV=1 | 7 |
| TNRC6B | Trinucleotide repeat-containing gene 6B protein OS=Homo sapiens OX=9606 GN=TNRC6B PE=1 SV=4 | 6 |
| PRKDC | DNA-dependent protein kinase catalytic subunit OS=Homo sapiens OX=9606 GN=PRKDC PE=1 SV=3 | 9 |
| PSMA1 | Proteasome subunit alpha type-1 OS=Homo sapiens OX=9606 GN=PSMA1 PE=1 SV=1 | 17 |
| SKIV2L | SKI2 subunit of superkiller complex protein OS=Homo sapiens OX=9606 GN=SKIC2 PE=1 SV=3 | 2 |
| MT-CO2 | Cytochrome c oxidase subunit 2 OS=Homo sapiens OX=9606 GN=MT-CO2 PE=1 SV=1 | 4 |
| RPUSD4 | Pseudouridylate synthase RPUSD4, mitochondrial OS=Homo sapiens OX=9606 GN=RPUSD4 PE=1 SV=1 | 1 |
| BPNT1 | 3'(2'),5'-bisphosphate nucleotidase 1 OS=Homo sapiens OX=9606 GN=BPNT1 PE=1 SV=1 | 5 |
| IPO9 | Importin-9 OS=Homo sapiens OX=9606 GN=IPO9 PE=1 SV=3 | 3 |
| TNPO3 | Transportin-3 OS=Homo sapiens OX=9606 GN=TNPO3 PE=1 SV=3 | 1 |
| AASDHPPT | L-aminoadipate-semialdehyde dehydrogenase-phosphopantetheinyl transferase OS=Homo sapiens OX=9606 GN=AASDHPPT PE=1 SV=2 | 2 |
| PSMA7 | Proteasome subunit alpha type-7 OS=Homo sapiens OX=9606 GN=PSMA7 PE=1 SV=1 | 13 |
| SAA4 | Serum amyloid A-4 protein OS=Homo sapiens OX=9606 GN=SAA4 PE=1 SV=2 | 2 |
| MRRF | Ribosome-recycling factor, mitochondrial OS=Homo sapiens OX=9606 GN=MRRF PE=1 SV=1 | 1 |
| ARPC4 | Actin-related protein 2/3 complex subunit 4 OS=Homo sapiens OX=9606 GN=ARPC4 PE=1 SV=3 | 7 |
| STAT1 | Signal transducer and activator of transcription 1-alpha/beta OS=Homo sapiens OX=9606 GN=STAT1 PE=1 SV=2 | 10 |
| PSMB7 | Proteasome subunit beta type-7 OS=Homo sapiens OX=9606 GN=PSMB7 PE=1 SV=1 | 5 |
| FGGY | FGGY carbohydrate kinase domain-containing protein OS=Homo sapiens OX=9606 GN=FGGY PE=1 SV=2 | 3 |
| PTBP2 | Polypyrimidine tract-binding protein 2 OS=Homo sapiens OX=9606 GN=PTBP2 PE=1 SV=1 | 8 |
| COQ3 | Ubiquinone biosynthesis O-methyltransferase, mitochondrial OS=Homo sapiens OX=9606 GN=COQ3 PE=1 SV=3 | 5 |
| U2AF2 | Splicing factor U2AF 65 kDa subunit OS=Homo sapiens OX=9606 GN=U2AF2 PE=1 SV=4 | 3 |
| SF3B6 | Splicing factor 3B subunit 6 OS=Homo sapiens OX=9606 GN=SF3B6 PE=1 SV=1 | 2 |
| ABCC4 | ATP-binding cassette sub-family C member 4 OS=Homo sapiens OX=9606 GN=ABCC4 PE=1 SV=3 | 2 |
| ANKLE2 | Ankyrin repeat and LEM domain-containing protein 2 OS=Homo sapiens OX=9606 GN=ANKLE2 PE=1 SV=4 | 3 |
| SPTAN1 | Spectrin alpha chain, non-erythrocytic 1 OS=Homo sapiens OX=9606 GN=SPTAN1 PE=1 SV=3 | 26 |
| WDR61 | SKI8 subunit of superkiller complex protein OS=Homo sapiens OX=9606 GN=SKIC8 PE=1 SV=1 | 9 |
| NAA35 | N-alpha-acetyltransferase 35, NatC auxiliary subunit OS=Homo sapiens OX=9606 GN=NAA35 PE=1 SV=1 | 2 |
| VAMP8 | Vesicle-associated membrane protein 8 OS=Homo sapiens OX=9606 GN=VAMP8 PE=1 SV=1 | 5 |
| CPSF1 | Cleavage and polyadenylation specificity factor subunit 1 OS=Homo sapiens OX=9606 GN=CPSF1 PE=1 SV=2 | 1 |
| JMY | Junction-mediating and -regulatory protein OS=Homo sapiens OX=9606 GN=JMY PE=1 SV=2 | 1 |
| IRGQ | Immunity-related GTPase family Q protein OS=Homo sapiens OX=9606 GN=IRGQ PE=1 SV=1 | 1 |
| GPR89A | Golgi pH regulator A OS=Homo sapiens OX=9606 GN=GPR89A PE=1 SV=2 | 1 |
| FUT3 | 3-galactosyl-N-acetylglucosaminide 4-alpha-L-fucosyltransferase FUT3 OS=Homo sapiens OX=9606 GN=FUT3 PE=1 SV=1 | 1 |
| PSPH | Phosphoserine phosphatase OS=Homo sapiens OX=9606 GN=PSPH PE=1 SV=2 | 1 |
| RRM2B | Ribonucleoside-diphosphate reductase subunit M2 B OS=Homo sapiens OX=9606 GN=RRM2B PE=1 SV=1 | 5 |
| MAPKAPK5 | MAP kinase-activated protein kinase 5 OS=Homo sapiens OX=9606 GN=MAPKAPK5 PE=1 SV=2 | 1 |
| RPS15 | 40S ribosomal protein S15 OS=Homo sapiens OX=9606 GN=RPS15 PE=1 SV=2 | 1 |
| NHLRC2 | NHL repeat-containing protein 2 OS=Homo sapiens OX=9606 GN=NHLRC2 PE=1 SV=1 | 4 |
| WDR17 | WD repeat-containing protein 17 OS=Homo sapiens OX=9606 GN=WDR17 PE=2 SV=2 | 1 |
| PSMA3 | Proteasome subunit alpha type-3 OS=Homo sapiens OX=9606 GN=PSMA3 PE=1 SV=2 | 7 |
| WDR44 | WD repeat-containing protein 44 OS=Homo sapiens OX=9606 GN=WDR44 PE=1 SV=1 | 24 |
| IGFBP2 | Insulin-like growth factor-binding protein 2 OS=Homo sapiens OX=9606 GN=IGFBP2 PE=1 SV=2 | 10 |
| SERPINI1 | Neuroserpin OS=Homo sapiens OX=9606 GN=SERPINI1 PE=1 SV=1 | 2 |
| HECTD3 | E3 ubiquitin-protein ligase HECTD3 OS=Homo sapiens OX=9606 GN=HECTD3 PE=1 SV=1 | 1 |
| MRFAP1 | MORF4 family-associated protein 1 OS=Homo sapiens OX=9606 GN=MRFAP1 PE=1 SV=1 | 3 |
| EHD4 | EH domain-containing protein 4 OS=Homo sapiens OX=9606 GN=EHD4 PE=1 SV=1 | 5 |
| NEDD1 | Protein NEDD1 OS=Homo sapiens OX=9606 GN=NEDD1 PE=1 SV=1 | 4 |
| EIF3M | Eukaryotic translation initiation factor 3 subunit M OS=Homo sapiens OX=9606 GN=EIF3M PE=1 SV=1 | 7 |
| TIGAR | Fructose-2,6-bisphosphatase TIGAR OS=Homo sapiens OX=9606 GN=TIGAR PE=1 SV=1 | 2 |
| MRPS7 | 28S ribosomal protein S7, mitochondrial OS=Homo sapiens OX=9606 GN=MRPS7 PE=1 SV=2 | 4 |
| PLB1 | Phospholipase B1, membrane-associated OS=Homo sapiens OX=9606 GN=PLB1 PE=1 SV=3 | 3 |
| RACK1 | Receptor of activated protein C kinase 1 OS=Homo sapiens OX=9606 GN=RACK1 PE=1 SV=3 | 15 |
| EPCAM | Epithelial cell adhesion molecule OS=Homo sapiens OX=9606 GN=EPCAM PE=1 SV=2 | 1 |
| CCDC134 | Coiled-coil domain-containing protein 134 OS=Homo sapiens OX=9606 GN=CCDC134 PE=1 SV=1 | 2 |
| MACO1 | Macoilin OS=Homo sapiens OX=9606 GN=MACO1 PE=1 SV=1 | 3 |
| CTNNA1 | Catenin alpha-1 OS=Homo sapiens OX=9606 GN=CTNNA1 PE=1 SV=1 | 19 |
| USP15 | Ubiquitin carboxyl-terminal hydrolase 15 OS=Homo sapiens OX=9606 GN=USP15 PE=1 SV=3 | 22 |
| C9 | Complement component C9 OS=Homo sapiens OX=9606 GN=C9 PE=1 SV=2 | 5 |
| NMES1 | Normal mucosa of esophagus-specific gene 1 protein OS=Homo sapiens OX=9606 GN=NMES1 PE=1 SV=1 | 5 |
| CD47 | Leukocyte surface antigen CD47 OS=Homo sapiens OX=9606 GN=CD47 PE=1 SV=1 | 2 |
| VCAM1 | Vascular cell adhesion protein 1 OS=Homo sapiens OX=9606 GN=VCAM1 PE=1 SV=1 | 1 |
| NDUFB4 | NADH dehydrogenase [ubiquinone] 1 beta subcomplex subunit 4 OS=Homo sapiens OX=9606 GN=NDUFB4 PE=1 SV=3 | 2 |
| LRATD2 | Protein LRATD2 OS=Homo sapiens OX=9606 GN=LRATD2 PE=1 SV=1 | 1 |
| COL6A2 | Collagen alpha-2(VI) chain OS=Homo sapiens OX=9606 GN=COL6A2 PE=1 SV=4 | 4 |
| COTL1 | Coactosin-like protein OS=Homo sapiens OX=9606 GN=COTL1 PE=1 SV=3 | 7 |
| PRKRA | Interferon-inducible double-stranded RNA-dependent protein kinase activator A OS=Homo sapiens OX=9606 GN=PRKRA PE=1 SV=1 | 5 |
| MED25 | Mediator of RNA polymerase II transcription subunit 25 OS=Homo sapiens OX=9606 GN=MED25 PE=1 SV=2 | 1 |
| POC5 | Centrosomal protein POC5 OS=Homo sapiens OX=9606 GN=POC5 PE=1 SV=2 | 1 |
| ANO10 | Anoctamin-10 OS=Homo sapiens OX=9606 GN=ANO10 PE=1 SV=2 | 7 |
| COQ5 | 2-methoxy-6-polyprenyl-1,4-benzoquinol methylase, mitochondrial OS=Homo sapiens OX=9606 GN=COQ5 PE=1 SV=2 | 8 |
| AKAP13 | A-kinase anchor protein 13 OS=Homo sapiens OX=9606 GN=AKAP13 PE=1 SV=2 | 2 |
| UQCRFS1 | Cytochrome b-c1 complex subunit Rieske, mitochondrial OS=Homo sapiens OX=9606 GN=UQCRFS1 PE=1 SV=2 | 11 |
| WDR18 | WD repeat-containing protein 18 OS=Homo sapiens OX=9606 GN=WDR18 PE=1 SV=2 | 5 |
| COX7A2L | Cytochrome c oxidase subunit 7A-related protein, mitochondrial OS=Homo sapiens OX=9606 GN=COX7A2L PE=1 SV=2 | 3 |
| PREP | Prolyl endopeptidase OS=Homo sapiens OX=9606 GN=PREP PE=1 SV=2 | 15 |
| VCL | Vinculin OS=Homo sapiens OX=9606 GN=VCL PE=1 SV=4 | 44 |
| ATP6V1D | V-type proton ATPase subunit D OS=Homo sapiens OX=9606 GN=ATP6V1D PE=1 SV=1 | 3 |
| SMC2 | Structural maintenance of chromosomes protein 2 OS=Homo sapiens OX=9606 GN=SMC2 PE=1 SV=2 | 4 |
| FBXO7 | F-box only protein 7 OS=Homo sapiens OX=9606 GN=FBXO7 PE=1 SV=1 | 1 |
| MRM1 | rRNA methyltransferase 1, mitochondrial OS=Homo sapiens OX=9606 GN=MRM1 PE=1 SV=1 | 1 |
| ZNF207 | BUB3-interacting and GLEBS motif-containing protein ZNF207 OS=Homo sapiens OX=9606 GN=ZNF207 PE=1 SV=1 | 1 |
| PPP1R9A | Neurabin-1 OS=Homo sapiens OX=9606 GN=PPP1R9A PE=1 SV=2 | 1 |
| EWSR1 | RNA-binding protein EWS OS=Homo sapiens OX=9606 GN=EWSR1 PE=1 SV=1 | 3 |
| LHFPL4 | LHFPL tetraspan subfamily member 4 protein OS=Homo sapiens OX=9606 GN=LHFPL4 PE=1 SV=1 | 1 |
| SNRPD1 | Small nuclear ribonucleoprotein Sm D1 OS=Homo sapiens OX=9606 GN=SNRPD1 PE=1 SV=1 | 1 |
| TIAL1 | Nucleolysin TIAR OS=Homo sapiens OX=9606 GN=TIAL1 PE=1 SV=1 | 7 |
| NIT1 | Deaminated glutathione amidase OS=Homo sapiens OX=9606 GN=NIT1 PE=1 SV=2 | 9 |
| PA2G4 | Proliferation-associated protein 2G4 OS=Homo sapiens OX=9606 GN=PA2G4 PE=1 SV=3 | 14 |
| TM9SF3 | Transmembrane 9 superfamily member 3 OS=Homo sapiens OX=9606 GN=TM9SF3 PE=1 SV=2 | 2 |
| PGM2 | Phosphopentomutase OS=Homo sapiens OX=9606 GN=PGM2 PE=1 SV=4 | 13 |
| SLC39A6 | Zinc transporter ZIP6 OS=Homo sapiens OX=9606 GN=SLC39A6 PE=1 SV=3 | 1 |
| RCBTB2 | RCC1 and BTB domain-containing protein 2 OS=Homo sapiens OX=9606 GN=RCBTB2 PE=1 SV=1 | 1 |
| NUP43 | Nucleoporin Nup43 OS=Homo sapiens OX=9606 GN=NUP43 PE=1 SV=1 | 3 |
| ITGA4 | Integrin alpha-4 OS=Homo sapiens OX=9606 GN=ITGA4 PE=1 SV=3 | 1 |
| TKT | Transketolase OS=Homo sapiens OX=9606 GN=TKT PE=1 SV=3 | 23 |
| PANX1 | Pannexin-1 OS=Homo sapiens OX=9606 GN=PANX1 PE=1 SV=4 | 2 |
| RPL30 | 60S ribosomal protein L30 OS=Homo sapiens OX=9606 GN=RPL30 PE=1 SV=2 | 4 |
| ZC3HC1 | Zinc finger C3HC-type protein 1 OS=Homo sapiens OX=9606 GN=ZC3HC1 PE=1 SV=1 | 1 |
| CIP2A | Protein CIP2A OS=Homo sapiens OX=9606 GN=CIP2A PE=1 SV=2 | 5 |
| DRG1 | Developmentally-regulated GTP-binding protein 1 OS=Homo sapiens OX=9606 GN=DRG1 PE=1 SV=1 | 6 |
| CSNK1A1 | Casein kinase I isoform alpha OS=Homo sapiens OX=9606 GN=CSNK1A1 PE=1 SV=2 | 2 |
| TMCO1 | Calcium load-activated calcium channel OS=Homo sapiens OX=9606 GN=TMCO1 PE=1 SV=2 | 2 |
| CBX1 | Chromobox protein homolog 1 OS=Homo sapiens OX=9606 GN=CBX1 PE=1 SV=1 | 5 |
| KLHL25 | Kelch-like protein 25 OS=Homo sapiens OX=9606 GN=KLHL25 PE=1 SV=1 | 4 |
| PIP4K2A | Phosphatidylinositol 5-phosphate 4-kinase type-2 alpha OS=Homo sapiens OX=9606 GN=PIP4K2A PE=1 SV=2 | 2 |
| NAA50 | N-alpha-acetyltransferase 50 OS=Homo sapiens OX=9606 GN=NAA50 PE=1 SV=1 | 9 |
| MRC1 | Macrophage mannose receptor 1 OS=Homo sapiens OX=9606 GN=MRC1 PE=1 SV=1 | 3 |
| C6orf52 | Putative uncharacterized protein C6orf52 OS=Homo sapiens OX=9606 GN=C6orf52 PE=2 SV=2 | 2 |
| LMAN1 | Protein ERGIC-53 OS=Homo sapiens OX=9606 GN=LMAN1 PE=1 SV=2 | 5 |
| PRPF39 | Pre-mRNA-processing factor 39 OS=Homo sapiens OX=9606 GN=PRPF39 PE=1 SV=3 | 4 |
| HAUS4 | HAUS augmin-like complex subunit 4 OS=Homo sapiens OX=9606 GN=HAUS4 PE=1 SV=1 | 1 |
| ABCF1 | ATP-binding cassette sub-family F member 1 OS=Homo sapiens OX=9606 GN=ABCF1 PE=1 SV=2 | 7 |
| FLII | Protein flightless-1 homolog OS=Homo sapiens OX=9606 GN=FLII PE=1 SV=2 | 1 |
| TSC1 | Hamartin OS=Homo sapiens OX=9606 GN=TSC1 PE=1 SV=2 | 1 |
| PSMB5 | Proteasome subunit beta type-5 OS=Homo sapiens OX=9606 GN=PSMB5 PE=1 SV=3 | 10 |
| SLC30A9 | Proton-coupled zinc antiporter SLC30A9, mitochondrial OS=Homo sapiens OX=9606 GN=SLC30A9 PE=1 SV=1 | 1 |
| FAHD1 | Acylpyruvase FAHD1, mitochondrial OS=Homo sapiens OX=9606 GN=FAHD1 PE=1 SV=2 | 4 |
| S100Z | Protein S100-Z OS=Homo sapiens OX=9606 GN=S100Z PE=1 SV=4 | 1 |
| PFKFB2 | 6-phosphofructo-2-kinase/fructose-2,6-bisphosphatase 2 OS=Homo sapiens OX=9606 GN=PFKFB2 PE=1 SV=2 | 20 |
| ESRP1 | Epithelial splicing regulatory protein 1 OS=Homo sapiens OX=9606 GN=ESRP1 PE=1 SV=2 | 4 |
| GEMIN4 | Gem-associated protein 4 OS=Homo sapiens OX=9606 GN=GEMIN4 PE=1 SV=2 | 1 |
| ZBTB33 | Transcriptional regulator Kaiso OS=Homo sapiens OX=9606 GN=ZBTB33 PE=1 SV=2 | 1 |
| MRPS34 | 28S ribosomal protein S34, mitochondrial OS=Homo sapiens OX=9606 GN=MRPS34 PE=1 SV=2 | 3 |
| SLC27A6 | Long-chain fatty acid transport protein 6 OS=Homo sapiens OX=9606 GN=SLC27A6 PE=2 SV=1 | 4 |
| SYT6 | Synaptotagmin-6 OS=Homo sapiens OX=9606 GN=SYT6 PE=1 SV=3 | 1 |
| SNRPF | Small nuclear ribonucleoprotein F OS=Homo sapiens OX=9606 GN=SNRPF PE=1 SV=1 | 1 |
| SERPINC1 | Antithrombin-III OS=Homo sapiens OX=9606 GN=SERPINC1 PE=1 SV=1 | 14 |
| SKP2 | S-phase kinase-associated protein 2 OS=Homo sapiens OX=9606 GN=SKP2 PE=1 SV=2 | 5 |
| NDUFB5 | NADH dehydrogenase [ubiquinone] 1 beta subcomplex subunit 5, mitochondrial OS=Homo sapiens OX=9606 GN=NDUFB5 PE=1 SV=1 | 3 |
| ARFIP2 | Arfaptin-2 OS=Homo sapiens OX=9606 GN=ARFIP2 PE=1 SV=1 | 5 |
| METTL21A | Protein N-lysine methyltransferase METTL21A OS=Homo sapiens OX=9606 GN=METTL21A PE=1 SV=2 | 3 |
| CDS2 | Phosphatidate cytidylyltransferase 2 OS=Homo sapiens OX=9606 GN=CDS2 PE=1 SV=1 | 2 |
| PEF1 | Peflin OS=Homo sapiens OX=9606 GN=PEF1 PE=1 SV=1 | 3 |
| KCTD2 | BTB/POZ domain-containing protein KCTD2 OS=Homo sapiens OX=9606 GN=KCTD2 PE=1 SV=3 | 2 |
| IRF2BPL | Probable E3 ubiquitin-protein ligase IRF2BPL OS=Homo sapiens OX=9606 GN=IRF2BPL PE=1 SV=1 | 2 |
| RABL3 | Rab-like protein 3 OS=Homo sapiens OX=9606 GN=RABL3 PE=1 SV=1 | 3 |
| RBM26 | RNA-binding protein 26 OS=Homo sapiens OX=9606 GN=RBM26 PE=1 SV=3 | 4 |
| NDUFC2 | NADH dehydrogenase [ubiquinone] 1 subunit C2 OS=Homo sapiens OX=9606 GN=NDUFC2 PE=1 SV=1 | 3 |
| RBM46 | Probable RNA-binding protein 46 OS=Homo sapiens OX=9606 GN=RBM46 PE=1 SV=1 | 2 |
| MRPL39 | 39S ribosomal protein L39, mitochondrial OS=Homo sapiens OX=9606 GN=MRPL39 PE=1 SV=3 | 9 |
| CKS2 | Cyclin-dependent kinases regulatory subunit 2 OS=Homo sapiens OX=9606 GN=CKS2 PE=1 SV=1 | 1 |
| POLR2C | DNA-directed RNA polymerase II subunit RPB3 OS=Homo sapiens OX=9606 GN=POLR2C PE=1 SV=2 | 3 |
| MDM1 | Nuclear protein MDM1 OS=Homo sapiens OX=9606 GN=MDM1 PE=1 SV=2 | 2 |
| TMEM256 | Transmembrane protein 256 OS=Homo sapiens OX=9606 GN=TMEM256 PE=3 SV=1 | 1 |
| SERPINA3 | Alpha-1-antichymotrypsin OS=Homo sapiens OX=9606 GN=SERPINA3 PE=1 SV=2 | 9 |
| EMD | Emerin OS=Homo sapiens OX=9606 GN=EMD PE=1 SV=1 | 2 |
| MCM7 | DNA replication licensing factor MCM7 OS=Homo sapiens OX=9606 GN=MCM7 PE=1 SV=4 | 3 |
| HNRNPM | Heterogeneous nuclear ribonucleoprotein M OS=Homo sapiens OX=9606 GN=HNRNPM PE=1 SV=3 | 14 |
| CPA1 | Carboxypeptidase A1 OS=Homo sapiens OX=9606 GN=CPA1 PE=1 SV=2 | 1 |
| POLR2K | DNA-directed RNA polymerases I, II, and III subunit RPABC4 OS=Homo sapiens OX=9606 GN=POLR2K PE=1 SV=1 | 1 |
| SPATA33 | Spermatogenesis-associated protein 33 OS=Homo sapiens OX=9606 GN=SPATA33 PE=1 SV=1 | 2 |
| TRAPPC2L | Trafficking protein particle complex subunit 2-like protein OS=Homo sapiens OX=9606 GN=TRAPPC2L PE=1 SV=1 | 4 |
| MRPL18 | 39S ribosomal protein L18, mitochondrial OS=Homo sapiens OX=9606 GN=MRPL18 PE=1 SV=1 | 3 |
| SFPQ | Splicing factor, proline- and glutamine-rich OS=Homo sapiens OX=9606 GN=SFPQ PE=1 SV=2 | 8 |
| UPF1 | Regulator of nonsense transcripts 1 OS=Homo sapiens OX=9606 GN=UPF1 PE=1 SV=2 | 10 |
| HDHD5 | Haloacid dehalogenase-like hydrolase domain-containing 5 OS=Homo sapiens OX=9606 GN=HDHD5 PE=1 SV=1 | 4 |
| NDUFB2 | NADH dehydrogenase [ubiquinone] 1 beta subcomplex subunit 2, mitochondrial OS=Homo sapiens OX=9606 GN=NDUFB2 PE=1 SV=1 | 1 |
| MAGED2 | Melanoma-associated antigen D2 OS=Homo sapiens OX=9606 GN=MAGED2 PE=1 SV=2 | 2 |
| CPM | Carboxypeptidase M OS=Homo sapiens OX=9606 GN=CPM PE=1 SV=2 | 6 |
| B2M | Beta-2-microglobulin OS=Homo sapiens OX=9606 GN=B2M PE=1 SV=1 | 1 |
| RAB14 | Ras-related protein Rab-14 OS=Homo sapiens OX=9606 GN=RAB14 PE=1 SV=4 | 7 |
| LAP3 | Cytosol aminopeptidase OS=Homo sapiens OX=9606 GN=LAP3 PE=1 SV=3 | 27 |
| C5 | Complement C5 OS=Homo sapiens OX=9606 GN=C5 PE=1 SV=4 | 73 |
| SORT1 | Sortilin OS=Homo sapiens OX=9606 GN=SORT1 PE=1 SV=3 | 5 |
| ZAR1 | Zygote arrest protein 1 OS=Homo sapiens OX=9606 GN=ZAR1 PE=1 SV=1 | 11 |
| TTC5 | Tetratricopeptide repeat protein 5 OS=Homo sapiens OX=9606 GN=TTC5 PE=1 SV=2 | 3 |
| LSM12 | Protein LSM12 OS=Homo sapiens OX=9606 GN=LSM12 PE=1 SV=2 | 1 |
| TDRKH | Tudor and KH domain-containing protein OS=Homo sapiens OX=9606 GN=TDRKH PE=1 SV=2 | 20 |
| PTGDS | Prostaglandin-H2 D-isomerase OS=Homo sapiens OX=9606 GN=PTGDS PE=1 SV=1 | 6 |
| RNF14 | E3 ubiquitin-protein ligase RNF14 OS=Homo sapiens OX=9606 GN=RNF14 PE=1 SV=1 | 4 |
| STX8 | Syntaxin-8 OS=Homo sapiens OX=9606 GN=STX8 PE=1 SV=2 | 1 |
| TSPAN4 | Tetraspanin-4 OS=Homo sapiens OX=9606 GN=TSPAN4 PE=1 SV=1 | 1 |
| UPF2 | Regulator of nonsense transcripts 2 OS=Homo sapiens OX=9606 GN=UPF2 PE=1 SV=1 | 3 |
| APEX1 | DNA-(apurinic or apyrimidinic site) endonuclease OS=Homo sapiens OX=9606 GN=APEX1 PE=1 SV=2 | 1 |
| MCAM | Cell surface glycoprotein MUC18 OS=Homo sapiens OX=9606 GN=MCAM PE=1 SV=2 | 10 |
| PSMB6 | Proteasome subunit beta type-6 OS=Homo sapiens OX=9606 GN=PSMB6 PE=1 SV=4 | 7 |
| PPP2R5C | Serine/threonine-protein phosphatase 2A 56 kDa regulatory subunit gamma isoform OS=Homo sapiens OX=9606 GN=PPP2R5C PE=1 SV=3 | 1 |
| CTSC | Dipeptidyl peptidase 1 OS=Homo sapiens OX=9606 GN=CTSC PE=1 SV=2 | 7 |
| PSMB3 | Proteasome subunit beta type-3 OS=Homo sapiens OX=9606 GN=PSMB3 PE=1 SV=2 | 6 |
| USP4 | Ubiquitin carboxyl-terminal hydrolase 4 OS=Homo sapiens OX=9606 GN=USP4 PE=1 SV=3 | 1 |
| ANGEL2 | Protein angel homolog 2 OS=Homo sapiens OX=9606 GN=ANGEL2 PE=1 SV=1 | 3 |
| ARHGEF16 | Rho guanine nucleotide exchange factor 16 OS=Homo sapiens OX=9606 GN=ARHGEF16 PE=1 SV=1 | 5 |
| EML1 | Echinoderm microtubule-associated protein-like 1 OS=Homo sapiens OX=9606 GN=EML1 PE=1 SV=3 | 3 |
| YTHDF3 | YTH domain-containing family protein 3 OS=Homo sapiens OX=9606 GN=YTHDF3 PE=1 SV=1 | 3 |
| CRYM | Ketimine reductase mu-crystallin OS=Homo sapiens OX=9606 GN=CRYM PE=1 SV=1 | 13 |
| ABL2 | Tyrosine-protein kinase ABL2 OS=Homo sapiens OX=9606 GN=ABL2 PE=1 SV=1 | 1 |
| AIM2 | Interferon-inducible protein AIM2 OS=Homo sapiens OX=9606 GN=AIM2 PE=1 SV=1 | 1 |
| ALAS1 | 5-aminolevulinate synthase, non-specific, mitochondrial OS=Homo sapiens OX=9606 GN=ALAS1 PE=1 SV=2 | 1 |
| GNB2 | Guanine nucleotide-binding protein G(I)/G(S)/G(T) subunit beta-2 OS=Homo sapiens OX=9606 GN=GNB2 PE=1 SV=3 | 3 |
| SNRNP70 | U1 small nuclear ribonucleoprotein 70 kDa OS=Homo sapiens OX=9606 GN=SNRNP70 PE=1 SV=2 | 4 |
| TAX1BP1 | Tax1-binding protein 1 OS=Homo sapiens OX=9606 GN=TAX1BP1 PE=1 SV=2 | 12 |
| RAB35 | Ras-related protein Rab-35 OS=Homo sapiens OX=9606 GN=RAB35 PE=1 SV=1 | 3 |
| GAS6 | Growth arrest-specific protein 6 OS=Homo sapiens OX=9606 GN=GAS6 PE=1 SV=3 | 9 |
| BCAS3 | BCAS3 microtubule associated cell migration factor OS=Homo sapiens OX=9606 GN=BCAS3 PE=1 SV=3 | 4 |
| GUSB | Beta-glucuronidase OS=Homo sapiens OX=9606 GN=GUSB PE=1 SV=2 | 1 |
| CARHSP1 | Calcium-regulated heat-stable protein 1 OS=Homo sapiens OX=9606 GN=CARHSP1 PE=1 SV=2 | 4 |
| STK24 | Serine/threonine-protein kinase 24 OS=Homo sapiens OX=9606 GN=STK24 PE=1 SV=1 | 5 |
| AAMDC | Mth938 domain-containing protein OS=Homo sapiens OX=9606 GN=AAMDC PE=1 SV=1 | 4 |
| PTRH2 | Peptidyl-tRNA hydrolase 2, mitochondrial OS=Homo sapiens OX=9606 GN=PTRH2 PE=1 SV=1 | 2 |
| MRPS10 | 28S ribosomal protein S10, mitochondrial OS=Homo sapiens OX=9606 GN=MRPS10 PE=1 SV=2 | 2 |
| STAM | Signal transducing adapter molecule 1 OS=Homo sapiens OX=9606 GN=STAM PE=1 SV=3 | 3 |
| C15orf40 | UPF0235 protein C15orf40 OS=Homo sapiens OX=9606 GN=C15orf40 PE=1 SV=2 | 1 |
| NPEPL1 | Probable aminopeptidase NPEPL1 OS=Homo sapiens OX=9606 GN=NPEPL1 PE=1 SV=3 | 1 |
| MTDH | Protein LYRIC OS=Homo sapiens OX=9606 GN=MTDH PE=1 SV=2 | 3 |
| NUDT1 | Oxidized purine nucleoside triphosphate hydrolase OS=Homo sapiens OX=9606 GN=NUDT1 PE=1 SV=4 | 2 |
| MRPS28 | 28S ribosomal protein S28, mitochondrial OS=Homo sapiens OX=9606 GN=MRPS28 PE=1 SV=1 | 3 |
| VKORC1 | Vitamin K epoxide reductase complex subunit 1 OS=Homo sapiens OX=9606 GN=VKORC1 PE=1 SV=1 | 1 |
| MSH2 | DNA mismatch repair protein Msh2 OS=Homo sapiens OX=9606 GN=MSH2 PE=1 SV=1 | 2 |
| PRSS2 | Trypsin-2 OS=Homo sapiens OX=9606 GN=PRSS2 PE=1 SV=1 | 2 |
| PTBP1 | Polypyrimidine tract-binding protein 1 OS=Homo sapiens OX=9606 GN=PTBP1 PE=1 SV=2 | 1 |
| ATXN3 | Ataxin-3 OS=Homo sapiens OX=9606 GN=ATXN3 PE=1 SV=5 | 1 |
| KLHDC4 | Kelch domain-containing protein 4 OS=Homo sapiens OX=9606 GN=KLHDC4 PE=1 SV=1 | 1 |
| MCTS1 | Malignant T-cell-amplified sequence 1 OS=Homo sapiens OX=9606 GN=MCTS1 PE=1 SV=1 | 5 |
| KRT4 | Keratin, type II cytoskeletal 4 OS=Homo sapiens OX=9606 GN=KRT4 PE=1 SV=5 | 23 |
| AKAP7 | A-kinase anchor protein 7 isoform gamma OS=Homo sapiens OX=9606 GN=AKAP7 PE=1 SV=2 | 2 |
| AMOTL2 | Angiomotin-like protein 2 OS=Homo sapiens OX=9606 GN=AMOTL2 PE=1 SV=3 | 1 |
| HDLBP | Vigilin OS=Homo sapiens OX=9606 GN=HDLBP PE=1 SV=2 | 2 |
| CEMIP2 | Cell surface hyaluronidase OS=Homo sapiens OX=9606 GN=CEMIP2 PE=1 SV=1 | 2 |
| "HBA1 | Hemoglobin subunit alpha OS=Homo sapiens OX=9606 GN=HBA2 PE=1 SV=2 | 5 |
| NAPRT | Nicotinate phosphoribosyltransferase OS=Homo sapiens OX=9606 GN=NAPRT PE=1 SV=2 | 11 |
| CCDC43 | Coiled-coil domain-containing protein 43 OS=Homo sapiens OX=9606 GN=CCDC43 PE=1 SV=2 | 1 |
| UFD1 | Ubiquitin recognition factor in ER-associated degradation protein 1 OS=Homo sapiens OX=9606 GN=UFD1 PE=1 SV=3 | 3 |
| EIF4EBP1 | Eukaryotic translation initiation factor 4E-binding protein 1 OS=Homo sapiens OX=9606 GN=EIF4EBP1 PE=1 SV=3 | 1 |
| F10 | Coagulation factor X OS=Homo sapiens OX=9606 GN=F10 PE=1 SV=2 | 1 |
| RSU1 | Ras suppressor protein 1 OS=Homo sapiens OX=9606 GN=RSU1 PE=1 SV=3 | 3 |
| LMNB2 | Lamin-B2 OS=Homo sapiens OX=9606 GN=LMNB2 PE=1 SV=4 | 7 |
| TMED7 | Transmembrane emp24 domain-containing protein 7 OS=Homo sapiens OX=9606 GN=TMED7 PE=1 SV=2 | 4 |
| CEP43 | Centrosomal protein 43 OS=Homo sapiens OX=9606 GN=CEP43 PE=1 SV=1 | 2 |
| TMED3 | Transmembrane emp24 domain-containing protein 3 OS=Homo sapiens OX=9606 GN=TMED3 PE=1 SV=1 | 3 |
| B3GAT3 | Galactosylgalactosylxylosylprotein 3-beta-glucuronosyltransferase 3 OS=Homo sapiens OX=9606 GN=B3GAT3 PE=1 SV=2 | 1 |
| TOMM6 | Mitochondrial import receptor subunit TOM6 homolog OS=Homo sapiens OX=9606 GN=TOMM6 PE=1 SV=1 | 1 |
| EEA1 | Early endosome antigen 1 OS=Homo sapiens OX=9606 GN=EEA1 PE=1 SV=2 | 1 |
| EXOSC9 | Exosome complex component RRP45 OS=Homo sapiens OX=9606 GN=EXOSC9 PE=1 SV=3 | 1 |
| MTMR9 | Myotubularin-related protein 9 OS=Homo sapiens OX=9606 GN=MTMR9 PE=1 SV=1 | 2 |
| MRPL11 | 39S ribosomal protein L11, mitochondrial OS=Homo sapiens OX=9606 GN=MRPL11 PE=1 SV=1 | 3 |
| KCTD5 | BTB/POZ domain-containing protein KCTD5 OS=Homo sapiens OX=9606 GN=KCTD5 PE=1 SV=1 | 2 |
| HSPA14 | Heat shock 70 kDa protein 14 OS=Homo sapiens OX=9606 GN=HSPA14 PE=1 SV=1 | 1 |
| MICU1 | Calcium uptake protein 1, mitochondrial OS=Homo sapiens OX=9606 GN=MICU1 PE=1 SV=1 | 1 |
| GSTA4 | Glutathione S-transferase A4 OS=Homo sapiens OX=9606 GN=GSTA4 PE=1 SV=1 | 1 |
| PDK3 | [Pyruvate dehydrogenase (acetyl-transferring)] kinase isozyme 3, mitochondrial OS=Homo sapiens OX=9606 GN=PDK3 PE=1 SV=1 | 3 |
| SCO1 | Protein SCO1 homolog, mitochondrial OS=Homo sapiens OX=9606 GN=SCO1 PE=1 SV=1 | 1 |
| IRF6 | Interferon regulatory factor 6 OS=Homo sapiens OX=9606 GN=IRF6 PE=1 SV=1 | 9 |
| VLDLR | Very low-density lipoprotein receptor OS=Homo sapiens OX=9606 GN=VLDLR PE=1 SV=1 | 5 |
| SNRPB | Small nuclear ribonucleoprotein-associated proteins B and B' OS=Homo sapiens OX=9606 GN=SNRPB PE=1 SV=2 | 2 |
| ASS1 | Argininosuccinate synthase OS=Homo sapiens OX=9606 GN=ASS1 PE=1 SV=2 | 1 |
| MGMT | Methylated-DNA--protein-cysteine methyltransferase OS=Homo sapiens OX=9606 GN=MGMT PE=1 SV=1 | 6 |
| RAB25 | Ras-related protein Rab-25 OS=Homo sapiens OX=9606 GN=RAB25 PE=1 SV=2 | 1 |
| CDH17 | Cadherin-17 OS=Homo sapiens OX=9606 GN=CDH17 PE=1 SV=3 | 5 |
| CYBC1 | Cytochrome b-245 chaperone 1 OS=Homo sapiens OX=9606 GN=CYBC1 PE=1 SV=1 | 1 |
| PTGES2 | Prostaglandin E synthase 2 OS=Homo sapiens OX=9606 GN=PTGES2 PE=1 SV=1 | 3 |
| FBXL13 | Dynein regulatory complex subunit 6 OS=Homo sapiens OX=9606 GN=FBXL13 PE=2 SV=3 | 1 |
| ENO1 | Alpha-enolase OS=Homo sapiens OX=9606 GN=ENO1 PE=1 SV=2 | 31 |
| CPNE3 | Copine-3 OS=Homo sapiens OX=9606 GN=CPNE3 PE=1 SV=1 | 10 |
| ILK | Integrin-linked protein kinase OS=Homo sapiens OX=9606 GN=ILK PE=1 SV=2 | 2 |
| GPR107 | Protein GPR107 OS=Homo sapiens OX=9606 GN=GPR107 PE=1 SV=1 | 7 |
| DMBT1 | Deleted in malignant brain tumors 1 protein OS=Homo sapiens OX=9606 GN=DMBT1 PE=1 SV=2 | 2 |
| CRPPA | D-ribitol-5-phosphate cytidylyltransferase OS=Homo sapiens OX=9606 GN=CRPPA PE=1 SV=2 | 5 |
| XXYLT1 | Xyloside xylosyltransferase 1 OS=Homo sapiens OX=9606 GN=XXYLT1 PE=1 SV=1 | 2 |
| ASAH1 | Acid ceramidase OS=Homo sapiens OX=9606 GN=ASAH1 PE=1 SV=5 | 10 |
| CYP20A1 | Cytochrome P450 20A1 OS=Homo sapiens OX=9606 GN=CYP20A1 PE=1 SV=1 | 2 |
| DTD2 | D-aminoacyl-tRNA deacylase 2 OS=Homo sapiens OX=9606 GN=DTD2 PE=1 SV=1 | 1 |
| GZF1 | GDNF-inducible zinc finger protein 1 OS=Homo sapiens OX=9606 GN=GZF1 PE=1 SV=1 | 1 |
| RPL8 | 60S ribosomal protein L8 OS=Homo sapiens OX=9606 GN=RPL8 PE=1 SV=2 | 6 |
| FGA | Fibrinogen alpha chain OS=Homo sapiens OX=9606 GN=FGA PE=1 SV=2 | 18 |
| YBX3 | Y-box-binding protein 3 OS=Homo sapiens OX=9606 GN=YBX3 PE=1 SV=4 | 3 |
| TOMM20 | Mitochondrial import receptor subunit TOM20 homolog OS=Homo sapiens OX=9606 GN=TOMM20 PE=1 SV=1 | 2 |
| ELMO2 | Engulfment and cell motility protein 2 OS=Homo sapiens OX=9606 GN=ELMO2 PE=1 SV=2 | 6 |
| SF3B2 | Splicing factor 3B subunit 2 OS=Homo sapiens OX=9606 GN=SF3B2 PE=1 SV=2 | 1 |
| PSMB4 | Proteasome subunit beta type-4 OS=Homo sapiens OX=9606 GN=PSMB4 PE=1 SV=4 | 7 |
| QSOX1 | Sulfhydryl oxidase 1 OS=Homo sapiens OX=9606 GN=QSOX1 PE=1 SV=3 | 2 |
| LACRT | Extracellular glycoprotein lacritin OS=Homo sapiens OX=9606 GN=LACRT PE=1 SV=1 | 3 |
| HDGFL2 | Hepatoma-derived growth factor-related protein 2 OS=Homo sapiens OX=9606 GN=HDGFL2 PE=1 SV=1 | 1 |
| ALDH2 | Aldehyde dehydrogenase, mitochondrial OS=Homo sapiens OX=9606 GN=ALDH2 PE=1 SV=2 | 4 |
| YIPF6 | Protein YIPF6 OS=Homo sapiens OX=9606 GN=YIPF6 PE=1 SV=2 | 1 |
| LSS | Lanosterol synthase OS=Homo sapiens OX=9606 GN=LSS PE=1 SV=1 | 2 |
| ITIH2 | Inter-alpha-trypsin inhibitor heavy chain H2 OS=Homo sapiens OX=9606 GN=ITIH2 PE=1 SV=2 | 20 |
| LMO7 | LIM domain only protein 7 OS=Homo sapiens OX=9606 GN=LMO7 PE=1 SV=3 | 3 |
| GFM2 | Ribosome-releasing factor 2, mitochondrial OS=Homo sapiens OX=9606 GN=GFM2 PE=1 SV=1 | 6 |
| C1orf122 | Uncharacterized protein C1orf122 OS=Homo sapiens OX=9606 GN=C1orf122 PE=4 SV=2 | 1 |
| ATP6V0D2 | V-type proton ATPase subunit d 2 OS=Homo sapiens OX=9606 GN=ATP6V0D2 PE=1 SV=1 | 1 |
| GM2A | Ganglioside GM2 activator OS=Homo sapiens OX=9606 GN=GM2A PE=1 SV=4 | 6 |
| CNPY4 | Protein canopy homolog 4 OS=Homo sapiens OX=9606 GN=CNPY4 PE=1 SV=1 | 2 |
| NAA25 | N-alpha-acetyltransferase 25, NatB auxiliary subunit OS=Homo sapiens OX=9606 GN=NAA25 PE=1 SV=1 | 1 |
| PYCR3 | Pyrroline-5-carboxylate reductase 3 OS=Homo sapiens OX=9606 GN=PYCR3 PE=1 SV=3 | 2 |
| GRN | Progranulin OS=Homo sapiens OX=9606 GN=GRN PE=1 SV=2 | 8 |
| HTATIP2 | Oxidoreductase HTATIP2 OS=Homo sapiens OX=9606 GN=HTATIP2 PE=1 SV=2 | 3 |
| GNAT2 | Guanine nucleotide-binding protein G(t) subunit alpha-2 OS=Homo sapiens OX=9606 GN=GNAT2 PE=1 SV=4 | 3 |
| DCD | Dermcidin OS=Homo sapiens OX=9606 GN=DCD PE=1 SV=2 | 6 |
| METAP2 | Methionine aminopeptidase 2 OS=Homo sapiens OX=9606 GN=METAP2 PE=1 SV=1 | 5 |
| CPT1A | Carnitine O-palmitoyltransferase 1, liver isoform OS=Homo sapiens OX=9606 GN=CPT1A PE=1 SV=2 | 1 |
| ILF2 | Interleukin enhancer-binding factor 2 OS=Homo sapiens OX=9606 GN=ILF2 PE=1 SV=2 | 4 |
| FUT8 | Alpha-(1,6)-fucosyltransferase OS=Homo sapiens OX=9606 GN=FUT8 PE=1 SV=2 | 1 |
| KHDRBS1 | KH domain-containing, RNA-binding, signal transduction-associated protein 1 OS=Homo sapiens OX=9606 GN=KHDRBS1 PE=1 SV=1 | 2 |
| MTO1 | Protein MTO1 homolog, mitochondrial OS=Homo sapiens OX=9606 GN=MTO1 PE=1 SV=2 | 2 |
| MRPS18B | 28S ribosomal protein S18b, mitochondrial OS=Homo sapiens OX=9606 GN=MRPS18B PE=1 SV=1 | 1 |
| RPL35 | 60S ribosomal protein L35 OS=Homo sapiens OX=9606 GN=RPL35 PE=1 SV=2 | 2 |
| MTFR1L | Mitochondrial fission regulator 1-like OS=Homo sapiens OX=9606 GN=MTFR1L PE=1 SV=2 | 1 |
| DEGS1 | Sphingolipid delta(4)-desaturase DES1 OS=Homo sapiens OX=9606 GN=DEGS1 PE=1 SV=1 | 1 |
| MED15 | Mediator of RNA polymerase II transcription subunit 15 OS=Homo sapiens OX=9606 GN=MED15 PE=1 SV=2 | 1 |
| PTN | Pleiotrophin OS=Homo sapiens OX=9606 GN=PTN PE=1 SV=1 | 1 |
| GDA | Guanine deaminase OS=Homo sapiens OX=9606 GN=GDA PE=1 SV=1 | 16 |
| MLLT11 | Protein AF1q OS=Homo sapiens OX=9606 GN=MLLT11 PE=1 SV=1 | 3 |
| CSRP2 | Cysteine and glycine-rich protein 2 OS=Homo sapiens OX=9606 GN=CSRP2 PE=1 SV=3 | 1 |
| SYNRG | Synergin gamma OS=Homo sapiens OX=9606 GN=SYNRG PE=1 SV=2 | 1 |
| COG7 | Conserved oligomeric Golgi complex subunit 7 OS=Homo sapiens OX=9606 GN=COG7 PE=1 SV=1 | 2 |
| ITPK1 | Inositol-tetrakisphosphate 1-kinase OS=Homo sapiens OX=9606 GN=ITPK1 PE=1 SV=2 | 1 |
| MRPL52 | 39S ribosomal protein L52, mitochondrial OS=Homo sapiens OX=9606 GN=MRPL52 PE=1 SV=2 | 1 |
| IGLC2 | Immunoglobulin lambda constant 2 OS=Homo sapiens OX=9606 GN=IGLC2 PE=1 SV=1 | 4 |
| IVL | Involucrin OS=Homo sapiens OX=9606 GN=IVL PE=1 SV=2 | 4 |
| ELAVL2 | ELAV-like protein 2 OS=Homo sapiens OX=9606 GN=ELAVL2 PE=1 SV=2 | 3 |
| WDR91 | WD repeat-containing protein 91 OS=Homo sapiens OX=9606 GN=WDR91 PE=1 SV=2 | 2 |
| COPRS | Coordinator of PRMT5 and differentiation stimulator OS=Homo sapiens OX=9606 GN=COPRS PE=1 SV=3 | 1 |
| LYPLA2 | Acyl-protein thioesterase 2 OS=Homo sapiens OX=9606 GN=LYPLA2 PE=1 SV=1 | 5 |
| TSC22D1 | TSC22 domain family protein 1 OS=Homo sapiens OX=9606 GN=TSC22D1 PE=1 SV=3 | 3 |
| CPEB1 | Cytoplasmic polyadenylation element-binding protein 1 OS=Homo sapiens OX=9606 GN=CPEB1 PE=1 SV=1 | 6 |
| ANXA4 | Annexin A4 OS=Homo sapiens OX=9606 GN=ANXA4 PE=1 SV=4 | 11 |
| DLG1 | Disks large homolog 1 OS=Homo sapiens OX=9606 GN=DLG1 PE=1 SV=2 | 1 |
| LDHB | L-lactate dehydrogenase B chain OS=Homo sapiens OX=9606 GN=LDHB PE=1 SV=2 | 7 |
| JAM3 | Junctional adhesion molecule C OS=Homo sapiens OX=9606 GN=JAM3 PE=1 SV=1 | 2 |
| IMPA2 | Inositol monophosphatase 2 OS=Homo sapiens OX=9606 GN=IMPA2 PE=1 SV=1 | 4 |
| PSG1 | Pregnancy-specific beta-1-glycoprotein 1 OS=Homo sapiens OX=9606 GN=PSG1 PE=1 SV=1 | 1 |
| FSCN1 | Fascin OS=Homo sapiens OX=9606 GN=FSCN1 PE=1 SV=3 | 15 |
| GAN | Gigaxonin OS=Homo sapiens OX=9606 GN=GAN PE=1 SV=1 | 3 |
| EIPR1 | EARP and GARP complex-interacting protein 1 OS=Homo sapiens OX=9606 GN=EIPR1 PE=1 SV=2 | 2 |
| TGM7 | Protein-glutamine gamma-glutamyltransferase Z OS=Homo sapiens OX=9606 GN=TGM7 PE=1 SV=1 | 2 |
| NDRG3 | Protein NDRG3 OS=Homo sapiens OX=9606 GN=NDRG3 PE=1 SV=2 | 3 |
| DAZAP1 | DAZ-associated protein 1 OS=Homo sapiens OX=9606 GN=DAZAP1 PE=1 SV=1 | 2 |
| STRA8 | Stimulated by retinoic acid gene 8 protein homolog OS=Homo sapiens OX=9606 GN=STRA8 PE=1 SV=1 | 1 |
| -.22 | Translation machinery-associated protein 7B OS=Homo sapiens OX=9606 GN=TMA7B PE=3 SV=1 | 2 |
| MPV17 | Protein Mpv17 OS=Homo sapiens OX=9606 GN=MPV17 PE=1 SV=1 | 1 |
| TBC1D10A | TBC1 domain family member 10A OS=Homo sapiens OX=9606 GN=TBC1D10A PE=1 SV=1 | 1 |
| UQCR10 | Cytochrome b-c1 complex subunit 9 OS=Homo sapiens OX=9606 GN=UQCR10 PE=1 SV=3 | 2 |
| GATD1 | Glutamine amidotransferase-like class 1 domain-containing protein 1 OS=Homo sapiens OX=9606 GN=GATD1 PE=1 SV=1 | 1 |
| PNLDC1 | Poly(A)-specific ribonuclease PNLDC1 OS=Homo sapiens OX=9606 GN=PNLDC1 PE=1 SV=2 | 8 |
| PLEC | Plectin OS=Homo sapiens OX=9606 GN=PLEC PE=1 SV=3 | 38 |
| IGF1R | Insulin-like growth factor 1 receptor OS=Homo sapiens OX=9606 GN=IGF1R PE=1 SV=1 | 4 |
| LARP4 | La-related protein 4 OS=Homo sapiens OX=9606 GN=LARP4 PE=1 SV=3 | 2 |
| MT2A | Metallothionein-2 OS=Homo sapiens OX=9606 GN=MT2A PE=1 SV=1 | 1 |
| ARID1A | AT-rich interactive domain-containing protein 1A OS=Homo sapiens OX=9606 GN=ARID1A PE=1 SV=3 | 1 |
| NCAPH | Condensin complex subunit 2 OS=Homo sapiens OX=9606 GN=NCAPH PE=1 SV=3 | 3 |
| CTSV | Cathepsin L2 OS=Homo sapiens OX=9606 GN=CTSV PE=1 SV=2 | 2 |
| SNU13 | NHP2-like protein 1 OS=Homo sapiens OX=9606 GN=SNU13 PE=1 SV=3 | 2 |
| CD151 | CD151 antigen OS=Homo sapiens OX=9606 GN=CD151 PE=1 SV=3 | 2 |
| ARPC1A | Actin-related protein 2/3 complex subunit 1A OS=Homo sapiens OX=9606 GN=ARPC1A PE=1 SV=2 | 1 |
| CCDC115 | Coiled-coil domain-containing protein 115 OS=Homo sapiens OX=9606 GN=CCDC115 PE=1 SV=1 | 2 |
| IGKV4-1 | Immunoglobulin kappa variable 4-1 OS=Homo sapiens OX=9606 GN=IGKV4-1 PE=1 SV=1 | 1 |
| WDR12 | Ribosome biogenesis protein WDR12 OS=Homo sapiens OX=9606 GN=WDR12 PE=1 SV=2 | 1 |
| GMPR2 | GMP reductase 2 OS=Homo sapiens OX=9606 GN=GMPR2 PE=1 SV=1 | 3 |
| NT5C | 5'(3')-deoxyribonucleotidase, cytosolic type OS=Homo sapiens OX=9606 GN=NT5C PE=1 SV=2 | 3 |
| TUBGCP6 | Gamma-tubulin complex component 6 OS=Homo sapiens OX=9606 GN=TUBGCP6 PE=1 SV=3 | 1 |
| TF | Serotransferrin OS=Homo sapiens OX=9606 GN=TF PE=1 SV=4 | 47 |
| TUBB | Tubulin beta chain OS=Homo sapiens OX=9606 GN=TUBB PE=1 SV=2 | 10 |
| CCDC88A | Girdin OS=Homo sapiens OX=9606 GN=CCDC88A PE=1 SV=2 | 2 |
| ITIH3 | Inter-alpha-trypsin inhibitor heavy chain H3 OS=Homo sapiens OX=9606 GN=ITIH3 PE=1 SV=2 | 10 |
| BTF3 | Transcription factor BTF3 OS=Homo sapiens OX=9606 GN=BTF3 PE=1 SV=1 | 1 |
| EIF4E3 | Eukaryotic translation initiation factor 4E type 3 OS=Homo sapiens OX=9606 GN=EIF4E3 PE=1 SV=4 | 1 |
| CTSA | Lysosomal protective protein OS=Homo sapiens OX=9606 GN=CTSA PE=1 SV=2 | 5 |
| LCN2 | Neutrophil gelatinase-associated lipocalin OS=Homo sapiens OX=9606 GN=LCN2 PE=1 SV=2 | 4 |
| SYT16 | Synaptotagmin-16 OS=Homo sapiens OX=9606 GN=SYT16 PE=1 SV=2 | 2 |
| UAP1 | UDP-N-acetylhexosamine pyrophosphorylase OS=Homo sapiens OX=9606 GN=UAP1 PE=1 SV=3 | 1 |
| FGD3 | FYVE, RhoGEF and PH domain-containing protein 3 OS=Homo sapiens OX=9606 GN=FGD3 PE=1 SV=1 | 1 |
| MRPS6 | 28S ribosomal protein S6, mitochondrial OS=Homo sapiens OX=9606 GN=MRPS6 PE=1 SV=3 | 1 |
| SERPINB7 | Serpin B7 OS=Homo sapiens OX=9606 GN=SERPINB7 PE=1 SV=1 | 12 |
| NRDC | Nardilysin OS=Homo sapiens OX=9606 GN=NRDC PE=1 SV=3 | 4 |
| SERPINB4 | Serpin B4 OS=Homo sapiens OX=9606 GN=SERPINB4 PE=1 SV=2 | 2 |
| IGKV3D-11 | Immunoglobulin kappa variable 3D-11 OS=Homo sapiens OX=9606 GN=IGKV3D-11 PE=3 SV=6 | 1 |
| SMAP | Small acidic protein OS=Homo sapiens OX=9606 GN=SMAP PE=1 SV=1 | 3 |
| AMY1B | Alpha-amylase 1B OS=Homo sapiens OX=9606 GN=AMY1B PE=1 SV=1 | 1 |
| AGBL3 | Cytosolic carboxypeptidase 3 OS=Homo sapiens OX=9606 GN=AGBL3 PE=2 SV=2 | 1 |
| IGKC | Immunoglobulin kappa constant OS=Homo sapiens OX=9606 GN=IGKC PE=1 SV=2 | 7 |
| PLEKHA5 | Pleckstrin homology domain-containing family A member 5 OS=Homo sapiens OX=9606 GN=PLEKHA5 PE=1 SV=1 | 1 |
| TXNL4A | Thioredoxin-like protein 4A OS=Homo sapiens OX=9606 GN=TXNL4A PE=1 SV=1 | 1 |
| SF3B3 | Splicing factor 3B subunit 3 OS=Homo sapiens OX=9606 GN=SF3B3 PE=1 SV=4 | 1 |
| CEP192 | Centrosomal protein of 192 kDa OS=Homo sapiens OX=9606 GN=CEP192 PE=1 SV=3 | 1 |
| MAOB | Amine oxidase [flavin-containing] B OS=Homo sapiens OX=9606 GN=MAOB PE=1 SV=3 | 2 |
| CLIC1 | Chloride intracellular channel protein 1 OS=Homo sapiens OX=9606 GN=CLIC1 PE=1 SV=4 | 4 |
| CTNNBIP1 | Beta-catenin-interacting protein 1 OS=Homo sapiens OX=9606 GN=CTNNBIP1 PE=1 SV=1 | 1 |
| HSPB1 | Heat shock protein beta-1 OS=Homo sapiens OX=9606 GN=HSPB1 PE=1 SV=2 | 22 |
| TIMM22 | Mitochondrial import inner membrane translocase subunit Tim22 OS=Homo sapiens OX=9606 GN=TIMM22 PE=1 SV=2 | 1 |
| CHMP5 | Charged multivesicular body protein 5 OS=Homo sapiens OX=9606 GN=CHMP5 PE=1 SV=1 | 1 |
| AHNAK | Neuroblast differentiation-associated protein AHNAK OS=Homo sapiens OX=9606 GN=AHNAK PE=1 SV=2 | 17 |
| THBS4 | Thrombospondin-4 OS=Homo sapiens OX=9606 GN=THBS4 PE=1 SV=2 | 1 |
| RPE | Ribulose-phosphate 3-epimerase OS=Homo sapiens OX=9606 GN=RPE PE=1 SV=1 | 1 |
| GSDMD | Gasdermin-D OS=Homo sapiens OX=9606 GN=GSDMD PE=1 SV=1 | 1 |
| CAPG | Macrophage-capping protein OS=Homo sapiens OX=9606 GN=CAPG PE=1 SV=2 | 2 |
| GOLGA7 | Golgin subfamily A member 7 OS=Homo sapiens OX=9606 GN=GOLGA7 PE=1 SV=2 | 2 |
| AKR1B15 | Aldo-keto reductase family 1 member B15 OS=Homo sapiens OX=9606 GN=AKR1B15 PE=1 SV=2 | 1 |
| CBWD6 | Zinc-regulated GTPase metalloprotein activator 1F OS=Homo sapiens OX=9606 GN=ZNG1F PE=3 SV=1 | 2 |
| APOL1 | Apolipoprotein L1 OS=Homo sapiens OX=9606 GN=APOL1 PE=1 SV=5 | 1 |
| NOTCH2 | Neurogenic locus notch homolog protein 2 OS=Homo sapiens OX=9606 GN=NOTCH2 PE=1 SV=3 | 1 |
| RBMX | RNA-binding motif protein, X chromosome OS=Homo sapiens OX=9606 GN=RBMX PE=1 SV=3 | 1 |
| SNAPIN | SNARE-associated protein Snapin OS=Homo sapiens OX=9606 GN=SNAPIN PE=1 SV=1 | 1 |
| WDR45B | WD repeat domain phosphoinositide-interacting protein 3 OS=Homo sapiens OX=9606 GN=WDR45B PE=1 SV=2 | 1 |
| SIDT2 | SID1 transmembrane family member 2 OS=Homo sapiens OX=9606 GN=SIDT2 PE=1 SV=2 | 1 |
| CYGB | Cytoglobin OS=Homo sapiens OX=9606 GN=CYGB PE=1 SV=1 | 1 |
| PARD6G | Partitioning defective 6 homolog gamma OS=Homo sapiens OX=9606 GN=PARD6G PE=1 SV=1 | 1 |
| IFITM1 | Interferon-induced transmembrane protein 1 OS=Homo sapiens OX=9606 GN=IFITM1 PE=1 SV=3 | 1 |
| MYH13 | Myosin-13 OS=Homo sapiens OX=9606 GN=MYH13 PE=2 SV=2 | 2 |
| SLC66A3 | Solute carrier family 66 member 3 OS=Homo sapiens OX=9606 GN=SLC66A3 PE=2 SV=1 | 2 |
| ATP6V0C | V-type proton ATPase 16 kDa proteolipid subunit c OS=Homo sapiens OX=9606 GN=ATP6V0C PE=1 SV=1 | 1 |
| CDIPT | CDP-diacylglycerol--inositol 3-phosphatidyltransferase OS=Homo sapiens OX=9606 GN=CDIPT PE=1 SV=1 | 1 |
| SEH1L | Nucleoporin SEH1 OS=Homo sapiens OX=9606 GN=SEH1L PE=1 SV=3 | 3 |
| DHCR7 | 7-dehydrocholesterol reductase OS=Homo sapiens OX=9606 GN=DHCR7 PE=1 SV=1 | 1 |
| CKAP4 | Cytoskeleton-associated protein 4 OS=Homo sapiens OX=9606 GN=CKAP4 PE=1 SV=2 | 3 |
| ALAD | Delta-aminolevulinic acid dehydratase OS=Homo sapiens OX=9606 GN=ALAD PE=1 SV=1 | 3 |
| DNAH9 | Dynein axonemal heavy chain 9 OS=Homo sapiens OX=9606 GN=DNAH9 PE=1 SV=3 | 1 |
| DNAJB6 | DnaJ homolog subfamily B member 6 OS=Homo sapiens OX=9606 GN=DNAJB6 PE=1 SV=2 | 1 |
| DHX9 | ATP-dependent RNA helicase A OS=Homo sapiens OX=9606 GN=DHX9 PE=1 SV=4 | 2 |
| LAMTOR4 | Ragulator complex protein LAMTOR4 OS=Homo sapiens OX=9606 GN=LAMTOR4 PE=1 SV=1 | 1 |
| OXCT1 | Succinyl-CoA:3-ketoacid coenzyme A transferase 1, mitochondrial OS=Homo sapiens OX=9606 GN=OXCT1 PE=1 SV=1 | 16 |
| CR2 | Complement receptor type 2 OS=Homo sapiens OX=9606 GN=CR2 PE=1 SV=2 | 2 |
| CRABP1 | Cellular retinoic acid-binding protein 1 OS=Homo sapiens OX=9606 GN=CRABP1 PE=1 SV=2 | 9 |
| NUP50 | Nuclear pore complex protein Nup50 OS=Homo sapiens OX=9606 GN=NUP50 PE=1 SV=2 | 1 |
| MEMO1 | Protein MEMO1 OS=Homo sapiens OX=9606 GN=MEMO1 PE=1 SV=1 | 2 |
| VASP | Vasodilator-stimulated phosphoprotein OS=Homo sapiens OX=9606 GN=VASP PE=1 SV=3 | 3 |
| RASAL1 | RasGAP-activating-like protein 1 OS=Homo sapiens OX=9606 GN=RASAL1 PE=1 SV=3 | 3 |
| CBFB | Core-binding factor subunit beta OS=Homo sapiens OX=9606 GN=CBFB PE=1 SV=2 | 1 |
| TPP1 | Tripeptidyl-peptidase 1 OS=Homo sapiens OX=9606 GN=TPP1 PE=1 SV=2 | 2 |
| CHD7 | Chromodomain-helicase-DNA-binding protein 7 OS=Homo sapiens OX=9606 GN=CHD7 PE=1 SV=3 | 1 |
| TYMP | Thymidine phosphorylase OS=Homo sapiens OX=9606 GN=TYMP PE=1 SV=2 | 6 |
| PLG | Plasminogen OS=Homo sapiens OX=9606 GN=PLG PE=1 SV=2 | 6 |
| IER5 | Immediate early response gene 5 protein OS=Homo sapiens OX=9606 GN=IER5 PE=1 SV=3 | 1 |
| CP | Ceruloplasmin OS=Homo sapiens OX=9606 GN=CP PE=1 SV=1 | 20 |
| IGHG1 | Immunoglobulin heavy constant gamma 1 OS=Homo sapiens OX=9606 GN=IGHG1 PE=1 SV=1 | 11 |
| HK1 | Hexokinase-1 OS=Homo sapiens OX=9606 GN=HK1 PE=1 SV=3 | 4 |
| RRM2 | Ribonucleoside-diphosphate reductase subunit M2 OS=Homo sapiens OX=9606 GN=RRM2 PE=1 SV=1 | 2 |
| TPM2 | Tropomyosin beta chain OS=Homo sapiens OX=9606 GN=TPM2 PE=1 SV=1 | 2 |
| MRPL48 | 39S ribosomal protein L48, mitochondrial OS=Homo sapiens OX=9606 GN=MRPL48 PE=1 SV=2 | 1 |
| HSPB6 | Heat shock protein beta-6 OS=Homo sapiens OX=9606 GN=HSPB6 PE=1 SV=2 | 1 |
| CTSD | Cathepsin D OS=Homo sapiens OX=9606 GN=CTSD PE=1 SV=1 | 16 |
| PHGDH | D-3-phosphoglycerate dehydrogenase OS=Homo sapiens OX=9606 GN=PHGDH PE=1 SV=4 | 5 |
| NCOA5 | Nuclear receptor coactivator 5 OS=Homo sapiens OX=9606 GN=NCOA5 PE=1 SV=2 | 1 |
| CHAT | Choline O-acetyltransferase OS=Homo sapiens OX=9606 GN=CHAT PE=1 SV=4 | 3 |
| GLYATL1 | Glycine N-acyltransferase-like protein 1 OS=Homo sapiens OX=9606 GN=GLYATL1 PE=1 SV=1 | 2 |
| PLS3 | Plastin-3 OS=Homo sapiens OX=9606 GN=PLS3 PE=1 SV=4 | 8 |
| MUCL1 | Mucin-like protein 1 OS=Homo sapiens OX=9606 GN=MUCL1 PE=1 SV=1 | 2 |
| AMER3 | APC membrane recruitment protein 3 OS=Homo sapiens OX=9606 GN=AMER3 PE=1 SV=2 | 1 |
| DHX34 | Probable ATP-dependent RNA helicase DHX34 OS=Homo sapiens OX=9606 GN=DHX34 PE=1 SV=2 | 1 |
| IGKV2-28 | Immunoglobulin kappa variable 2-28 OS=Homo sapiens OX=9606 GN=IGKV2-28 PE=3 SV=1 | 3 |
| SERPINB13 | Serpin B13 OS=Homo sapiens OX=9606 GN=SERPINB13 PE=1 SV=2 | 10 |
| AMBRA1 | Activating molecule in BECN1-regulated autophagy protein 1 OS=Homo sapiens OX=9606 GN=AMBRA1 PE=1 SV=2 | 1 |
| LDHA | L-lactate dehydrogenase A chain OS=Homo sapiens OX=9606 GN=LDHA PE=1 SV=2 | 14 |
| SERPINB8 | Serpin B8 OS=Homo sapiens OX=9606 GN=SERPINB8 PE=1 SV=2 | 8 |
| C6 | Complement component C6 OS=Homo sapiens OX=9606 GN=C6 PE=1 SV=3 | 2 |
| CAPN1 | Calpain-1 catalytic subunit OS=Homo sapiens OX=9606 GN=CAPN1 PE=1 SV=1 | 15 |
| ANAPC7 | Anaphase-promoting complex subunit 7 OS=Homo sapiens OX=9606 GN=ANAPC7 PE=1 SV=5 | 1 |
| JCHAIN | Immunoglobulin J chain OS=Homo sapiens OX=9606 GN=JCHAIN PE=1 SV=4 | 2 |
| FABP5 | Fatty acid-binding protein 5 OS=Homo sapiens OX=9606 GN=FABP5 PE=1 SV=3 | 12 |
| FAM184B | Protein FAM184B OS=Homo sapiens OX=9606 GN=FAM184B PE=2 SV=3 | 2 |
| EIF6 | Eukaryotic translation initiation factor 6 OS=Homo sapiens OX=9606 GN=EIF6 PE=1 SV=1 | 3 |
| CALML3 | Calmodulin-like protein 3 OS=Homo sapiens OX=9606 GN=CALML3 PE=1 SV=2 | 6 |
| IDE | Insulin-degrading enzyme OS=Homo sapiens OX=9606 GN=IDE PE=1 SV=4 | 46 |
| CALML5 | Calmodulin-like protein 5 OS=Homo sapiens OX=9606 GN=CALML5 PE=1 SV=2 | 11 |
| TTBK1 | Tau-tubulin kinase 1 OS=Homo sapiens OX=9606 GN=TTBK1 PE=1 SV=2 | 1 |
| CFB | Complement factor B OS=Homo sapiens OX=9606 GN=CFB PE=1 SV=2 | 8 |
| CST1 | Cystatin-SN OS=Homo sapiens OX=9606 GN=CST1 PE=1 SV=3 | 2 |
| RBM4 | RNA-binding protein 4 OS=Homo sapiens OX=9606 GN=RBM4 PE=1 SV=1 | 2 |
| IGHG2 | Immunoglobulin heavy constant gamma 2 OS=Homo sapiens OX=9606 GN=IGHG2 PE=1 SV=2 | 7 |
| AZGP1 | Zinc-alpha-2-glycoprotein OS=Homo sapiens OX=9606 GN=AZGP1 PE=1 SV=2 | 18 |
| A2ML1 | Alpha-2-macroglobulin-like protein 1 OS=Homo sapiens OX=9606 GN=A2ML1 PE=1 SV=3 | 22 |
| BLMH | Bleomycin hydrolase OS=Homo sapiens OX=9606 GN=BLMH PE=1 SV=1 | 18 |
| LMNA | Prelamin-A/C OS=Homo sapiens OX=9606 GN=LMNA PE=1 SV=1 | 37 |
| SERPINB5 | Serpin B5 OS=Homo sapiens OX=9606 GN=SERPINB5 PE=1 SV=2 | 10 |
| AGA | N(4)-(beta-N-acetylglucosaminyl)-L-asparaginase OS=Homo sapiens OX=9606 GN=AGA PE=1 SV=2 | 1 |
| TXLNG | Gamma-taxilin OS=Homo sapiens OX=9606 GN=TXLNG PE=1 SV=2 | 2 |
| DENND1B | DENN domain-containing protein 1B OS=Homo sapiens OX=9606 GN=DENND1B PE=1 SV=2 | 1 |
| MAGEE1 | Melanoma-associated antigen E1 OS=Homo sapiens OX=9606 GN=MAGEE1 PE=1 SV=2 | 1 |
| LCN1 | Lipocalin-1 OS=Homo sapiens OX=9606 GN=LCN1 PE=1 SV=1 | 3 |
| MAGI1 | Membrane-associated guanylate kinase, WW and PDZ domain-containing protein 1 OS=Homo sapiens OX=9606 GN=MAGI1 PE=1 SV=3 | 1 |
| ACP2 | Lysosomal acid phosphatase OS=Homo sapiens OX=9606 GN=ACP2 PE=1 SV=3 | 1 |
| KLK5 | Kallikrein-5 OS=Homo sapiens OX=9606 GN=KLK5 PE=1 SV=3 | 3 |
| KRT85 | Keratin, type II cuticular Hb5 OS=Homo sapiens OX=9606 GN=KRT85 PE=1 SV=1 | 16 |
| GGCT | Gamma-glutamylcyclotransferase OS=Homo sapiens OX=9606 GN=GGCT PE=1 SV=1 | 10 |
| IGKV3-20 | Immunoglobulin kappa variable 3-20 OS=Homo sapiens OX=9606 GN=IGKV3-20 PE=1 SV=2 | 1 |
| CNOT7 | CCR4-NOT transcription complex subunit 7 OS=Homo sapiens OX=9606 GN=CNOT7 PE=1 SV=3 | 1 |
| S100A16 | Protein S100-A16 OS=Homo sapiens OX=9606 GN=S100A16 PE=1 SV=1 | 3 |
| DOT1L | Histone-lysine N-methyltransferase, H3 lysine-79 specific OS=Homo sapiens OX=9606 GN=DOT1L PE=1 SV=3 | 2 |
| RBM3 | RNA-binding protein 3 OS=Homo sapiens OX=9606 GN=RBM3 PE=1 SV=1 | 2 |
| ATP2B3 | Plasma membrane calcium-transporting ATPase 3 OS=Homo sapiens OX=9606 GN=ATP2B3 PE=1 SV=3 | 2 |
| TXN | Thioredoxin OS=Homo sapiens OX=9606 GN=TXN PE=1 SV=3 | 9 |
| "DEFA1B | Neutrophil defensin 1 OS=Homo sapiens OX=9606 GN=DEFA1B PE=1 SV=1 | 2 |
| PLBD2 | Putative phospholipase B-like 2 OS=Homo sapiens OX=9606 GN=PLBD2 PE=1 SV=2 | 1 |
| ANXA8 | Annexin A8 OS=Homo sapiens OX=9606 GN=ANXA8 PE=1 SV=3 | 3 |
| FLG | Filaggrin OS=Homo sapiens OX=9606 GN=FLG PE=1 SV=3 | 18 |
| PCBD1 | Pterin-4-alpha-carbinolamine dehydratase OS=Homo sapiens OX=9606 GN=PCBD1 PE=1 SV=2 | 1 |
| SERPINA12 | Serpin A12 OS=Homo sapiens OX=9606 GN=SERPINA12 PE=1 SV=1 | 13 |
| DST | Dystonin OS=Homo sapiens OX=9606 GN=DST PE=1 SV=4 | 1 |
| GGA3 | ADP-ribosylation factor-binding protein GGA3 OS=Homo sapiens OX=9606 GN=GGA3 PE=1 SV=1 | 1 |
| IGHM | Immunoglobulin heavy constant mu OS=Homo sapiens OX=9606 GN=IGHM PE=1 SV=4 | 11 |
| CPE | Carboxypeptidase E OS=Homo sapiens OX=9606 GN=CPE PE=1 SV=1 | 1 |
| SNRPD3 | Small nuclear ribonucleoprotein Sm D3 OS=Homo sapiens OX=9606 GN=SNRPD3 PE=1 SV=1 | 3 |
| STX19 | Syntaxin-19 OS=Homo sapiens OX=9606 GN=STX19 PE=1 SV=1 | 1 |
| TRIR | Telomerase RNA component interacting RNase OS=Homo sapiens OX=9606 GN=TRIR PE=1 SV=1 | 1 |
| IGHV3-49 | Immunoglobulin heavy variable 3-49 OS=Homo sapiens OX=9606 GN=IGHV3-49 PE=3 SV=1 | 2 |
| RAMAC | RNA guanine-N7 methyltransferase activating subunit OS=Homo sapiens OX=9606 GN=RAMAC PE=1 SV=1 | 1 |
| HRAS | GTPase HRas OS=Homo sapiens OX=9606 GN=HRAS PE=1 SV=1 | 1 |
| PI3 | Elafin OS=Homo sapiens OX=9606 GN=PI3 PE=1 SV=3 | 1 |
| SCPEP1 | Retinoid-inducible serine carboxypeptidase OS=Homo sapiens OX=9606 GN=SCPEP1 PE=1 SV=1 | 2 |
| DENND1A | DENN domain-containing protein 1A OS=Homo sapiens OX=9606 GN=DENND1A PE=1 SV=2 | 1 |
| ADCK5 | Uncharacterized aarF domain-containing protein kinase 5 OS=Homo sapiens OX=9606 GN=ADCK5 PE=1 SV=3 | 2 |
| ESYT1 | Extended synaptotagmin-1 OS=Homo sapiens OX=9606 GN=ESYT1 PE=1 SV=1 | 1 |
| ACTN3 | Alpha-actinin-3 OS=Homo sapiens OX=9606 GN=ACTN3 PE=1 SV=2 | 9 |
| SERINC5 | Serine incorporator 5 OS=Homo sapiens OX=9606 GN=SERINC5 PE=2 SV=1 | 1 |
| APOM | Apolipoprotein M OS=Homo sapiens OX=9606 GN=APOM PE=1 SV=2 | 1 |
| MTERF3 | Transcription termination factor 3, mitochondrial OS=Homo sapiens OX=9606 GN=MTERF3 PE=1 SV=2 | 7 |
| IGHV3OR16-9 | Immunoglobulin heavy variable 3/OR16-9 (non-functional) (Fragment) OS=Homo sapiens OX=9606 GN=IGHV3OR16-9 PE=1 SV=1 | 3 |
| GDPD3 | Lysophospholipase D GDPD3 OS=Homo sapiens OX=9606 GN=GDPD3 PE=1 SV=3 | 7 |
| CAT | Catalase OS=Homo sapiens OX=9606 GN=CAT PE=1 SV=3 | 23 |
| APOA1 | Apolipoprotein A-I OS=Homo sapiens OX=9606 GN=APOA1 PE=1 SV=1 | 20 |
| APOA4 | Apolipoprotein A-IV OS=Homo sapiens OX=9606 GN=APOA4 PE=1 SV=4 | 8 |
| IGHA1 | Immunoglobulin heavy constant alpha 1 OS=Homo sapiens OX=9606 GN=IGHA1 PE=1 SV=2 | 12 |
| ACP3 | Prostatic acid phosphatase OS=Homo sapiens OX=9606 GN=ACP3 PE=1 SV=3 | 5 |
| PIGR | Polymeric immunoglobulin receptor OS=Homo sapiens OX=9606 GN=PIGR PE=1 SV=4 | 7 |
| ANXA1 | Annexin A1 OS=Homo sapiens OX=9606 GN=ANXA1 PE=1 SV=2 | 8 |
| SFN | 14-3-3 protein sigma OS=Homo sapiens OX=9606 GN=SFN PE=1 SV=1 | 6 |
| STS | Steryl-sulfatase OS=Homo sapiens OX=9606 GN=STS PE=1 SV=2 | 2 |
| NCKAP1 | Nck-associated protein 1 OS=Homo sapiens OX=9606 GN=NCKAP1 PE=1 SV=1 | 2 |
| S100A7 | Protein S100-A7 OS=Homo sapiens OX=9606 GN=S100A7 PE=1 SV=4 | 11 |
| S100P | Protein S100-P OS=Homo sapiens OX=9606 GN=S100P PE=1 SV=2 | 2 |
| KRT3 | Keratin, type II cytoskeletal 3 OS=Homo sapiens OX=9606 GN=KRT3 PE=1 SV=3 | 7 |
| THUMPD1 | THUMP domain-containing protein 1 OS=Homo sapiens OX=9606 GN=THUMPD1 PE=1 SV=2 | 2 |
| CST6 | Cystatin-M OS=Homo sapiens OX=9606 GN=CST6 PE=1 SV=1 | 4 |
| TGM5 | Protein-glutamine gamma-glutamyltransferase 5 OS=Homo sapiens OX=9606 GN=TGM5 PE=1 SV=4 | 9 |
| PIP | Prolactin-inducible protein OS=Homo sapiens OX=9606 GN=PIP PE=1 SV=1 | 6 |
| ZFPL1 | Zinc finger protein-like 1 OS=Homo sapiens OX=9606 GN=ZFPL1 PE=1 SV=2 | 1 |
| DUSP14 | Dual specificity protein phosphatase 14 OS=Homo sapiens OX=9606 GN=DUSP14 PE=1 SV=1 | 3 |
| PNP | Purine nucleoside phosphorylase OS=Homo sapiens OX=9606 GN=PNP PE=1 SV=2 | 10 |
| A2M | Alpha-2-macroglobulin OS=Homo sapiens OX=9606 GN=A2M PE=1 SV=3 | 16 |
| PRKAB1 | 5'-AMP-activated protein kinase subunit beta-1 OS=Homo sapiens OX=9606 GN=PRKAB1 PE=1 SV=4 | 1 |
| KLK10 | Kallikrein-10 OS=Homo sapiens OX=9606 GN=KLK10 PE=1 SV=3 | 4 |
| C3 | Complement C3 OS=Homo sapiens OX=9606 GN=C3 PE=1 SV=2 | 78 |
| DSC3 | Desmocollin-3 OS=Homo sapiens OX=9606 GN=DSC3 PE=1 SV=3 | 15 |
| FGB | Fibrinogen beta chain OS=Homo sapiens OX=9606 GN=FGB PE=1 SV=2 | 17 |
| GIT1 | ARF GTPase-activating protein GIT1 OS=Homo sapiens OX=9606 GN=GIT1 PE=1 SV=2 | 1 |
| IL37 | Interleukin-37 OS=Homo sapiens OX=9606 GN=IL37 PE=1 SV=1 | 4 |
| IGHV3-15 | Immunoglobulin heavy variable 3-15 OS=Homo sapiens OX=9606 GN=IGHV3-15 PE=3 SV=1 | 1 |
| KRT13 | Keratin, type I cytoskeletal 13 OS=Homo sapiens OX=9606 GN=KRT13 PE=1 SV=4 | 22 |
| SBSN | Suprabasin OS=Homo sapiens OX=9606 GN=SBSN PE=1 SV=2 | 20 |
| KRT31 | Keratin, type I cuticular Ha1 OS=Homo sapiens OX=9606 GN=KRT31 PE=1 SV=3 | 25 |
| EPPK1 | Epiplakin OS=Homo sapiens OX=9606 GN=EPPK1 PE=1 SV=3 | 30 |
| S100A6 | Protein S100-A6 OS=Homo sapiens OX=9606 GN=S100A6 PE=1 SV=1 | 1 |
| CSTA | Cystatin-A OS=Homo sapiens OX=9606 GN=CSTA PE=1 SV=1 | 12 |
| PON1 | Serum paraoxonase/arylesterase 1 OS=Homo sapiens OX=9606 GN=PON1 PE=1 SV=3 | 3 |
| S100A10 | Protein S100-A10 OS=Homo sapiens OX=9606 GN=S100A10 PE=1 SV=2 | 1 |
| CASP14 | Caspase-14 OS=Homo sapiens OX=9606 GN=CASP14 PE=1 SV=2 | 24 |
| IGKV3D-20 | Immunoglobulin kappa variable 3D-20 OS=Homo sapiens OX=9606 GN=IGKV3D-20 PE=3 SV=1 | 2 |
| POF1B | Protein POF1B OS=Homo sapiens OX=9606 GN=POF1B PE=1 SV=3 | 27 |
| KRT2 | Keratin, type II cytoskeletal 2 epidermal OS=Homo sapiens OX=9606 GN=KRT2 PE=1 SV=2 | 52 |
| DMKN | Dermokine OS=Homo sapiens OX=9606 GN=DMKN PE=1 SV=3 | 4 |
| KRT86 | Keratin, type II cuticular Hb6 OS=Homo sapiens OX=9606 GN=KRT86 PE=1 SV=1 | 27 |
| LTF | Lactotransferrin OS=Homo sapiens OX=9606 GN=LTF PE=1 SV=6 | 24 |
| NCCRP1 | F-box only protein 50 OS=Homo sapiens OX=9606 GN=NCCRP1 PE=1 SV=1 | 7 |
| "LGALS7B | Galectin-7 OS=Homo sapiens OX=9606 GN=LGALS7B PE=1 SV=2 | 11 |
| KLK7 | Kallikrein-7 OS=Homo sapiens OX=9606 GN=KLK7 PE=1 SV=1 | 7 |
| FGG | Fibrinogen gamma chain OS=Homo sapiens OX=9606 GN=FGG PE=1 SV=3 | 15 |
| ZG16B | Zymogen granule protein 16 homolog B OS=Homo sapiens OX=9606 GN=ZG16B PE=1 SV=3 | 3 |
| IL1RN | Interleukin-1 receptor antagonist protein OS=Homo sapiens OX=9606 GN=IL1RN PE=1 SV=1 | 3 |
| PKP1 | Plakophilin-1 OS=Homo sapiens OX=9606 GN=PKP1 PE=1 SV=2 | 34 |
| EVPL | Envoplakin OS=Homo sapiens OX=9606 GN=EVPL PE=1 SV=3 | 2 |
| CAMK2D | Calcium/calmodulin-dependent protein kinase type II subunit delta OS=Homo sapiens OX=9606 GN=CAMK2D PE=1 SV=3 | 1 |
| SERPINB3 | Serpin B3 OS=Homo sapiens OX=9606 GN=SERPINB3 PE=1 SV=2 | 26 |
| CDSN | Corneodesmosin OS=Homo sapiens OX=9606 GN=CDSN PE=1 SV=3 | 4 |
| CCDC88C | Protein Daple OS=Homo sapiens OX=9606 GN=CCDC88C PE=1 SV=3 | 1 |
| RAB20 | Ras-related protein Rab-20 OS=Homo sapiens OX=9606 GN=RAB20 PE=1 SV=1 | 1 |
| PLBD1 | Phospholipase B-like 1 OS=Homo sapiens OX=9606 GN=PLBD1 PE=1 SV=2 | 16 |
| S100A14 | Protein S100-A14 OS=Homo sapiens OX=9606 GN=S100A14 PE=1 SV=1 | 5 |
| LGALSL | Galectin-related protein OS=Homo sapiens OX=9606 GN=LGALSL PE=1 SV=2 | 2 |
| MYO15A | Unconventional myosin-XV OS=Homo sapiens OX=9606 GN=MYO15A PE=1 SV=2 | 1 |
| TUBB2A | Tubulin beta-2A chain OS=Homo sapiens OX=9606 GN=TUBB2A PE=1 SV=1 | 1 |
| ANXA2 | Annexin A2 OS=Homo sapiens OX=9606 GN=ANXA2 PE=1 SV=2 | 28 |
| S100A8 | Protein S100-A8 OS=Homo sapiens OX=9606 GN=S100A8 PE=1 SV=1 | 12 |
| LYZ | Lysozyme C OS=Homo sapiens OX=9606 GN=LYZ PE=1 SV=1 | 10 |
| CENPH | Centromere protein H OS=Homo sapiens OX=9606 GN=CENPH PE=1 SV=1 | 1 |
| CTSG | Cathepsin G OS=Homo sapiens OX=9606 GN=CTSG PE=1 SV=2 | 1 |
| YOD1 | Ubiquitin thioesterase OTU1 OS=Homo sapiens OX=9606 GN=YOD1 PE=1 SV=1 | 8 |
| KRT15 | Keratin, type I cytoskeletal 15 OS=Homo sapiens OX=9606 GN=KRT15 PE=1 SV=3 | 6 |
| PSAPL1 | Proactivator polypeptide-like 1 OS=Homo sapiens OX=9606 GN=PSAPL1 PE=2 SV=2 | 3 |
| S100A9 | Protein S100-A9 OS=Homo sapiens OX=9606 GN=S100A9 PE=1 SV=1 | 9 |
| DDAH2 | N(G),N(G)-dimethylarginine dimethylaminohydrolase 2 OS=Homo sapiens OX=9606 GN=DDAH2 PE=1 SV=1 | 7 |
| HAL | Histidine ammonia-lyase OS=Homo sapiens OX=9606 GN=HAL PE=1 SV=1 | 15 |
| KRT10 | Keratin, type I cytoskeletal 10 OS=Homo sapiens OX=9606 GN=KRT10 PE=1 SV=6 | 49 |
| IGHV3-7 | Immunoglobulin heavy variable 3-7 OS=Homo sapiens OX=9606 GN=IGHV3-7 PE=1 SV=2 | 2 |
| KRT76 | Keratin, type II cytoskeletal 2 oral OS=Homo sapiens OX=9606 GN=KRT76 PE=1 SV=2 | 14 |
| DSP | Desmoplakin OS=Homo sapiens OX=9606 GN=DSP PE=1 SV=3 | 241 |
| SCGB2A1 | Mammaglobin-B OS=Homo sapiens OX=9606 GN=SCGB2A1 PE=1 SV=1 | 3 |
| S100A11 | Protein S100-A11 OS=Homo sapiens OX=9606 GN=S100A11 PE=1 SV=2 | 3 |
| LMO4 | LIM domain transcription factor LMO4 OS=Homo sapiens OX=9606 GN=LMO4 PE=1 SV=1 | 1 |
| SERPINB2 | Plasminogen activator inhibitor 2 OS=Homo sapiens OX=9606 GN=SERPINB2 PE=1 SV=2 | 9 |
| NEU2 | Sialidase-2 OS=Homo sapiens OX=9606 GN=NEU2 PE=1 SV=2 | 13 |
| WFDC12 | WAP four-disulfide core domain protein 12 OS=Homo sapiens OX=9606 GN=WFDC12 PE=1 SV=1 | 3 |
| KRT75 | Keratin, type II cytoskeletal 75 OS=Homo sapiens OX=9606 GN=KRT75 PE=1 SV=2 | 1 |
| ALOX12B | Arachidonate 12-lipoxygenase, 12R-type OS=Homo sapiens OX=9606 GN=ALOX12B PE=1 SV=1 | 15 |
| CPA4 | Carboxypeptidase A4 OS=Homo sapiens OX=9606 GN=CPA4 PE=1 SV=2 | 13 |
| LGALS3 | Galectin-3 OS=Homo sapiens OX=9606 GN=LGALS3 PE=1 SV=5 | 5 |
| FLG2 | Filaggrin-2 OS=Homo sapiens OX=9606 GN=FLG2 PE=1 SV=1 | 25 |
| DSC1 | Desmocollin-1 OS=Homo sapiens OX=9606 GN=DSC1 PE=1 SV=2 | 31 |
| SCGB2A2 | Mammaglobin-A OS=Homo sapiens OX=9606 GN=SCGB2A2 PE=1 SV=1 | 1 |
| ME1 | NADP-dependent malic enzyme OS=Homo sapiens OX=9606 GN=ME1 PE=1 SV=1 | 5 |
| JUP | Junction plakoglobin OS=Homo sapiens OX=9606 GN=JUP PE=1 SV=3 | 42 |
| C1QC | Complement C1q subcomponent subunit C OS=Homo sapiens OX=9606 GN=C1QC PE=1 SV=3 | 1 |
| ALOXE3 | Hydroperoxide isomerase ALOXE3 OS=Homo sapiens OX=9606 GN=ALOXE3 PE=1 SV=1 | 5 |
| TGM3 | Protein-glutamine gamma-glutamyltransferase E OS=Homo sapiens OX=9606 GN=TGM3 PE=1 SV=4 | 34 |
| KRT23 | Keratin, type I cytoskeletal 23 OS=Homo sapiens OX=9606 GN=KRT23 PE=1 SV=2 | 20 |
| KRT80 | Keratin, type II cytoskeletal 80 OS=Homo sapiens OX=9606 GN=KRT80 PE=1 SV=2 | 30 |
| KRT9 | Keratin, type I cytoskeletal 9 OS=Homo sapiens OX=9606 GN=KRT9 PE=1 SV=3 | 49 |
| CNFN | Cornifelin OS=Homo sapiens OX=9606 GN=CNFN PE=1 SV=2 | 1 |
| ARG1 | Arginase-1 OS=Homo sapiens OX=9606 GN=ARG1 PE=1 SV=2 | 21 |
| CACNA1S | Voltage-dependent L-type calcium channel subunit alpha-1S OS=Homo sapiens OX=9606 GN=CACNA1S PE=1 SV=4 | 1 |
| KRT1 | Keratin, type II cytoskeletal 1 OS=Homo sapiens OX=9606 GN=KRT1 PE=1 SV=6 | 63 |
| IL21 | Interleukin-21 OS=Homo sapiens OX=9606 GN=IL21 PE=1 SV=3 | 1 |
| ECM1 | Extracellular matrix protein 1 OS=Homo sapiens OX=9606 GN=ECM1 PE=1 SV=2 | 15 |
| BUD23 | Probable 18S rRNA (guanine-N(7))-methyltransferase OS=Homo sapiens OX=9606 GN=BUD23 PE=1 SV=2 | 1 |
| MYCBP2 | E3 ubiquitin-protein ligase MYCBP2 OS=Homo sapiens OX=9606 GN=MYCBP2 PE=1 SV=4 | 1 |
| IL36G | Interleukin-36 gamma OS=Homo sapiens OX=9606 GN=IL36G PE=1 SV=1 | 8 |
| KRT77 | Keratin, type II cytoskeletal 1b OS=Homo sapiens OX=9606 GN=KRT77 PE=1 SV=3 | 31 |
| KRT19 | Keratin, type I cytoskeletal 19 OS=Homo sapiens OX=9606 GN=KRT19 PE=1 SV=4 | 5 |
| KRT78 | Keratin, type II cytoskeletal 78 OS=Homo sapiens OX=9606 GN=KRT78 PE=1 SV=2 | 29 |
| SERPINB12 | Serpin B12 OS=Homo sapiens OX=9606 GN=SERPINB12 PE=1 SV=1 | 21 |
| KRT34 | Keratin, type I cuticular Ha4 OS=Homo sapiens OX=9606 GN=KRT34 PE=1 SV=2 | 11 |
| EGFL6 | Epidermal growth factor-like protein 6 OS=Homo sapiens OX=9606 GN=EGFL6 PE=1 SV=1 | 1 |
| LORICRIN | Loricrin OS=Homo sapiens OX=9606 GN=LORICRIN PE=1 SV=2 | 4 |
| RNASE7 | Ribonuclease 7 OS=Homo sapiens OX=9606 GN=RNASE7 PE=1 SV=2 | 5 |
| XP32 | Skin-specific protein 32 OS=Homo sapiens OX=9606 GN=XP32 PE=1 SV=1 | 8 |
| CIC | Protein capicua homolog OS=Homo sapiens OX=9606 GN=CIC PE=1 SV=2 | 1 |
| RPS6KA2 | Ribosomal protein S6 kinase alpha-2 OS=Homo sapiens OX=9606 GN=RPS6KA2 PE=1 SV=2 | 1 |
| DSG1 | Desmoglein-1 OS=Homo sapiens OX=9606 GN=DSG1 PE=1 SV=2 | 36 |
| KRT5 | Keratin, type II cytoskeletal 5 OS=Homo sapiens OX=9606 GN=KRT5 PE=1 SV=3 | 62 |
| TGM1 | Protein-glutamine gamma-glutamyltransferase K OS=Homo sapiens OX=9606 GN=TGM1 PE=1 SV=4 | 31 |
| CCDC194 | Coiled-coil domain-containing protein 194 OS=Homo sapiens OX=9606 GN=CCDC194 PE=3 SV=1 | 1 |
| LY6G6C | Lymphocyte antigen 6 complex locus protein G6c OS=Homo sapiens OX=9606 GN=LY6G6C PE=1 SV=1 | 2 |
| KRT72 | Keratin, type II cytoskeletal 72 OS=Homo sapiens OX=9606 GN=KRT72 PE=1 SV=2 | 5 |
| NES | Nestin OS=Homo sapiens OX=9606 GN=NES PE=1 SV=2 | 1 |
| PRCP | Lysosomal Pro-X carboxypeptidase OS=Homo sapiens OX=9606 GN=PRCP PE=1 SV=1 | 3 |
| SELENBP1 | Methanethiol oxidase OS=Homo sapiens OX=9606 GN=SELENBP1 PE=1 SV=2 | 1 |
| SPRR2D | Small proline-rich protein 2D OS=Homo sapiens OX=9606 GN=SPRR2D PE=2 SV=2 | 3 |
| KRT14 | Keratin, type I cytoskeletal 14 OS=Homo sapiens OX=9606 GN=KRT14 PE=1 SV=4 | 54 |
| SPRR2G | Small proline-rich protein 2G OS=Homo sapiens OX=9606 GN=SPRR2G PE=3 SV=1 | 4 |
| HRNR | Hornerin OS=Homo sapiens OX=9606 GN=HRNR PE=1 SV=2 | 27 |
| SLPI | Antileukoproteinase OS=Homo sapiens OX=9606 GN=SLPI PE=1 SV=2 | 1 |
| SPTBN2 | Spectrin beta chain, non-erythrocytic 2 OS=Homo sapiens OX=9606 GN=SPTBN2 PE=1 SV=3 | 1 |
| KRT12 | Keratin, type I cytoskeletal 12 OS=Homo sapiens OX=9606 GN=KRT12 PE=1 SV=1 | 2 |
| SPRR1B | Cornifin-B OS=Homo sapiens OX=9606 GN=SPRR1B PE=1 SV=2 | 5 |
| KRT17 | Keratin, type I cytoskeletal 17 OS=Homo sapiens OX=9606 GN=KRT17 PE=1 SV=2 | 30 |
| SDR9C7 | Short-chain dehydrogenase/reductase family 9C member 7 OS=Homo sapiens OX=9606 GN=SDR9C7 PE=1 SV=1 | 7 |
| KPRP | Keratinocyte proline-rich protein OS=Homo sapiens OX=9606 GN=KPRP PE=1 SV=1 | 21 |
| KRT33B | Keratin, type I cuticular Ha3-II OS=Homo sapiens OX=9606 GN=KRT33B PE=1 SV=3 | 4 |
| GSDMA | Gasdermin-A OS=Homo sapiens OX=9606 GN=GSDMA PE=1 SV=4 | 15 |
| ABCB6 | ATP-binding cassette sub-family B member 6 OS=Homo sapiens OX=9606 GN=ABCB6 PE=1 SV=1 | 1 |
| KRT16 | Keratin, type I cytoskeletal 16 OS=Homo sapiens OX=9606 GN=KRT16 PE=1 SV=4 | 34 |
| MXRA5 | Matrix-remodeling-associated protein 5 OS=Homo sapiens OX=9606 GN=MXRA5 PE=1 SV=3 | 1 |
| KRT6B | Keratin, type II cytoskeletal 6B OS=Homo sapiens OX=9606 GN=KRT6B PE=1 SV=5 | 10 |
| GPNMB | Transmembrane glycoprotein NMB OS=Homo sapiens OX=9606 GN=GPNMB PE=1 SV=2 | 1 |
| ASPRV1 | Retroviral-like aspartic protease 1 OS=Homo sapiens OX=9606 GN=ASPRV1 PE=1 SV=1 | 2 |
| TEP1 | Telomerase protein component 1 OS=Homo sapiens OX=9606 GN=TEP1 PE=1 SV=2 | 1 |
| SRCIN1 | SRC kinase signaling inhibitor 1 OS=Homo sapiens OX=9606 GN=SRCIN1 PE=1 SV=4 | 1 |
| KRT6A | Keratin, type II cytoskeletal 6A OS=Homo sapiens OX=9606 GN=KRT6A PE=1 SV=3 | 42 |
| PWP2 | Periodic tryptophan protein 2 homolog OS=Homo sapiens OX=9606 GN=PWP2 PE=1 SV=2 | 1 |
| SPRR2F | Small proline-rich protein 2F OS=Homo sapiens OX=9606 GN=SPRR2F PE=3 SV=1 | 3 |
| WDR75 | WD repeat-containing protein 75 OS=Homo sapiens OX=9606 GN=WDR75 PE=1 SV=1 | 1 |
| SENP1 | Sentrin-specific protease 1 OS=Homo sapiens OX=9606 GN=SENP1 PE=1 SV=2 | 1 |
| HAS1 | Hyaluronan synthase 1 OS=Homo sapiens OX=9606 GN=HAS1 PE=1 SV=2 | 1 |

| **Supplementary Table 9. Differential abundant proteins (DAPs) in ffEVs-treated oocytes (FDR<0.1%)** | | | |
| --- | --- | --- | --- |
| **Name** | **description** | **Fold change** | **P-value adj.** |
| HAS1 | Hyaluronan synthase 1 OS=Homo sapiens OX=9606 GN=HAS1 PE=1 SV=2 | 6.55 | 0.0925 |
| SPRR2F | Small proline-rich protein 2F OS=Homo sapiens OX=9606 GN=SPRR2F PE=3 SV=1 | 4.48 | 0.0013 |
| SRCIN1 | SRC kinase signaling inhibitor 1 OS=Homo sapiens OX=9606 GN=SRCIN1 PE=1 SV=4 | 3.64 | 0.0566 |
| ASPRV1 | Retroviral-like aspartic protease 1 OS=Homo sapiens OX=9606 GN=ASPRV1 PE=1 SV=1 | 3.61 | 0.0114 |
| GPNMB | Transmembrane glycoprotein NMB OS=Homo sapiens OX=9606 GN=GPNMB PE=1 SV=2 | 3.40 | 0.0043 |
| GSDMA | Gasdermin-A OS=Homo sapiens OX=9606 GN=GSDMA PE=1 SV=4 | 3.25 | 0.0021 |
| KPRP | Keratinocyte proline-rich protein OS=Homo sapiens OX=9606 GN=KPRP PE=1 SV=1 | 3.20 | 0.0444 |
| SDR9C7 | Short-chain dehydrogenase/reductase family 9C member 7 OS=Homo sapiens OX=9606 GN=SDR9C7 PE=1 SV=1 | 3.17 | 0.0086 |
| SLPI | Antileukoproteinase OS=Homo sapiens OX=9606 GN=SLPI PE=1 SV=2 | 2.93 | 0.0988 |
| SPRR2D | Small proline-rich protein 2D OS=Homo sapiens OX=9606 GN=SPRR2D PE=2 SV=2 | 2.79 | 0.0983 |
| SELENBP1 | Methanethiol oxidase OS=Homo sapiens OX=9606 GN=SELENBP1 PE=1 SV=2 | 2.79 | 0.0358 |
| PRCP | Lysosomal Pro-X carboxypeptidase OS=Homo sapiens OX=9606 GN=PRCP PE=1 SV=1 | 2.76 | 0.0359 |
| TGM1 | Protein-glutamine gamma-glutamyltransferase K OS=Homo sapiens OX=9606 GN=TGM1 PE=1 SV=4 | 2.71 | 0.0028 |
| CIC | Protein capicua homolog OS=Homo sapiens OX=9606 GN=CIC PE=1 SV=2 | 2.67 | 0.0000 |
| XP32 | Skin-specific protein 32 OS=Homo sapiens OX=9606 GN=XP32 PE=1 SV=1 | 2.66 | 0.0998 |
| RNASE7 | Ribonuclease 7 OS=Homo sapiens OX=9606 GN=RNASE7 PE=1 SV=2 | 2.65 | 0.0857 |
| SERPINB12 | Serpin B12 OS=Homo sapiens OX=9606 GN=SERPINB12 PE=1 SV=1 | 2.61 | 0.0477 |
| KRT78 | Keratin, type II cytoskeletal 78 OS=Homo sapiens OX=9606 GN=KRT78 PE=1 SV=2 | 2.59 | 0.0777 |
| KRT77 | Keratin, type II cytoskeletal 1b OS=Homo sapiens OX=9606 GN=KRT77 PE=1 SV=3 | 2.58 | 0.0830 |
| MYCBP2 | E3 ubiquitin-protein ligase MYCBP2 OS=Homo sapiens OX=9606 GN=MYCBP2 PE=1 SV=4 | 2.56 | 0.0322 |
| CNFN | Cornifelin OS=Homo sapiens OX=9606 GN=CNFN PE=1 SV=2 | 2.45 | 0.0972 |
| KRT80 | Keratin, type II cytoskeletal 80 OS=Homo sapiens OX=9606 GN=KRT80 PE=1 SV=2 | 2.42 | 0.0996 |
| TGM3 | Protein-glutamine gamma-glutamyltransferase E OS=Homo sapiens OX=9606 GN=TGM3 PE=1 SV=4 | 2.42 | 0.0876 |
| ALOXE3 | Hydroperoxide isomerase ALOXE3 OS=Homo sapiens OX=9606 GN=ALOXE3 PE=1 SV=1 | 2.41 | 0.0375 |
| SCGB2A2 | Mammaglobin-A OS=Homo sapiens OX=9606 GN=SCGB2A2 PE=1 SV=1 | 2.36 | 0.0991 |
| FLG2 | Filaggrin-2 OS=Homo sapiens OX=9606 GN=FLG2 PE=1 SV=1 | 2.33 | 0.0976 |
| CPA4 | Carboxypeptidase A4 OS=Homo sapiens OX=9606 GN=CPA4 PE=1 SV=2 | 2.31 | 0.0035 |
| WFDC12 | WAP four-disulfide core domain protein 12 OS=Homo sapiens OX=9606 GN=WFDC12 PE=1 SV=1 | 2.29 | 0.0523 |
| SERPINB2 | Plasminogen activator inhibitor 2 OS=Homo sapiens OX=9606 GN=SERPINB2 PE=1 SV=2 | 2.28 | 0.0905 |
| PSAPL1 | Proactivator polypeptide-like 1 OS=Homo sapiens OX=9606 GN=PSAPL1 PE=2 SV=2 | 2.15 | 0.0928 |
| LYZ | Lysozyme C OS=Homo sapiens OX=9606 GN=LYZ PE=1 SV=1 | 2.11 | 0.0386 |
| TUBB2A | Tubulin beta-2A chain OS=Homo sapiens OX=9606 GN=TUBB2A PE=1 SV=1 | 2.09 | 0.0591 |
| RAB20 | Ras-related protein Rab-20 OS=Homo sapiens OX=9606 GN=RAB20 PE=1 SV=1 | 2.07 | 0.0591 |
| SERPINB3 | Serpin B3 OS=Homo sapiens OX=9606 GN=SERPINB3 PE=1 SV=2 | 2.06 | 0.0952 |
| IL1RN | Interleukin-1 receptor antagonist protein OS=Homo sapiens OX=9606 GN=IL1RN PE=1 SV=1 | 2.02 | 0.0675 |
| KLK7 | Kallikrein-7 OS=Homo sapiens OX=9606 GN=KLK7 PE=1 SV=1 | 2.00 | 0.0980 |
| LTF | Lactotransferrin OS=Homo sapiens OX=9606 GN=LTF PE=1 SV=6 | 1.96 | 0.0223 |
| MTERF3 | Transcription termination factor 3, mitochondrial OS=Homo sapiens OX=9606 GN=MTERF3 PE=1 SV=2 | 1.46 | 0.0942 |
| ESYT1 | Extended synaptotagmin-1 OS=Homo sapiens OX=9606 GN=ESYT1 PE=1 SV=1 | 1.42 | 0.0821 |
| SNRPD3 | Small nuclear ribonucleoprotein Sm D3 OS=Homo sapiens OX=9606 GN=SNRPD3 PE=1 SV=1 | 1.33 | 0.0000 |
| FUT8 | Alpha-(1,6)-fucosyltransferase OS=Homo sapiens OX=9606 GN=FUT8 PE=1 SV=2 | 0.63 | 0.0930 |
| IRF6 | Interferon regulatory factor 6 OS=Homo sapiens OX=9606 GN=IRF6 PE=1 SV=1 | 0.54 | 0.0860 |
| TPD52 | Tumor protein D52 OS=Homo sapiens OX=9606 GN=TPD52 PE=1 SV=2 | -0.58 | 0.0997 |
| GET3 | ATPase GET3 OS=Homo sapiens OX=9606 GN=GET3 PE=1 SV=2 | -0.60 | 0.1012 |
| CPVL | Probable serine carboxypeptidase CPVL OS=Homo sapiens OX=9606 GN=CPVL PE=1 SV=2 | -0.68 | 0.0257 |
| HTRA1 | Serine protease HTRA1 OS=Homo sapiens OX=9606 GN=HTRA1 PE=1 SV=1 | -0.73 | 0.0810 |
| CHMP2A | Charged multivesicular body protein 2a OS=Homo sapiens OX=9606 GN=CHMP2A PE=1 SV=1 | -0.81 | 0.0880 |
| DERL1 | Derlin-1 OS=Homo sapiens OX=9606 GN=DERL1 PE=1 SV=1 | -0.83 | 0.0822 |
| CENPS | Centromere protein S OS=Homo sapiens OX=9606 GN=CENPS PE=1 SV=1 | -0.87 | 0.0906 |
| DMAC2L | ATP synthase subunit s, mitochondrial OS=Homo sapiens OX=9606 GN=DMAC2L PE=1 SV=3 | -1.11 | 0.0858 |
| SRP19 | Signal recognition particle 19 kDa protein OS=Homo sapiens OX=9606 GN=SRP19 PE=1 SV=3 | -1.31 | 0.0171 |
| SLC27A1 | Long-chain fatty acid transport protein 1 OS=Homo sapiens OX=9606 GN=SLC27A1 PE=1 SV=1 | -2.07 | 0.0790 |
| PKP2 | Plakophilin-2 OS=Homo sapiens OX=9606 GN=PKP2 PE=1 SV=2 | -2.13 | 0.0369 |
| UTP20 | Small subunit processome component 20 homolog OS=Homo sapiens OX=9606 GN=UTP20 PE=1 SV=3 | -2.53 | 0.0743 |
| PPIE | Peptidyl-prolyl cis-trans isomerase E OS=Homo sapiens OX=9606 GN=PPIE PE=1 SV=1 | -2.66 | 0.0009 |
| BRI3BP | BRI3-binding protein OS=Homo sapiens OX=9606 GN=BRI3BP PE=1 SV=1 | -2.67 | 0.0325 |

| **Supplementary Table 10. Differential abundant proteins (DAPs) in ffEVs-treated oocytes (FDR<0.2%)** | | | |
| --- | --- | --- | --- |
| **Name** | **description** | **Fold change** | **P-value adj.** |
| HAS1 | Hyaluronan synthase 1 OS=Homo sapiens OX=9606 GN=HAS1 PE=1 SV=2 | 6.55 | 0.0925 |
| SPRR2F | Small proline-rich protein 2F OS=Homo sapiens OX=9606 GN=SPRR2F PE=3 SV=1 | 4.48 | 0.0013 |
| KRT6A | Keratin, type II cytoskeletal 6A OS=Homo sapiens OX=9606 GN=KRT6A PE=1 SV=3 | 3.88 | 0.1627 |
| SRCIN1 | SRC kinase signaling inhibitor 1 OS=Homo sapiens OX=9606 GN=SRCIN1 PE=1 SV=4 | 3.64 | 0.0566 |
| TEP1 | Telomerase protein component 1 OS=Homo sapiens OX=9606 GN=TEP1 PE=1 SV=2 | 3.64 | 0.1944 |
| ASPRV1 | Retroviral-like aspartic protease 1 OS=Homo sapiens OX=9606 GN=ASPRV1 PE=1 SV=1 | 3.61 | 0.0114 |
| GPNMB | Transmembrane glycoprotein NMB OS=Homo sapiens OX=9606 GN=GPNMB PE=1 SV=2 | 3.40 | 0.0043 |
| GSDMA | Gasdermin-A OS=Homo sapiens OX=9606 GN=GSDMA PE=1 SV=4 | 3.25 | 0.0021 |
| KPRP | Keratinocyte proline-rich protein OS=Homo sapiens OX=9606 GN=KPRP PE=1 SV=1 | 3.20 | 0.0444 |
| SDR9C7 | Short-chain dehydrogenase/reductase family 9C member 7 OS=Homo sapiens OX=9606 GN=SDR9C7 PE=1 SV=1 | 3.17 | 0.0086 |
| SLPI | Antileukoproteinase OS=Homo sapiens OX=9606 GN=SLPI PE=1 SV=2 | 2.93 | 0.0988 |
| SPRR2D | Small proline-rich protein 2D OS=Homo sapiens OX=9606 GN=SPRR2D PE=2 SV=2 | 2.79 | 0.0983 |
| SELENBP1 | Methanethiol oxidase OS=Homo sapiens OX=9606 GN=SELENBP1 PE=1 SV=2 | 2.79 | 0.0358 |
| PRCP | Lysosomal Pro-X carboxypeptidase OS=Homo sapiens OX=9606 GN=PRCP PE=1 SV=1 | 2.76 | 0.0359 |
| KRT72 | Keratin, type II cytoskeletal 72 OS=Homo sapiens OX=9606 GN=KRT72 PE=1 SV=2 | 2.72 | 0.1806 |
| TGM1 | Protein-glutamine gamma-glutamyltransferase K OS=Homo sapiens OX=9606 GN=TGM1 PE=1 SV=4 | 2.71 | 0.0028 |
| KRT5 | Keratin, type II cytoskeletal 5 OS=Homo sapiens OX=9606 GN=KRT5 PE=1 SV=3 | 2.70 | 0.1530 |
| DSG1 | Desmoglein-1 OS=Homo sapiens OX=9606 GN=DSG1 PE=1 SV=2 | 2.70 | 0.1615 |
| CIC | Protein capicua homolog OS=Homo sapiens OX=9606 GN=CIC PE=1 SV=2 | 2.67 | 0.0000 |
| XP32 | Skin-specific protein 32 OS=Homo sapiens OX=9606 GN=XP32 PE=1 SV=1 | 2.66 | 0.0998 |
| RNASE7 | Ribonuclease 7 OS=Homo sapiens OX=9606 GN=RNASE7 PE=1 SV=2 | 2.65 | 0.0857 |
| LORICRIN | Loricrin OS=Homo sapiens OX=9606 GN=LORICRIN PE=1 SV=2 | 2.64 | 0.1588 |
| SERPINB12 | Serpin B12 OS=Homo sapiens OX=9606 GN=SERPINB12 PE=1 SV=1 | 2.61 | 0.0477 |
| KRT78 | Keratin, type II cytoskeletal 78 OS=Homo sapiens OX=9606 GN=KRT78 PE=1 SV=2 | 2.59 | 0.0777 |
| KRT77 | Keratin, type II cytoskeletal 1b OS=Homo sapiens OX=9606 GN=KRT77 PE=1 SV=3 | 2.58 | 0.0830 |
| MYCBP2 | E3 ubiquitin-protein ligase MYCBP2 OS=Homo sapiens OX=9606 GN=MYCBP2 PE=1 SV=4 | 2.56 | 0.0322 |
| ECM1 | Extracellular matrix protein 1 OS=Homo sapiens OX=9606 GN=ECM1 PE=1 SV=2 | 2.51 | 0.1058 |
| CACNA1S | Voltage-dependent L-type calcium channel subunit alpha-1S OS=Homo sapiens OX=9606 GN=CACNA1S PE=1 SV=4 | 2.46 | 0.1260 |
| ARG1 | Arginase-1 OS=Homo sapiens OX=9606 GN=ARG1 PE=1 SV=2 | 2.45 | 0.1200 |
| CNFN | Cornifelin OS=Homo sapiens OX=9606 GN=CNFN PE=1 SV=2 | 2.45 | 0.0972 |
| KRT80 | Keratin, type II cytoskeletal 80 OS=Homo sapiens OX=9606 GN=KRT80 PE=1 SV=2 | 2.42 | 0.0996 |
| KRT23 | Keratin, type I cytoskeletal 23 OS=Homo sapiens OX=9606 GN=KRT23 PE=1 SV=2 | 2.42 | 0.1085 |
| TGM3 | Protein-glutamine gamma-glutamyltransferase E OS=Homo sapiens OX=9606 GN=TGM3 PE=1 SV=4 | 2.42 | 0.0876 |
| ALOXE3 | Hydroperoxide isomerase ALOXE3 OS=Homo sapiens OX=9606 GN=ALOXE3 PE=1 SV=1 | 2.41 | 0.0375 |
| C1QC | Complement C1q subcomponent subunit C OS=Homo sapiens OX=9606 GN=C1QC PE=1 SV=3 | 2.41 | 0.1206 |
| SCGB2A2 | Mammaglobin-A OS=Homo sapiens OX=9606 GN=SCGB2A2 PE=1 SV=1 | 2.36 | 0.0991 |
| FLG2 | Filaggrin-2 OS=Homo sapiens OX=9606 GN=FLG2 PE=1 SV=1 | 2.33 | 0.0976 |
| CPA4 | Carboxypeptidase A4 OS=Homo sapiens OX=9606 GN=CPA4 PE=1 SV=2 | 2.31 | 0.0035 |
| WFDC12 | WAP four-disulfide core domain protein 12 OS=Homo sapiens OX=9606 GN=WFDC12 PE=1 SV=1 | 2.29 | 0.0523 |
| NEU2 | Sialidase-2 OS=Homo sapiens OX=9606 GN=NEU2 PE=1 SV=2 | 2.28 | 0.1642 |
| SERPINB2 | Plasminogen activator inhibitor 2 OS=Homo sapiens OX=9606 GN=SERPINB2 PE=1 SV=2 | 2.28 | 0.0905 |
| IGHV3-7 | Immunoglobulin heavy variable 3-7 OS=Homo sapiens OX=9606 GN=IGHV3-7 PE=1 SV=2 | 2.20 | 0.1912 |
| HAL | Histidine ammonia-lyase OS=Homo sapiens OX=9606 GN=HAL PE=1 SV=1 | 2.17 | 0.1023 |
| PSAPL1 | Proactivator polypeptide-like 1 OS=Homo sapiens OX=9606 GN=PSAPL1 PE=2 SV=2 | 2.15 | 0.0928 |
| KRT15 | Keratin, type I cytoskeletal 15 OS=Homo sapiens OX=9606 GN=KRT15 PE=1 SV=3 | 2.15 | 0.1999 |
| CTSG | Cathepsin G OS=Homo sapiens OX=9606 GN=CTSG PE=1 SV=2 | 2.12 | 0.1592 |
| CENPH | Centromere protein H OS=Homo sapiens OX=9606 GN=CENPH PE=1 SV=1 | 2.11 | 0.1721 |
| LYZ | Lysozyme C OS=Homo sapiens OX=9606 GN=LYZ PE=1 SV=1 | 2.11 | 0.0386 |
| ANXA2 | Annexin A2 OS=Homo sapiens OX=9606 GN=ANXA2 PE=1 SV=2 | 2.10 | 0.1025 |
| TUBB2A | Tubulin beta-2A chain OS=Homo sapiens OX=9606 GN=TUBB2A PE=1 SV=1 | 2.09 | 0.0591 |
| PLBD1 | Phospholipase B-like 1 OS=Homo sapiens OX=9606 GN=PLBD1 PE=1 SV=2 | 2.08 | 0.1621 |
| RAB20 | Ras-related protein Rab-20 OS=Homo sapiens OX=9606 GN=RAB20 PE=1 SV=1 | 2.07 | 0.0591 |
| SERPINB3 | Serpin B3 OS=Homo sapiens OX=9606 GN=SERPINB3 PE=1 SV=2 | 2.06 | 0.0952 |
| EVPL | Envoplakin OS=Homo sapiens OX=9606 GN=EVPL PE=1 SV=3 | 2.03 | 0.1148 |
| IL1RN | Interleukin-1 receptor antagonist protein OS=Homo sapiens OX=9606 GN=IL1RN PE=1 SV=1 | 2.02 | 0.0675 |
| KLK7 | Kallikrein-7 OS=Homo sapiens OX=9606 GN=KLK7 PE=1 SV=1 | 2.00 | 0.0980 |
| NCCRP1 | F-box only protein 50 OS=Homo sapiens OX=9606 GN=NCCRP1 PE=1 SV=1 | 1.98 | 0.1875 |
| LTF | Lactotransferrin OS=Homo sapiens OX=9606 GN=LTF PE=1 SV=6 | 1.96 | 0.0223 |
| S100A10 | Protein S100-A10 OS=Homo sapiens OX=9606 GN=S100A10 PE=1 SV=2 | 1.90 | 0.1447 |
| IL37 | Interleukin-37 OS=Homo sapiens OX=9606 GN=IL37 PE=1 SV=1 | 1.81 | 0.1926 |
| PIP | Prolactin-inducible protein OS=Homo sapiens OX=9606 GN=PIP PE=1 SV=1 | 1.71 | 0.1932 |
| PIGR | Polymeric immunoglobulin receptor OS=Homo sapiens OX=9606 GN=PIGR PE=1 SV=4 | 1.53 | 0.1815 |
| APOA4 | Apolipoprotein A-IV OS=Homo sapiens OX=9606 GN=APOA4 PE=1 SV=4 | 1.47 | 0.1445 |
| MTERF3 | Transcription termination factor 3, mitochondrial OS=Homo sapiens OX=9606 GN=MTERF3 PE=1 SV=2 | 1.46 | 0.0942 |
| ESYT1 | Extended synaptotagmin-1 OS=Homo sapiens OX=9606 GN=ESYT1 PE=1 SV=1 | 1.42 | 0.0821 |
| SNRPD3 | Small nuclear ribonucleoprotein Sm D3 OS=Homo sapiens OX=9606 GN=SNRPD3 PE=1 SV=1 | 1.33 | 0.0000 |
| SERPINB5 | Serpin B5 OS=Homo sapiens OX=9606 GN=SERPINB5 PE=1 SV=2 | 1.18 | 0.1898 |
| BLMH | Bleomycin hydrolase OS=Homo sapiens OX=9606 GN=BLMH PE=1 SV=1 | 1.17 | 0.1231 |
| FUT8 | Alpha-(1,6)-fucosyltransferase OS=Homo sapiens OX=9606 GN=FUT8 PE=1 SV=2 | 0.63 | 0.0930 |
| ELMO2 | Engulfment and cell motility protein 2 OS=Homo sapiens OX=9606 GN=ELMO2 PE=1 SV=2 | 0.60 | 0.1686 |
| RPL8 | 60S ribosomal protein L8 OS=Homo sapiens OX=9606 GN=RPL8 PE=1 SV=2 | 0.59 | 0.1141 |
| IRF6 | Interferon regulatory factor 6 OS=Homo sapiens OX=9606 GN=IRF6 PE=1 SV=1 | 0.54 | 0.0860 |
| TPD52 | Tumor protein D52 OS=Homo sapiens OX=9606 GN=TPD52 PE=1 SV=2 | -0.58 | 0.0997 |
| GET3 | ATPase GET3 OS=Homo sapiens OX=9606 GN=GET3 PE=1 SV=2 | -0.60 | 0.1012 |
| LGALS8 | Galectin-8 OS=Homo sapiens OX=9606 GN=LGALS8 PE=1 SV=4 | -0.60 | 0.1871 |
| RAB8B | Ras-related protein Rab-8B OS=Homo sapiens OX=9606 GN=RAB8B PE=1 SV=2 | -0.62 | 0.1912 |
| CSRP1 | Cysteine and glycine-rich protein 1 OS=Homo sapiens OX=9606 GN=CSRP1 PE=1 SV=3 | -0.62 | 0.1464 |
| CPVL | Probable serine carboxypeptidase CPVL OS=Homo sapiens OX=9606 GN=CPVL PE=1 SV=2 | -0.68 | 0.0257 |
| HTRA1 | Serine protease HTRA1 OS=Homo sapiens OX=9606 GN=HTRA1 PE=1 SV=1 | -0.73 | 0.0810 |
| AGPAT3 | 1-acyl-sn-glycerol-3-phosphate acyltransferase gamma OS=Homo sapiens OX=9606 GN=AGPAT3 PE=1 SV=1 | -0.79 | 0.1660 |
| CHMP2A | Charged multivesicular body protein 2a OS=Homo sapiens OX=9606 GN=CHMP2A PE=1 SV=1 | -0.81 | 0.0880 |
| DERL1 | Derlin-1 OS=Homo sapiens OX=9606 GN=DERL1 PE=1 SV=1 | -0.83 | 0.0822 |
| CENPS | Centromere protein S OS=Homo sapiens OX=9606 GN=CENPS PE=1 SV=1 | -0.87 | 0.0906 |
| ATP5MF | ATP synthase subunit f, mitochondrial OS=Homo sapiens OX=9606 GN=ATP5MF PE=1 SV=3 | -0.88 | 0.1369 |
| DMAC2L | ATP synthase subunit s, mitochondrial OS=Homo sapiens OX=9606 GN=DMAC2L PE=1 SV=3 | -1.11 | 0.0858 |
| MECP2 | Methyl-CpG-binding protein 2 OS=Homo sapiens OX=9606 GN=MECP2 PE=1 SV=1 | -1.20 | 0.1319 |
| SRP19 | Signal recognition particle 19 kDa protein OS=Homo sapiens OX=9606 GN=SRP19 PE=1 SV=3 | -1.31 | 0.0171 |
| SLC27A1 | Long-chain fatty acid transport protein 1 OS=Homo sapiens OX=9606 GN=SLC27A1 PE=1 SV=1 | -2.07 | 0.0790 |
| PKP2 | Plakophilin-2 OS=Homo sapiens OX=9606 GN=PKP2 PE=1 SV=2 | -2.13 | 0.0369 |
| UTP20 | Small subunit processome component 20 homolog OS=Homo sapiens OX=9606 GN=UTP20 PE=1 SV=3 | -2.53 | 0.0743 |
| PPIE | Peptidyl-prolyl cis-trans isomerase E OS=Homo sapiens OX=9606 GN=PPIE PE=1 SV=1 | -2.66 | 0.0009 |
| HGH1 | Protein HGH1 homolog OS=Homo sapiens OX=9606 GN=HGH1 PE=1 SV=1 | -2.66 | 0.1758 |
| BRI3BP | BRI3-binding protein OS=Homo sapiens OX=9606 GN=BRI3BP PE=1 SV=1 | -2.67 | 0.0325 |

| **Supplementary Table 11. Functional enrichment analysis with g:Profiler of DAPs (FDR<0.2%) specific to ffEVs-treated oocytes** | | | | |
| --- | --- | --- | --- | --- |
| **GO.ID** | **Description** | **p.Val** | **FDR** | **Genes** |
| GO:0030280 | structural constituent of skin epidermis | 8.67E-08 | 8.67E-08 | KRT6A,KRT72,KRT5,LORICRIN,KRT78,KRT77,KRT80 |
| GO:0004867 | serine-type endopeptidase inhibitor activity | 0.000162732 | 0.000162732 | SLPI,SERPINB12,WFDC12,SERPINB2,ANXA2,SERPINB3,SERPINB5 |
| GO:0004866 | endopeptidase inhibitor activity | 0.000348695 | 0.000348695 | SLPI,SERPINB12,WFDC12,SERPINB2,ANXA2,SERPINB3,LTF,SERPINB5 |
| GO:0030414 | peptidase inhibitor activity | 0.000454729 | 0.000454729 | SLPI,SERPINB12,WFDC12,SERPINB2,ANXA2,SERPINB3,LTF,SERPINB5 |
| GO:0061135 | endopeptidase regulator activity | 0.000692632 | 0.000692632 | SLPI,SERPINB12,WFDC12,SERPINB2,ANXA2,SERPINB3,LTF,SERPINB5 |
| GO:0061134 | peptidase regulator activity | 0.003290001 | 0.003290001 | SLPI,SERPINB12,WFDC12,SERPINB2,ANXA2,SERPINB3,LTF,SERPINB5 |
| GO:0005198 | structural molecule activity | 0.00936865 | 0.00936865 | KRT6A,KRT72,KRT5,LORICRIN,KRT78,KRT77,ECM1,KRT80,KRT23,TGM3,FLG2,KRT15,TUBB2A,EVPL,RPL8,CSRP1 |
| GO:0004180 | carboxypeptidase activity | 0.025674092 | 0.025674092 | PRCP,CPA4,BLMH,CPVL |
| GO:0031424 | keratinization | 4.70E-14 | 4.70E-14 | SPRR2F,KRT6A,SPRR2D,KRT72,TGM1,KRT5,LORICRIN,KRT78,KRT77,CNFN,KRT80,TGM3,EVPL |
| GO:0008544 | epidermis development | 7.64E-12 | 7.64E-12 | SPRR2F,KRT6A,SPRR2D,KRT72,TGM1,KRT5,XP32,LORICRIN,KRT78,KRT77,CNFN,KRT80,TGM3,ALOXE3,FLG2,KRT15,EVPL,KLK7,IRF6 |
| GO:0030216 | keratinocyte differentiation | 5.16E-11 | 5.16E-11 | SPRR2F,KRT6A,SPRR2D,KRT72,TGM1,KRT5,LORICRIN,KRT78,KRT77,CNFN,KRT80,TGM3,EVPL,IRF6 |
| GO:0043588 | skin development | 5.80E-11 | 5.80E-11 | SPRR2F,KRT6A,ASPRV1,SPRR2D,KRT72,TGM1,KRT5,LORICRIN,KRT78,KRT77,CNFN,KRT80,TGM3,ALOXE3,FLG2,EVPL,IRF6 |
| GO:0009913 | epidermal cell differentiation | 6.43E-09 | 6.43E-09 | SPRR2F,KRT6A,SPRR2D,KRT72,TGM1,KRT5,LORICRIN,KRT78,KRT77,CNFN,KRT80,TGM3,EVPL,IRF6 |
| GO:0045104 | intermediate filament cytoskeleton organization | 1.76E-08 | 1.76E-08 | KRT6A,KRT72,KRT5,KRT78,KRT77,KRT80,KRT23,KRT15,EVPL,PKP2 |
| GO:0045103 | intermediate filament-based process | 1.96E-08 | 1.96E-08 | KRT6A,KRT72,KRT5,KRT78,KRT77,KRT80,KRT23,KRT15,EVPL,PKP2 |
| GO:0045109 | intermediate filament organization | 5.33E-08 | 5.33E-08 | KRT6A,KRT72,KRT5,KRT78,KRT77,KRT80,KRT23,KRT15,PKP2 |
| GO:0009888 | tissue development | 1.80426E-05 | 1.80426E-05 | SPRR2F,KRT6A,SPRR2D,KRT72,TGM1,KRT5,XP32,LORICRIN,KRT78,KRT77,ECM1,CACNA1S,CNFN,KRT80,KRT23,TGM3,ALOXE3,FLG2,PSAPL1,KRT15,SERPINB3,EVPL,KLK7,LTF,SERPINB5,IRF6,CSRP1,PKP2 |
| GO:0030855 | epithelial cell differentiation | 2.73325E-05 | 2.73325E-05 | SPRR2F,KRT6A,SPRR2D,KRT72,TGM1,KRT5,LORICRIN,KRT78,KRT77,CNFN,KRT80,KRT23,TGM3,PSAPL1,KRT15,EVPL,IRF6 |
| GO:0019730 | antimicrobial humoral response | 0.000201226 | 0.000201226 | KRT6A,SLPI,RNASE7,WFDC12,CTSG,LYZ,KLK7,LTF |
| GO:0060429 | epithelium development | 0.000346429 | 0.000346429 | SPRR2F,KRT6A,SPRR2D,KRT72,TGM1,KRT5,LORICRIN,KRT78,KRT77,CNFN,KRT80,KRT23,TGM3,ALOXE3,FLG2,PSAPL1,KRT15,EVPL,SERPINB5,IRF6 |
| GO:0019731 | antibacterial humoral response | 0.001022204 | 0.001022204 | SLPI,RNASE7,WFDC12,CTSG,KLK7,LTF |
| GO:0006959 | humoral immune response | 0.00309644 | 0.00309644 | KRT6A,SLPI,RNASE7,C1QC,WFDC12,CTSG,LYZ,KLK7,LTF |
| GO:0048856 | anatomical structure development | 0.004934914 | 0.004934914 | SPRR2F,KRT6A,SRCIN1,ASPRV1,GPNMB,SPRR2D,PRCP,KRT72,TGM1,KRT5,CIC,XP32,LORICRIN,SERPINB12,KRT78,KRT77,MYCBP2,ECM1,CACNA1S,CNFN,KRT80,KRT23,TGM3,ALOXE3,C1QC,FLG2,PSAPL1,KRT15,ANXA2,TUBB2A,SERPINB3,EVPL,IL1RN,KLK7,LTF,S100A10,APOA4,SERPINB5,FUT8,IRF6,TPD52,LGALS8,CSRP1,HTRA1,MECP2,PKP2 |
| GO:0141061 | disruption of cell in another organism | 0.006140337 | 0.006140337 | KRT6A,RNASE7,ARG1,CTSG,LYZ,LTF |
| GO:0031640 | killing of cells of another organism | 0.006140337 | 0.006140337 | KRT6A,RNASE7,ARG1,CTSG,LYZ,LTF |
| GO:0141060 | disruption of anatomical structure in another organism | 0.007408509 | 0.007408509 | KRT6A,RNASE7,ARG1,CTSG,LYZ,LTF |
| GO:0007010 | cytoskeleton organization | 0.009340654 | 0.009340654 | KRT6A,SRCIN1,KRT72,KRT5,LORICRIN,KRT78,KRT77,MYCBP2,KRT80,KRT23,KRT15,CENPH,TUBB2A,EVPL,S100A10,ELMO2,CSRP1,CHMP2A,MECP2,PKP2 |
| GO:0018149 | peptide cross-linking | 0.013515265 | 0.013515265 | TGM1,LORICRIN,TGM3,EVPL |
| GO:0042742 | defense response to bacterium | 0.023019045 | 0.023019045 | KRT6A,GSDMA,SLPI,RNASE7,WFDC12,CTSG,LYZ,KLK7,LTF |
| GO:0097435 | supramolecular fiber organization | 0.027221242 | 0.027221242 | KRT6A,KRT72,KRT5,KRT78,KRT77,KRT80,KRT23,KRT15,ANXA2,S100A10,ELMO2,CSRP1,MECP2,PKP2 |
| GO:0048513 | animal organ development | 0.028176533 | 0.028176533 | SPRR2F,KRT6A,ASPRV1,SPRR2D,KRT72,TGM1,KRT5,CIC,LORICRIN,KRT78,KRT77,ECM1,CACNA1S,CNFN,KRT80,TGM3,ALOXE3,FLG2,PSAPL1,ANXA2,TUBB2A,SERPINB3,EVPL,LTF,SERPINB5,IRF6,HTRA1,MECP2,PKP2 |
| GO:0001533 | cornified envelope | 4.38E-21 | 4.38E-21 | SPRR2F,SPRR2D,TGM1,DSG1,LORICRIN,SERPINB12,KRT77,CNFN,FLG2,SERPINB2,ANXA2,EVPL,KLK7,SERPINB5,PKP2 |
| GO:0005615 | extracellular space | 1.39E-16 | 1.39E-16 | KRT6A,KPRP,SLPI,SELENBP1,PRCP,KRT72,TGM1,KRT5,RNASE7,SERPINB12,KRT78,KRT77,ECM1,ARG1,TGM3,C1QC,SCGB2A2,CPA4,WFDC12,SERPINB2,IGHV3-7,KRT15,CTSG,LYZ,ANXA2,TUBB2A,PLBD1,SERPINB3,EVPL,IL1RN,KLK7,NCCRP1,LTF,S100A10,IL37,PIP,PIGR,APOA4,SERPINB5,BLMH,FUT8,IRF6,GET3,LGALS8,RAB8B,CSRP1,CPVL,HTRA1,CHMP2A,MECP2 |
| GO:0005576 | extracellular region | 2.18E-14 | 2.18E-14 | KRT6A,KPRP,SLPI,SELENBP1,PRCP,KRT72,TGM1,KRT5,RNASE7,SERPINB12,KRT78,KRT77,ECM1,ARG1,TGM3,C1QC,SCGB2A2,FLG2,CPA4,WFDC12,SERPINB2,IGHV3-7,PSAPL1,KRT15,CTSG,LYZ,ANXA2,TUBB2A,PLBD1,SERPINB3,EVPL,IL1RN,KLK7,NCCRP1,LTF,S100A10,IL37,PIP,PIGR,APOA4,SERPINB5,BLMH,FUT8,IRF6,GET3,LGALS8,RAB8B,CSRP1,CPVL,HTRA1,CHMP2A,MECP2,PPIE |
| GO:0070062 | extracellular exosome | 4.28E-12 | 4.28E-12 | KRT6A,KPRP,SLPI,SELENBP1,PRCP,KRT72,TGM1,KRT5,KRT78,KRT77,ECM1,TGM3,IGHV3-7,KRT15,CTSG,LYZ,ANXA2,TUBB2A,SERPINB3,EVPL,IL1RN,NCCRP1,LTF,S100A10,PIP,PIGR,APOA4,BLMH,FUT8,IRF6,GET3,RAB8B,CSRP1,CPVL,HTRA1,CHMP2A |
| GO:1903561 | extracellular vesicle | 6.06E-12 | 6.06E-12 | KRT6A,KPRP,SLPI,SELENBP1,PRCP,KRT72,TGM1,KRT5,KRT78,KRT77,ECM1,TGM3,IGHV3-7,KRT15,CTSG,LYZ,ANXA2,TUBB2A,SERPINB3,EVPL,IL1RN,NCCRP1,LTF,S100A10,PIP,PIGR,APOA4,BLMH,FUT8,IRF6,GET3,RAB8B,CSRP1,CPVL,HTRA1,CHMP2A |
| GO:0065010 | extracellular membrane-bounded organelle | 6.15E-12 | 6.15E-12 | KRT6A,KPRP,SLPI,SELENBP1,PRCP,KRT72,TGM1,KRT5,KRT78,KRT77,ECM1,TGM3,IGHV3-7,KRT15,CTSG,LYZ,ANXA2,TUBB2A,SERPINB3,EVPL,IL1RN,NCCRP1,LTF,S100A10,PIP,PIGR,APOA4,BLMH,FUT8,IRF6,GET3,RAB8B,CSRP1,CPVL,HTRA1,CHMP2A |
| GO:0043230 | extracellular organelle | 6.15E-12 | 6.15E-12 | KRT6A,KPRP,SLPI,SELENBP1,PRCP,KRT72,TGM1,KRT5,KRT78,KRT77,ECM1,TGM3,IGHV3-7,KRT15,CTSG,LYZ,ANXA2,TUBB2A,SERPINB3,EVPL,IL1RN,NCCRP1,LTF,S100A10,PIP,PIGR,APOA4,BLMH,FUT8,IRF6,GET3,RAB8B,CSRP1,CPVL,HTRA1,CHMP2A |
| GO:0031982 | vesicle | 3.90E-10 | 3.90E-10 | KRT6A,GPNMB,KPRP,SLPI,SELENBP1,PRCP,KRT72,TGM1,KRT5,DSG1,SERPINB12,KRT78,KRT77,ECM1,ARG1,TGM3,FLG2,IGHV3-7,KRT15,CTSG,LYZ,ANXA2,TUBB2A,RAB20,SERPINB3,EVPL,IL1RN,KLK7,NCCRP1,LTF,S100A10,PIP,PIGR,APOA4,BLMH,FUT8,IRF6,GET3,LGALS8,RAB8B,CSRP1,CPVL,HTRA1,CHMP2A,DERL1,PPIE |
| GO:0005737 | cytoplasm | 3.17E-09 | 3.17E-09 | HAS1,SPRR2F,KRT6A,SRCIN1,TEP1,GPNMB,GSDMA,KPRP,SDR9C7,SLPI,SPRR2D,SELENBP1,PRCP,KRT72,TGM1,KRT5,DSG1,RNASE7,LORICRIN,SERPINB12,KRT78,KRT77,MYCBP2,ECM1,CACNA1S,ARG1,CNFN,KRT80,KRT23,TGM3,ALOXE3,FLG2,NEU2,SERPINB2,HAL,PSAPL1,KRT15,CTSG,CENPH,LYZ,ANXA2,TUBB2A,PLBD1,RAB20,SERPINB3,EVPL,IL1RN,KLK7,NCCRP1,LTF,S100A10,IL37,PIGR,APOA4,MTERF3,ESYT1,SERPINB5,BLMH,FUT8,ELMO2,RPL8,IRF6,TPD52,GET3,LGALS8,RAB8B,CSRP1,HTRA1,AGPAT3,CHMP2A,DERL1,ATP5MF,DMAC2L,MECP2,SRP19,SLC27A1,PKP2,UTP20,PPIE,BRI3BP |
| GO:0005882 | intermediate filament | 6.26658E-06 | 6.26658E-06 | KRT6A,KRT72,KRT5,KRT78,KRT77,KRT80,KRT23,KRT15,EVPL,PKP2 |
| GO:0045111 | intermediate filament cytoskeleton | 2.64336E-05 | 2.64336E-05 | KRT6A,KRT72,KRT5,KRT78,KRT77,KRT80,KRT23,KRT15,EVPL,PKP2 |
| GO:0045095 | keratin filament | 0.000921814 | 0.000921814 | KRT6A,KRT72,KRT5,KRT78,KRT77,KRT80 |
| GO:0030141 | secretory granule | 0.000960308 | 0.000960308 | SLPI,PRCP,DSG1,SERPINB12,ECM1,ARG1,FLG2,CTSG,LYZ,ANXA2,SERPINB3,KLK7,LTF,PIGR,PPIE |
| GO:0005766 | primary lysosome | 0.00098359 | 0.00098359 | PRCP,ARG1,CTSG,LYZ,ANXA2,SERPINB3,PIGR |
| GO:0042582 | azurophil granule | 0.00098359 | 0.00098359 | PRCP,ARG1,CTSG,LYZ,ANXA2,SERPINB3,PIGR |
| GO:0005829 | cytosol | 0.001701251 | 0.001701251 | SPRR2F,KRT6A,GSDMA,SPRR2D,SELENBP1,KRT72,TGM1,KRT5,DSG1,LORICRIN,KRT78,KRT77,MYCBP2,ARG1,KRT80,KRT23,ALOXE3,NEU2,HAL,PSAPL1,KRT15,CTSG,CENPH,ANXA2,PLBD1,SERPINB3,EVPL,IL1RN,NCCRP1,IL37,APOA4,MTERF3,BLMH,ELMO2,RPL8,IRF6,LGALS8,HTRA1,CHMP2A,MECP2,SRP19,PPIE |
| GO:0099503 | secretory vesicle | 0.001812327 | 0.001812327 | SLPI,PRCP,DSG1,SERPINB12,ECM1,ARG1,FLG2,CTSG,LYZ,ANXA2,SERPINB3,KLK7,LTF,PIGR,RAB8B,PPIE |
| GO:0034774 | secretory granule lumen | 0.001939164 | 0.001939164 | SLPI,ECM1,ARG1,CTSG,LYZ,ANXA2,SERPINB3,LTF,PPIE |
| GO:0060205 | cytoplasmic vesicle lumen | 0.002090435 | 0.002090435 | SLPI,ECM1,ARG1,CTSG,LYZ,ANXA2,SERPINB3,LTF,PPIE |
| GO:0031983 | vesicle lumen | 0.002143055 | 0.002143055 | SLPI,ECM1,ARG1,CTSG,LYZ,ANXA2,SERPINB3,LTF,PPIE |
| GO:1990665 | AnxA2-p11 complex | 0.005032316 | 0.005032316 | ANXA2,S100A10 |
| GO:0035578 | azurophil granule lumen | 0.010846181 | 0.010846181 | ARG1,CTSG,LYZ,ANXA2,SERPINB3 |
| GO:0070820 | tertiary granule | 0.016824884 | 0.016824884 | PRCP,DSG1,SERPINB12,FLG2,LYZ,LTF |
| GO:0062023 | collagen-containing extracellular matrix | 0.018116801 | 0.018116801 | SLPI,SERPINB12,ECM1,C1QC,CTSG,ANXA2,S100A10,APOA4,HTRA1 |
| GO:0099512 | supramolecular fiber | 0.027129256 | 0.027129256 | KRT6A,KRT72,KRT5,KRT78,KRT77,CACNA1S,KRT80,KRT23,KRT15,TUBB2A,EVPL,CSRP1,CHMP2A,PKP2 |
| GO:0099081 | supramolecular polymer | 0.029392355 | 0.029392355 | KRT6A,KRT72,KRT5,KRT78,KRT77,CACNA1S,KRT80,KRT23,KRT15,TUBB2A,EVPL,CSRP1,CHMP2A,PKP2 |
| GO:0099513 | polymeric cytoskeletal fiber | 0.031926937 | 0.031926937 | KRT6A,KRT72,KRT5,KRT78,KRT77,KRT80,KRT23,KRT15,TUBB2A,EVPL,CHMP2A,PKP2 |
| GO:0035580 | specific granule lumen | 0.03449424 | 0.03449424 | SLPI,ARG1,LYZ,LTF |
| KEGG:05150 | Staphylococcus aureus infection | 0.044687588 | 0.044687588 | DSG1,KRT23,C1QC,KRT15 |
| REAC:R-HSA-6809371 | Formation of the cornified envelope | 7.20E-13 | 7.20E-13 | SPRR2F,KRT6A,SPRR2D,KRT72,TGM1,KRT5,DSG1,LORICRIN,KRT78,KRT77,KRT80,KRT23,KRT15,EVPL,PKP2 |
| REAC:R-HSA-6805567 | Keratinization | 1.62E-09 | 1.62E-09 | SPRR2F,KRT6A,SPRR2D,KRT72,TGM1,KRT5,DSG1,LORICRIN,KRT78,KRT77,KRT80,KRT23,KRT15,EVPL,PKP2 |
| REAC:R-HSA-6798695 | Neutrophil degranulation | 0.003232762 | 0.003232762 | SLPI,PRCP,DSG1,SERPINB12,ARG1,FLG2,CTSG,LYZ,ANXA2,SERPINB3,LTF,PIGR,PPIE |
| REAC:R-HSA-75205 | Dissolution of Fibrin Clot | 0.028088045 | 0.028088045 | SERPINB2,ANXA2,S100A10 |
| WP:WP5390 | Pancreatic cancer subtypes | 0.010255944 | 0.010255944 | KRT6A,KRT15,LYZ,SERPINB3 |
| TF:M12021_1 | Factor: T-box; motif: NASGTGTNAN; match class: 1 | 0.027395241 | 0.027395241 | SPRR2F,KRT78,KRT77,ARG1,ALOXE3,SCGB2A2,LYZ,NCCRP1,LTF,CSRP1,MECP2,PPIE |
| HPA:0461401 | skin 1; cells in corneal layer[‚â•Low] | 8.48E-14 | 8.48E-14 | KRT6A,KPRP,TGM1,DSG1,LORICRIN,ECM1,CNFN,C1QC,HAL,KRT15,SERPINB3,KLK7,SERPINB5,BLMH |
| HPA:0461402 | skin 1; cells in corneal layer[‚â•Medium] | 3.43E-11 | 3.43E-11 | KRT6A,KPRP,TGM1,DSG1,LORICRIN,ECM1,KRT15,SERPINB3,KLK7,SERPINB5,BLMH |
| HPA:0471402 | skin 2; cells in corneal layer[‚â•Medium] | 1.69E-09 | 1.69E-09 | KRT6A,ASPRV1,TGM1,DSG1,LORICRIN,ECM1,CNFN,KRT15,SERPINB3,KLK7,SERPINB5 |
| HPA:0471401 | skin 2; cells in corneal layer[‚â•Low] | 1.04E-08 | 1.04E-08 | KRT6A,ASPRV1,TGM1,DSG1,LORICRIN,ECM1,CNFN,HAL,KRT15,SERPINB3,KLK7,SERPINB5 |
| HPA:0460181 | skin 1; cells in granular layer[‚â•Low] | 9.79261E-06 | 9.79261E-06 | KRT6A,ASPRV1,KPRP,DSG1,LORICRIN,ECM1,SERPINB2,HAL,KRT15,TUBB2A,SERPINB3,EVPL,IL1RN,KLK7,SERPINB5,BLMH,RPL8,TPD52,CHMP2A |
| HPA:0460182 | skin 1; cells in granular layer[‚â•Medium] | 5.61062E-05 | 5.61062E-05 | KRT6A,ASPRV1,KPRP,DSG1,LORICRIN,HAL,KRT15,TUBB2A,SERPINB3,EVPL,KLK7,SERPINB5,BLMH,RPL8,TPD52,CHMP2A |
| HPA:0470181 | skin 2; cells in granular layer[‚â•Low] | 6.47558E-05 | 6.47558E-05 | KRT6A,ASPRV1,KPRP,DSG1,LORICRIN,CNFN,SERPINB2,HAL,KRT15,TUBB2A,SERPINB3,EVPL,IL1RN,KLK7,SERPINB5,RPL8,TPD52,CHMP2A |
| HPA:0471411 | skin 2; cells in spinous layer[‚â•Low] | 0.000232572 | 0.000232572 | KRT6A,KRT5,DSG1,LORICRIN,CNFN,C1QC,SERPINB2,HAL,KRT15,ANXA2,TUBB2A,SERPINB3,EVPL,IL1RN,S100A10,SERPINB5,RPL8,TPD52,CHMP2A |
| HPA:0470182 | skin 2; cells in granular layer[‚â•Medium] | 0.000354013 | 0.000354013 | KRT6A,ASPRV1,KPRP,LORICRIN,CNFN,SERPINB2,HAL,KRT15,TUBB2A,SERPINB3,EVPL,KLK7,SERPINB5,RPL8,CHMP2A |
| HPA:0471412 | skin 2; cells in spinous layer[‚â•Medium] | 0.000820318 | 0.000820318 | KRT6A,KRT5,DSG1,LORICRIN,CNFN,HAL,KRT15,ANXA2,TUBB2A,SERPINB3,EVPL,S100A10,SERPINB5,RPL8,TPD52,CHMP2A |
| HPA:0461411 | skin 1; cells in spinous layer[‚â•Low] | 0.003197219 | 0.003197219 | KRT6A,KRT5,DSG1,LORICRIN,HAL,KRT15,ANXA2,TUBB2A,SERPINB3,EVPL,IL1RN,S100A10,SERPINB5,RPL8,TPD52,CHMP2A |
| HPA:0461412 | skin 1; cells in spinous layer[‚â•Medium] | 0.003372016 | 0.003372016 | KRT6A,KRT5,DSG1,LORICRIN,HAL,KRT15,ANXA2,TUBB2A,SERPINB3,EVPL,SERPINB5,RPL8,TPD52,CHMP2A |
| HPA:0470183 | skin 2; cells in granular layer[High] | 0.005301454 | 0.005301454 | KRT6A,ASPRV1,KPRP,LORICRIN,CNFN,HAL,TUBB2A,SERPINB3,SERPINB5 |
| HPA:0461403 | skin 1; cells in corneal layer[High] | 0.01190623 | 0.01190623 | KRT6A,DSG1,SERPINB3,BLMH |
| HPA:0450000 | skin | 0.015227881 | 0.015227881 | GSDMA,KRT77,KRT15 |
| HPA:0471403 | skin 2; cells in corneal layer[High] | 0.016898528 | 0.016898528 | KRT6A,ASPRV1,DSG1,SERPINB3 |
| HPA:0460183 | skin 1; cells in granular layer[High] | 0.035611376 | 0.035611376 | KRT6A,ASPRV1,HAL,TUBB2A,SERPINB3,EVPL,SERPINB5,BLMH |
| HP:0000962 | Hyperkeratosis | 6.89413E-05 | 6.89413E-05 | KRT6A,ASPRV1,SDR9C7,TGM1,KRT5,DSG1,LORICRIN,ECM1,ALOXE3,IL1RN,ELMO2 |
| HP:0011122 | Abnormality of skin physiology | 0.000346966 | 0.000346966 | ASPRV1,GPNMB,SDR9C7,TGM1,KRT5,DSG1,CIC,ECM1,ALOXE3,C1QC,FLG2,LYZ,IL1RN,MECP2,SRP19 |
| HP:0011123 | Inflammatory abnormality of the skin | 0.000480912 | 0.000480912 | ASPRV1,SDR9C7,TGM1,KRT5,DSG1,CIC,ECM1,ALOXE3,C1QC,FLG2,LYZ,IL1RN,MECP2,SRP19 |
| HP:0011368 | Epidermal thickening | 0.001227836 | 0.001227836 | KRT6A,ASPRV1,SDR9C7,TGM1,KRT5,DSG1,LORICRIN,ECM1,ALOXE3,FLG2,IL1RN,ELMO2 |
| HP:0011124 | Abnormal epidermal morphology | 0.001560308 | 0.001560308 | TGM1,DSG1,LORICRIN,ECM1,ALOXE3,FLG2,IL1RN |
| HP:0001596 | Alopecia | 0.002026871 | 0.002026871 | KRT6A,SDR9C7,TGM1,KRT5,LORICRIN,ECM1,TGM3,ALOXE3,HTRA1,MECP2 |
| HP:0000982 | Palmoplantar keratoderma | 0.00430591 | 0.00430591 | KRT6A,SDR9C7,TGM1,KRT5,DSG1,LORICRIN,ALOXE3 |
| HP:0001072 | Thickened skin | 0.004651786 | 0.004651786 | KRT6A,ASPRV1,SDR9C7,TGM1,KRT5,DSG1,LORICRIN,ECM1,ALOXE3,FLG2,IL1RN,ELMO2 |
| HP:0025114 | Hypergranulosis | 0.009148125 | 0.009148125 | SDR9C7,DSG1,LORICRIN,ALOXE3 |
| HP:0000972 | Palmoplantar hyperkeratosis | 0.01028304 | 0.01028304 | KRT6A,SDR9C7,TGM1,KRT5,DSG1,LORICRIN,ALOXE3 |
| HP:0010765 | Palmar hyperkeratosis | 0.011445809 | 0.011445809 | KRT6A,SDR9C7,TGM1,KRT5,DSG1,LORICRIN,ALOXE3 |
| HP:0007556 | Plantar hyperkeratosis | 0.011445809 | 0.011445809 | KRT6A,SDR9C7,TGM1,KRT5,DSG1,LORICRIN,ALOXE3 |
| HP:0007479 | Congenital nonbullous ichthyosiform erythroderma | 0.013417209 | 0.013417209 | ASPRV1,TGM1,LORICRIN,ALOXE3 |
| HP:0025092 | Epidermal acanthosis | 0.018973752 | 0.018973752 | TGM1,DSG1,LORICRIN,ALOXE3,IL1RN |
| HP:0000989 | Pruritus | 0.031789725 | 0.031789725 | ASPRV1,GPNMB,SDR9C7,TGM1,KRT5,ALOXE3,FLG2 |
| HP:0007431 | Congenital ichthyosiform erythroderma | 0.046038971 | 0.046038971 | ASPRV1,TGM1,LORICRIN,ALOXE3 |
| HP:0007446 | Palmoplantar blistering | 0.046467022 | 0.046467022 | KRT6A,KRT5,DSG1 |
